# Protein Structure Prediction with Expectation Reflection

**DOI:** 10.1101/2022.07.12.499755

**Authors:** Evan Cresswell-Clay, Danh-Tai Hoang, Joe McKenna, Chris Yang, Eric Zhang, Vipul Periwal

## Abstract

Sequence covariation in multiple sequence alignments of homologous proteins has been used extensively to obtain insights into protein structure. However, global statistical inference is required in order to ascertain direct relationships between amino acid positions in these sequences that are not simply secondary correlations induced by interactions with a third residue. Methods for statistical inference of such covariation have been developed to exploit the growing availability of sequence data. These hints about the folded protein structure provide critical *a priori* information for more detailed 3D predictions by neural networks. We present a novel method for protein structure inference using an iterative parameter-free model estimator which uses the formalism of statistical physics. With no tunable learning rate, our method scales to large system sizes while providing improved performance in the regime of small sample sizes. We apply this method to 40974 PDB structures and compare its performance to that of other methods. Our method outperforms existing methods for 76% of analysed proteins.

## 1 Introduction

Proteins are central actors in biological function so understanding protein function is of significant importance in pharmaceutical research [Uhlig et al., 2014, Patel et al., 2007], drug design [Craik et al., 2013, Fosgerau & Hoffmann, 2015], disease diagnosis [Stalmach et al., 2014, Gautam et al., 2014], and therapy [Vlieghe et al., 2010, Lau & Dunn, 2018]. Protein structure prediction has been important since Kendrew and Perut’s research teams established the relation between function and structure through investigation of myoglobin and hemoglobin tertiary structures [Perutz et al., 1960, Kendrew et al., 1958]. Thus, if a given protein’s function is closely tied to its three-dimensional native structure, deducing this structure is key to such medical applications.

As organisms reproduce, non-synonymous mutations in DNA lead to altered protein sequences that sample the space of possible sequences. Therefore the compilation of evolutionary protein sequence records such as PFAM [Mistry et al., 2020] gives a large sampling of sequence space for a given evolutionary function of the protein family [Altschuh et al., 1987]. As a protein evolves, those aspects of the spatial proximity of amino acids (AA) that are required for function are encoded as co-evolution signals in the protein residue sequence [Altschuh et al., 1987, Miller & Eisenberg, 2008]. Therefore the phylogenetic analysis of proteins can provide insights into the functionally important contacts in the protein’s network of interacting residues. This concept led to the exploitation of correlations in amino acid residues for determining connections in protein structure [Altschuh et al., 1988, Göbel et al., 1994, Atchley et al., 2000, Fodor & Aldrich, 2004, Fariselli et al., 2001].

However, simply using correlation as a marker for spatial proximity is confounded by the existence of evolutionary constraints which are not related to spatial information [Dunn et al., 2008, Burger & Van Nimwegen, 2010, Lapedes et al., 2002]. This problem resulted in the development of methods to sift spatial information from evolutionary sequence records beginning with the maximum entropy approach, introduced by [Giraud et al., 1999, Lapedes et al., 2002] and expanded using several varying approaches in implementation [Weigt et al., 2009, Burger & Van Nimwegen, 2008, Mora et al., 2010]. In addition, global statistical inference is required in order to infer direct relationships between amino acid positions in these sequences that are not simply correlations that could arise as a result of third party influence. This potential issue led to the most recent advance in protein contact predictions: the implementation of direct coupling analysis (DCA) giving inferred AA-pair contacts through the calculation of direct information (DI) leading to greatly improved contact prediction [Morcos et al., 2011] and through that prediction, a more efficient 3D protein structure prediction by constraining the search of possible protein conformations [Marks et al., 2011]. Several variants of DCA have emerged through different approaches to inferring the model parameters. Two popular variants are the original mean field model (MF) and a later variant using pseudo-likelihood maximization (PLM).

The evolutionary constraints generated by DCA help to reduce the conformation space considered in 3D structure prediction. Indeed, current neural network PSP systems rely on the output of the DCA family of approaches as one of the inputs into machine learning the tertiary protein structure [Jones et al., 2015, Wang et al., 2017, Golkov et al., 2016, Liu et al., 2018, Senior et al., 2020, AlQuraishi, 2019]. Therefore tertiary structure prediction remains an important problem for all scales of protein structure prediction. This work is concerned with improving the contact prediction in protein tertiary structure, building on the work cited above.

### 1.1 Protein Structure Prediction using Expectation Reflection

We present a novel variant of DCA (DCA-ER) which also uses sequence covariation to infer the network of interacting residues in proteins. The novelty in our approach lies in the specific algorithm used to compute the pairwise residue interaction coefficients in the same formalism of statistical physics used in previous work [Lapedes et al., 2002, Morcos et al., 2011, Ekeberg et al., 2014]. Thus, we define a formal free energy of observations from a partition function with an energy function as usual. However, we use a multiplicative iteration to update the interaction coefficients that avoids the pitfalls of cost minimization. This is a key step because, even with the plethora of sequence information available now, the number of sequences for any family of proteins is still minuscule relative to the available configuration space of all sequences. Our method has no tunable learning rate, scales to large system sizes, and is particularly suited to sparse high-dimensional datasets to infer networks of such interacting residues [Hoang et al., 2019a, Hoang et al., 2019b]. We analyze 40974 PDB structures and demonstrate that our method outperforms DCA and PLM methods across all ranges of protein family size and sequence length. When comparing the true-positive versus false-positive rates of contact prediction of our method with the predictions of other DCA variants, our method provided the most accurate contact prediction for a significant majority of structures.

## 2 Results

### 2.1 Single Protein Example: 1ZDR

We begin our analysis by presenting the contact predictions of a single protein structure (PDB-id: 1ZDR, Pfam: PF00186). Running Expectation Reflection to infer DCA model couplings and then calculate DI, we compute a list of ordered amino acid column pairs. In Table 1, we summarize these pairs by listing the top 5 pairs with their DCA-calculated Direct Information (DI).

**Table 1:** Top 5 amino acid pair contacts by direct information.

| Position 1 | Position 2 | DI |
| --- | --- | --- |
| 12 | 121 | 0.2996 |
| 49 | 50 | 0.1966 |
| 35 | 55 | 0.1689 |
| 57 | 72 | 0.1632 |
| 135 | 138 | 0.1475 |

We can visualize the spatial information in our calculated DI by plotting the values against the distance between the associated AA-pair in Figure 2. In Figure 2, it is important to note the location of strongly linked amino acid pairs. Pairs with larger DI values generally show spatial proximity and as one increases AA-pair distance this direct information decreases.

**Figure 1:**
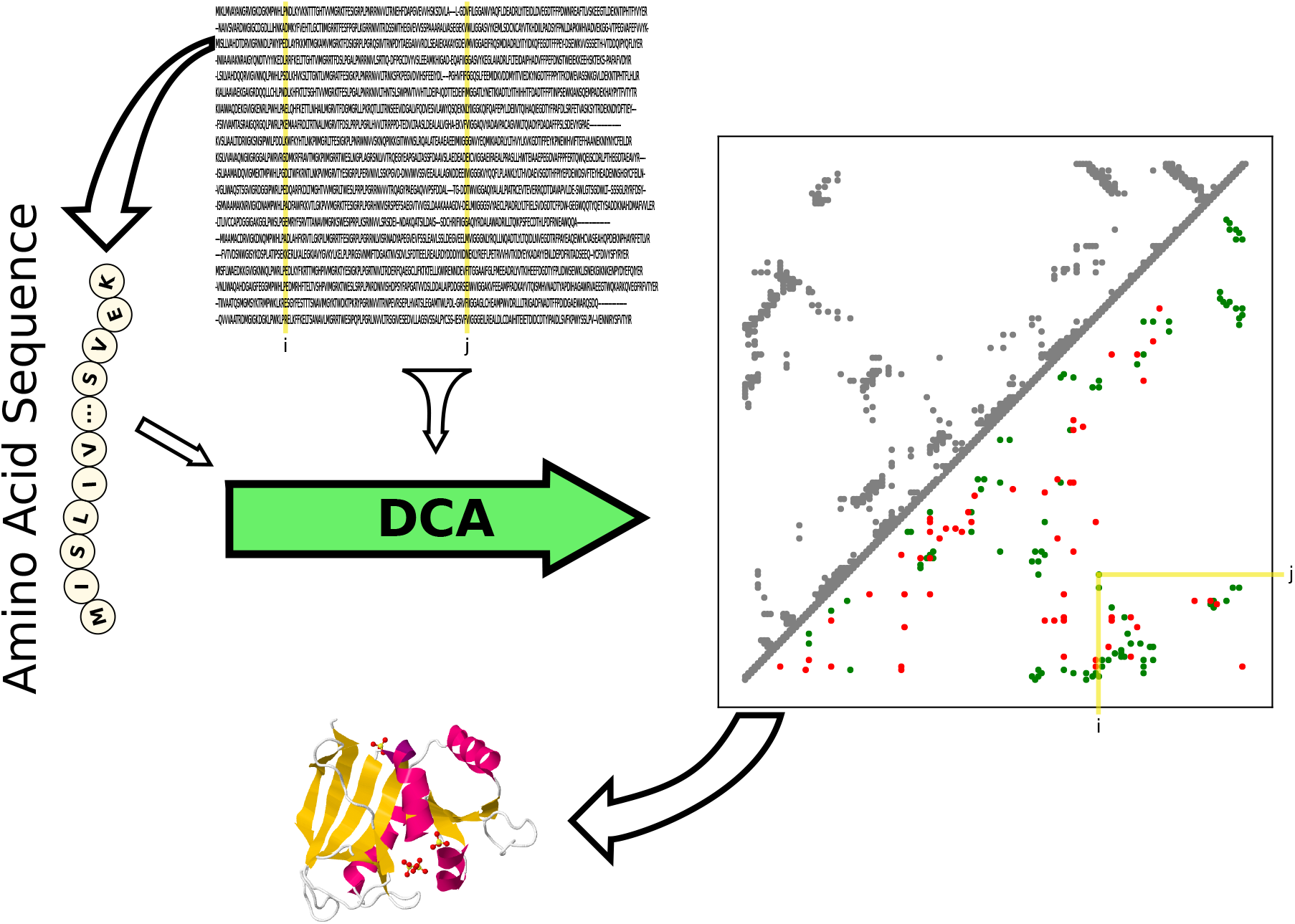
Schematic of Protein Contact Prediction using Co-Evolution. Direct Coupling Analysis uses co-evolution of amino acid position pairs across aligned sets of homologous protein sequences to infer contacts in the native structure.

**Figure 2:**
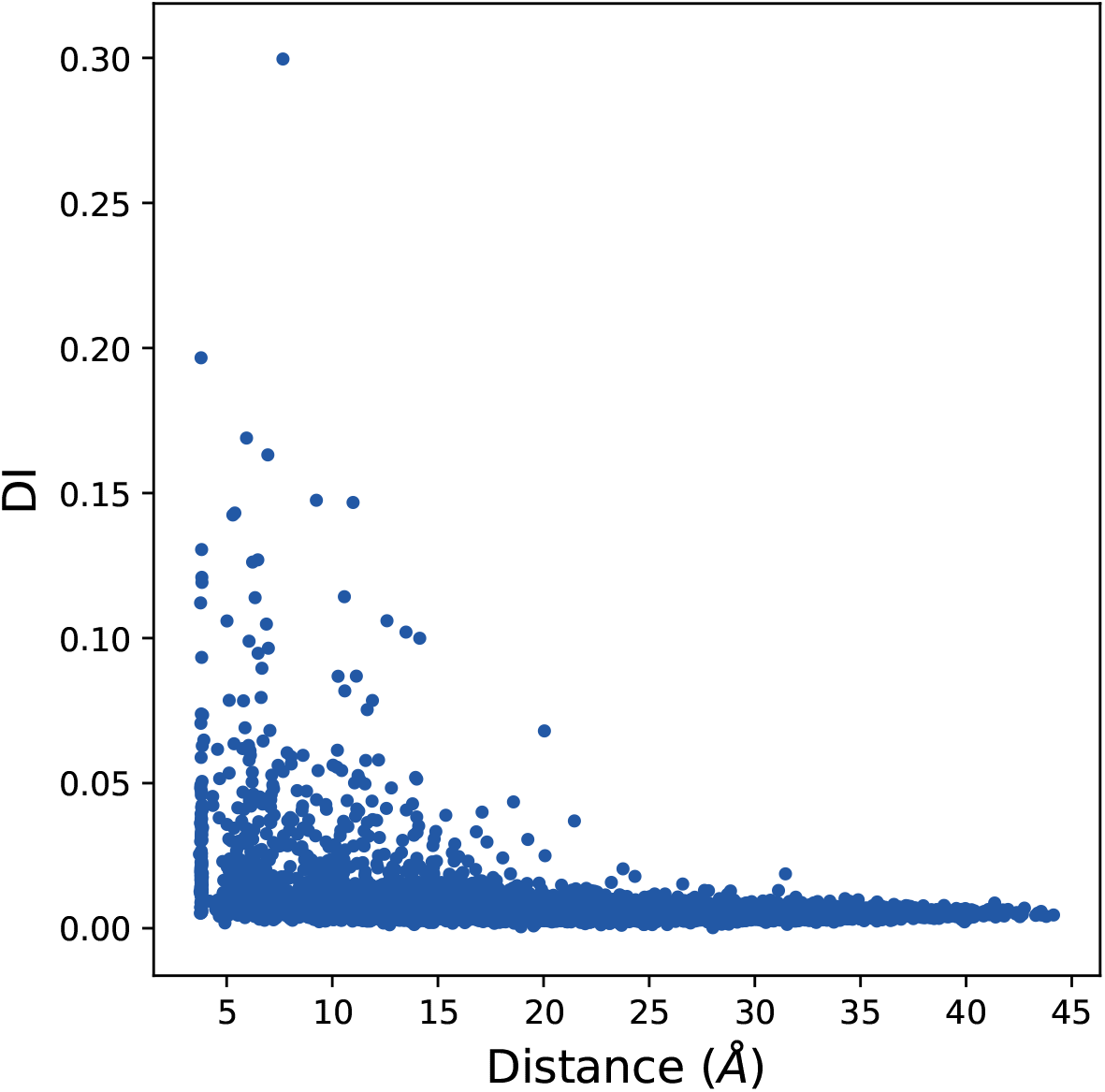
AA-Pair Distance vs Direct Information. In order to visualize the spatial information encoded in our calculated DI pairs we plot the AA-pair DI against the distance between the AA-pair

#### 2.1.1 Prediction Accuracy

Given our ER-calculated values of DI we can analyse the accuracy of the resulting contact predictions by considering the true positive rate of these predictions. In Figure 3 true positive rate (blue) is plotted against the rank to judge the accuracy of the model. Perfect accuracy is plotted as well (black) for comparison. We also visualize the predicted tertiary structure on a 2D mapping in Figure 4, with correct predictions plotted in green and incorrect predictions plotted in red (lower triangle) and actual contacts from the PDB structure in grey (upper triangle).

**Figure 3:**
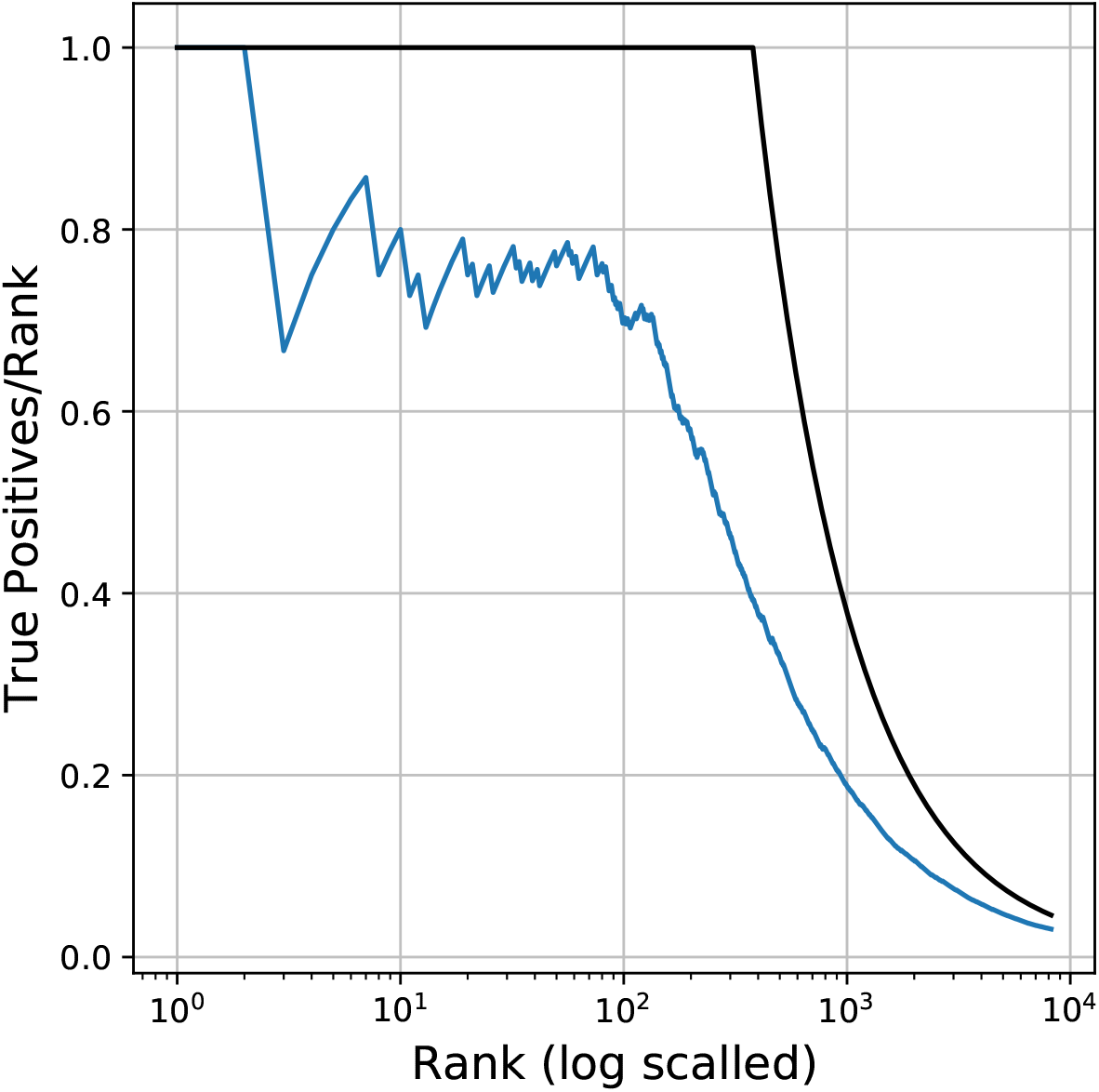
True Positive Rate of ER Contacts. In order to visualize the accuracy of direct information calculated with Expectation Reflection, we plot the true positive(TP) rate of contact predictions with DCA-ER.

**Figure 4:**
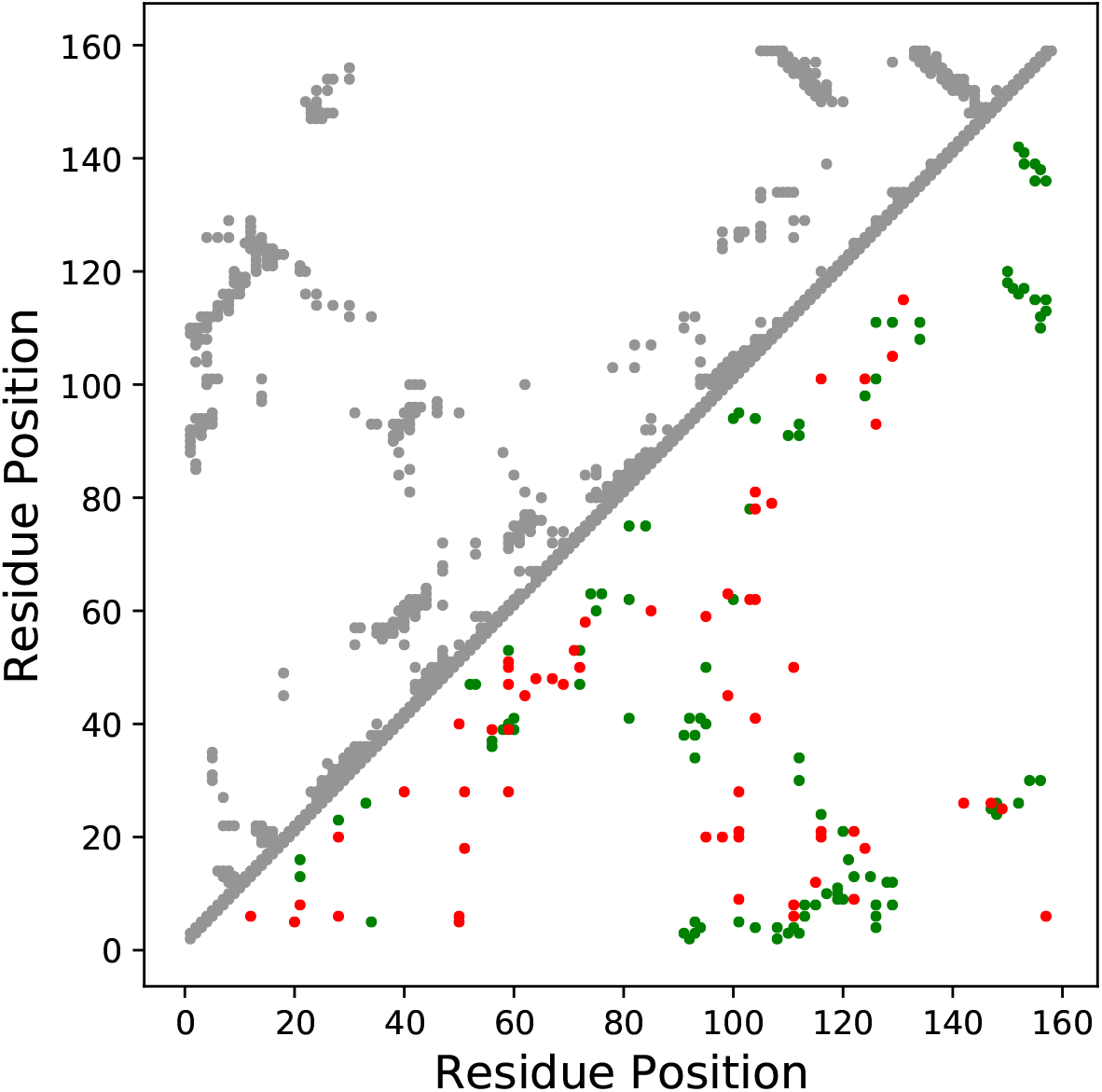
Contact Map of ER Contacts. In order to visualize the tertiary structure of contacts predicted with Expectation Reflection, we plot the contact map with the PDB-contacts (grey) in the upper-triangular region and the true (green) and false (red) DCA-ER contact predictions in the lower triangular region.

#### 2.1.2 Comparing with other methods

It is important to consider the accuracy of the ER DCA variant contact prediction in the context of existing methods. In order to verify the efficacy of our method we compare it against two existing popular variants which differ in their method of calculation of underlying couplings in the Potts model (see Section 4.1.2),

1. Mean-Field (MF) [Morcos et al., 2011]
2. Pseudo-Likelihood Maximization (PLM) [Ekeberg et al., 2014]

We use the implementation of these methods in the PyDCA package [Zerihun et al., 2020]. We begin our comparison by considering the example PDB structure 1ZDR from the previous section. We plot the DI AA-pair contact predictions of ER against the other methods where green lists successful predictions and red represents unsuccessful predictions in Figure 5. From Figure 5, we see a skew towards improved predictions by ER. Though the scale of the DI cannot be compared between methods, if we follow any diagonal split of the two axes (method DI strengths) we see that successful contacts appear weighted in favor of ER DI-strength. This visualization is helpful for intuition but we must extend our analysis, as before, to consider the TP rate. In Figure 3 we show the TP rate curve of the ER derived AA contact predictions. We add the TP rate curves of PLM- and MF-derived AA contacts in Figure 6 for comparison. Because of the noisy nature of the low-rank TP rate we also present the Receiver Operating Characteristic (ROC) curve which provides a smoother metric for comparing the three methods. From the ROC curves we can also quantify the differences between different methods by measuring the area under the curve of each method (AUC). Returning to our example of 1ZDR, the AUC values of ER and other methods are given in Table 2, showing that ER outperforms existing DCA variants.

**Figure 5:**
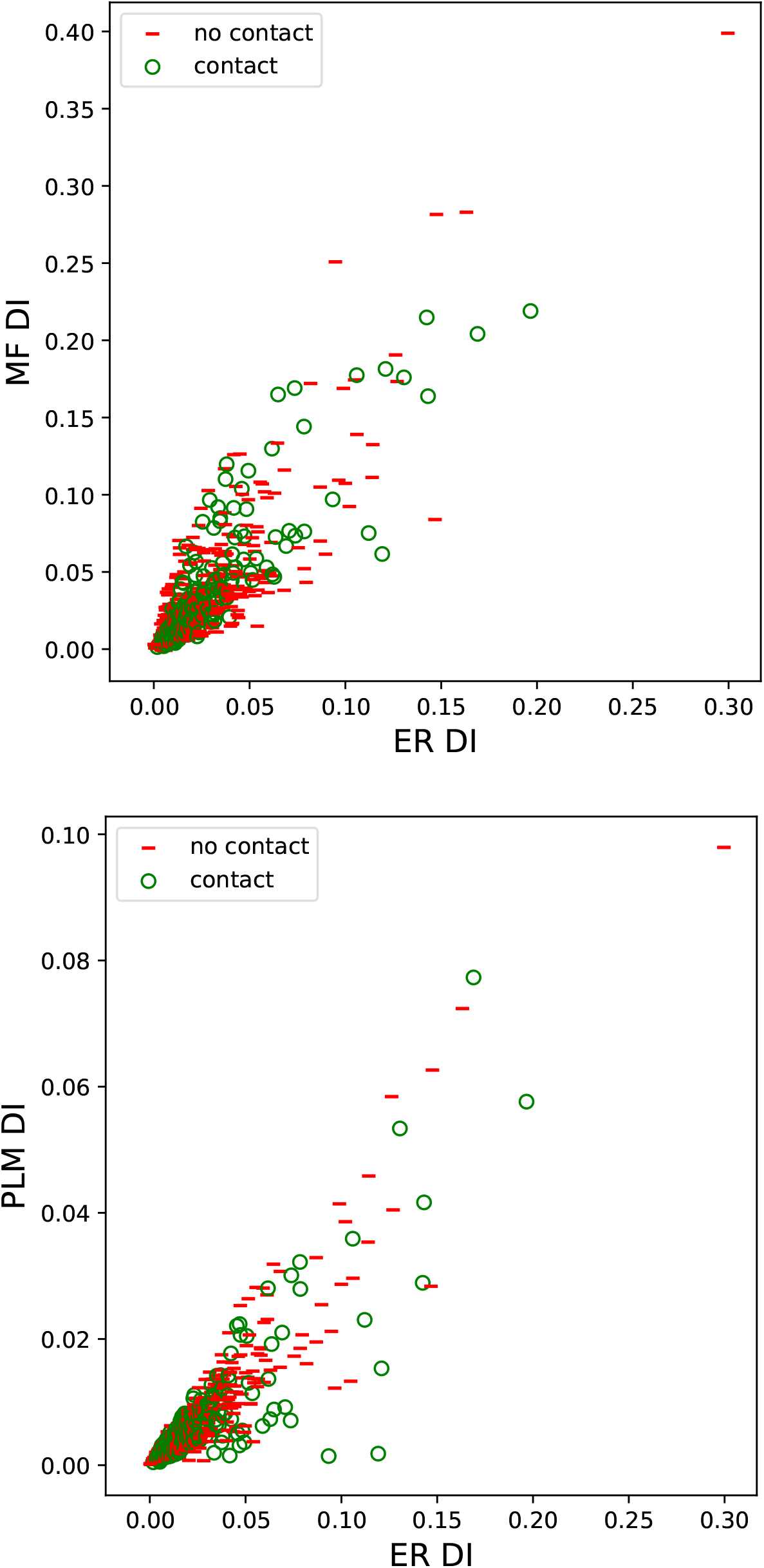
DI Contact Predictions of 1ZDR. In order to visualize differences in DI between DCA-ER and other DCA variants, we plot ER DI values against the MF (top) and PLM (bottom) variant DI values with green circles representing successful contact predictions and red dashes representing unsuccessful predictions.

**Figure 6:**
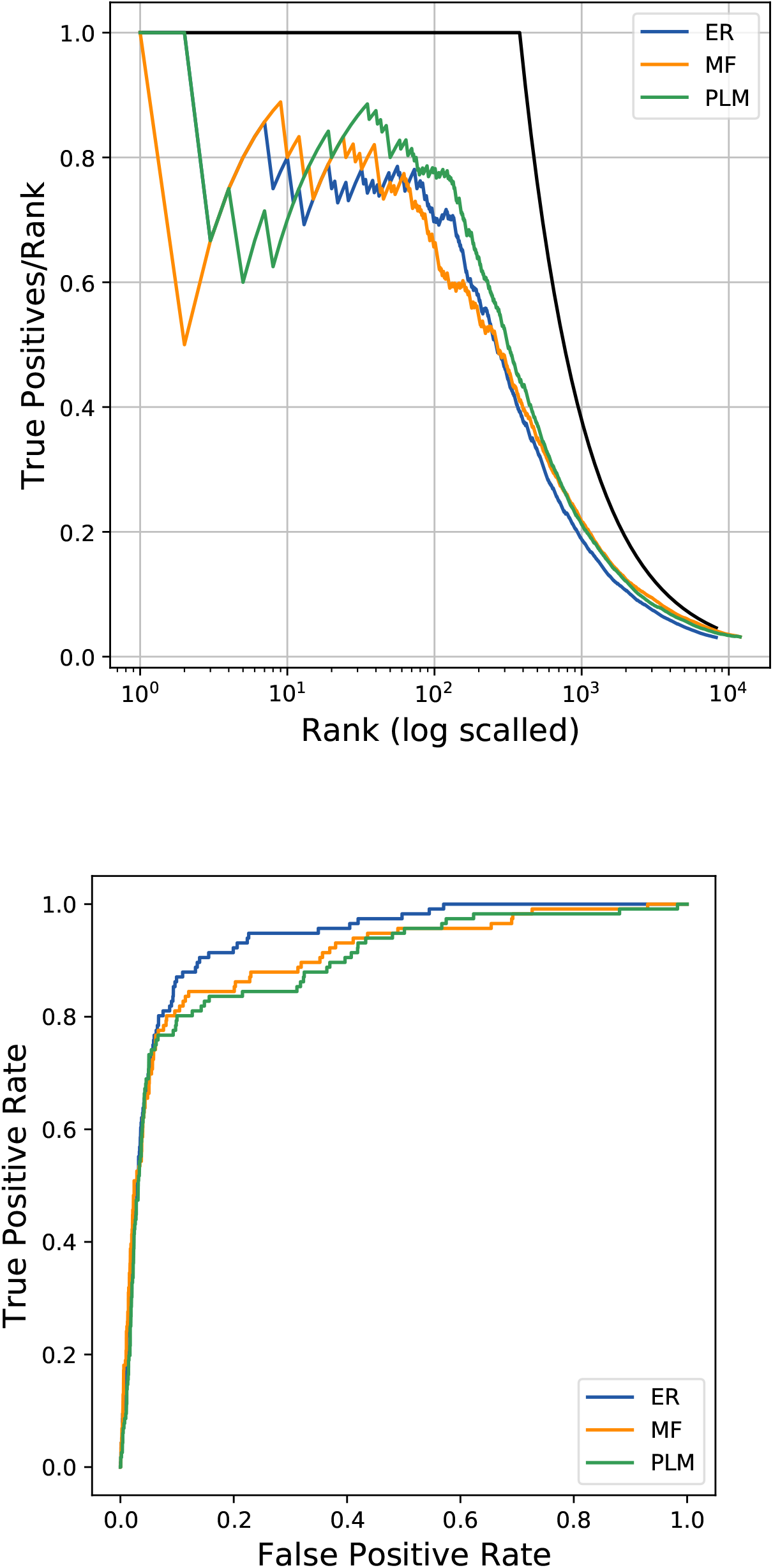
Receiver Operating Characteristic of 1ZDR Predictions. In order to visualize the accuracy the direct information calculated by DCA-ER, we plot the ROC curve of contact predictions.

**Table 2:** ROC curve AUC of each method.

| Method | AUC |
| --- | --- |
| ER | .936 |
| MF | .906 |
| PLM | .895 |

#### 2.1.3 Verifying a Clear Difference in Prediction

As seen above for this specific PDB-ID, ER improves on the other methods. However, we want to ensure that any improvement in accuracy is not marginal. In order to verify a clear difference in PSP we utilise a two-sided Kolmogorov-Smirnov test [Massey, 1951] with the null hypothesis that the two prediction samples are the same. For further explanation see Section 4.2.

From Table 3, we can establish a statistical significance for the difference between the ROC of Expection Reflection’s prediction and those of the other methods by calculating the critical value for which the hypothesis that two given ROC AUCs are the same can be rejected. In this case the critical value as calculated in Equation 8 in Section 4.2 is *D*_*crit*_ = .016. Comparing *D*_*crit*_ against the KS statistics in Table 3 shows that the methods are providing distinguishable results from each other. Therefore in this example we can assert that ER outperforms the other methods in a statistically significant way.

**Table 3:** Kolmogorov-Smirnov Statistics comparing Results of different methods.

| Method 1 | Method 2 | KS Statistic | p-value |
| --- | --- | --- | --- |
| ER | MF | 0.0862 | 2.119e-28 |
| ER | PLM | 0.1379 | 3.330e-72 |
| MF | PLM | 0.1552 | 2.280e-91 |

### 2.2 Method Comparison of 40974 Proteins

We have introduced our analysis for individual proteins and compared Expectation Reflection against other popular methods. In order to ensure that the improvement seen in ER is not merely a sampling fluke, we now extend the same analysis to 40974 proteins in order to see how ER contact predictions compare against the other methods on a larger scale. We can begin by considering several contact maps generated across proteins with various MSA sizes in Figure 7 In Figure 8 we bin the area under the curve (AUC) of the ROC curves of all three methods for each of the 40974 protein. From this histogram we can immediately see that the AUC histogram for Expectation Reflection is farther shifted to the right than the compared methods. This demonstrates the increased efficacy of ER in contact predictions.

**Figure 7:**
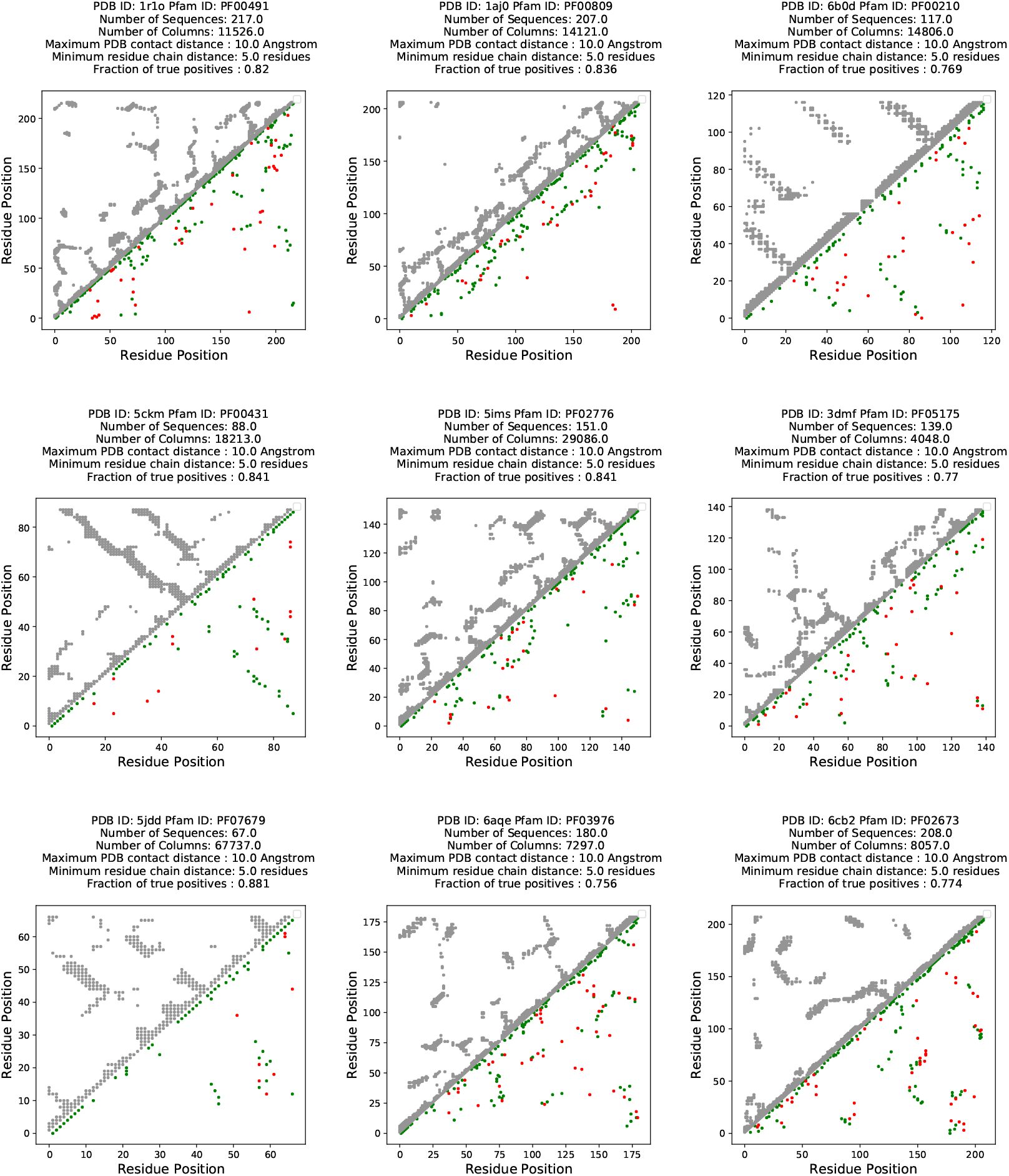
ER Prediction Contact Maps. Of the 40974 analysed proteins with DCA-ER, we plot nine example contact maps with PDB contacts (grey) compared against true positive (green) and false positive (red) predictions.

**Figure 8:**
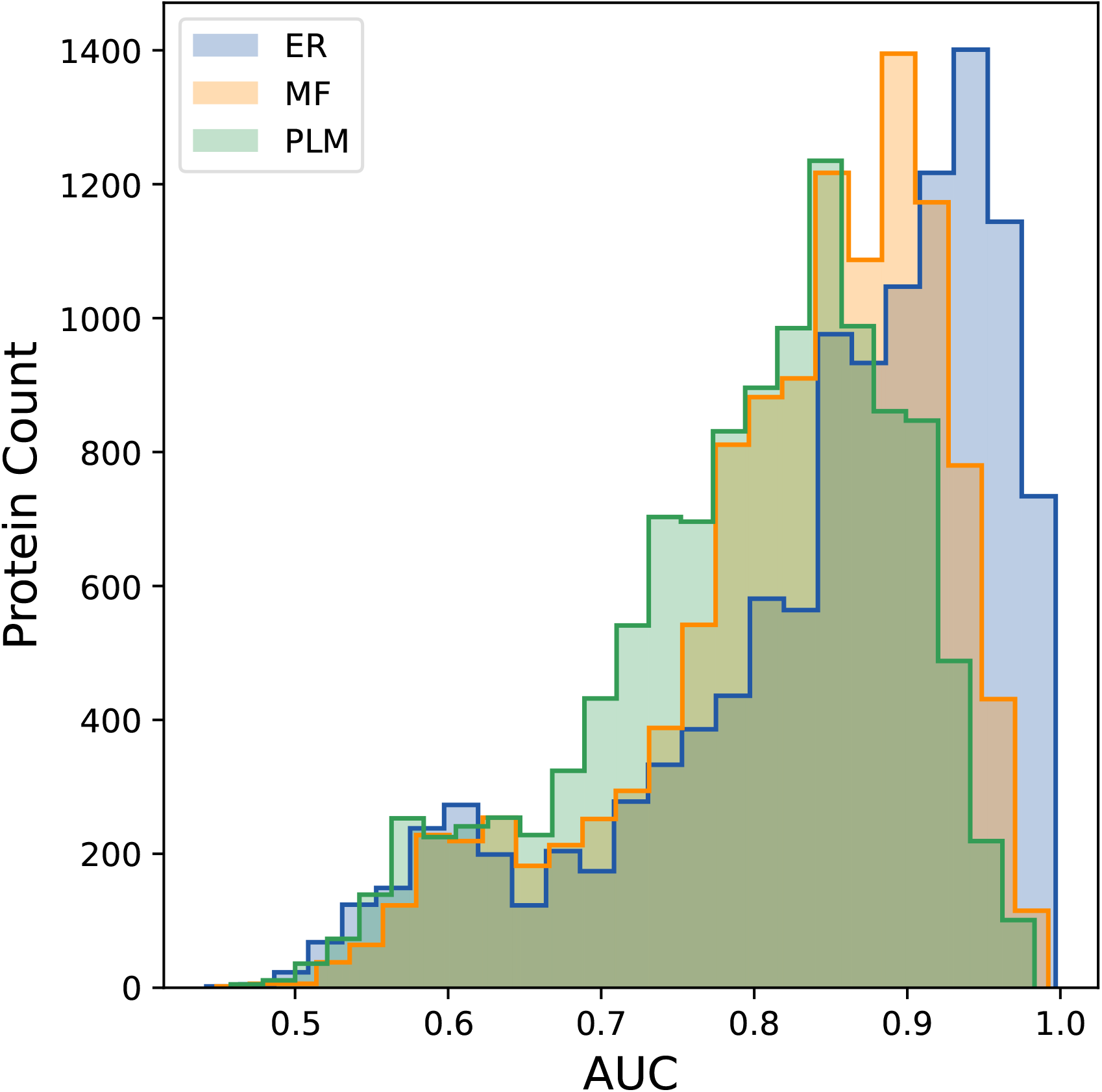
AUC Values. We bin the AUC values for the ROC curve for all 40974 proteins for each DCA variant.

#### 2.2.1 Finding the Best Method

To better visualize this comparison, we use the ROC-AUC value for every protein to identify which method provided the best contact prediction. In Figure 9 we plot the number of proteins where each method gave the best prediction. From the inlay in Figure 9 we see that ER outperforms MF and PLM for 76% of the 40974 proteins considered.

**Figure 9:**
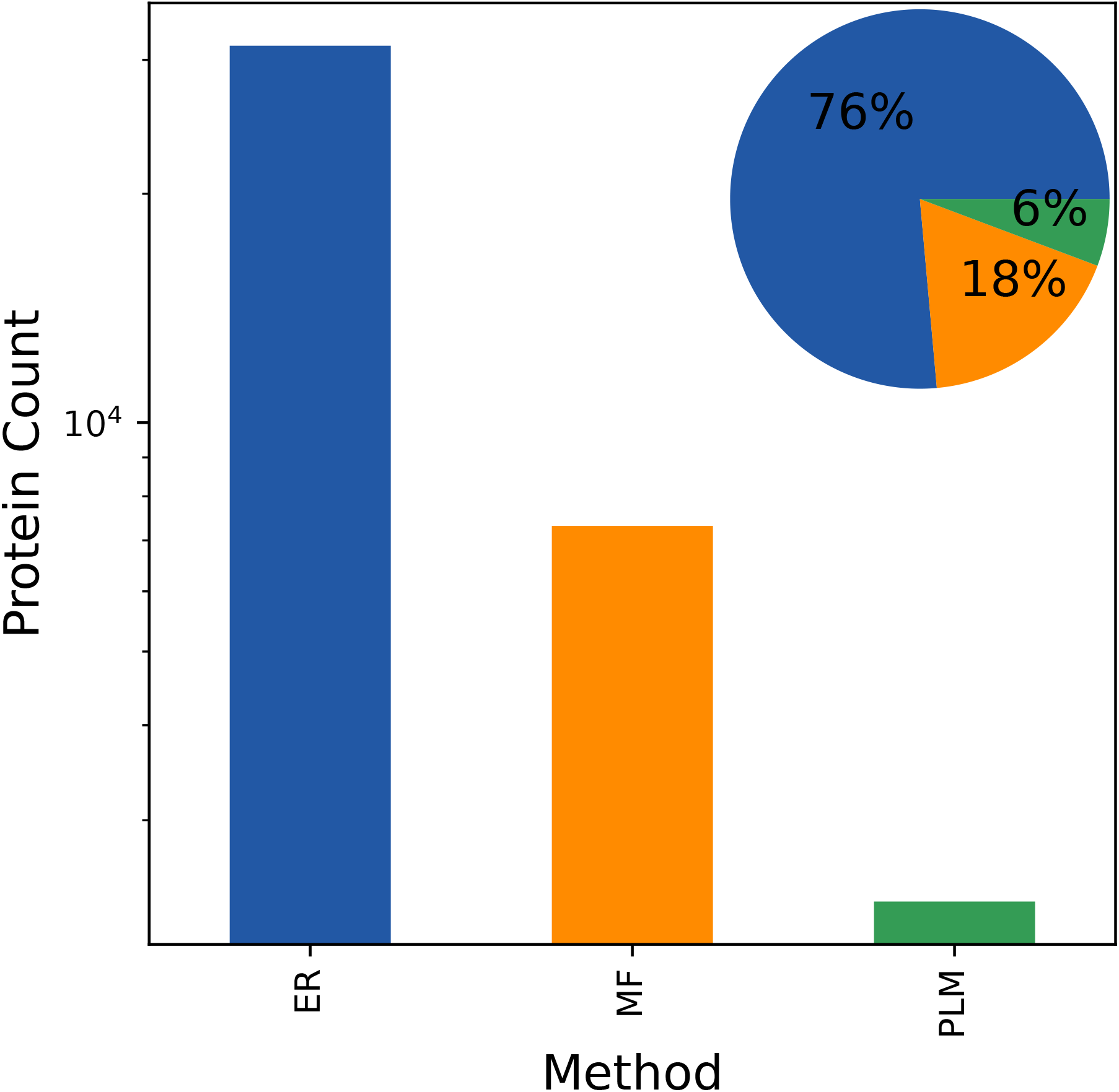
Counts of Best Method by ROC-AUC. We compare the three methods across all 40974 proteins by ROC-AUC values with ER providing the best contact prediction for 76% of tested proteins.

For further analysis in Figures 10 and 11 we stratify the method comparison across a given protein’s effective number of homologous sequences and the sequence length respectively. Figures 10 and 11 show that ER outperforms MF and PLM methods in contact predictions over all ranges of both the size of protein MSA and the length of protein sequences. From Figures 10 and 11 it is evident that ER outperforms existing methods across the full gamut of protein family sequences in sequence number and in sequence length.

**Figure 10:**
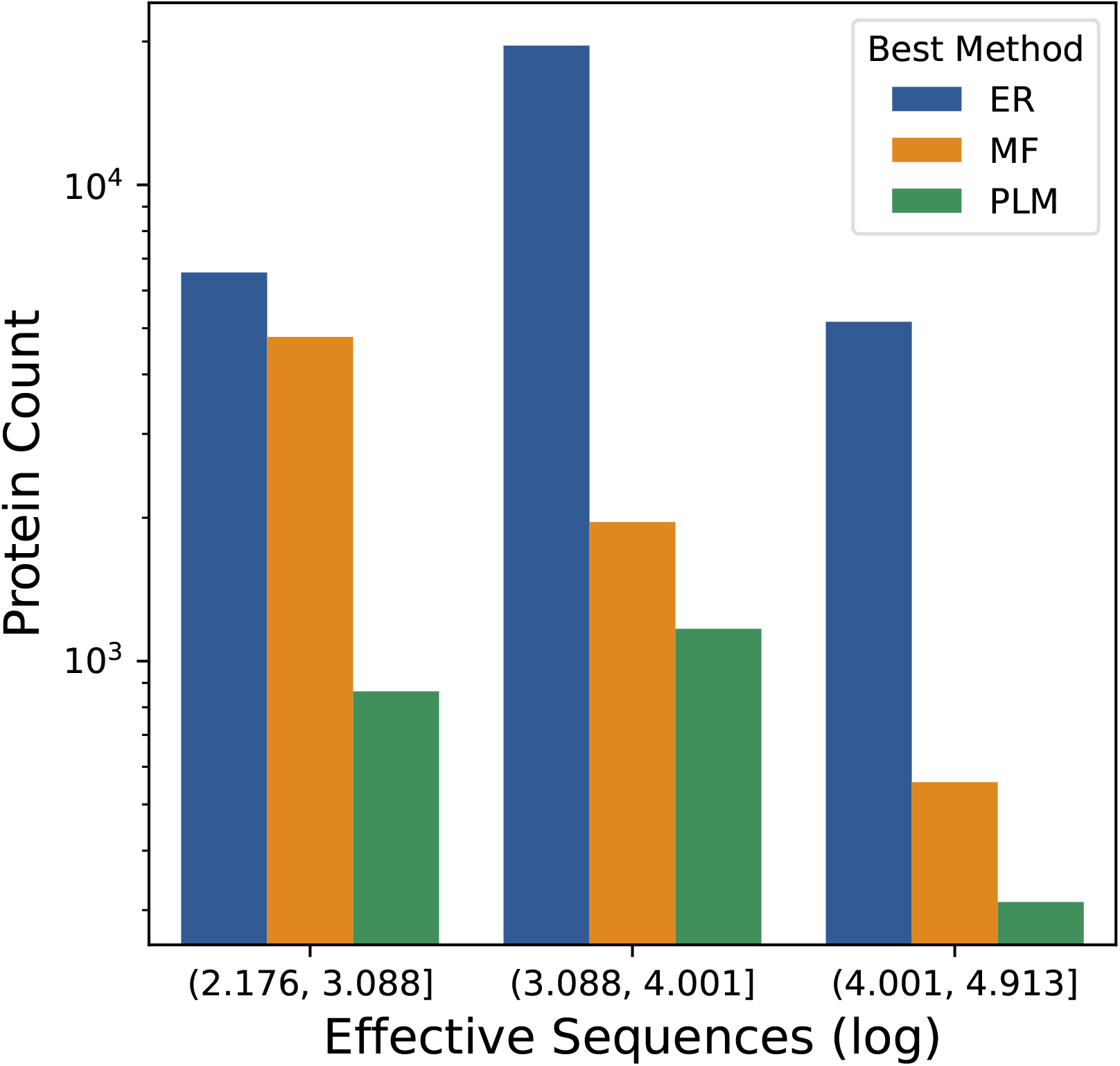
Best Method Stratified by Number of Sequences. We compare the three DCA variants across all 40974 proteins by ROC-AUC values. We then group proteins by the number of effective sequences in the MSA and compare all three methods for the different groups.

**Figure 11:**
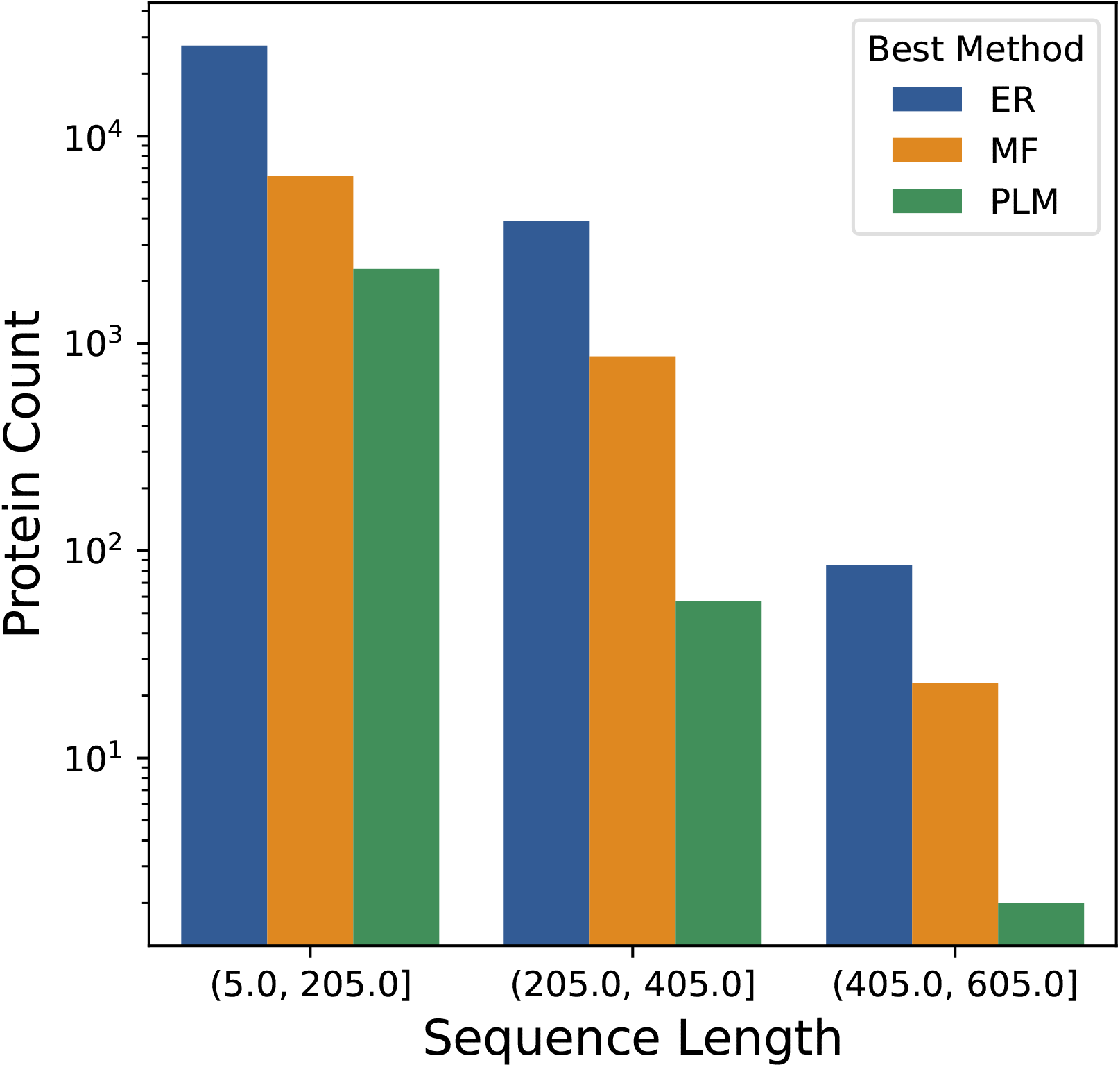
Best Method Stratified by Sequence Length. We compare the three DCA variants across all 40974 proteins by ROC-AUC values. We then group proteins by the length of the protein sequence (ie number of columns) and compare all three methods for the different groups.

## 3 Discussion

In this work we have presented a novel variant of the DCA family of methods for tertiary protein structure prediction. We formulated a Potts model coupling calculation in terms of a novel multiplicative update for the model’s interaction coefficients. As there is no closed form solution to such logistic regression problems, all methods for finding coefficients are iterative. Our method (ER) has been shown in previous work to be successful in under-determined inference problems with a large state space [Hoang et al., 2019a, Hoang et al., 2019b]. As we demonstrated, our method provides marked improvements on existing techniques for DCA model inference.

Beginning with the analysis and visualization of an individual protein (1ZDR) we gained some intuition on the extent to which direct information calculated using the Expectation Reflection variant of Direct Coupling Analysis (DCA-ER) can provide insight into protein tertiary structure. Importantly, we compared our method against two popular existing methods and found our method to improve on these methods for a large majority of analyzed proteins. It is important to note differences in methodology aside from the method used to compute the couplings of the statistical energy model. For one, we applied more inclusive data processing compared to that of existing methodology. In order to understand the impact of this approach to data processing, we implemented a version of MF with our own data processing and compared against the traditional data processing approach in Section 4.3.1 and saw little to distinguish the predicted results. In addition, any differences should not yield increased efficacy as our data processing was less stringent than previous approaches.

### 3.1 Future Directions

Future extensions of this analysis provide several avenues of investigation. As discussed previously, tertiary structure prediction is important input into existing state of the art machine learning protein prediction. Therefore, due to DCA-ER’s clear improvement on existing methods, it would be of interest to incorporate it into that analysis pipeline of existing neural network predictors. Of course, some existing neural network predictors, such as AlphaFold, incorporate co-evolution internally so it may not be useful to provide a redundant co-evolution contact pair prediction. A more interesting avenue to explore is to disentangle structural interactions from functional interactions, by comparing accurate 3D native structure predictions with those obtained from neural network predictors on sequences randomly evolved from native sequences by application of point mutation matrices such as the PAM or BLOSUM families.

## 4 Materials and Methods

### 4.1 Direct Coupling Analysis

The principle that structural and functional information can be extracted from evolutionary data is well supported [Taylor & Hatrick, 1994, Göbel et al., 1994, Neher, 1994]. With this motivation, a series of methods were developed to exploit evolutionary data using maximum entropy-based inference of such relationships, both in gene networks [Lezon et al., 2006, Locasale & Wolf-Yadlin, 2009], and protein contact prediction [Marks et al., 2011, Morcos et al., 2011, Jones et al., 2012, Ekeberg et al., 2014]. We will outline this methodology here. A thorough review of biological data inference is provided in [Stein et al., 2015]. We begin with the assumption that protein residues which are close in space within the folded protein will mutate in a correlated manner. Transitive correlations can be the result of statistical noise, phylogenetic sampling bias or residue interactions which are not a result of protein folding [Dunn et al., 2008, Burger & Van Nimwegen, 2010]. In this section we describe the methodology for inferring protein contact relationships from evolutionary data in a way the minimizes the effect of such secondary correlations.

#### 4.1.1 Amino Acid Sequences as categorical variables

We begin with a characterization of the data used to infer connections and then transition to a description of the general DCA model as well as the Expectation Reflection variant. Consider a set of *N* observed sequence configurations 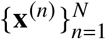 of homologous protein sequences where **x**^(*n*)^ is the *n*^th^ sequence. This amino acid sequence can be considered as a jointly distributed categorical variable **x** = (*x*_1_, · · ·, *x*_*L*_)^*T*^ where each variable *x*_*i*_ is in the finite set Ω = {A, C, D, E, F, G, H, I, K, L, M, N, P, Q, R, S, T, V, W, Y, -}. Therefore a given sequence of length *L* is defined as **x** = (*x*_1_, · · ·, *x*_*L*_)^*T*^ ∈ Ω^*L*^.

##### Binary Embedding

In order to utilize a formulation of pairwise maximum-entropy distribution we use one-hot encoding. **x** is cast in a binary embedding which specifies any *L*-vector as a binary *Lq*-vector σ, where *q* = 21 is the number of elements in our finite set of amino acid states Ω. This expresses a given residue *x*_*i*_ into *q*-subvector binary representation,

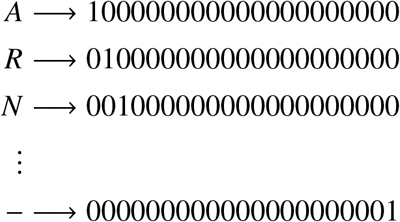

which allows us to convert the protein sequence set into a binary sequence set. Therefore the **x** becomes a binary sequence σ(**x**) = OneHot(**x**), with the well-known constraint that the last of the one-hot sequence values is determined if we know all the rest.

#### 4.1.2 DCA Methodology

Because of the potential influence of transitive correlations, it is important to consider the influence of all protein residues on the observed relationship between a given pair of co-evolving positions. This relationship can be characterized by a global statistical measure of evolutionary information, direct information (DI). The direct information between two positions *i* and *j* (as in Figure 1) in **x** can be described as

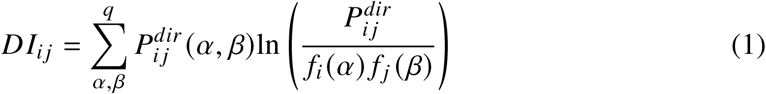

where 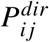 is a pairwise probability model and *α, β* ∈ Ω represent one of 21 possible residue states at positions *i* and *j* respectively. This is a marginalization of some general probability that captures lower order interactions between positions as measured in existing observations while also predicting future frequencies. Therefore we want the simplest probabilistic approach constrained to recreate characteristics of the observed data such that

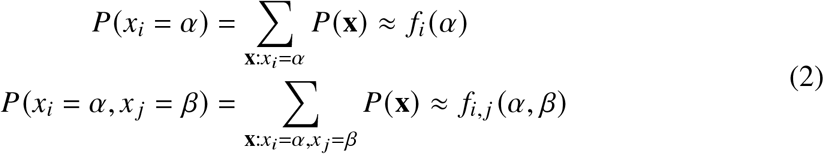

The derivation of such a probability has been characterized as a 21-state Potts model, with the initial investigation utilizing an Ising model analogy for protein sequences by Lapedes et al. [Lapedes et al., 2002, Stein et al., 2015].

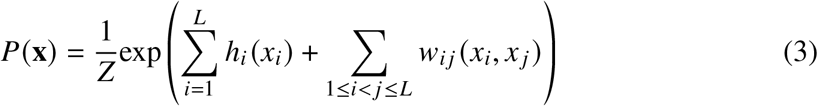

Where 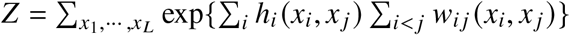. This distribution gives a maximum-entropy inference as presented in previous work [Weigt et al., 2009, Marks et al., 2011, Morcos et al., 2011, Ekeberg et al., 2014]. In order to infer the model parameters {**h, w**}, we proceed to maximize likelihood, by minimizing negative log-likelihood *l*,

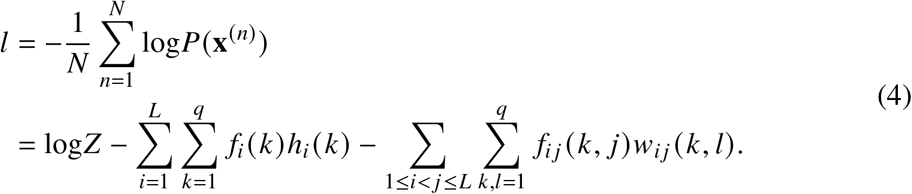

Because *l* is differentiable, we can minimize by finding a parameter set such that 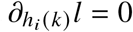 and 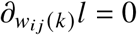, which is our goal for model inference,

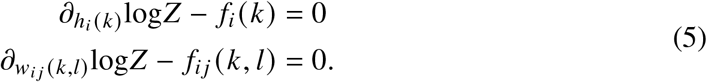

To find this minimization computationally we need to calculate the partition function *Z* for a given set of the parameters {**h, w**} and this is computationally intractable. Therefore, approaches to evaluate these parameters have been developed by many groups with small refinements on how the couplings of the model are actually approximated [Weigt et al., 2009, Marks et al., 2011, Morcos et al., 2011, Ekeberg et al., 2014]. The inference of the parameters is also where we introduce our novel variant, by using Expectation Reflection to infer the model couplings for DCA (DCA-ER).

#### 4.1.3 Potts model inference with Expectation Reflection

We will infer our model couplings **w** from the multiple sequence alignment of homologous sequences. Given a set of observed sequence configurations 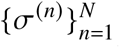 where a protein sequence state configured as a binary variable σ^(^*n*) representing the *n*^*th*^ sequence such that *n* ∈ [1, *N* = *N*_sequences_] represents all possible sequences, or states, protein-family poly-peptide can take. An individual sequence has the form σ = (σ_1_, σ_2_, · · ·, σ_*L*_*q*) where *Lq* is the number of binary variables as described in Section 4.1.1. We can then assume that given a current sequence σ^(*n*)^, an individual position in the sequence is determined stochastically according to the following conditional probability,

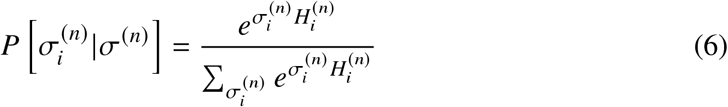

where the local field, 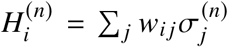 encodes the effect of all other positions on the residue at position *i*, and *w*_*i j*_ represents the interaction between positions *i* and *j*. 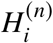 is a function of the rest of the current sequence state 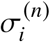 and expresses the influence of all other sequence positions on the state of position *i*. This conditional relationship allows us to iteratively search for the optimal **w** using all *N* available sequences. The method, and resulting algorithm, is discussed in further detail in previous work [Hoang et al., 2019a, Hoang et al., 2019b].

Once we have used the entire MSA to infer our position interactions **w** we can go from the coupling coefficient matrix to a measure of interaction strength between sequence position pairs, described in Equation 1.

### 4.2 Kolmogorov-Smirnov Test for Comparing Contact Predictions

Given a pair of CDF’s, the standard two-sample KS test is used to verify whether the predictions are drawn from the same underlying distribution. We use the two-sample Kolmogorov-Smirnov (KS) test to determine whether the predictions of two given methods are significantly different. The TP rate of each method’s predictions serves as a proxy for a cumulative distribution function for our problem. Using this rate, we can plot all KS values where the higher values are a stronger indicator of rejecting the null hypothesis (i.e., that the predictions are too similar).

In the case of comparing protein structure predictions, given a sample size of *N* contact predictions, we calculate the critical value for our KS-statistic as follows:

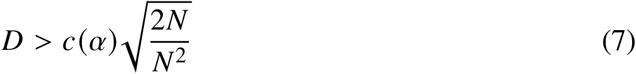

where *c*(*α*) is the inverse of the KS distribution at *α*. Therefore, different levels of confidence provide a condition for corresponding predictions. We utilize a confidence of *α* = .001 which gives *c*(*α*) = 1.949. Therefore given a protein with N amino acids the critical KS threshold becomes:

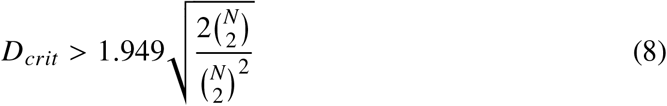

### 4.3 Data acquisition and processing

To test our method over the widest possible variety of protein structures we created a data acquisition pipeline which uses a given PDB-ID (e.g. 1ZDR) and finds the best corresponding homologous protein sequence set from the Pfam database [Mistry et al., 2020] using the Biopython [Cock et al., 2009] and ProDy [Bakan et al., 2011] packages for sequence pairing and database searches.

#### Algorithm 1

PDB → MSA Database Search

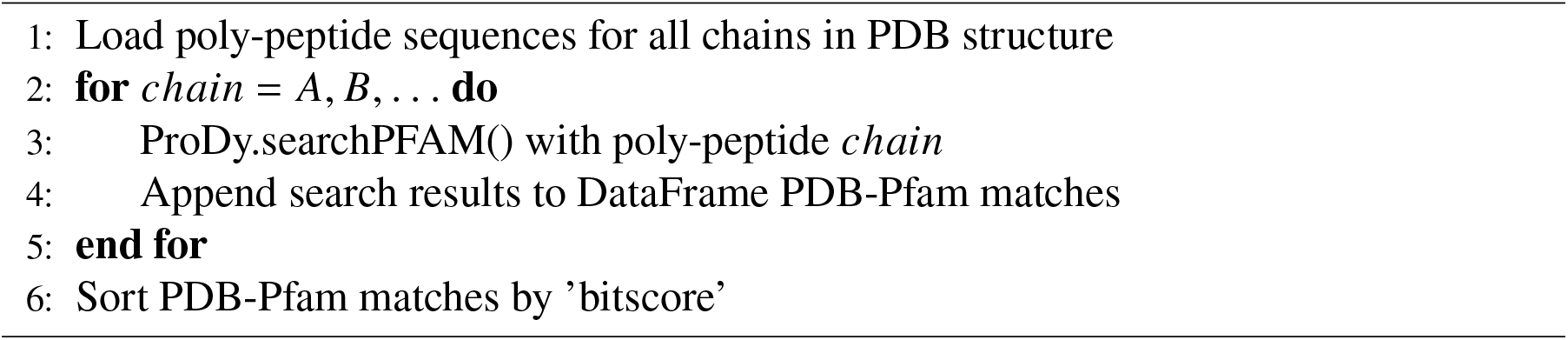

#### Algorithm 2

Data Processing

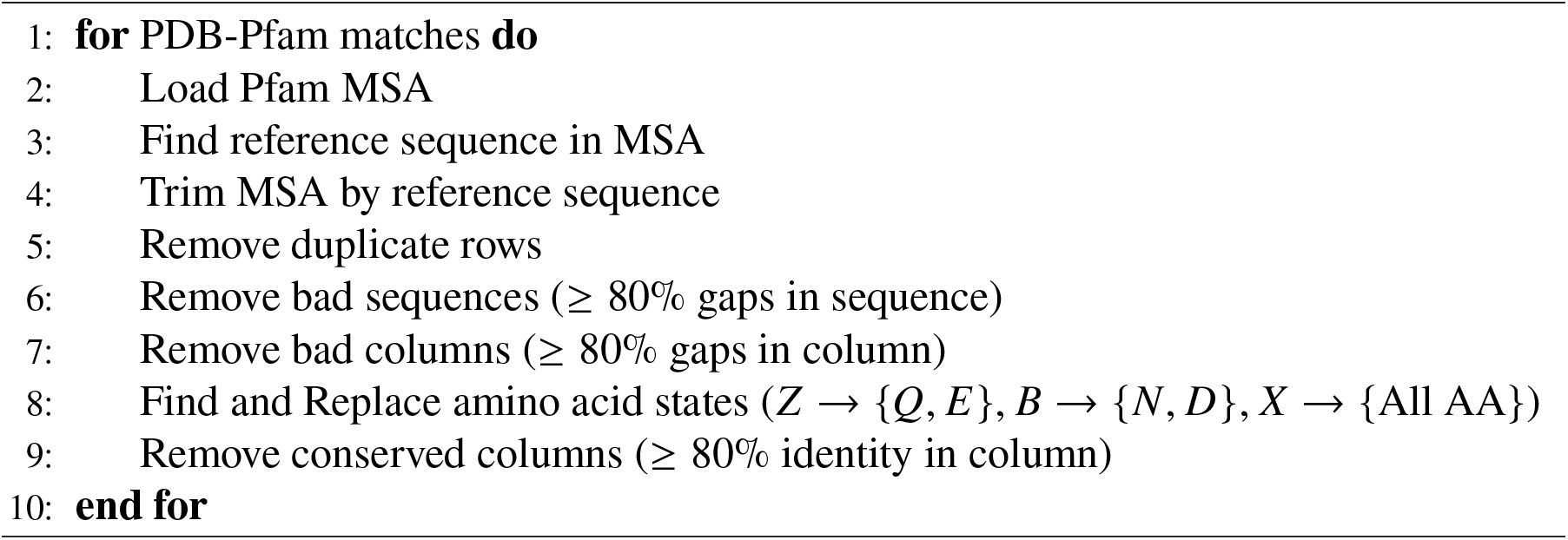

#### 4.3.1 Differences in Data Processing

Notably, this represents changes to previous data processing techniques. Namely, in the methods with which we compare our variant, all sequences were kept regardless of gap percentage in sequence. Additionally more stringent criteria were used for removing gap columns, with previous methods removing columns with ≥ 80% gaps. Finally, previous data processing neglected to substitute inconclusive amino acid representations (step 9 in Algorithm 2), opting to simply discard these sequences. This discrepancy yields a negligible difference in resulting analytics as we show in Figure 13, where we use the KS statistic to infer the effect of data processing. We consider the prediction difference between the MF and PLM variant as well as the prediction difference between the MF variant with two types of data processing. The distribution of KS statistics comparing MF and PLM with the same approach to data processing is skewed much higher than the distribution comparing MF with the two different approaches to data processing. This implies that the effect of data processing on final predictions is minimal relative to the differences yielded by different DCA variants.

**Figure 12:**
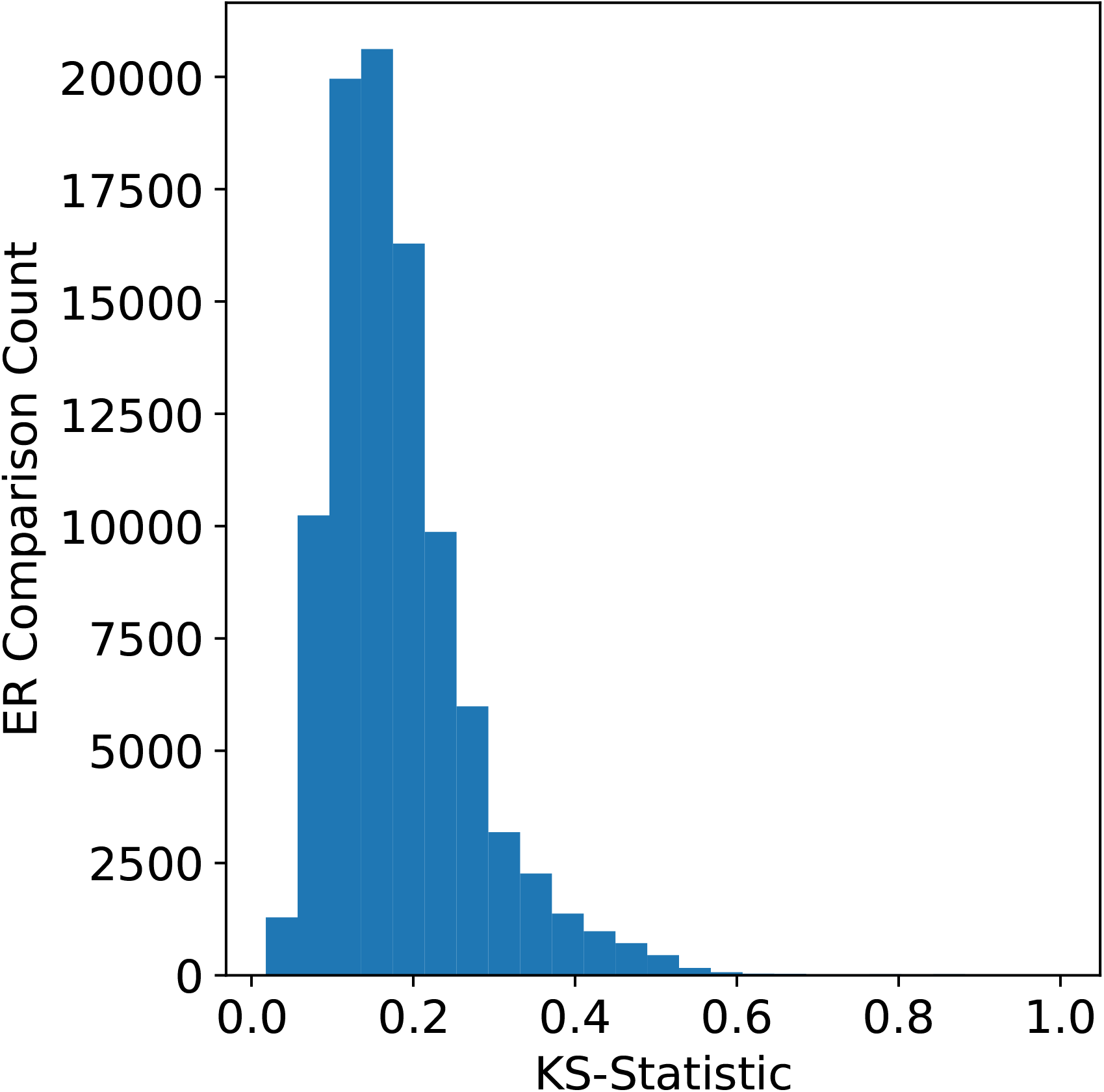
ER KS-Statistics. We plot The Kolmogorov-Smirnov statistic, determining whether the contact predictions from ER is significantly different from that of either MF or PLM, for all proteins.

**Figure 13:**
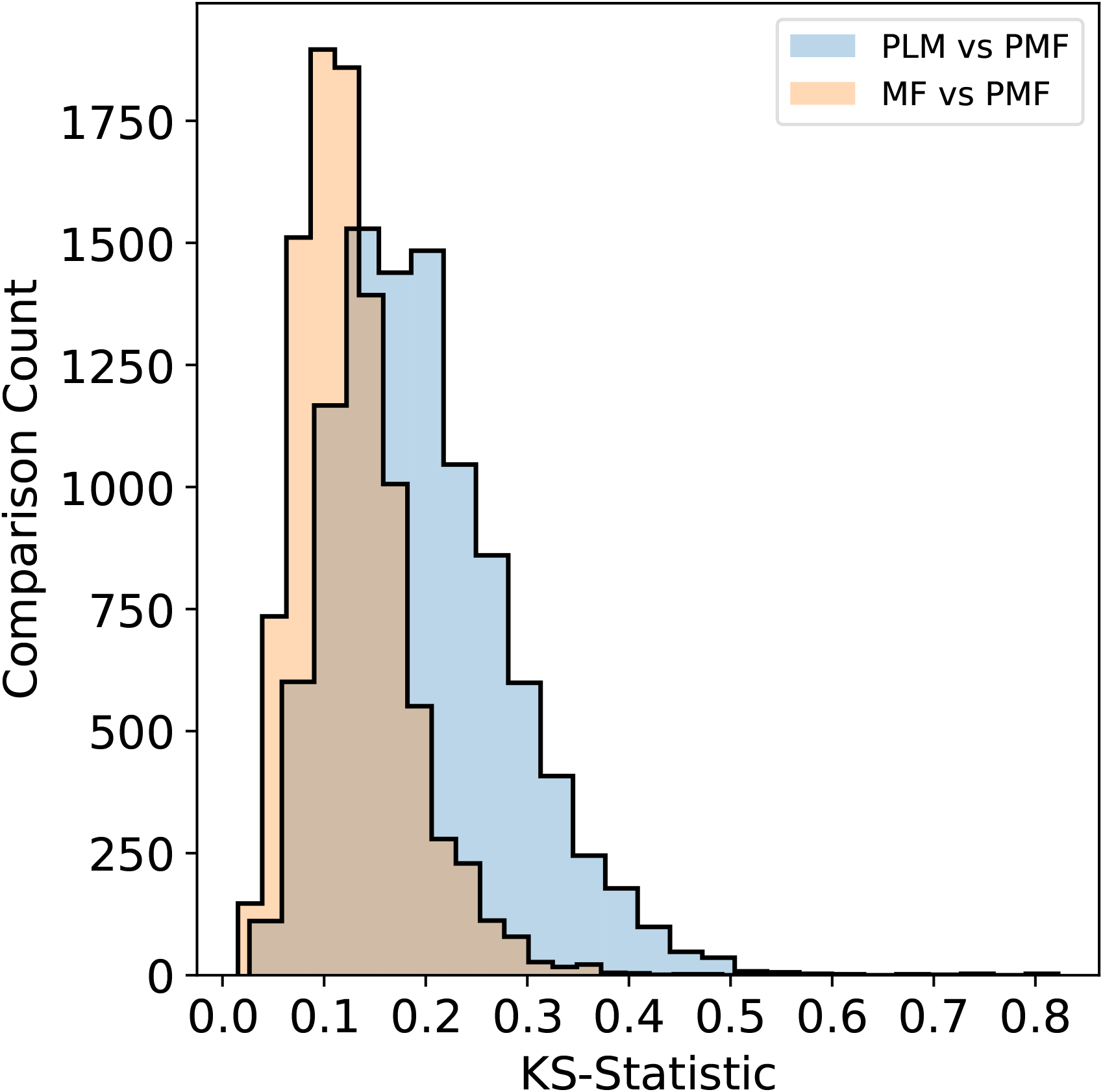
Data Processing Comparison. We plot The Kolmogorov-Smirnov statistic, to visualize the effect of data processing on contact predictions. We first compare the results of MF using both approaches to data processing (MF vs PMF). We then compare MF and PLM variants using the same approach to data processing (PLM vs PMF).

### 4.4 Code and Data

The processed data and results are available upon request. The code and instructions for the analysis of individual proteins is available on Github at https://github.com/nihcompmed/ER-PSP. We also provide the individual analysis plots (as in Section 2.1) of 265 proteins as supplementary data, with 100 selected from each grouping of MSA size in Figure 10.

## Supporting information

Supplemental Contact Maps

## 5 Acknowledgements

The authors would like to thank the HPC staff, Wolfgang Resch and Gennady Denisov for computational support using Biowulf. This work utilized the computational resources of the NIH HPC Biowulf cluster (http://hpc.nih.gov). This work was supported by the Intramural Research Program of the National Institute of Diabetes and Digestive and Kidney Diseases.

## 6 Competing Interests

The authors declare that they have no conflict of interest.

## Notes

https://github.com/nihcompmed/ER-PSP

### Competing Interest Statement

The authors have declared no competing interest.

