## Supplemental Contact Maps for "Protein Structure Prediction with Expectation Reflection": 3seq_225col_5any_PF01589_plots.pdf

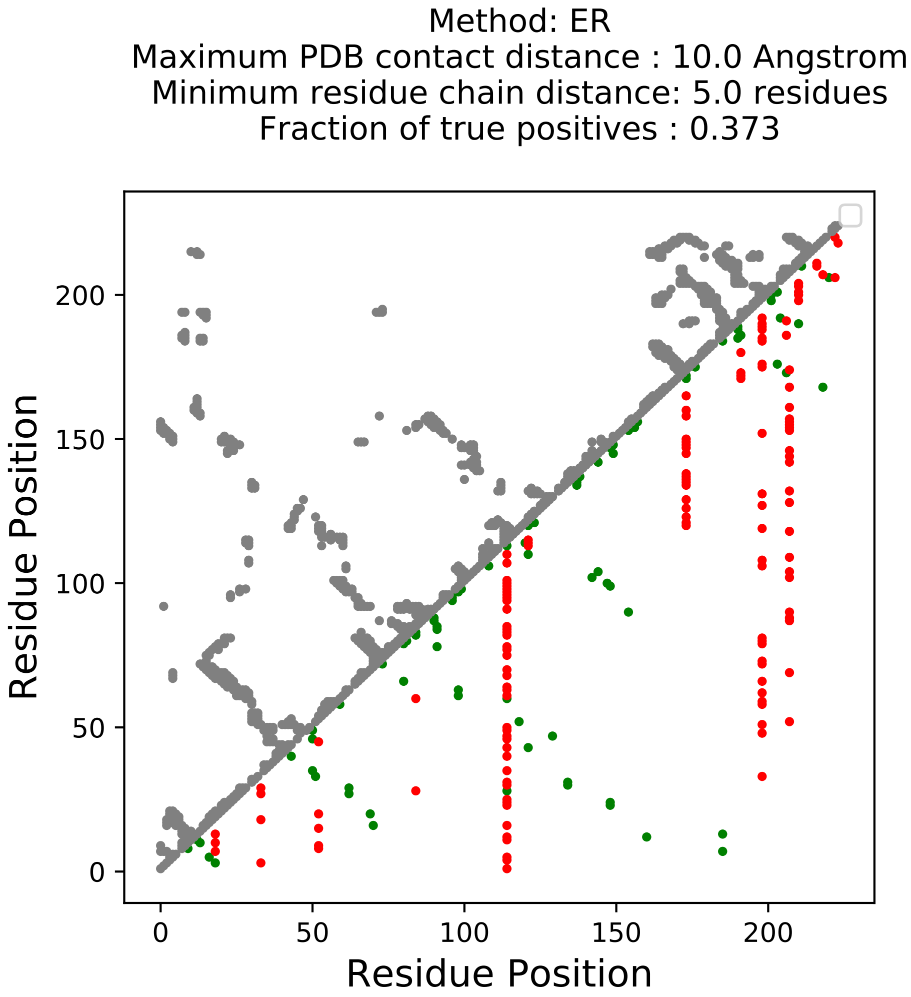

Method: ER
Maximum PDB contact distance : 10.0 Angstrom
Minimum residue chain distance: 5.0 residues
Fraction of true positives : 0.373
Residue Position
Residue Position
0
50
100
150
200
0
50
100
150
200

Method: MF
Maximum PDB contact distance : 10.0 Angstrom
Minimum residue chain distance: 5.0 residues
Fraction of true positives : 0.4

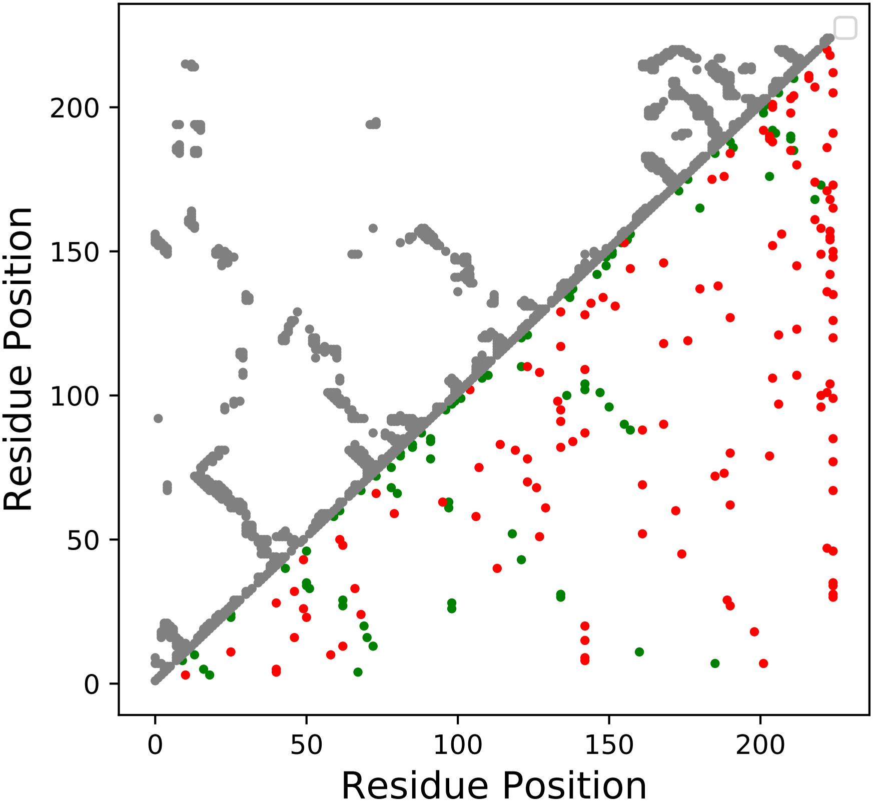

Method: PLM
Maximum PDB contact distance : 10.0 Angstrom
Minimum residue chain distance: 5.0 residues
Fraction of true positives : 0.347

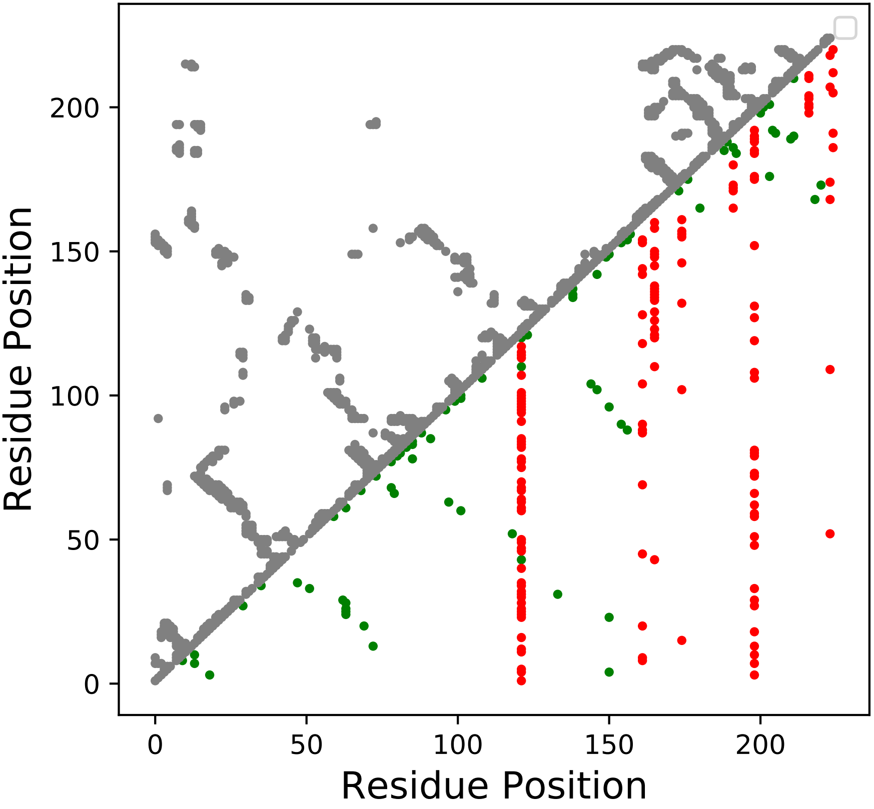

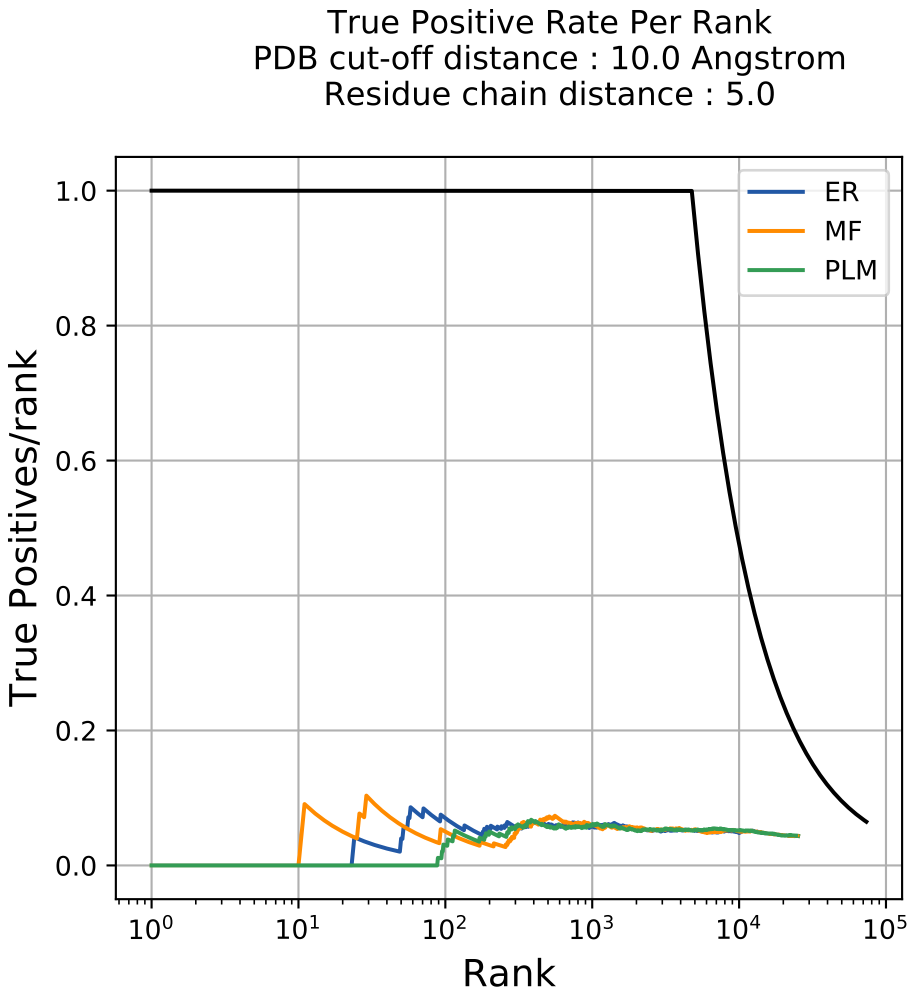

True Positive Rate Per Rank
PDB cut-off distance : 10.0 Angstrom
Residue chain distance : 5.0
ER
MF
PLM
True Positives/rank
1.0
0.8
0.6
0.4
0.2
0.0
Rank
$10^0$
$10^1$
$10^2$
$10^3$
$10^4$
$10^5$

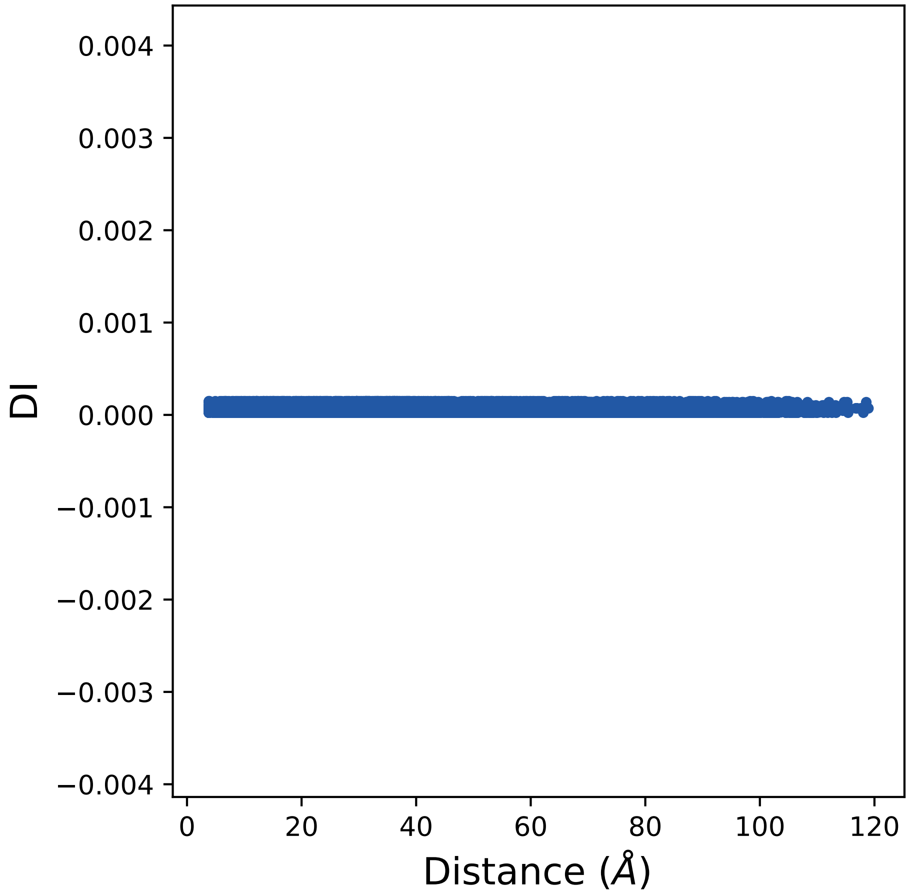

0.004
0.003
0.002
0.001
0.000
−0.001
−0.002
−0.003
−0.004
DI
0
20
40
60
80
100
120
Distance (Å)

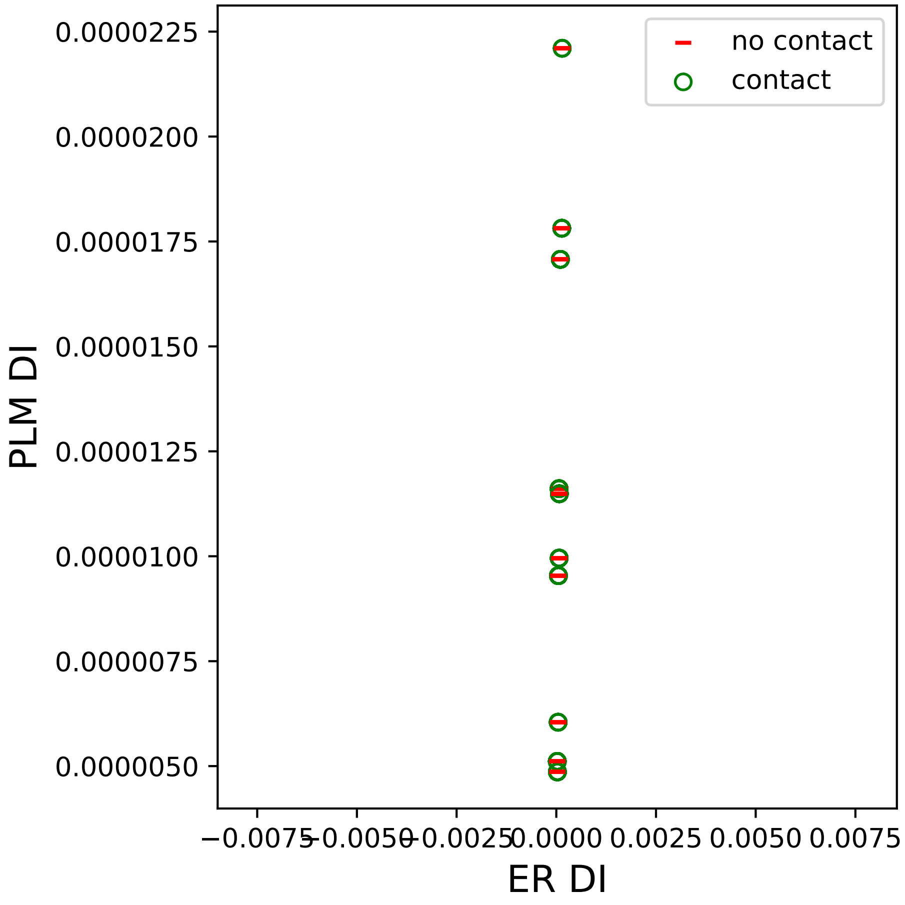

0.0000225
0.0000200
0.0000175
0.0000150
0.0000125
0.0000100
0.0000075
0.0000050
PLM DI
−0.0075
−0.0050
−0.0025
0.0000
0.0025
0.0050
0.0075
ER DI
no contact
contact

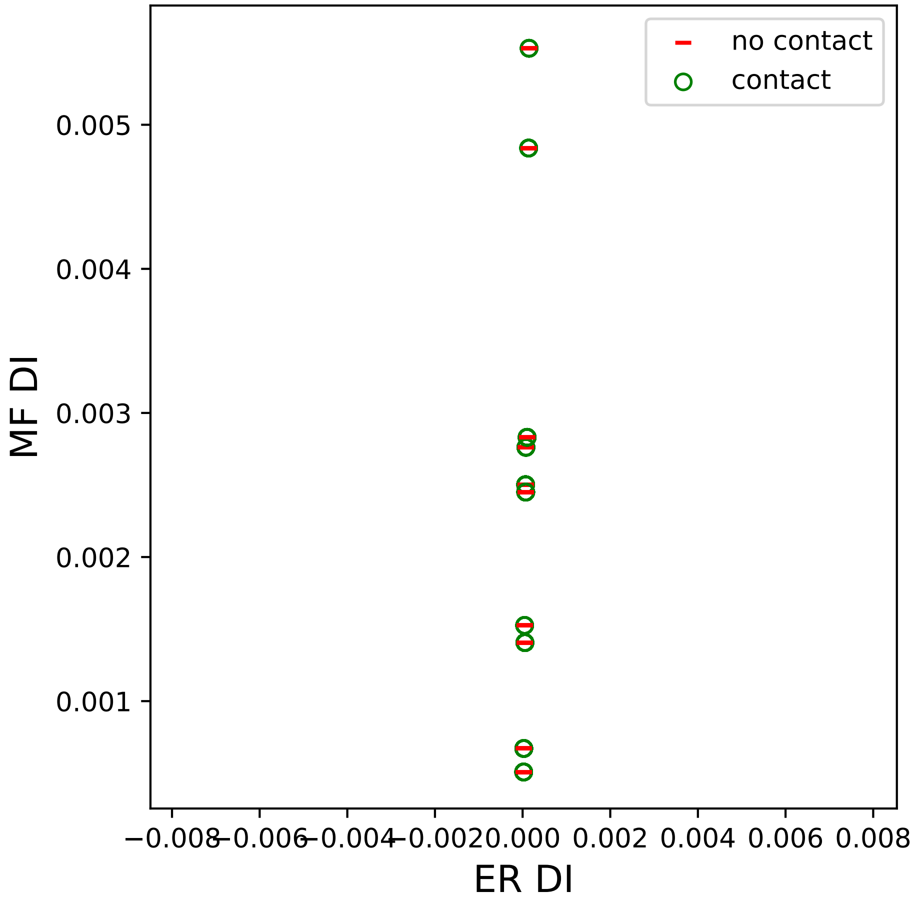

no contact
contact
0.005
0.004
0.003
0.002
0.001
MF DI
−0.008
−0.006
−0.004
−0.002
0.000
0.002
0.004
0.006
0.008
ER DI

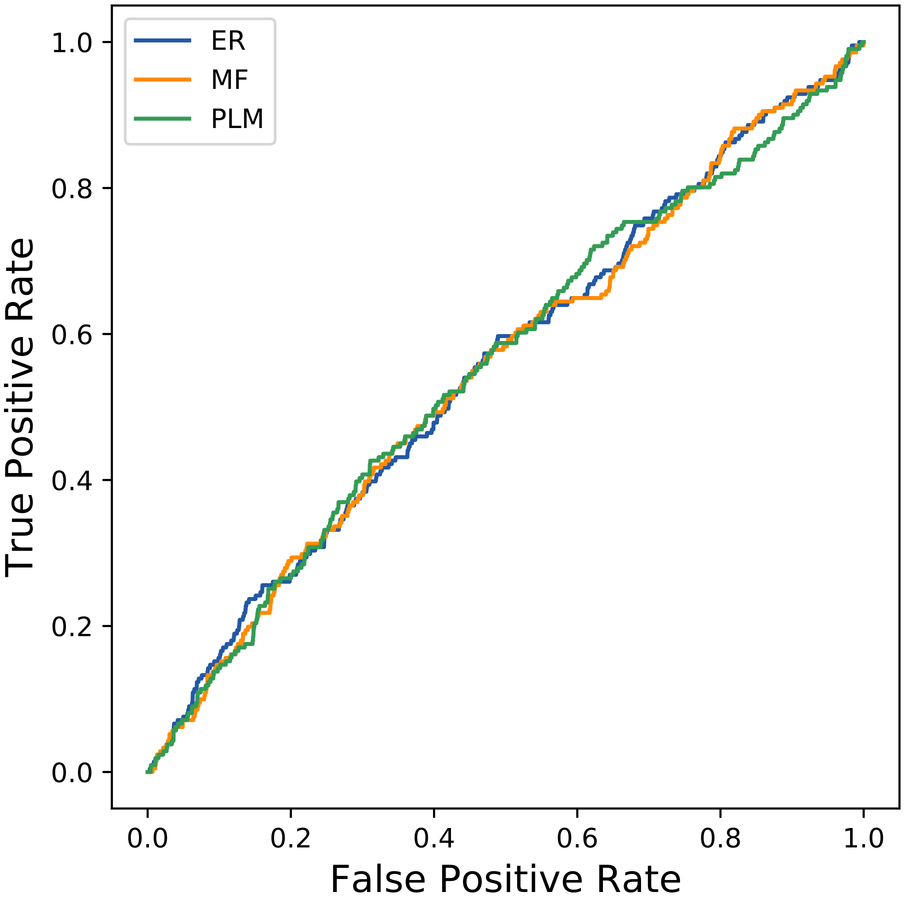

ER
MF
PLM
True Positive Rate
False Positive Rate
0.0
0.2
0.4
0.6
0.8
1.0
