## Supplemental Contact Maps for "Protein Structure Prediction with Expectation Reflection": 18seq_23col_5by1_PF00600_plots.pdf

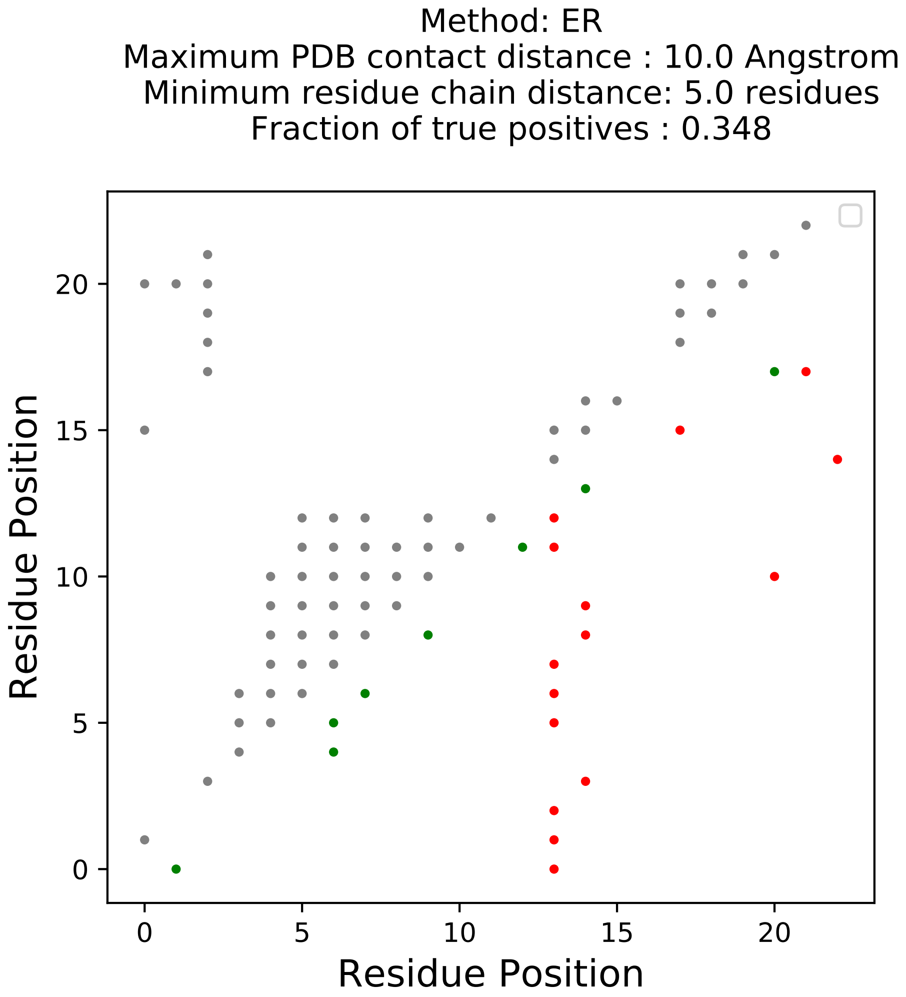

Method: ER
Maximum PDB contact distance : 10.0 Angstrom
Minimum residue chain distance: 5.0 residues
Fraction of true positives : 0.348
Residue Position
Residue Position
0
5
10
15
20

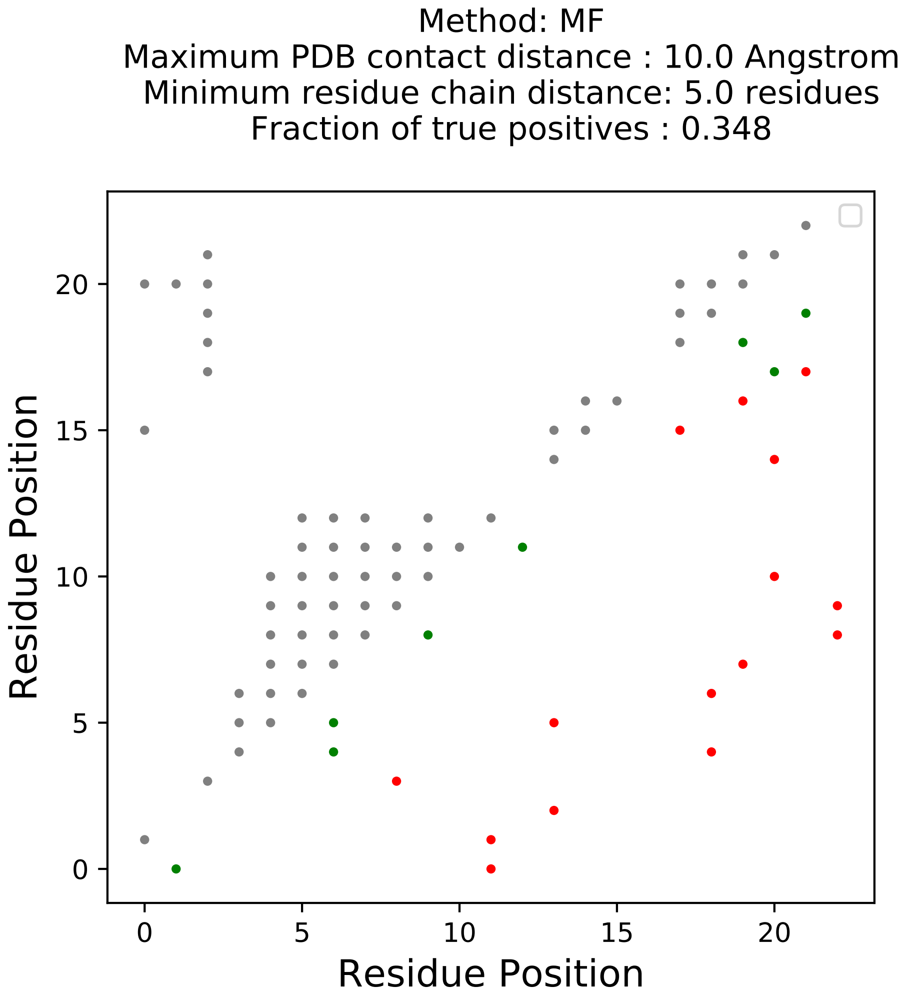
Method: MF
Maximum PDB contact distance : 10.0 Angstrom
Minimum residue chain distance: 5.0 residues
Fraction of true positives : 0.348
Residue Position
Residue Position
0
5
10
15
20

Method: PLM
Maximum PDB contact distance : 10.0 Angstrom
Minimum residue chain distance: 5.0 residues
Fraction of true positives : 0.435

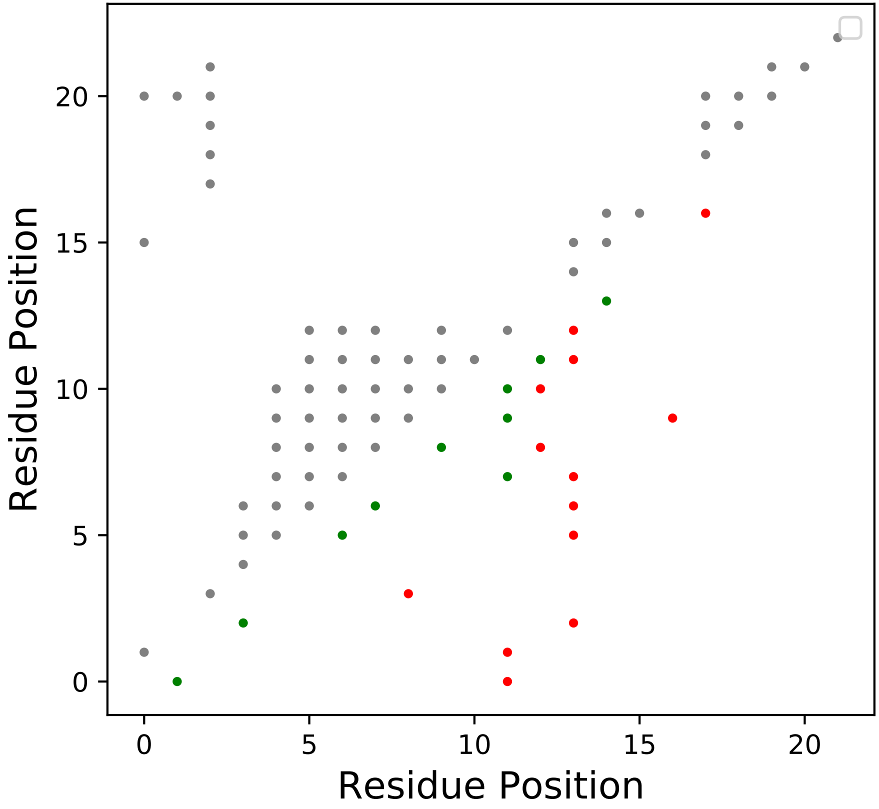

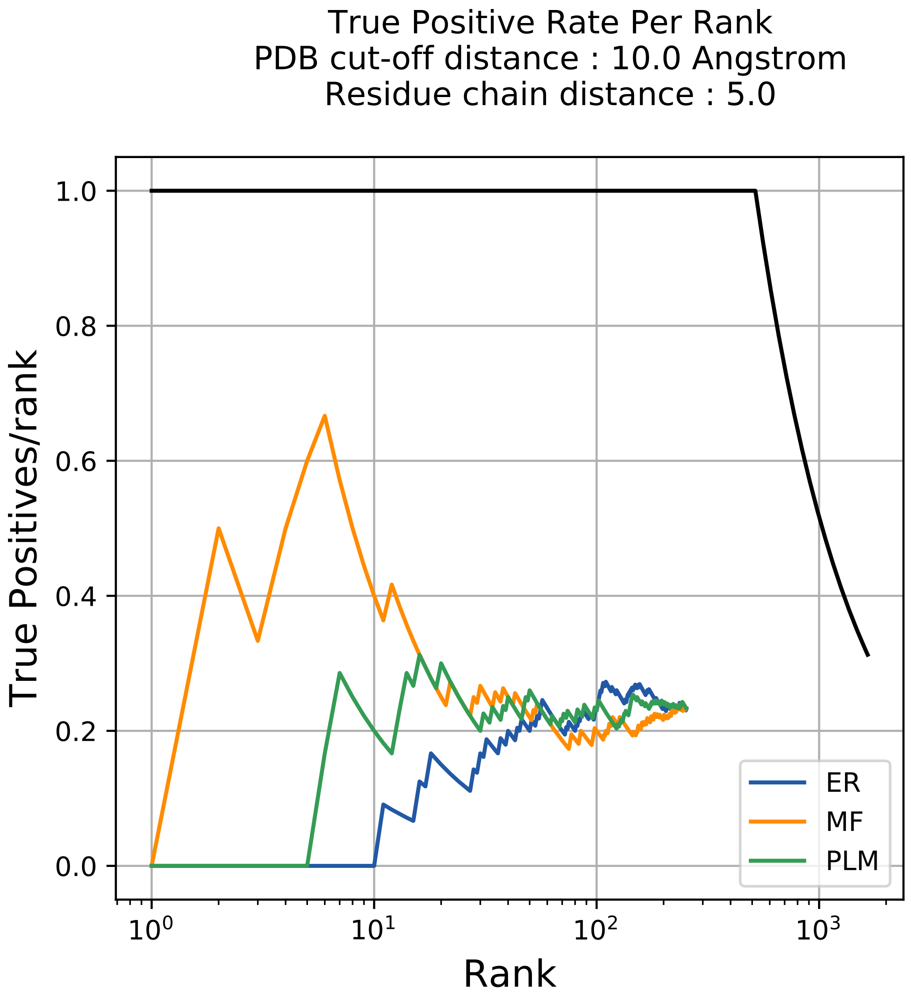
True Positive Rate Per Rank
PDB cut-off distance : 10.0 Angstrom
Residue chain distance : 5.0
True Positives/rank
Rank
1.0
0.8
0.6
0.4
0.2
0.0
$10^0$
$10^1$
$10^2$
$10^3$
ER
MF
PLM

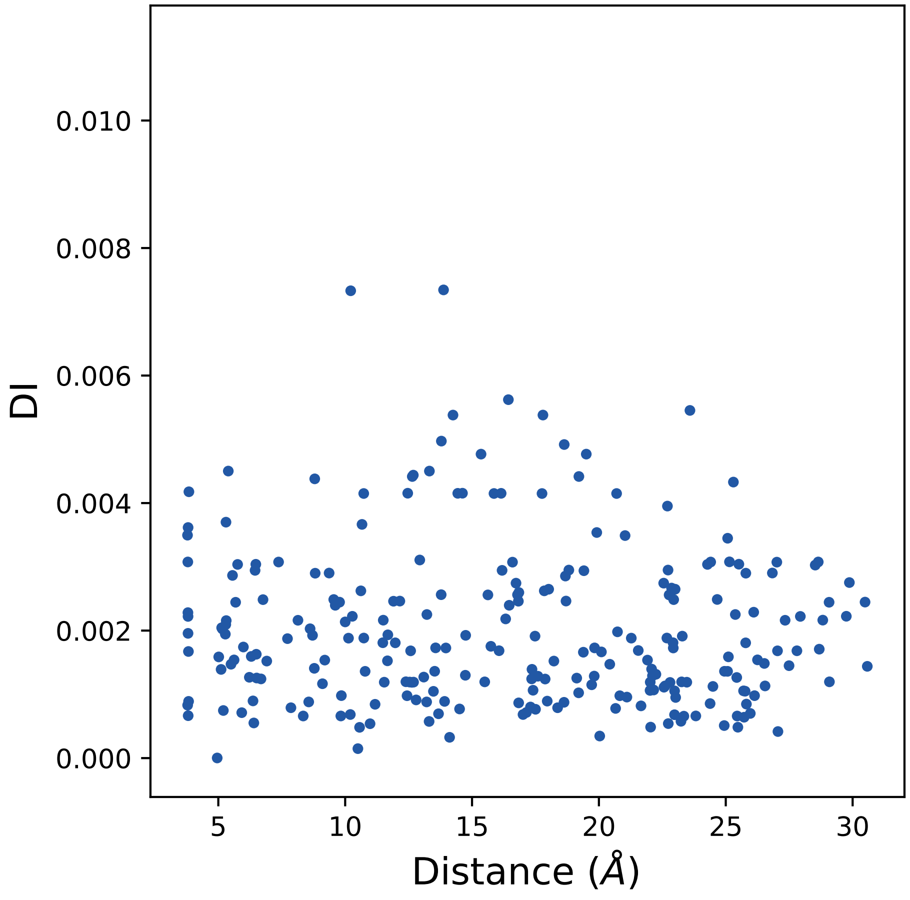

DI
0.000
0.002
0.004
0.006
0.008
0.010
5
10
15
20
25
30
Distance (Å)

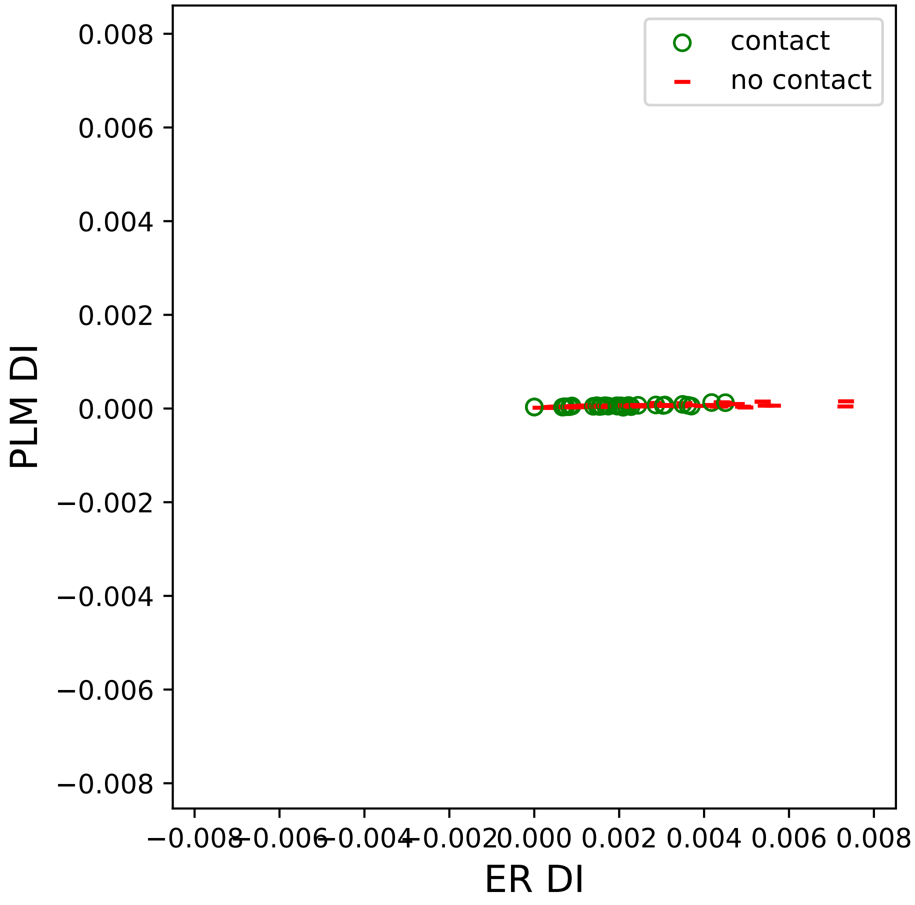

contact
no contact
PLM DI
ER DI
0.008
0.006
0.004
0.002
0.000
−0.002
−0.004
−0.006
−0.008
−0.008
−0.006
−0.004
−0.002
0.000
0.002
0.004
0.006
0.008

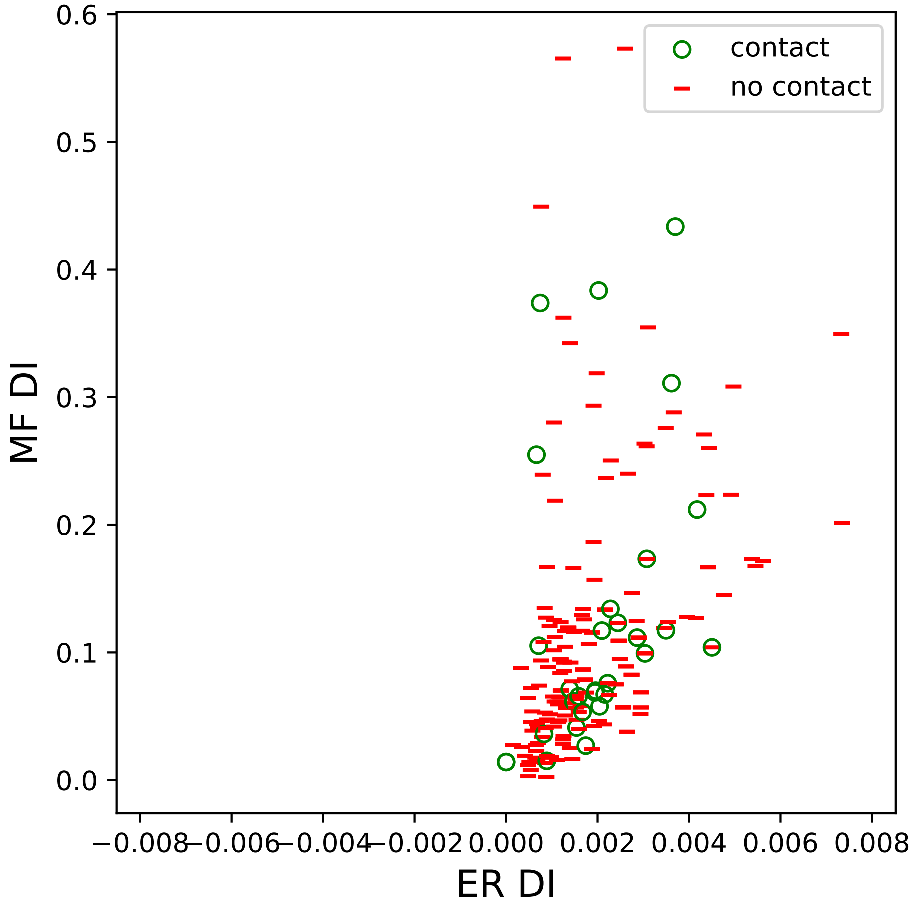

contact
no contact
0.6
0.5
0.4
0.3
0.2
0.1
0.0
MF DI
−0.008
−0.006
−0.004
−0.002
0.000
0.002
0.004
0.006
0.008
ER DI

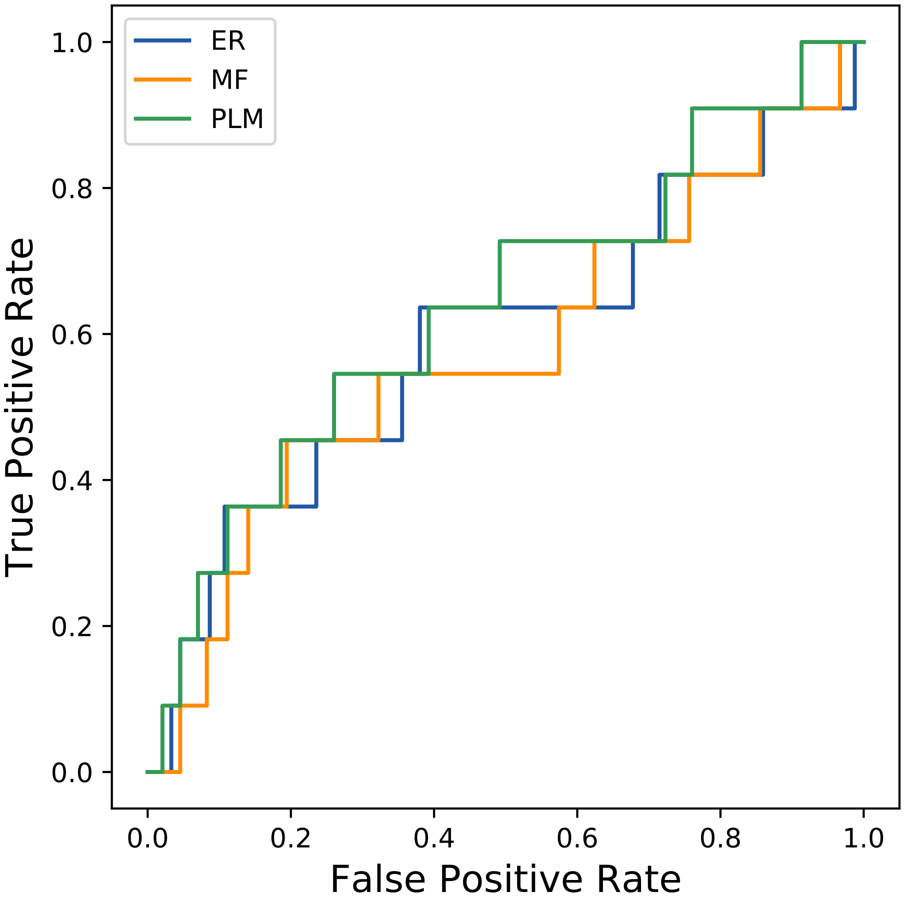

ER
MF
PLM
1.0
0.8
0.6
0.4
0.2
0.0
True Positive Rate
0.0
0.2
0.4
0.6
0.8
1.0
False Positive Rate
