## Supplemental Contact Maps for "Protein Structure Prediction with Expectation Reflection": 18seq_40col_6j1c_PF01353_plots.pdf

Method: ER
Maximum PDB contact distance : 10.0 Angstrom
Minimum residue chain distance: 5.0 residues
Fraction of true positives : 0.425

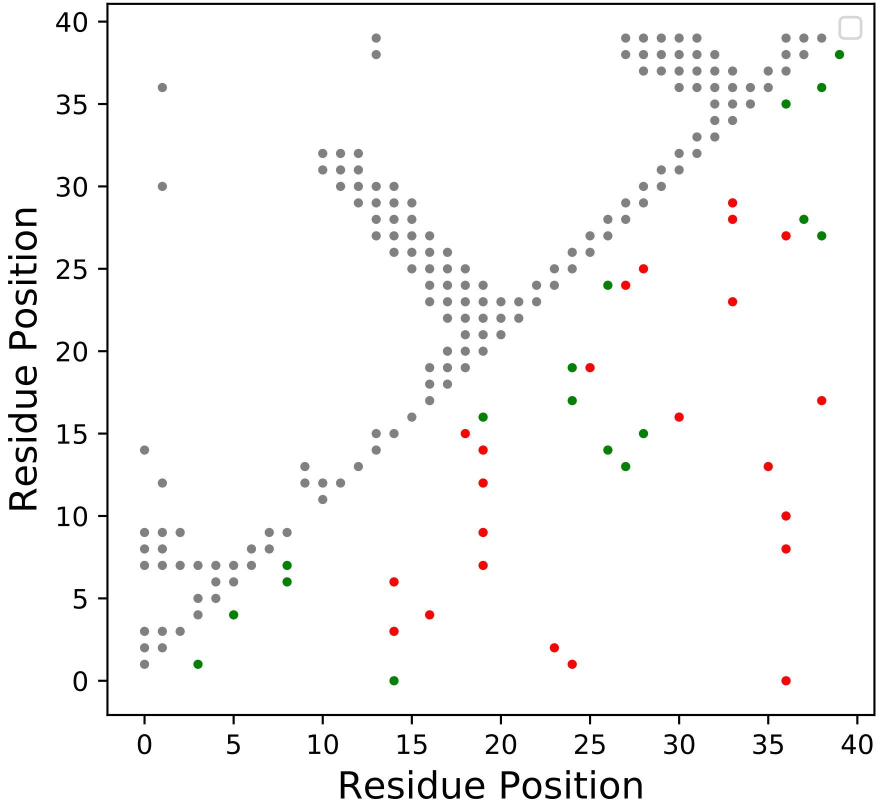

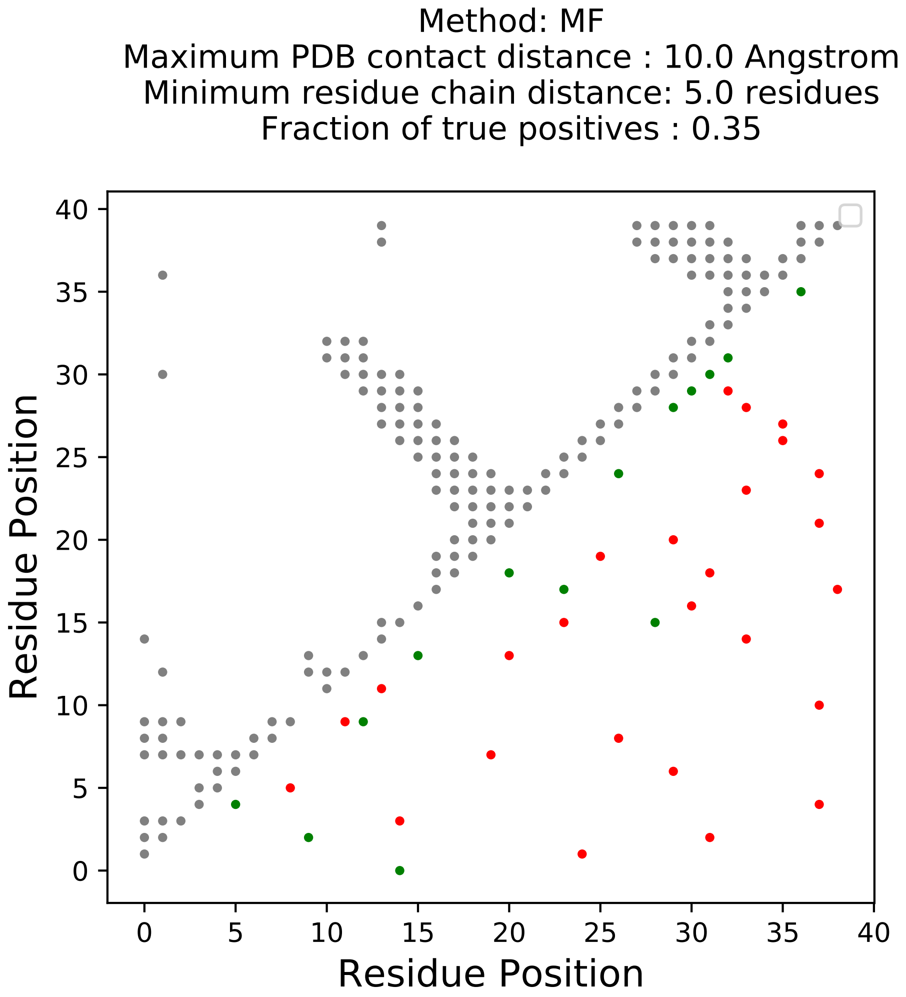
Method: MF
Maximum PDB contact distance : 10.0 Angstrom
Minimum residue chain distance: 5.0 residues
Fraction of true positives : 0.35
Residue Position
Residue Position
0
5
10
15
20
25
30
35
40

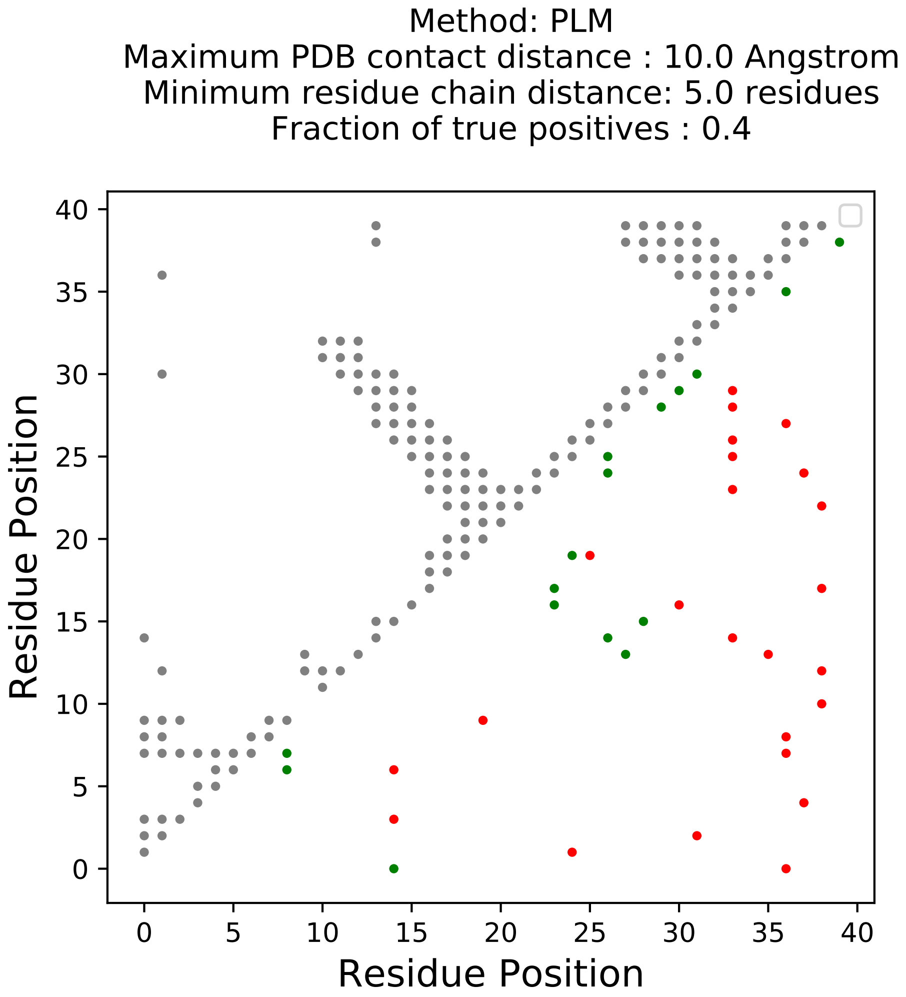

Method: PLM
Maximum PDB contact distance : 10.0 Angstrom
Minimum residue chain distance: 5.0 residues
Fraction of true positives : 0.4
Residue Position
Residue Position
0
5
10
15
20
25
30
35
40

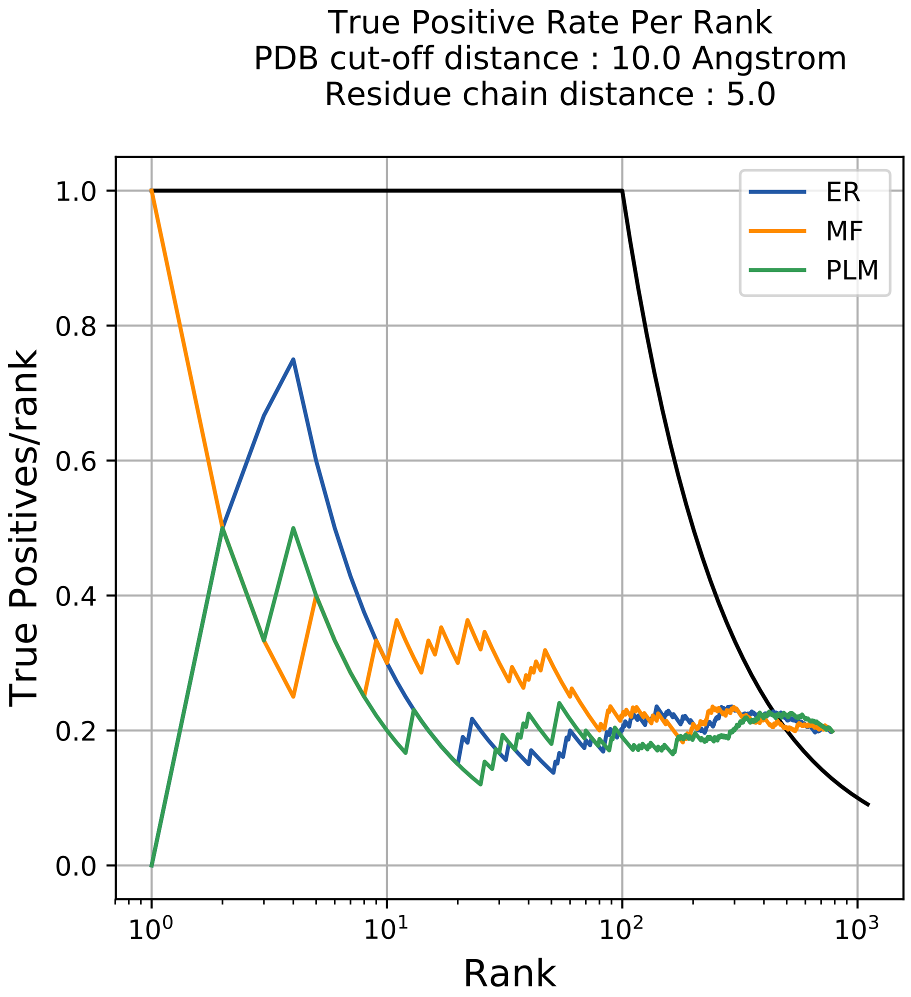

True Positive Rate Per Rank
PDB cut-off distance : 10.0 Angstrom
Residue chain distance : 5.0
ER
MF
PLM
True Positives/rank
1.0
0.8
0.6
0.4
0.2
0.0
$10^0$
$10^1$
$10^2$
$10^3$
Rank

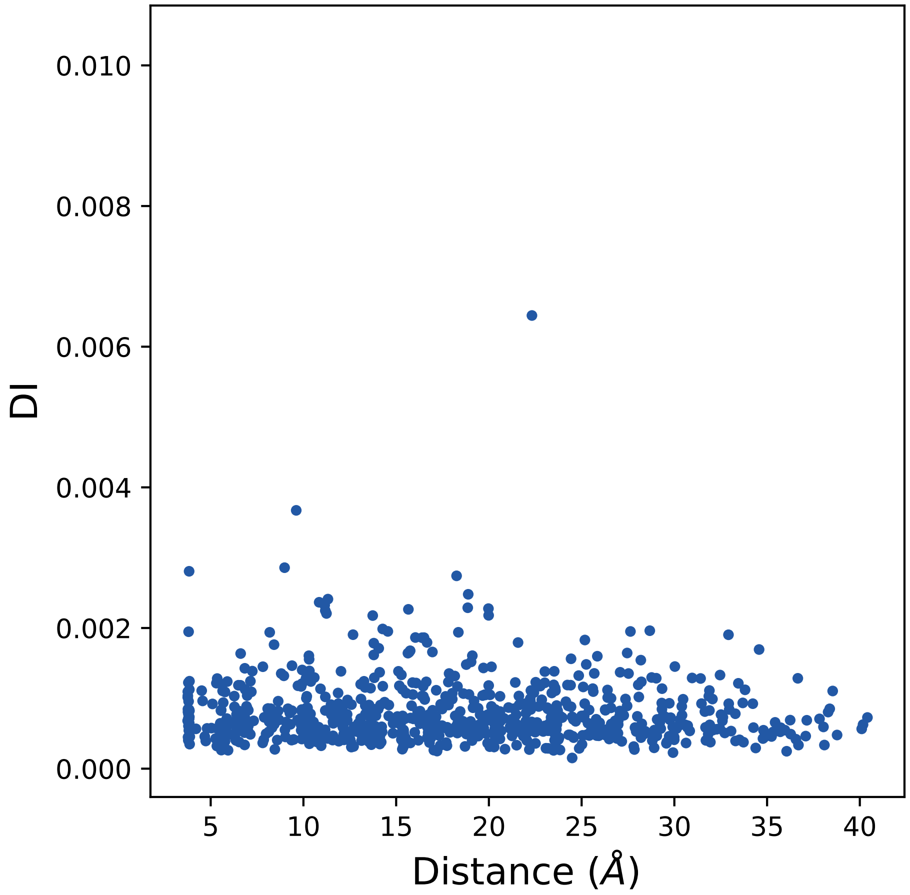
DI
0.000
0.002
0.004
0.006
0.008
0.010
5
10
15
20
25
30
35
40
Distance (Å)

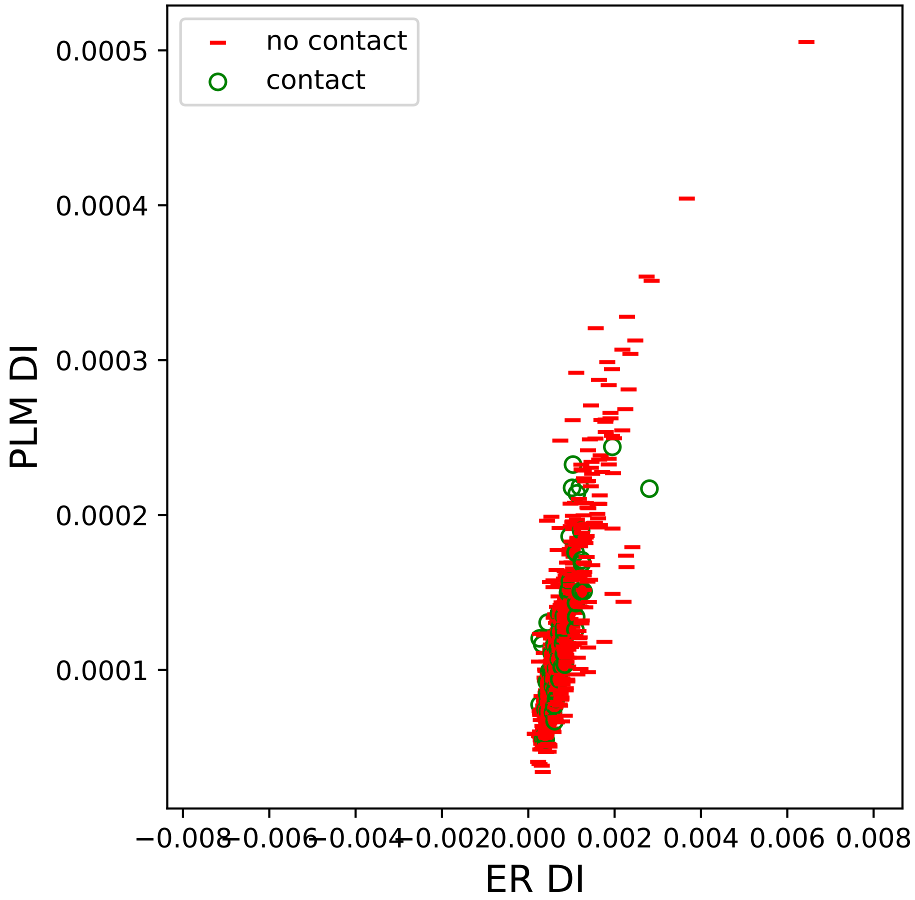

no contact
contact
0.0005
0.0004
0.0003
0.0002
0.0001
PLM DI
−0.008
−0.006
−0.004
−0.002
0.000
0.002
0.004
0.006
0.008
ER DI

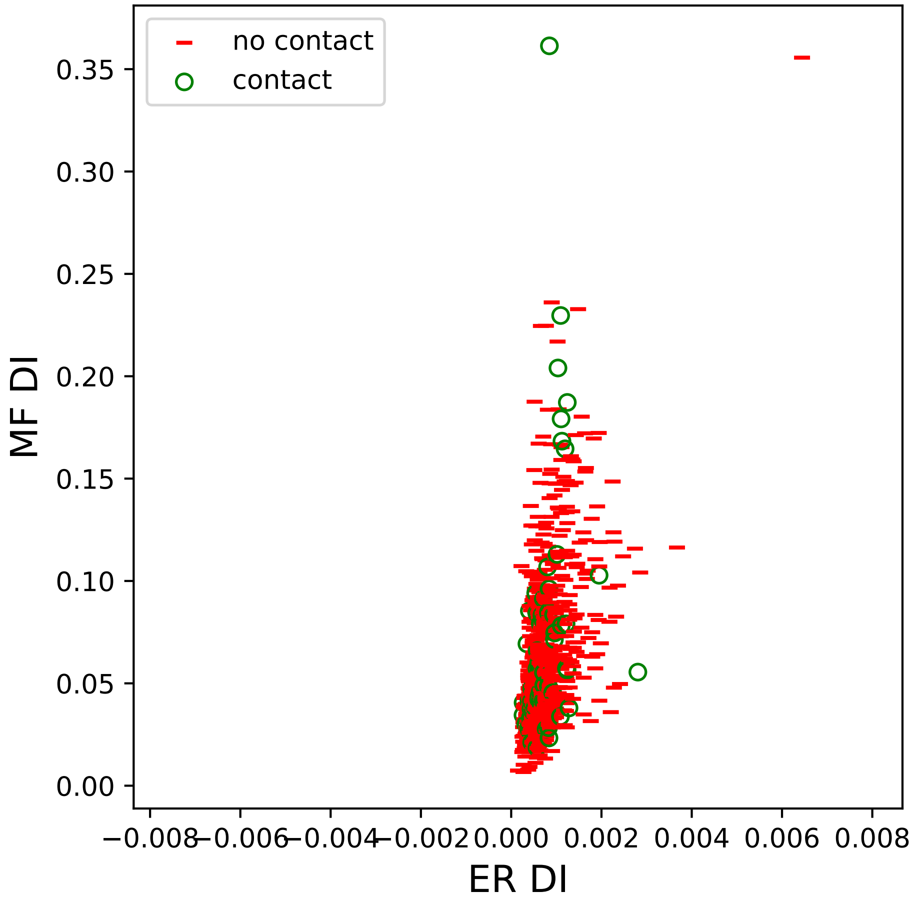

no contact
contact
0.35
0.30
0.25
0.20
0.15
0.10
0.05
0.00
MF DI
−0.008
−0.006
−0.004
−0.002
0.000
0.002
0.004
0.006
0.008
ER DI

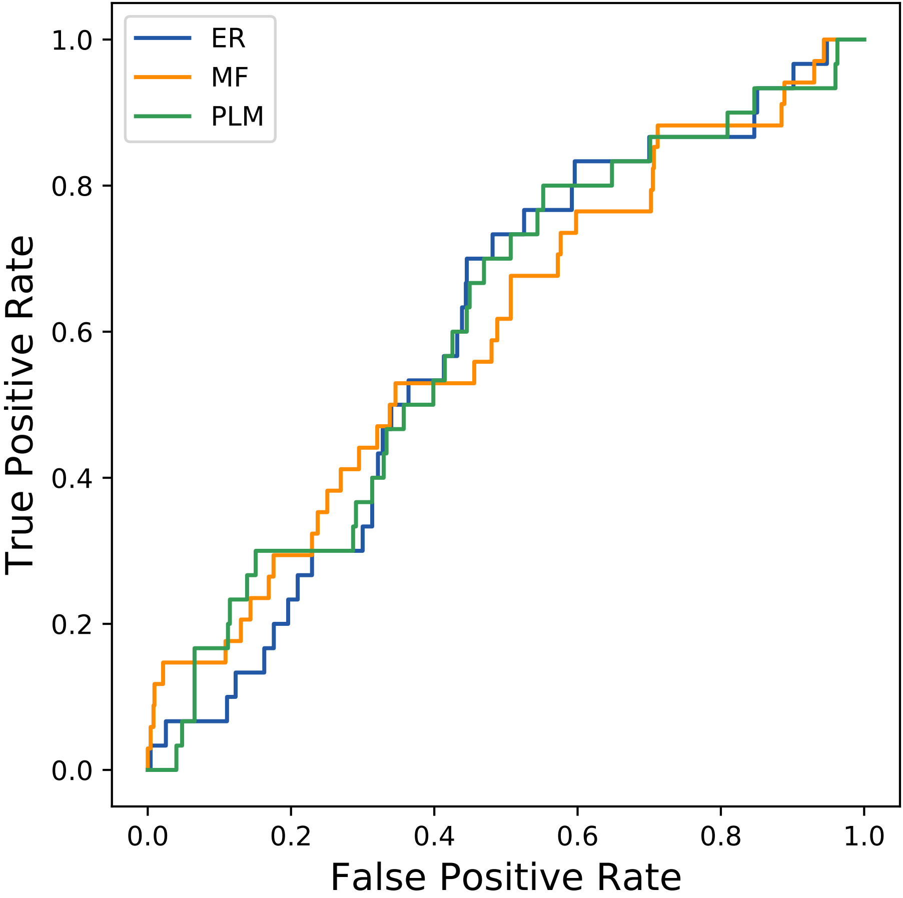

ER
MF
PLM
1.0
0.8
0.6
0.4
0.2
0.0
True Positive Rate
0.0
0.2
0.4
0.6
0.8
1.0
False Positive Rate
