## Supplemental Contact Maps for "Protein Structure Prediction with Expectation Reflection": 27seq_48col_4dvt_PF00516_plots.pdf

Method: ER
Maximum PDB contact distance : 10.0 Angstrom
Minimum residue chain distance: 5.0 residues
Fraction of true positives : 0.396

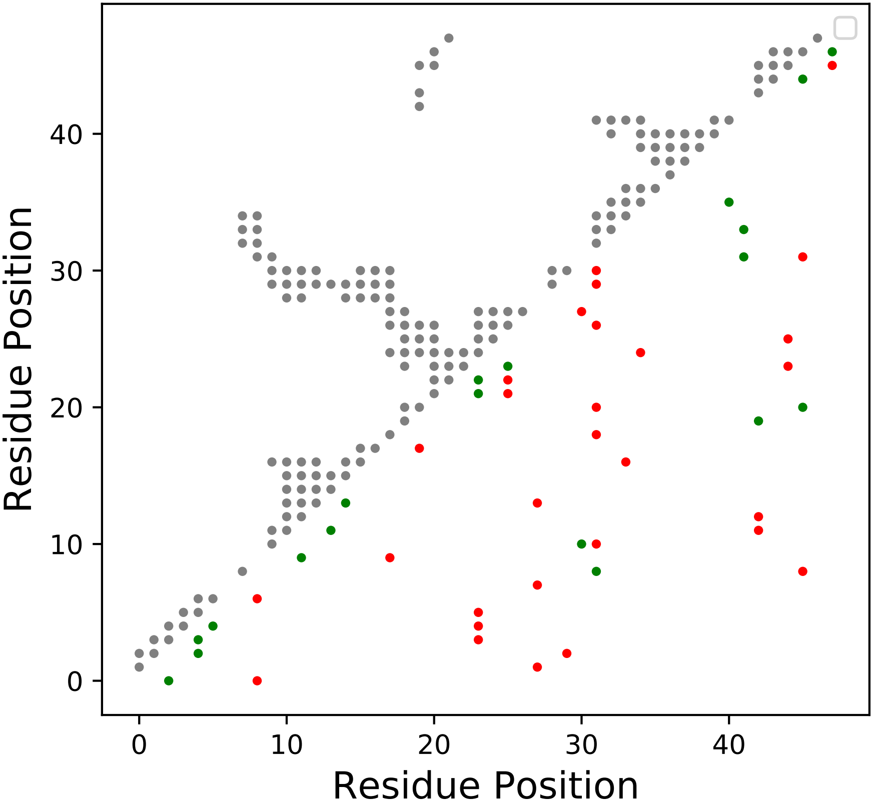

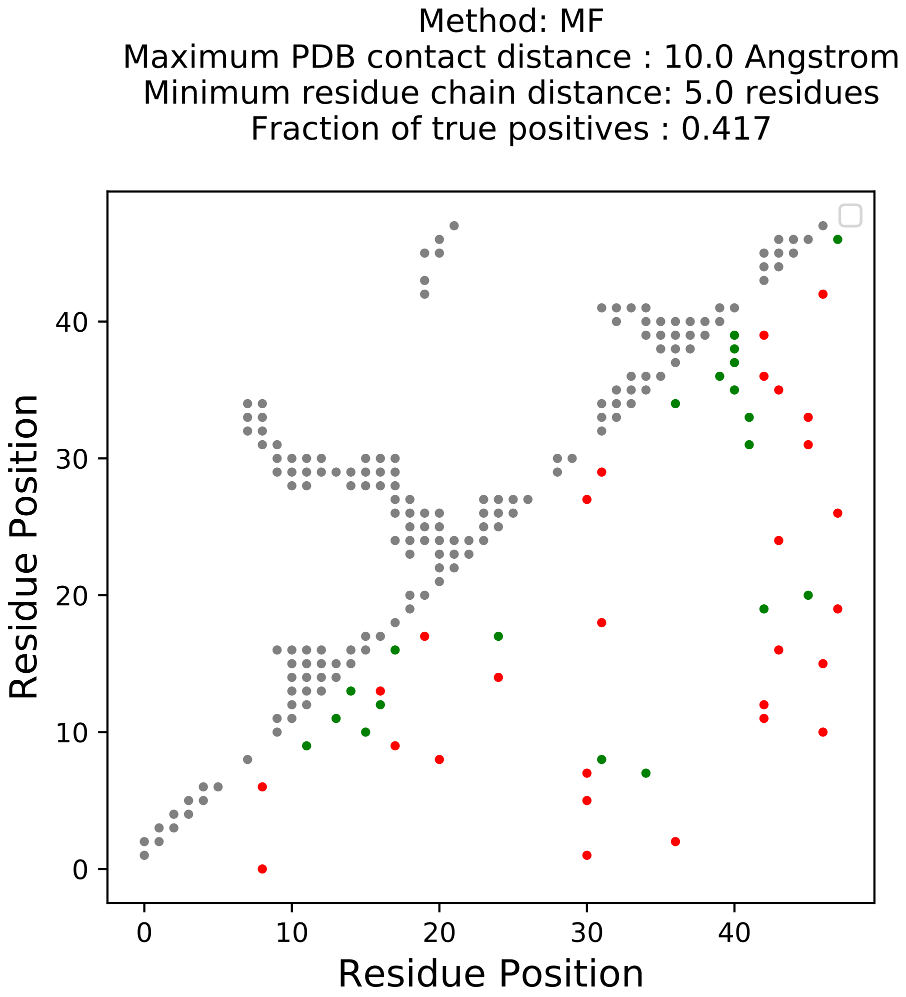
Method: MF
Maximum PDB contact distance : 10.0 Angstrom
Minimum residue chain distance: 5.0 residues
Fraction of true positives : 0.417
Residue Position
Residue Position
0
10
20
30
40

Method: PLM
Maximum PDB contact distance : 10.0 Angstrom
Minimum residue chain distance: 5.0 residues
Fraction of true positives : 0.396

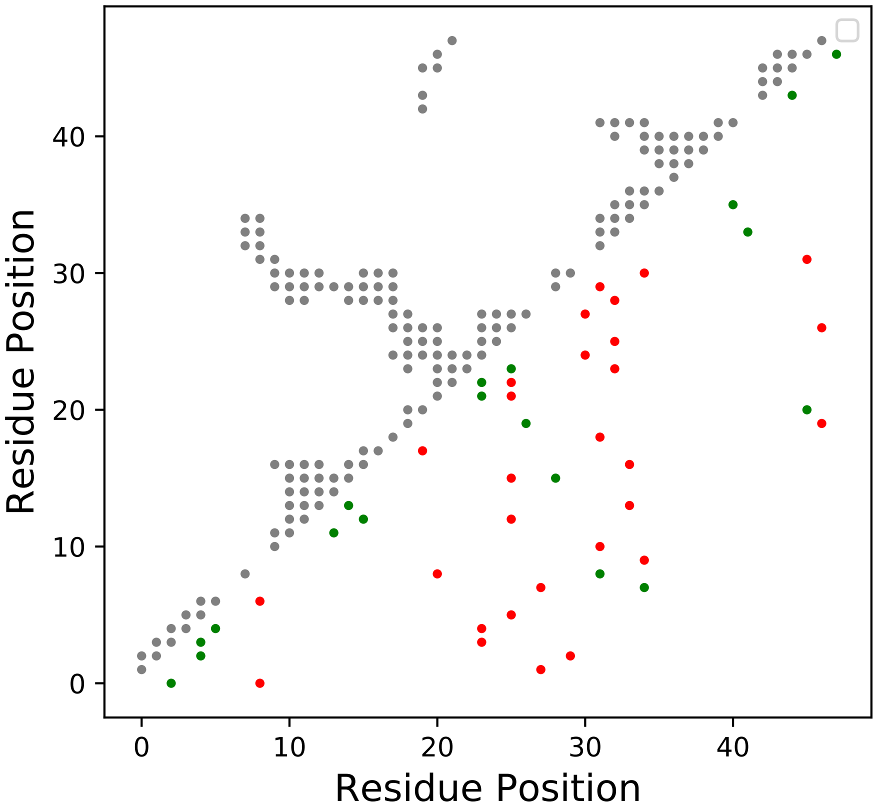

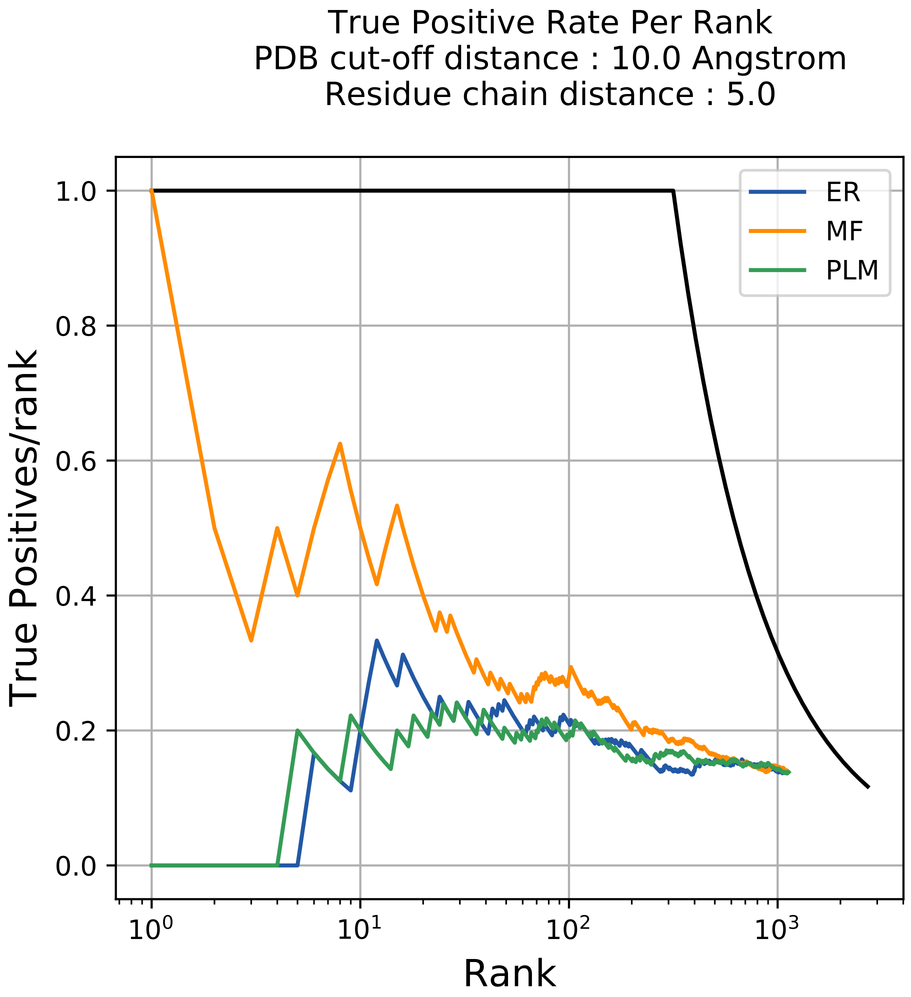
True Positive Rate Per Rank
PDB cut-off distance : 10.0 Angstrom
Residue chain distance : 5.0
True Positives/rank
Rank
ER
MF
PLM
1.0
0.8
0.6
0.4
0.2
0.0
$10^0$
$10^1$
$10^2$
$10^3$

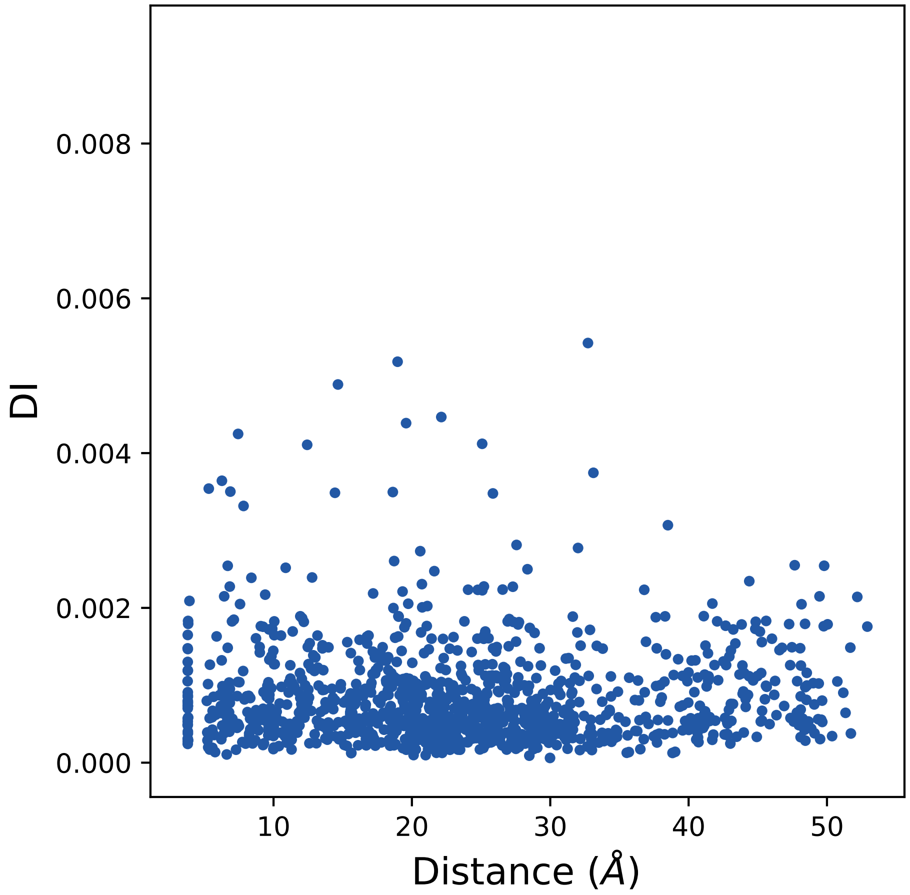

DI
0.000
0.002
0.004
0.006
0.008
10
20
30
40
50
Distance ($\AA$)

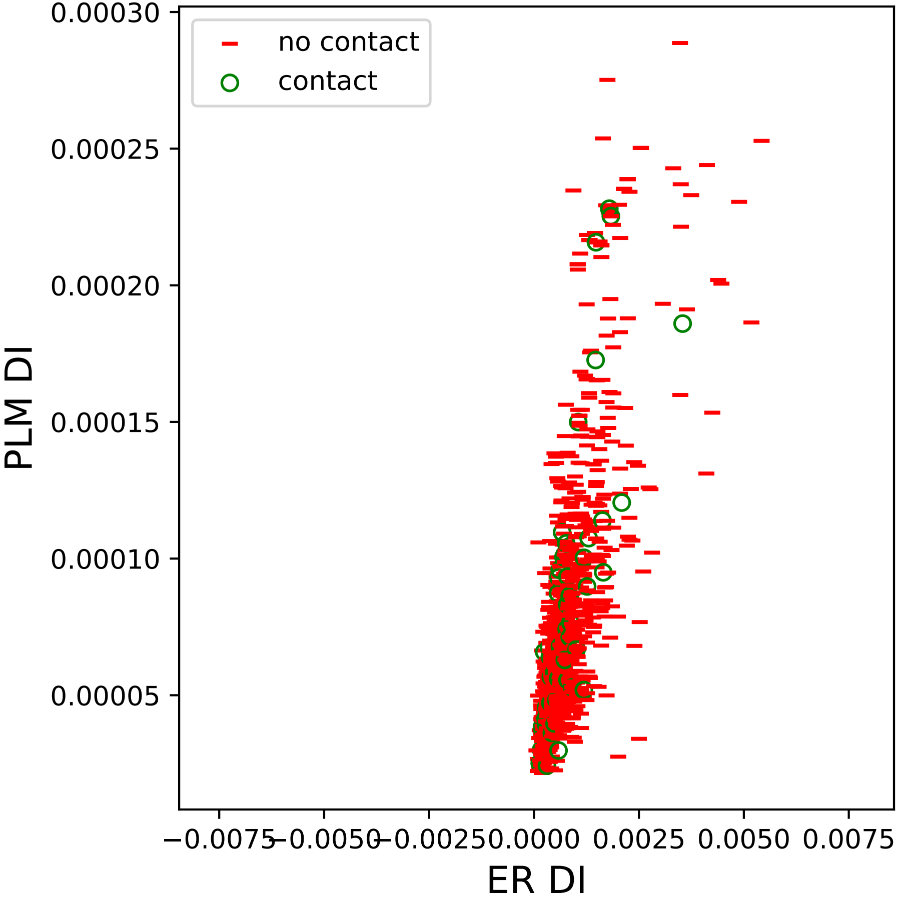

no contact
contact
PLM DI
ER DI
0.00030
0.00025
0.00020
0.00015
0.00010
0.00005
−0.0075
−0.0050
−0.0025
0.0000
0.0025
0.0050
0.0075

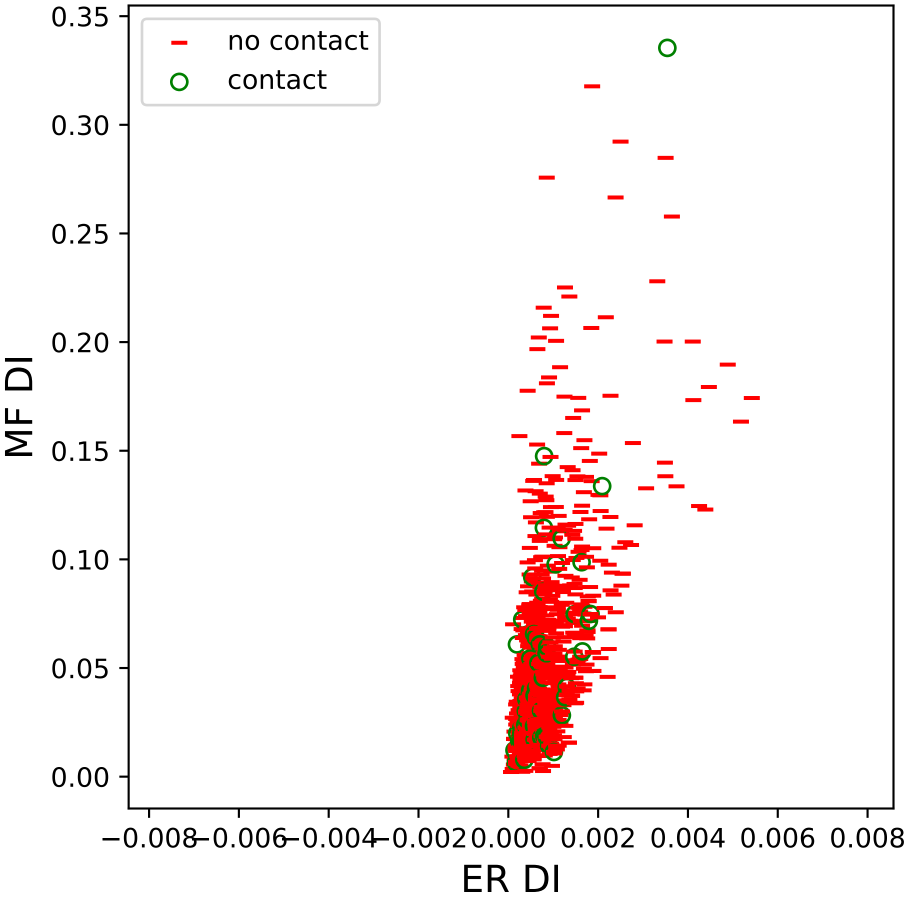
no contact
contact
0.35
0.30
0.25
0.20
0.15
0.10
0.05
0.00
MF DI
−0.008
−0.006
−0.004
−0.002
0.000
0.002
0.004
0.006
0.008
ER DI

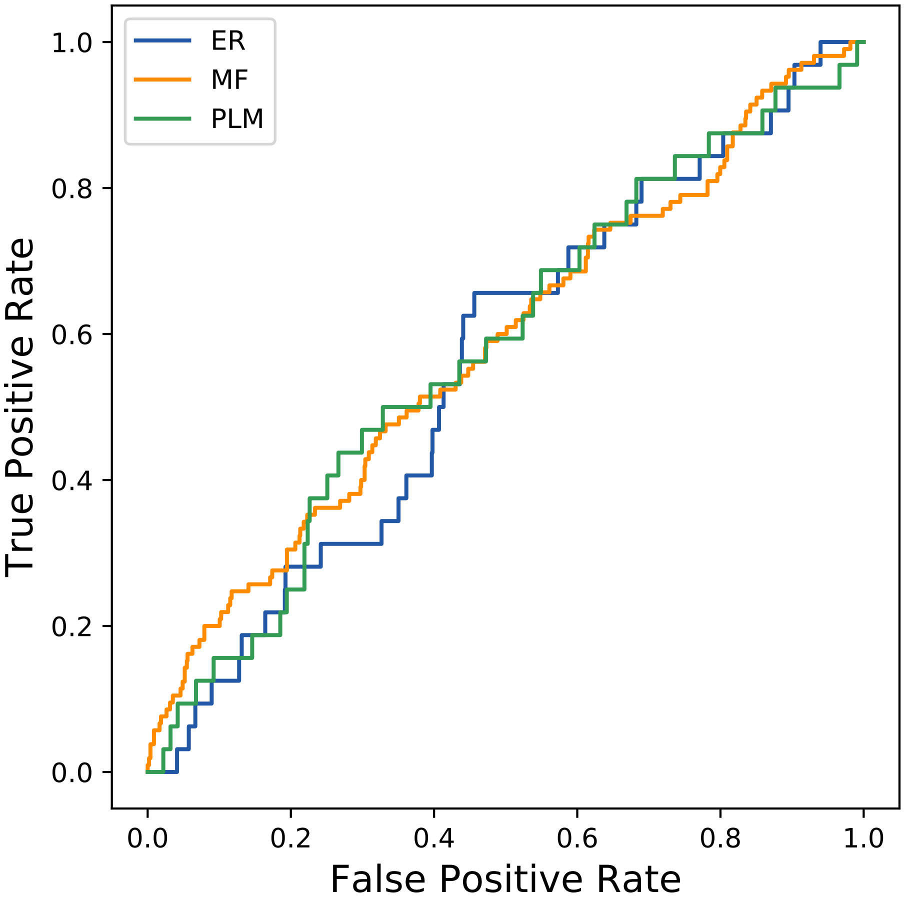

ER
MF
PLM
1.0
0.8
0.6
0.4
0.2
0.0
True Positive Rate
0.0
0.2
0.4
0.6
0.8
1.0
False Positive Rate
