## Supplemental Contact Maps for "Protein Structure Prediction with Expectation Reflection": 28seq_139col_4jes_PF06438_plots.pdf

Method: ER
Maximum PDB contact distance : 10.0 Angstrom
Minimum residue chain distance: 5.0 residues
Fraction of true positives : 0.381

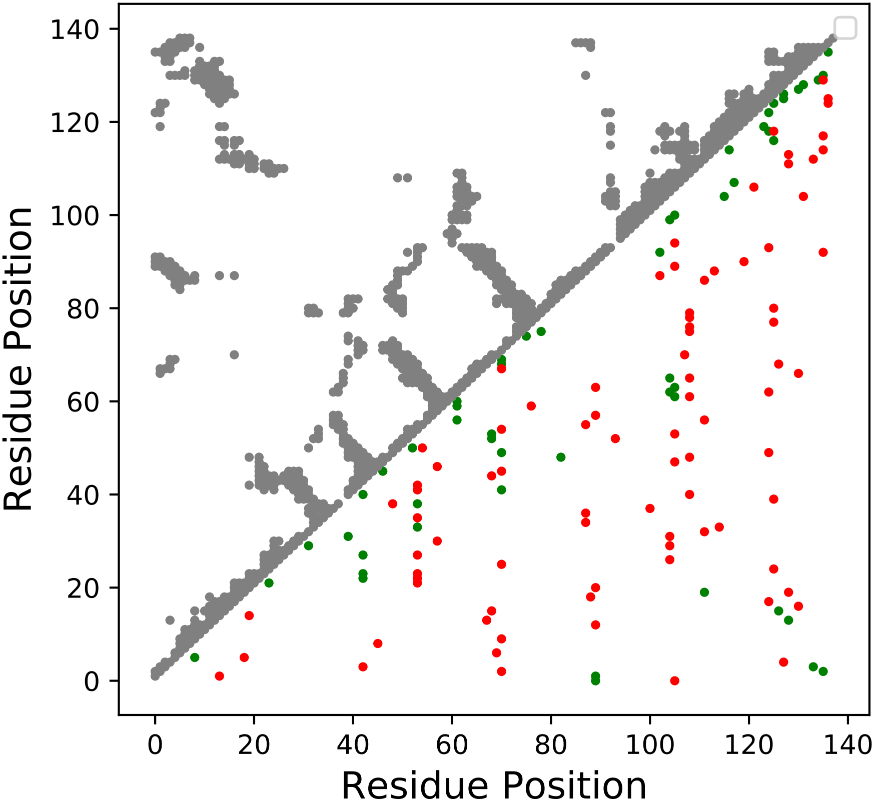

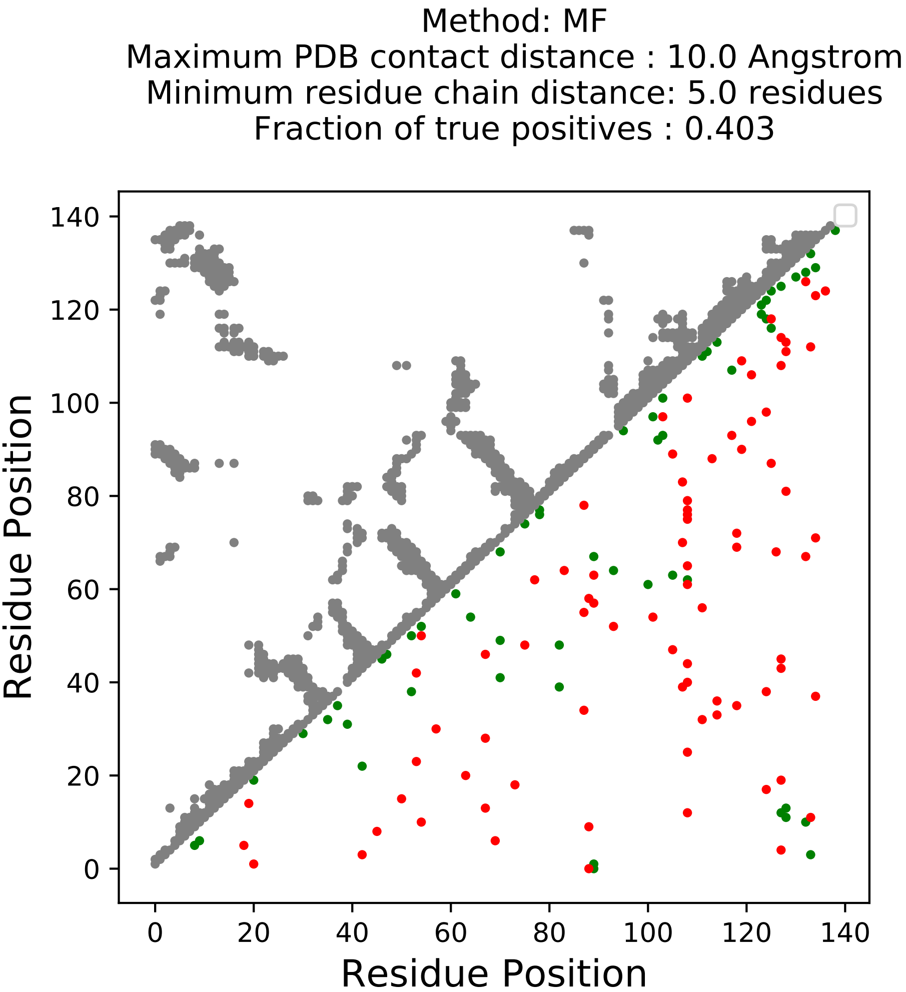
Method: MF
Maximum PDB contact distance : 10.0 Angstrom
Minimum residue chain distance: 5.0 residues
Fraction of true positives : 0.403
Residue Position
Residue Position
0
20
40
60
80
100
120
140

Method: PLM
Maximum PDB contact distance : 10.0 Angstrom
Minimum residue chain distance: 5.0 residues
Fraction of true positives : 0.417

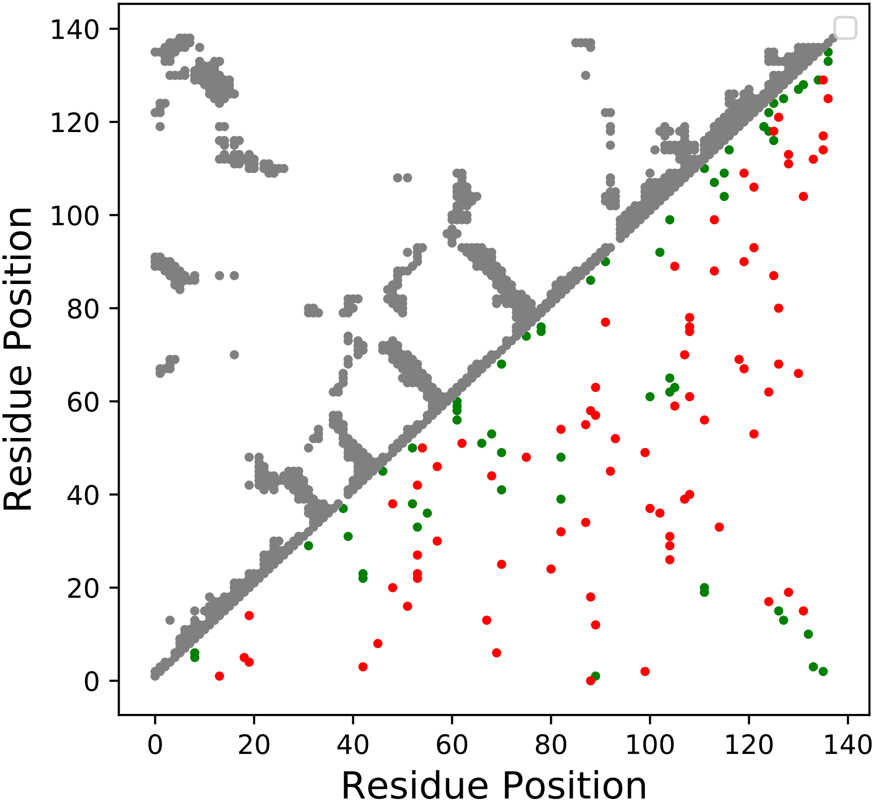

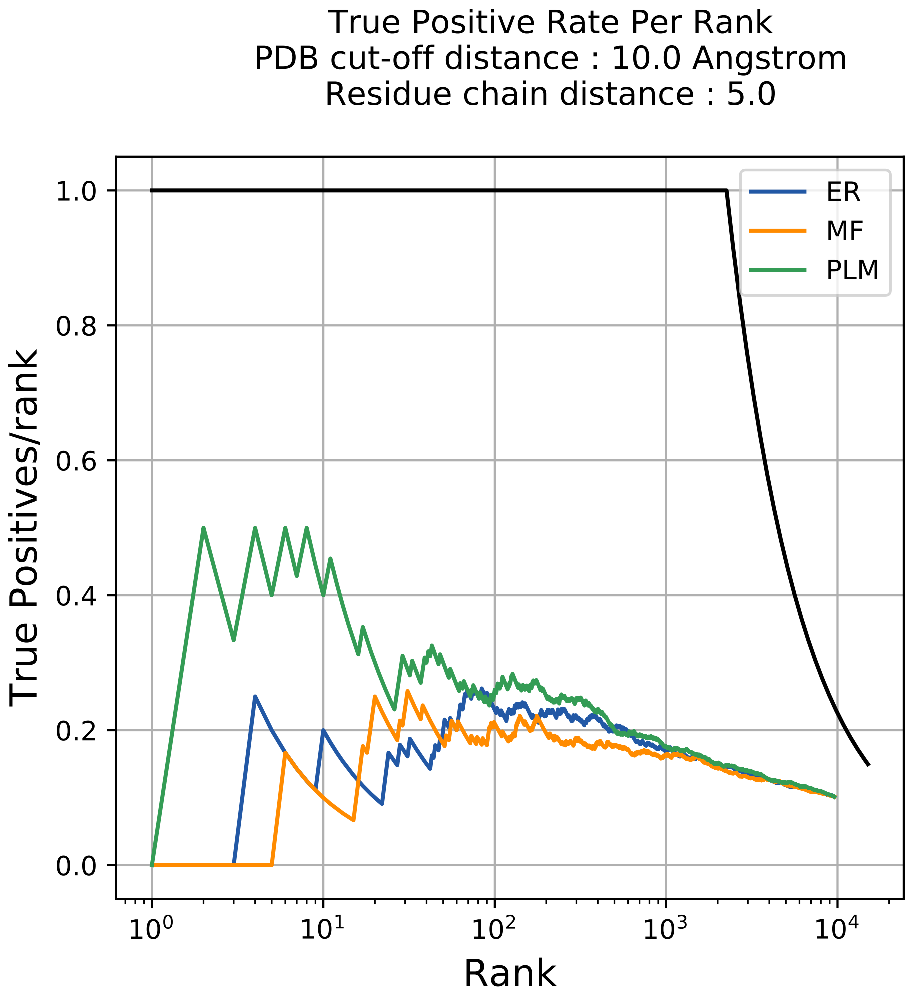

True Positive Rate Per Rank
PDB cut-off distance : 10.0 Angstrom
Residue chain distance : 5.0
ER
MF
PLM
1.0
0.8
0.6
0.4
0.2
0.0
True Positives/rank
$10^0$
$10^1$
$10^2$
$10^3$
$10^4$
Rank

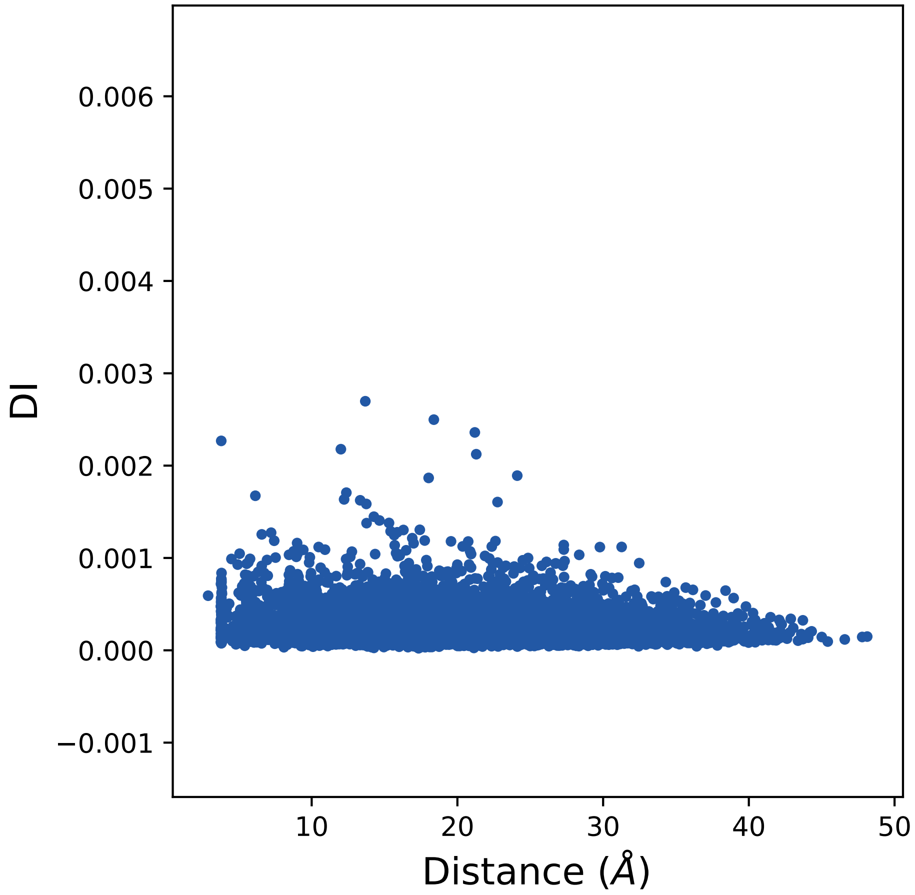

0.006
0.005
0.004
0.003
0.002
0.001
0.000
−0.001
DI
10
20
30
40
50
Distance (Å)

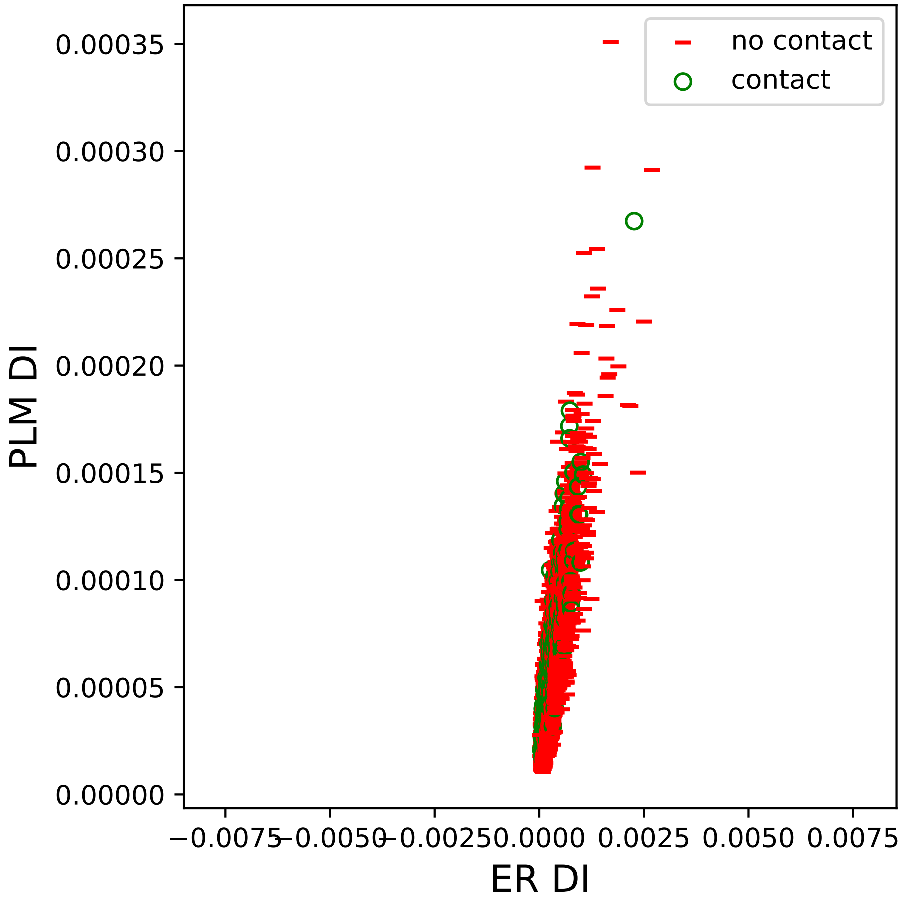

no contact
contact
PLM DI
0.00035
0.00030
0.00025
0.00020
0.00015
0.00010
0.00005
0.00000
−0.0075
−0.0050
−0.0025
0.0000
0.0025
0.0050
0.0075
ER DI

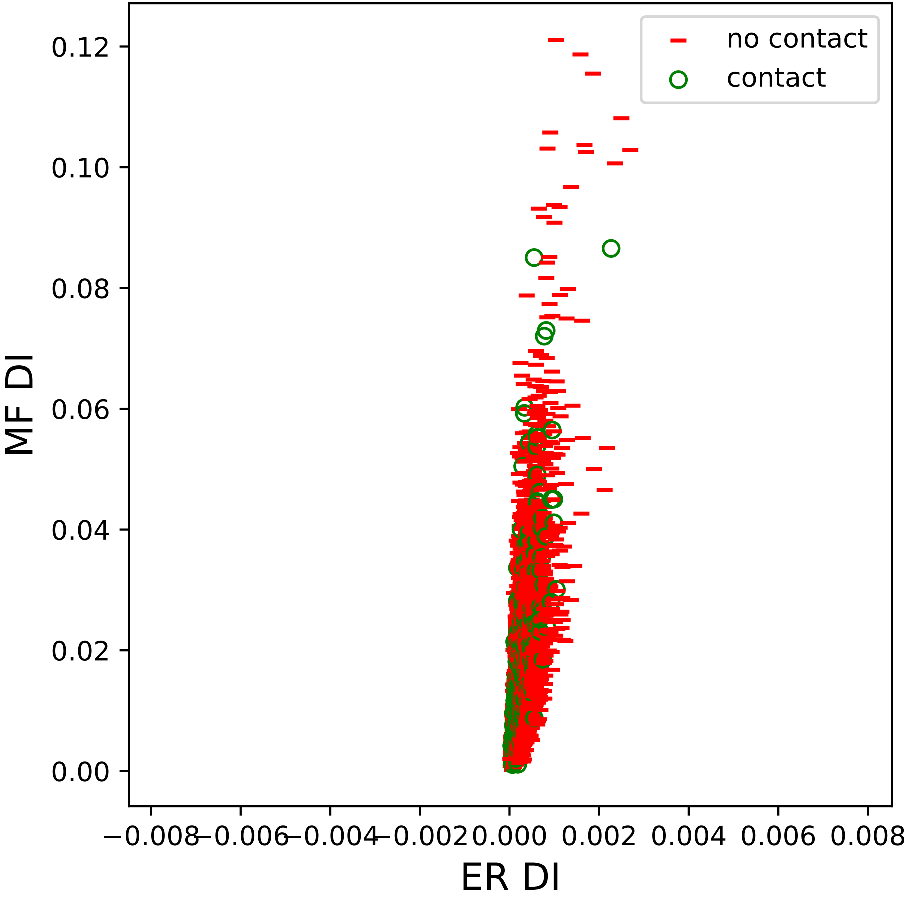

no contact
contact
MF DI
ER DI
0.12
0.10
0.08
0.06
0.04
0.02
0.00
−0.008
−0.006
−0.004
−0.002
0.000
0.002
0.004
0.006
0.008

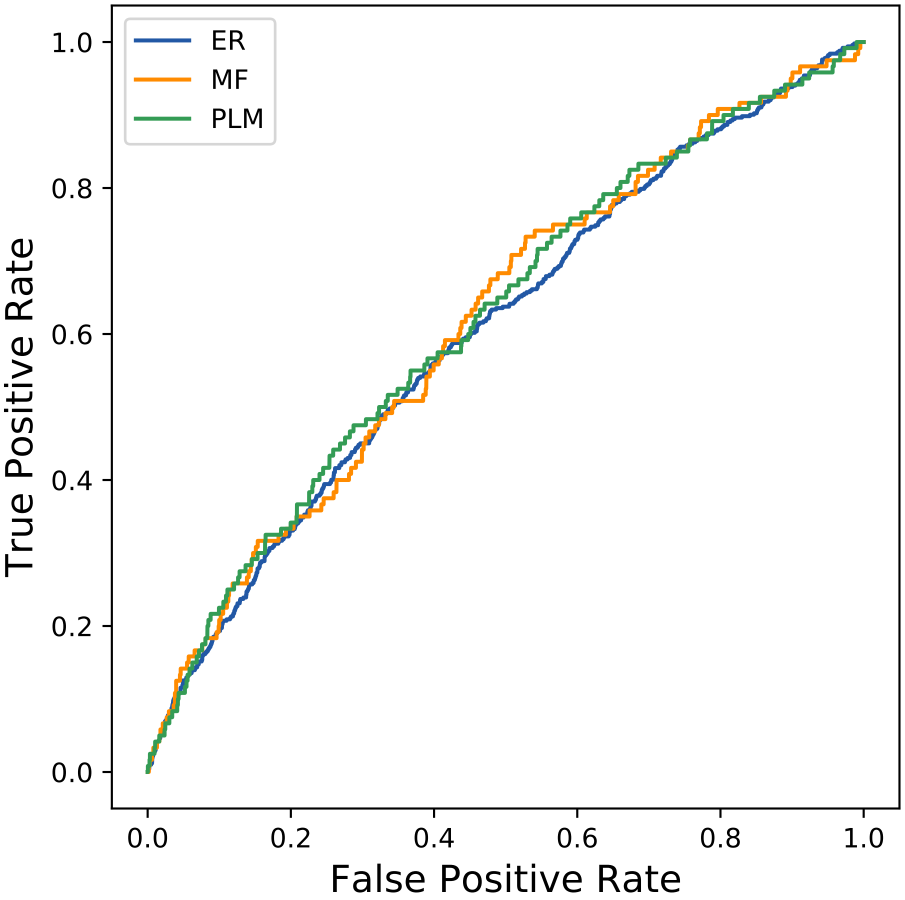

ER
MF
PLM
1.0
0.8
0.6
0.4
0.2
0.0
True Positive Rate
0.0
0.2
0.4
0.6
0.8
1.0
False Positive Rate
