## Supplemental Contact Maps for "Protein Structure Prediction with Expectation Reflection": 45seq_207col_6cam_PF08363_plots.pdf

Method: ER
Maximum PDB contact distance : 10.0 Angstrom
Minimum residue chain distance: 5.0 residues
Fraction of true positives : 0.343

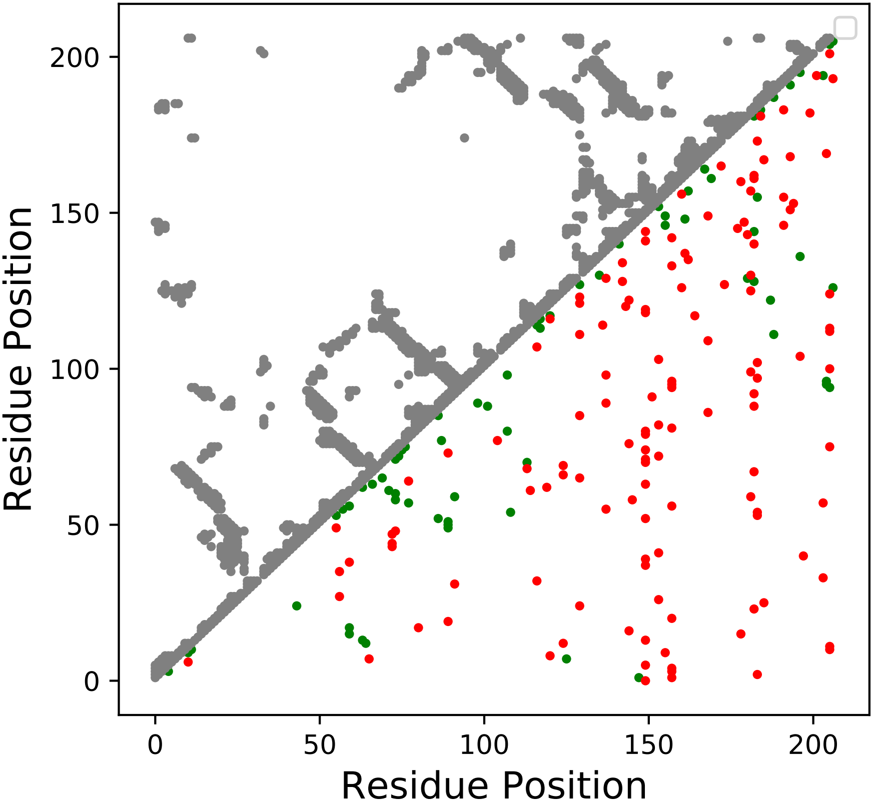

Method: MF
Maximum PDB contact distance : 10.0 Angstrom
Minimum residue chain distance: 5.0 residues
Fraction of true positives : 0.362

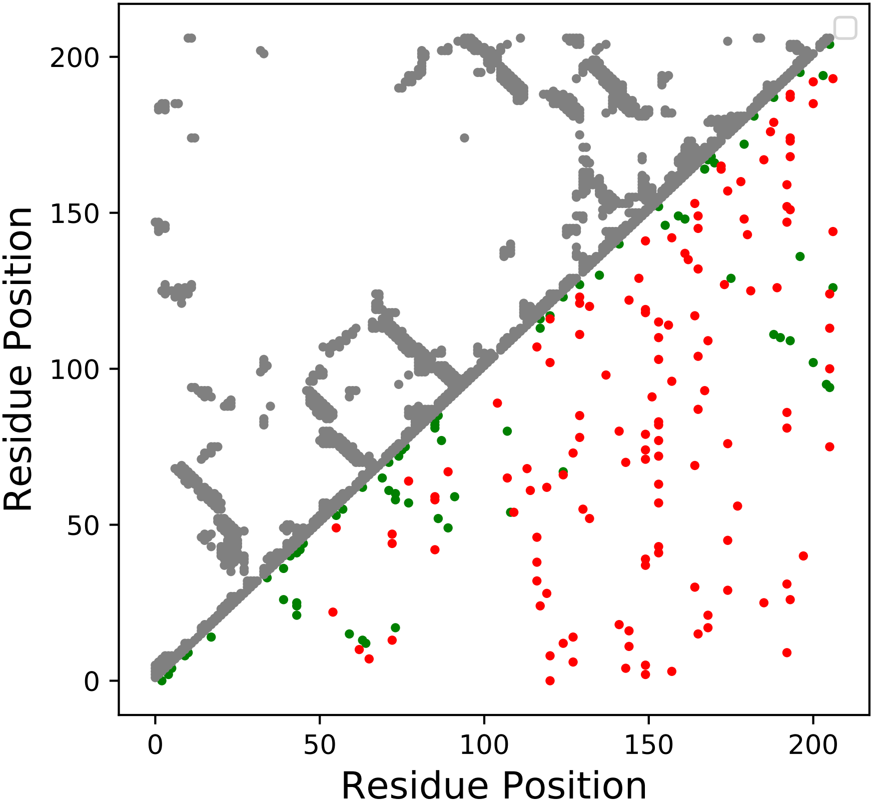

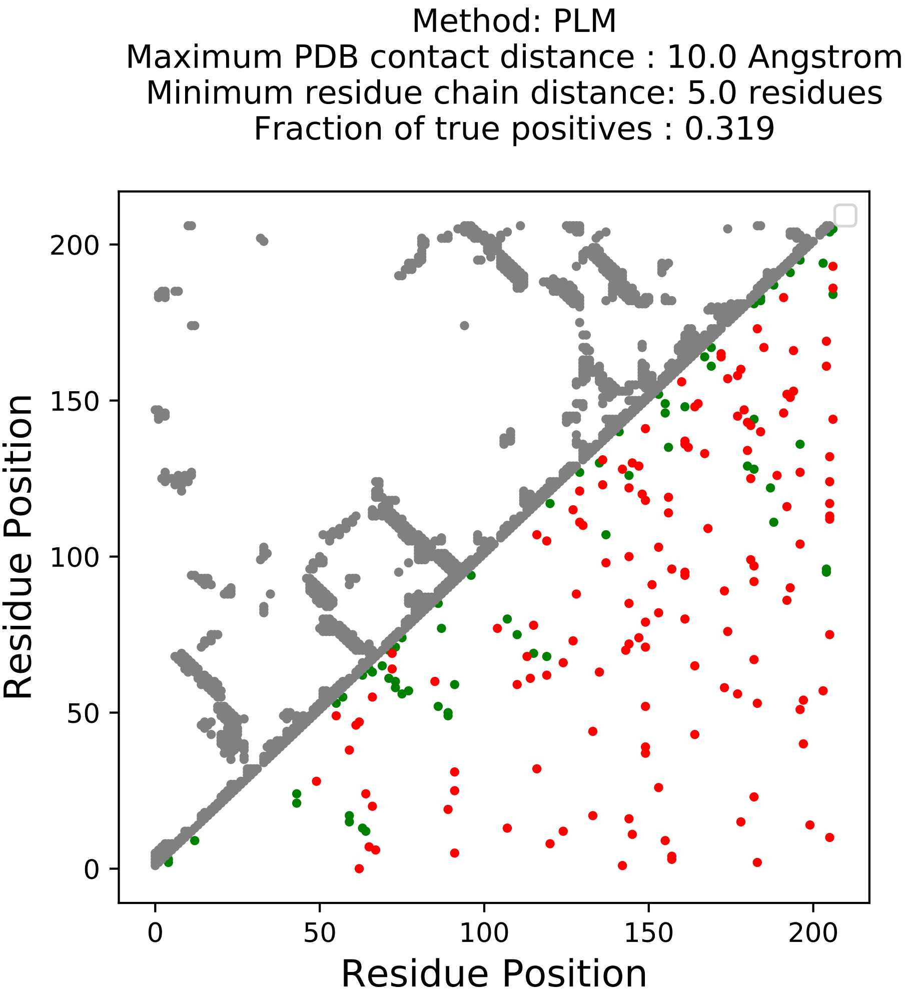
Method: PLM
Maximum PDB contact distance : 10.0 Angstrom
Minimum residue chain distance: 5.0 residues
Fraction of true positives : 0.319
Residue Position
Residue Position
0
50
100
150
200

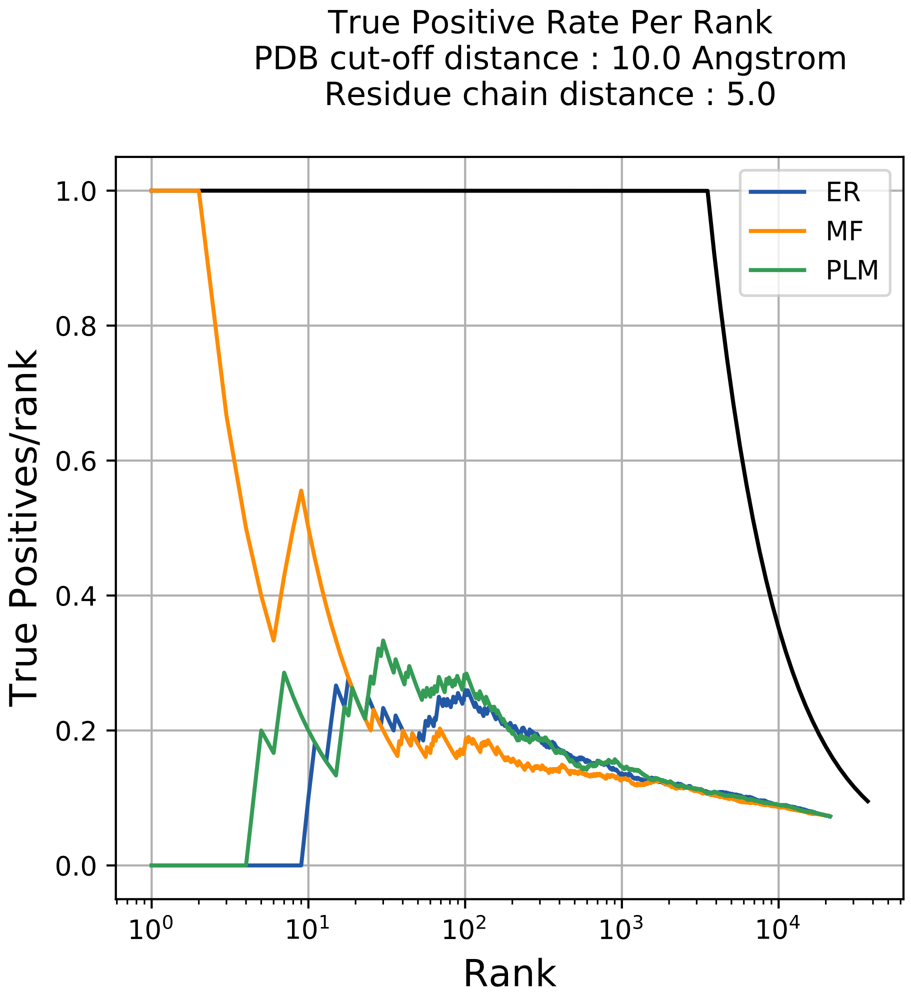
True Positive Rate Per Rank
PDB cut-off distance : 10.0 Angstrom
Residue chain distance : 5.0
ER
MF
PLM
True Positives/rank
1.0
0.8
0.6
0.4
0.2
0.0
$10^0$
$10^1$
$10^2$
$10^3$
$10^4$
Rank

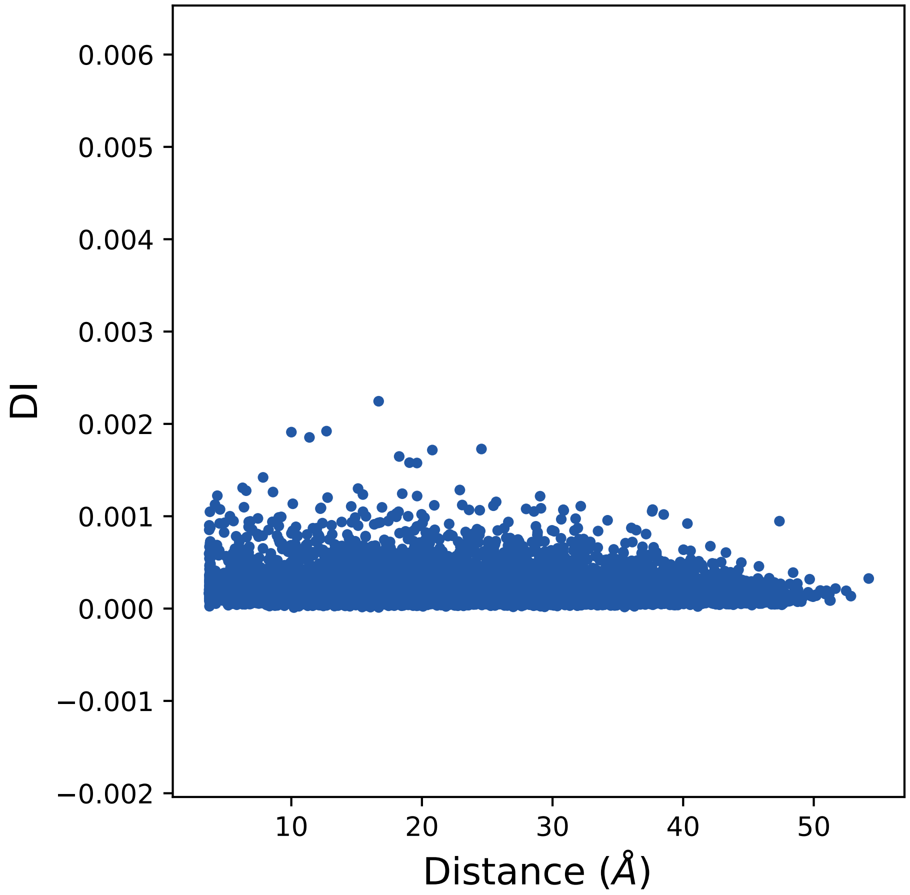

0.006
0.005
0.004
0.003
0.002
0.001
0.000
−0.001
−0.002
DI
10
20
30
40
50
Distance (Å)

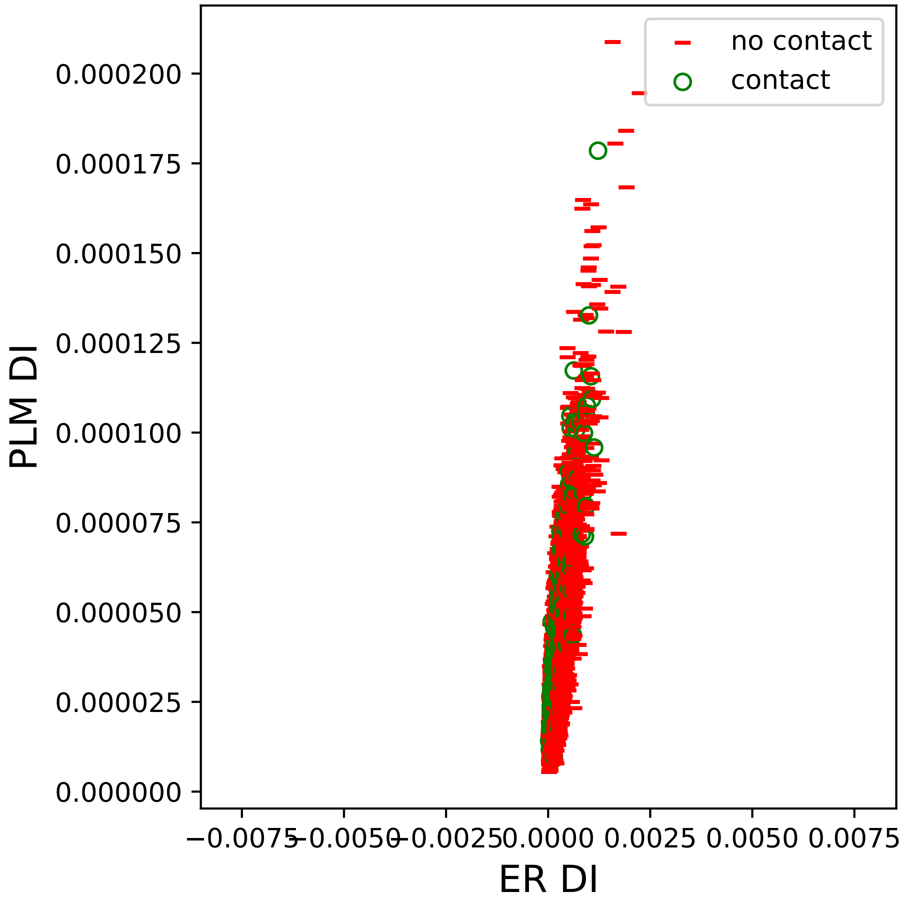
no contact
contact
PLM DI
ER DI
0.000200
0.000175
0.000150
0.000125
0.000100
0.000075
0.000050
0.000025
0.000000
−0.0075
−0.0050
−0.0025
0.0000
0.0025
0.0050
0.0075

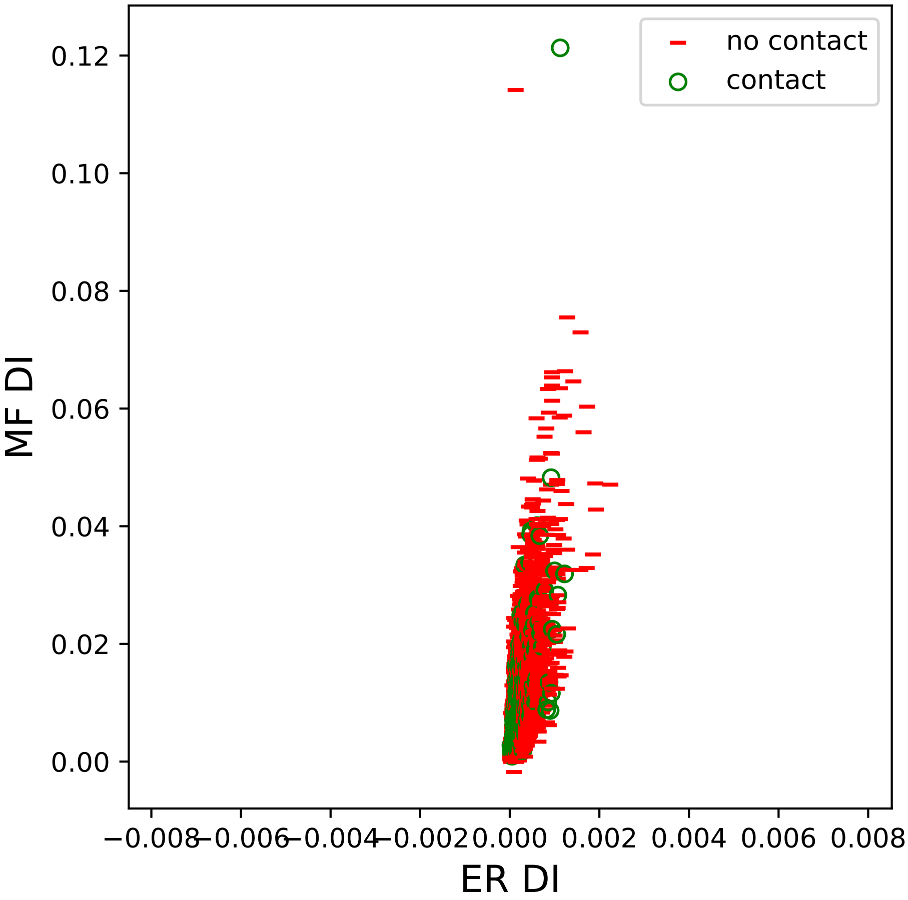

no contact
contact
MF DI
ER DI
0.12
0.10
0.08
0.06
0.04
0.02
0.00
−0.008
−0.006
−0.004
−0.002
0.000
0.002
0.004
0.006
0.008

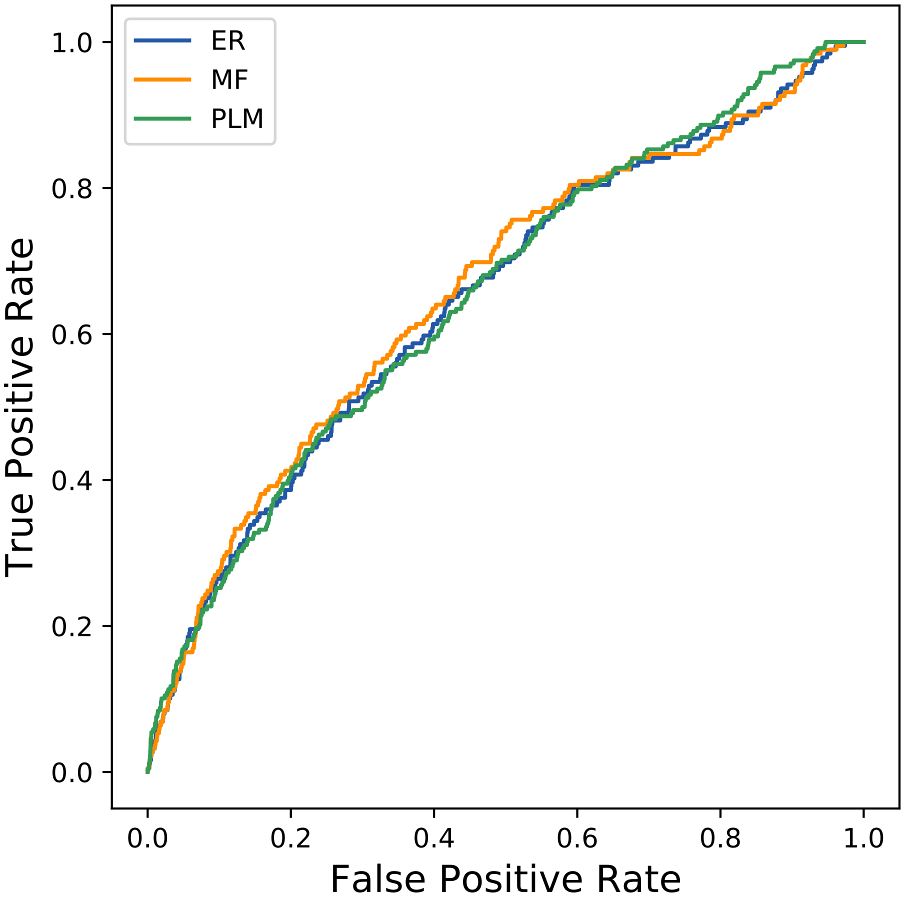

ER
MF
PLM
1.0
0.8
0.6
0.4
0.2
0.0
0.0
0.2
0.4
0.6
0.8
1.0
True Positive Rate
False Positive Rate
