## Supplemental Contact Maps for "Protein Structure Prediction with Expectation Reflection": 212seq_71col_4pz1_PF14131_plots.pdf

Method: ER
Maximum PDB contact distance : 10.0 Angstrom
Minimum residue chain distance: 5.0 residues
Fraction of true positives : 0.423

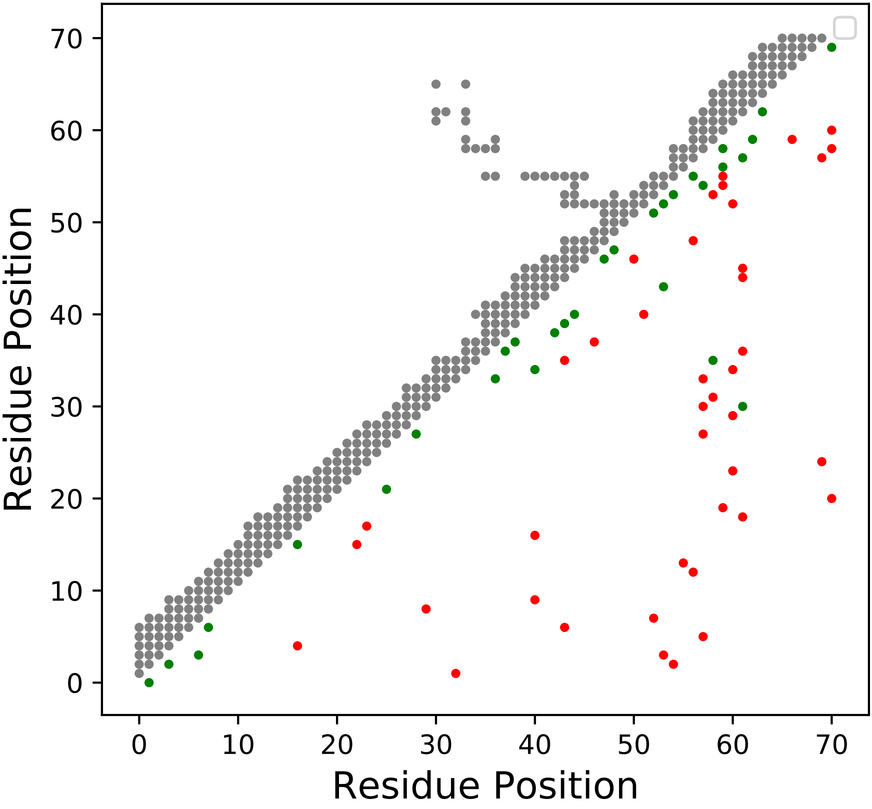

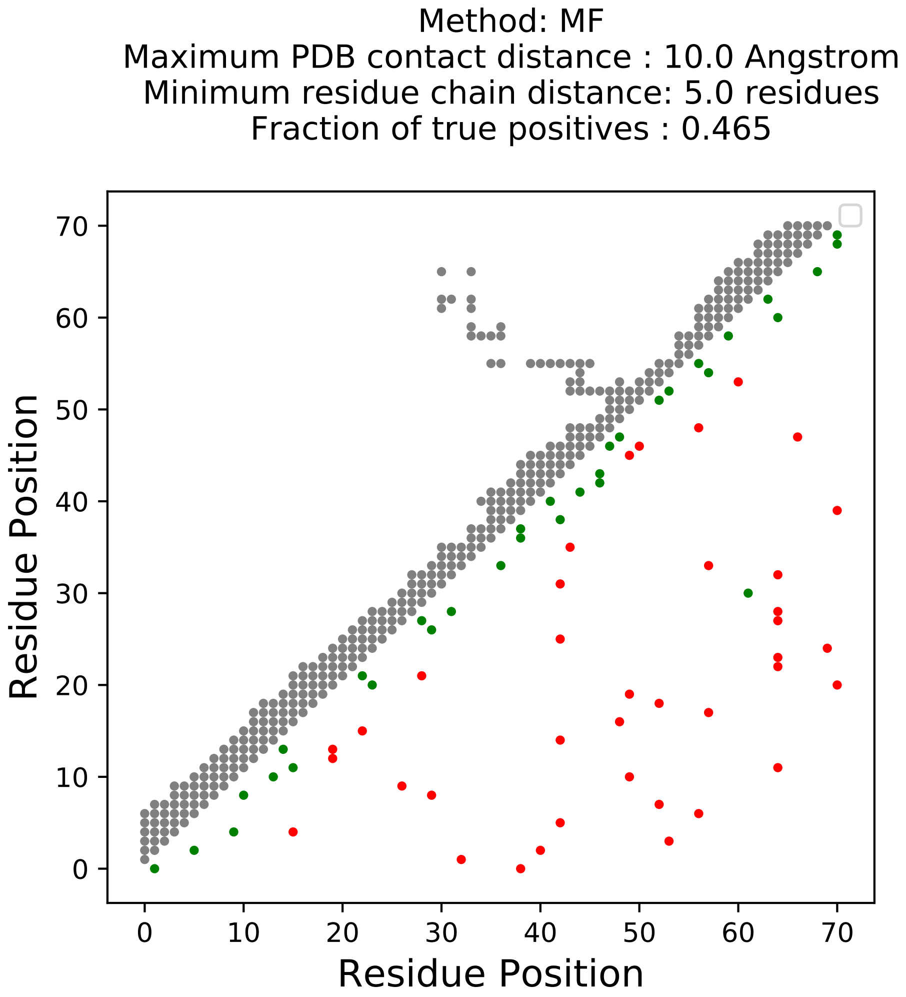
Method: MF
Maximum PDB contact distance : 10.0 Angstrom
Minimum residue chain distance: 5.0 residues
Fraction of true positives : 0.465
Residue Position
Residue Position
0
10
20
30
40
50
60
70

Method: PLM
Maximum PDB contact distance : 10.0 Angstrom
Minimum residue chain distance: 5.0 residues
Fraction of true positives : 0.437

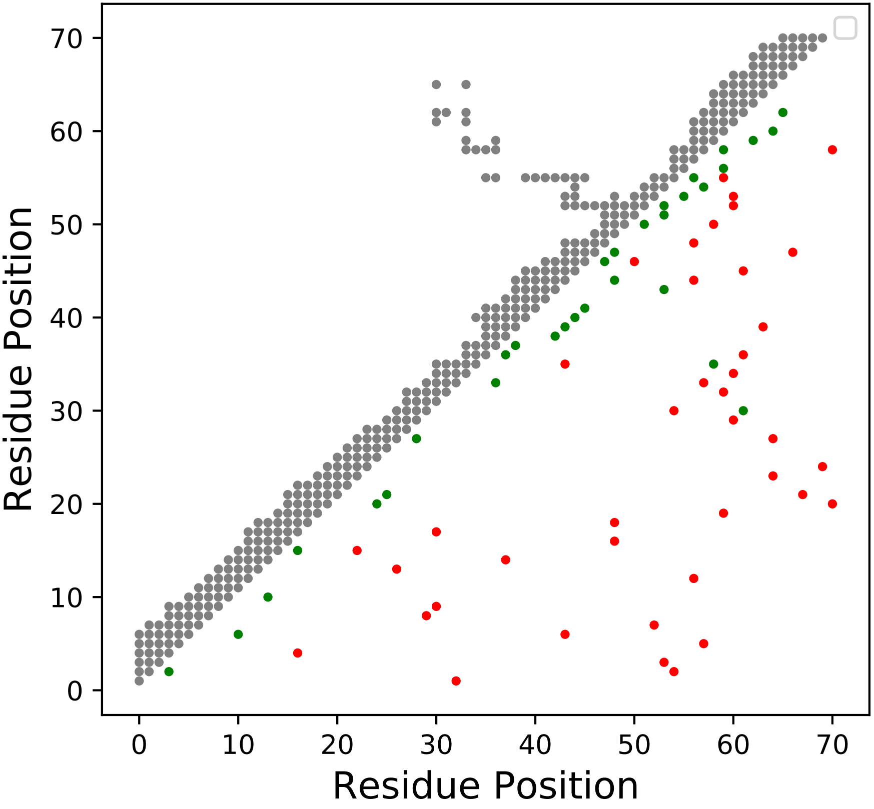

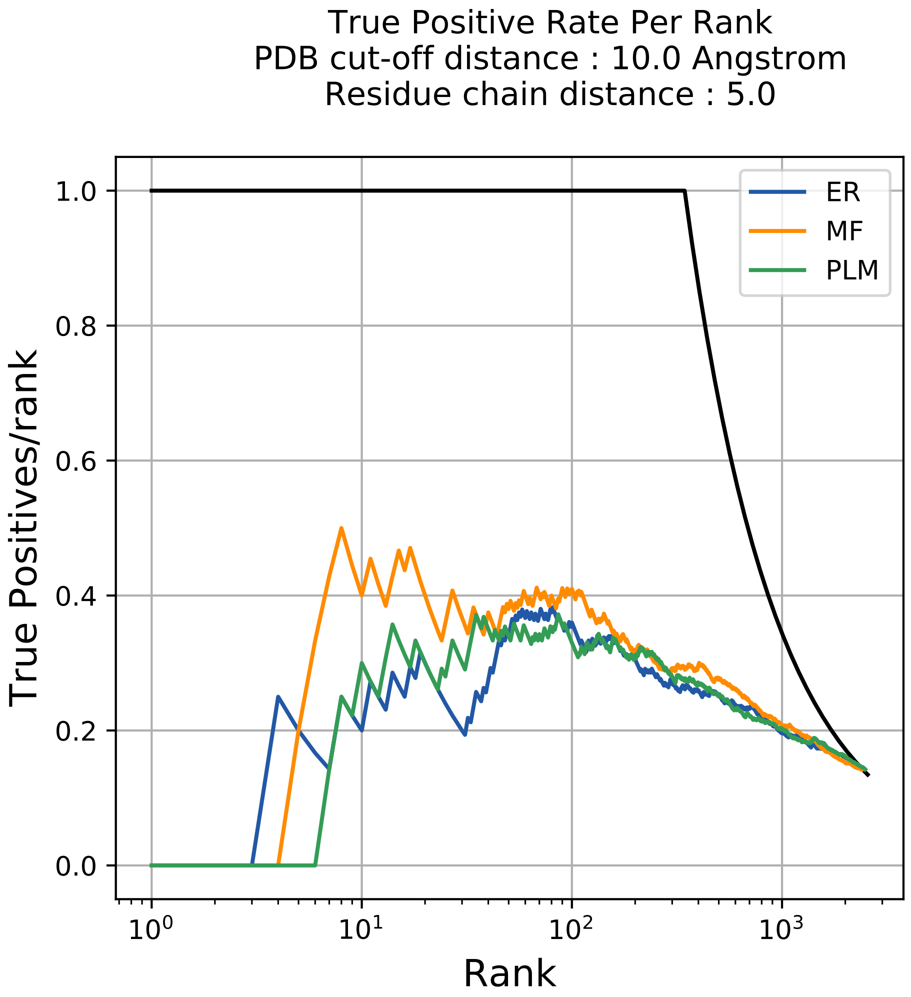

True Positive Rate Per Rank
PDB cut-off distance : 10.0 Angstrom
Residue chain distance : 5.0
True Positives/rank
Rank
ER
MF
PLM
0.0
0.2
0.4
0.6
0.8
1.0
$10^0$
$10^1$
$10^2$
$10^3$

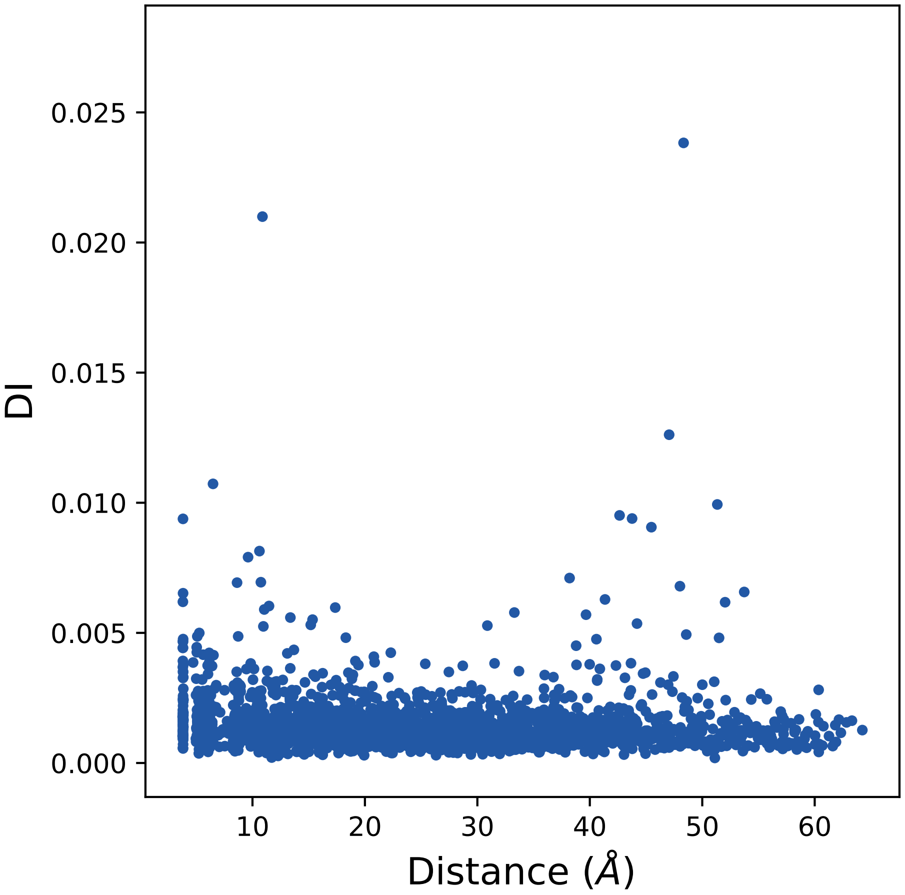

0.025
0.020
0.015
0.010
0.005
0.000
DI
10
20
30
40
50
60
Distance (Å)

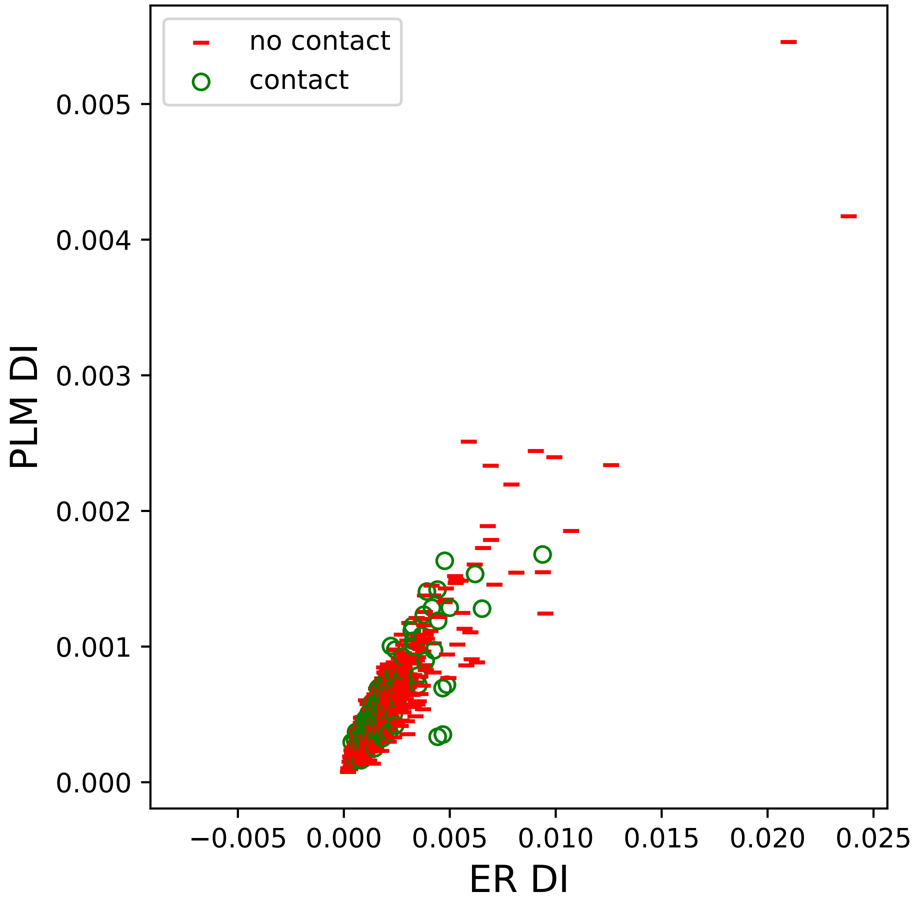

no contact
contact
0.005
0.004
0.003
0.002
0.001
0.000
PLM DI
−0.005
0.000
0.005
0.010
0.015
0.020
0.025
ER DI

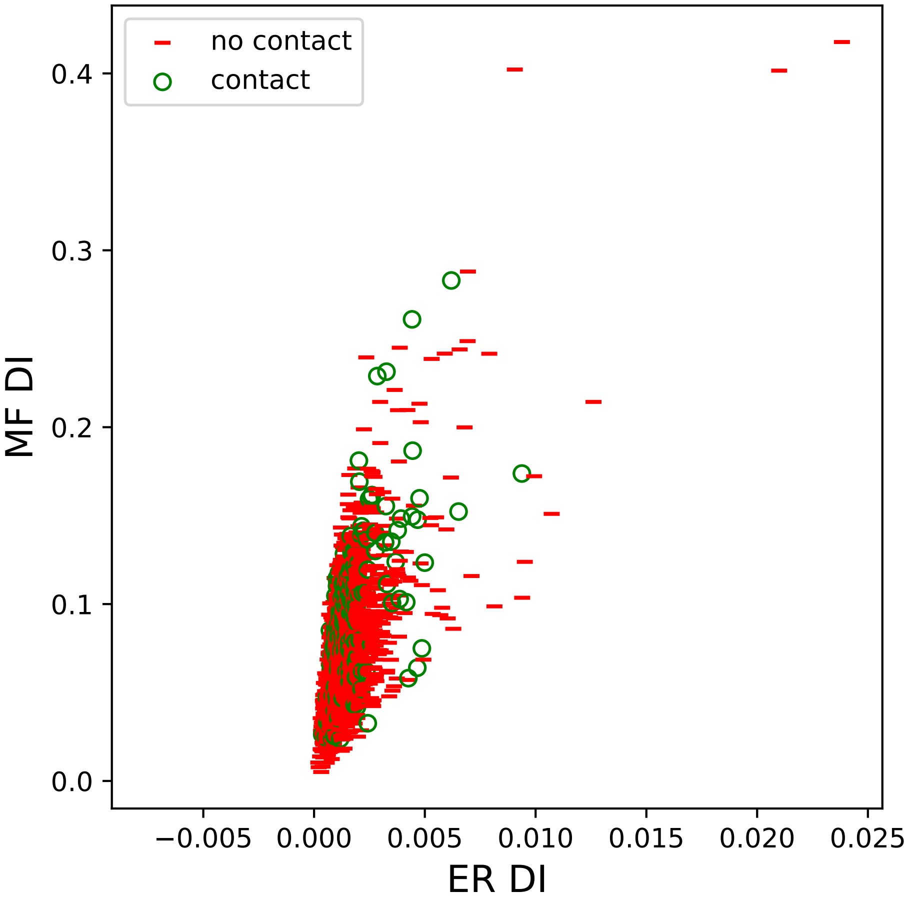
no contact
contact
MF DI
ER DI
0.0
0.1
0.2
0.3
0.4
−0.005
0.000
0.005
0.010
0.015
0.020
0.025

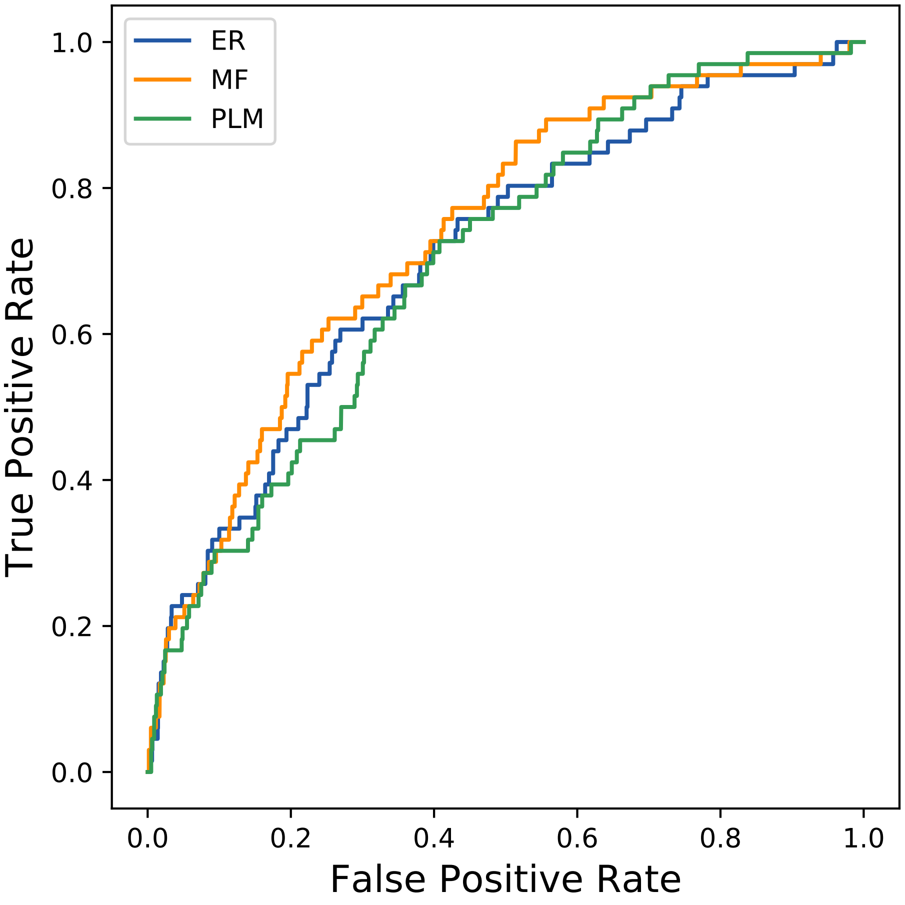
ER
MF
PLM
True Positive Rate
False Positive Rate
0.0
0.2
0.4
0.6
0.8
1.0
