## Supplemental Contact Maps for "Protein Structure Prediction with Expectation Reflection": 216seq_23col_1jgk_PF00087_plots.pdf

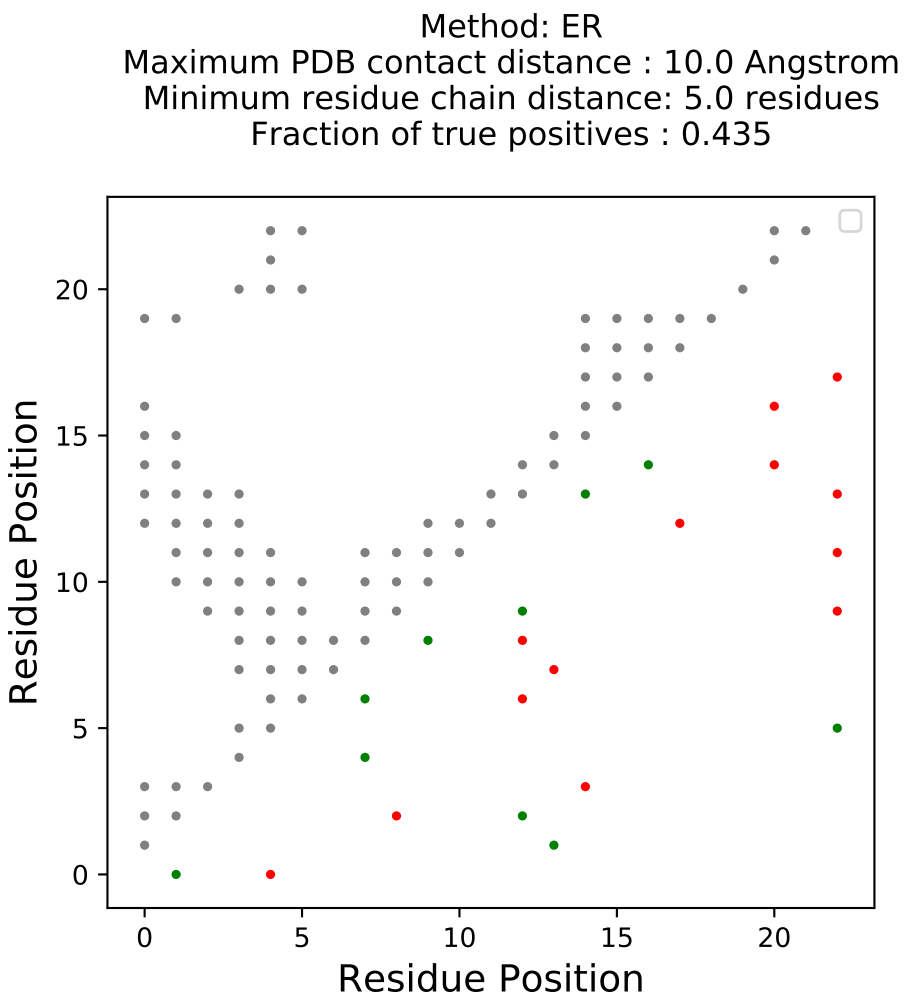
Method: ER
Maximum PDB contact distance : 10.0 Angstrom
Minimum residue chain distance: 5.0 residues
Fraction of true positives : 0.435
Residue Position
Residue Position

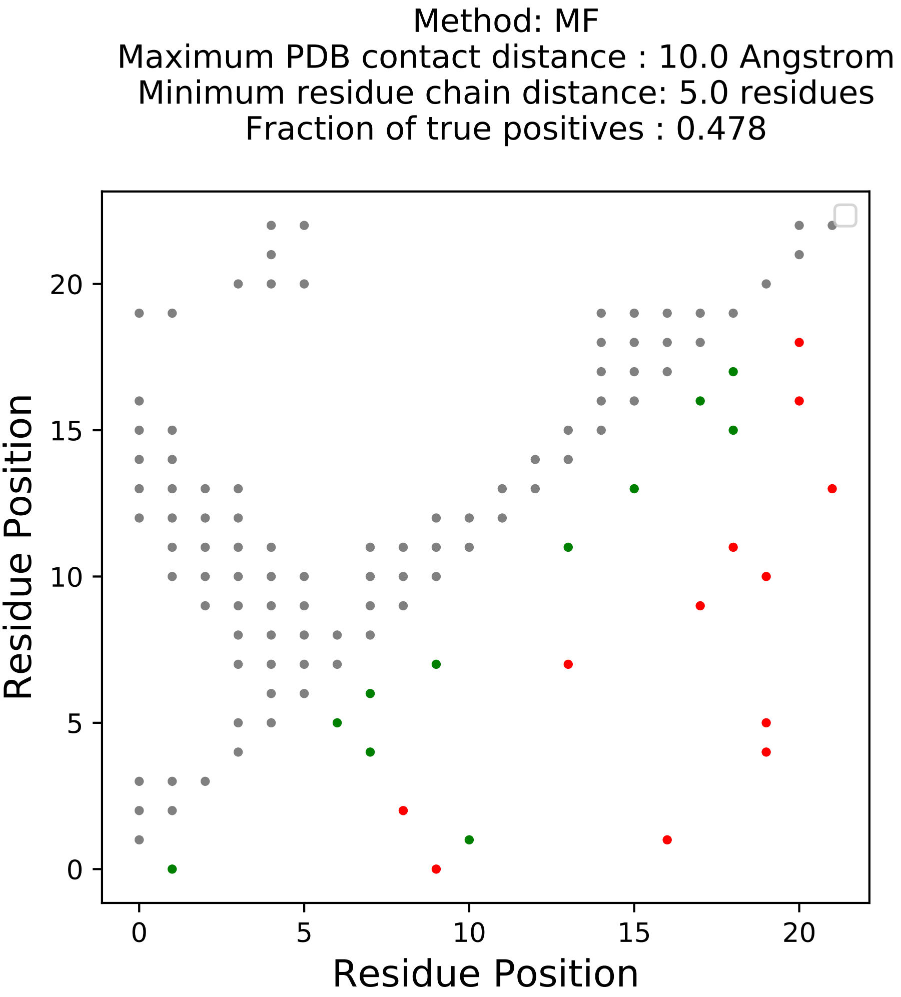
Method: MF
Maximum PDB contact distance : 10.0 Angstrom
Minimum residue chain distance: 5.0 residues
Fraction of true positives : 0.478
Residue Position
Residue Position
0
5
10
15
20

Method: PLM
Maximum PDB contact distance : 10.0 Angstrom
Minimum residue chain distance: 5.0 residues
Fraction of true positives : 0.478

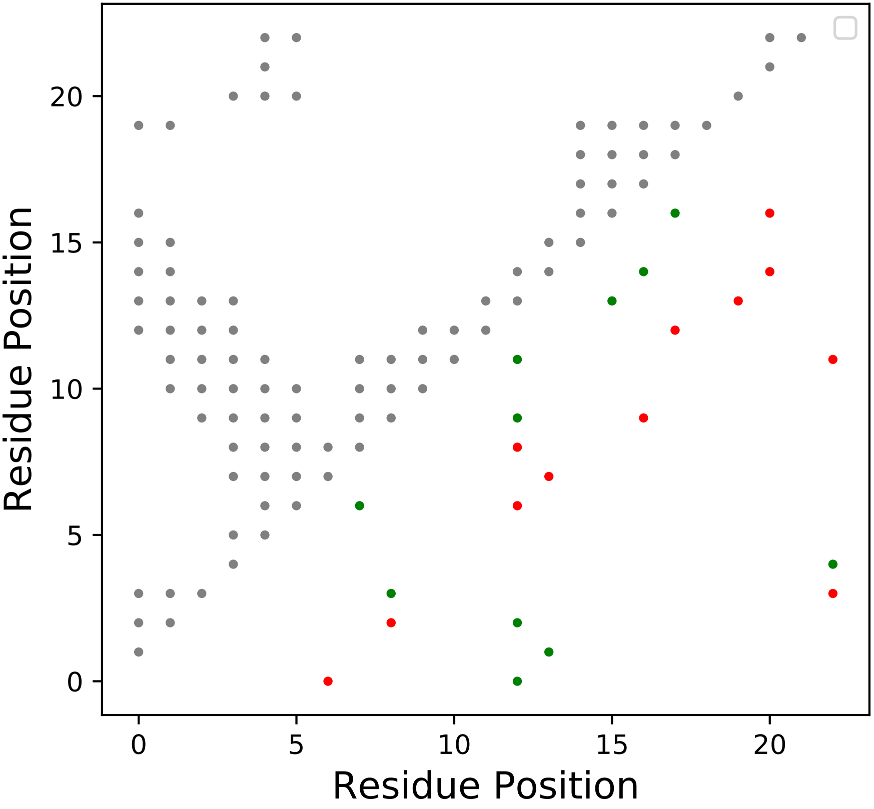

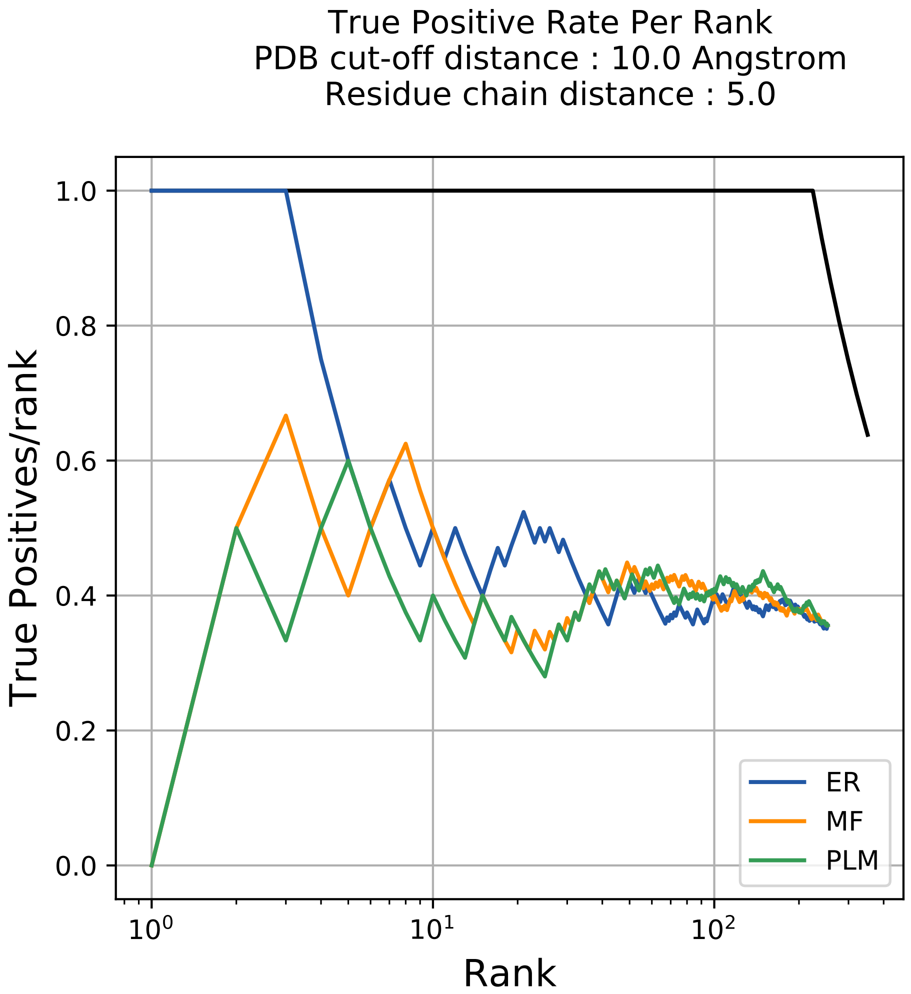
True Positive Rate Per Rank
PDB cut-off distance : 10.0 Angstrom
Residue chain distance : 5.0
True Positives/rank
Rank
1.0
0.8
0.6
0.4
0.2
0.0
10^0
10^1
10^2
ER
MF
PLM

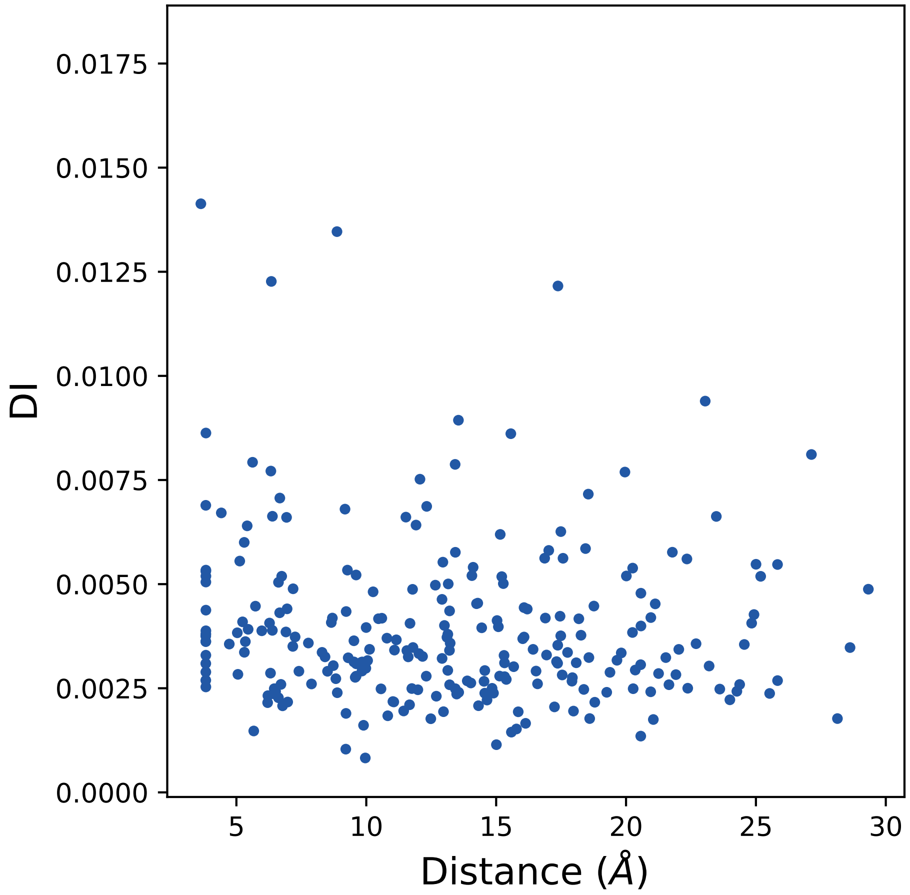
DI
Distance (Å)
0.0000
0.0025
0.0050
0.0075
0.0100
0.0125
0.0150
0.0175
5
10
15
20
25
30

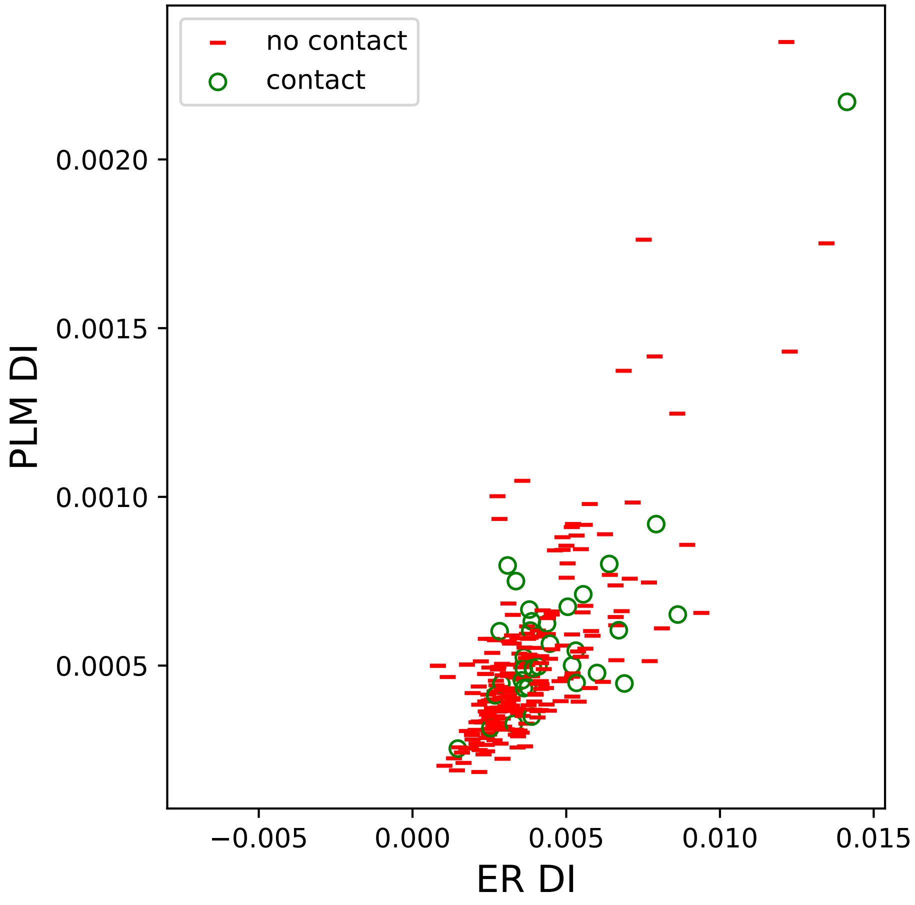

no contact
contact
0.0020
0.0015
0.0010
0.0005
PLM DI
−0.005
0.000
0.005
0.010
0.015
ER DI

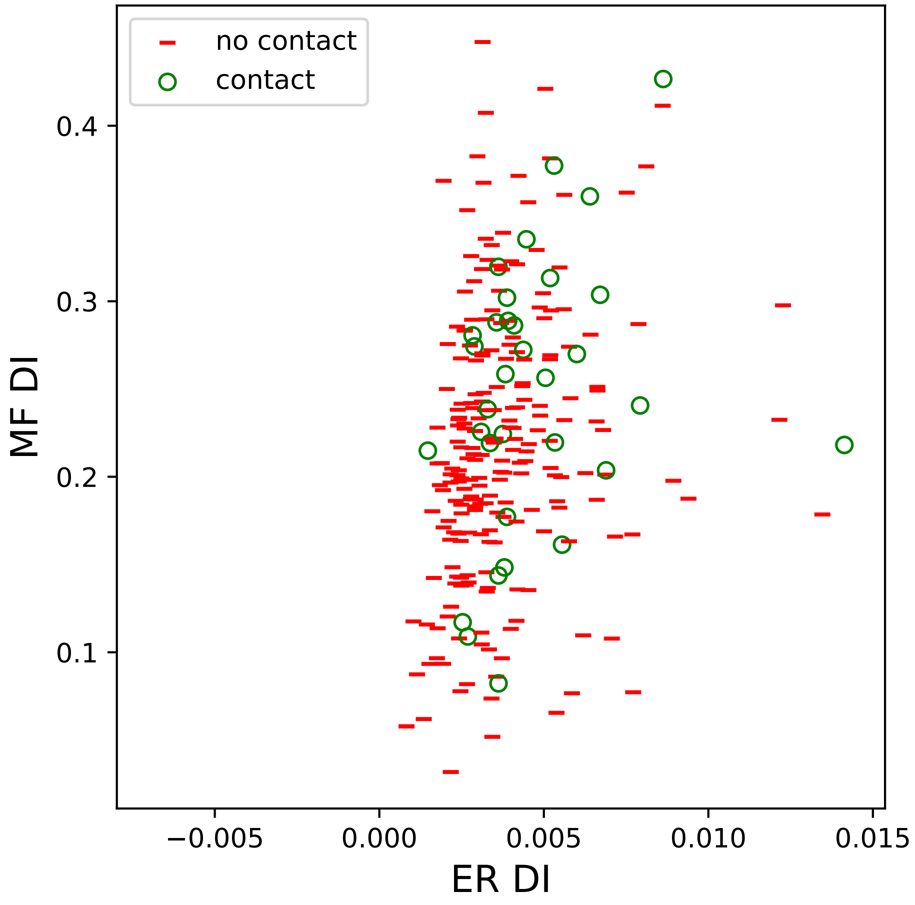
no contact
contact
0.1
0.2
0.3
0.4
MF DI
−0.005
0.000
0.005
0.010
0.015
ER DI

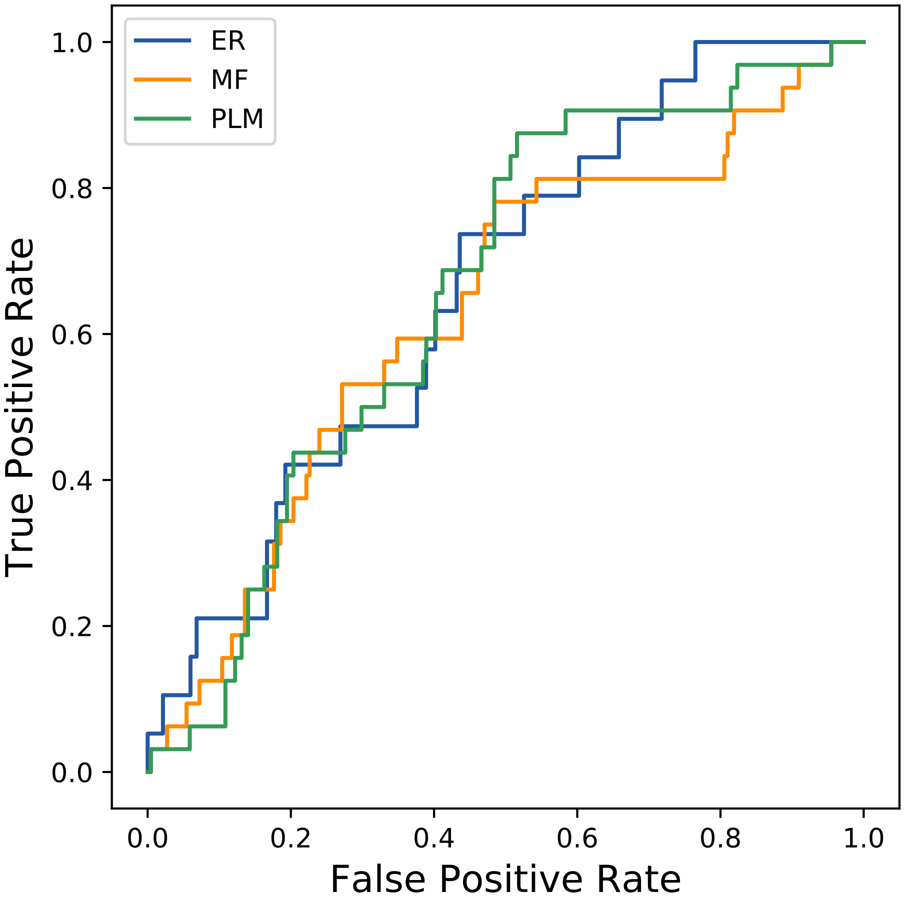

ER
MF
PLM
True Positive Rate
False Positive Rate
0.0
0.2
0.4
0.6
0.8
1.0
