## Supplemental Contact Maps for "Protein Structure Prediction with Expectation Reflection": 222seq_54col_6bp8_PF11978_plots.pdf

Method: ER
Maximum PDB contact distance : 10.0 Angstrom
Minimum residue chain distance: 5.0 residues
Fraction of true positives : 0.352

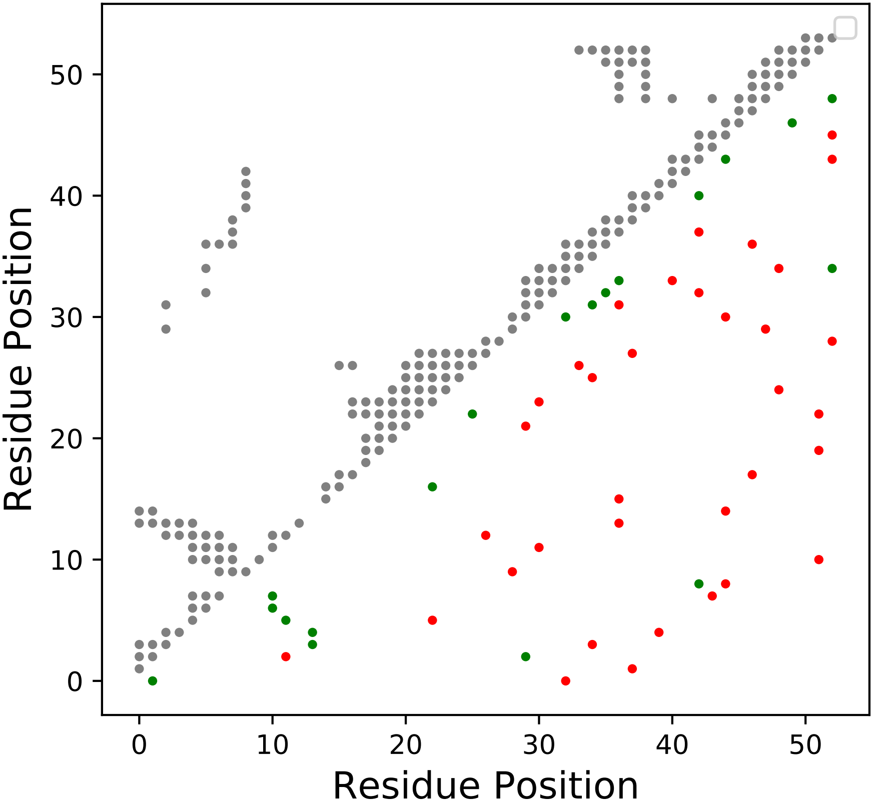

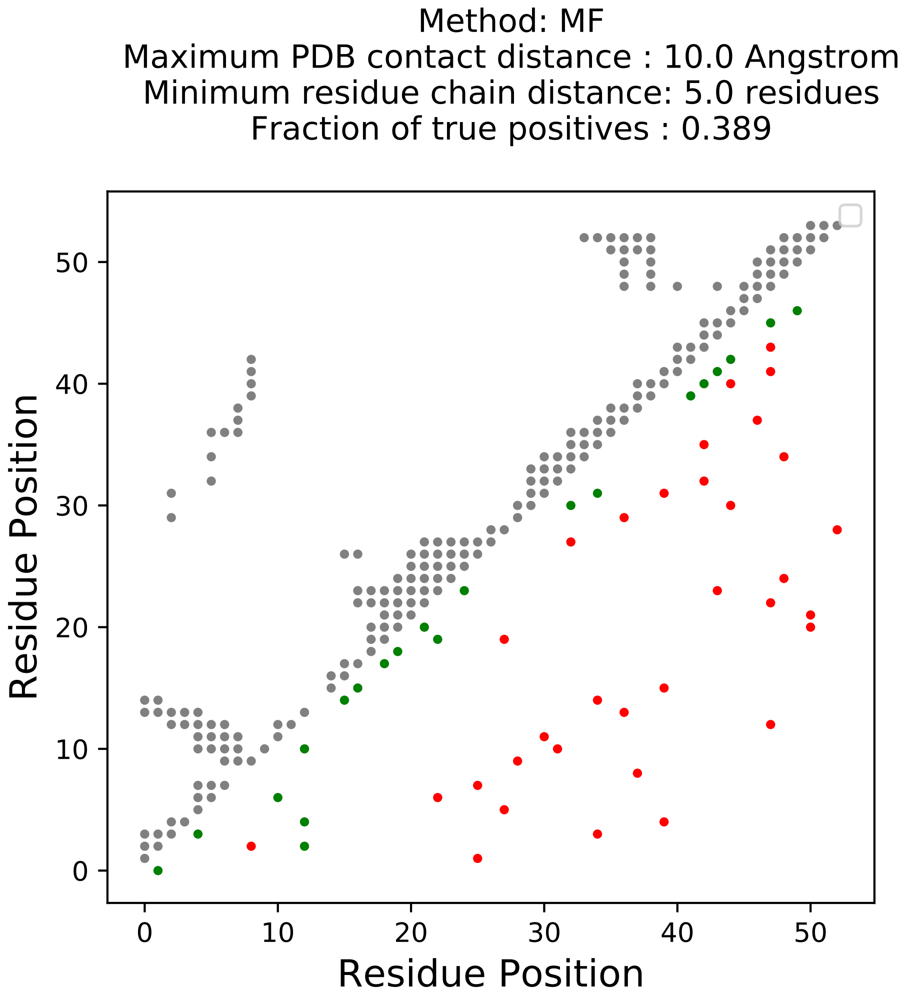
Method: MF
Maximum PDB contact distance : 10.0 Angstrom
Minimum residue chain distance: 5.0 residues
Fraction of true positives : 0.389
Residue Position
Residue Position
0
10
20
30
40
50
0
10
20
30
40
50

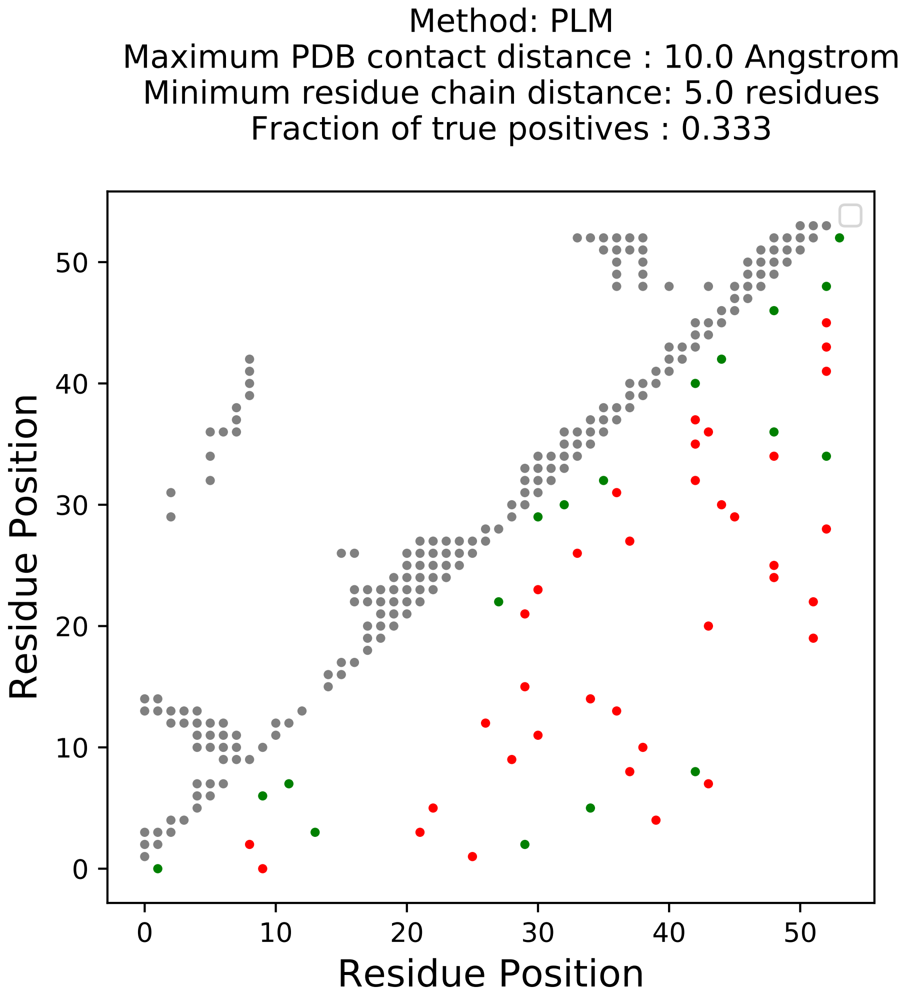
Method: PLM
Maximum PDB contact distance : 10.0 Angstrom
Minimum residue chain distance: 5.0 residues
Fraction of true positives : 0.333
Residue Position
Residue Position
0
10
20
30
40
50

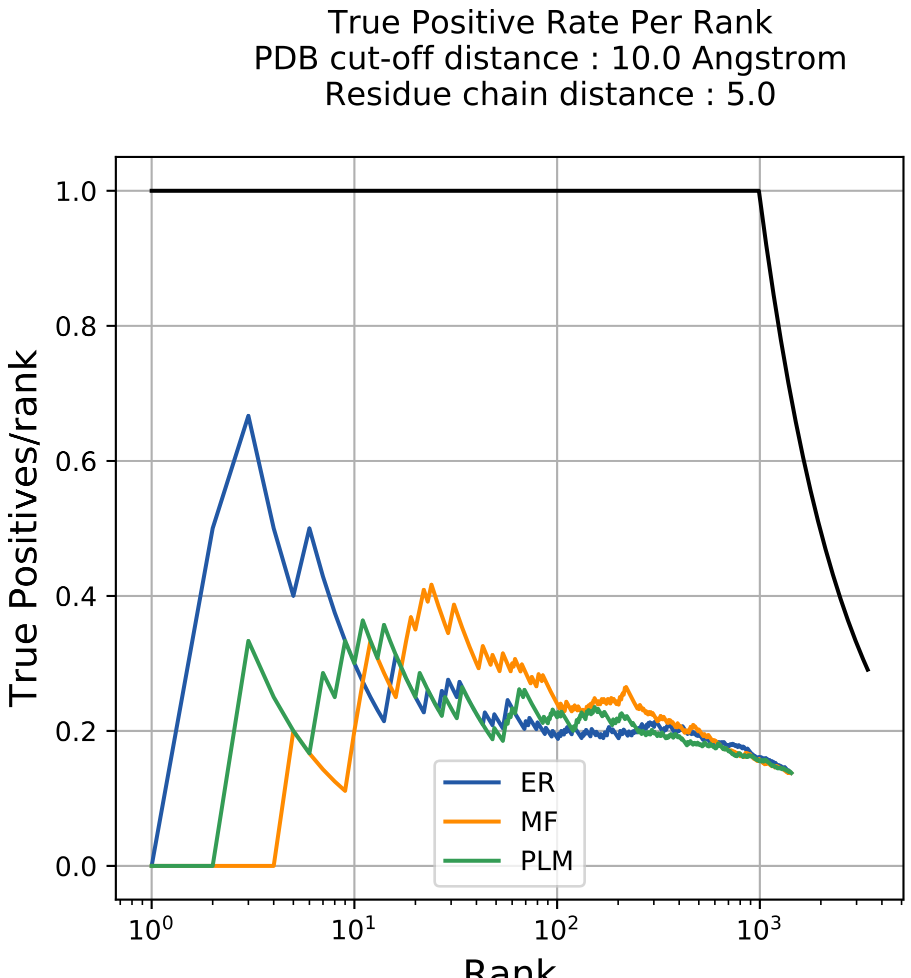

True Positive Rate Per Rank
PDB cut-off distance : 10.0 Angstrom
Residue chain distance : 5.0
True Positives/rank
1.0
0.8
0.6
0.4
0.2
0.0
$10^0$
$10^1$
$10^2$
$10^3$
Rank
ER
MF
PLM

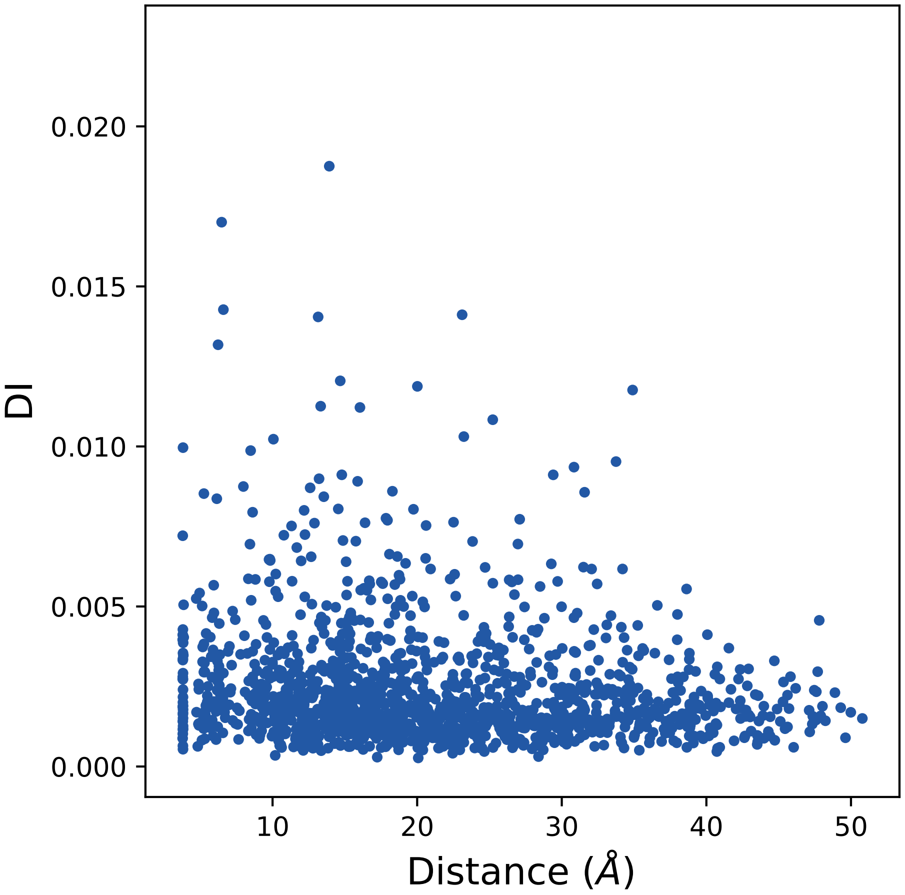

0.020
0.015
0.010
0.005
0.000
DI
10
20
30
40
50
Distance (Å)

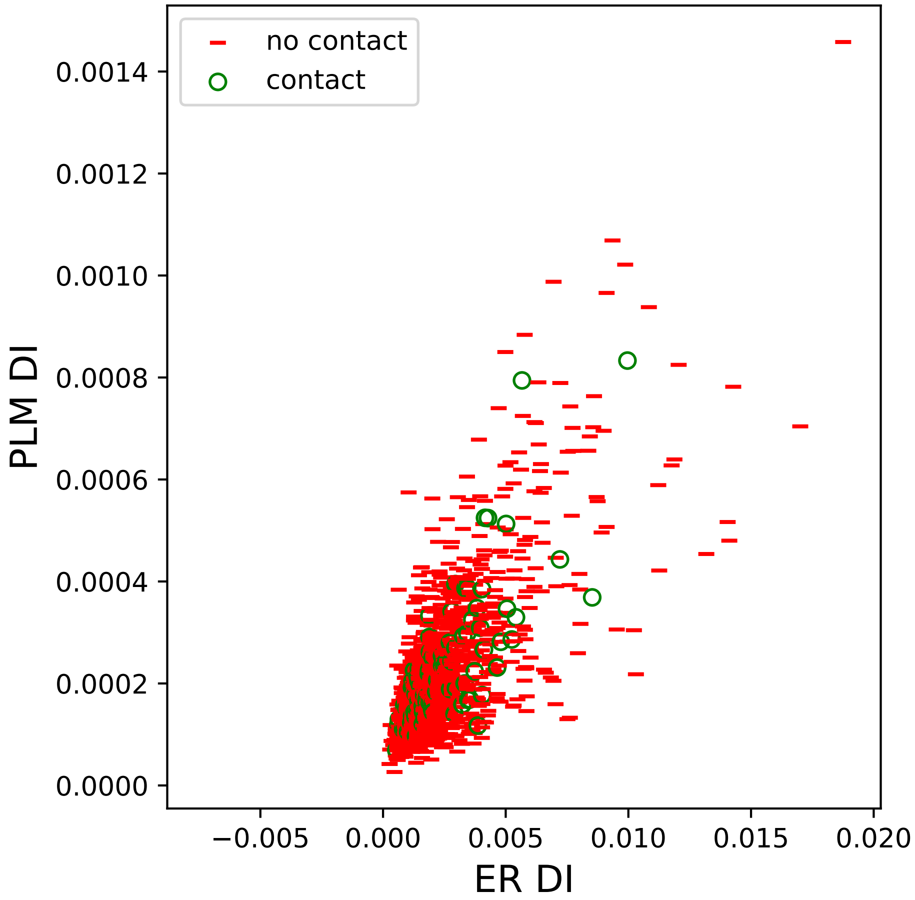

no contact
contact
0.0014
0.0012
0.0010
0.0008
0.0006
0.0004
0.0002
0.0000
PLM DI
−0.005
0.000
0.005
0.010
0.015
0.020
ER DI

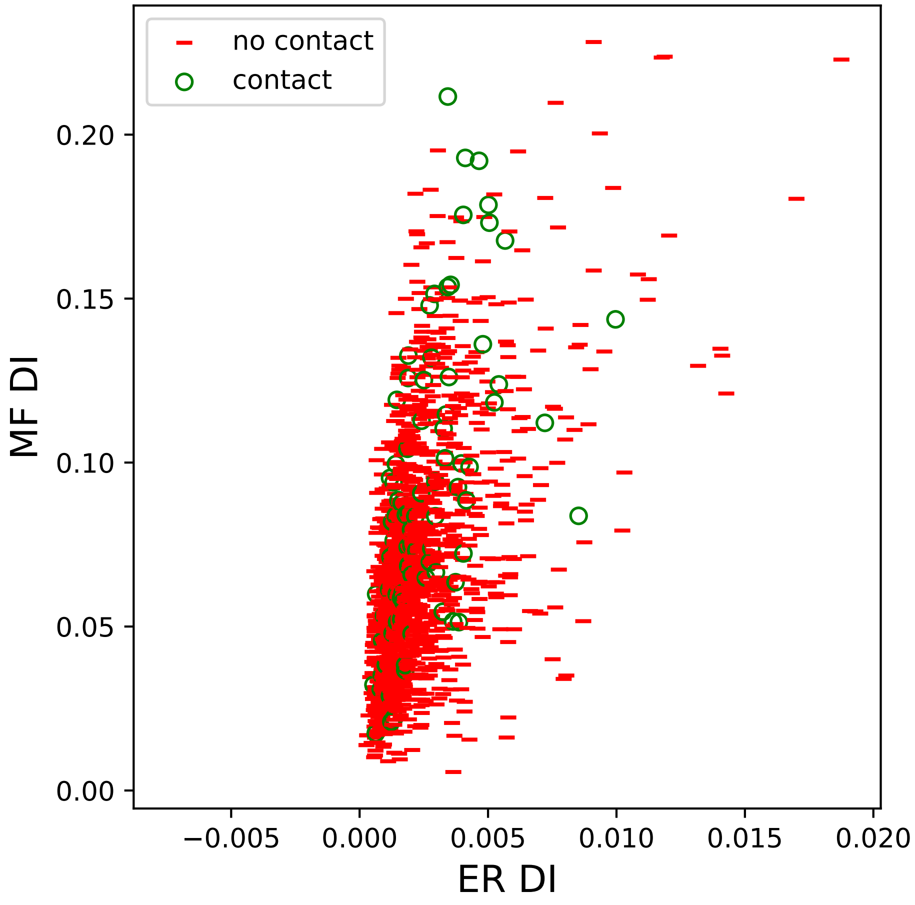

no contact
contact
0.20
0.15
0.10
0.05
0.00
MF DI
−0.005
0.000
0.005
0.010
0.015
0.020
ER DI

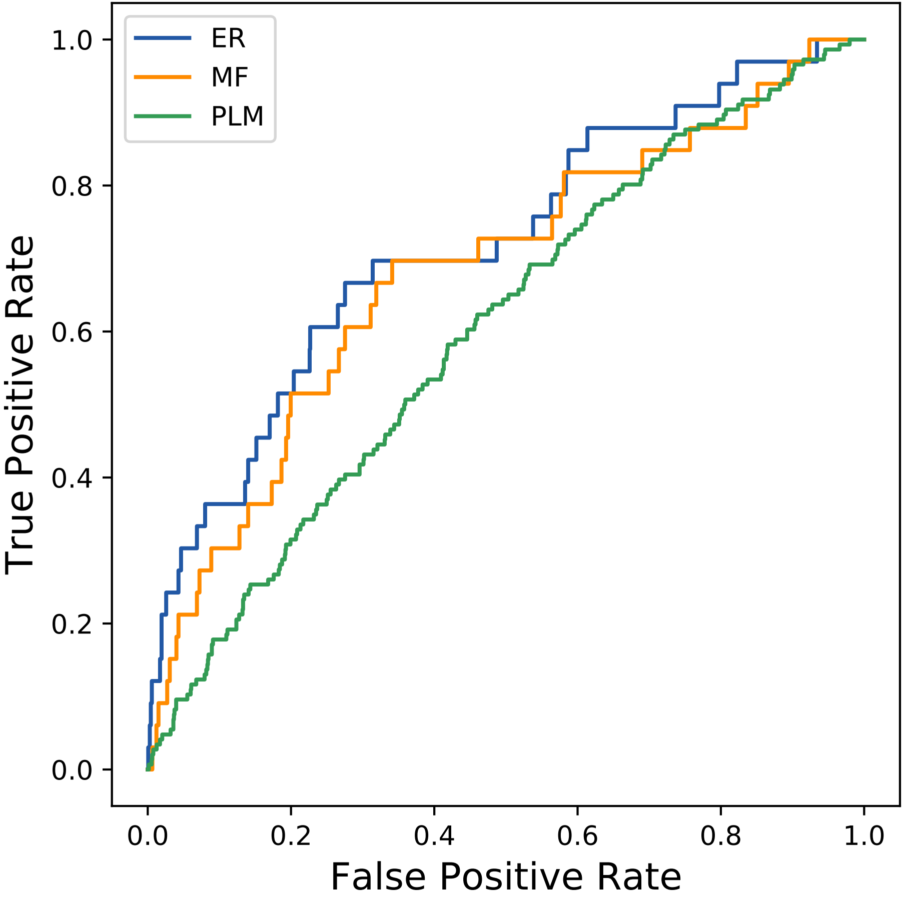

ER
MF
PLM
True Positive Rate
False Positive Rate
0.0
0.2
0.4
0.6
0.8
1.0
