## Supplemental Contact Maps for "Protein Structure Prediction with Expectation Reflection": 246seq_62col_5zu4_PF03318_plots.pdf

Method: ER
Maximum PDB contact distance : 10.0 Angstrom
Minimum residue chain distance: 5.0 residues
Fraction of true positives : 0.419

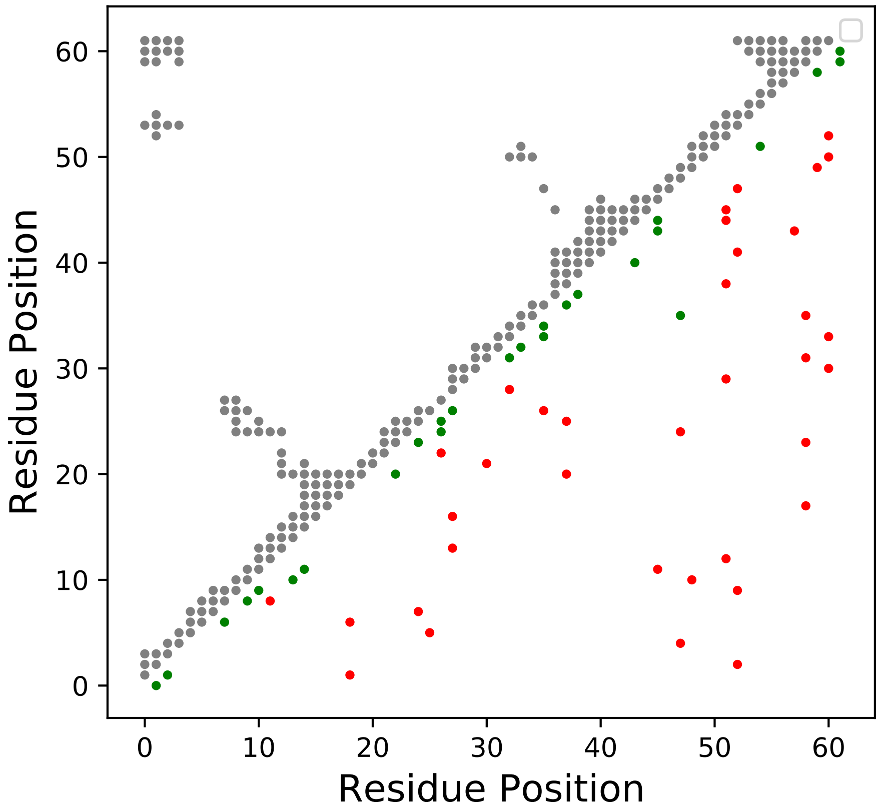

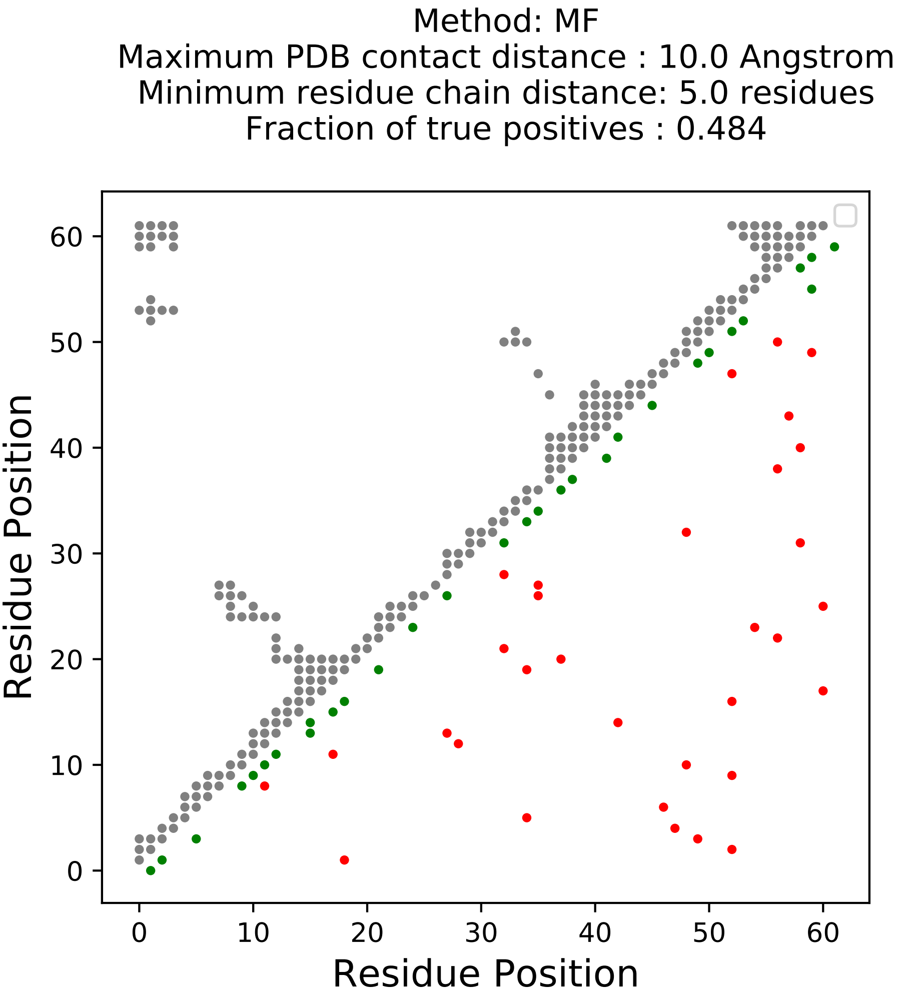
Method: MF
Maximum PDB contact distance : 10.0 Angstrom
Minimum residue chain distance: 5.0 residues
Fraction of true positives : 0.484
Residue Position
Residue Position
0
10
20
30
40
50
60

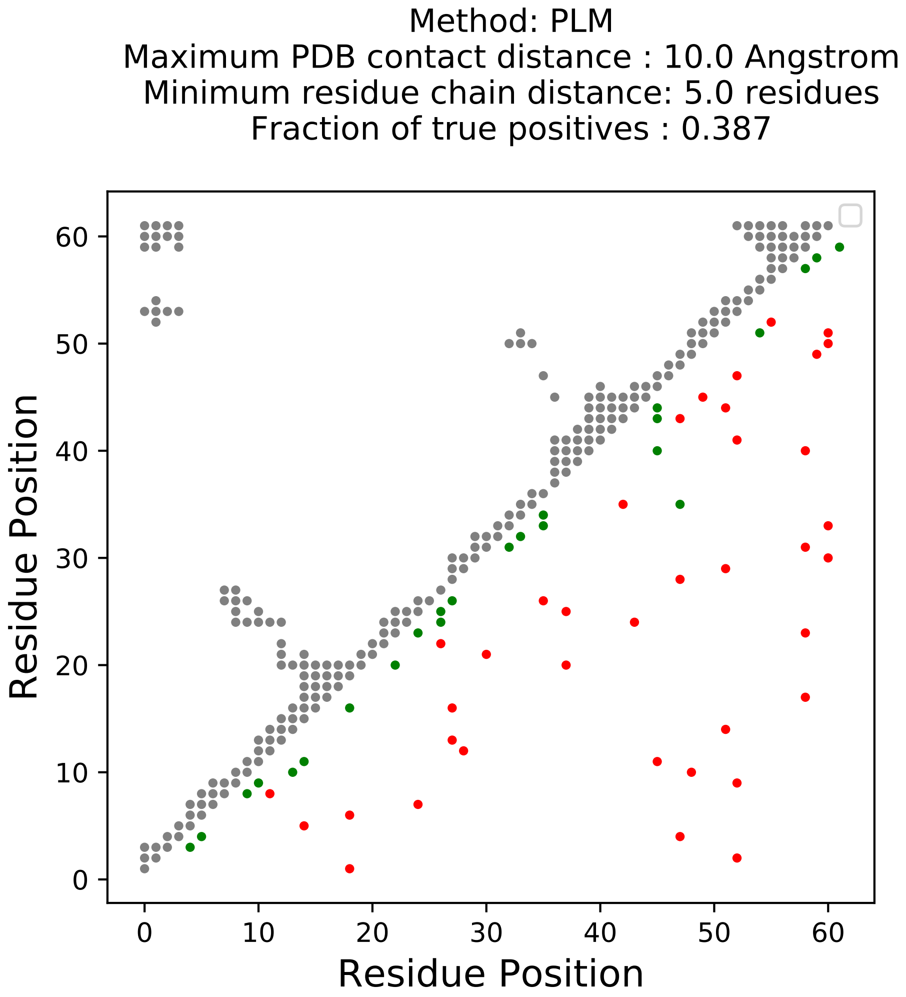
Method: PLM
Maximum PDB contact distance : 10.0 Angstrom
Minimum residue chain distance: 5.0 residues
Fraction of true positives : 0.387
Residue Position
Residue Position
0
10
20
30
40
50
60

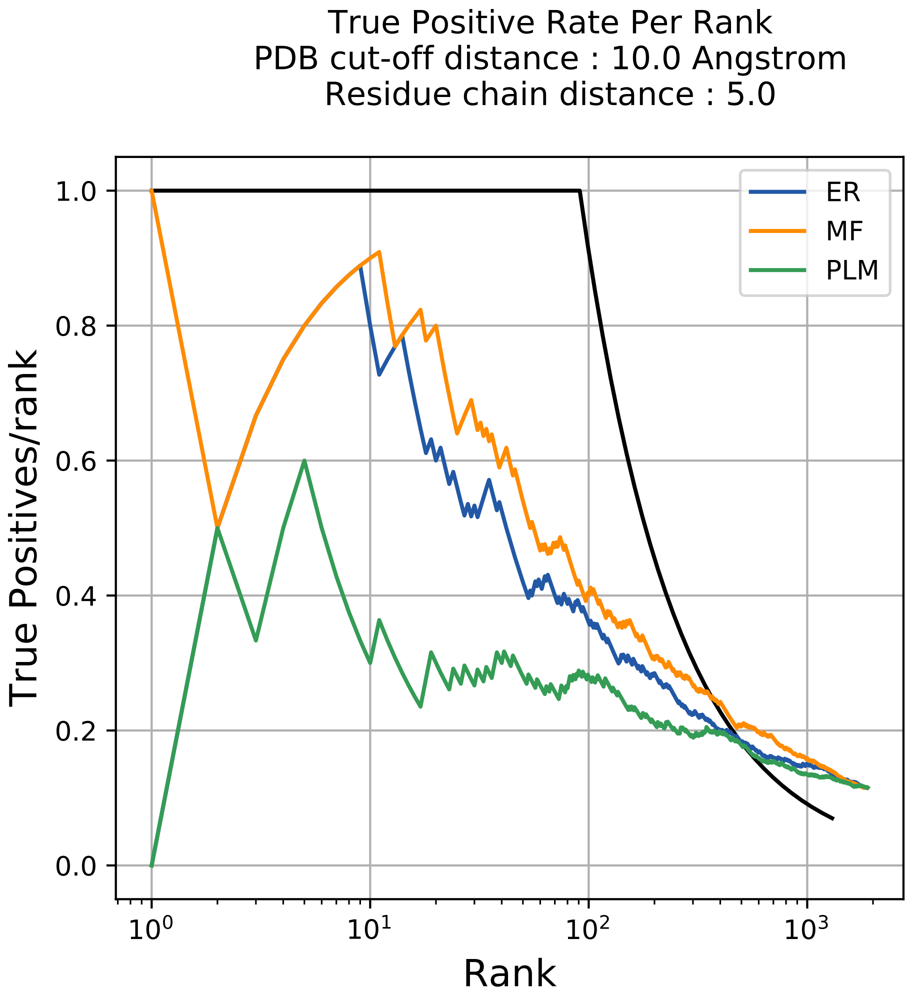
True Positive Rate Per Rank
PDB cut-off distance : 10.0 Angstrom
Residue chain distance : 5.0
True Positives/rank
Rank
ER
MF
PLM
1.0
0.8
0.6
0.4
0.2
0.0
$10^0$
$10^1$
$10^2$
$10^3$

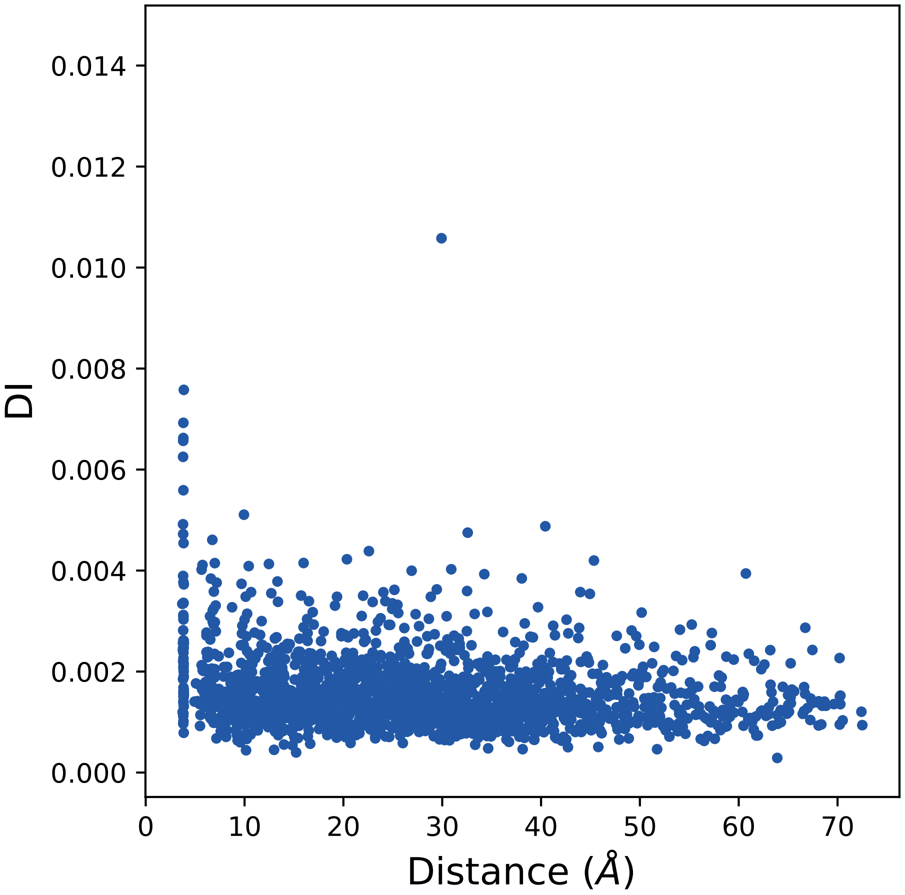

DI
Distance (Å)
0.000
0.002
0.004
0.006
0.008
0.010
0.012
0.014
0
10
20
30
40
50
60
70

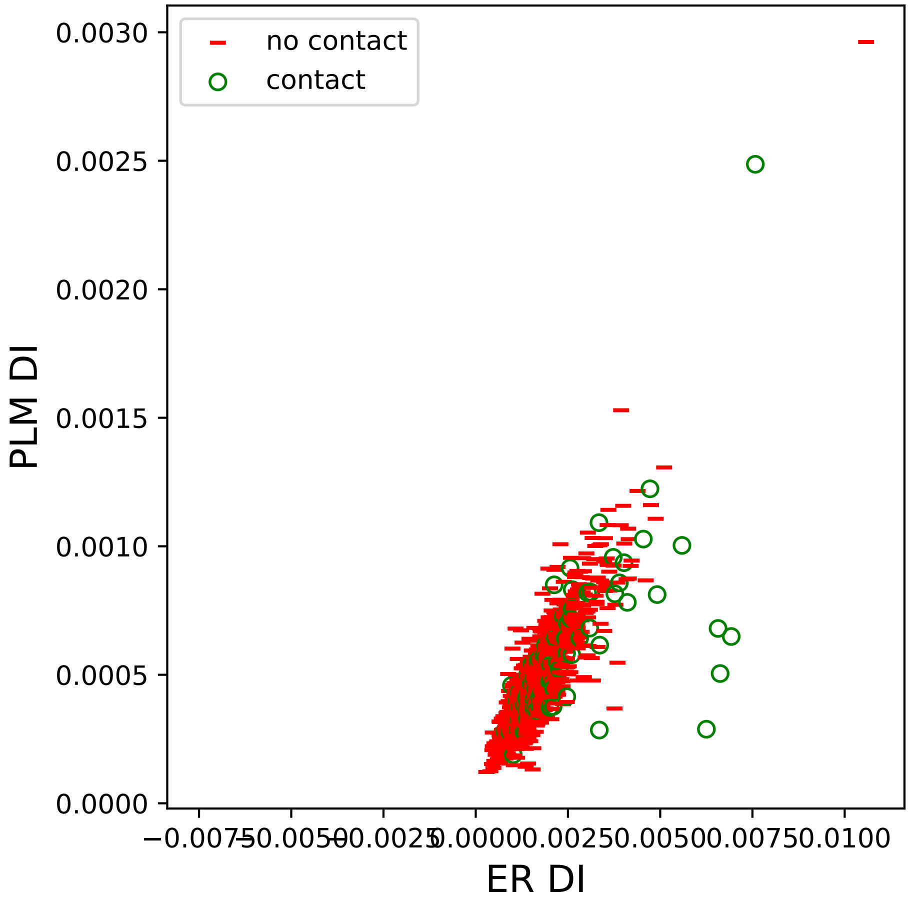

no contact
contact
0.0030
0.0025
0.0020
0.0015
0.0010
0.0005
0.0000
PLM DI
−0.0075
−0.0050
−0.0025
0.0000
0.0025
0.0050
0.0075
0.0100
ER DI

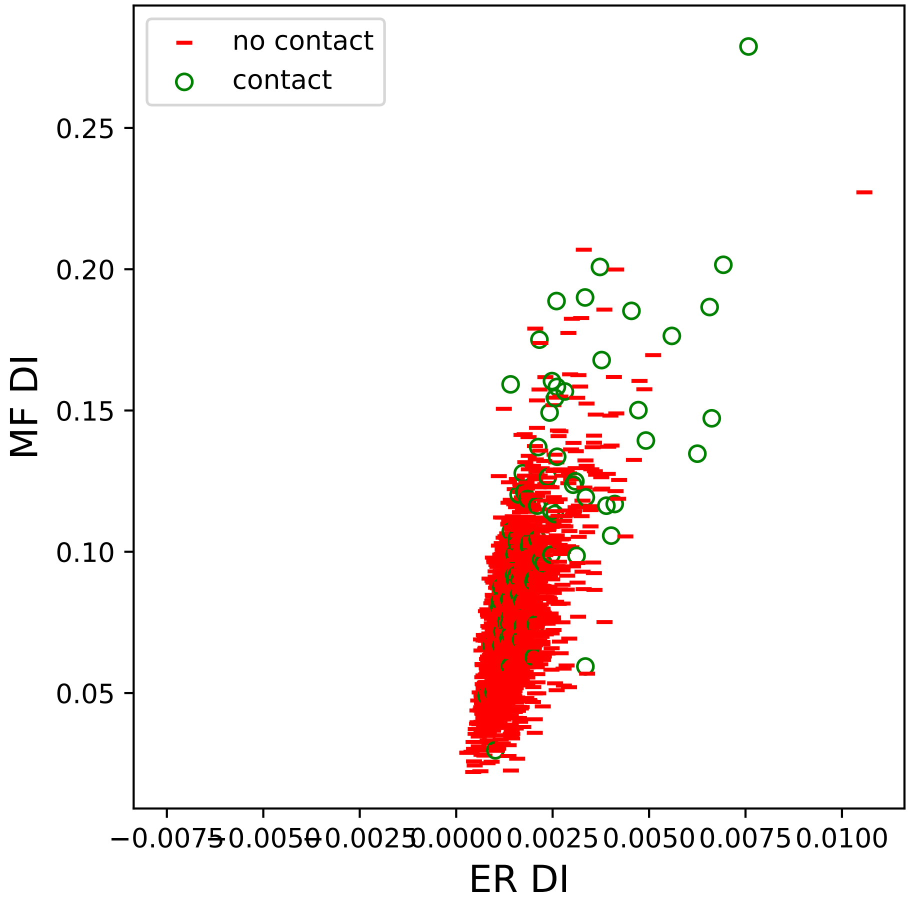
no contact
contact
0.25
0.20
0.15
0.10
0.05
MF DI
−0.0075
−0.0050
−0.0025
0.0000
0.0025
0.0050
0.0075
0.0100
ER DI

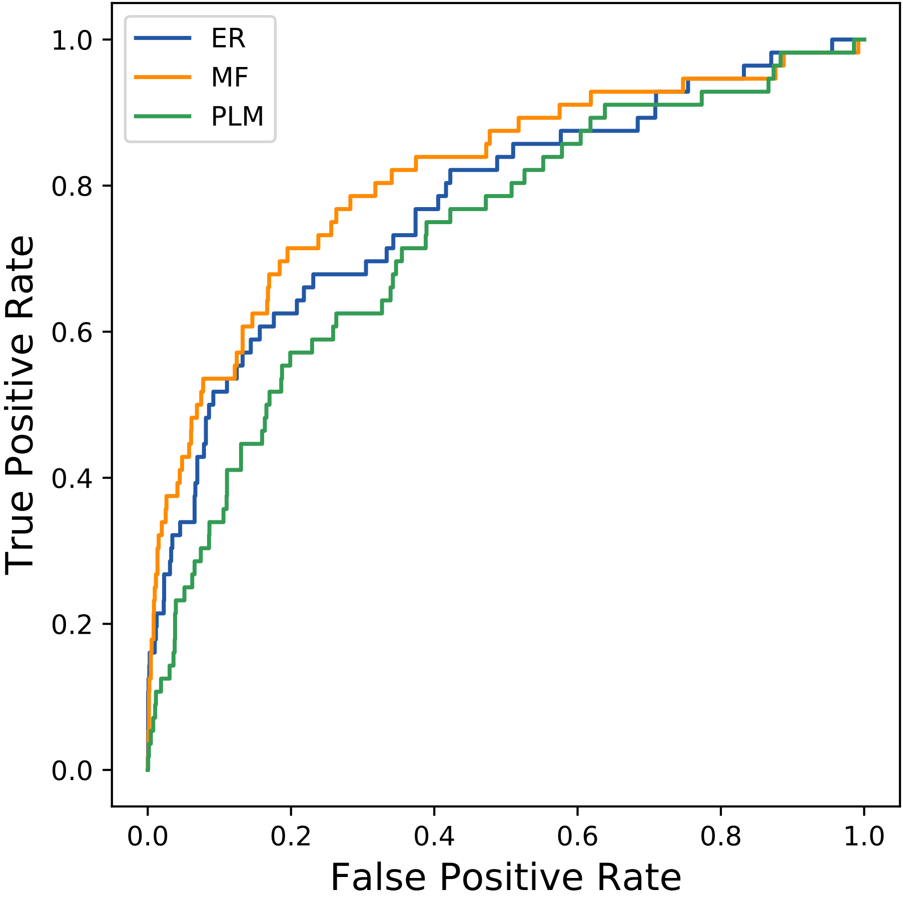
ER
MF
PLM
1.0
0.8
0.6
0.4
0.2
0.0
True Positive Rate
0.0
0.2
0.4
0.6
0.8
1.0
False Positive Rate
