## Supplemental Contact Maps for "Protein Structure Prediction with Expectation Reflection": 278seq_94col_6a2u_PF12318_plots.pdf

Method: ER
Maximum PDB contact distance : 10.0 Angstrom
Minimum residue chain distance: 5.0 residues
Fraction of true positives : 0.351

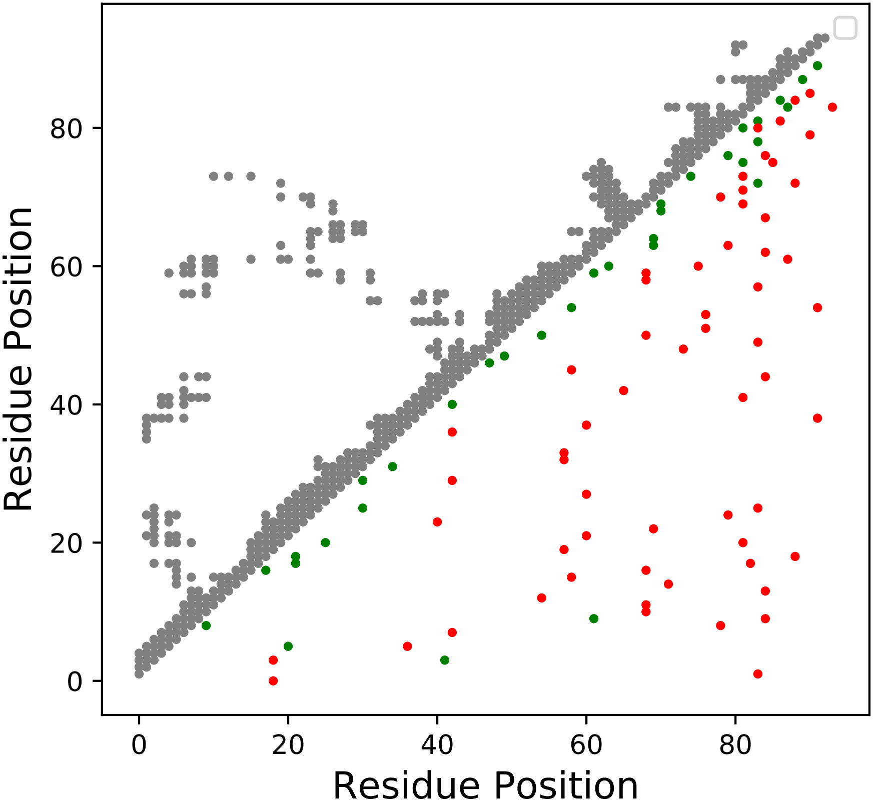

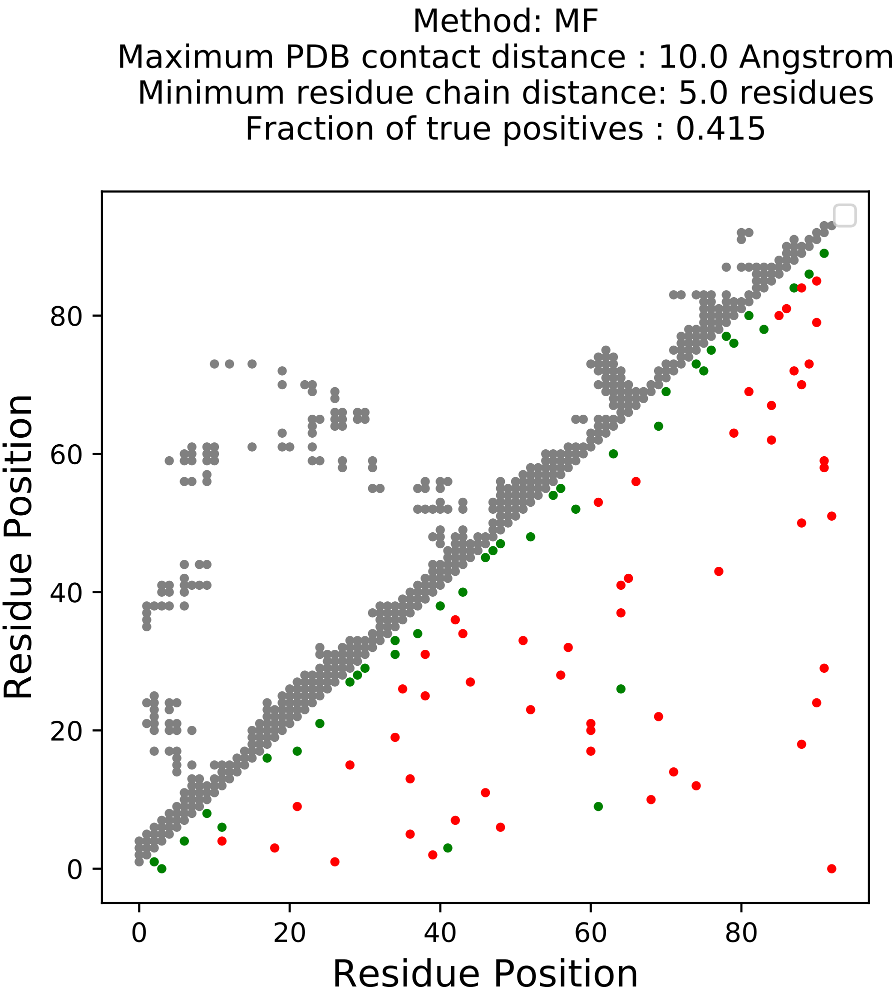
Method: MF
Maximum PDB contact distance : 10.0 Angstrom
Minimum residue chain distance: 5.0 residues
Fraction of true positives : 0.415
Residue Position
Residue Position
0
20
40
60
80
0
20
40
60
80

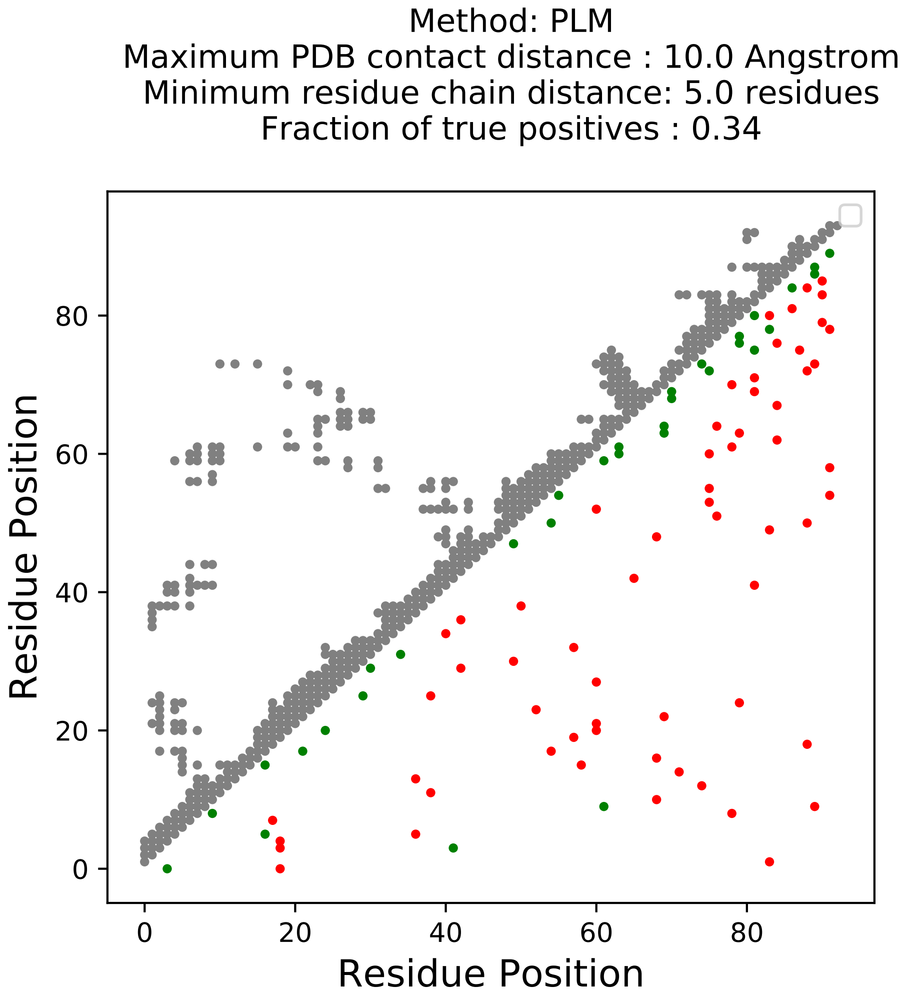
Method: PLM
Maximum PDB contact distance : 10.0 Angstrom
Minimum residue chain distance: 5.0 residues
Fraction of true positives : 0.34
Residue Position
Residue Position
0
20
40
60
80
0
20
40
60
80

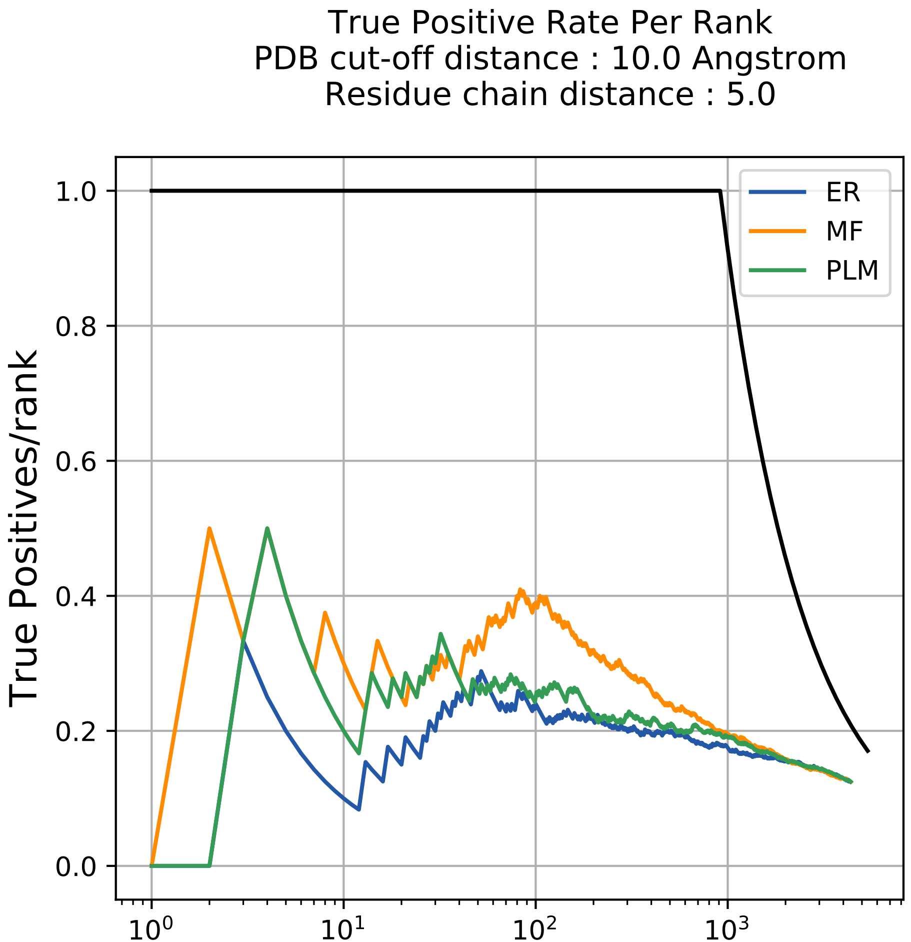

True Positive Rate Per Rank
PDB cut-off distance : 10.0 Angstrom
Residue chain distance : 5.0
True Positives/rank
Rank
ER
MF
PLM
1.0
0.8
0.6
0.4
0.2
0.0
$10^0$
$10^1$
$10^2$
$10^3$

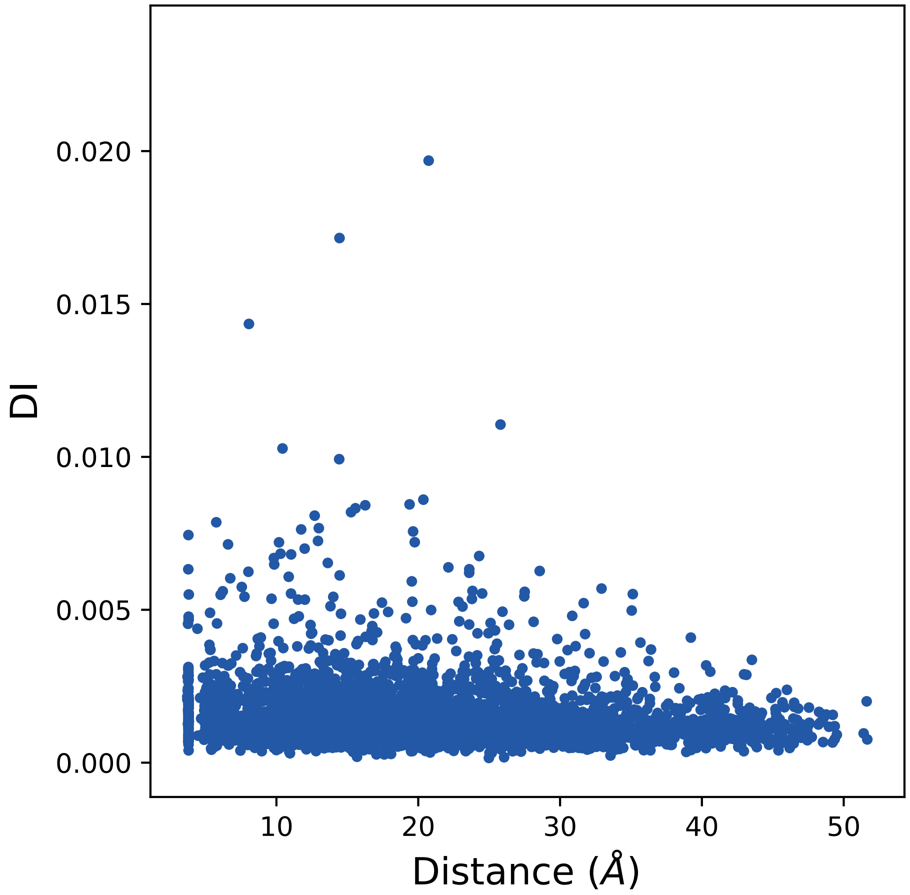
0.020
0.015
0.010
0.005
0.000
DI
10
20
30
40
50
Distance (Å)

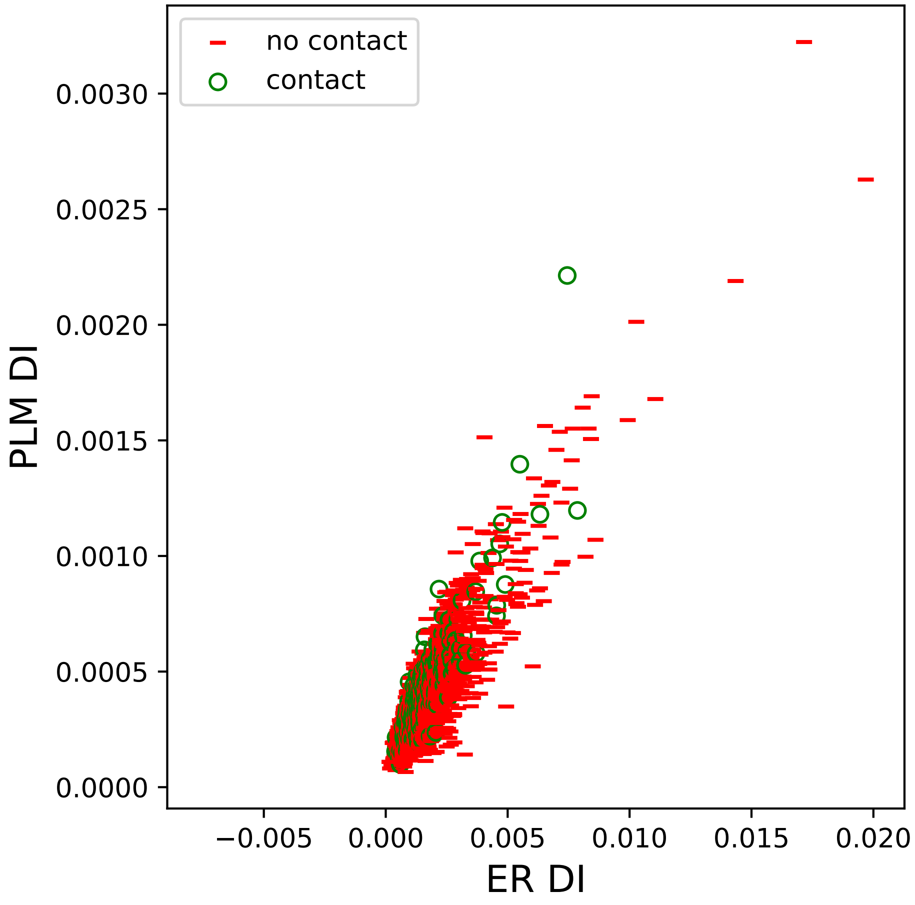

no contact
contact
PLM DI
ER DI
0.0000
0.0005
0.0010
0.0015
0.0020
0.0025
0.0030
−0.005
0.000
0.005
0.010
0.015
0.020

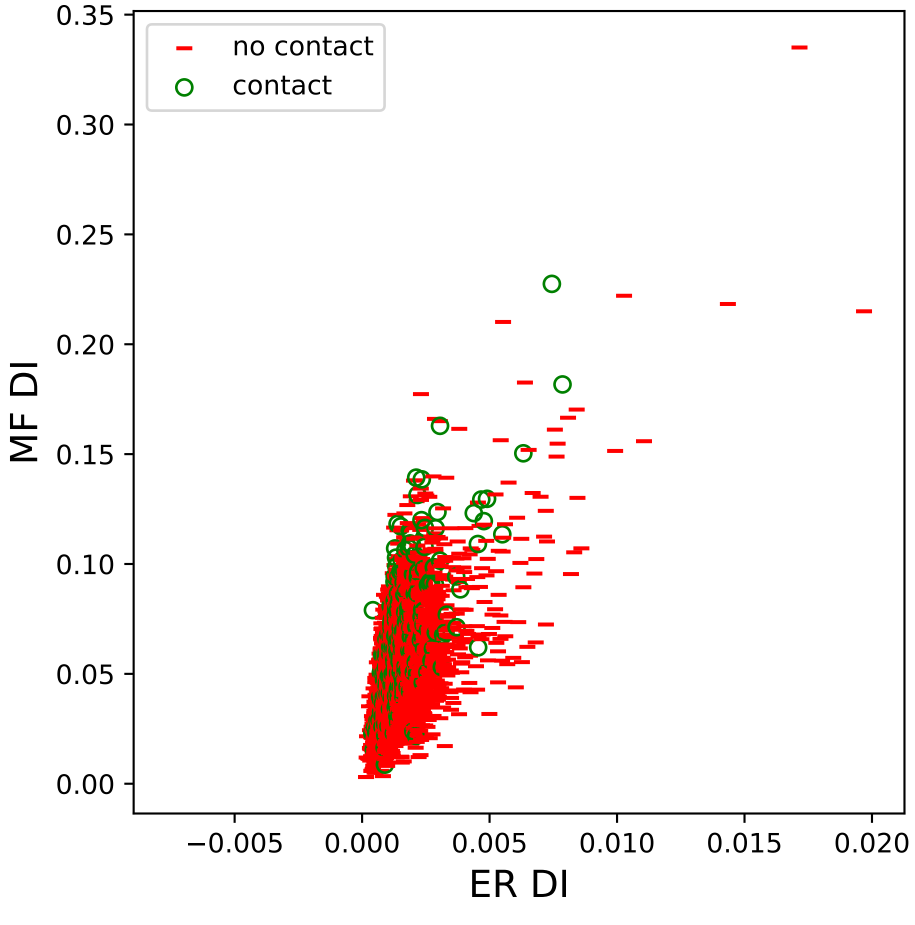

no contact
contact
MF DI
ER DI
0.00
0.05
0.10
0.15
0.20
0.25
0.30
0.35
−0.005
0.000
0.005
0.010
0.015
0.020

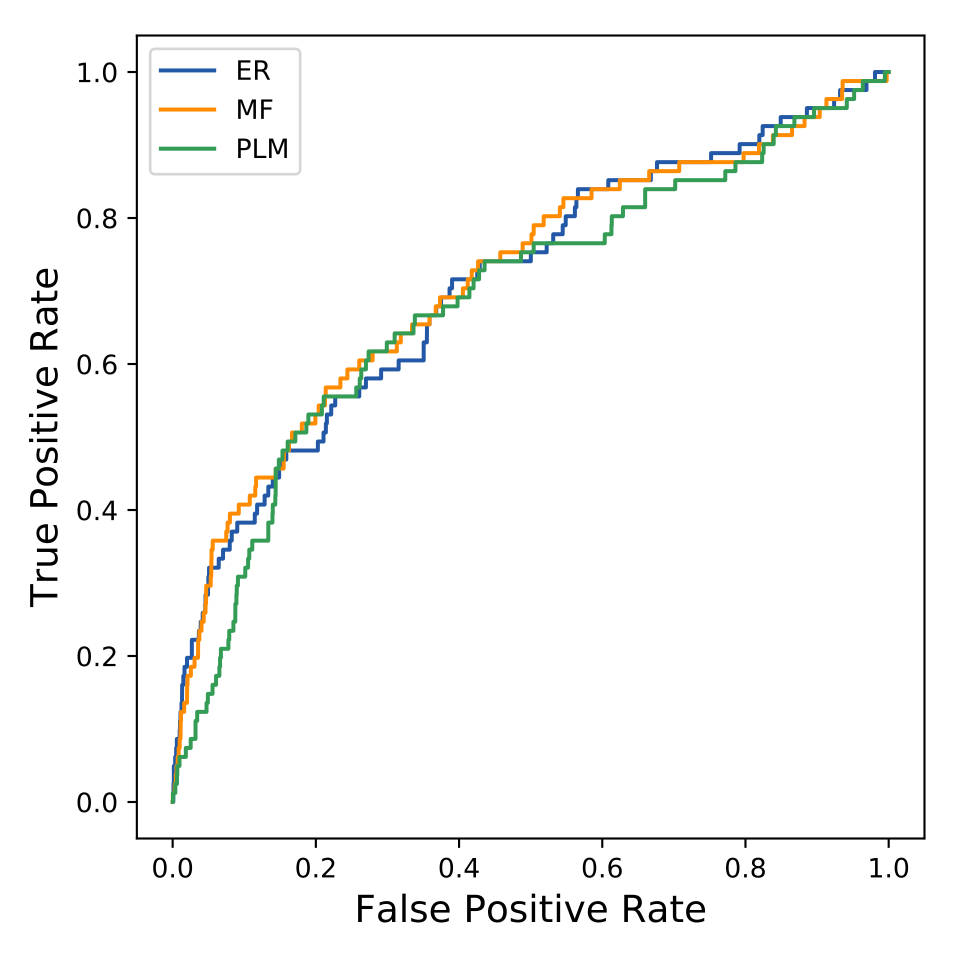
ER
MF
PLM
True Positive Rate
False Positive Rate
0.0
0.2
0.4
0.6
0.8
1.0
0.0
0.2
0.4
0.6
0.8
1.0
