## Supplemental Contact Maps for "Protein Structure Prediction with Expectation Reflection": 393seq_85col_1j0s_PF00340_plots.pdf

Method: ER
Maximum PDB contact distance : 10.0 Angstrom
Minimum residue chain distance: 5.0 residues
Fraction of true positives : 0.424

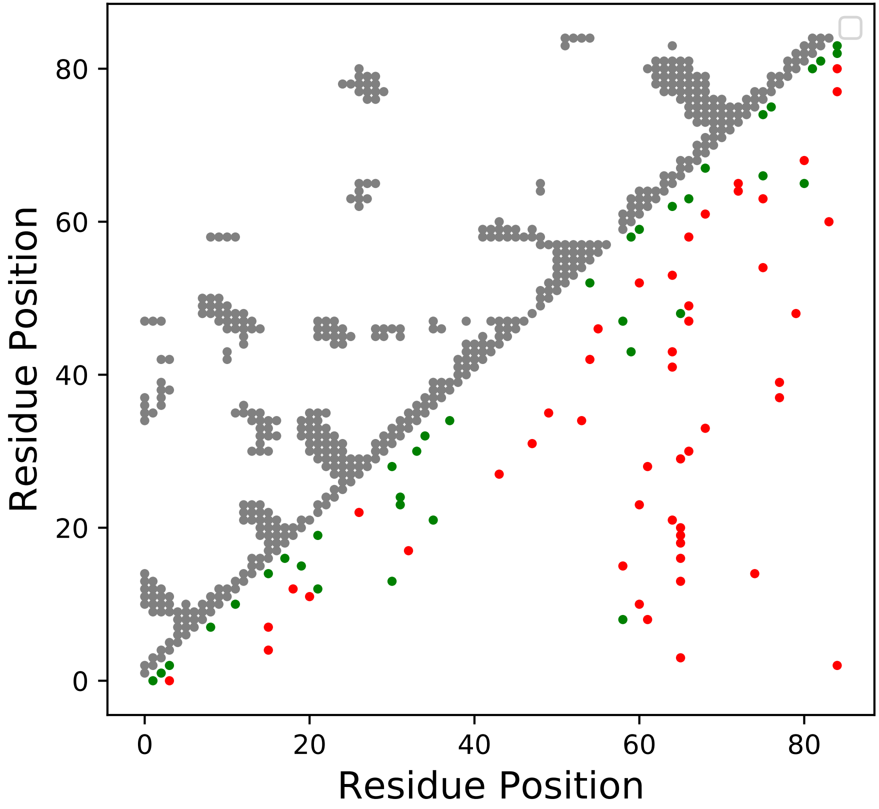

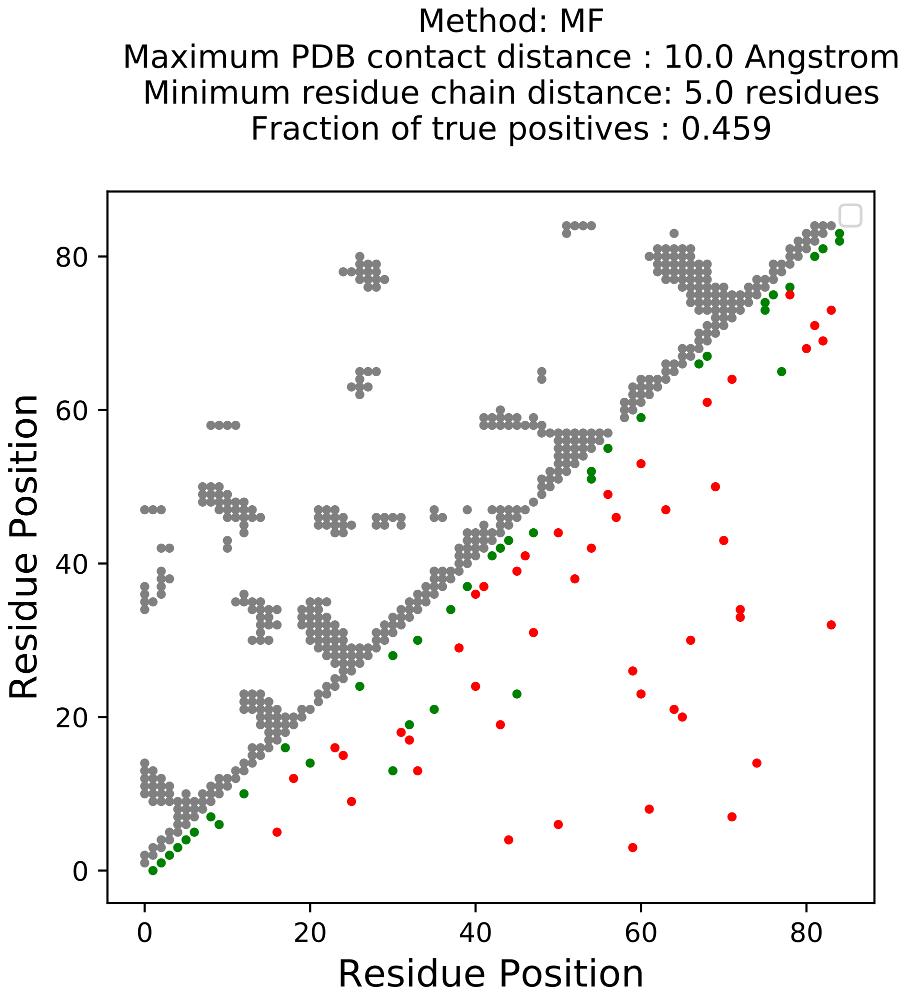
Method: MF
Maximum PDB contact distance : 10.0 Angstrom
Minimum residue chain distance: 5.0 residues
Fraction of true positives : 0.459
Residue Position
Residue Position
0
20
40
60
80
0
20
40
60
80

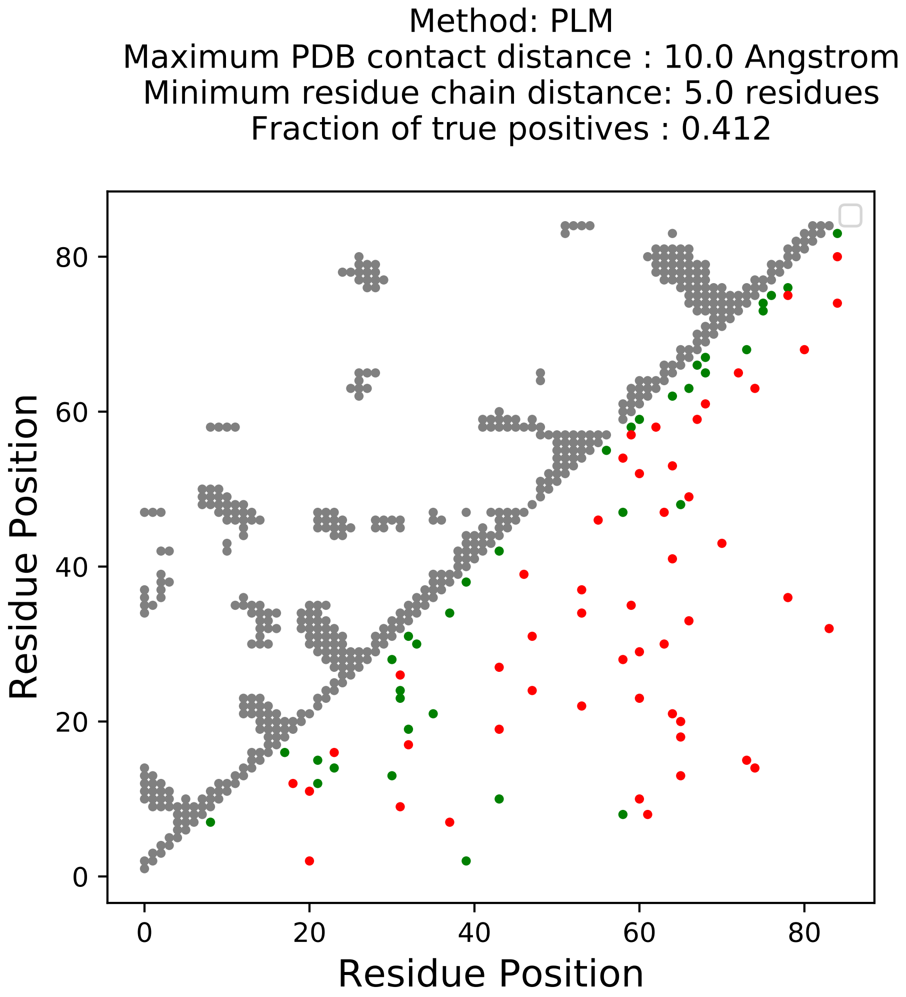

Method: PLM
Maximum PDB contact distance : 10.0 Angstrom
Minimum residue chain distance: 5.0 residues
Fraction of true positives : 0.412
Residue Position
Residue Position
0
20
40
60
80

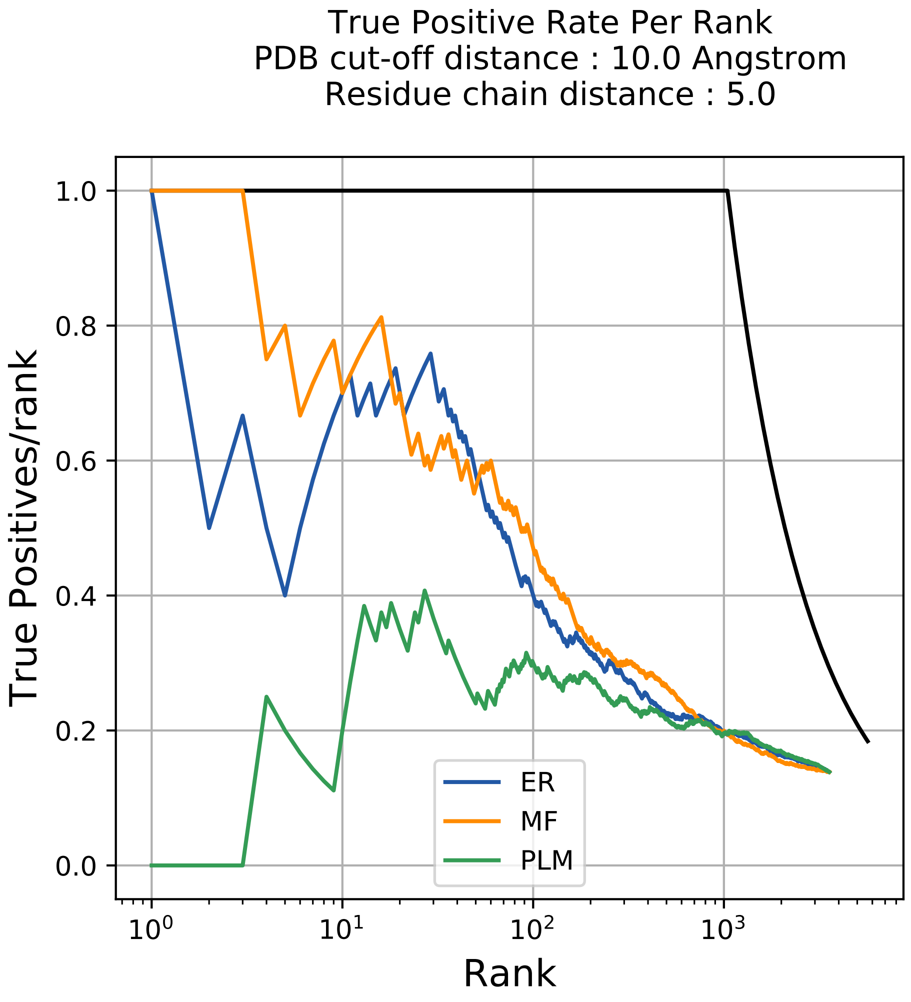
True Positive Rate Per Rank
PDB cut-off distance : 10.0 Angstrom
Residue chain distance : 5.0
True Positives/rank
Rank
1.0
0.8
0.6
0.4
0.2
0.0
$10^0$
$10^1$
$10^2$
$10^3$
ER
MF
PLM

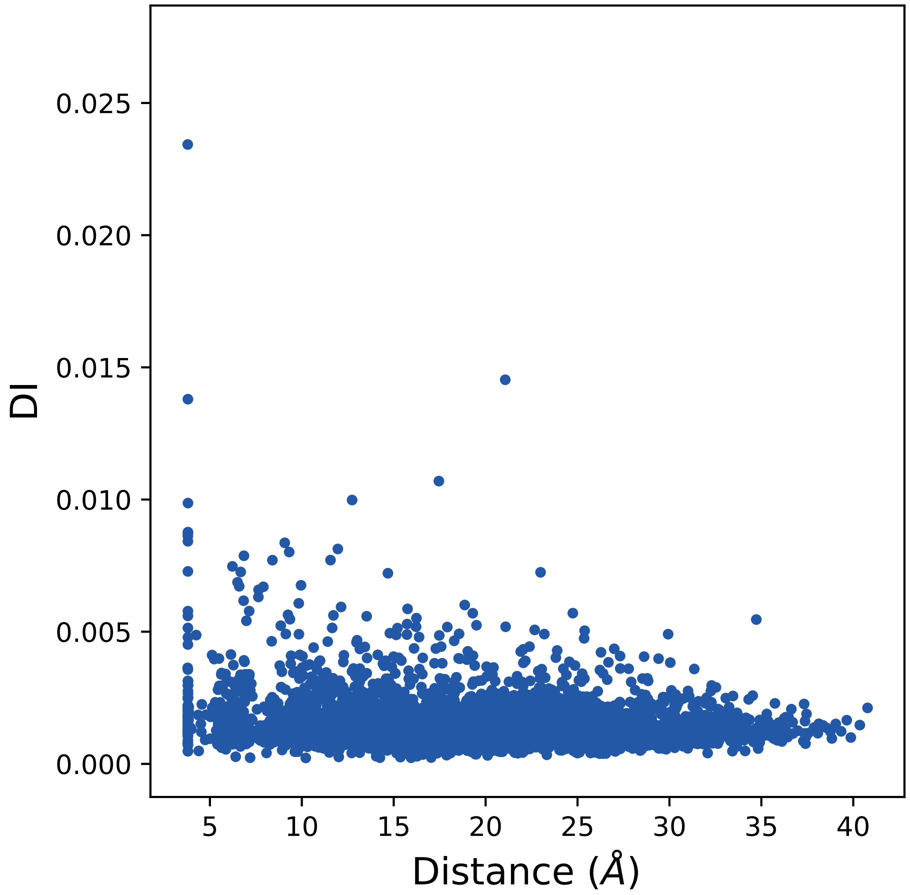

0.025
0.020
0.015
0.010
0.005
0.000
DI
5
10
15
20
25
30
35
40
Distance (Å)

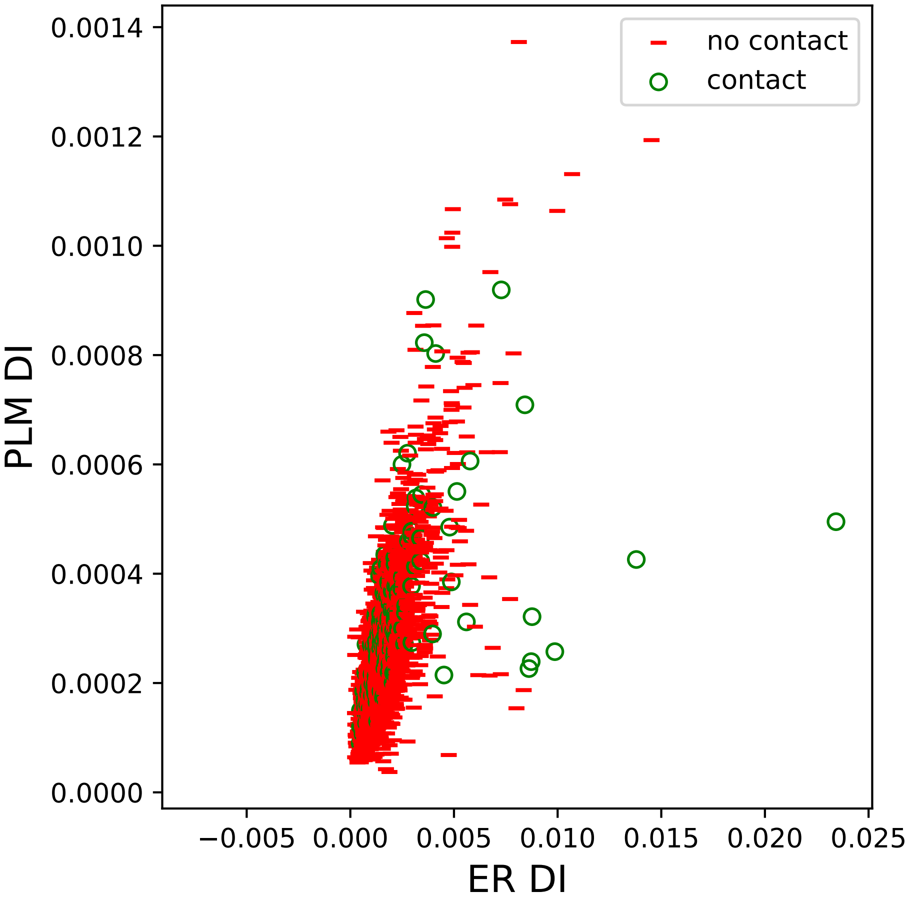

no contact
contact
PLM DI
ER DI
0.0014
0.0012
0.0010
0.0008
0.0006
0.0004
0.0002
0.0000
−0.005
0.000
0.005
0.010
0.015
0.020
0.025

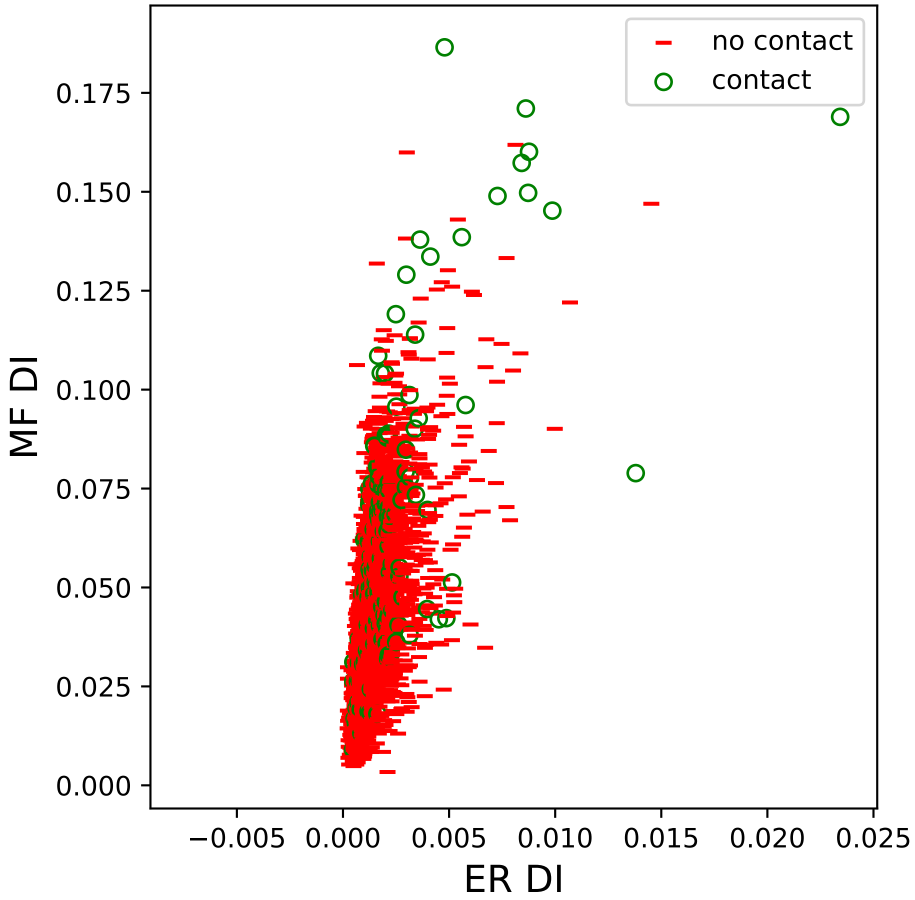

no contact
contact
MF DI
ER DI
0.000
0.025
0.050
0.075
0.100
0.125
0.150
0.175
−0.005
0.000
0.005
0.010
0.015
0.020
0.025

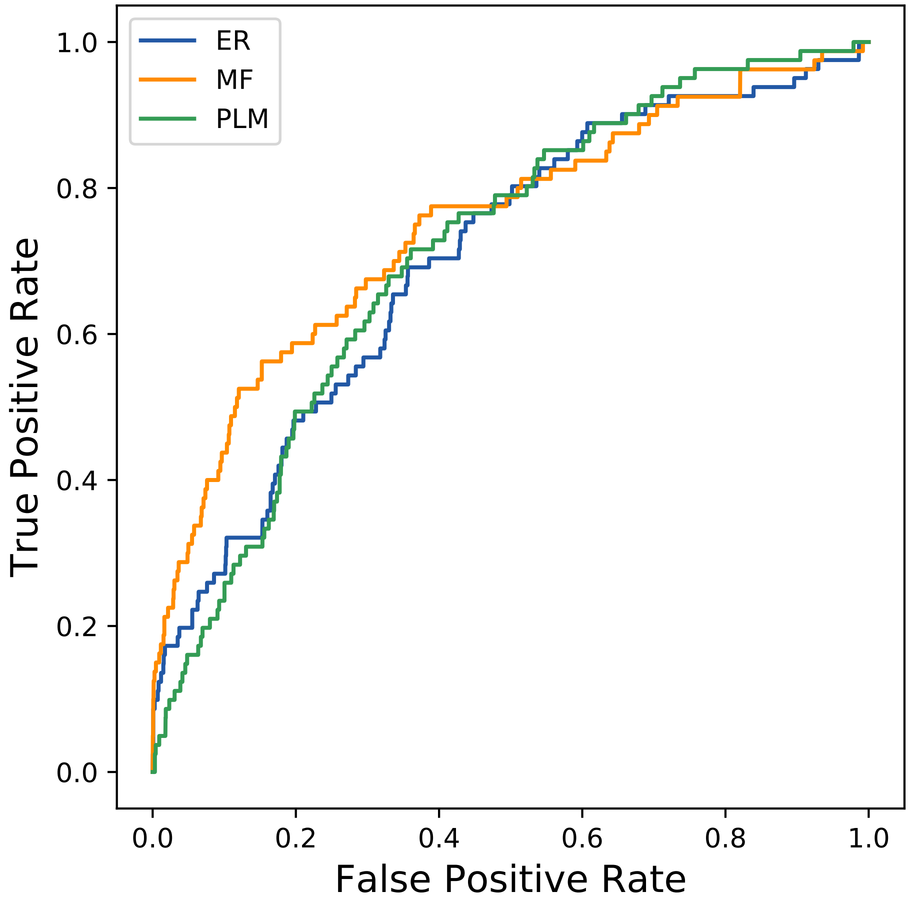

ER
MF
PLM
1.0
0.8
0.6
0.4
0.2
0.0
True Positive Rate
0.0
0.2
0.4
0.6
0.8
1.0
False Positive Rate
