## Supplemental Contact Maps for "Protein Structure Prediction with Expectation Reflection": 400seq_80col_2j0b_PF02434_plots.pdf

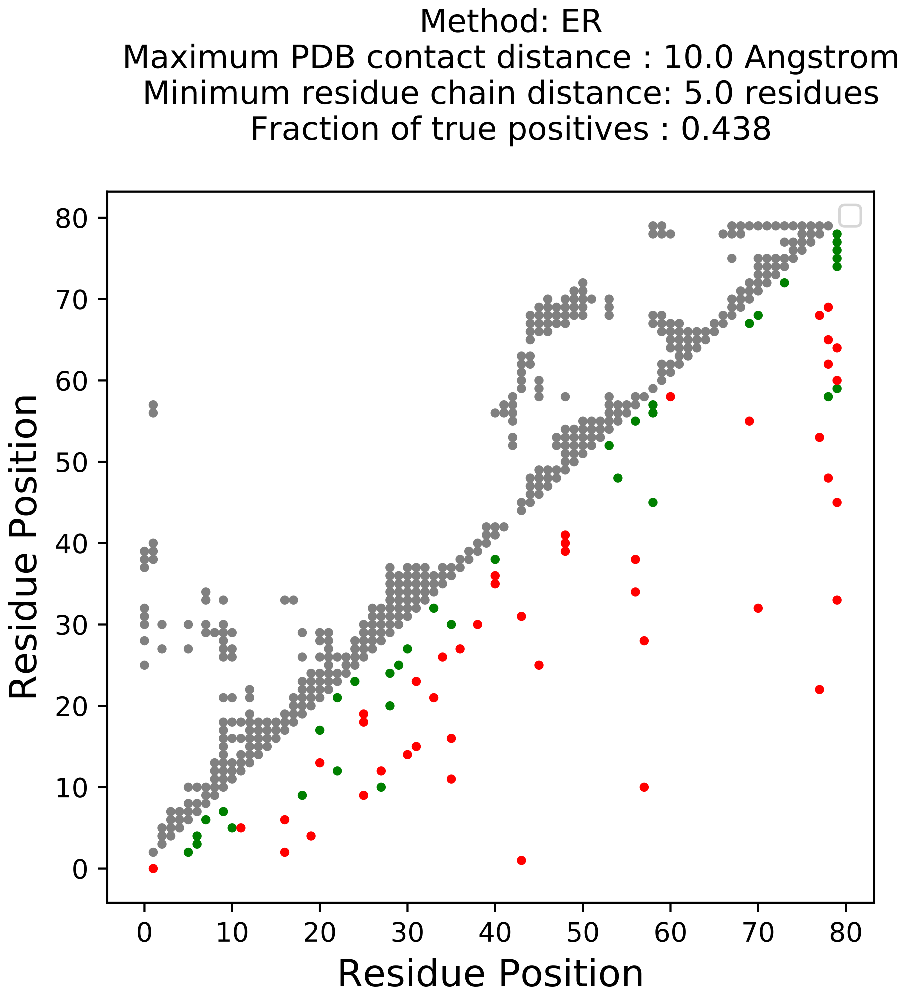

Method: ER
Maximum PDB contact distance : 10.0 Angstrom
Minimum residue chain distance: 5.0 residues
Fraction of true positives : 0.438
Residue Position
Residue Position
0
10
20
30
40
50
60
70
80

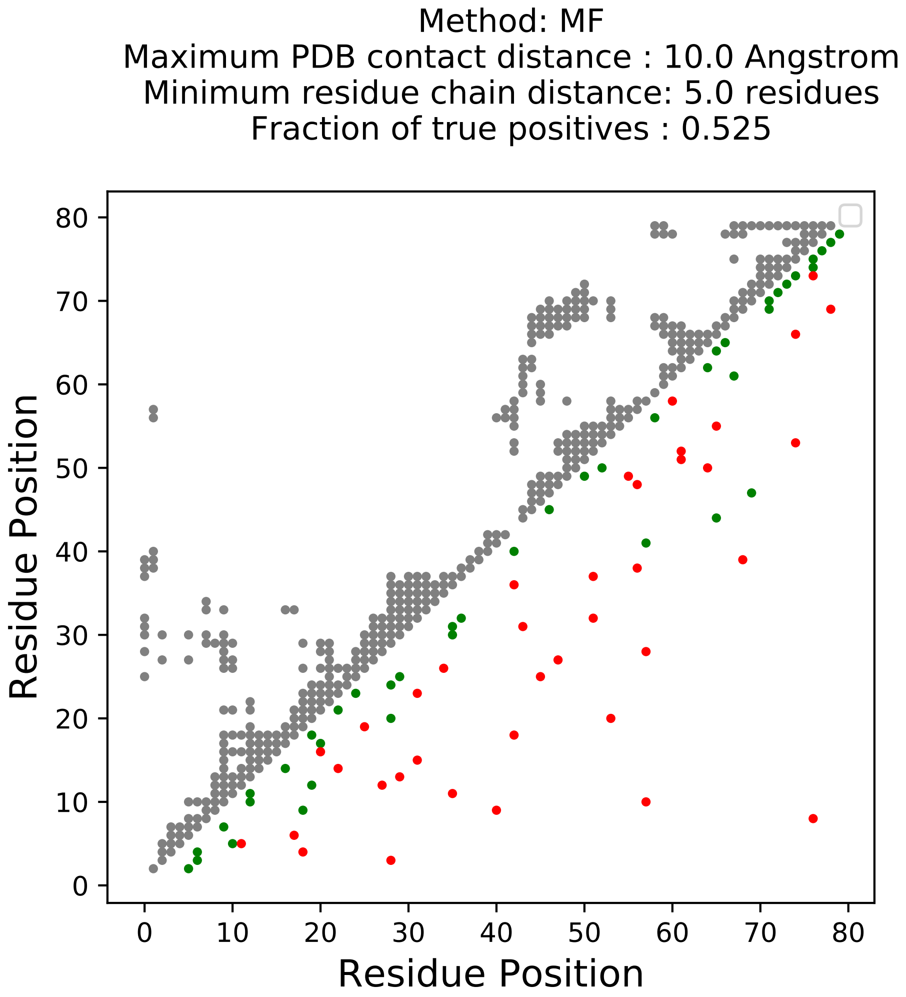
Method: MF
Maximum PDB contact distance : 10.0 Angstrom
Minimum residue chain distance: 5.0 residues
Fraction of true positives : 0.525
Residue Position
Residue Position
0
10
20
30
40
50
60
70
80

Method: PLM
Maximum PDB contact distance : 10.0 Angstrom
Minimum residue chain distance: 5.0 residues
Fraction of true positives : 0.4

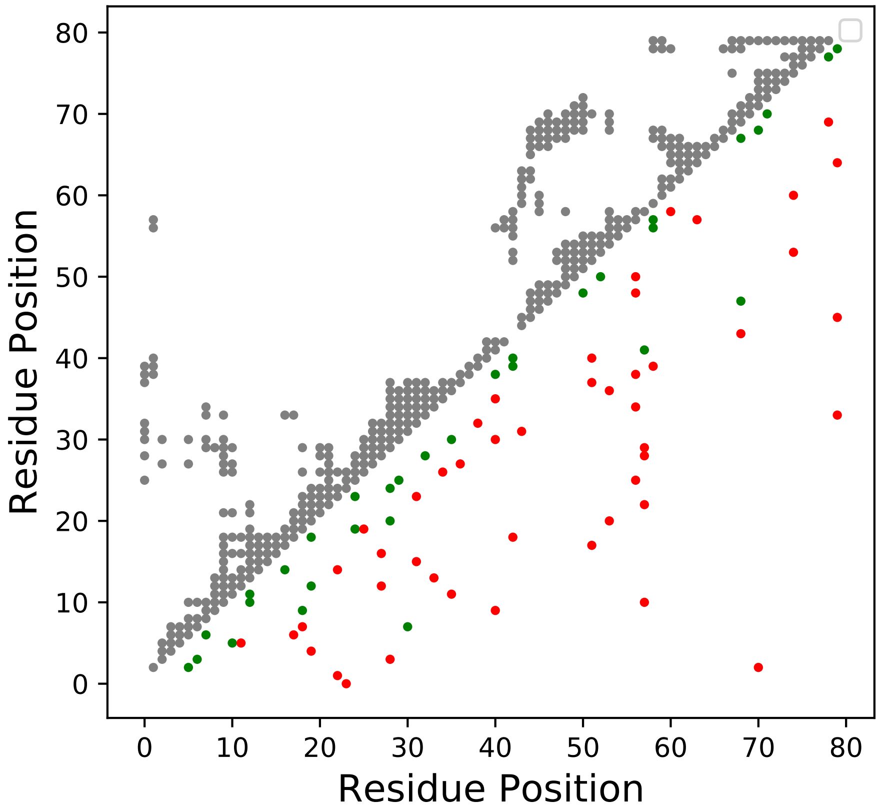

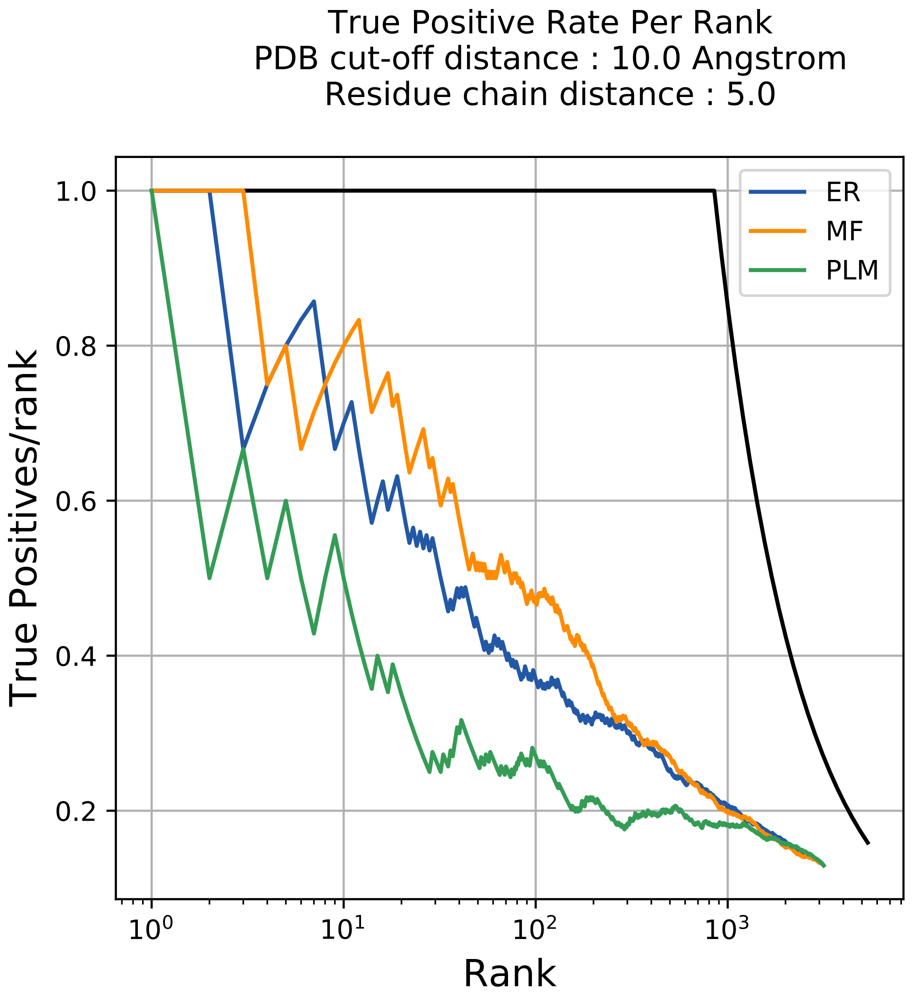

True Positive Rate Per Rank
PDB cut-off distance : 10.0 Angstrom
Residue chain distance : 5.0
True Positives/rank
Rank
1.0
0.8
0.6
0.4
0.2
$10^0$
$10^1$
$10^2$
$10^3$
ER
MF
PLM

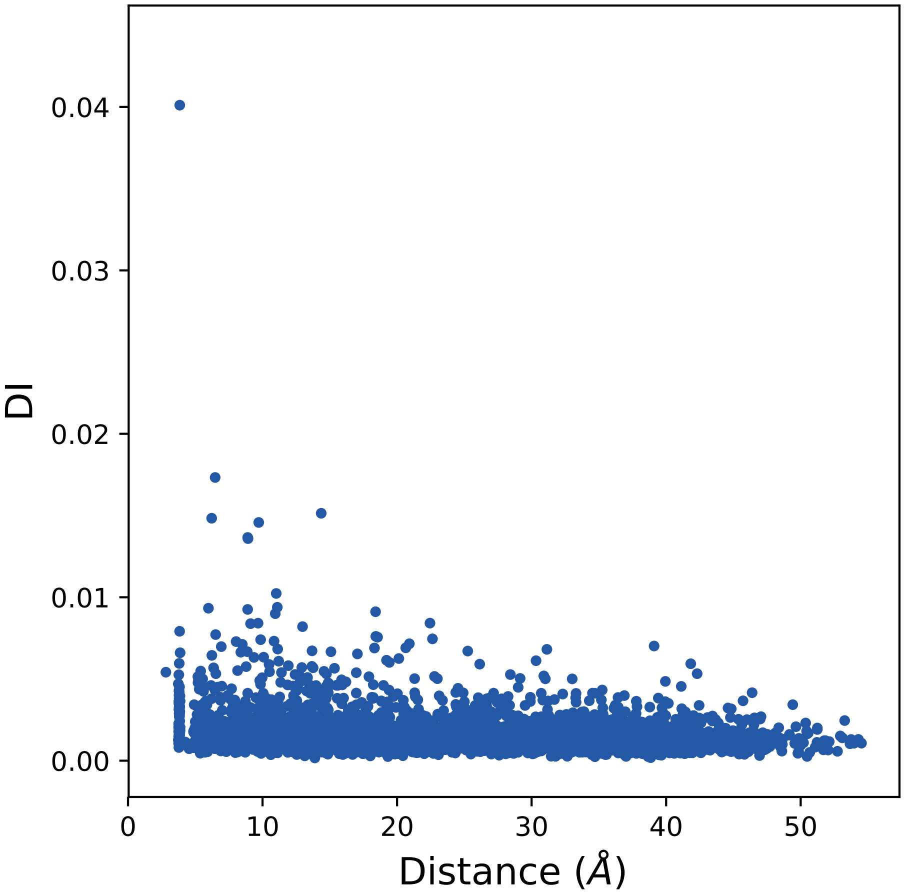
0.04
0.03
0.02
0.01
0.00
DI
0
10
20
30
40
50
Distance (Å)

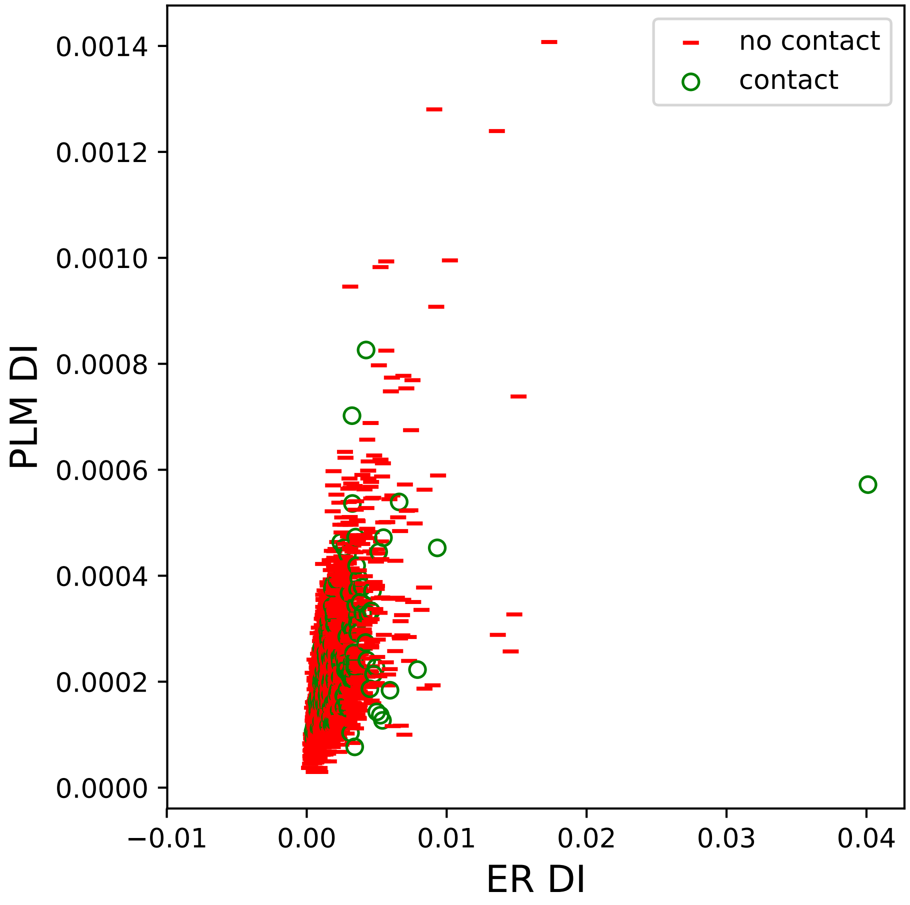

no contact
contact
PLM DI
0.0014
0.0012
0.0010
0.0008
0.0006
0.0004
0.0002
0.0000
−0.01
0.00
0.01
0.02
0.03
0.04
ER DI

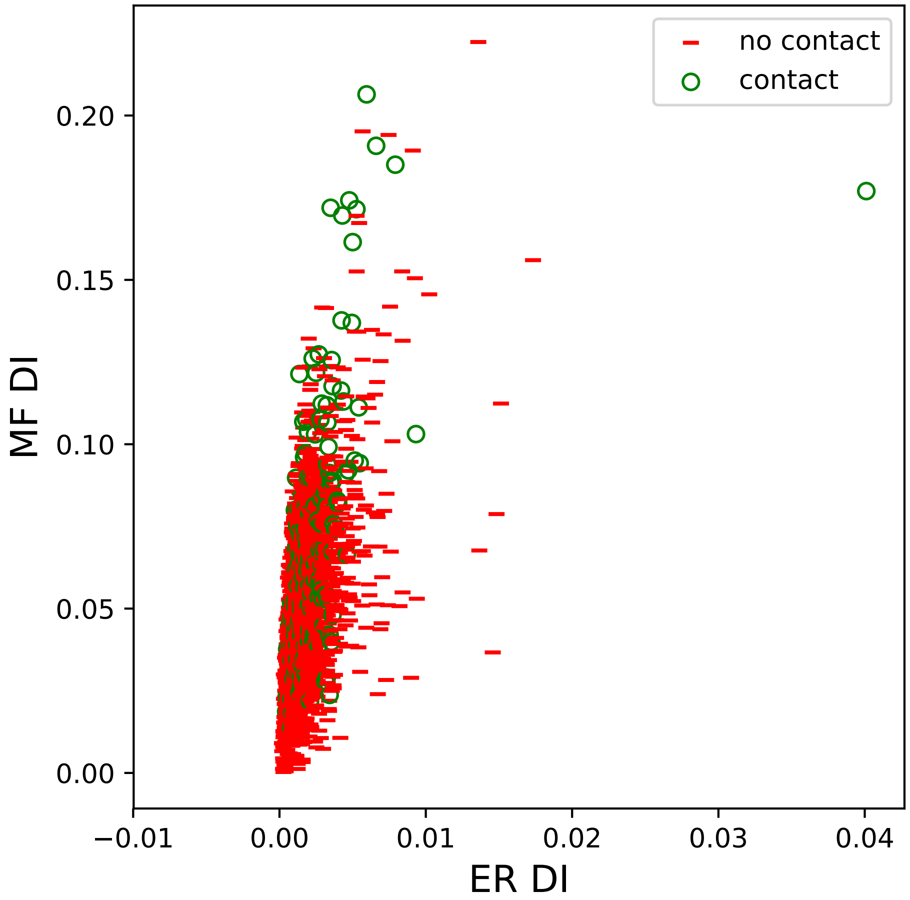

no contact
contact
0.20
0.15
0.10
0.05
0.00
MF DI
−0.01
0.00
0.01
0.02
0.03
0.04
ER DI

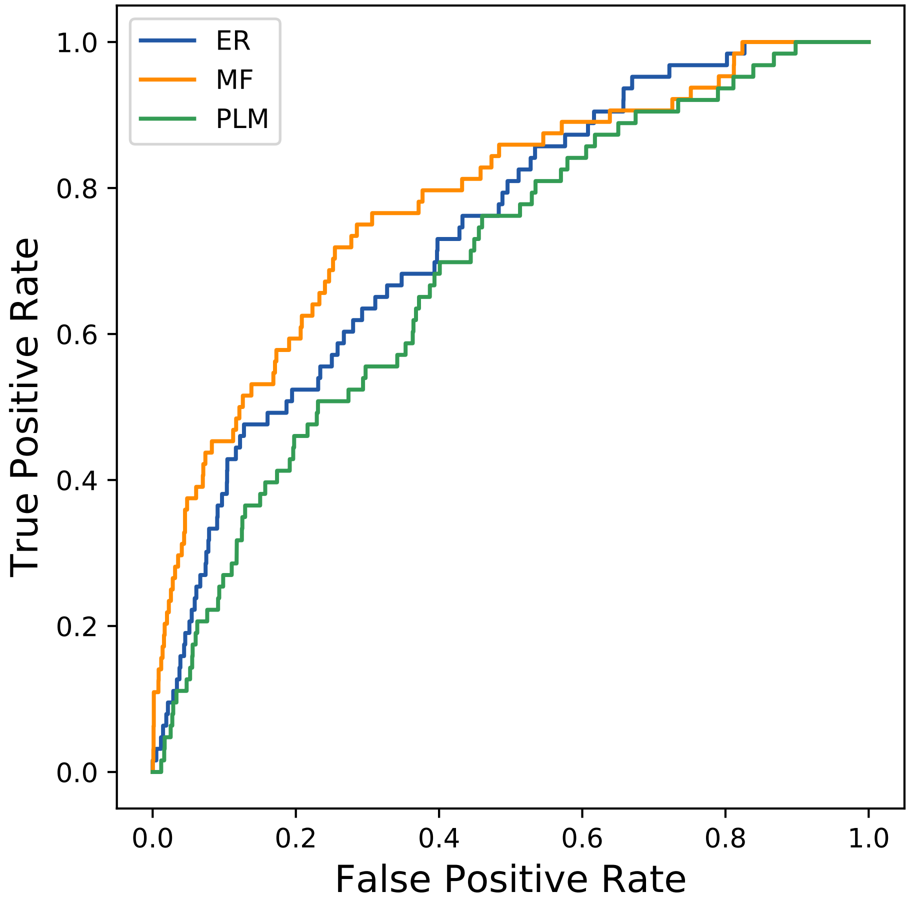
ER
MF
PLM
1.0
0.8
0.6
0.4
0.2
0.0
True Positive Rate
0.0
0.2
0.4
0.6
0.8
1.0
False Positive Rate
