## Supplemental Contact Maps for "Protein Structure Prediction with Expectation Reflection": 414seq_88col_8i1b_PF00340_plots.pdf

Method: ER
Maximum PDB contact distance : 10.0 Angstrom
Minimum residue chain distance: 5.0 residues
Fraction of true positives : 0.42

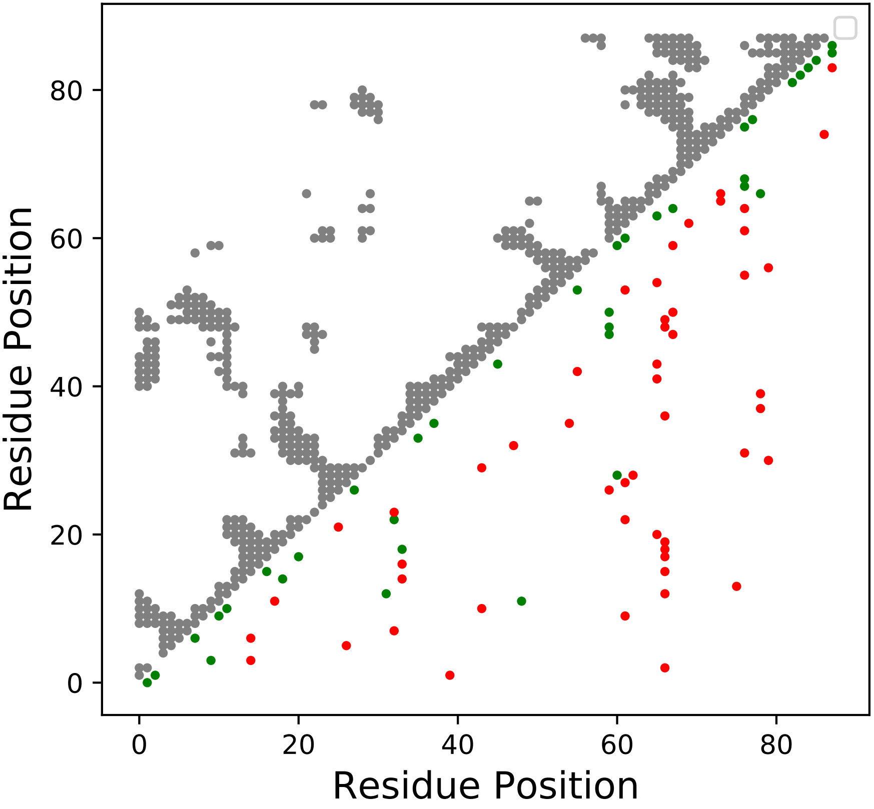

Method: MF
Maximum PDB contact distance : 10.0 Angstrom
Minimum residue chain distance: 5.0 residues
Fraction of true positives : 0.42

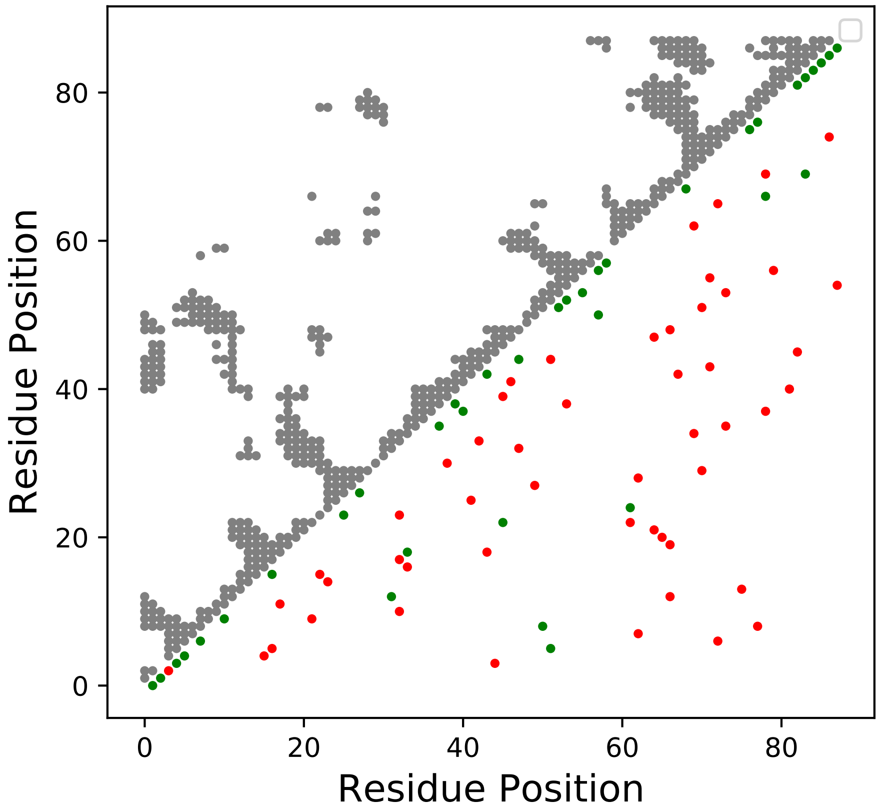

Method: PLM
Maximum PDB contact distance : 10.0 Angstrom
Minimum residue chain distance: 5.0 residues
Fraction of true positives : 0.398

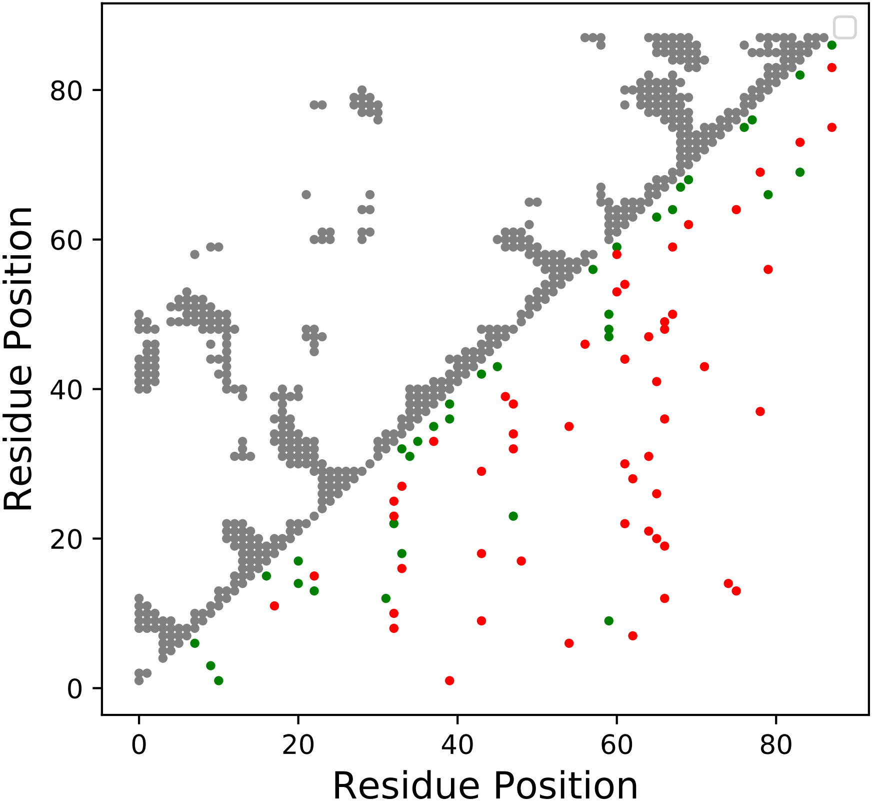

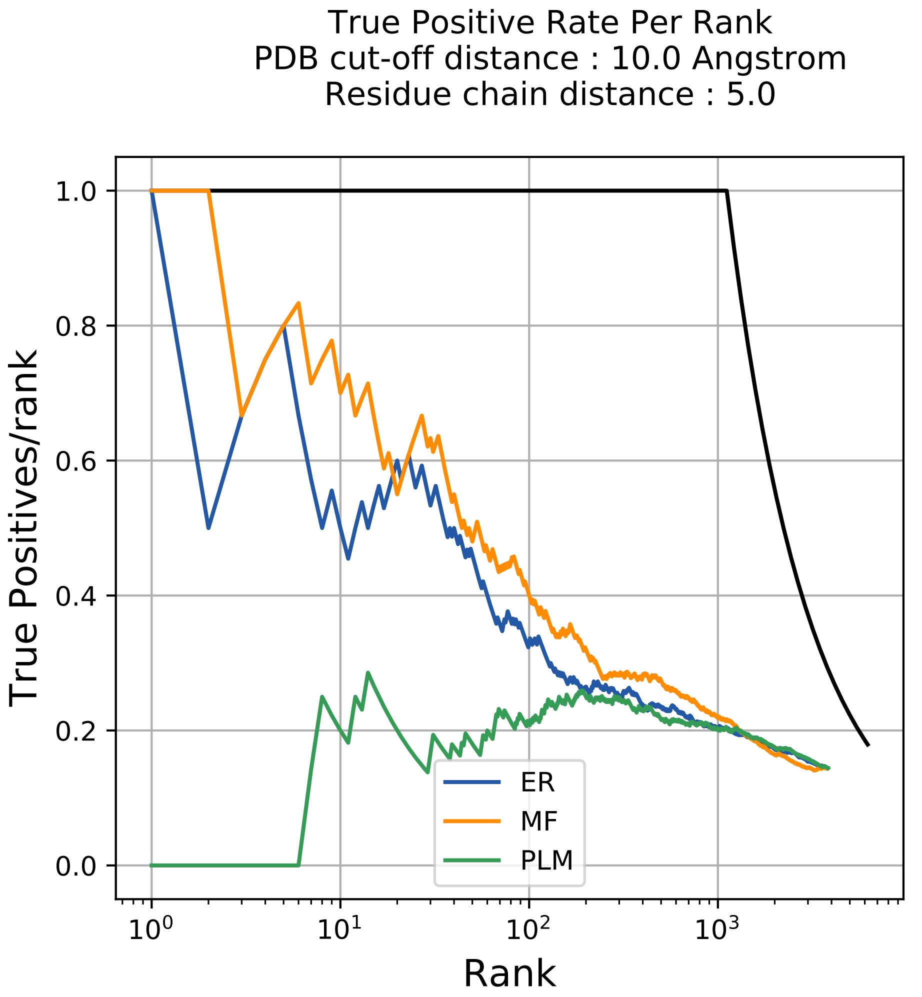

True Positive Rate Per Rank
PDB cut-off distance : 10.0 Angstrom
Residue chain distance : 5.0
True Positives/rank
Rank
1.0
0.8
0.6
0.4
0.2
0.0
$10^0$
$10^1$
$10^2$
$10^3$
ER
MF
PLM

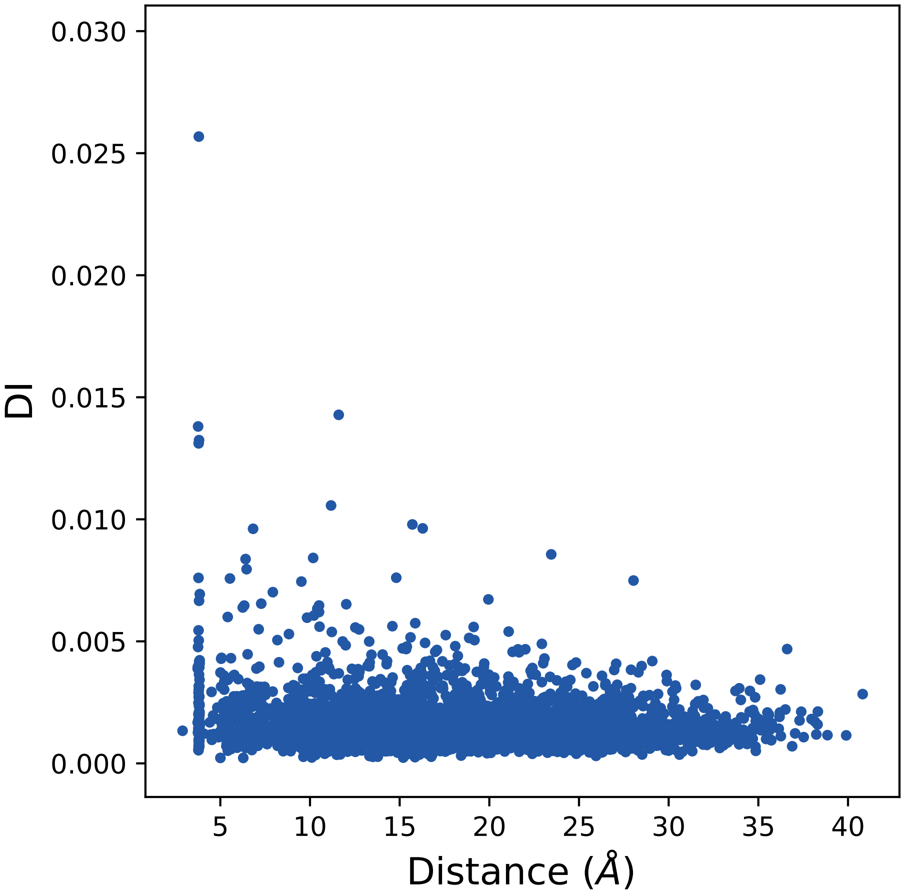
0.030
0.025
0.020
0.015
0.010
0.005
0.000
DI
5
10
15
20
25
30
35
40
Distance (Å)

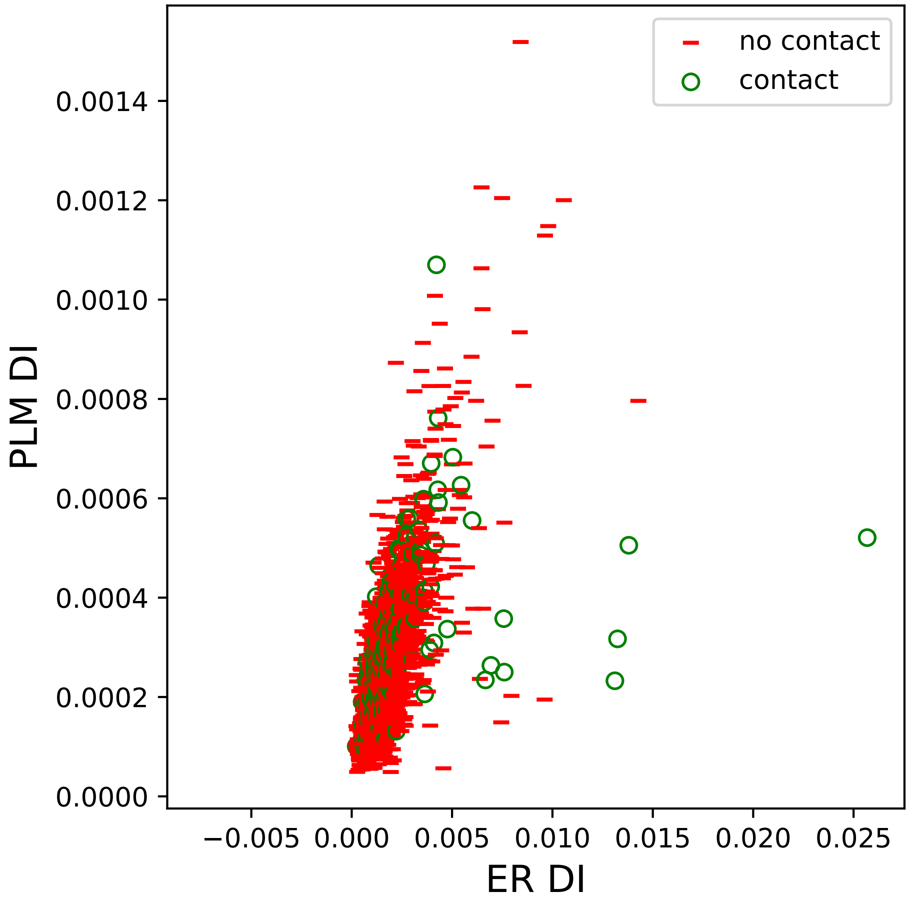
no contact
contact
0.0014
0.0012
0.0010
0.0008
0.0006
0.0004
0.0002
0.0000
PLM DI
−0.005
0.000
0.005
0.010
0.015
0.020
0.025
ER DI

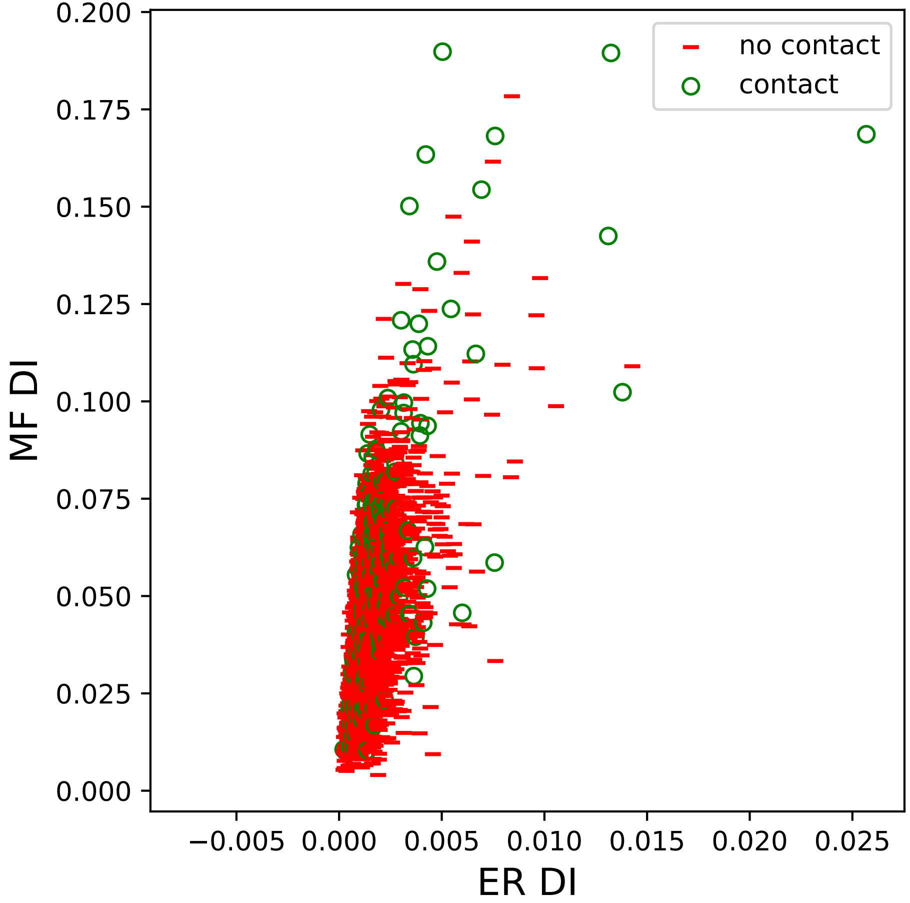
no contact
contact
MF DI
ER DI
0.200
0.175
0.150
0.125
0.100
0.075
0.050
0.025
0.000
−0.005
0.000
0.005
0.010
0.015
0.020
0.025

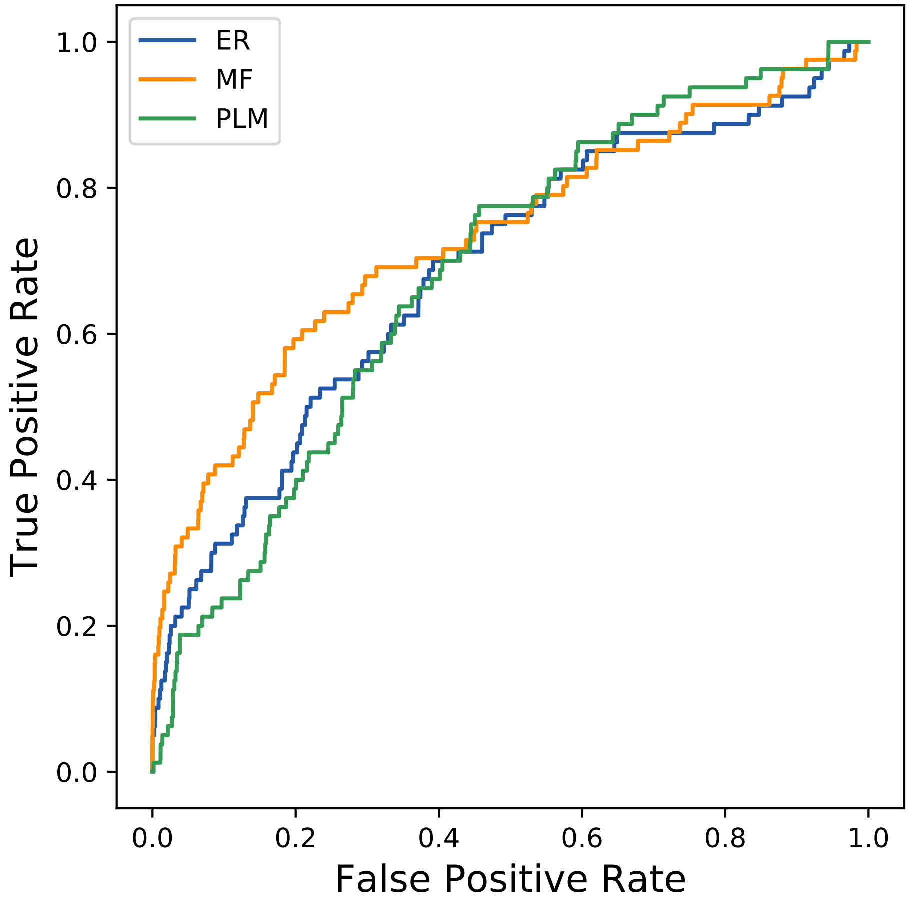

ER
MF
PLM
True Positive Rate
False Positive Rate
0.0
0.2
0.4
0.6
0.8
1.0
