## Supplemental Contact Maps for "Protein Structure Prediction with Expectation Reflection": 851seq_56col_3zu8_PF00942_plots.pdf

Method: ER
Maximum PDB contact distance : 10.0 Angstrom
Minimum residue chain distance: 5.0 residues
Fraction of true positives : 0.5

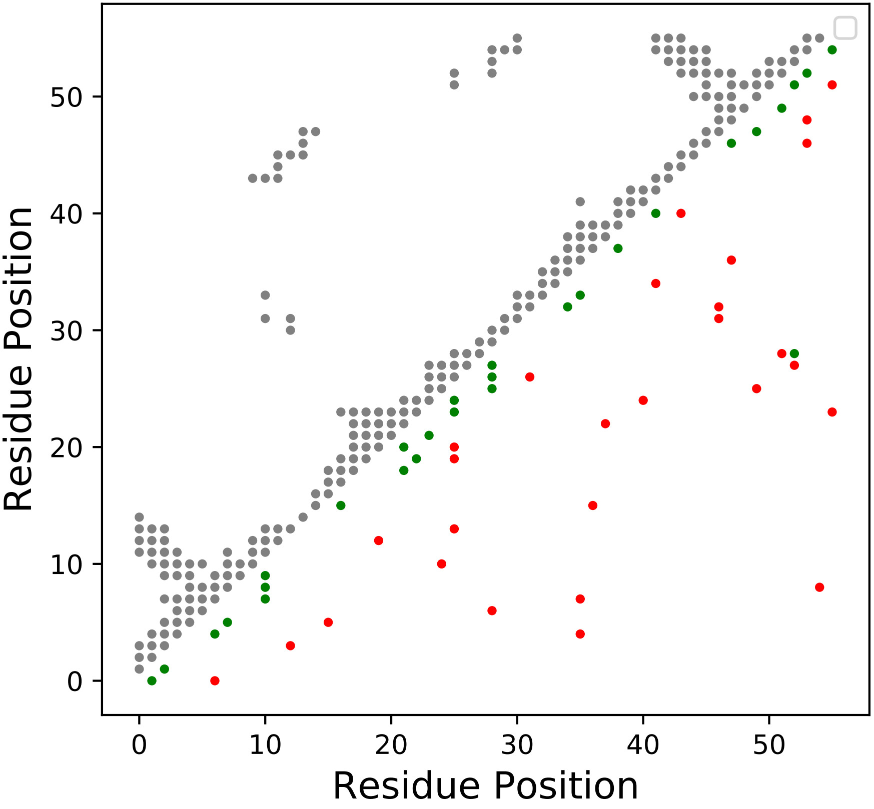

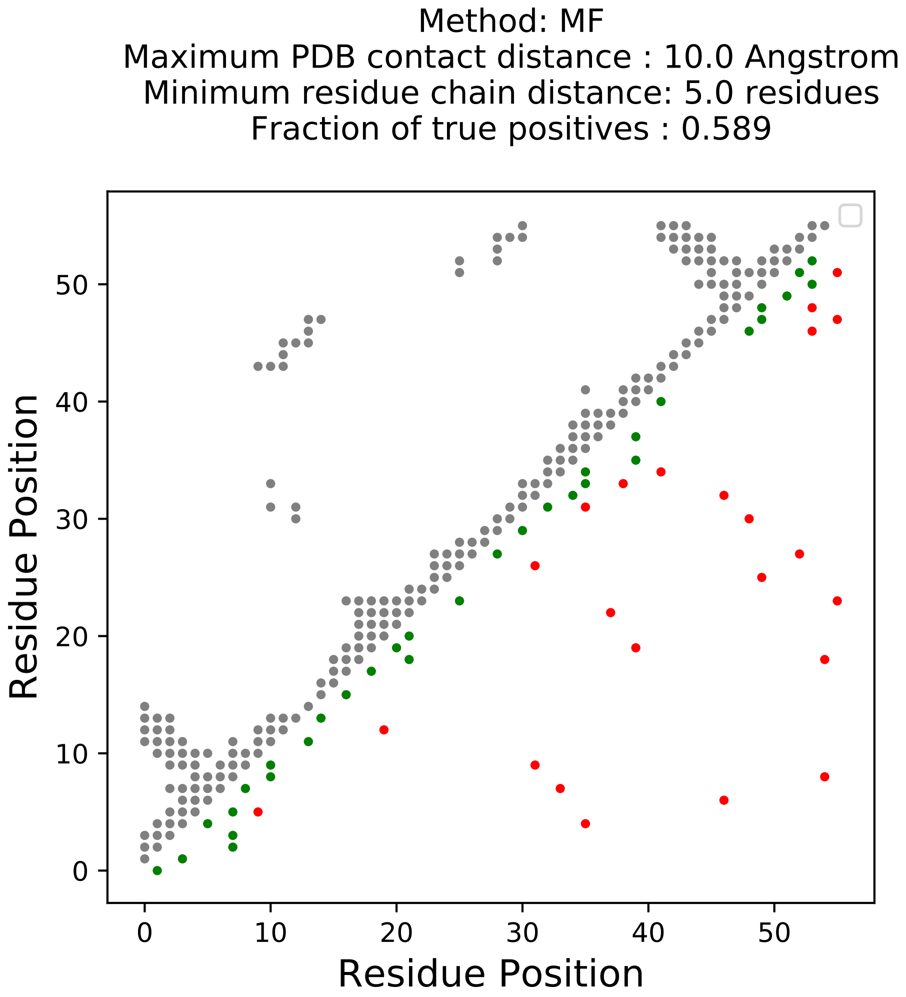
Method: MF
Maximum PDB contact distance : 10.0 Angstrom
Minimum residue chain distance: 5.0 residues
Fraction of true positives : 0.589
Residue Position
Residue Position
0
10
20
30
40
50

Method: PLM
Maximum PDB contact distance : 10.0 Angstrom
Minimum residue chain distance: 5.0 residues
Fraction of true positives : 0.536

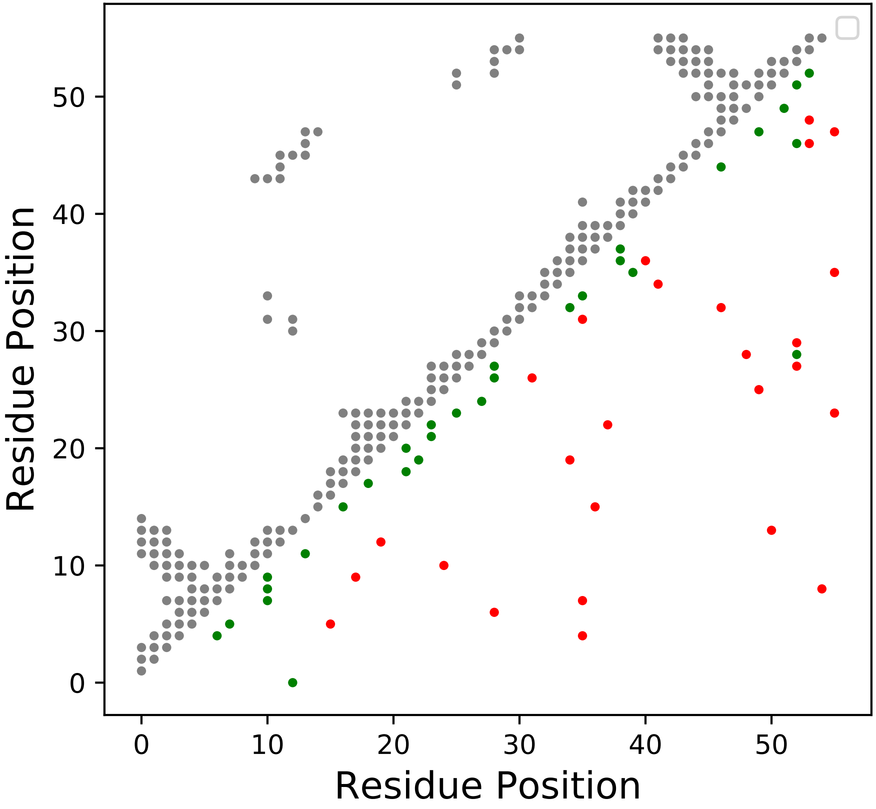

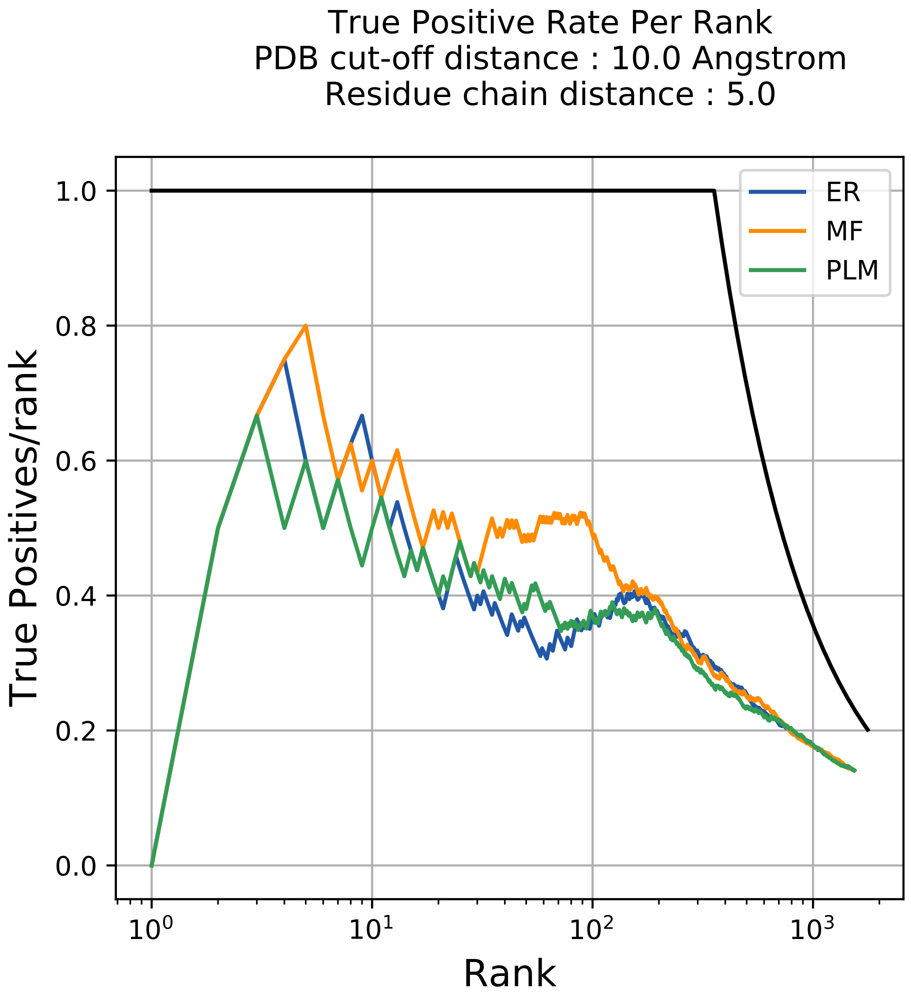
True Positive Rate Per Rank
PDB cut-off distance : 10.0 Angstrom
Residue chain distance : 5.0
True Positives/rank
Rank
ER
MF
PLM
1.0
0.8
0.6
0.4
0.2
0.0
$10^0$
$10^1$
$10^2$
$10^3$

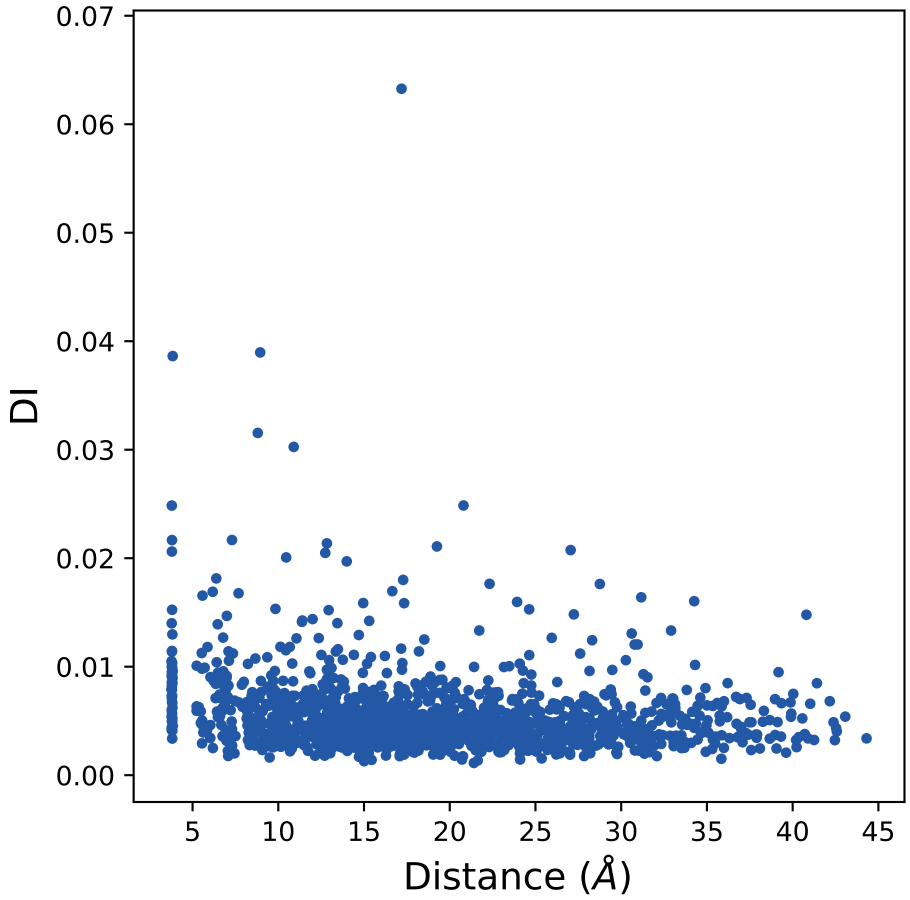
0.07
0.06
0.05
0.04
0.03
0.02
0.01
0.00
DI
5
10
15
20
25
30
35
40
45
Distance (Å)

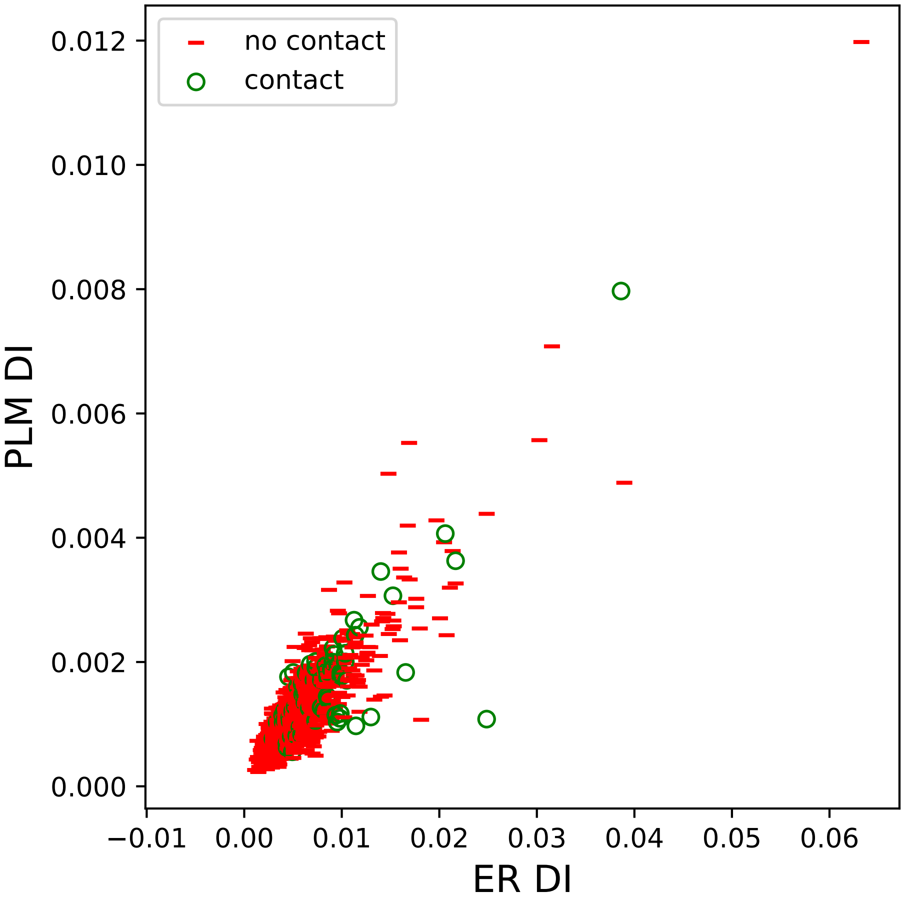
no contact
contact
PLM DI
ER DI
0.000
0.002
0.004
0.006
0.008
0.010
0.012
−0.01
0.00
0.01
0.02
0.03
0.04
0.05
0.06

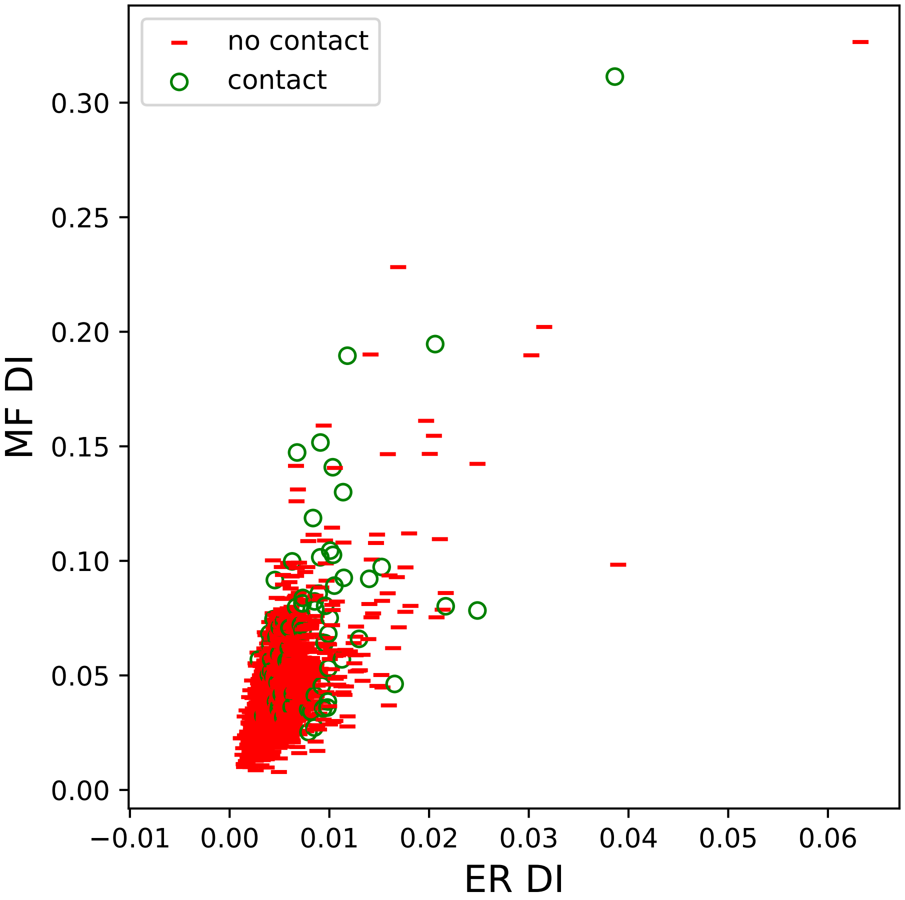

no contact
contact
0.30
0.25
0.20
0.15
0.10
0.05
0.00
MF DI
−0.01
0.00
0.01
0.02
0.03
0.04
0.05
0.06
ER DI

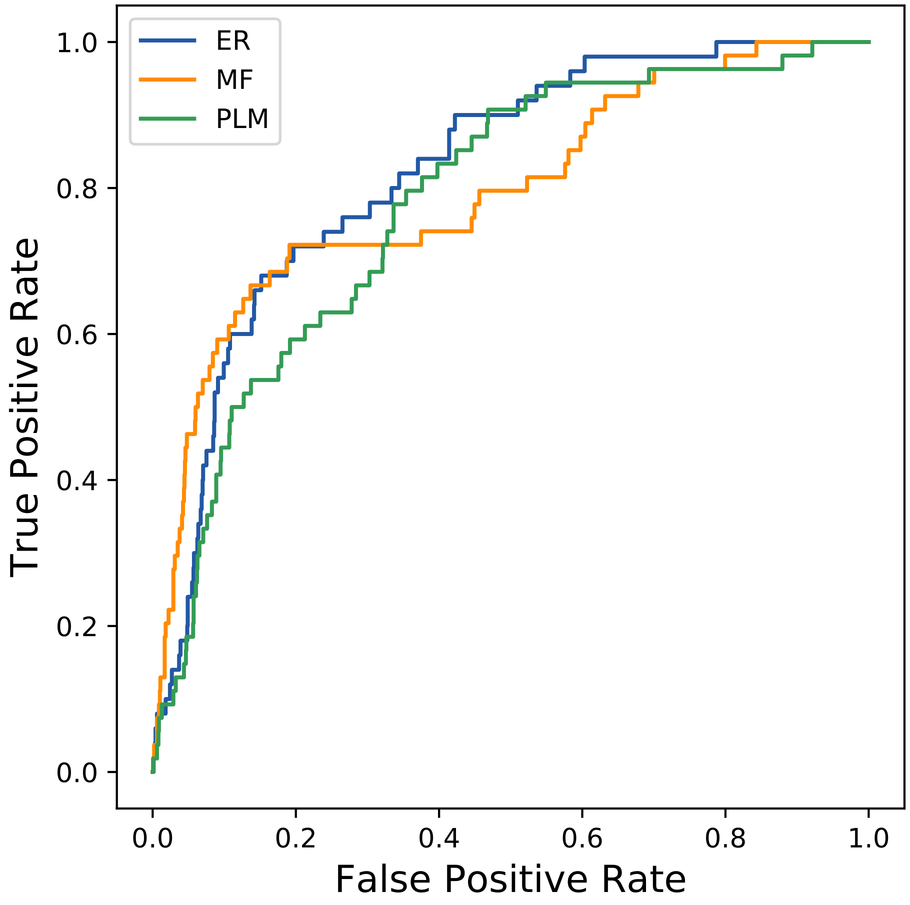

ER
MF
PLM
1.0
0.8
0.6
0.4
0.2
0.0
True Positive Rate
0.0
0.2
0.4
0.6
0.8
1.0
False Positive Rate
