## Supplemental Contact Maps for "Protein Structure Prediction with Expectation Reflection": 932seq_90col_1ckd_PF00062_plots.pdf

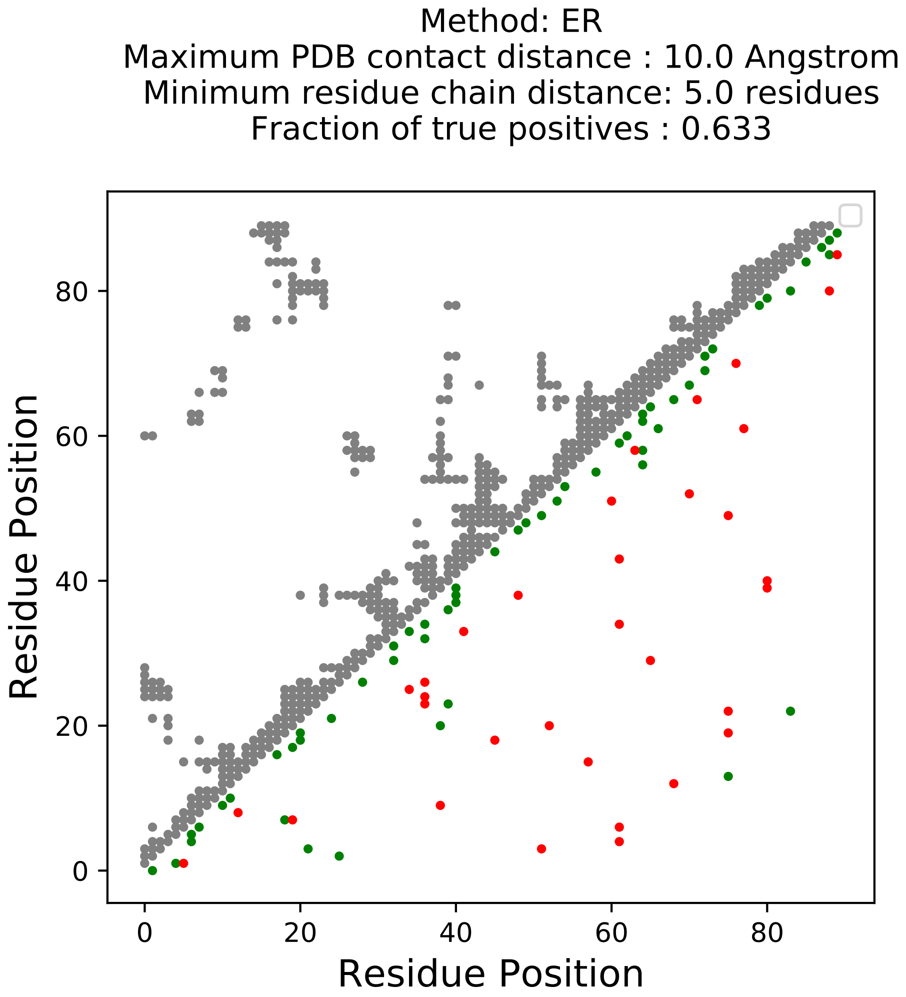
Method: ER
Maximum PDB contact distance : 10.0 Angstrom
Minimum residue chain distance: 5.0 residues
Fraction of true positives : 0.633
Residue Position
Residue Position
0
20
40
60
80

Method: MF
Maximum PDB contact distance : 10.0 Angstrom
Minimum residue chain distance: 5.0 residues
Fraction of true positives : 0.589

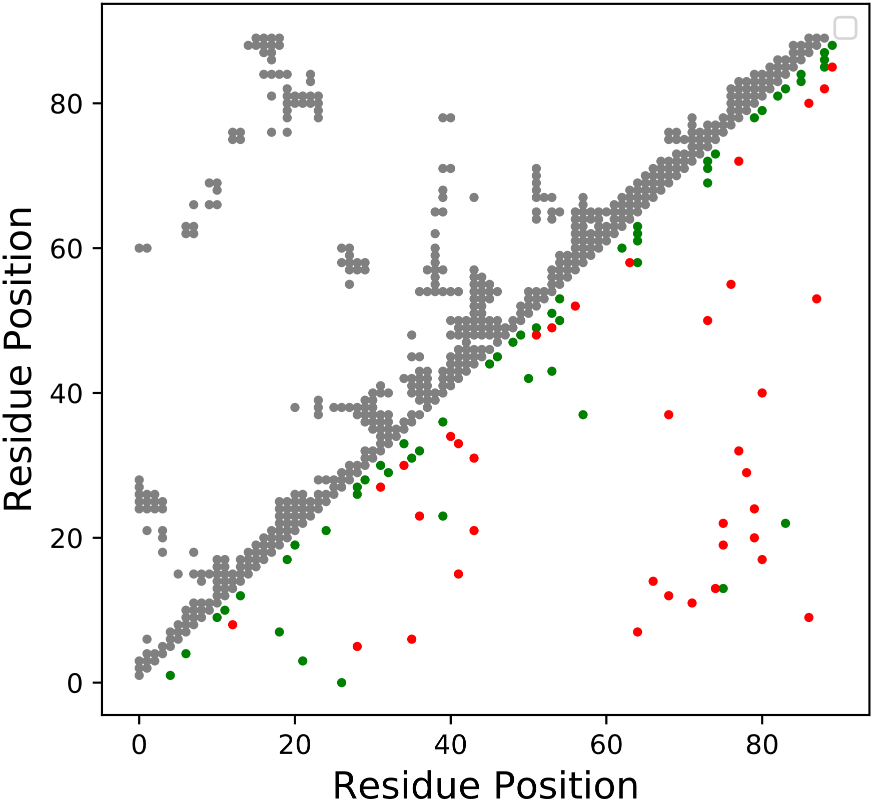

Method: PLM
Maximum PDB contact distance : 10.0 Angstrom
Minimum residue chain distance: 5.0 residues
Fraction of true positives : 0.556

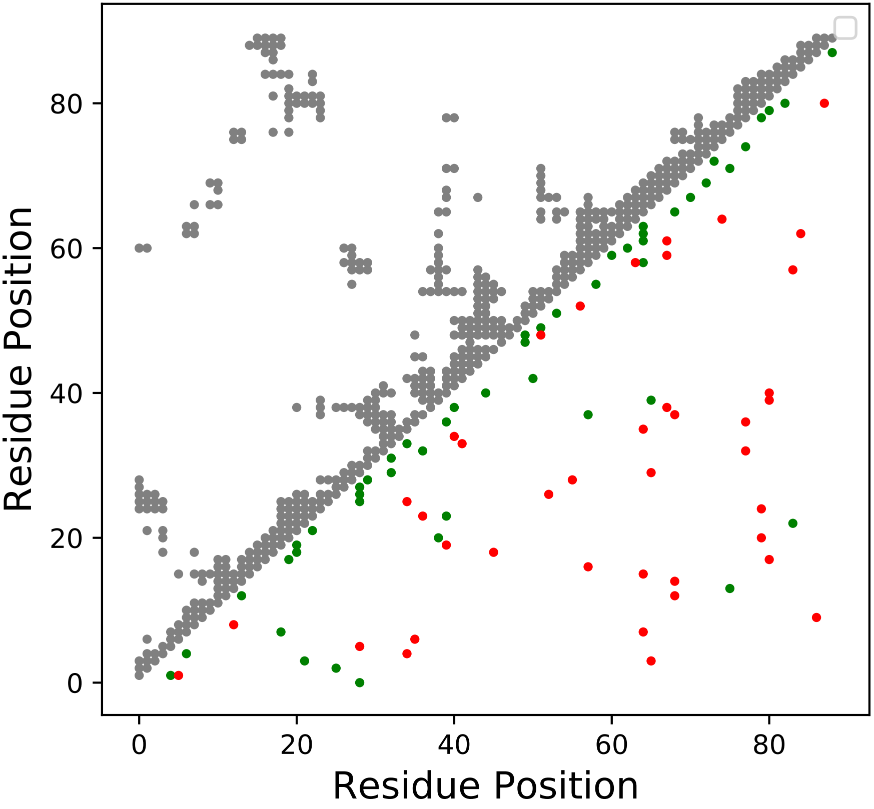

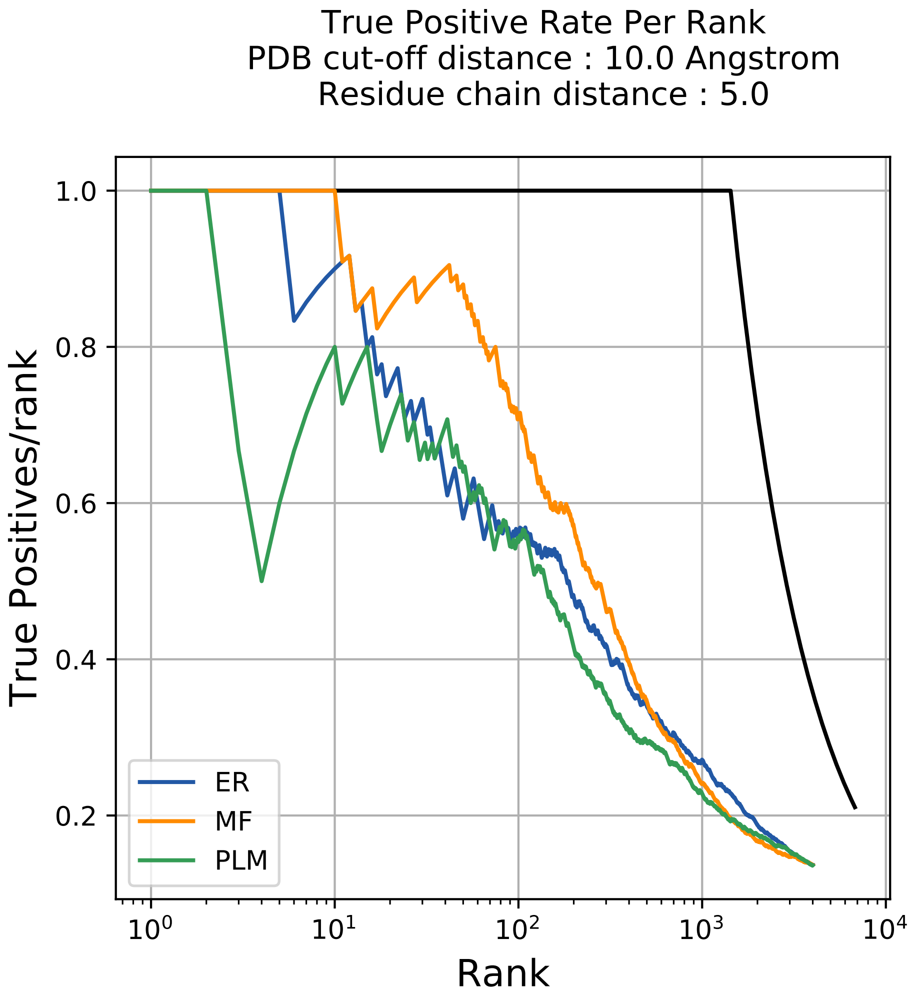

True Positive Rate Per Rank
PDB cut-off distance : 10.0 Angstrom
Residue chain distance : 5.0
True Positives/rank
Rank
1.0
0.8
0.6
0.4
0.2
$10^0$
$10^1$
$10^2$
$10^3$
$10^4$
ER
MF
PLM

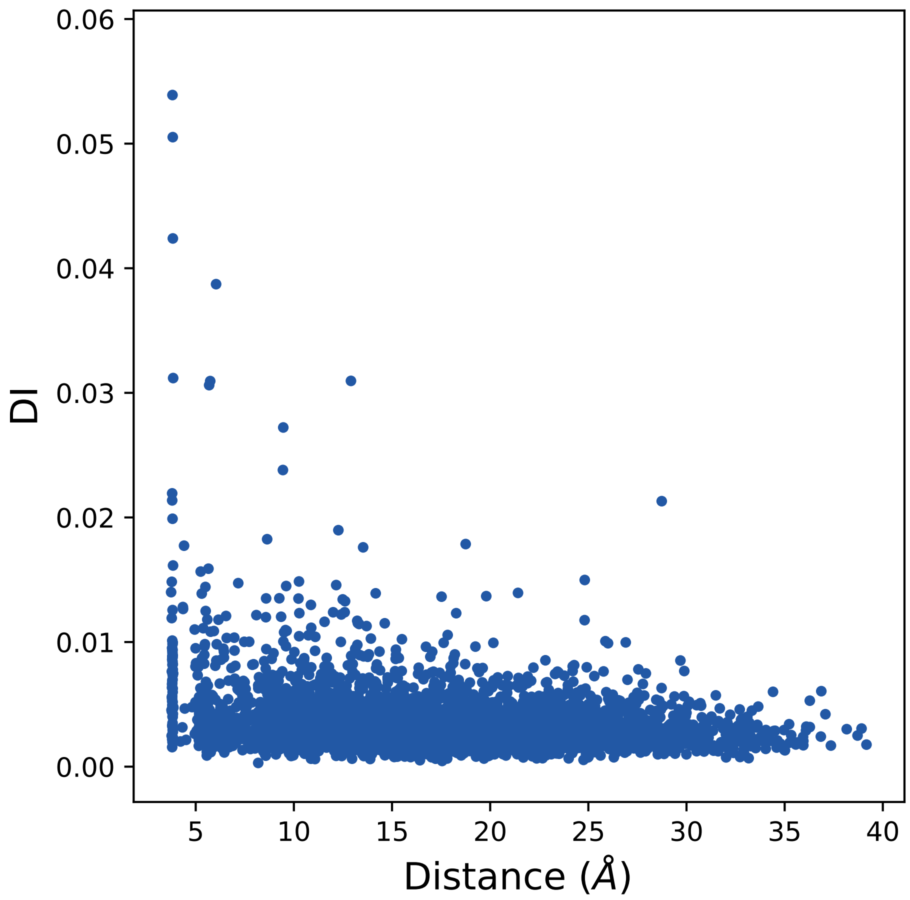

0.06
0.05
0.04
0.03
0.02
0.01
0.00
DI
5
10
15
20
25
30
35
40
Distance (Å)

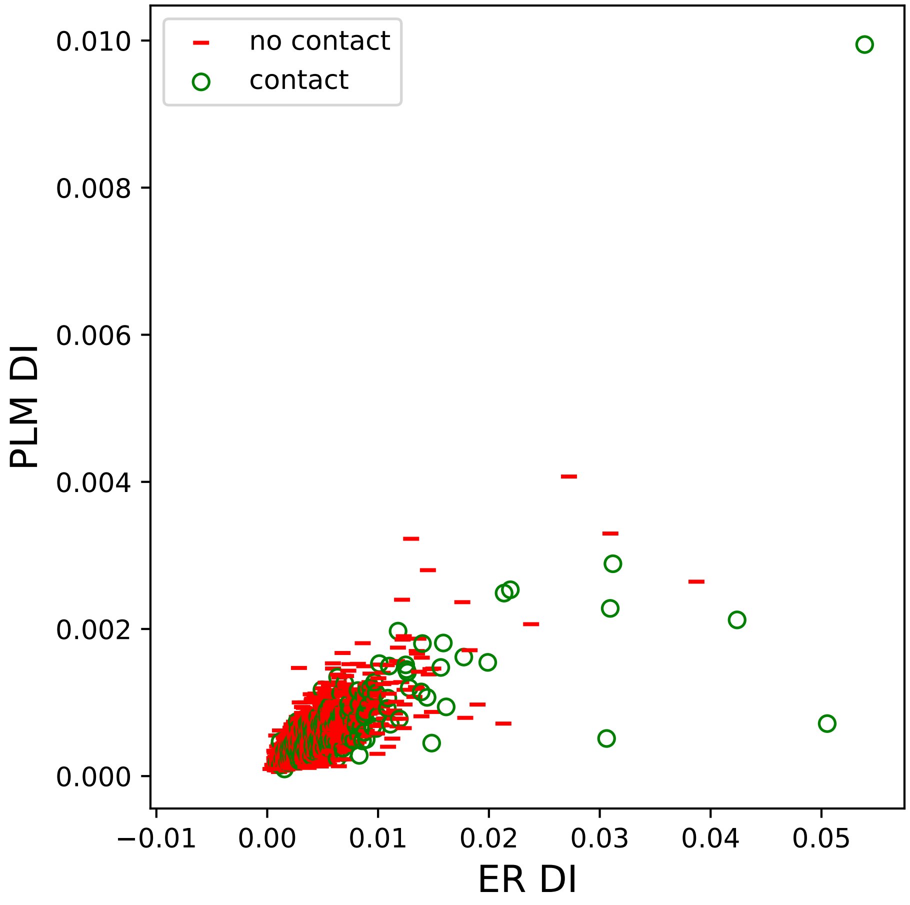

no contact
contact
0.010
0.008
0.006
0.004
0.002
0.000
PLM DI
−0.01
0.00
0.01
0.02
0.03
0.04
0.05
ER DI

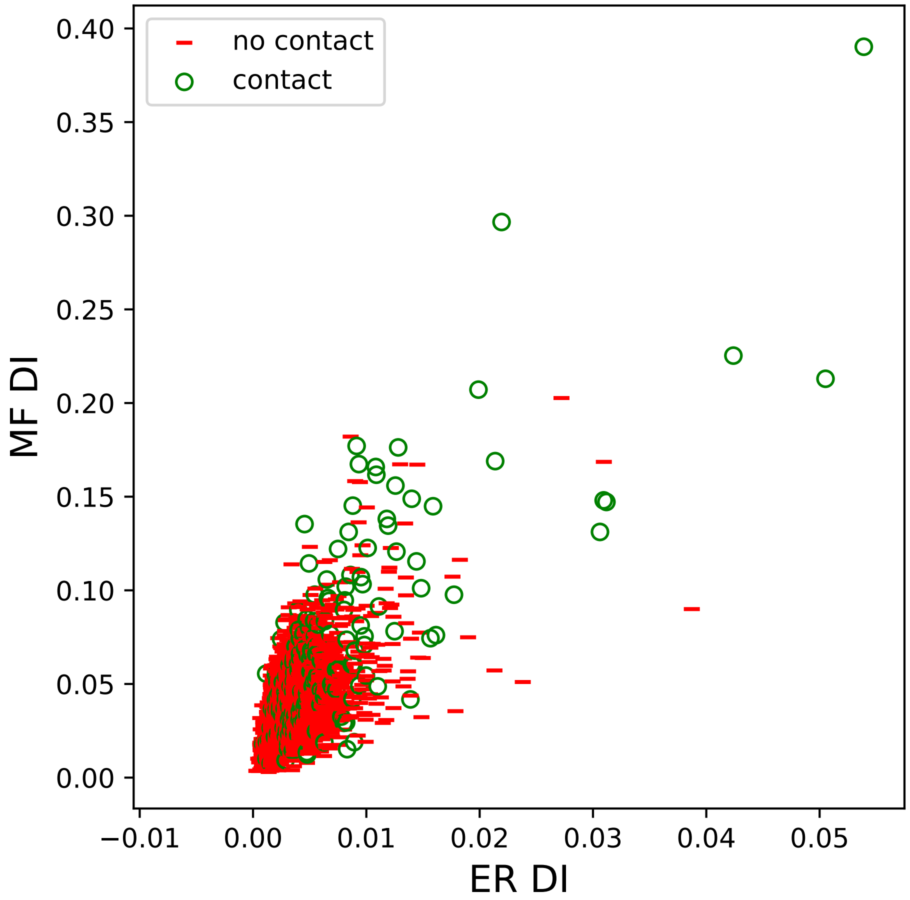

no contact
contact
MF DI
ER DI
0.00
0.05
0.10
0.15
0.20
0.25
0.30
0.35
0.40
−0.01
0.00
0.01
0.02
0.03
0.04
0.05

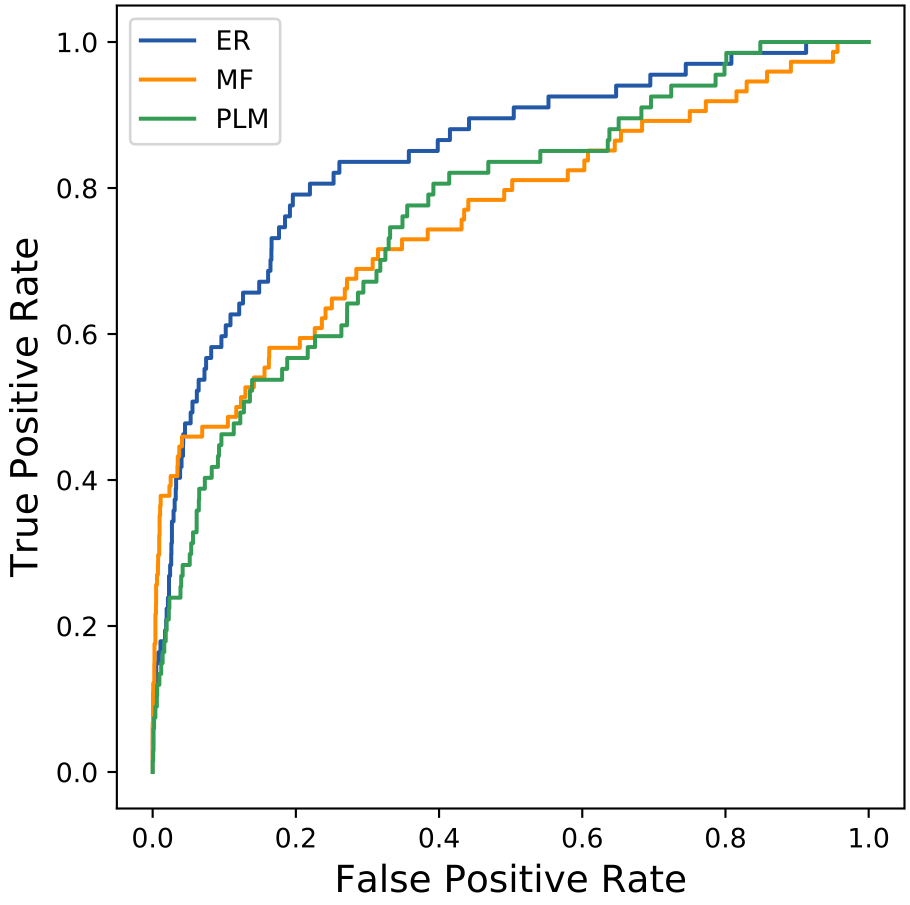
ER
MF
PLM
1.0
0.8
0.6
0.4
0.2
0.0
True Positive Rate
0.0
0.2
0.4
0.6
0.8
1.0
False Positive Rate
