## Supplemental Contact Maps for "Protein Structure Prediction with Expectation Reflection": 1090seq_155col_5jai_PF12241_plots.pdf

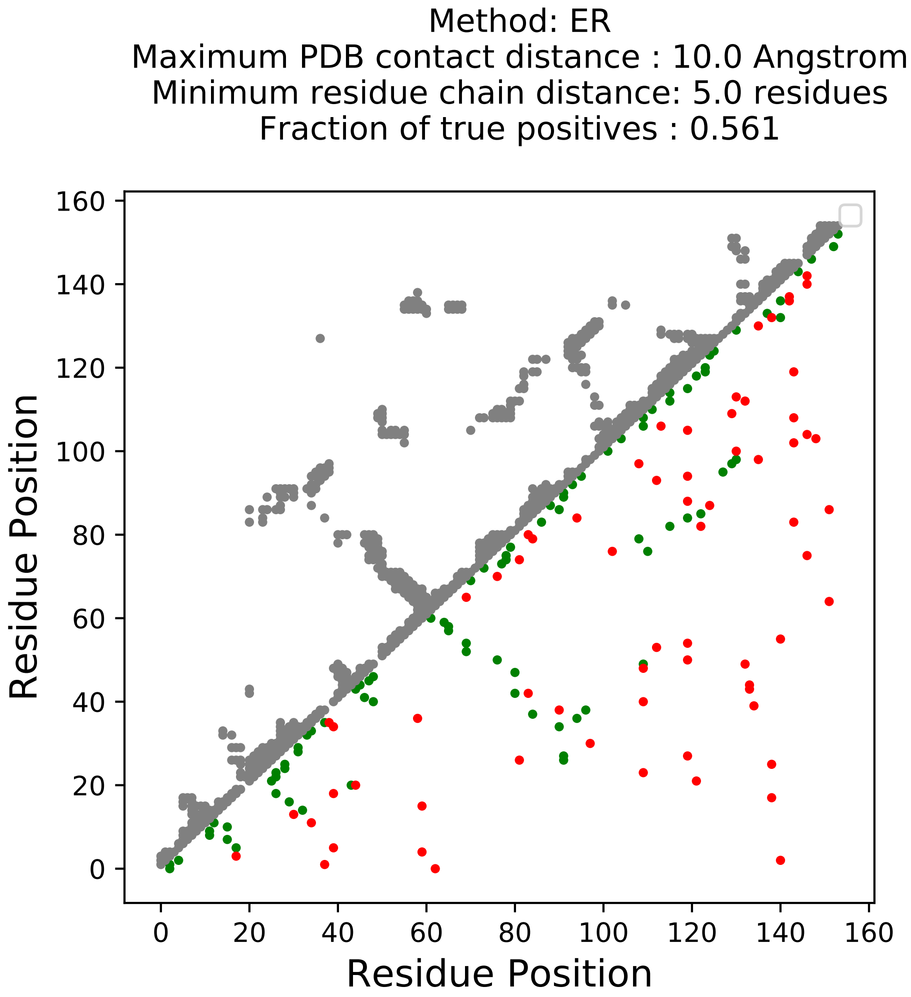

Method: ER
Maximum PDB contact distance : 10.0 Angstrom
Minimum residue chain distance: 5.0 residues
Fraction of true positives : 0.561
Residue Position
Residue Position
0
20
40
60
80
100
120
140
160

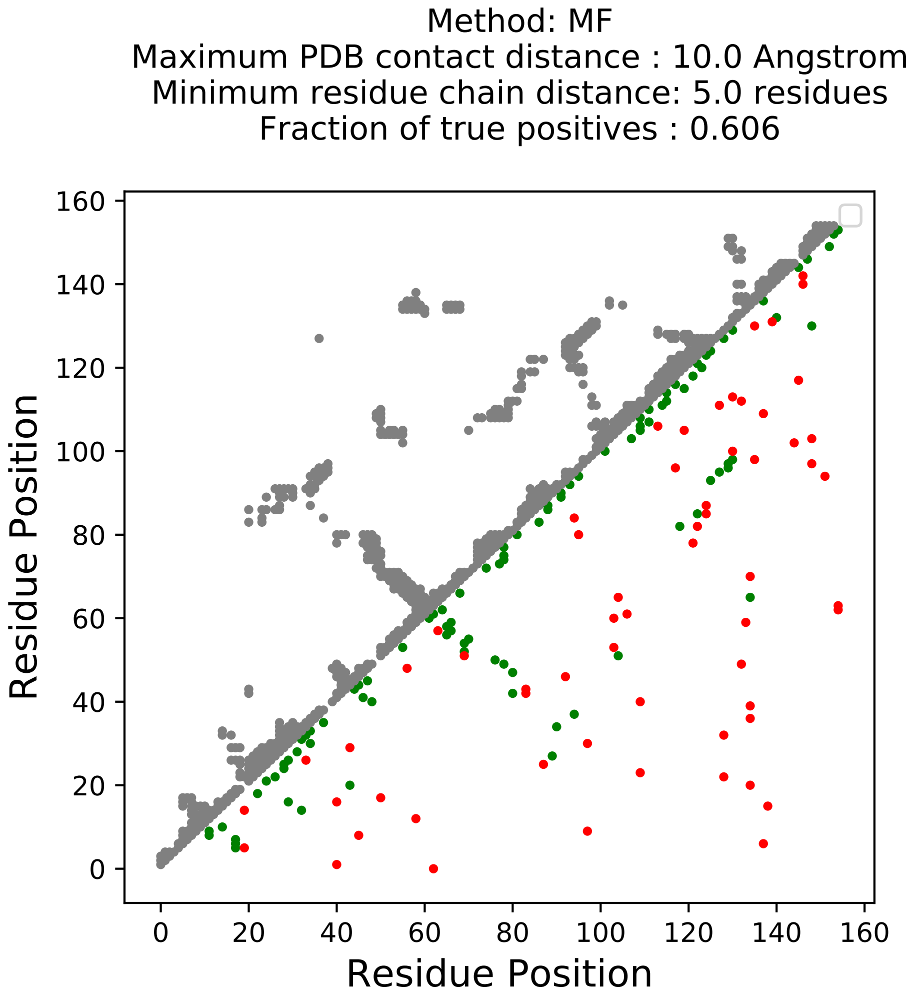

Method: MF
Maximum PDB contact distance : 10.0 Angstrom
Minimum residue chain distance: 5.0 residues
Fraction of true positives : 0.606
Residue Position
Residue Position
0
20
40
60
80
100
120
140
160

Method: PLM
Maximum PDB contact distance : 10.0 Angstrom
Minimum residue chain distance: 5.0 residues
Fraction of true positives : 0.529

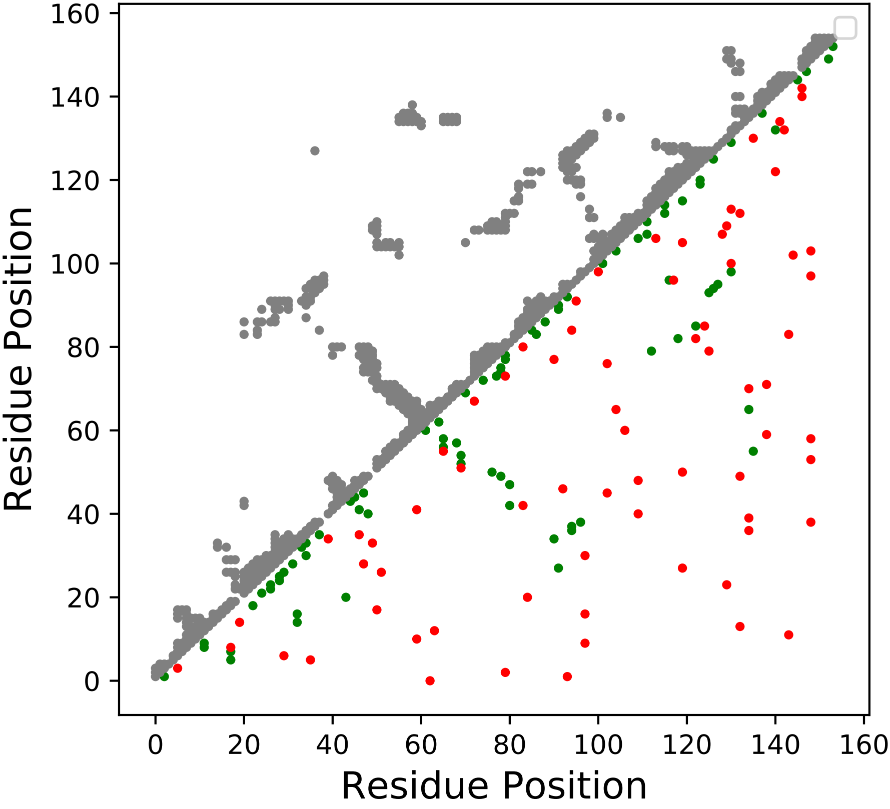

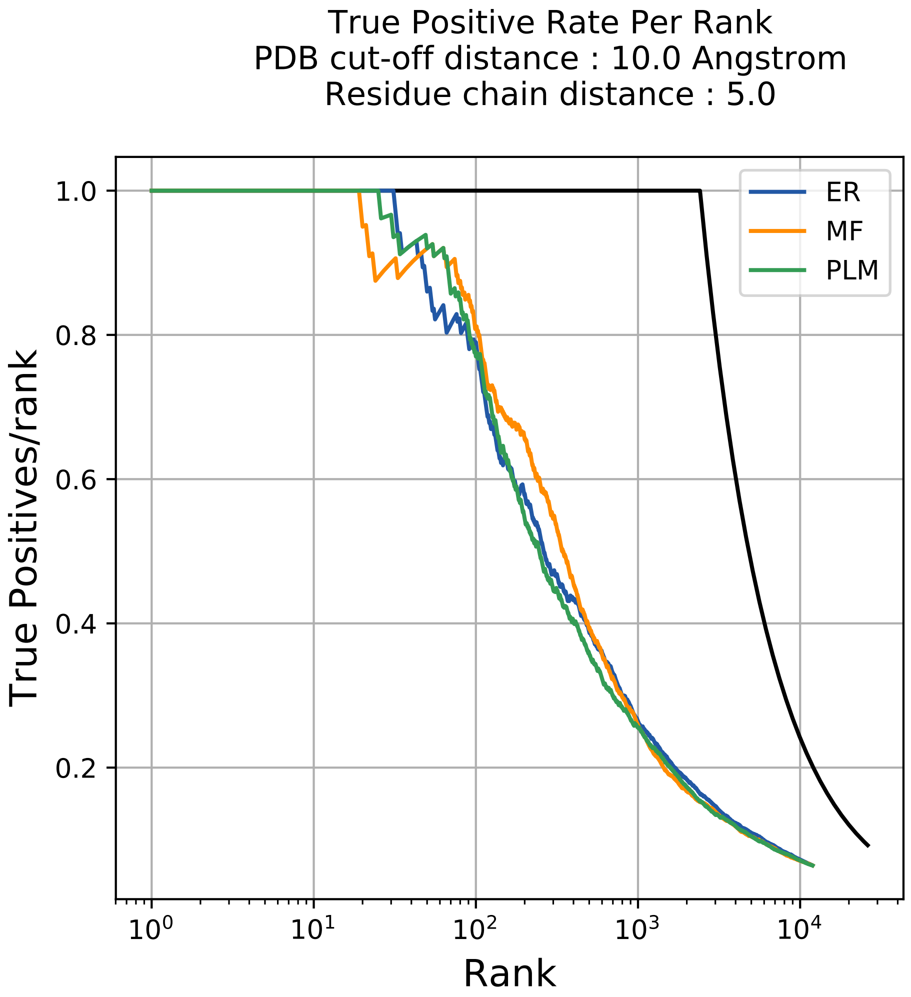
True Positive Rate Per Rank
PDB cut-off distance : 10.0 Angstrom
Residue chain distance : 5.0
ER
MF
PLM
1.0
0.8
0.6
0.4
0.2
True Positives/rank
$10^0$
$10^1$
$10^2$
$10^3$
$10^4$
Rank

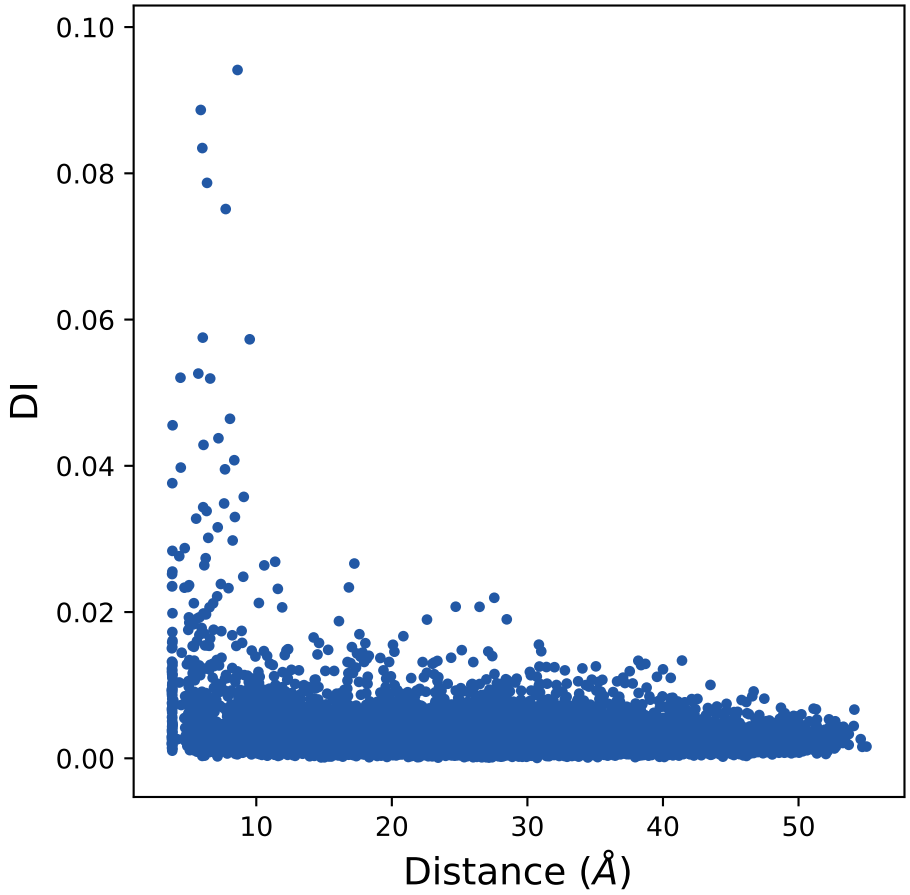

0.10
0.08
0.06
0.04
0.02
0.00
DI
10
20
30
40
50
Distance (Å)

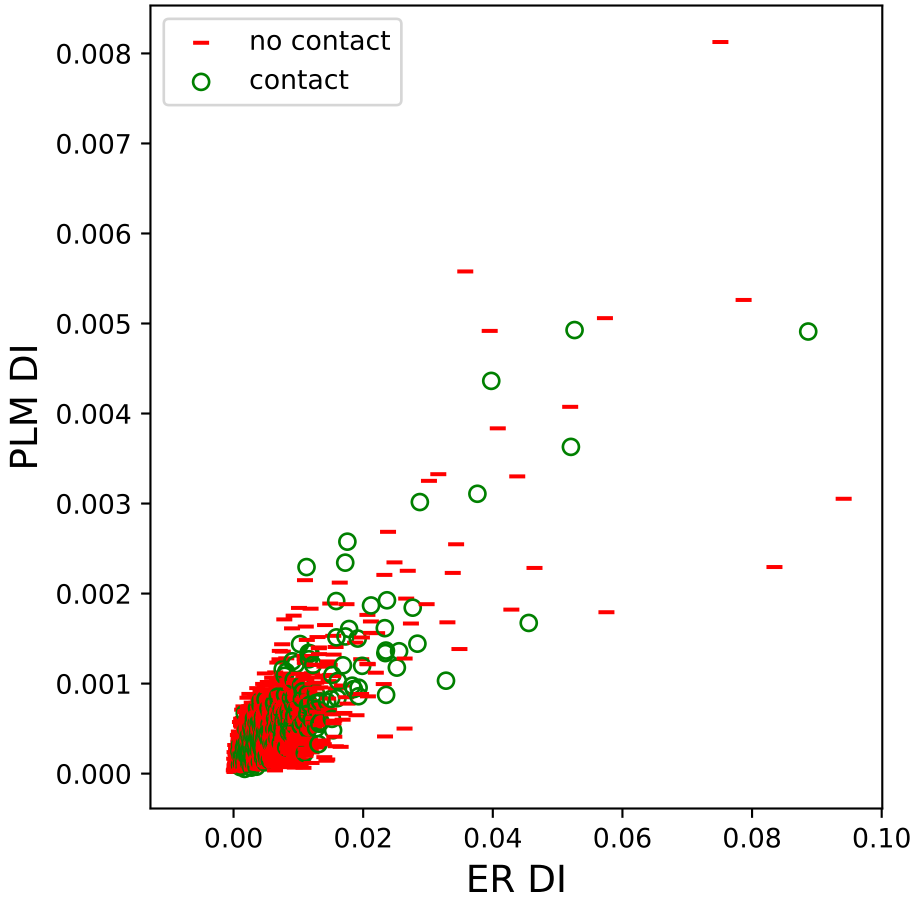

no contact
contact
PLM DI
ER DI
0.000
0.001
0.002
0.003
0.004
0.005
0.006
0.007
0.008
0.00
0.02
0.04
0.06
0.08
0.10

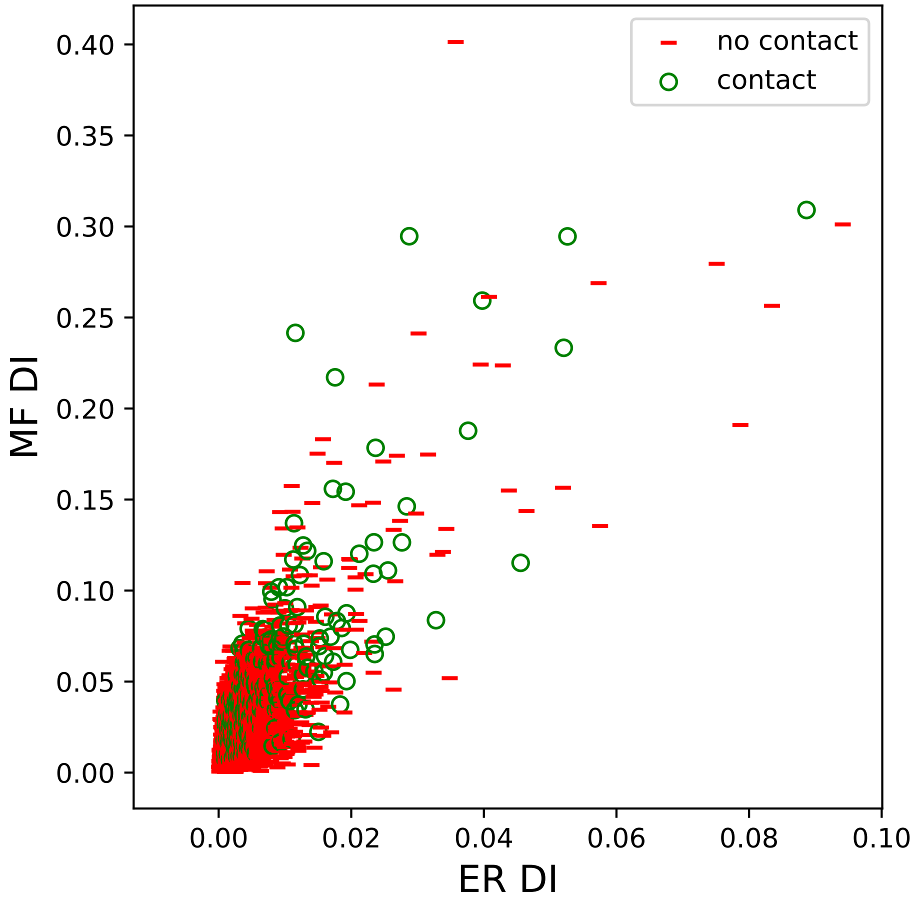

no contact
contact
MF DI
ER DI
0.00
0.05
0.10
0.15
0.20
0.25
0.30
0.35
0.40
0.00
0.02
0.04
0.06
0.08
0.10

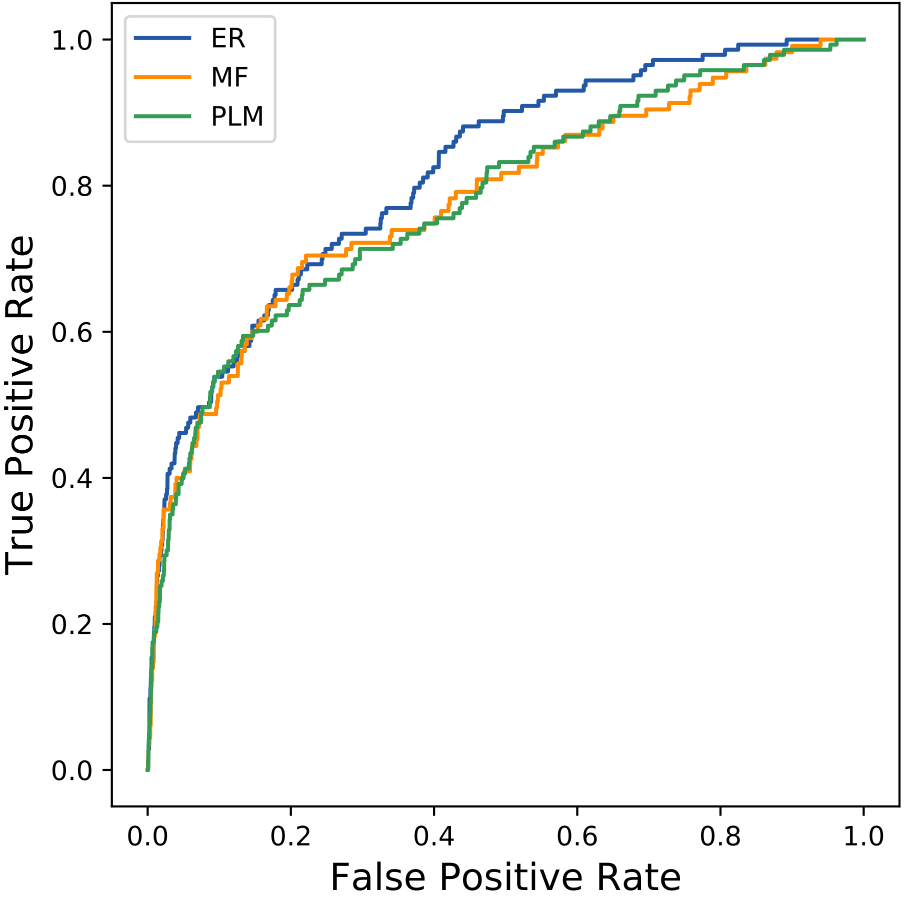
ER
MF
PLM
1.0
0.8
0.6
0.4
0.2
0.0
True Positive Rate
0.0
0.2
0.4
0.6
0.8
1.0
False Positive Rate
