## Supplemental Contact Maps for "Protein Structure Prediction with Expectation Reflection": 1228seq_141col_3b7m_PF01670_plots.pdf

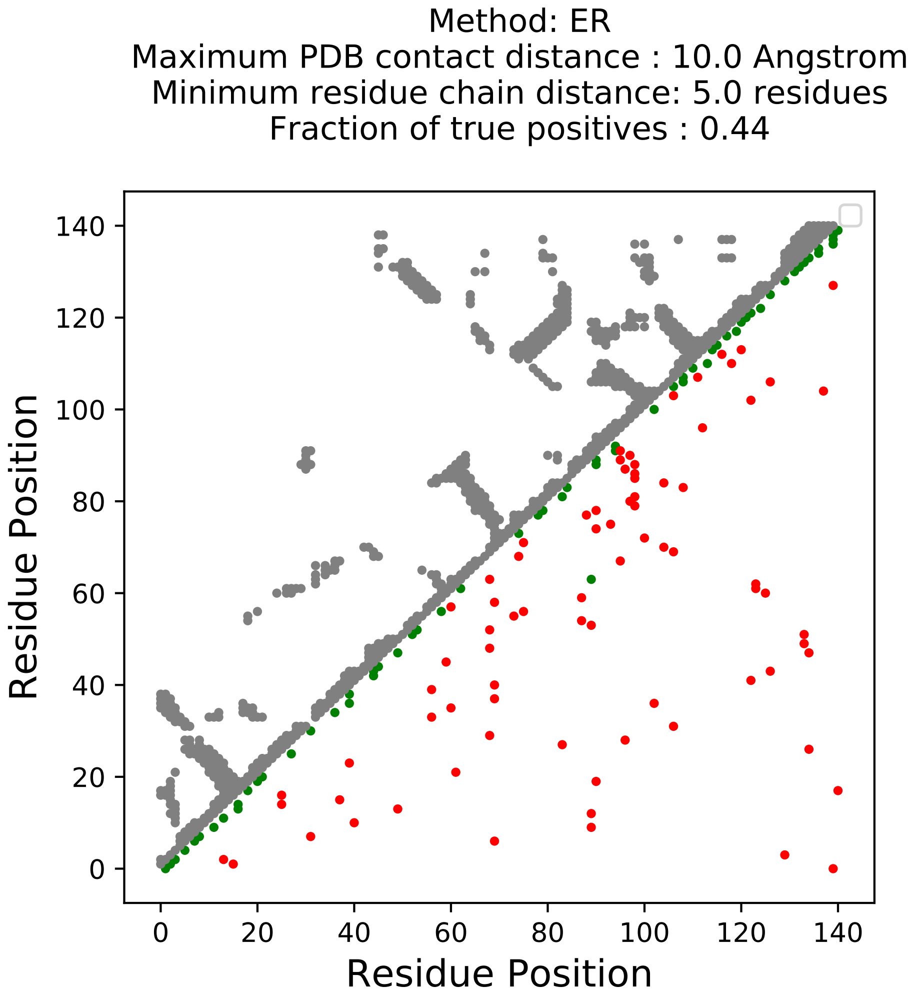
Method: ER
Maximum PDB contact distance : 10.0 Angstrom
Minimum residue chain distance: 5.0 residues
Fraction of true positives : 0.44
Residue Position
Residue Position
0
20
40
60
80
100
120
140

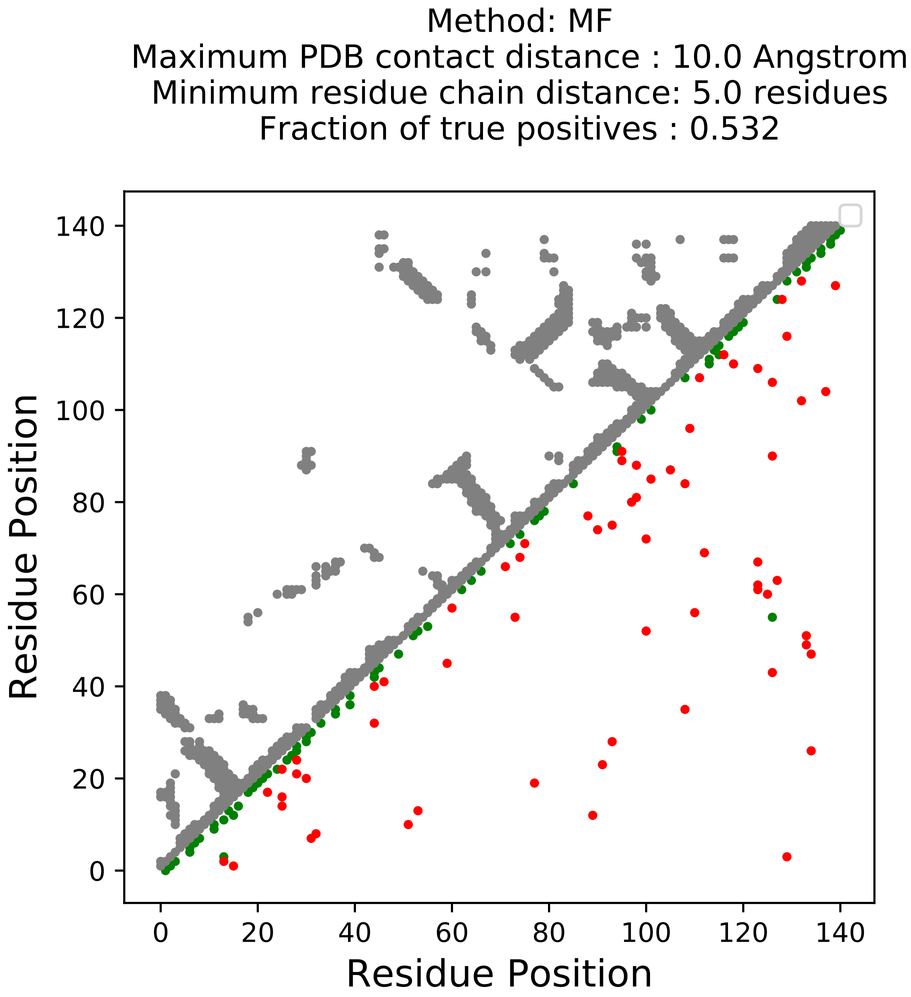
Method: MF
Maximum PDB contact distance : 10.0 Angstrom
Minimum residue chain distance: 5.0 residues
Fraction of true positives : 0.532
Residue Position
Residue Position
0
20
40
60
80
100
120
140

Method: PLM
Maximum PDB contact distance : 10.0 Angstrom
Minimum residue chain distance: 5.0 residues
Fraction of true positives : 0.461

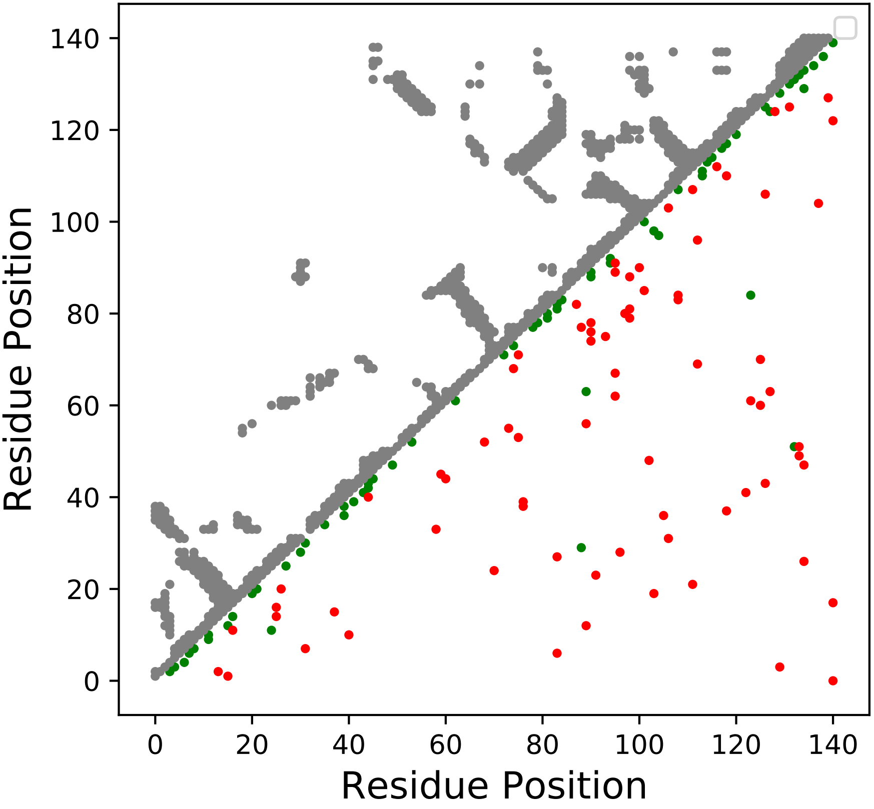

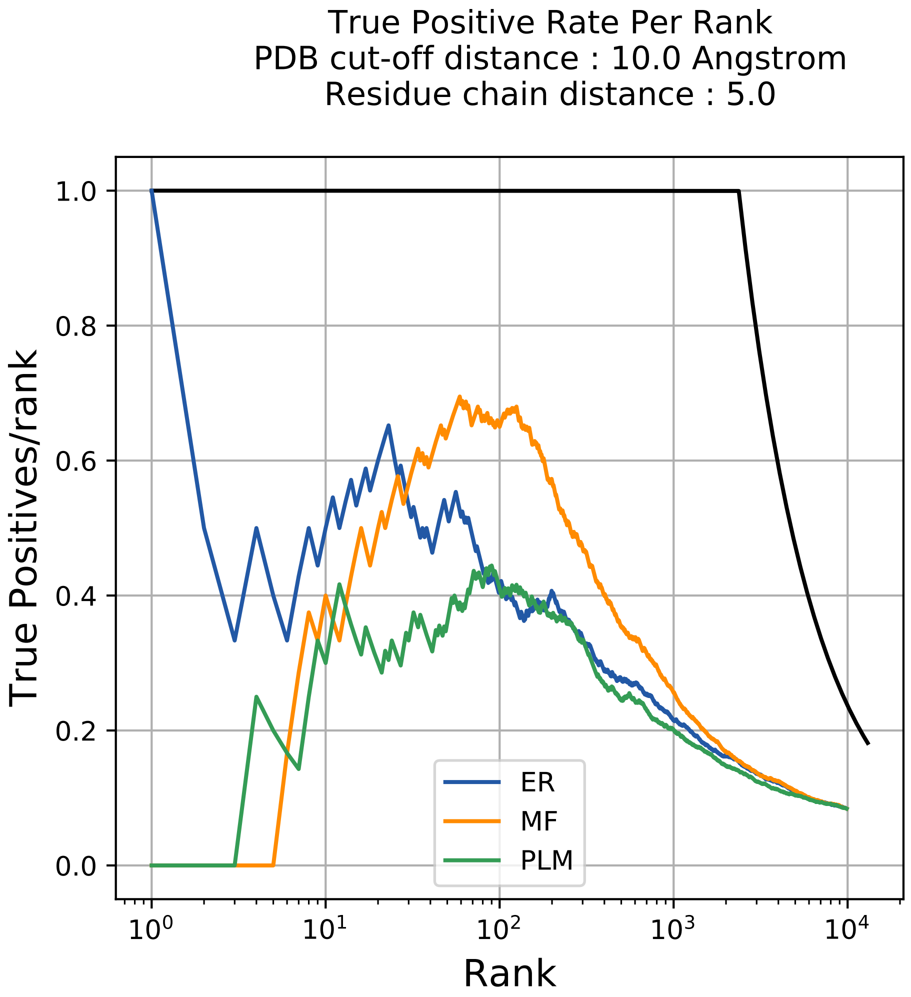

True Positive Rate Per Rank
PDB cut-off distance : 10.0 Angstrom
Residue chain distance : 5.0
True Positives/rank
Rank
1.0
0.8
0.6
0.4
0.2
0.0
$10^0$
$10^1$
$10^2$
$10^3$
$10^4$
ER
MF
PLM

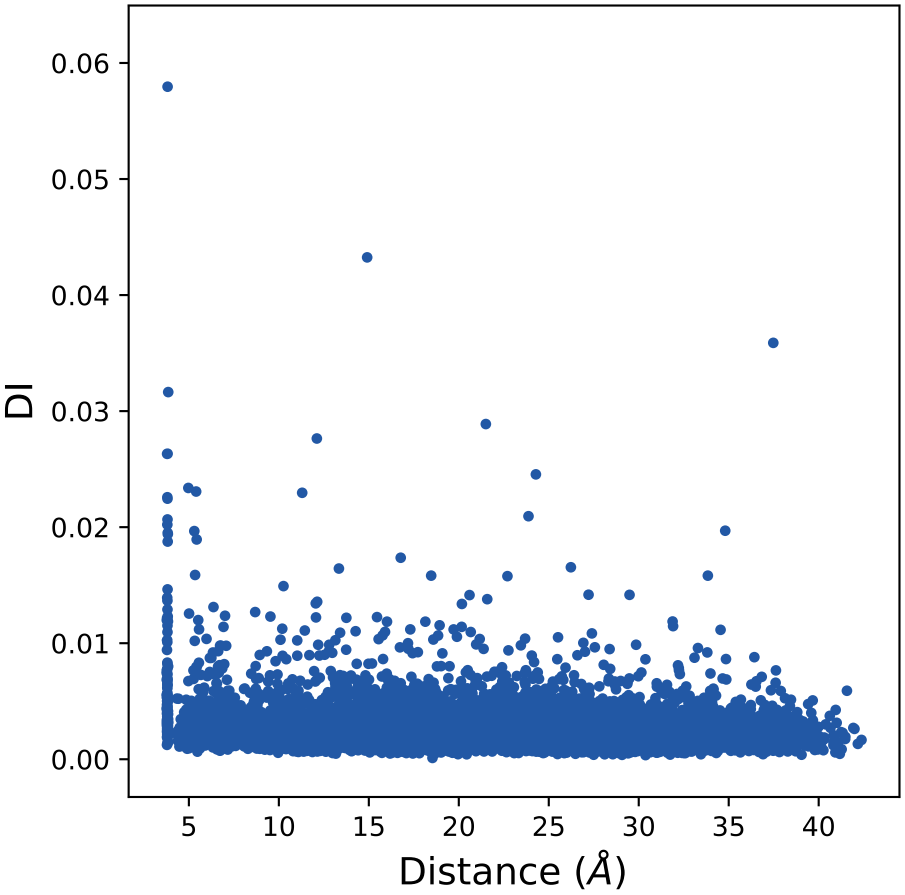
0.06
0.05
0.04
0.03
0.02
0.01
0.00
DI
5
10
15
20
25
30
35
40
Distance (Å)

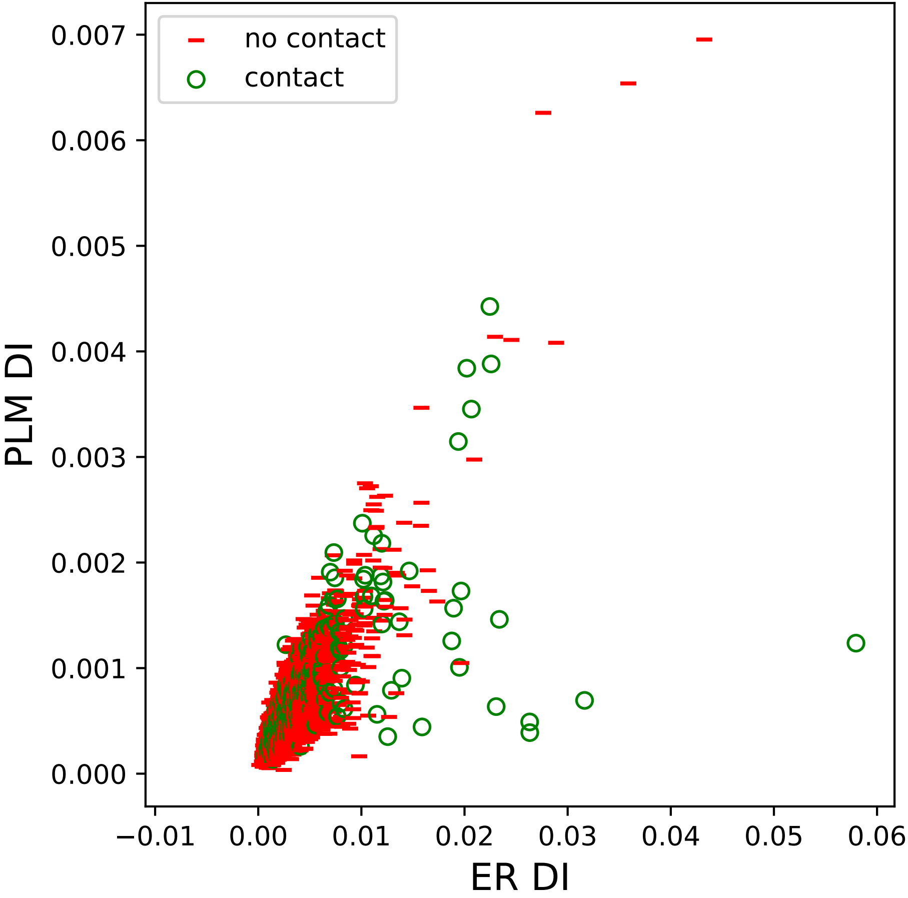

no contact
contact
PLM DI
ER DI
0.000
0.001
0.002
0.003
0.004
0.005
0.006
0.007
−0.01
0.00
0.01
0.02
0.03
0.04
0.05
0.06

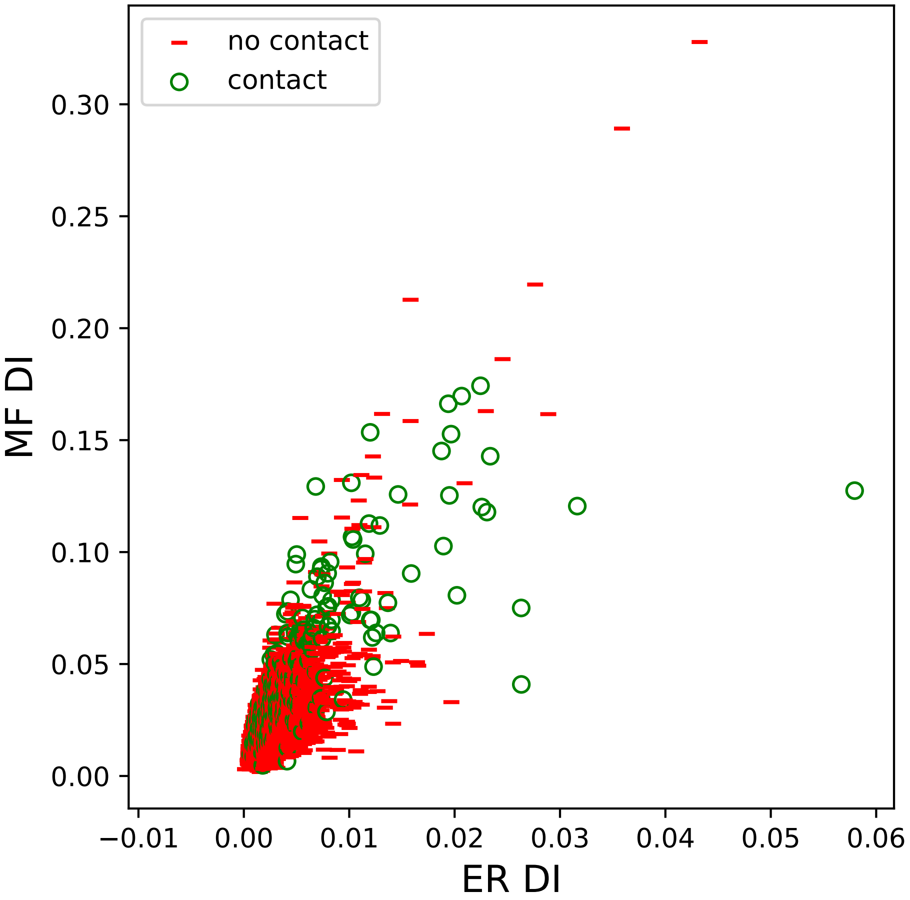
no contact
contact
0.00
0.05
0.10
0.15
0.20
0.25
0.30
MF DI
−0.01
0.00
0.01
0.02
0.03
0.04
0.05
0.06
ER DI

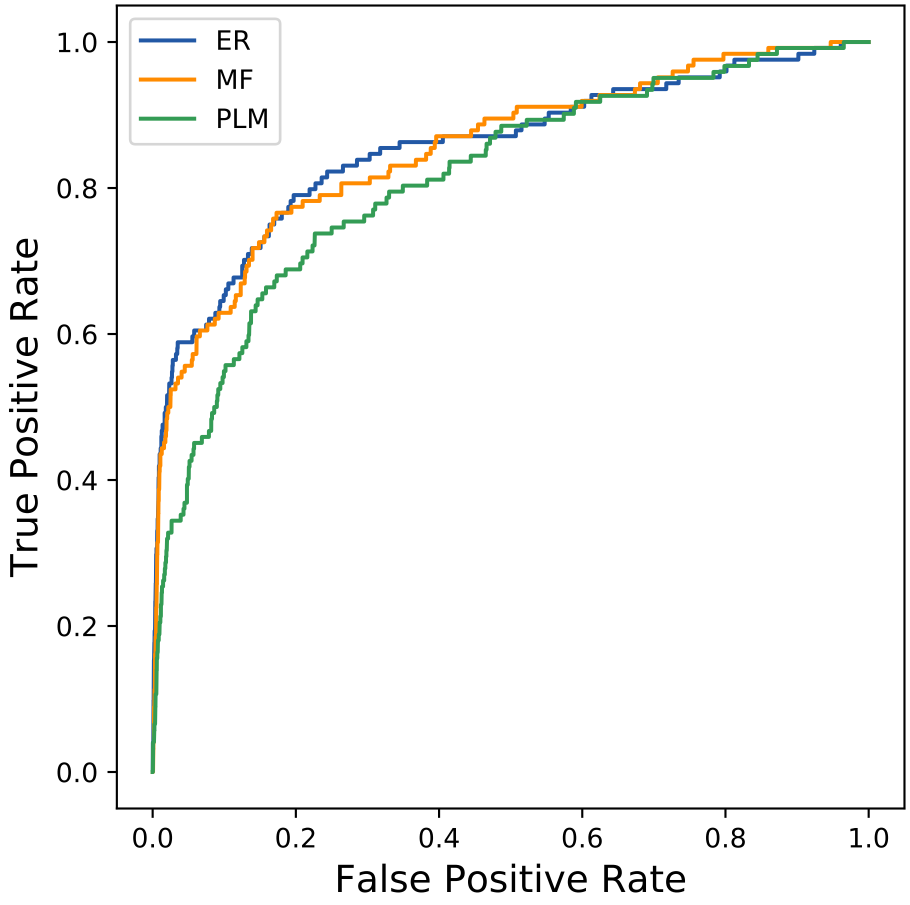

ER
MF
PLM
True Positive Rate
False Positive Rate
0.0
0.2
0.4
0.6
0.8
1.0
0.0
0.2
0.4
0.6
0.8
1.0
