## Supplemental Contact Maps for "Protein Structure Prediction with Expectation Reflection": 1258seq_101col_4jq0_PF00167_plots.pdf

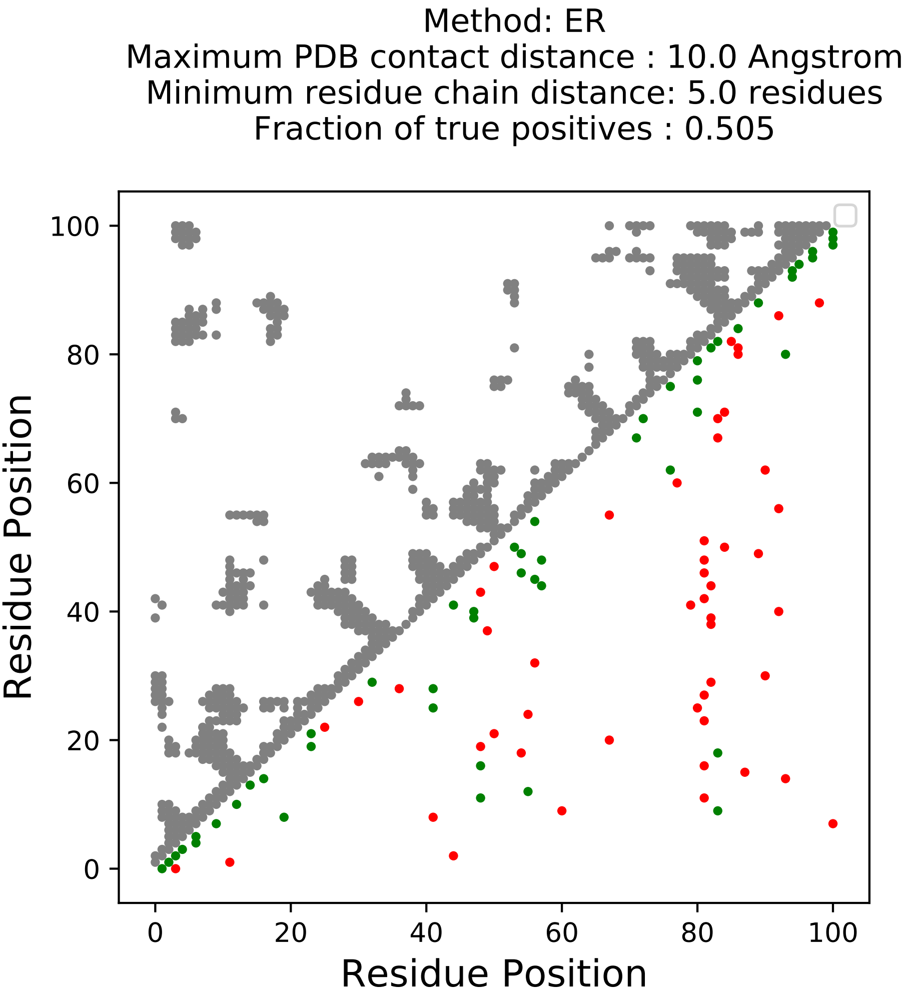
Method: ER
Maximum PDB contact distance : 10.0 Angstrom
Minimum residue chain distance: 5.0 residues
Fraction of true positives : 0.505
Residue Position
Residue Position
0
20
40
60
80
100

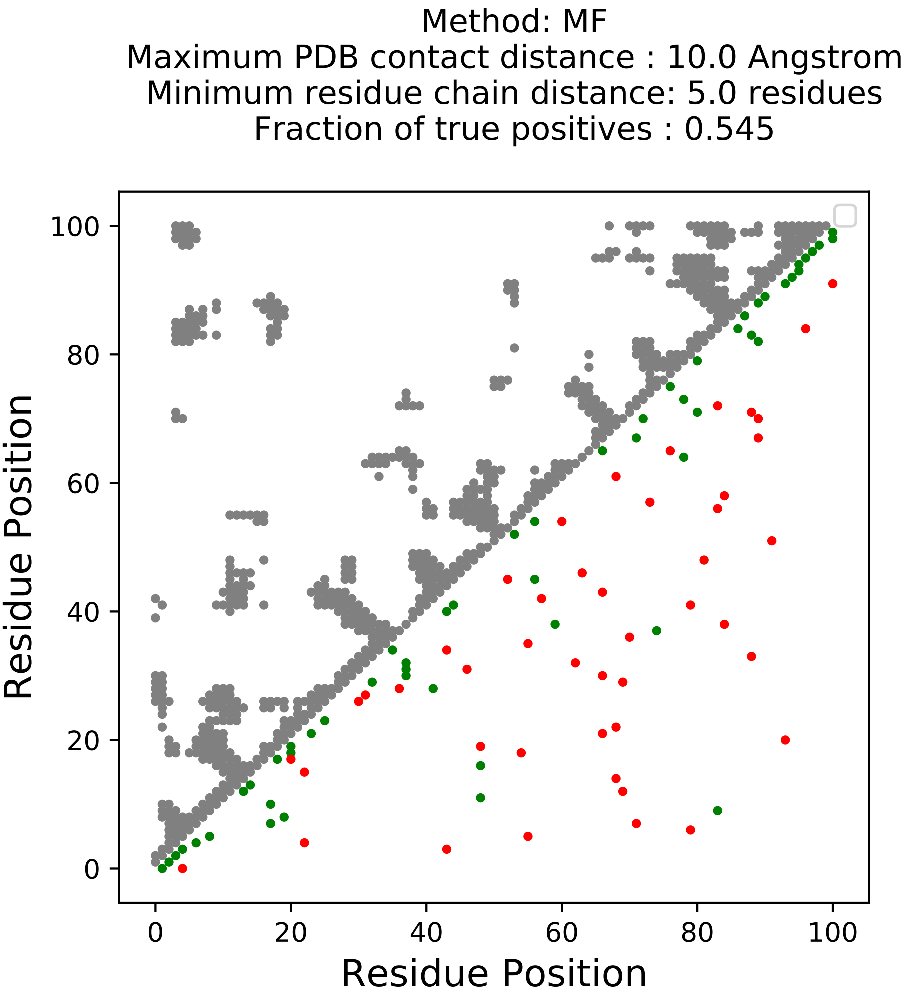
Method: MF
Maximum PDB contact distance : 10.0 Angstrom
Minimum residue chain distance: 5.0 residues
Fraction of true positives : 0.545
Residue Position
Residue Position
0
20
40
60
80
100
0
20
40
60
80
100

Method: PLM
Maximum PDB contact distance : 10.0 Angstrom
Minimum residue chain distance: 5.0 residues
Fraction of true positives : 0.465

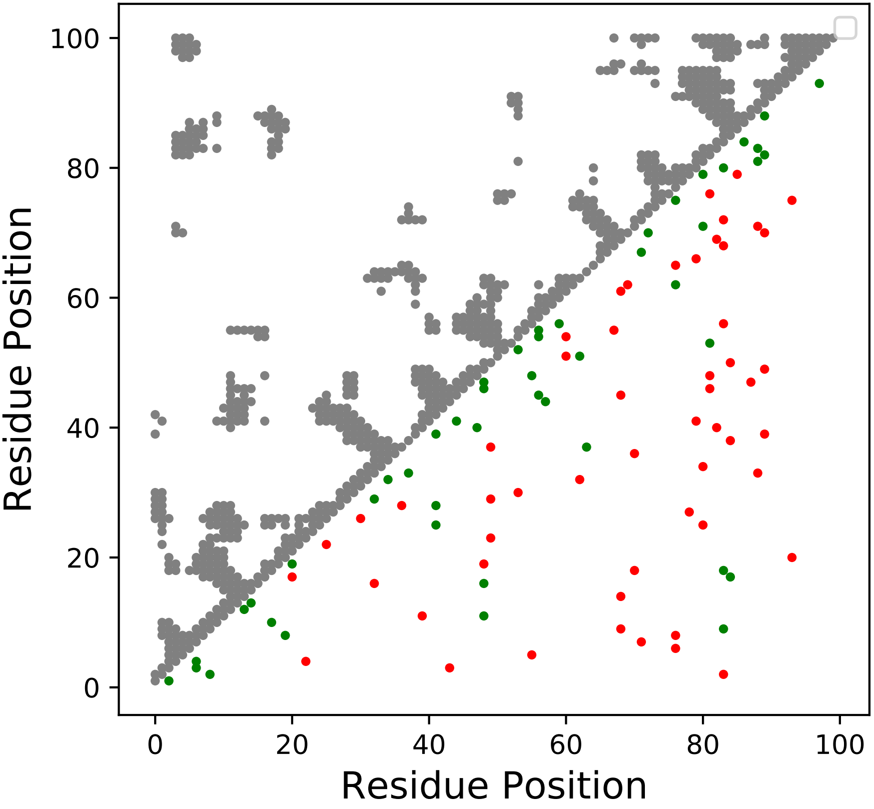

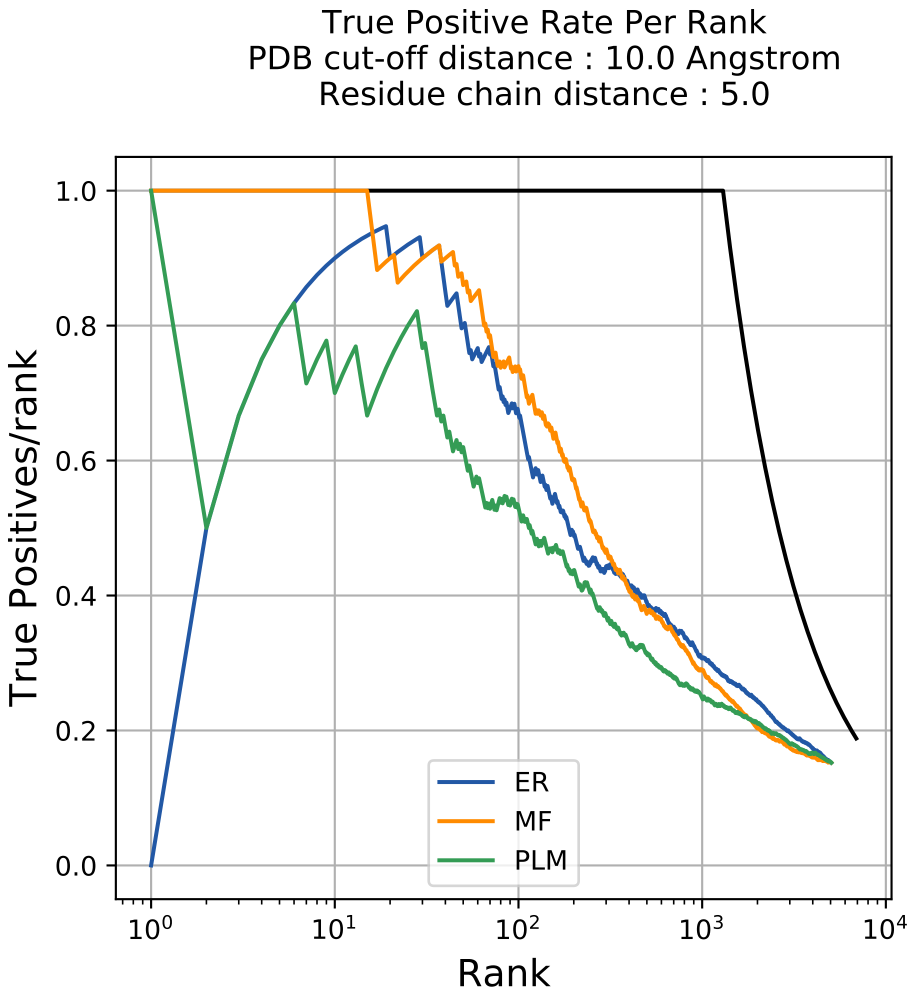
True Positive Rate Per Rank
PDB cut-off distance : 10.0 Angstrom
Residue chain distance : 5.0
True Positives/rank
Rank
1.0
0.8
0.6
0.4
0.2
0.0
$10^0$
$10^1$
$10^2$
$10^3$
$10^4$
ER
MF
PLM

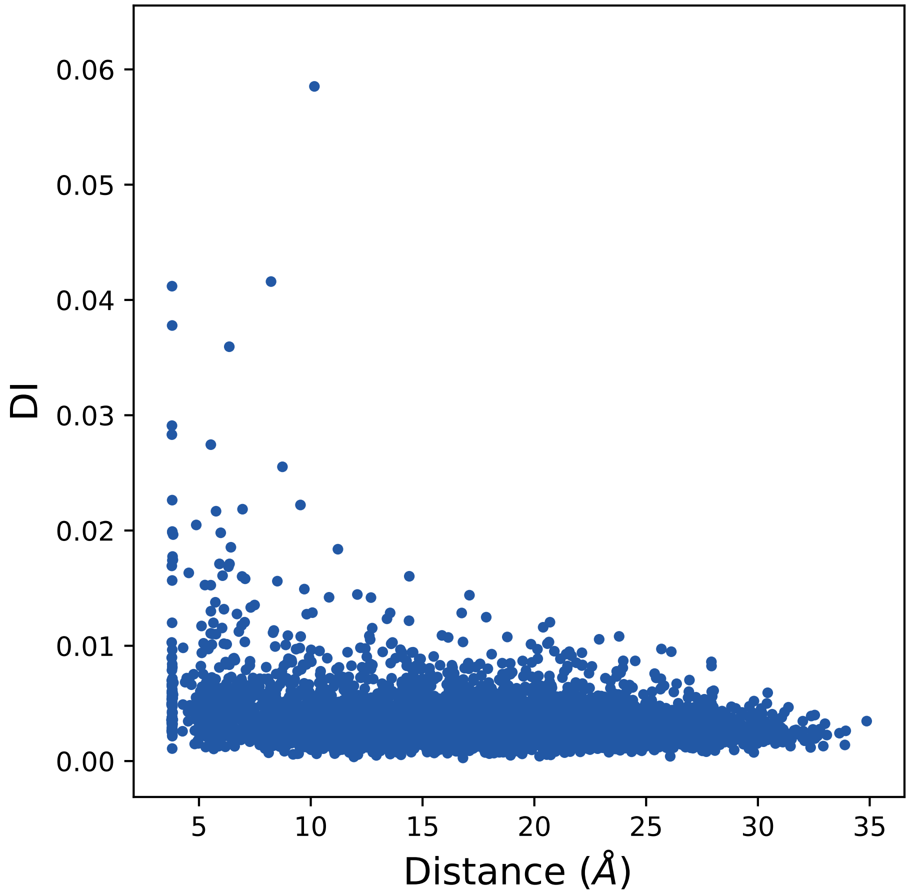
0.06
0.05
0.04
0.03
0.02
0.01
0.00
DI
5
10
15
20
25
30
35
Distance (Å)

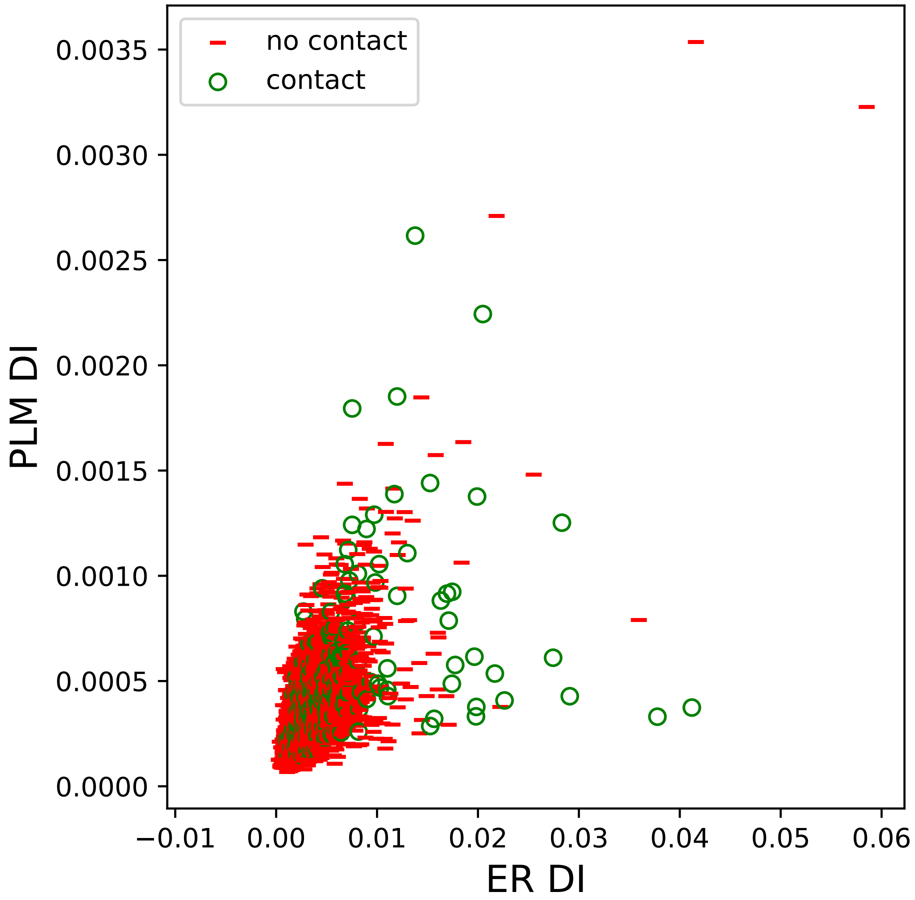

no contact
contact
PLM DI
ER DI
0.0000
0.0005
0.0010
0.0015
0.0020
0.0025
0.0030
0.0035
−0.01
0.00
0.01
0.02
0.03
0.04
0.05
0.06

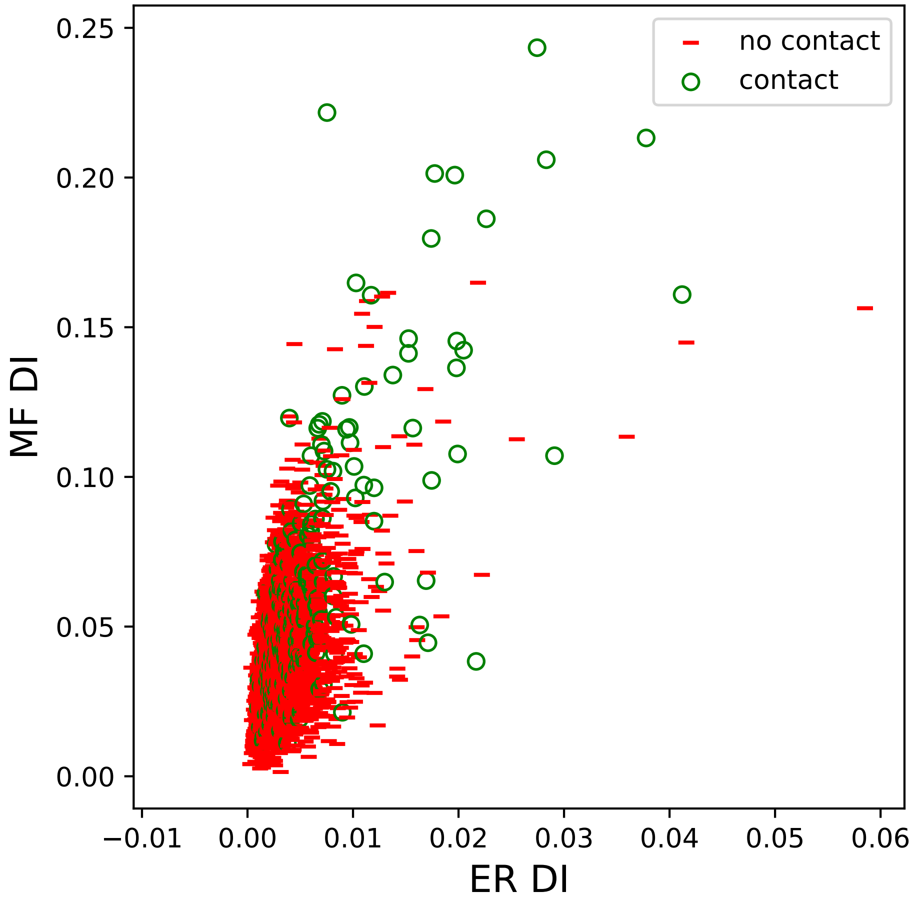

no contact
contact
MF DI
ER DI
0.25
0.20
0.15
0.10
0.05
0.00
−0.01
0.00
0.01
0.02
0.03
0.04
0.05
0.06

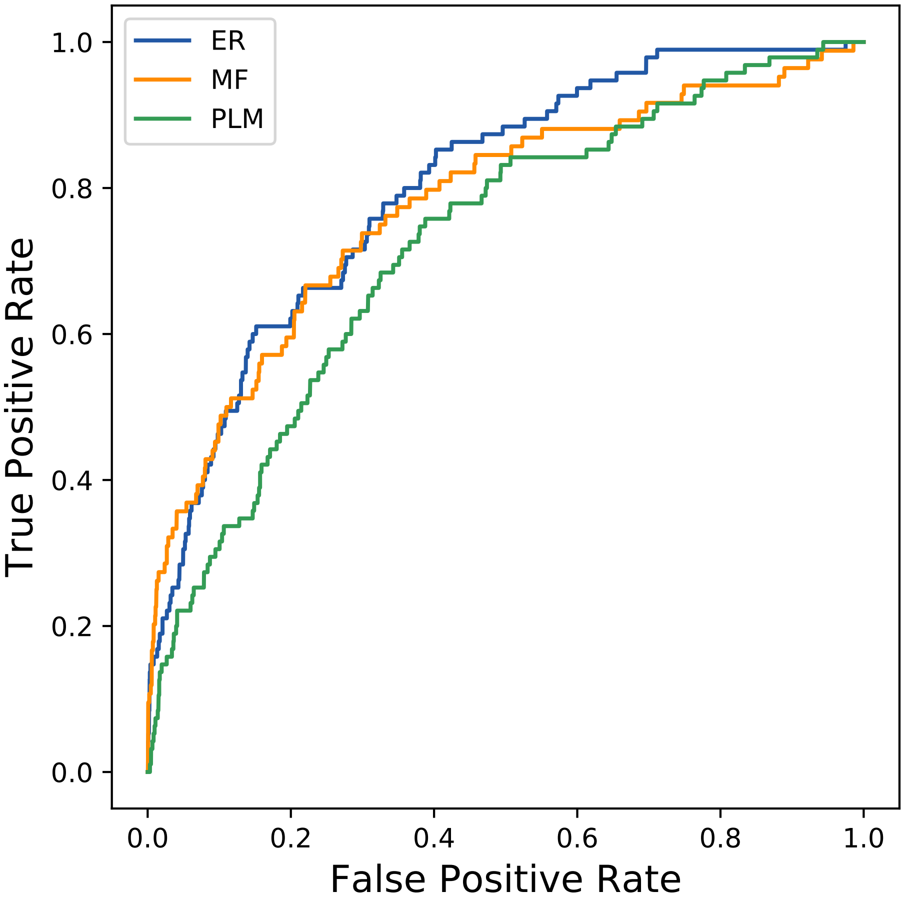
ER
MF
PLM
True Positive Rate
0.0
0.2
0.4
0.6
0.8
1.0
False Positive Rate
0.0
0.2
0.4
0.6
0.8
1.0
