## Supplemental Contact Maps for "Protein Structure Prediction with Expectation Reflection": 1386seq_72col_7jn1_PF00029_plots.pdf

Method: ER
Maximum PDB contact distance : 10.0 Angstrom
Minimum residue chain distance: 5.0 residues
Fraction of true positives : 0.486

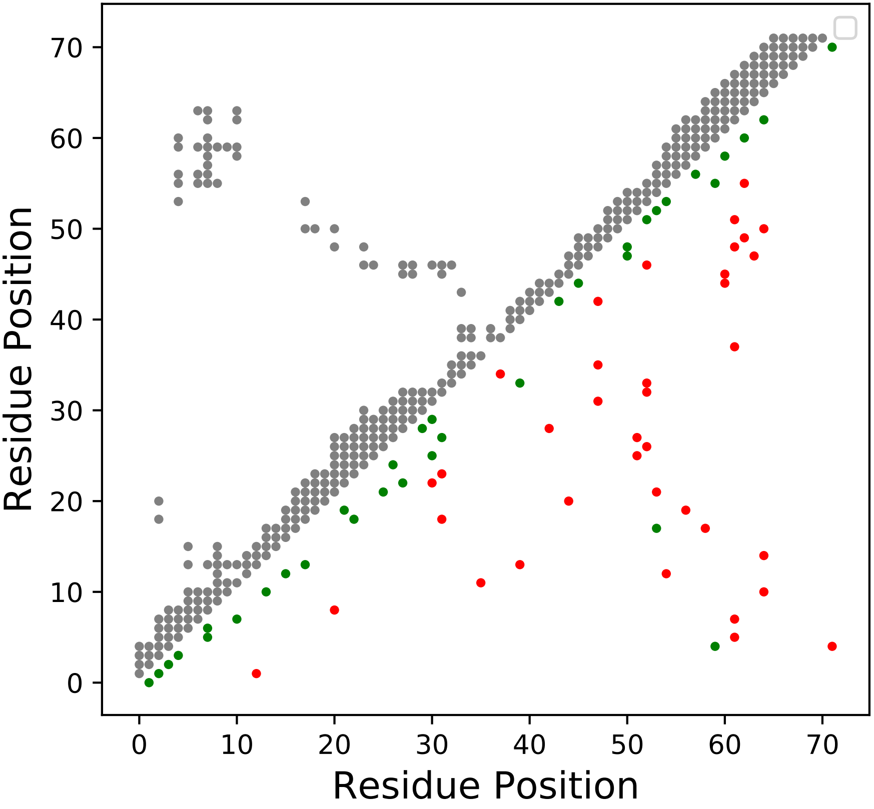

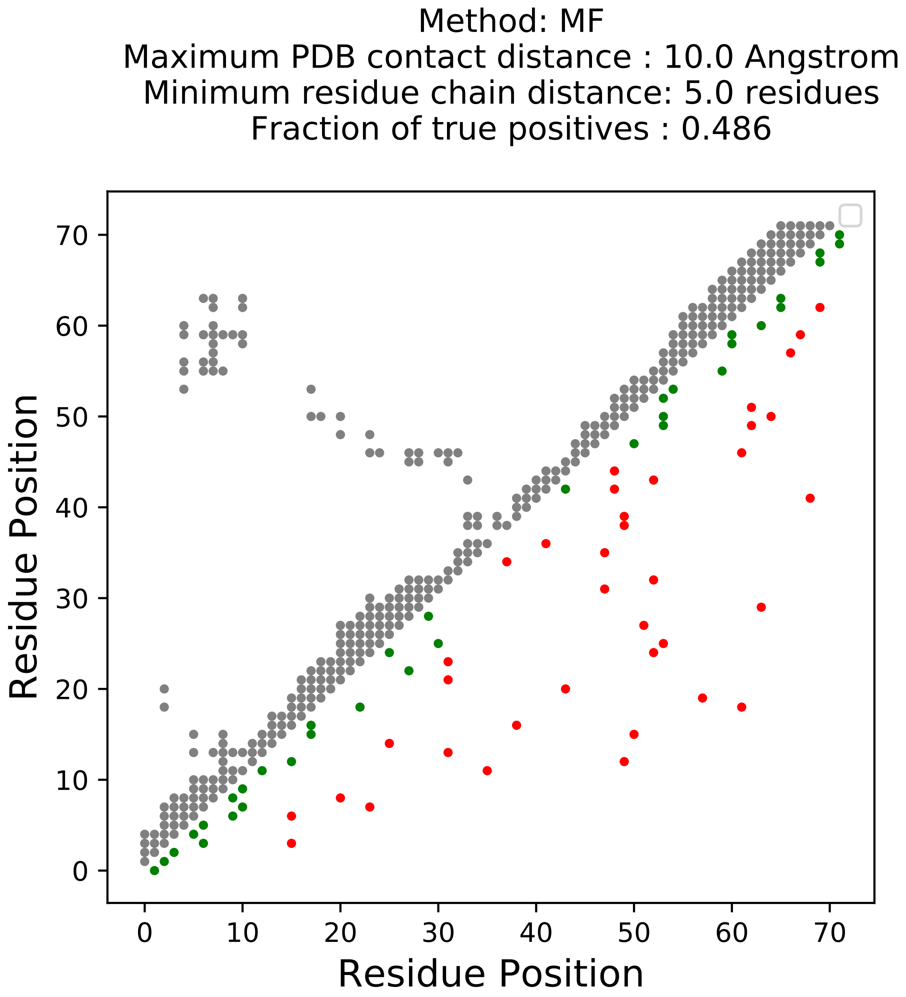
Method: MF
Maximum PDB contact distance : 10.0 Angstrom
Minimum residue chain distance: 5.0 residues
Fraction of true positives : 0.486
Residue Position
Residue Position
0
10
20
30
40
50
60
70

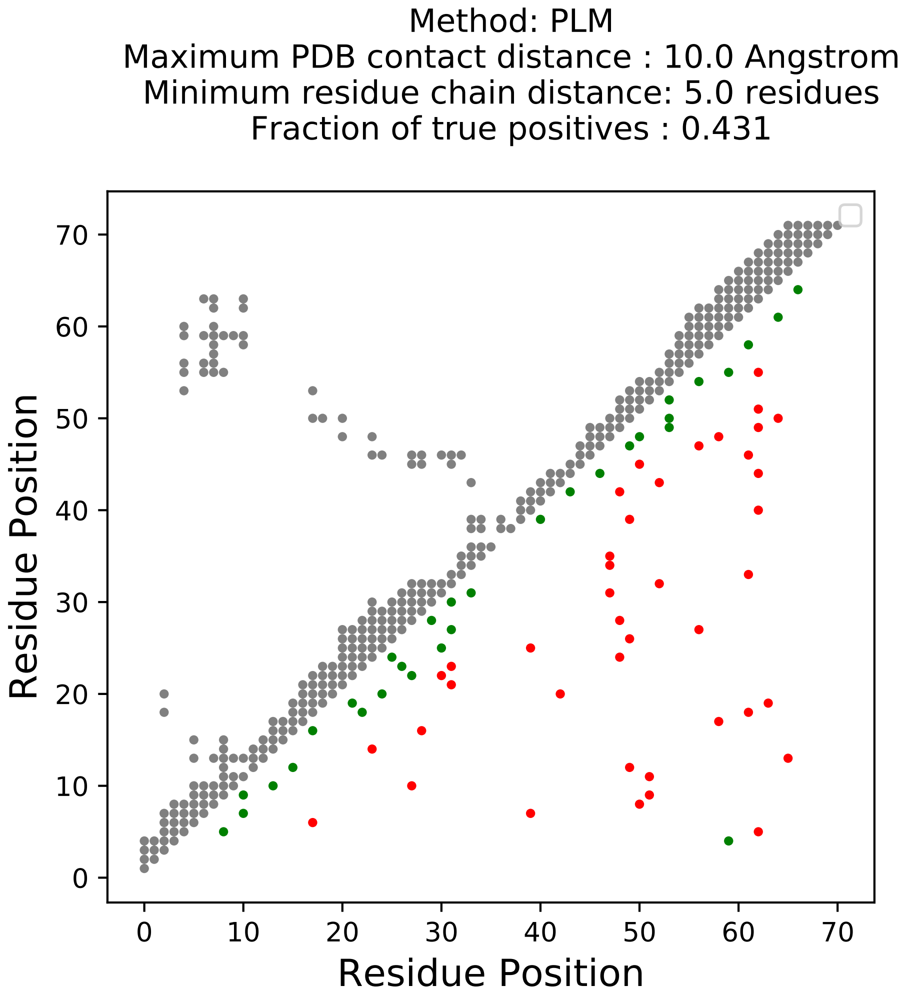

Method: PLM
Maximum PDB contact distance : 10.0 Angstrom
Minimum residue chain distance: 5.0 residues
Fraction of true positives : 0.431
Residue Position
Residue Position
0
10
20
30
40
50
60
70

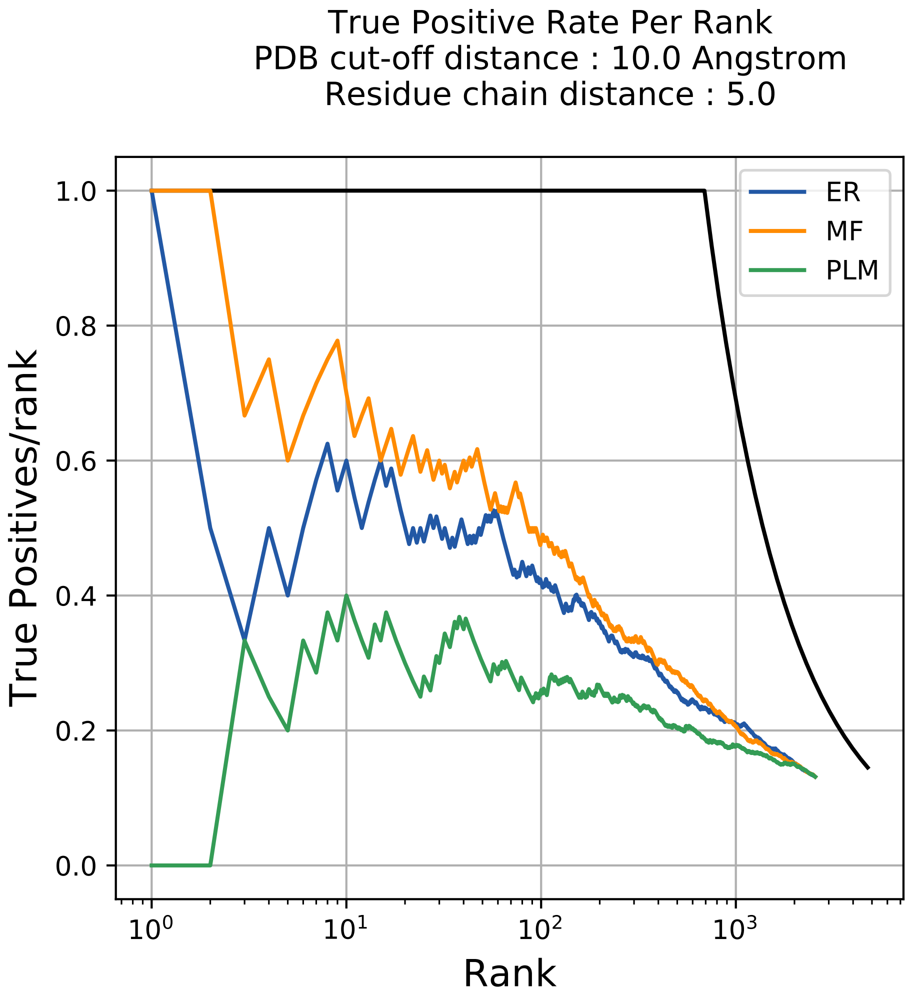
True Positive Rate Per Rank
PDB cut-off distance : 10.0 Angstrom
Residue chain distance : 5.0
True Positives/rank
Rank
ER
MF
PLM
1.0
0.8
0.6
0.4
0.2
0.0
$10^0$
$10^1$
$10^2$
$10^3$

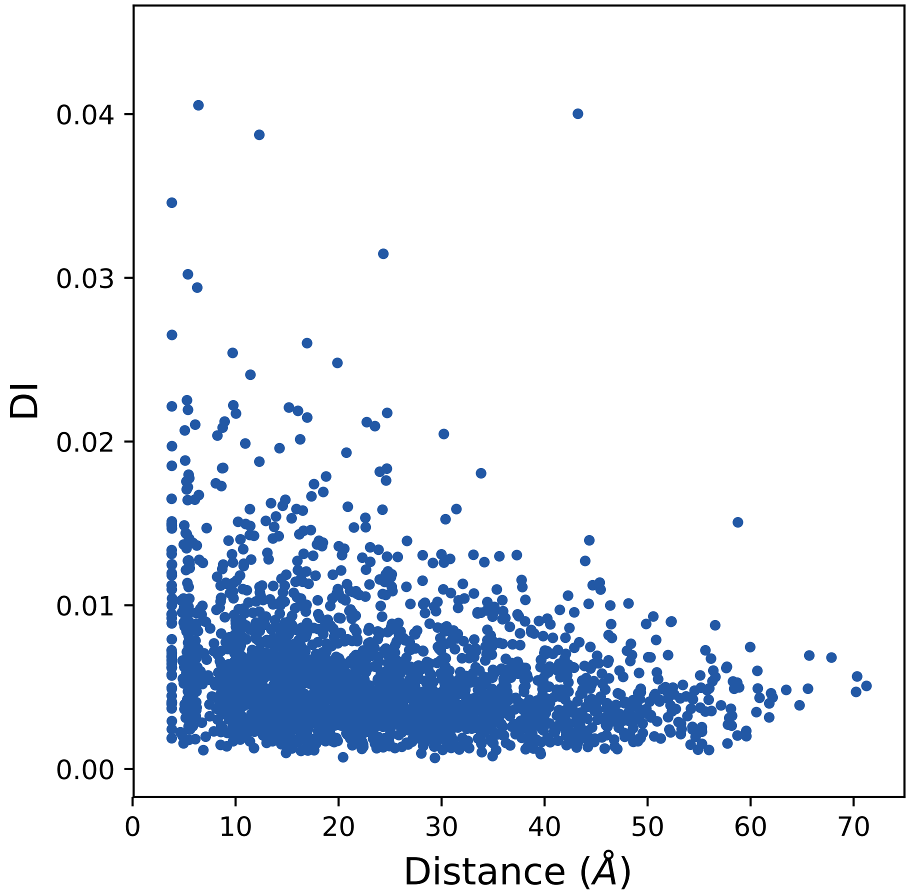

0.04
0.03
0.02
0.01
0.00
DI
0
10
20
30
40
50
60
70
Distance (Å)

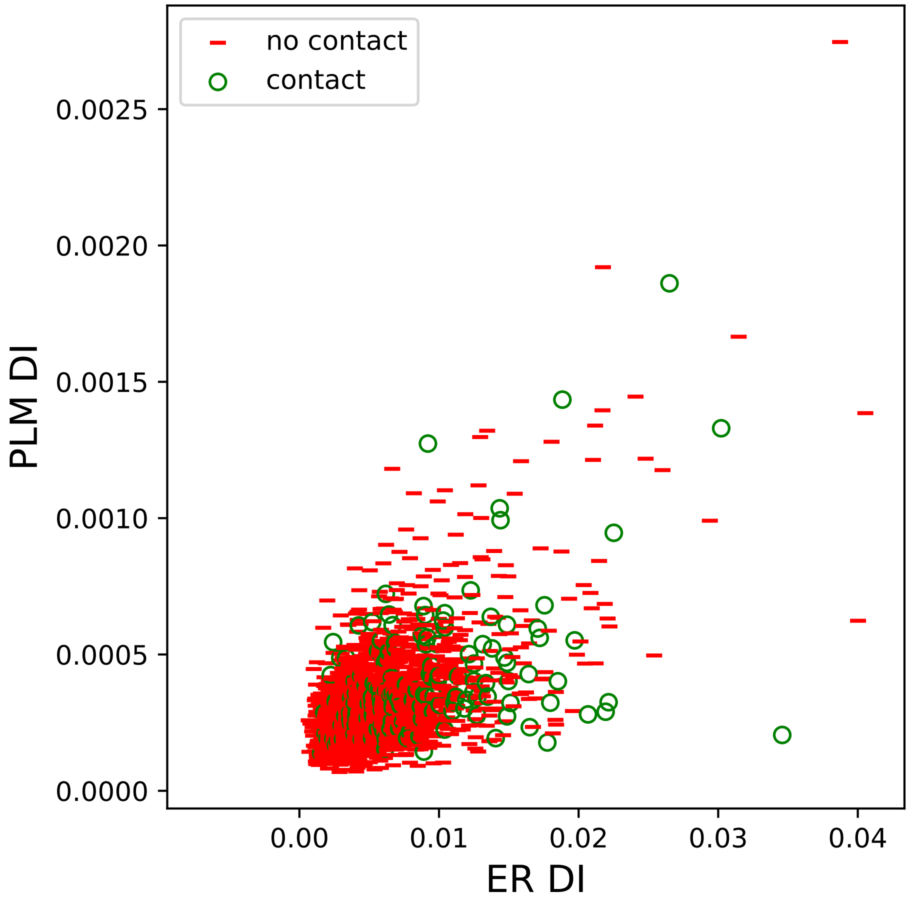
no contact
contact
0.0025
0.0020
0.0015
0.0010
0.0005
0.0000
PLM DI
0.00
0.01
0.02
0.03
0.04
ER DI

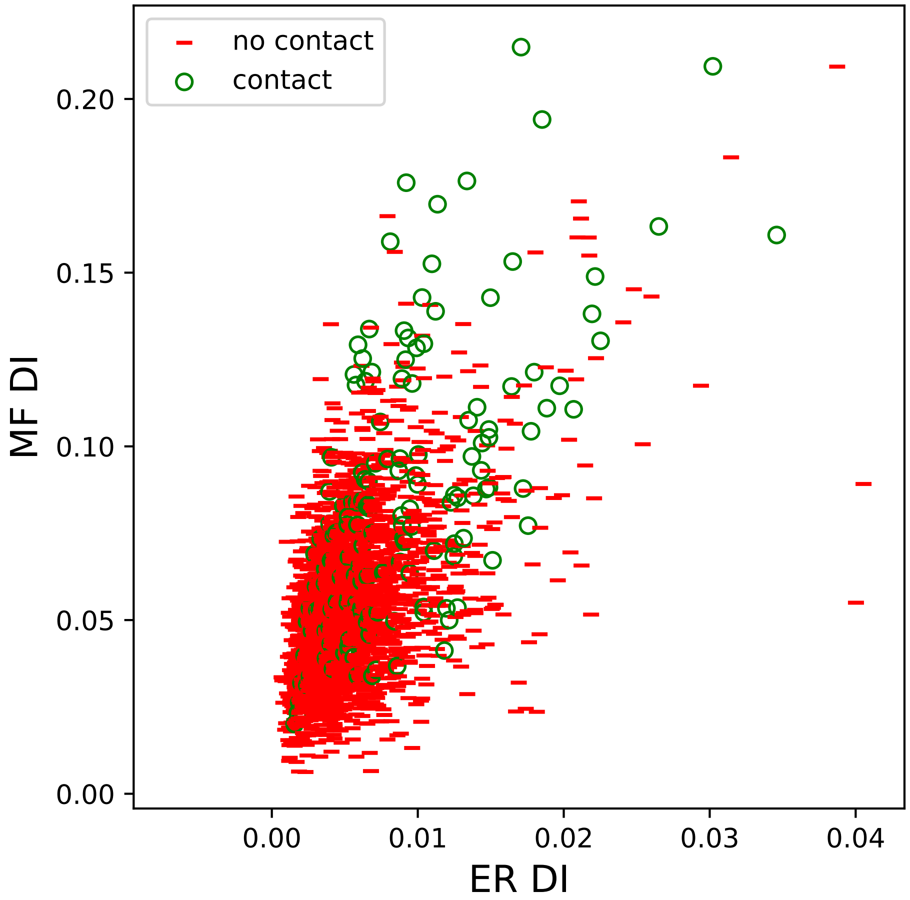
no contact
contact
0.00
0.05
0.10
0.15
0.20
MF DI
0.00
0.01
0.02
0.03
0.04
ER DI

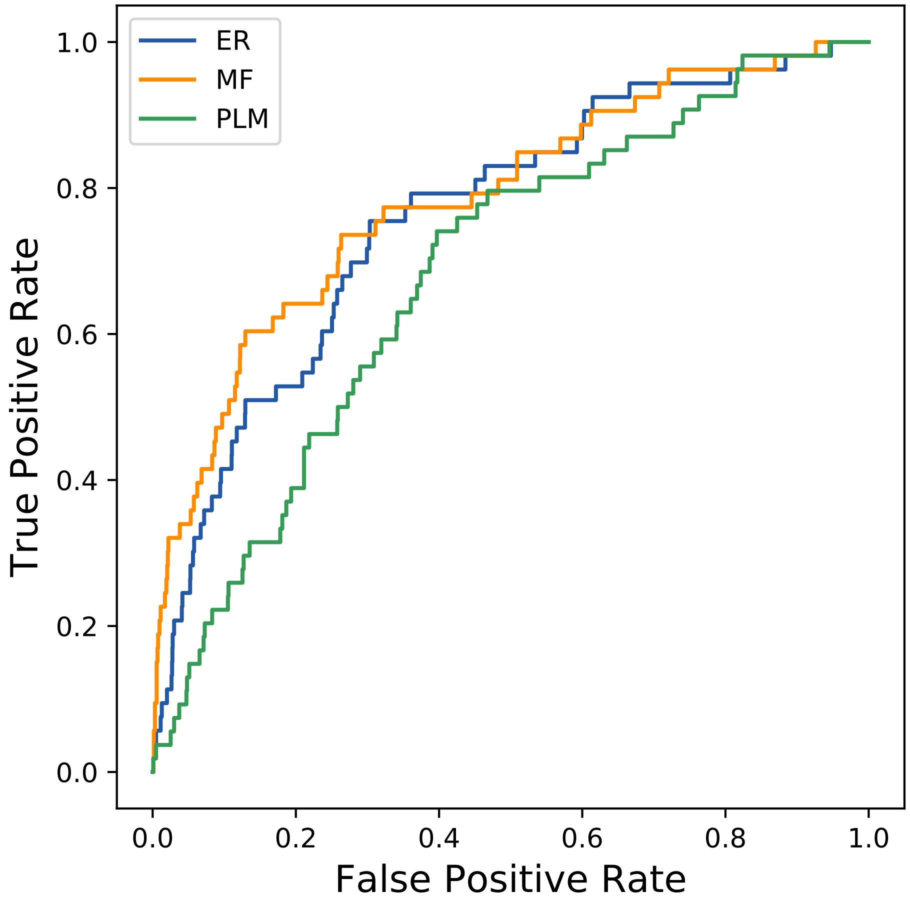

ER
MF
PLM
1.0
0.8
0.6
0.4
0.2
0.0
True Positive Rate
0.0
0.2
0.4
0.6
0.8
1.0
False Positive Rate
