## Supplemental Contact Maps for "Protein Structure Prediction with Expectation Reflection": 1397seq_136col_3pwu_PF00129_plots.pdf

Method: ER
Maximum PDB contact distance : 10.0 Angstrom
Minimum residue chain distance: 5.0 residues
Fraction of true positives : 0.551

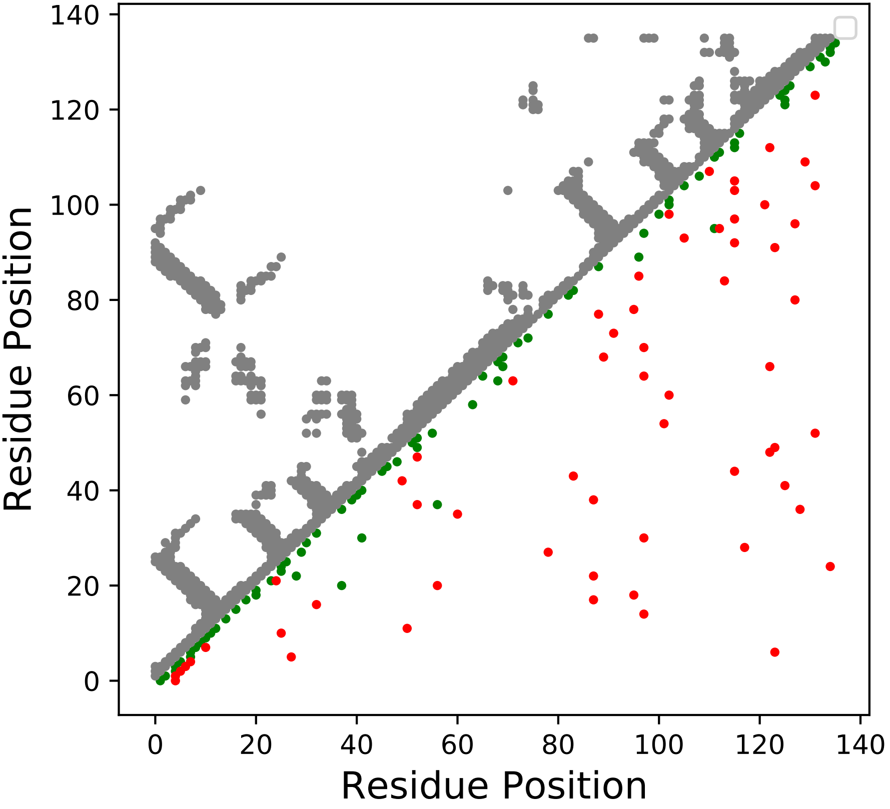

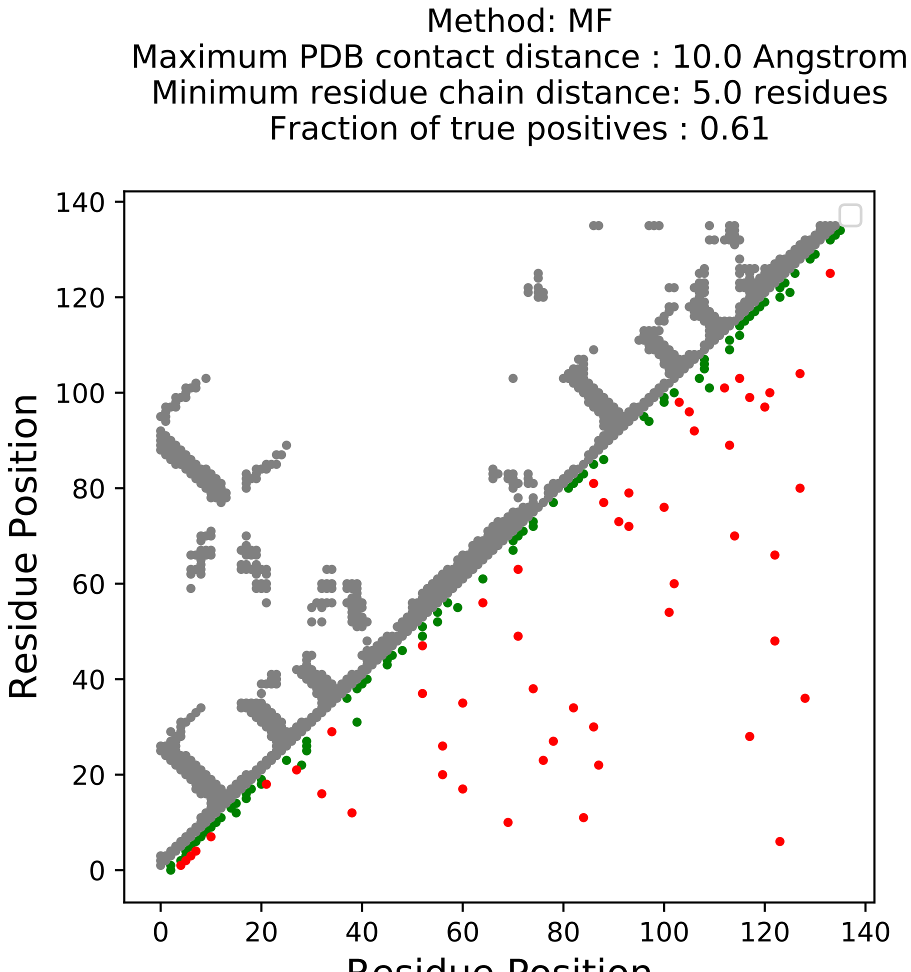
Method: MF
Maximum PDB contact distance : 10.0 Angstrom
Minimum residue chain distance: 5.0 residues
Fraction of true positives : 0.61
Residue Position
Residue Position
0
20
40
60
80
100
120
140

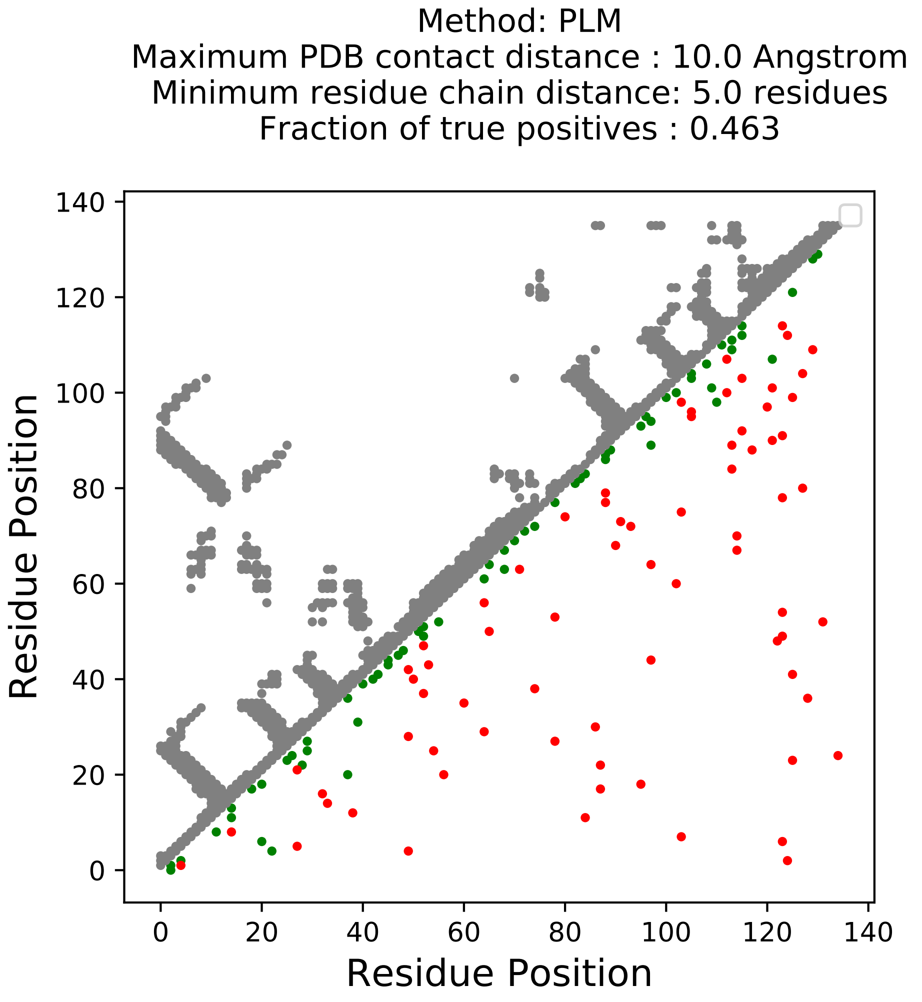
Method: PLM
Maximum PDB contact distance : 10.0 Angstrom
Minimum residue chain distance: 5.0 residues
Fraction of true positives : 0.463
Residue Position
Residue Position
0
20
40
60
80
100
120
140

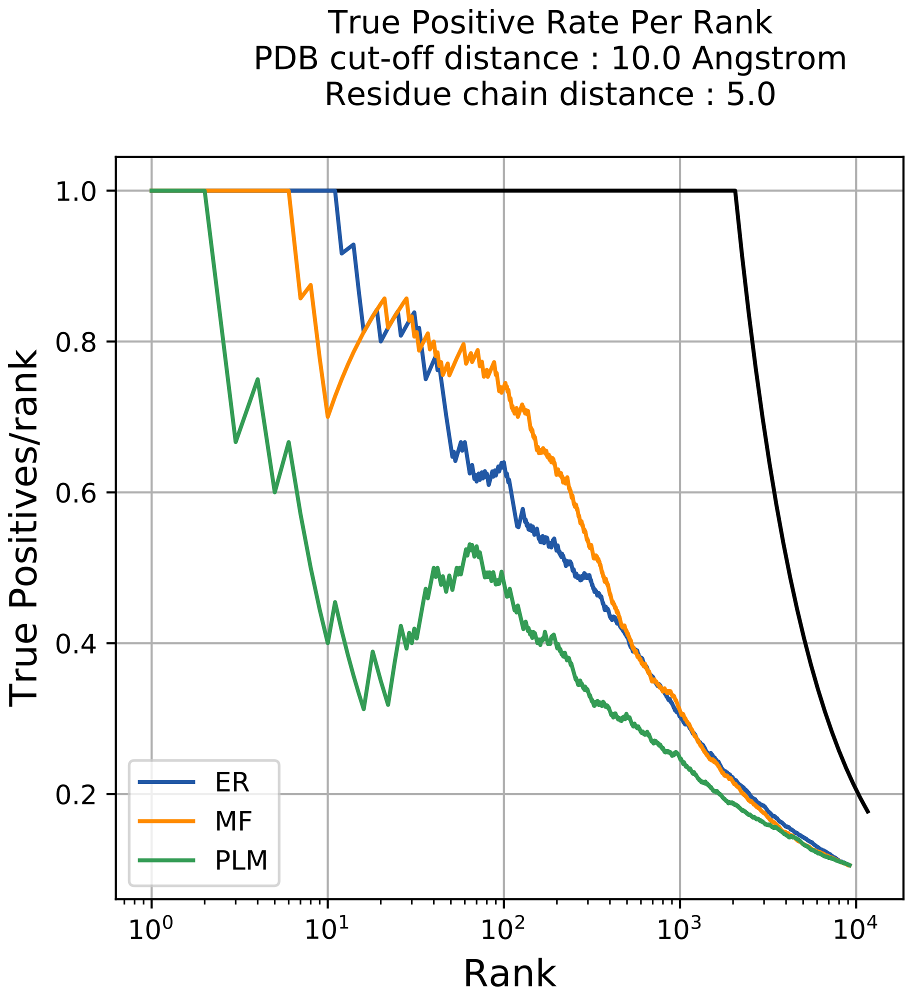
True Positive Rate Per Rank
PDB cut-off distance : 10.0 Angstrom
Residue chain distance : 5.0
True Positives/rank
Rank
1.0
0.8
0.6
0.4
0.2
$10^0$
$10^1$
$10^2$
$10^3$
$10^4$
ER
MF
PLM

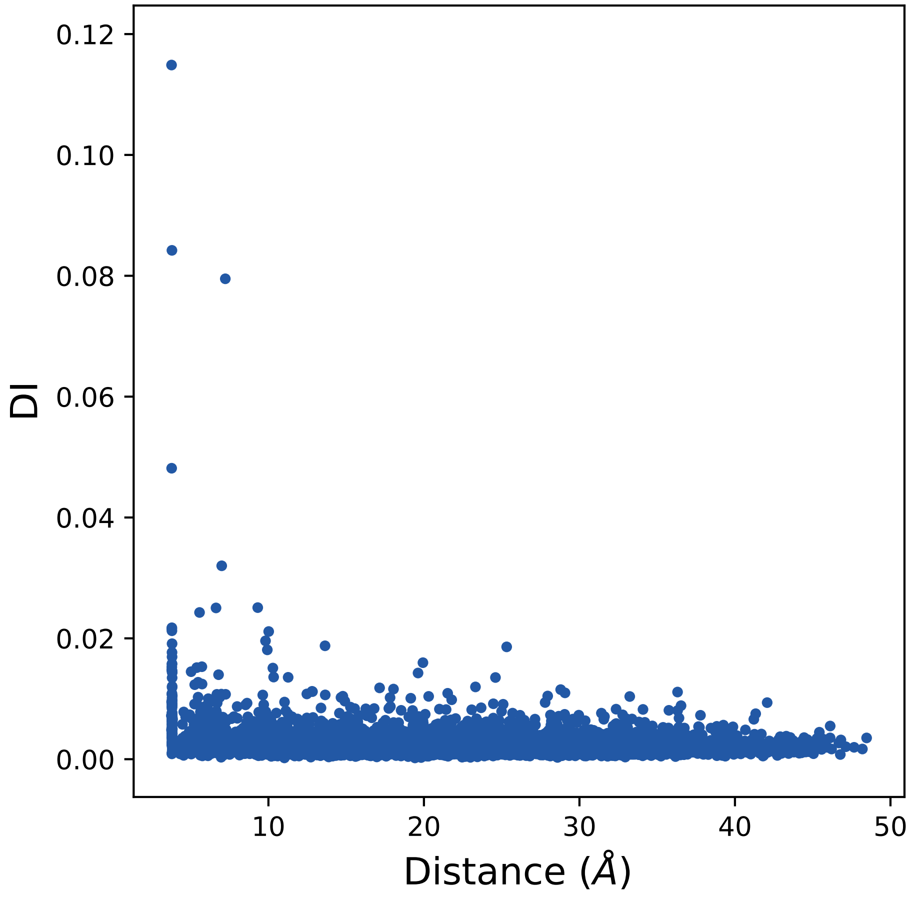
DI
0.12
0.10
0.08
0.06
0.04
0.02
0.00
10
20
30
40
50
Distance (Å)

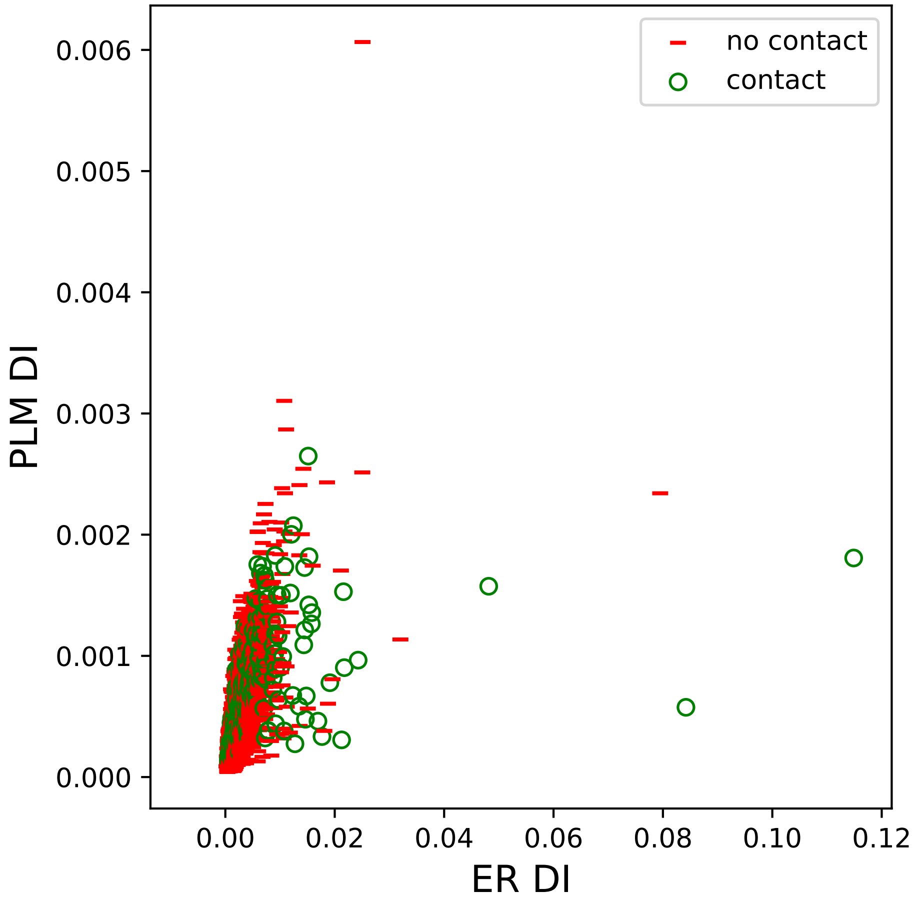

no contact
contact
0.006
0.005
0.004
0.003
0.002
0.001
0.000
PLM DI
0.00
0.02
0.04
0.06
0.08
0.10
0.12
ER DI

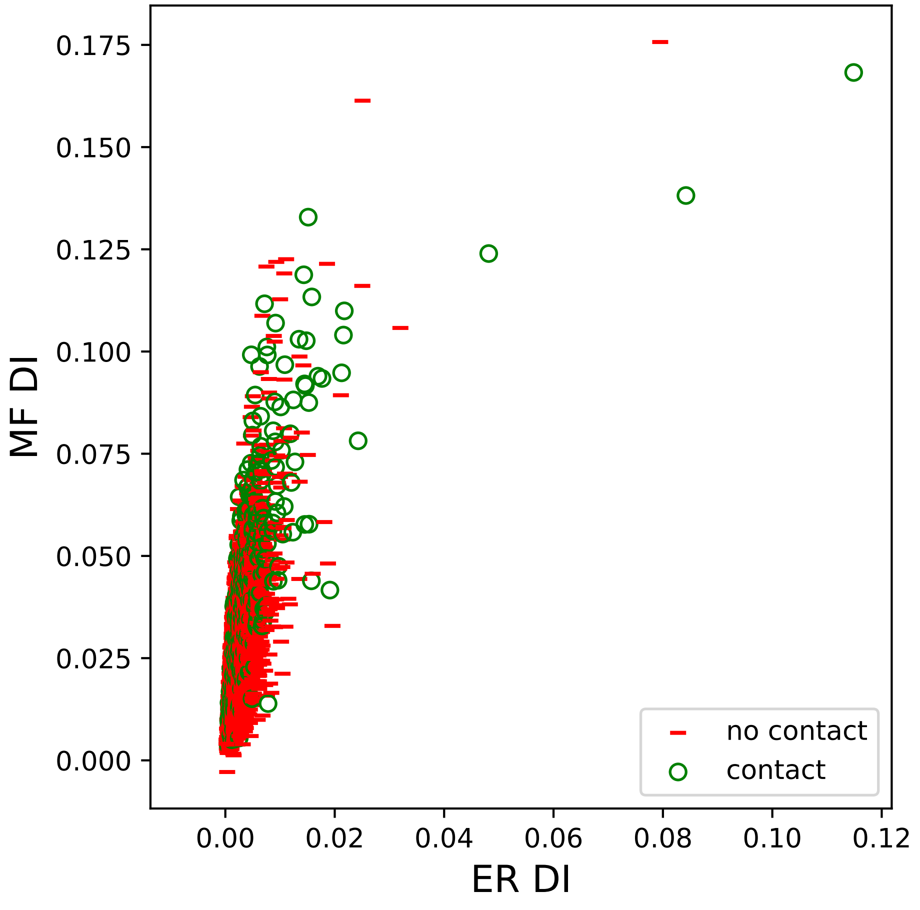

MF DI
ER DI
0.000
0.025
0.050
0.075
0.100
0.125
0.150
0.175
0.00
0.02
0.04
0.06
0.08
0.10
0.12
no contact
contact

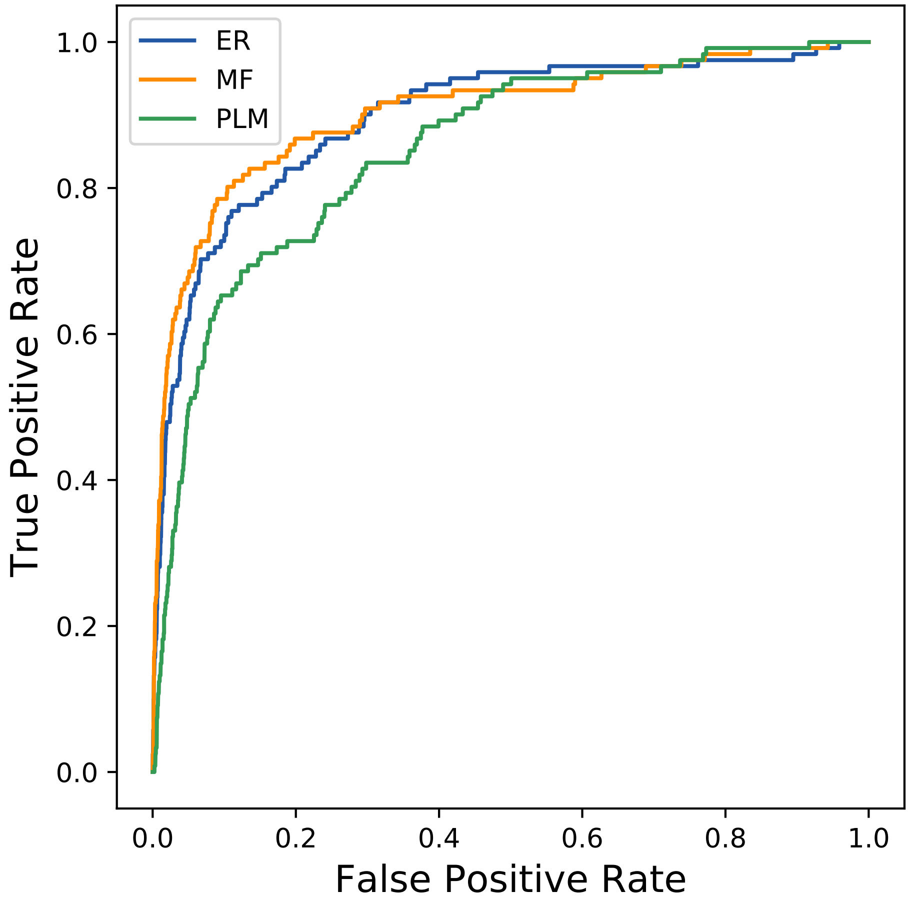
ER
MF
PLM
True Positive Rate
False Positive Rate
0.0
0.2
0.4
0.6
0.8
1.0
0.0
0.2
0.4
0.6
0.8
1.0
