## Supplemental Contact Maps for "Protein Structure Prediction with Expectation Reflection": 1423seq_156col_1i1f_PF00129_plots.pdf

Method: ER
Maximum PDB contact distance : 10.0 Angstrom
Minimum residue chain distance: 5.0 residues
Fraction of true positives : 0.609

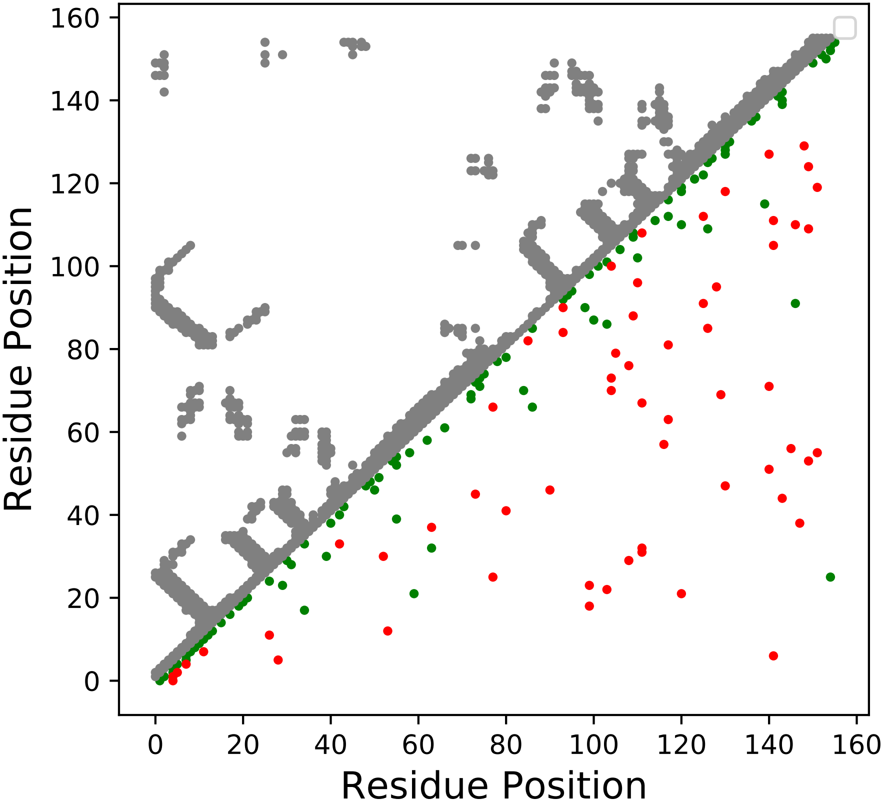

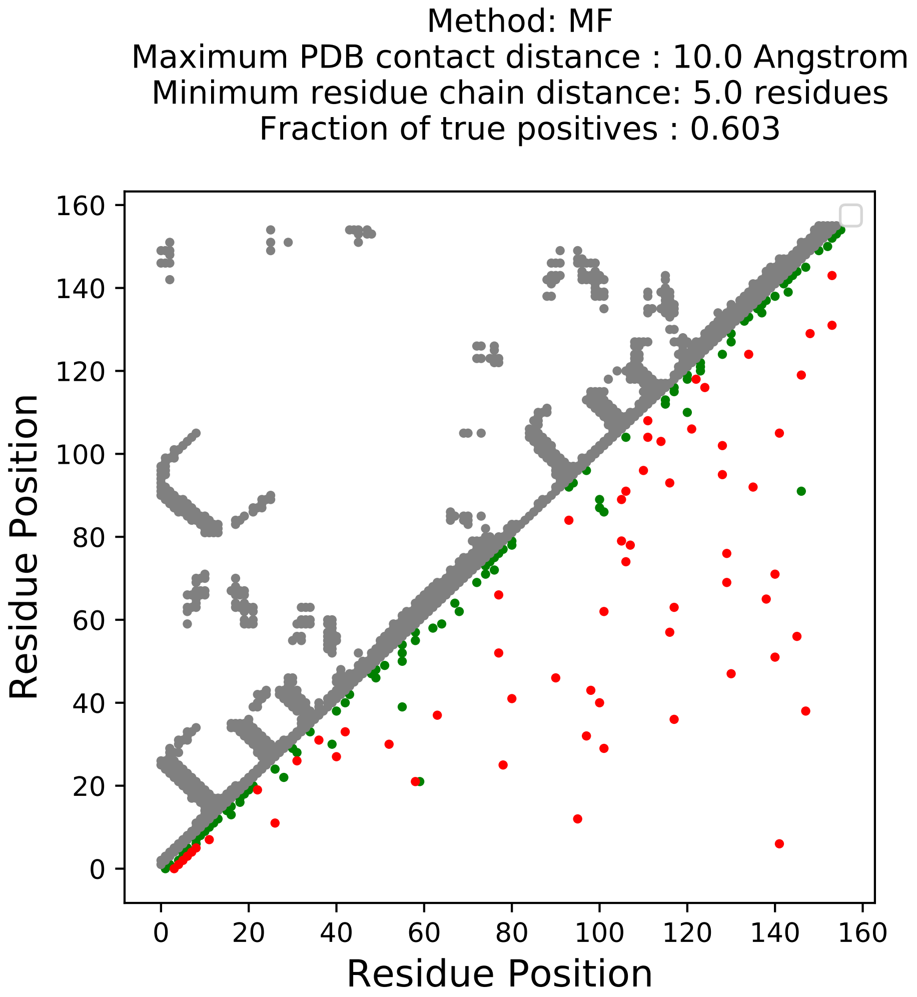
Method: MF
Maximum PDB contact distance : 10.0 Angstrom
Minimum residue chain distance: 5.0 residues
Fraction of true positives : 0.603
Residue Position
Residue Position
0
20
40
60
80
100
120
140
160

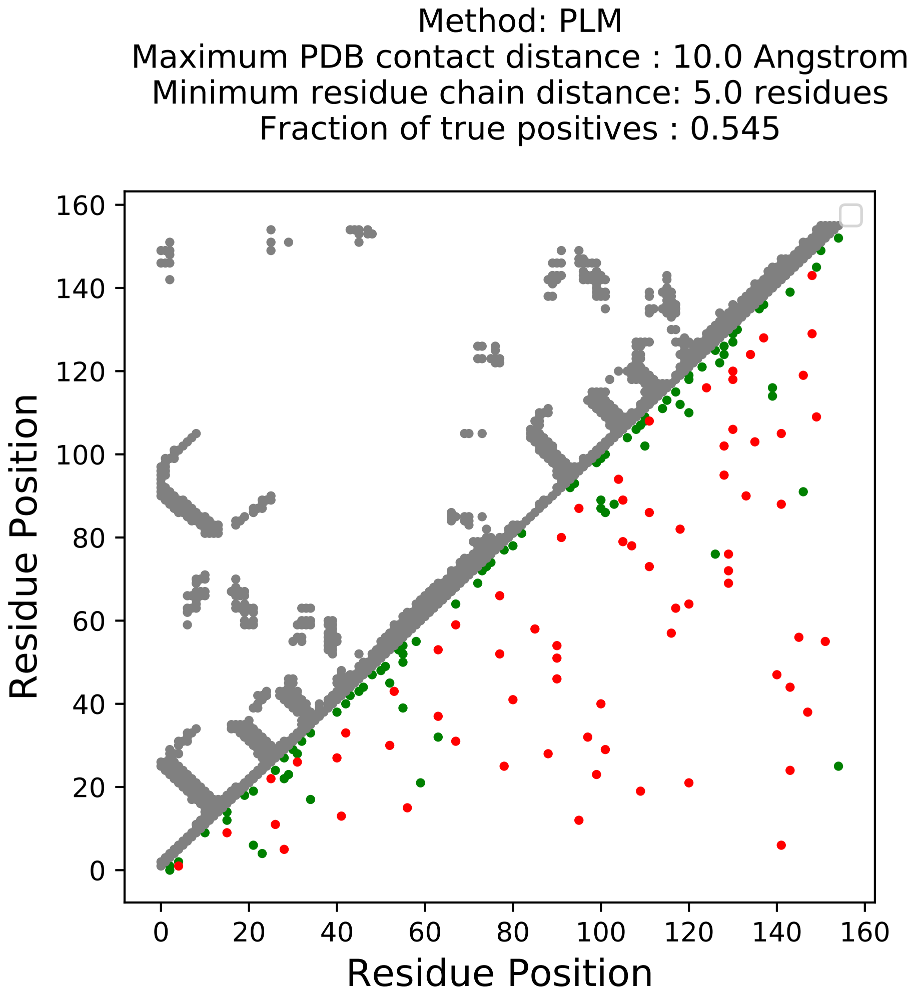
Method: PLM
Maximum PDB contact distance : 10.0 Angstrom
Minimum residue chain distance: 5.0 residues
Fraction of true positives : 0.545
Residue Position
Residue Position
0
20
40
60
80
100
120
140
160

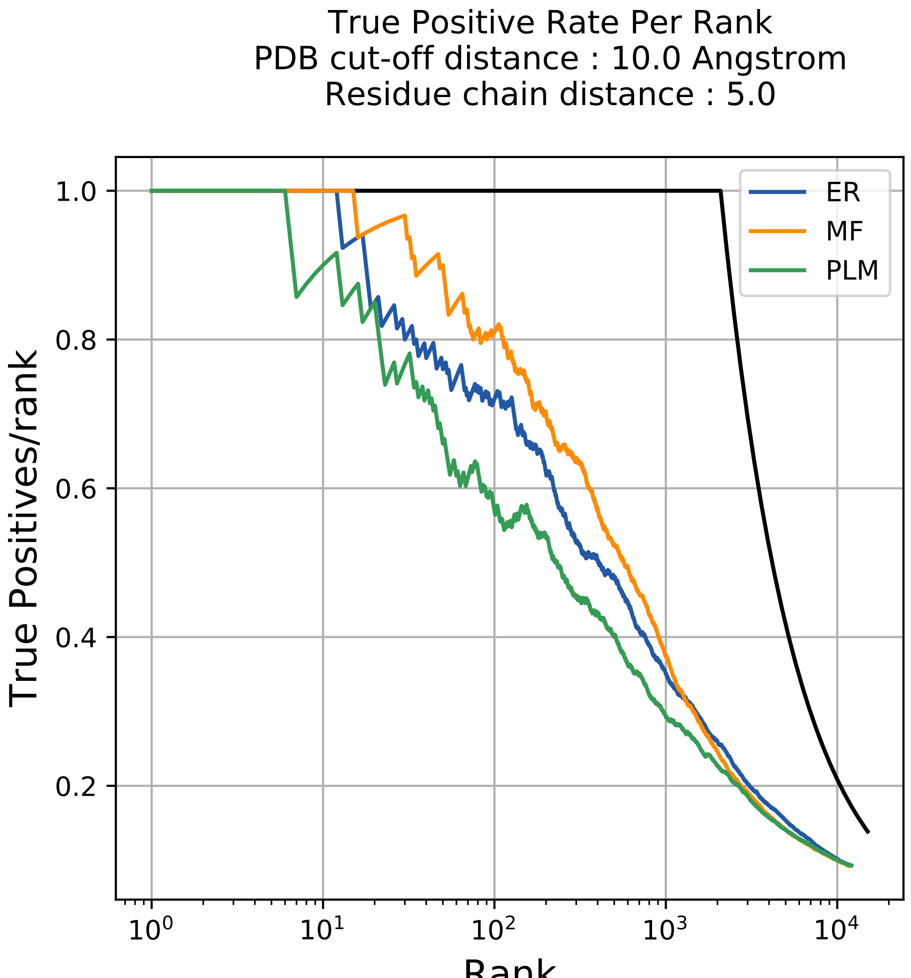
True Positive Rate Per Rank
PDB cut-off distance : 10.0 Angstrom
Residue chain distance : 5.0
True Positives/rank
Rank
1.0
0.8
0.6
0.4
0.2
$10^0$
$10^1$
$10^2$
$10^3$
$10^4$
ER
MF
PLM

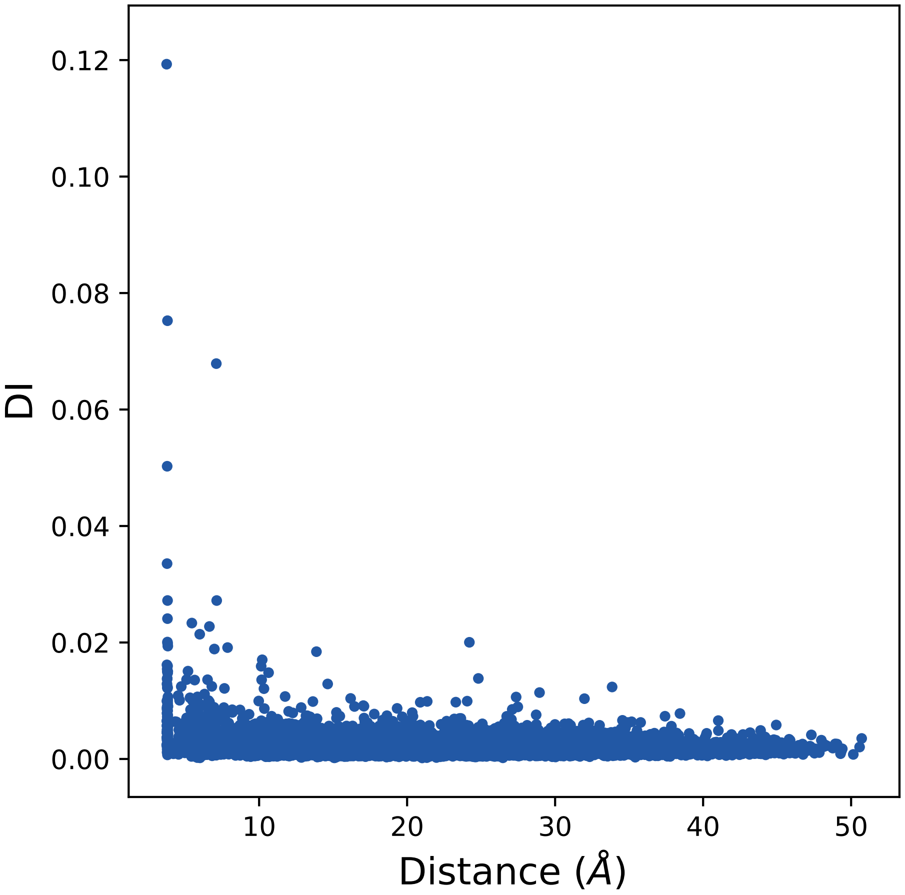
0.12
0.10
0.08
0.06
0.04
0.02
0.00
DI
10
20
30
40
50
Distance (Å)

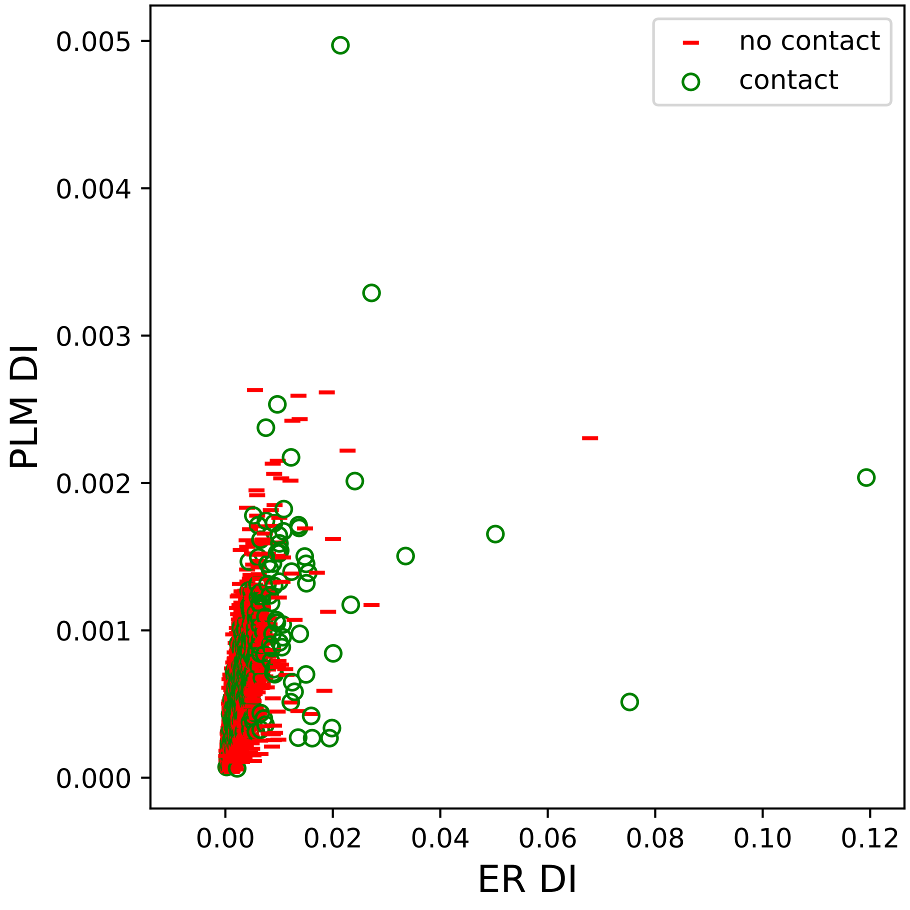
no contact
contact
0.005
0.004
0.003
0.002
0.001
0.000
PLM DI
0.00
0.02
0.04
0.06
0.08
0.10
0.12
ER DI

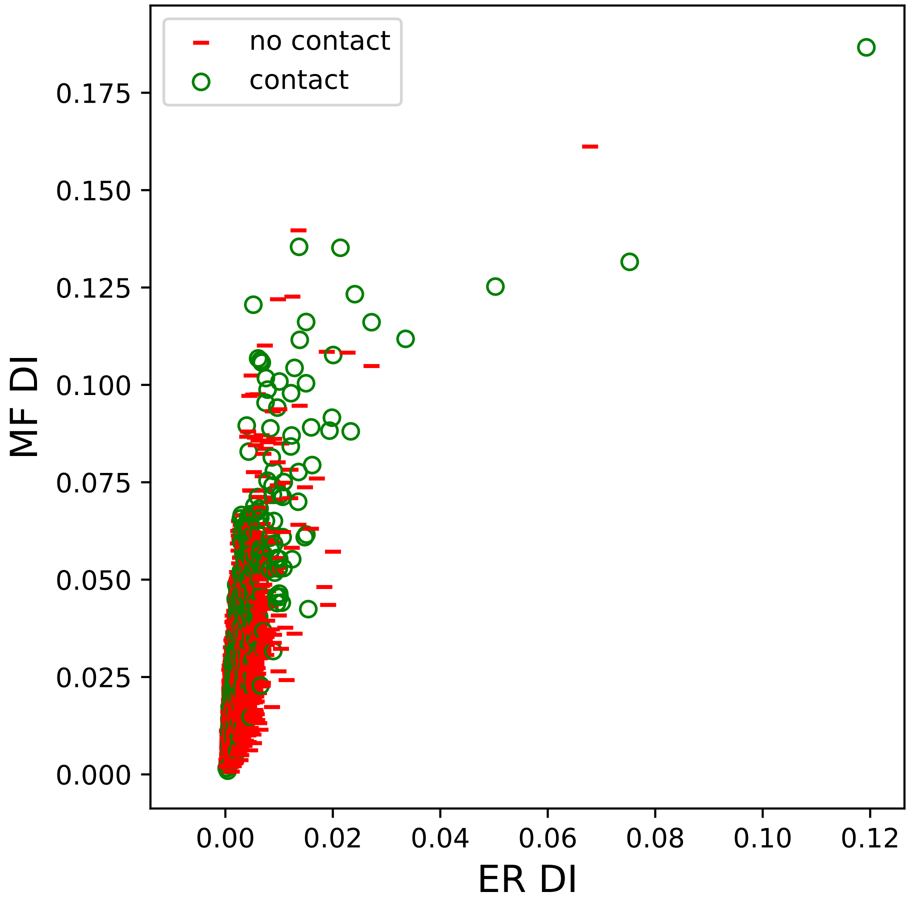
no contact
contact
MF DI
ER DI
0.000
0.025
0.050
0.075
0.100
0.125
0.150
0.175
0.00
0.02
0.04
0.06
0.08
0.10
0.12

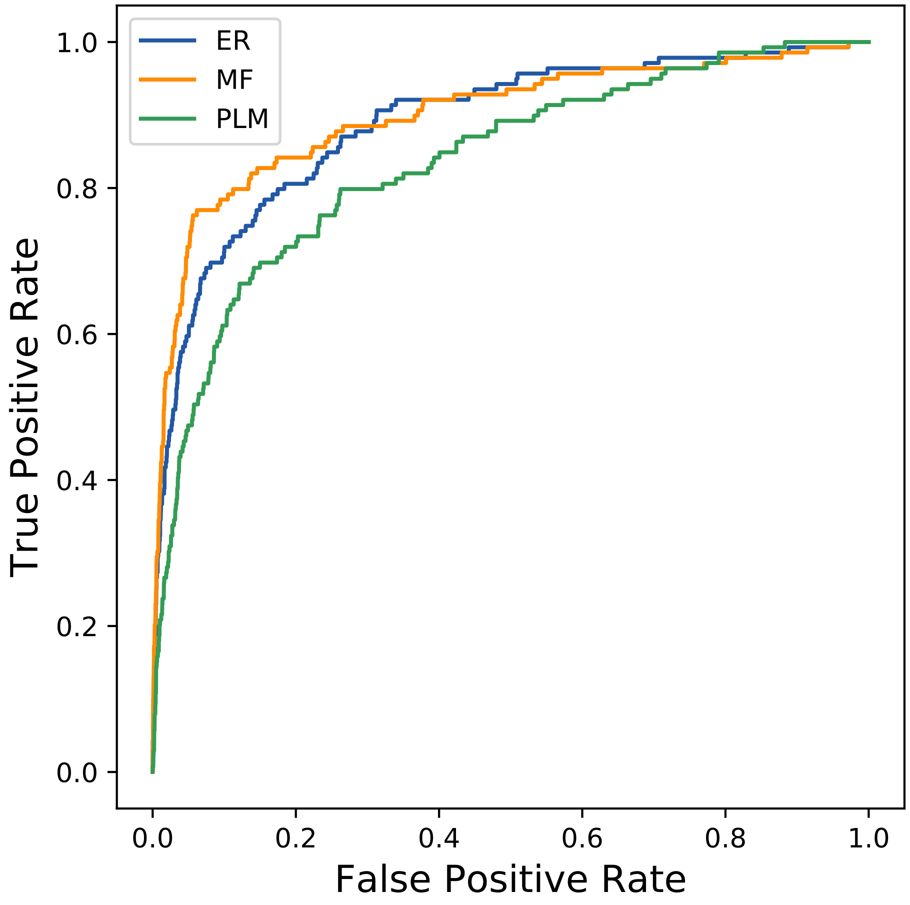
ER
MF
PLM
1.0
0.8
0.6
0.4
0.2
0.0
True Positive Rate
0.0
0.2
0.4
0.6
0.8
1.0
False Positive Rate
