## Supplemental Contact Maps for "Protein Structure Prediction with Expectation Reflection": 1484seq_79col_3sb9_PF00959_plots.pdf

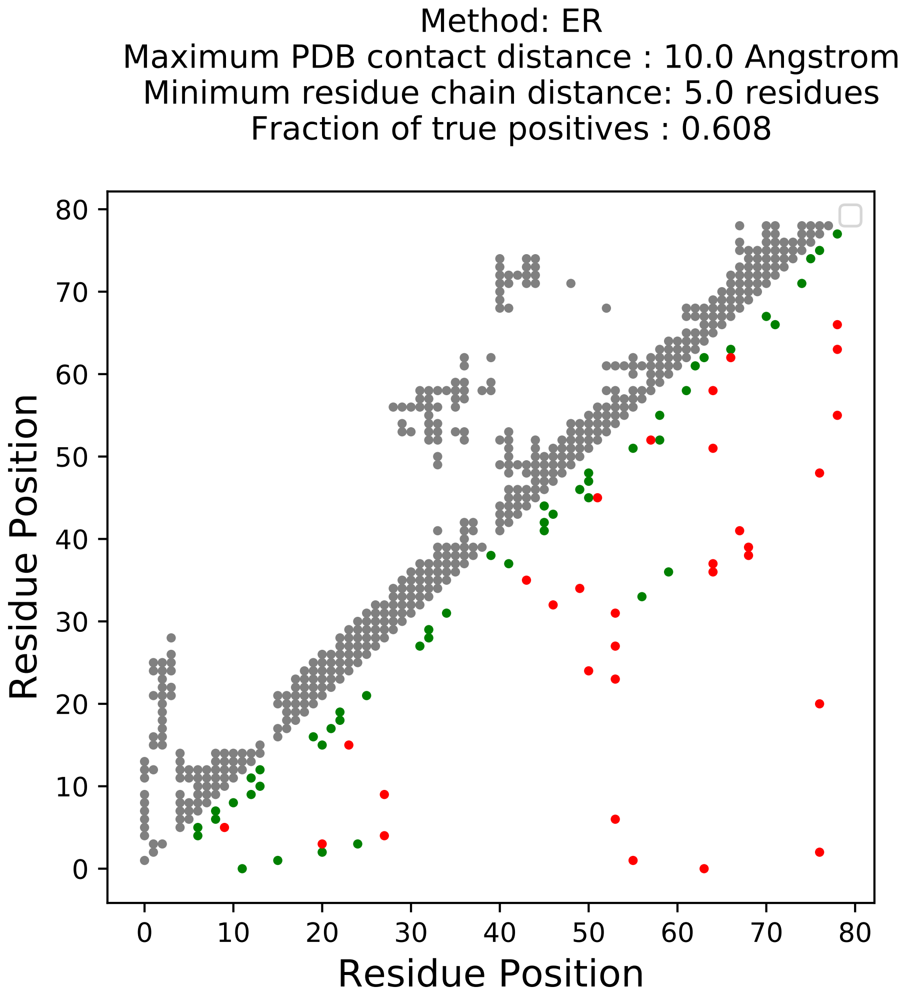
Method: ER
Maximum PDB contact distance : 10.0 Angstrom
Minimum residue chain distance: 5.0 residues
Fraction of true positives : 0.608
Residue Position
Residue Position
0
10
20
30
40
50
60
70
80

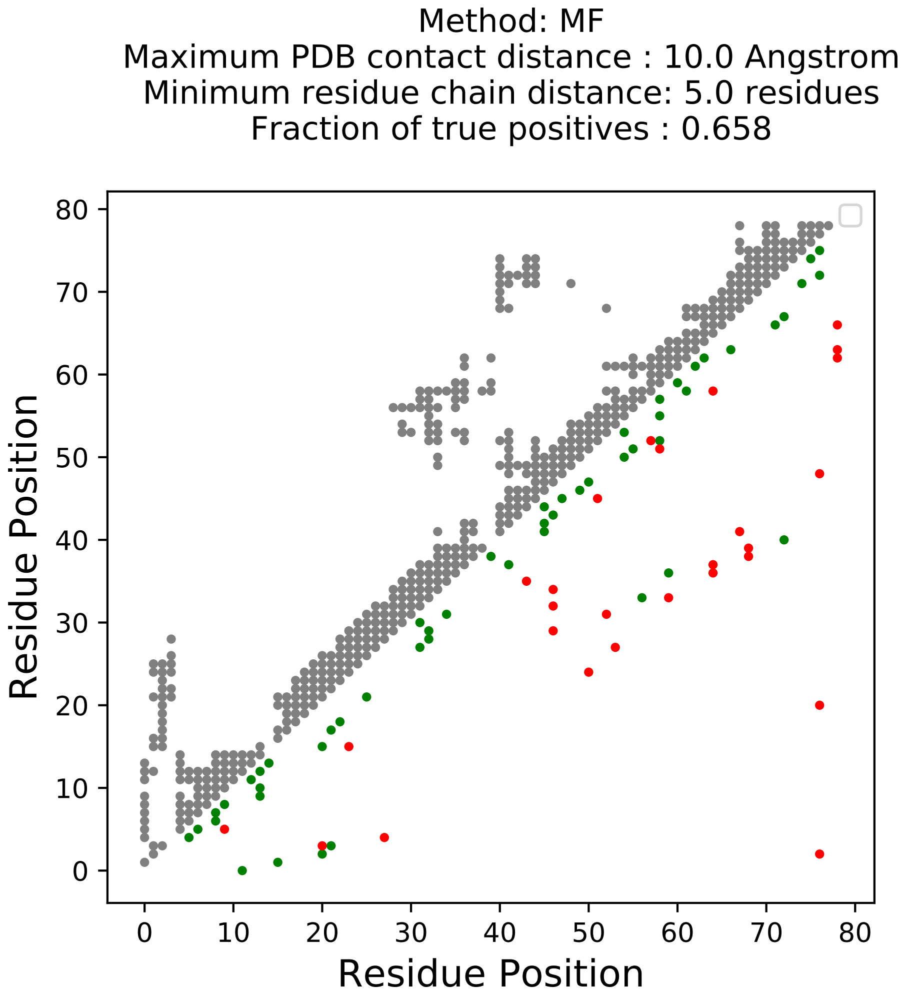
Method: MF
Maximum PDB contact distance : 10.0 Angstrom
Minimum residue chain distance: 5.0 residues
Fraction of true positives : 0.658
Residue Position
Residue Position
0
10
20
30
40
50
60
70
80

Method: PLM
Maximum PDB contact distance : 10.0 Angstrom
Minimum residue chain distance: 5.0 residues
Fraction of true positives : 0.532

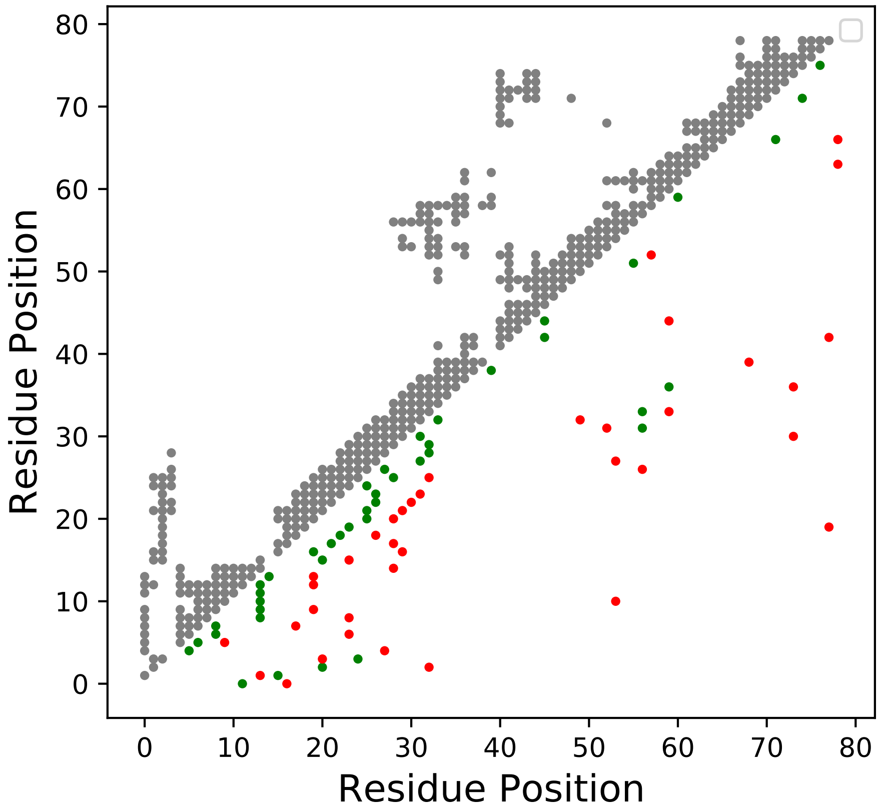

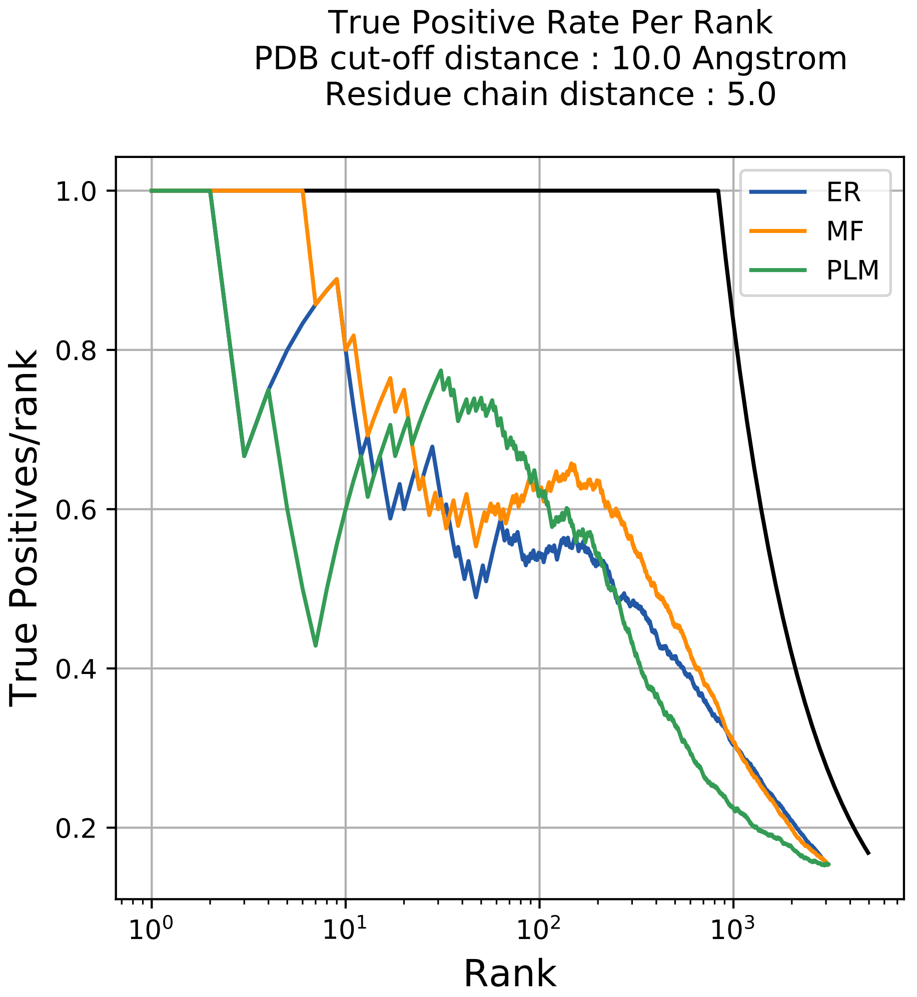
True Positive Rate Per Rank
PDB cut-off distance : 10.0 Angstrom
Residue chain distance : 5.0
True Positives/rank
Rank
1.0
0.8
0.6
0.4
0.2
$10^0$
$10^1$
$10^2$
$10^3$
ER
MF
PLM

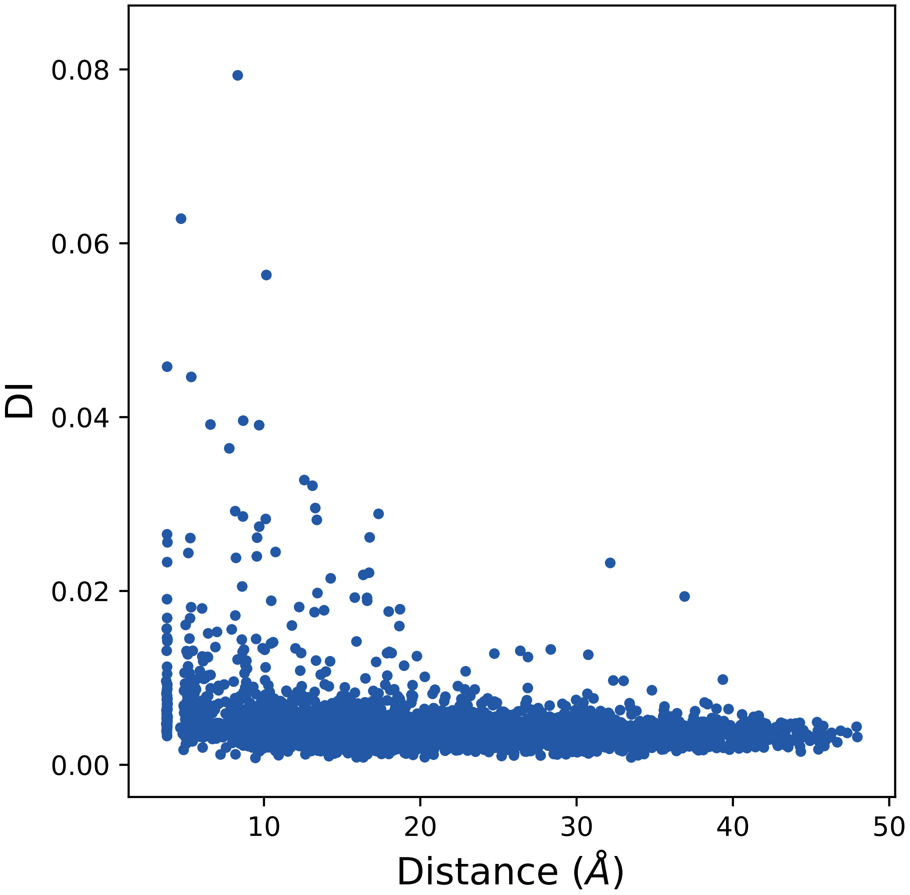
0.08
0.06
0.04
0.02
0.00
DI
10
20
30
40
50
Distance (Å)

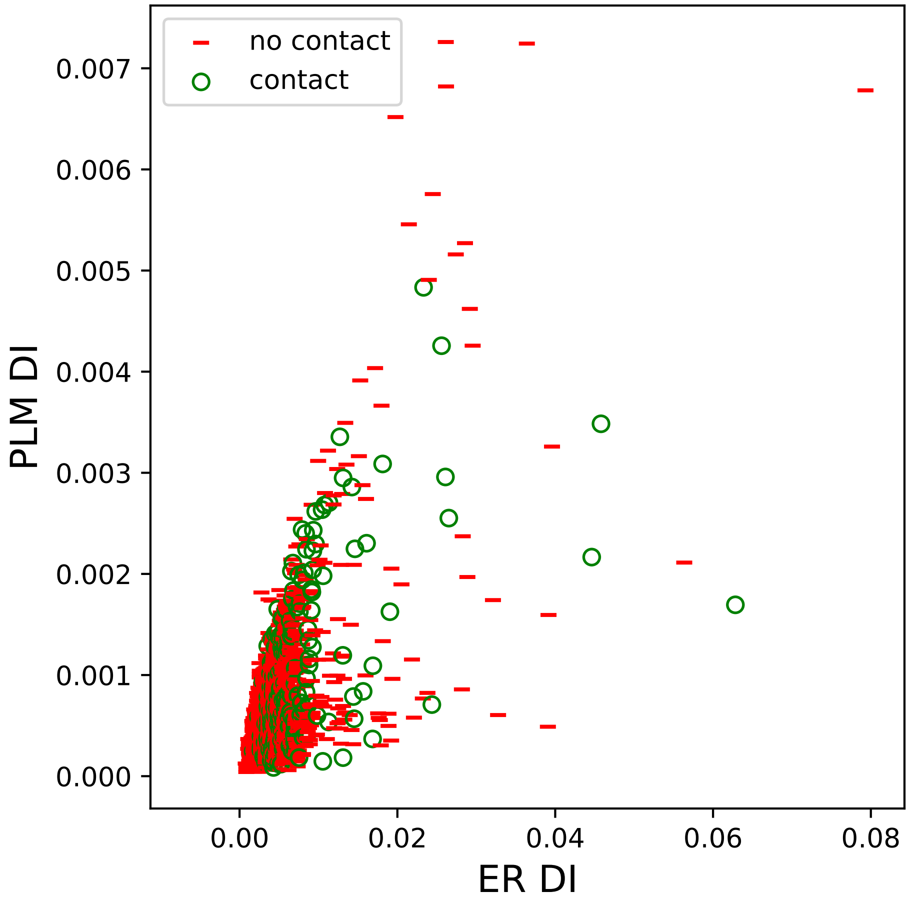
no contact
contact
PLM DI
ER DI
0.007
0.006
0.005
0.004
0.003
0.002
0.001
0.000
0.00
0.02
0.04
0.06
0.08

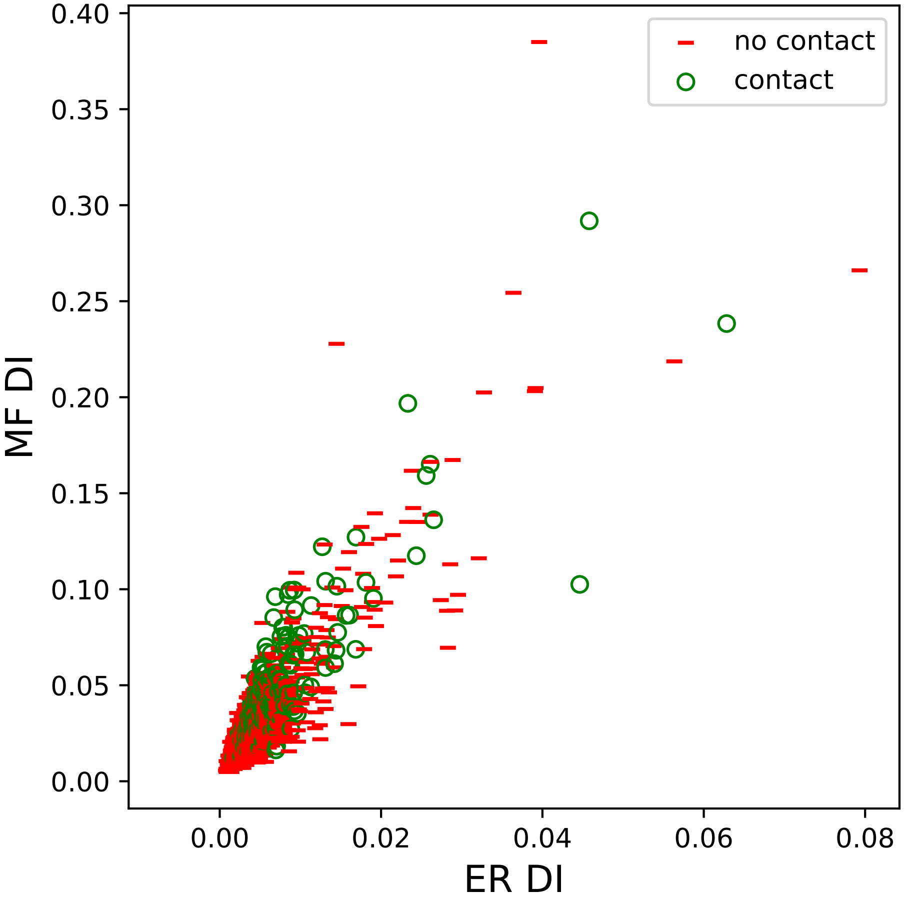

0.40
0.35
0.30
0.25
0.20
0.15
0.10
0.05
0.00
MF DI
0.00
0.02
0.04
0.06
0.08
ER DI
no contact
contact

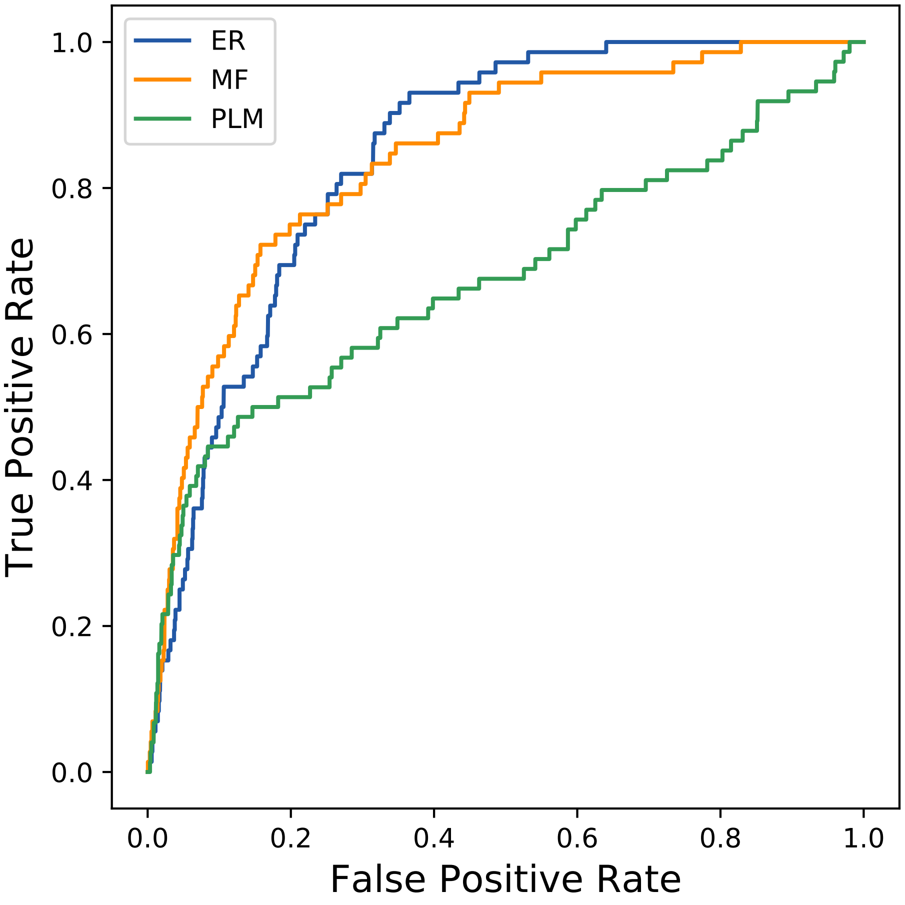
ER
MF
PLM
1.0
0.8
0.6
0.4
0.2
0.0
True Positive Rate
0.0
0.2
0.4
0.6
0.8
1.0
False Positive Rate
