## Supplemental Contact Maps for "Protein Structure Prediction with Expectation Reflection": 1658seq_95col_7rsp_PF00613_plots.pdf

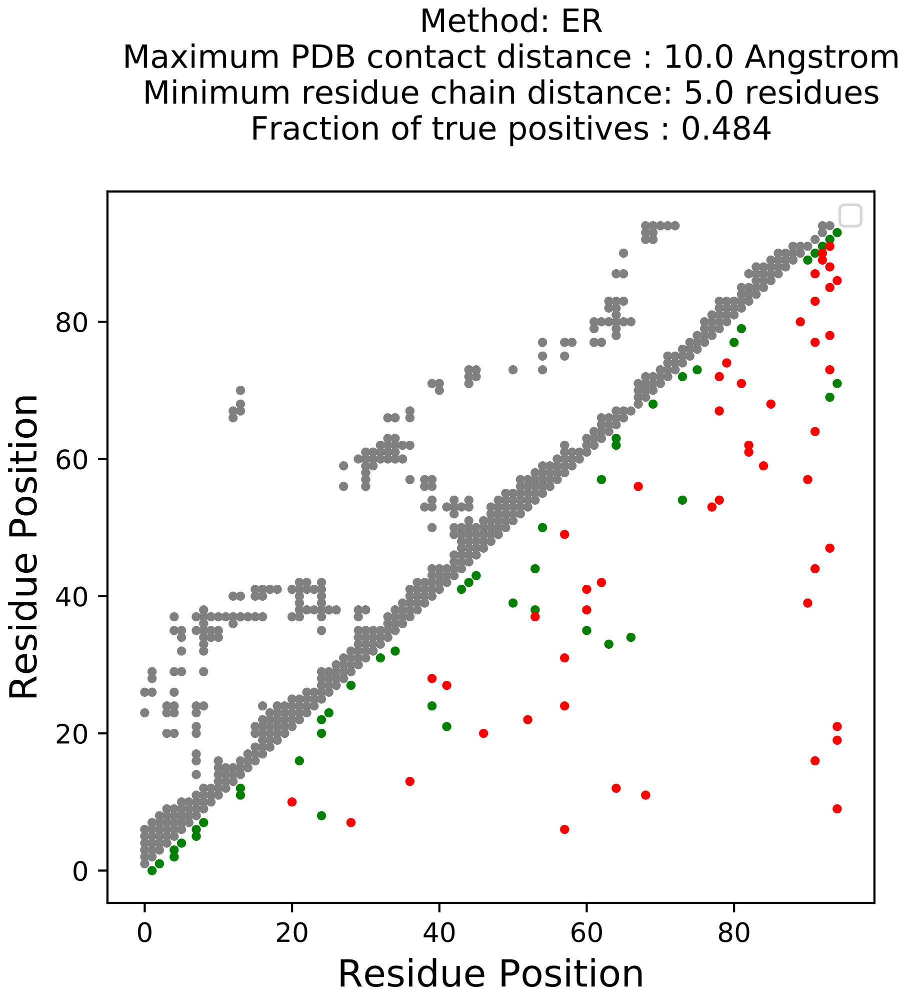

Method: ER
Maximum PDB contact distance : 10.0 Angstrom
Minimum residue chain distance: 5.0 residues
Fraction of true positives : 0.484
Residue Position
Residue Position
0
20
40
60
80

Method: MF
Maximum PDB contact distance : 10.0 Angstrom
Minimum residue chain distance: 5.0 residues
Fraction of true positives : 0.495

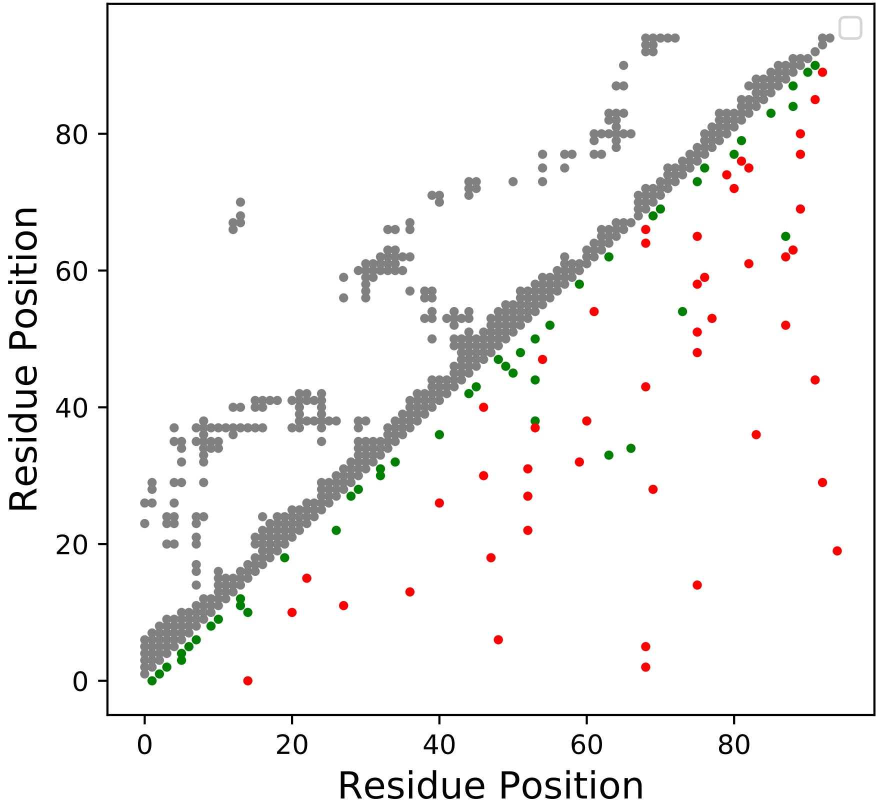

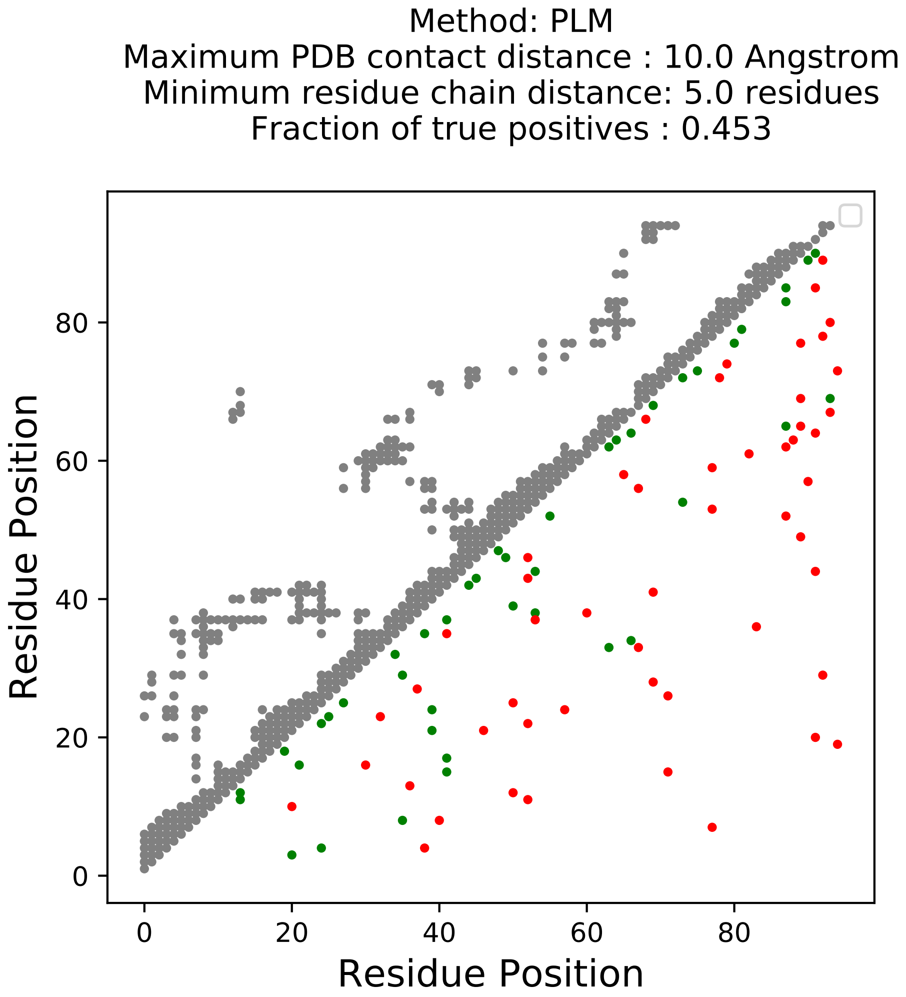
Method: PLM
Maximum PDB contact distance : 10.0 Angstrom
Minimum residue chain distance: 5.0 residues
Fraction of true positives : 0.453
Residue Position
Residue Position
0
20
40
60
80

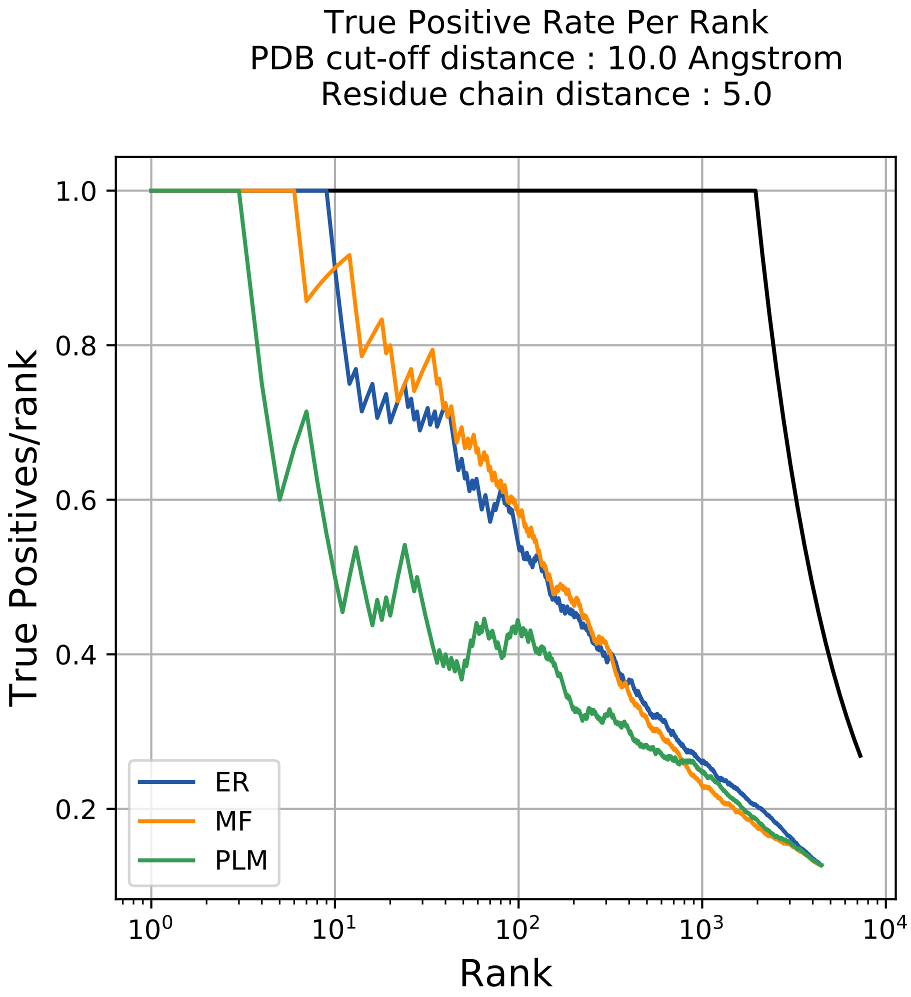

True Positive Rate Per Rank
PDB cut-off distance : 10.0 Angstrom
Residue chain distance : 5.0
True Positives/rank
1.0
0.8
0.6
0.4
0.2
$10^0$
$10^1$
$10^2$
$10^3$
$10^4$
Rank
ER
MF
PLM

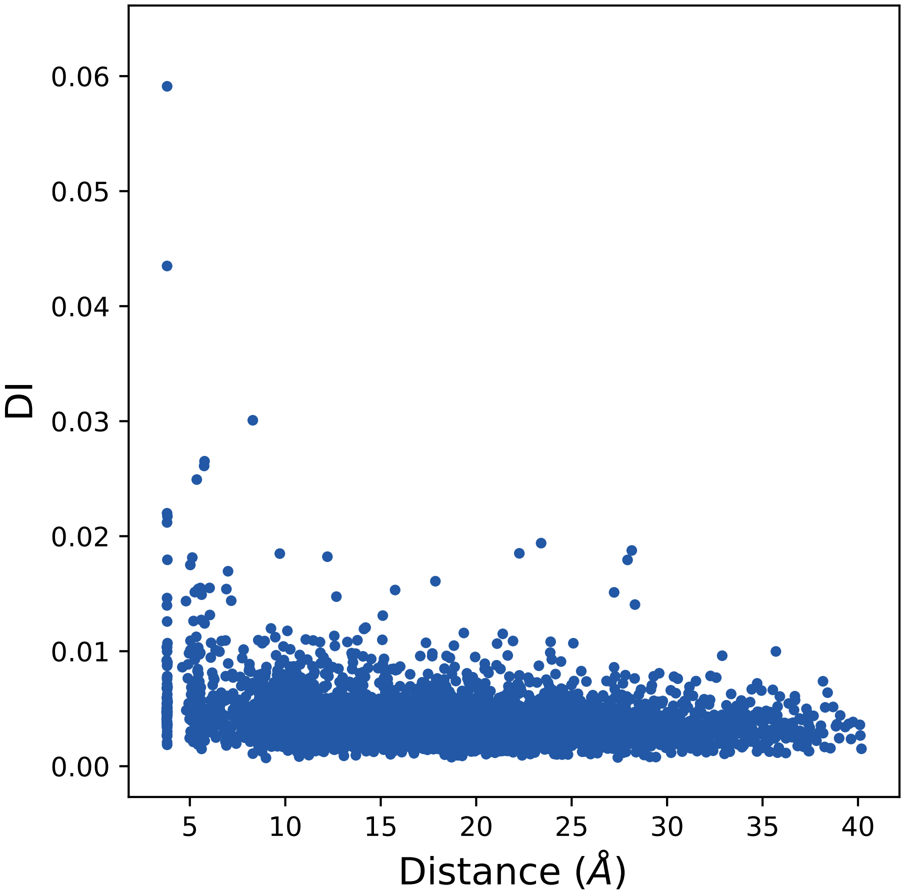
0.06
0.05
0.04
0.03
0.02
0.01
0.00
DI
5
10
15
20
25
30
35
40
Distance (Å)

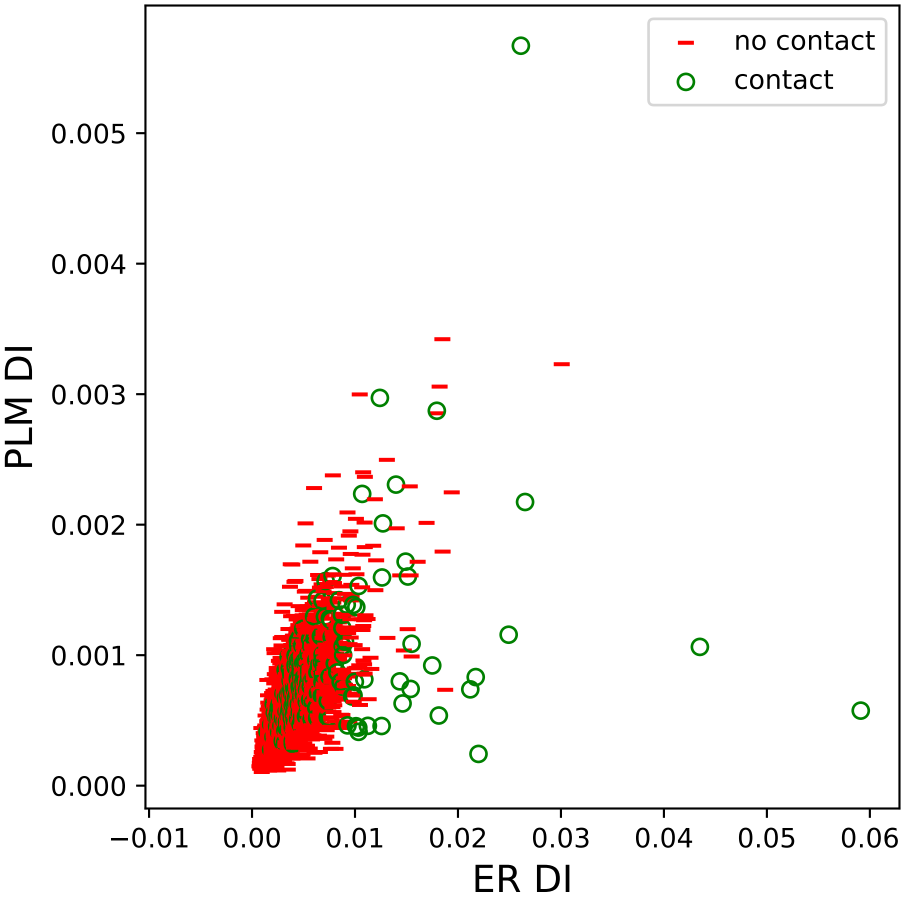

no contact
contact
0.005
0.004
0.003
0.002
0.001
0.000
PLM DI
−0.01
0.00
0.01
0.02
0.03
0.04
0.05
0.06
ER DI

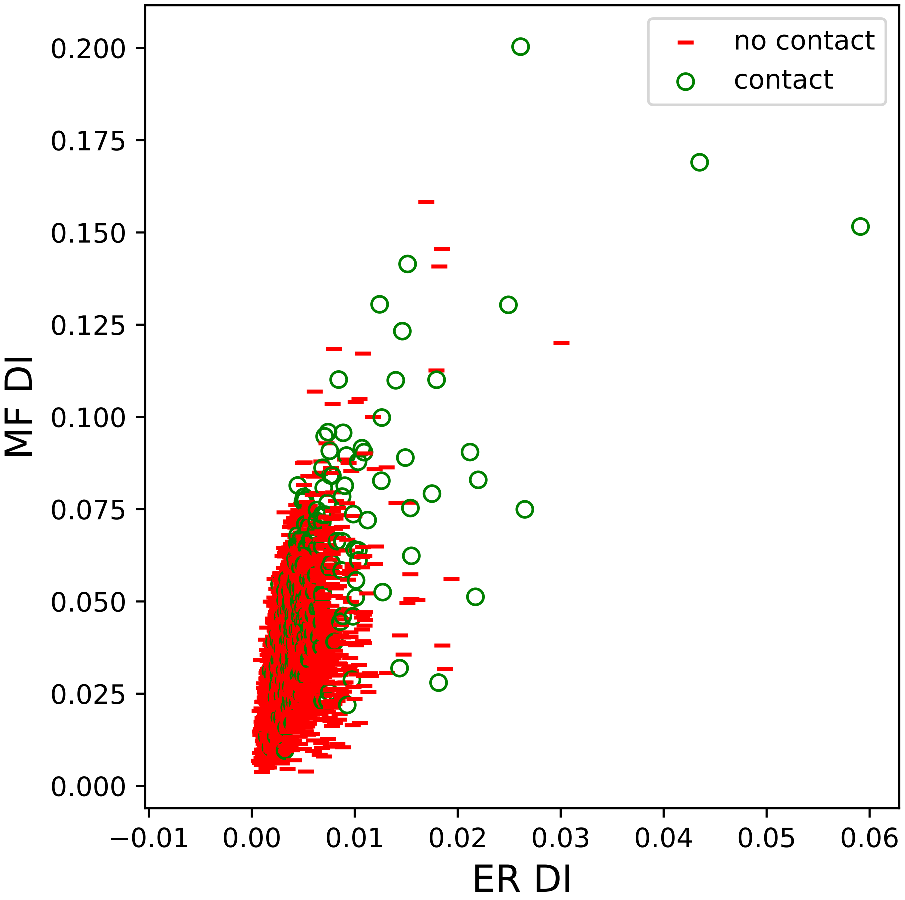

no contact
contact
MF DI
ER DI
0.200
0.175
0.150
0.125
0.100
0.075
0.050
0.025
0.000
−0.01
0.00
0.01
0.02
0.03
0.04
0.05
0.06

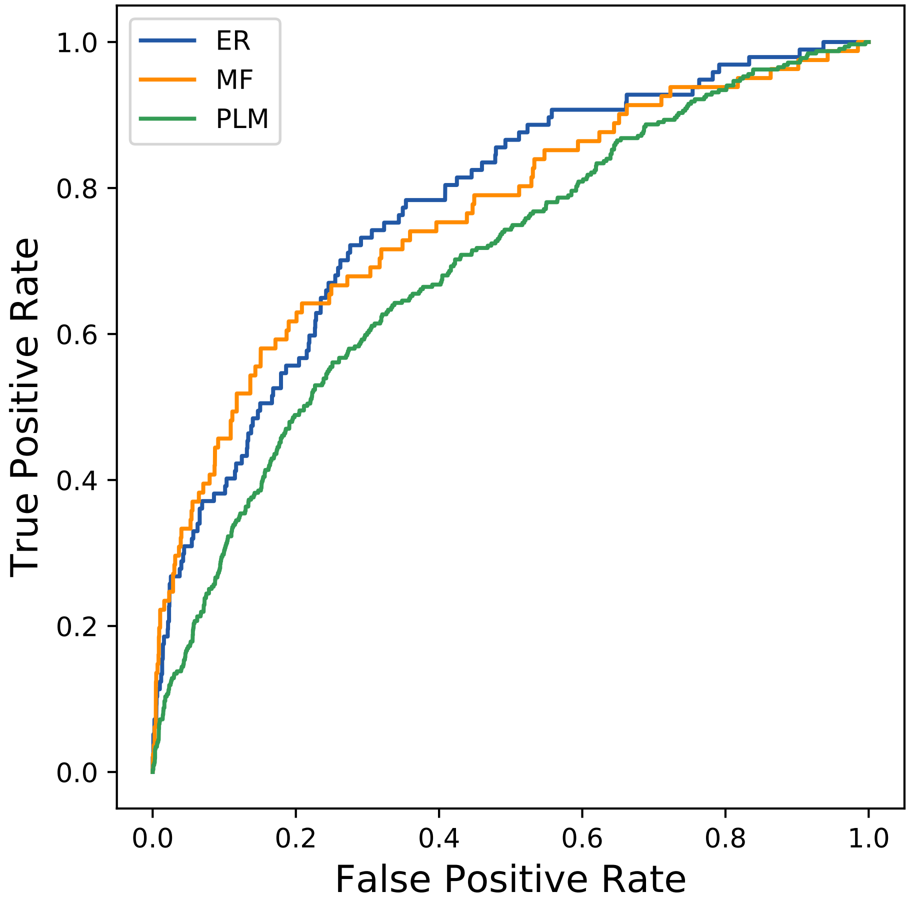
ER
MF
PLM
1.0
0.8
0.6
0.4
0.2
0.0
True Positive Rate
0.0
0.2
0.4
0.6
0.8
1.0
False Positive Rate
