## Supplemental Contact Maps for "Protein Structure Prediction with Expectation Reflection": 2182seq_90col_4qde_PF11969_plots.pdf

Method: ER
Maximum PDB contact distance : 10.0 Angstrom
Minimum residue chain distance: 5.0 residues
Fraction of true positives : 0.667

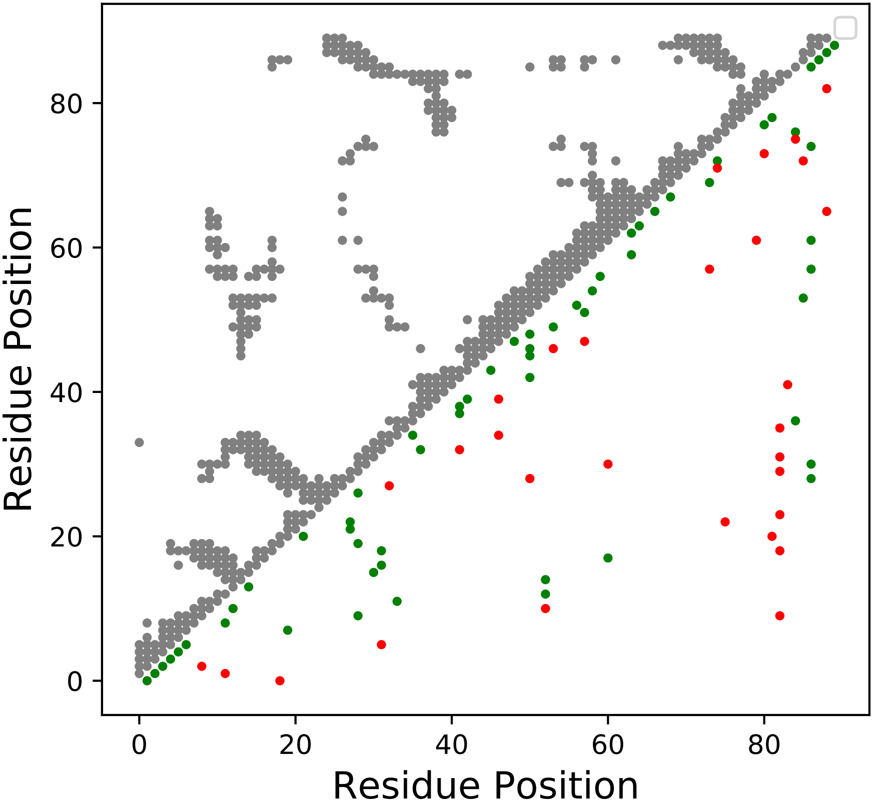

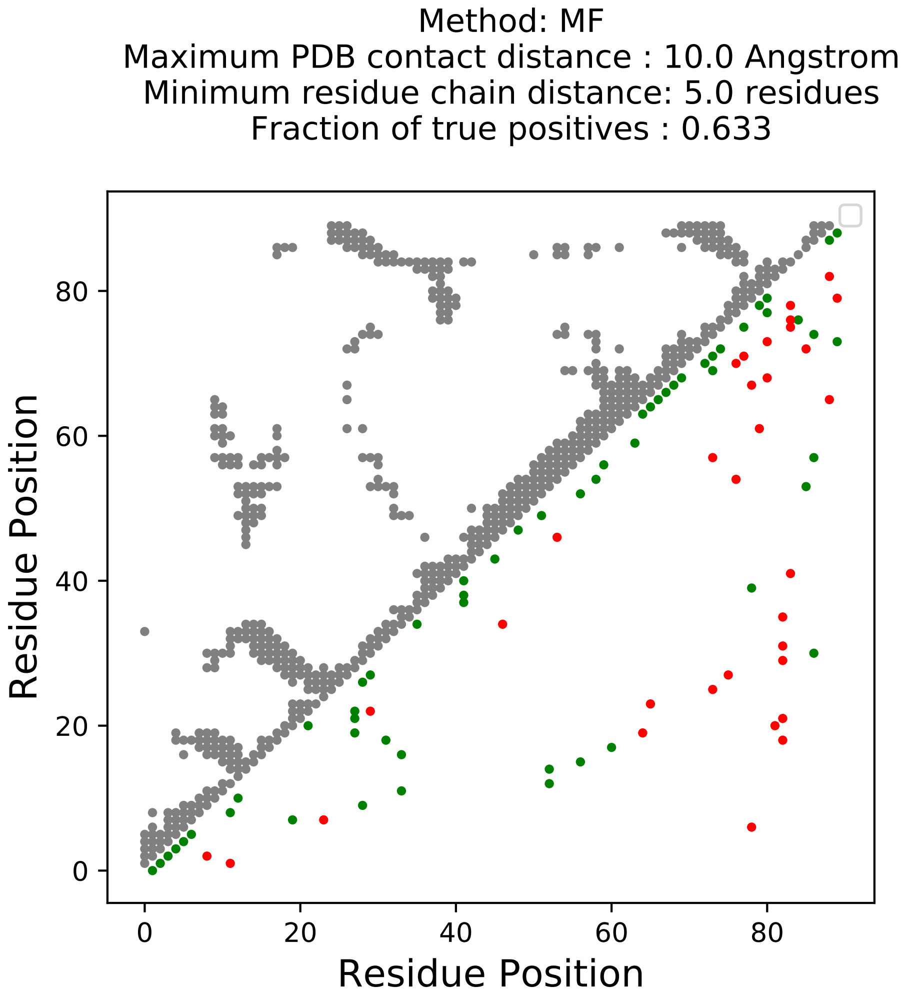

Method: MF
Maximum PDB contact distance : 10.0 Angstrom
Minimum residue chain distance: 5.0 residues
Fraction of true positives : 0.633
Residue Position
Residue Position
0
20
40
60
80

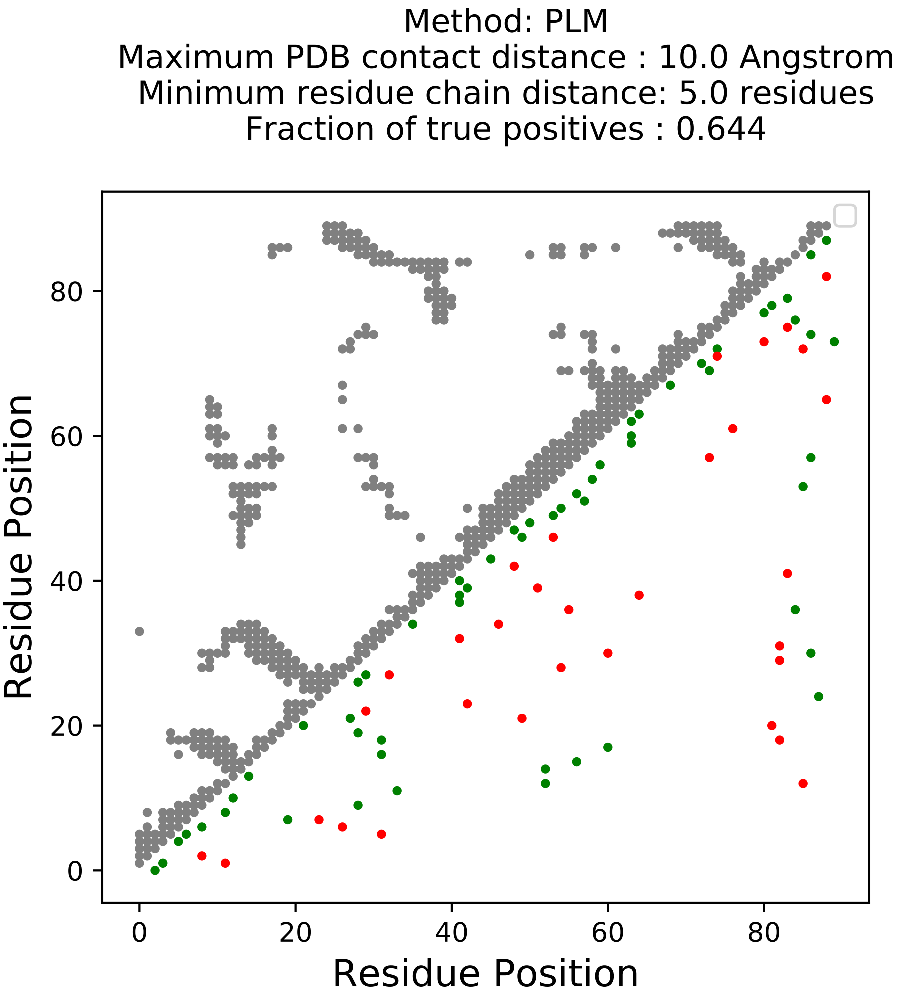

Method: PLM
Maximum PDB contact distance : 10.0 Angstrom
Minimum residue chain distance: 5.0 residues
Fraction of true positives : 0.644
Residue Position
Residue Position
0
20
40
60
80

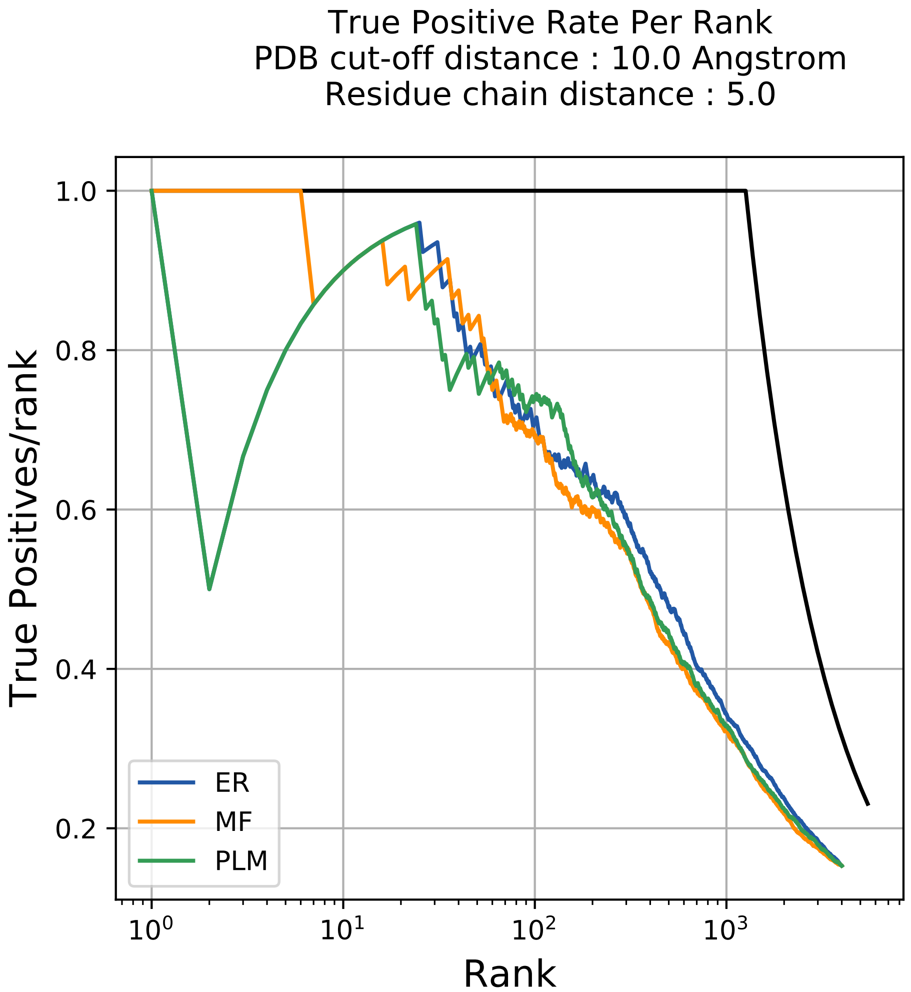

True Positive Rate Per Rank
PDB cut-off distance : 10.0 Angstrom
Residue chain distance : 5.0
True Positives/rank
1.0
0.8
0.6
0.4
0.2
$10^0$
$10^1$
$10^2$
$10^3$
Rank
ER
MF
PLM

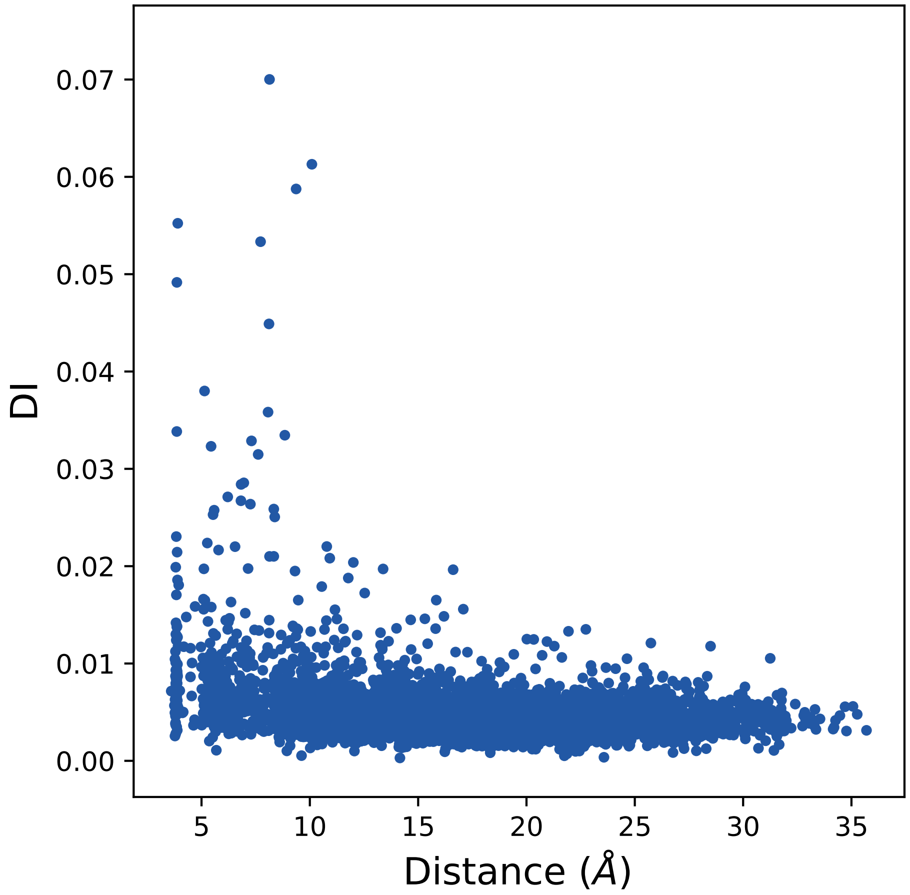
0.07
0.06
0.05
0.04
0.03
0.02
0.01
0.00
DI
5
10
15
20
25
30
35
Distance (Å)

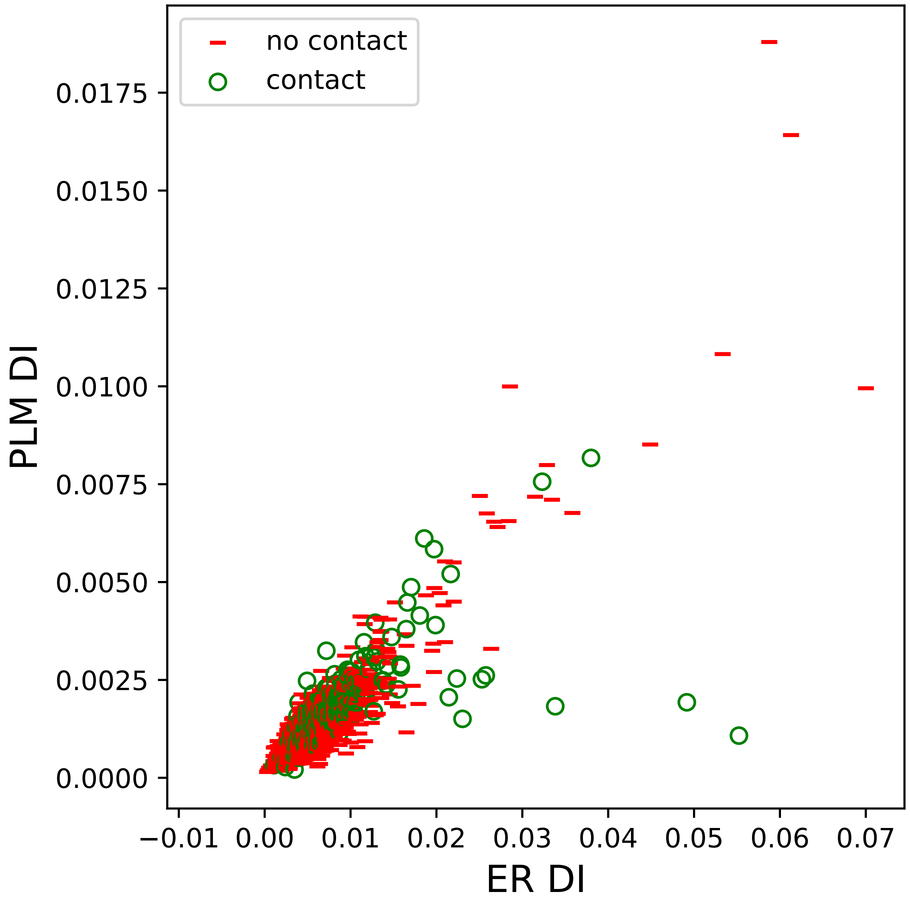

no contact
contact
0.0175
0.0150
0.0125
0.0100
0.0075
0.0050
0.0025
0.0000
PLM DI
−0.01
0.00
0.01
0.02
0.03
0.04
0.05
0.06
0.07
ER DI

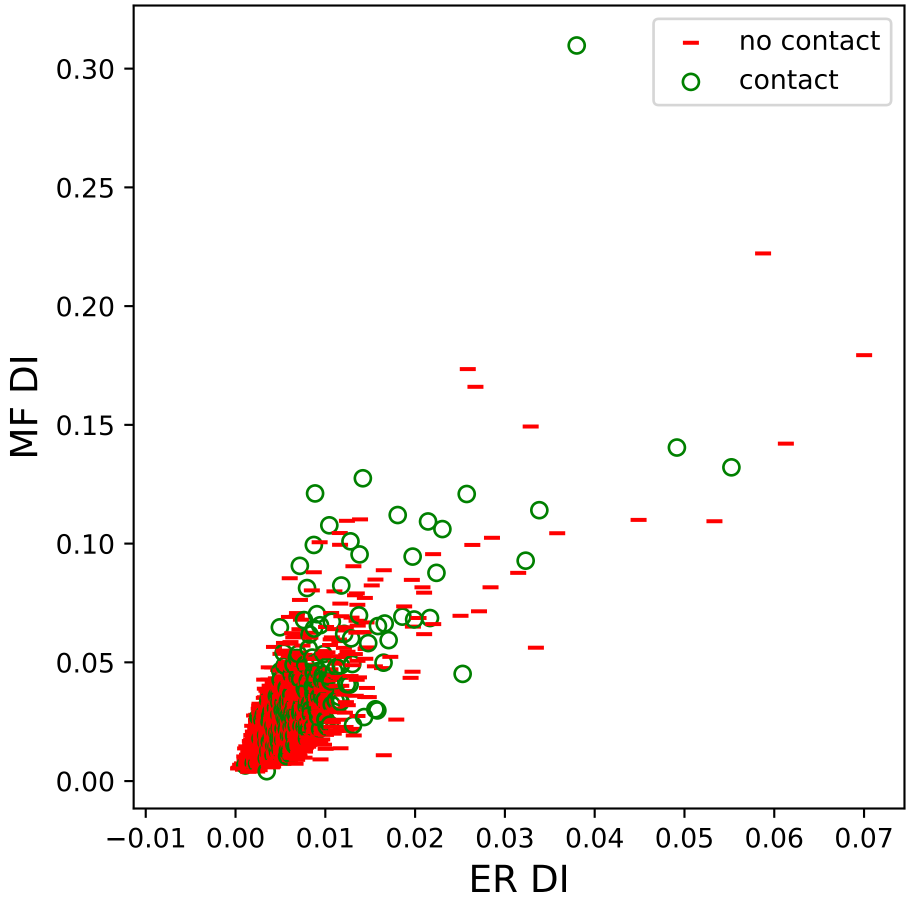

no contact
contact
MF DI
ER DI
0.30
0.25
0.20
0.15
0.10
0.05
0.00
−0.01
0.00
0.01
0.02
0.03
0.04
0.05
0.06
0.07

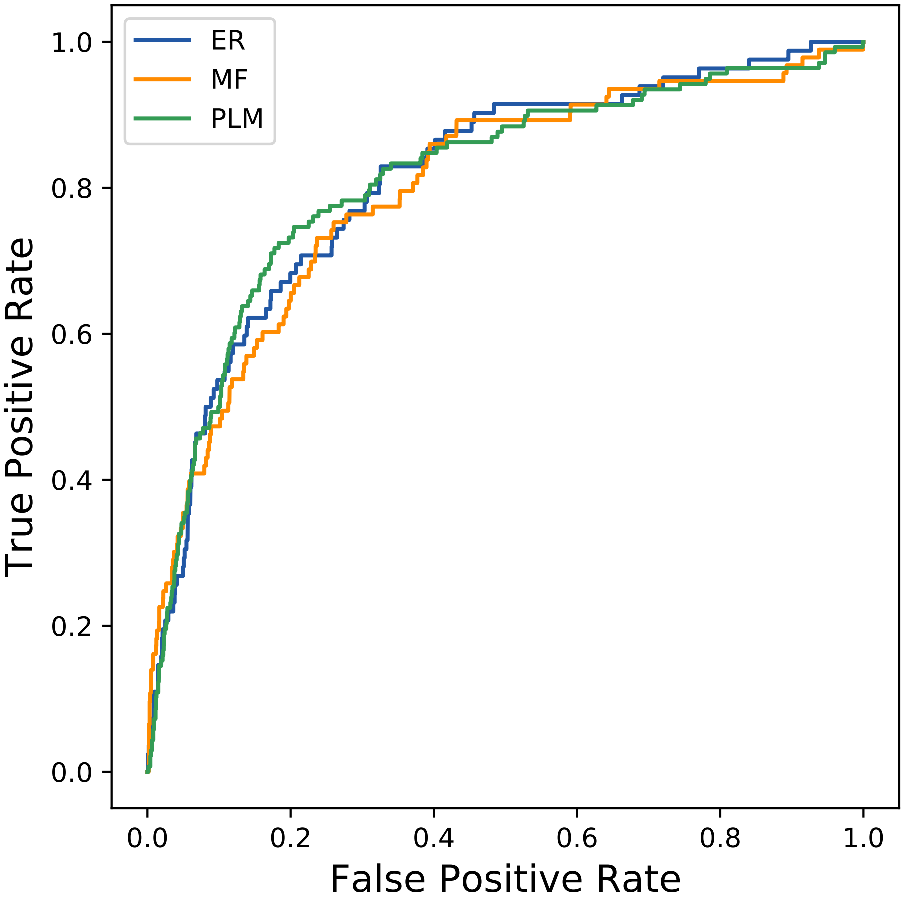
ER
MF
PLM
True Positive Rate
False Positive Rate
0.0
0.2
0.4
0.6
0.8
1.0
