## Supplemental Contact Maps for "Protein Structure Prediction with Expectation Reflection": 2594seq_277col_6q0w_PF10433_plots.pdf

Method: ER
Maximum PDB contact distance : 10.0 Angstrom
Minimum residue chain distance: 5.0 residues
Fraction of true positives : 0.596

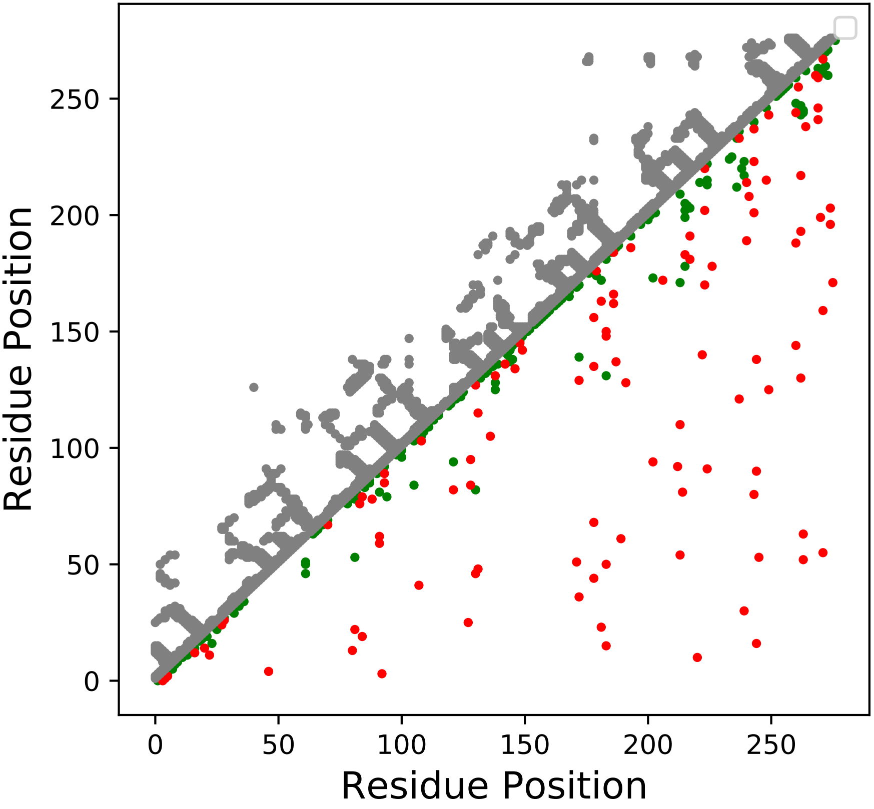

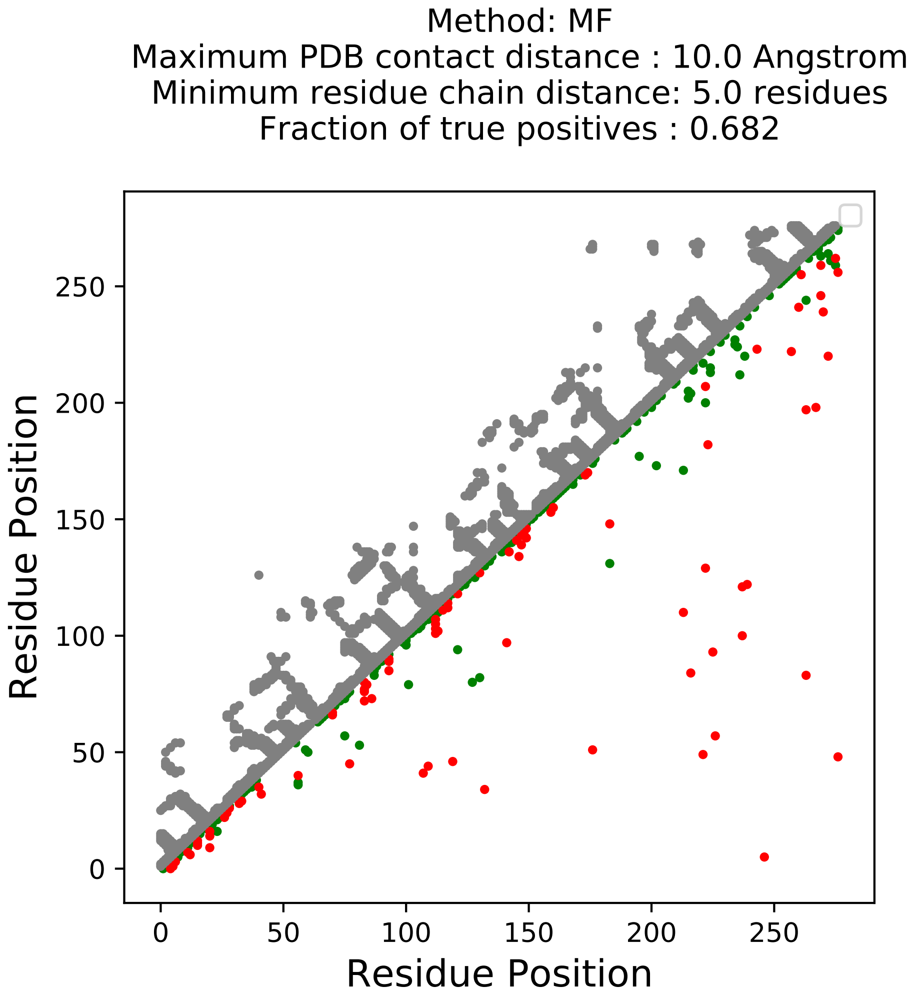
Method: MF
Maximum PDB contact distance : 10.0 Angstrom
Minimum residue chain distance: 5.0 residues
Fraction of true positives : 0.682
Residue Position
Residue Position
0
50
100
150
200
250
0
50
100
150
200
250

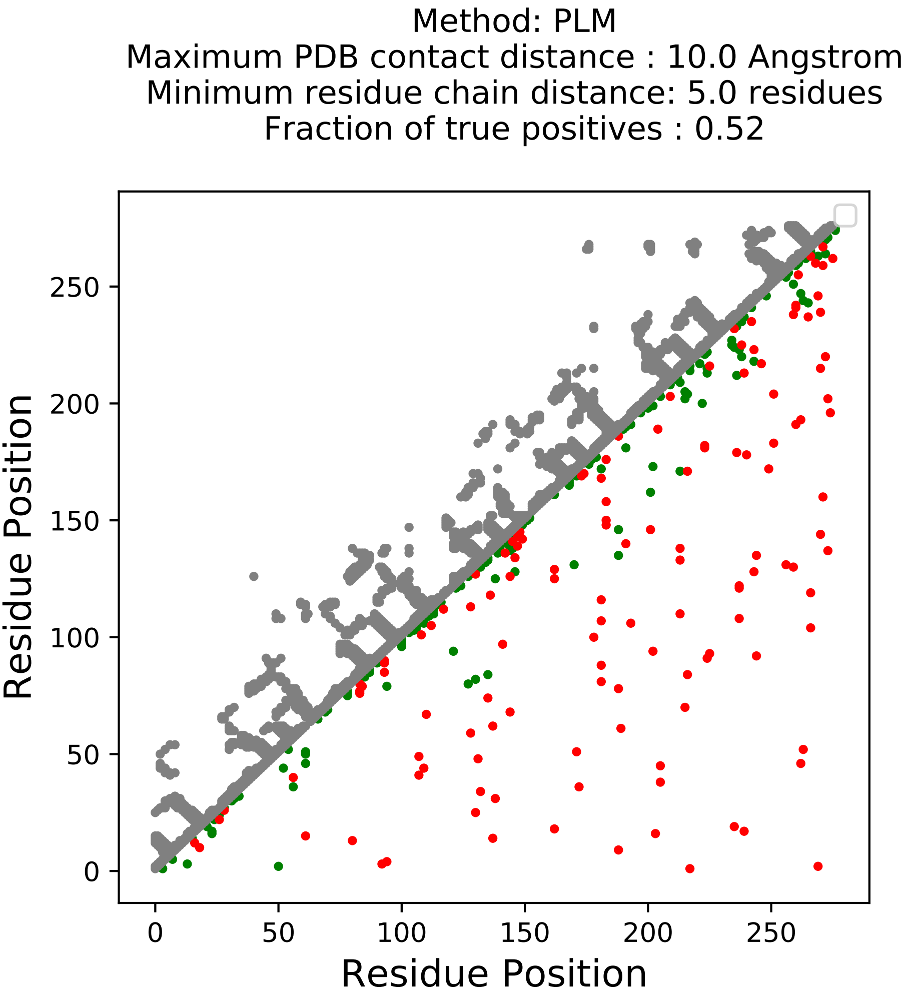
Method: PLM
Maximum PDB contact distance : 10.0 Angstrom
Minimum residue chain distance: 5.0 residues
Fraction of true positives : 0.52
Residue Position
Residue Position
0
50
100
150
200
250

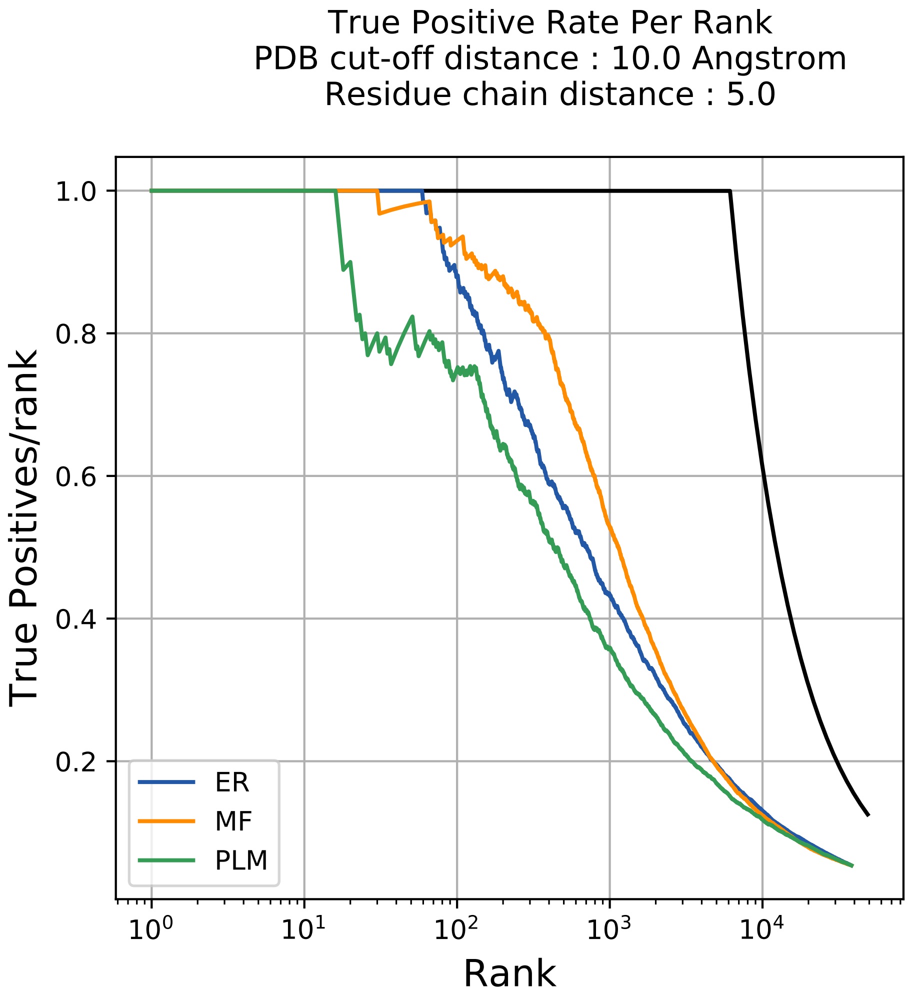
True Positive Rate Per Rank
PDB cut-off distance : 10.0 Angstrom
Residue chain distance : 5.0
1.0
0.8
0.6
0.4
0.2
True Positives/rank
$10^0$
$10^1$
$10^2$
$10^3$
$10^4$
Rank
ER
MF
PLM

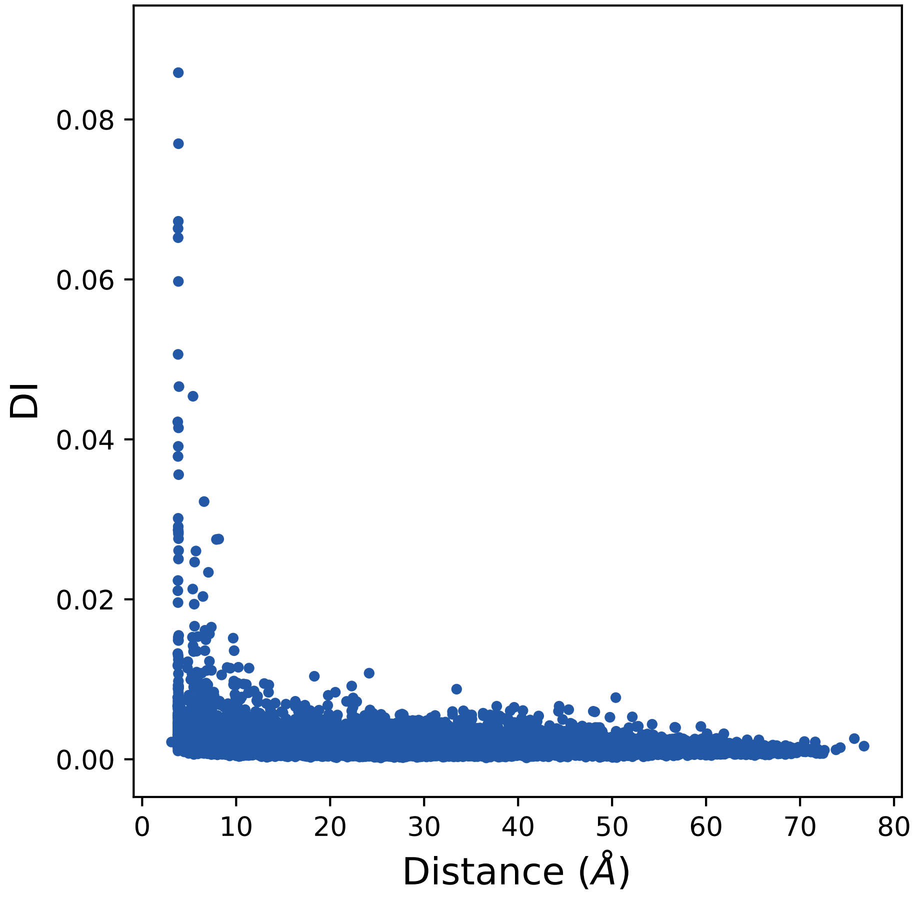

0.08
0.06
0.04
0.02
0.00
DI
0
10
20
30
40
50
60
70
80
Distance (Å)

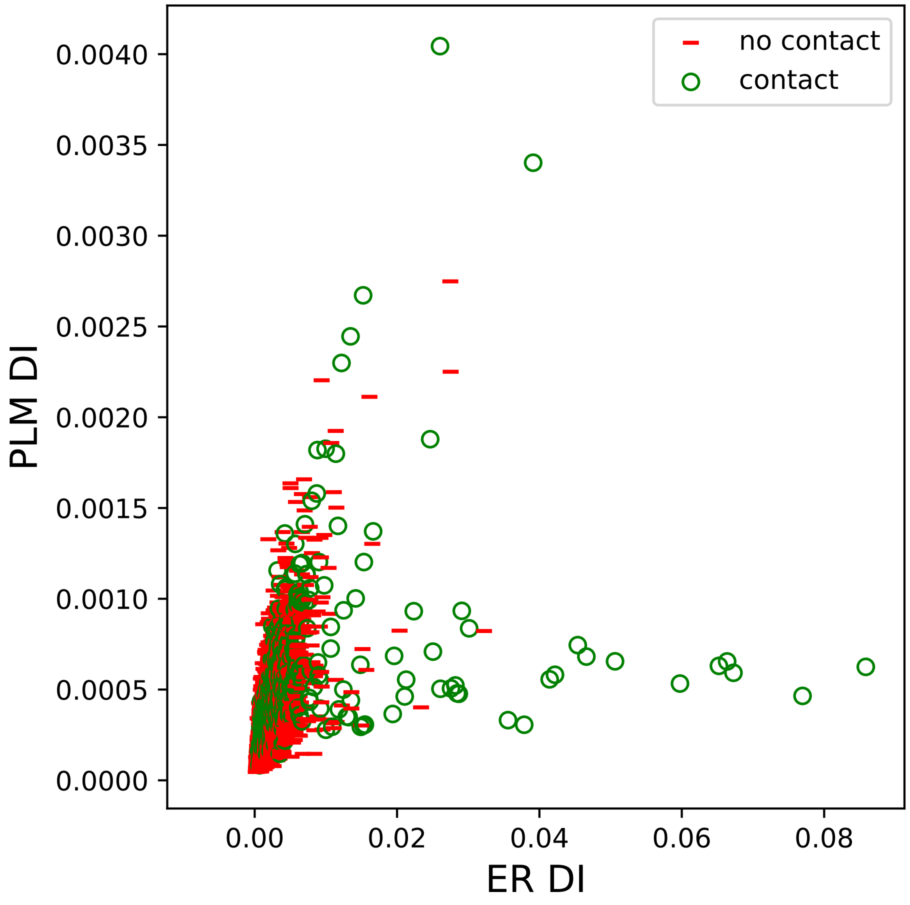

no contact
contact
0.0040
0.0035
0.0030
0.0025
0.0020
0.0015
0.0010
0.0005
0.0000
PLM DI
0.00
0.02
0.04
0.06
0.08
ER DI

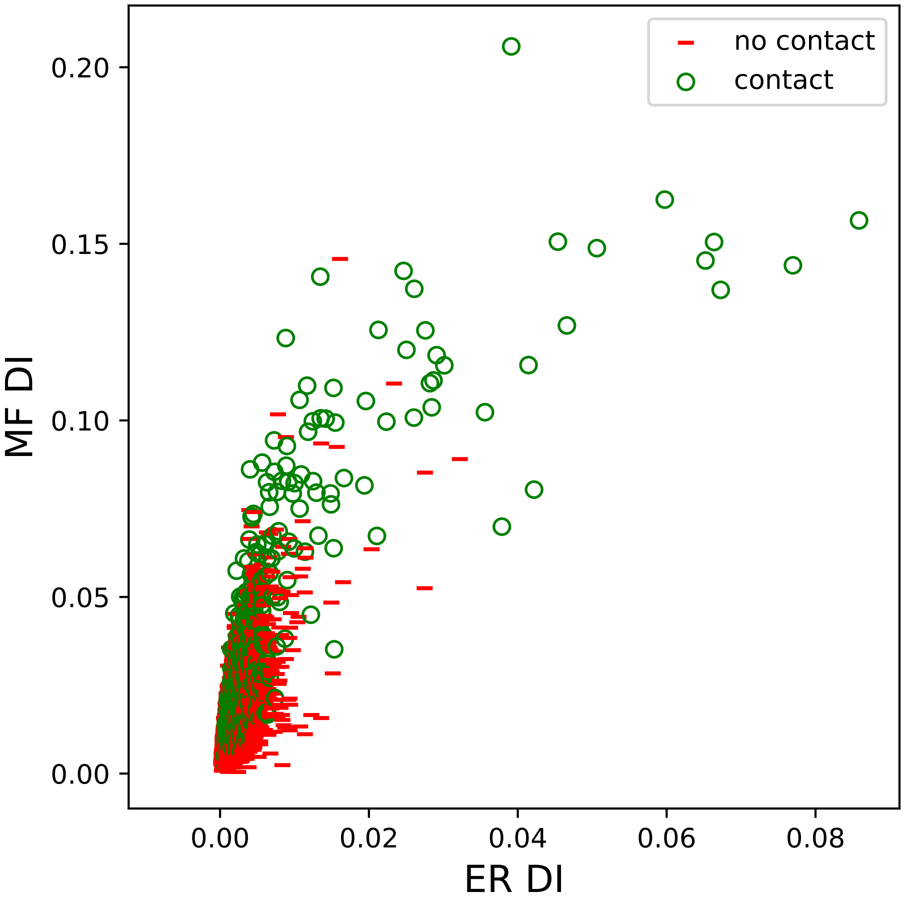

no contact
contact
0.20
0.15
0.10
0.05
0.00
MF DI
0.00
0.02
0.04
0.06
0.08
ER DI

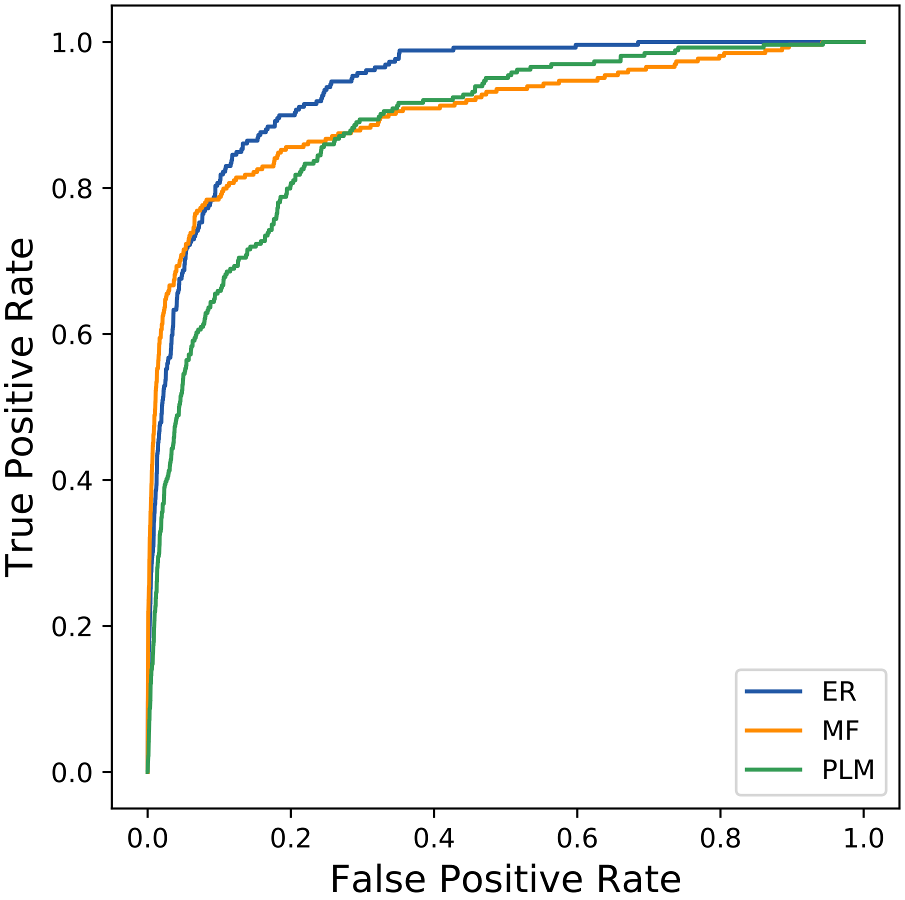

True Positive Rate
False Positive Rate
0.0
0.2
0.4
0.6
0.8
1.0
ER
MF
PLM
