## Supplemental Contact Maps for "Protein Structure Prediction with Expectation Reflection": 2790seq_81col_1ig3_PF04263_plots.pdf

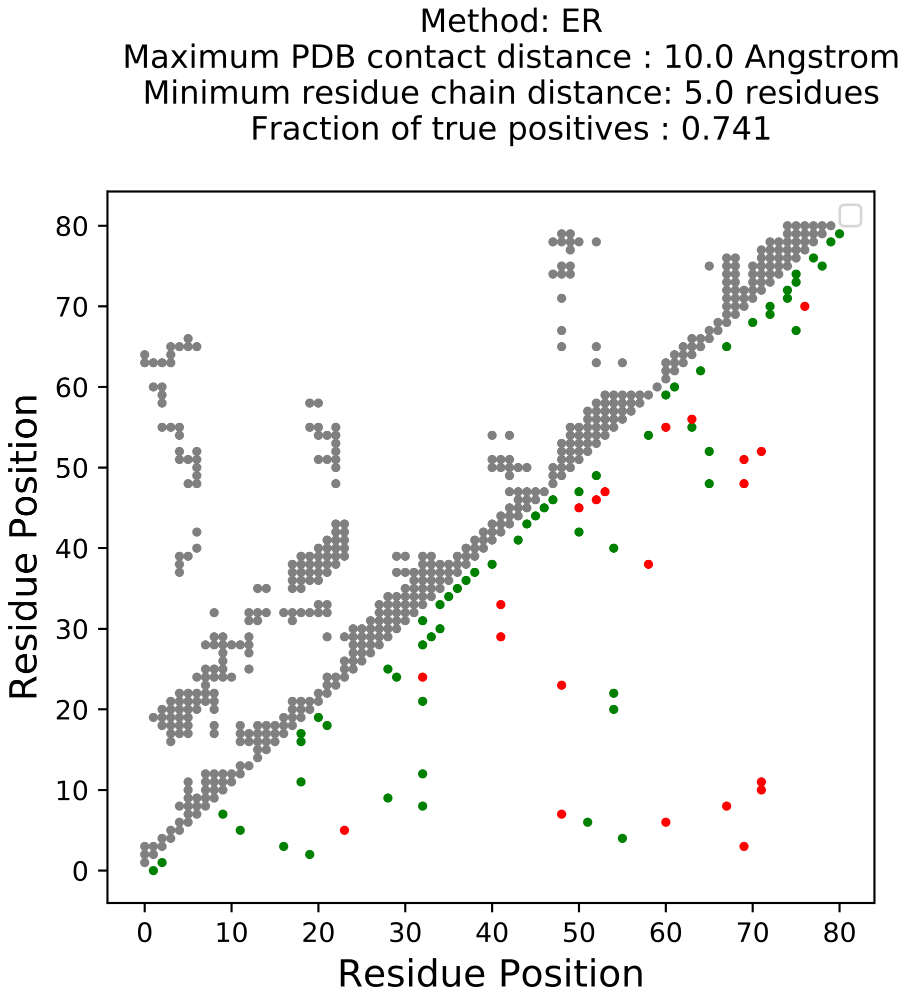

Method: ER
Maximum PDB contact distance : 10.0 Angstrom
Minimum residue chain distance: 5.0 residues
Fraction of true positives : 0.741
Residue Position
Residue Position
0
10
20
30
40
50
60
70
80

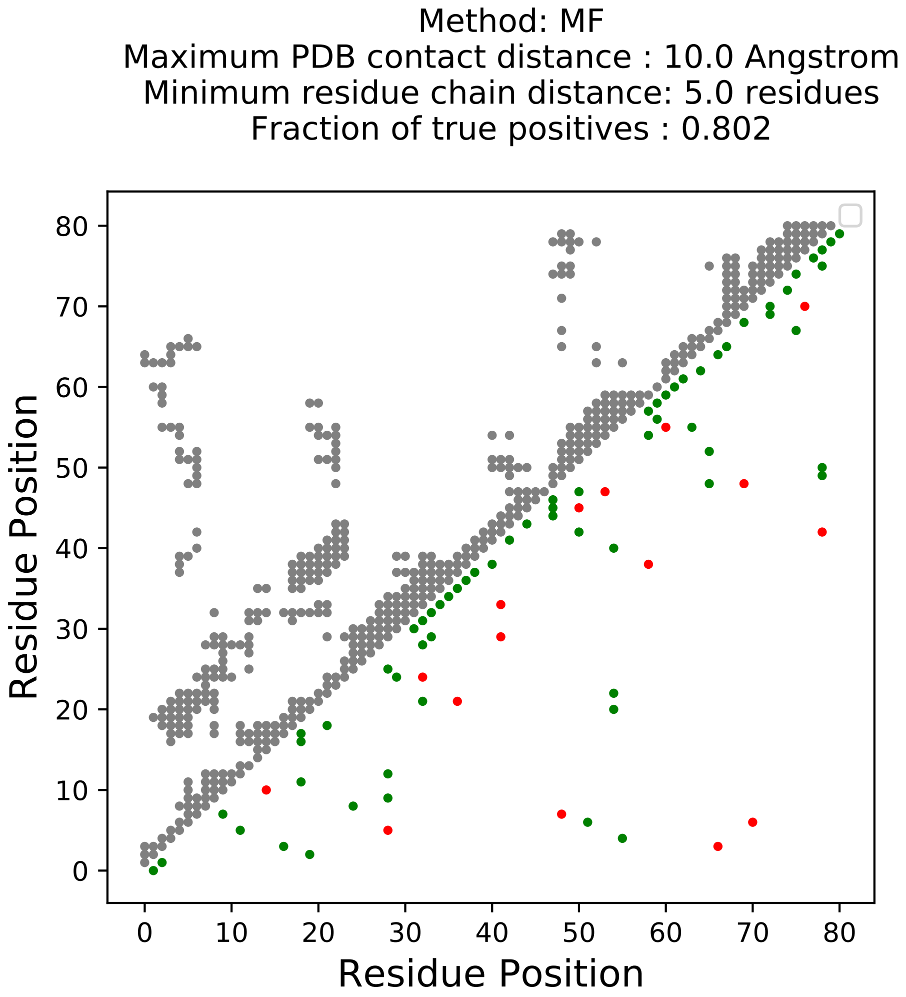

Method: MF
Maximum PDB contact distance : 10.0 Angstrom
Minimum residue chain distance: 5.0 residues
Fraction of true positives : 0.802
Residue Position
Residue Position
0
10
20
30
40
50
60
70
80

Method: PLM
Maximum PDB contact distance : 10.0 Angstrom
Minimum residue chain distance: 5.0 residues
Fraction of true positives : 0.728

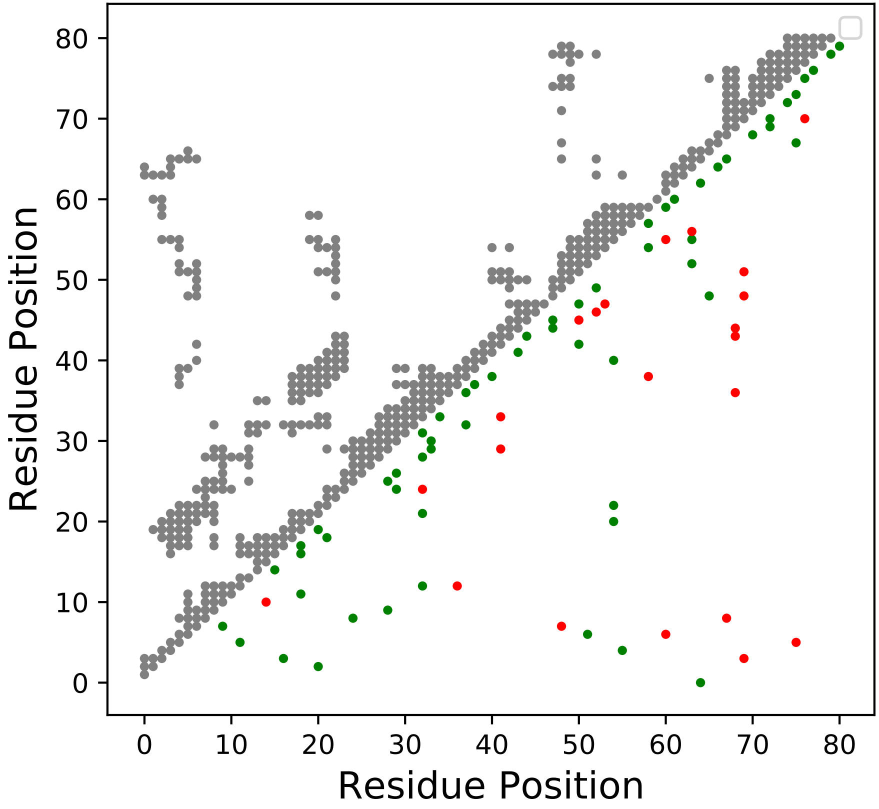

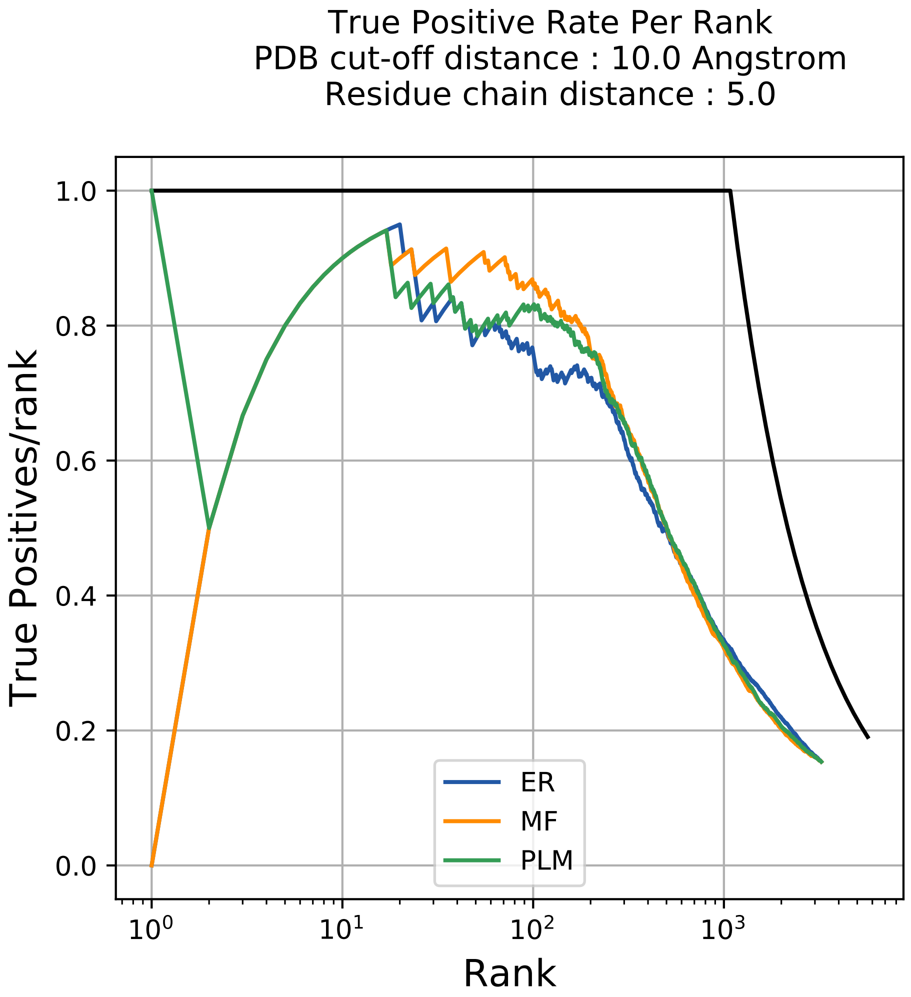
True Positive Rate Per Rank
PDB cut-off distance : 10.0 Angstrom
Residue chain distance : 5.0
True Positives/rank
Rank
1.0
0.8
0.6
0.4
0.2
0.0
$10^0$
$10^1$
$10^2$
$10^3$
ER
MF
PLM

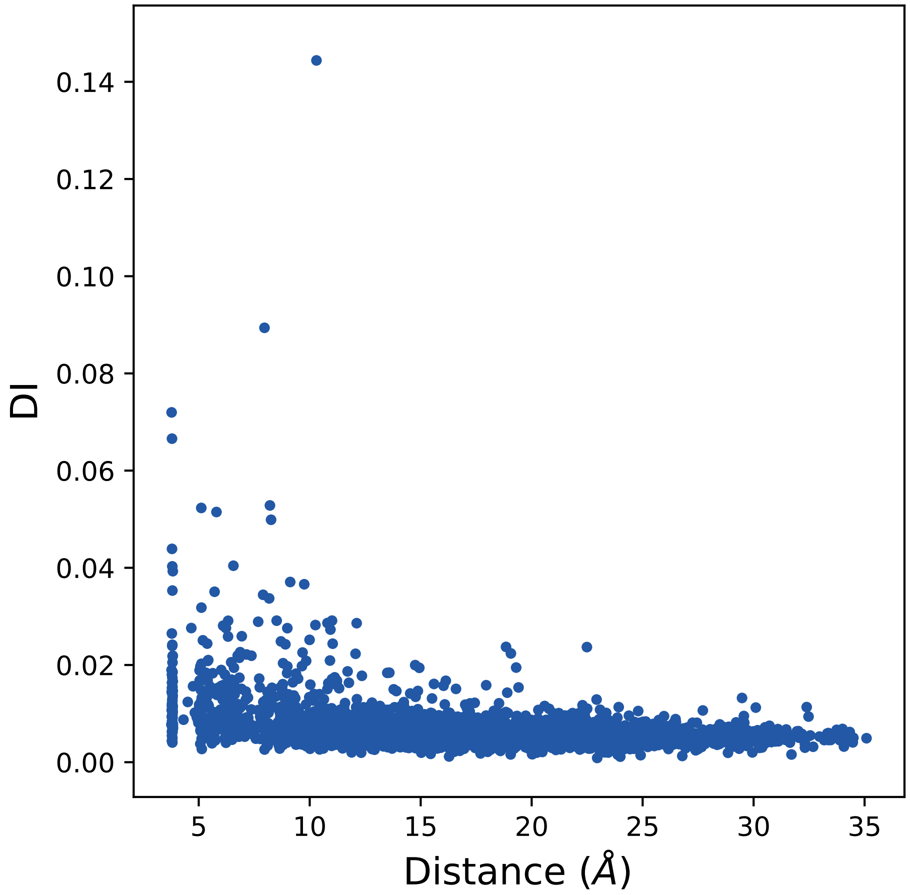

0.14
0.12
0.10
0.08
0.06
0.04
0.02
0.00
DI
5
10
15
20
25
30
35
Distance (Å)

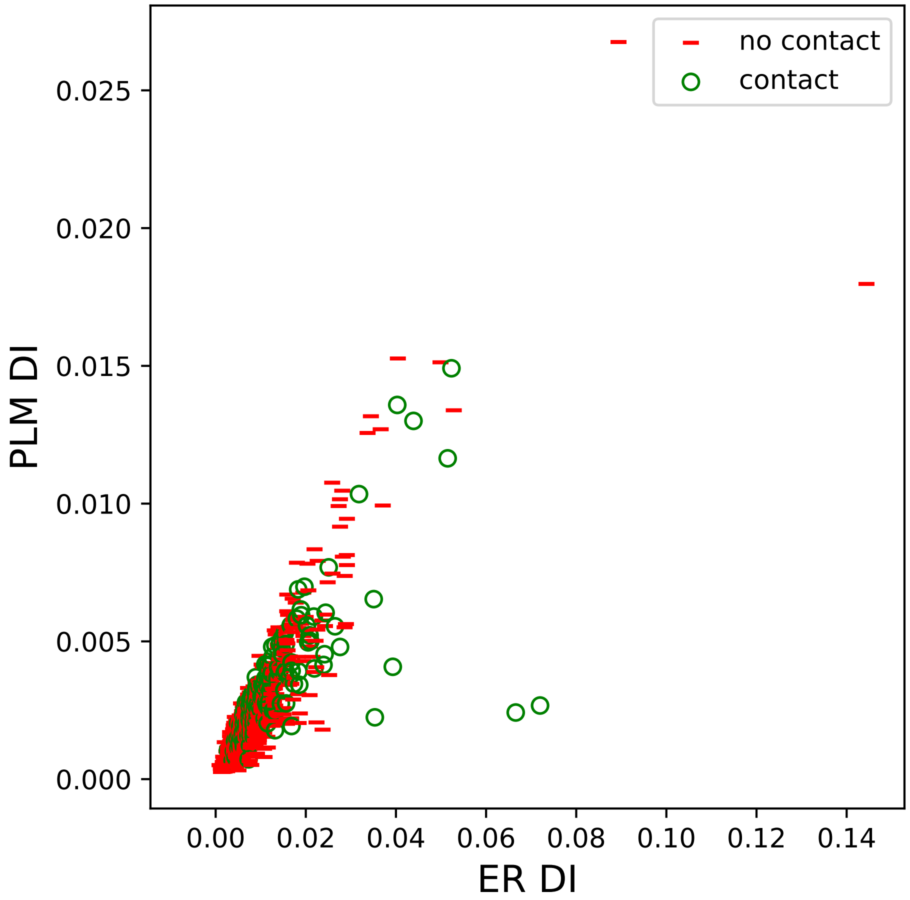
no contact
contact
PLM DI
ER DI
0.000
0.005
0.010
0.015
0.020
0.025
0.00
0.02
0.04
0.06
0.08
0.10
0.12
0.14

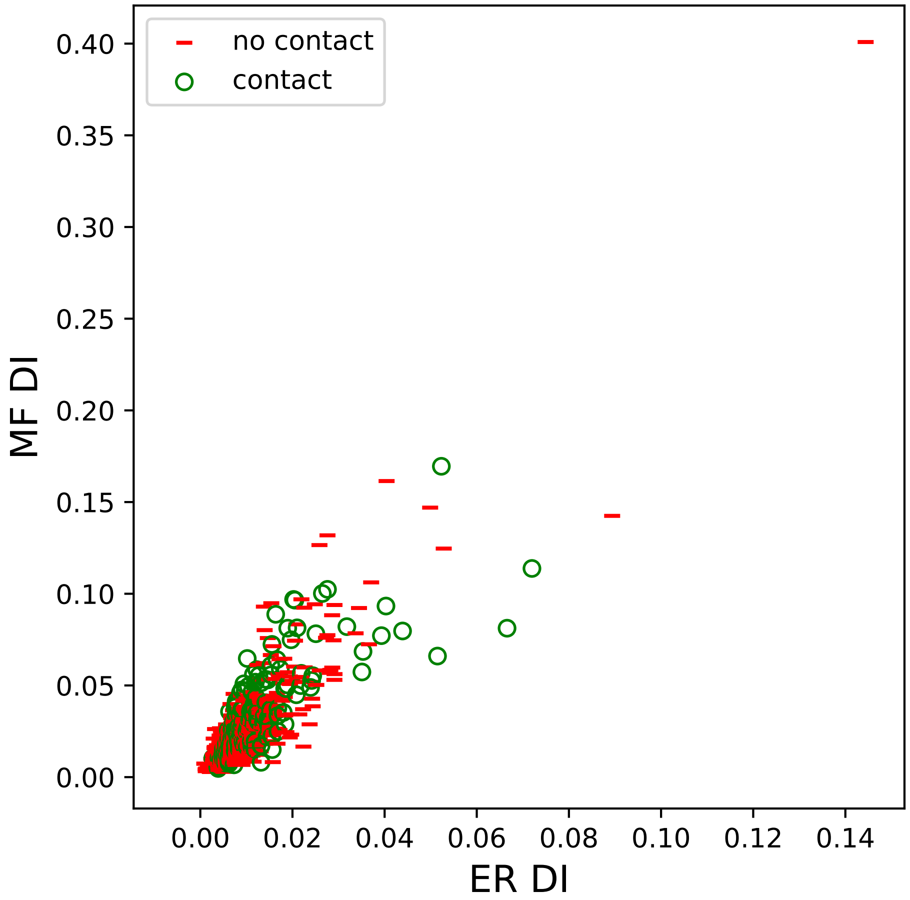

no contact
contact
MF DI
ER DI
0.00
0.05
0.10
0.15
0.20
0.25
0.30
0.35
0.40
0.00
0.02
0.04
0.06
0.08
0.10
0.12
0.14

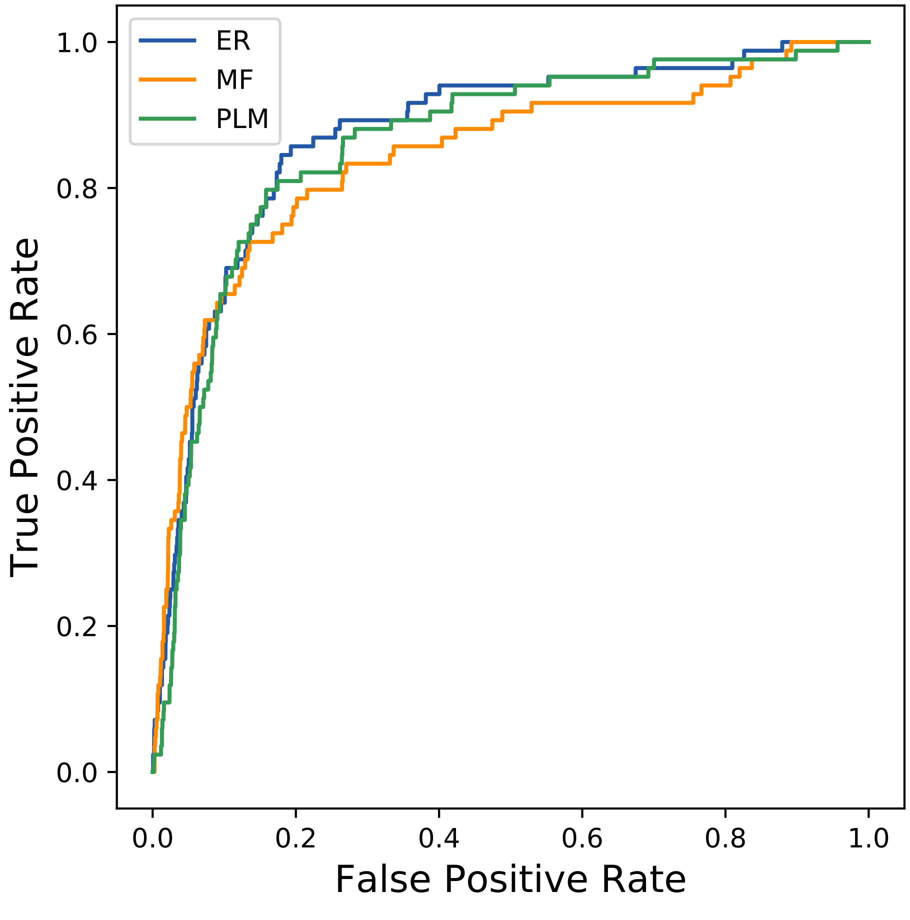
ER
MF
PLM
True Positive Rate
False Positive Rate
0.0
0.2
0.4
0.6
0.8
1.0
0.0
0.2
0.4
0.6
0.8
1.0
