## Supplemental Contact Maps for "Protein Structure Prediction with Expectation Reflection": 2996seq_116col_7jtb_PF00339_plots.pdf

Method: ER
Maximum PDB contact distance : 10.0 Angstrom
Minimum residue chain distance: 5.0 residues
Fraction of true positives : 0.638

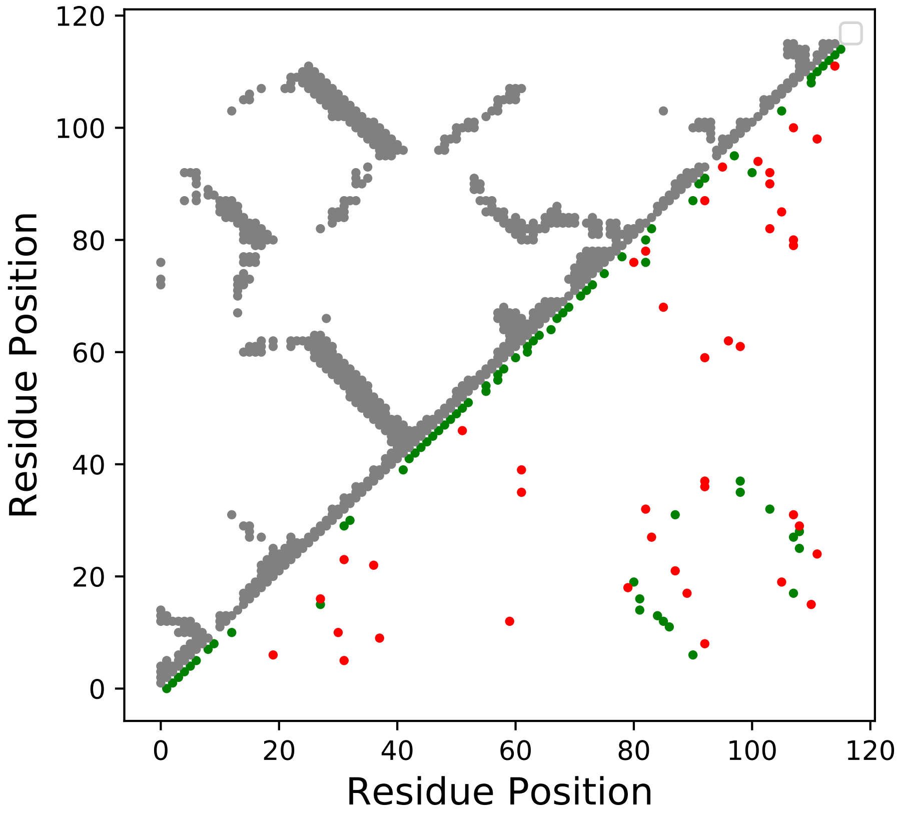

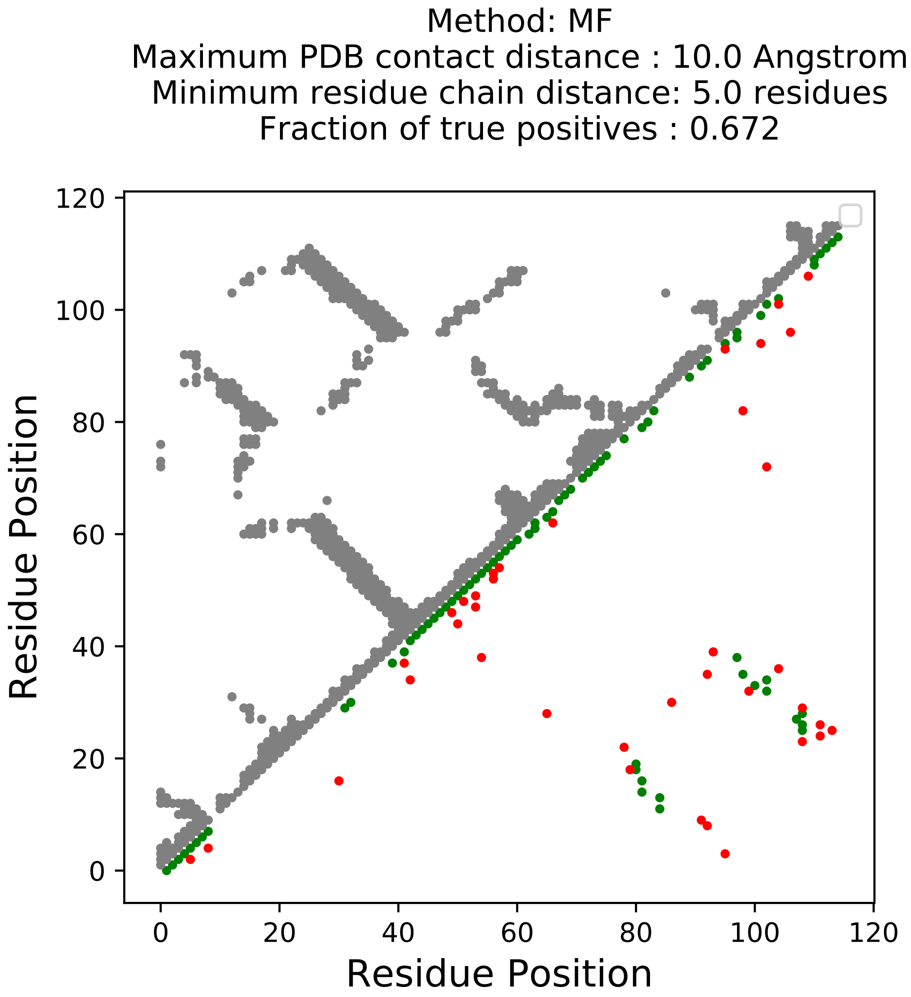
Method: MF
Maximum PDB contact distance : 10.0 Angstrom
Minimum residue chain distance: 5.0 residues
Fraction of true positives : 0.672
Residue Position
Residue Position
0
20
40
60
80
100
120

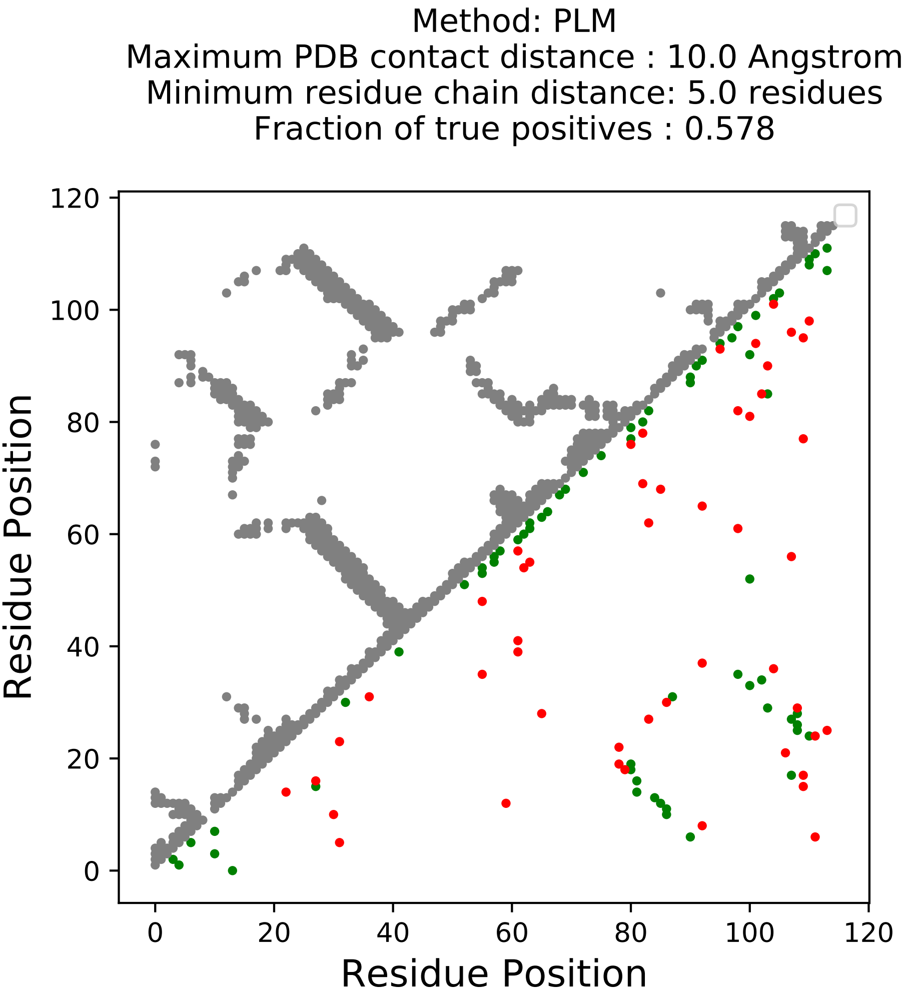
Method: PLM
Maximum PDB contact distance : 10.0 Angstrom
Minimum residue chain distance: 5.0 residues
Fraction of true positives : 0.578
Residue Position
Residue Position
0
20
40
60
80
100
120

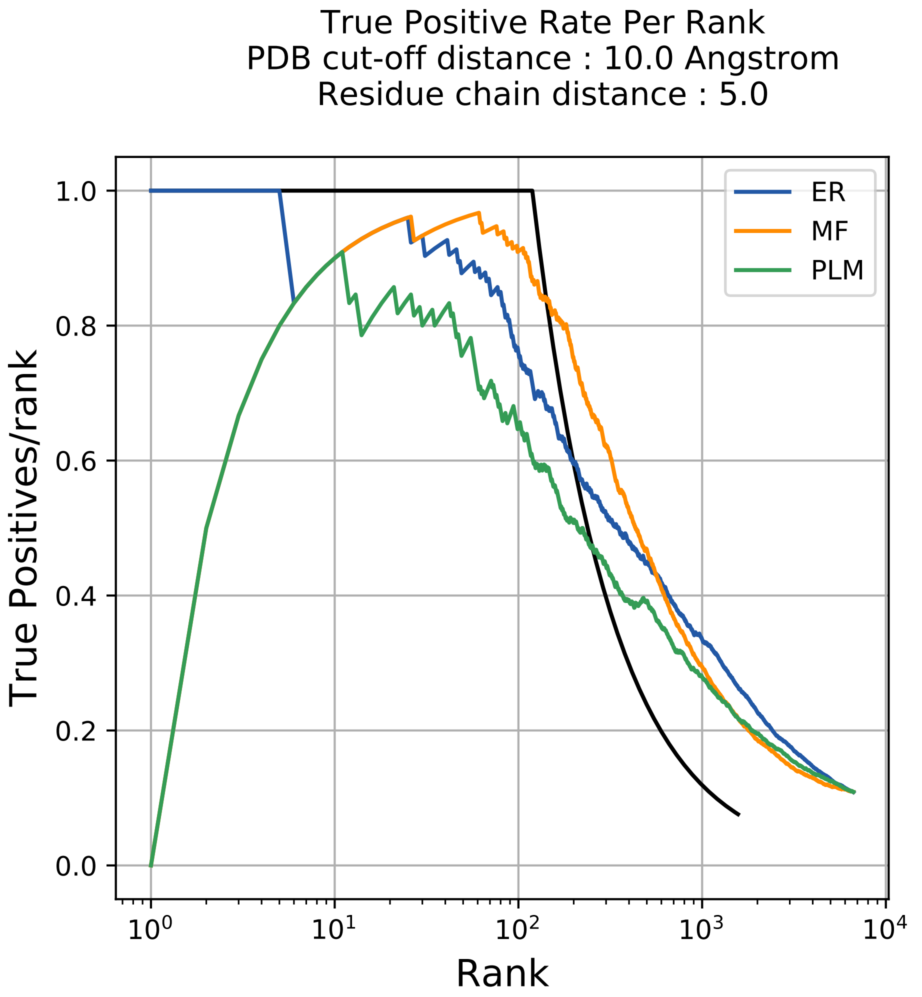
True Positive Rate Per Rank
PDB cut-off distance : 10.0 Angstrom
Residue chain distance : 5.0
True Positives/rank
Rank
1.0
0.8
0.6
0.4
0.2
0.0
$10^0$
$10^1$
$10^2$
$10^3$
$10^4$
ER
MF
PLM

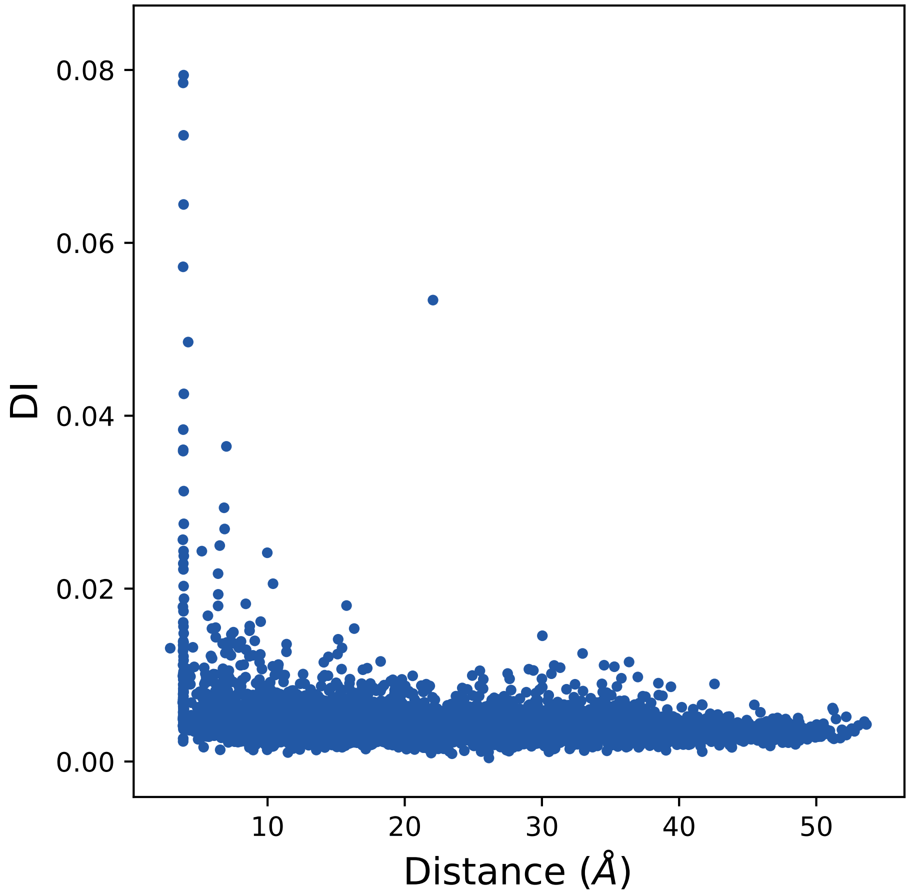

0.08
0.06
0.04
0.02
0.00
DI
10
20
30
40
50
Distance (Å)

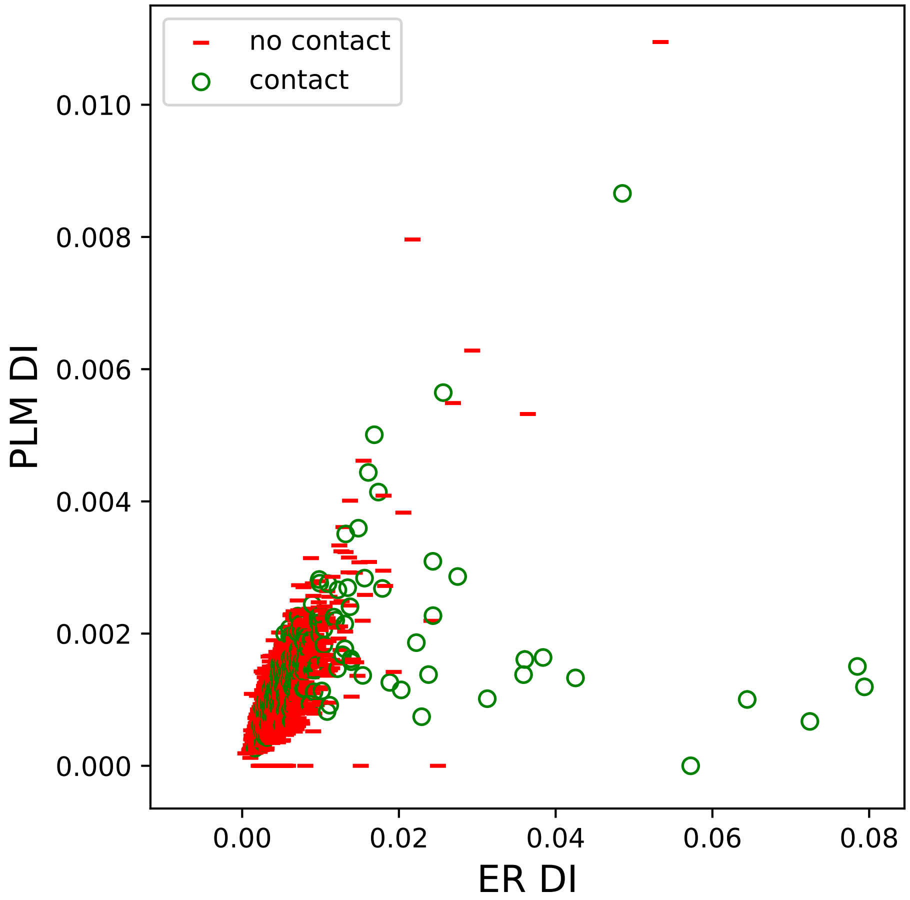
no contact
contact
0.010
0.008
0.006
0.004
0.002
0.000
PLM DI
0.00
0.02
0.04
0.06
0.08
ER DI

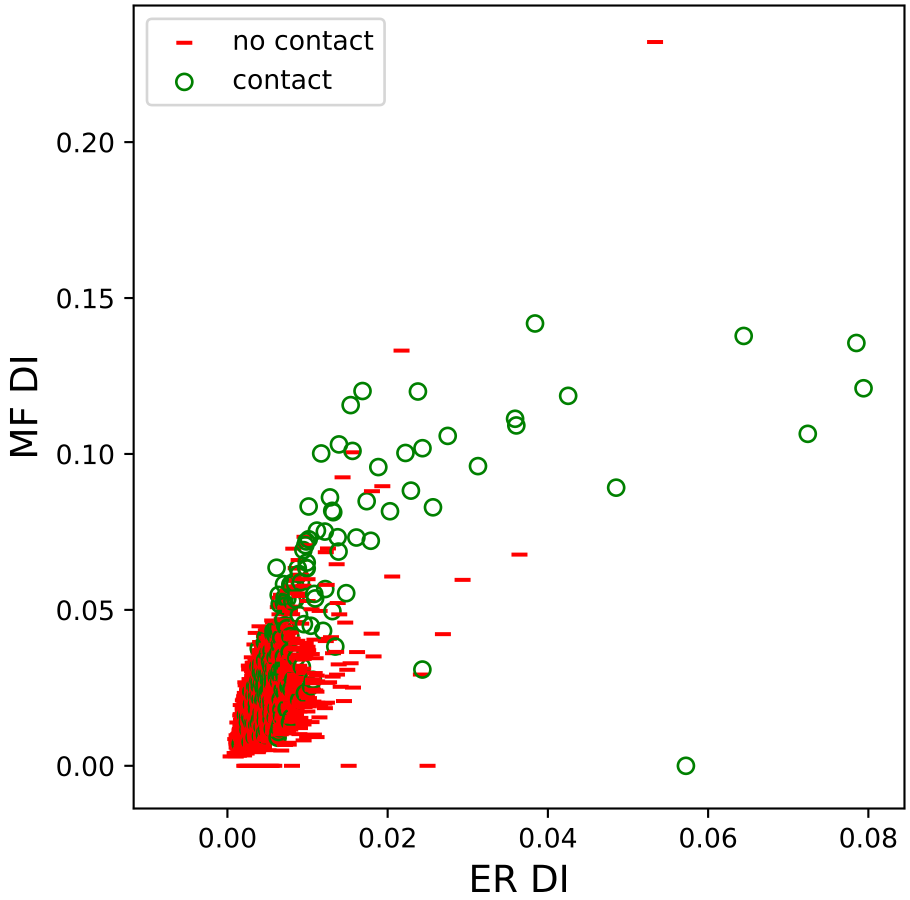

no contact
contact
MF DI
ER DI
0.00
0.05
0.10
0.15
0.20
0.00
0.02
0.04
0.06
0.08

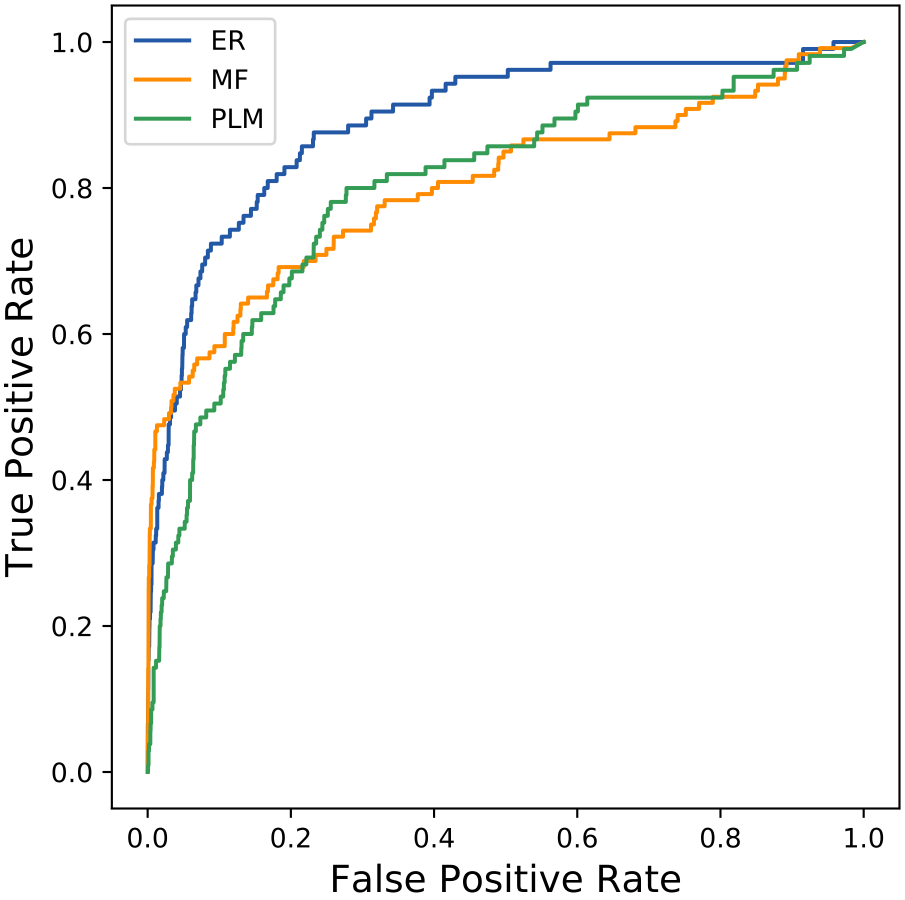

ER
MF
PLM
True Positive Rate
False Positive Rate
0.0
0.2
0.4
0.6
0.8
1.0
