## Supplemental Contact Maps for "Protein Structure Prediction with Expectation Reflection": 3139seq_161col_3zbo_PF08713_plots.pdf

Method: ER
Maximum PDB contact distance : 10.0 Angstrom
Minimum residue chain distance: 5.0 residues
Fraction of true positives : 0.665

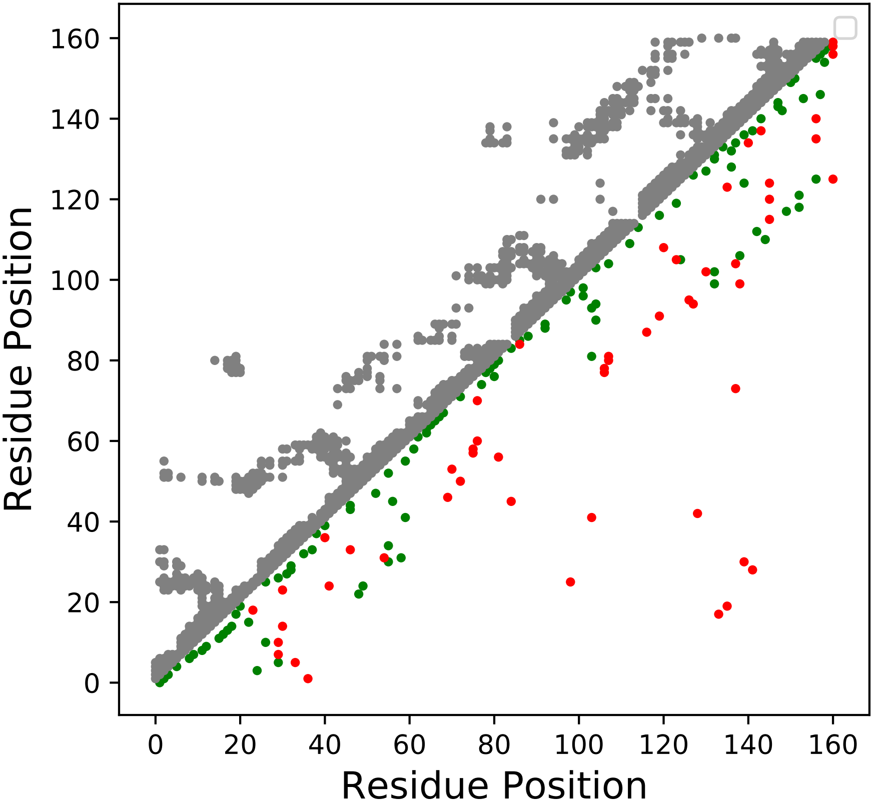

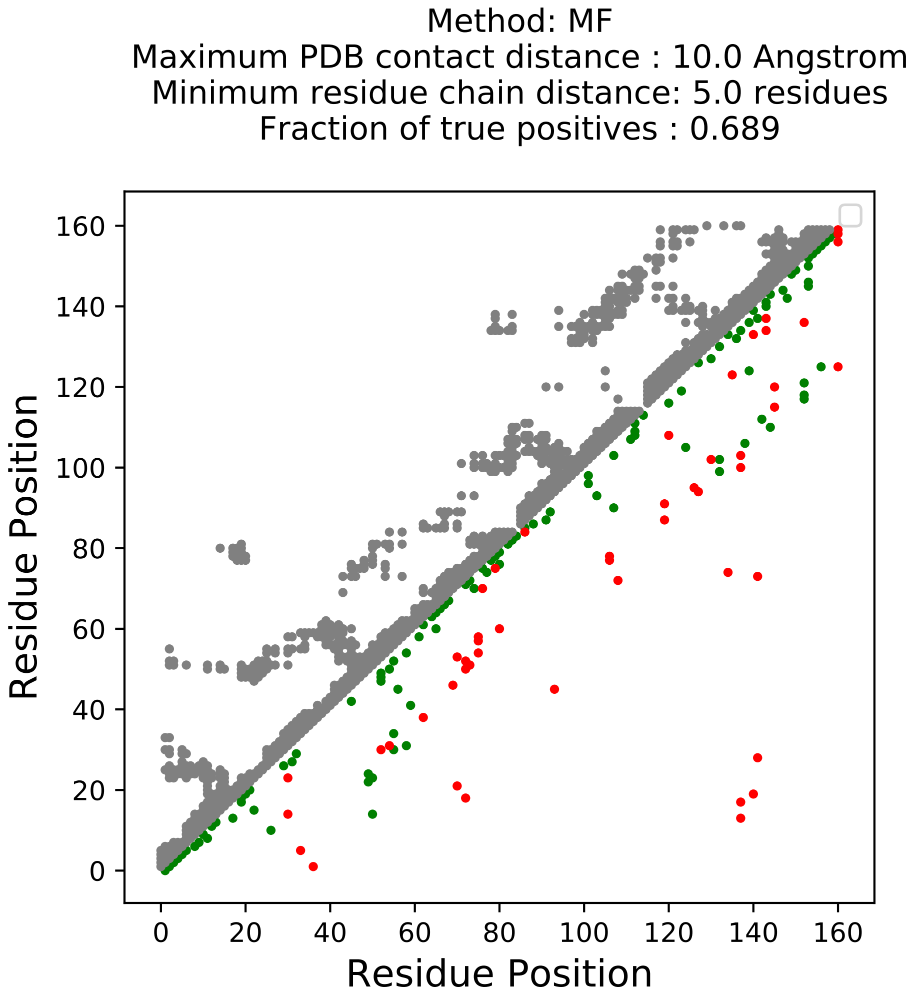
Method: MF
Maximum PDB contact distance : 10.0 Angstrom
Minimum residue chain distance: 5.0 residues
Fraction of true positives : 0.689
Residue Position
Residue Position
0
20
40
60
80
100
120
140
160

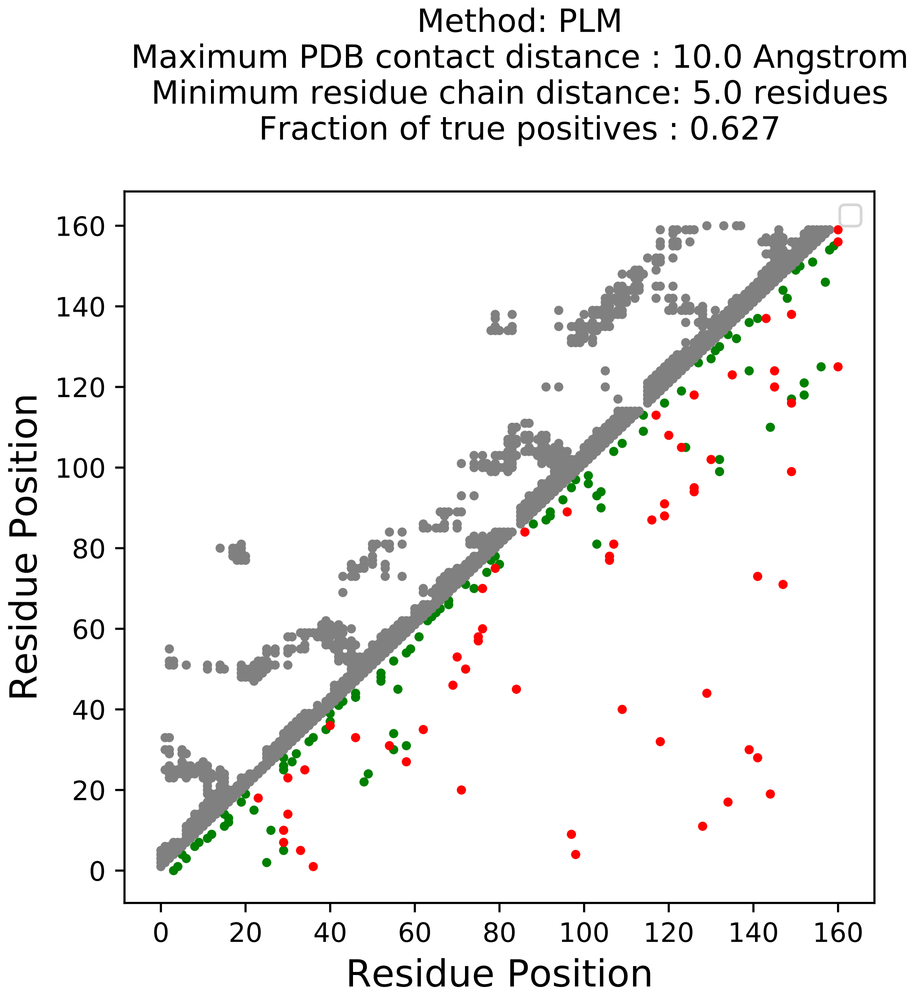
Method: PLM
Maximum PDB contact distance : 10.0 Angstrom
Minimum residue chain distance: 5.0 residues
Fraction of true positives : 0.627
Residue Position
Residue Position
0
20
40
60
80
100
120
140
160

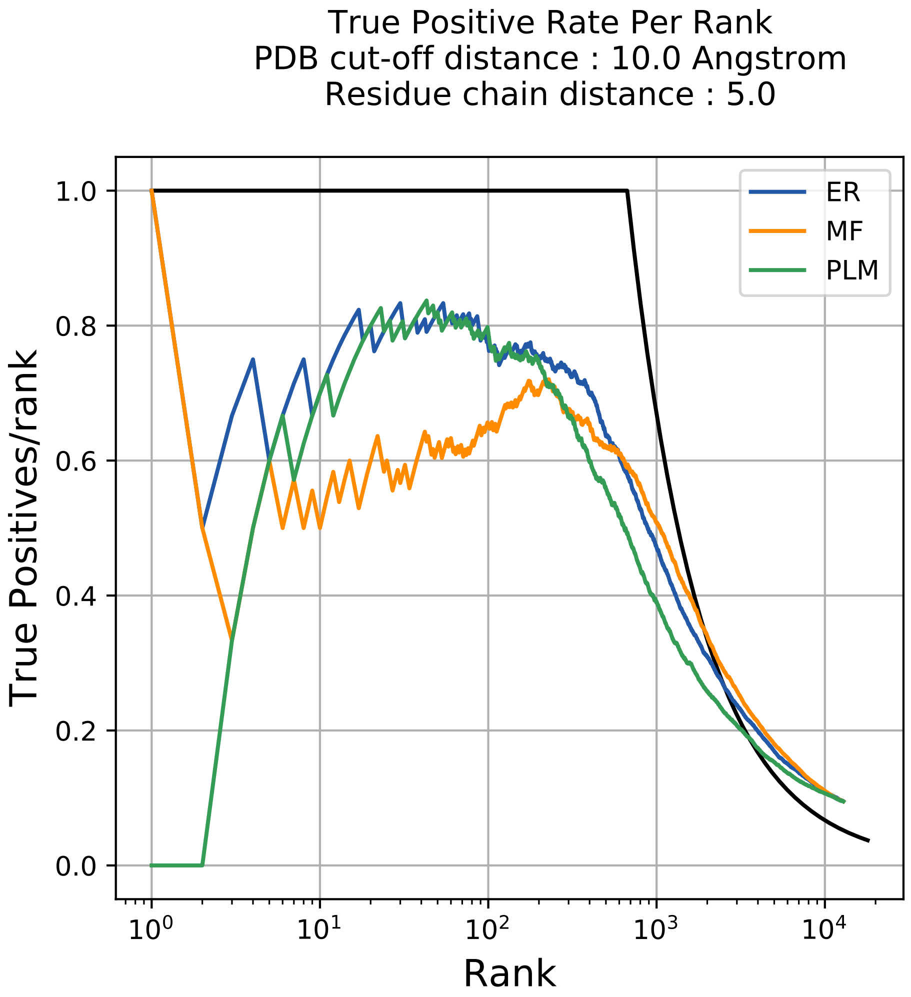

True Positive Rate Per Rank
PDB cut-off distance : 10.0 Angstrom
Residue chain distance : 5.0
True Positives/rank
Rank
ER
MF
PLM
1.0
0.8
0.6
0.4
0.2
0.0
$10^0$
$10^1$
$10^2$
$10^3$
$10^4$

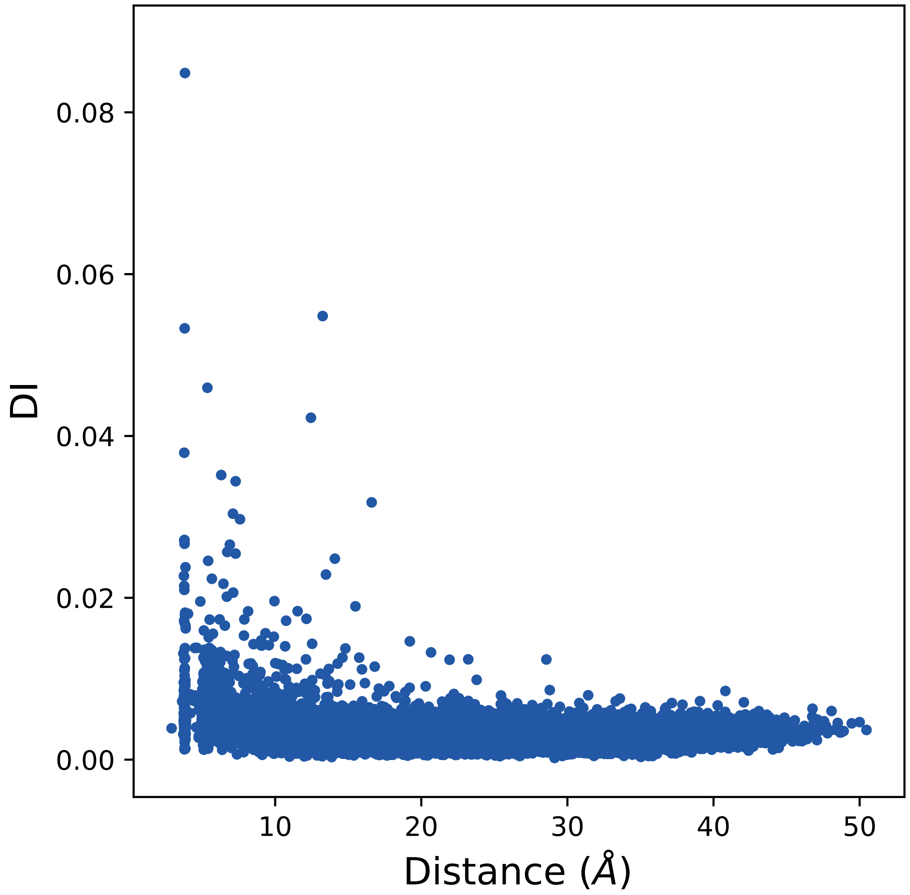

0.08
0.06
0.04
0.02
0.00
DI
10
20
30
40
50
Distance (Å)

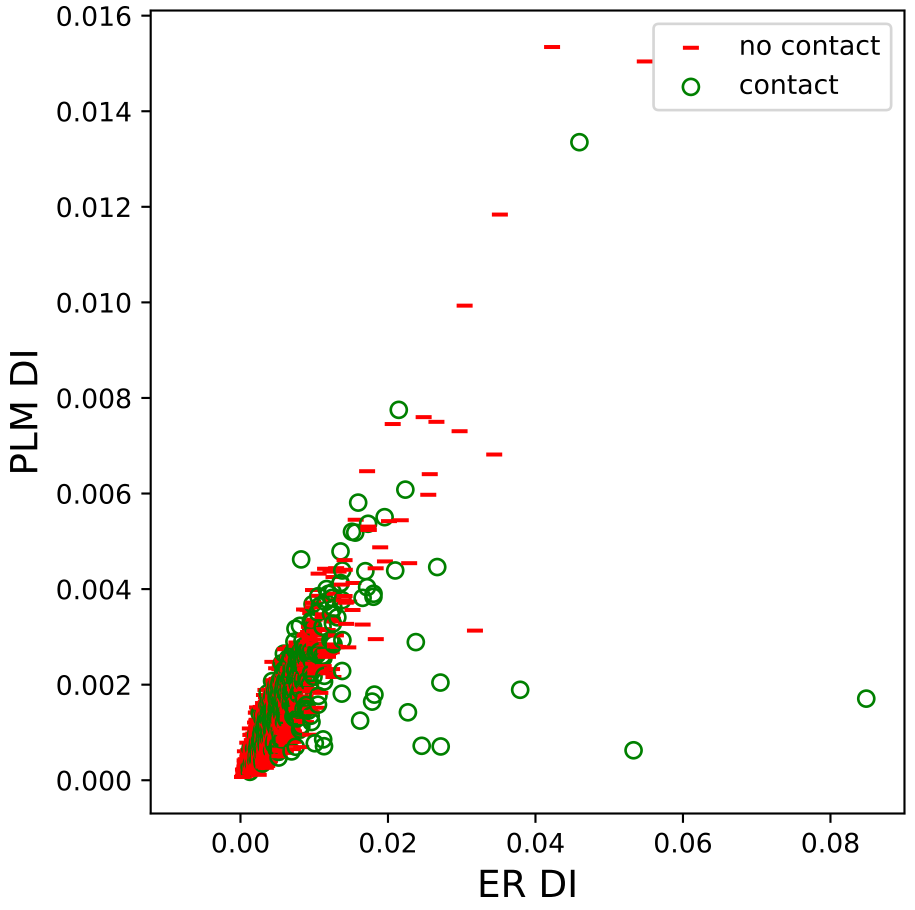

no contact
contact
PLM DI
ER DI
0.016
0.014
0.012
0.010
0.008
0.006
0.004
0.002
0.000
0.00
0.02
0.04
0.06
0.08

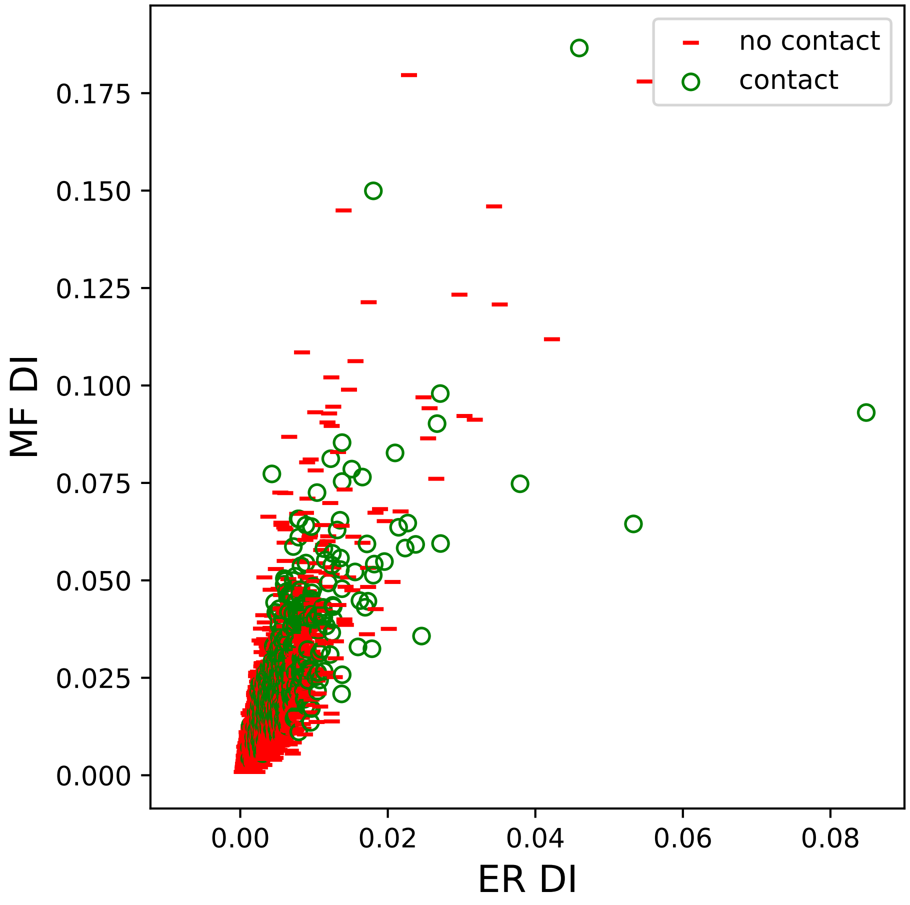

no contact
contact
MF DI
ER DI
0.000
0.025
0.050
0.075
0.100
0.125
0.150
0.175
0.00
0.02
0.04
0.06
0.08

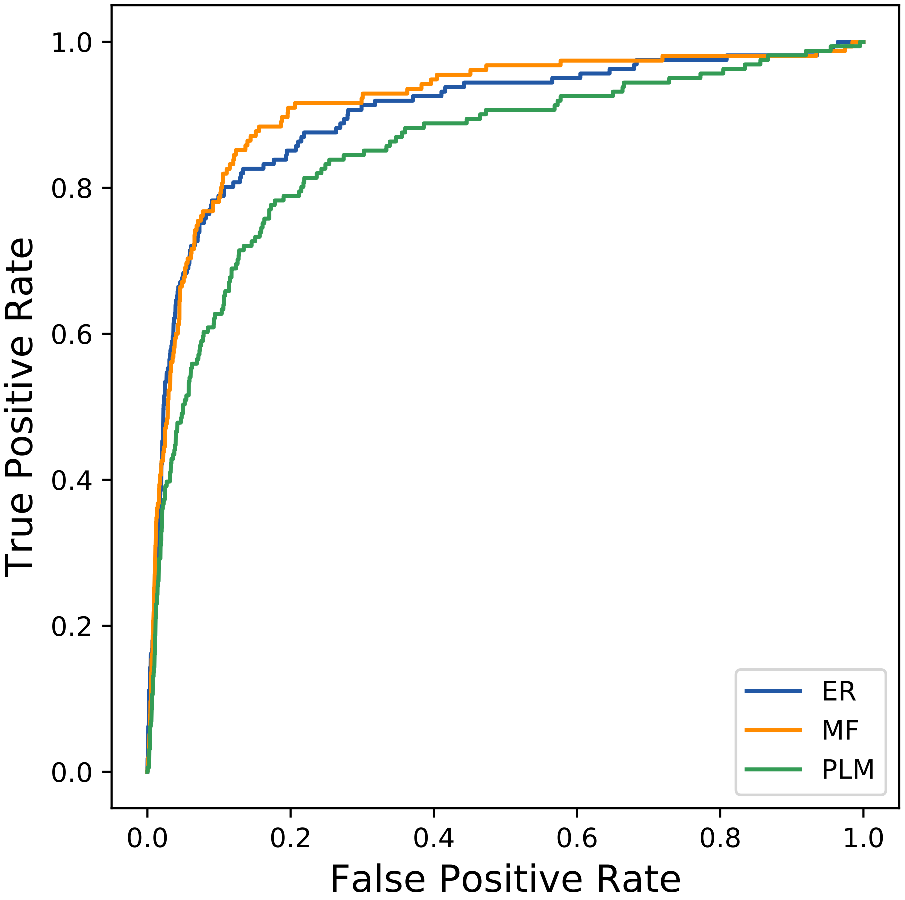

True Positive Rate
False Positive Rate
0.0
0.2
0.4
0.6
0.8
1.0
0.0
0.2
0.4
0.6
0.8
1.0
ER
MF
PLM
