## Supplemental Contact Maps for "Protein Structure Prediction with Expectation Reflection": 3207seq_91col_2zbt_PF01680_plots.pdf

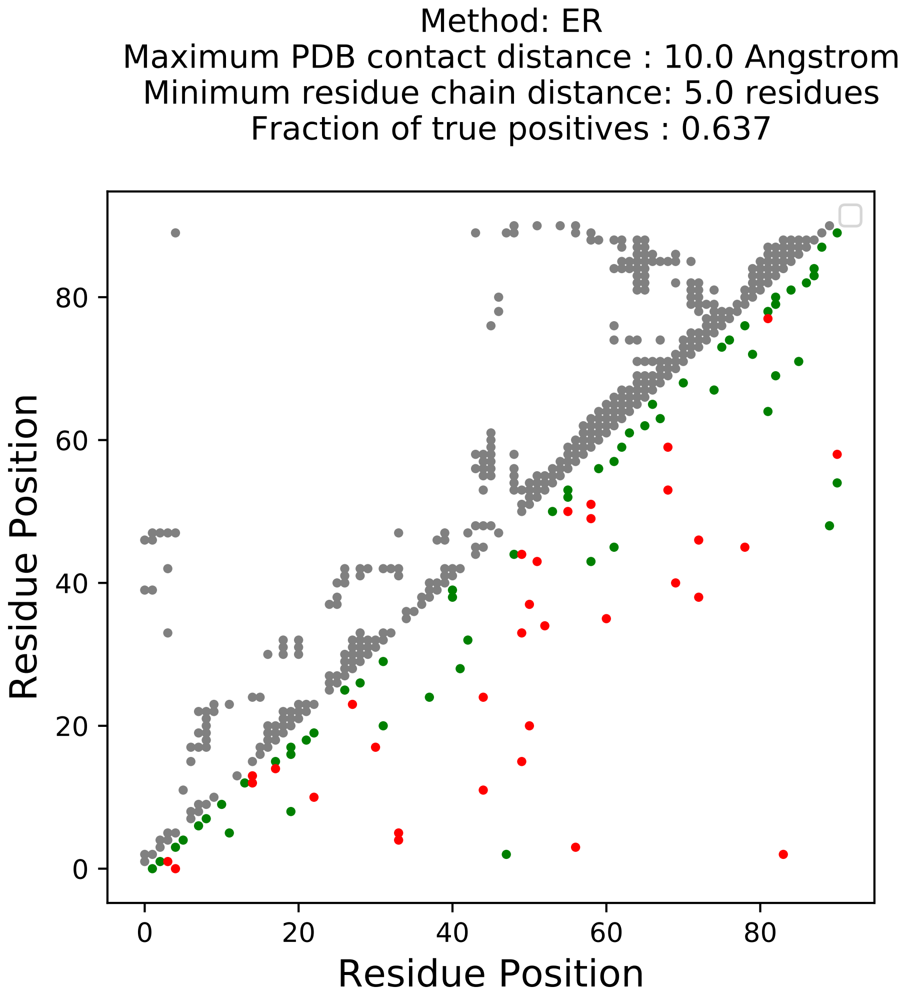
Method: ER
Maximum PDB contact distance : 10.0 Angstrom
Minimum residue chain distance: 5.0 residues
Fraction of true positives : 0.637
Residue Position
Residue Position
0
20
40
60
80
0
20
40
60
80

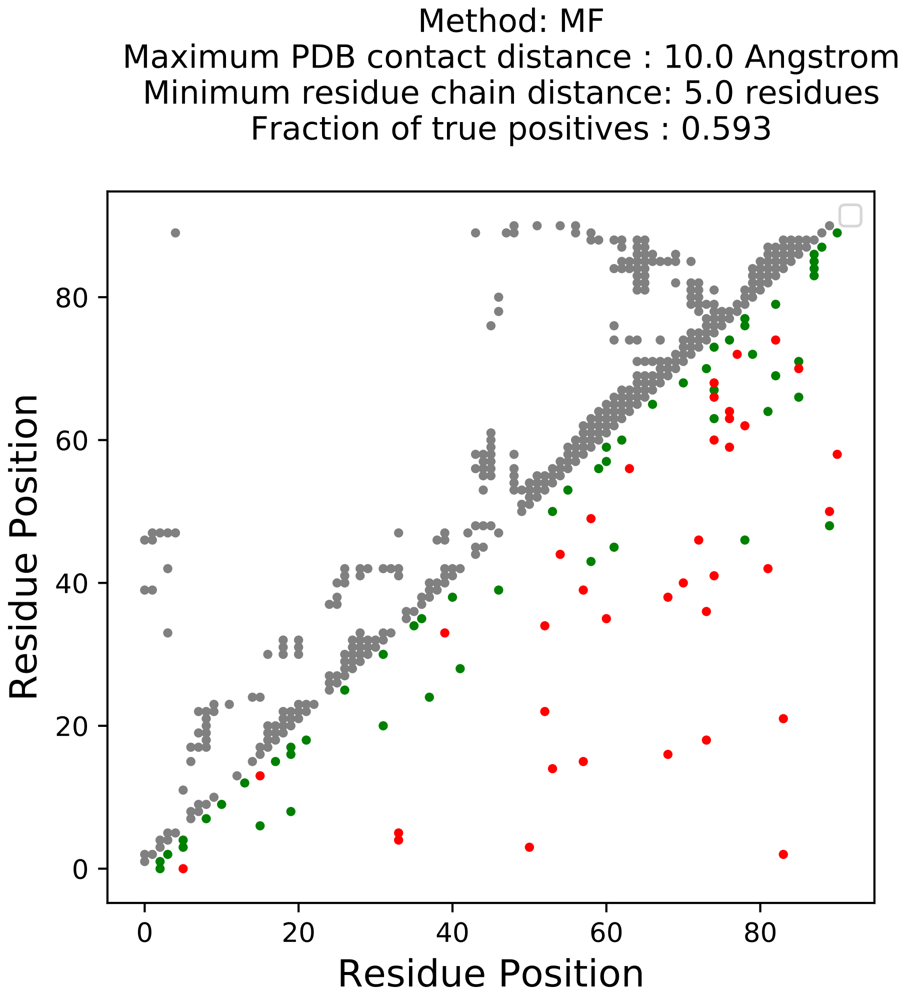
Method: MF
Maximum PDB contact distance : 10.0 Angstrom
Minimum residue chain distance: 5.0 residues
Fraction of true positives : 0.593
Residue Position
Residue Position
0
20
40
60
80

Method: PLM
Maximum PDB contact distance : 10.0 Angstrom
Minimum residue chain distance: 5.0 residues
Fraction of true positives : 0.495

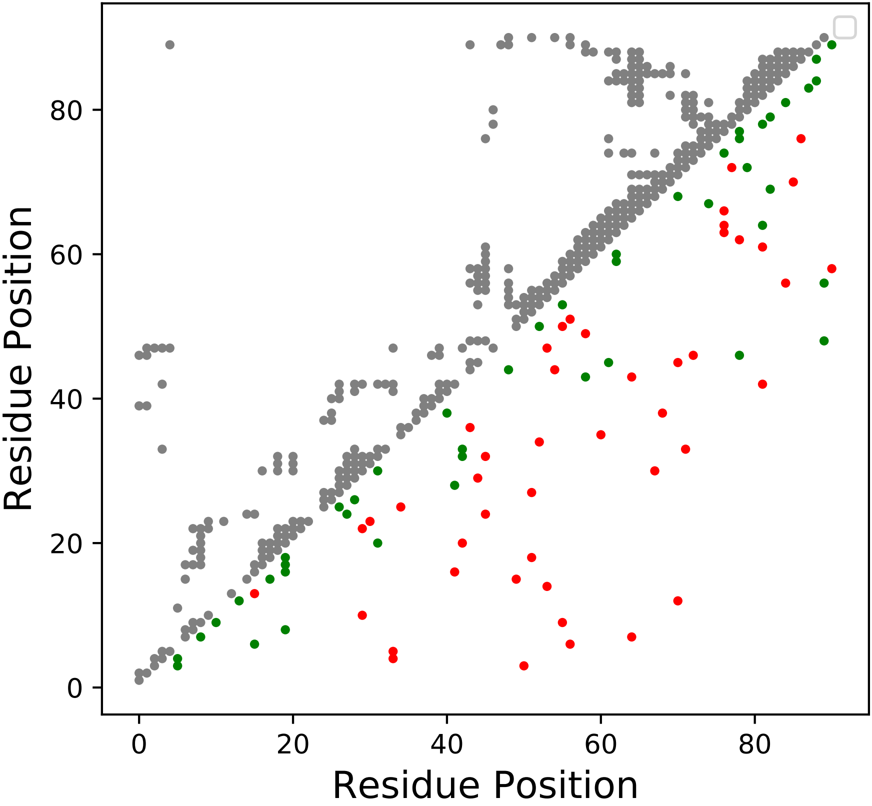

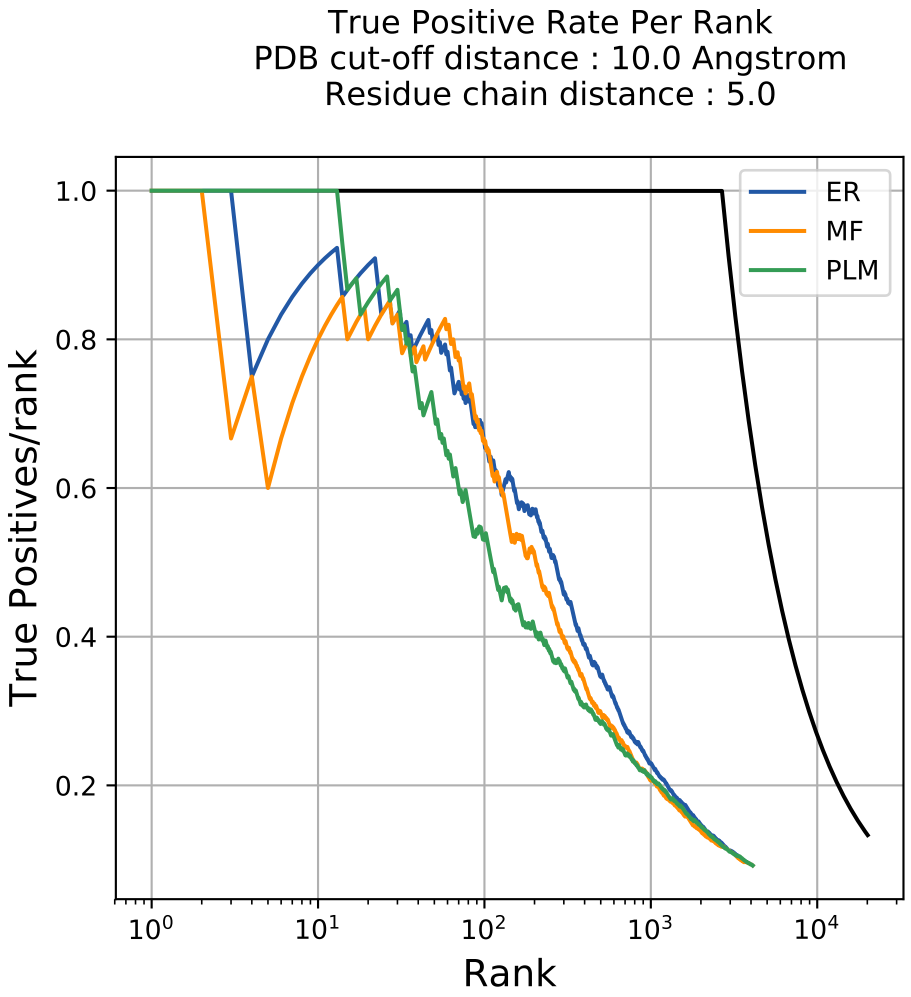

True Positive Rate Per Rank
PDB cut-off distance : 10.0 Angstrom
Residue chain distance : 5.0
True Positives/rank
Rank
ER
MF
PLM
1.0
0.8
0.6
0.4
0.2
$10^0$
$10^1$
$10^2$
$10^3$
$10^4$

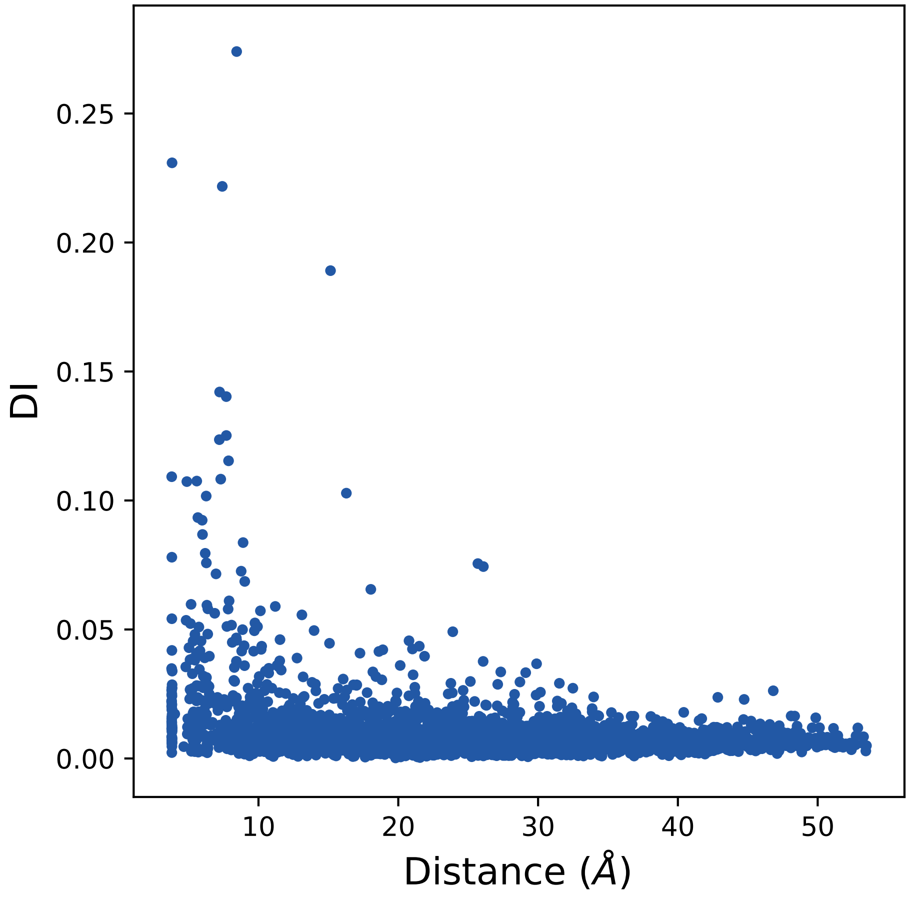

0.25
0.20
0.15
0.10
0.05
0.00
DI
10
20
30
40
50
Distance (Å)

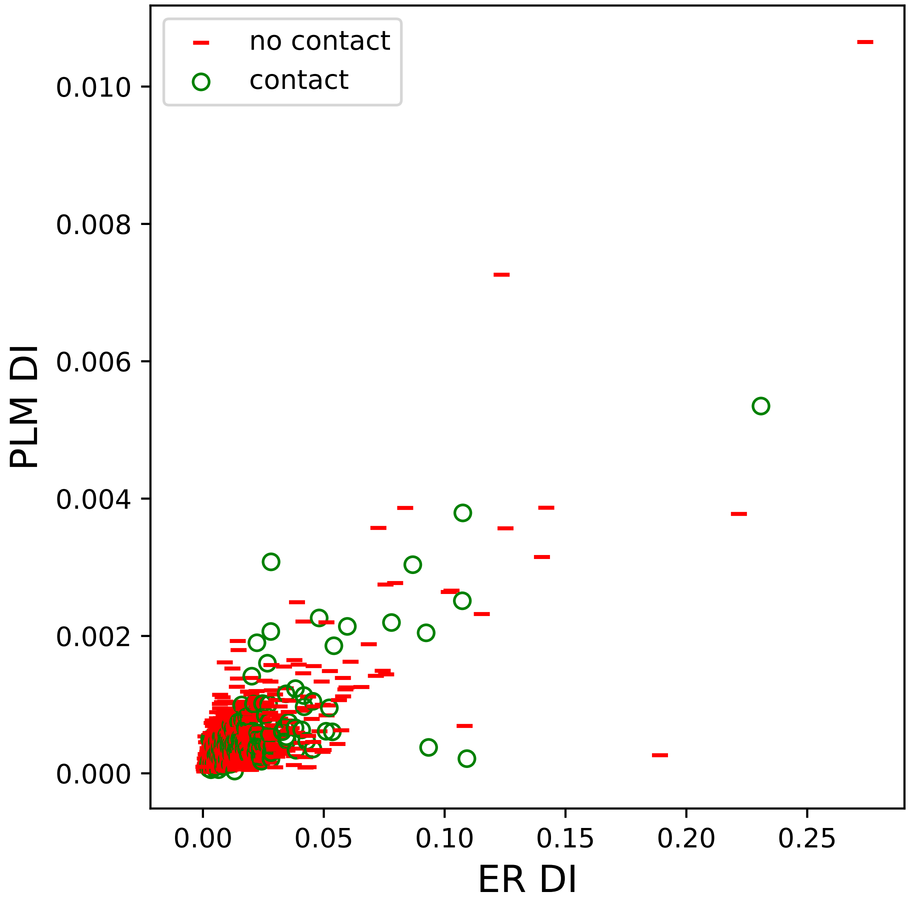

no contact
contact
0.010
0.008
0.006
0.004
0.002
0.000
PLM DI
0.00
0.05
0.10
0.15
0.20
0.25
ER DI

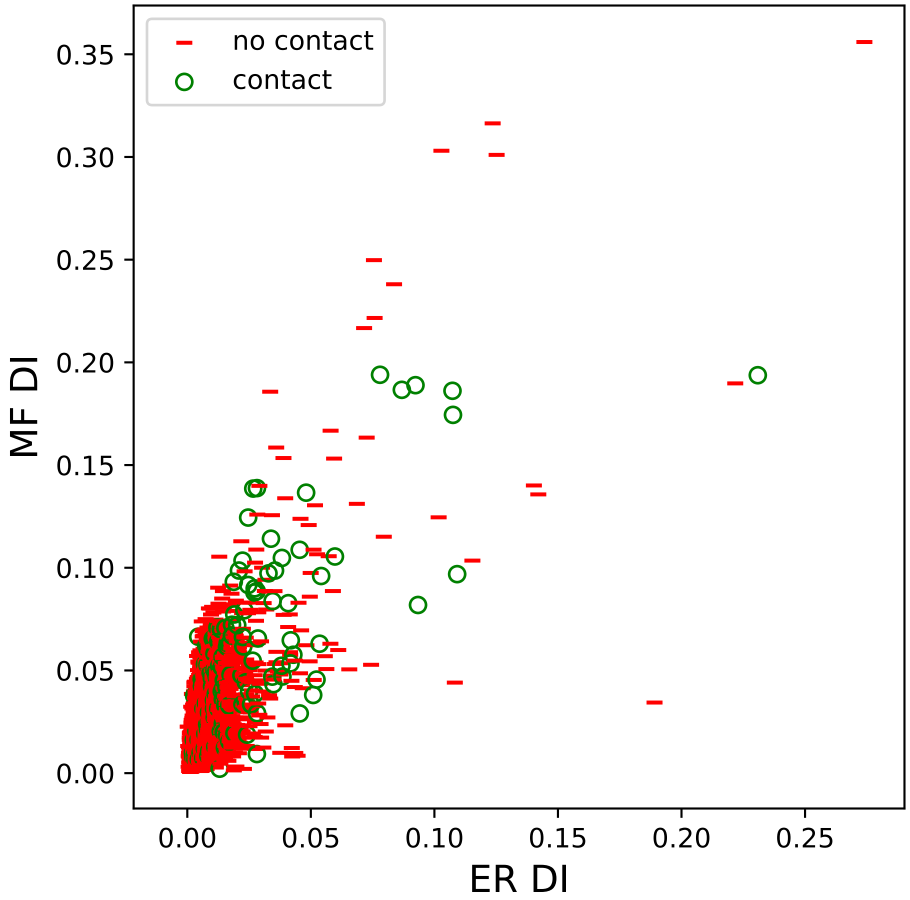
no contact
contact
0.35
0.30
0.25
0.20
0.15
0.10
0.05
0.00
MF DI
0.00
0.05
0.10
0.15
0.20
0.25
ER DI

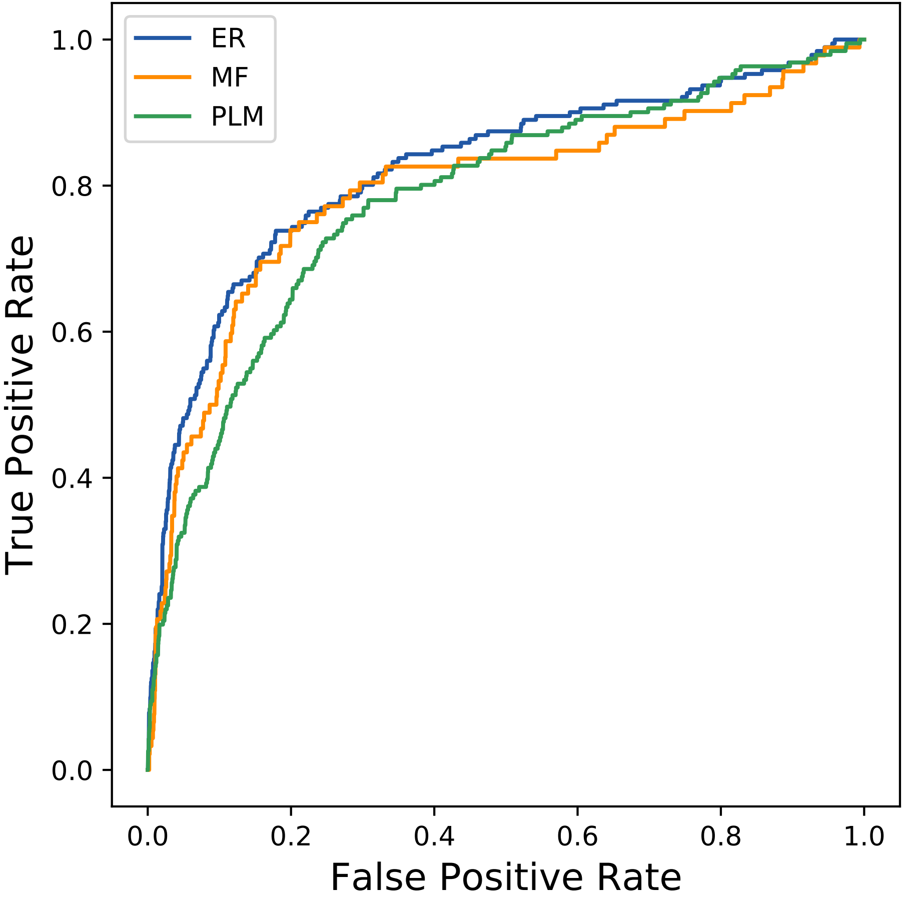
ER
MF
PLM
True Positive Rate
False Positive Rate
0.0
0.2
0.4
0.6
0.8
1.0
