## Supplemental Contact Maps for "Protein Structure Prediction with Expectation Reflection": 3426seq_76col_3iwu_PF00576_plots.pdf

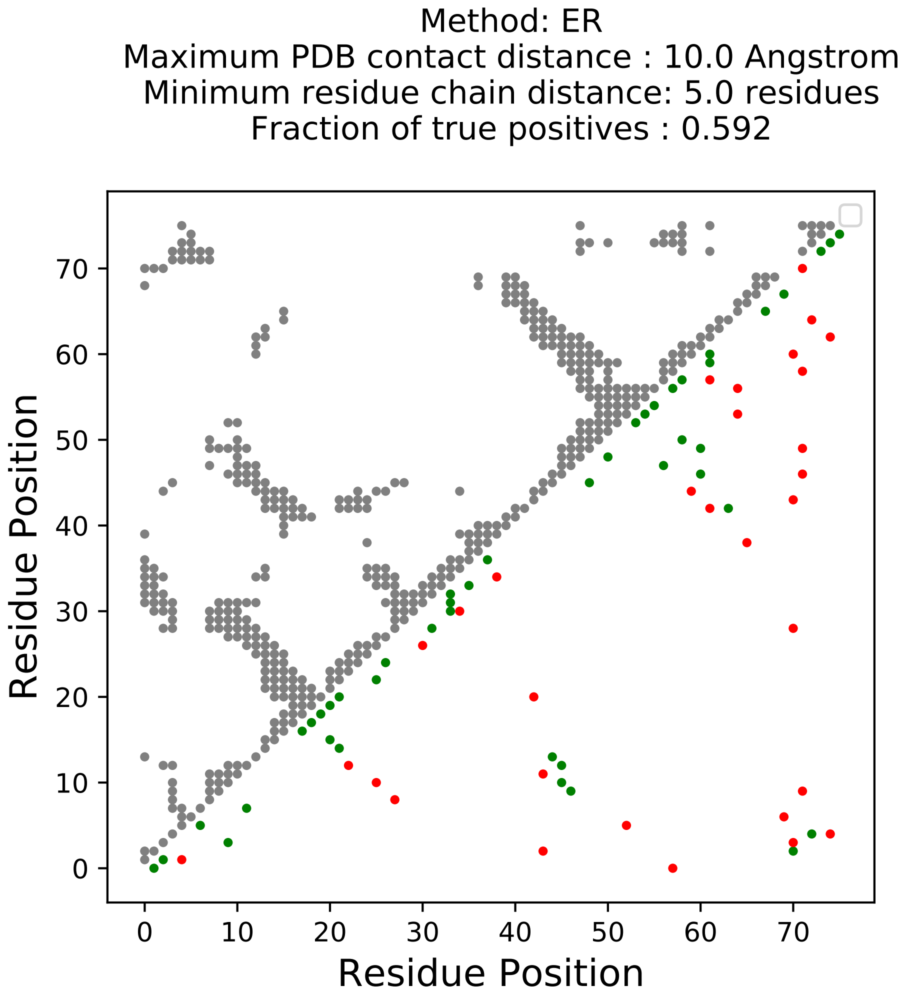
Method: ER
Maximum PDB contact distance : 10.0 Angstrom
Minimum residue chain distance: 5.0 residues
Fraction of true positives : 0.592
Residue Position
Residue Position
0
10
20
30
40
50
60
70

Method: MF
Maximum PDB contact distance : 10.0 Angstrom
Minimum residue chain distance: 5.0 residues
Fraction of true positives : 0.605

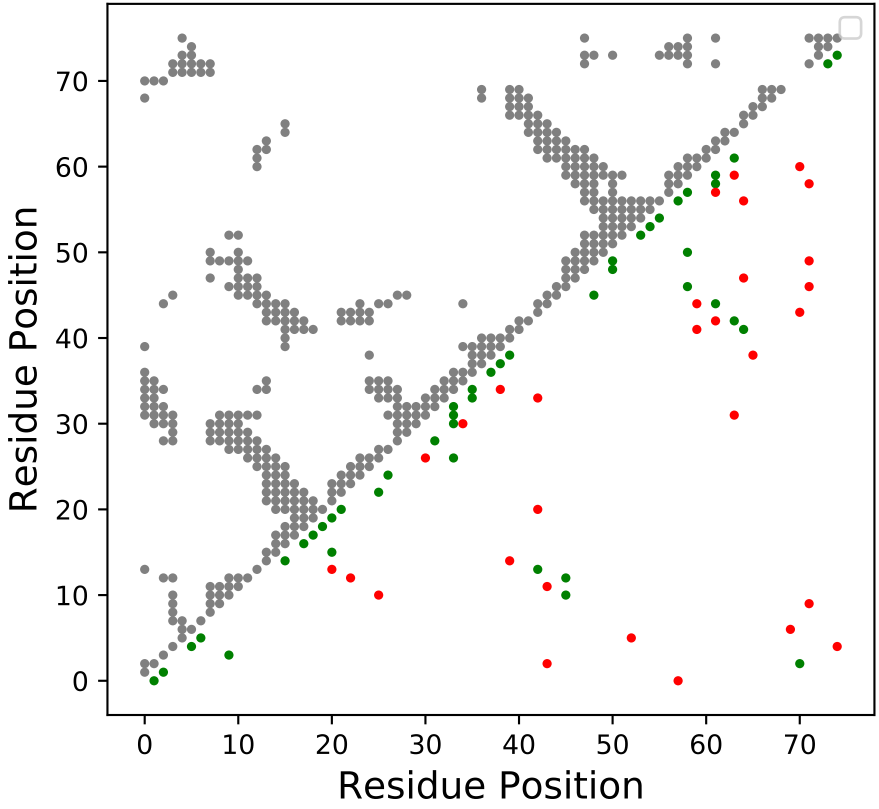

Method: PLM
Maximum PDB contact distance : 10.0 Angstrom
Minimum residue chain distance: 5.0 residues
Fraction of true positives : 0.618

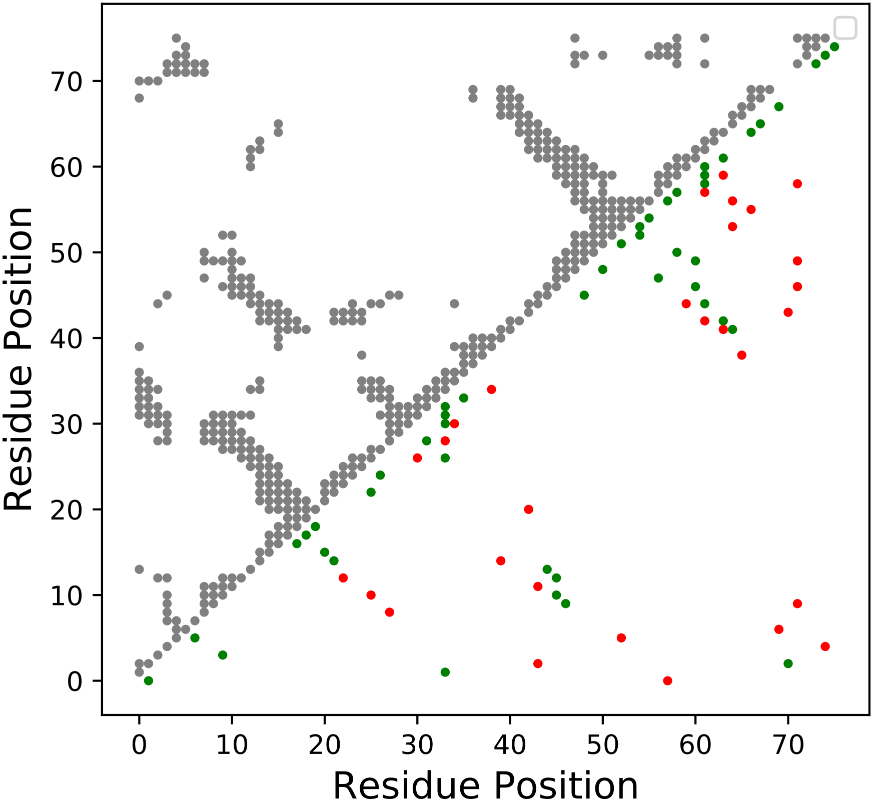

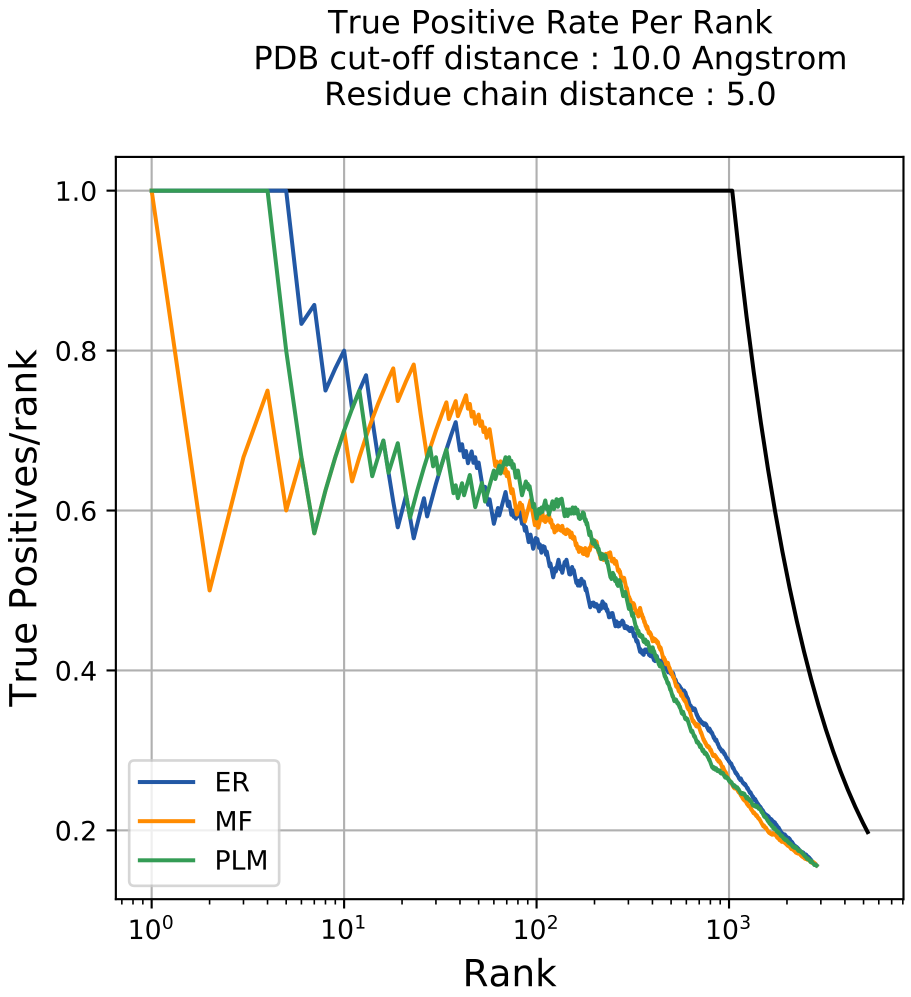

True Positive Rate Per Rank
PDB cut-off distance : 10.0 Angstrom
Residue chain distance : 5.0
True Positives/rank
Rank
1.0
0.8
0.6
0.4
0.2
$10^0$
$10^1$
$10^2$
$10^3$
ER
MF
PLM

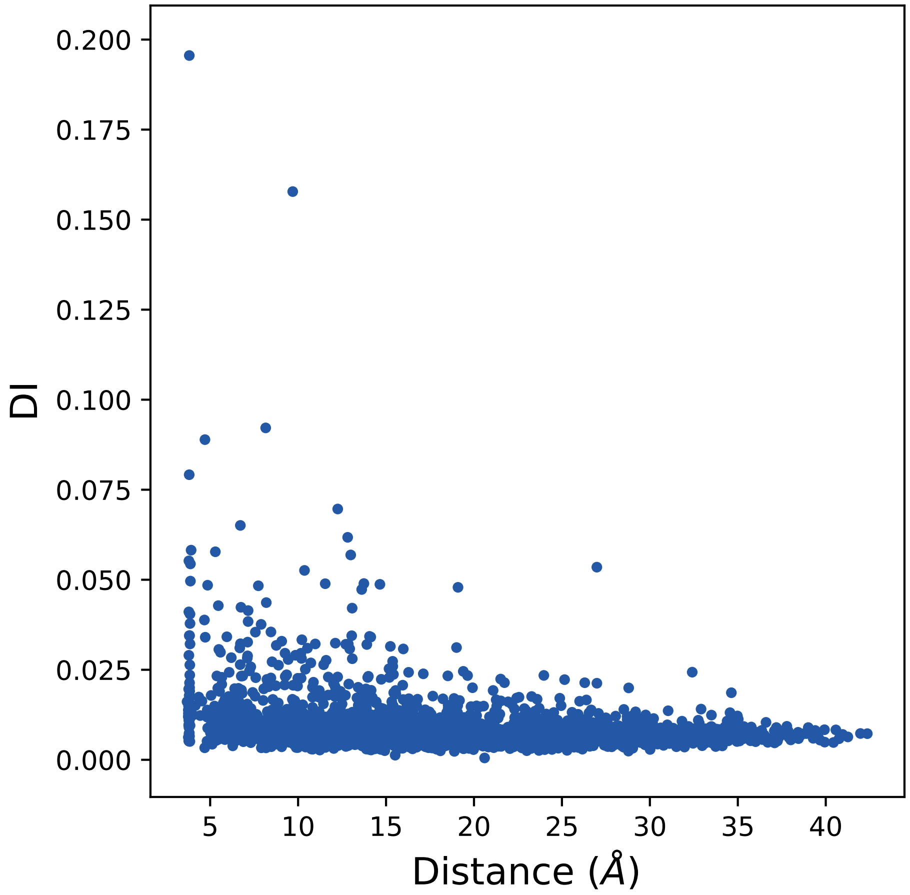

0.200
0.175
0.150
0.125
0.100
0.075
0.050
0.025
0.000
DI
5
10
15
20
25
30
35
40
Distance (Å)

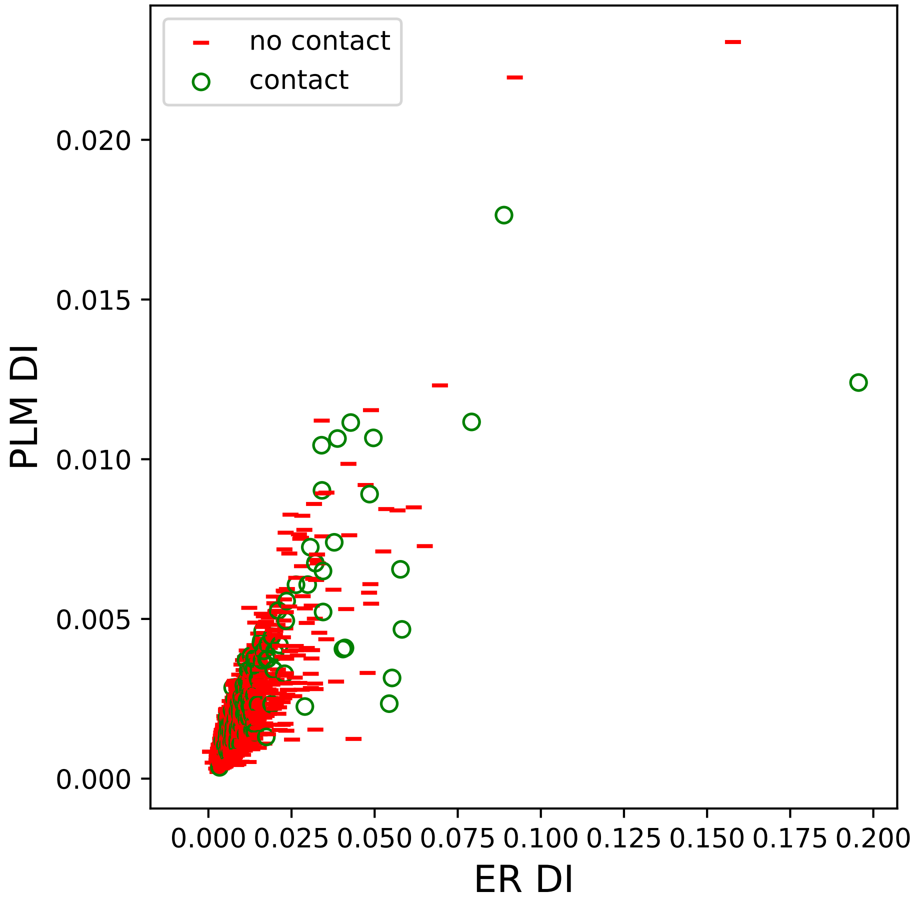
no contact
contact
0.000
0.005
0.010
0.015
0.020
PLM DI
0.000
0.025
0.050
0.075
0.100
0.125
0.150
0.175
0.200
ER DI

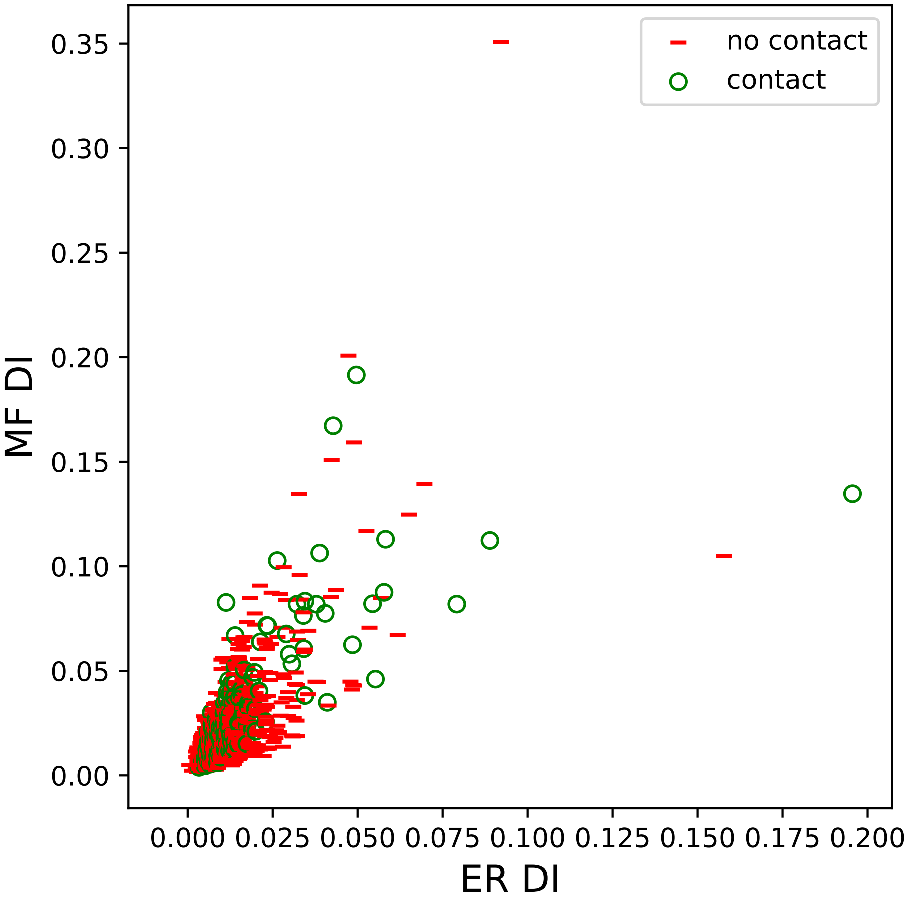

no contact
contact
MF DI
ER DI
0.00
0.05
0.10
0.15
0.20
0.25
0.30
0.35
0.000
0.025
0.050
0.075
0.100
0.125
0.150
0.175
0.200

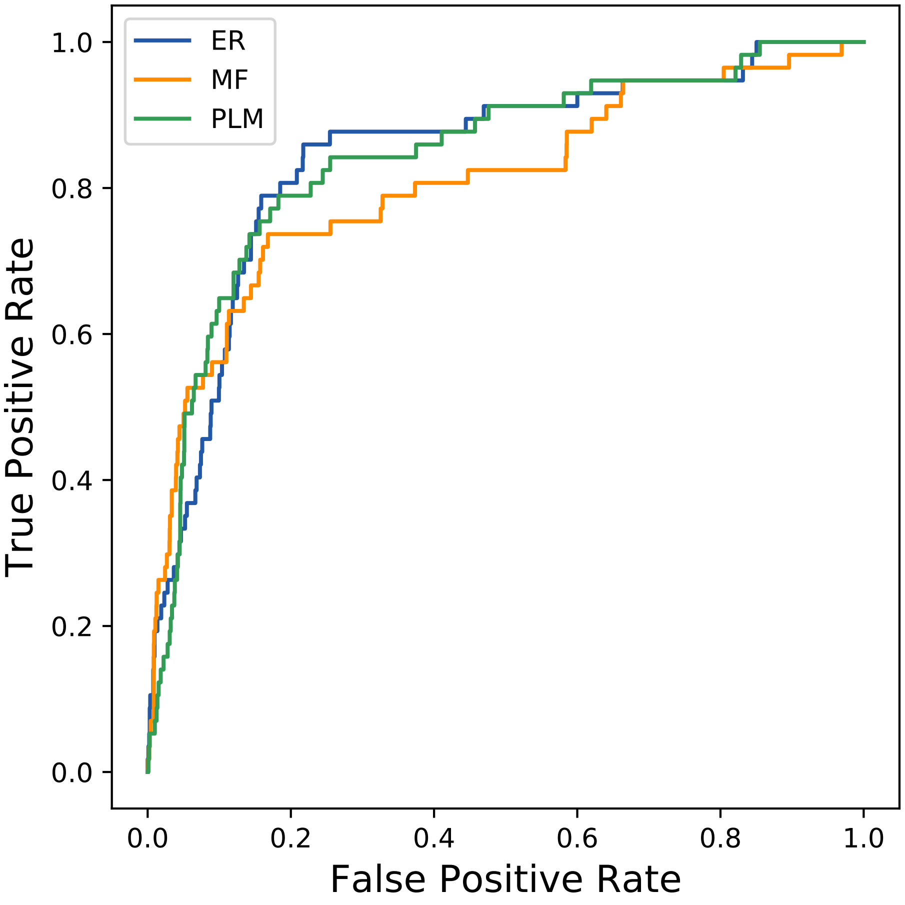

ER
MF
PLM
1.0
0.8
0.6
0.4
0.2
0.0
0.0
0.2
0.4
0.6
0.8
1.0
True Positive Rate
False Positive Rate
