## Supplemental Contact Maps for "Protein Structure Prediction with Expectation Reflection": 3912seq_35col_1s0y_PF01361_plots.pdf

Method: ER
Maximum PDB contact distance : 10.0 Angstrom
Minimum residue chain distance: 5.0 residues
Fraction of true positives : 0.629

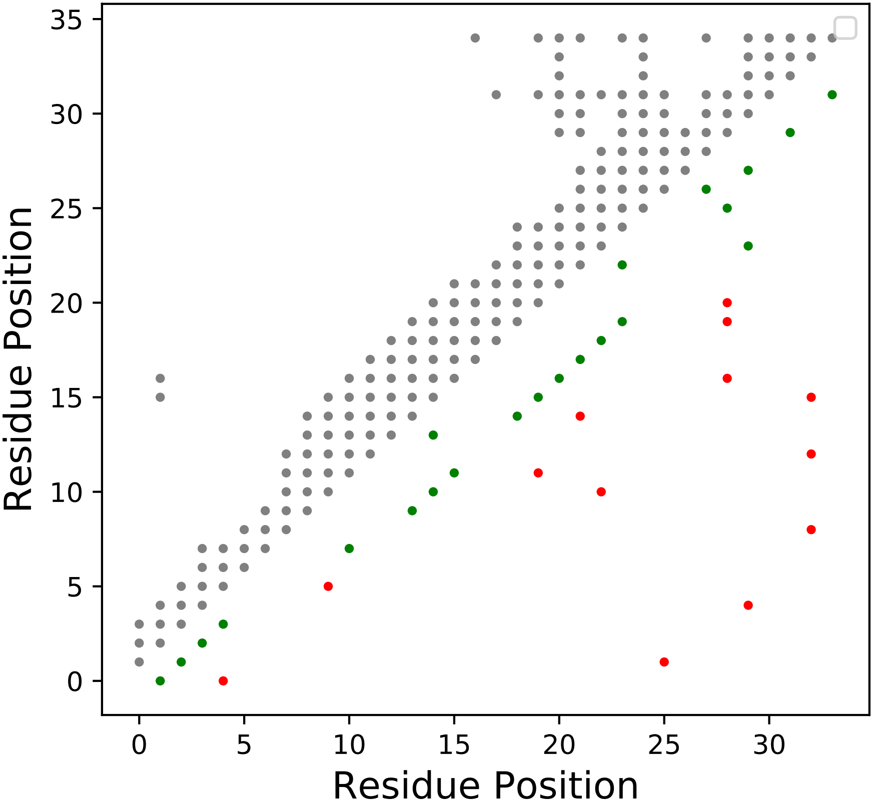

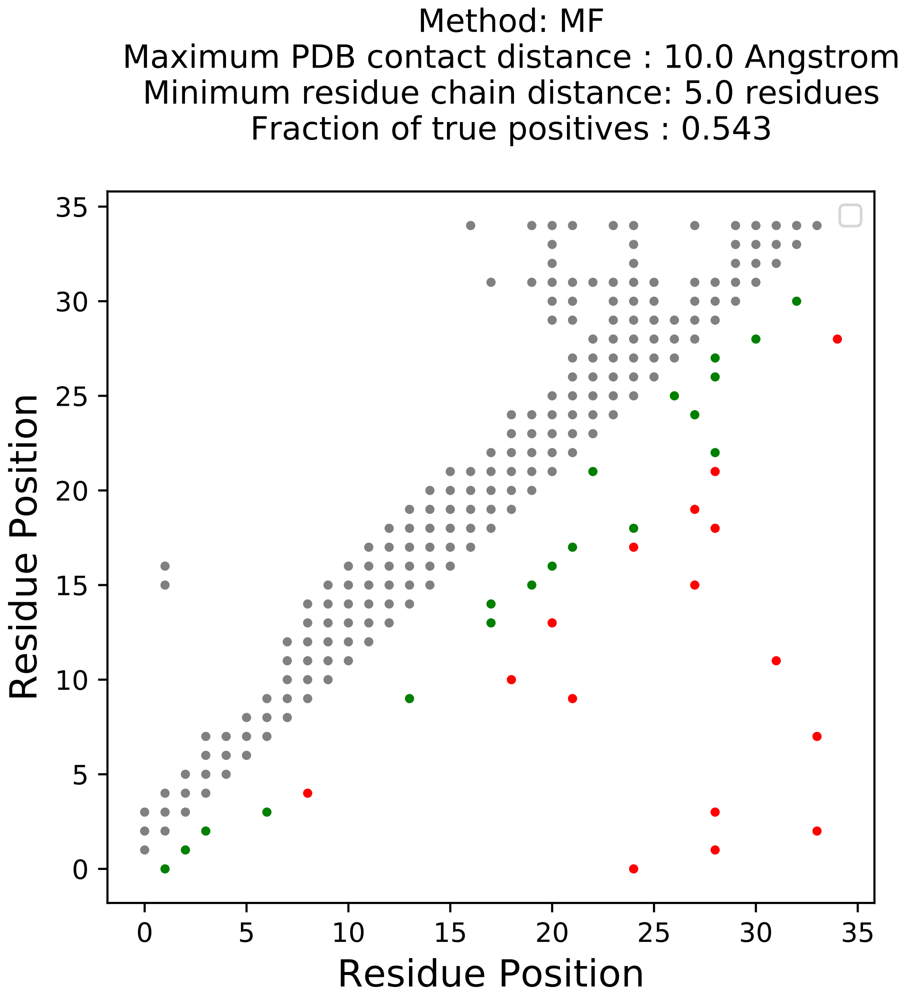
Method: MF
Maximum PDB contact distance : 10.0 Angstrom
Minimum residue chain distance: 5.0 residues
Fraction of true positives : 0.543
Residue Position
Residue Position
0
5
10
15
20
25
30
35

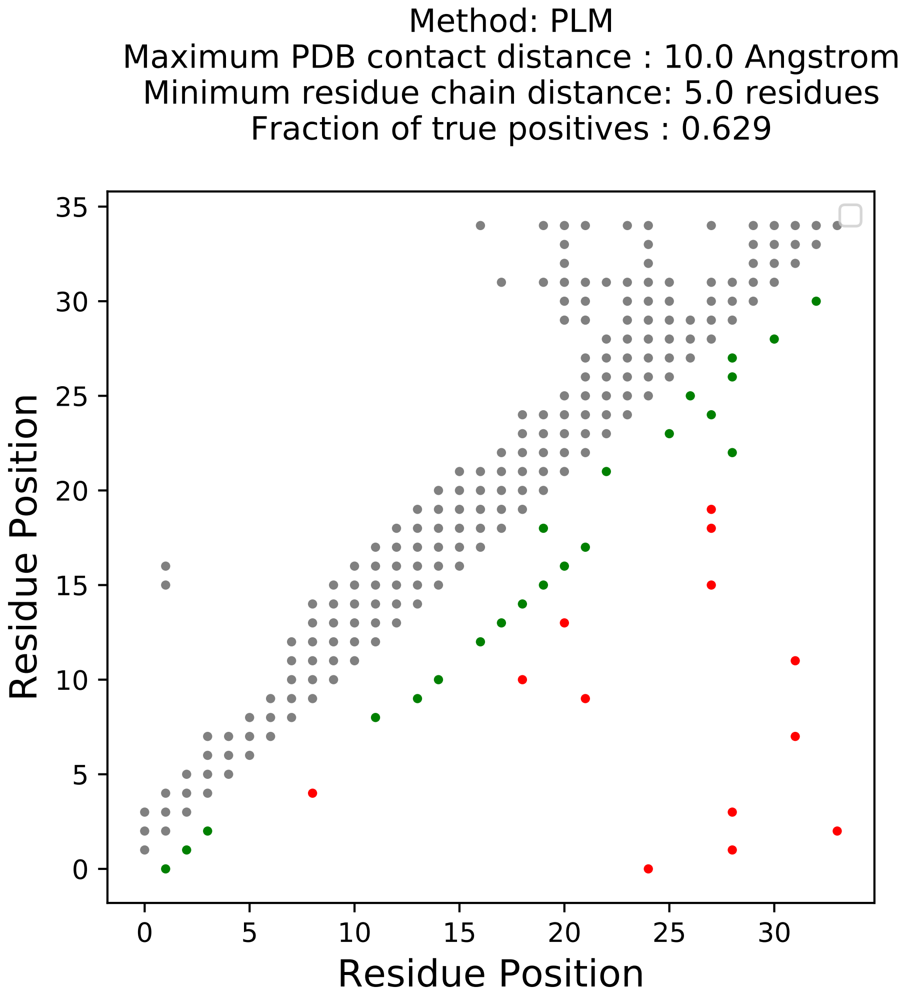
Method: PLM
Maximum PDB contact distance : 10.0 Angstrom
Minimum residue chain distance: 5.0 residues
Fraction of true positives : 0.629
Residue Position
Residue Position
0
5
10
15
20
25
30
35

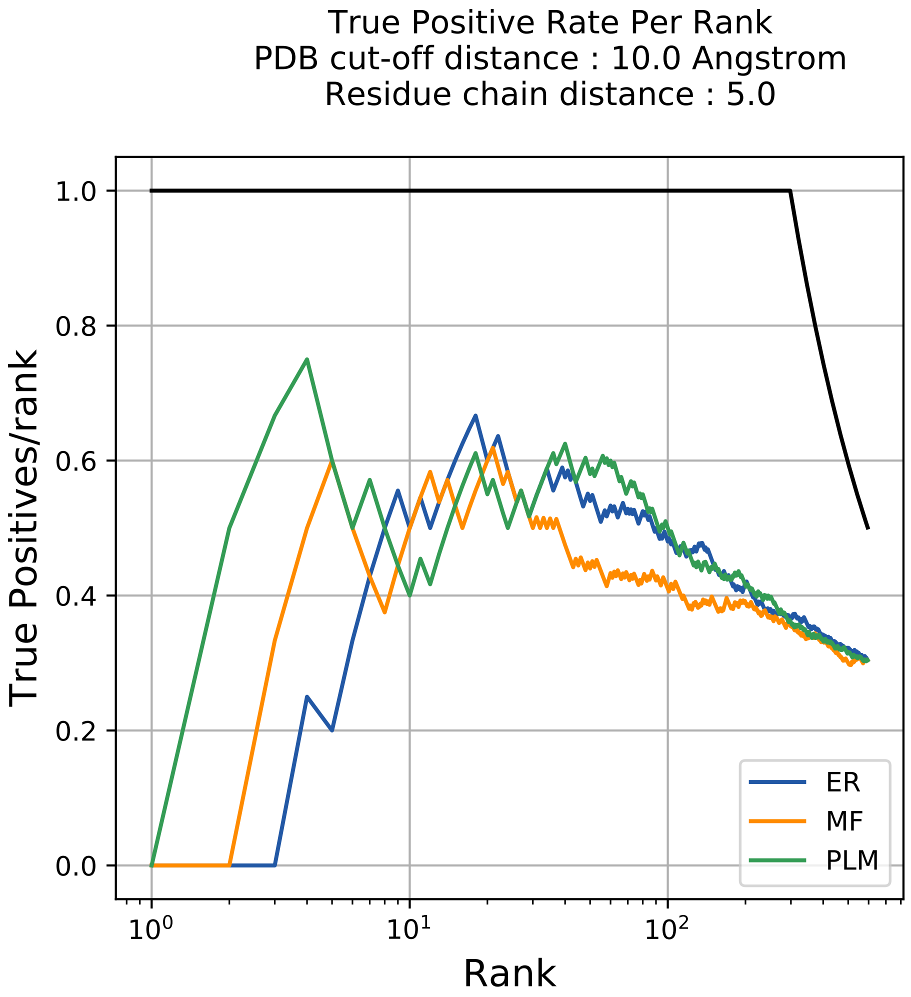

True Positive Rate Per Rank
PDB cut-off distance : 10.0 Angstrom
Residue chain distance : 5.0
True Positives/rank
Rank
1.0
0.8
0.6
0.4
0.2
0.0
$10^0$
$10^1$
$10^2$
ER
MF
PLM

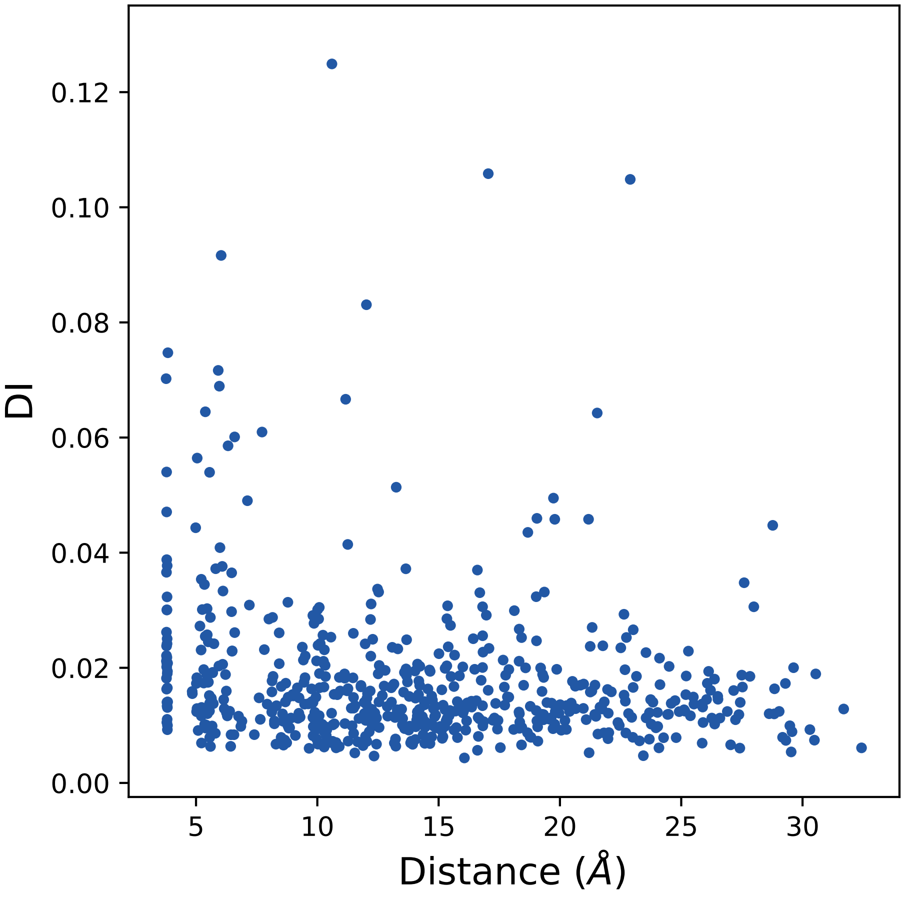

0.00
0.02
0.04
0.06
0.08
0.10
0.12
DI
5
10
15
20
25
30
Distance (Å)

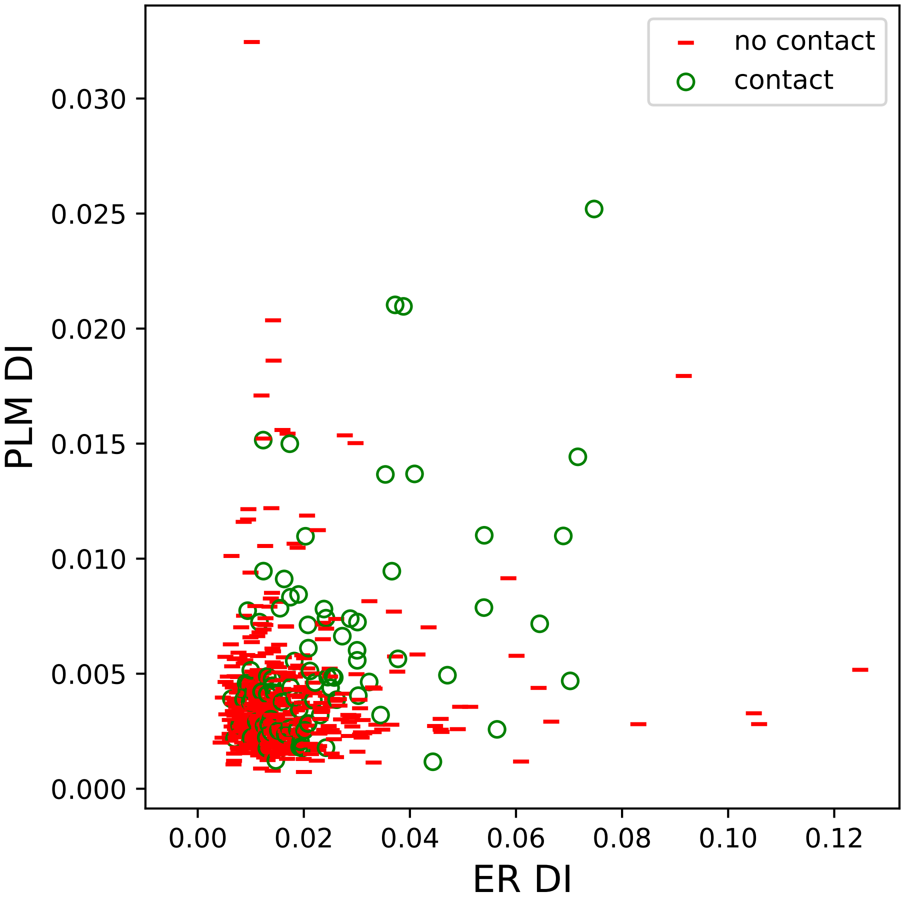
no contact
contact
0.030
0.025
0.020
0.015
0.010
0.005
0.000
PLM DI
0.00
0.02
0.04
0.06
0.08
0.10
0.12
ER DI

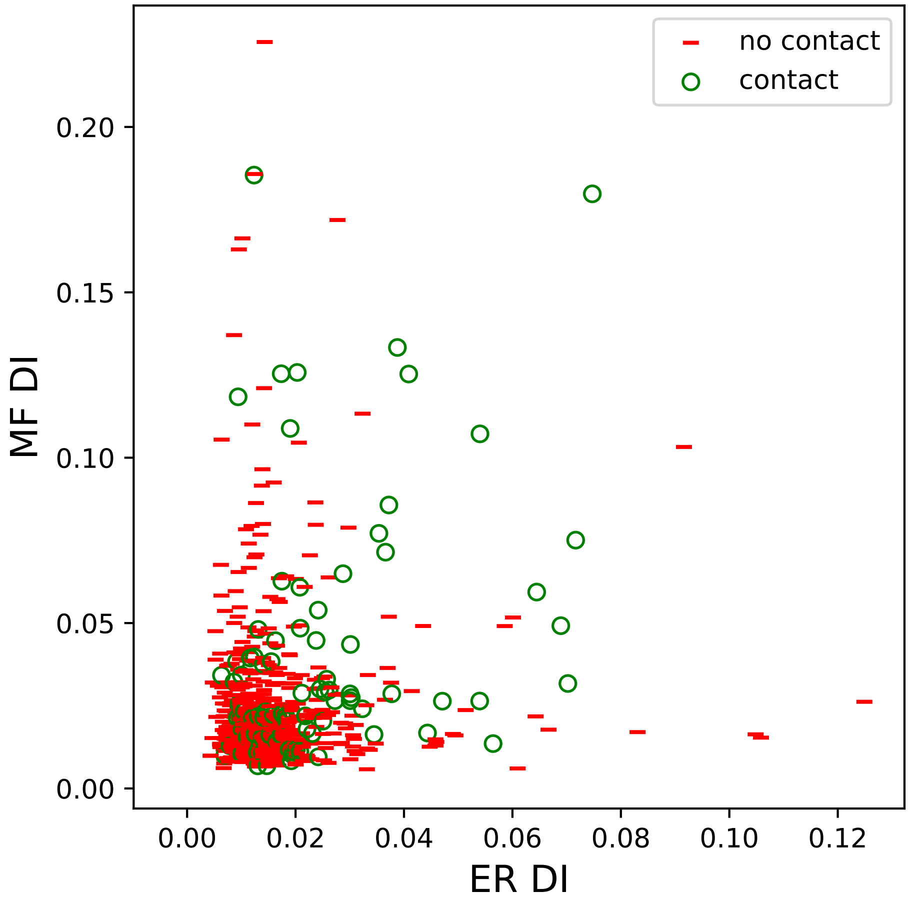

no contact
contact
0.00
0.05
0.10
0.15
0.20
MF DI
0.00
0.02
0.04
0.06
0.08
0.10
0.12
ER DI

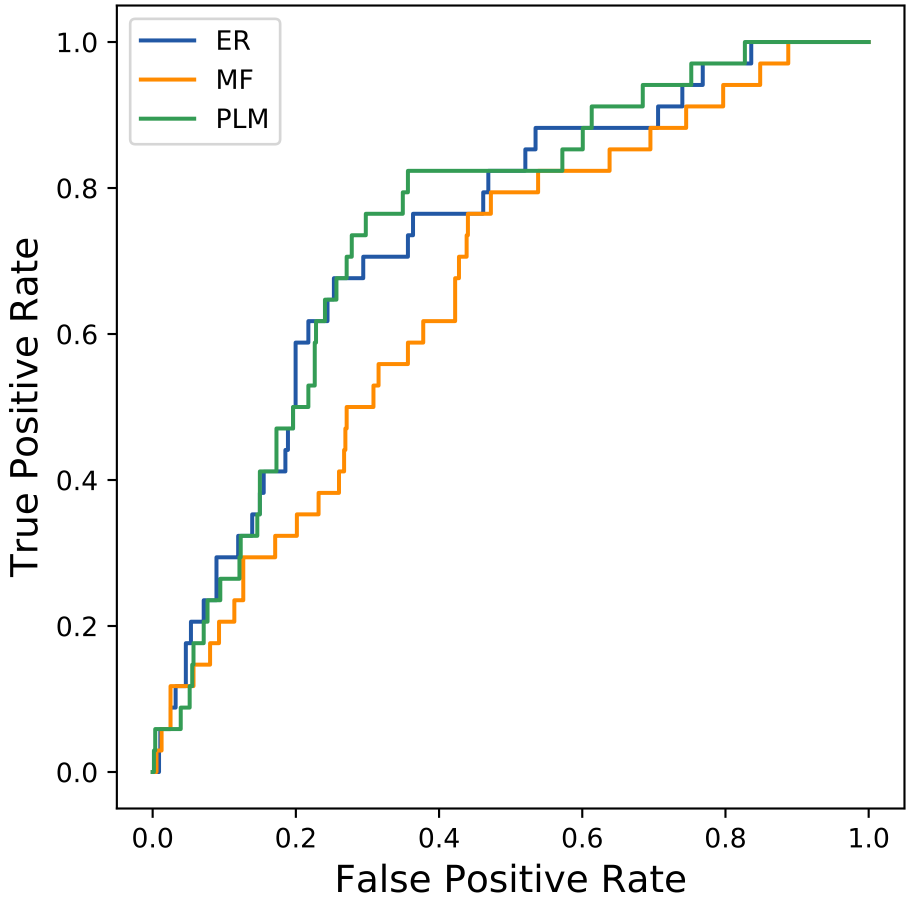
ER
MF
PLM
1.0
0.8
0.6
0.4
0.2
0.0
True Positive Rate
0.0
0.2
0.4
0.6
0.8
1.0
False Positive Rate
