## Supplemental Contact Maps for "Protein Structure Prediction with Expectation Reflection": 3933seq_206col_6iv2_PF01062_plots.pdf

Method: ER
Maximum PDB contact distance : 10.0 Angstrom
Minimum residue chain distance: 5.0 residues
Fraction of true positives : 0.553

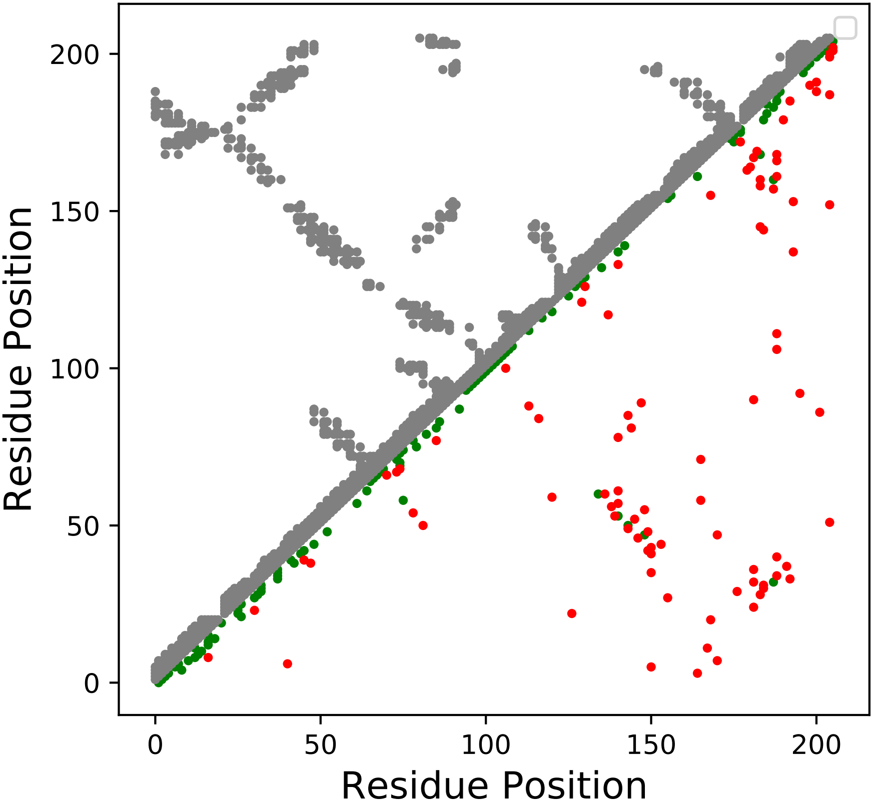

Method: MF
Maximum PDB contact distance : 10.0 Angstrom
Minimum residue chain distance: 5.0 residues
Fraction of true positives : 0.636

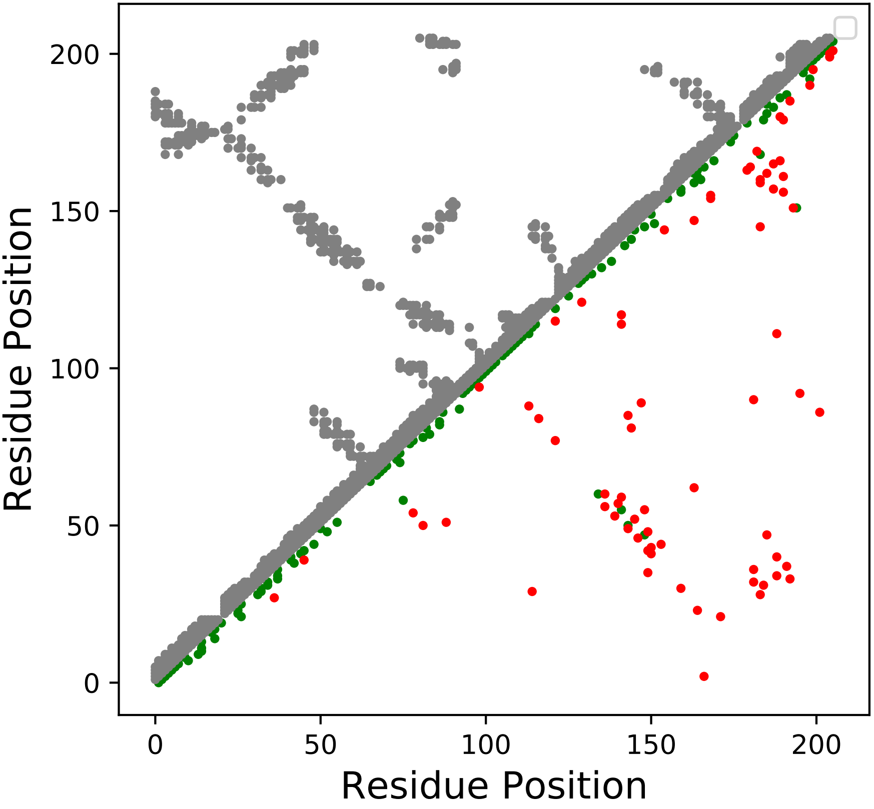

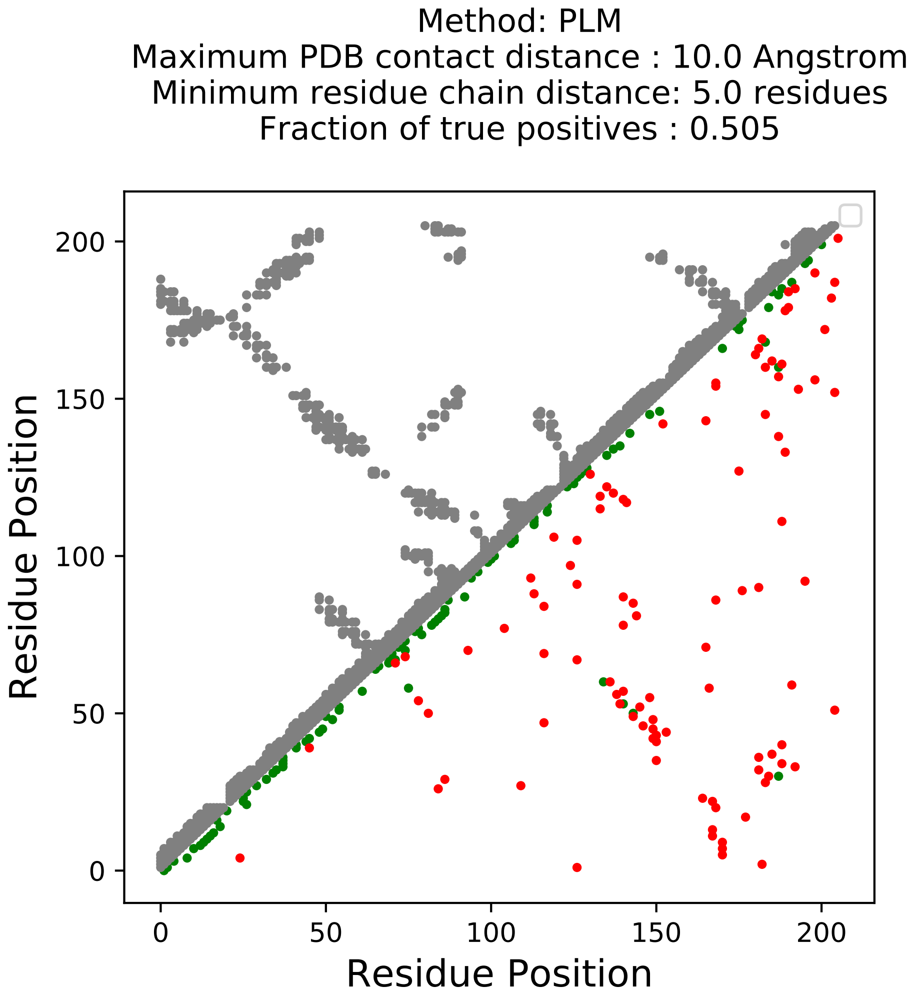
Method: PLM
Maximum PDB contact distance : 10.0 Angstrom
Minimum residue chain distance: 5.0 residues
Fraction of true positives : 0.505
Residue Position
Residue Position
0
50
100
150
200
0
50
100
150
200

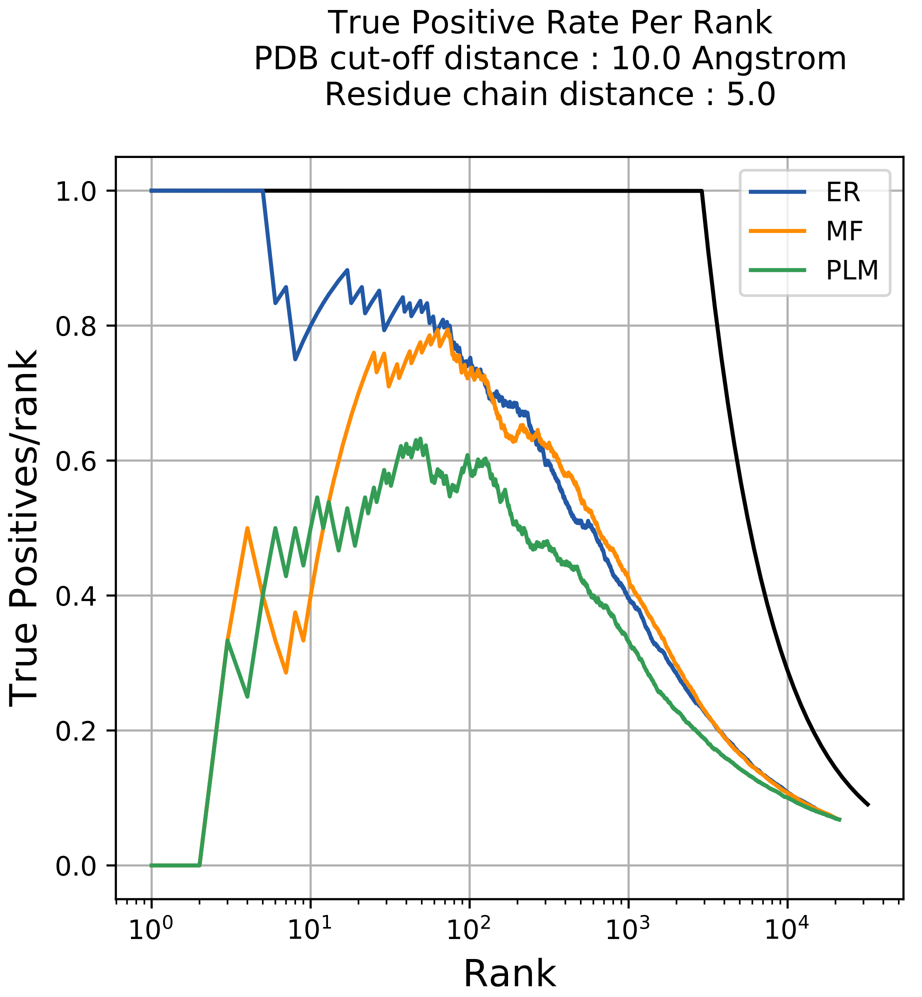
True Positive Rate Per Rank
PDB cut-off distance : 10.0 Angstrom
Residue chain distance : 5.0
True Positives/rank
Rank
ER
MF
PLM
1.0
0.8
0.6
0.4
0.2
0.0
$10^0$
$10^1$
$10^2$
$10^3$
$10^4$

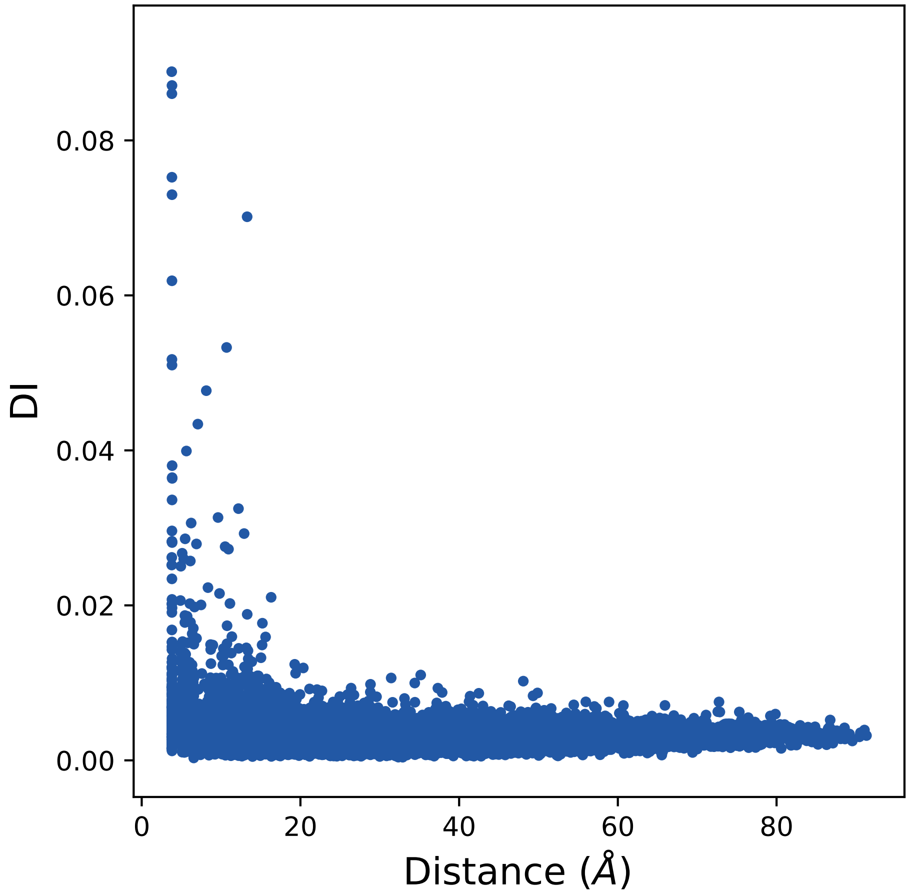
0.08
0.06
0.04
0.02
0.00
DI
0
20
40
60
80
Distance (Å)

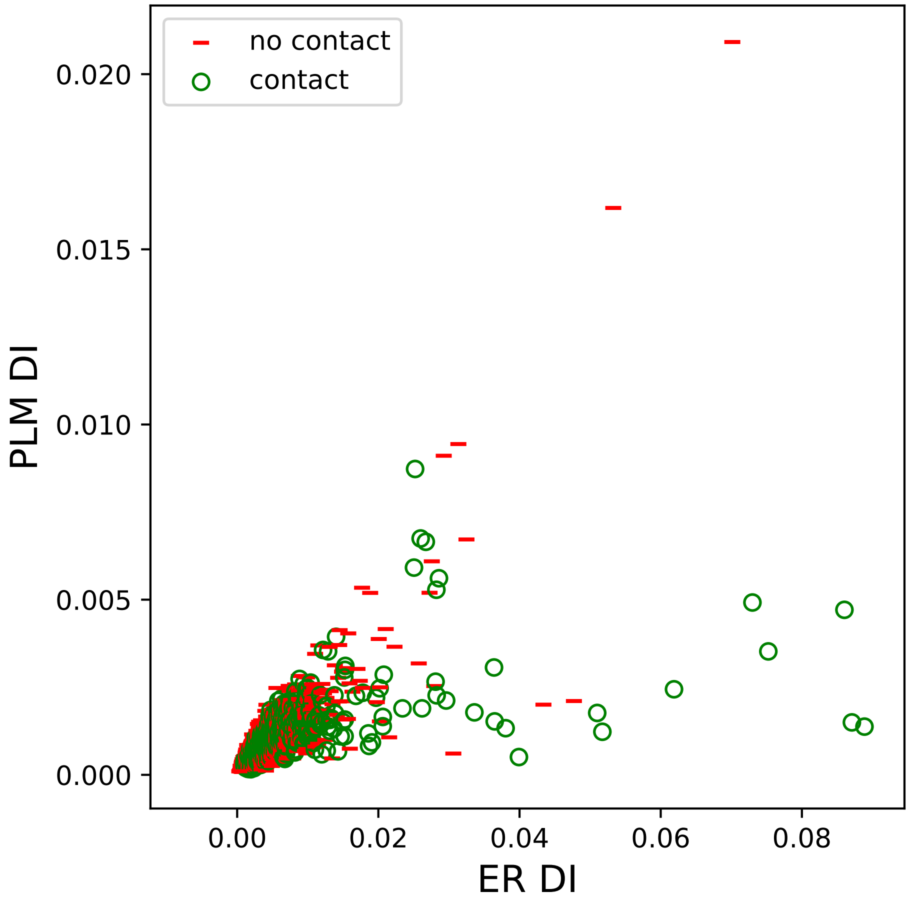

no contact
contact
0.020
0.015
0.010
0.005
0.000
PLM DI
0.00
0.02
0.04
0.06
0.08
ER DI

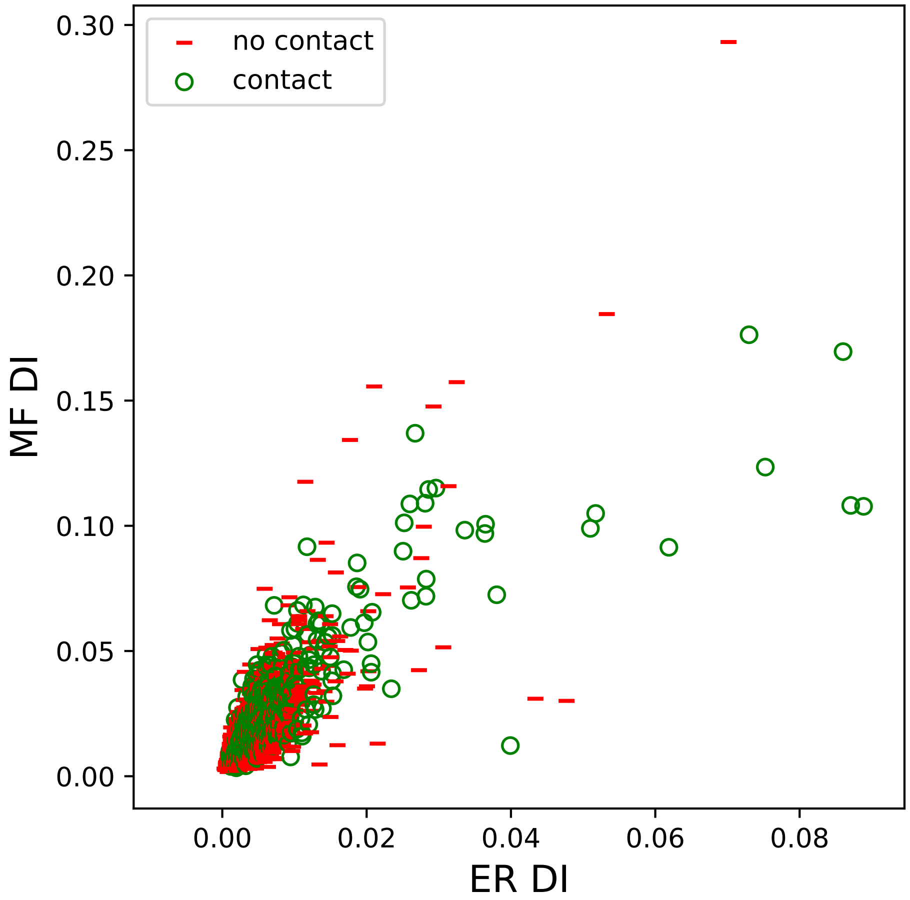

no contact
contact
MF DI
ER DI
0.00
0.05
0.10
0.15
0.20
0.25
0.30
0.00
0.02
0.04
0.06
0.08

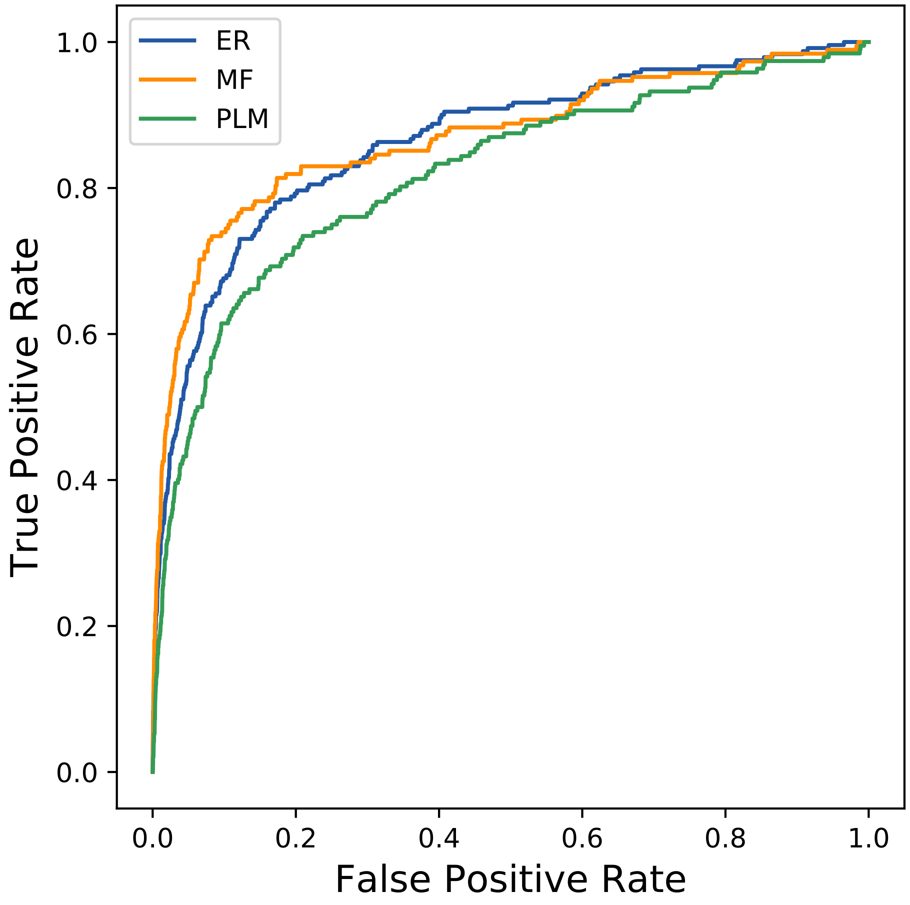
ER
MF
PLM
1.0
0.8
0.6
0.4
0.2
0.0
True Positive Rate
0.0
0.2
0.4
0.6
0.8
1.0
False Positive Rate
