## Supplemental Contact Maps for "Protein Structure Prediction with Expectation Reflection": 4413seq_62col_4ppd_PF00936_plots.pdf

Method: ER
Maximum PDB contact distance : 10.0 Angstrom
Minimum residue chain distance: 5.0 residues
Fraction of true positives : 0.645

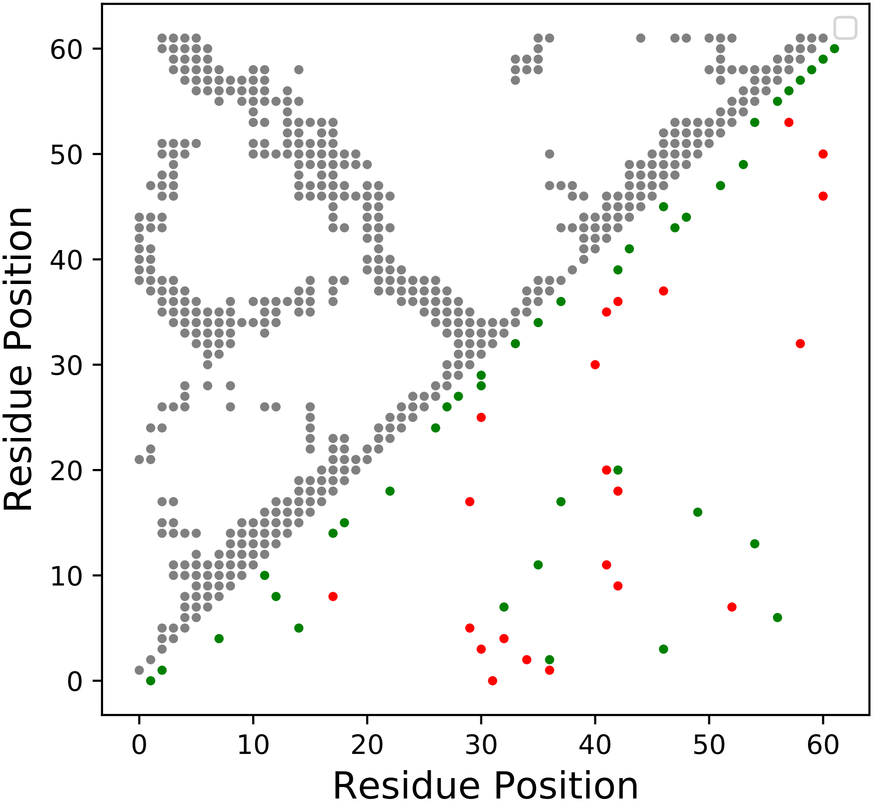

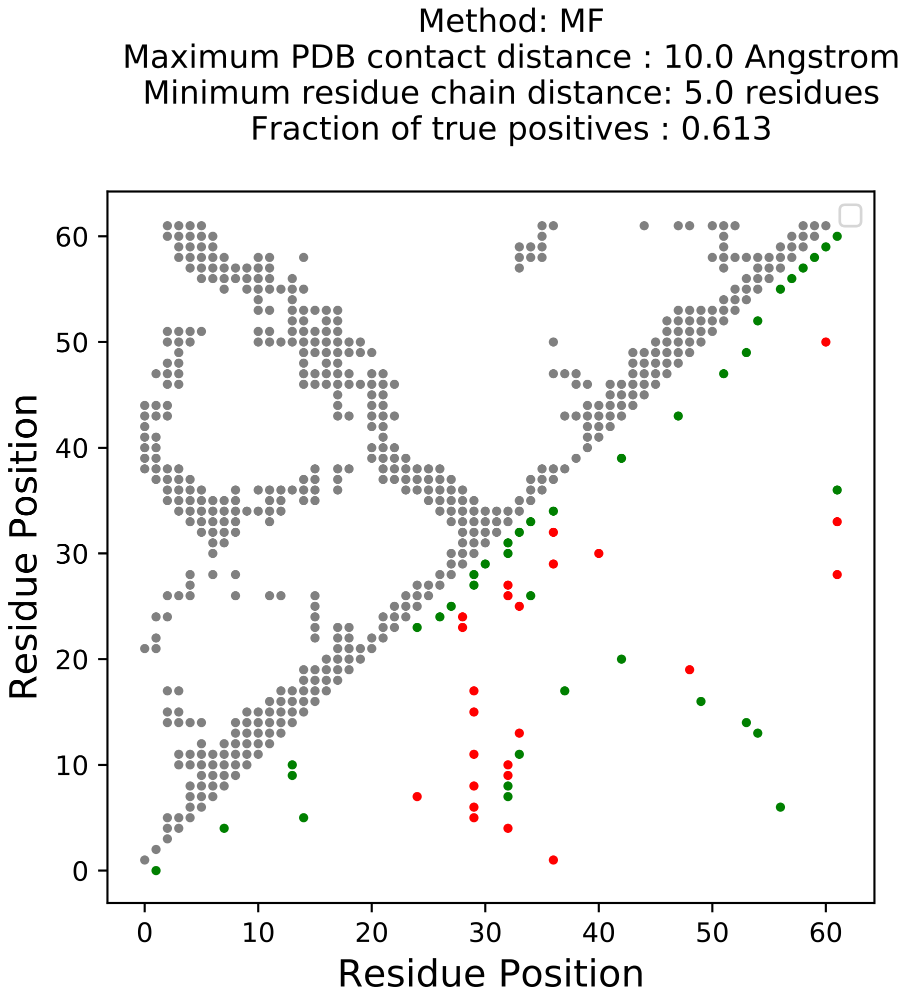

Method: MF
Maximum PDB contact distance : 10.0 Angstrom
Minimum residue chain distance: 5.0 residues
Fraction of true positives : 0.613
Residue Position
Residue Position
0
10
20
30
40
50
60

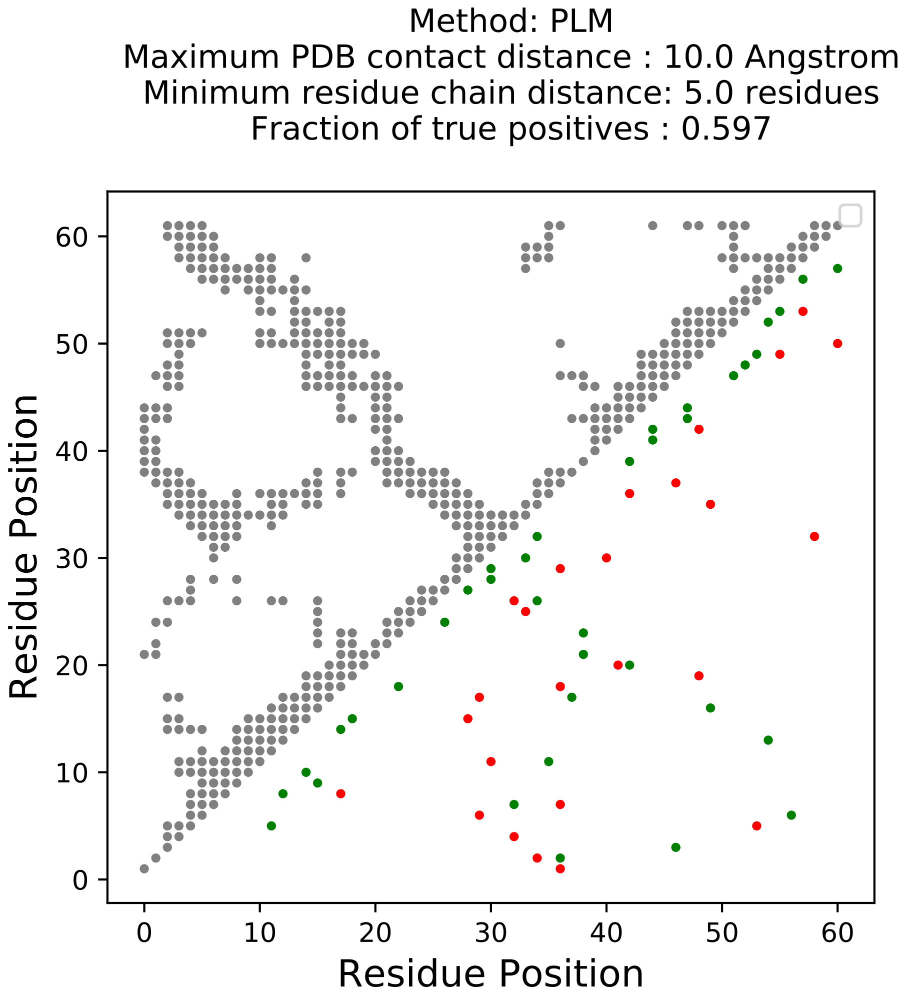
Method: PLM
Maximum PDB contact distance : 10.0 Angstrom
Minimum residue chain distance: 5.0 residues
Fraction of true positives : 0.597
Residue Position
Residue Position
0
10
20
30
40
50
60

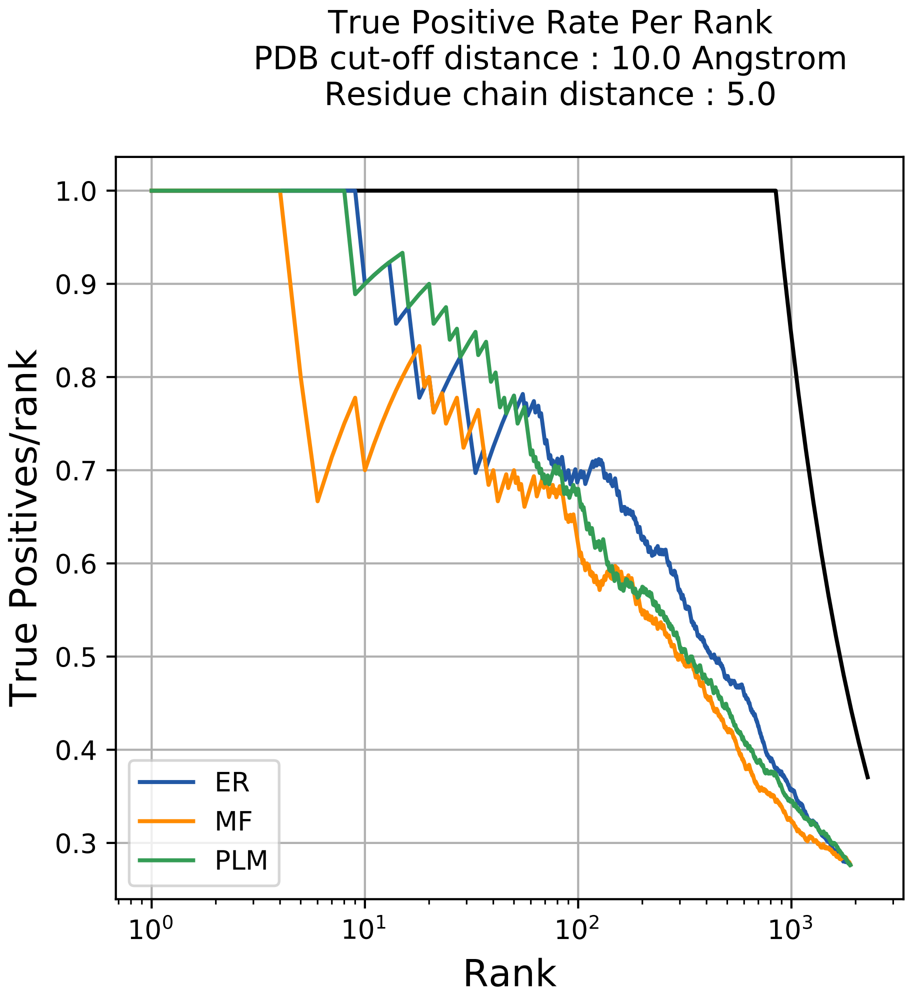
True Positive Rate Per Rank
PDB cut-off distance : 10.0 Angstrom
Residue chain distance : 5.0
True Positives/rank
Rank
1.0
0.9
0.8
0.7
0.6
0.5
0.4
0.3
$10^0$
$10^1$
$10^2$
$10^3$
ER
MF
PLM

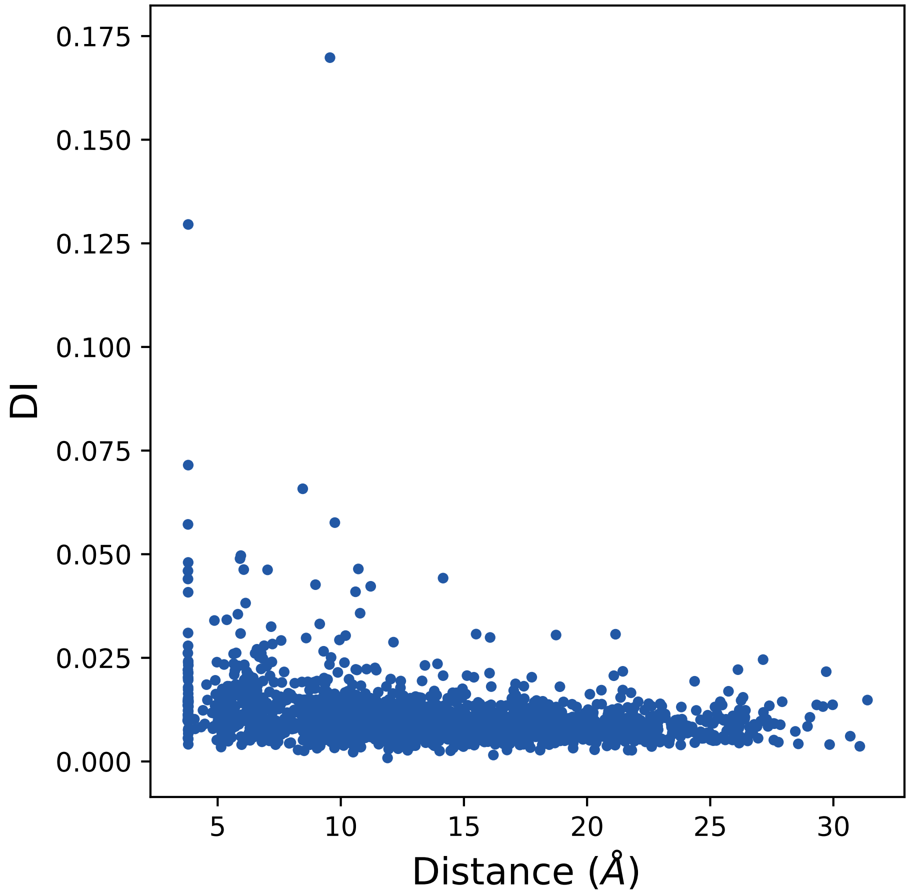
0.175
0.150
0.125
0.100
0.075
0.050
0.025
0.000
DI
5
10
15
20
25
30
Distance (Å)

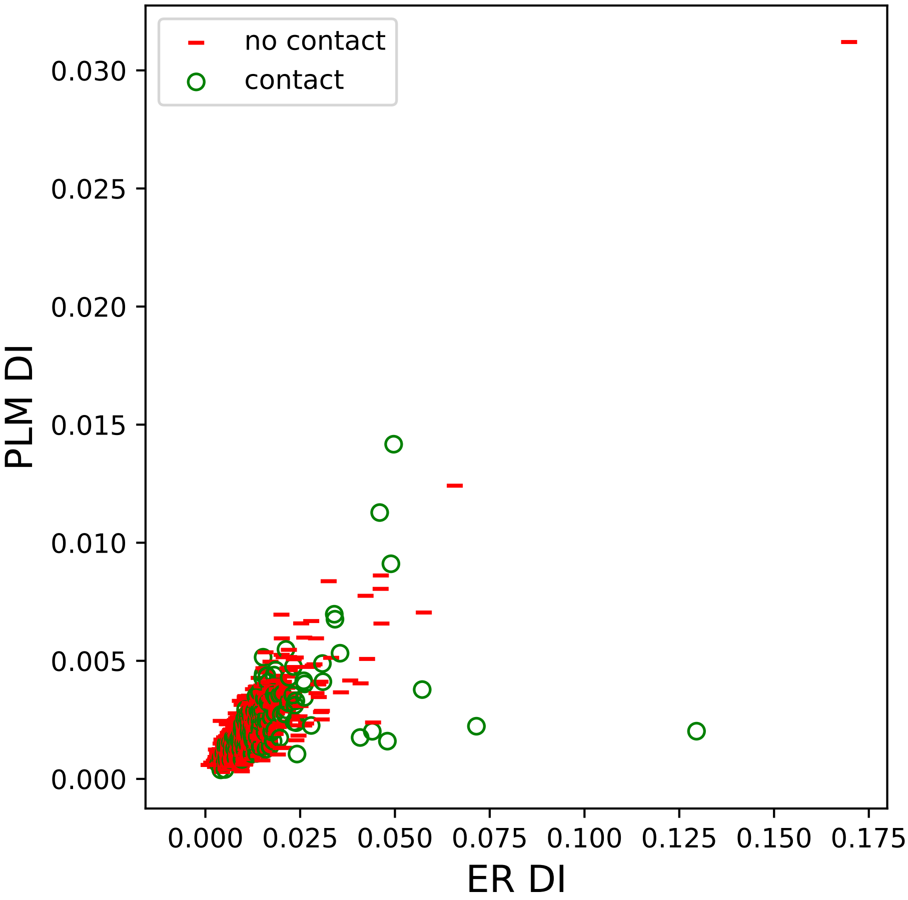
no contact
contact
0.030
0.025
0.020
0.015
0.010
0.005
0.000
PLM DI
0.000
0.025
0.050
0.075
0.100
0.125
0.150
0.175
ER DI

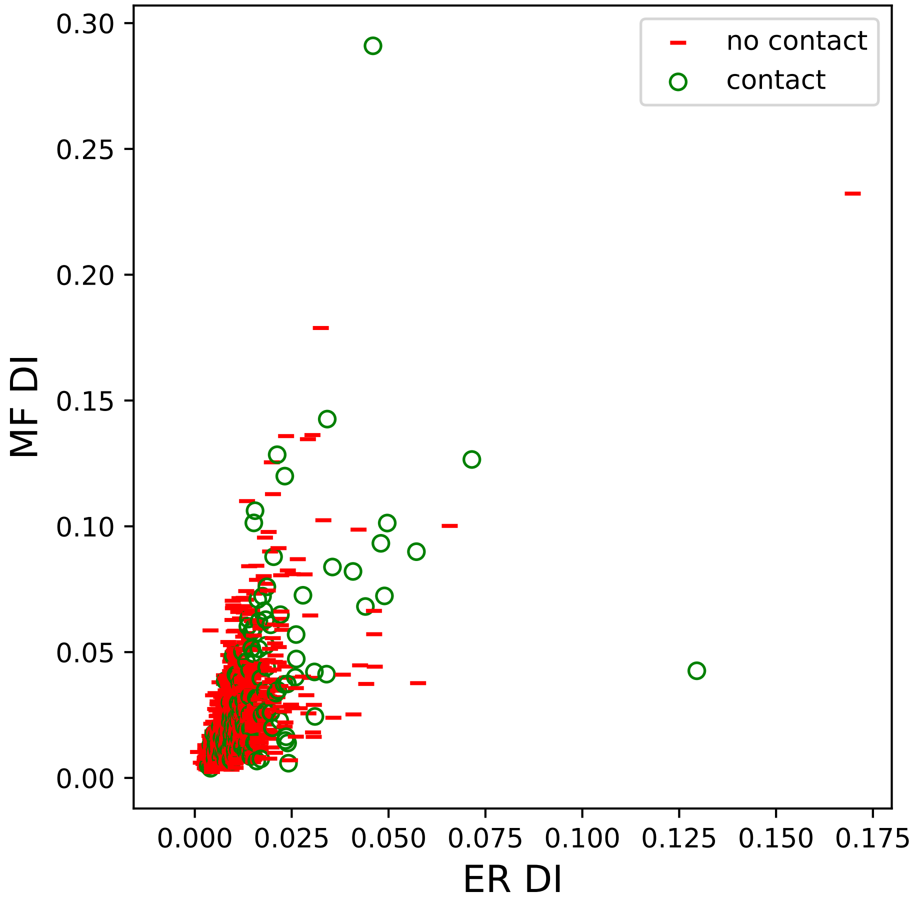

no contact
contact
MF DI
ER DI
0.00
0.05
0.10
0.15
0.20
0.25
0.30
0.000
0.025
0.050
0.075
0.100
0.125
0.150
0.175

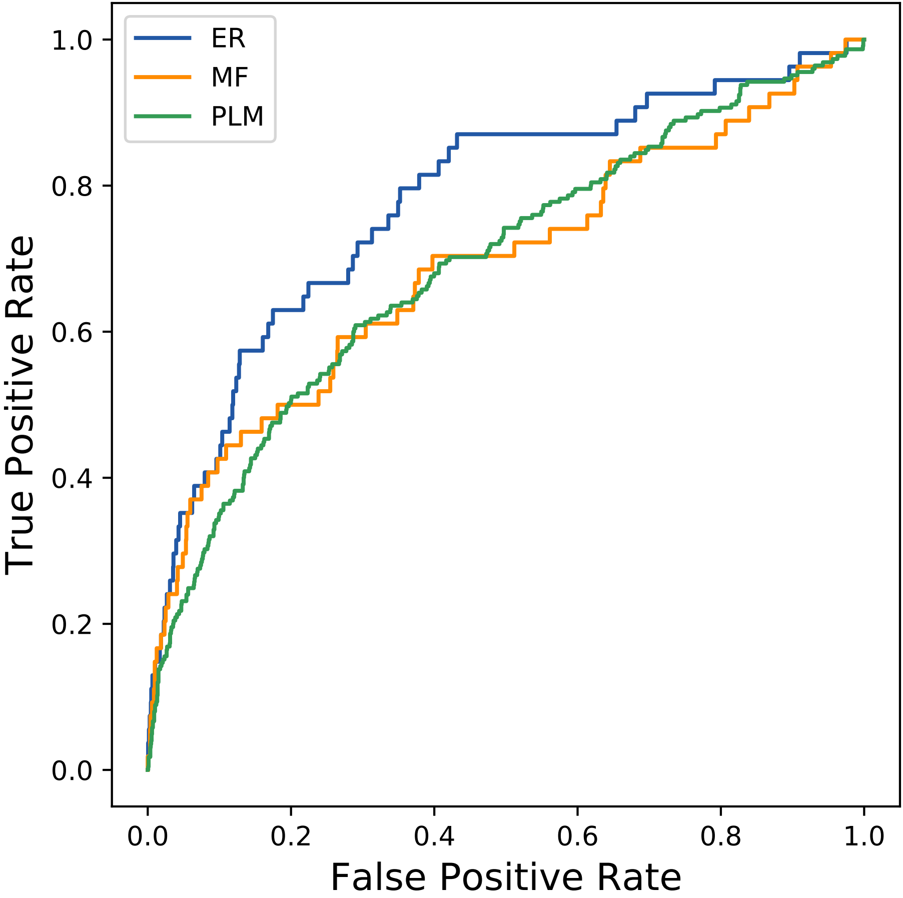

ER
MF
PLM
1.0
0.8
0.6
0.4
0.2
0.0
True Positive Rate
0.0
0.2
0.4
0.6
0.8
1.0
False Positive Rate
