## Supplemental Contact Maps for "Protein Structure Prediction with Expectation Reflection": 4458seq_63col_1etz_PF07654_plots.pdf

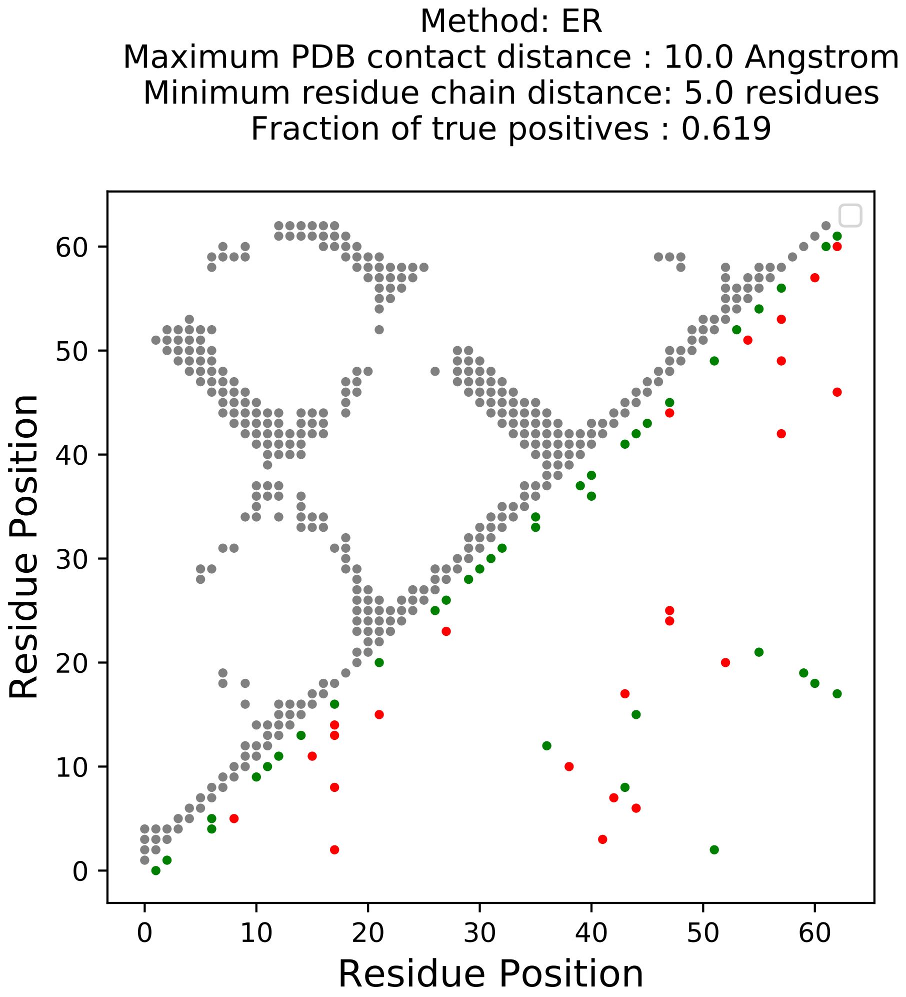
Method: ER
Maximum PDB contact distance : 10.0 Angstrom
Minimum residue chain distance: 5.0 residues
Fraction of true positives : 0.619
Residue Position
Residue Position

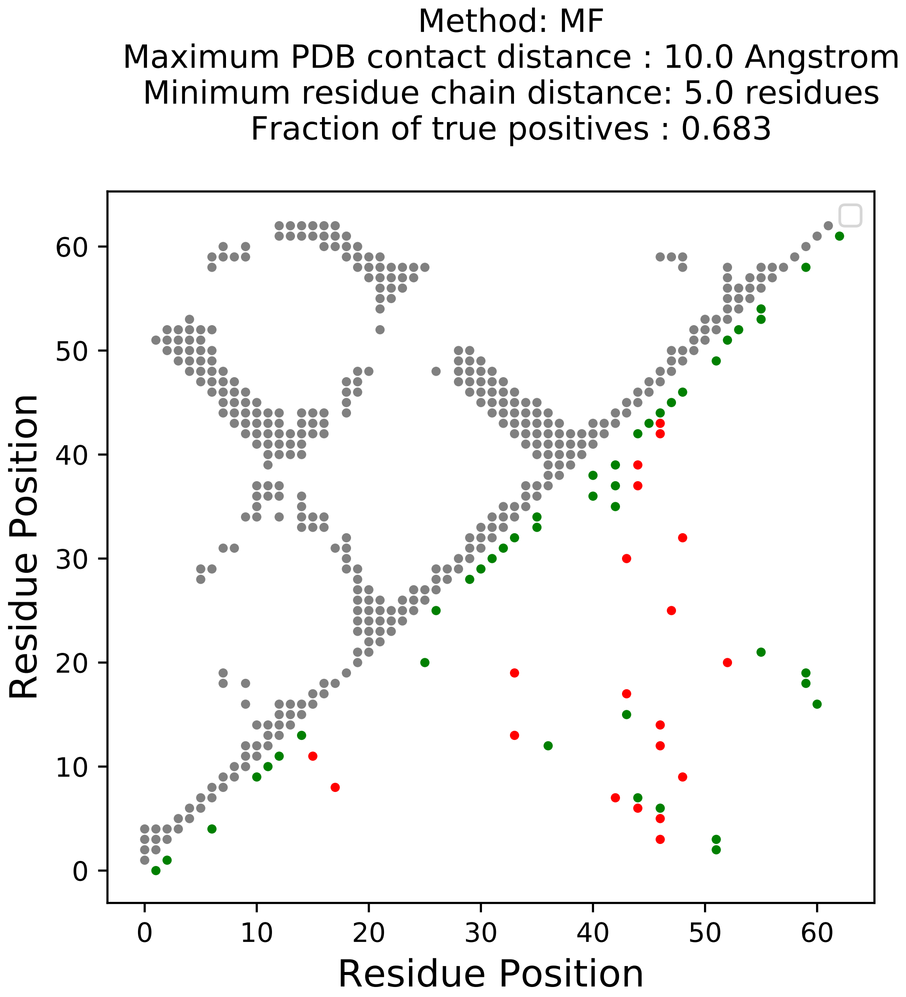

Method: MF
Maximum PDB contact distance : 10.0 Angstrom
Minimum residue chain distance: 5.0 residues
Fraction of true positives : 0.683
Residue Position
Residue Position
0
10
20
30
40
50
60

Method: PLM
Maximum PDB contact distance : 10.0 Angstrom
Minimum residue chain distance: 5.0 residues
Fraction of true positives : 0.651

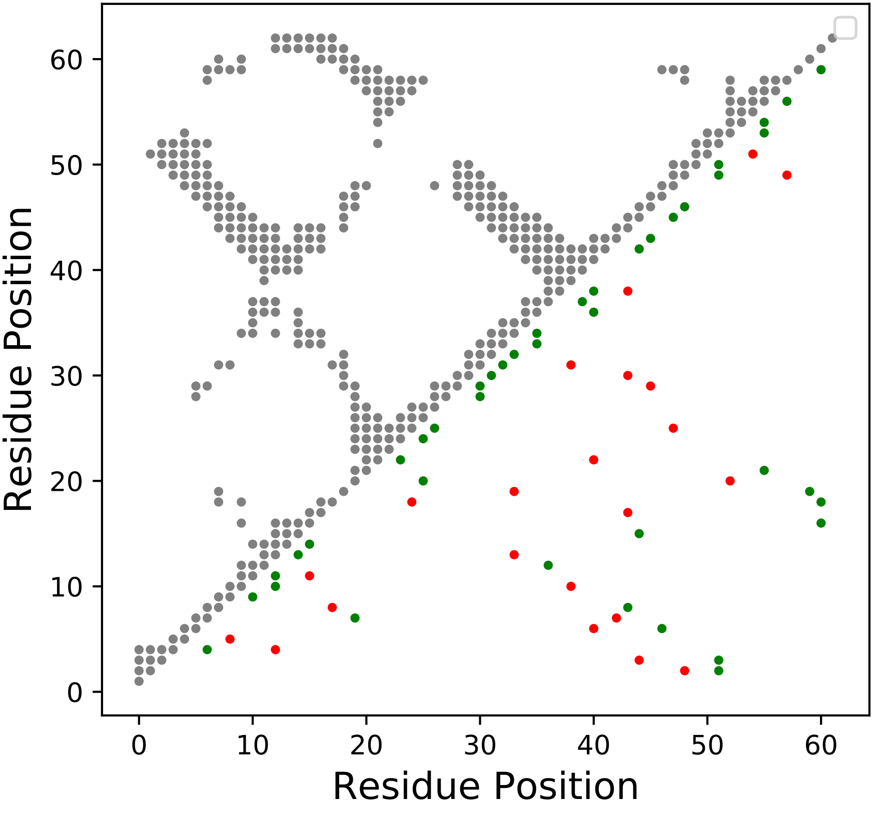

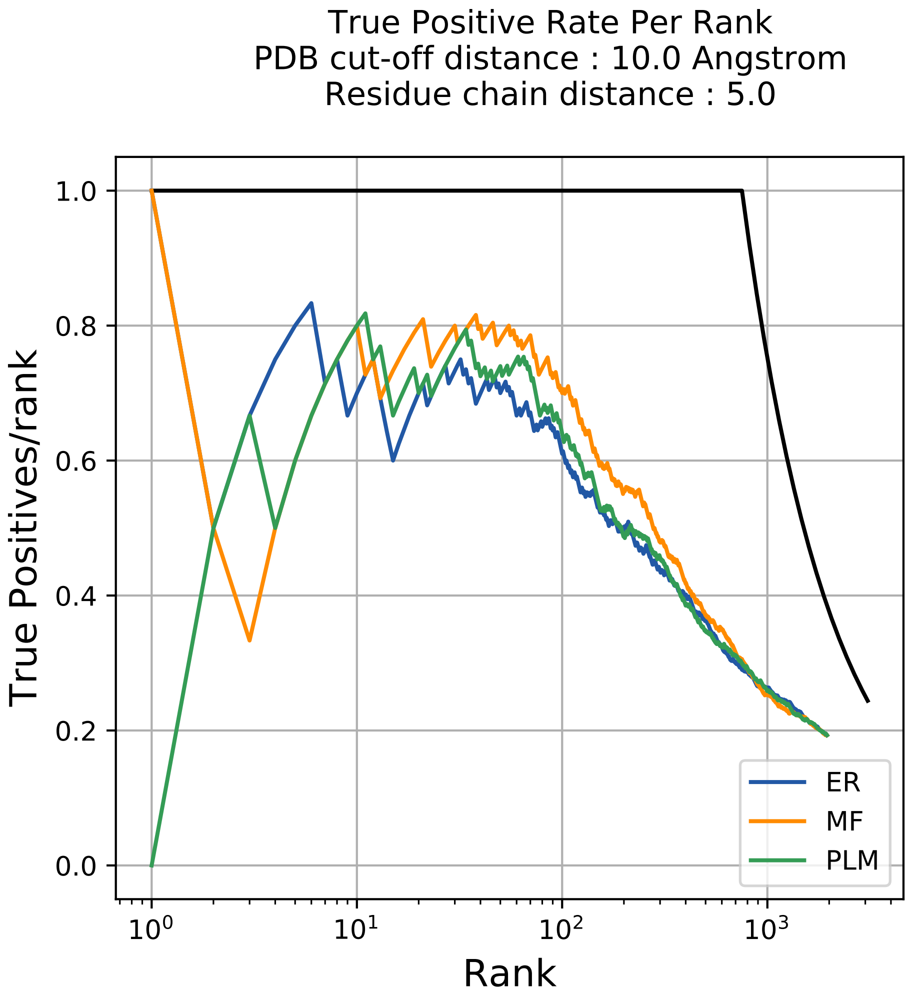

True Positive Rate Per Rank
PDB cut-off distance : 10.0 Angstrom
Residue chain distance : 5.0
True Positives/rank
1.0
0.8
0.6
0.4
0.2
0.0
$10^0$
$10^1$
$10^2$
$10^3$
Rank
ER
MF
PLM

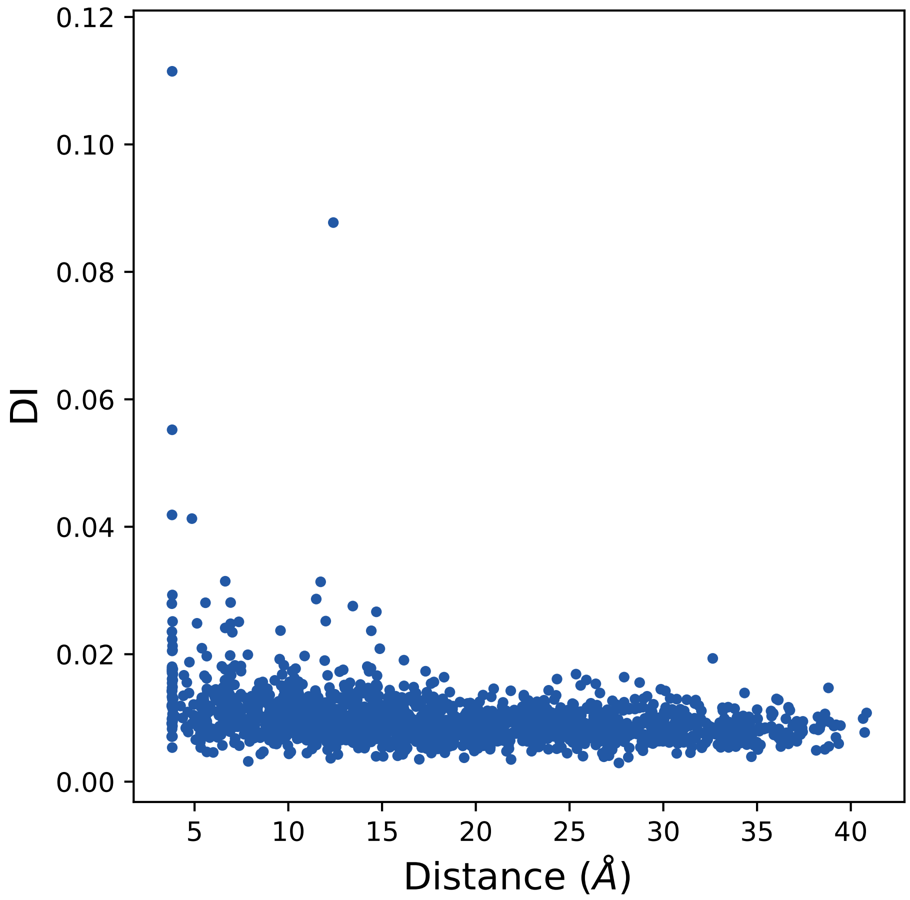

0.12
0.10
0.08
0.06
0.04
0.02
0.00
DI
5
10
15
20
25
30
35
40
Distance (Å)

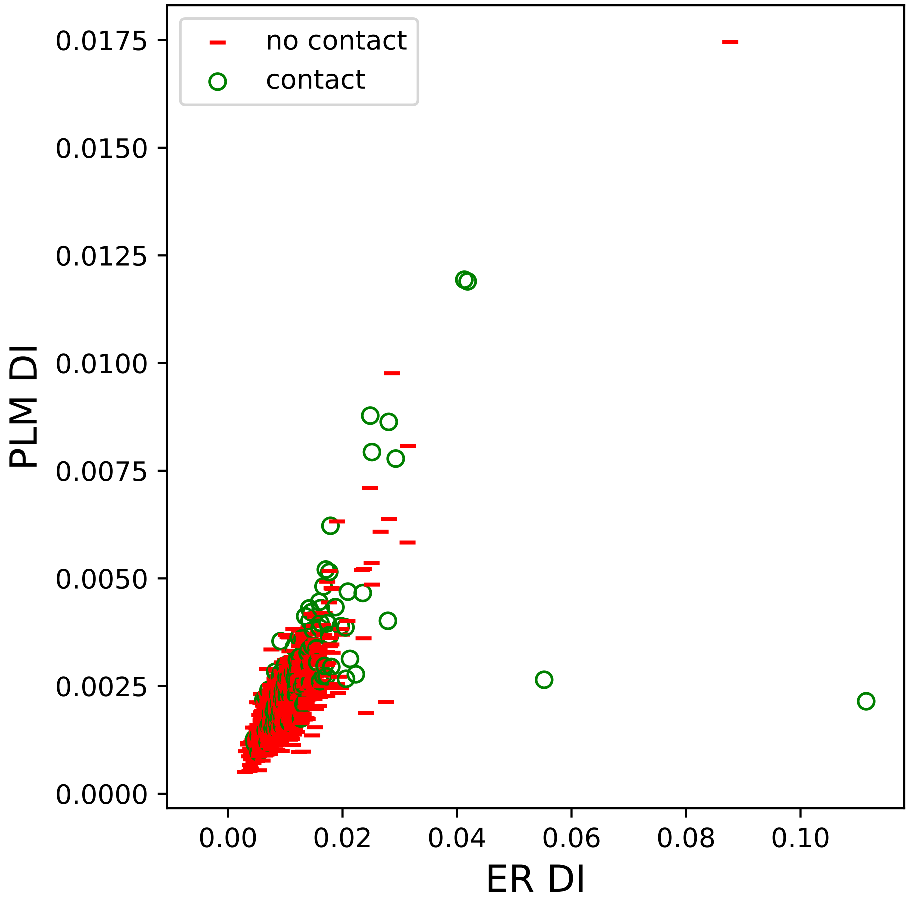
no contact
contact
PLM DI
ER DI
0.0000
0.0025
0.0050
0.0075
0.0100
0.0125
0.0150
0.0175
0.00
0.02
0.04
0.06
0.08
0.10

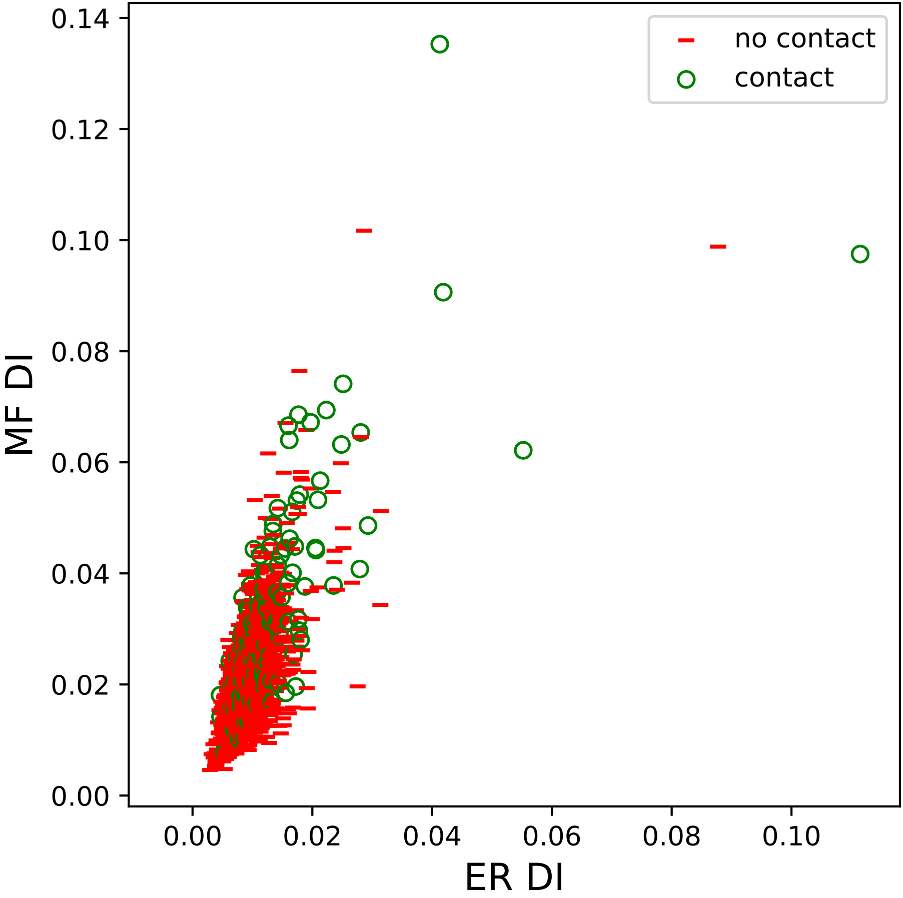

0.14
0.12
0.10
0.08
0.06
0.04
0.02
0.00
MF DI
0.00
0.02
0.04
0.06
0.08
0.10
ER DI
no contact
contact

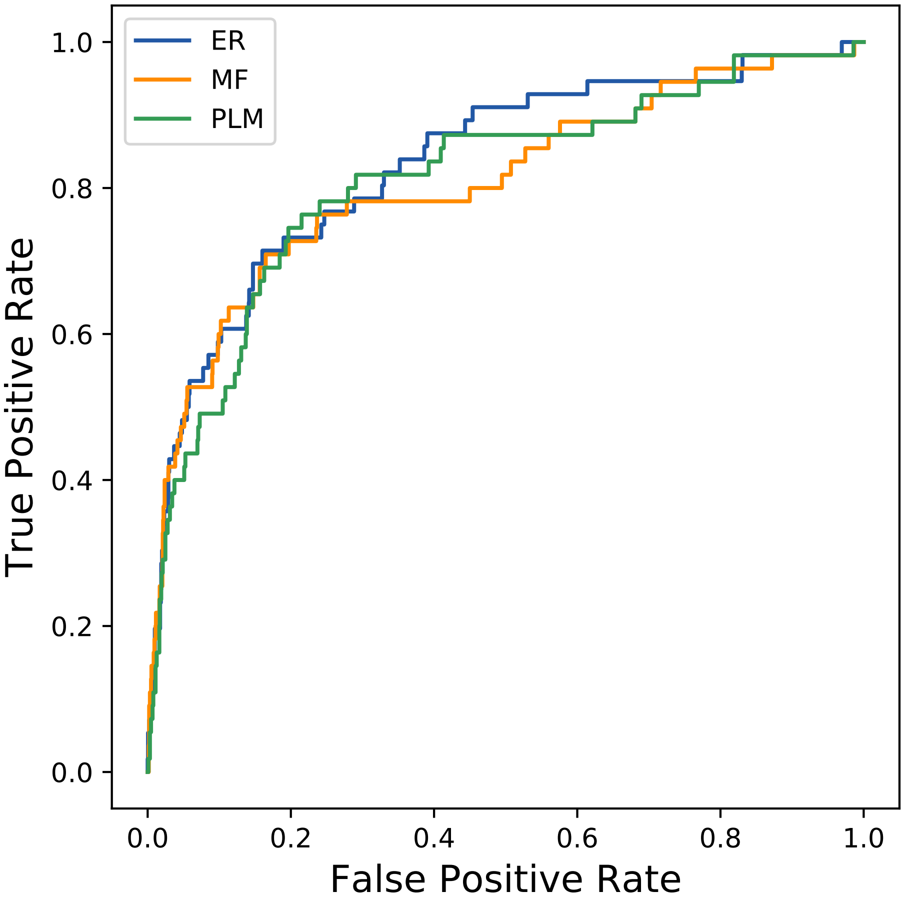

ER
MF
PLM
True Positive Rate
False Positive Rate
0.0
0.2
0.4
0.6
0.8
1.0
