## Supplemental Contact Maps for "Protein Structure Prediction with Expectation Reflection": 4765seq_180col_4bcw_PF00194_plots.pdf

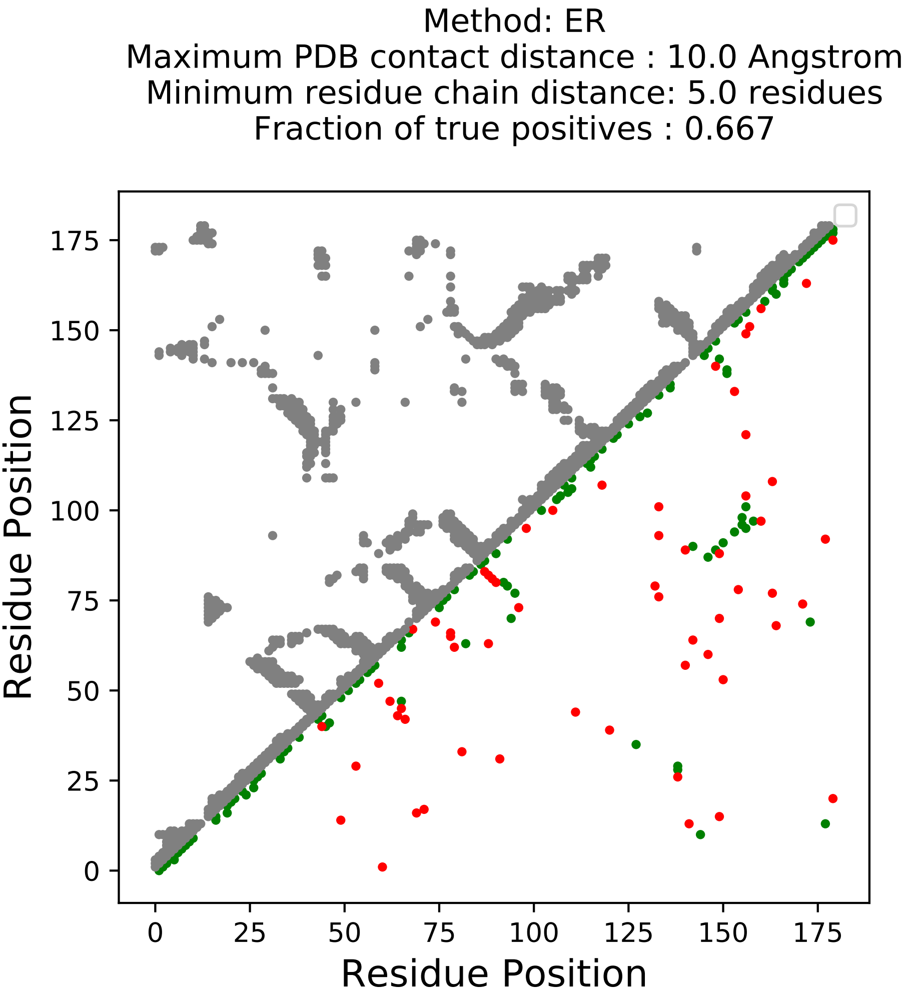
Method: ER
Maximum PDB contact distance : 10.0 Angstrom
Minimum residue chain distance: 5.0 residues
Fraction of true positives : 0.667
Residue Position
Residue Position
0
25
50
75
100
125
150
175

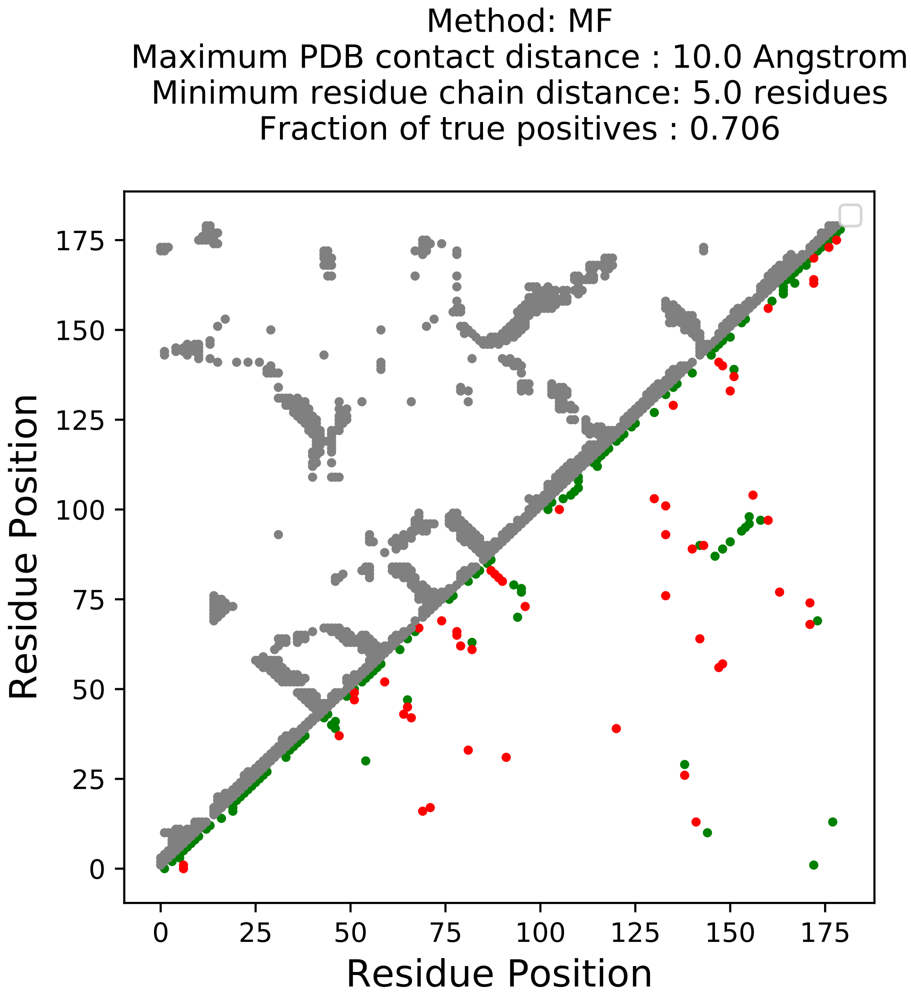
Method: MF
Maximum PDB contact distance : 10.0 Angstrom
Minimum residue chain distance: 5.0 residues
Fraction of true positives : 0.706
Residue Position
Residue Position
0
25
50
75
100
125
150
175

Method: PLM
Maximum PDB contact distance : 10.0 Angstrom
Minimum residue chain distance: 5.0 residues
Fraction of true positives : 0.611

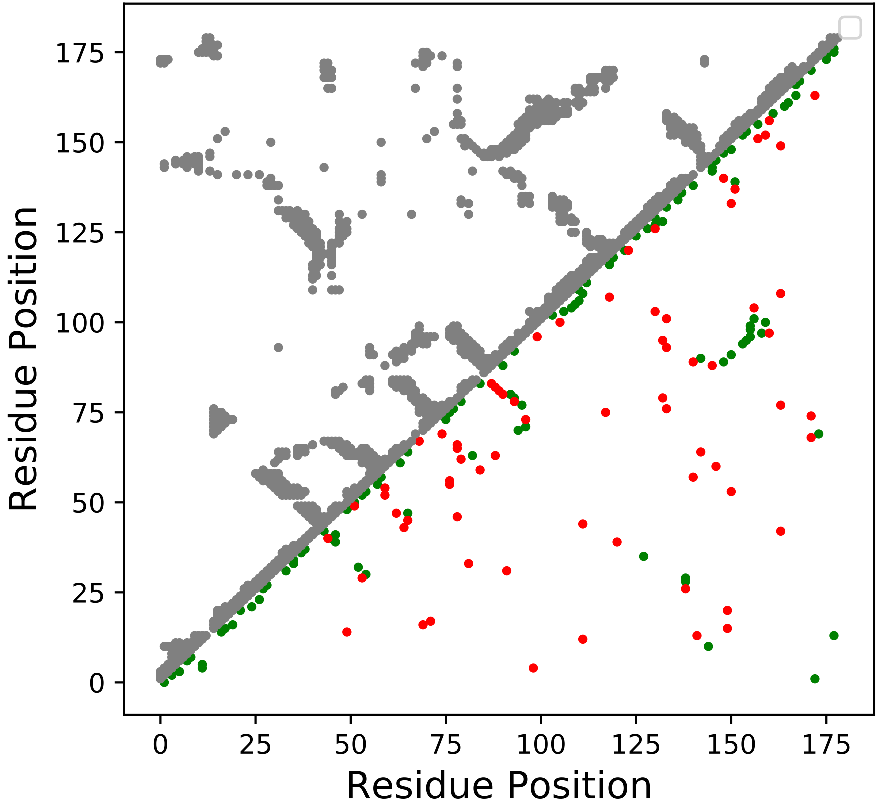

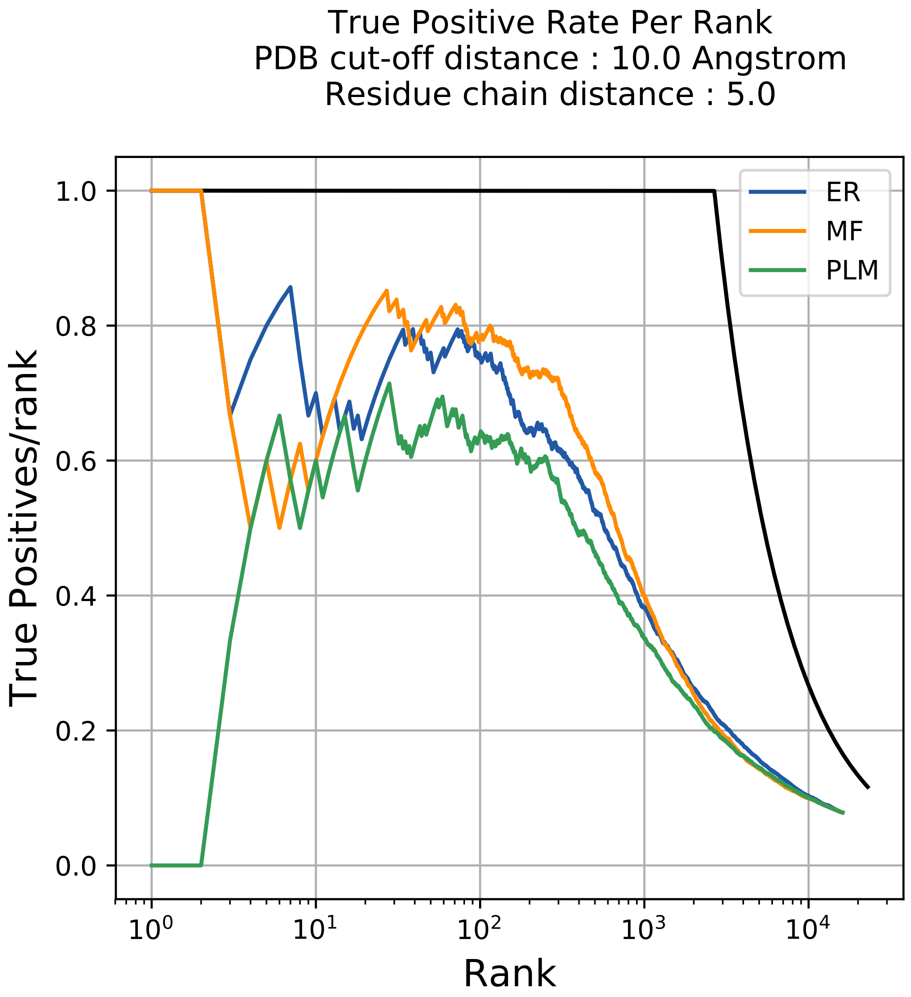
True Positive Rate Per Rank
PDB cut-off distance : 10.0 Angstrom
Residue chain distance : 5.0
True Positives/rank
Rank
1.0
0.8
0.6
0.4
0.2
0.0
$10^0$
$10^1$
$10^2$
$10^3$
$10^4$
ER
MF
PLM

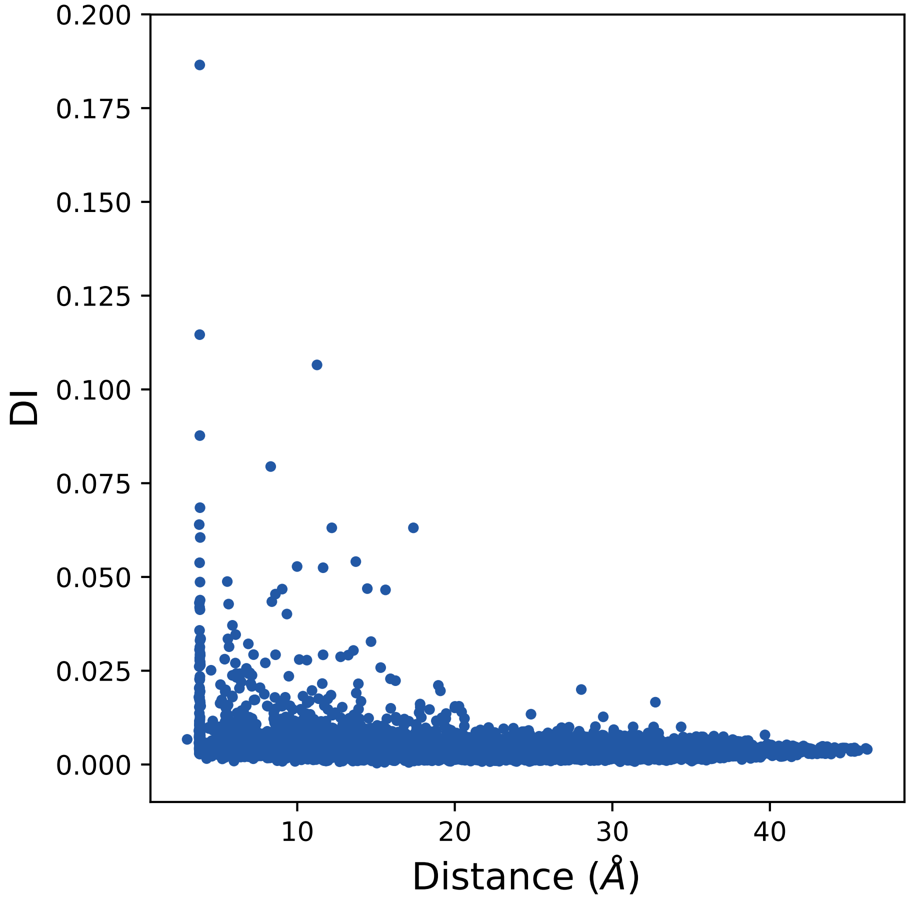
0.200
0.175
0.150
0.125
0.100
0.075
0.050
0.025
0.000
DI
10
20
30
40
Distance (Å)

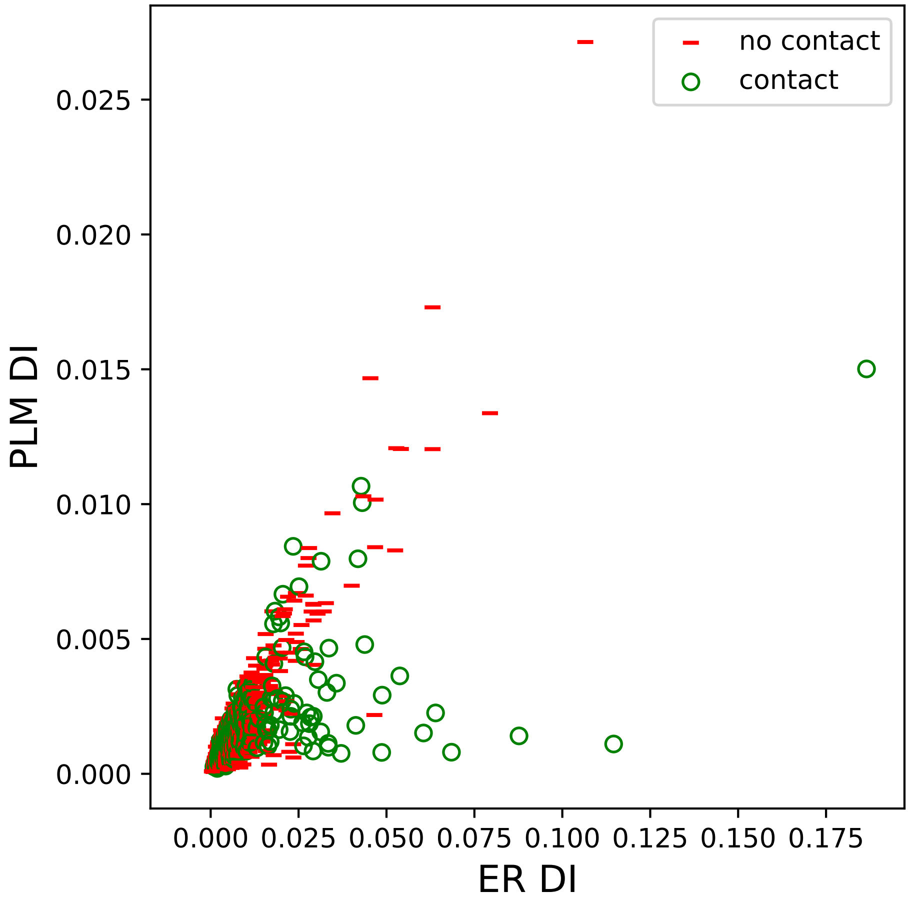
no contact
contact
PLM DI
ER DI
0.000
0.005
0.010
0.015
0.020
0.025
0.000
0.025
0.050
0.075
0.100
0.125
0.150
0.175

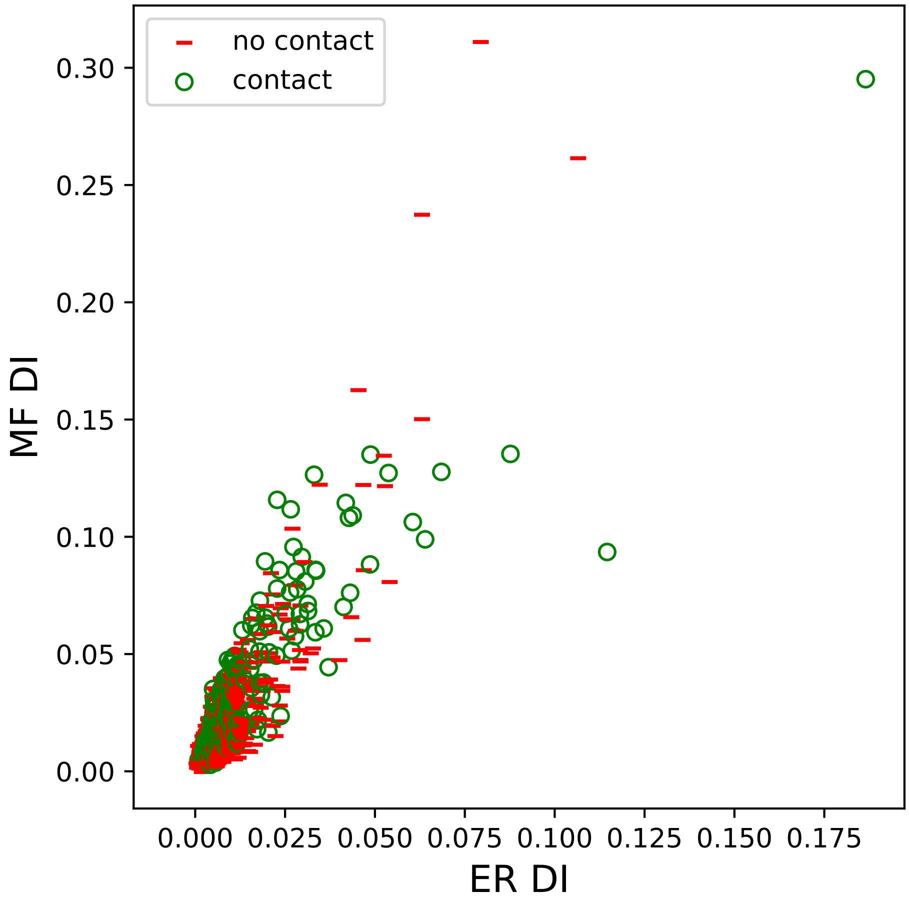
no contact
contact
MF DI
ER DI
0.00
0.05
0.10
0.15
0.20
0.25
0.30
0.000
0.025
0.050
0.075
0.100
0.125
0.150
0.175

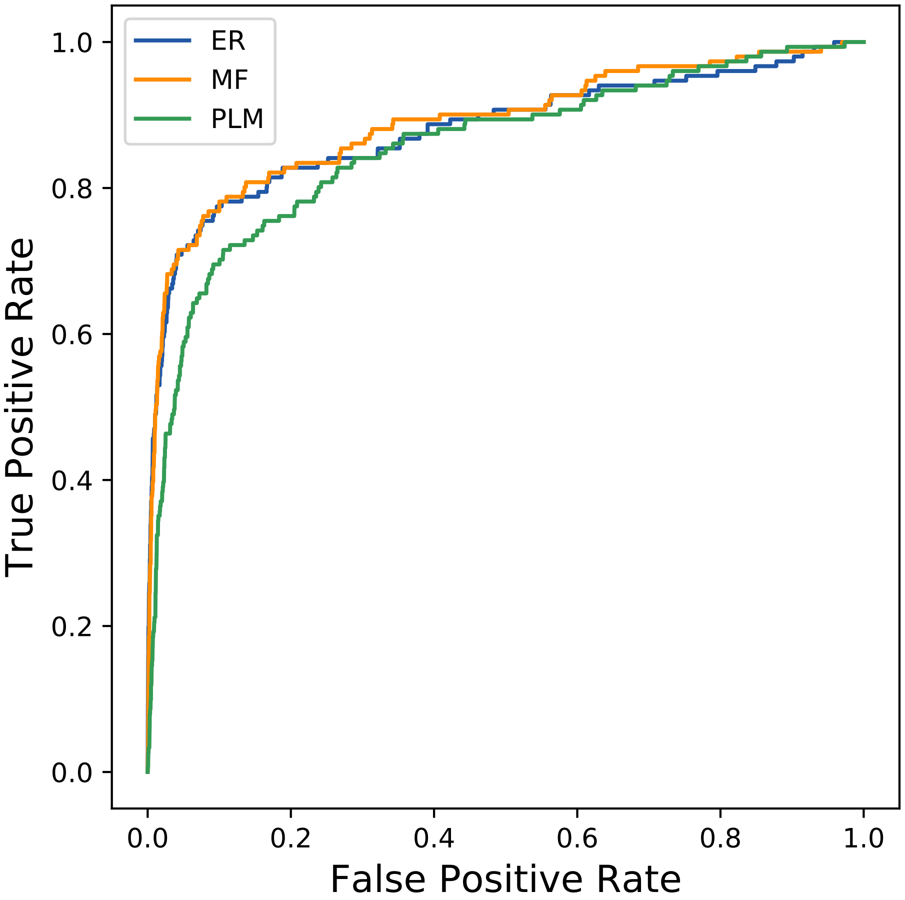
ER
MF
PLM
1.0
0.8
0.6
0.4
0.2
0.0
0.0
0.2
0.4
0.6
0.8
1.0
True Positive Rate
False Positive Rate
