## Supplemental Contact Maps for "Protein Structure Prediction with Expectation Reflection": 7932seq_237col_5i5g_PF00456_plots.pdf

Method: ER
Maximum PDB contact distance : 10.0 Angstrom
Minimum residue chain distance: 5.0 residues
Fraction of true positives : 0.797

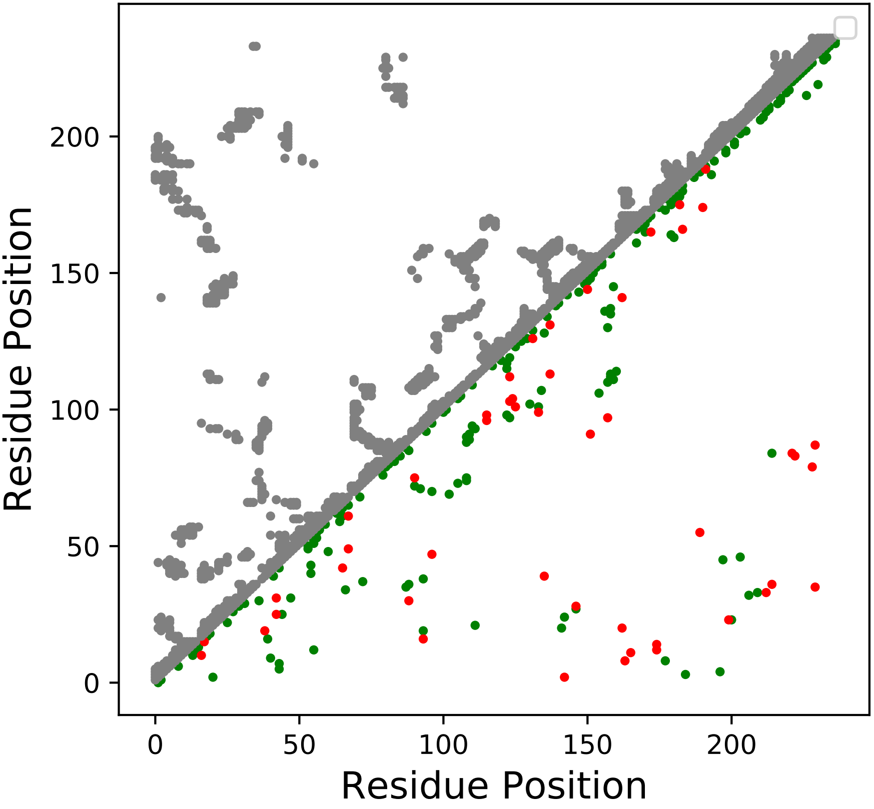

Method: MF
Maximum PDB contact distance : 10.0 Angstrom
Minimum residue chain distance: 5.0 residues
Fraction of true positives : 0.785

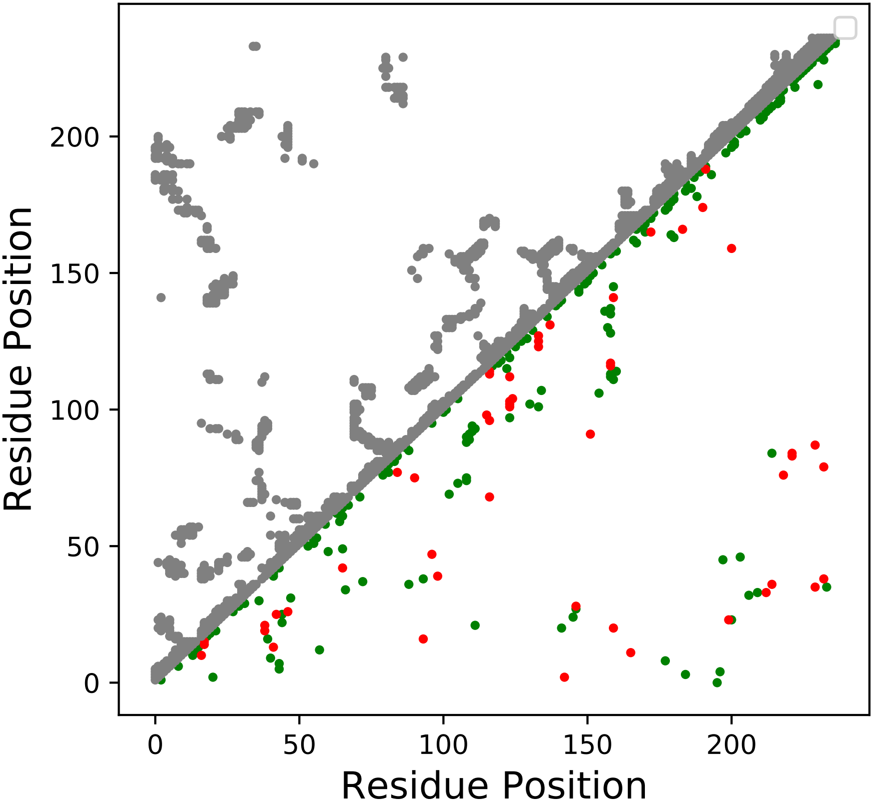

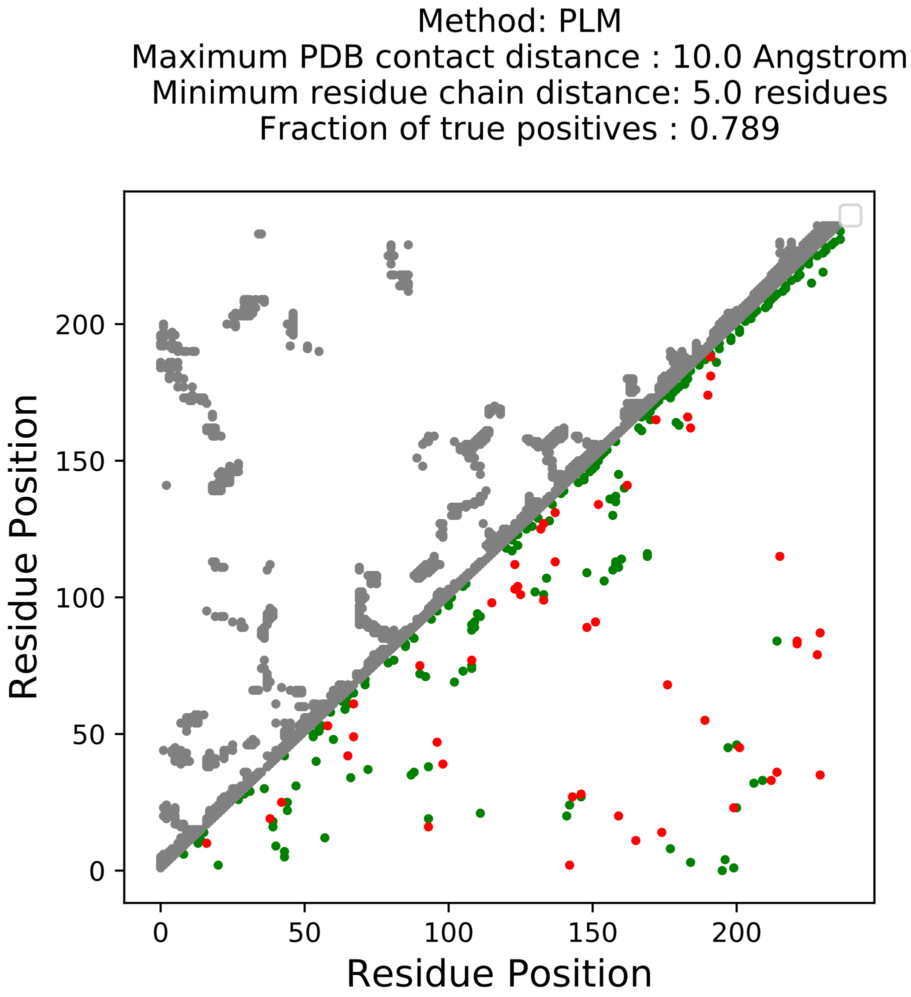
Method: PLM
Maximum PDB contact distance : 10.0 Angstrom
Minimum residue chain distance: 5.0 residues
Fraction of true positives : 0.789
Residue Position
Residue Position
0
50
100
150
200
0
50
100
150
200

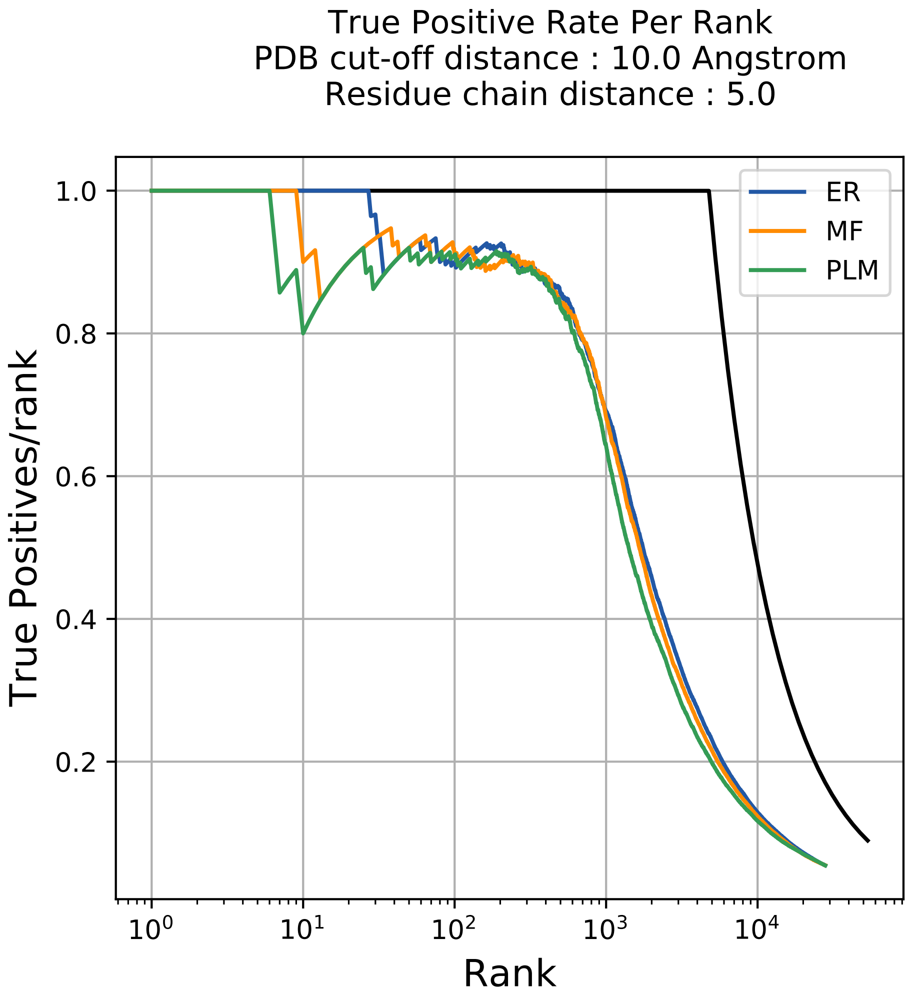

True Positive Rate Per Rank
PDB cut-off distance : 10.0 Angstrom
Residue chain distance : 5.0
True Positives/rank
Rank
ER
MF
PLM
1.0
0.8
0.6
0.4
0.2
$10^0$
$10^1$
$10^2$
$10^3$
$10^4$

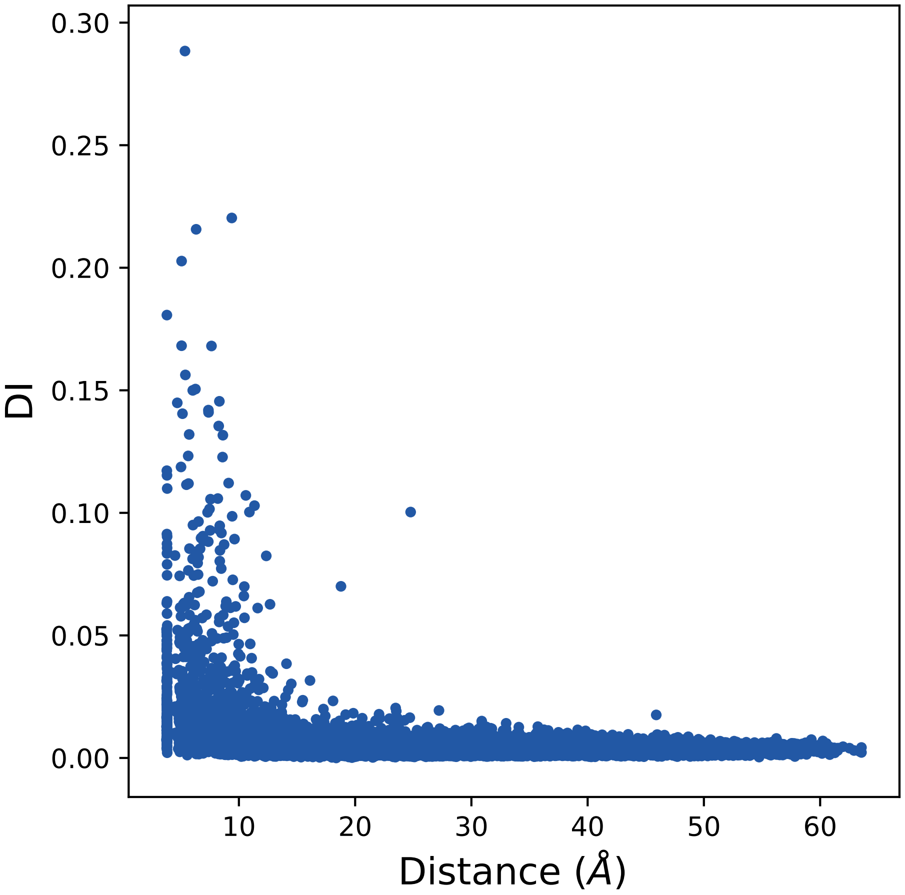

0.30
0.25
0.20
0.15
0.10
0.05
0.00
DI
10
20
30
40
50
60
Distance (Å)

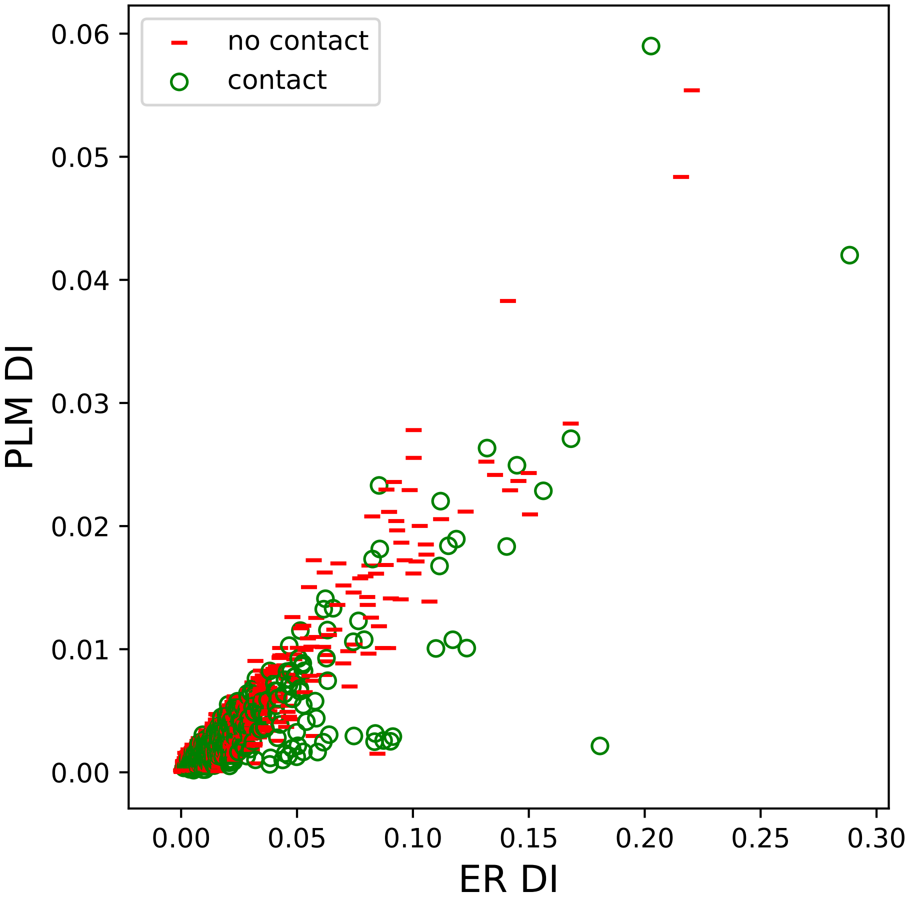
no contact
contact
PLM DI
ER DI
0.06
0.05
0.04
0.03
0.02
0.01
0.00
0.00
0.05
0.10
0.15
0.20
0.25
0.30

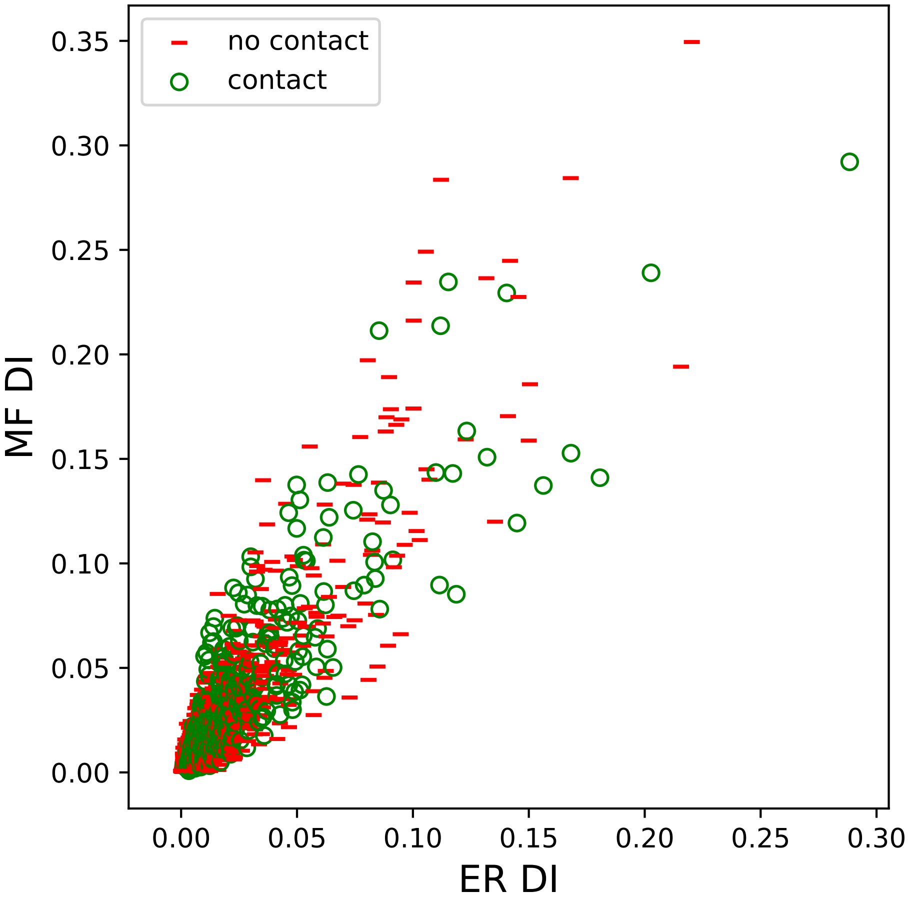

no contact
contact
0.35
0.30
0.25
0.20
0.15
0.10
0.05
0.00
MF DI
0.00
0.05
0.10
0.15
0.20
0.25
0.30
ER DI

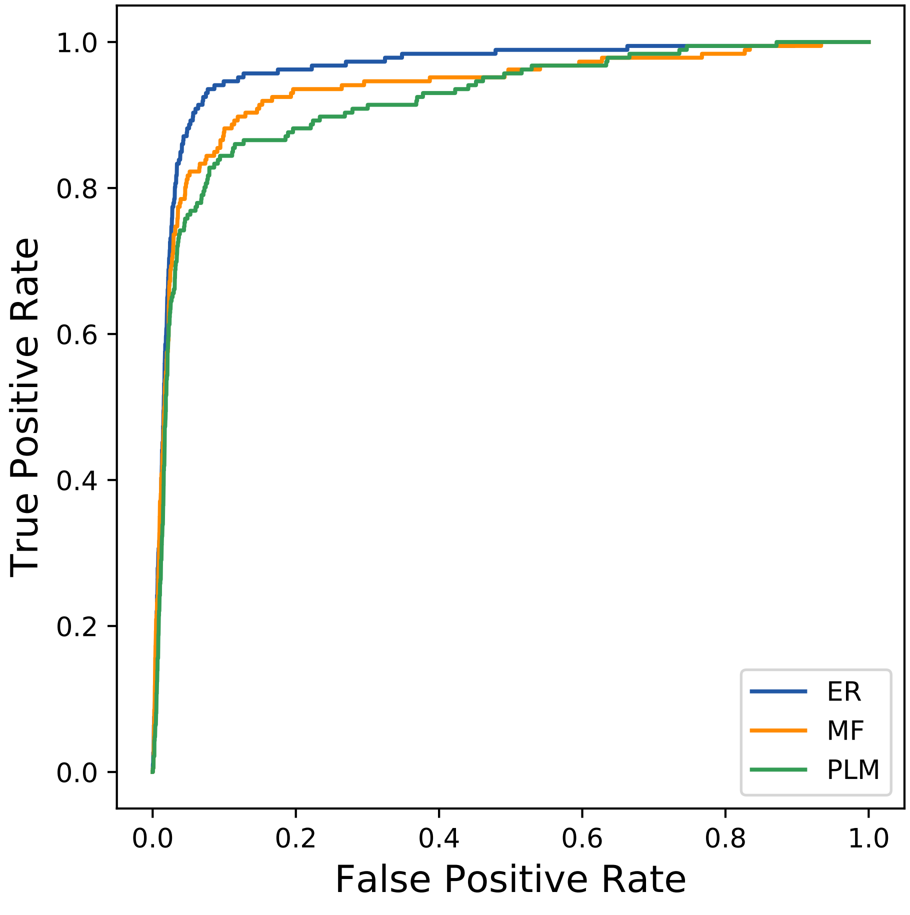
True Positive Rate
False Positive Rate
0.0
0.2
0.4
0.6
0.8
1.0
ER
MF
PLM
