## Supplemental Contact Maps for "Protein Structure Prediction with Expectation Reflection": 8057seq_208col_6cb2_PF02673_plots.pdf

Method: ER
Maximum PDB contact distance : 10.0 Angstrom
Minimum residue chain distance: 5.0 residues
Fraction of true positives : 0.774

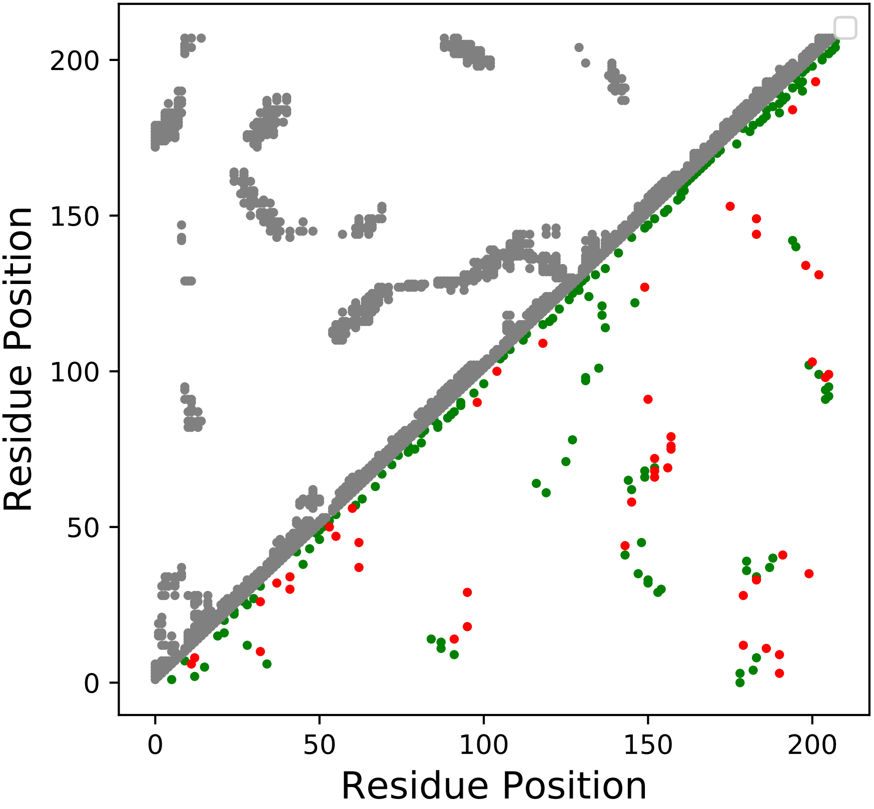

Method: MF
Maximum PDB contact distance : 10.0 Angstrom
Minimum residue chain distance: 5.0 residues
Fraction of true positives : 0.678

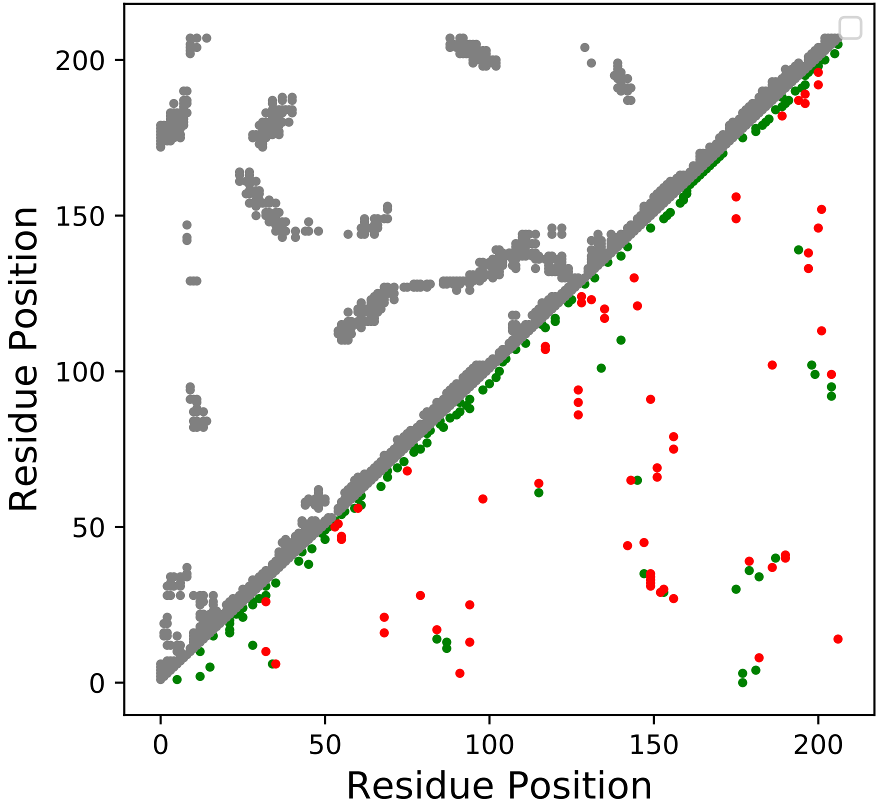

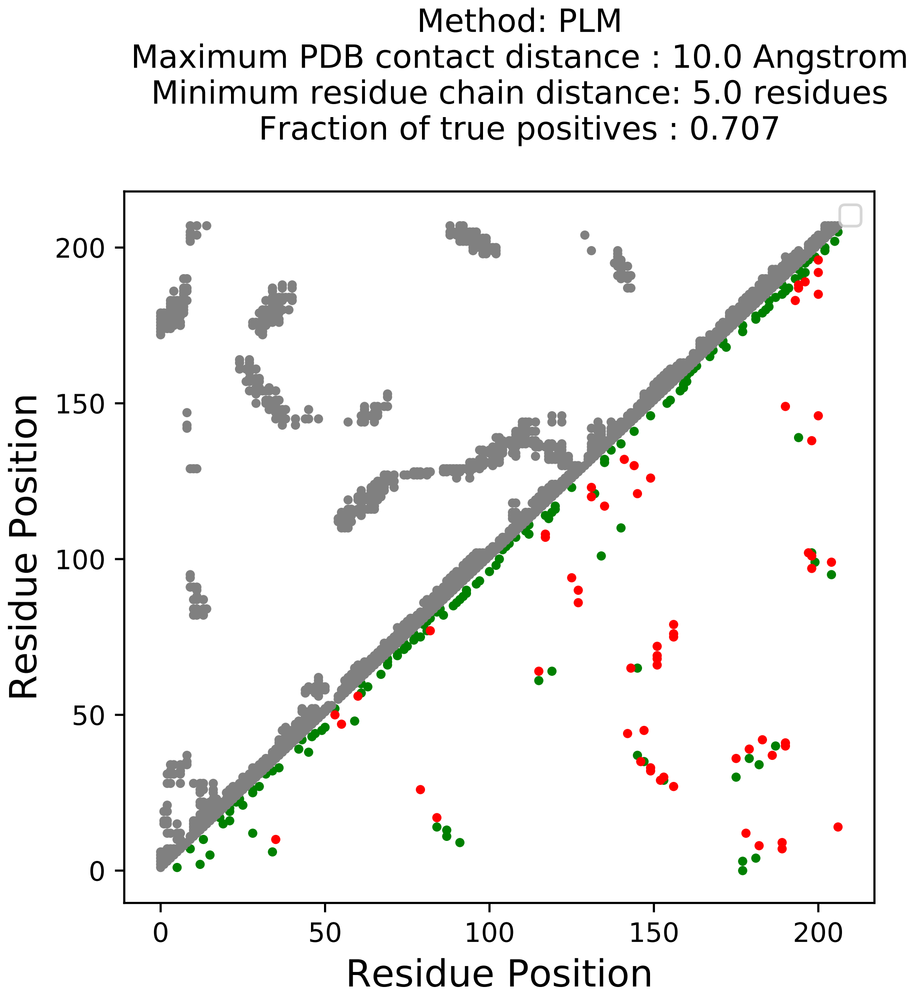

Method: PLM
Maximum PDB contact distance : 10.0 Angstrom
Minimum residue chain distance: 5.0 residues
Fraction of true positives : 0.707
Residue Position
Residue Position
0
50
100
150
200
0
50
100
150
200

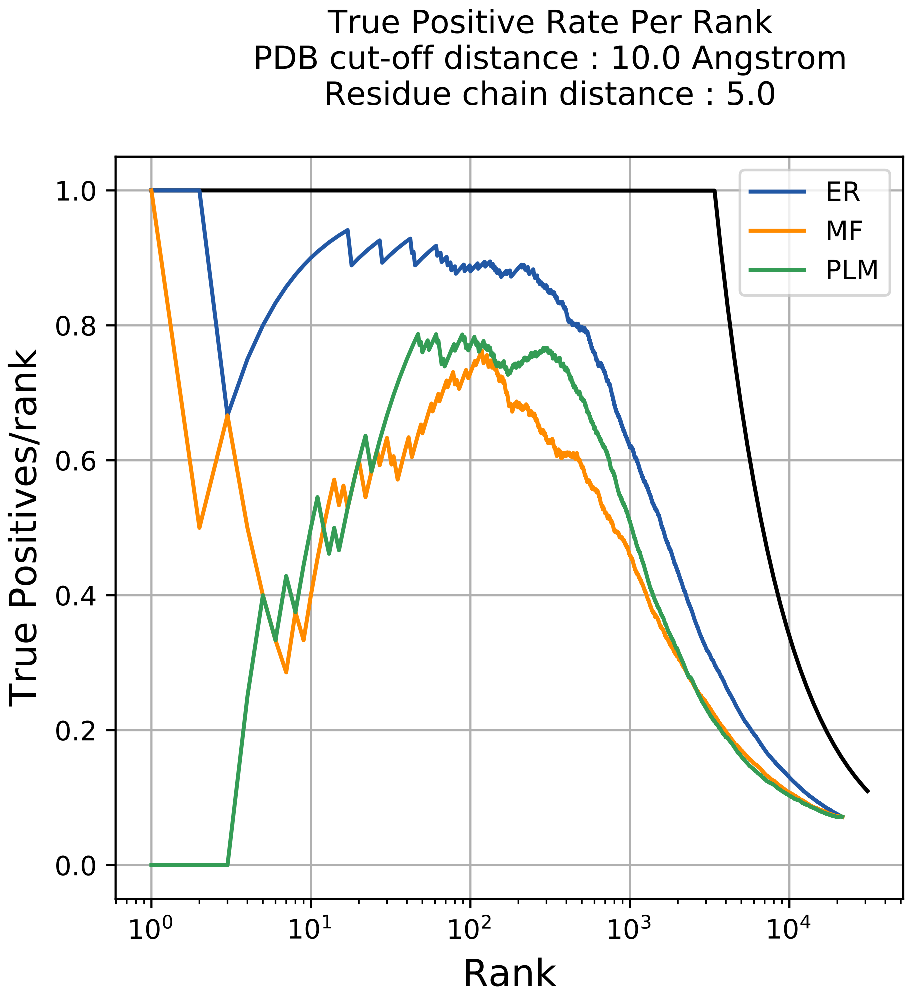

True Positive Rate Per Rank
PDB cut-off distance : 10.0 Angstrom
Residue chain distance : 5.0
True Positives/rank
Rank
1.0
0.8
0.6
0.4
0.2
0.0
$10^0$
$10^1$
$10^2$
$10^3$
$10^4$
ER
MF
PLM

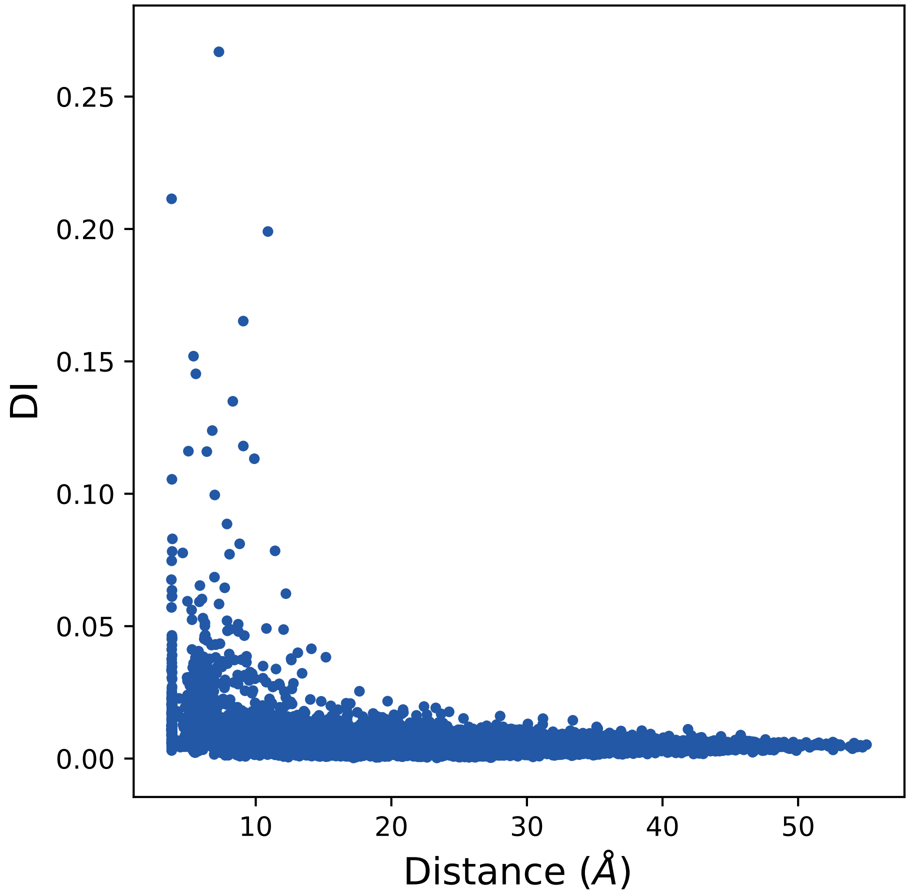

0.25
0.20
0.15
0.10
0.05
0.00
DI
10
20
30
40
50
Distance (Å)

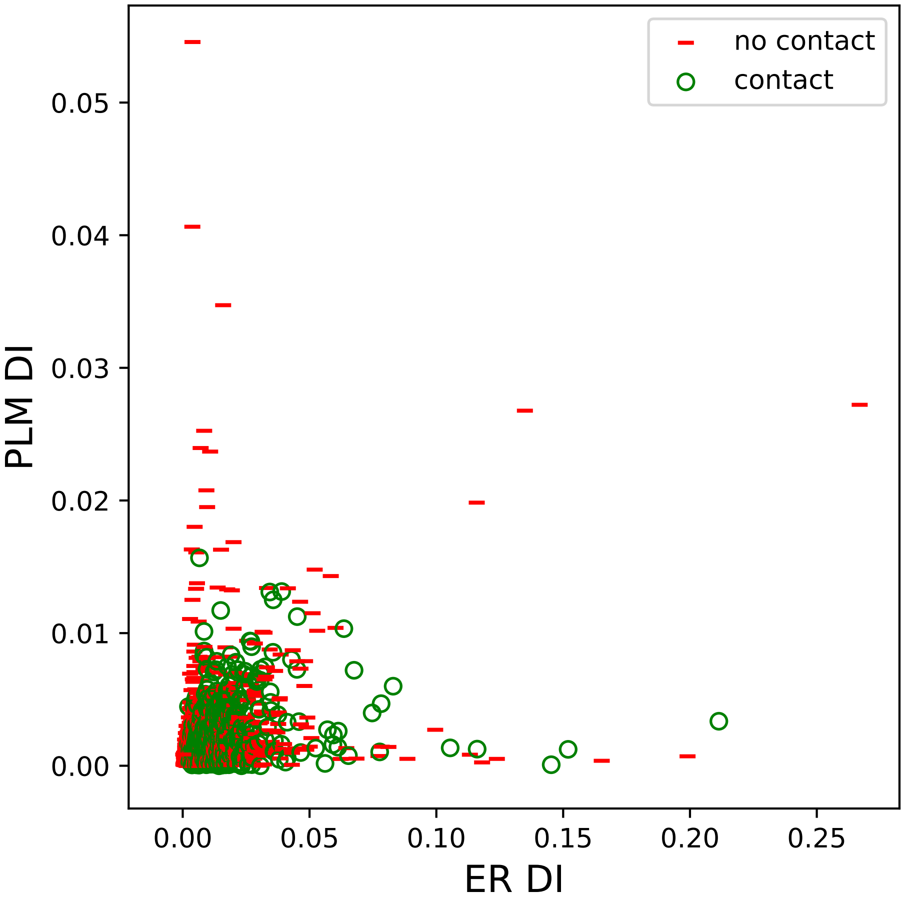

no contact
contact
0.05
0.04
0.03
0.02
0.01
0.00
PLM DI
0.00
0.05
0.10
0.15
0.20
0.25
ER DI

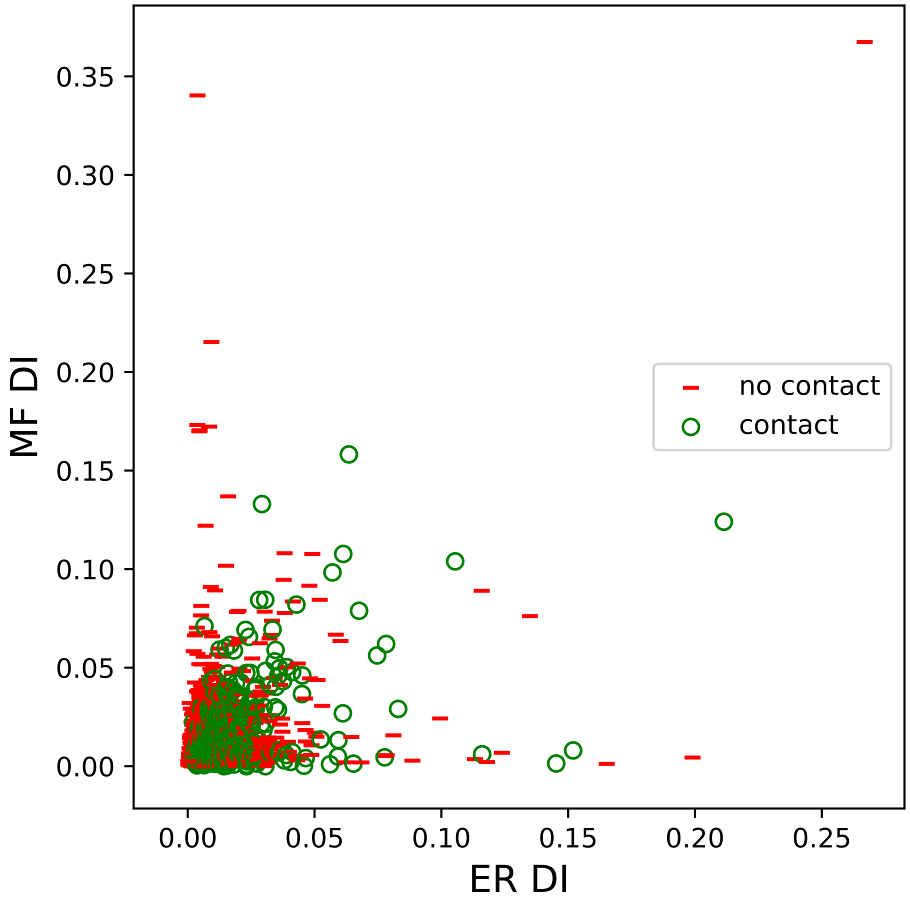

MF DI
ER DI
0.35
0.30
0.25
0.20
0.15
0.10
0.05
0.00
0.00
0.05
0.10
0.15
0.20
0.25
no contact
contact

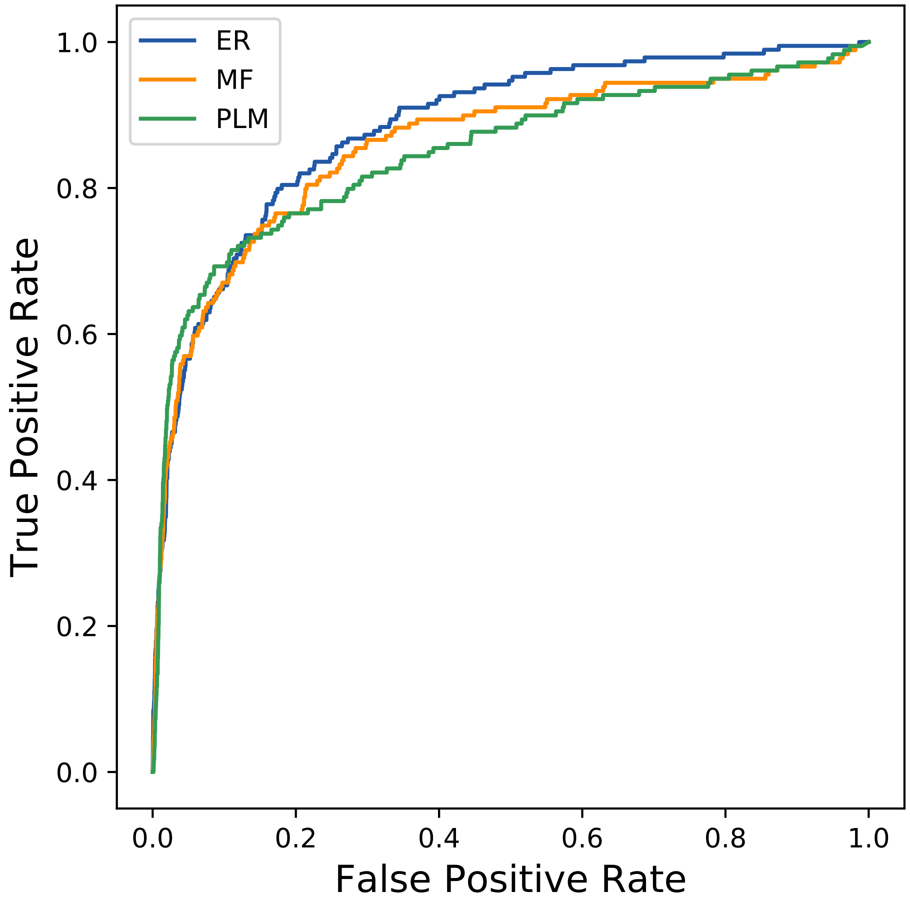
ER
MF
PLM
True Positive Rate
False Positive Rate
0.0
0.2
0.4
0.6
0.8
1.0
