## Supplemental Contact Maps for "Protein Structure Prediction with Expectation Reflection": 8363seq_67col_4ig4_PF13344_plots.pdf

Method: ER
Maximum PDB contact distance : 10.0 Angstrom
Minimum residue chain distance: 5.0 residues
Fraction of true positives : 0.582

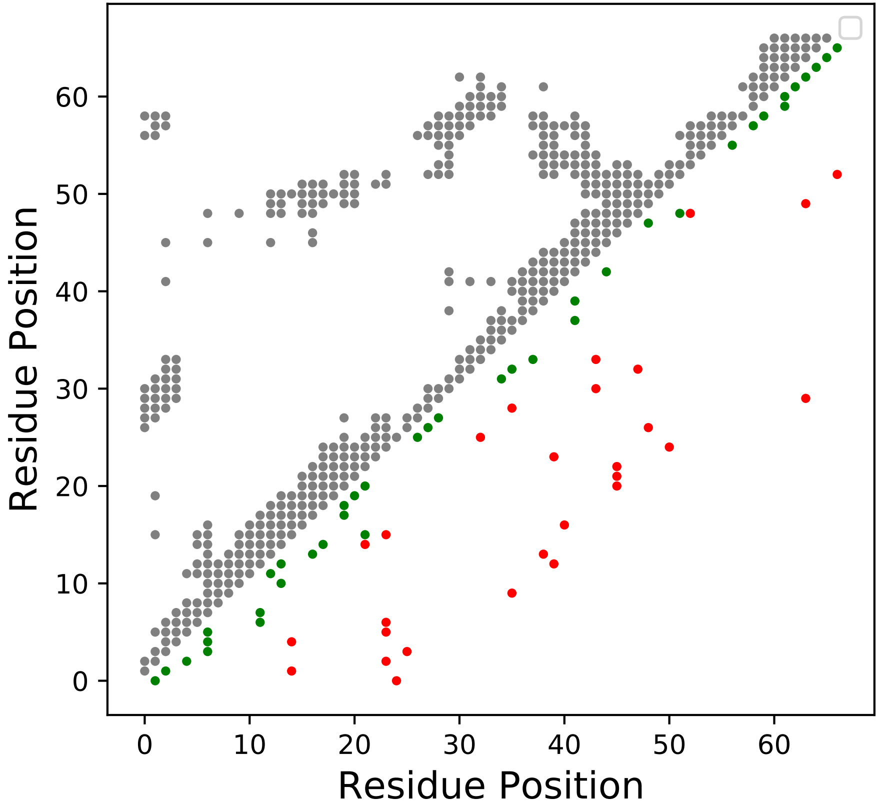

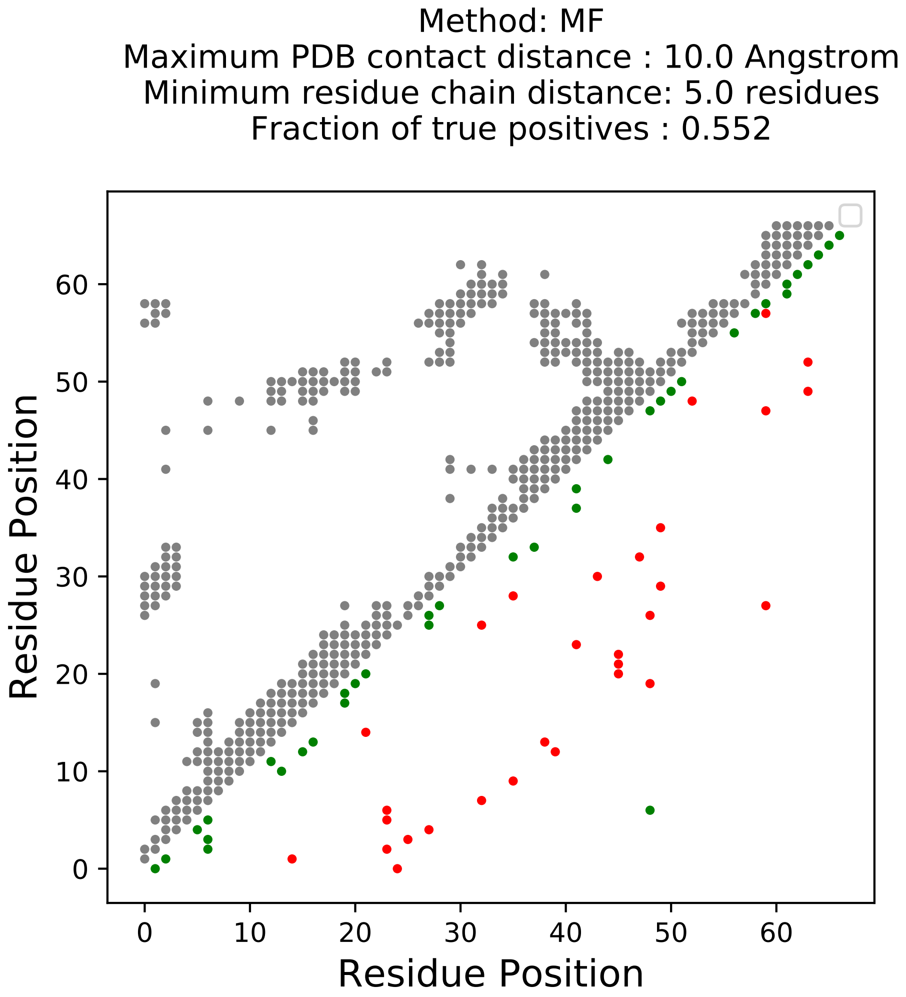

Method: MF
Maximum PDB contact distance : 10.0 Angstrom
Minimum residue chain distance: 5.0 residues
Fraction of true positives : 0.552
Residue Position
Residue Position
0
10
20
30
40
50
60

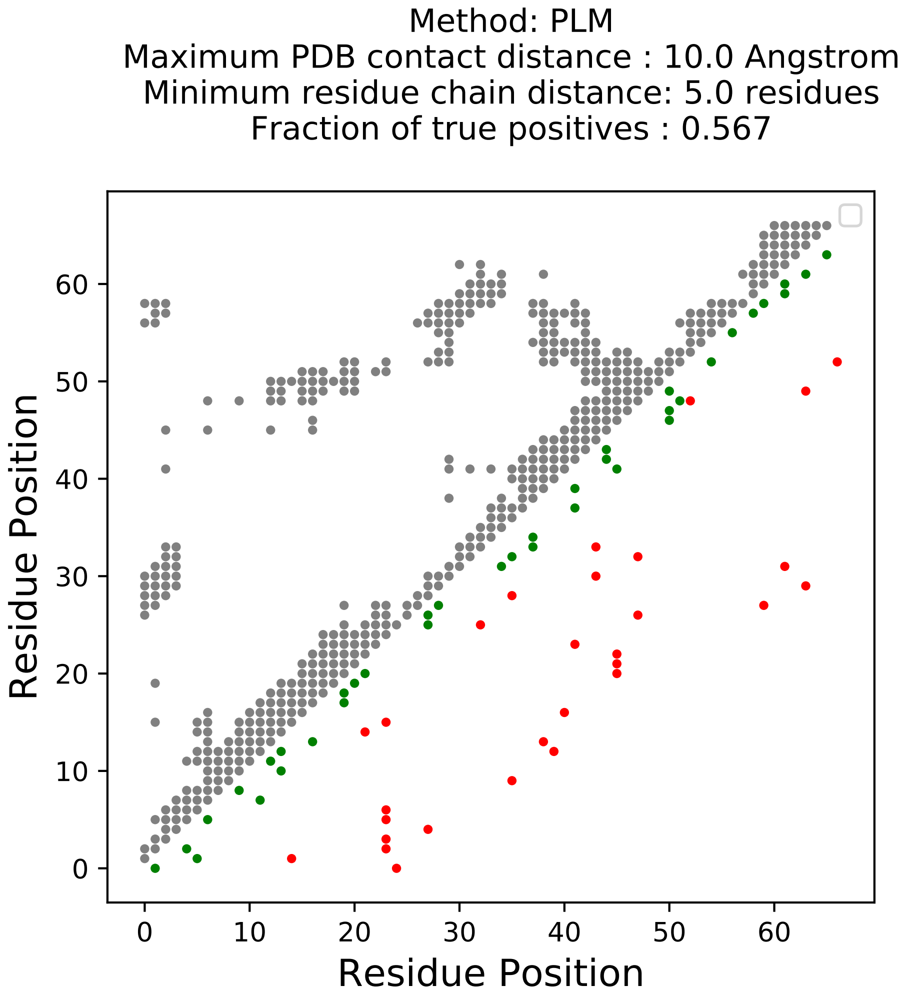

Method: PLM
Maximum PDB contact distance : 10.0 Angstrom
Minimum residue chain distance: 5.0 residues
Fraction of true positives : 0.567
Residue Position
Residue Position
0
10
20
30
40
50
60

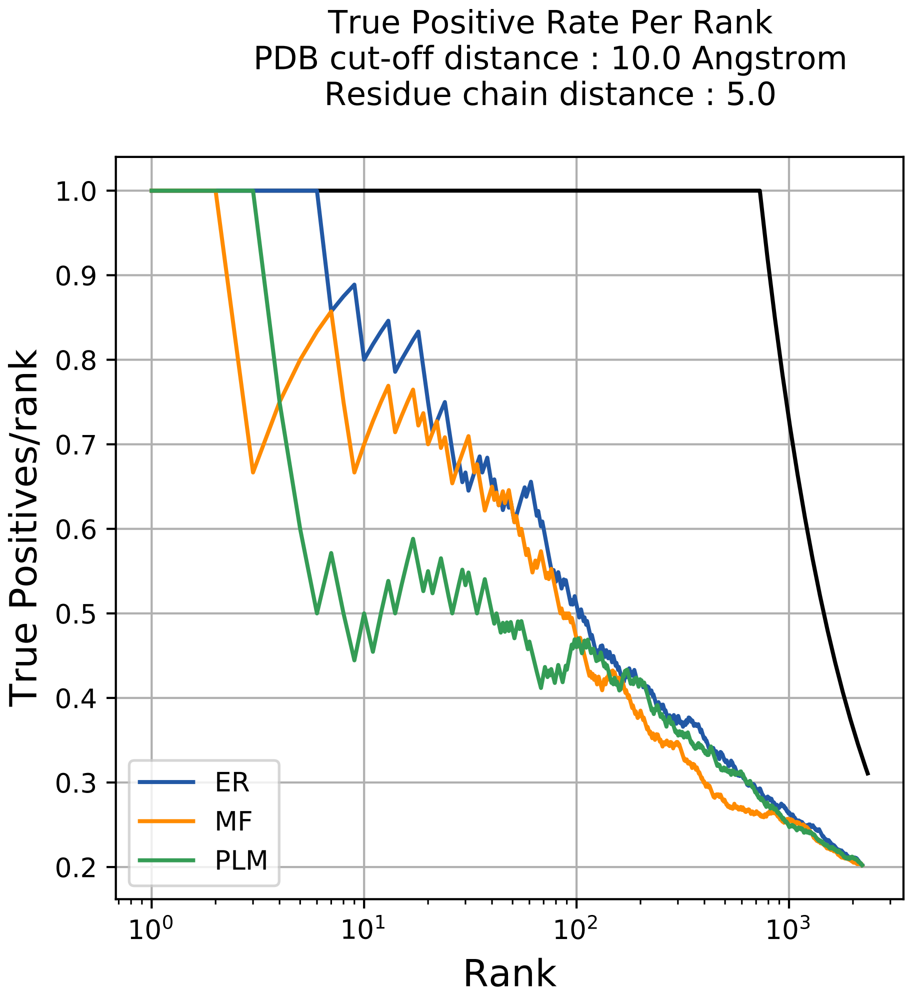
True Positive Rate Per Rank
PDB cut-off distance : 10.0 Angstrom
Residue chain distance : 5.0
True Positives/rank
1.0
0.9
0.8
0.7
0.6
0.5
0.4
0.3
0.2
$10^0$
$10^1$
$10^2$
$10^3$
Rank
ER
MF
PLM

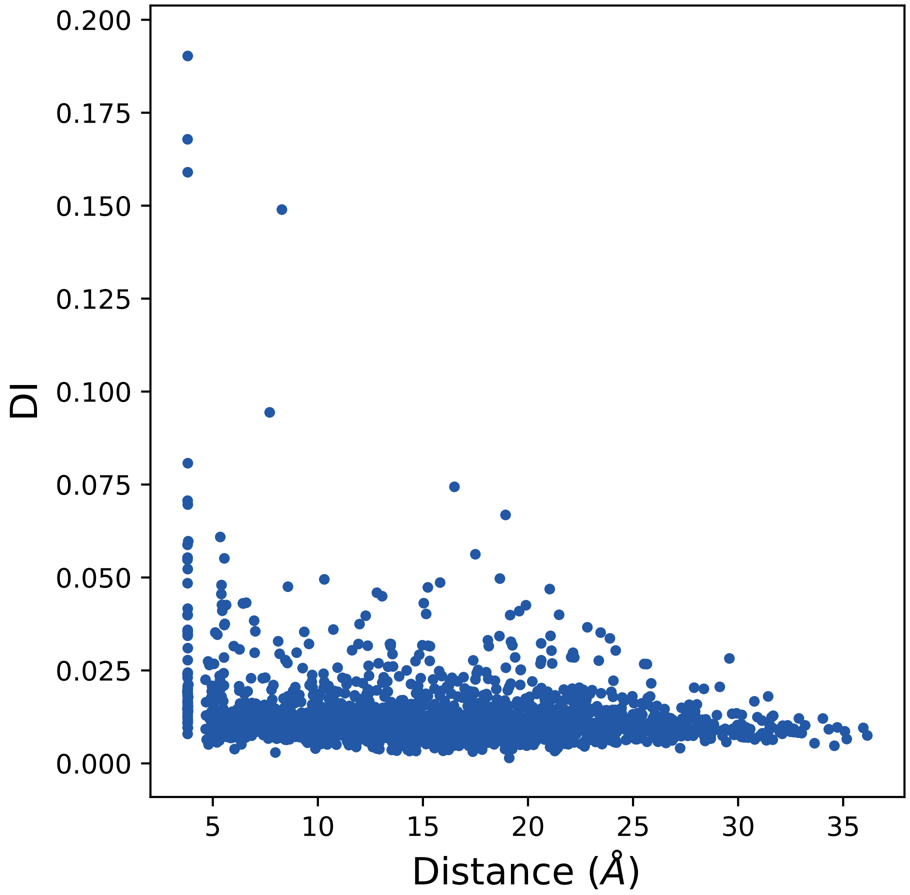
0.200
0.175
0.150
0.125
0.100
0.075
0.050
0.025
0.000
DI
5
10
15
20
25
30
35
Distance (Å)

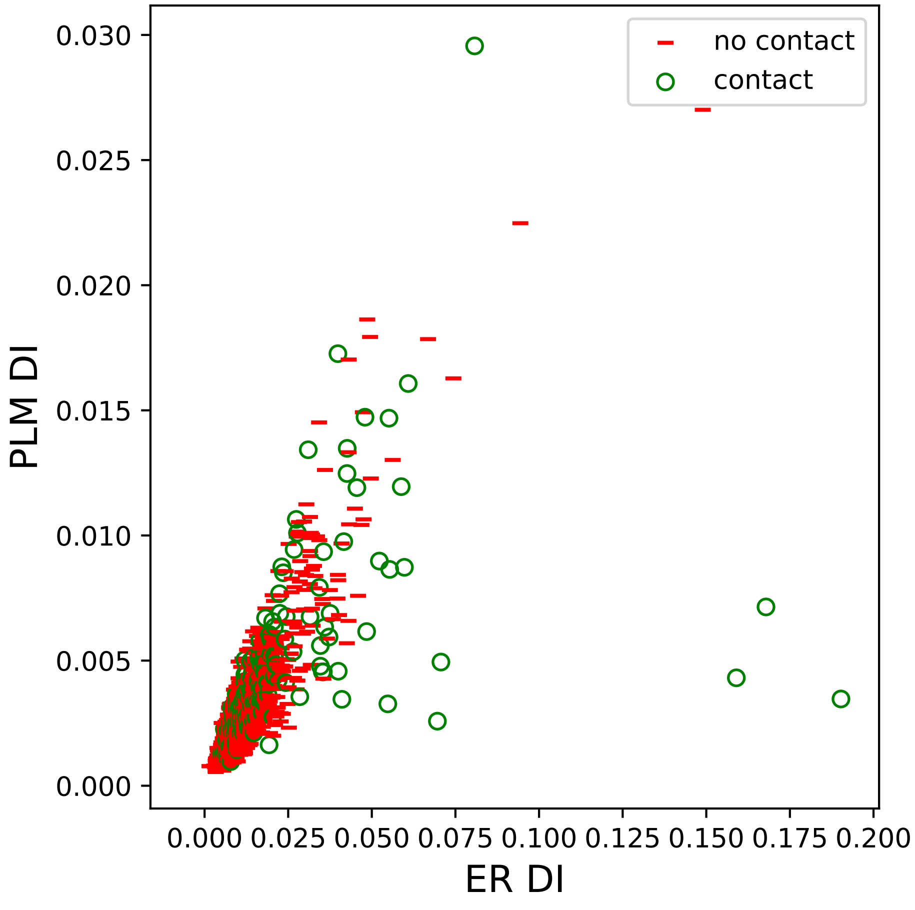
no contact
contact
PLM DI
ER DI
0.000
0.005
0.010
0.015
0.020
0.025
0.030
0.000 0.025 0.050 0.075 0.100 0.125 0.150 0.175 0.200

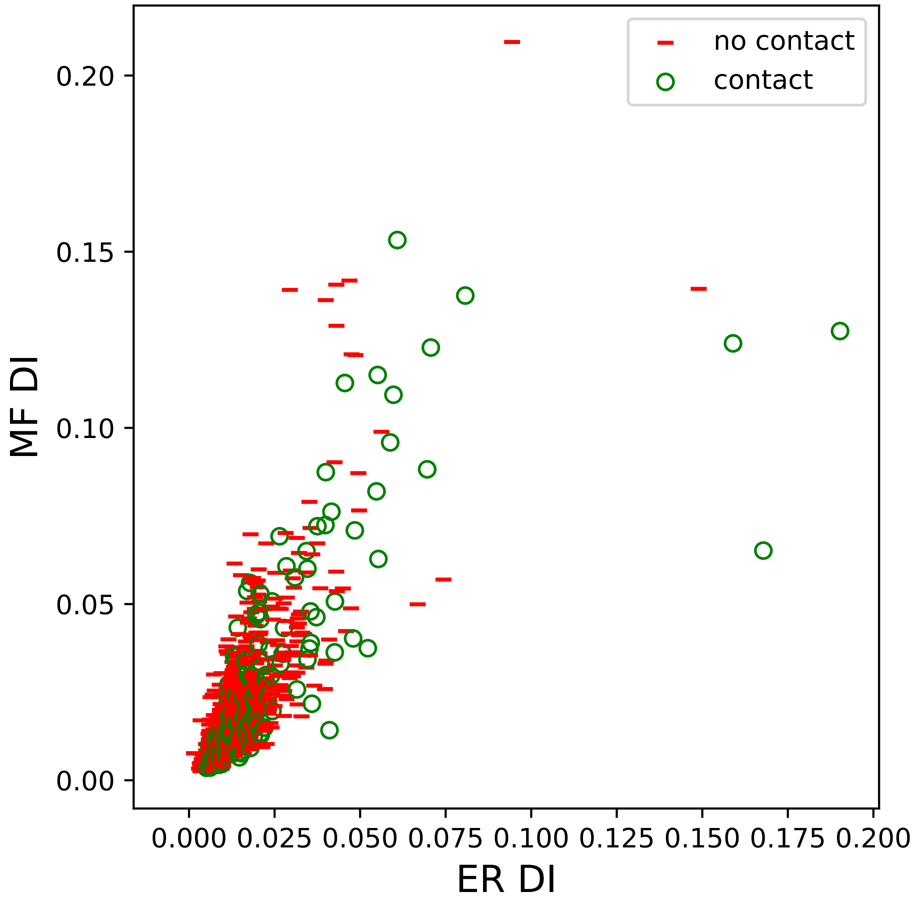

no contact
contact
MF DI
ER DI
0.00
0.05
0.10
0.15
0.20
0.000
0.025
0.050
0.075
0.100
0.125
0.150
0.175
0.200

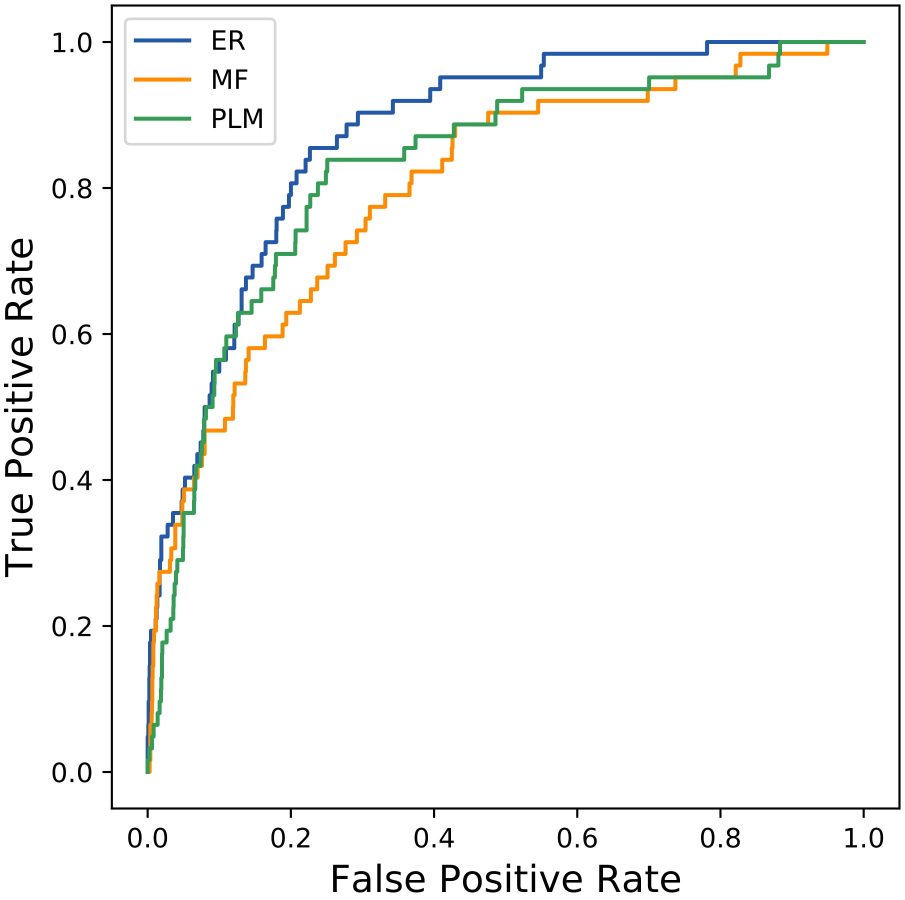

ER
MF
PLM
1.0
0.8
0.6
0.4
0.2
0.0
True Positive Rate
0.0
0.2
0.4
0.6
0.8
1.0
False Positive Rate
