## Supplemental Contact Maps for "Protein Structure Prediction with Expectation Reflection": 8791seq_107col_6e8y_PF07479_plots.pdf

Method: ER
Maximum PDB contact distance : 10.0 Angstrom
Minimum residue chain distance: 5.0 residues
Fraction of true positives : 0.729

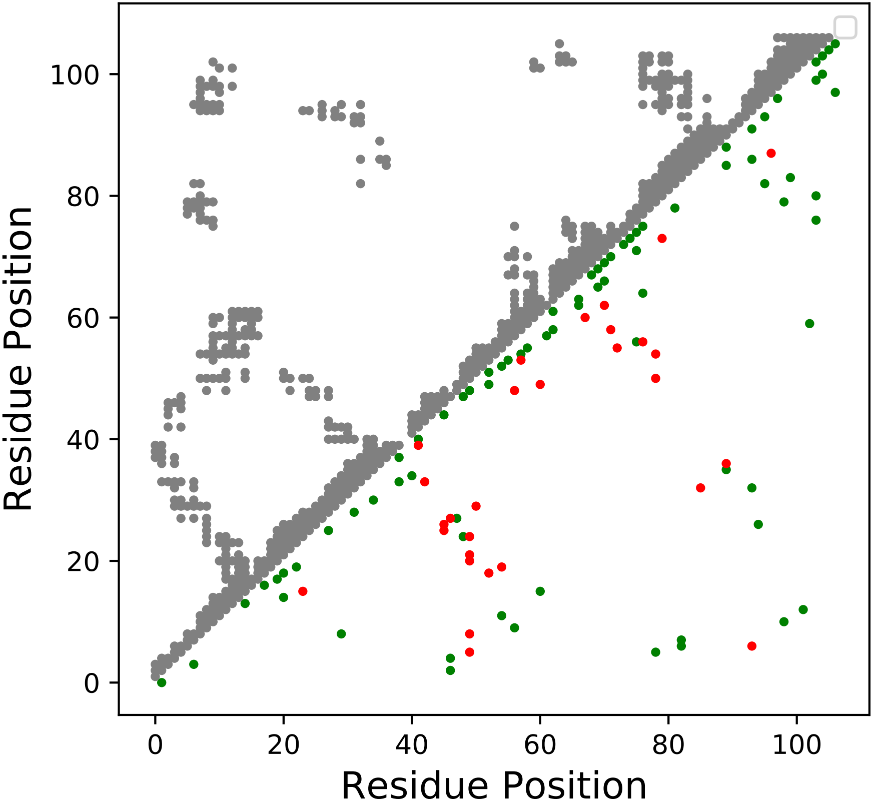

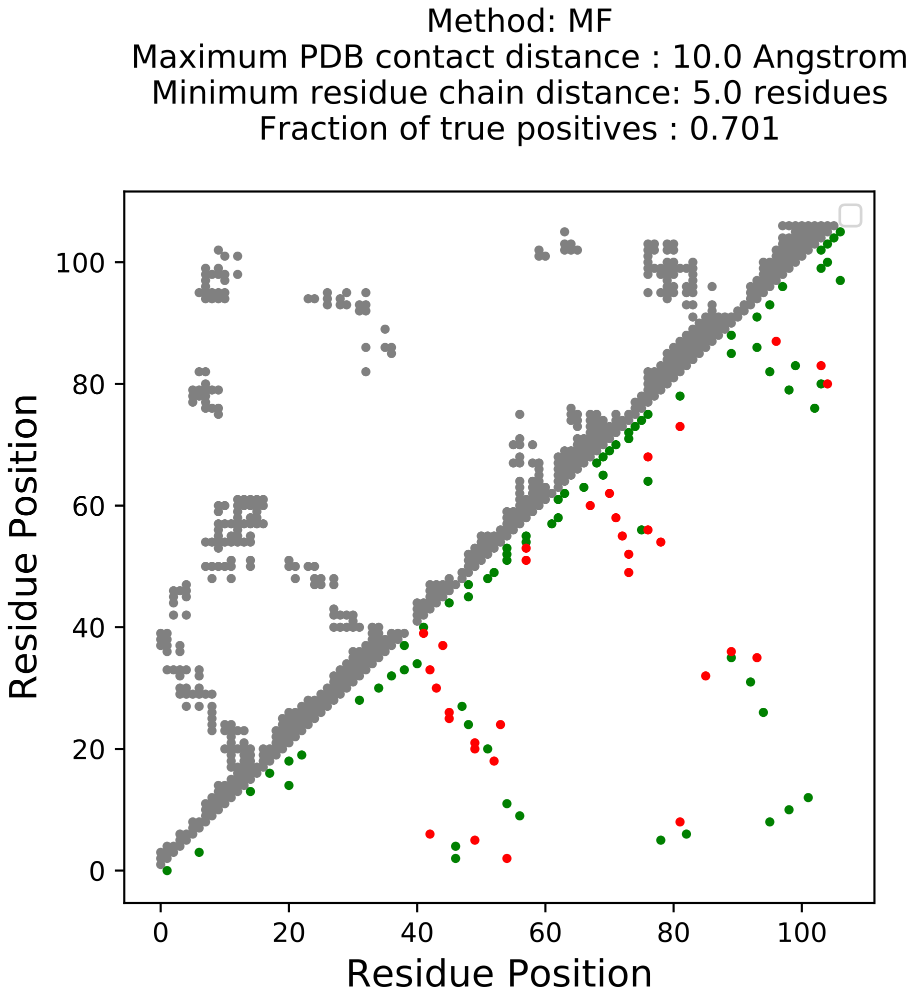

Method: MF
Maximum PDB contact distance : 10.0 Angstrom
Minimum residue chain distance: 5.0 residues
Fraction of true positives : 0.701
Residue Position
Residue Position
0
20
40
60
80
100

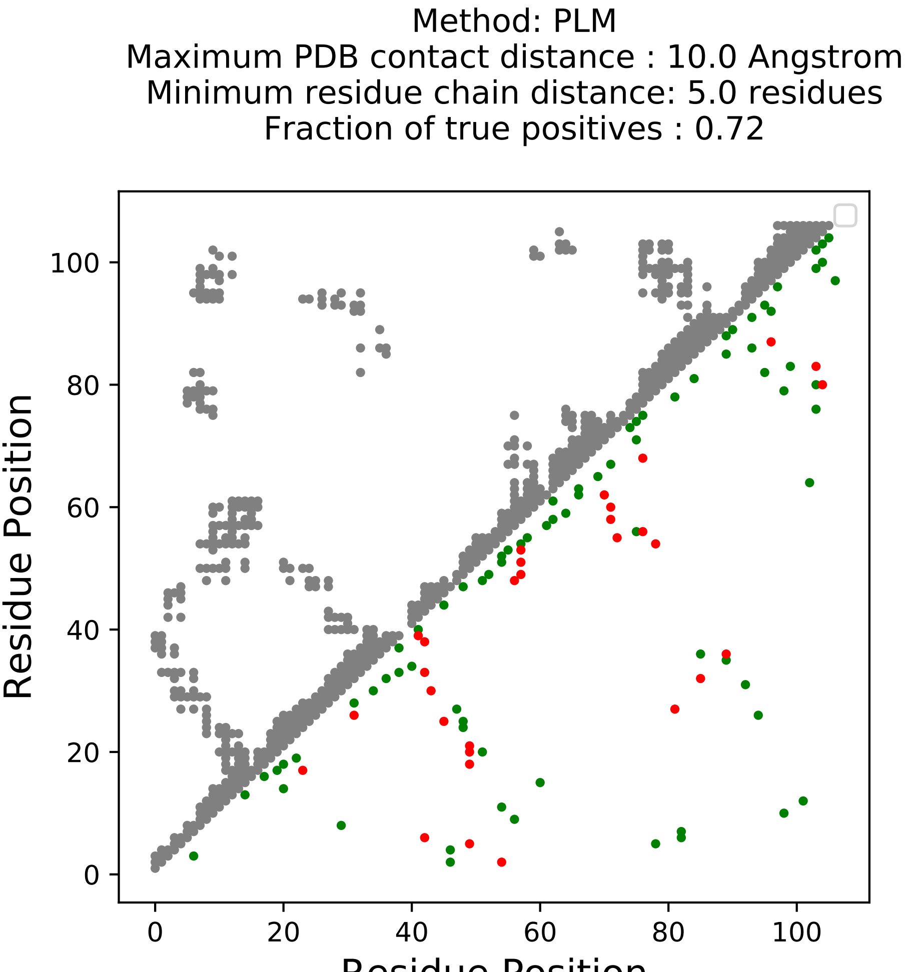
Method: PLM
Maximum PDB contact distance : 10.0 Angstrom
Minimum residue chain distance: 5.0 residues
Fraction of true positives : 0.72
Residue Position
Residue Position
0
20
40
60
80
100

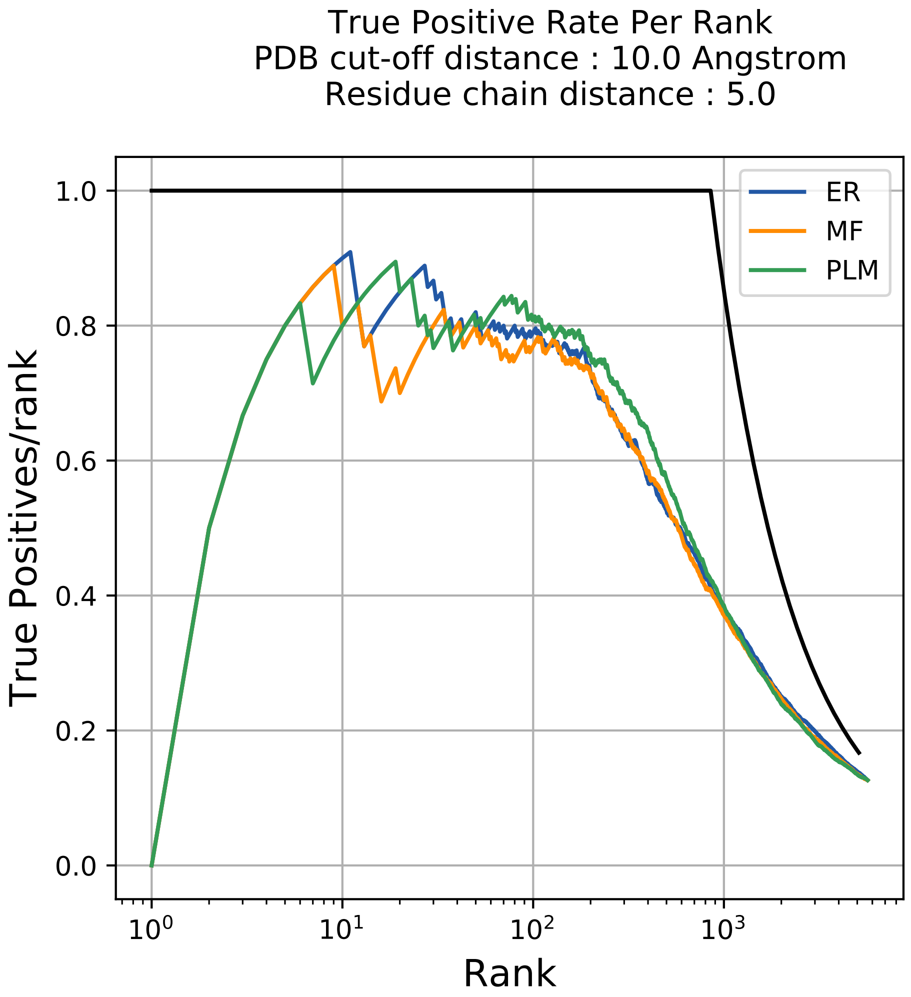
True Positive Rate Per Rank
PDB cut-off distance : 10.0 Angstrom
Residue chain distance : 5.0
ER
MF
PLM
1.0
0.8
0.6
0.4
0.2
0.0
True Positives/rank
$10^0$
$10^1$
$10^2$
$10^3$
Rank

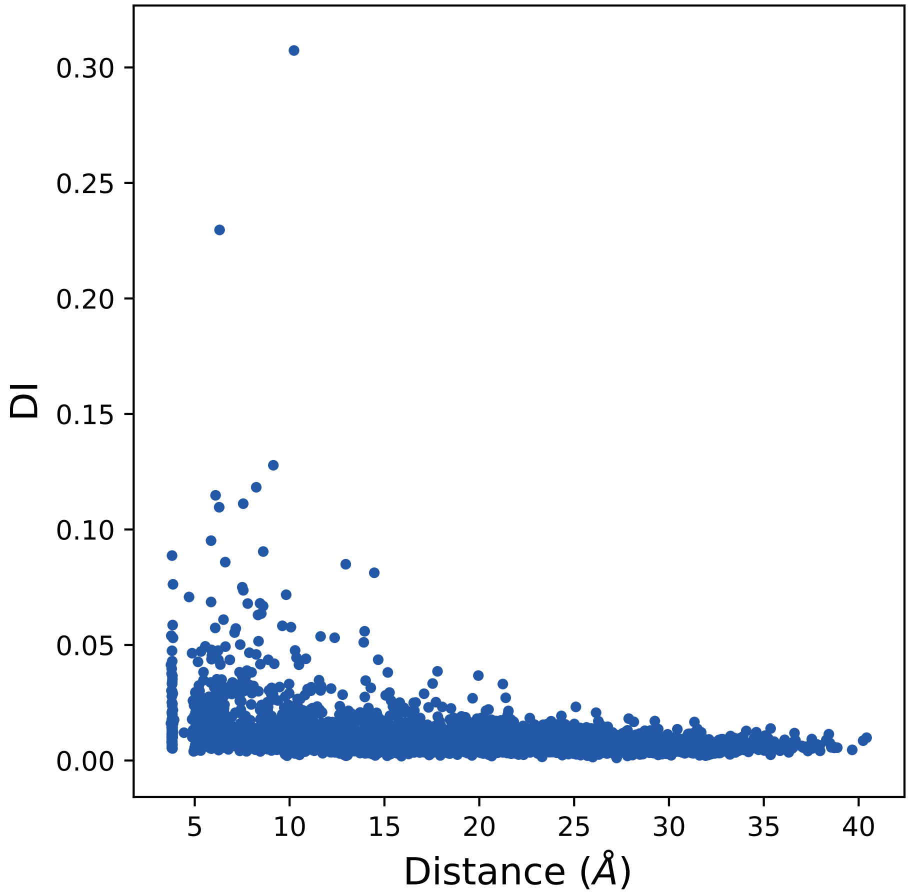

0.30
0.25
0.20
0.15
0.10
0.05
0.00
DI
5
10
15
20
25
30
35
40
Distance (Å)

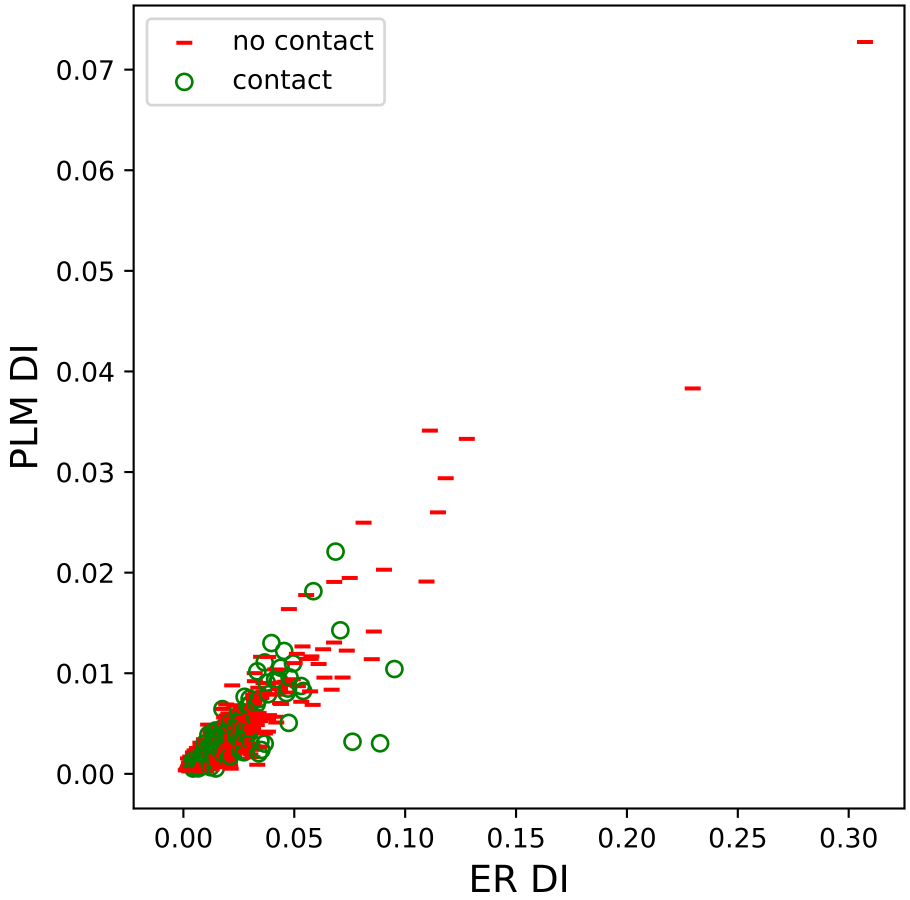

no contact
contact
PLM DI
0.07
0.06
0.05
0.04
0.03
0.02
0.01
0.00
0.00
0.05
0.10
0.15
0.20
0.25
0.30
ER DI

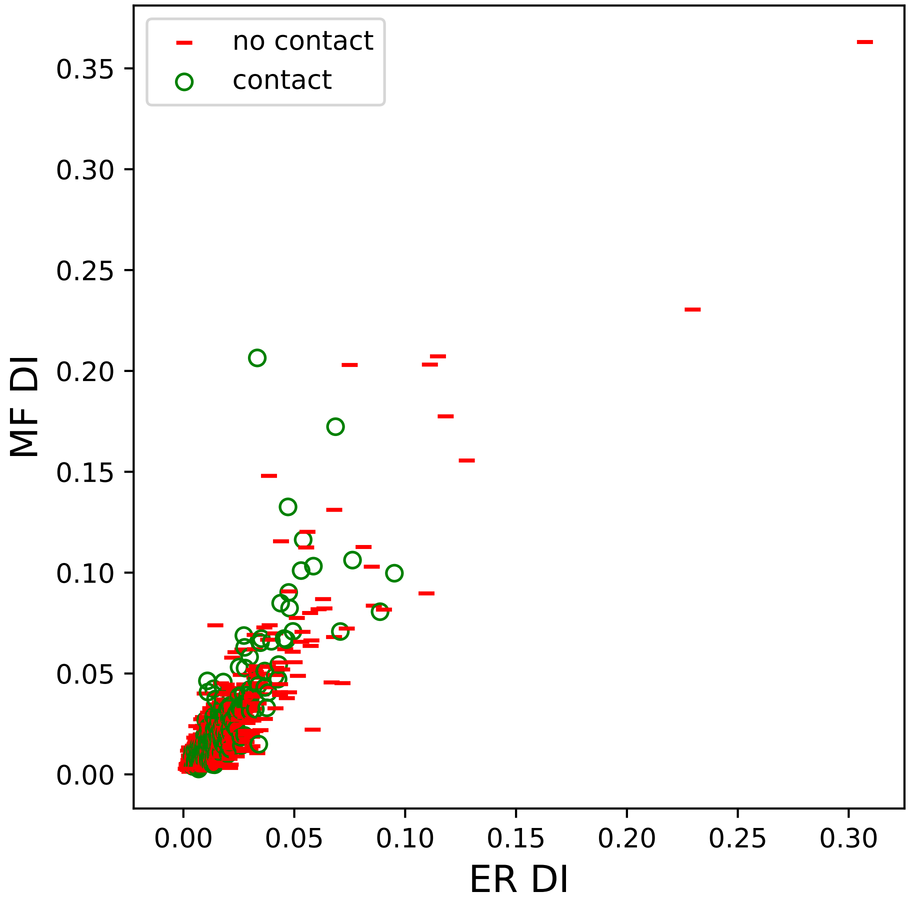

no contact
contact
MF DI
ER DI
0.00
0.05
0.10
0.15
0.20
0.25
0.30
0.35

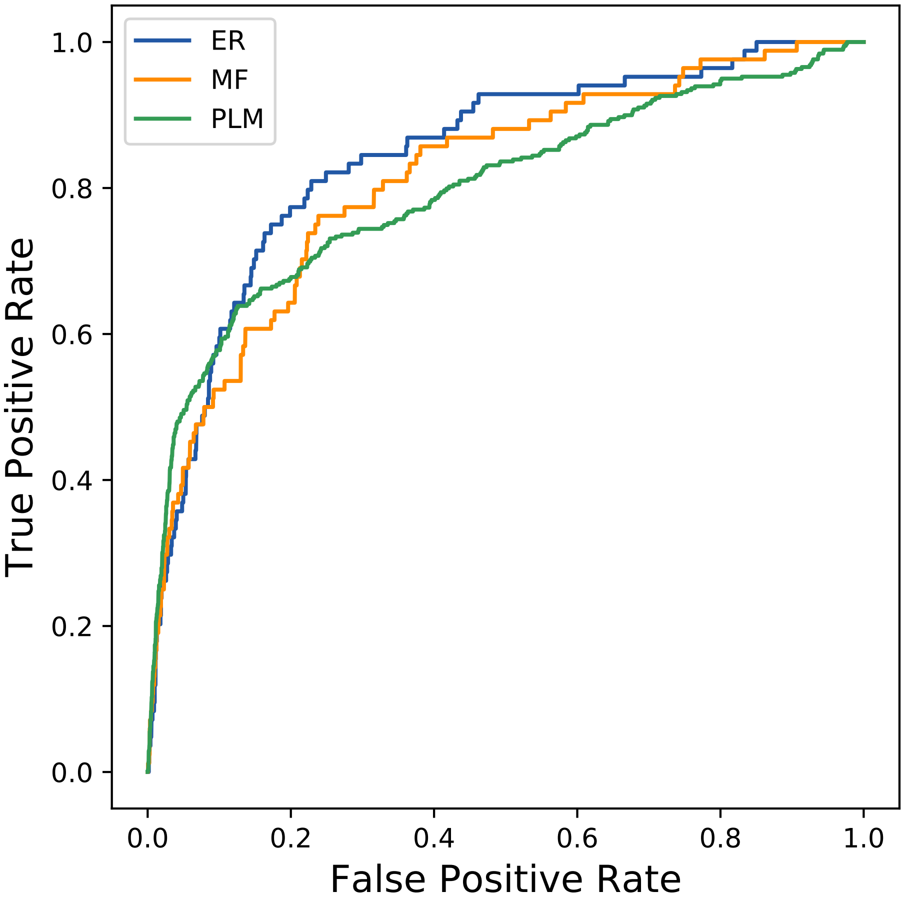
ER
MF
PLM
1.0
0.8
0.6
0.4
0.2
0.0
0.0
0.2
0.4
0.6
0.8
1.0
True Positive Rate
False Positive Rate
