## Supplemental Contact Maps for "Protein Structure Prediction with Expectation Reflection": 10139seq_94col_4aoo_PF00692_plots.pdf

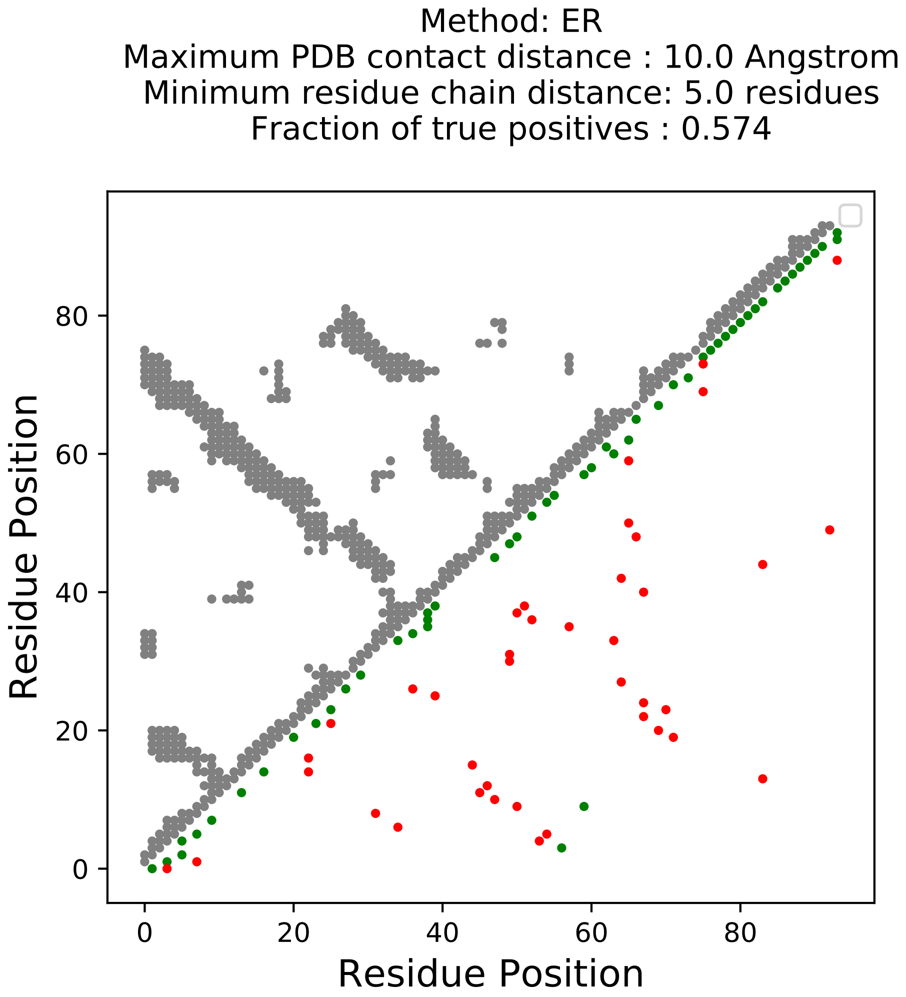
Method: ER
Maximum PDB contact distance : 10.0 Angstrom
Minimum residue chain distance: 5.0 residues
Fraction of true positives : 0.574
Residue Position
Residue Position
0
20
40
60
80
0
20
40
60
80

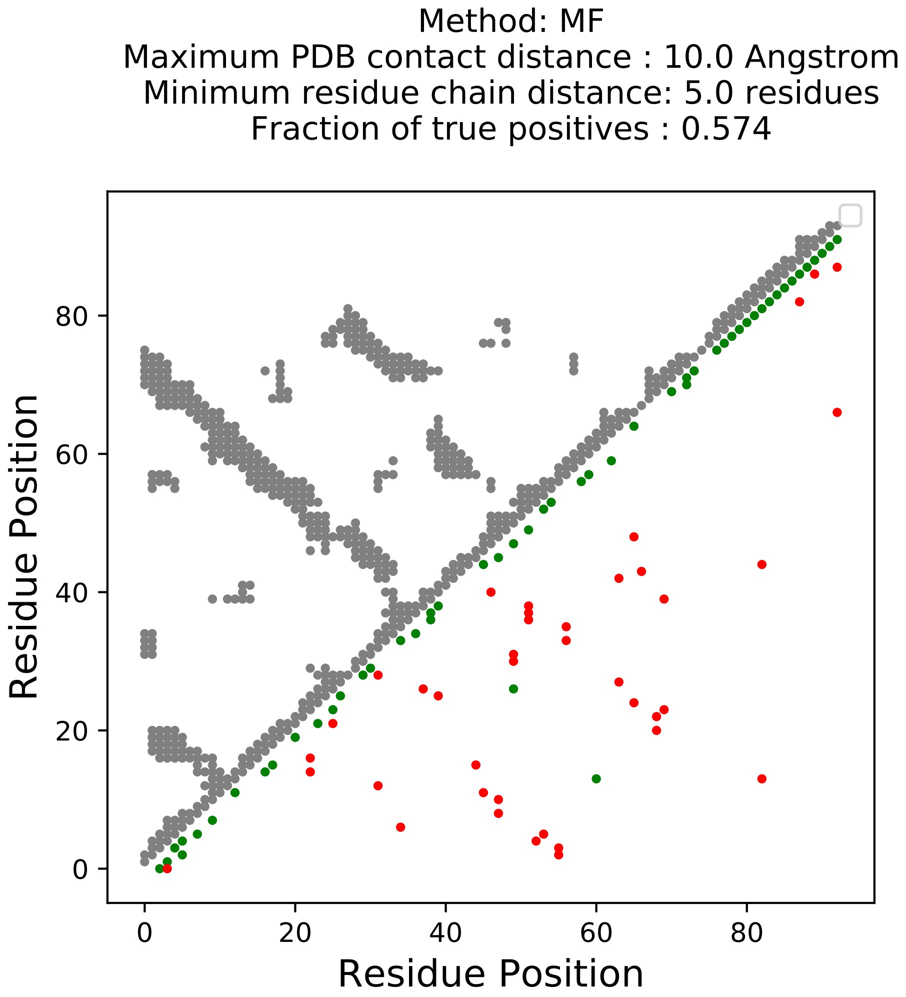
Method: MF
Maximum PDB contact distance : 10.0 Angstrom
Minimum residue chain distance: 5.0 residues
Fraction of true positives : 0.574
Residue Position
Residue Position
0
20
40
60
80
0
20
40
60
80

Method: PLM
Maximum PDB contact distance : 10.0 Angstrom
Minimum residue chain distance: 5.0 residues
Fraction of true positives : 0.521

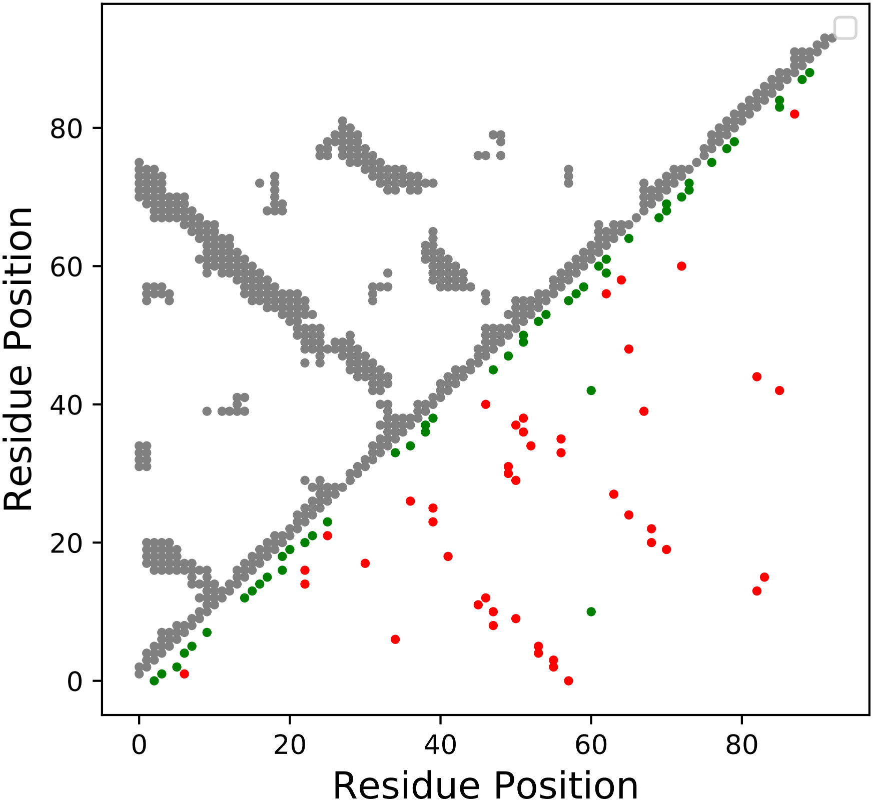

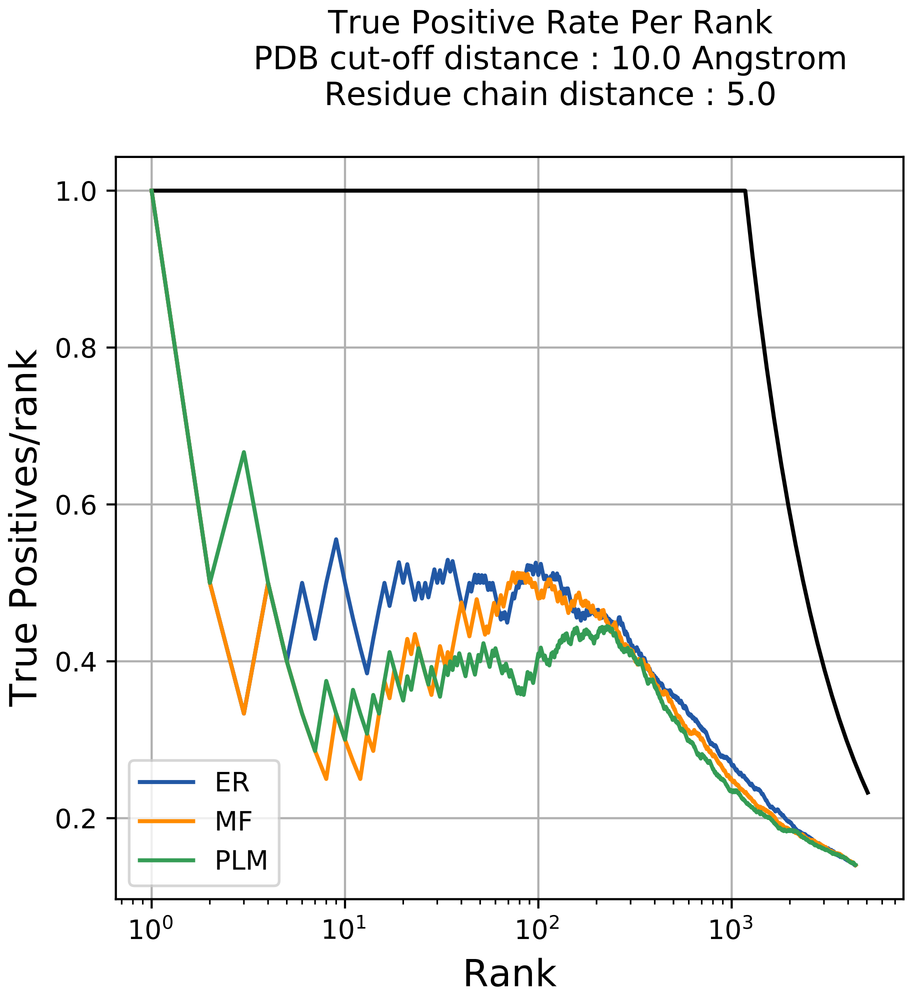
True Positive Rate Per Rank
PDB cut-off distance : 10.0 Angstrom
Residue chain distance : 5.0
True Positives/rank
1.0
0.8
0.6
0.4
0.2
Rank
$10^0$
$10^1$
$10^2$
$10^3$
ER
MF
PLM

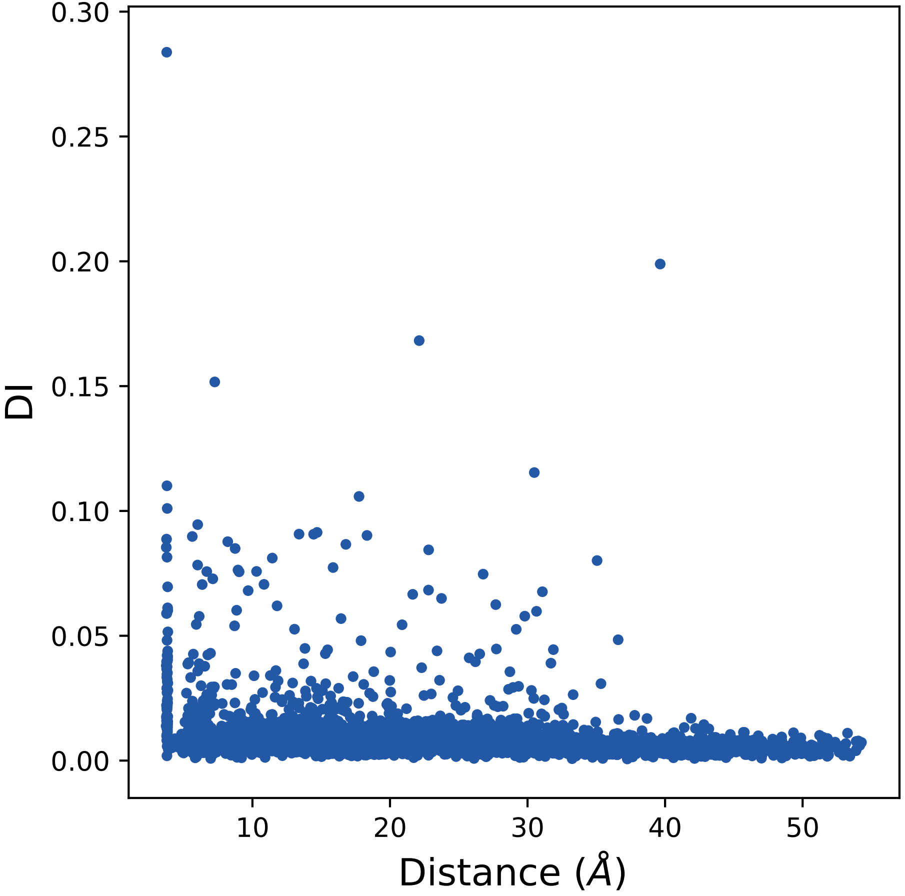
0.30
0.25
0.20
0.15
0.10
0.05
0.00
DI
10
20
30
40
50
Distance (Å)

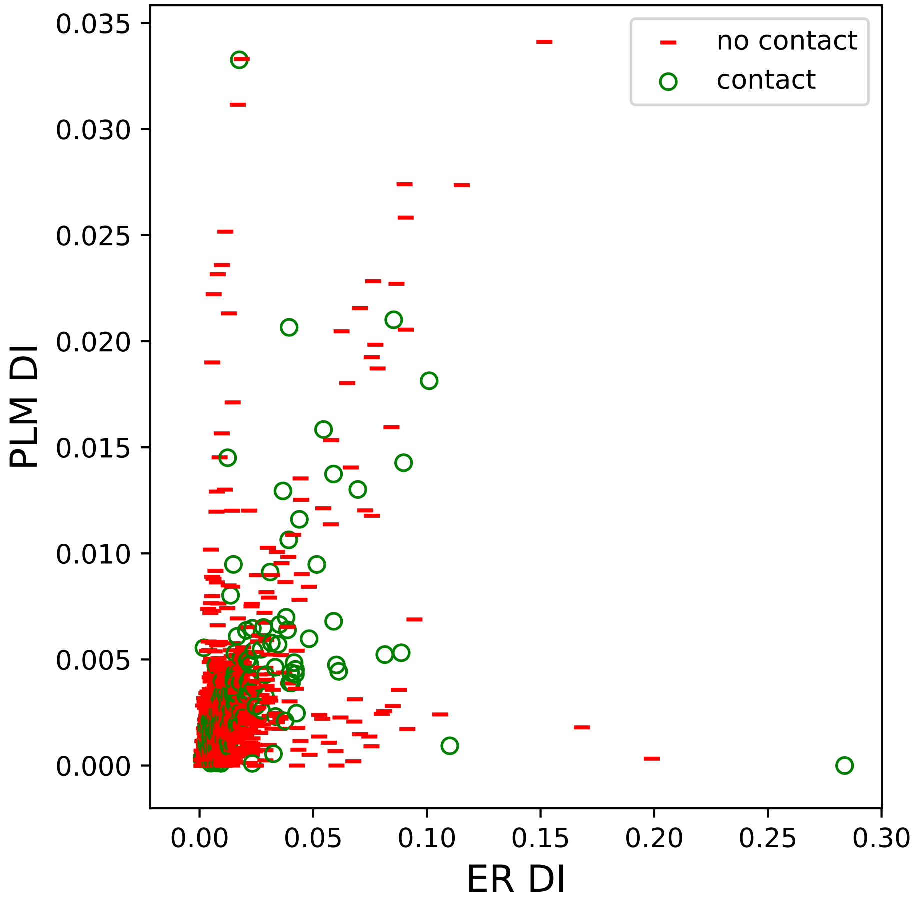

no contact
contact
PLM DI
ER DI
0.035
0.030
0.025
0.020
0.015
0.010
0.005
0.000
0.00
0.05
0.10
0.15
0.20
0.25
0.30

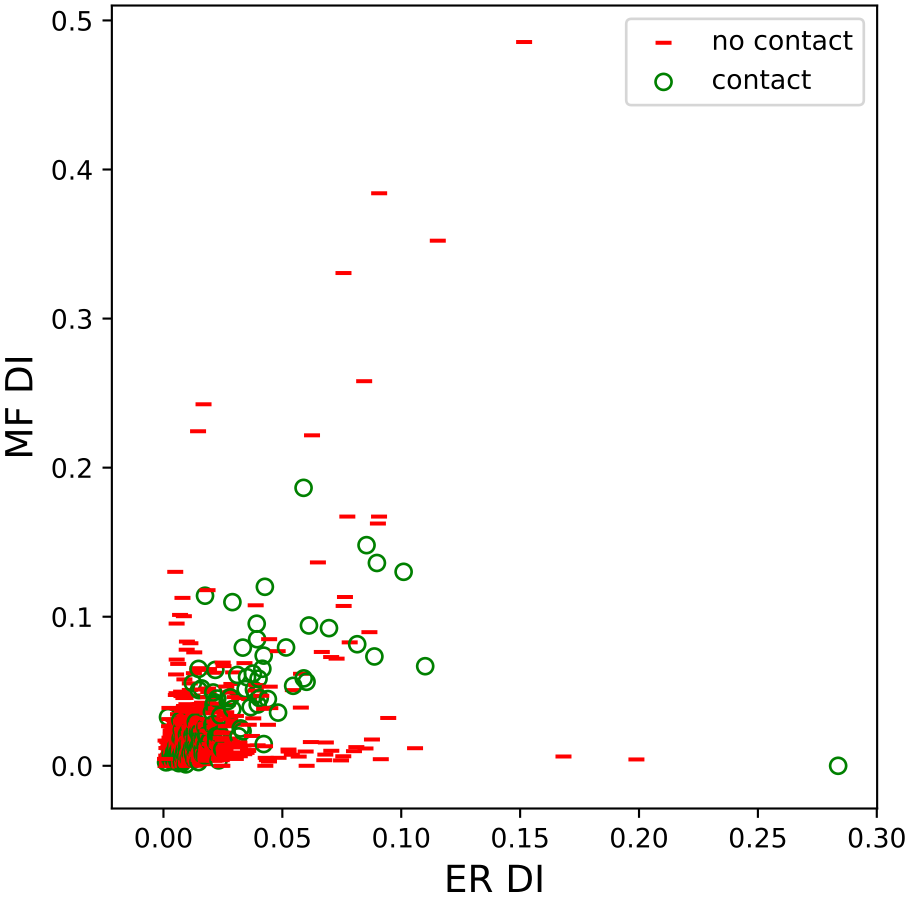
no contact
contact
MF DI
ER DI
0.0
0.1
0.2
0.3
0.4
0.5
0.00
0.05
0.10
0.15
0.20
0.25
0.30

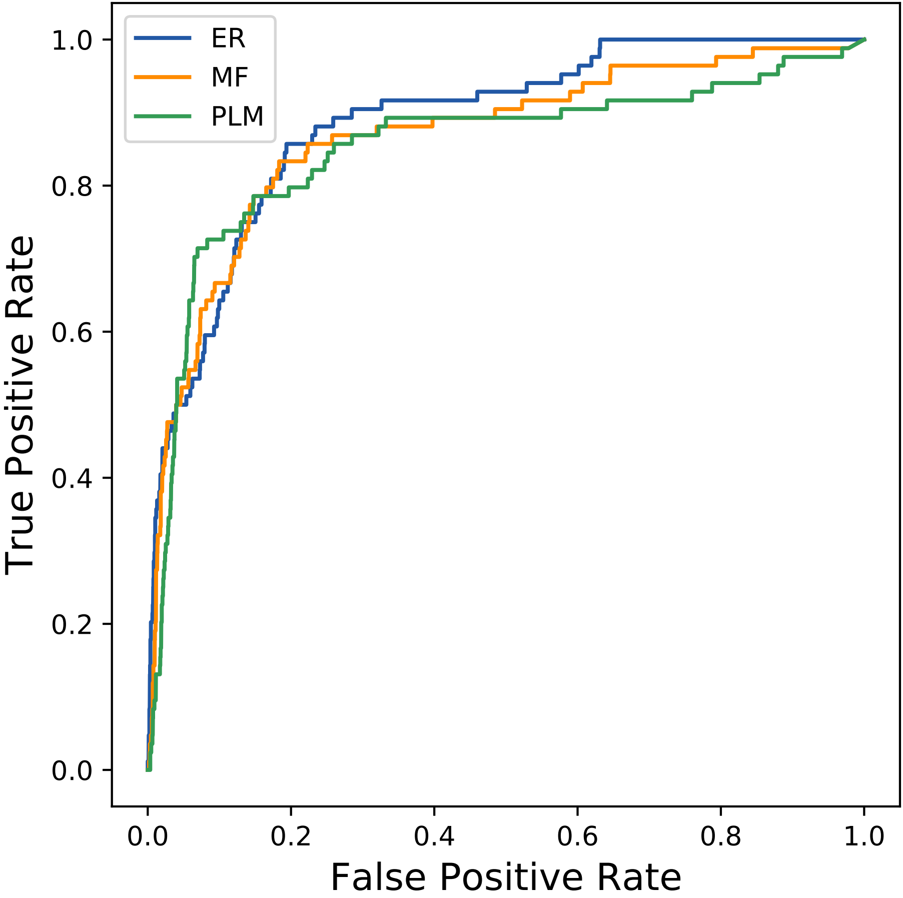

ER
MF
PLM
True Positive Rate
False Positive Rate
0.0
0.2
0.4
0.6
0.8
1.0
