## Supplemental Contact Maps for "Protein Structure Prediction with Expectation Reflection": 10315seq_70col_2q4o_PF03641_plots.pdf

Method: ER
Maximum PDB contact distance : 10.0 Angstrom
Minimum residue chain distance: 5.0 residues
Fraction of true positives : 0.8

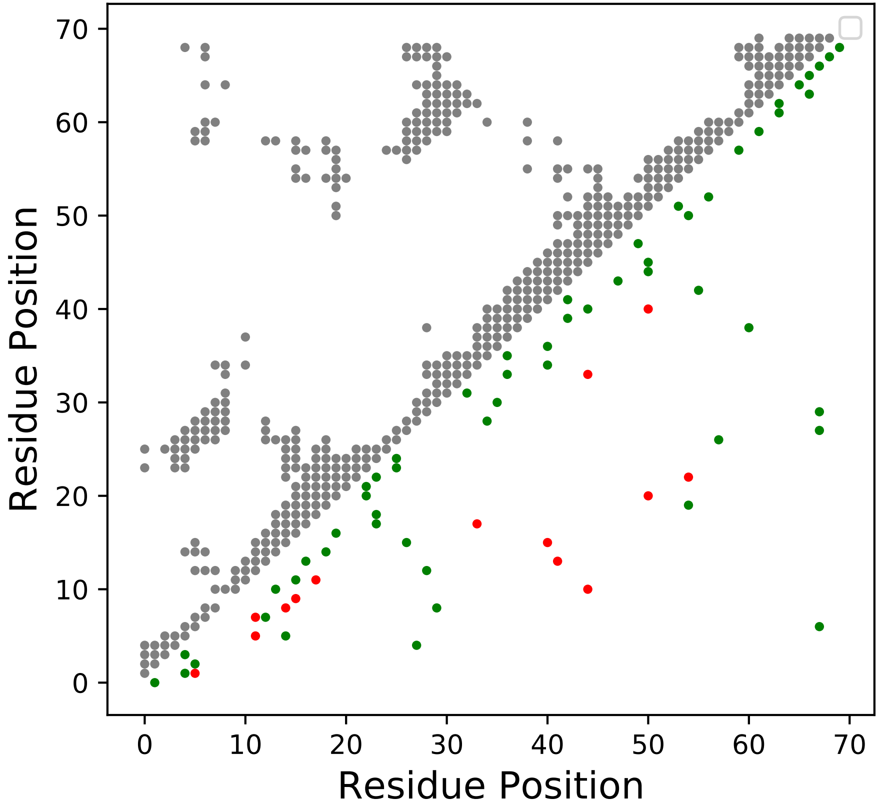

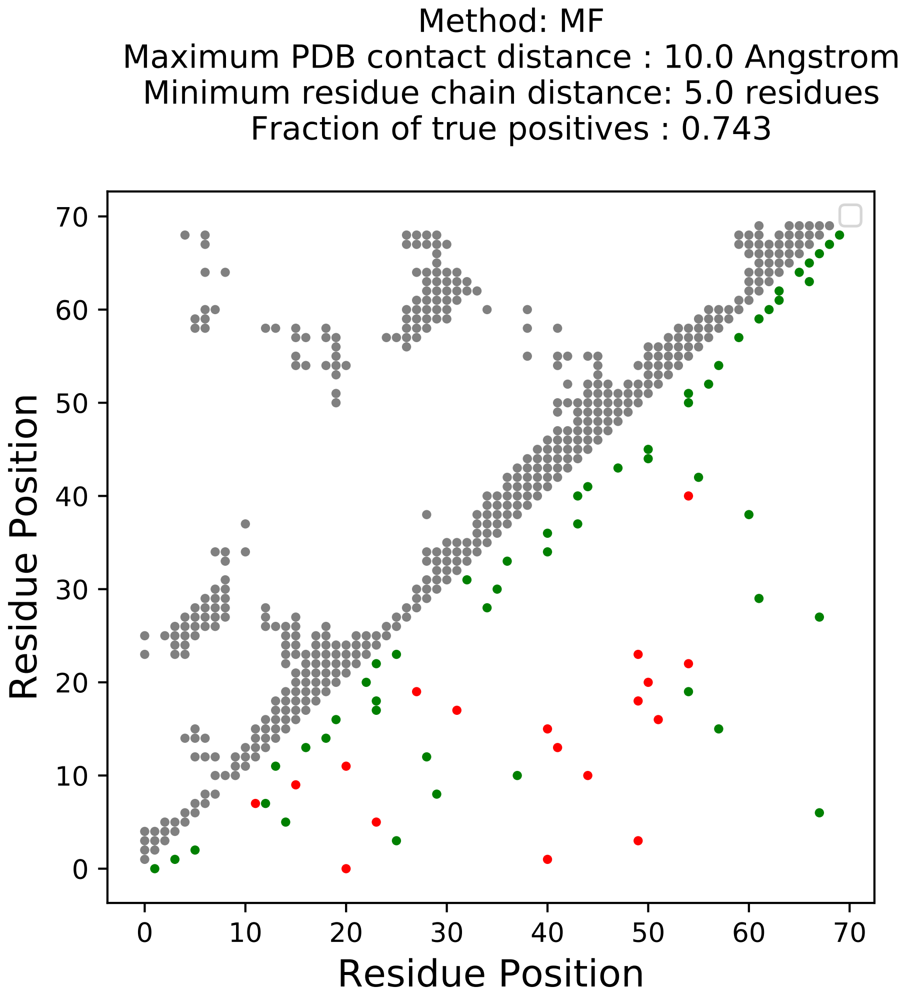

Method: MF
Maximum PDB contact distance : 10.0 Angstrom
Minimum residue chain distance: 5.0 residues
Fraction of true positives : 0.743
Residue Position
Residue Position
0
10
20
30
40
50
60
70

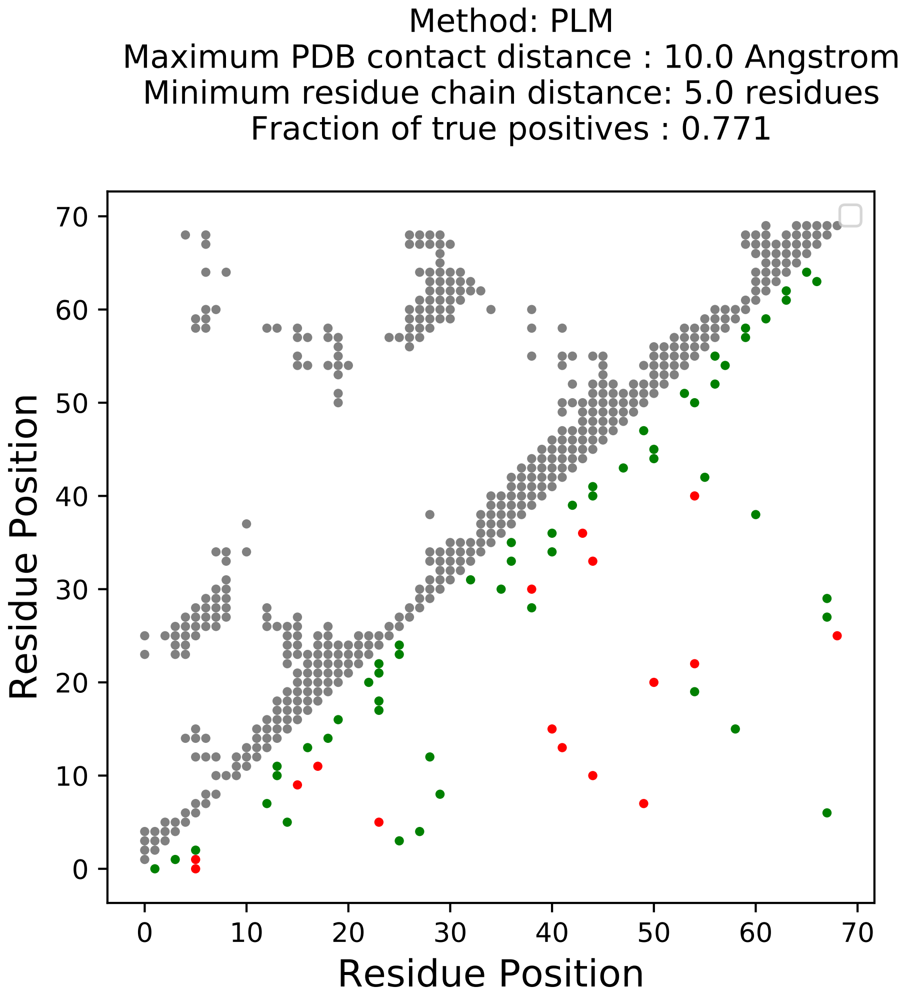
Method: PLM
Maximum PDB contact distance : 10.0 Angstrom
Minimum residue chain distance: 5.0 residues
Fraction of true positives : 0.771
Residue Position
Residue Position
0
10
20
30
40
50
60
70
0
10
20
30
40
50
60
70

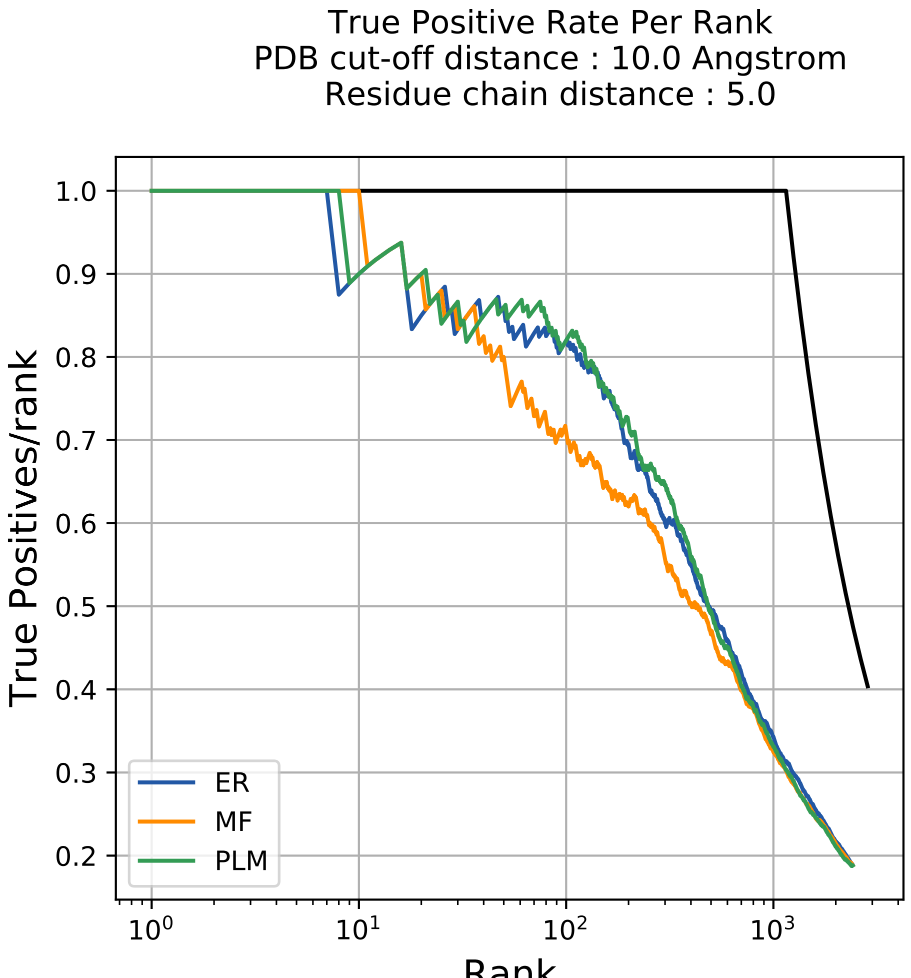
True Positive Rate Per Rank
PDB cut-off distance : 10.0 Angstrom
Residue chain distance : 5.0
True Positives/rank
Rank
1.0
0.9
0.8
0.7
0.6
0.5
0.4
0.3
0.2
$10^0$
$10^1$
$10^2$
$10^3$
ER
MF
PLM

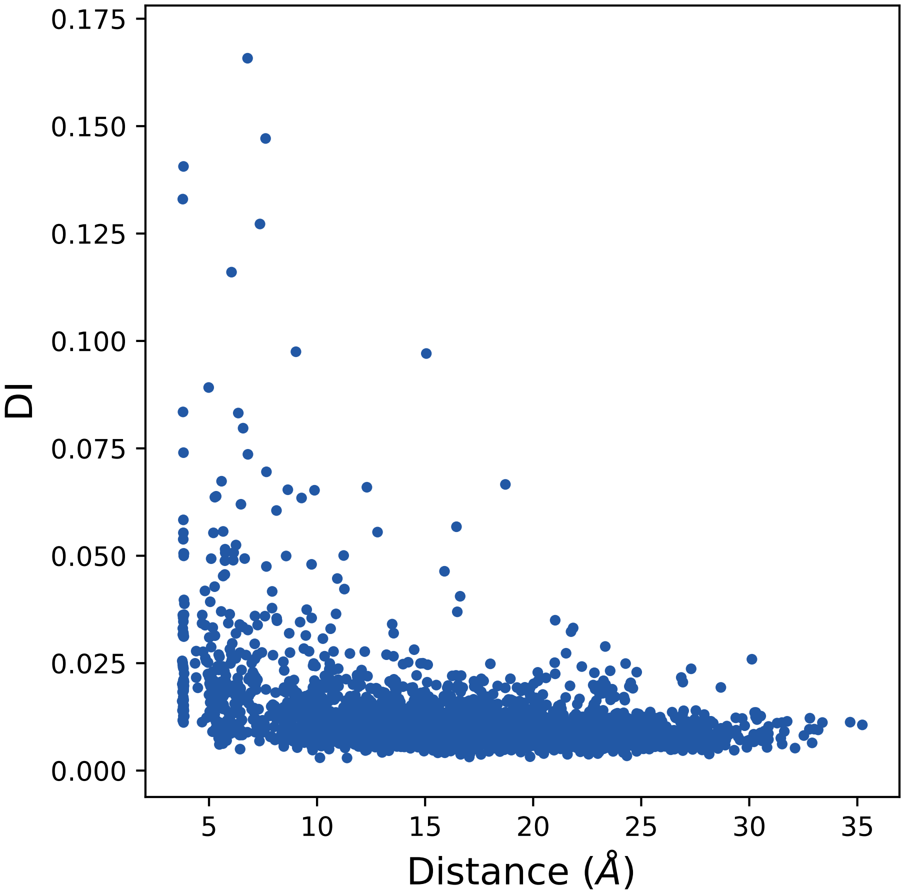

0.175
0.150
0.125
0.100
0.075
0.050
0.025
0.000
DI
5
10
15
20
25
30
35
Distance (Å)

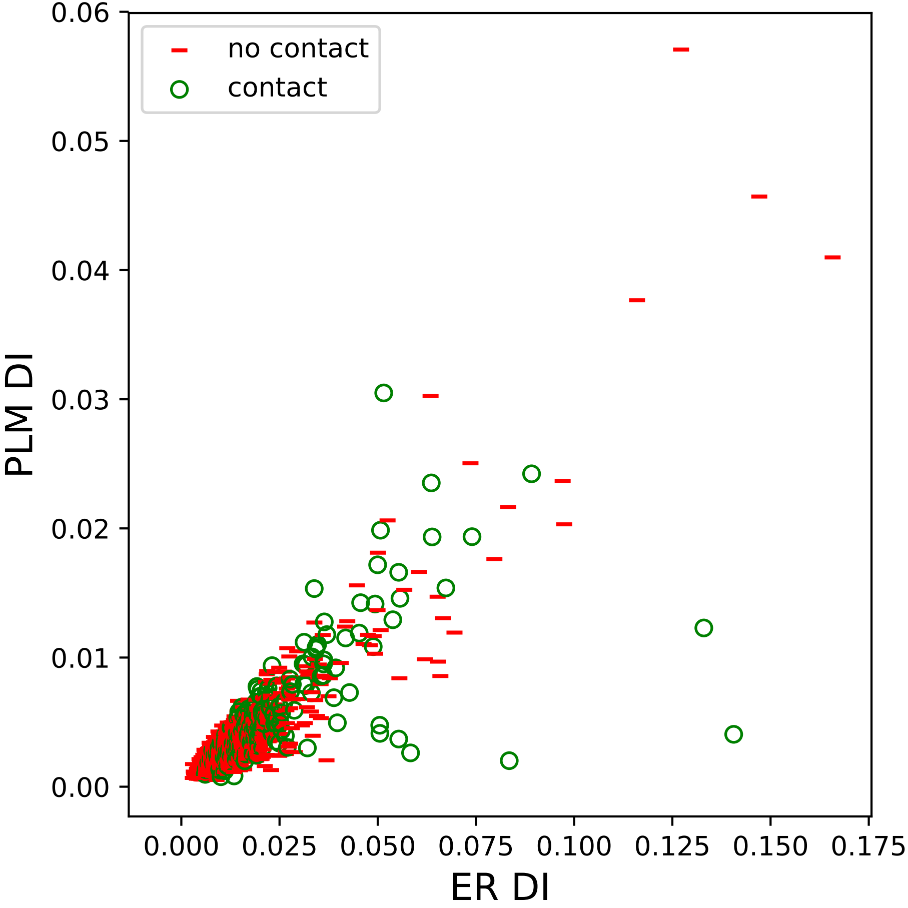

no contact
contact
PLM DI
ER DI
0.00
0.01
0.02
0.03
0.04
0.05
0.06
0.000
0.025
0.050
0.075
0.100
0.125
0.150
0.175

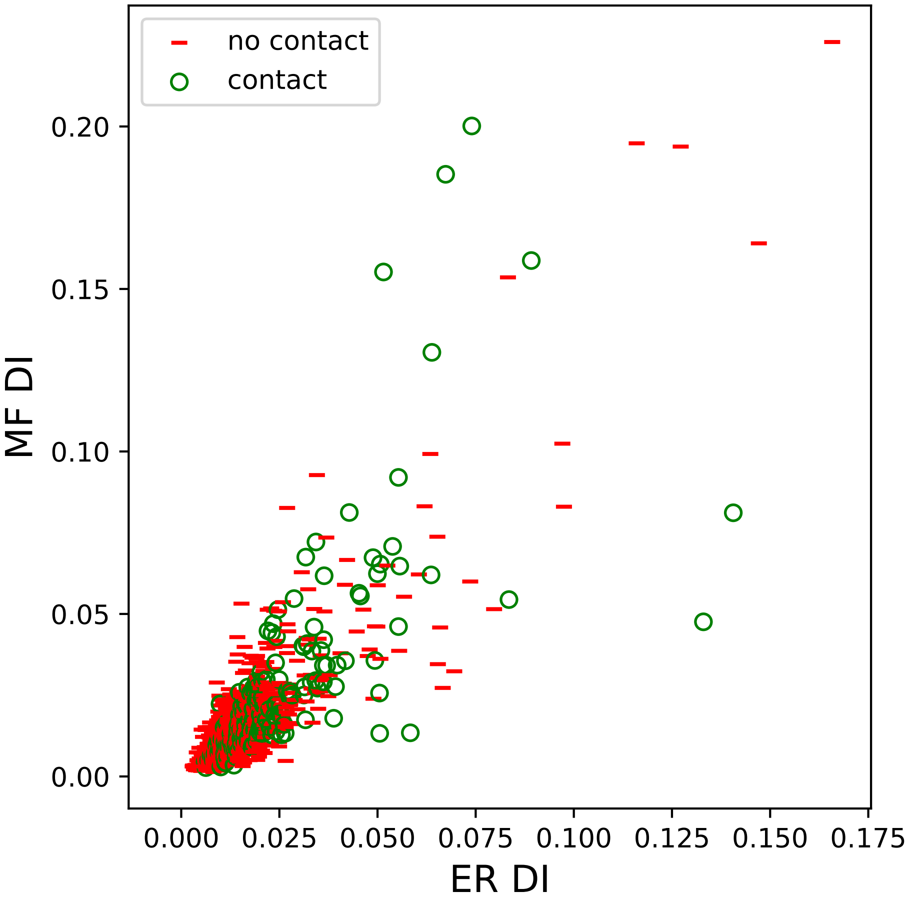

no contact
contact
0.20
0.15
0.10
0.05
0.00
MF DI
0.000
0.025
0.050
0.075
0.100
0.125
0.150
0.175
ER DI

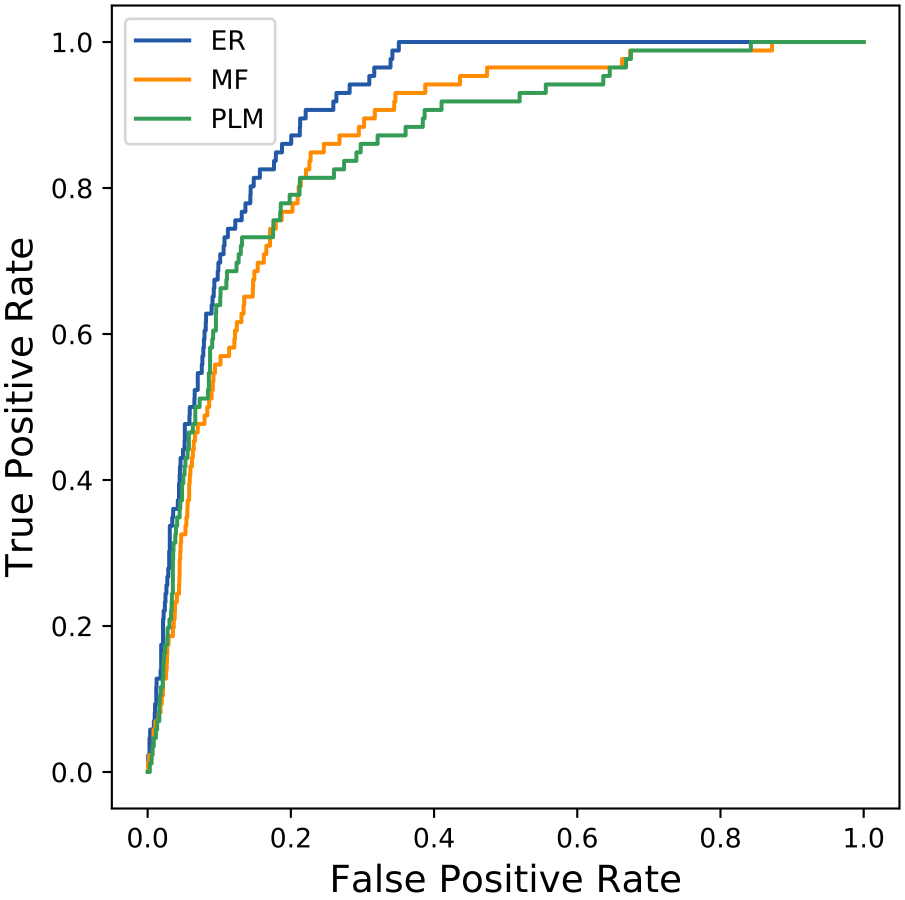
ER
MF
PLM
1.0
0.8
0.6
0.4
0.2
0.0
0.0
0.2
0.4
0.6
0.8
1.0
True Positive Rate
False Positive Rate
