## Supplemental Contact Maps for "Protein Structure Prediction with Expectation Reflection": 10968seq_111col_2qrj_PF05222_plots.pdf

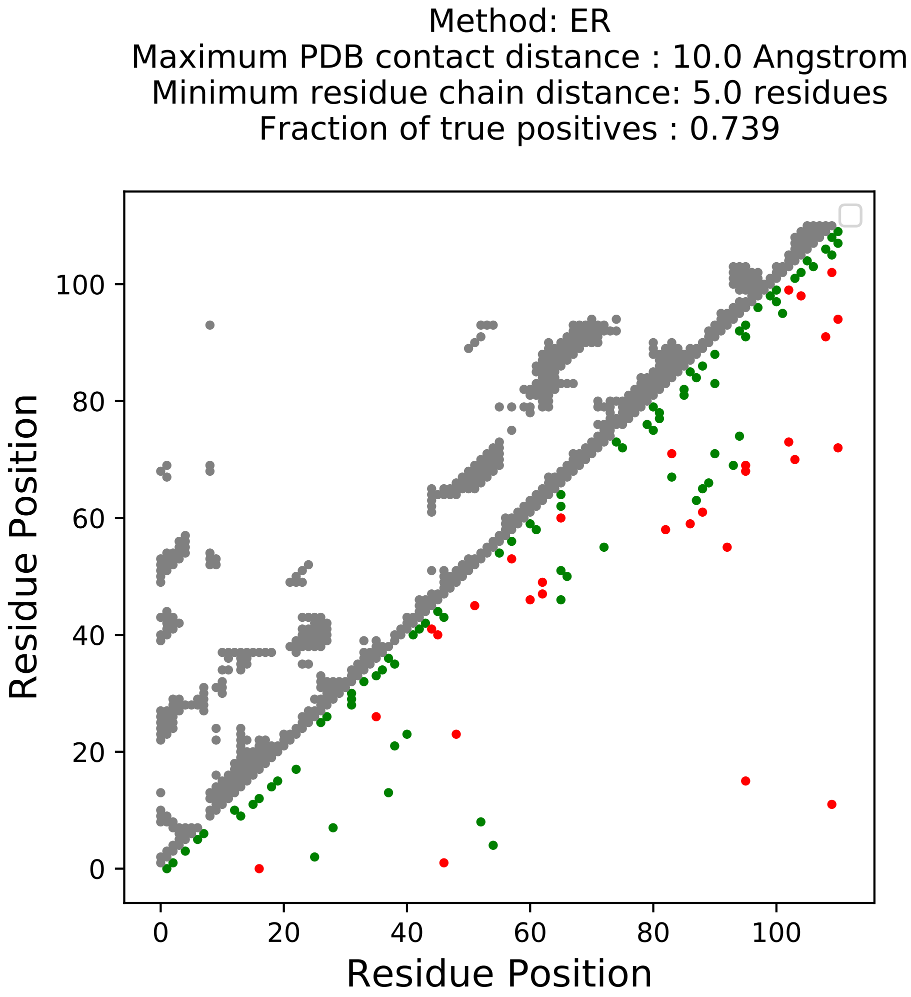
Method: ER
Maximum PDB contact distance : 10.0 Angstrom
Minimum residue chain distance: 5.0 residues
Fraction of true positives : 0.739
Residue Position
Residue Position
0
20
40
60
80
100

Method: MF
Maximum PDB contact distance : 10.0 Angstrom
Minimum residue chain distance: 5.0 residues
Fraction of true positives : 0.658

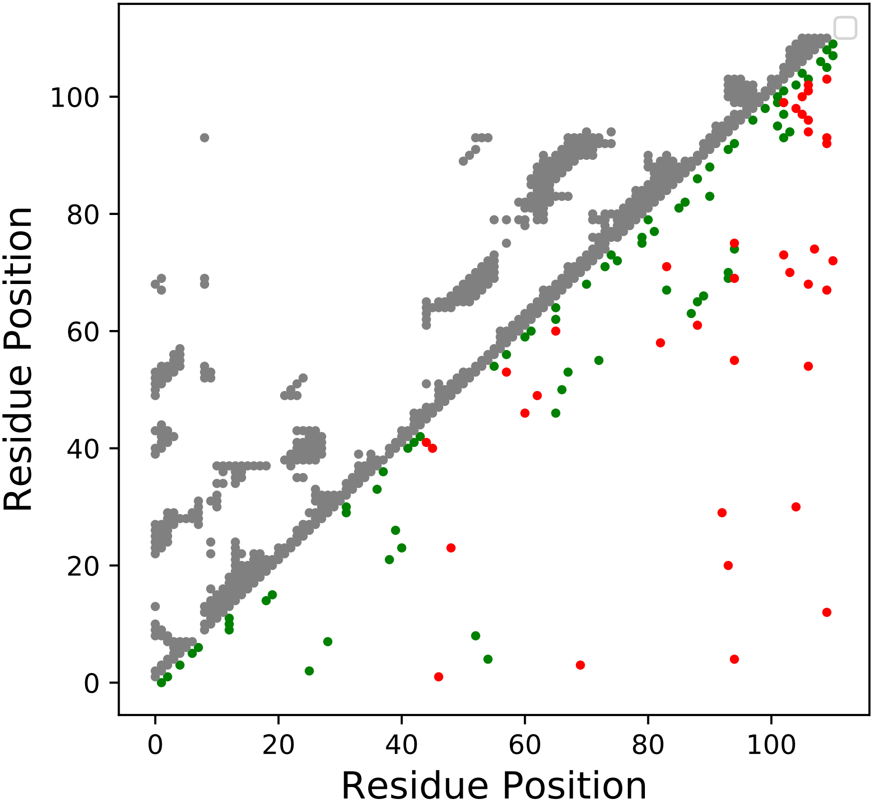

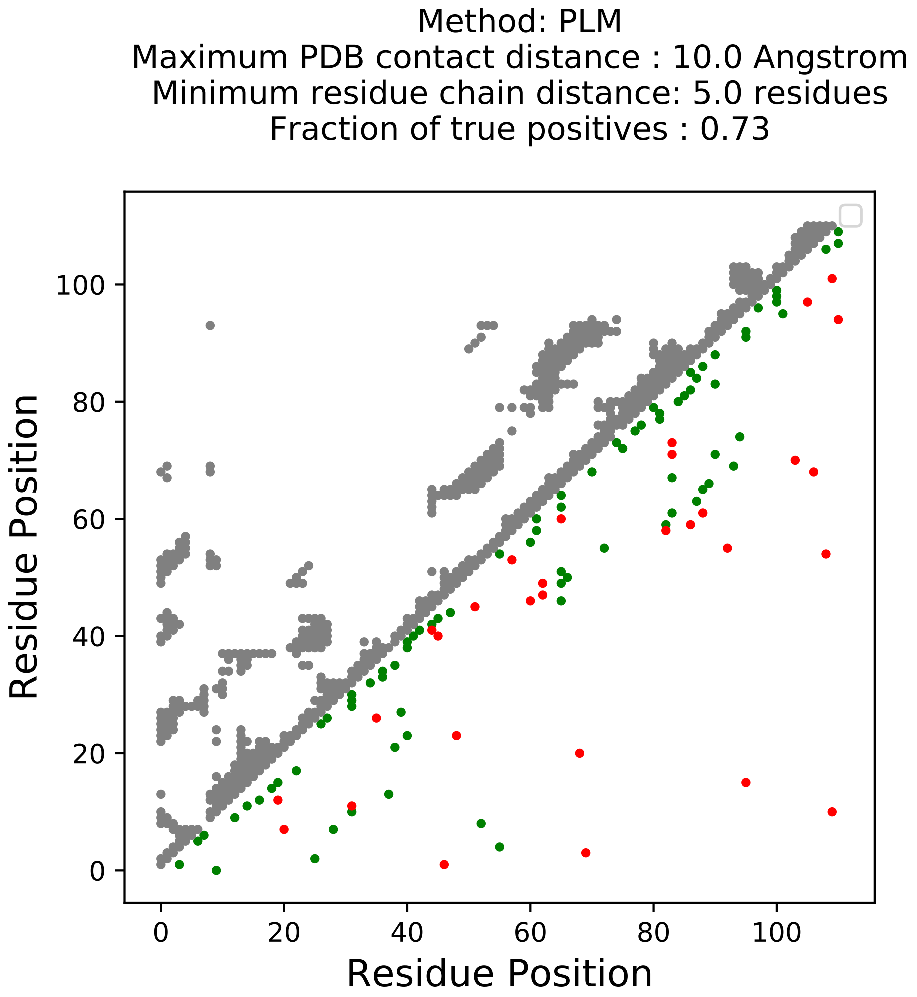
Method: PLM
Maximum PDB contact distance : 10.0 Angstrom
Minimum residue chain distance: 5.0 residues
Fraction of true positives : 0.73
Residue Position
Residue Position
0
20
40
60
80
100
0
20
40
60
80
100

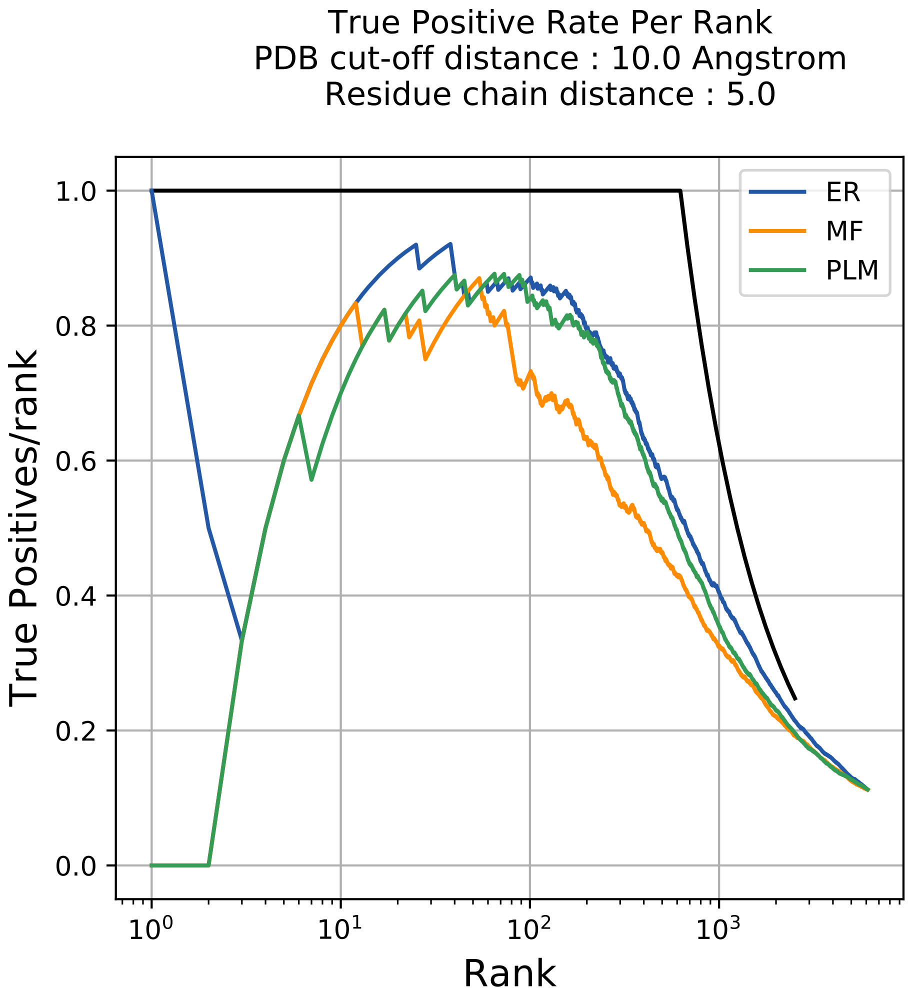
True Positive Rate Per Rank
PDB cut-off distance : 10.0 Angstrom
Residue chain distance : 5.0
True Positives/rank
Rank
1.0
0.8
0.6
0.4
0.2
0.0
$10^0$
$10^1$
$10^2$
$10^3$
ER
MF
PLM

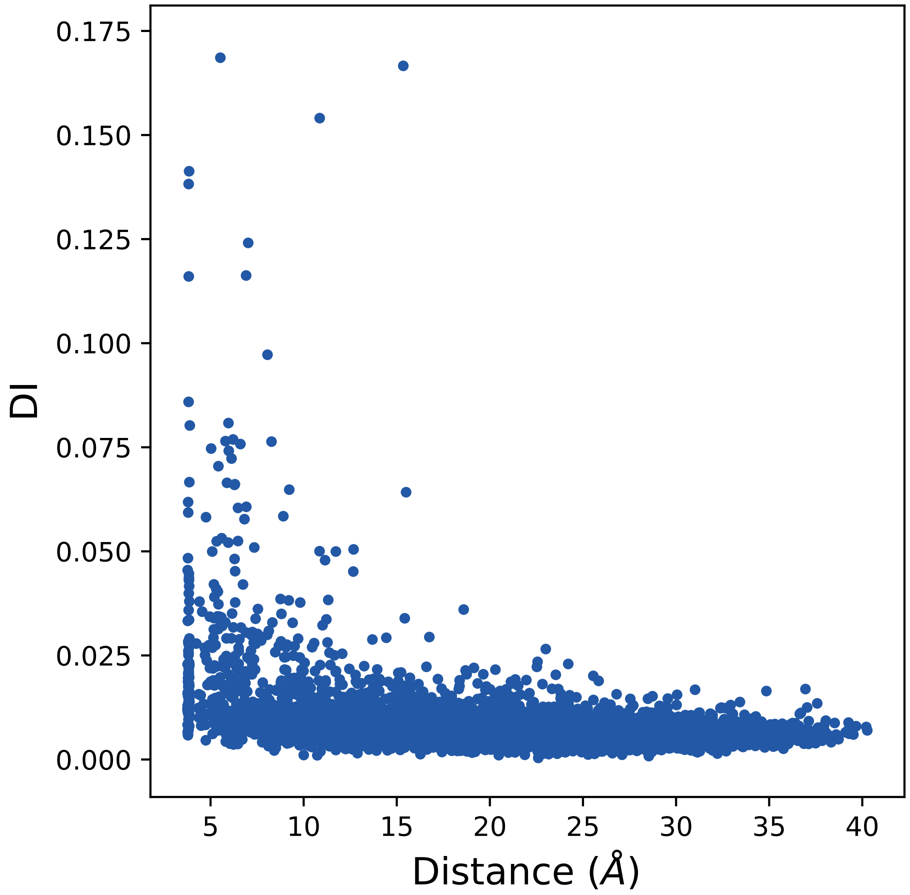

0.175
0.150
0.125
0.100
0.075
0.050
0.025
0.000
DI
5
10
15
20
25
30
35
40
Distance (Å)

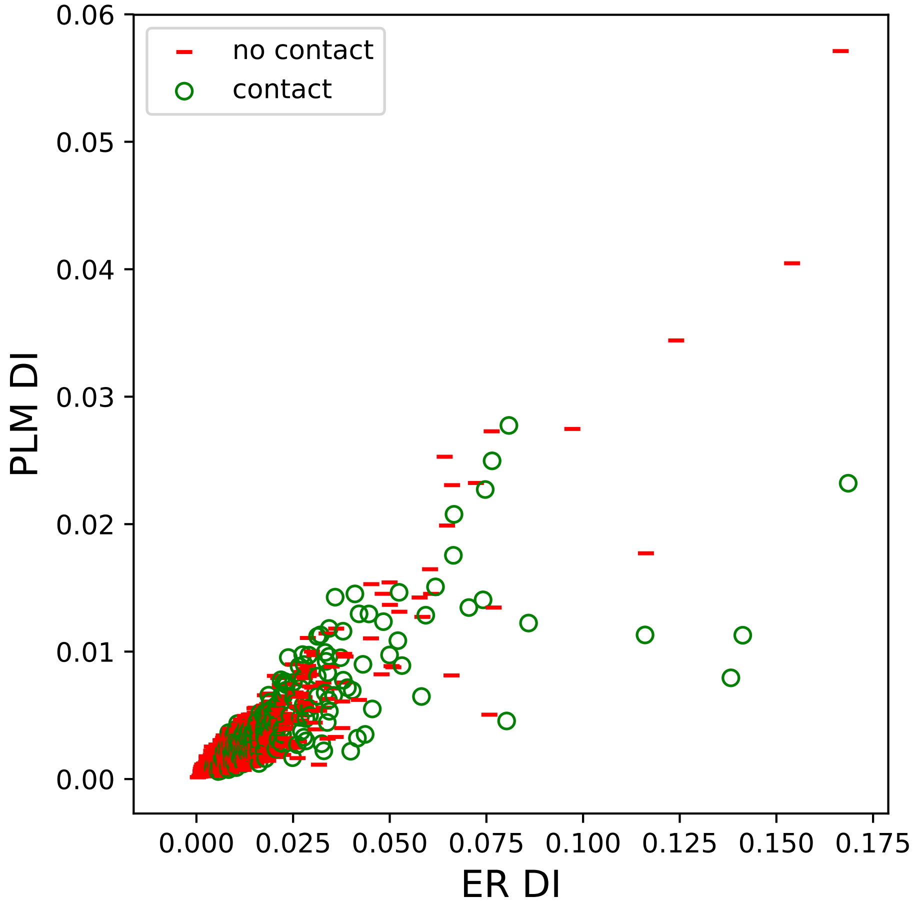

no contact
contact
PLM DI
ER DI
0.00
0.01
0.02
0.03
0.04
0.05
0.06
0.000
0.025
0.050
0.075
0.100
0.125
0.150
0.175

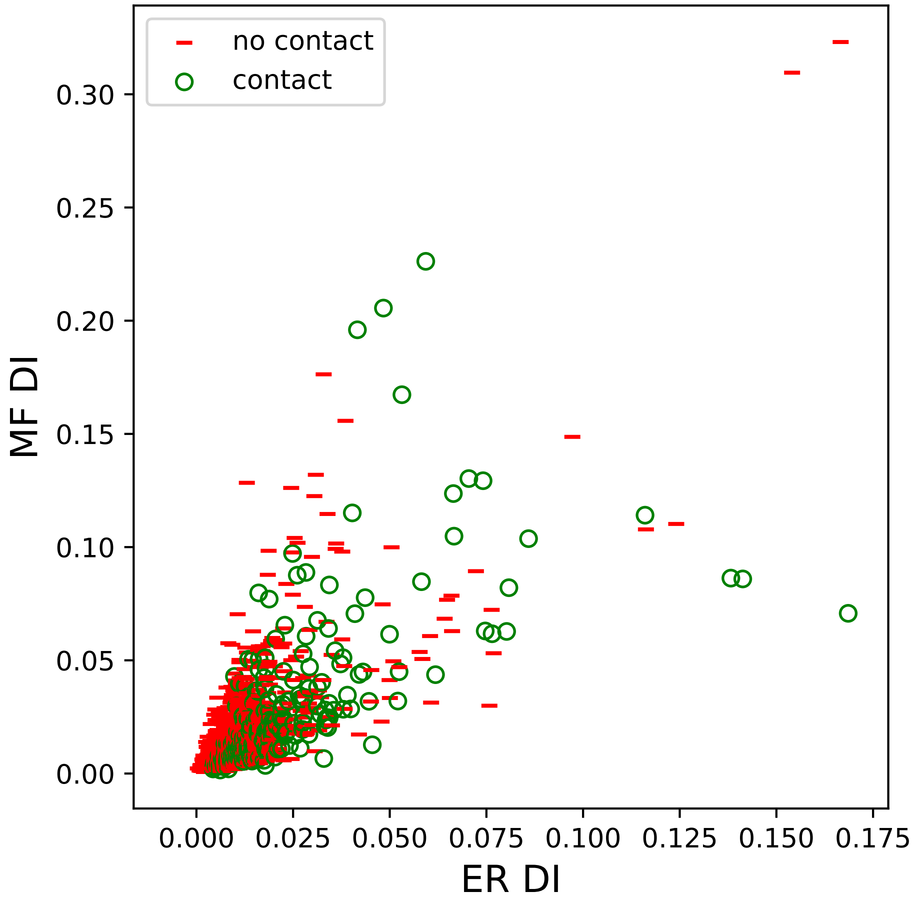
no contact
contact
MF DI
ER DI
0.00
0.05
0.10
0.15
0.20
0.25
0.30
0.000
0.025
0.050
0.075
0.100
0.125
0.150
0.175

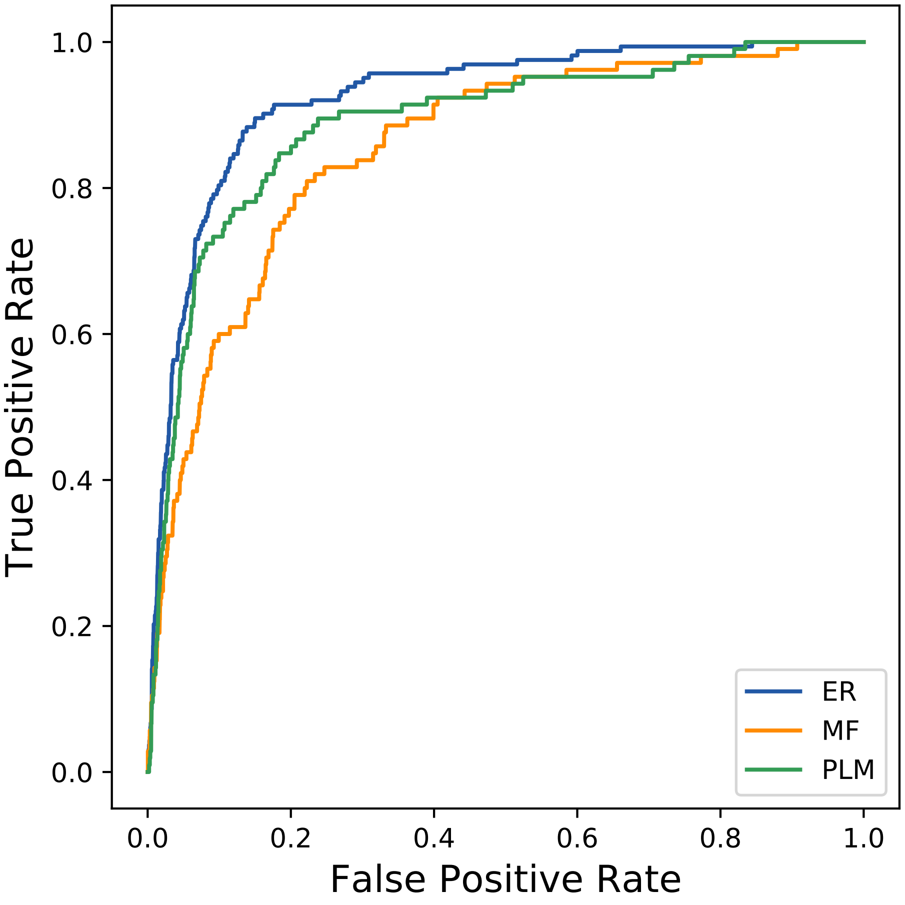

True Positive Rate
1.0
0.8
0.6
0.4
0.2
0.0
0.0
0.2
0.4
0.6
0.8
1.0
False Positive Rate
ER
MF
PLM
