## Supplemental Contact Maps for "Protein Structure Prediction with Expectation Reflection": 11117seq_76col_3qfi_PF03816_plots.pdf

Method: ER
Maximum PDB contact distance : 10.0 Angstrom
Minimum residue chain distance: 5.0 residues
Fraction of true positives : 0.789

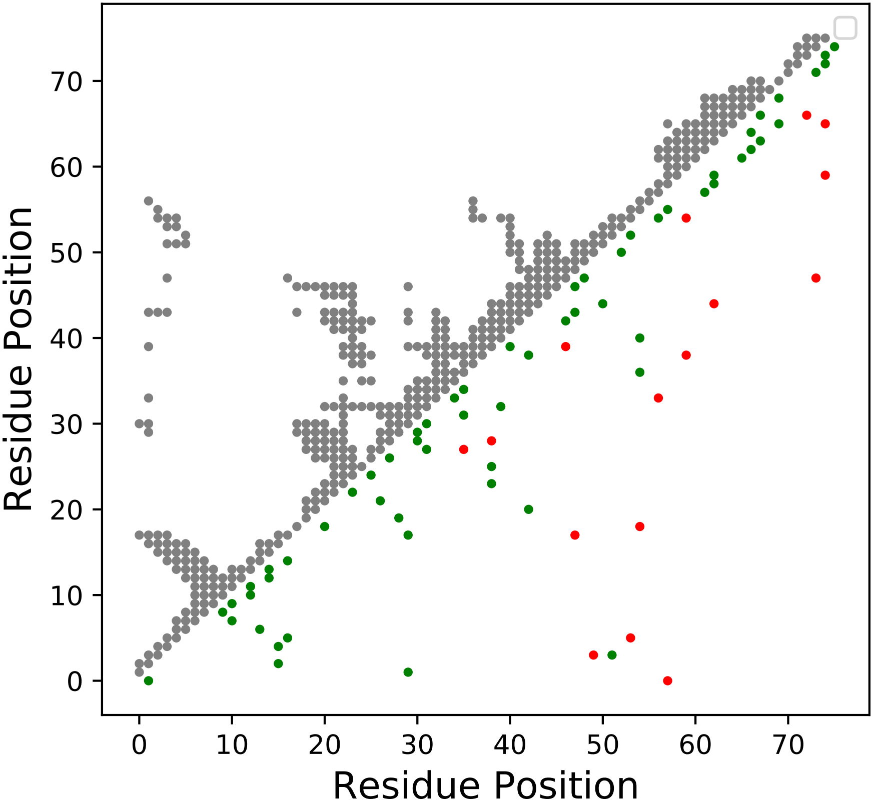

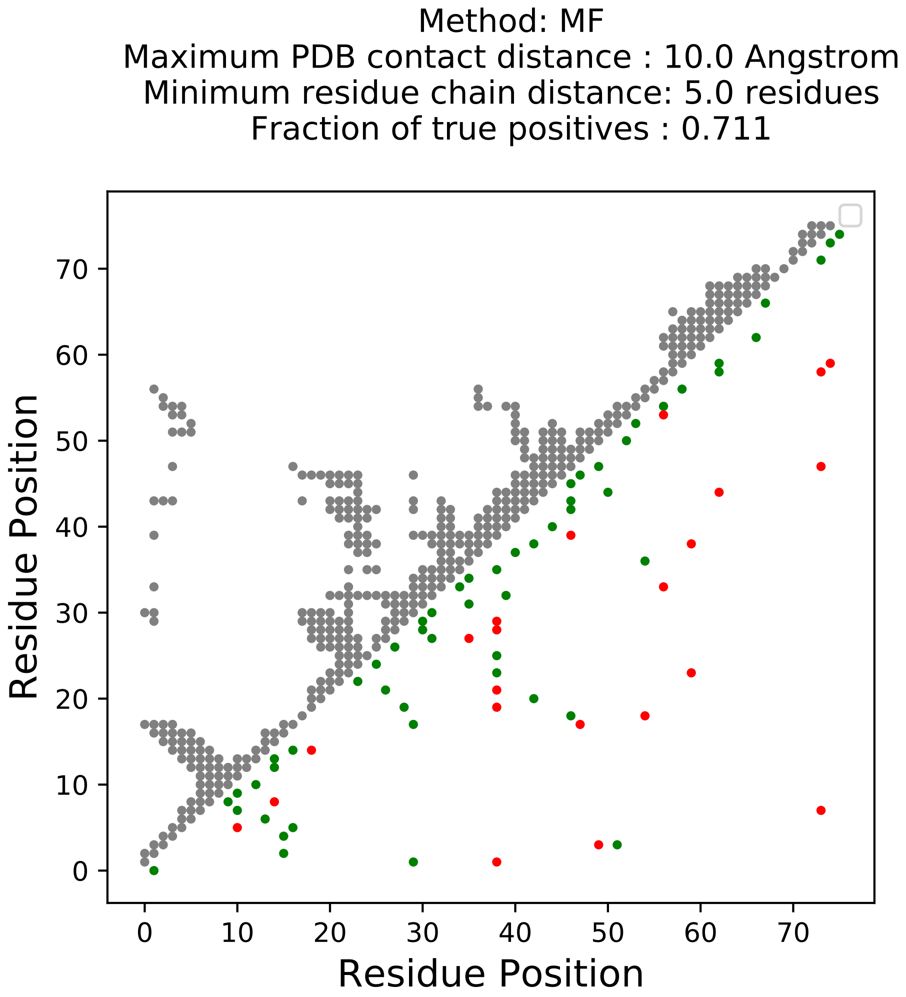

Method: MF
Maximum PDB contact distance : 10.0 Angstrom
Minimum residue chain distance: 5.0 residues
Fraction of true positives : 0.711
Residue Position
Residue Position
0
10
20
30
40
50
60
70

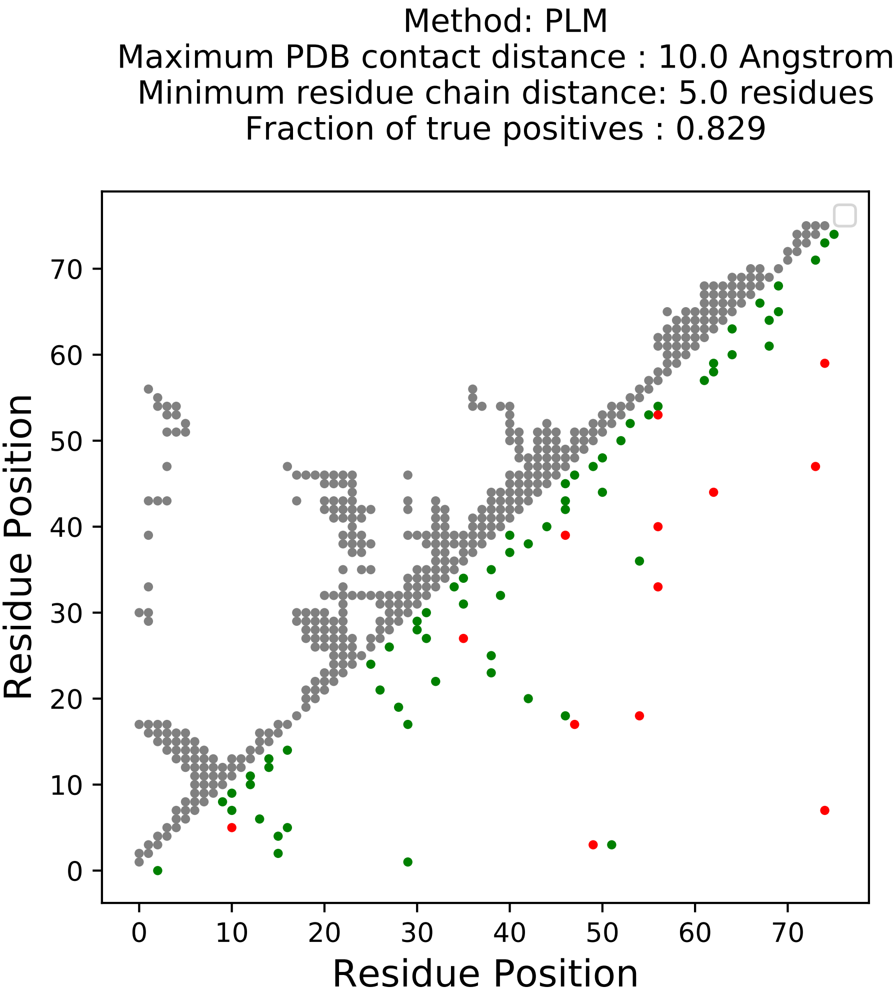
Method: PLM
Maximum PDB contact distance : 10.0 Angstrom
Minimum residue chain distance: 5.0 residues
Fraction of true positives : 0.829
Residue Position
Residue Position
0
10
20
30
40
50
60
70

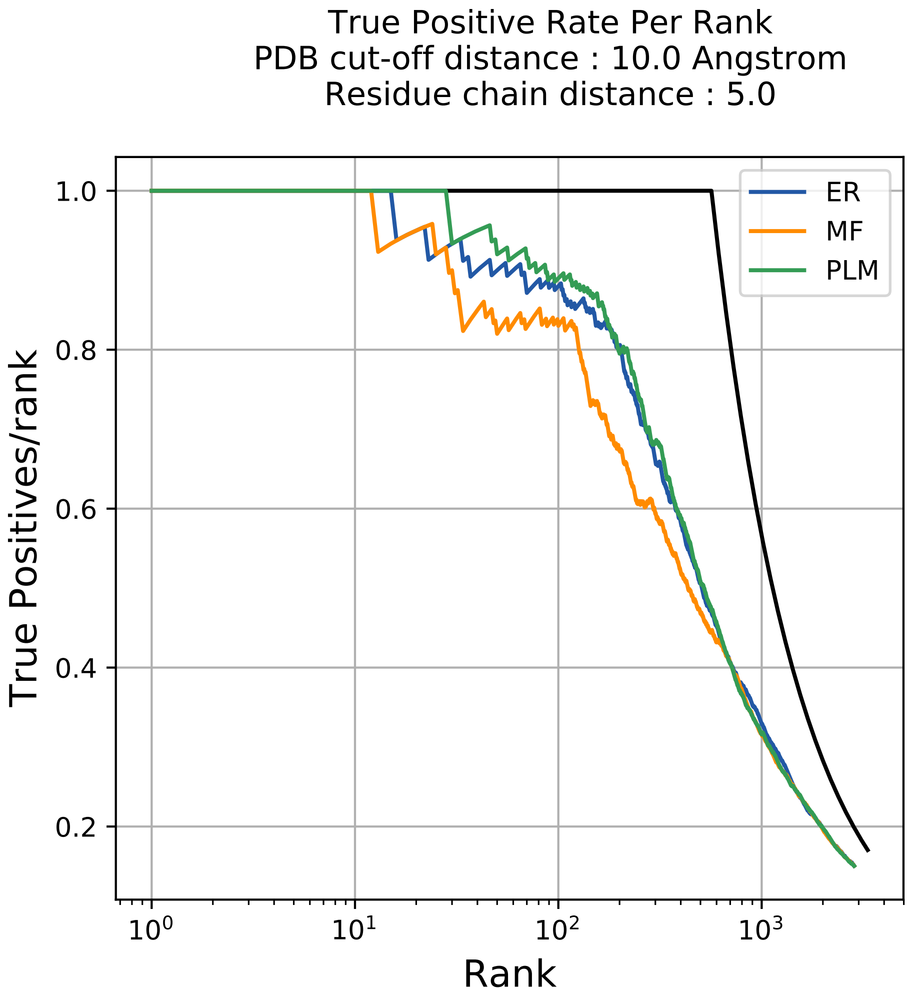
True Positive Rate Per Rank
PDB cut-off distance : 10.0 Angstrom
Residue chain distance : 5.0
ER
MF
PLM
1.0
0.8
0.6
0.4
0.2
True Positives/rank
$10^0$
$10^1$
$10^2$
$10^3$
Rank

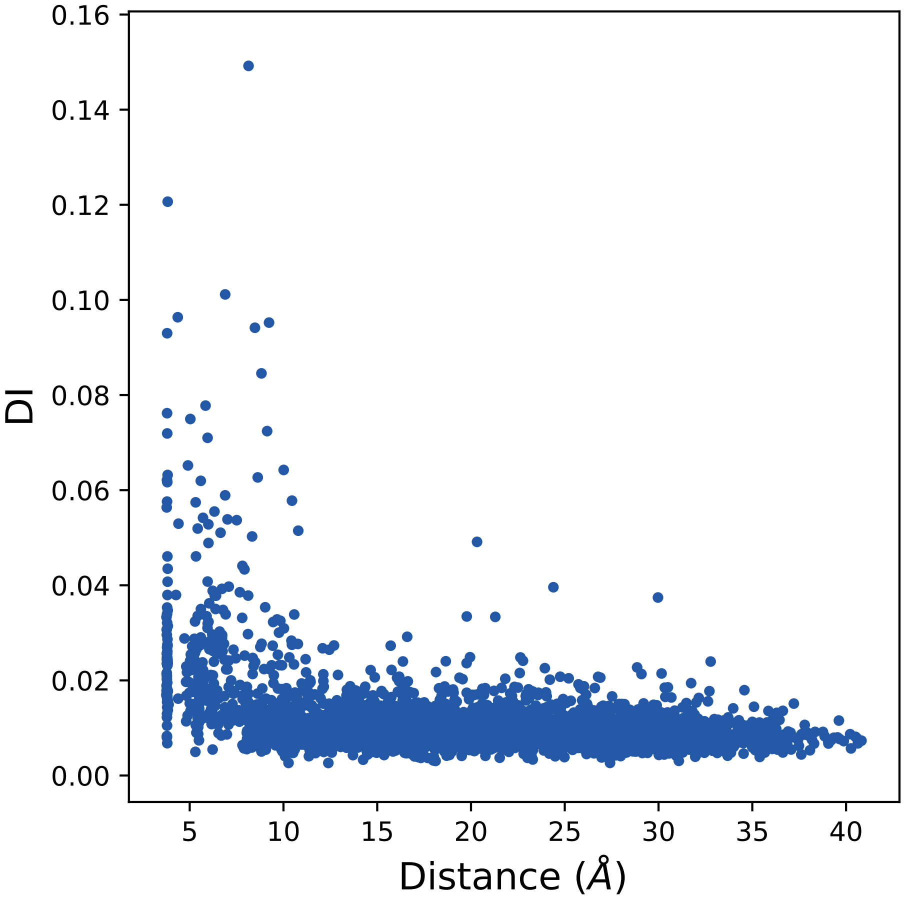
0.16
0.14
0.12
0.10
0.08
0.06
0.04
0.02
0.00
DI
5
10
15
20
25
30
35
40
Distance ($\AA$)

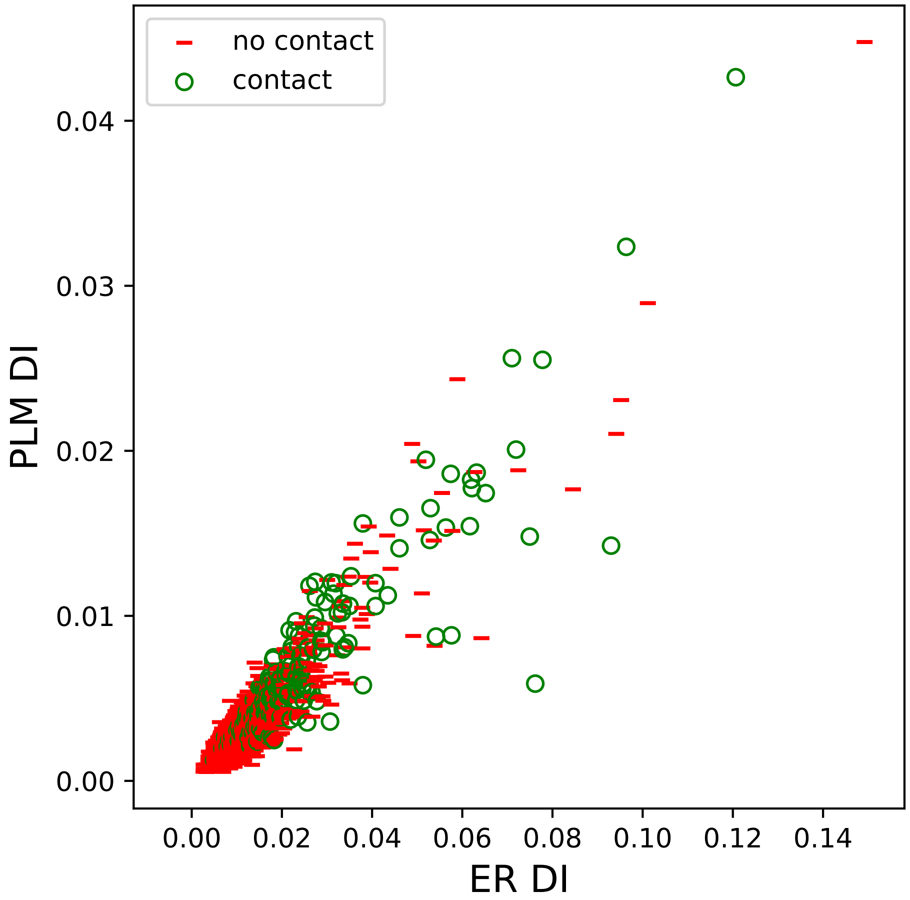
no contact
contact
0.00
0.01
0.02
0.03
0.04
PLM DI
0.00
0.02
0.04
0.06
0.08
0.10
0.12
0.14
ER DI

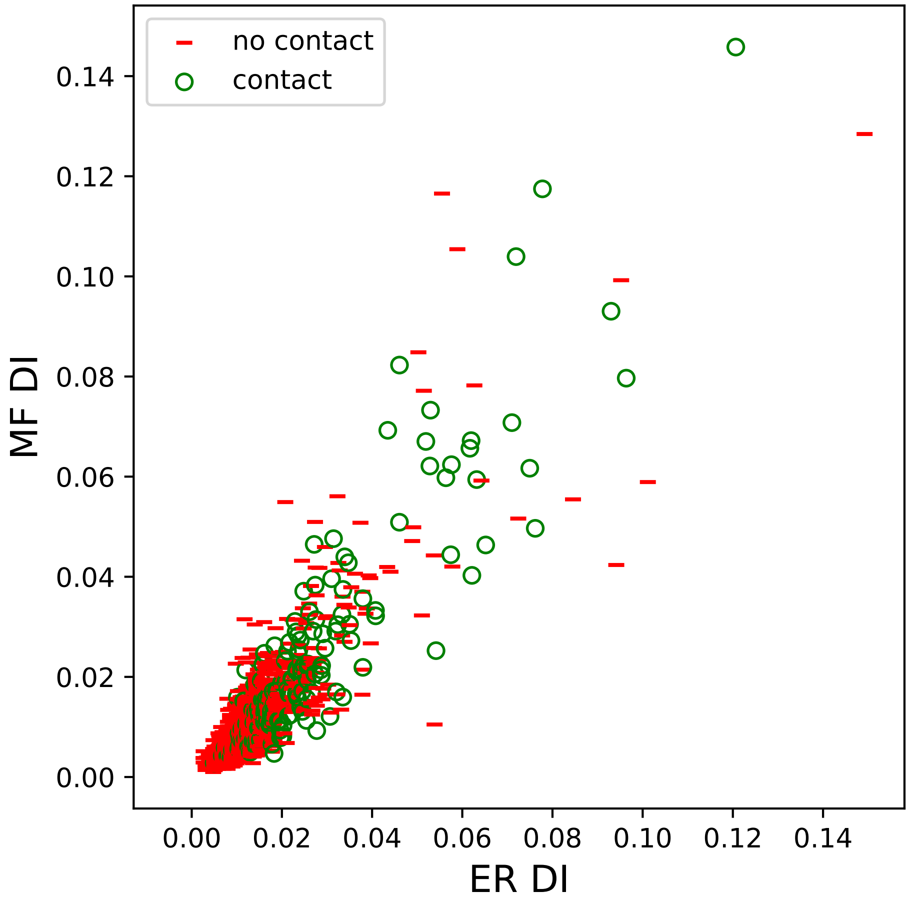
no contact
contact
MF DI
ER DI
0.00
0.02
0.04
0.06
0.08
0.10
0.12
0.14

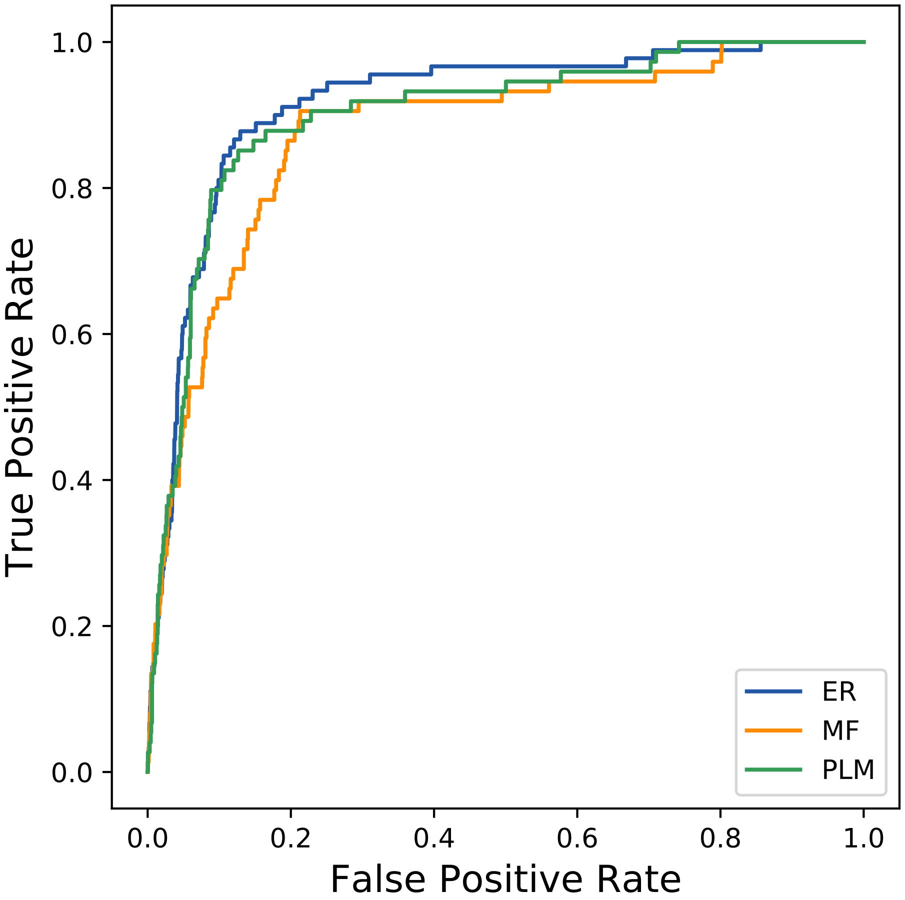
True Positive Rate
False Positive Rate
0.0
0.2
0.4
0.6
0.8
1.0
ER
MF
PLM
