## Supplemental Contact Maps for "Protein Structure Prediction with Expectation Reflection": 11526seq_217col_1r1o_PF00491_plots.pdf

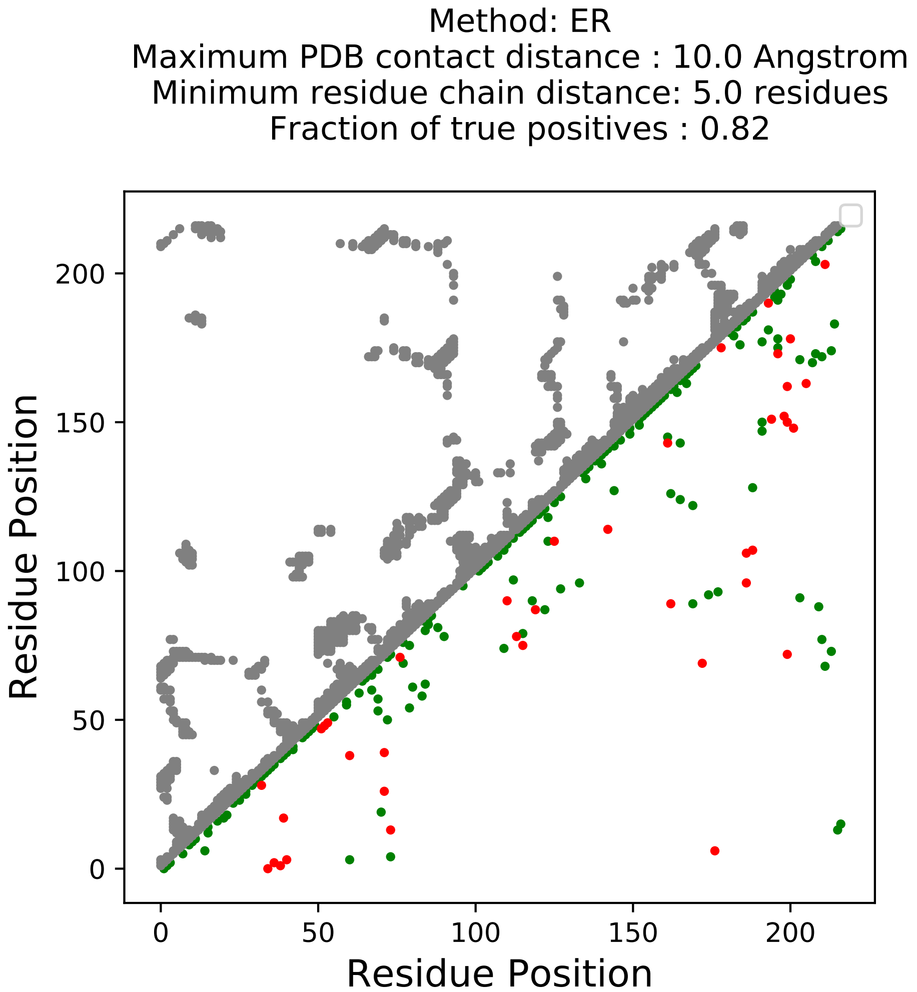
Method: ER
Maximum PDB contact distance : 10.0 Angstrom
Minimum residue chain distance: 5.0 residues
Fraction of true positives : 0.82
Residue Position
Residue Position
0
50
100
150
200
0
50
100
150
200

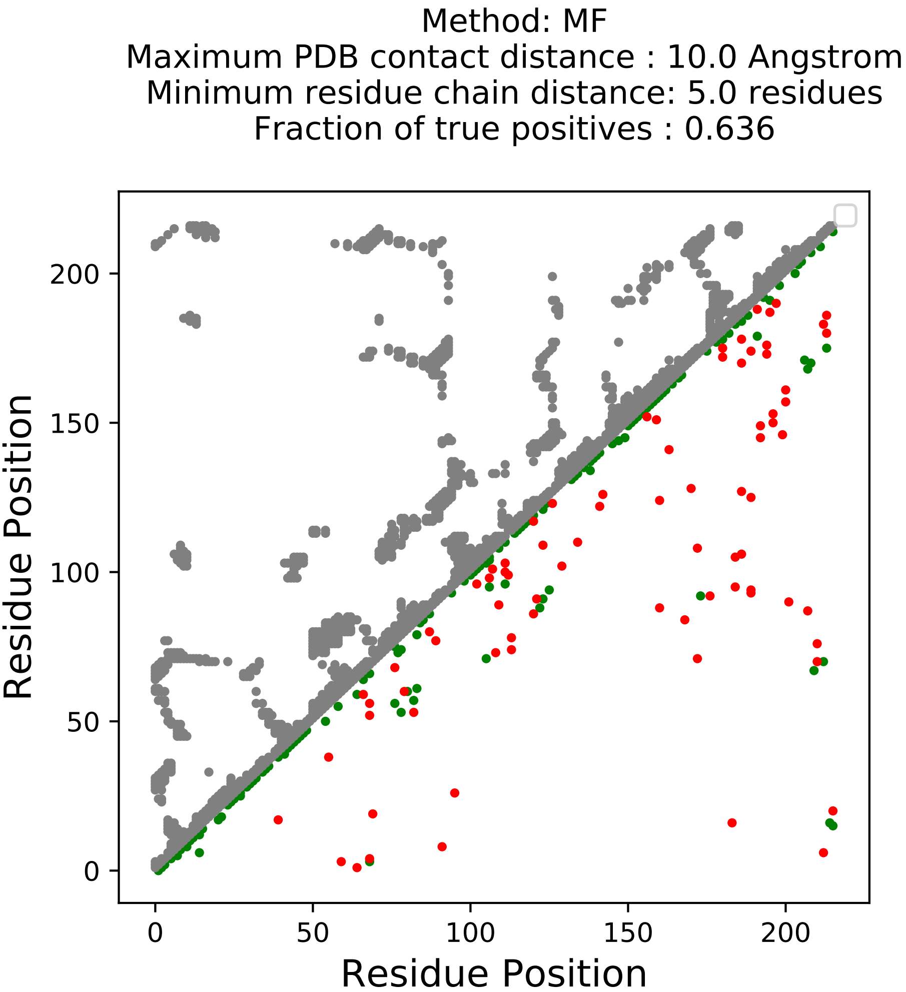
Method: MF
Maximum PDB contact distance : 10.0 Angstrom
Minimum residue chain distance: 5.0 residues
Fraction of true positives : 0.636
Residue Position
Residue Position
0
50
100
150
200
0
50
100
150
200

Method: PLM
Maximum PDB contact distance : 10.0 Angstrom
Minimum residue chain distance: 5.0 residues
Fraction of true positives : 0.618

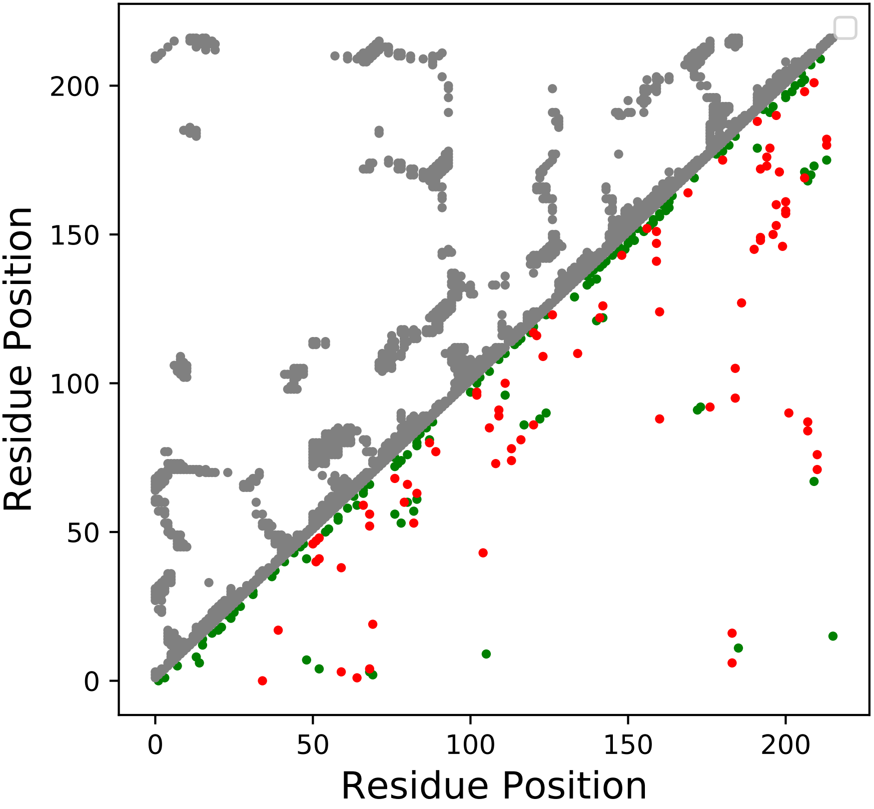

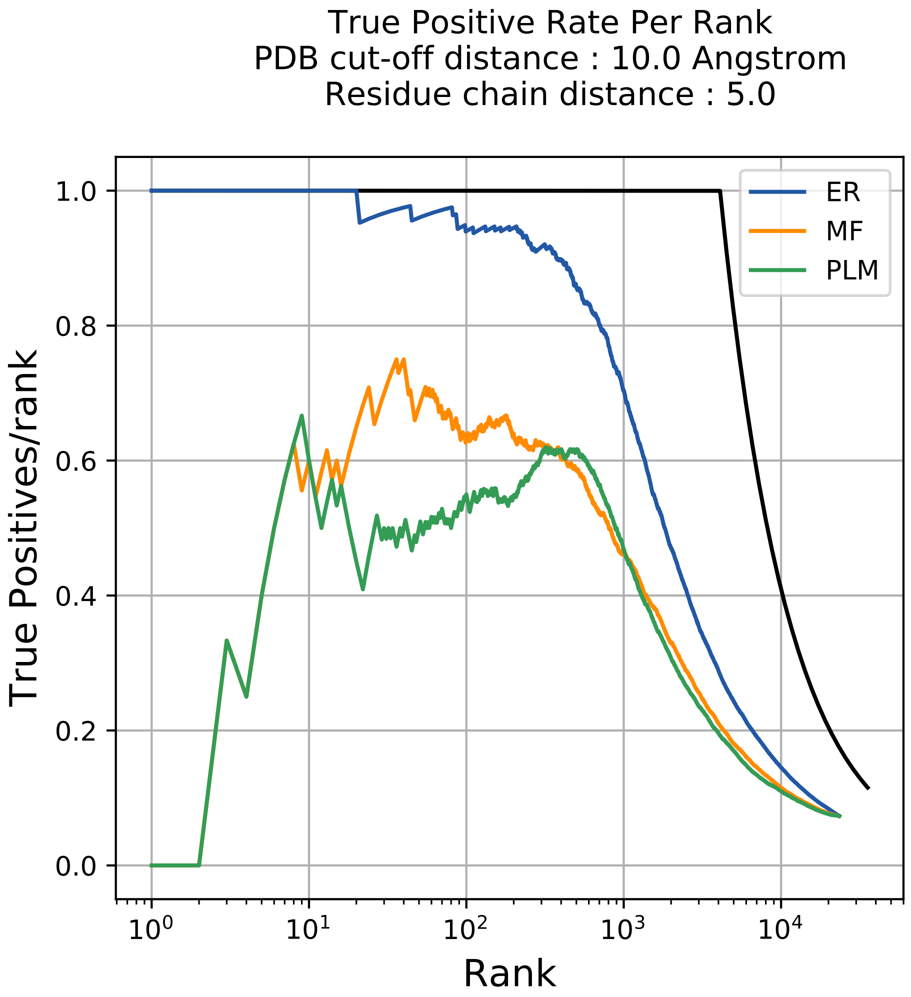
True Positive Rate Per Rank
PDB cut-off distance : 10.0 Angstrom
Residue chain distance : 5.0
ER
MF
PLM
True Positives/rank
Rank
1.0
0.8
0.6
0.4
0.2
0.0
$10^0$
$10^1$
$10^2$
$10^3$
$10^4$

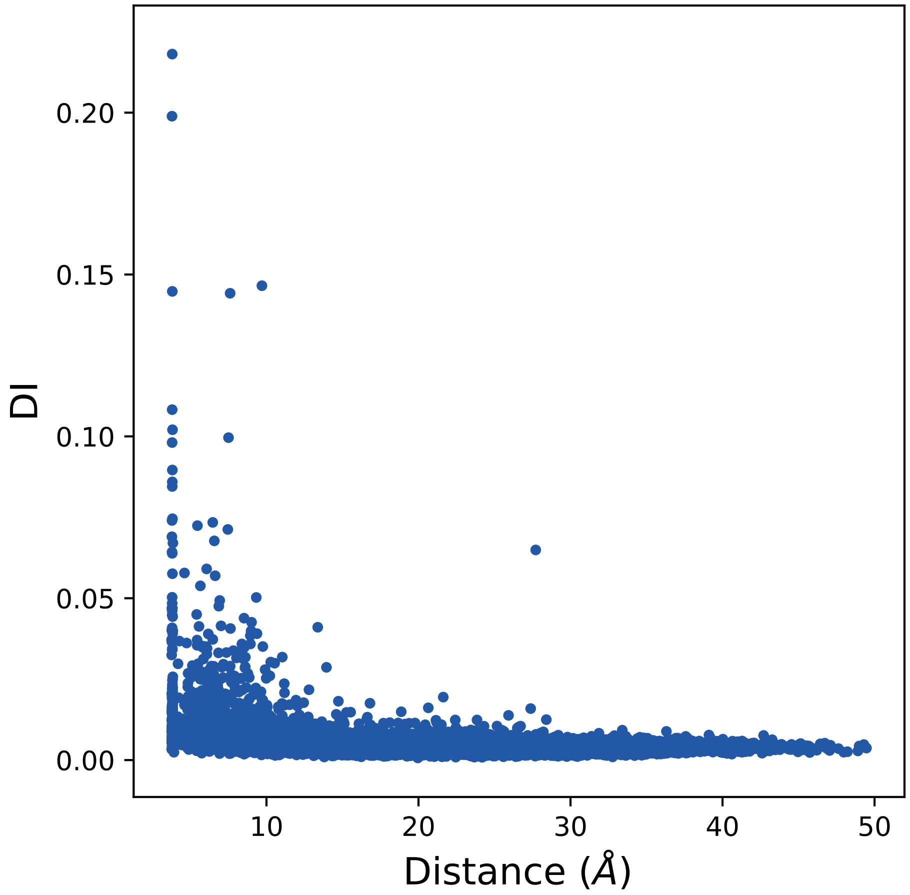
0.20
0.15
0.10
0.05
0.00
DI
10
20
30
40
50
Distance (Å)

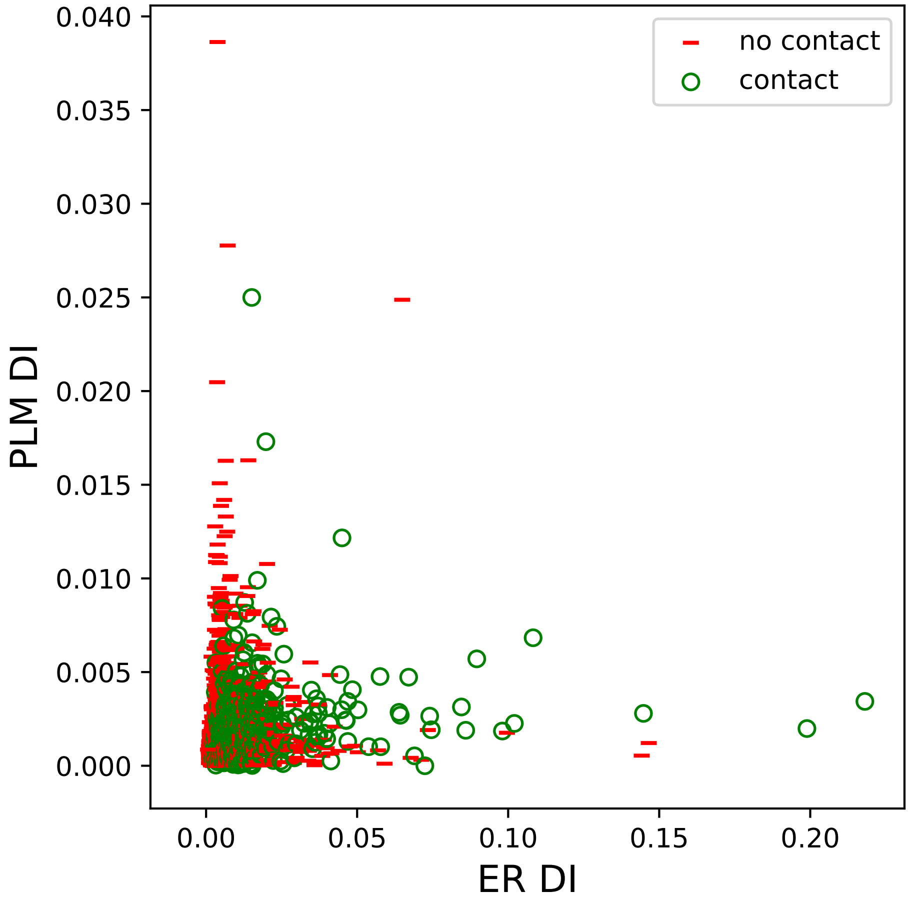
no contact
contact
PLM DI
ER DI
0.040
0.035
0.030
0.025
0.020
0.015
0.010
0.005
0.000
0.00
0.05
0.10
0.15
0.20

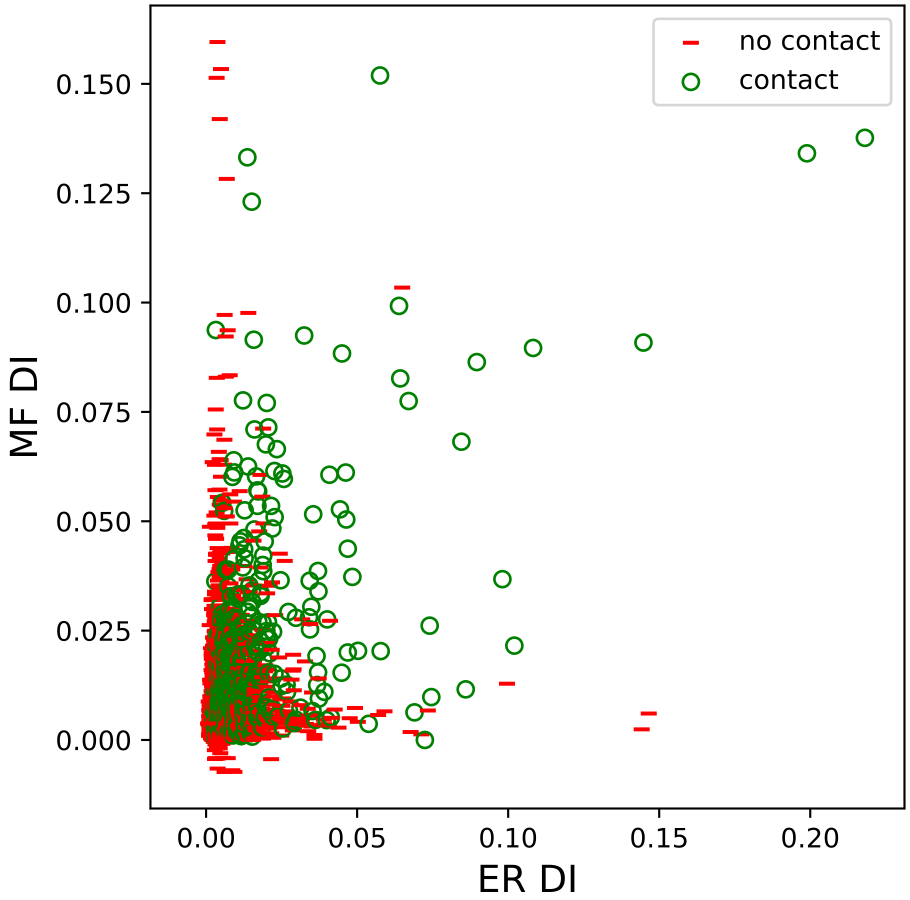

no contact
contact
MF DI
ER DI
0.000
0.025
0.050
0.075
0.100
0.125
0.150
0.00
0.05
0.10
0.15
0.20

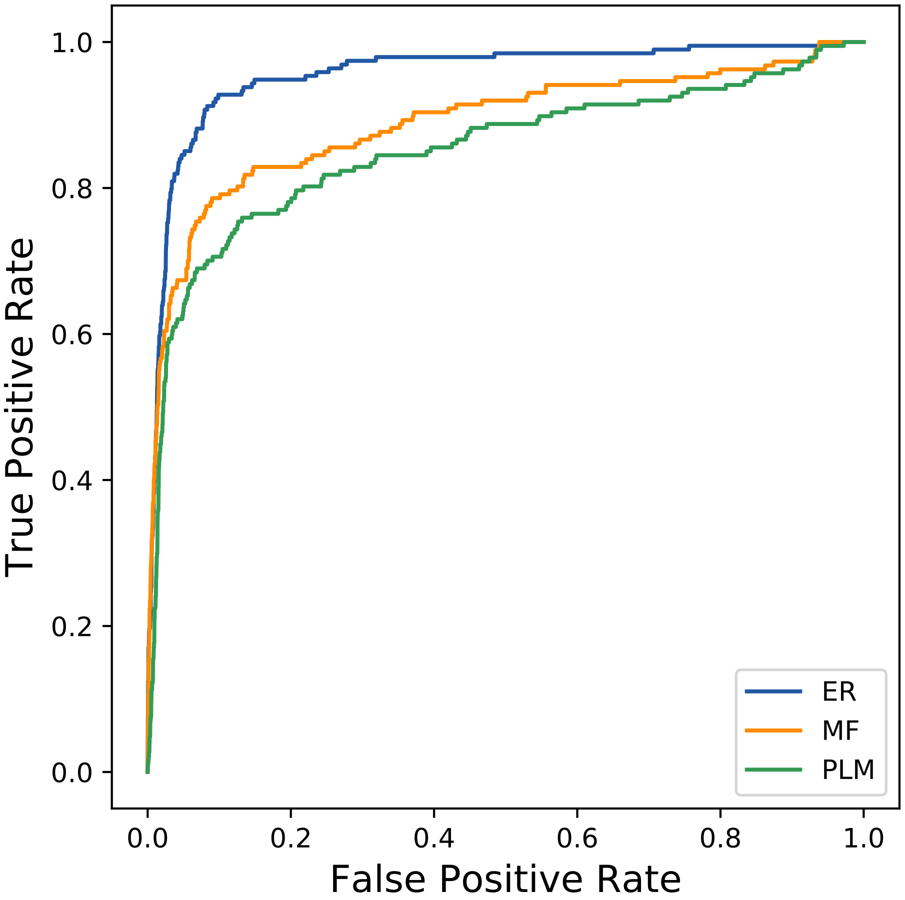

True Positive Rate
1.0
0.8
0.6
0.4
0.2
0.0
0.0
0.2
0.4
0.6
0.8
1.0
False Positive Rate
ER
MF
PLM
