## Supplemental Contact Maps for "Protein Structure Prediction with Expectation Reflection": 11708seq_168col_4rhq_PF00491_plots.pdf

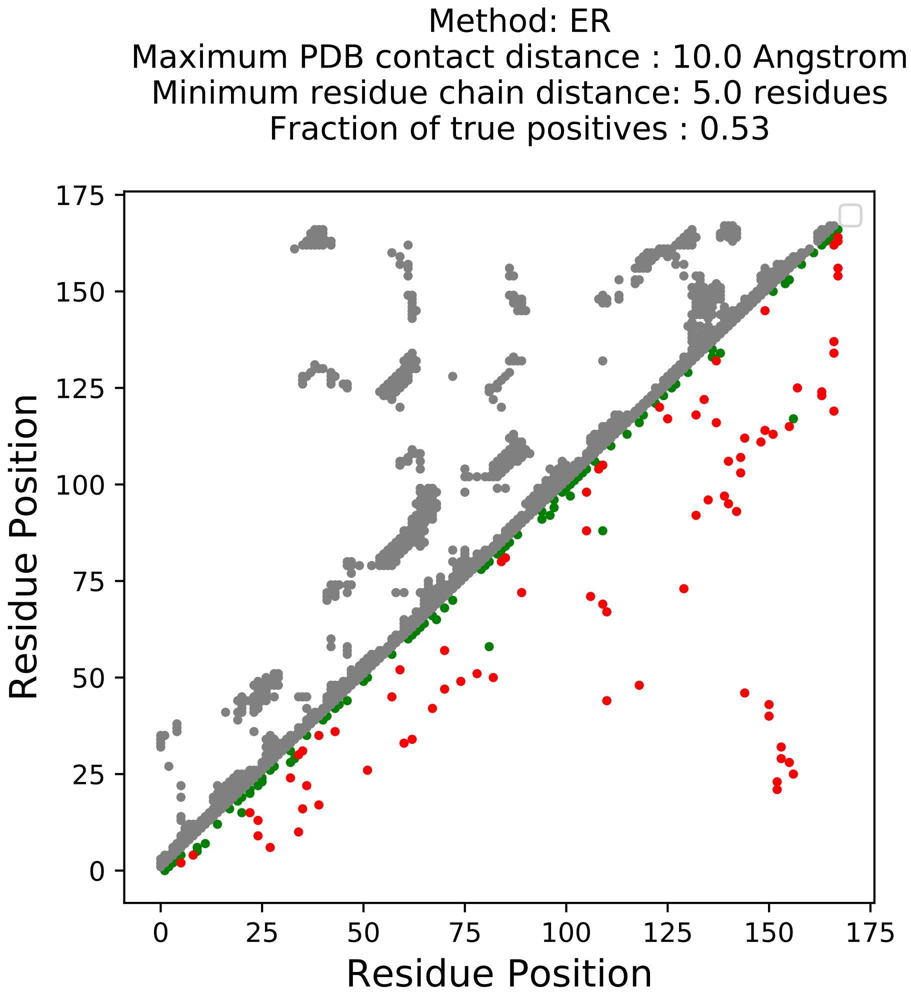

Method: ER
Maximum PDB contact distance : 10.0 Angstrom
Minimum residue chain distance: 5.0 residues
Fraction of true positives : 0.53
Residue Position
Residue Position
0
25
50
75
100
125
150
175

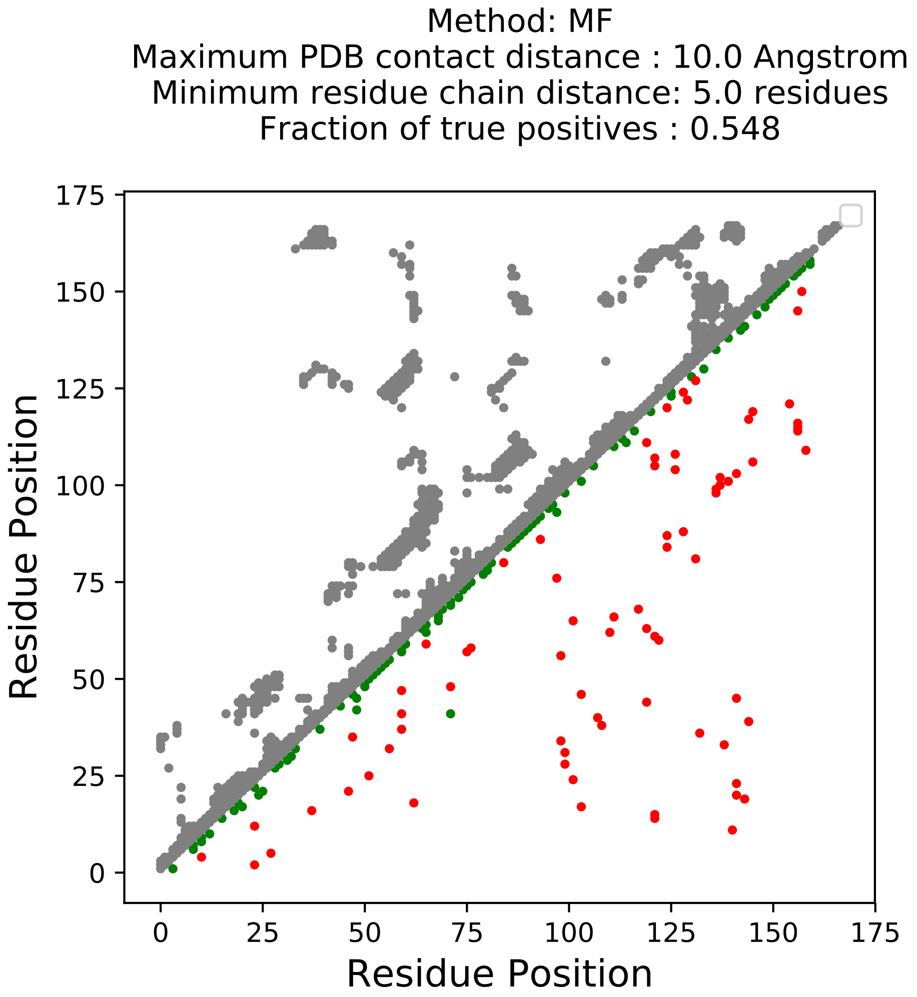
Method: MF
Maximum PDB contact distance : 10.0 Angstrom
Minimum residue chain distance: 5.0 residues
Fraction of true positives : 0.548
Residue Position
Residue Position
0
25
50
75
100
125
150
175

Method: PLM
Maximum PDB contact distance : 10.0 Angstrom
Minimum residue chain distance: 5.0 residues
Fraction of true positives : 0.476

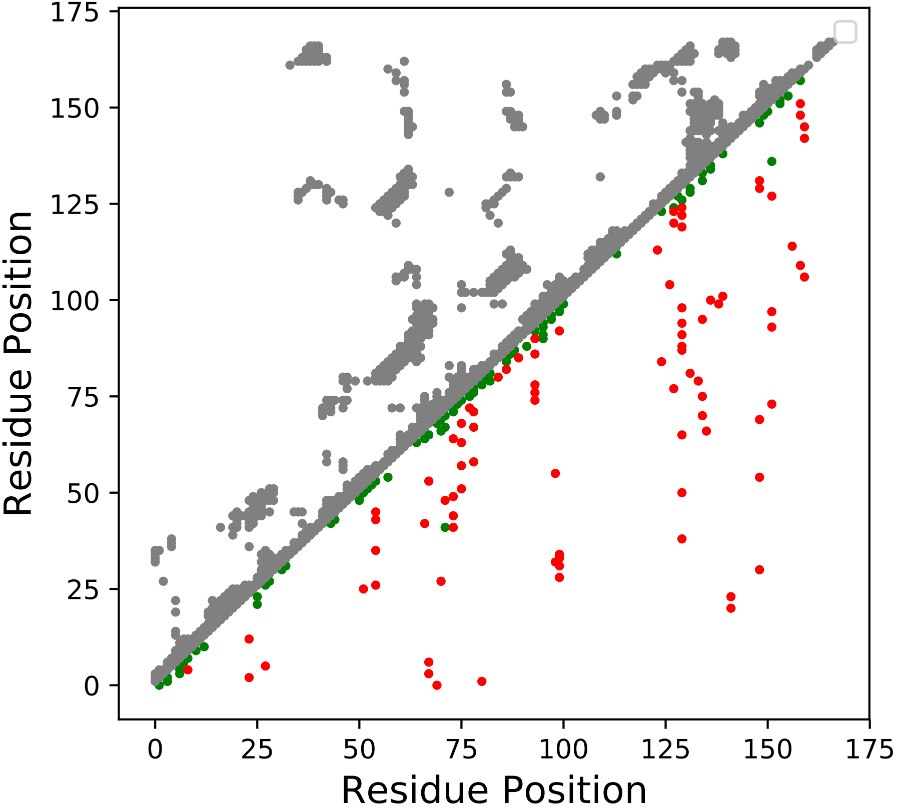

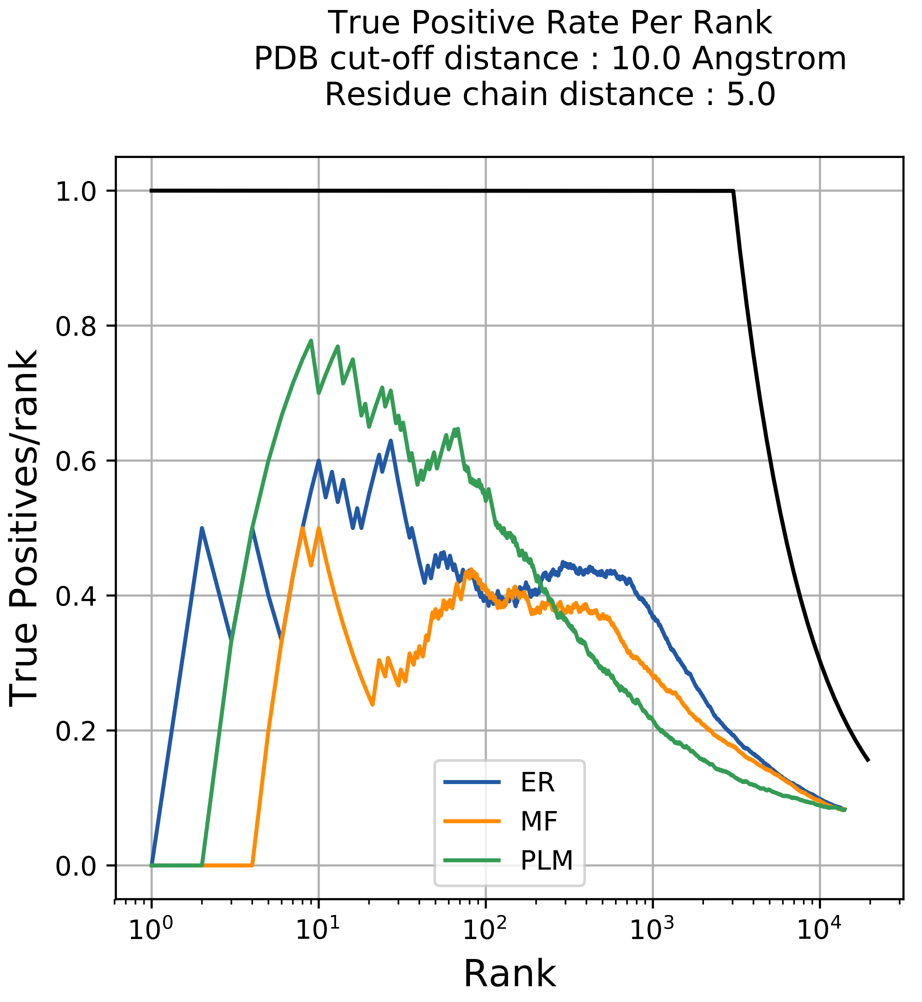
True Positive Rate Per Rank
PDB cut-off distance : 10.0 Angstrom
Residue chain distance : 5.0
True Positives/rank
Rank
1.0
0.8
0.6
0.4
0.2
0.0
$10^0$
$10^1$
$10^2$
$10^3$
$10^4$
ER
MF
PLM

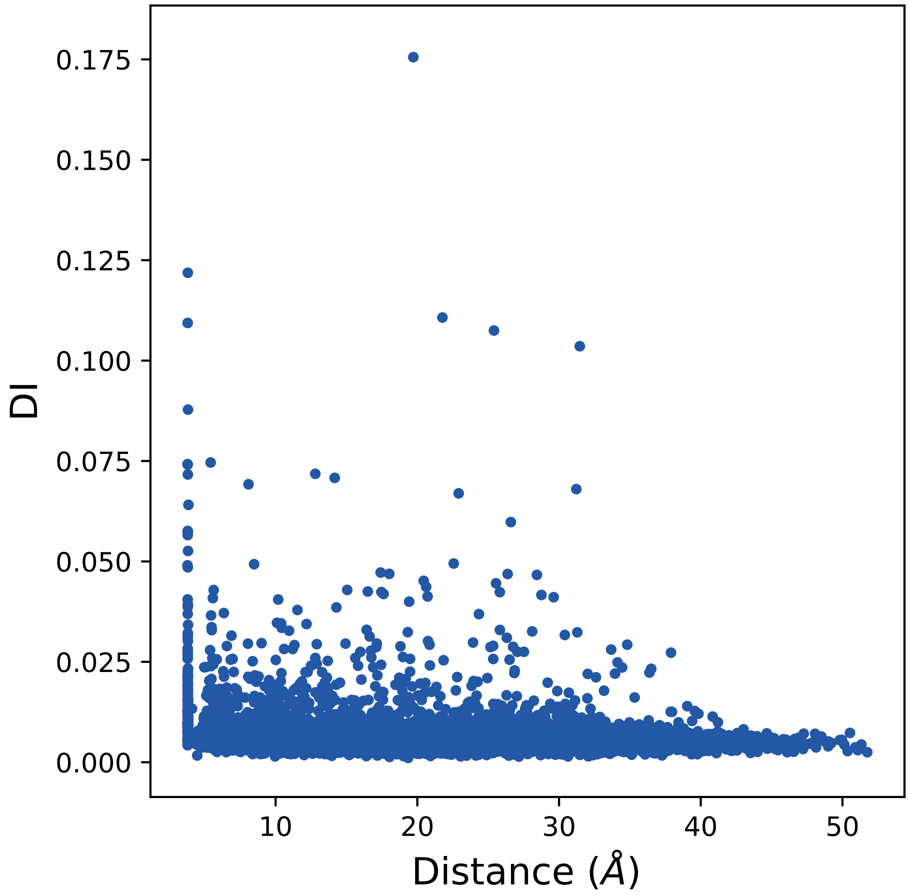

0.175
0.150
0.125
0.100
0.075
0.050
0.025
0.000
DI
10
20
30
40
50
Distance (Å)

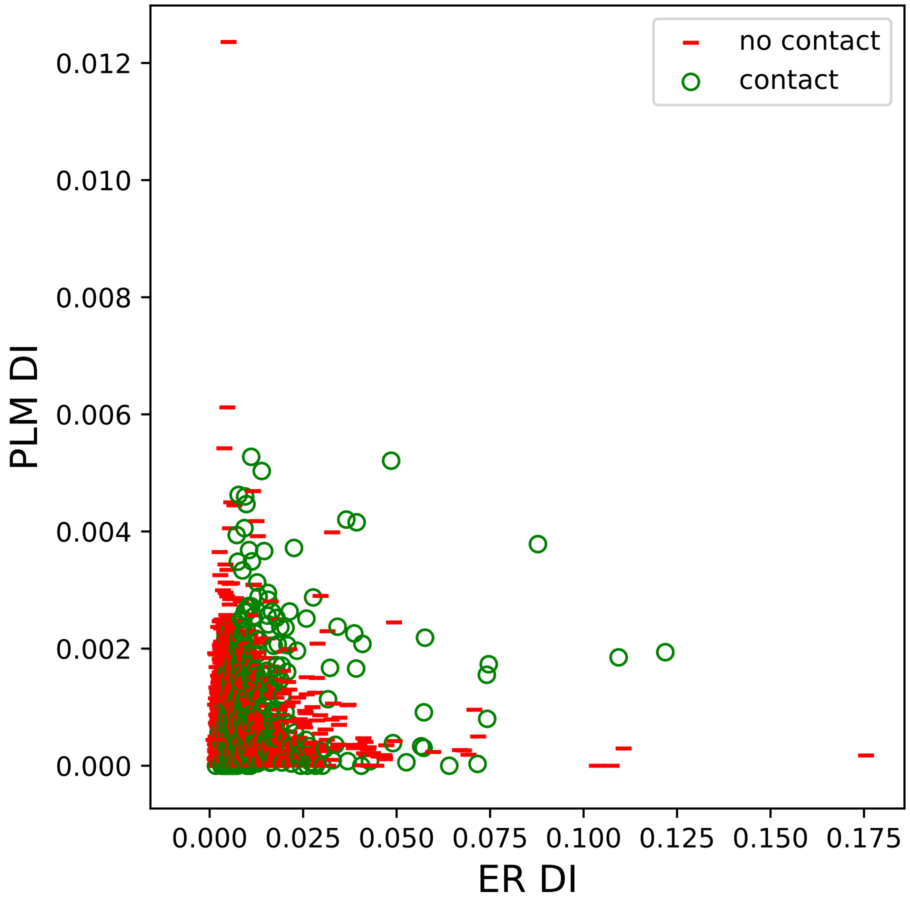

no contact
contact
PLM DI
ER DI
0.012
0.010
0.008
0.006
0.004
0.002
0.000
0.000
0.025
0.050
0.075
0.100
0.125
0.150
0.175

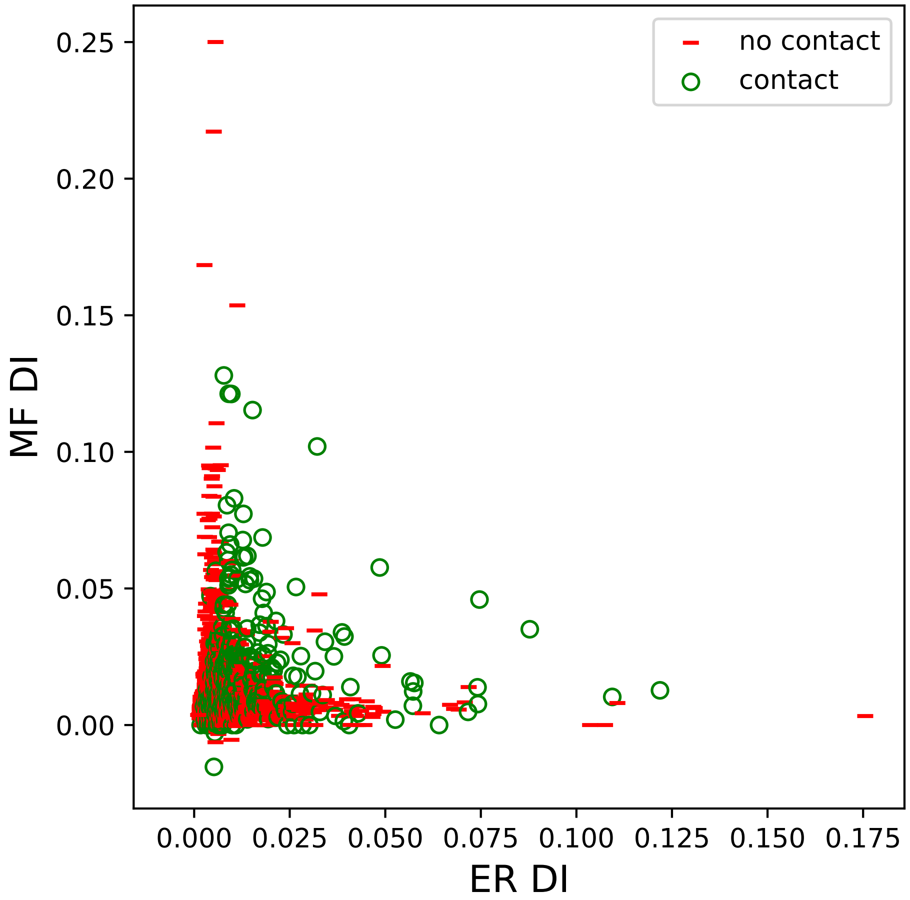

no contact
contact
MF DI
ER DI
0.25
0.20
0.15
0.10
0.05
0.00
0.000
0.025
0.050
0.075
0.100
0.125
0.150
0.175

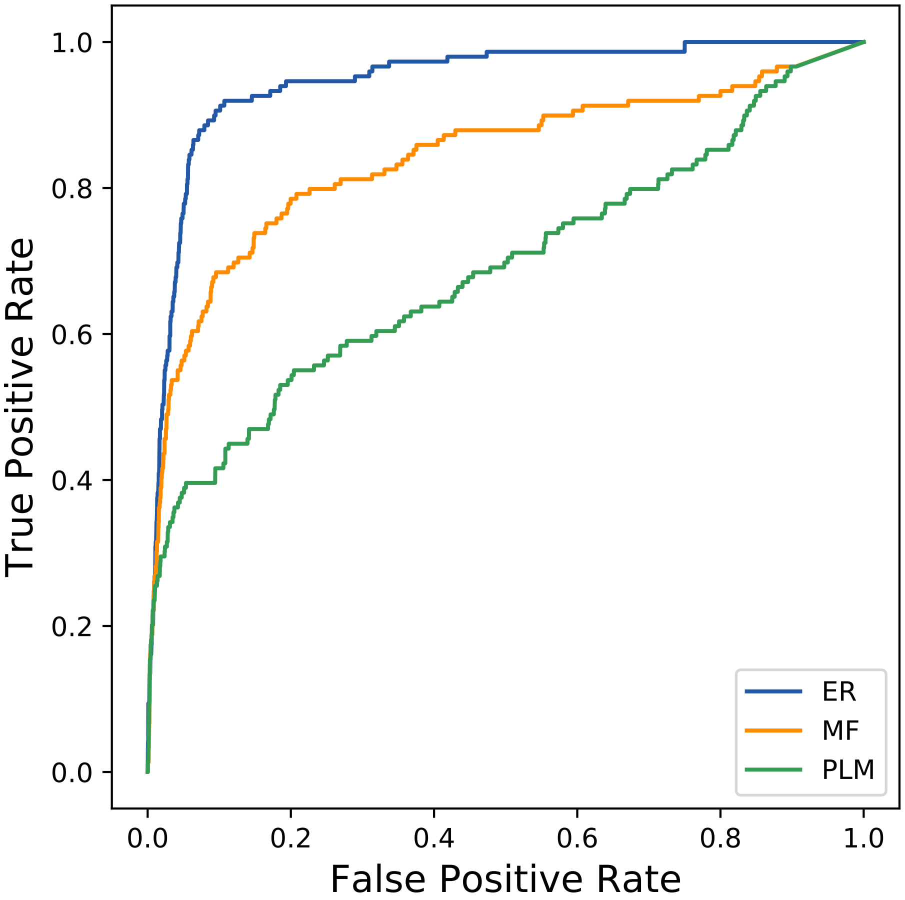

True Positive Rate
False Positive Rate
0.0
0.2
0.4
0.6
0.8
1.0
ER
MF
PLM
