## Supplemental Contact Maps for "Protein Structure Prediction with Expectation Reflection": 12187seq_111col_7epf_PF00258_plots.pdf

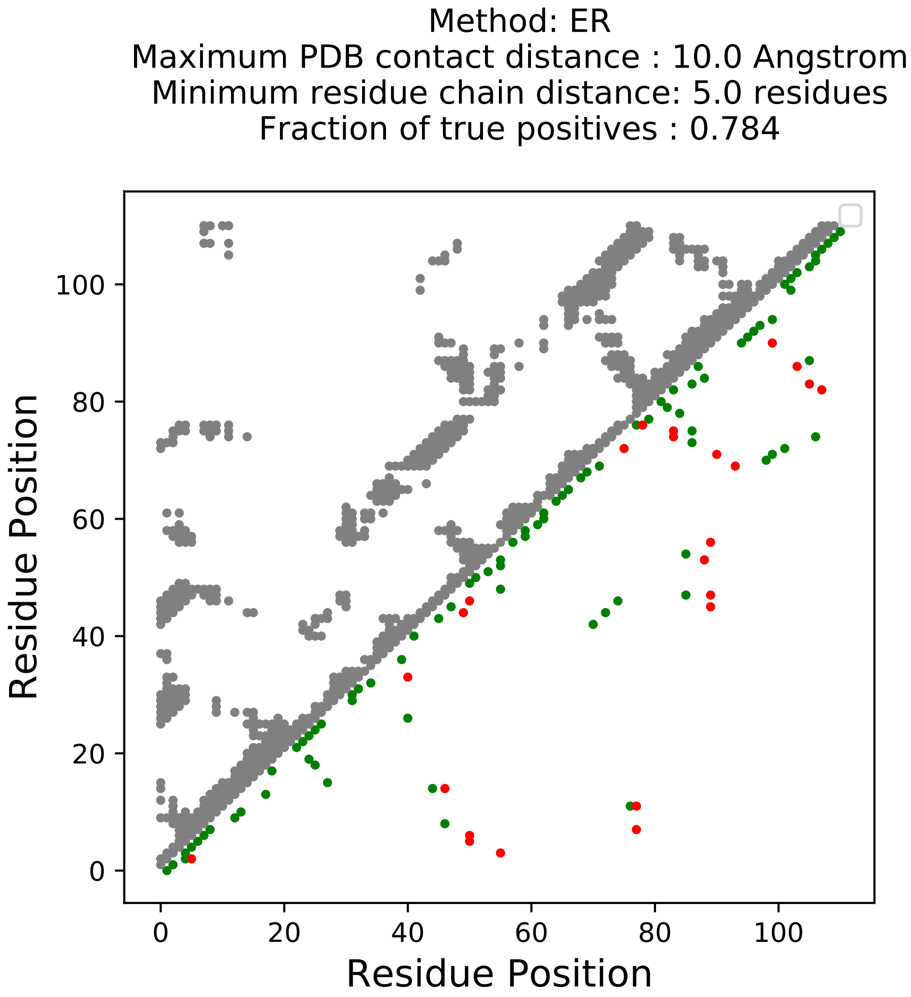
Method: ER
Maximum PDB contact distance : 10.0 Angstrom
Minimum residue chain distance: 5.0 residues
Fraction of true positives : 0.784
Residue Position
Residue Position
0
20
40
60
80
100

Method: MF
Maximum PDB contact distance : 10.0 Angstrom
Minimum residue chain distance: 5.0 residues
Fraction of true positives : 0.694

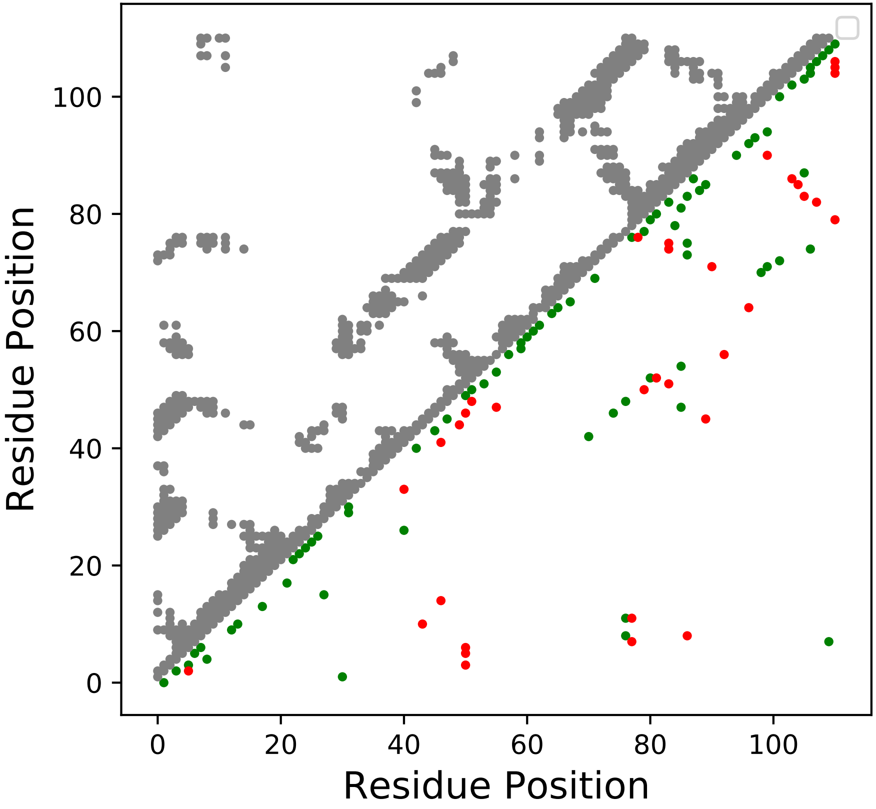

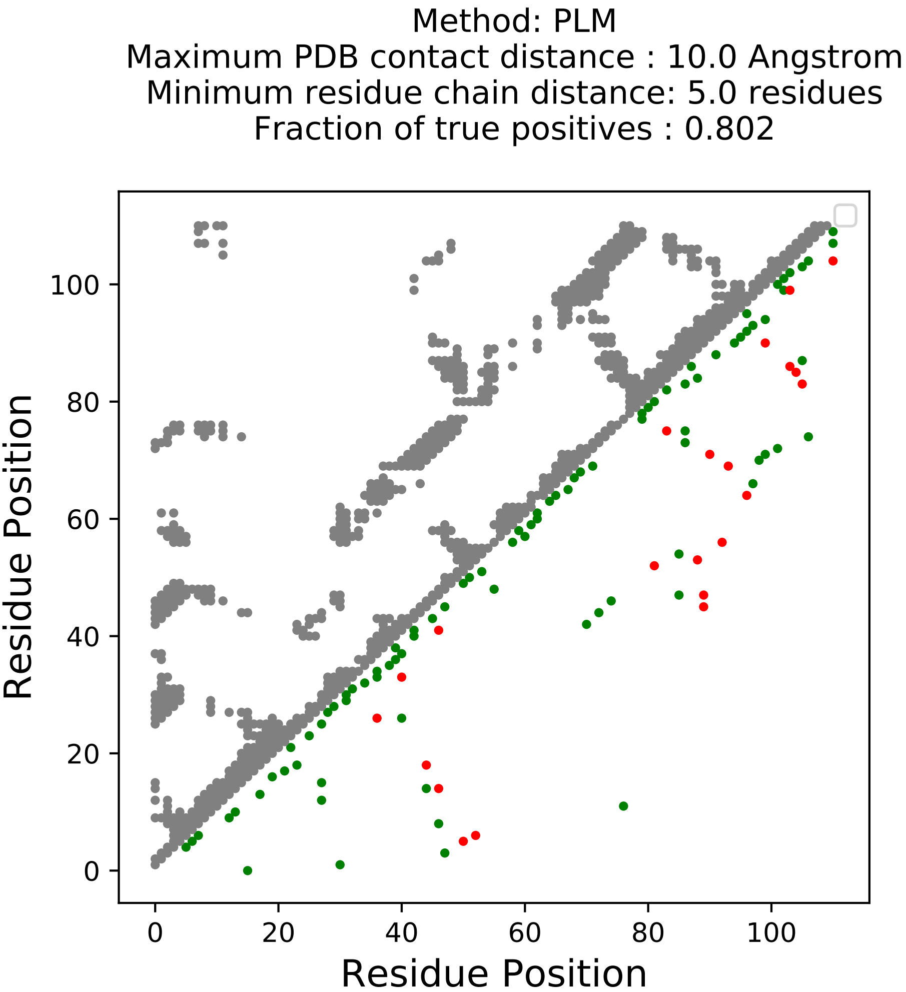
Method: PLM
Maximum PDB contact distance : 10.0 Angstrom
Minimum residue chain distance: 5.0 residues
Fraction of true positives : 0.802
Residue Position
Residue Position
0
20
40
60
80
100

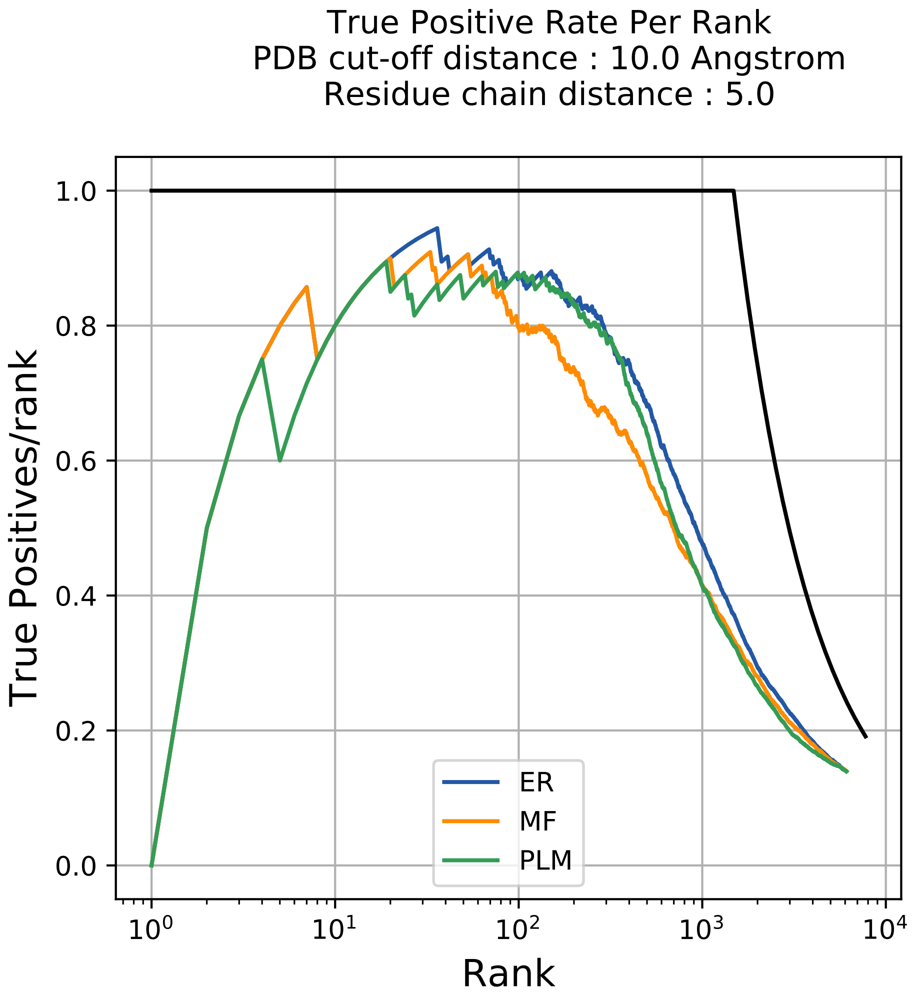

True Positive Rate Per Rank
PDB cut-off distance : 10.0 Angstrom
Residue chain distance : 5.0
True Positives/rank
Rank
ER
MF
PLM

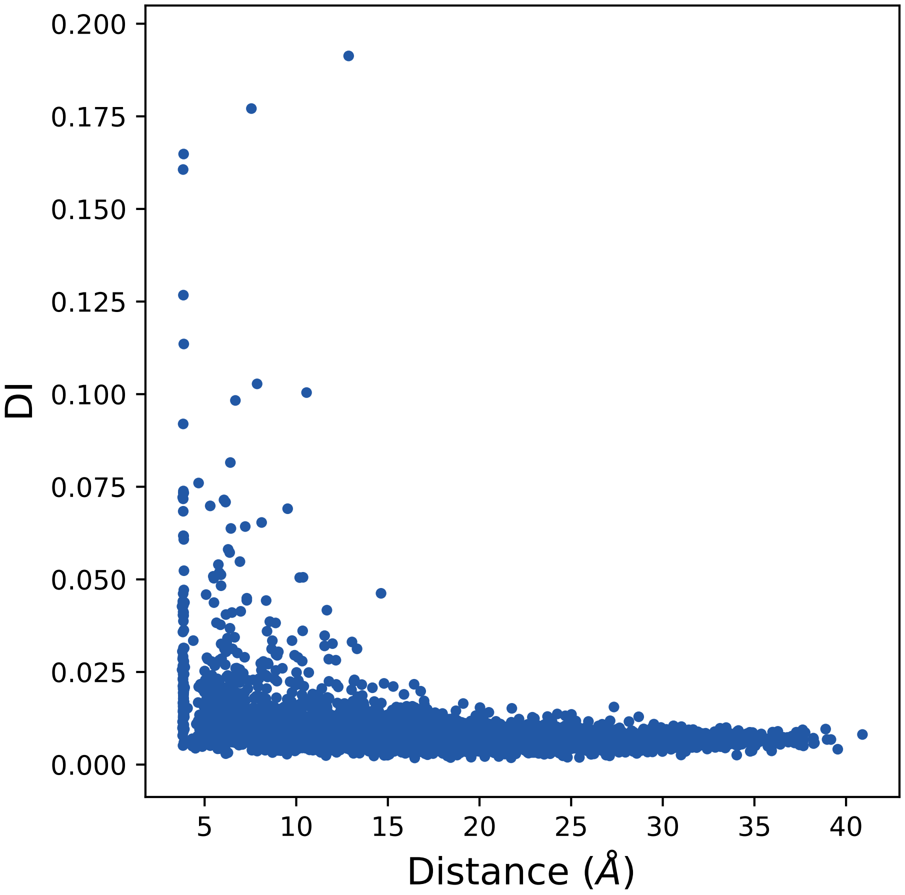

0.200
0.175
0.150
0.125
0.100
0.075
0.050
0.025
0.000
DI
5
10
15
20
25
30
35
40
Distance (Å)

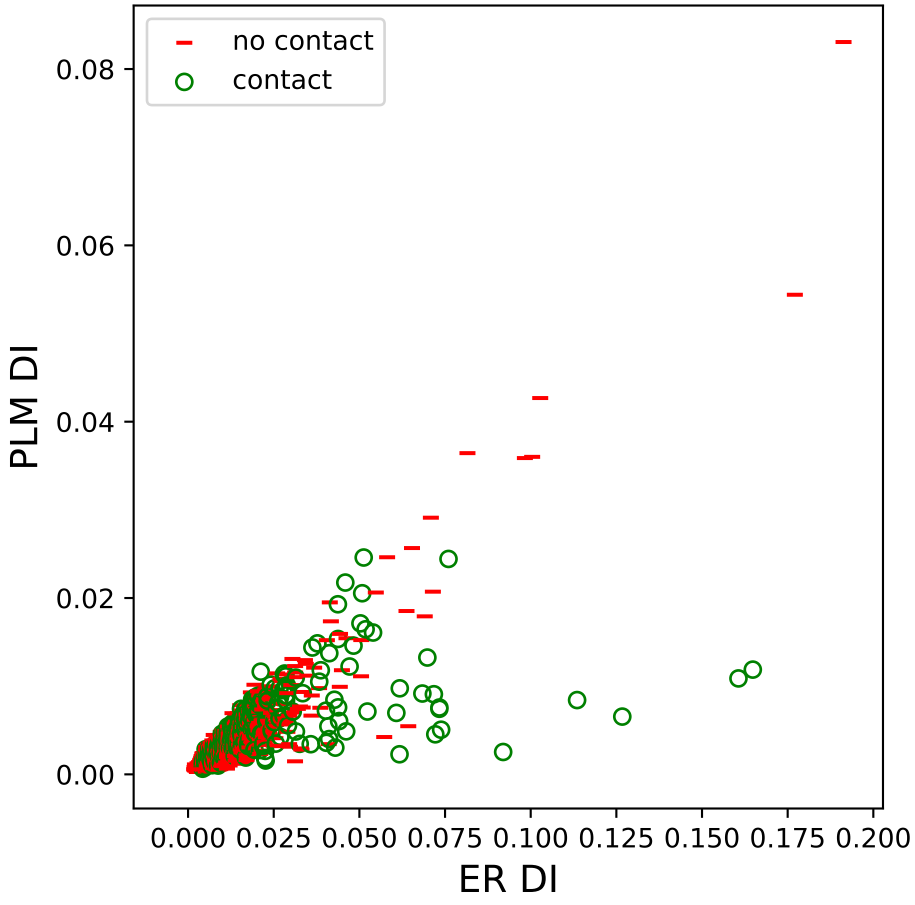
no contact
contact
PLM DI
ER DI
0.00
0.02
0.04
0.06
0.08
0.000
0.025
0.050
0.075
0.100
0.125
0.150
0.175
0.200

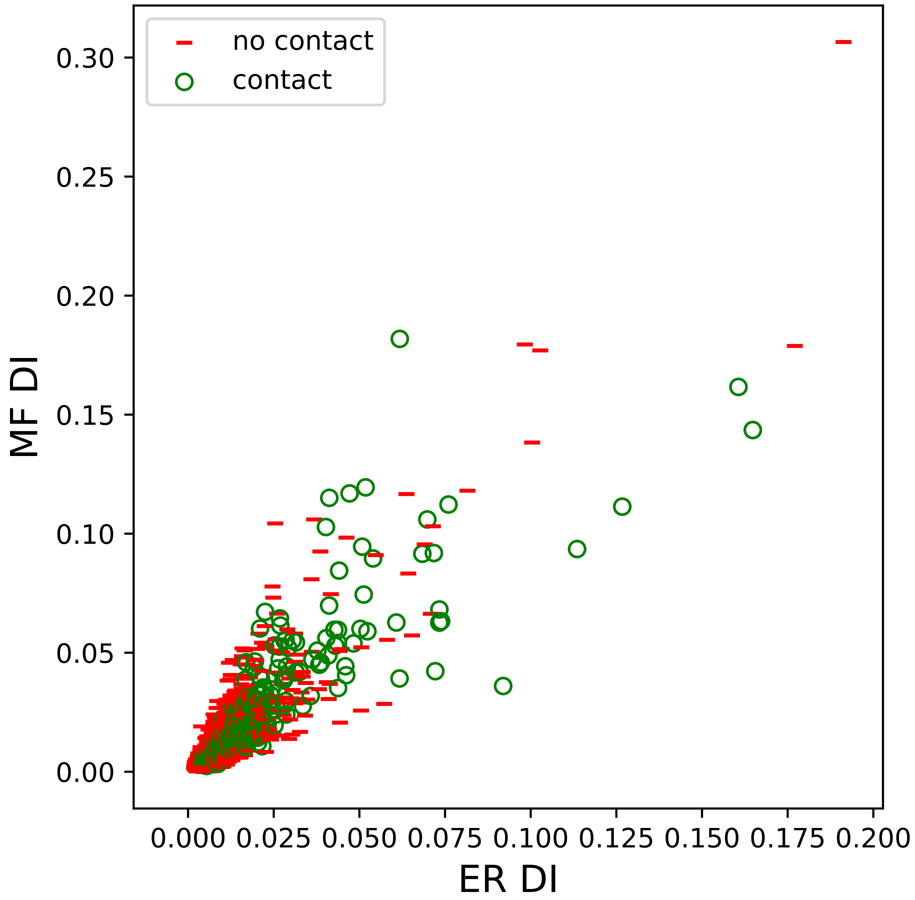

no contact
contact
0.30
0.25
0.20
0.15
0.10
0.05
0.00
MF DI
0.000
0.025
0.050
0.075
0.100
0.125
0.150
0.175
0.200
ER DI

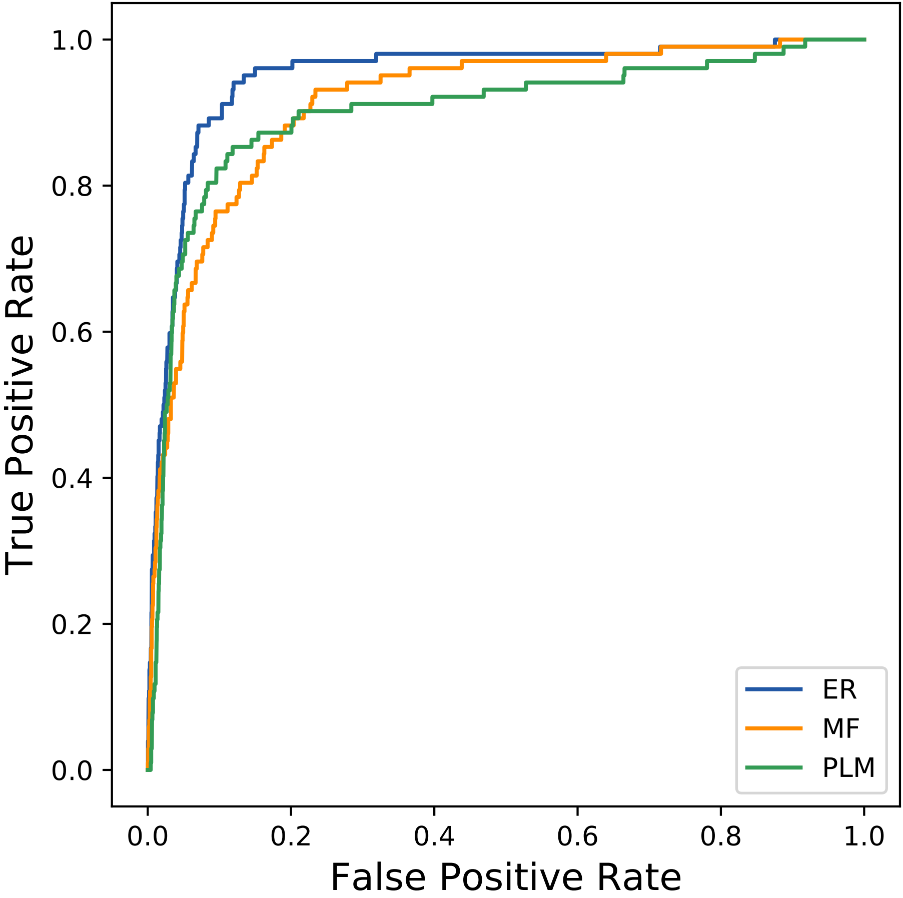

True Positive Rate
False Positive Rate
0.0
0.2
0.4
0.6
0.8
1.0
ER
MF
PLM
