## Supplemental Contact Maps for "Protein Structure Prediction with Expectation Reflection": 12334seq_108col_1zy1_PF01327_plots.pdf

Method: ER
Maximum PDB contact distance : 10.0 Angstrom
Minimum residue chain distance: 5.0 residues
Fraction of true positives : 0.843

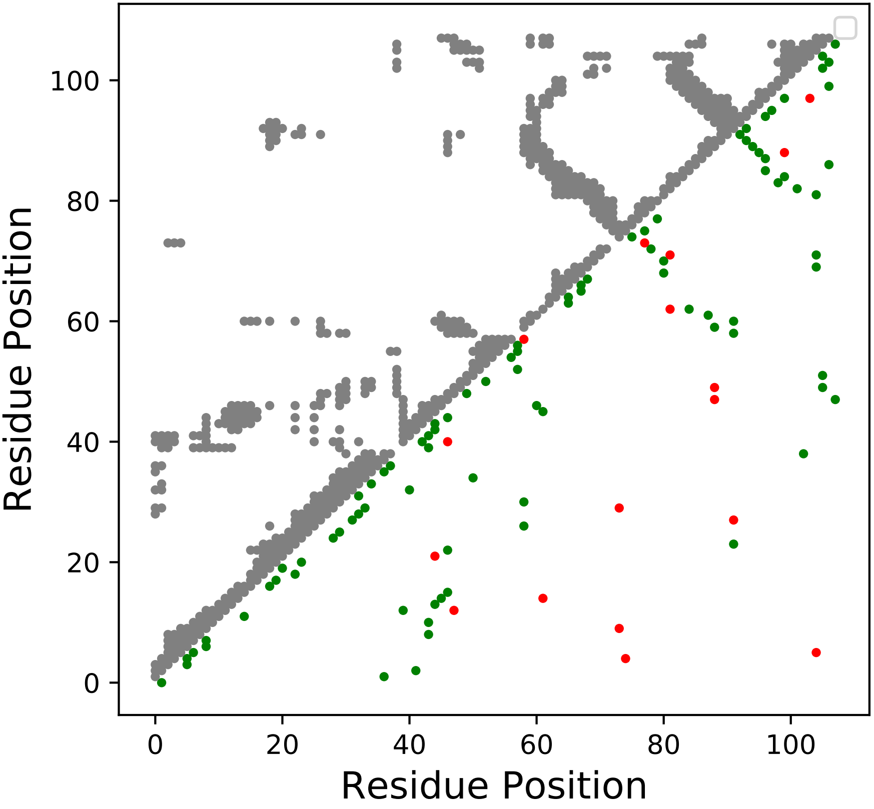

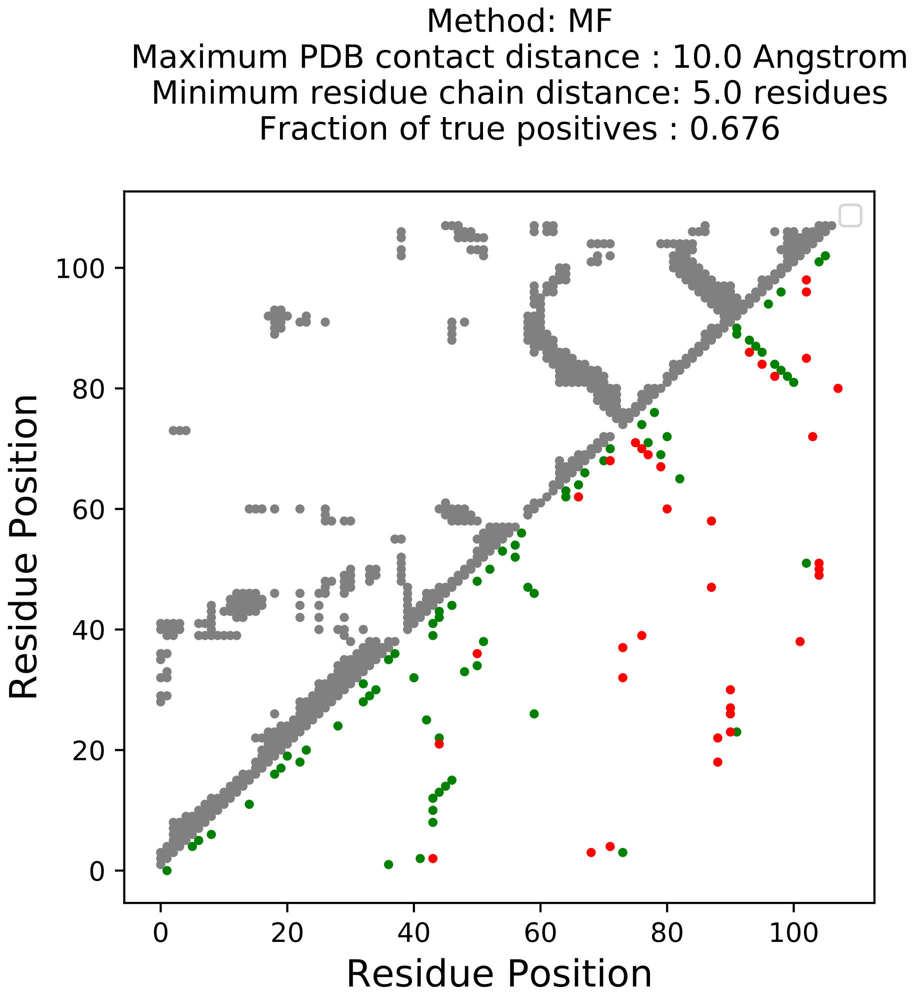
Method: MF
Maximum PDB contact distance : 10.0 Angstrom
Minimum residue chain distance: 5.0 residues
Fraction of true positives : 0.676
Residue Position
Residue Position
0
20
40
60
80
100

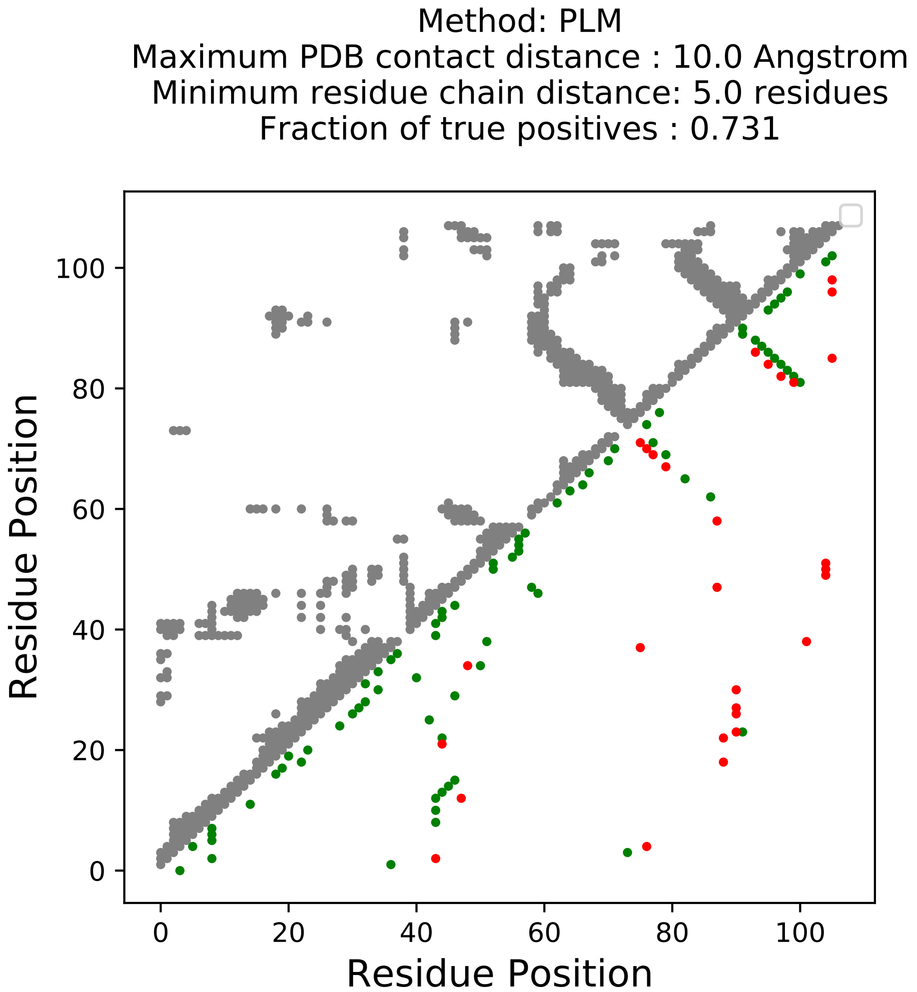

Method: PLM
Maximum PDB contact distance : 10.0 Angstrom
Minimum residue chain distance: 5.0 residues
Fraction of true positives : 0.731
Residue Position
Residue Position
0
20
40
60
80
100

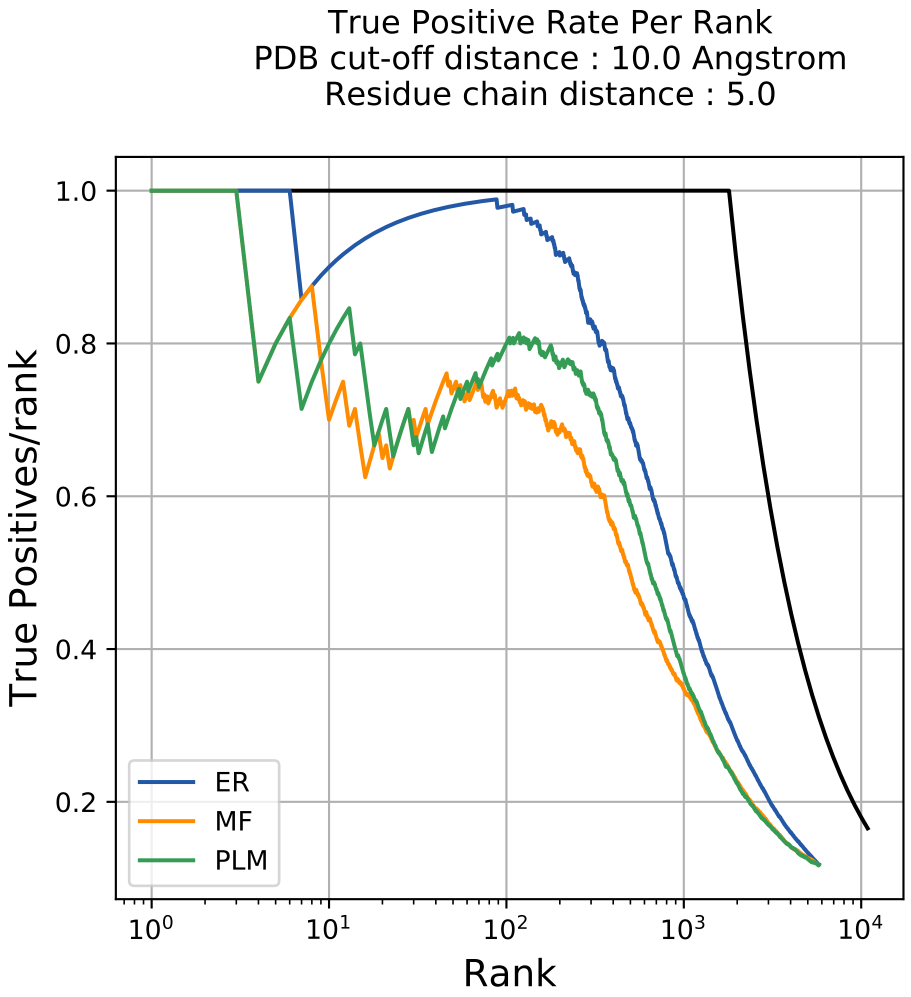
True Positive Rate Per Rank
PDB cut-off distance : 10.0 Angstrom
Residue chain distance : 5.0
True Positives/rank
Rank
1.0
0.8
0.6
0.4
0.2
$10^0$
$10^1$
$10^2$
$10^3$
$10^4$
ER
MF
PLM

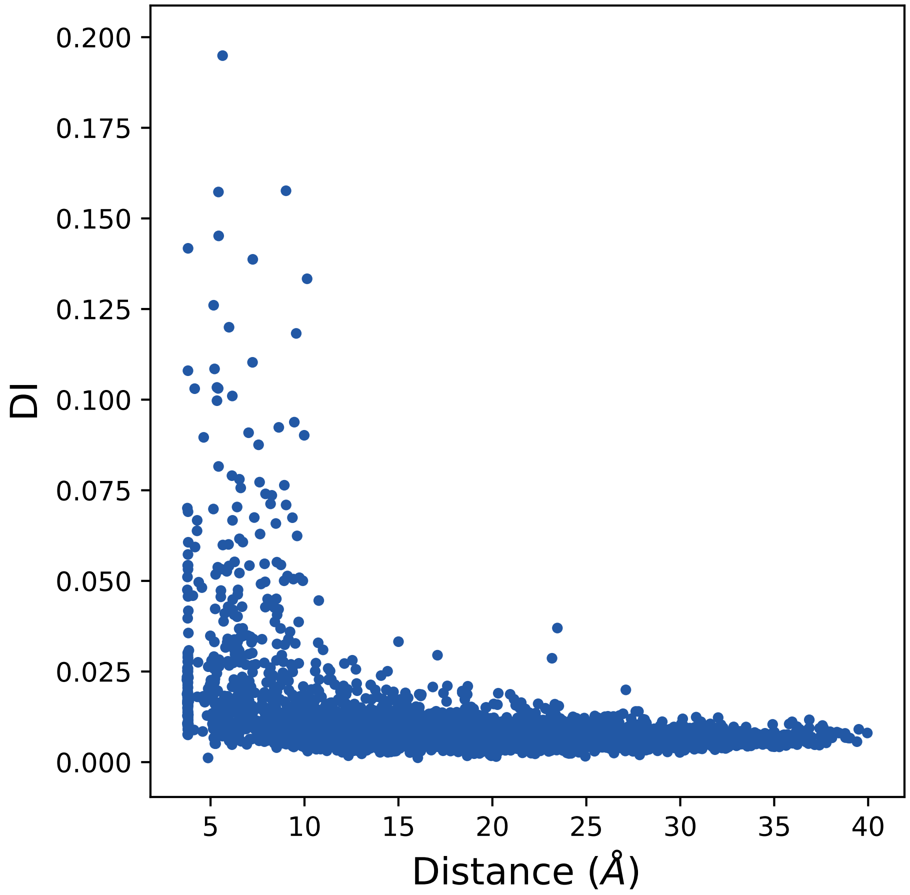

DI
Distance (Å)
0.000
0.025
0.050
0.075
0.100
0.125
0.150
0.175
0.200
5
10
15
20
25
30
35
40

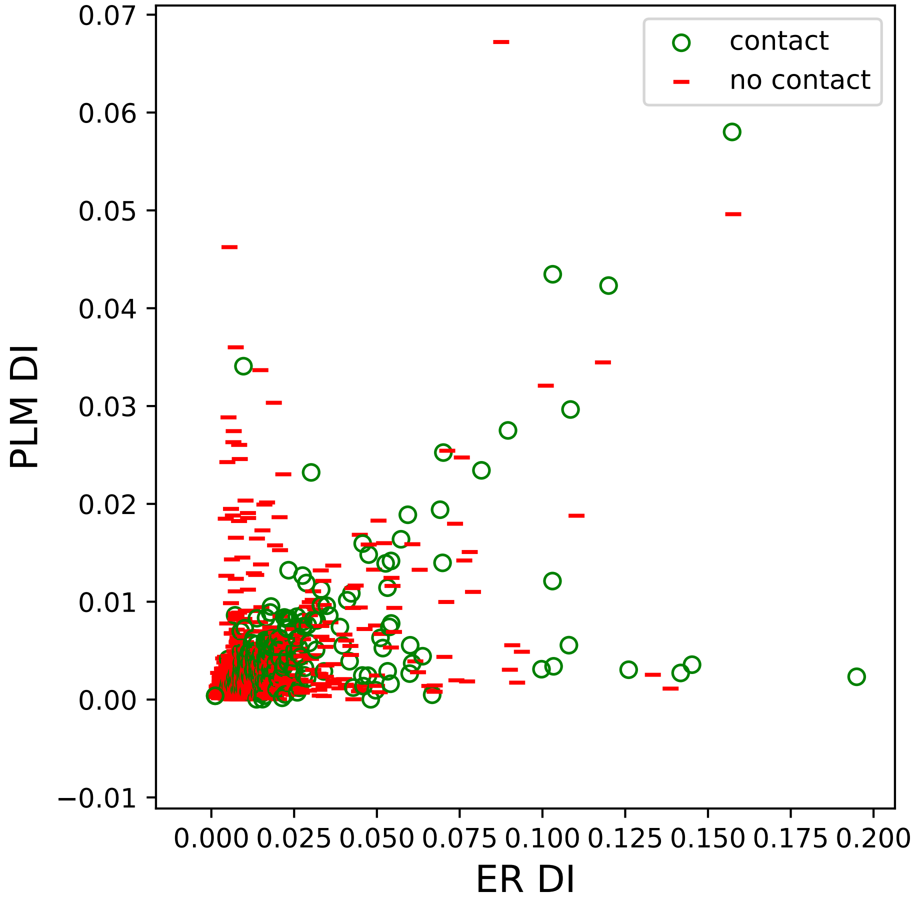

contact
no contact
PLM DI
ER DI
0.07
0.06
0.05
0.04
0.03
0.02
0.01
0.00
−0.01
0.000
0.025
0.050
0.075
0.100
0.125
0.150
0.175
0.200

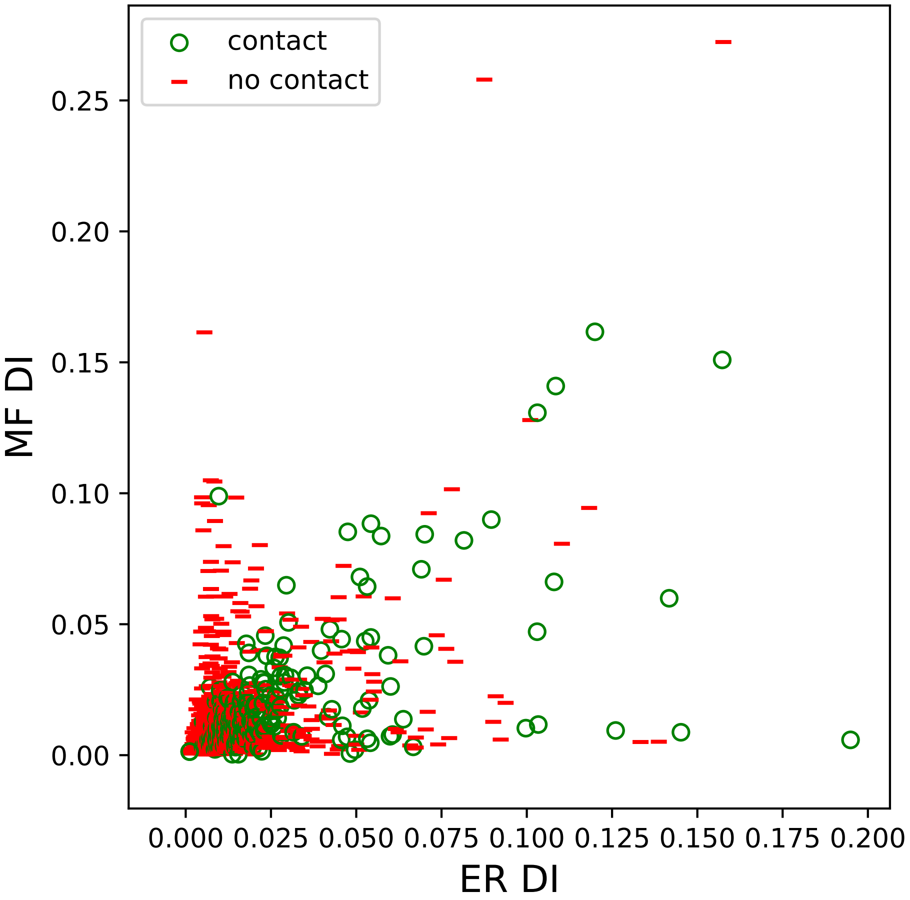
contact
no contact
0.25
0.20
0.15
0.10
0.05
0.00
MF DI
0.000
0.025
0.050
0.075
0.100
0.125
0.150
0.175
0.200
ER DI

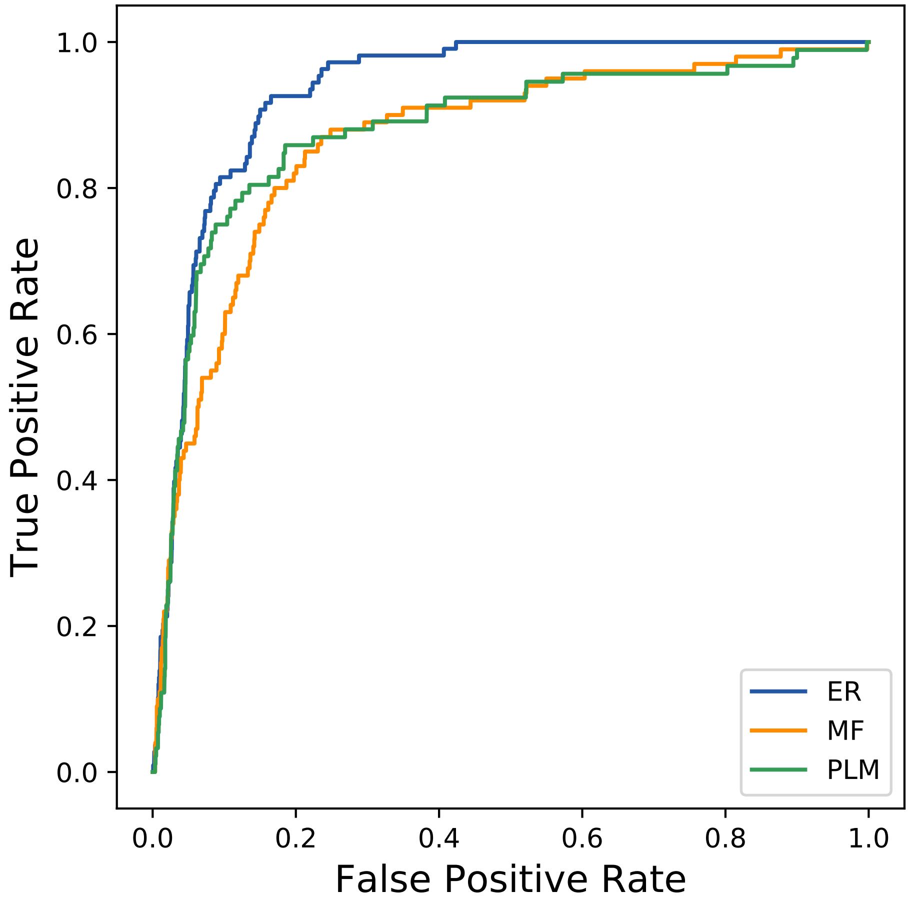
True Positive Rate
False Positive Rate
0.0
0.2
0.4
0.6
0.8
1.0
ER
MF
PLM
