## Supplemental Contact Maps for "Protein Structure Prediction with Expectation Reflection": 12971seq_32col_5px5_PF00439_plots.pdf

Method: ER
Maximum PDB contact distance : 10.0 Angstrom
Minimum residue chain distance: 5.0 residues
Fraction of true positives : 0.688

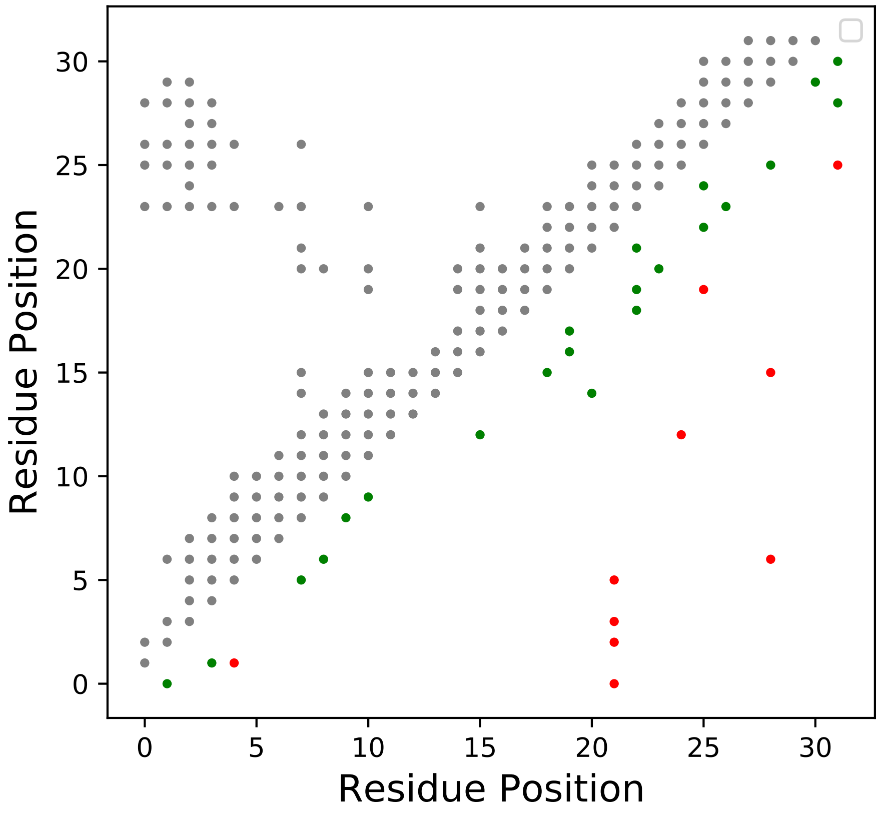

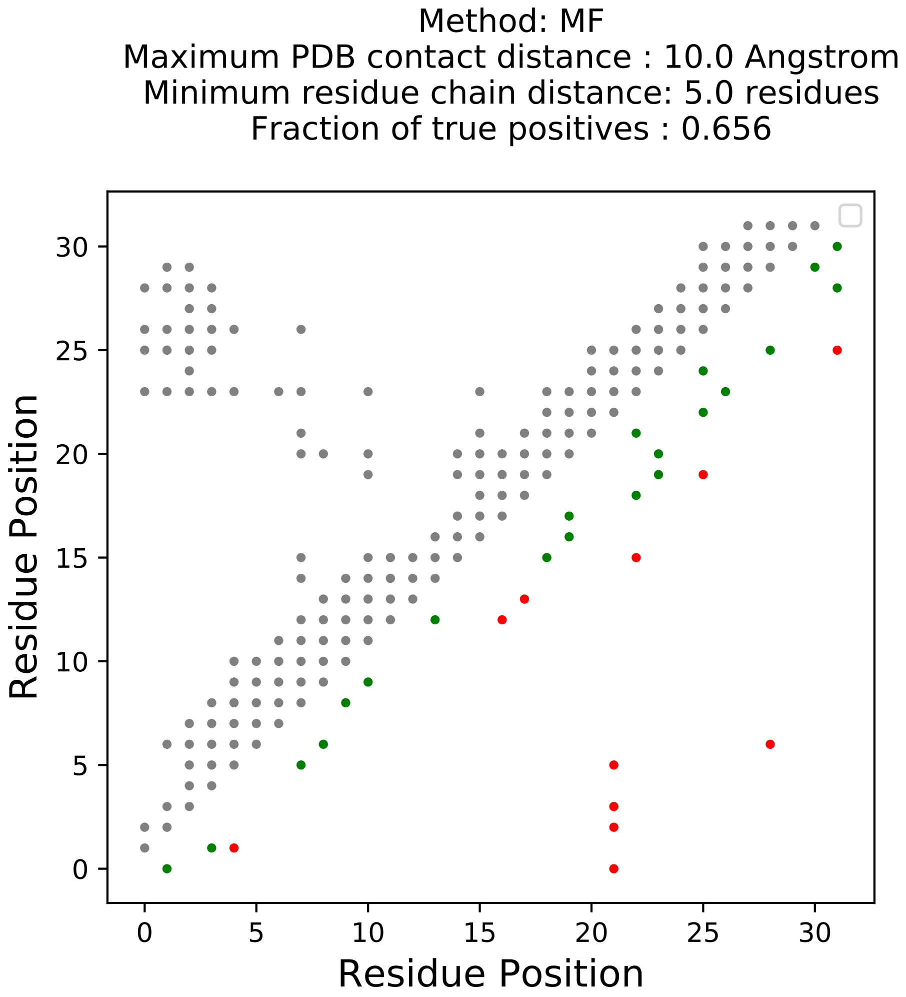
Method: MF
Maximum PDB contact distance : 10.0 Angstrom
Minimum residue chain distance: 5.0 residues
Fraction of true positives : 0.656
Residue Position
Residue Position
0
5
10
15
20
25
30

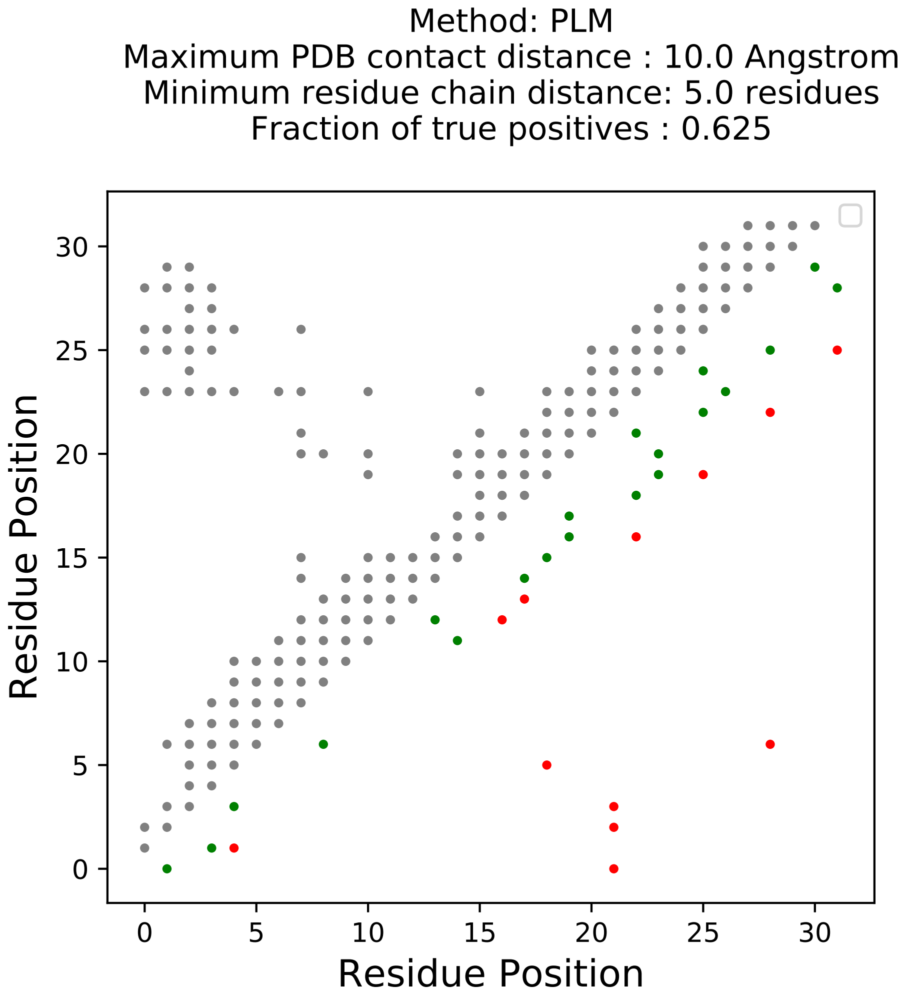
Method: PLM
Maximum PDB contact distance : 10.0 Angstrom
Minimum residue chain distance: 5.0 residues
Fraction of true positives : 0.625
Residue Position
Residue Position
0
5
10
15
20
25
30

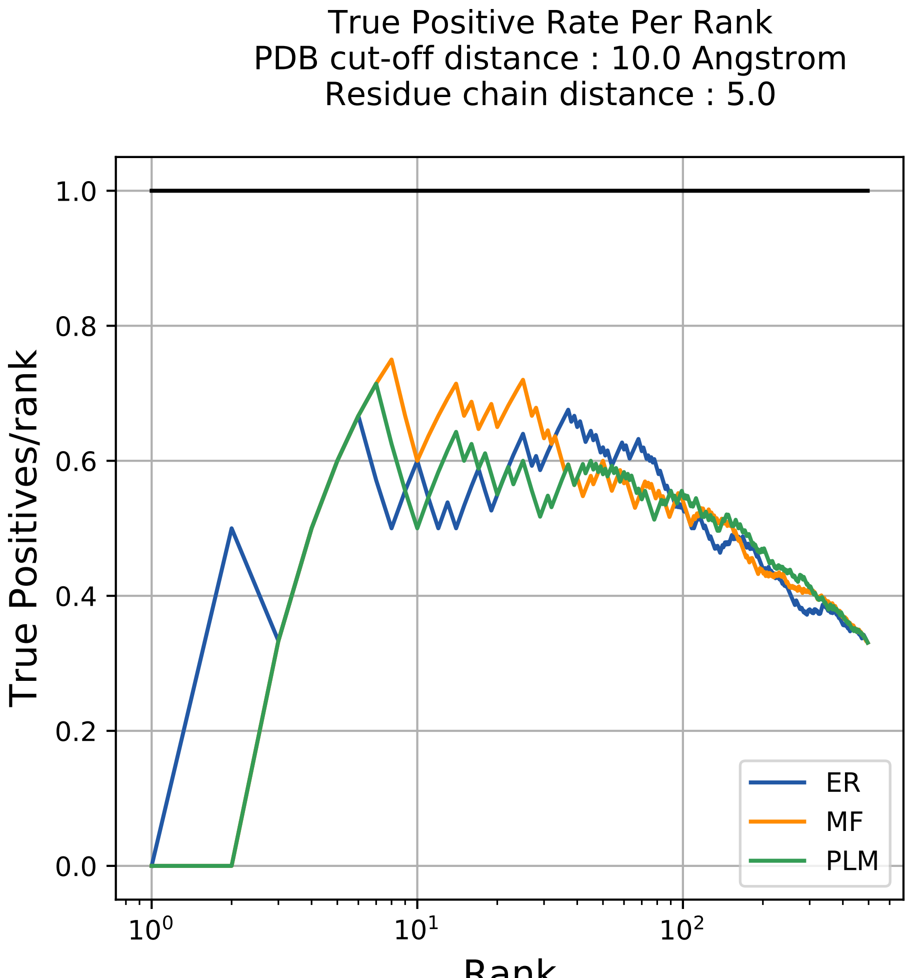
True Positive Rate Per Rank
PDB cut-off distance : 10.0 Angstrom
Residue chain distance : 5.0
True Positives/rank
Rank
1.0
0.8
0.6
0.4
0.2
0.0
$10^0$
$10^1$
$10^2$
ER
MF
PLM

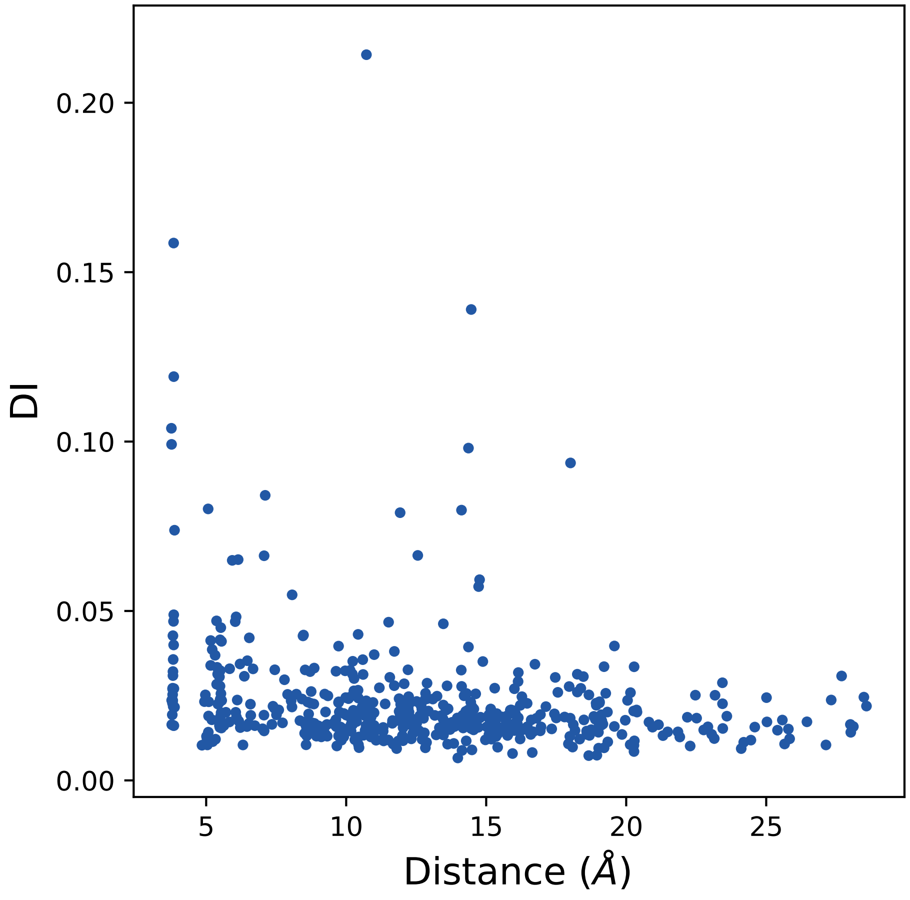

DI
Distance (Å)
0.00
0.05
0.10
0.15
0.20
5
10
15
20
25

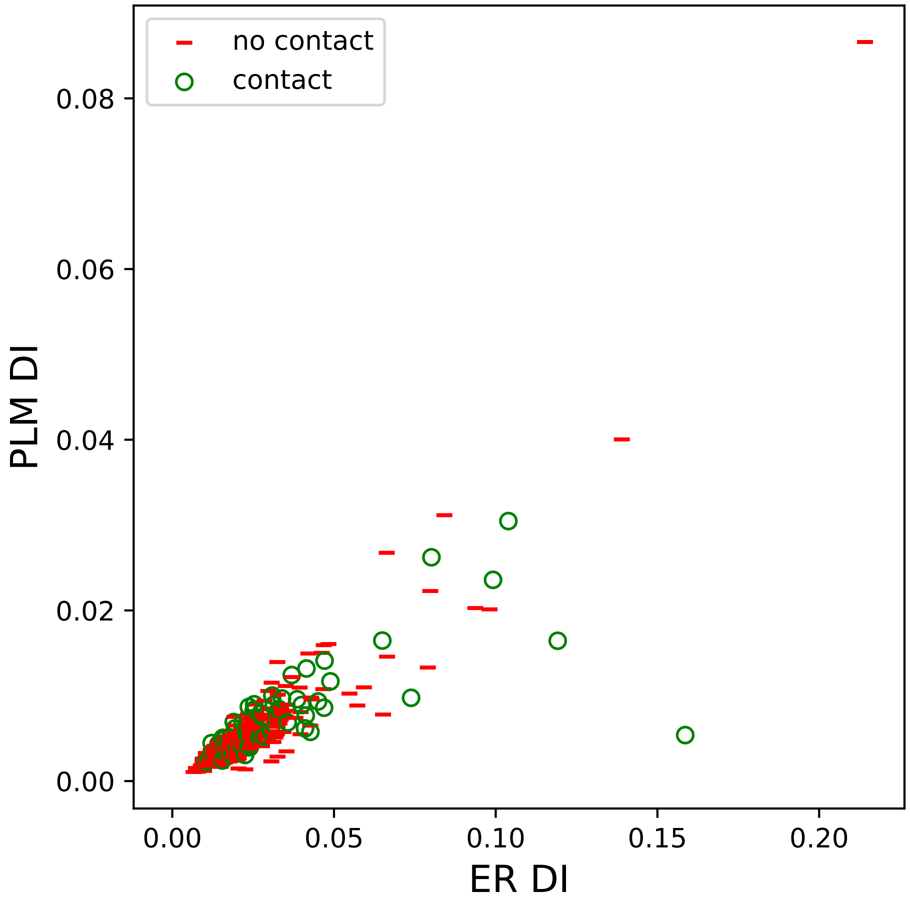

no contact
contact
0.08
0.06
0.04
0.02
0.00
PLM DI
0.00
0.05
0.10
0.15
0.20
ER DI

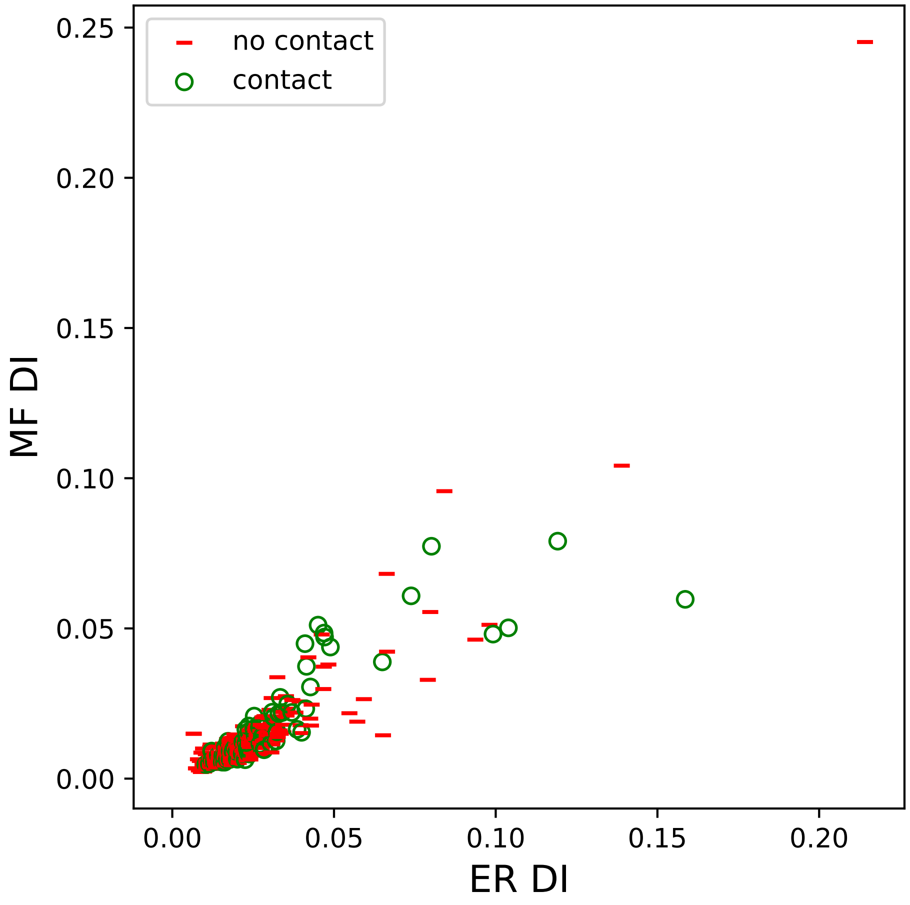

no contact
contact
MF DI
ER DI
0.00
0.05
0.10
0.15
0.20
0.25

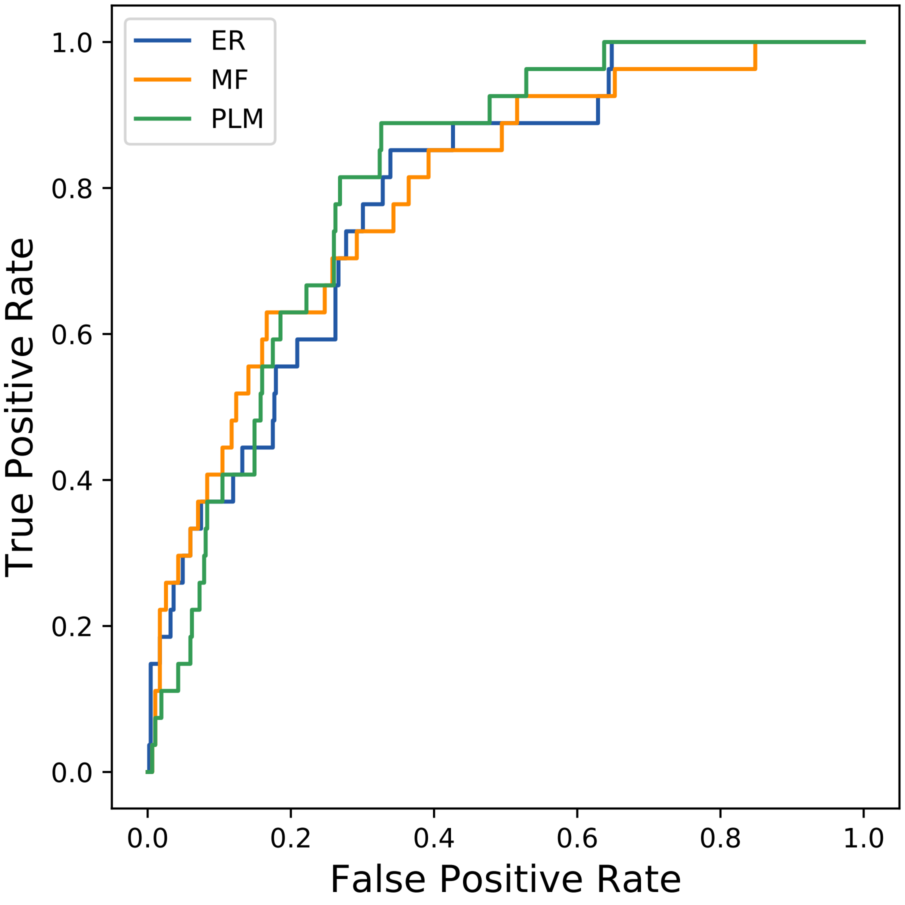
ER
MF
PLM
1.0
0.8
0.6
0.4
0.2
0.0
True Positive Rate
0.0
0.2
0.4
0.6
0.8
1.0
False Positive Rate
