## Supplemental Contact Maps for "Protein Structure Prediction with Expectation Reflection": 13816seq_58col_3dai_PF00439_plots.pdf

Method: ER
Maximum PDB contact distance : 10.0 Angstrom
Minimum residue chain distance: 5.0 residues
Fraction of true positives : 0.793

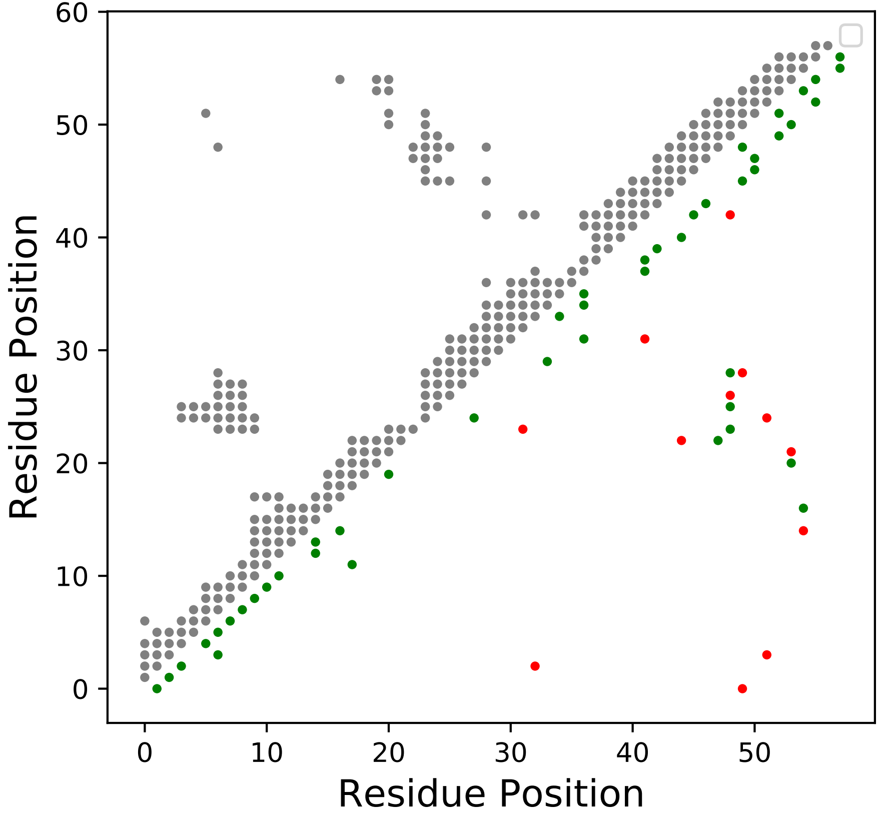

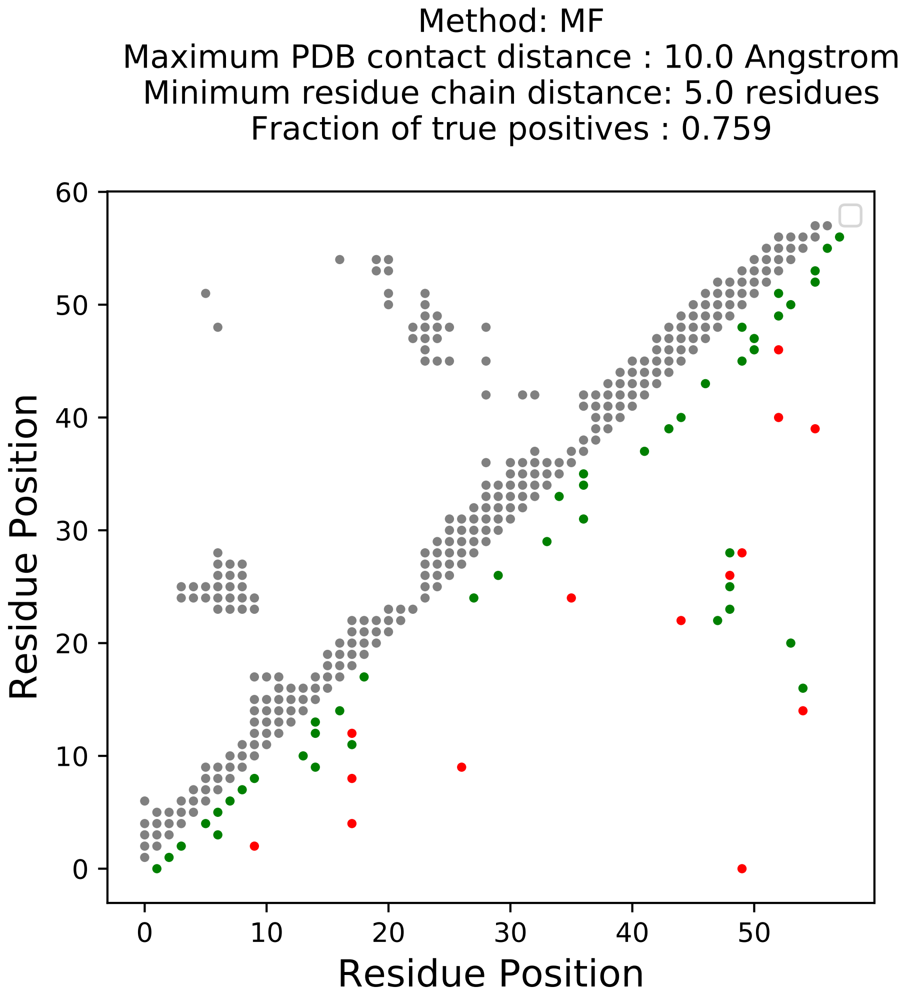
Method: MF
Maximum PDB contact distance : 10.0 Angstrom
Minimum residue chain distance: 5.0 residues
Fraction of true positives : 0.759
Residue Position
Residue Position
0
10
20
30
40
50
60

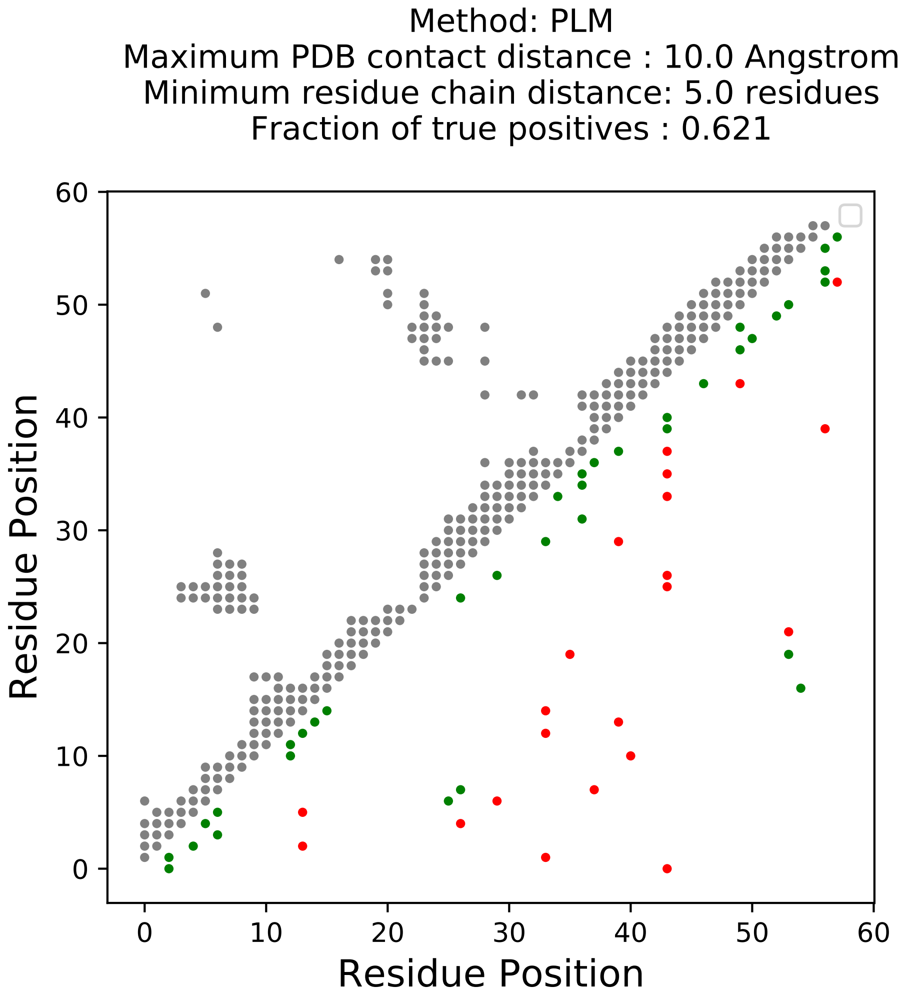
Method: PLM
Maximum PDB contact distance : 10.0 Angstrom
Minimum residue chain distance: 5.0 residues
Fraction of true positives : 0.621
Residue Position
Residue Position
0
10
20
30
40
50
60

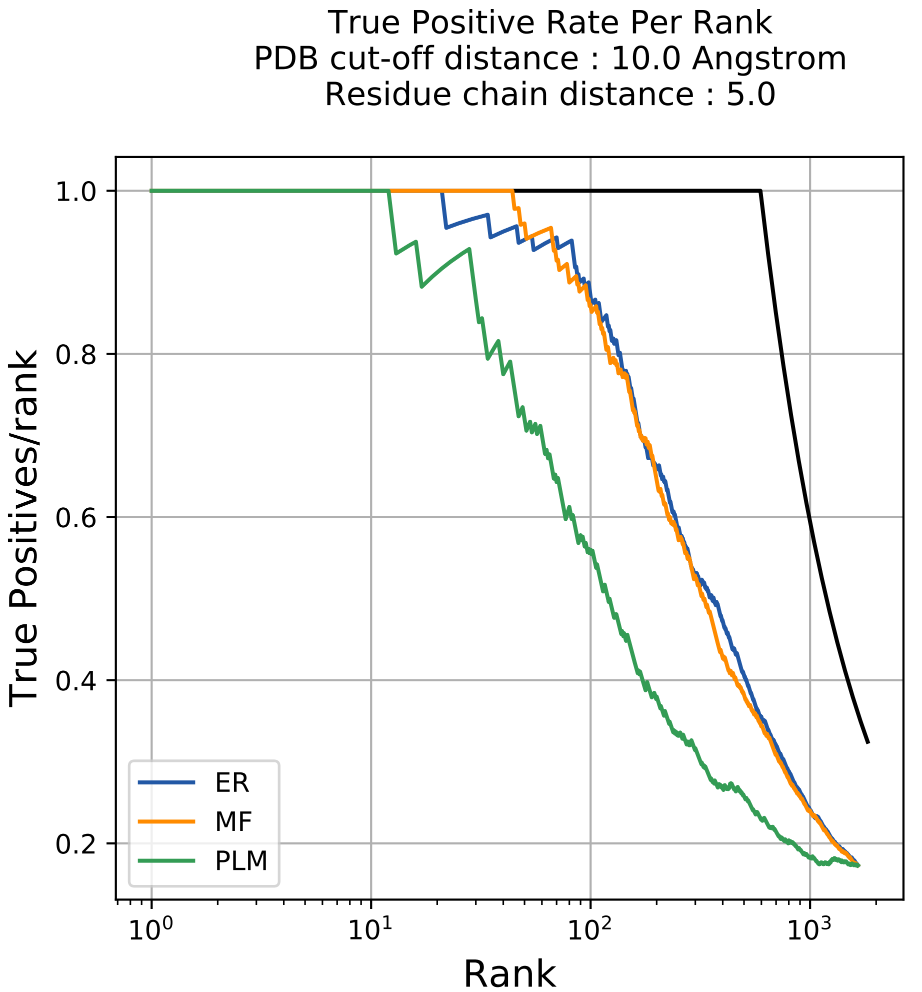

True Positive Rate Per Rank
PDB cut-off distance : 10.0 Angstrom
Residue chain distance : 5.0
True Positives/rank
Rank
1.0
0.8
0.6
0.4
0.2
$10^0$
$10^1$
$10^2$
$10^3$
ER
MF
PLM

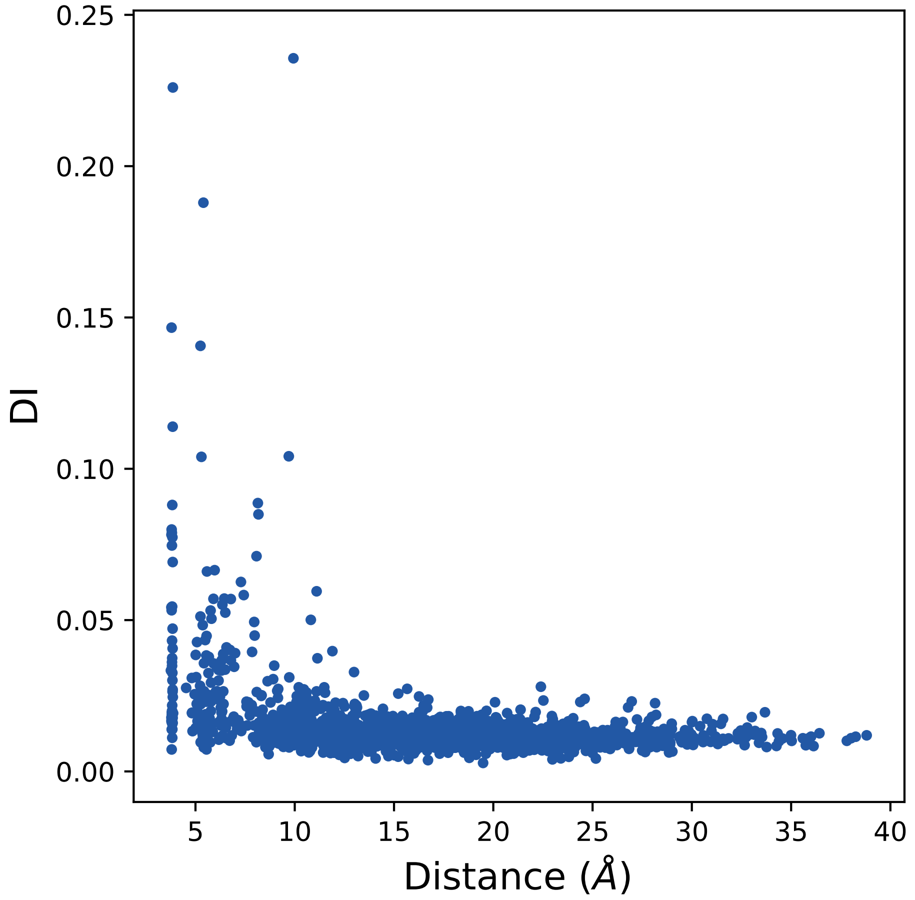

0.25
0.20
0.15
0.10
0.05
0.00
DI
5
10
15
20
25
30
35
40
Distance (Å)

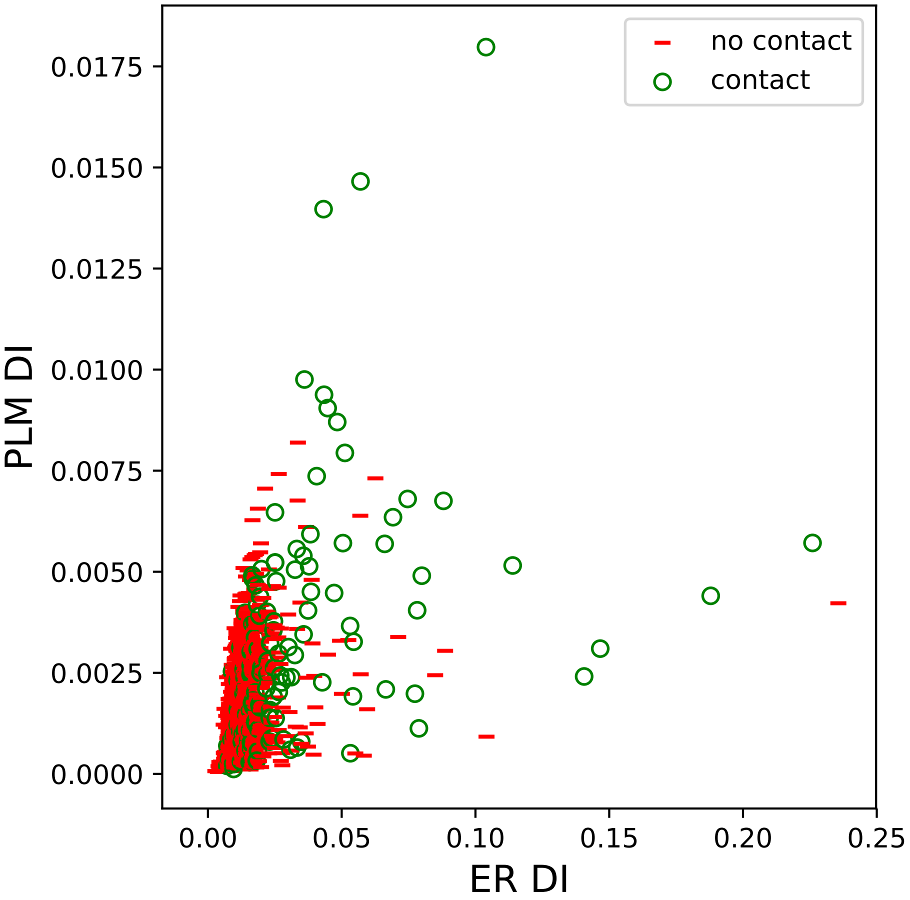

no contact
contact
PLM DI
ER DI
0.0000
0.0025
0.0050
0.0075
0.0100
0.0125
0.0150
0.0175
0.00
0.05
0.10
0.15
0.20
0.25

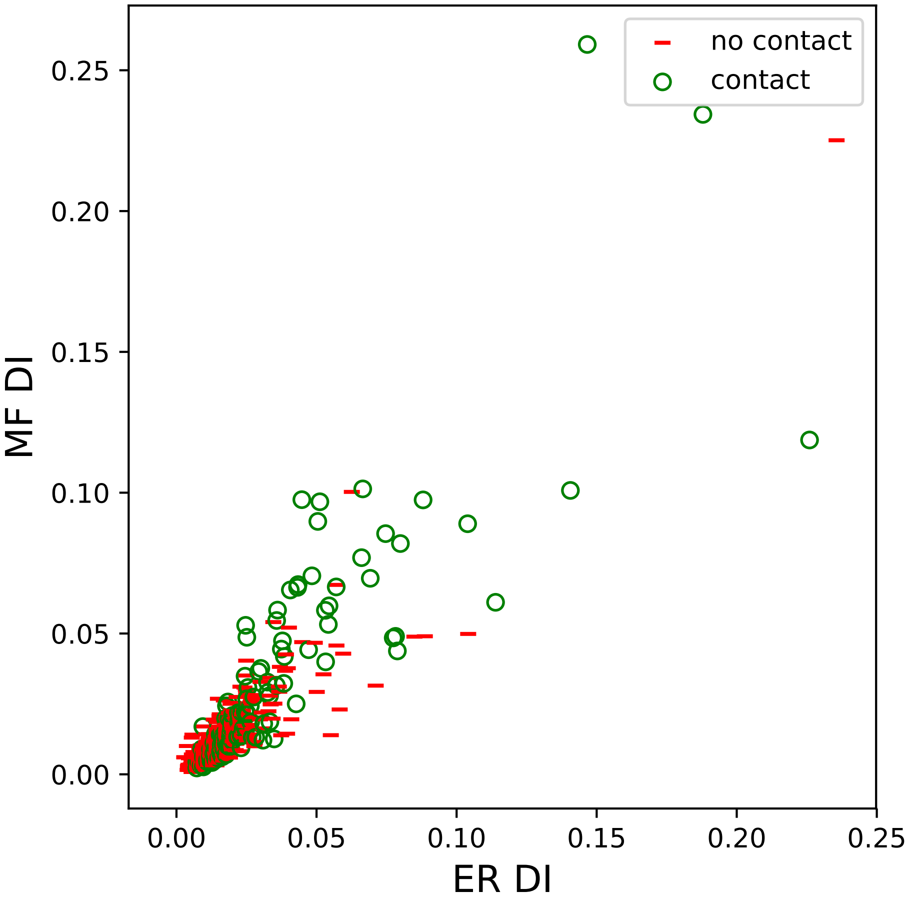

no contact
contact
MF DI
ER DI
0.00
0.05
0.10
0.15
0.20
0.25

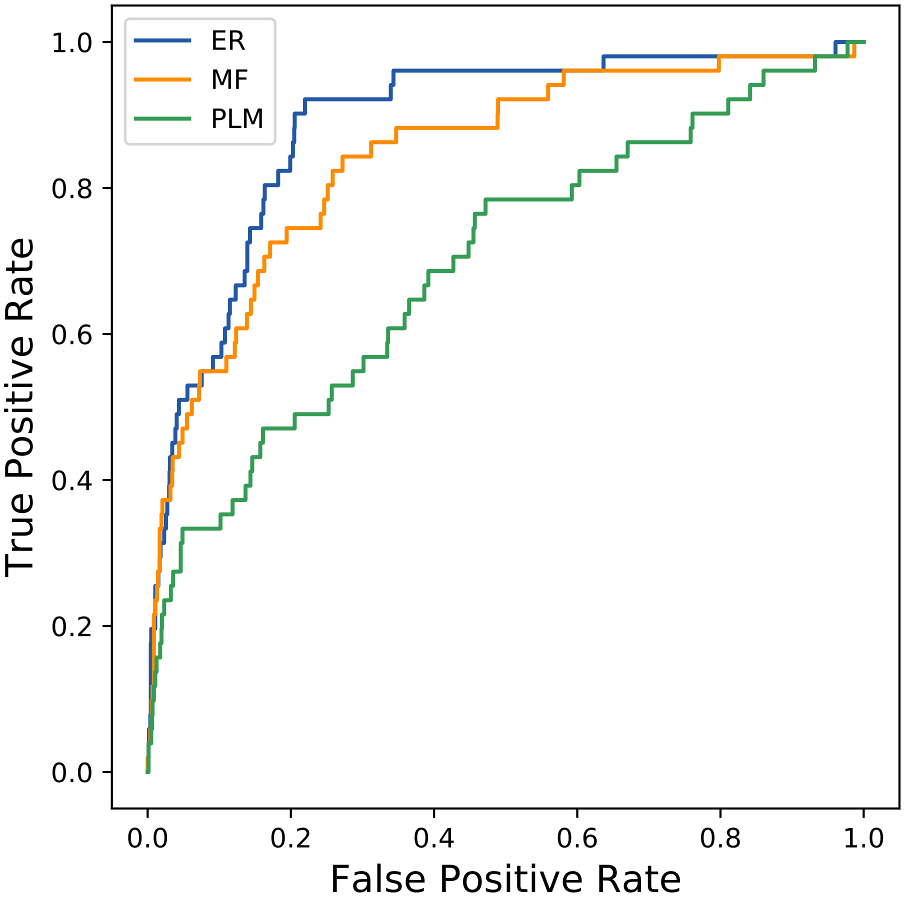

ER
MF
PLM
True Positive Rate
False Positive Rate
0.0
0.2
0.4
0.6
0.8
1.0
