## Supplemental Contact Maps for "Protein Structure Prediction with Expectation Reflection": 14408seq_75col_3il5_PF08541_plots.pdf

Method: ER
Maximum PDB contact distance : 10.0 Angstrom
Minimum residue chain distance: 5.0 residues
Fraction of true positives : 0.587

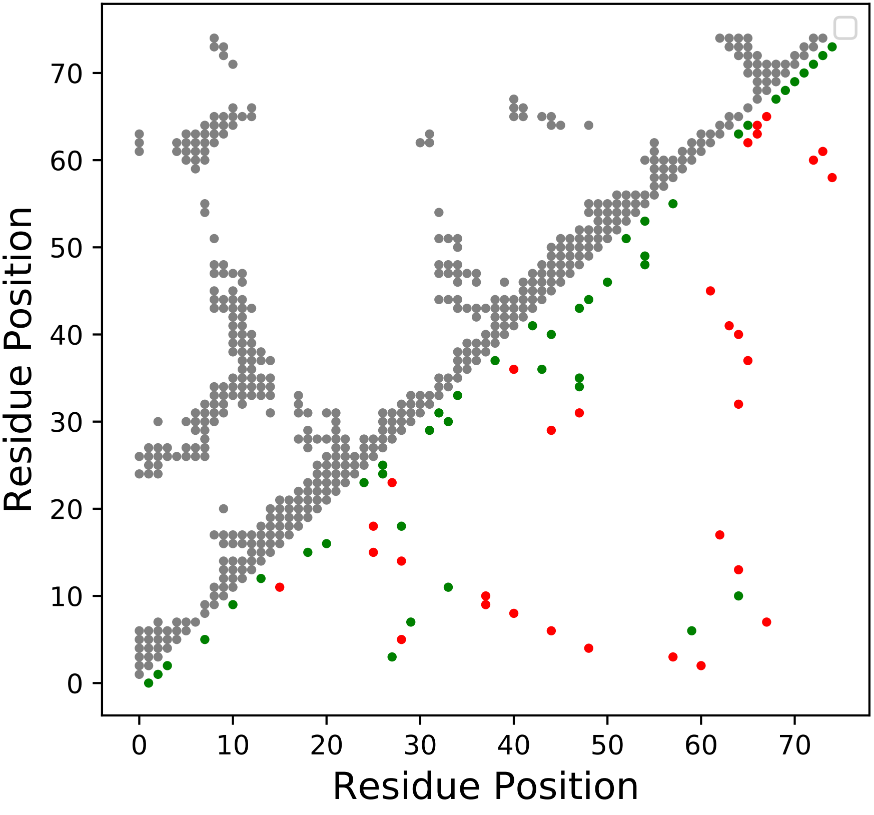

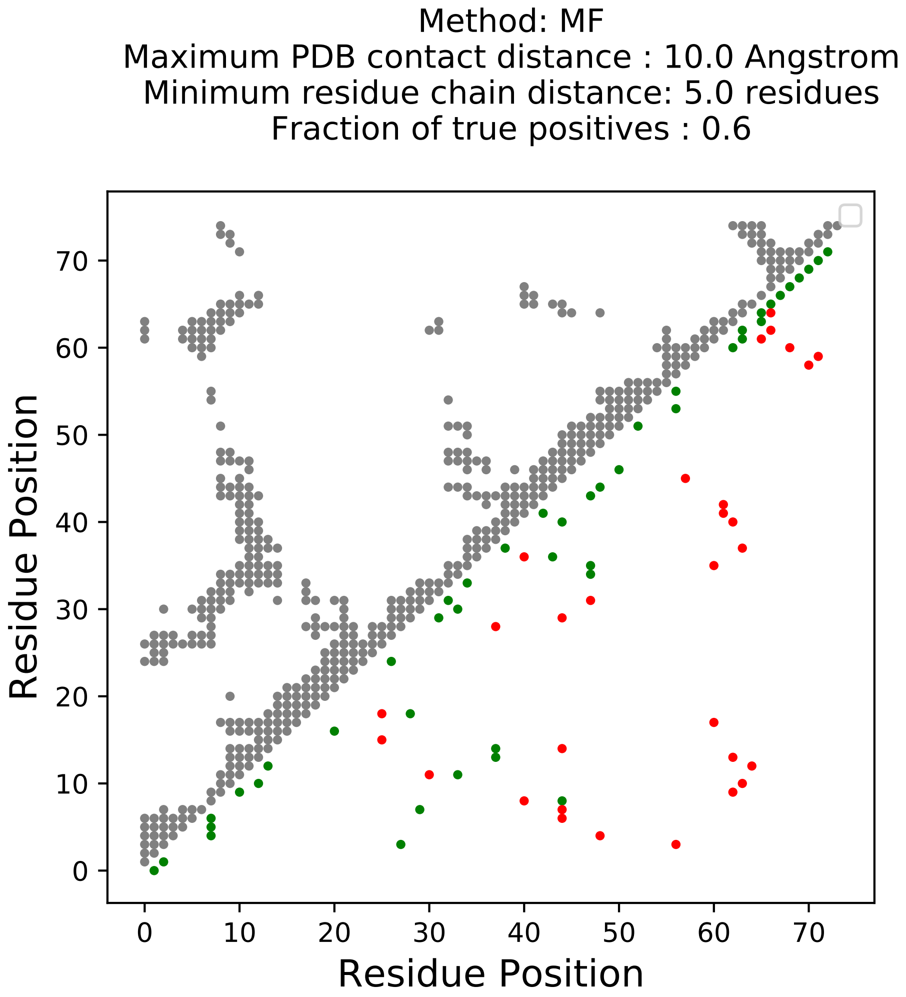
Method: MF
Maximum PDB contact distance : 10.0 Angstrom
Minimum residue chain distance: 5.0 residues
Fraction of true positives : 0.6
Residue Position
Residue Position
0
10
20
30
40
50
60
70

Method: PLM
Maximum PDB contact distance : 10.0 Angstrom
Minimum residue chain distance: 5.0 residues
Fraction of true positives : 0.627

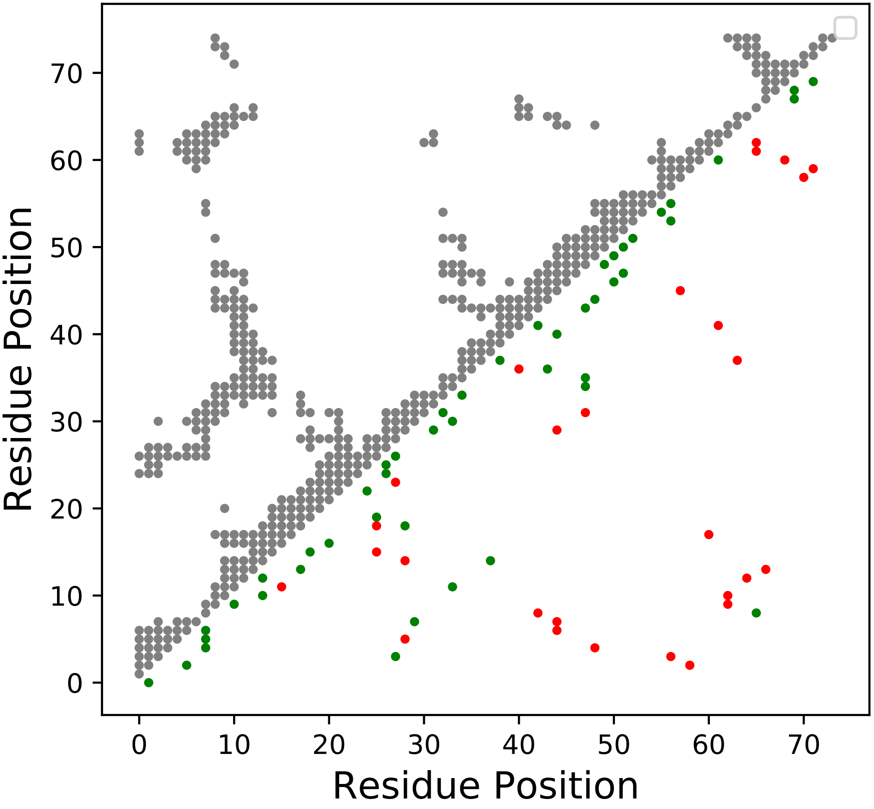

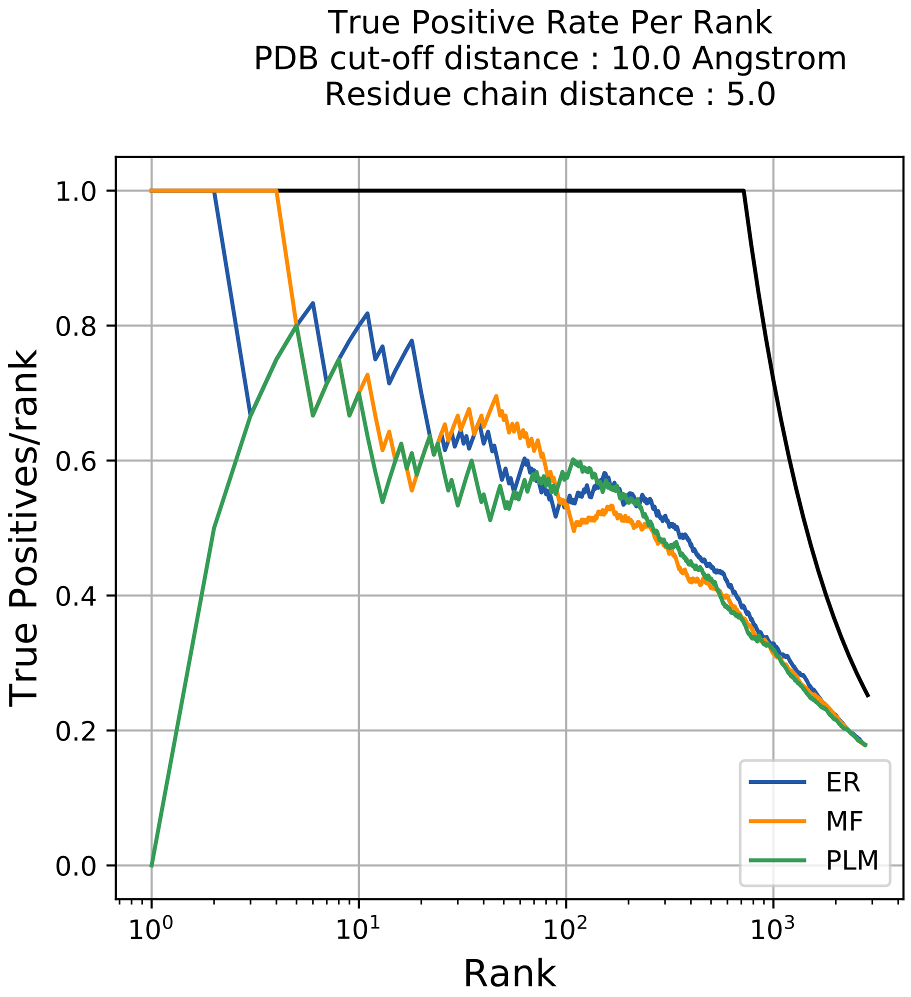
True Positive Rate Per Rank
PDB cut-off distance : 10.0 Angstrom
Residue chain distance : 5.0
True Positives/rank
Rank
1.0
0.8
0.6
0.4
0.2
0.0
$10^0$
$10^1$
$10^2$
$10^3$
ER
MF
PLM

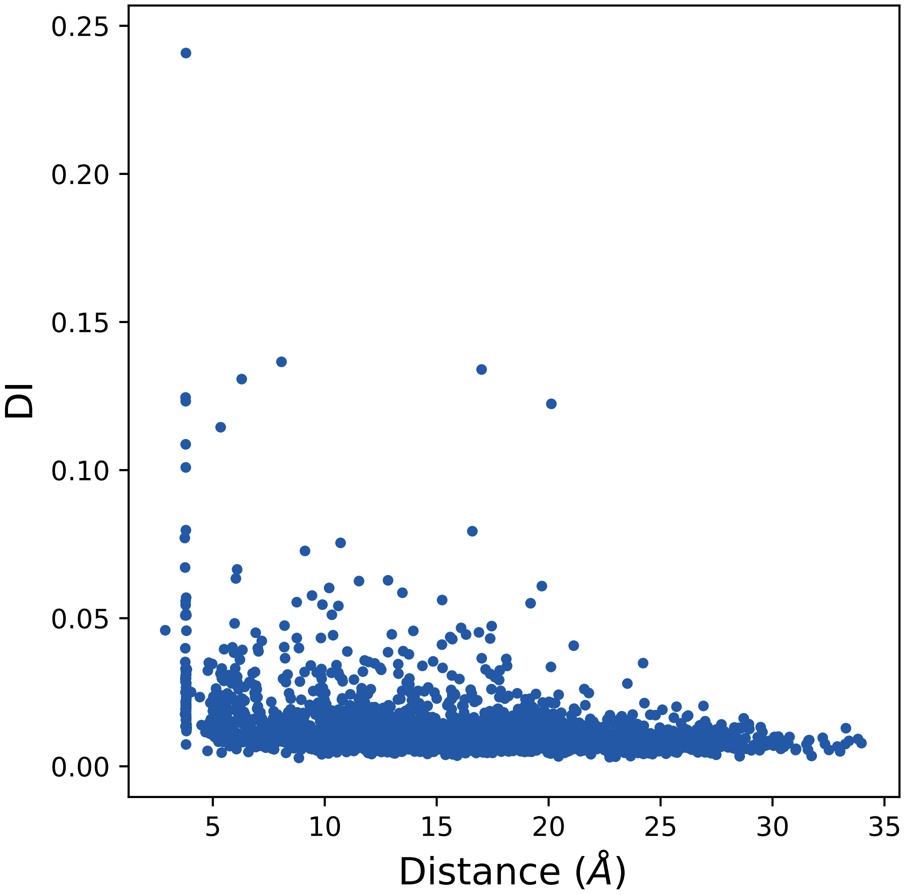
DI
Distance (Å)
0.00
0.05
0.10
0.15
0.20
0.25
5
10
15
20
25
30
35

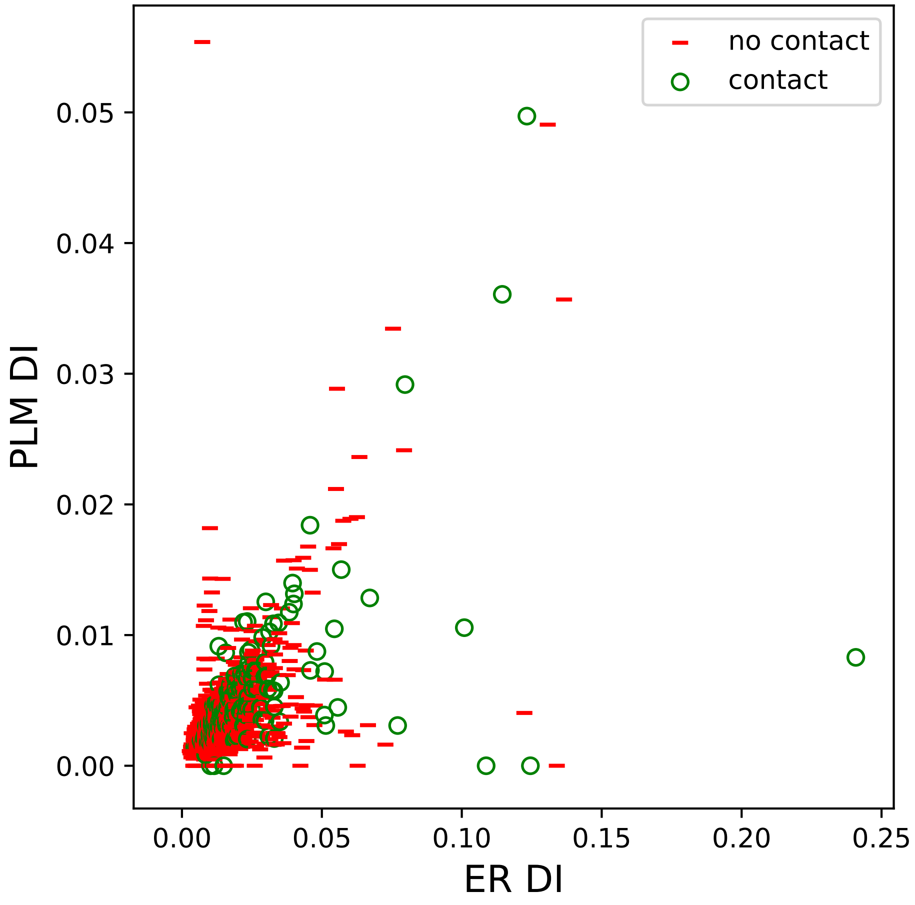

no contact
contact
PLM DI
ER DI
0.00
0.01
0.02
0.03
0.04
0.05
0.00
0.05
0.10
0.15
0.20
0.25

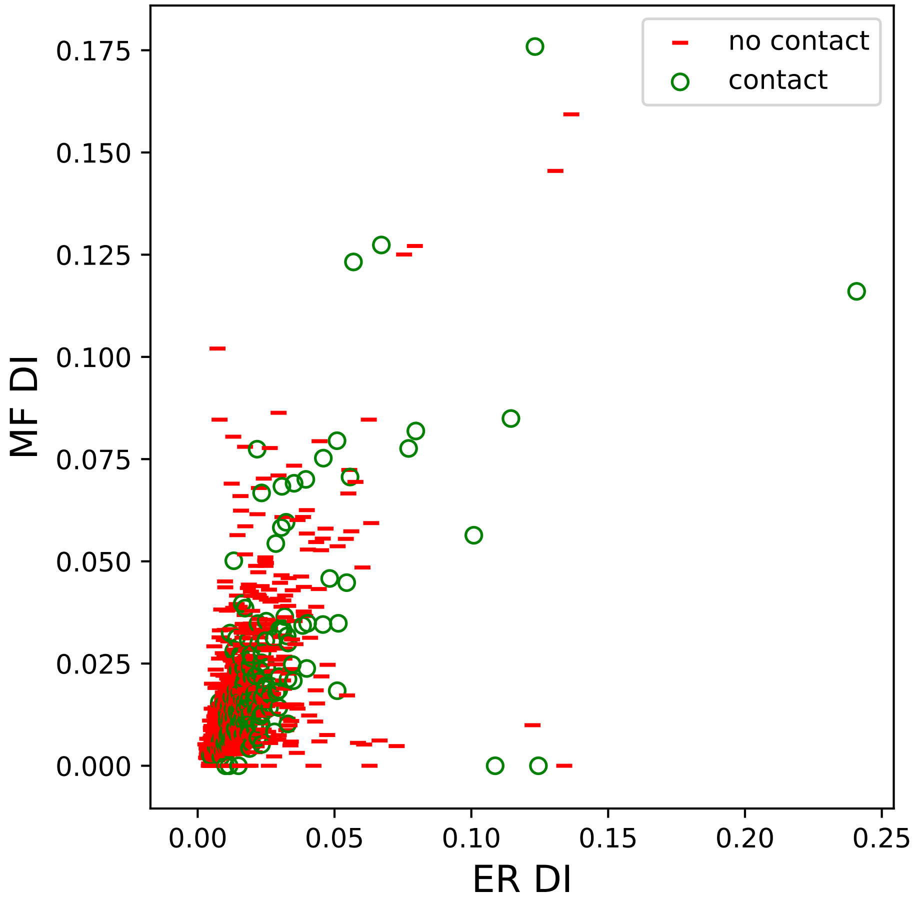

no contact
contact
MF DI
ER DI
0.000
0.025
0.050
0.075
0.100
0.125
0.150
0.175
0.00
0.05
0.10
0.15
0.20
0.25

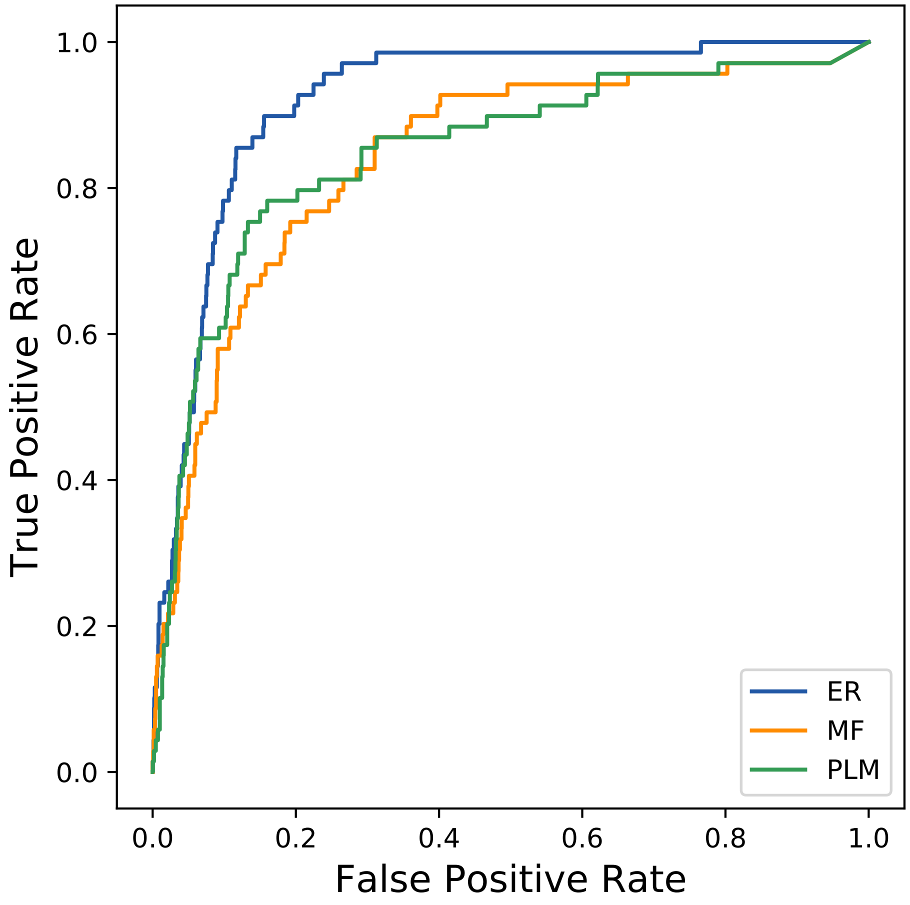
True Positive Rate
False Positive Rate
0.0
0.2
0.4
0.6
0.8
1.0
ER
MF
PLM
