## Supplemental Contact Maps for "Protein Structure Prediction with Expectation Reflection": 14704seq_125col_6io6_PF02800_plots.pdf

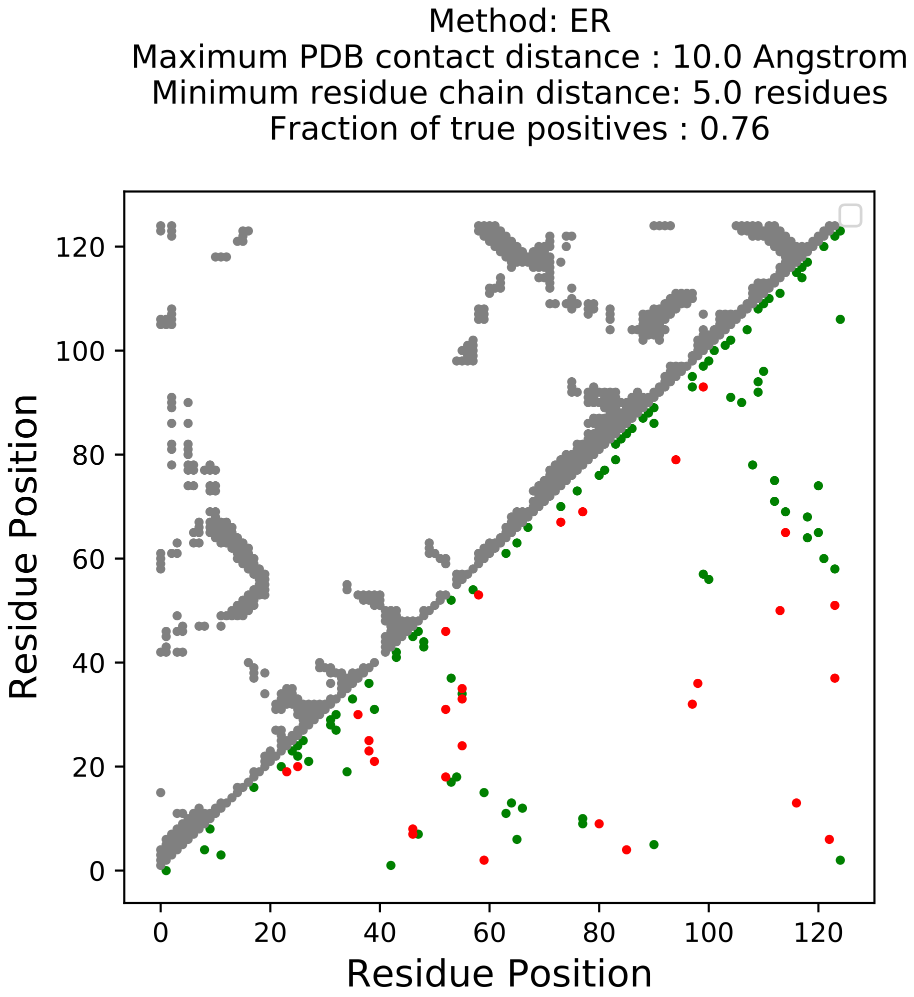
Method: ER
Maximum PDB contact distance : 10.0 Angstrom
Minimum residue chain distance: 5.0 residues
Fraction of true positives : 0.76
Residue Position
Residue Position
0
20
40
60
80
100
120

Method: MF
Maximum PDB contact distance : 10.0 Angstrom
Minimum residue chain distance: 5.0 residues
Fraction of true positives : 0.744

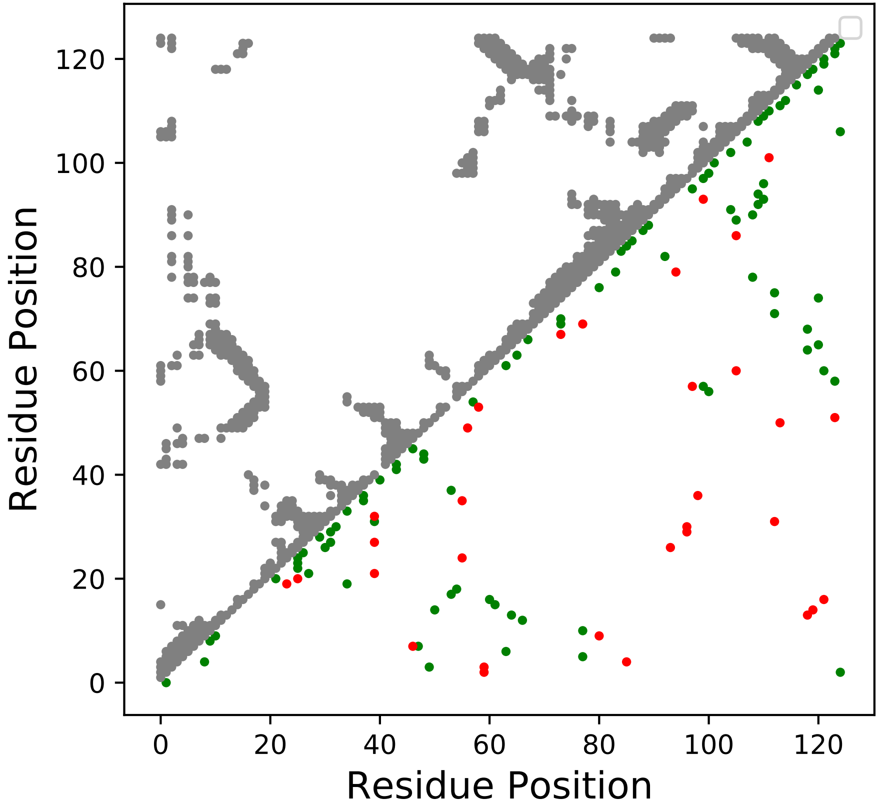

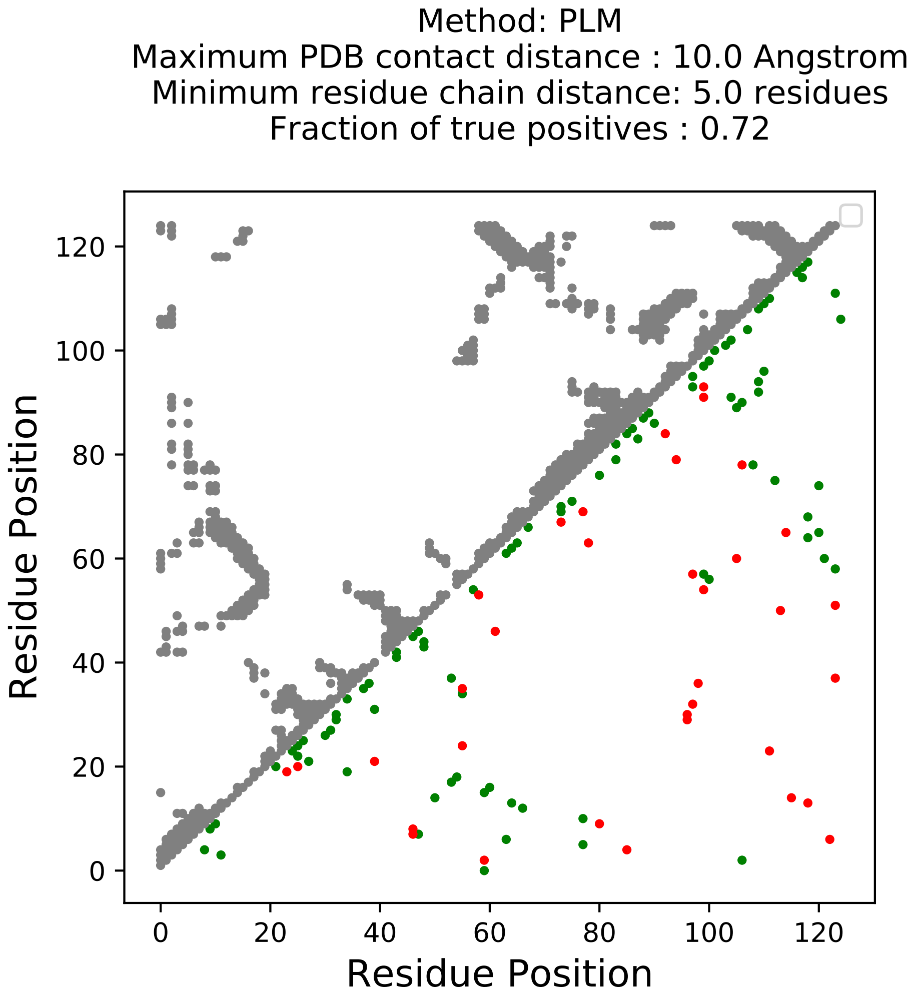
Method: PLM
Maximum PDB contact distance : 10.0 Angstrom
Minimum residue chain distance: 5.0 residues
Fraction of true positives : 0.72
Residue Position
Residue Position
0
20
40
60
80
100
120
0
20
40
60
80
100
120

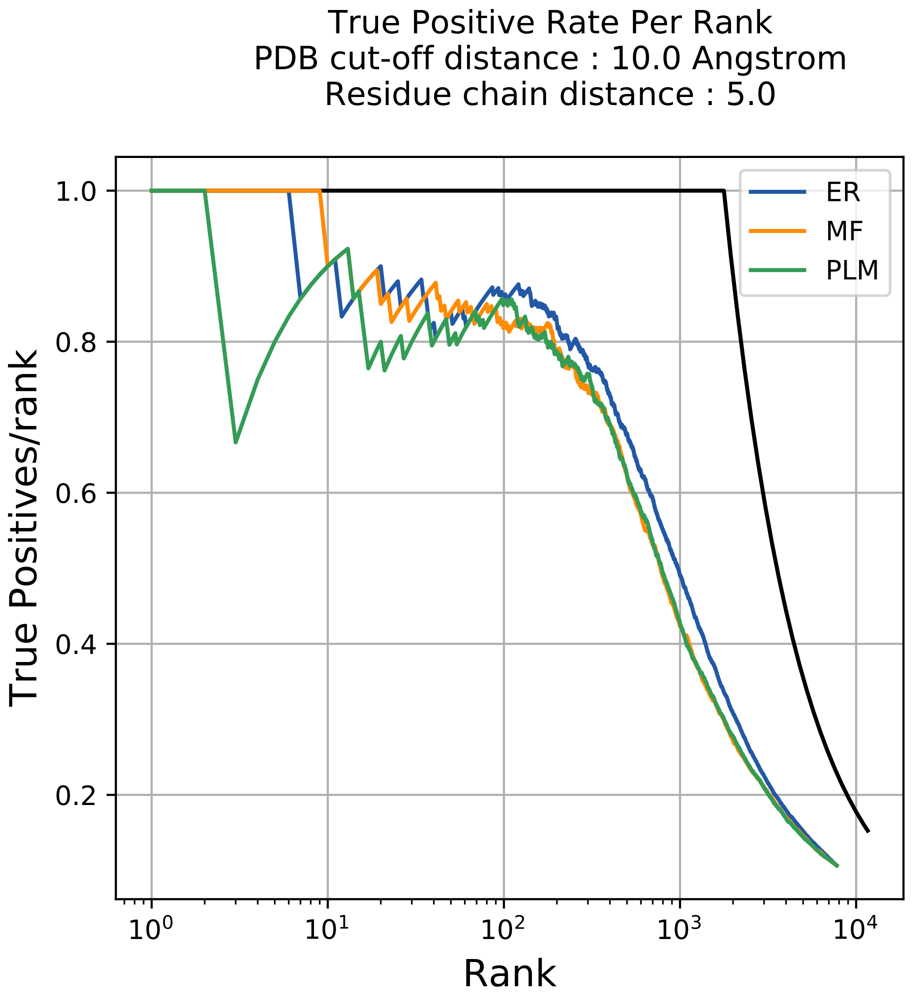
True Positive Rate Per Rank
PDB cut-off distance : 10.0 Angstrom
Residue chain distance : 5.0
ER
MF
PLM
1.0
0.8
0.6
0.4
0.2
True Positives/rank
$10^0$
$10^1$
$10^2$
$10^3$
$10^4$
Rank

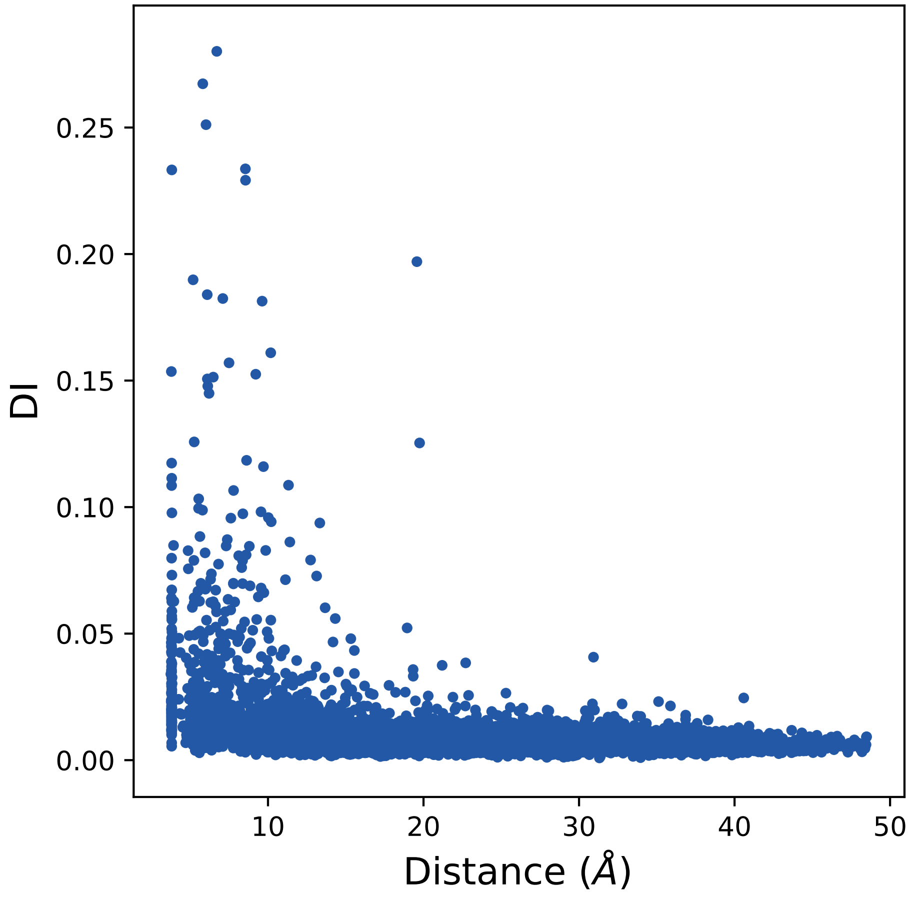
DI
0.25
0.20
0.15
0.10
0.05
0.00
10
20
30
40
50
Distance (Å)

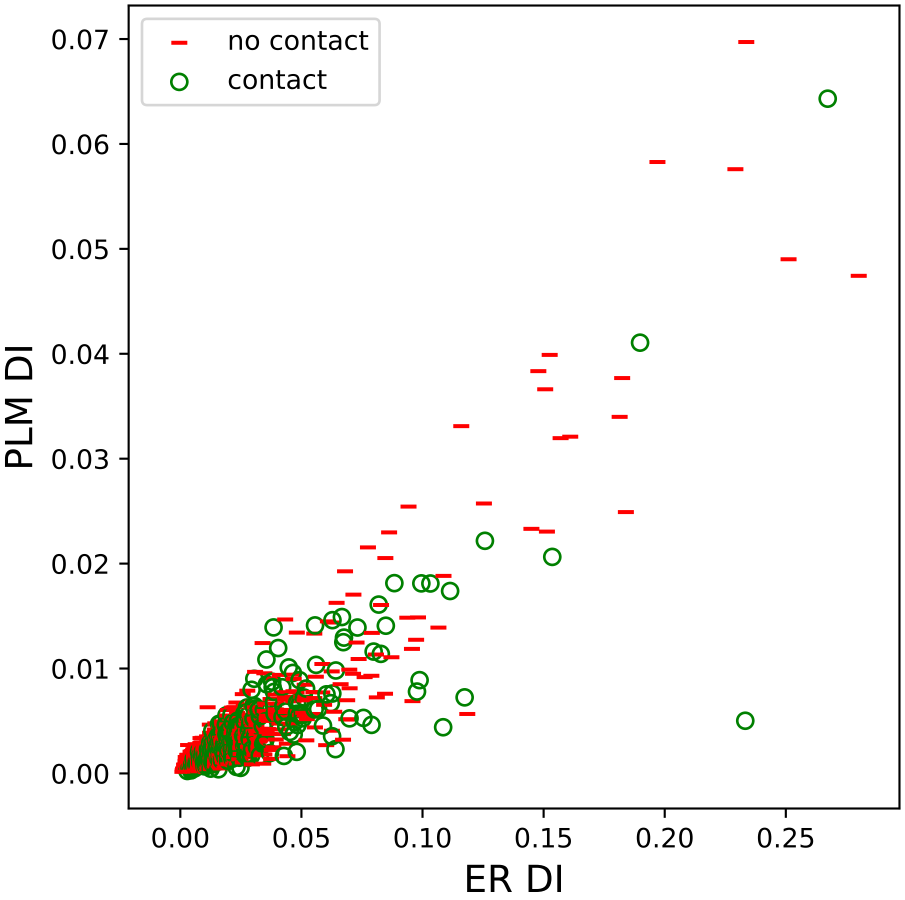

no contact
contact
PLM DI
ER DI
0.00
0.01
0.02
0.03
0.04
0.05
0.06
0.07
0.00
0.05
0.10
0.15
0.20
0.25

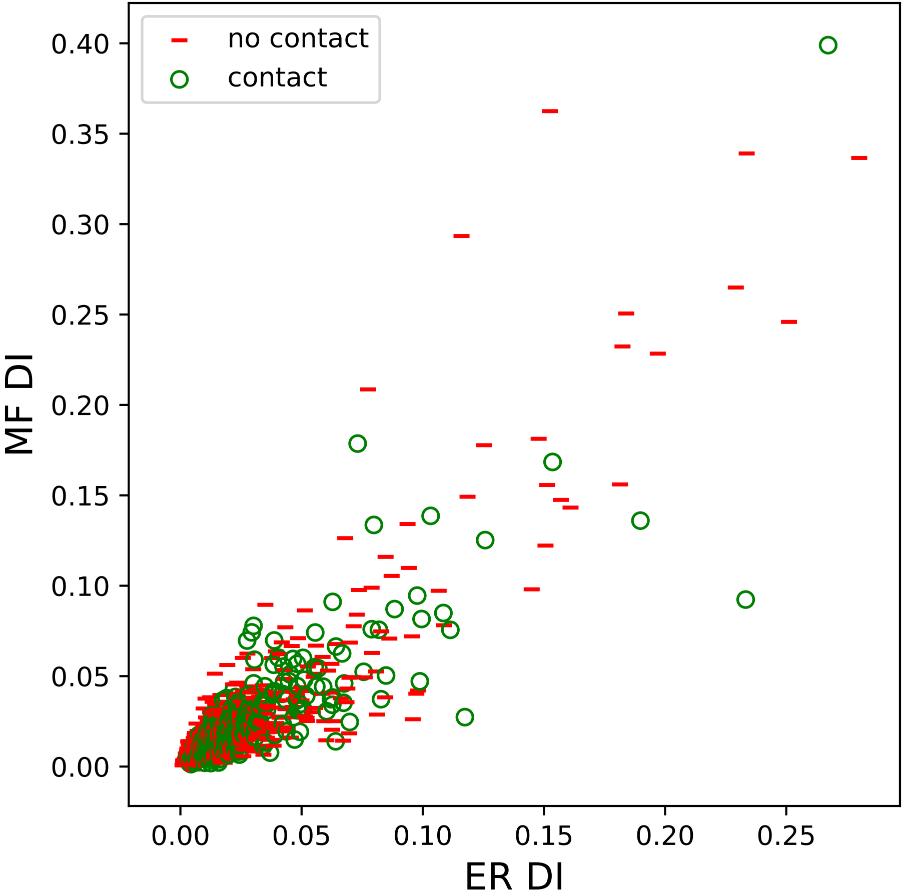

no contact
contact
0.40
0.35
0.30
0.25
0.20
0.15
0.10
0.05
0.00
MF DI
0.00
0.05
0.10
0.15
0.20
0.25
ER DI

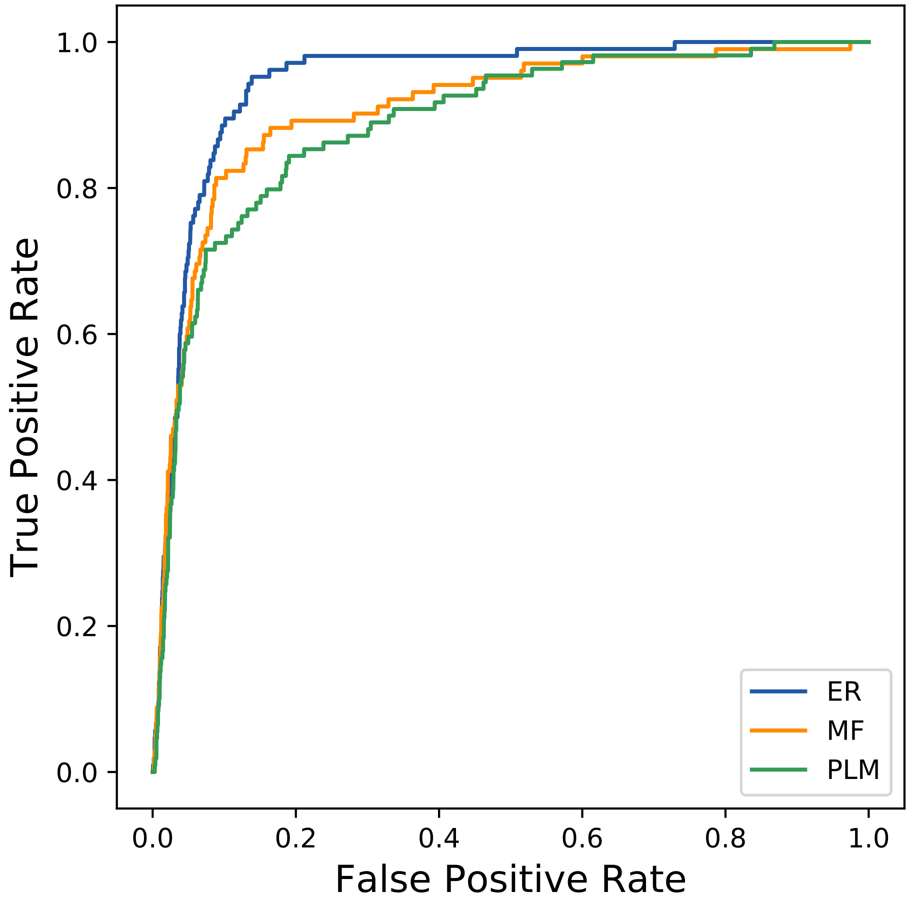

True Positive Rate
False Positive Rate
0.0
0.2
0.4
0.6
0.8
1.0
ER
MF
PLM
