## Supplemental Contact Maps for "Protein Structure Prediction with Expectation Reflection": 14787seq_179col_4rvb_PF01433_plots.pdf

Method: ER
Maximum PDB contact distance : 10.0 Angstrom
Minimum residue chain distance: 5.0 residues
Fraction of true positives : 0.816

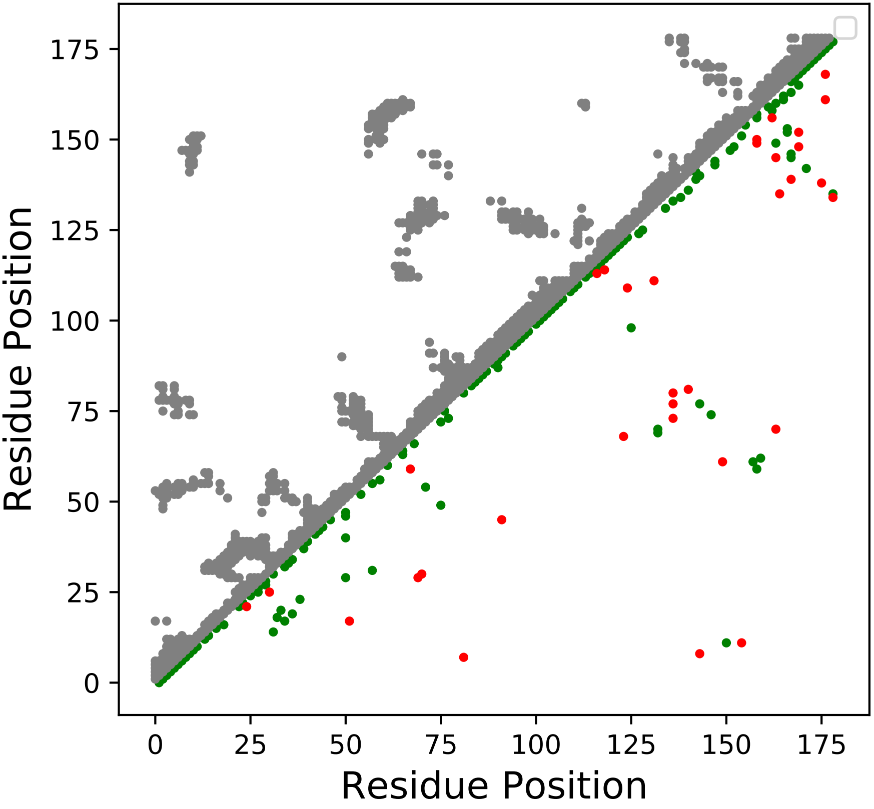

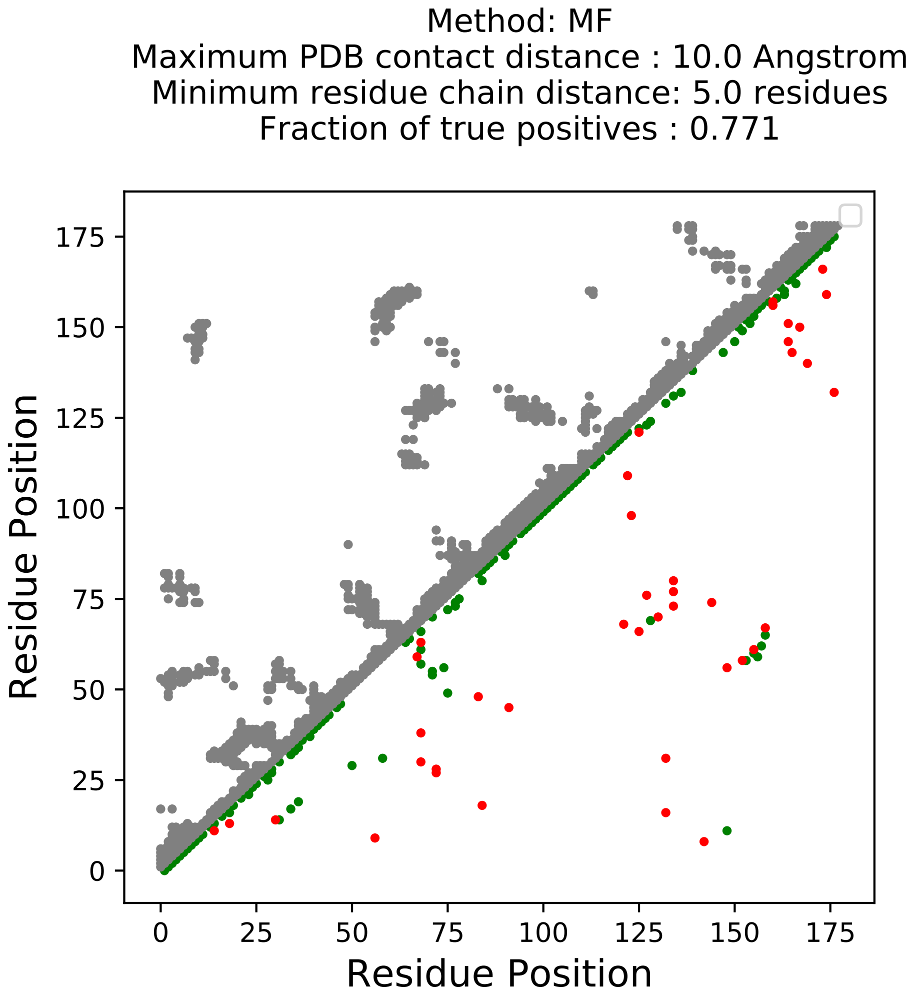
Method: MF
Maximum PDB contact distance : 10.0 Angstrom
Minimum residue chain distance: 5.0 residues
Fraction of true positives : 0.771
Residue Position
Residue Position
0
25
50
75
100
125
150
175

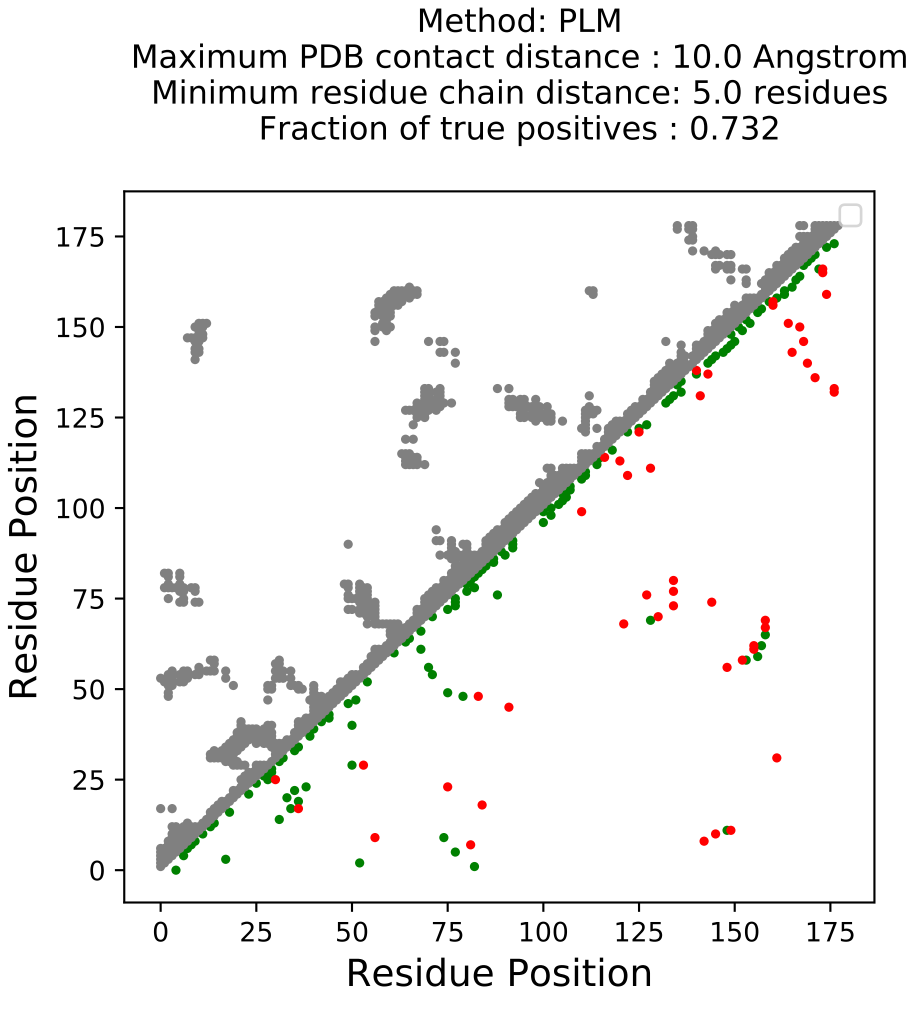
Method: PLM
Maximum PDB contact distance : 10.0 Angstrom
Minimum residue chain distance: 5.0 residues
Fraction of true positives : 0.732
Residue Position
Residue Position
0
25
50
75
100
125
150
175

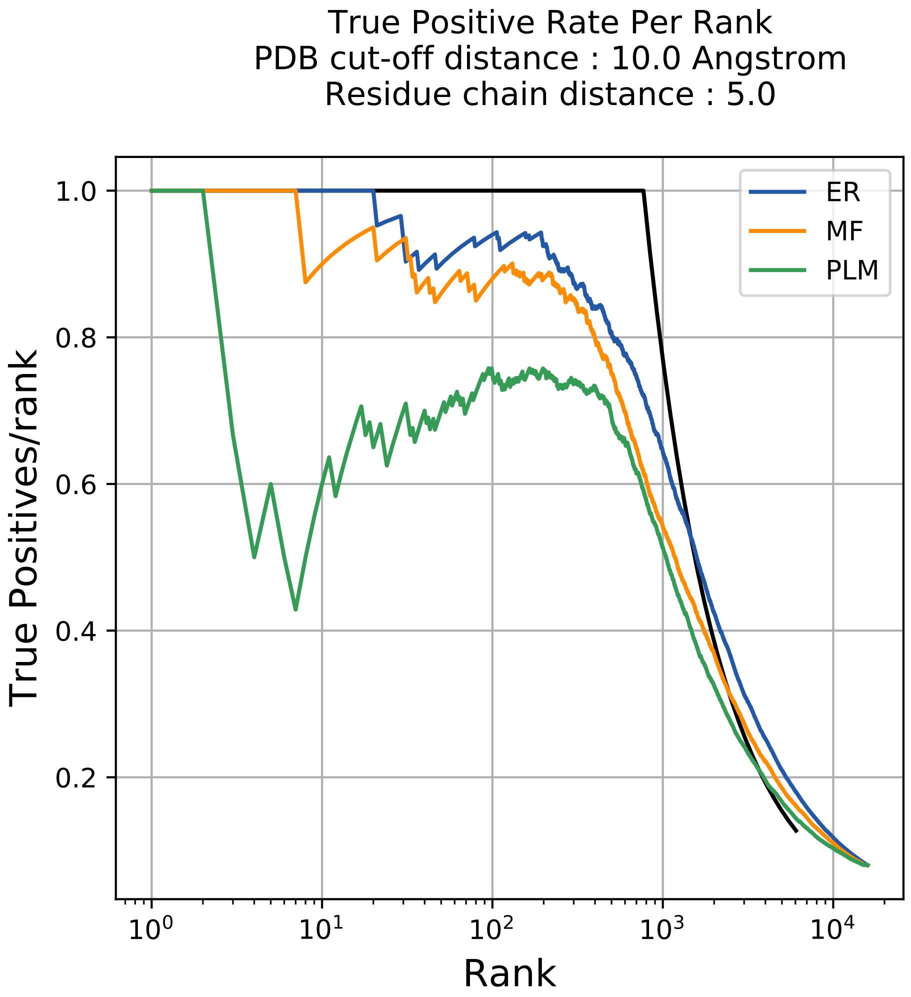
True Positive Rate Per Rank
PDB cut-off distance : 10.0 Angstrom
Residue chain distance : 5.0
ER
MF
PLM
1.0
0.8
0.6
0.4
0.2
True Positives/rank
$10^0$
$10^1$
$10^2$
$10^3$
$10^4$
Rank

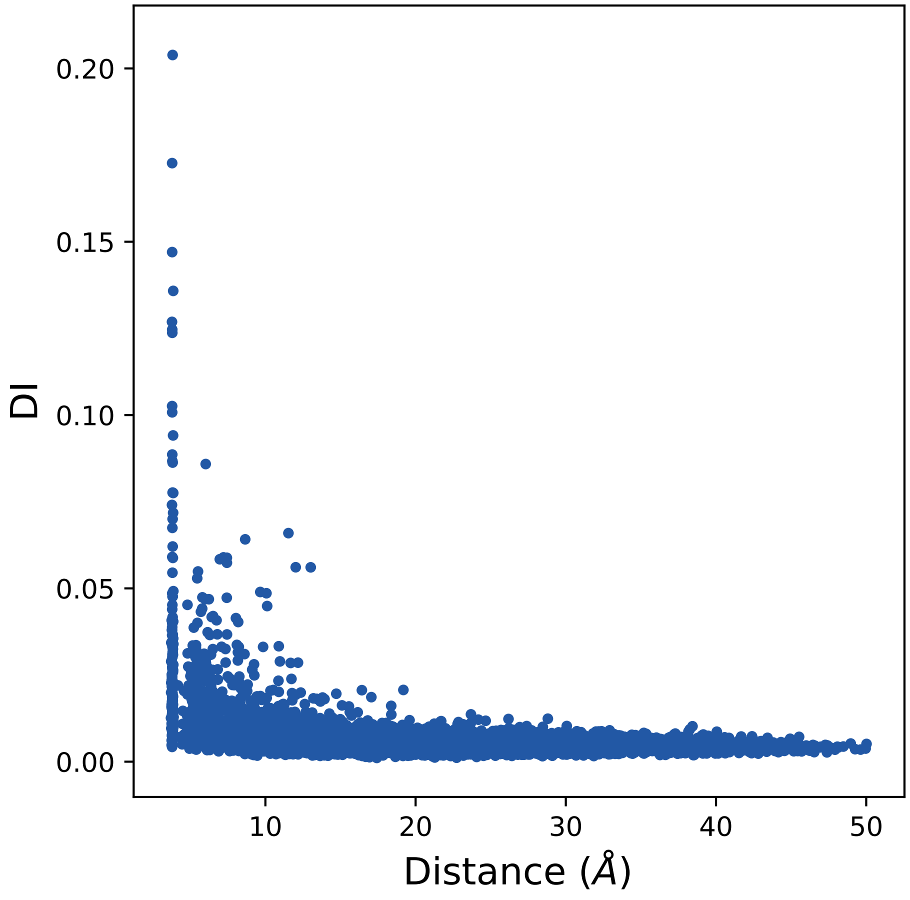

DI
Distance ($\AA$)
0.20
0.15
0.10
0.05
0.00
10
20
30
40
50

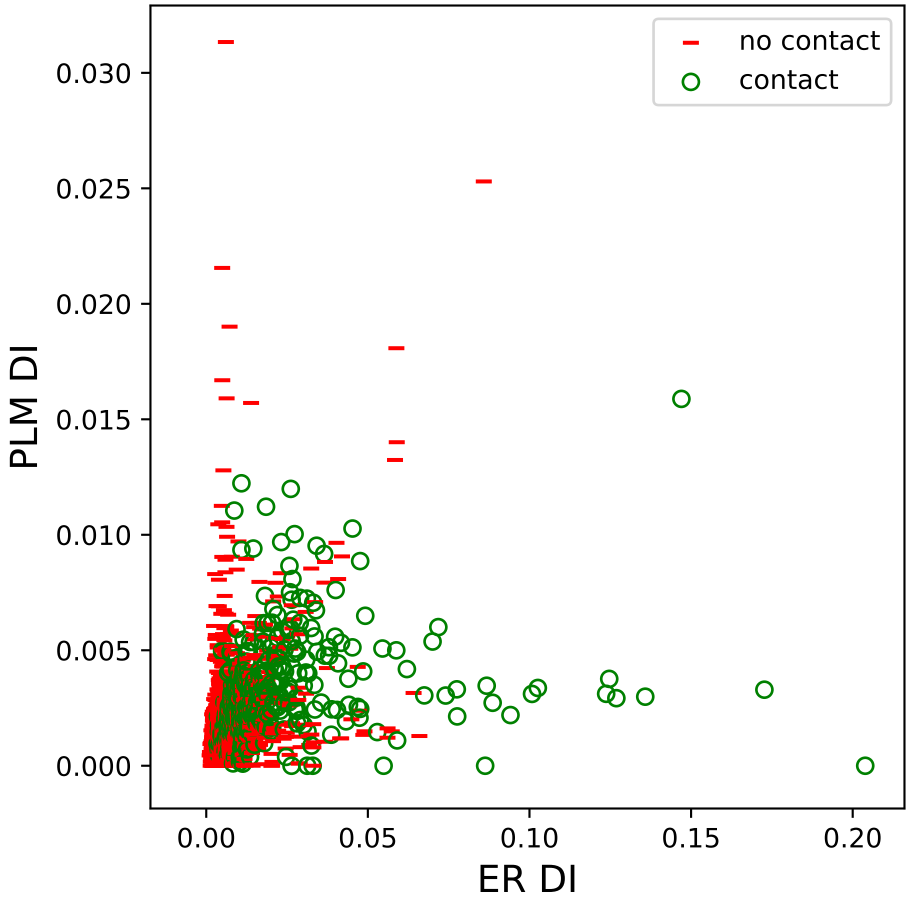

no contact
contact
PLM DI
ER DI
0.030
0.025
0.020
0.015
0.010
0.005
0.000
0.00
0.05
0.10
0.15
0.20

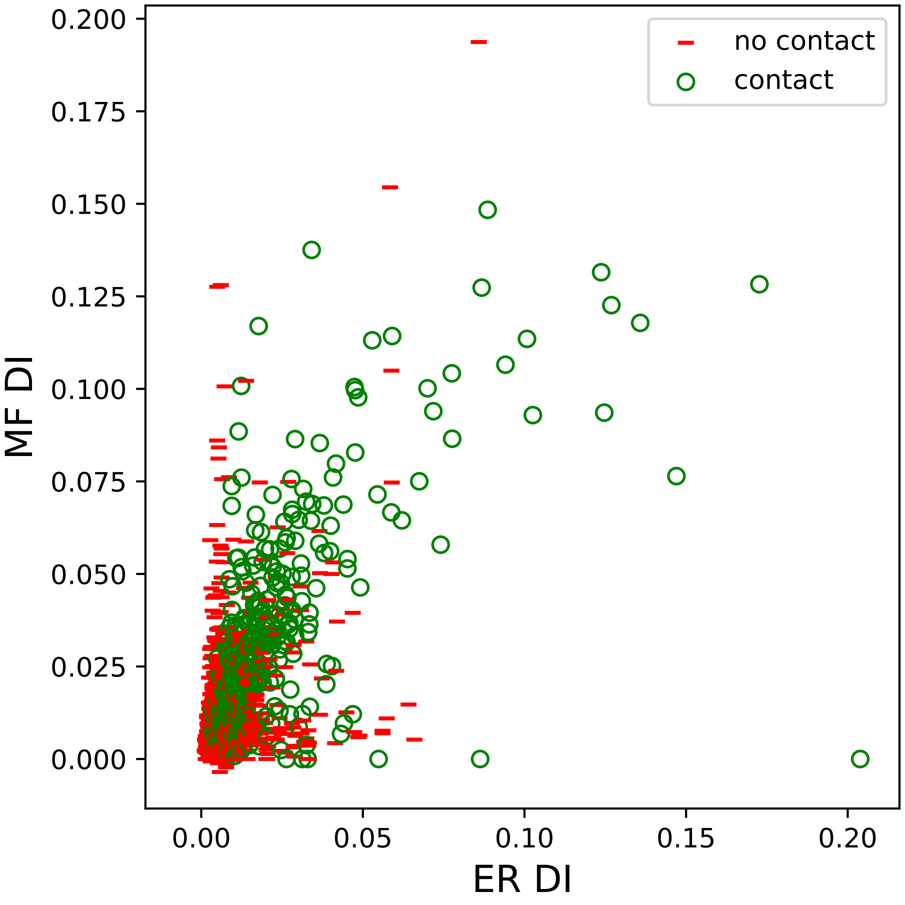

no contact
contact
MF DI
ER DI
0.000
0.025
0.050
0.075
0.100
0.125
0.150
0.175
0.200
0.00
0.05
0.10
0.15
0.20

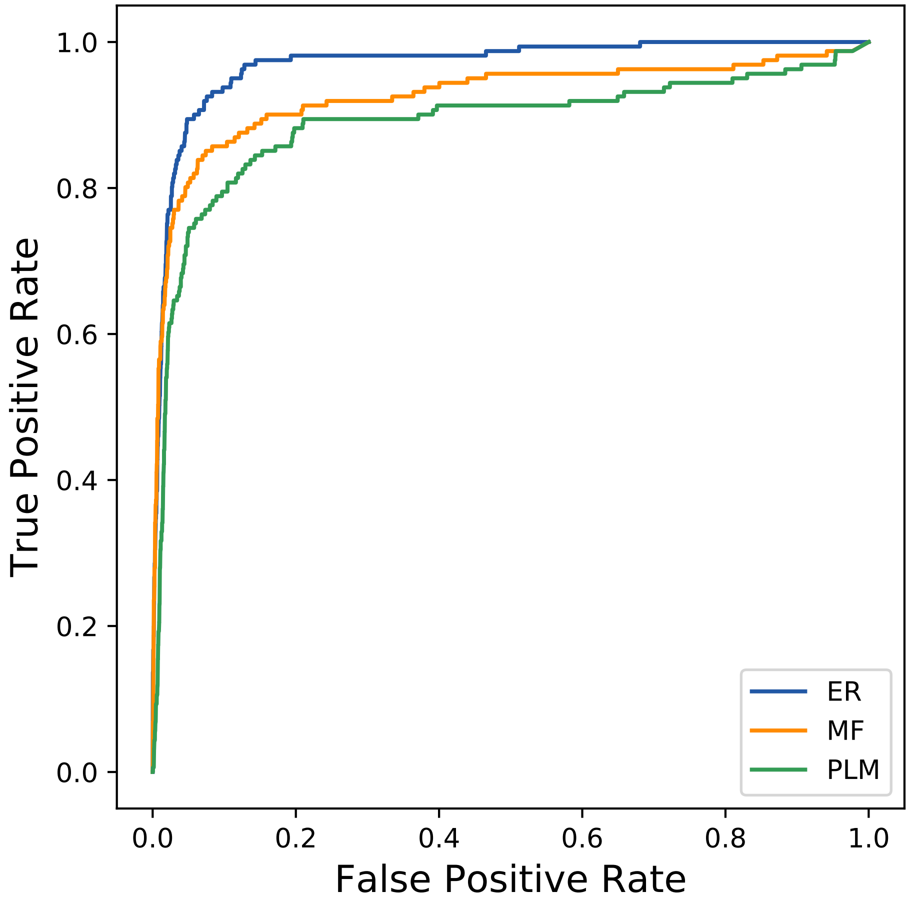

True Positive Rate
False Positive Rate
0.0
0.2
0.4
0.6
0.8
1.0
ER
MF
PLM
