## Supplemental Contact Maps for "Protein Structure Prediction with Expectation Reflection": 15484seq_248col_6r5i_PF00933_plots.pdf

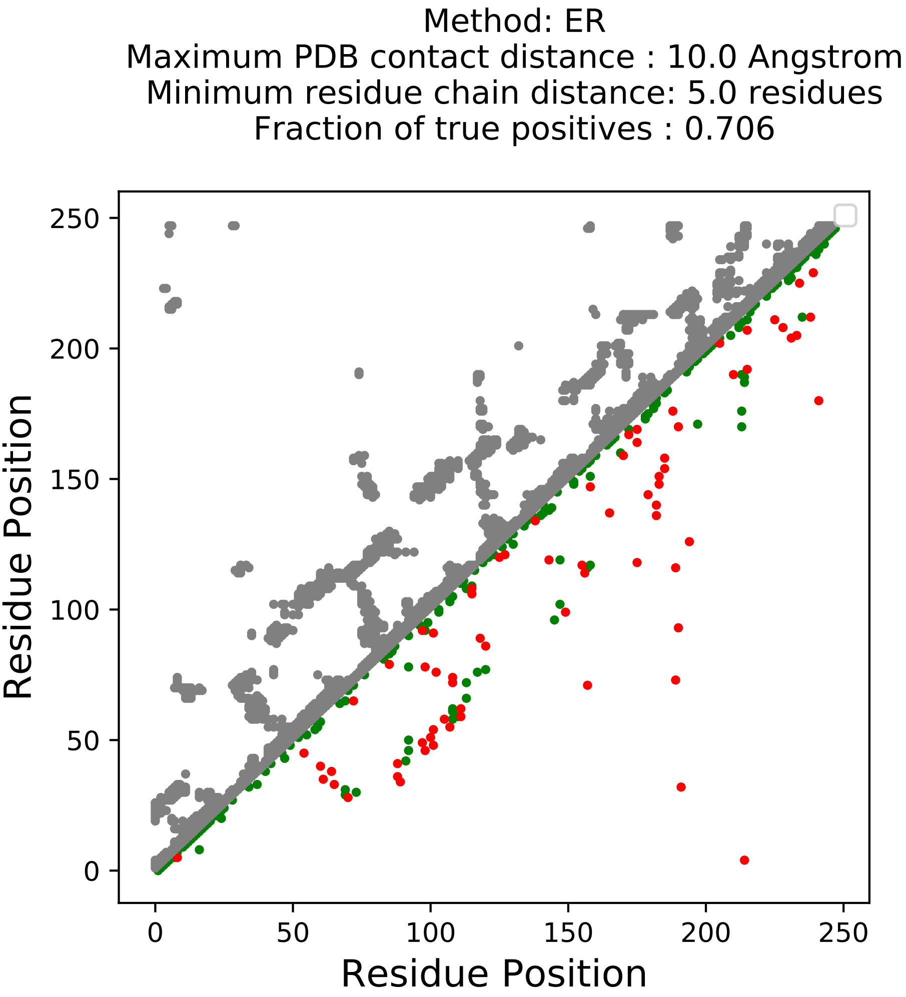
Method: ER
Maximum PDB contact distance : 10.0 Angstrom
Minimum residue chain distance: 5.0 residues
Fraction of true positives : 0.706
Residue Position
Residue Position
0
50
100
150
200
250
0
50
100
150
200
250

Method: MF
Maximum PDB contact distance : 10.0 Angstrom
Minimum residue chain distance: 5.0 residues
Fraction of true positives : 0.661

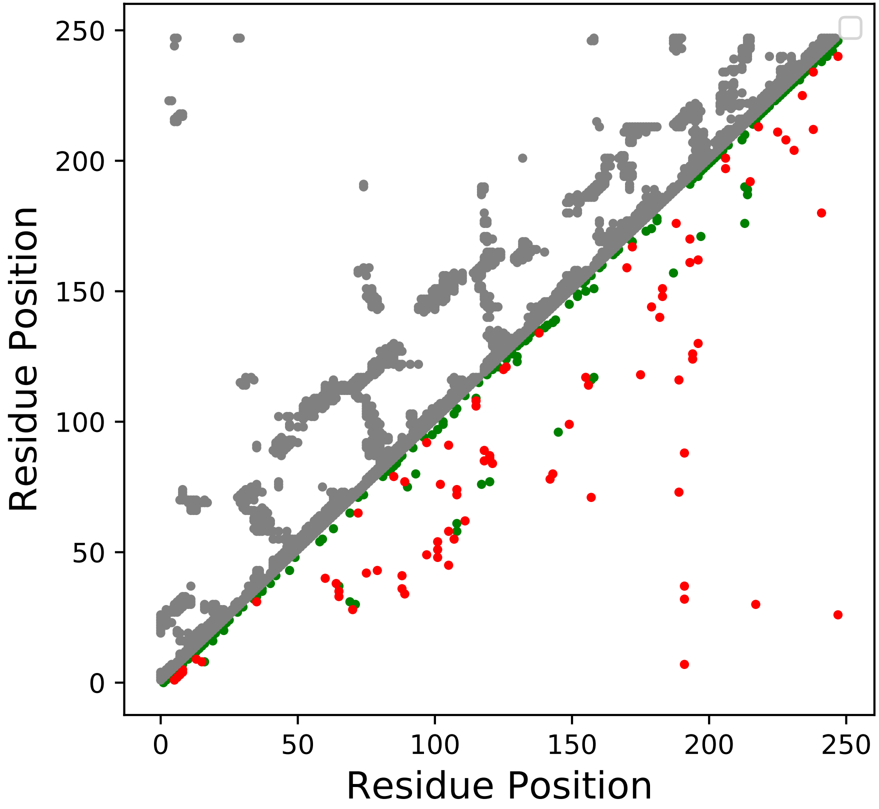

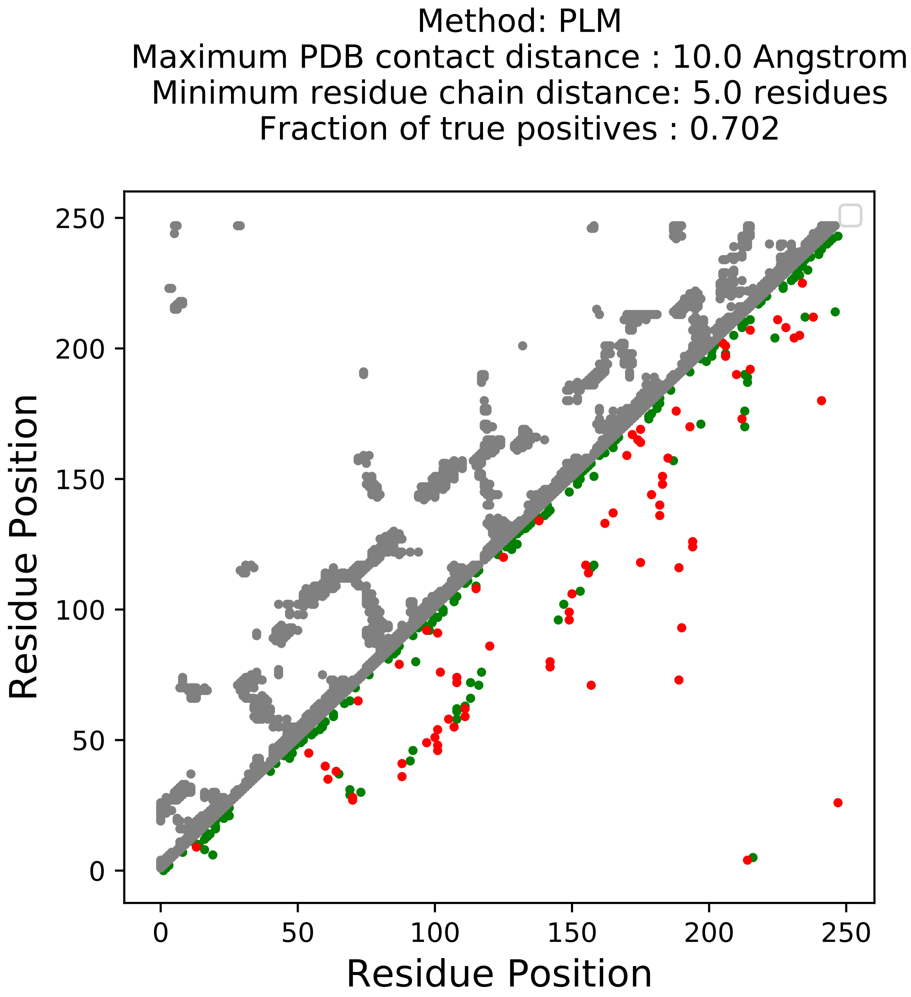
Method: PLM
Maximum PDB contact distance : 10.0 Angstrom
Minimum residue chain distance: 5.0 residues
Fraction of true positives : 0.702
Residue Position
Residue Position
0
50
100
150
200
250

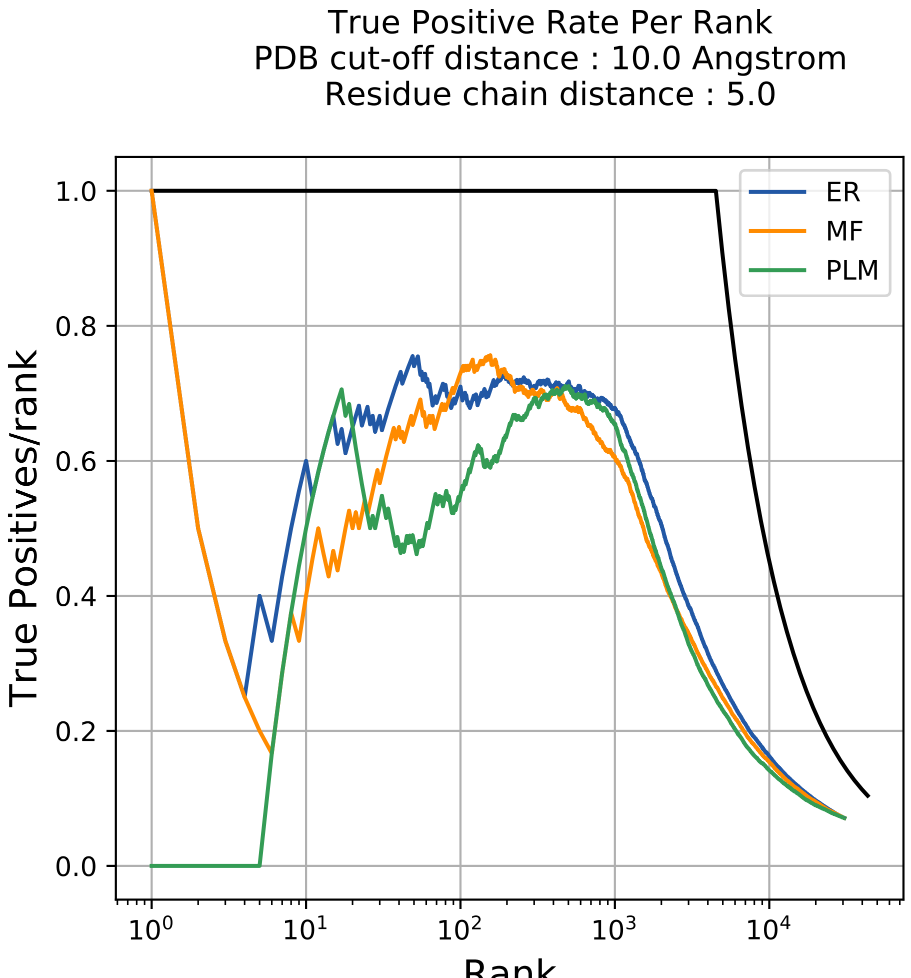
True Positive Rate Per Rank
PDB cut-off distance : 10.0 Angstrom
Residue chain distance : 5.0
True Positives/rank
Rank
ER
MF
PLM
1.0
0.8
0.6
0.4
0.2
0.0
$10^0$
$10^1$
$10^2$
$10^3$
$10^4$

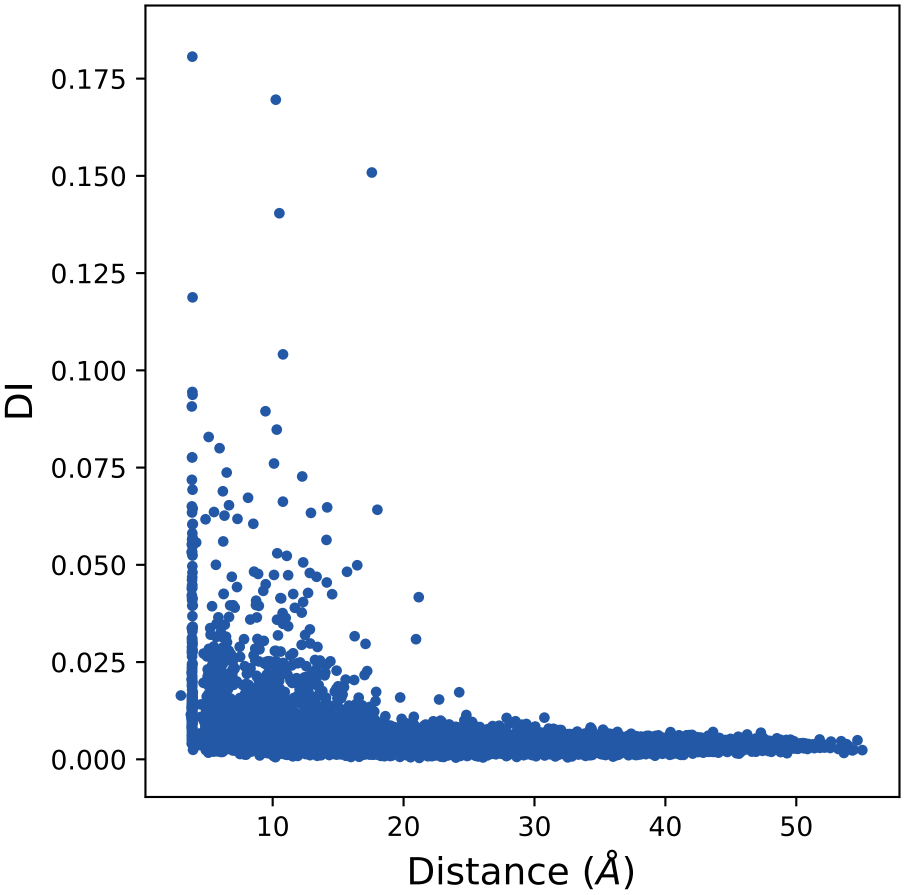

0.175
0.150
0.125
0.100
0.075
0.050
0.025
0.000
DI
10
20
30
40
50
Distance (Å)

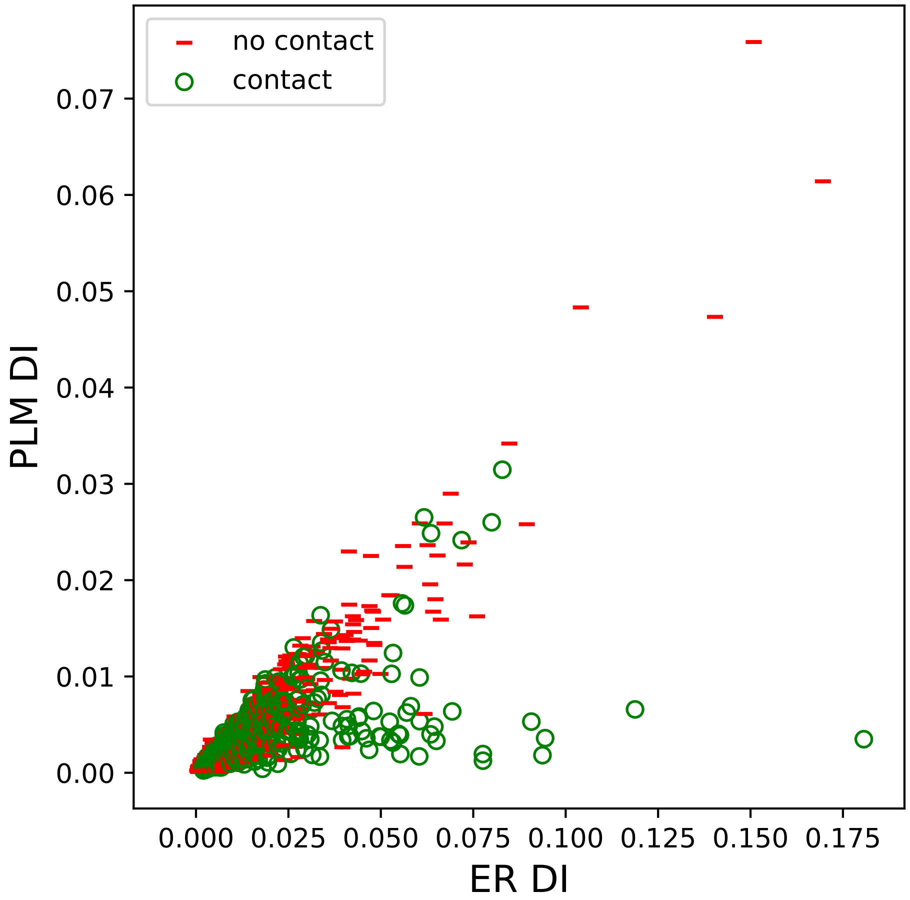
no contact
contact
PLM DI
ER DI
0.00
0.01
0.02
0.03
0.04
0.05
0.06
0.07
0.000
0.025
0.050
0.075
0.100
0.125
0.150
0.175

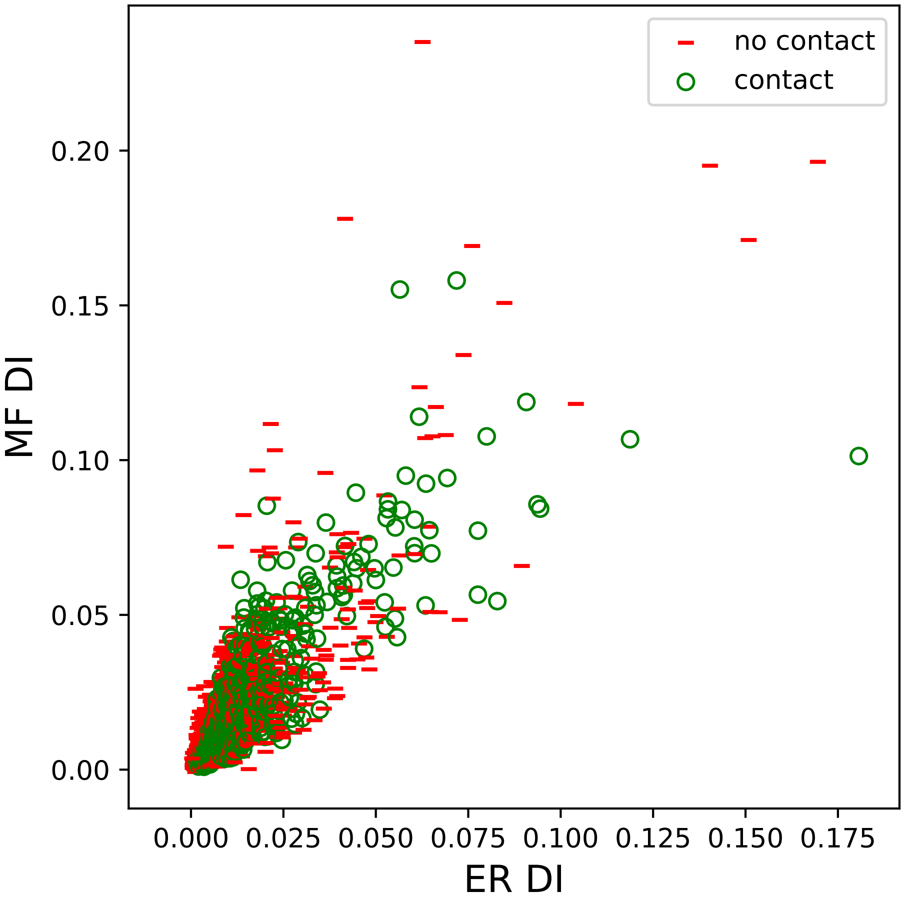

no contact
contact
0.20
0.15
0.10
0.05
0.00
MF DI
0.000
0.025
0.050
0.075
0.100
0.125
0.150
0.175
ER DI

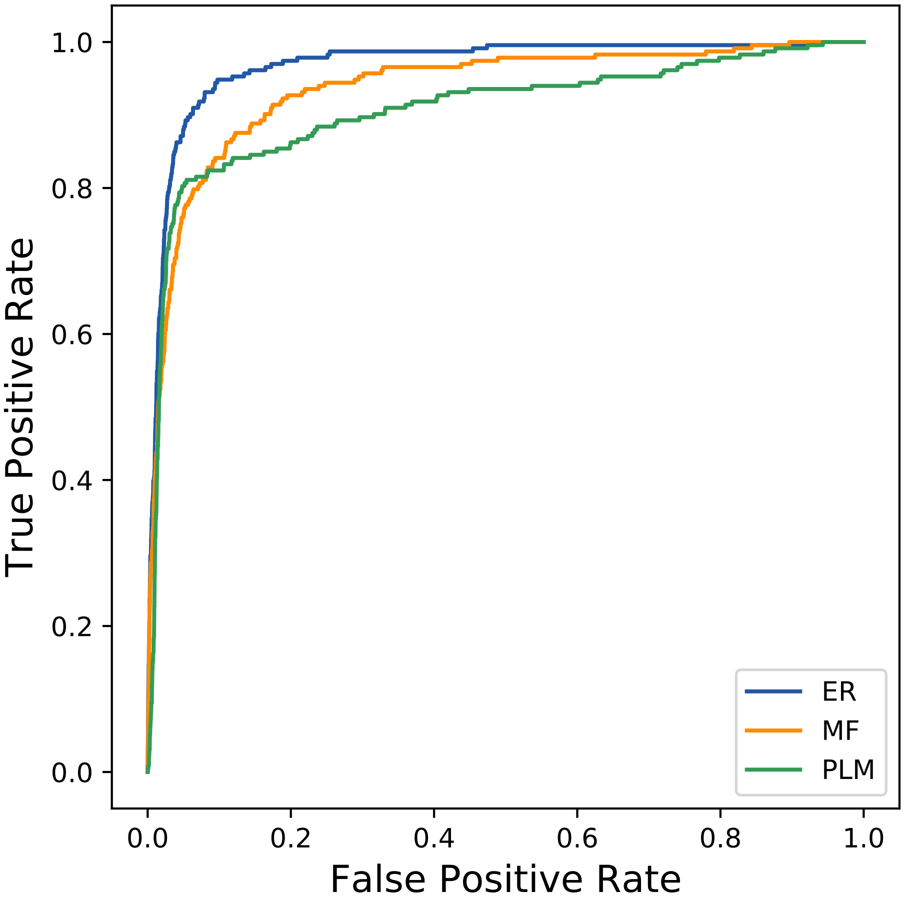

True Positive Rate
False Positive Rate
0.0
0.2
0.4
0.6
0.8
1.0
0.0
0.2
0.4
0.6
0.8
1.0
ER
MF
PLM
