## Supplemental Contact Maps for "Protein Structure Prediction with Expectation Reflection": 17807seq_33col_7afw_PF00514_plots.pdf

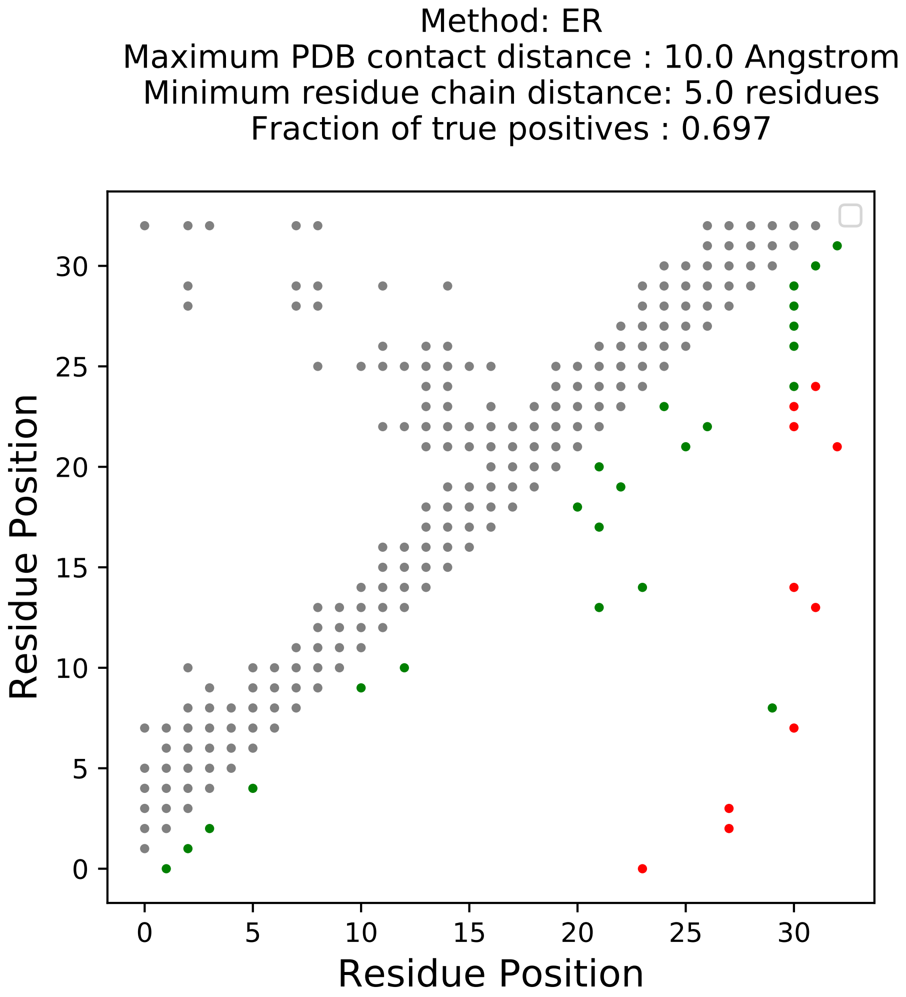
Method: ER
Maximum PDB contact distance : 10.0 Angstrom
Minimum residue chain distance: 5.0 residues
Fraction of true positives : 0.697
Residue Position
Residue Position
0
5
10
15
20
25
30

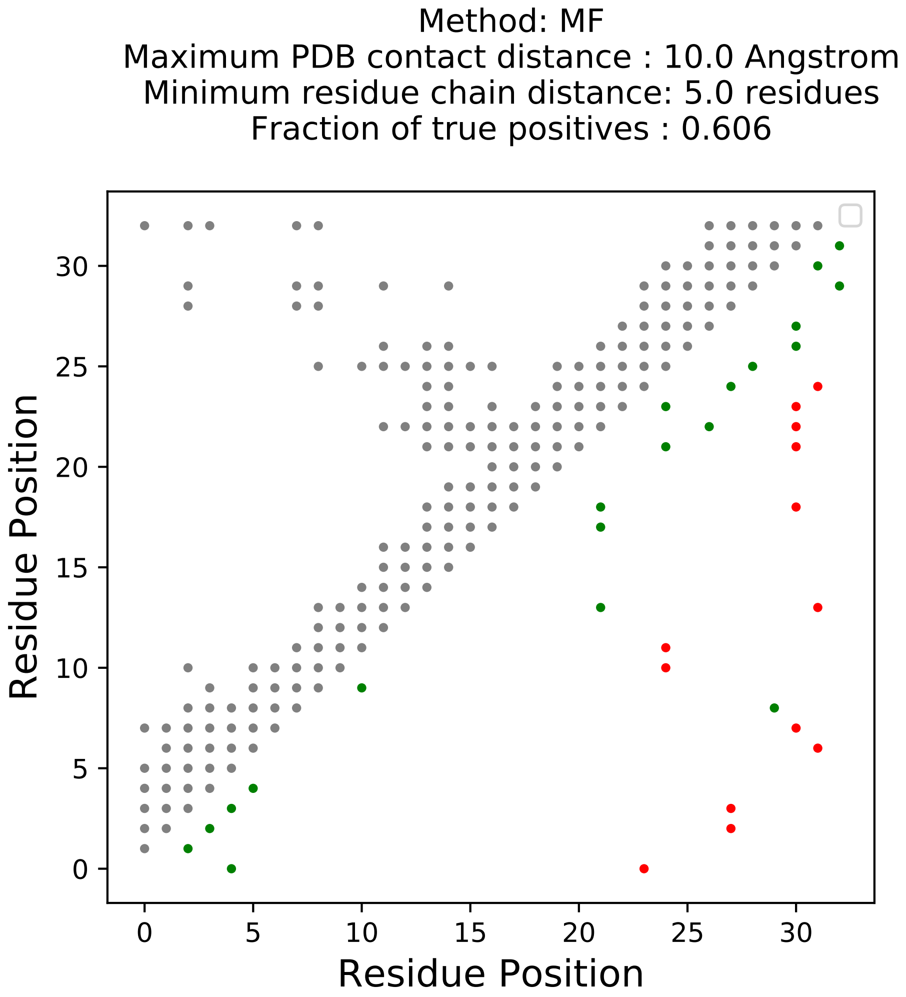
Method: MF
Maximum PDB contact distance : 10.0 Angstrom
Minimum residue chain distance: 5.0 residues
Fraction of true positives : 0.606
Residue Position
Residue Position
0
5
10
15
20
25
30

Method: PLM
Maximum PDB contact distance : 10.0 Angstrom
Minimum residue chain distance: 5.0 residues
Fraction of true positives : 0.727

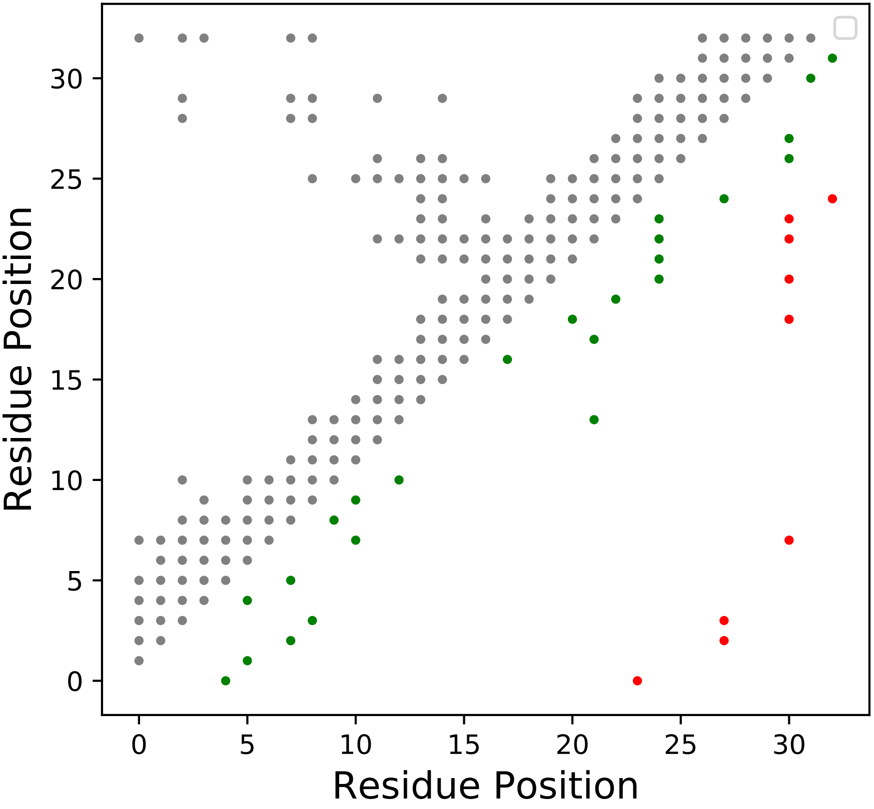

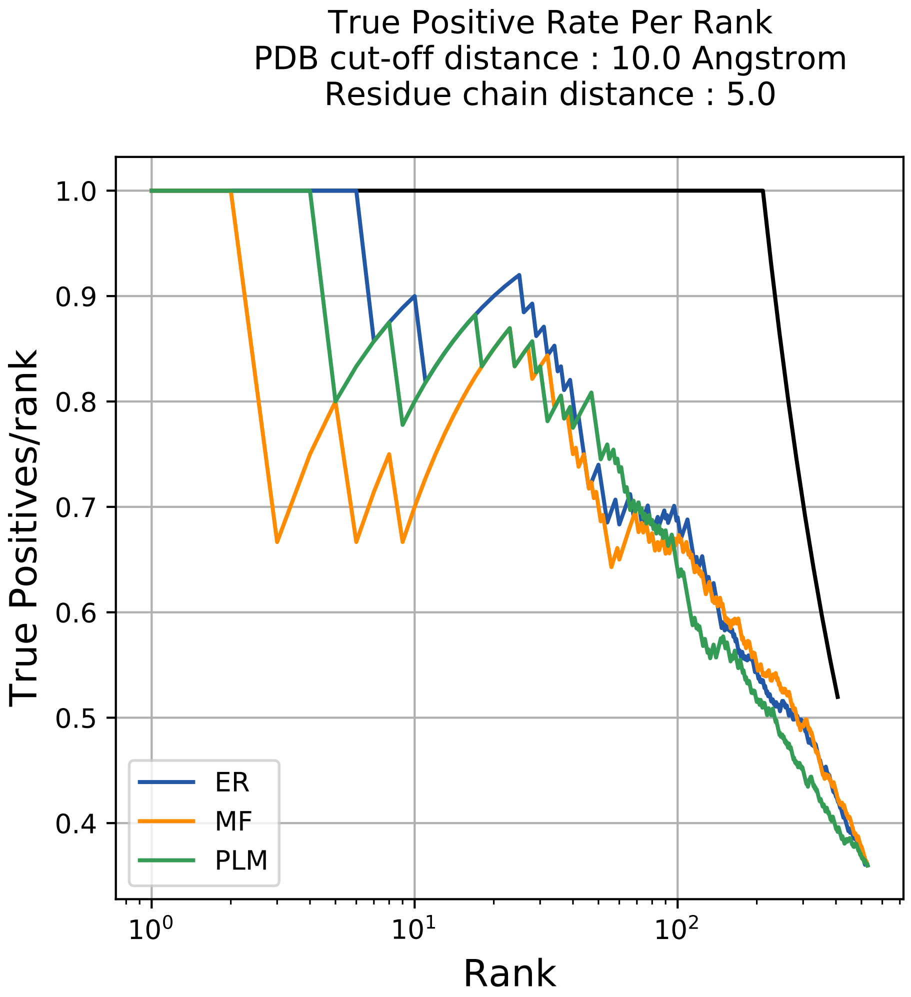
True Positive Rate Per Rank
PDB cut-off distance : 10.0 Angstrom
Residue chain distance : 5.0
True Positives/rank
Rank
1.0
0.9
0.8
0.7
0.6
0.5
0.4
$10^0$
$10^1$
$10^2$
ER
MF
PLM

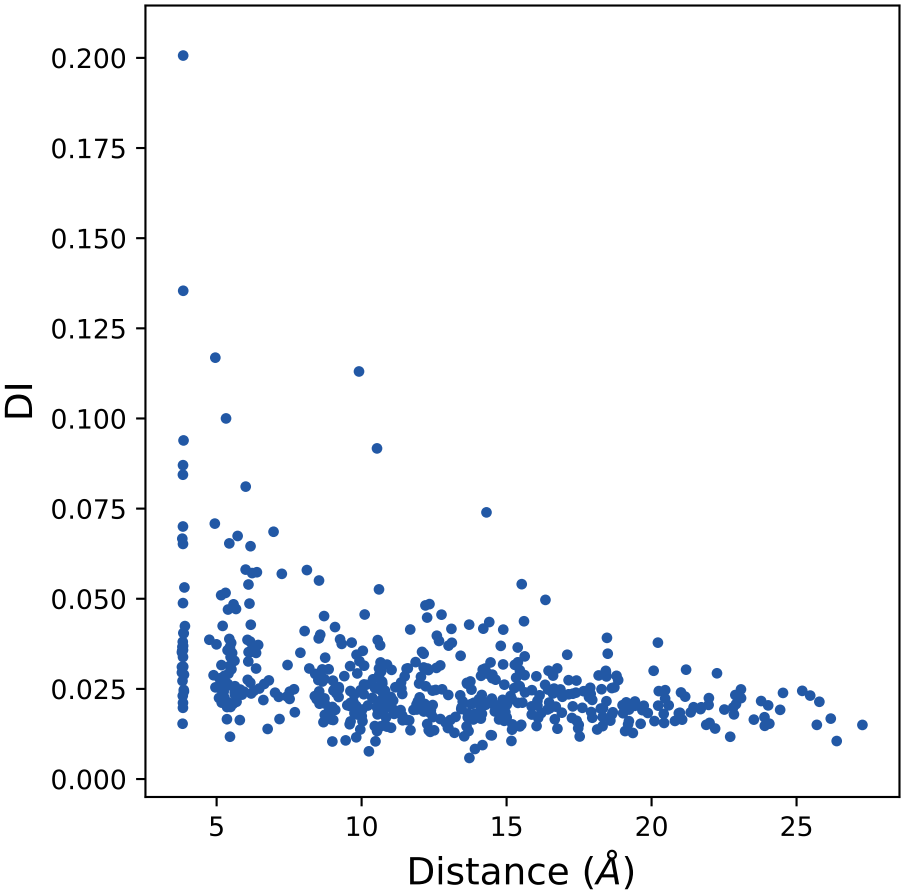

DI
Distance (Å)
0.200
0.175
0.150
0.125
0.100
0.075
0.050
0.025
0.000
5
10
15
20
25

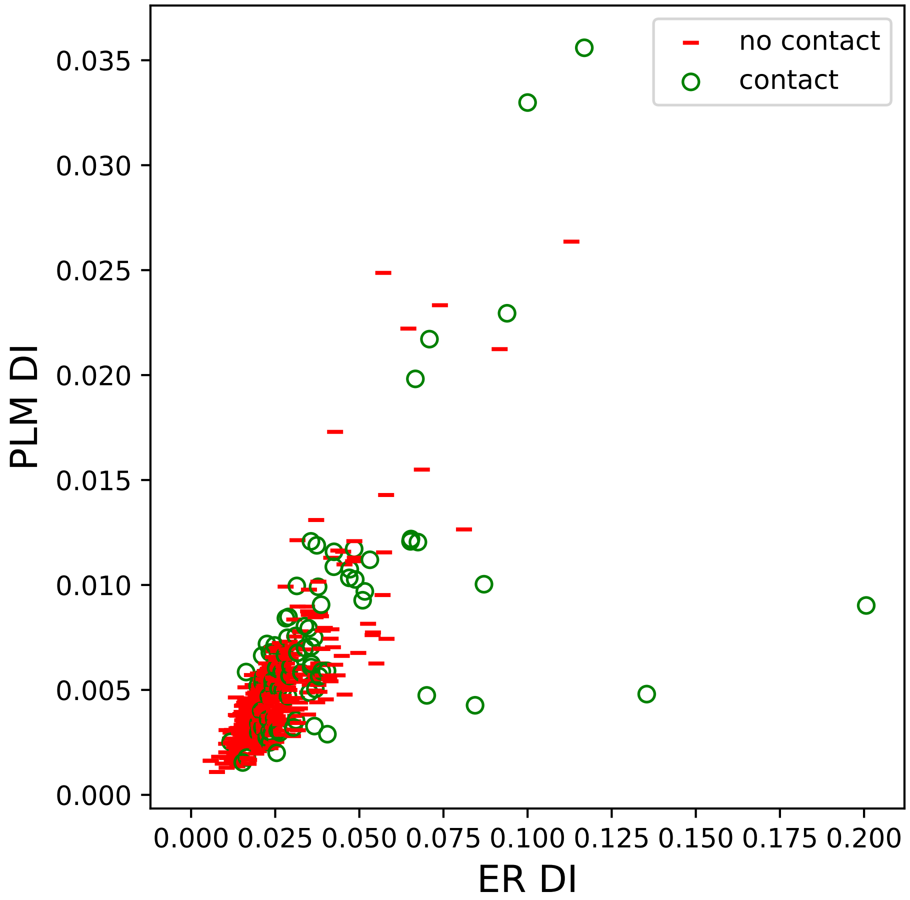

no contact
contact
PLM DI
ER DI
0.000
0.005
0.010
0.015
0.020
0.025
0.030
0.035
0.000 0.025 0.050 0.075 0.100 0.125 0.150 0.175 0.200

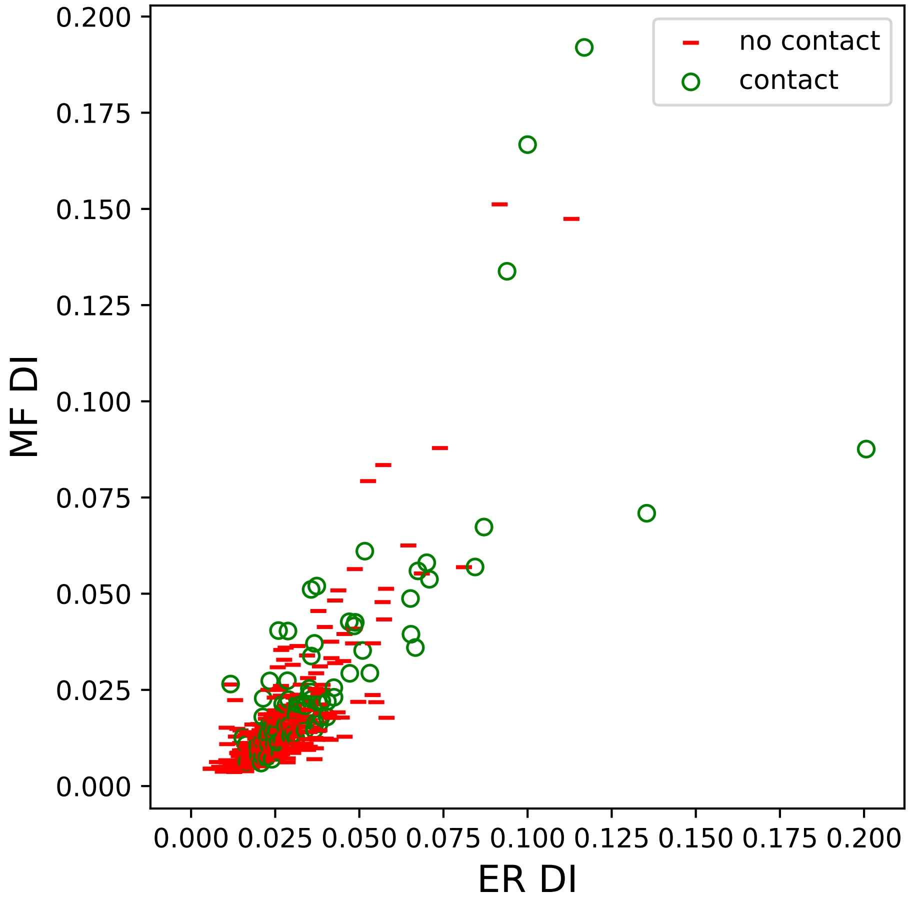

no contact
contact
MF DI
ER DI
0.000
0.025
0.050
0.075
0.100
0.125
0.150
0.175
0.200

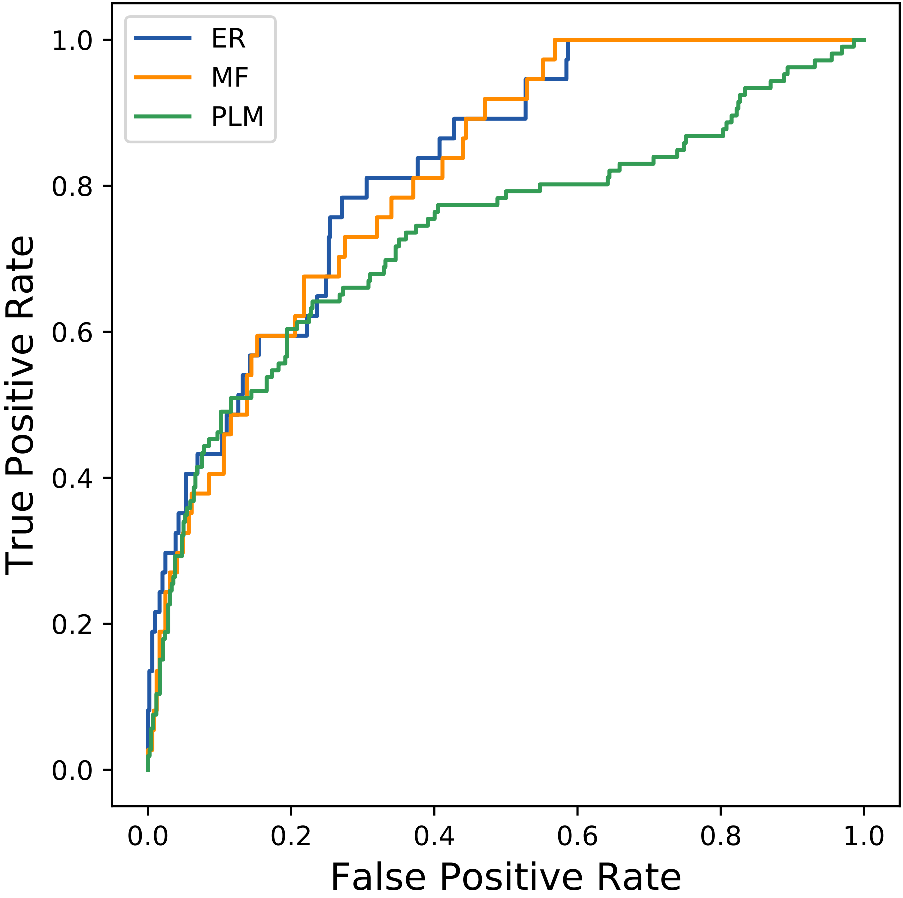

ER
MF
PLM
True Positive Rate
False Positive Rate
0.0
0.2
0.4
0.6
0.8
1.0
