## Supplemental Contact Maps for "Protein Structure Prediction with Expectation Reflection": 18192seq_199col_7sf6_PF01497_plots.pdf

Method: ER
Maximum PDB contact distance : 10.0 Angstrom
Minimum residue chain distance: 5.0 residues
Fraction of true positives : 0.844

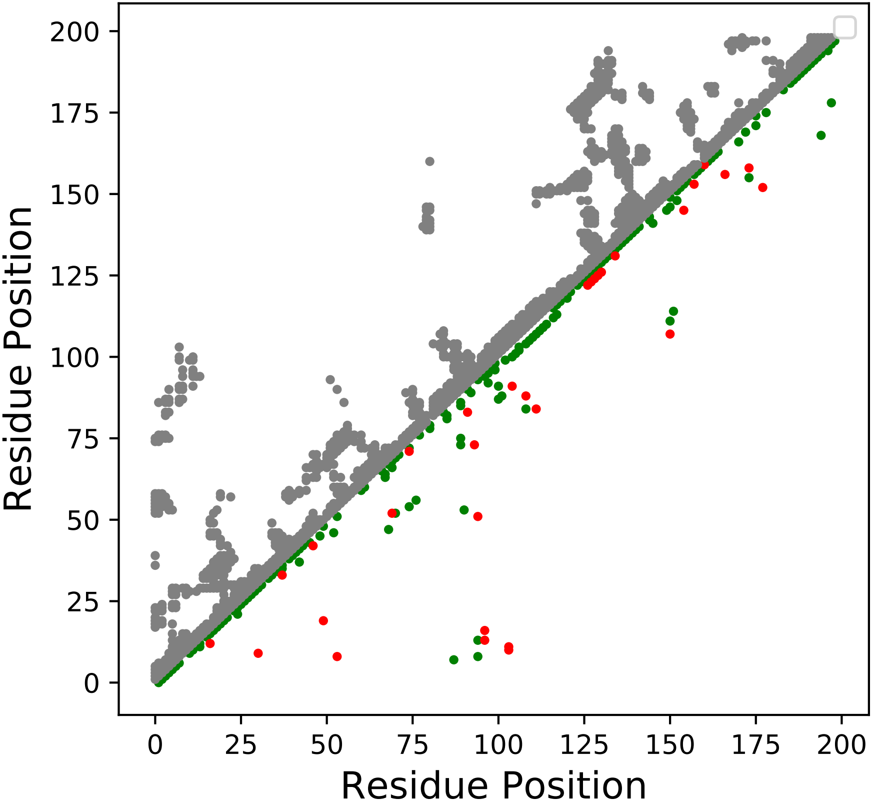

Method: MF
Maximum PDB contact distance : 10.0 Angstrom
Minimum residue chain distance: 5.0 residues
Fraction of true positives : 0.799

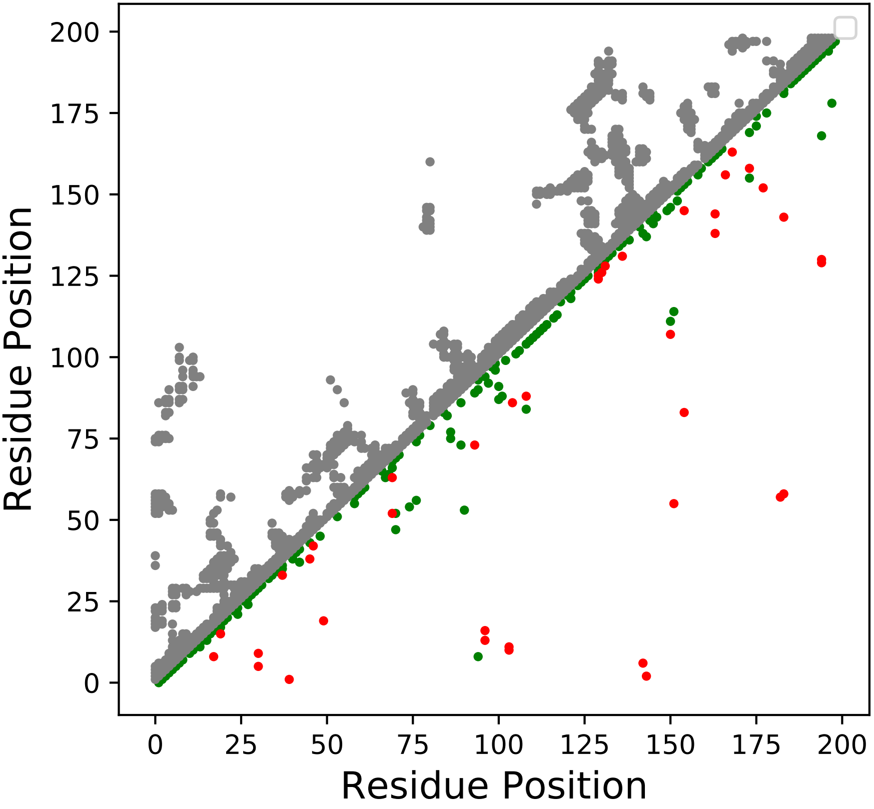

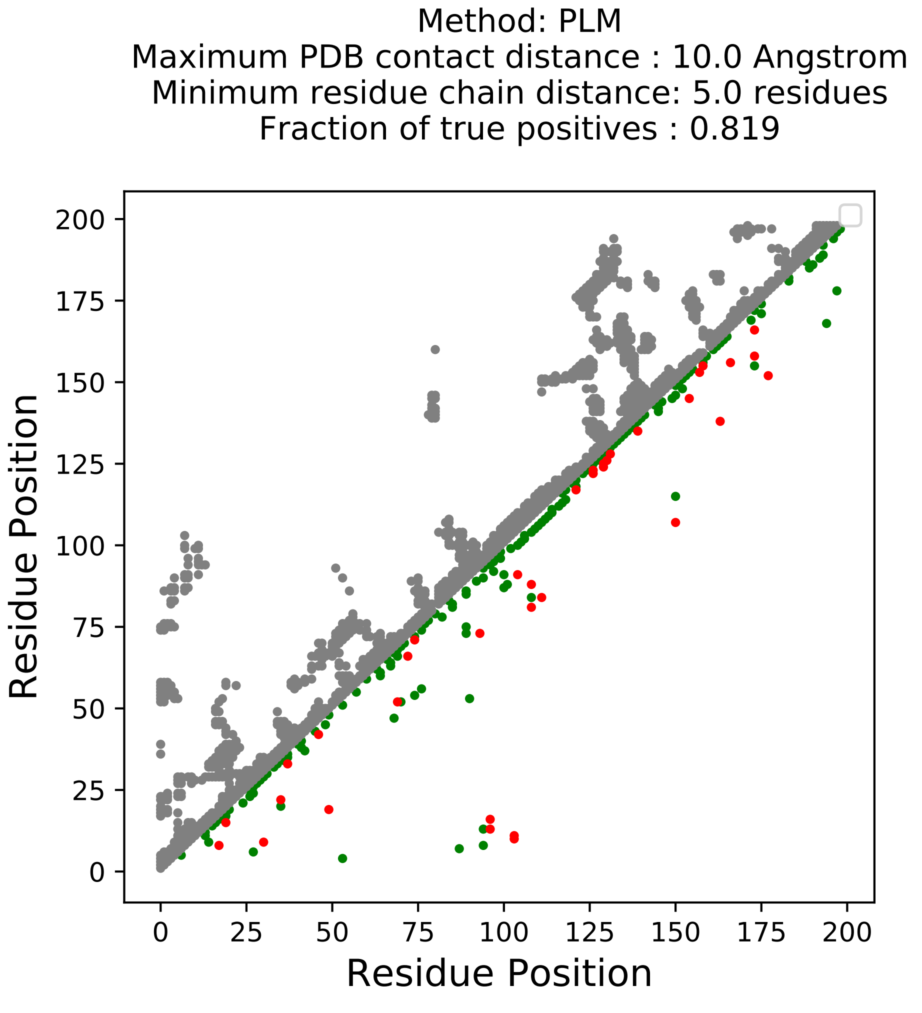

Method: PLM
Maximum PDB contact distance : 10.0 Angstrom
Minimum residue chain distance: 5.0 residues
Fraction of true positives : 0.819
Residue Position
Residue Position
0
25
50
75
100
125
150
175
200

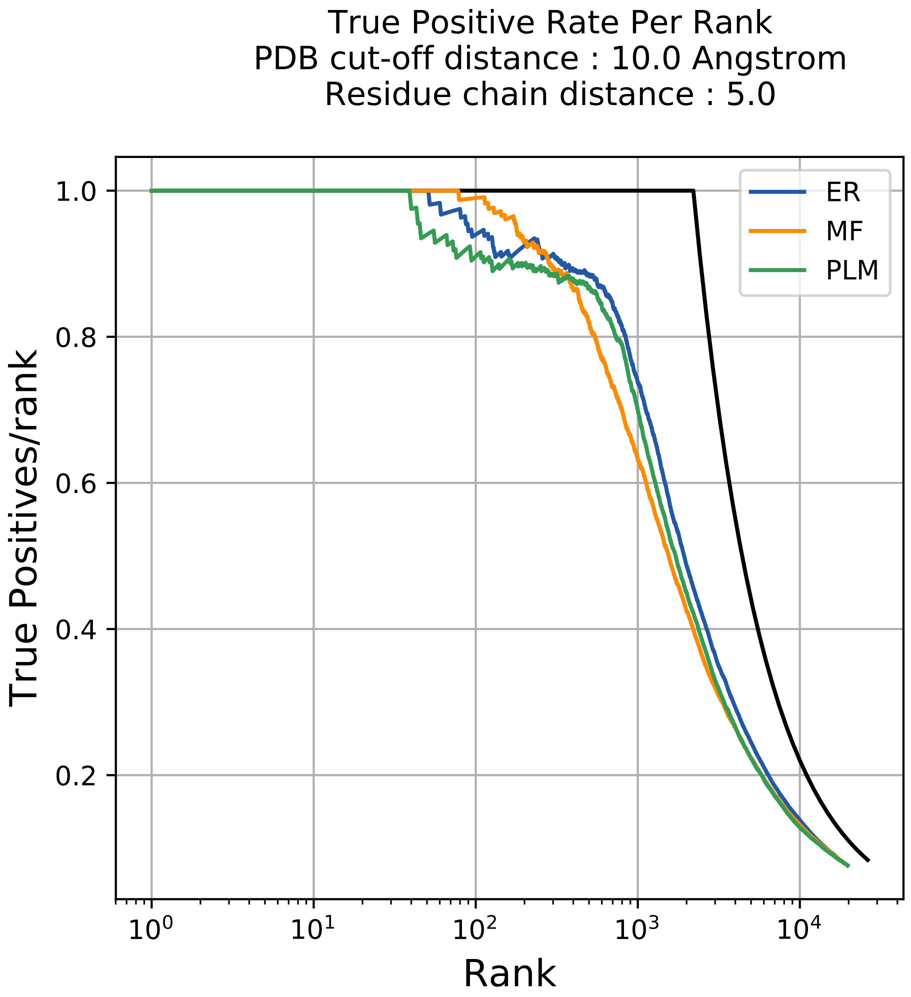

True Positive Rate Per Rank
PDB cut-off distance : 10.0 Angstrom
Residue chain distance : 5.0
True Positives/rank
Rank
ER
MF
PLM
1.0
0.8
0.6
0.4
0.2
$10^0$
$10^1$
$10^2$
$10^3$
$10^4$

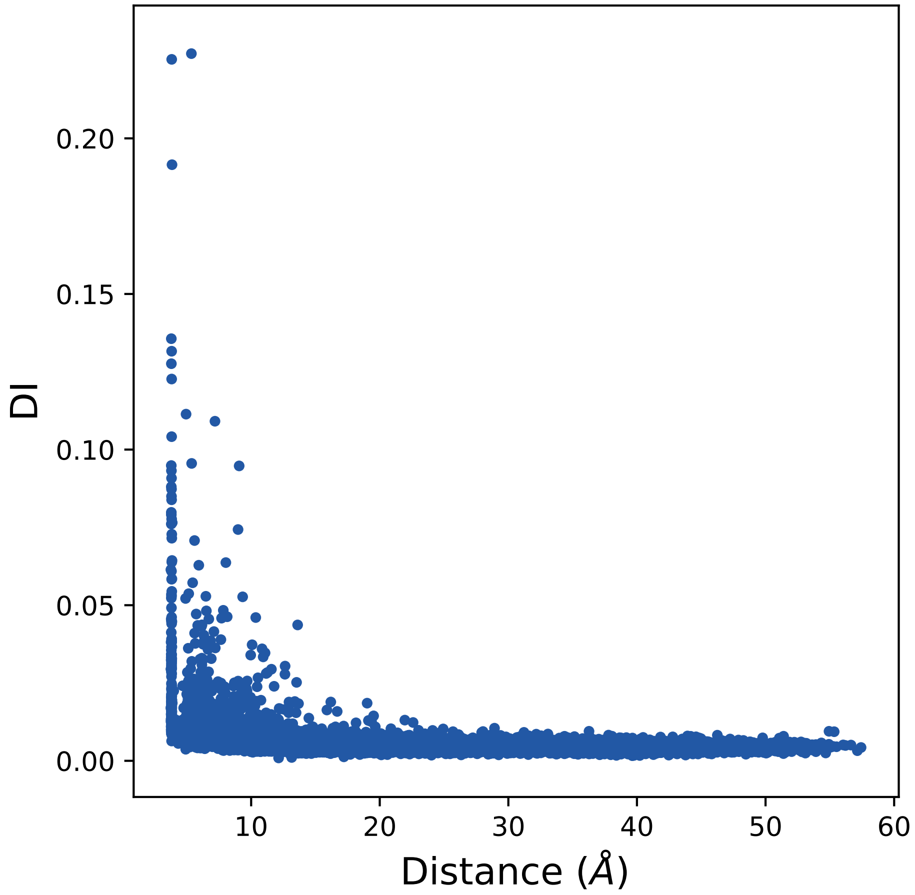

0.20
0.15
0.10
0.05
0.00
DI
10
20
30
40
50
60
Distance (Å)

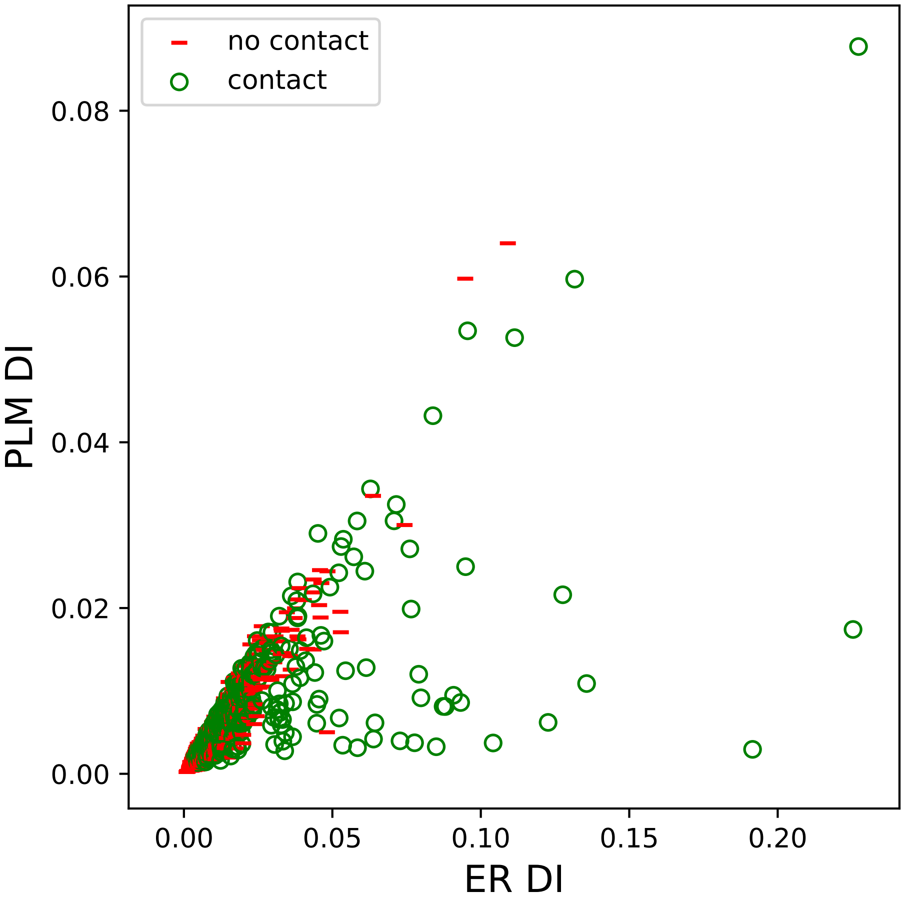

no contact
contact
0.08
0.06
0.04
0.02
0.00
PLM DI
0.00
0.05
0.10
0.15
0.20
ER DI

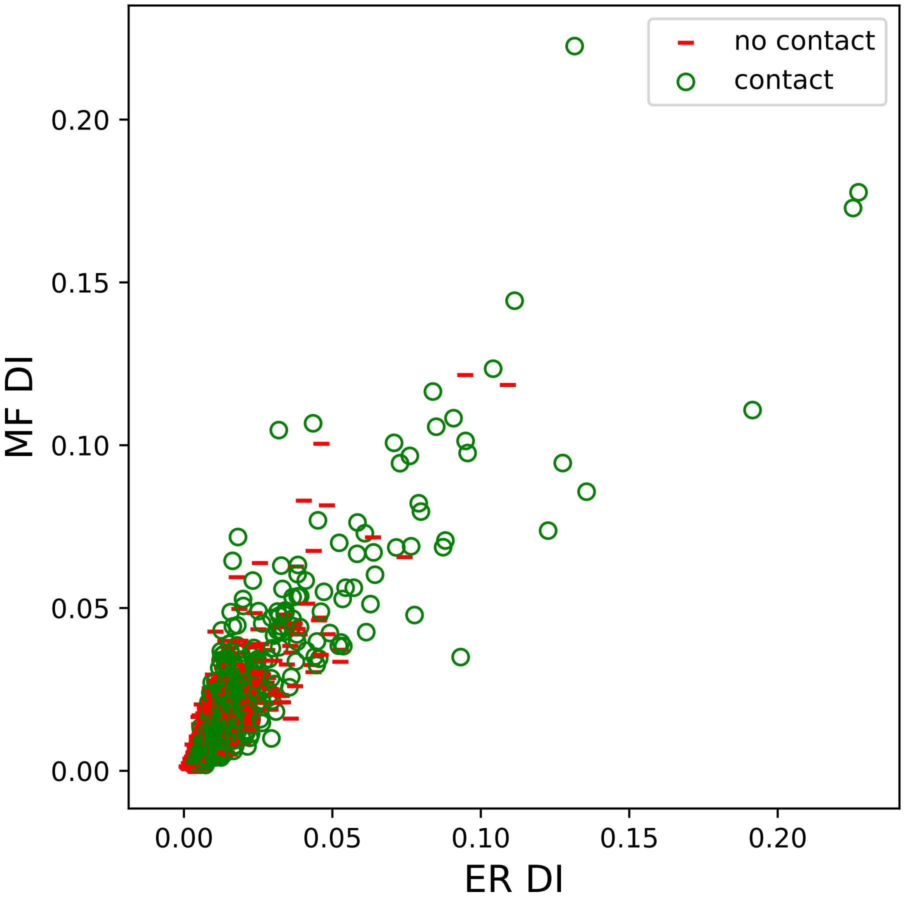

no contact
contact
MF DI
ER DI
0.00
0.05
0.10
0.15
0.20

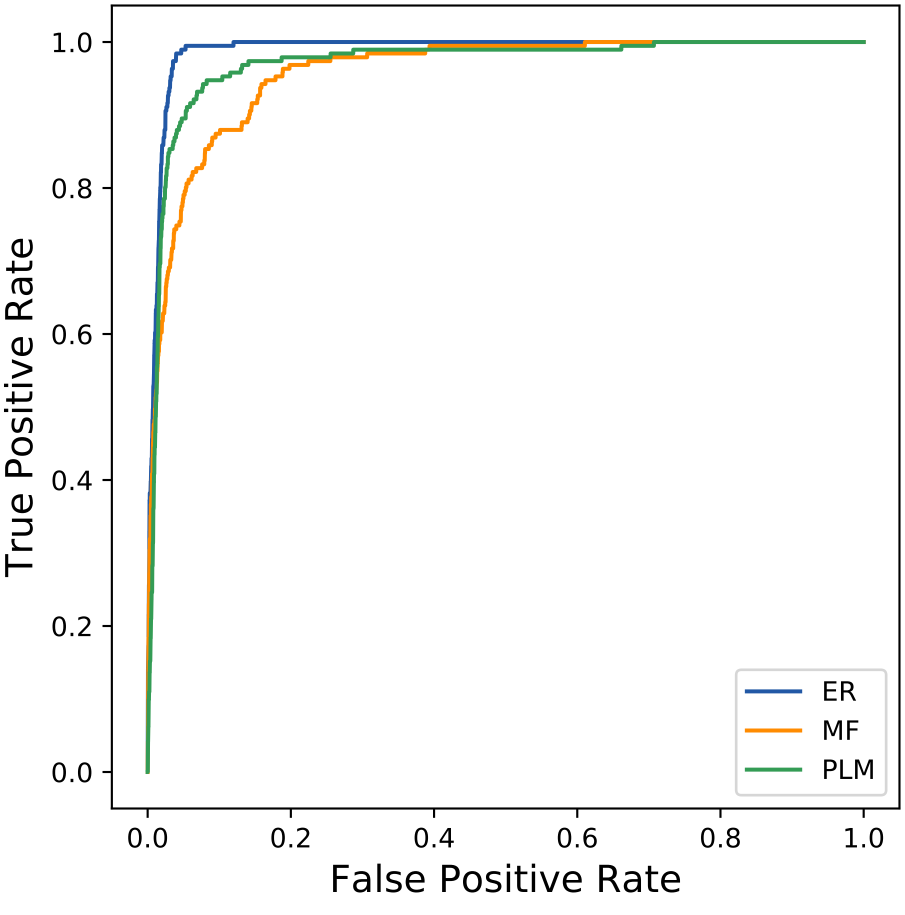

True Positive Rate
False Positive Rate
0.0
0.2
0.4
0.6
0.8
1.0
ER
MF
PLM
