## Supplemental Contact Maps for "Protein Structure Prediction with Expectation Reflection": 19378seq_256col_3ps7_PF00701_plots.pdf

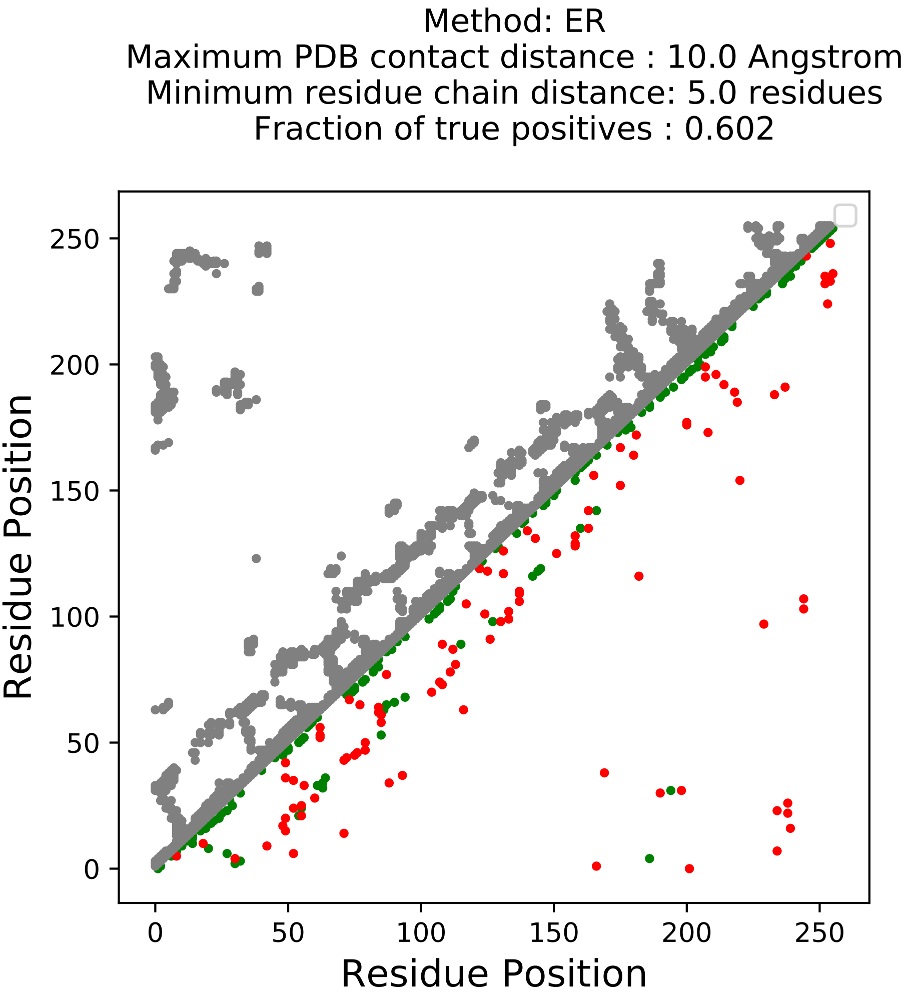
Method: ER
Maximum PDB contact distance : 10.0 Angstrom
Minimum residue chain distance: 5.0 residues
Fraction of true positives : 0.602
Residue Position
Residue Position
0
50
100
150
200
250
0
50
100
150
200
250

Method: MF
Maximum PDB contact distance : 10.0 Angstrom
Minimum residue chain distance: 5.0 residues
Fraction of true positives : 0.594

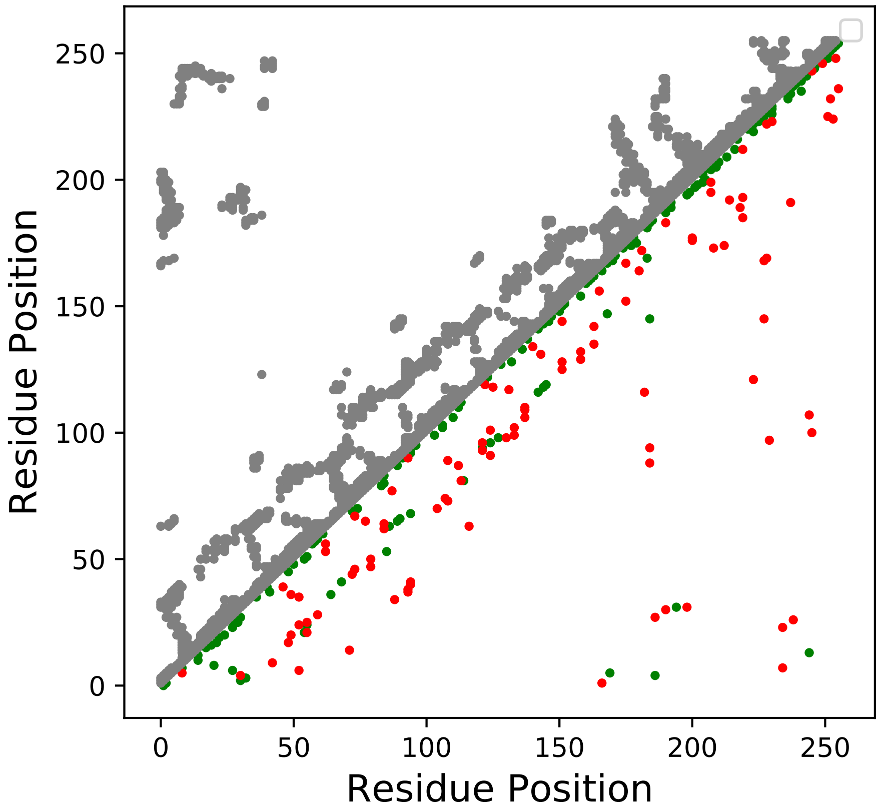

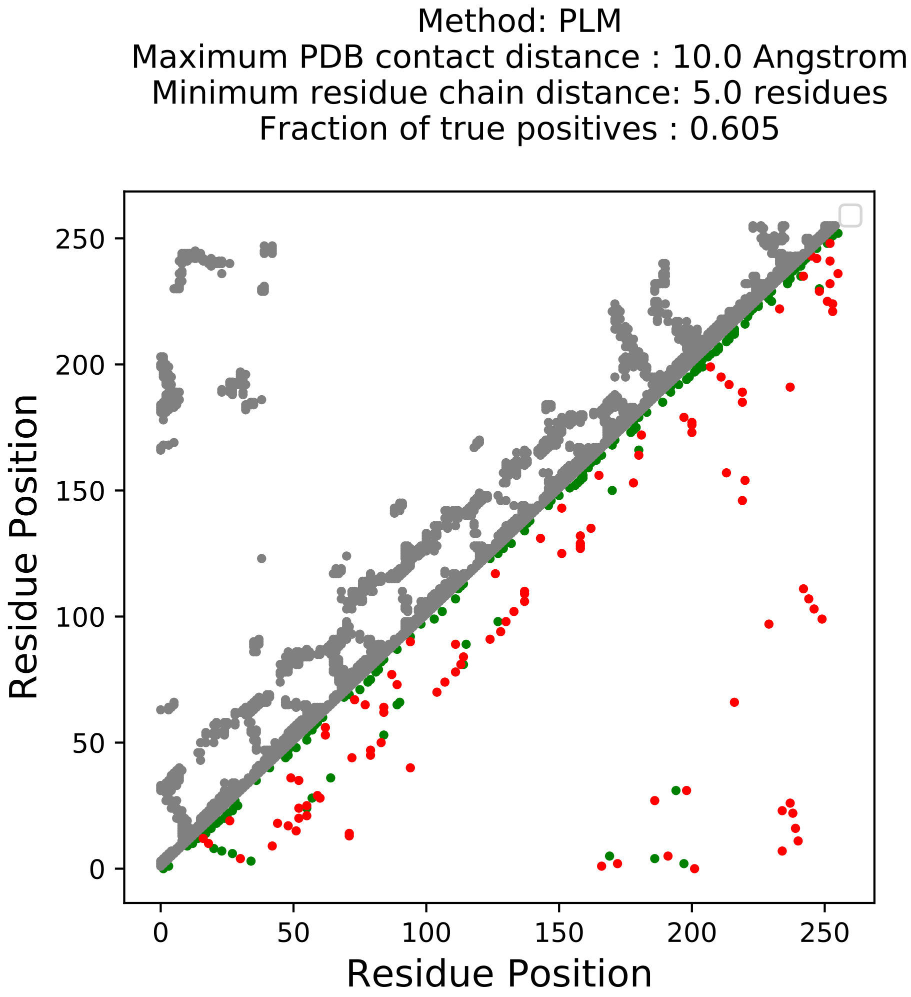
Method: PLM
Maximum PDB contact distance : 10.0 Angstrom
Minimum residue chain distance: 5.0 residues
Fraction of true positives : 0.605
Residue Position
Residue Position
0
50
100
150
200
250
0
50
100
150
200
250

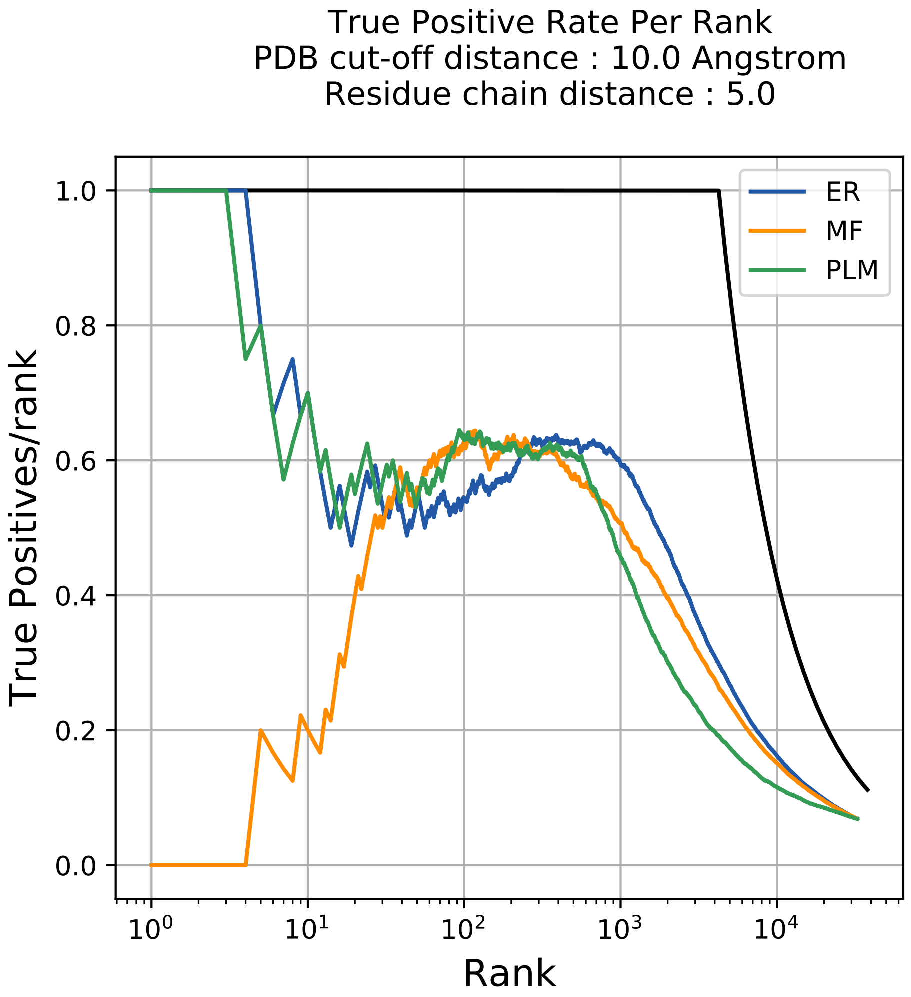

True Positive Rate Per Rank
PDB cut-off distance : 10.0 Angstrom
Residue chain distance : 5.0
True Positives/rank
Rank
1.0
0.8
0.6
0.4
0.2
0.0
$10^0$
$10^1$
$10^2$
$10^3$
$10^4$
ER
MF
PLM

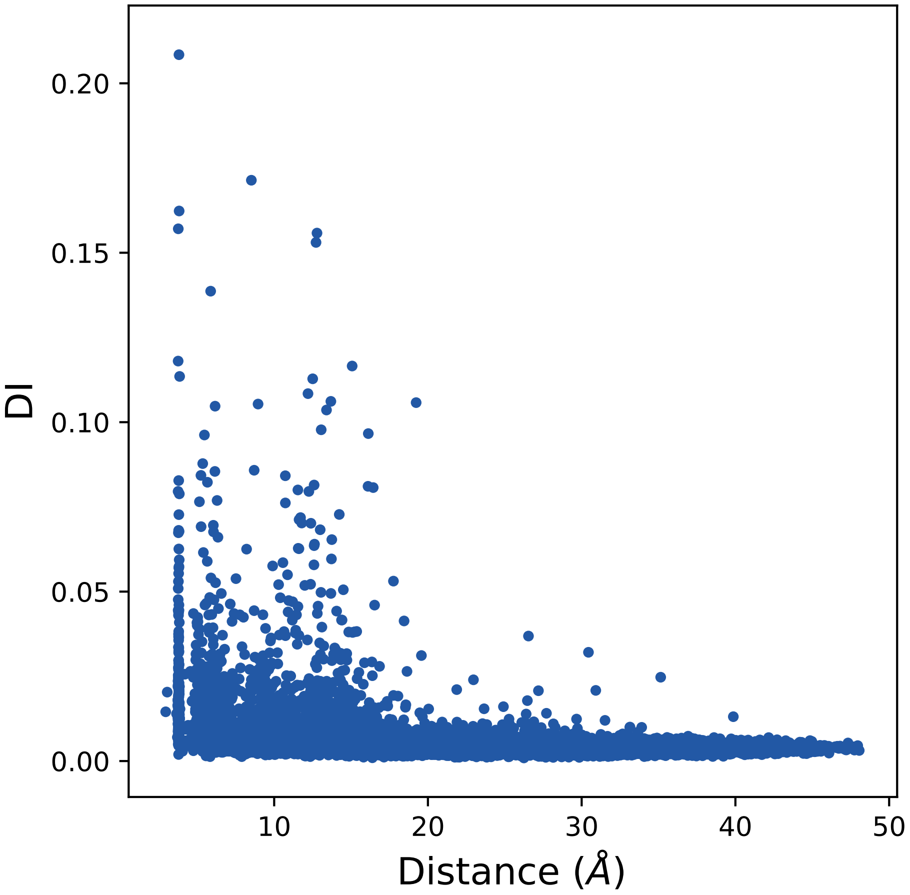

0.20
0.15
0.10
0.05
0.00
DI
10
20
30
40
50
Distance (Å)

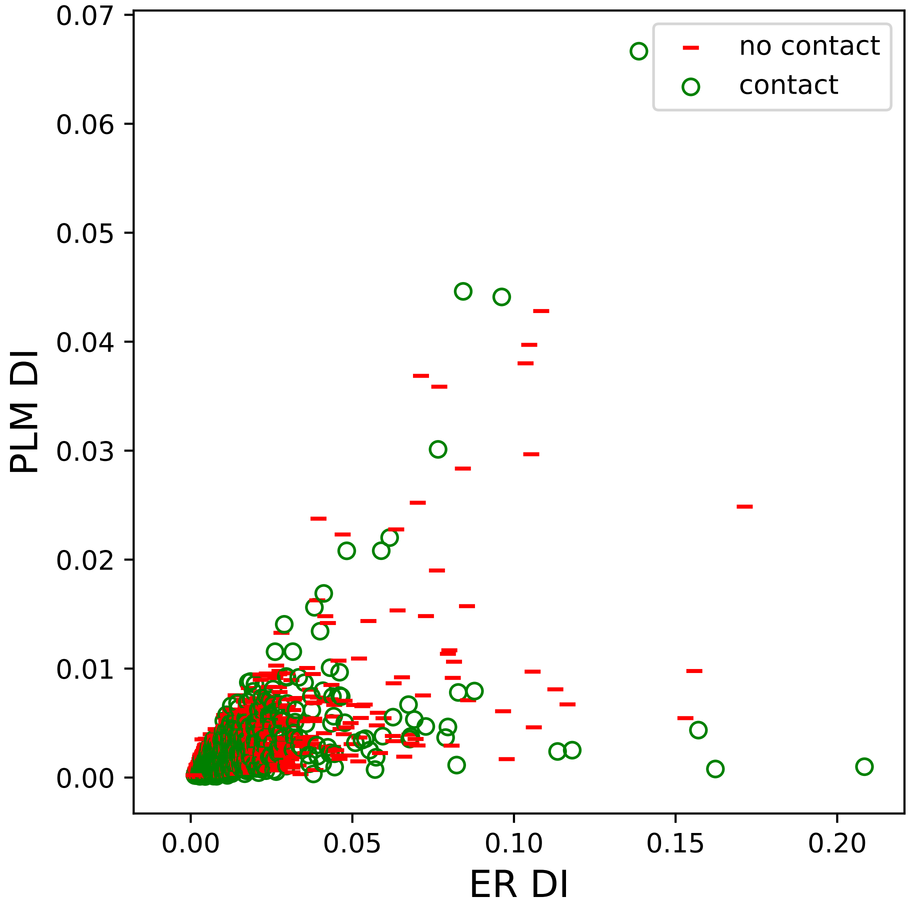

PLM DI
ER DI
0.00
0.01
0.02
0.03
0.04
0.05
0.06
0.07
0.00
0.05
0.10
0.15
0.20
no contact
contact

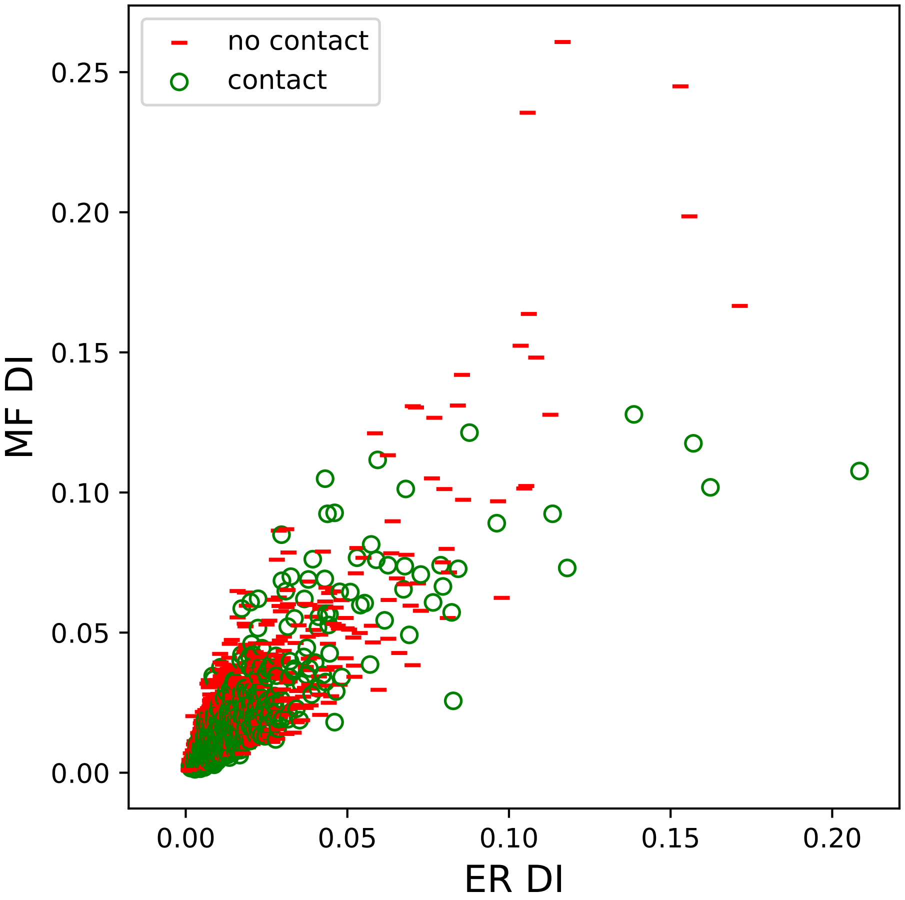
no contact
contact
MF DI
ER DI
0.00
0.05
0.10
0.15
0.20
0.25
0.00
0.05
0.10
0.15
0.20

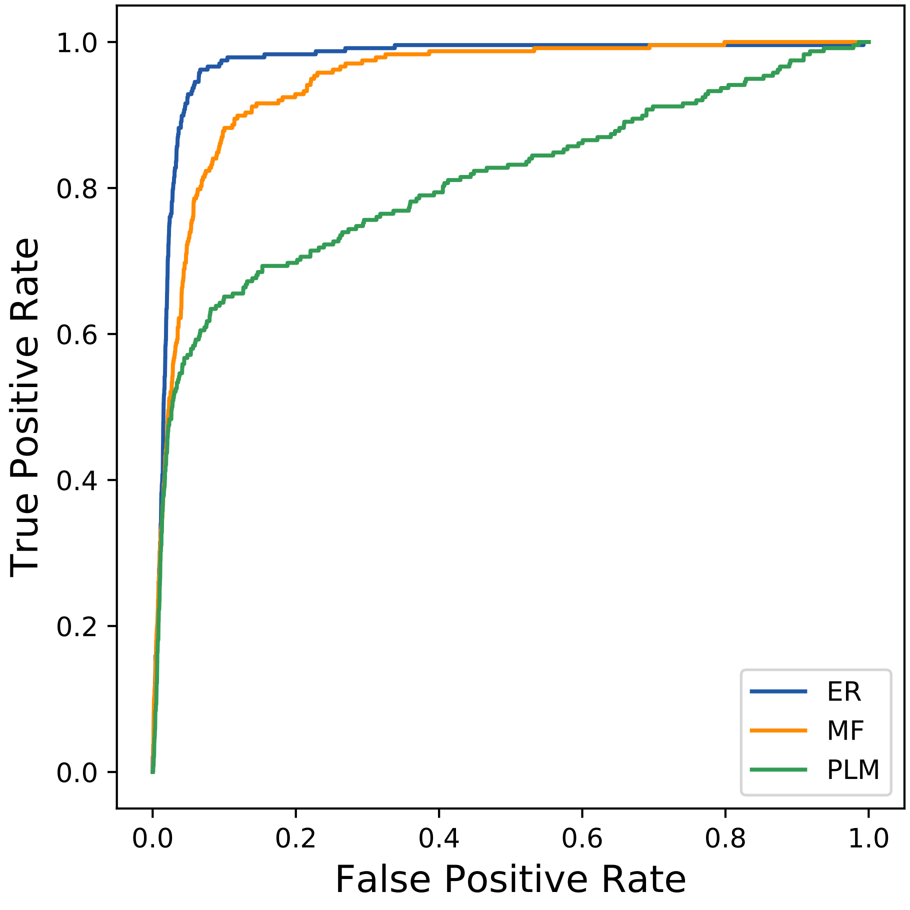

1.0
0.8
0.6
0.4
0.2
0.0
True Positive Rate
0.0
0.2
0.4
0.6
0.8
1.0
False Positive Rate
ER
MF
PLM
