## Supplemental Contact Maps for "Protein Structure Prediction with Expectation Reflection": 20130seq_145col_7jzb_PF00701_plots.pdf

Method: ER
Maximum PDB contact distance : 10.0 Angstrom
Minimum residue chain distance: 5.0 residues
Fraction of true positives : 0.621

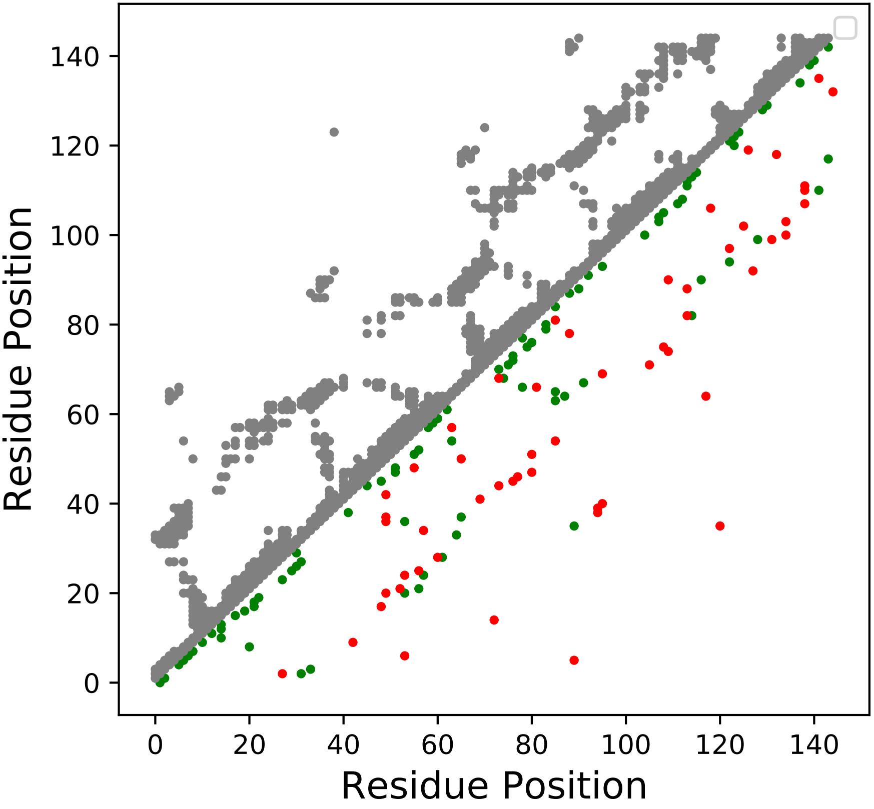

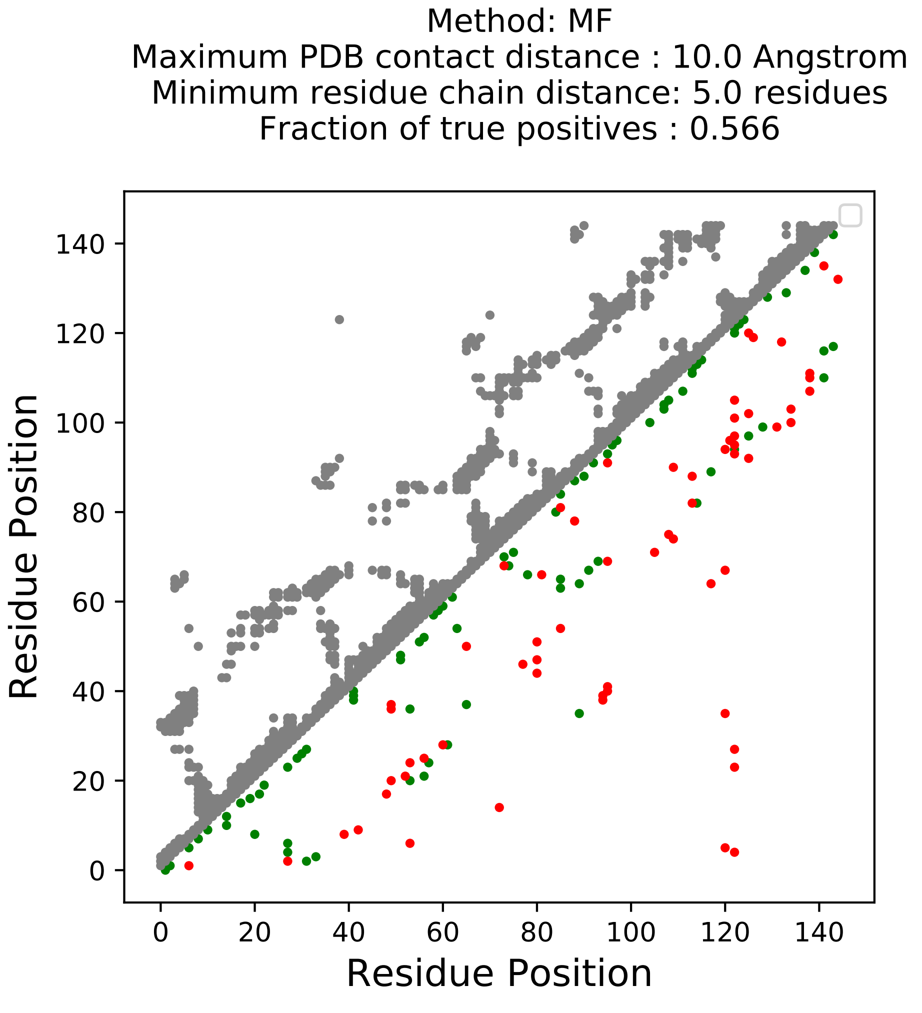
Method: MF
Maximum PDB contact distance : 10.0 Angstrom
Minimum residue chain distance: 5.0 residues
Fraction of true positives : 0.566
Residue Position
Residue Position
0
20
40
60
80
100
120
140

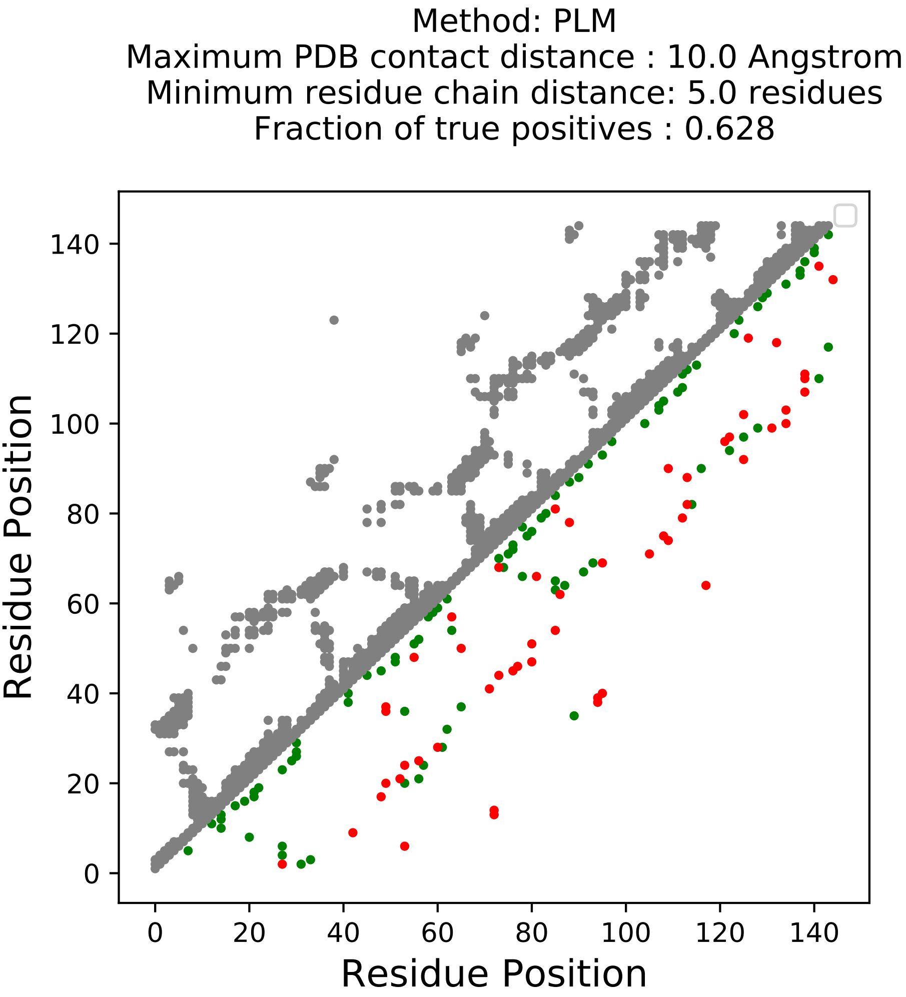
Method: PLM
Maximum PDB contact distance : 10.0 Angstrom
Minimum residue chain distance: 5.0 residues
Fraction of true positives : 0.628
Residue Position
Residue Position
0
20
40
60
80
100
120
140

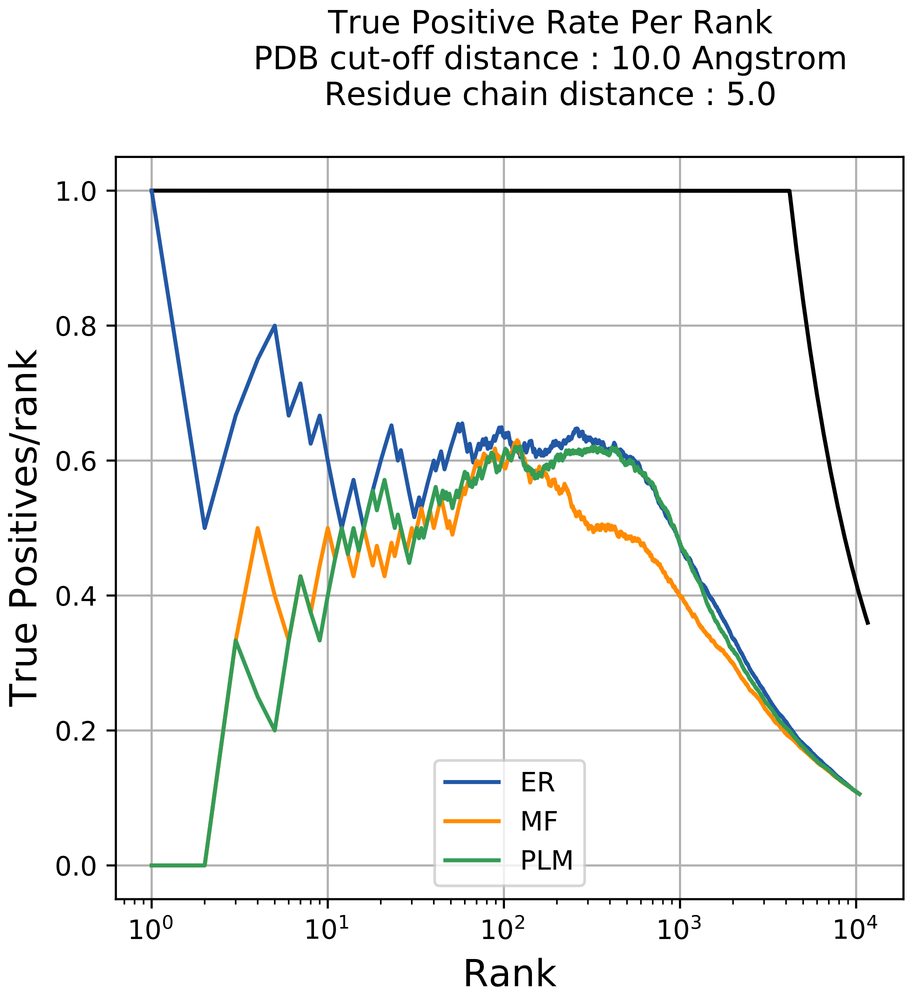
True Positive Rate Per Rank
PDB cut-off distance : 10.0 Angstrom
Residue chain distance : 5.0
True Positives/rank
Rank
1.0
0.8
0.6
0.4
0.2
0.0
$10^0$
$10^1$
$10^2$
$10^3$
$10^4$
ER
MF
PLM

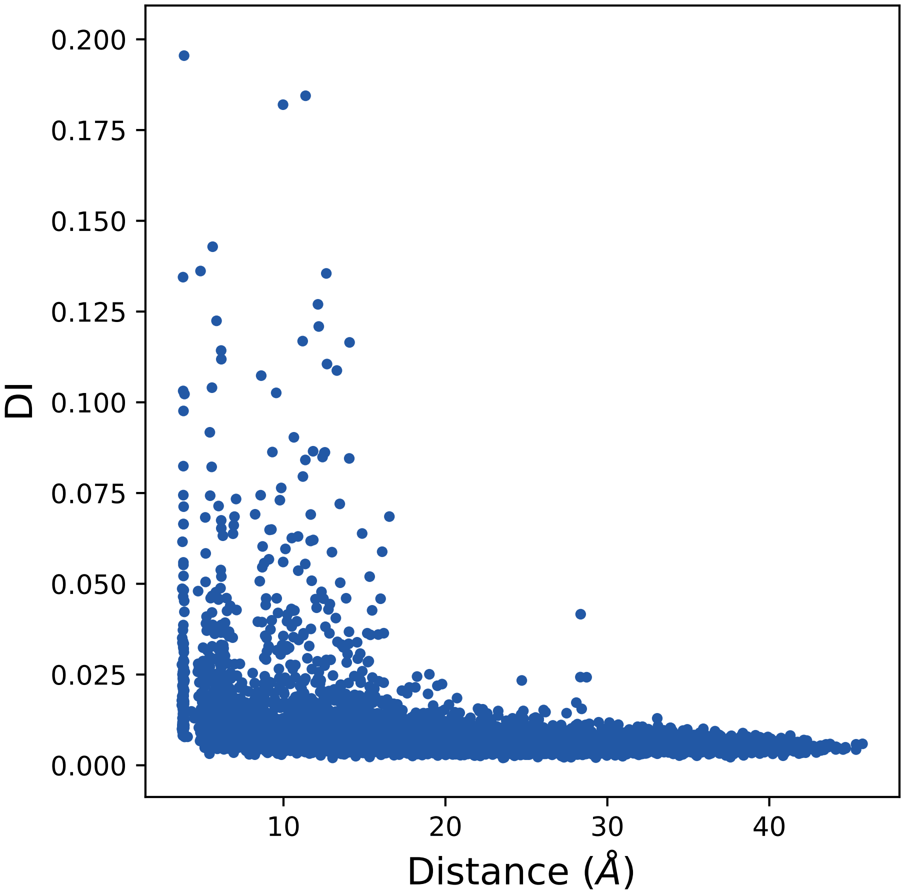

0.200
0.175
0.150
0.125
0.100
0.075
0.050
0.025
0.000
DI
10
20
30
40
Distance (Å)

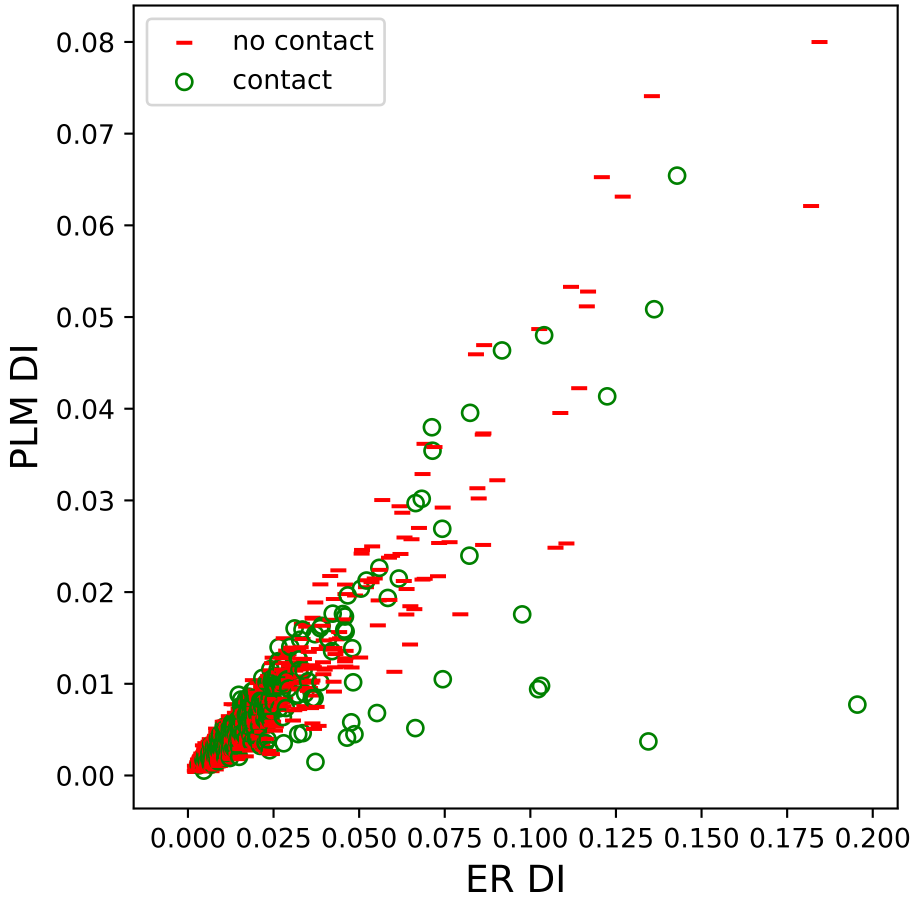

no contact
contact
PLM DI
ER DI
0.00
0.01
0.02
0.03
0.04
0.05
0.06
0.07
0.08
0.000
0.025
0.050
0.075
0.100
0.125
0.150
0.175
0.200

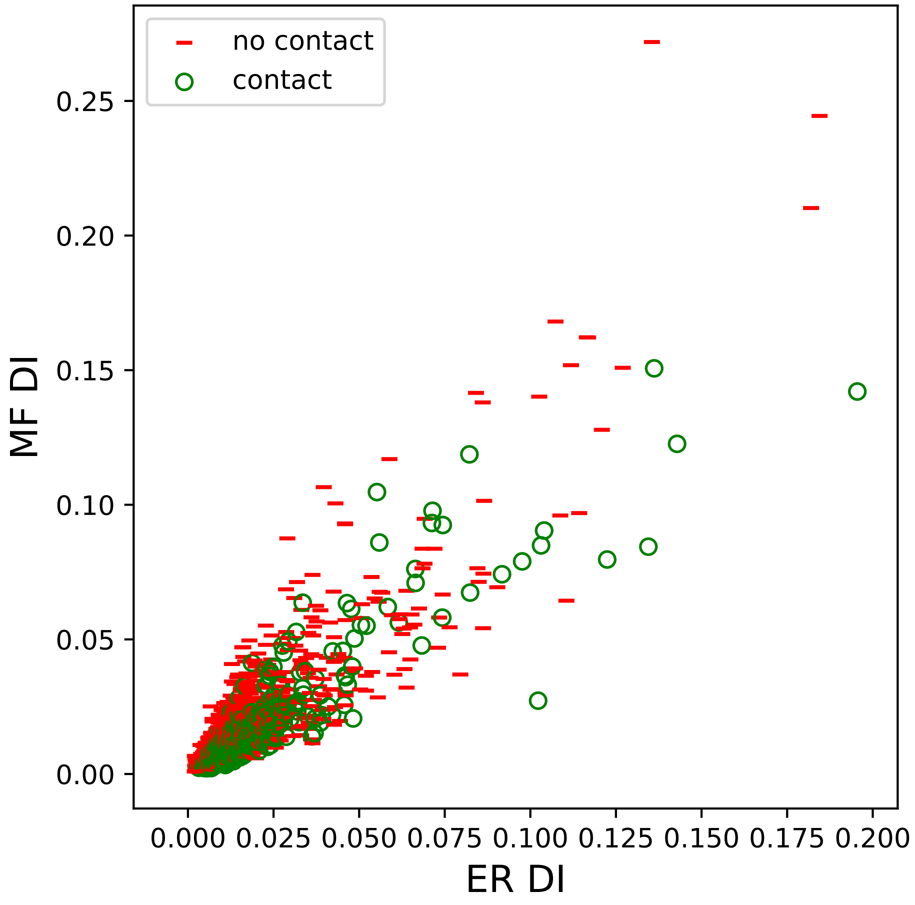
no contact
contact
MF DI
ER DI
0.00
0.05
0.10
0.15
0.20
0.25
0.000
0.025
0.050
0.075
0.100
0.125
0.150
0.175
0.200

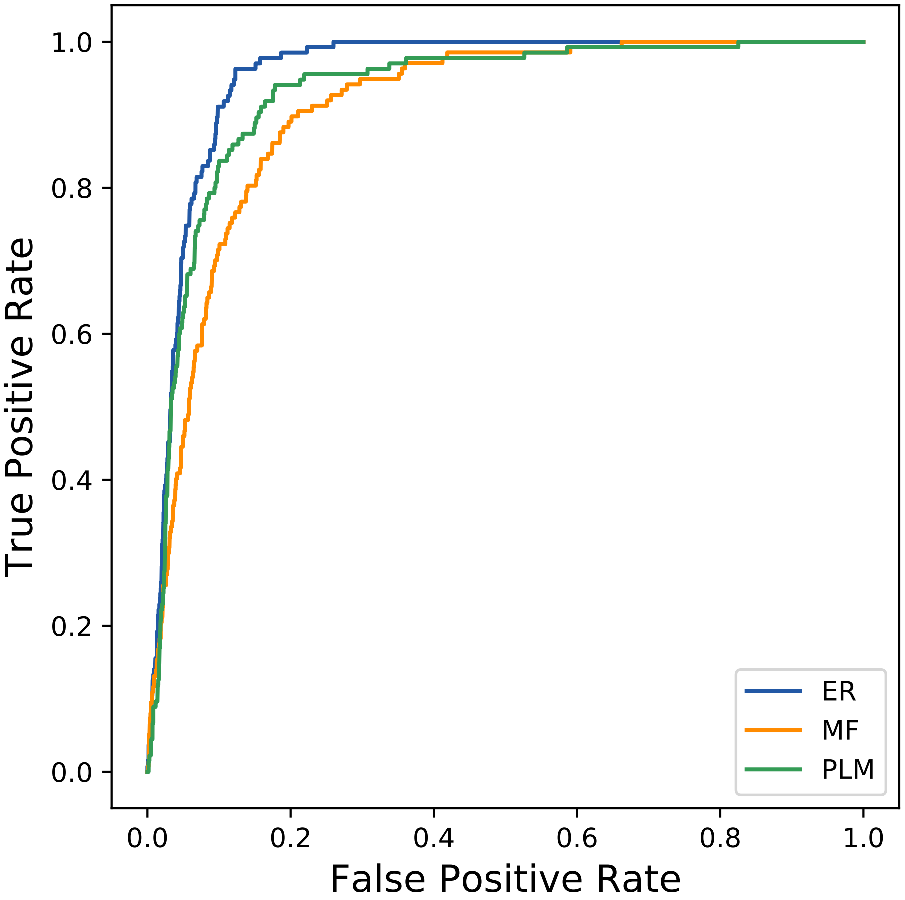
True Positive Rate
False Positive Rate
0.0
0.2
0.4
0.6
0.8
1.0
ER
MF
PLM
