## Supplemental Contact Maps for "Protein Structure Prediction with Expectation Reflection": 23176seq_67col_4zid_PF00034_plots.pdf

Method: ER
Maximum PDB contact distance : 10.0 Angstrom
Minimum residue chain distance: 5.0 residues
Fraction of true positives : 0.896

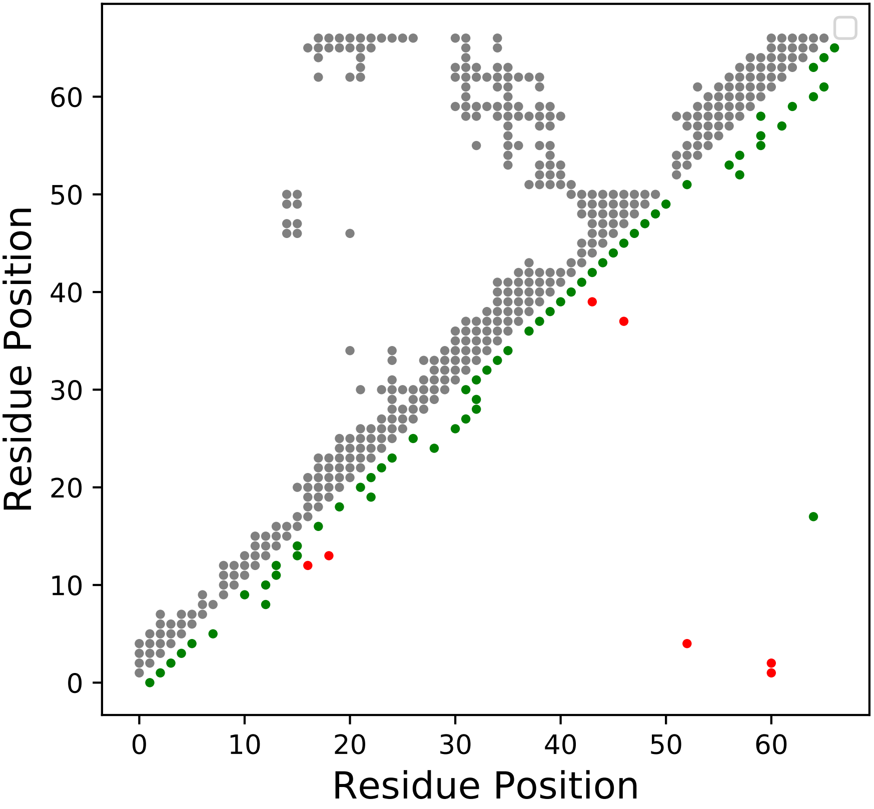

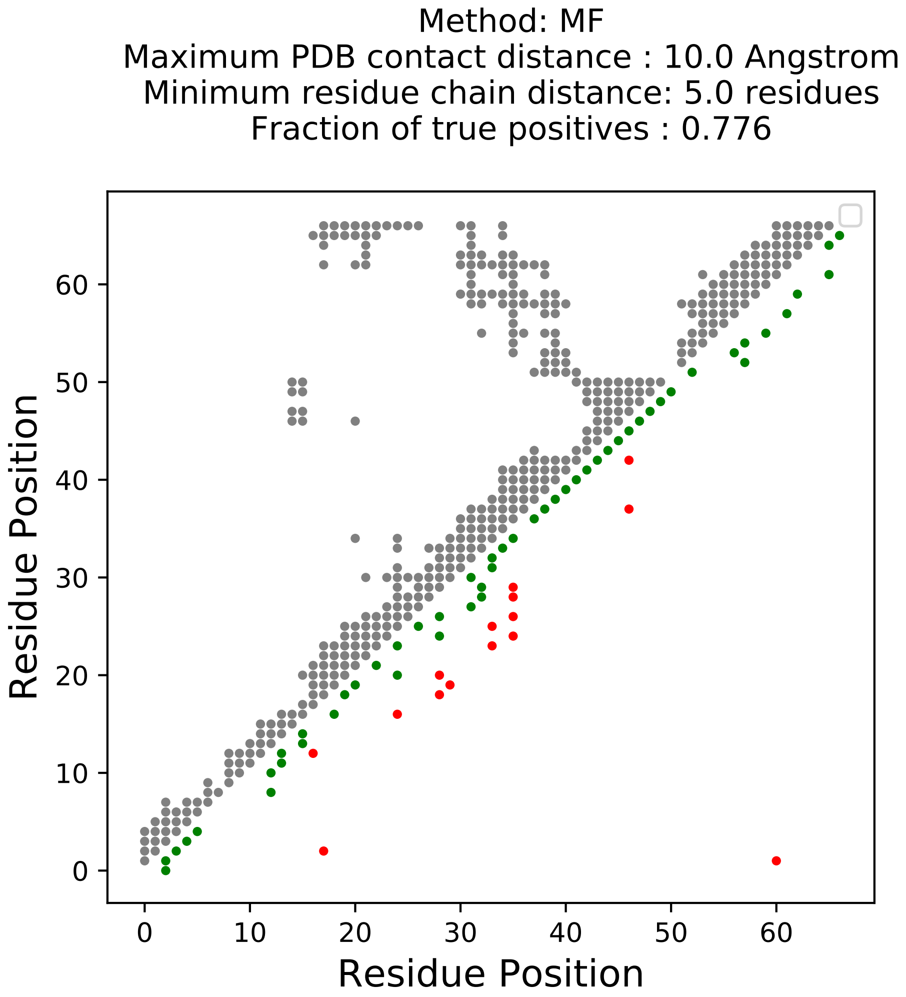

Method: MF
Maximum PDB contact distance : 10.0 Angstrom
Minimum residue chain distance: 5.0 residues
Fraction of true positives : 0.776
Residue Position
Residue Position
0
10
20
30
40
50
60

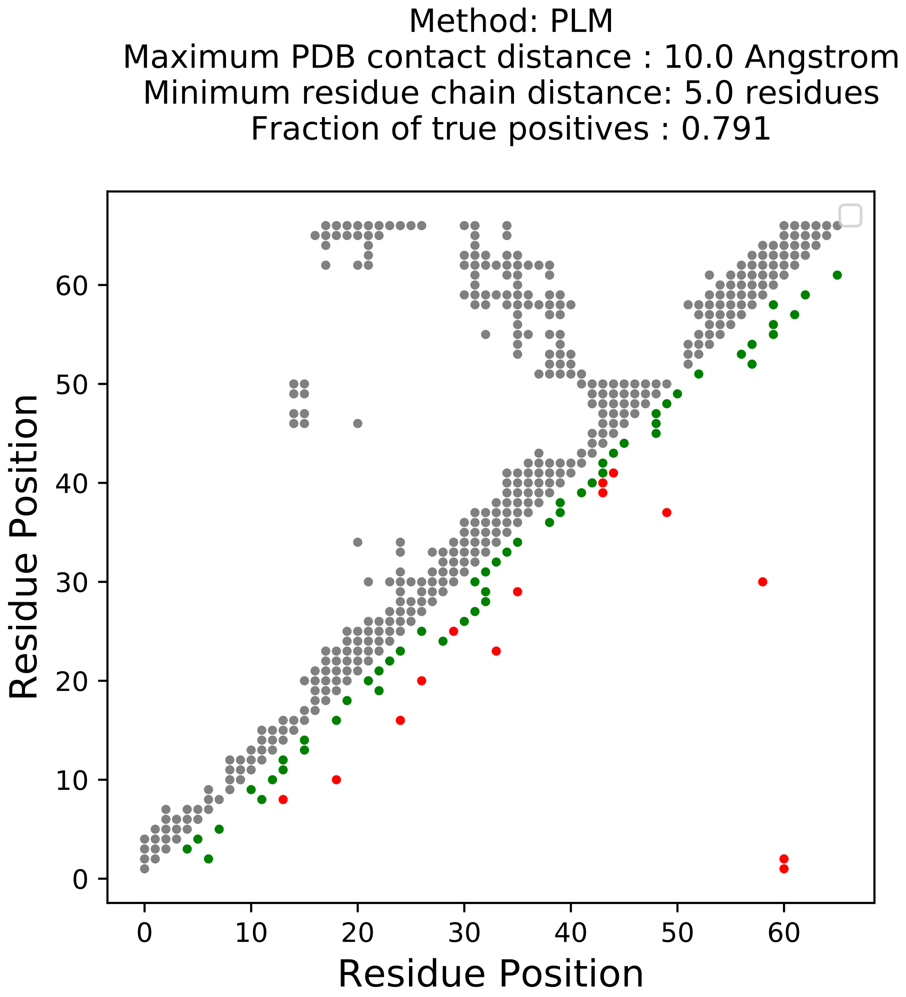

Method: PLM
Maximum PDB contact distance : 10.0 Angstrom
Minimum residue chain distance: 5.0 residues
Fraction of true positives : 0.791
Residue Position
Residue Position
0
10
20
30
40
50
60

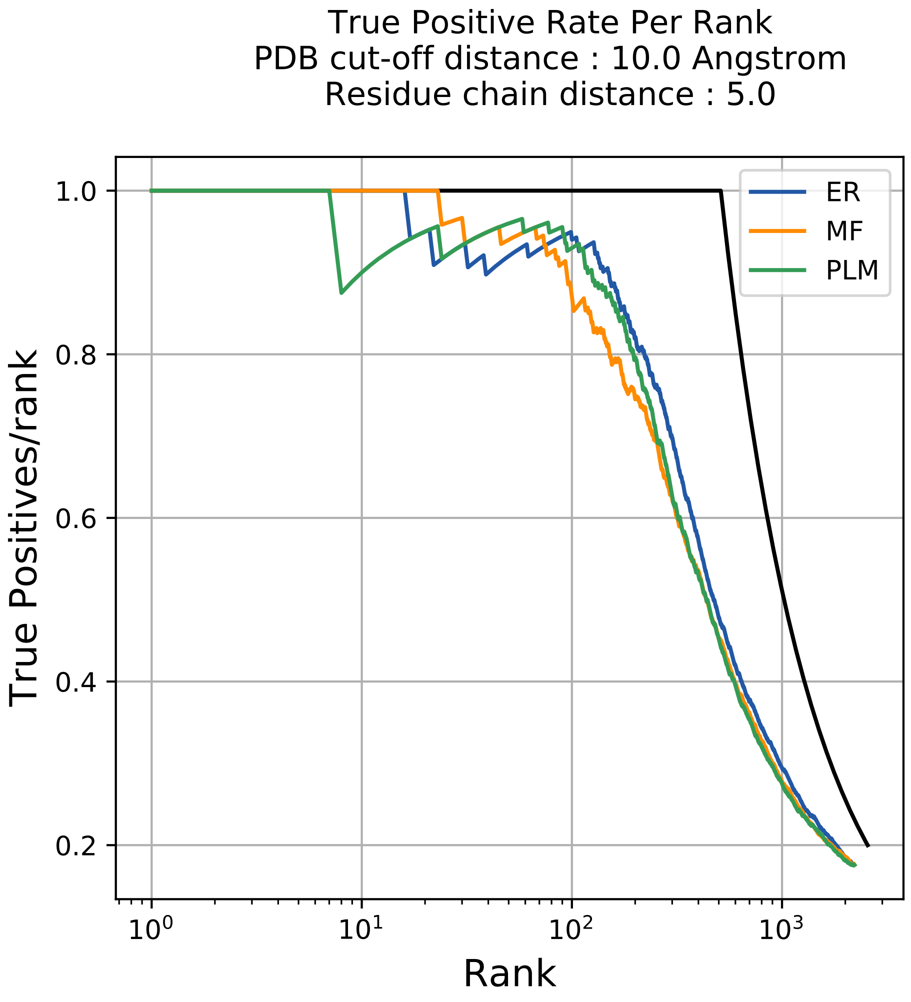
True Positive Rate Per Rank
PDB cut-off distance : 10.0 Angstrom
Residue chain distance : 5.0
True Positives/rank
Rank
1.0
0.8
0.6
0.4
0.2
$10^0$
$10^1$
$10^2$
$10^3$
ER
MF
PLM

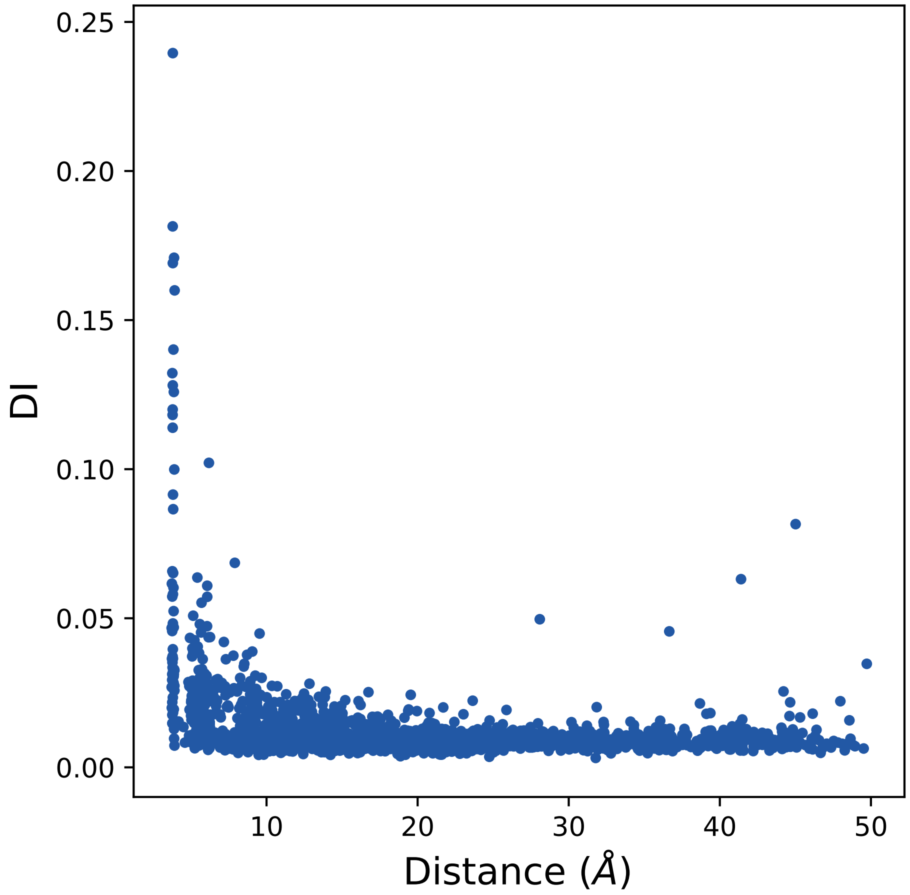
0.25
0.20
0.15
0.10
0.05
0.00
DI
10
20
30
40
50
Distance (Å)

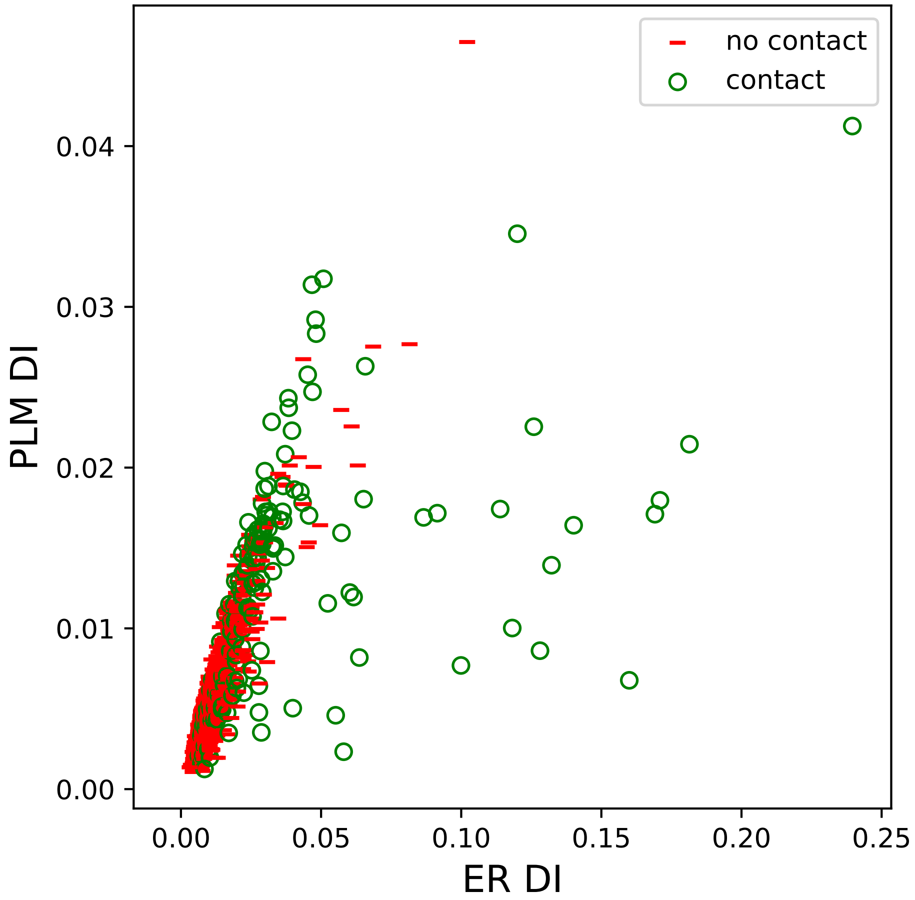

no contact
contact
0.04
0.03
0.02
0.01
0.00
PLM DI
0.00
0.05
0.10
0.15
0.20
0.25
ER DI

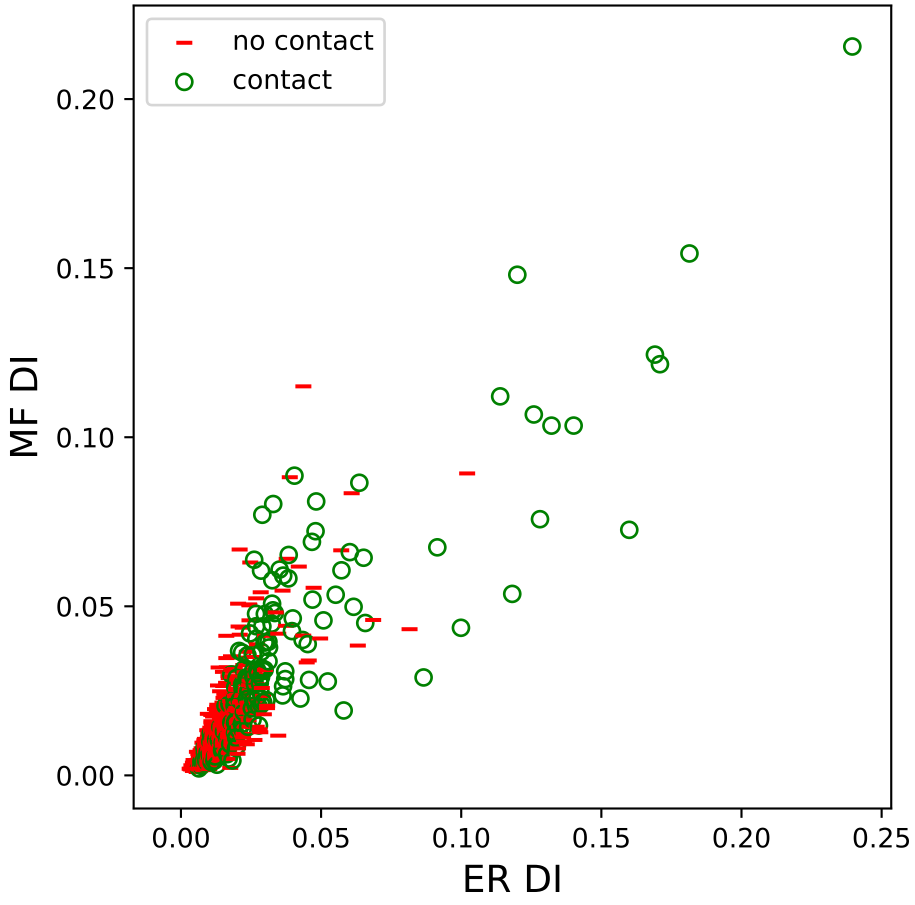
no contact
contact
MF DI
ER DI
0.00
0.05
0.10
0.15
0.20
0.25

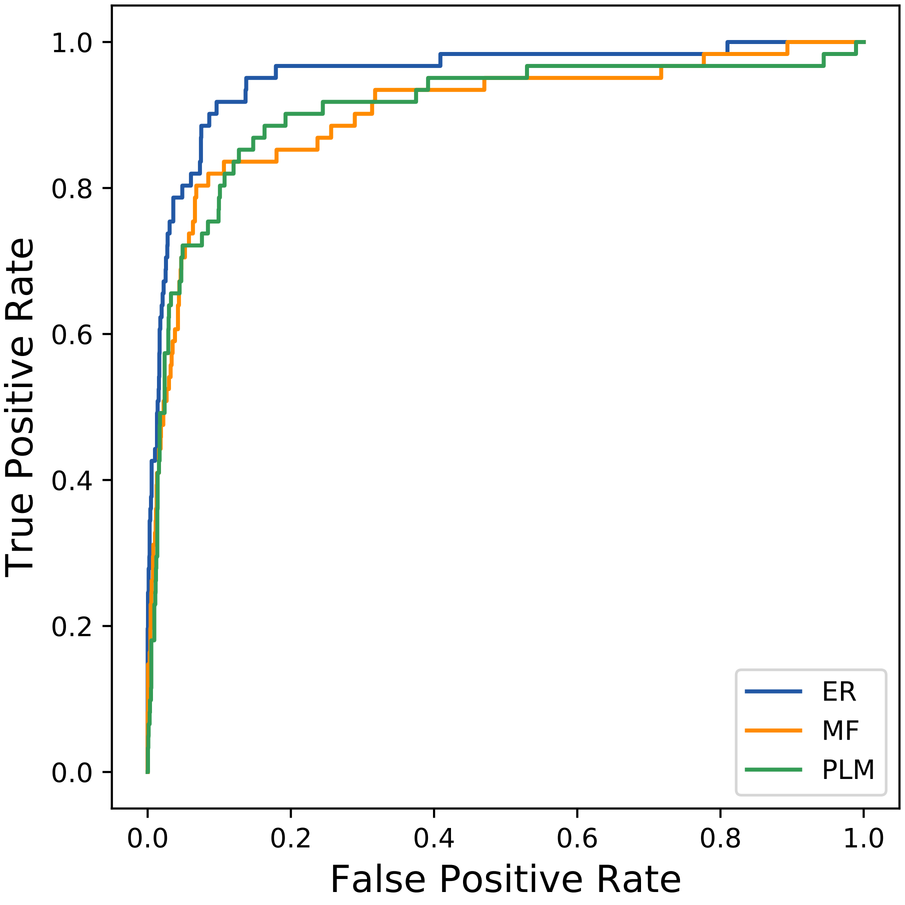

True Positive Rate
False Positive Rate
0.0
0.2
0.4
0.6
0.8
1.0
ER
MF
PLM
