## Supplemental Contact Maps for "Protein Structure Prediction with Expectation Reflection": 23319seq_103col_6s5e_PF02254_plots.pdf

Method: ER
Maximum PDB contact distance : 10.0 Angstrom
Minimum residue chain distance: 5.0 residues
Fraction of true positives : 0.786

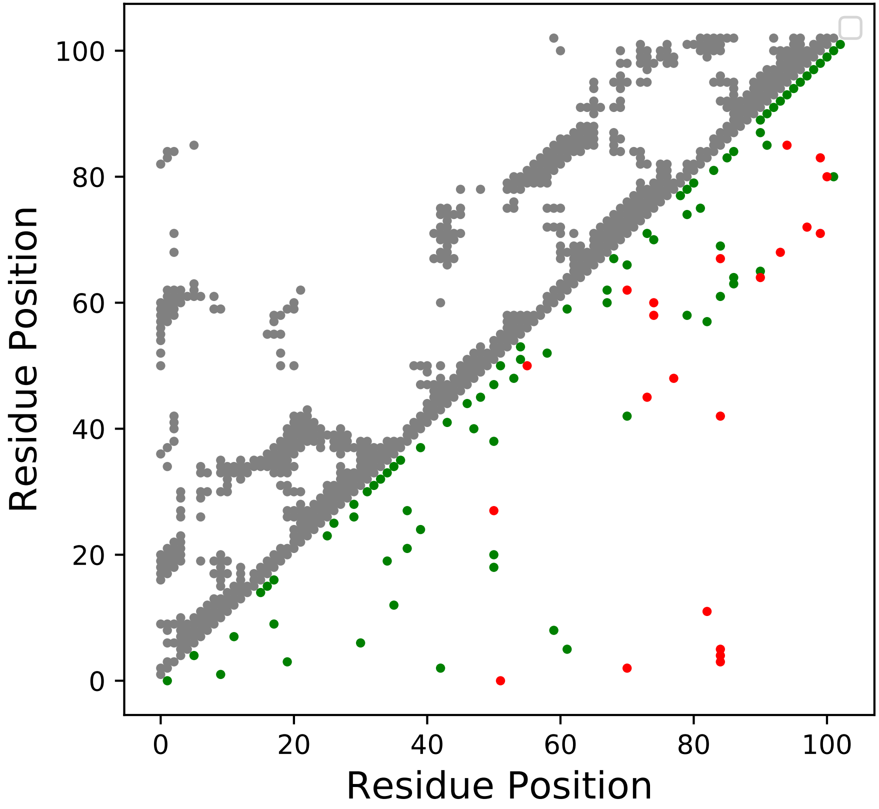

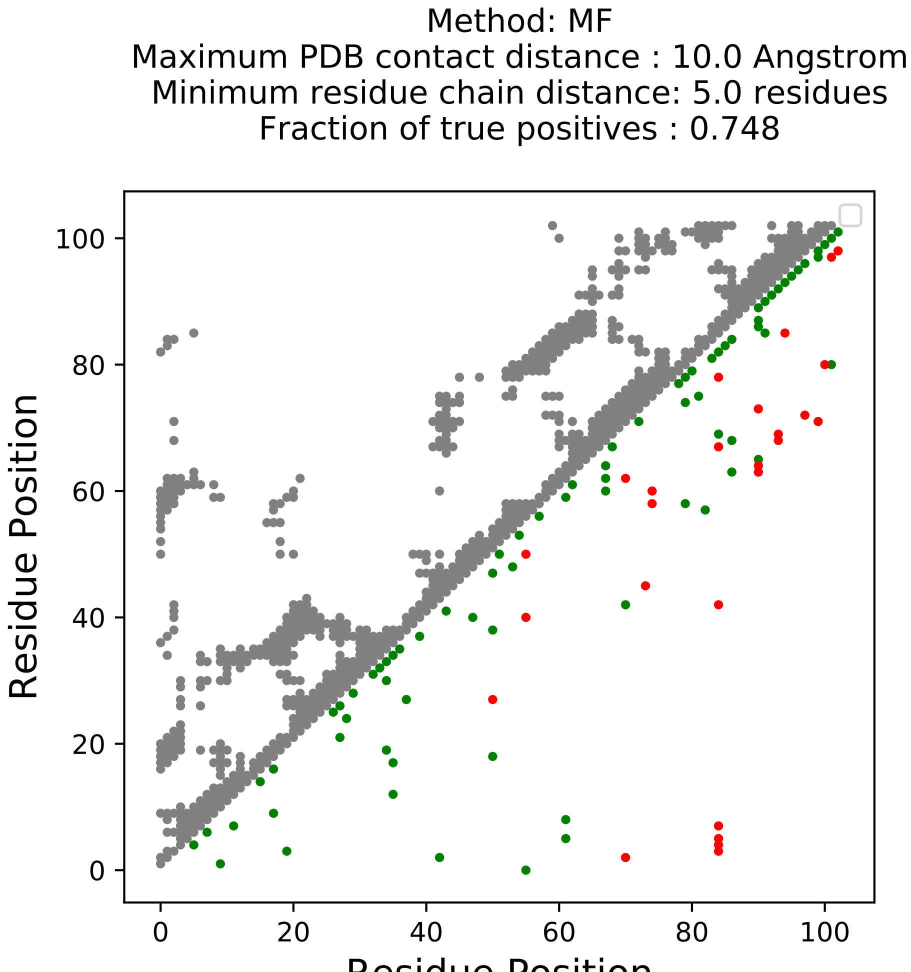
Method: MF
Maximum PDB contact distance : 10.0 Angstrom
Minimum residue chain distance: 5.0 residues
Fraction of true positives : 0.748
Residue Position
Residue Position
0
20
40
60
80
100

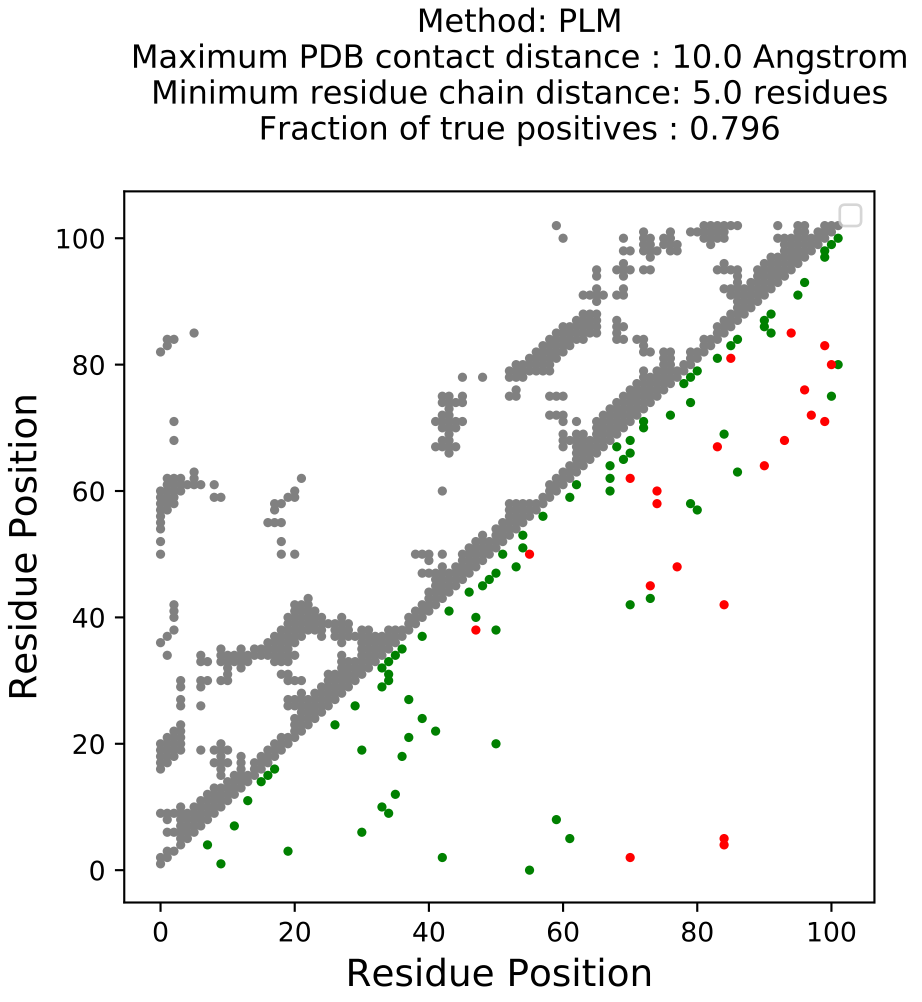

Method: PLM
Maximum PDB contact distance : 10.0 Angstrom
Minimum residue chain distance: 5.0 residues
Fraction of true positives : 0.796
Residue Position
Residue Position
0
20
40
60
80
100

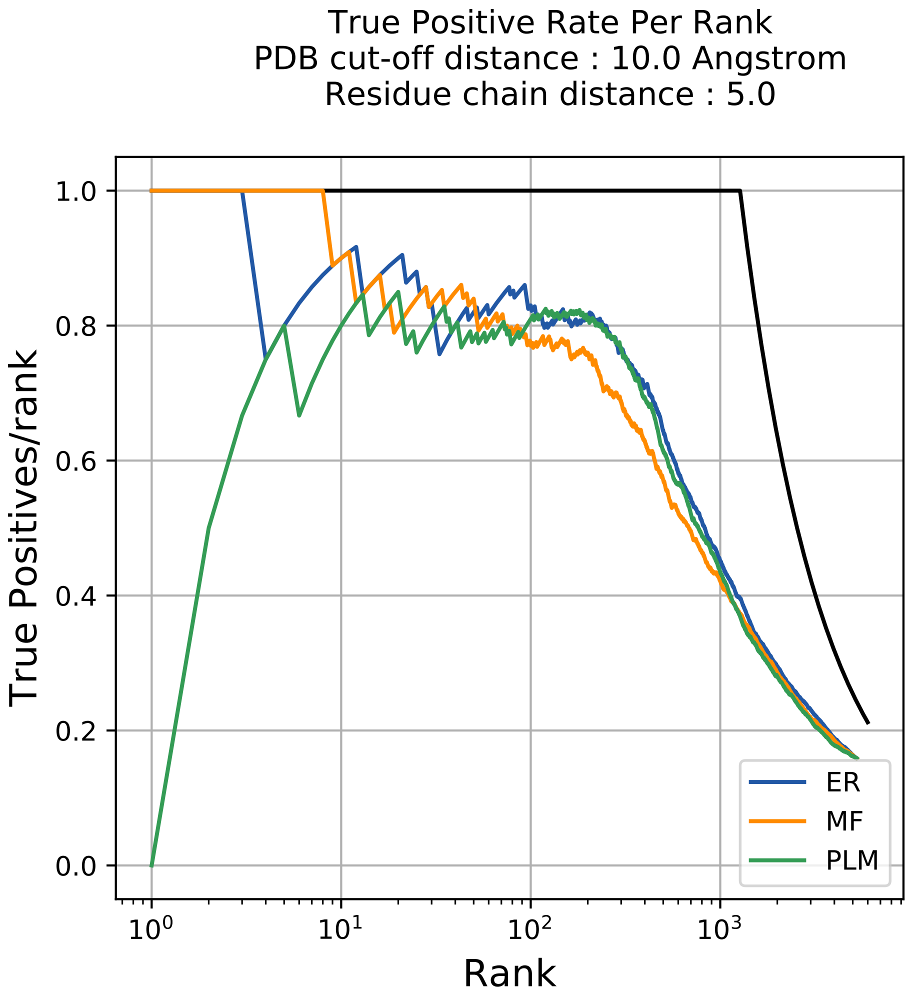

True Positive Rate Per Rank
PDB cut-off distance : 10.0 Angstrom
Residue chain distance : 5.0
True Positives/rank
Rank
1.0
0.8
0.6
0.4
0.2
0.0
$10^0$
$10^1$
$10^2$
$10^3$
ER
MF
PLM

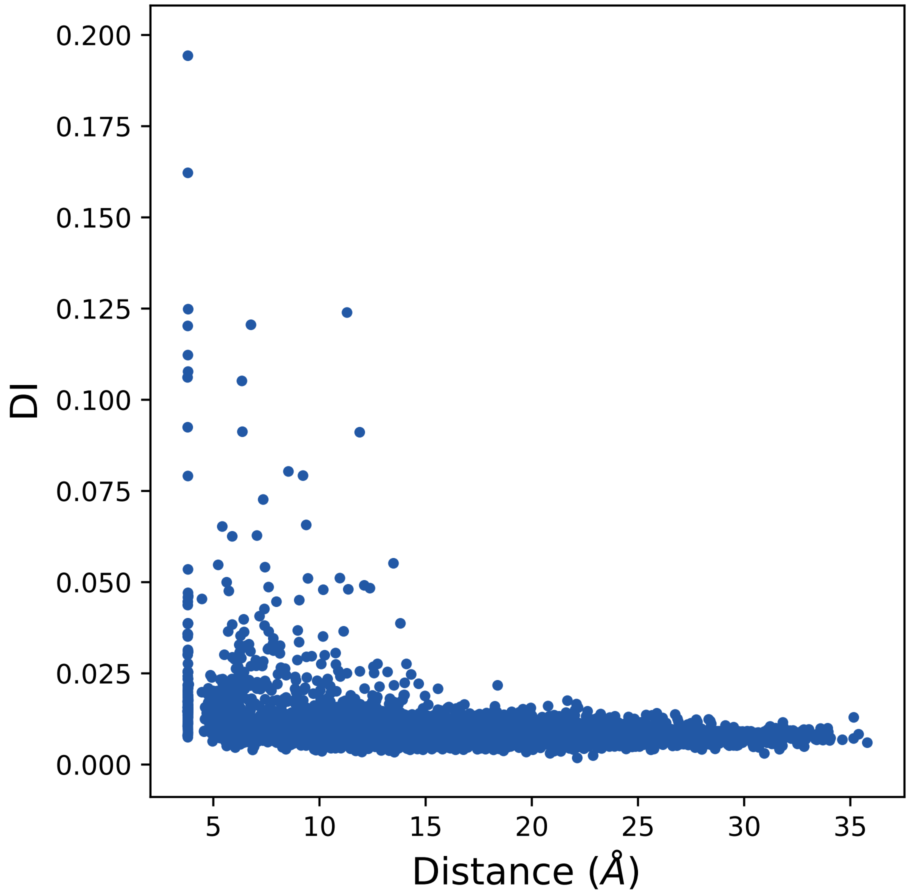

0.200
0.175
0.150
0.125
0.100
0.075
0.050
0.025
0.000
DI
5
10
15
20
25
30
35
Distance (Å)

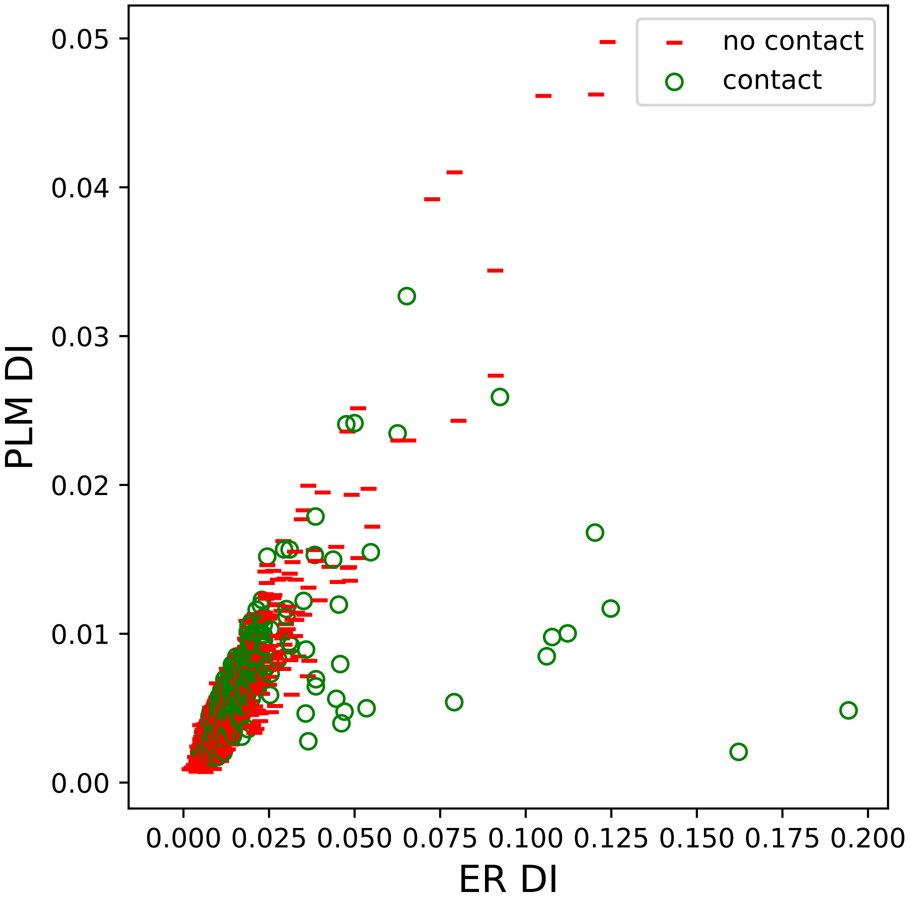

no contact
contact
PLM DI
ER DI
0.05
0.04
0.03
0.02
0.01
0.00
0.000
0.025
0.050
0.075
0.100
0.125
0.150
0.175
0.200

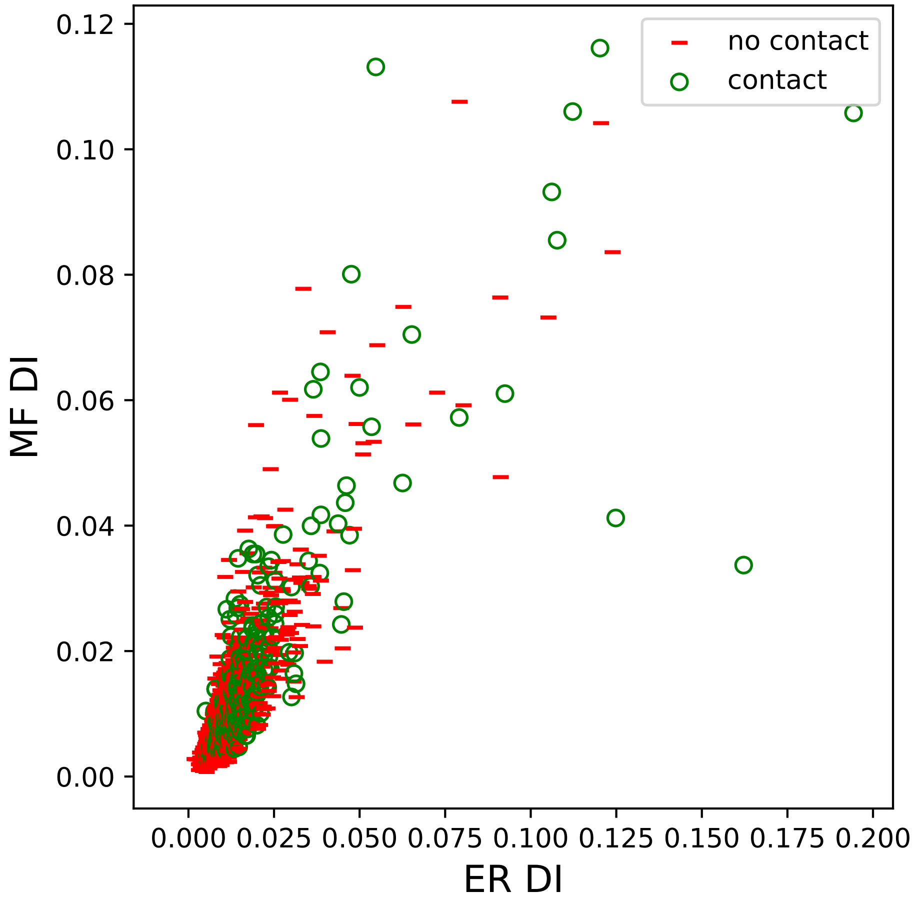

no contact
contact
MF DI
ER DI
0.00
0.02
0.04
0.06
0.08
0.10
0.12
0.000
0.025
0.050
0.075
0.100
0.125
0.150
0.175
0.200

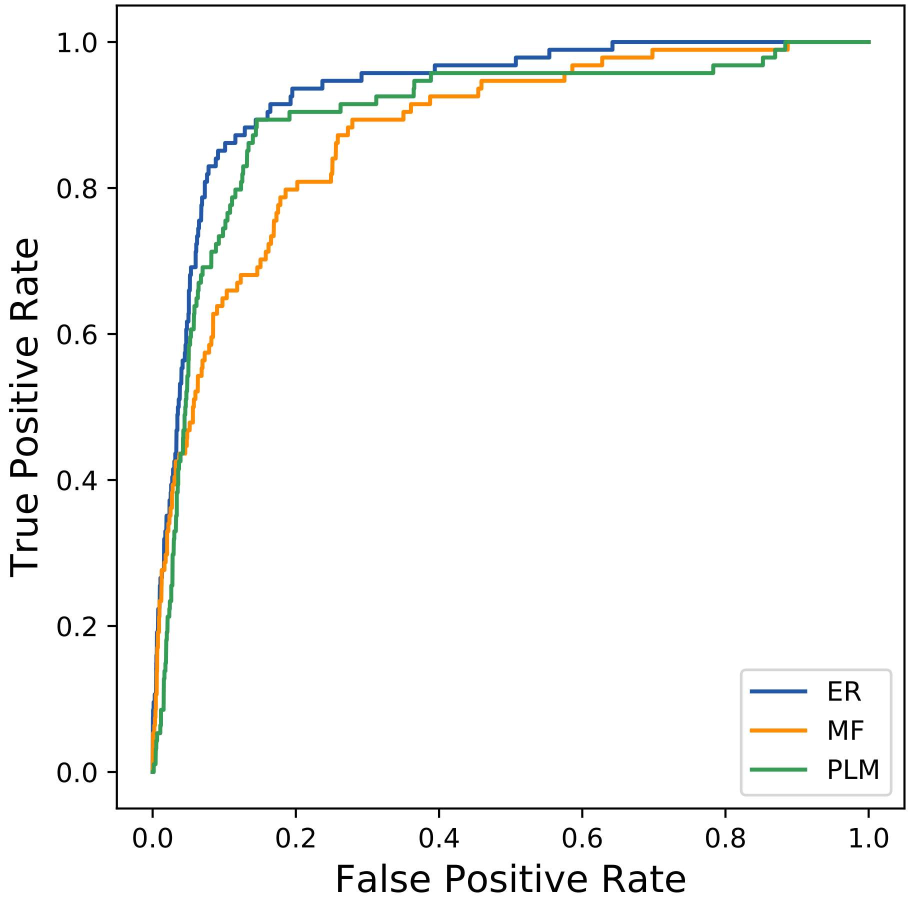

True Positive Rate
False Positive Rate
0.0
0.2
0.4
0.6
0.8
1.0
ER
MF
PLM
