## Supplemental Contact Maps for "Protein Structure Prediction with Expectation Reflection": 23496seq_59col_1rfk_PF00111_plots.pdf

Method: ER
Maximum PDB contact distance : 10.0 Angstrom
Minimum residue chain distance: 5.0 residues
Fraction of true positives : 0.746

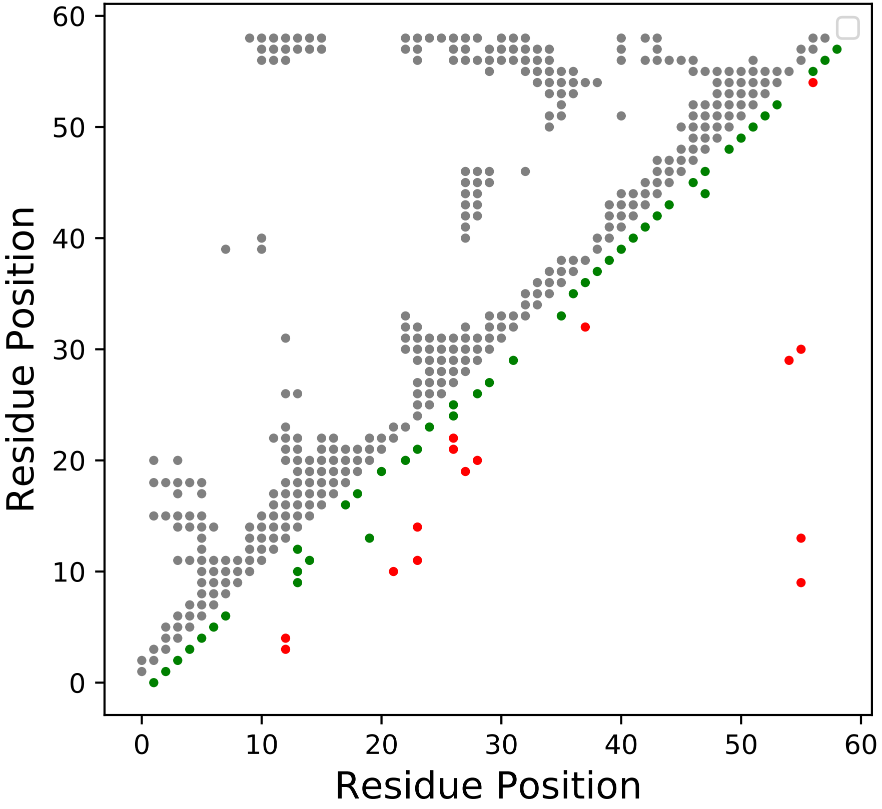

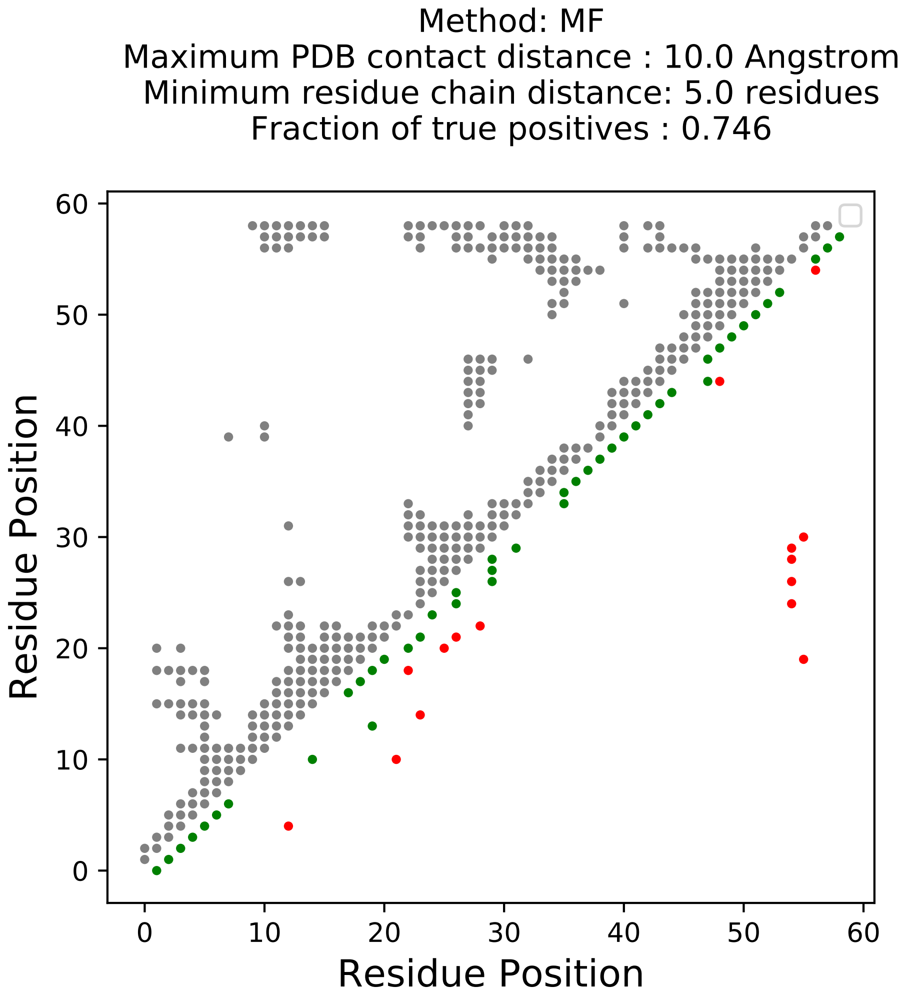
Method: MF
Maximum PDB contact distance : 10.0 Angstrom
Minimum residue chain distance: 5.0 residues
Fraction of true positives : 0.746
Residue Position
Residue Position
0
10
20
30
40
50
60

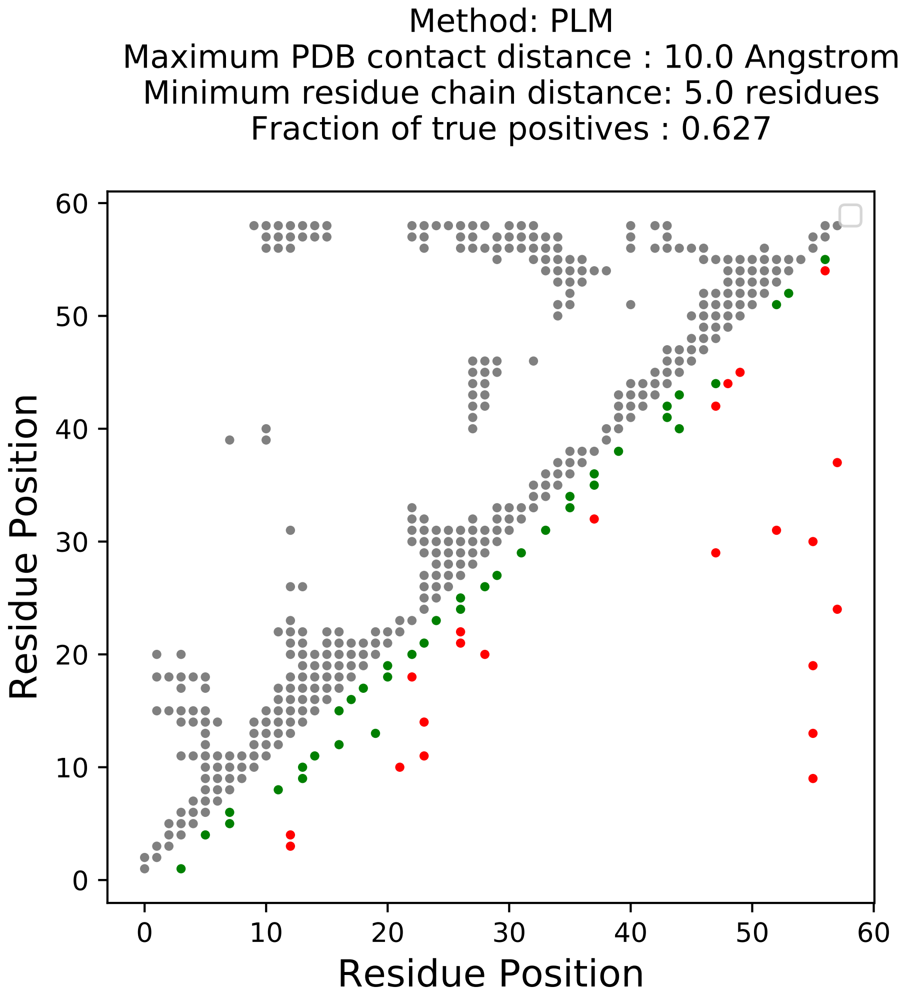
Method: PLM
Maximum PDB contact distance : 10.0 Angstrom
Minimum residue chain distance: 5.0 residues
Fraction of true positives : 0.627
Residue Position
Residue Position
0
10
20
30
40
50
60
0
10
20
30
40
50
60

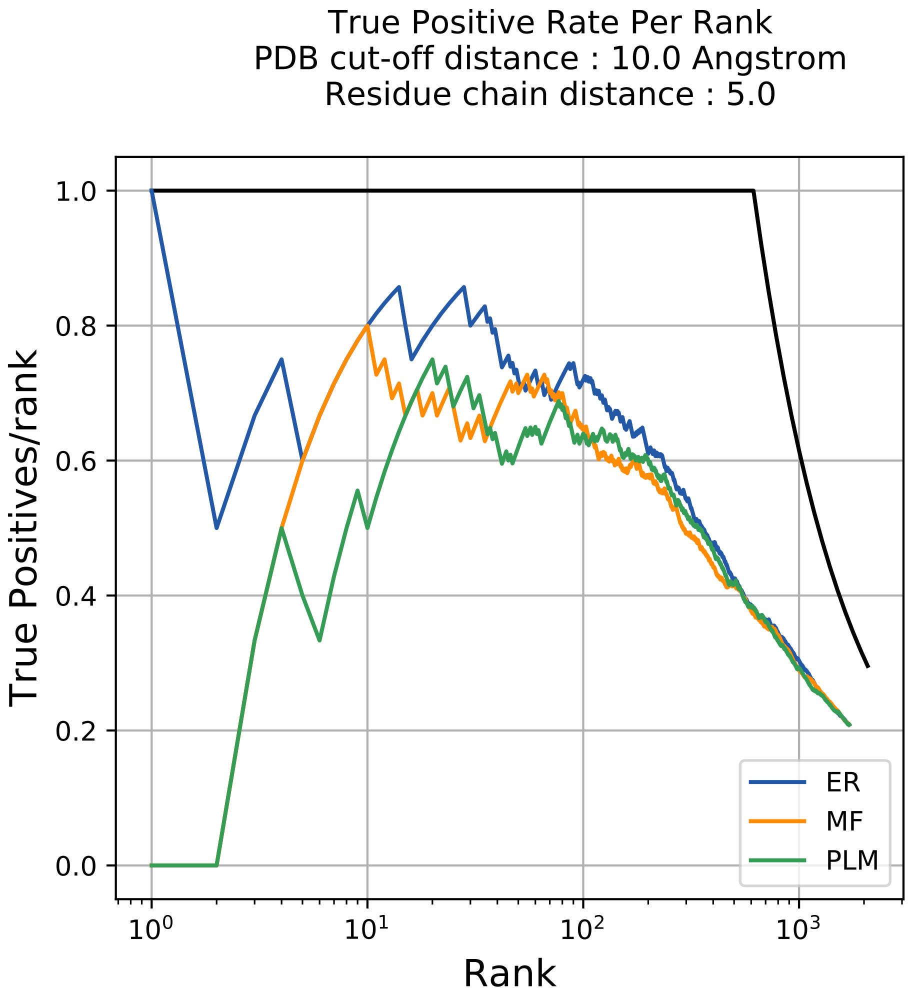
True Positive Rate Per Rank
PDB cut-off distance : 10.0 Angstrom
Residue chain distance : 5.0
True Positives/rank
Rank
1.0
0.8
0.6
0.4
0.2
0.0
$10^0$
$10^1$
$10^2$
$10^3$
ER
MF
PLM

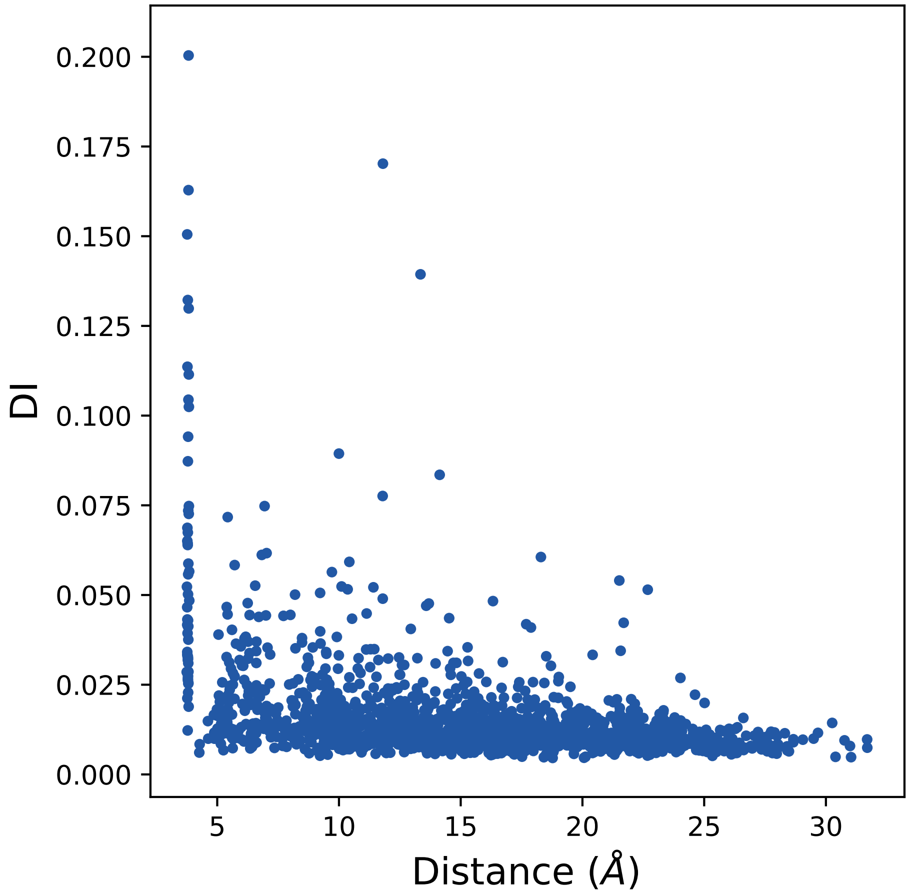

0.200
0.175
0.150
0.125
0.100
0.075
0.050
0.025
0.000
DI
5
10
15
20
25
30
Distance (Å)

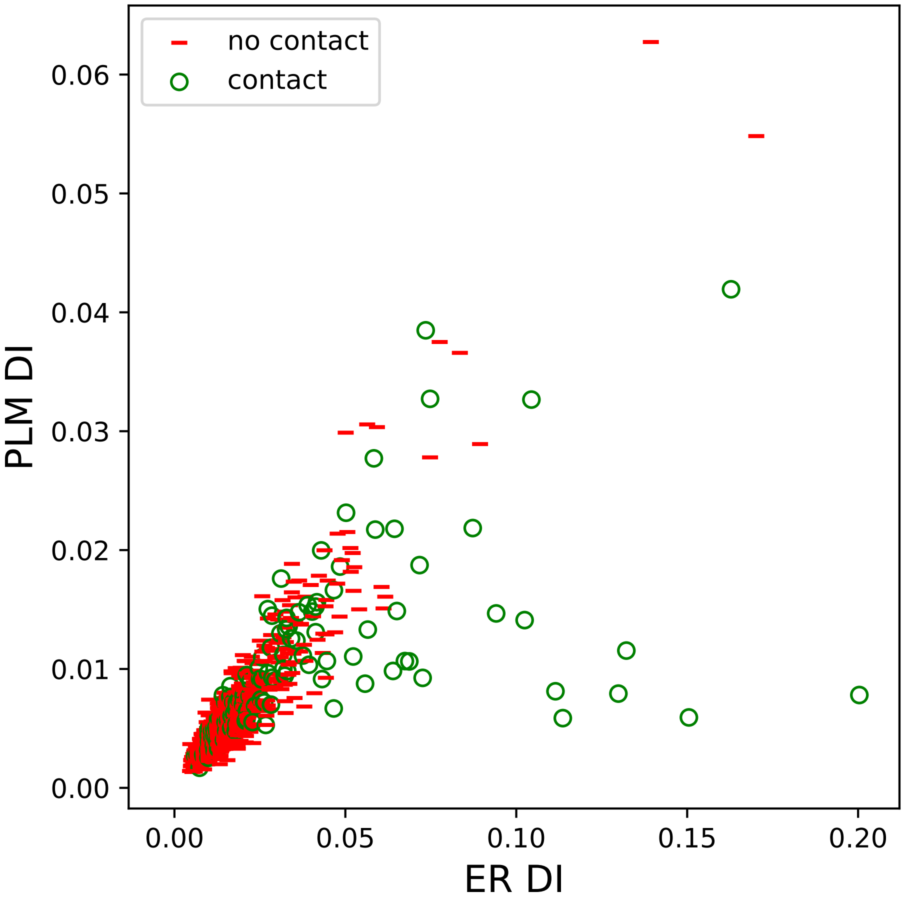

no contact
contact
0.06
0.05
0.04
0.03
0.02
0.01
0.00
PLM DI
0.00
0.05
0.10
0.15
0.20
ER DI

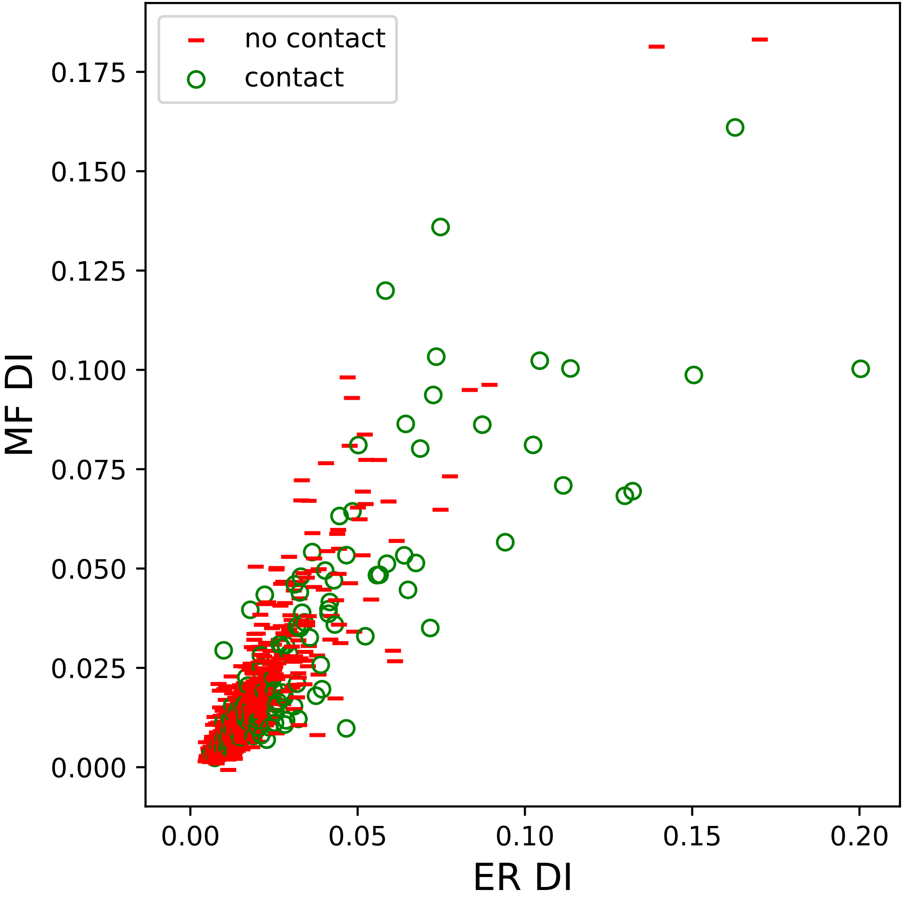
no contact
contact
0.175
0.150
0.125
0.100
0.075
0.050
0.025
0.000
MF DI
0.00
0.05
0.10
0.15
0.20
ER DI

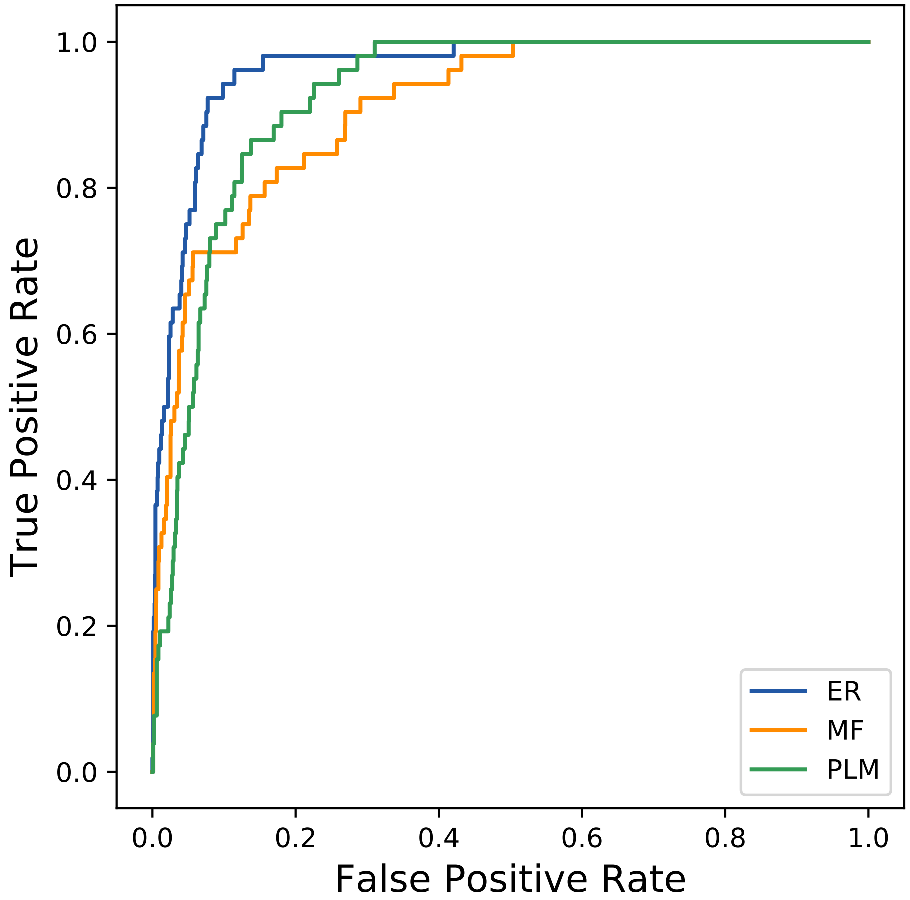

True Positive Rate
False Positive Rate
1.0
0.8
0.6
0.4
0.2
0.0
0.0
0.2
0.4
0.6
0.8
1.0
ER
MF
PLM
