## Supplemental Contact Maps for "Protein Structure Prediction with Expectation Reflection": 28072seq_142col_7c23_PF13472_plots.pdf

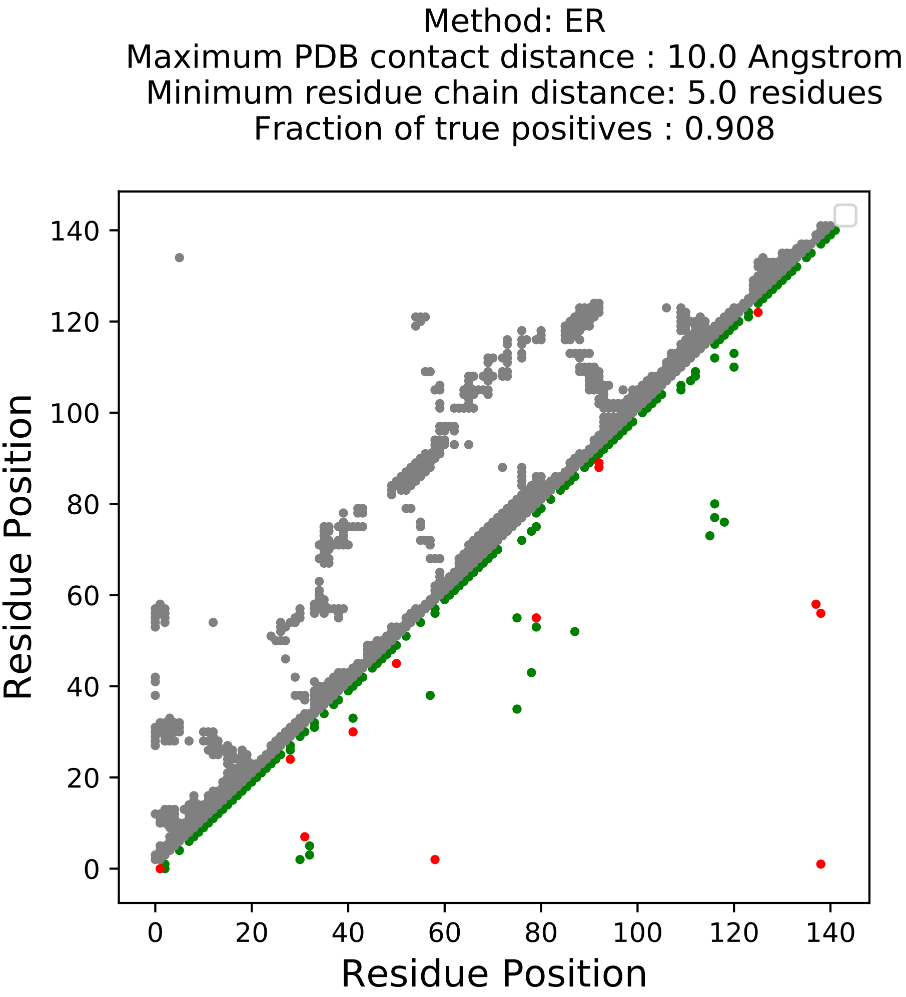
Method: ER
Maximum PDB contact distance : 10.0 Angstrom
Minimum residue chain distance: 5.0 residues
Fraction of true positives : 0.908
Residue Position
Residue Position
0
20
40
60
80
100
120
140

Method: MF
Maximum PDB contact distance : 10.0 Angstrom
Minimum residue chain distance: 5.0 residues
Fraction of true positives : 0.852

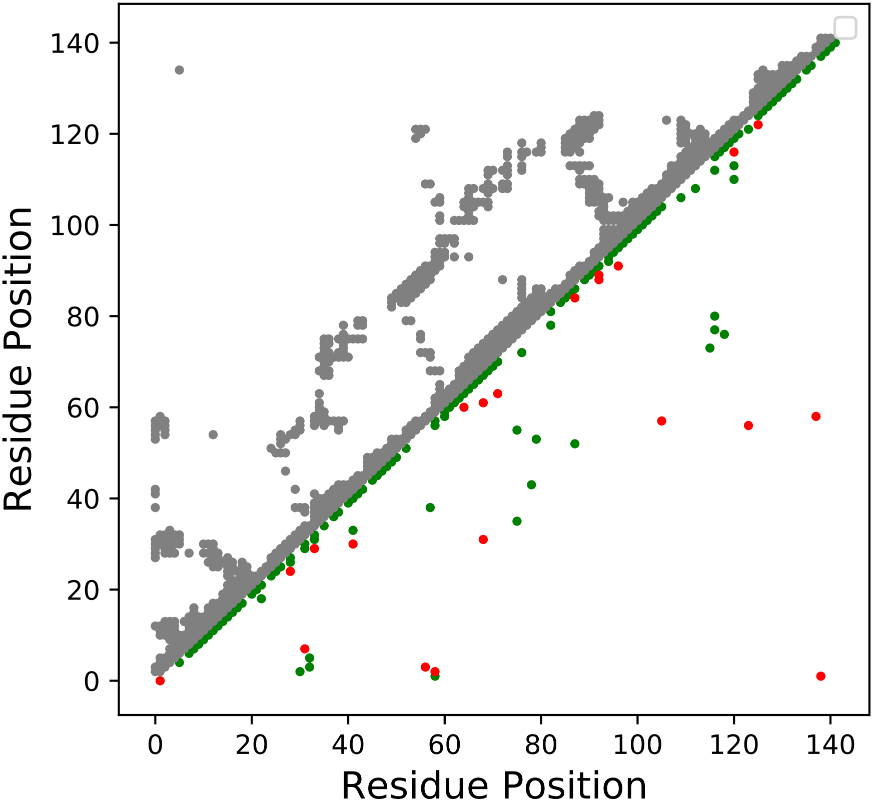

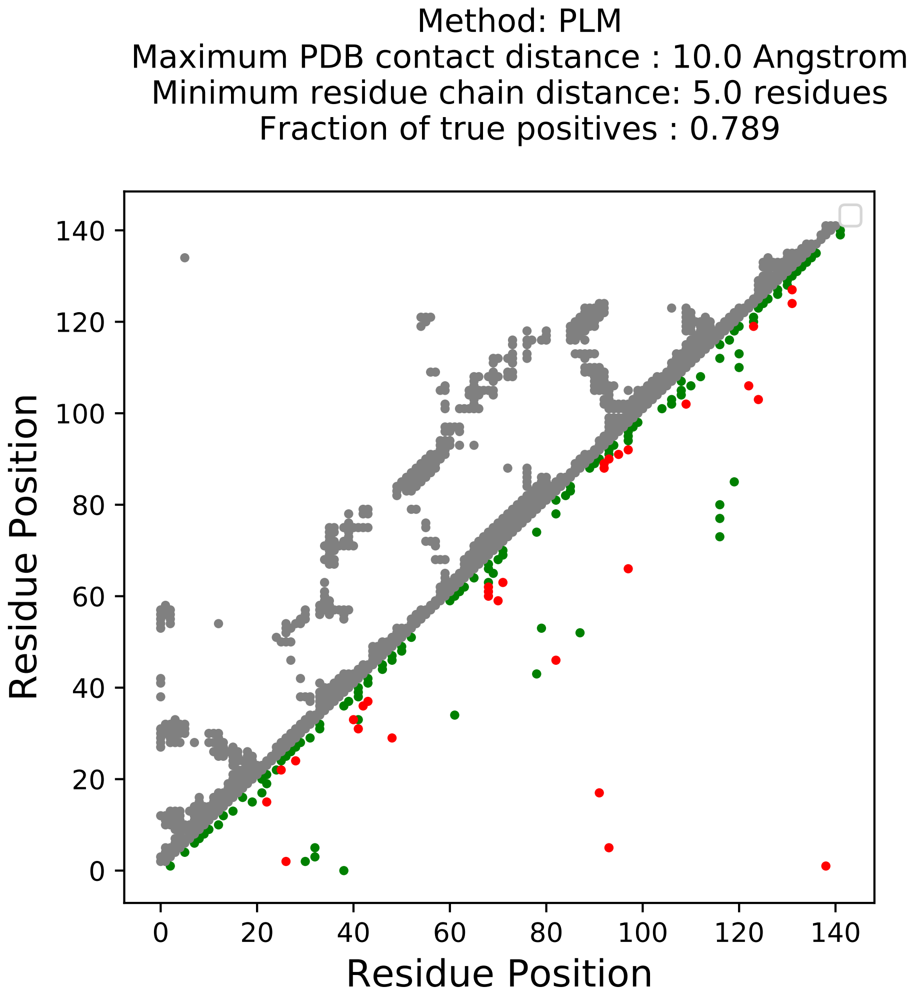
Method: PLM
Maximum PDB contact distance : 10.0 Angstrom
Minimum residue chain distance: 5.0 residues
Fraction of true positives : 0.789
Residue Position
Residue Position
0
20
40
60
80
100
120
140

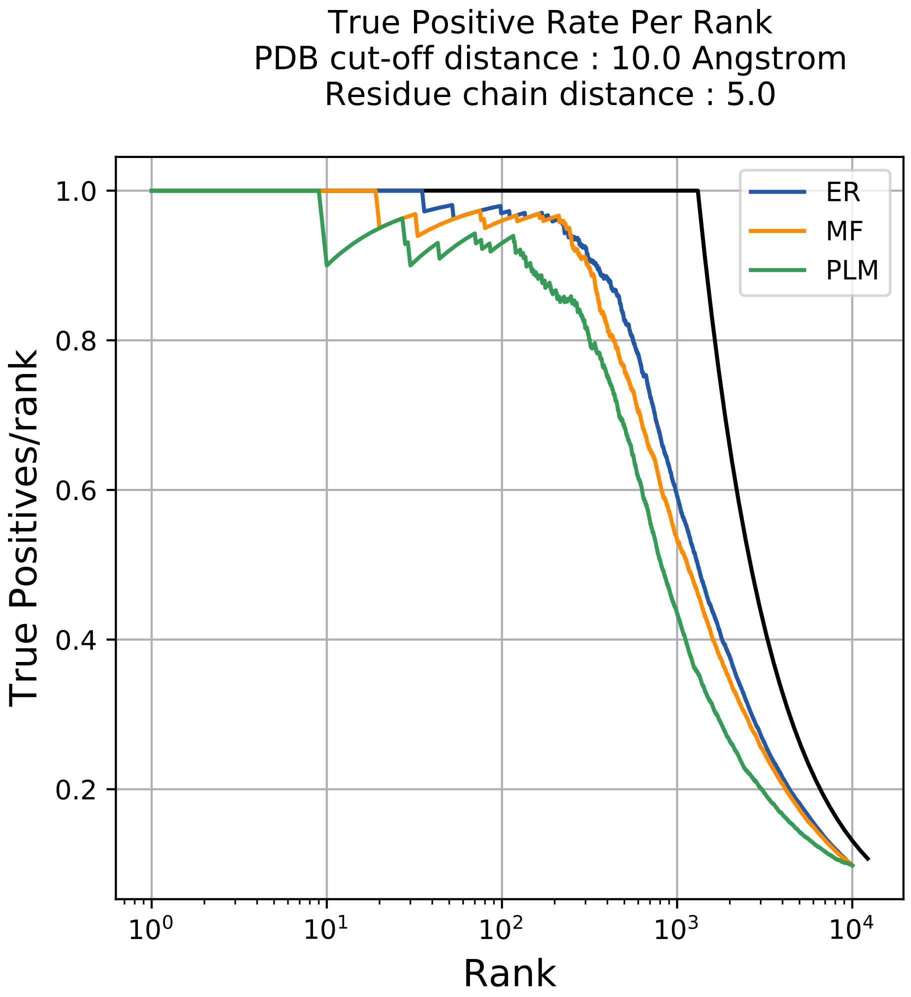

True Positive Rate Per Rank
PDB cut-off distance : 10.0 Angstrom
Residue chain distance : 5.0
True Positives/rank
Rank
ER
MF
PLM
1.0
0.8
0.6
0.4
0.2
$10^0$
$10^1$
$10^2$
$10^3$
$10^4$

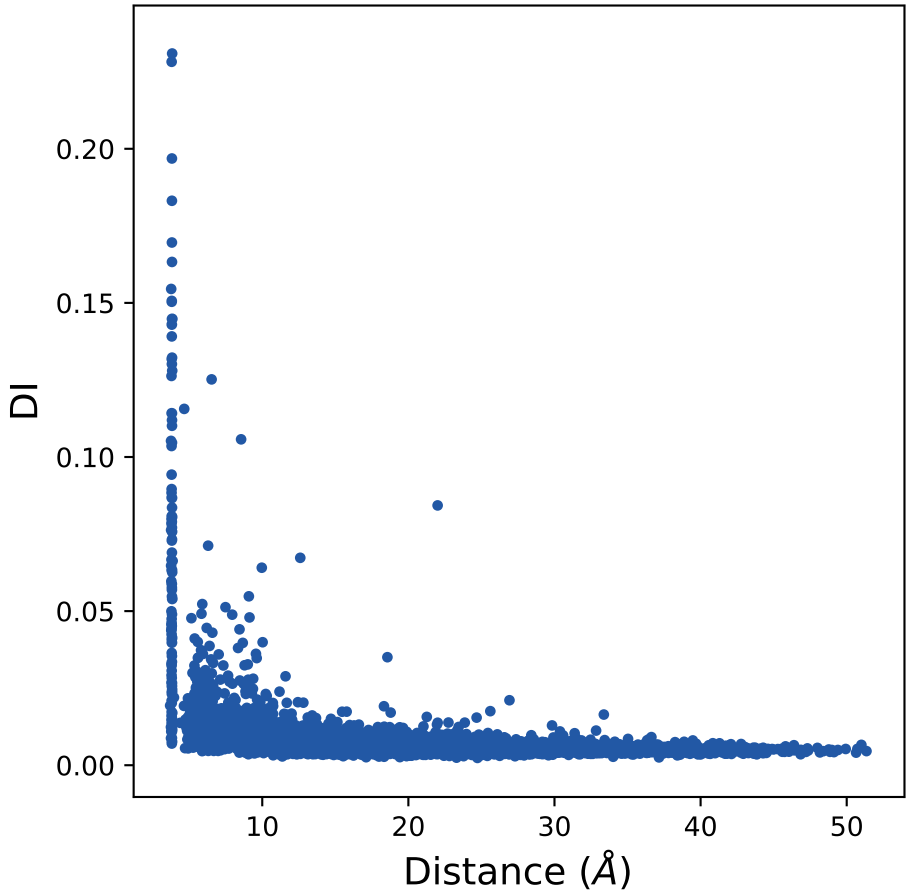

DI
0.00
0.05
0.10
0.15
0.20
10
20
30
40
50
Distance (Å)

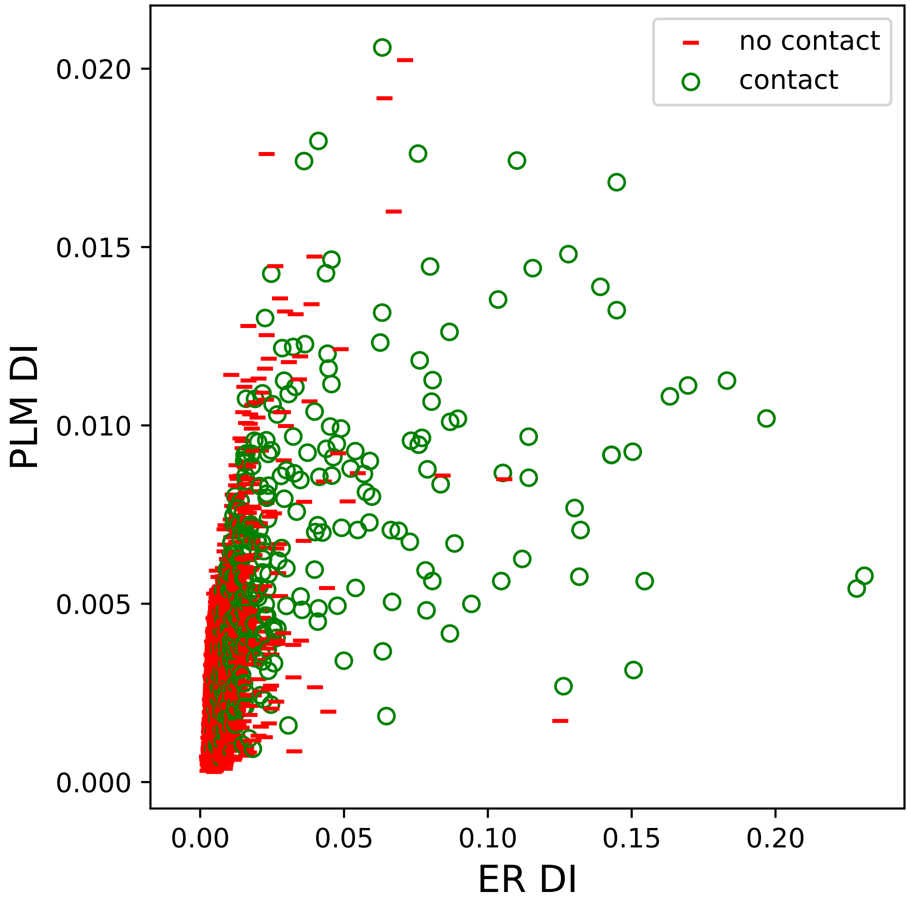
no contact
contact
0.020
0.015
0.010
0.005
0.000
PLM DI
0.00
0.05
0.10
0.15
0.20
ER DI

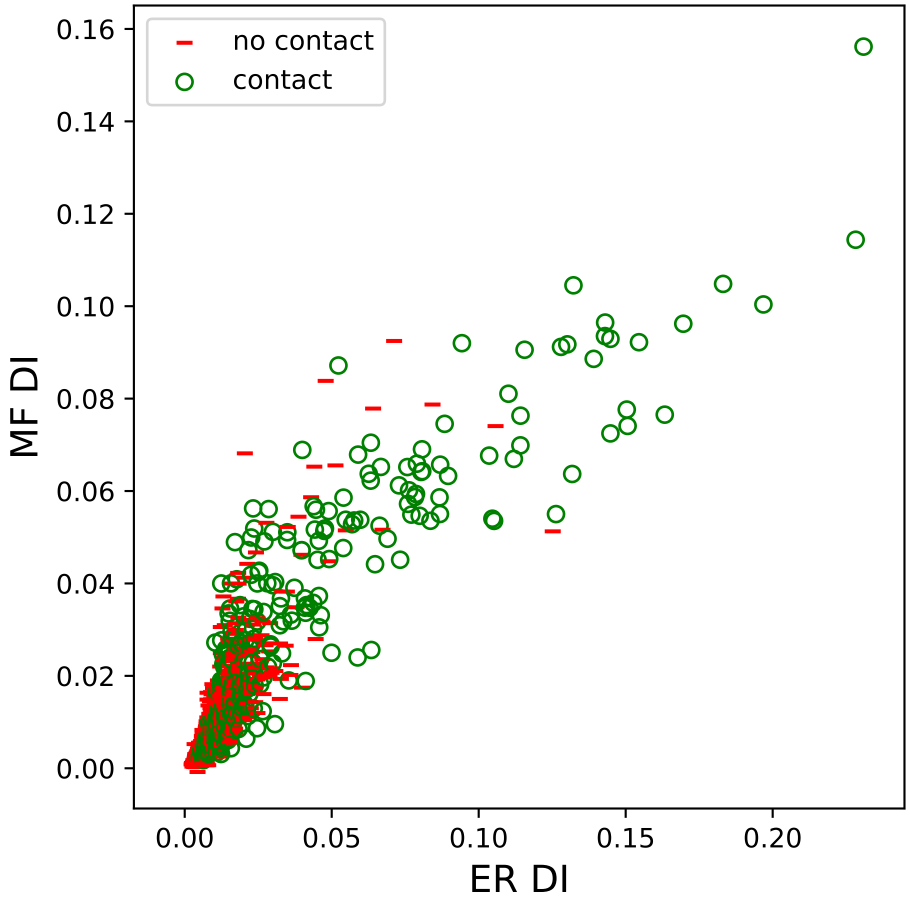

no contact
contact
0.16
0.14
0.12
0.10
0.08
0.06
0.04
0.02
0.00
MF DI
0.00
0.05
0.10
0.15
0.20
ER DI

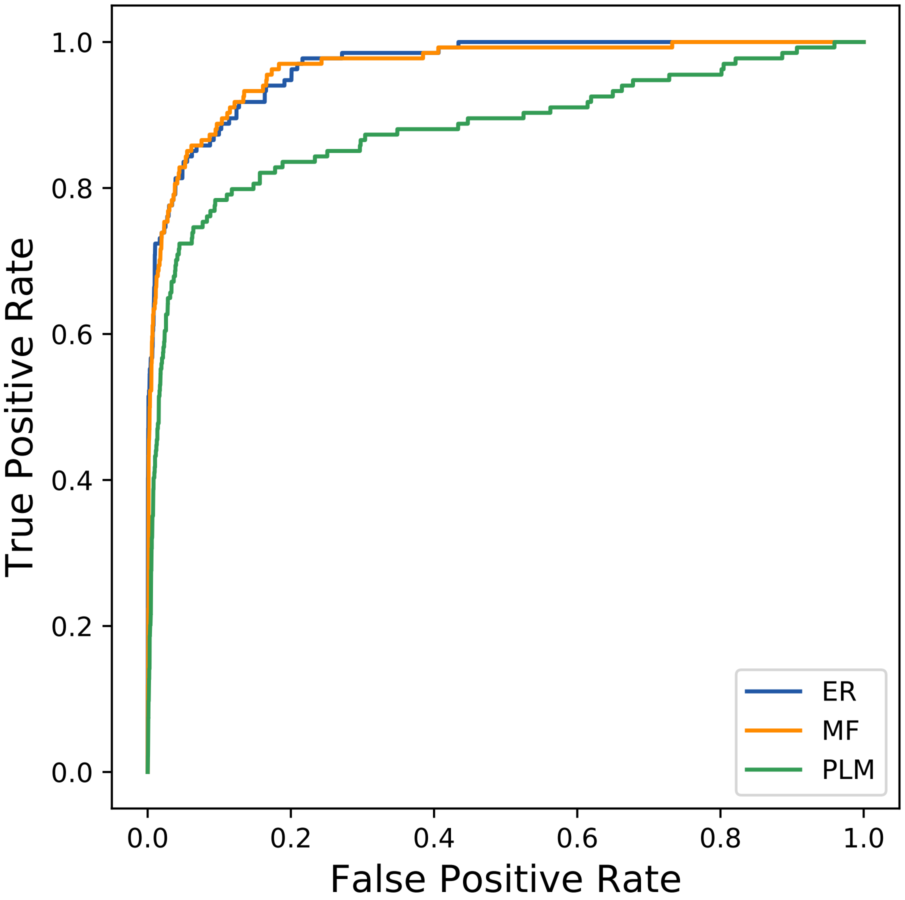

True Positive Rate
False Positive Rate
0.0
0.2
0.4
0.6
0.8
1.0
ER
MF
PLM
