## Supplemental Contact Maps for "Protein Structure Prediction with Expectation Reflection": 29086seq_151col_5ims_PF02776_plots.pdf

Method: ER
Maximum PDB contact distance : 10.0 Angstrom
Minimum residue chain distance: 5.0 residues
Fraction of true positives : 0.841

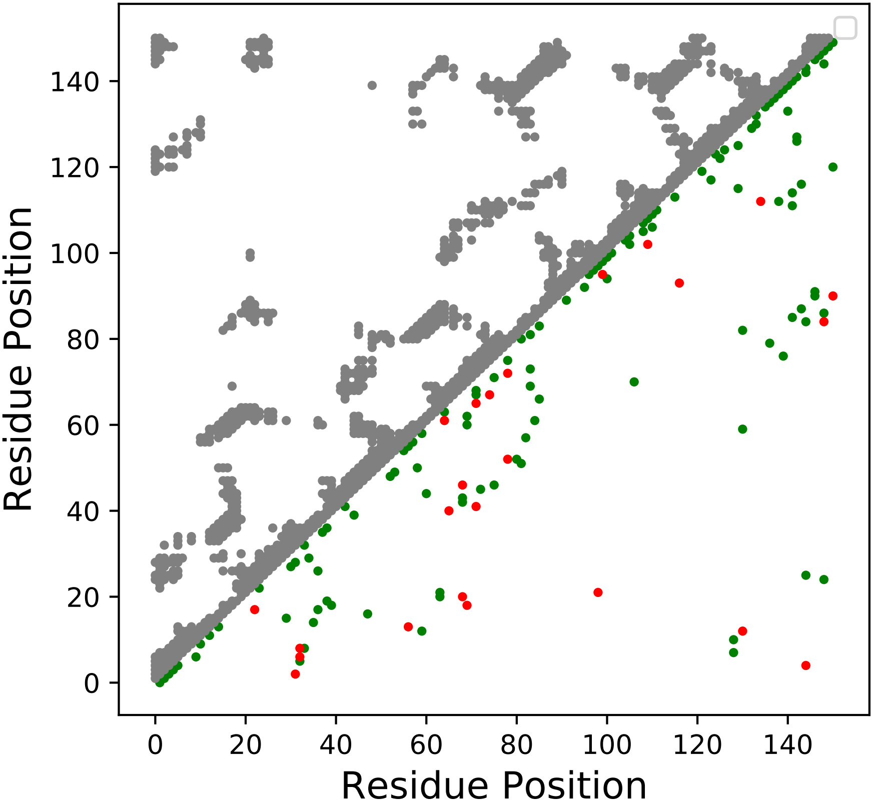

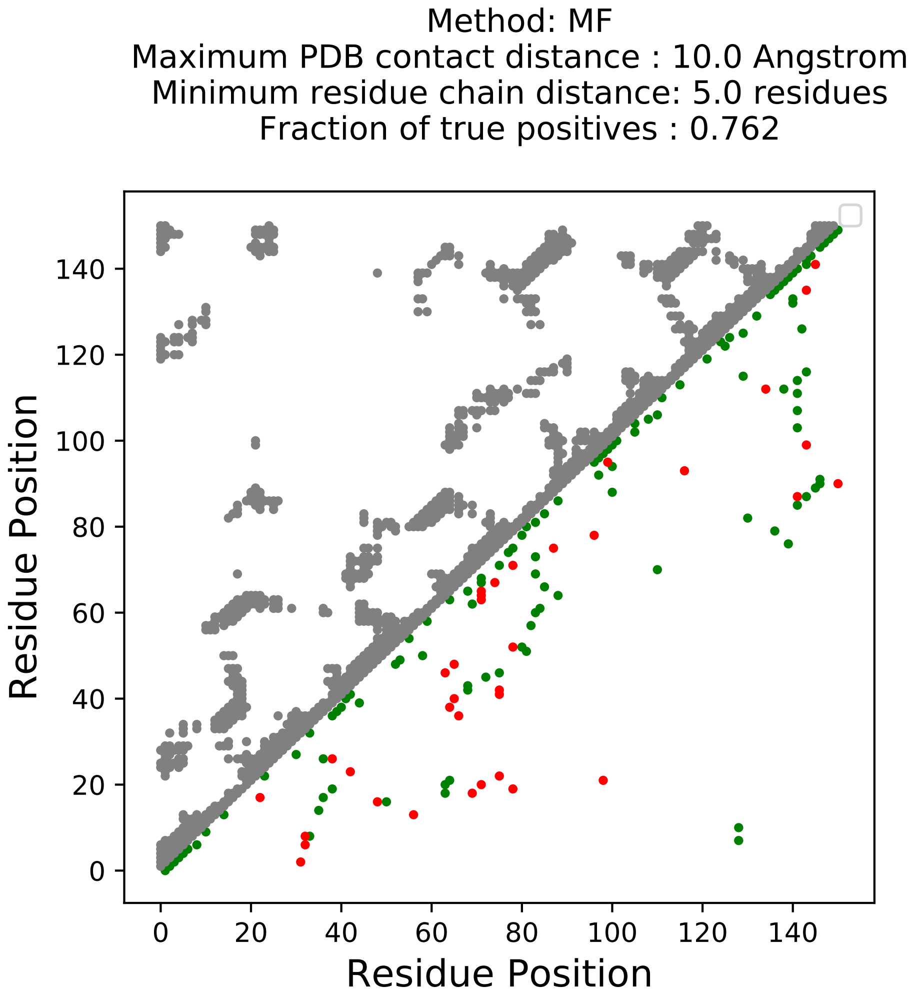

Method: MF
Maximum PDB contact distance : 10.0 Angstrom
Minimum residue chain distance: 5.0 residues
Fraction of true positives : 0.762
Residue Position
Residue Position
0
20
40
60
80
100
120
140

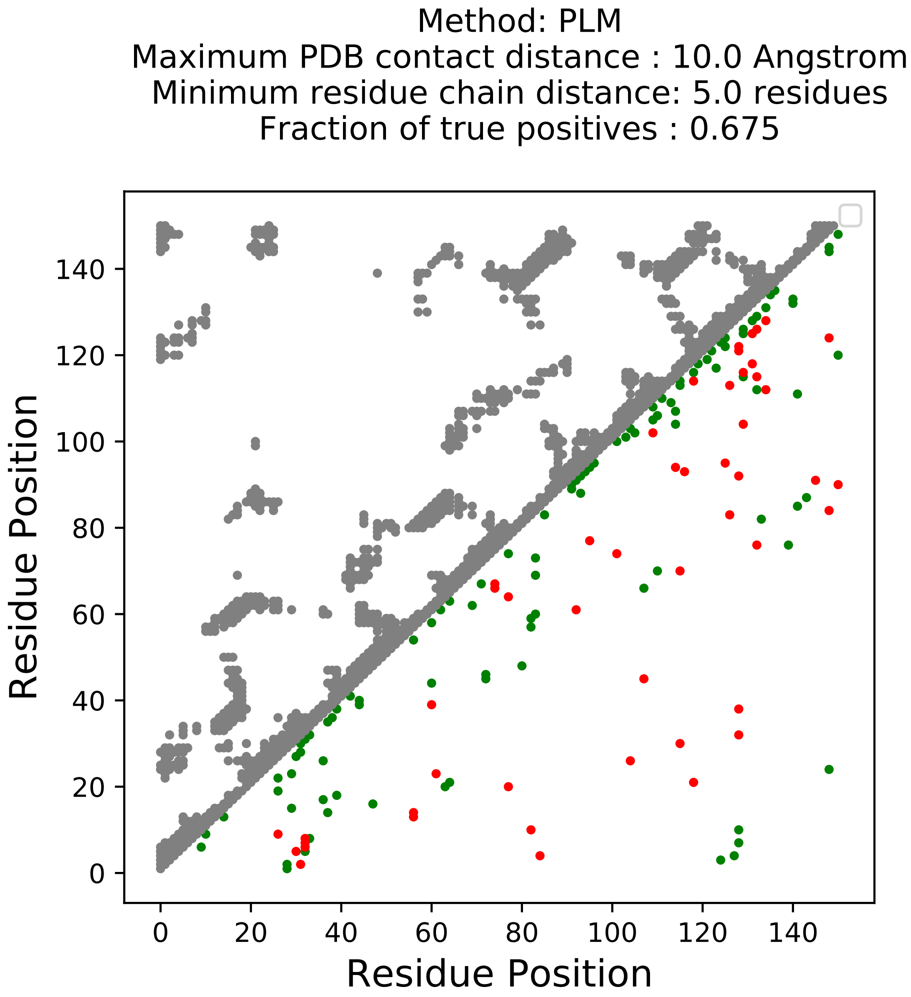
Method: PLM
Maximum PDB contact distance : 10.0 Angstrom
Minimum residue chain distance: 5.0 residues
Fraction of true positives : 0.675
Residue Position
Residue Position
0
20
40
60
80
100
120
140

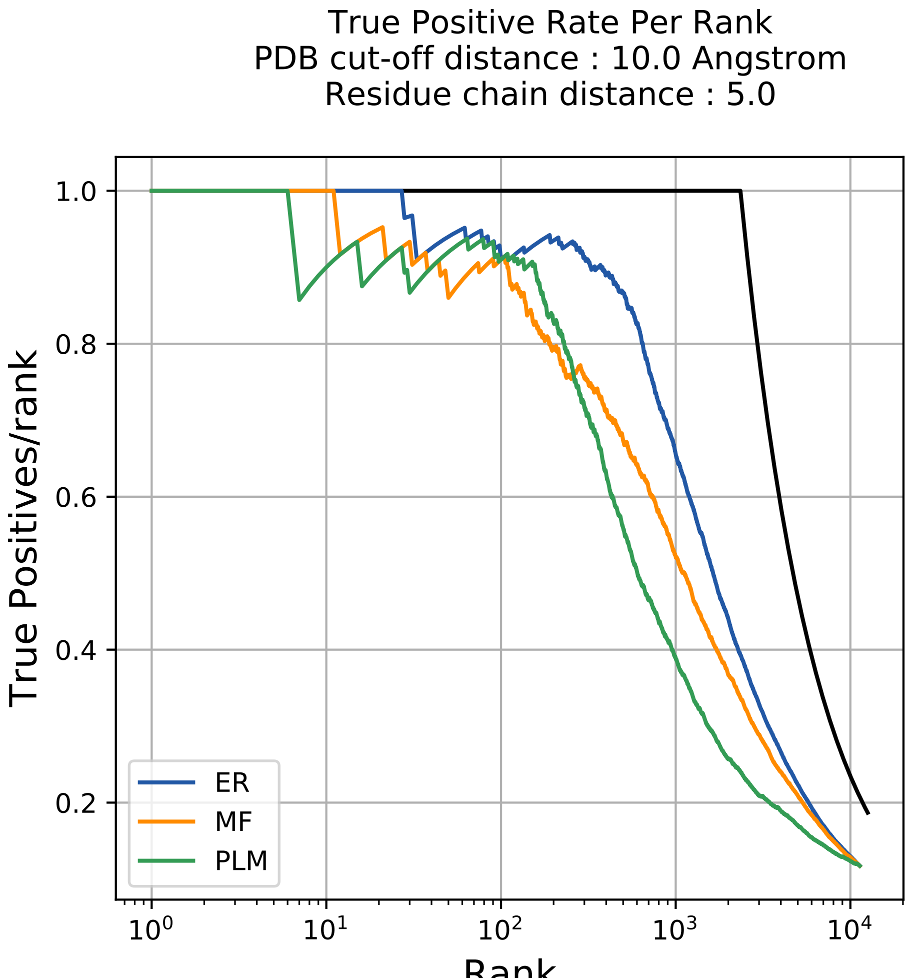

True Positive Rate Per Rank
PDB cut-off distance : 10.0 Angstrom
Residue chain distance : 5.0
True Positives/rank
Rank
1.0
0.8
0.6
0.4
0.2
$10^0$
$10^1$
$10^2$
$10^3$
$10^4$
ER
MF
PLM

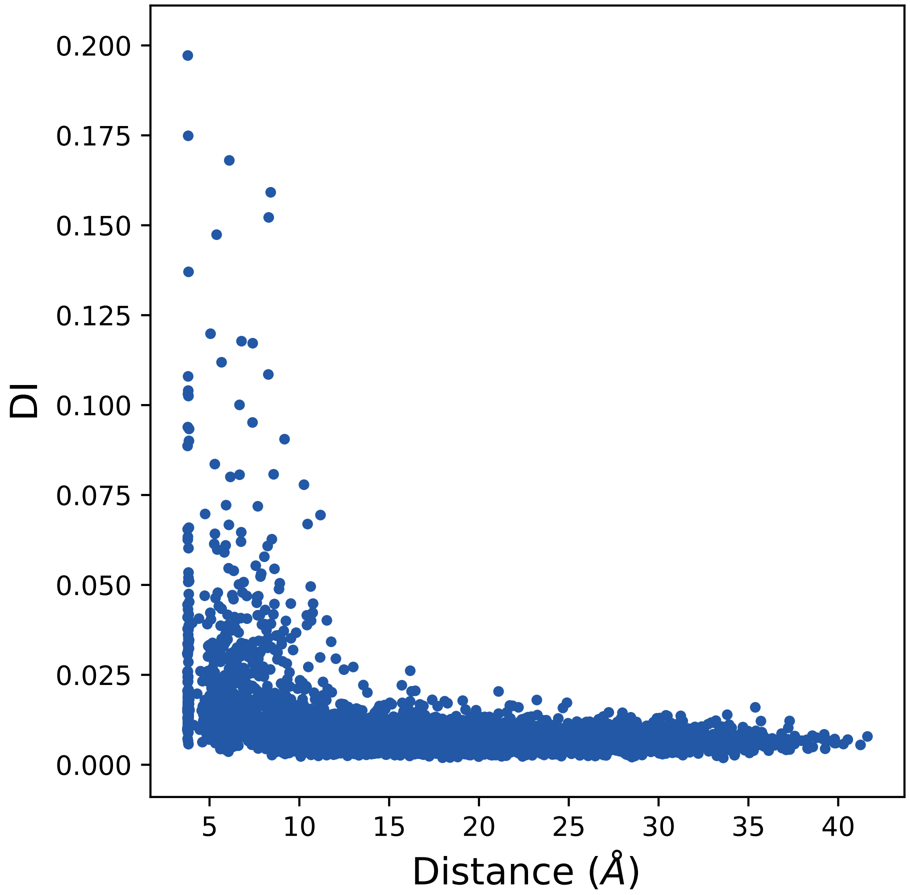
0.200
0.175
0.150
0.125
0.100
0.075
0.050
0.025
0.000
DI
5
10
15
20
25
30
35
40
Distance (Å)

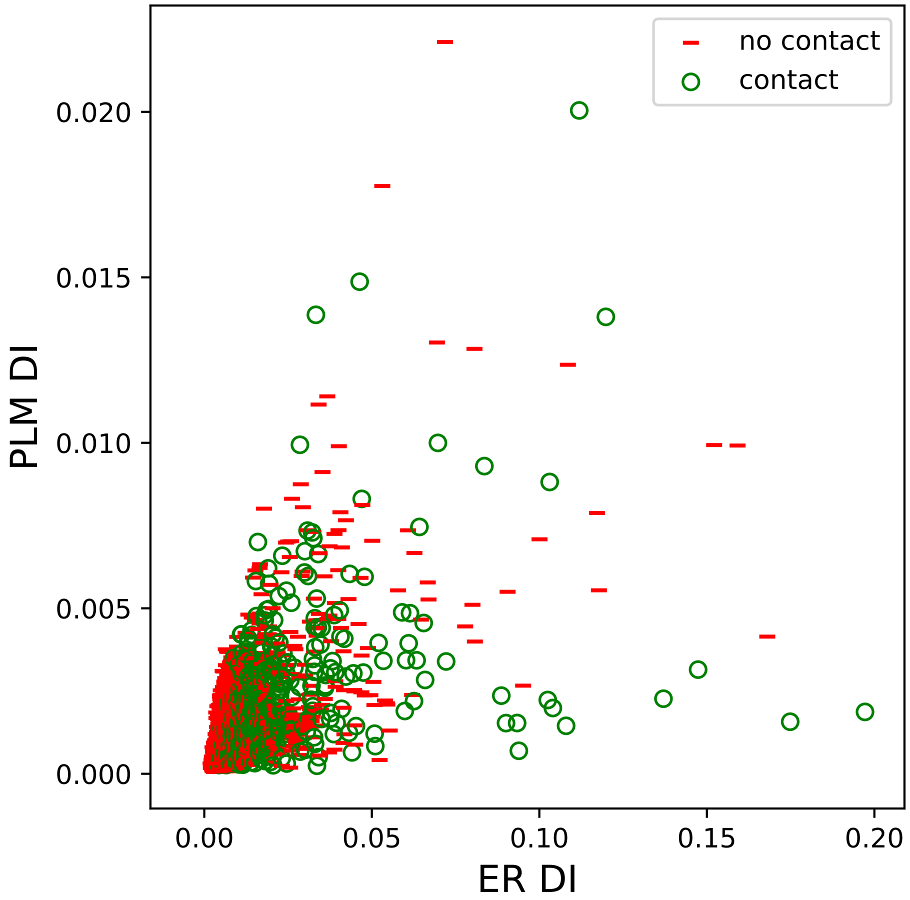

no contact
contact
0.020
0.015
0.010
0.005
0.000
PLM DI
0.00
0.05
0.10
0.15
0.20
ER DI

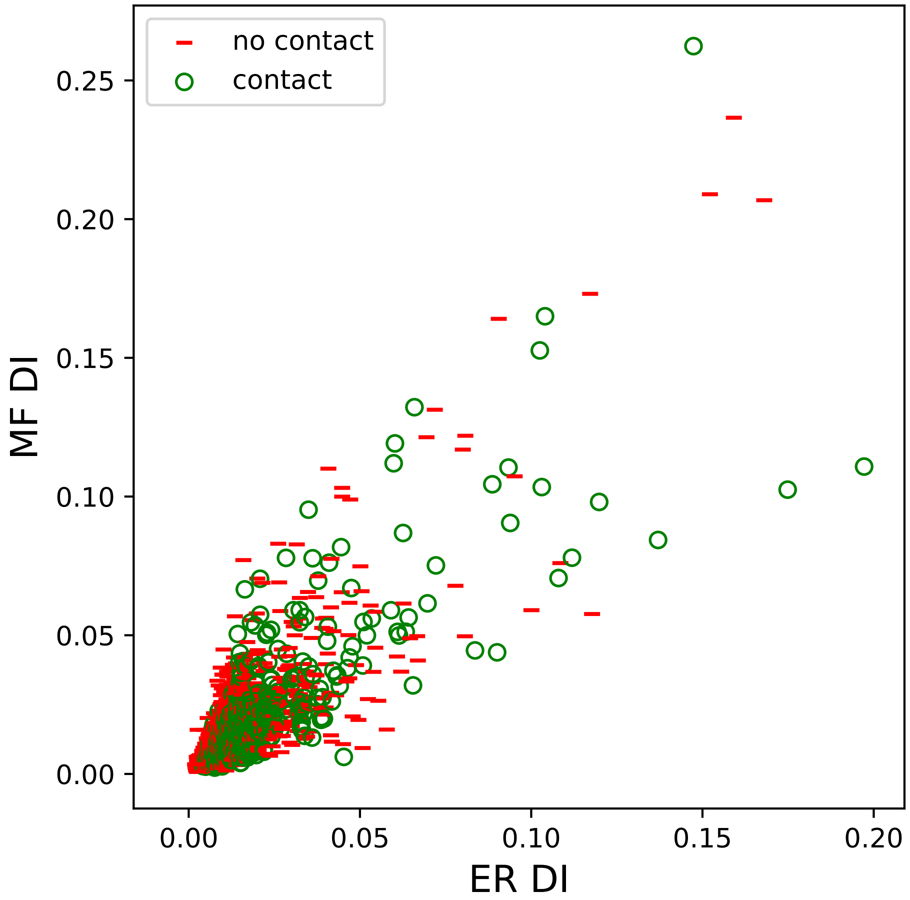

no contact
contact
0.25
0.20
0.15
0.10
0.05
0.00
MF DI
0.00
0.05
0.10
0.15
0.20
ER DI

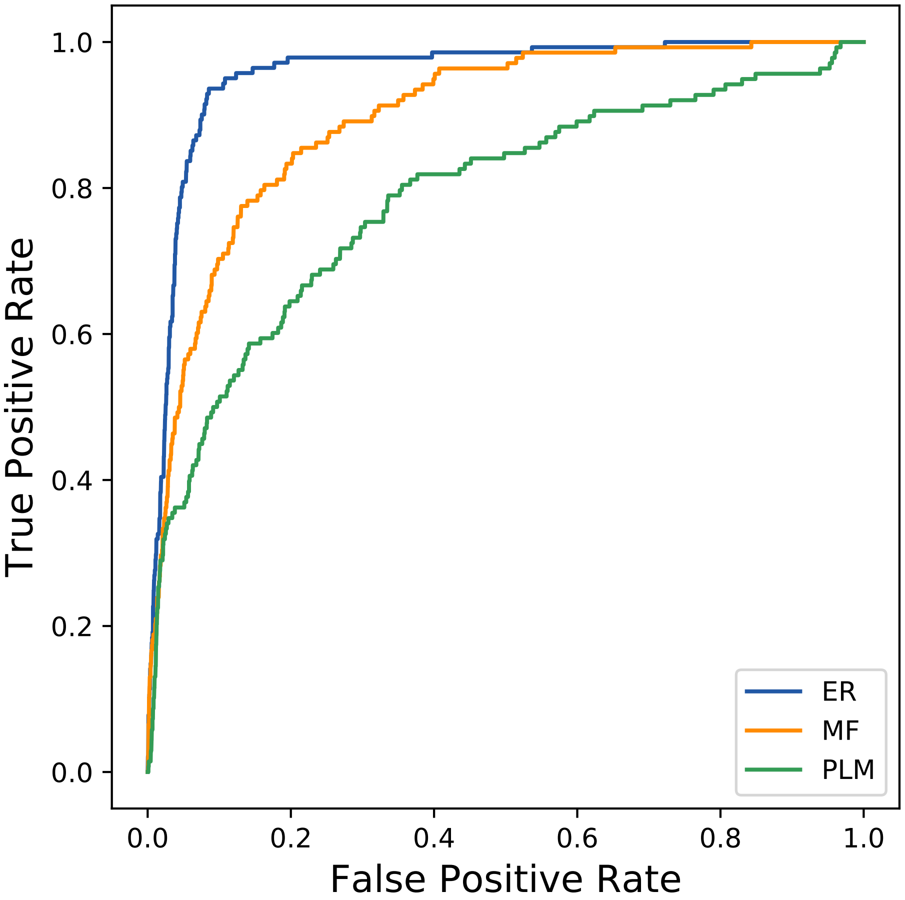

True Positive Rate
False Positive Rate
0.0
0.2
0.4
0.6
0.8
1.0
ER
MF
PLM
