## Supplemental Contact Maps for "Protein Structure Prediction with Expectation Reflection": 34481seq_143col_2a78_PF00071_plots.pdf

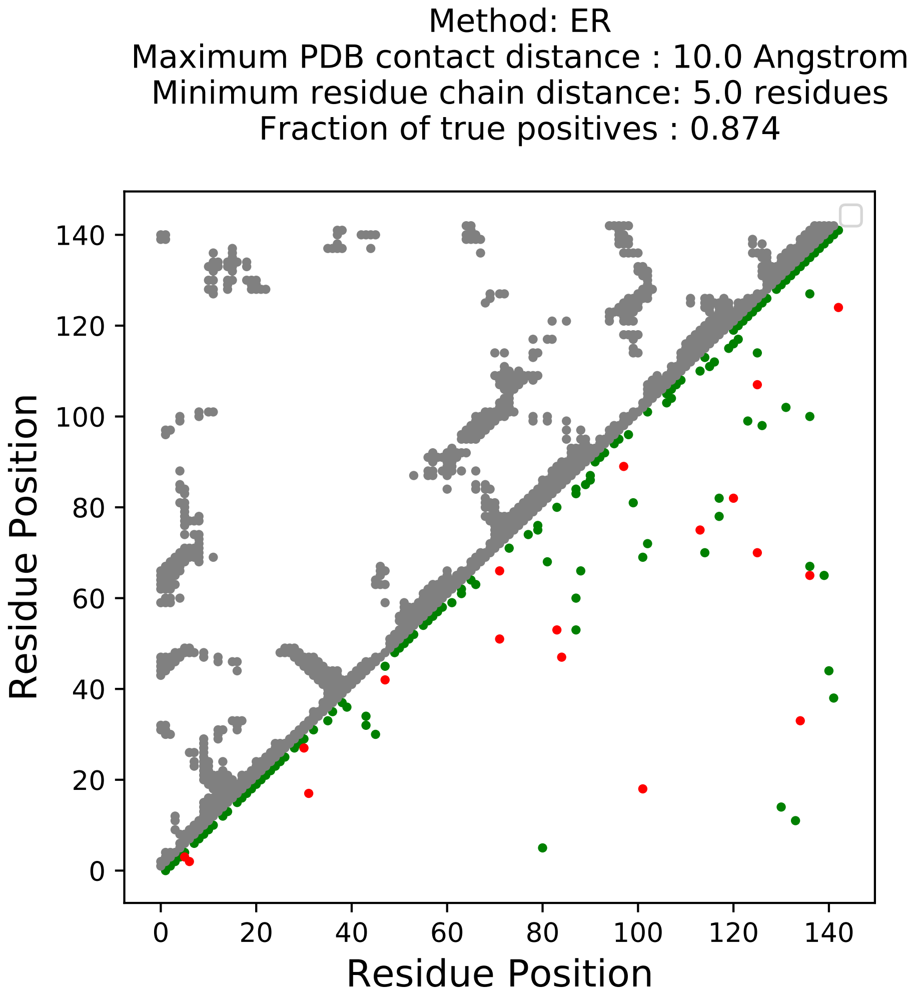
Method: ER
Maximum PDB contact distance : 10.0 Angstrom
Minimum residue chain distance: 5.0 residues
Fraction of true positives : 0.874
Residue Position
Residue Position
0
20
40
60
80
100
120
140
0
20
40
60
80
100
120
140

Method: MF
Maximum PDB contact distance : 10.0 Angstrom
Minimum residue chain distance: 5.0 residues
Fraction of true positives : 0.811

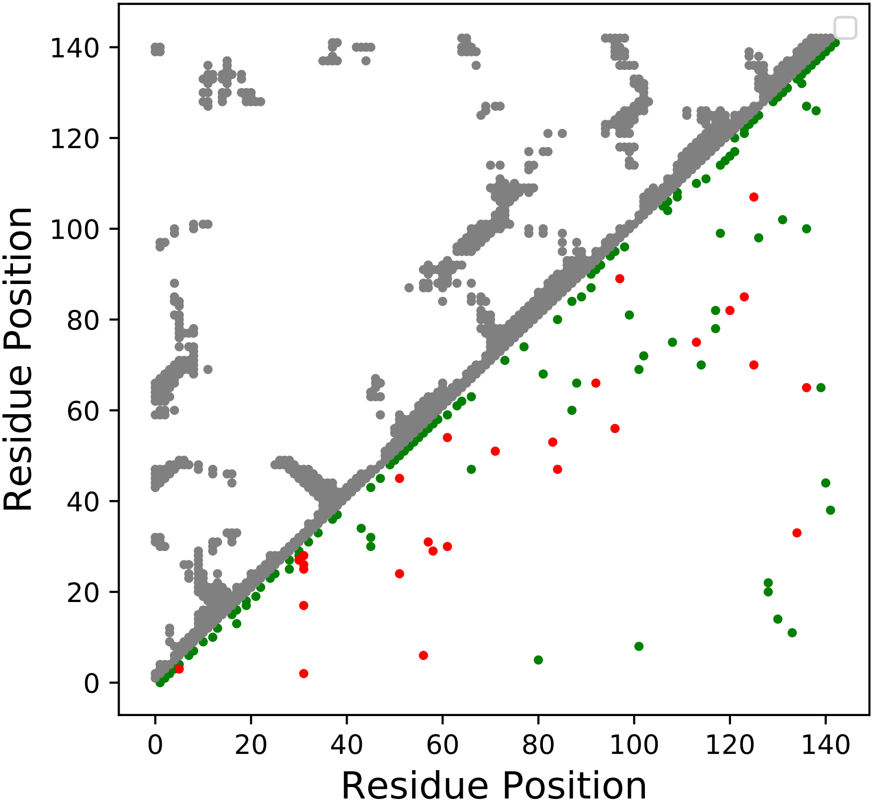

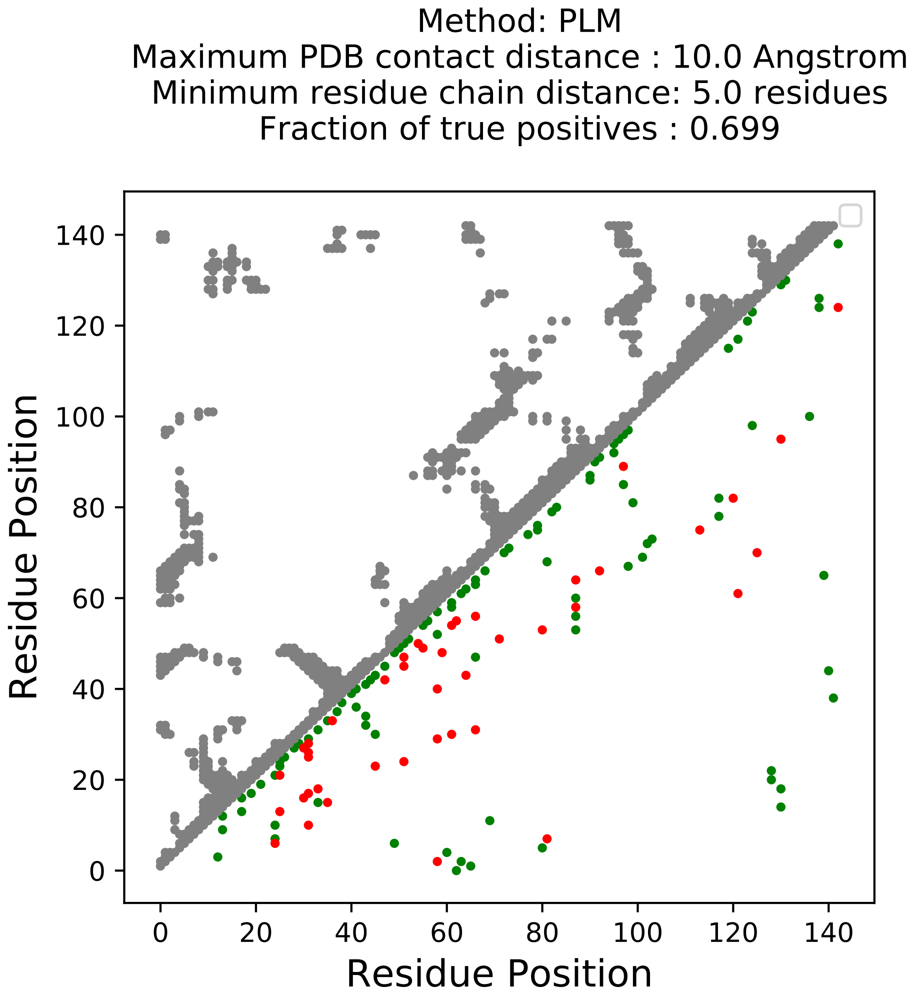

Method: PLM
Maximum PDB contact distance : 10.0 Angstrom
Minimum residue chain distance: 5.0 residues
Fraction of true positives : 0.699
Residue Position
Residue Position
0
20
40
60
80
100
120
140

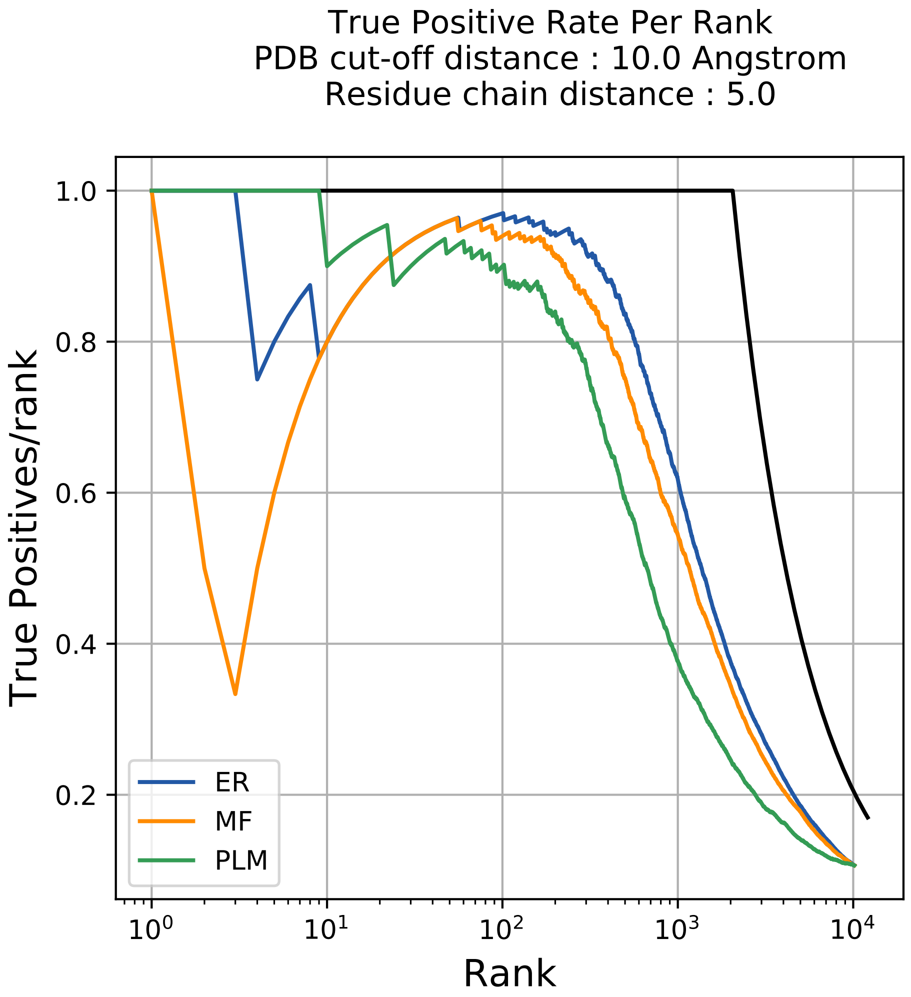
True Positive Rate Per Rank
PDB cut-off distance : 10.0 Angstrom
Residue chain distance : 5.0
True Positives/rank
Rank
1.0
0.8
0.6
0.4
0.2
$10^0$
$10^1$
$10^2$
$10^3$
$10^4$
ER
MF
PLM

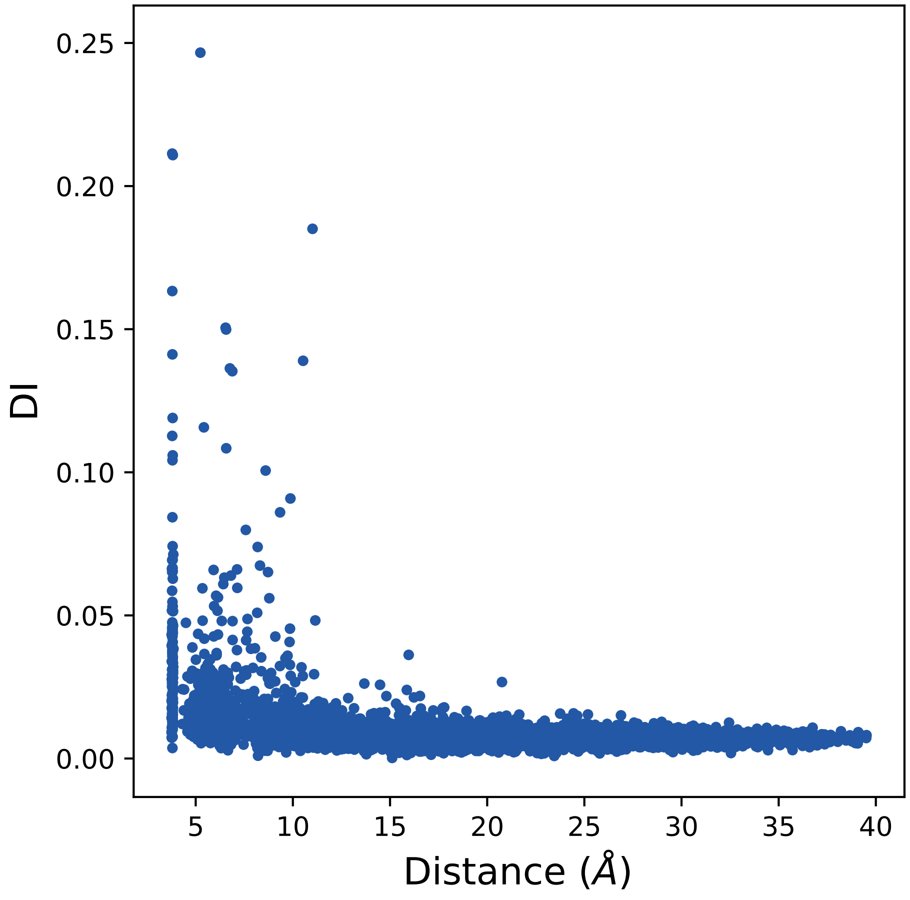

DI
0.25
0.20
0.15
0.10
0.05
0.00
5
10
15
20
25
30
35
40
Distance (Å)

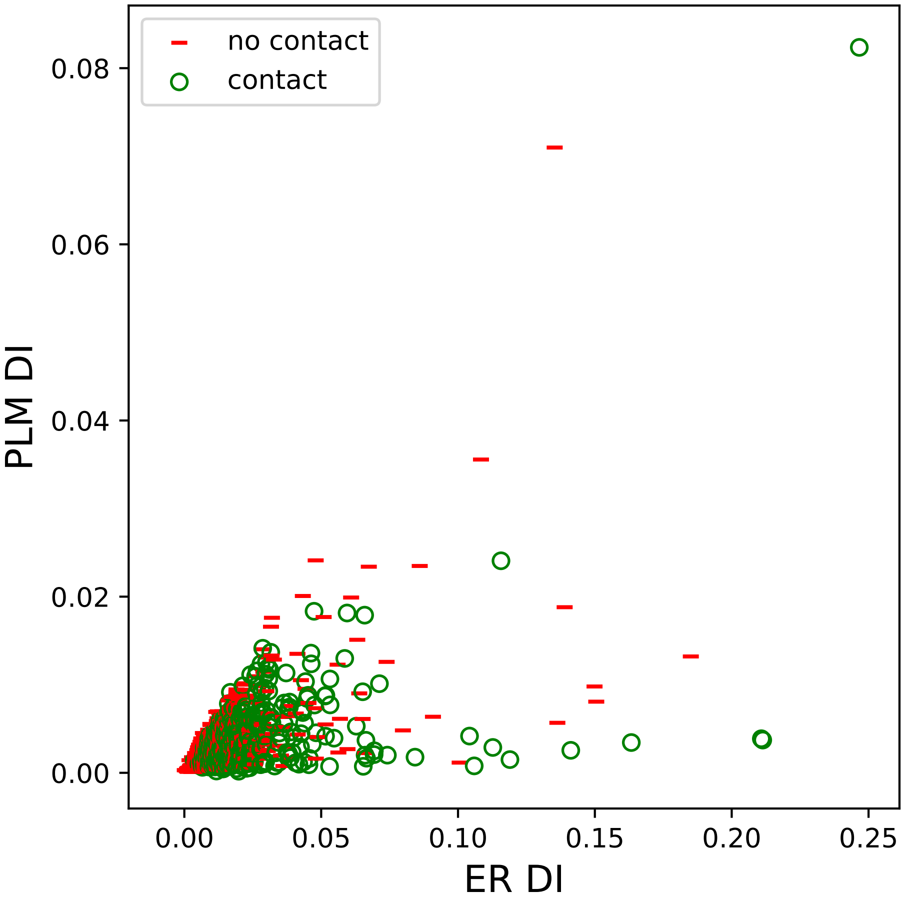

no contact
contact
PLM DI
0.08
0.06
0.04
0.02
0.00
ER DI
0.00
0.05
0.10
0.15
0.20
0.25

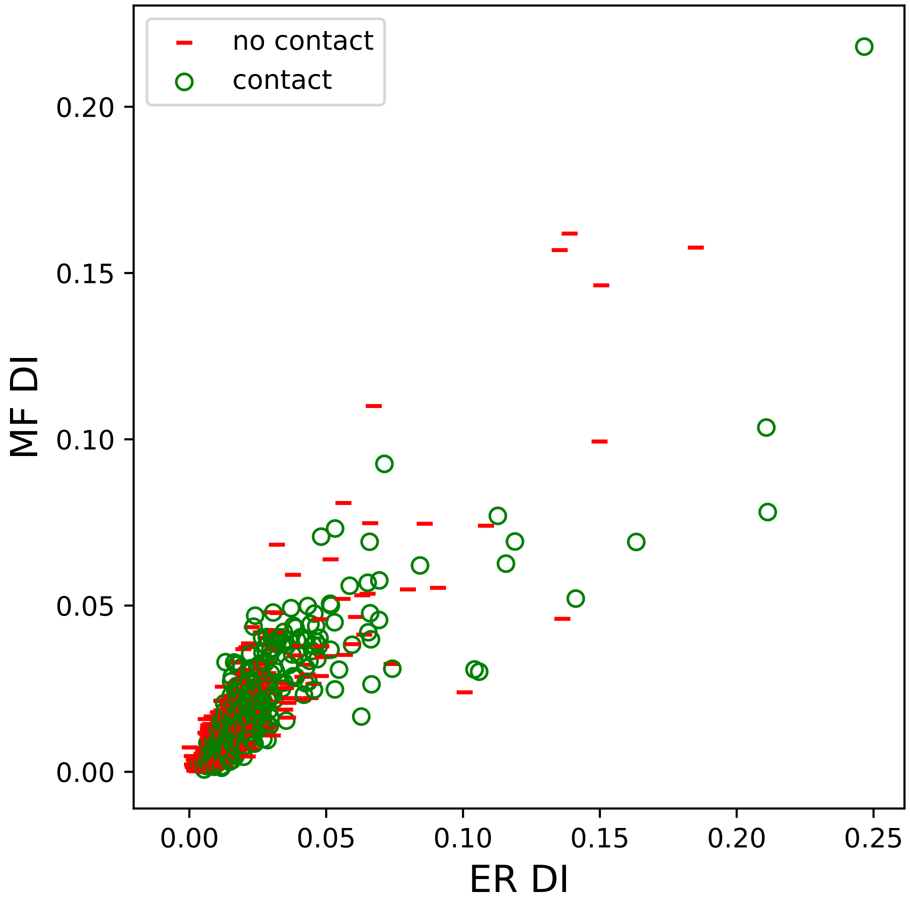

no contact
contact
MF DI
ER DI
0.00
0.05
0.10
0.15
0.20
0.25

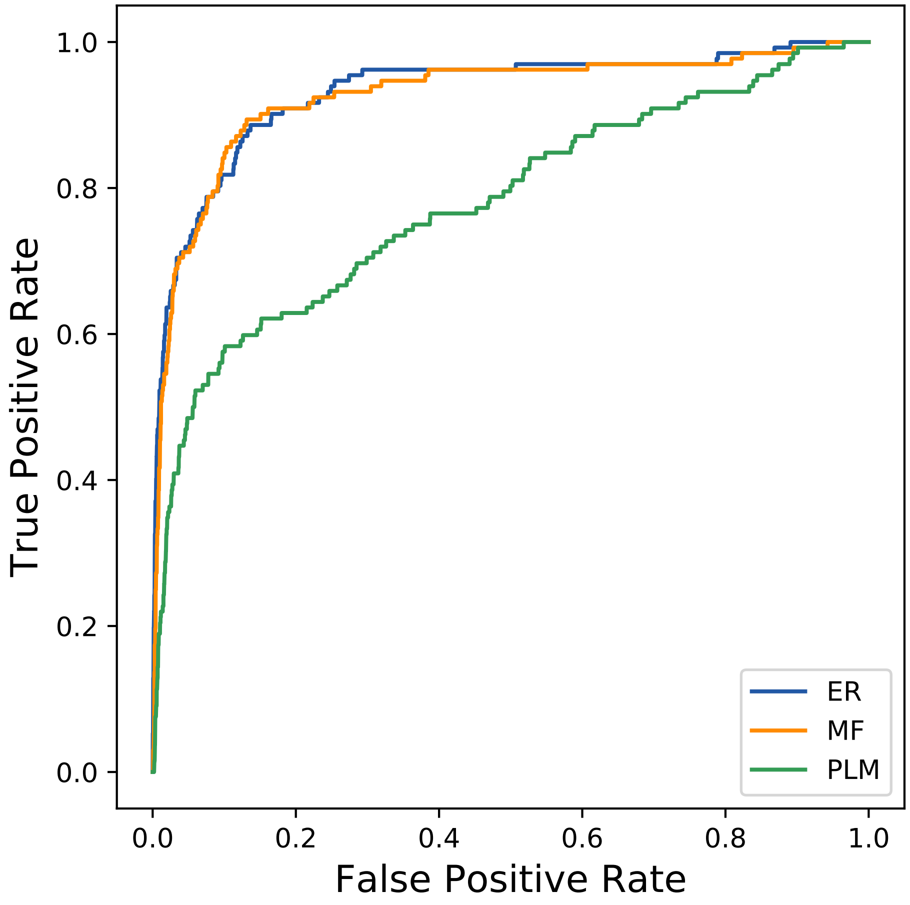
True Positive Rate
False Positive Rate
0.0
0.2
0.4
0.6
0.8
1.0
ER
MF
PLM
