## Supplemental Contact Maps for "Protein Structure Prediction with Expectation Reflection": 44031seq_117col_1jhz_PF13377_plots.pdf

Method: ER
Maximum PDB contact distance : 10.0 Angstrom
Minimum residue chain distance: 5.0 residues
Fraction of true positives : 0.718

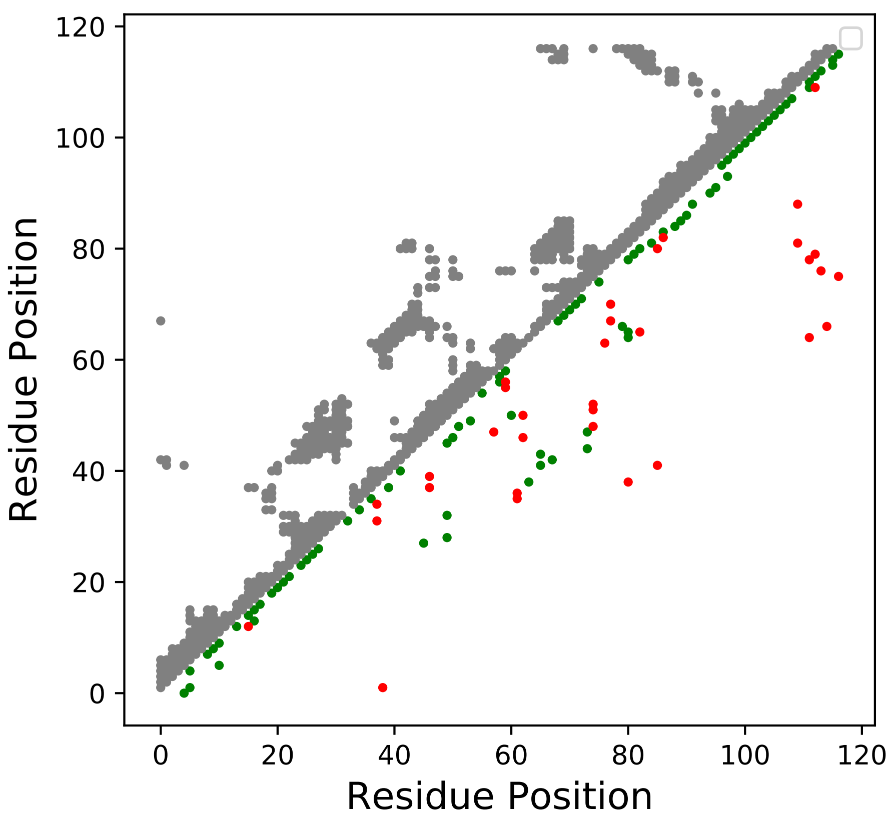

Method: MF
Maximum PDB contact distance : 10.0 Angstrom
Minimum residue chain distance: 5.0 residues
Fraction of true positives : 0.684

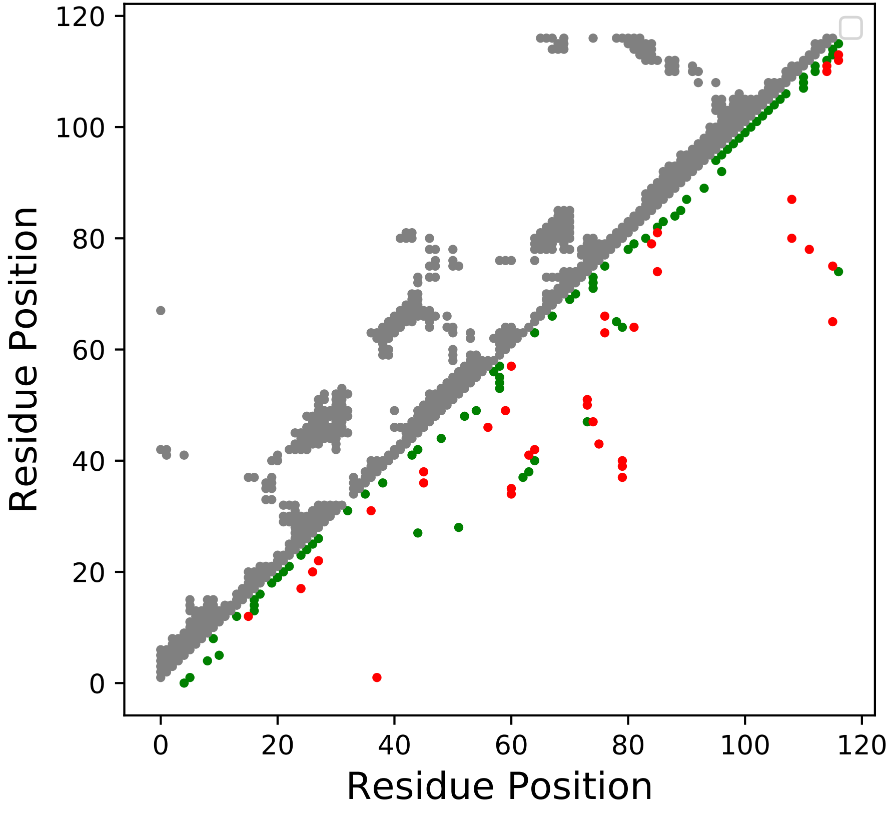

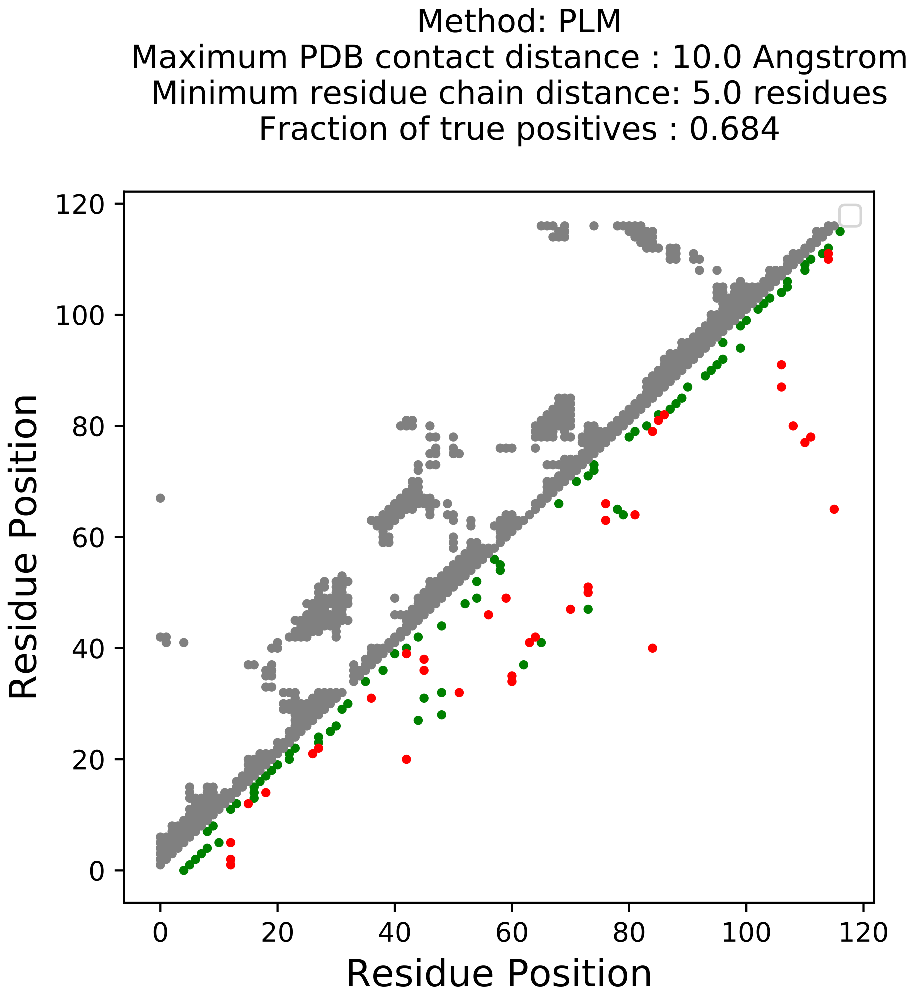
Method: PLM
Maximum PDB contact distance : 10.0 Angstrom
Minimum residue chain distance: 5.0 residues
Fraction of true positives : 0.684
Residue Position
Residue Position
0
20
40
60
80
100
120

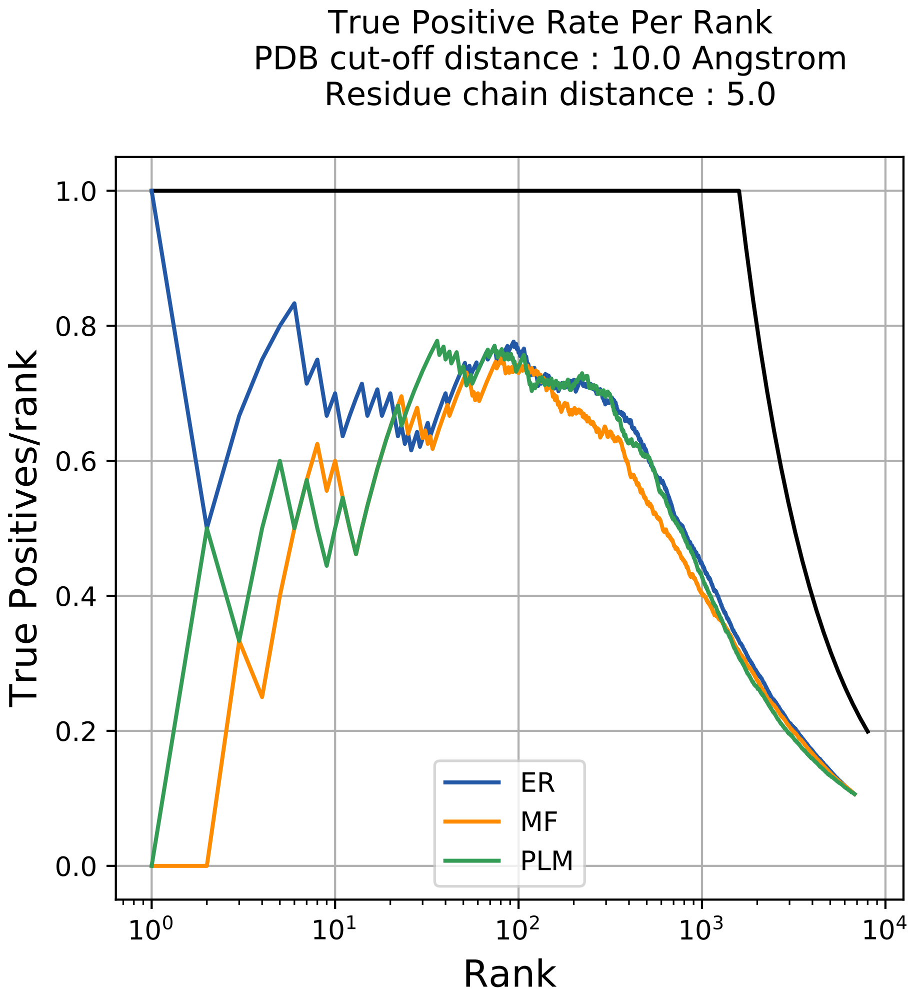

True Positive Rate Per Rank
PDB cut-off distance : 10.0 Angstrom
Residue chain distance : 5.0
True Positives/rank
Rank
1.0
0.8
0.6
0.4
0.2
0.0
$10^0$
$10^1$
$10^2$
$10^3$
$10^4$
ER
MF
PLM

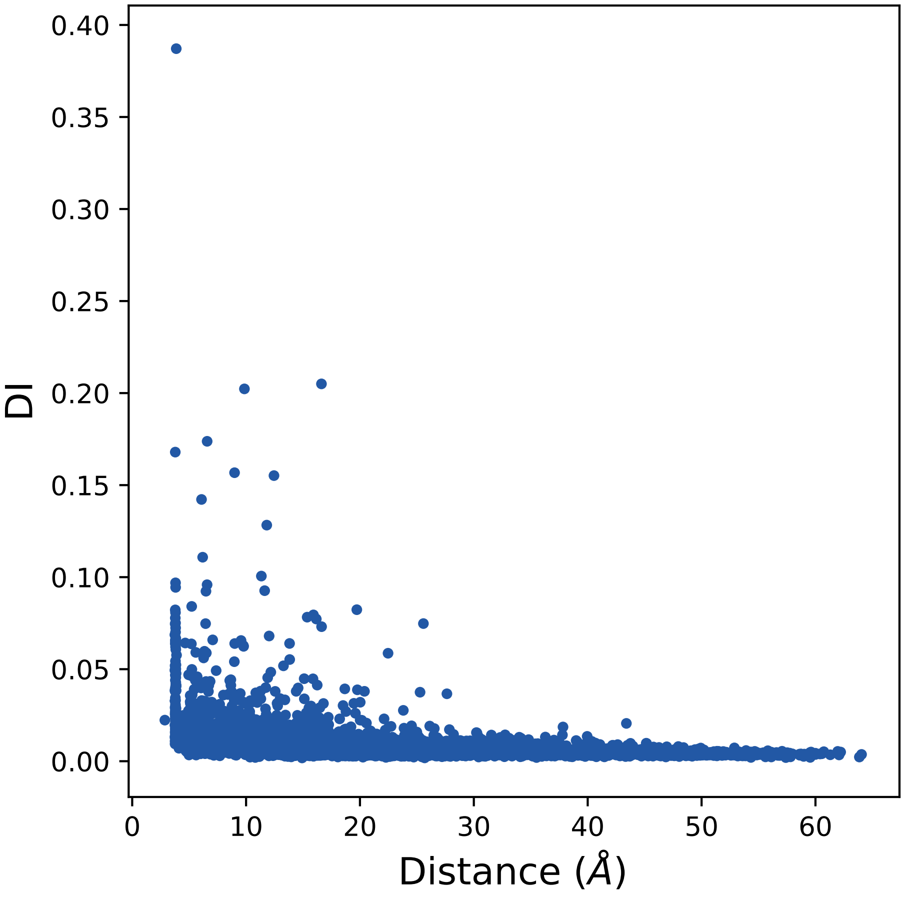

0.40
0.35
0.30
0.25
0.20
0.15
0.10
0.05
0.00
DI
0
10
20
30
40
50
60
Distance (Å)

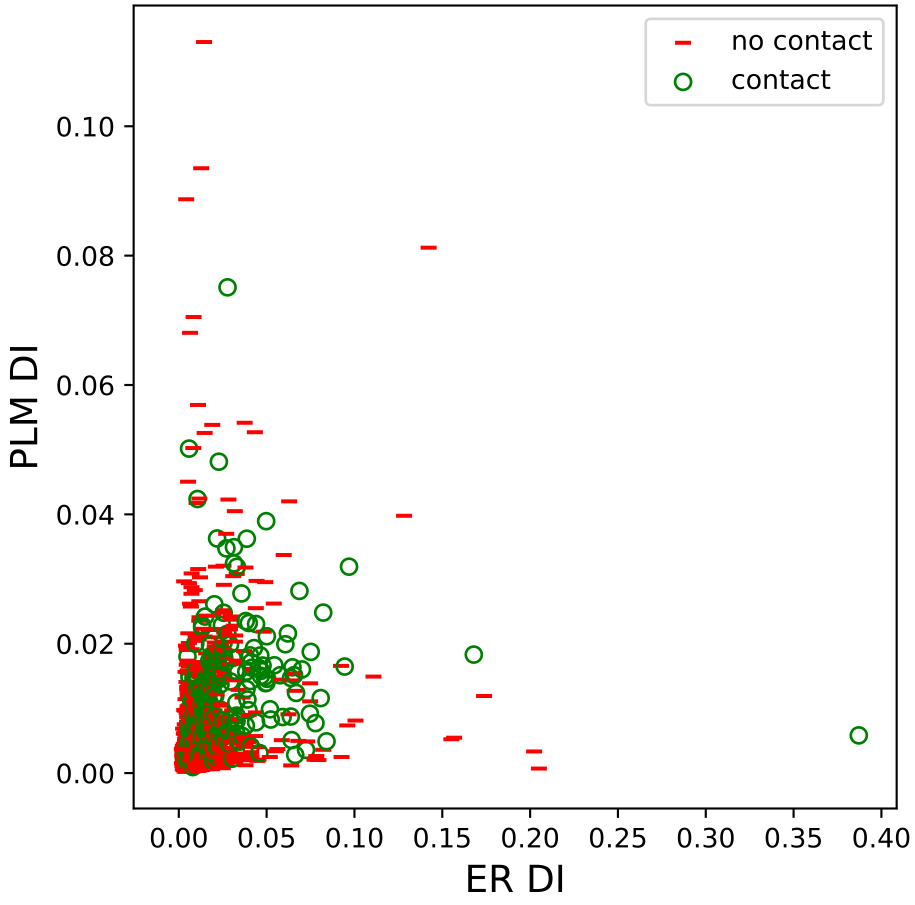
no contact
contact
PLM DI
ER DI
0.00
0.02
0.04
0.06
0.08
0.10
0.00
0.05
0.10
0.15
0.20
0.25
0.30
0.35
0.40

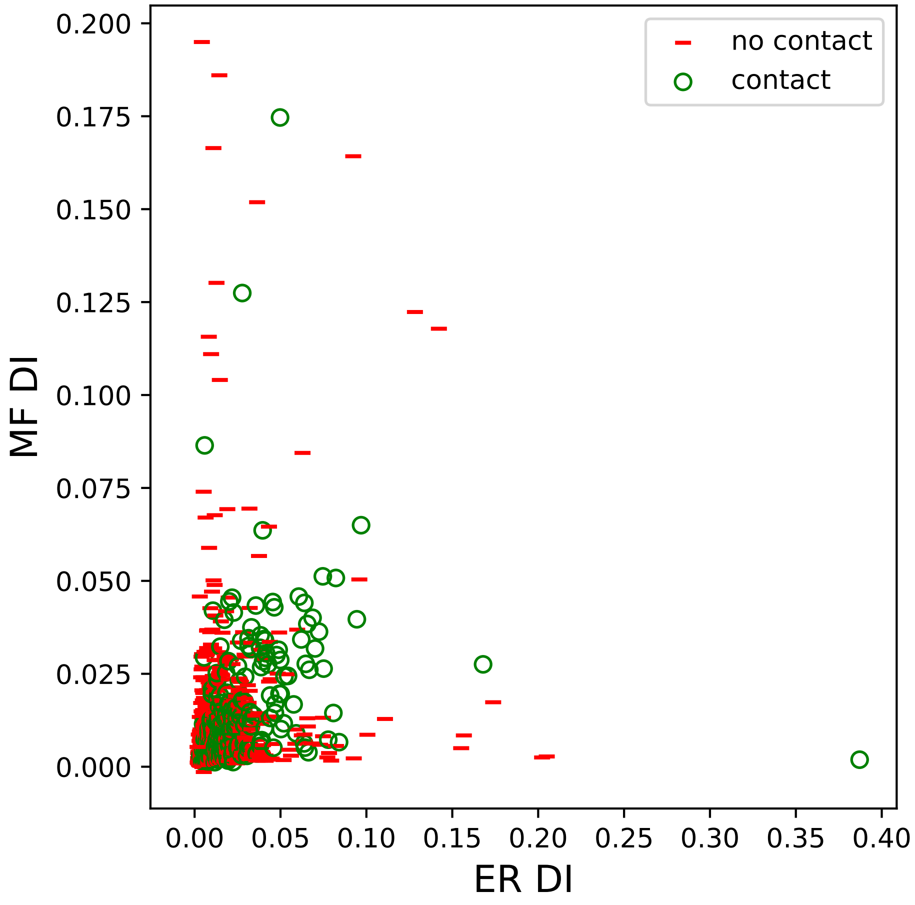

0.200
0.175
0.150
0.125
0.100
0.075
0.050
0.025
0.000
MF DI
0.00
0.05
0.10
0.15
0.20
0.25
0.30
0.35
0.40
ER DI
no contact
contact

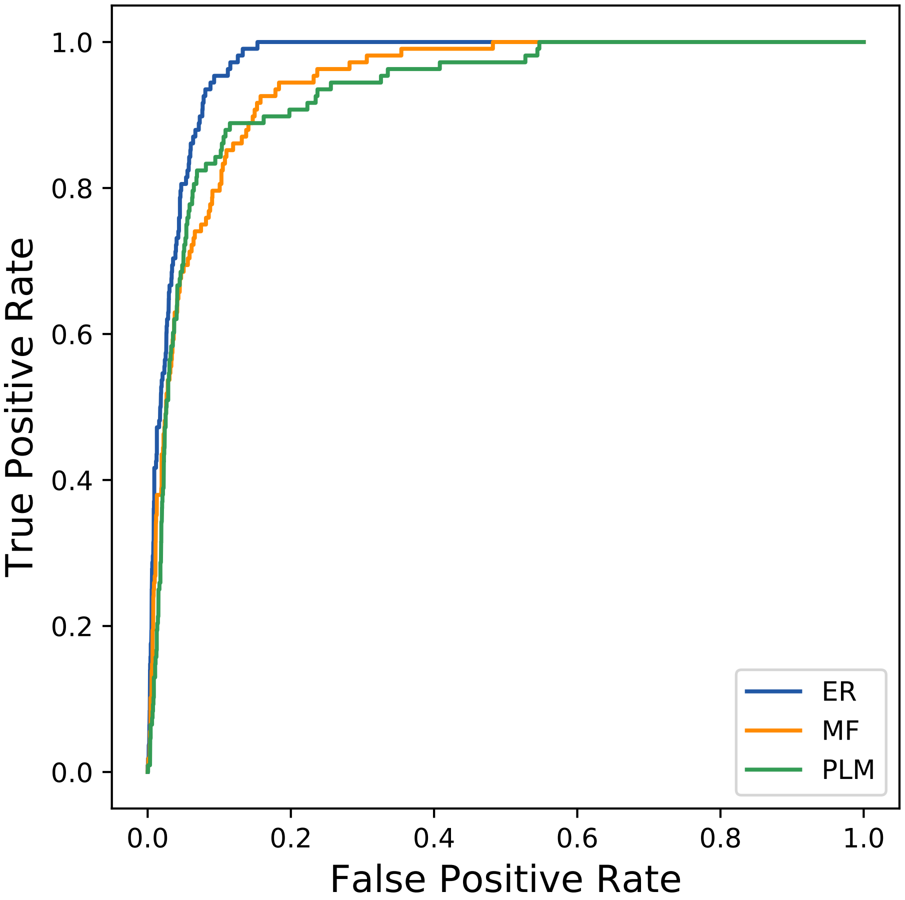

True Positive Rate
False Positive Rate
0.0
0.2
0.4
0.6
0.8
1.0
ER
MF
PLM
