## Supplementary figures and images for "Protein Structure Prediction with Expectation Reflection"

### 2seq_82col_4qpg_PF12944_plots.pdf

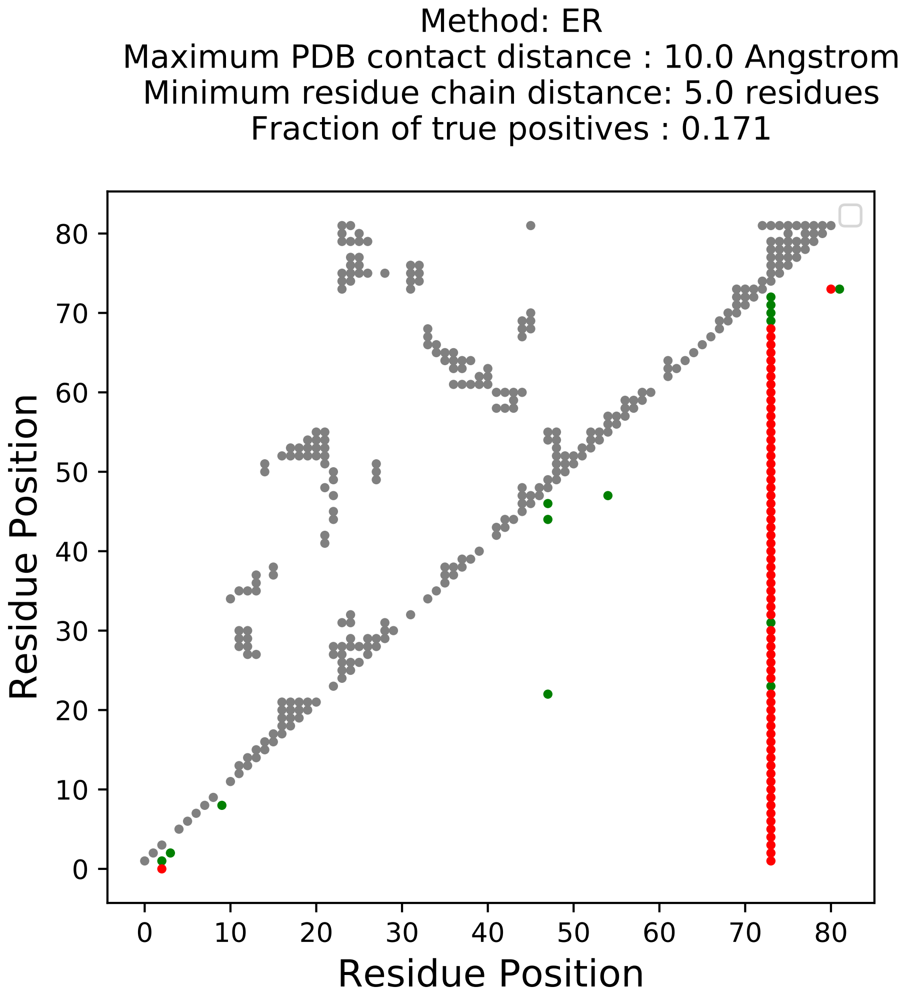

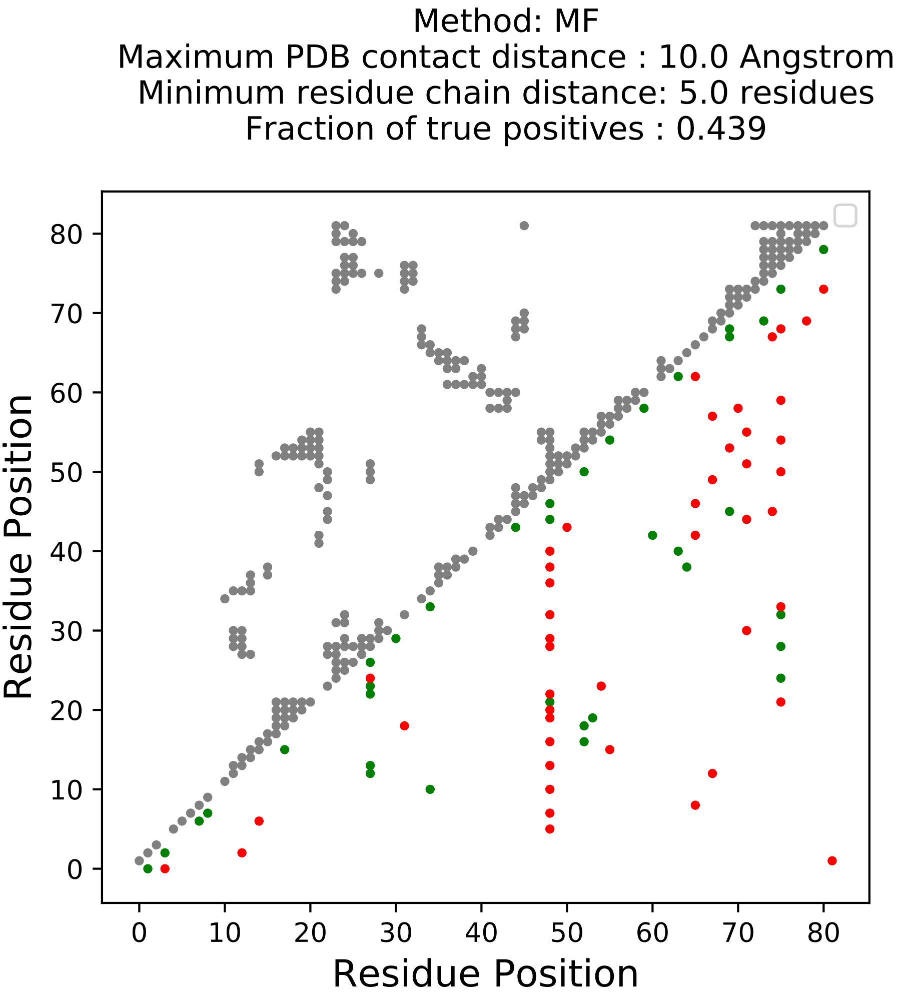

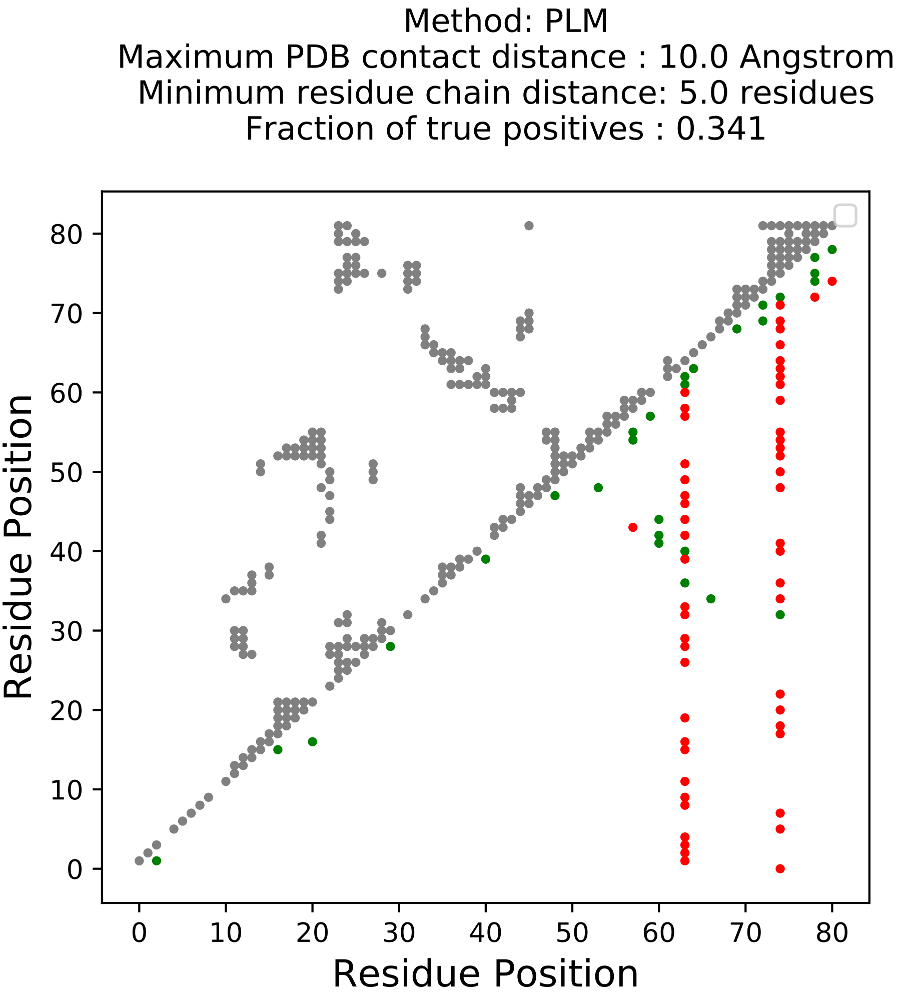

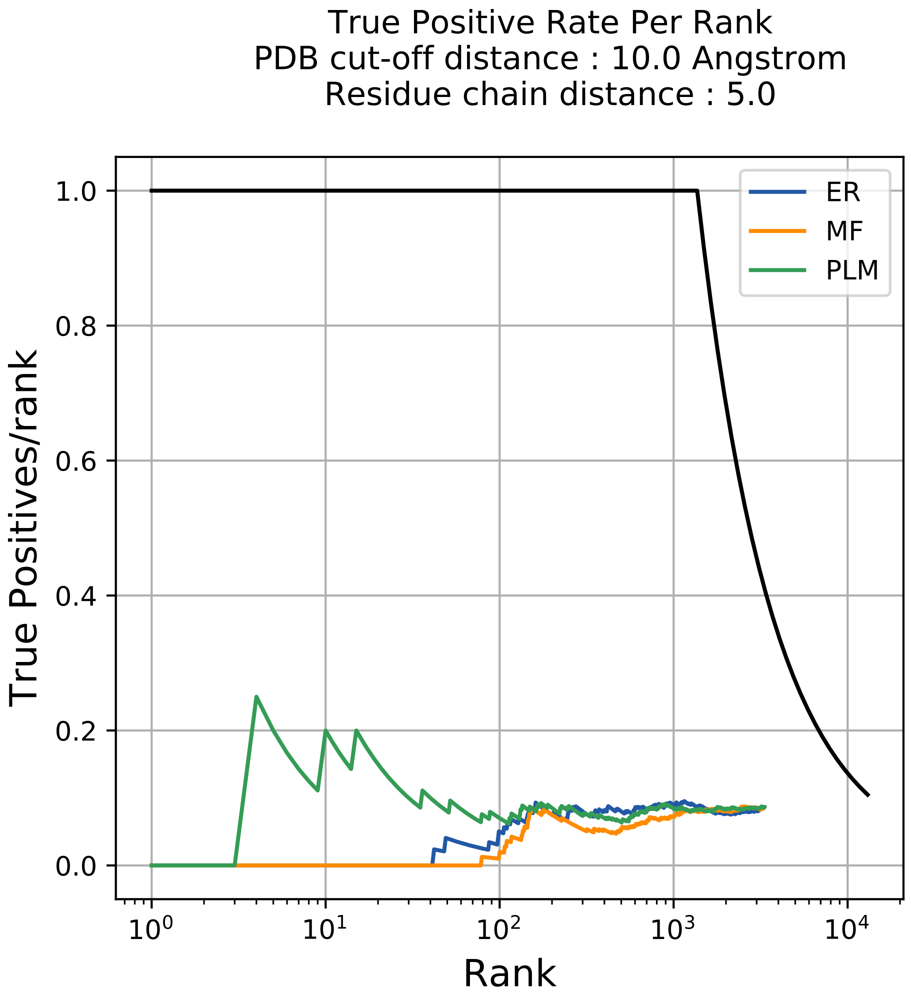

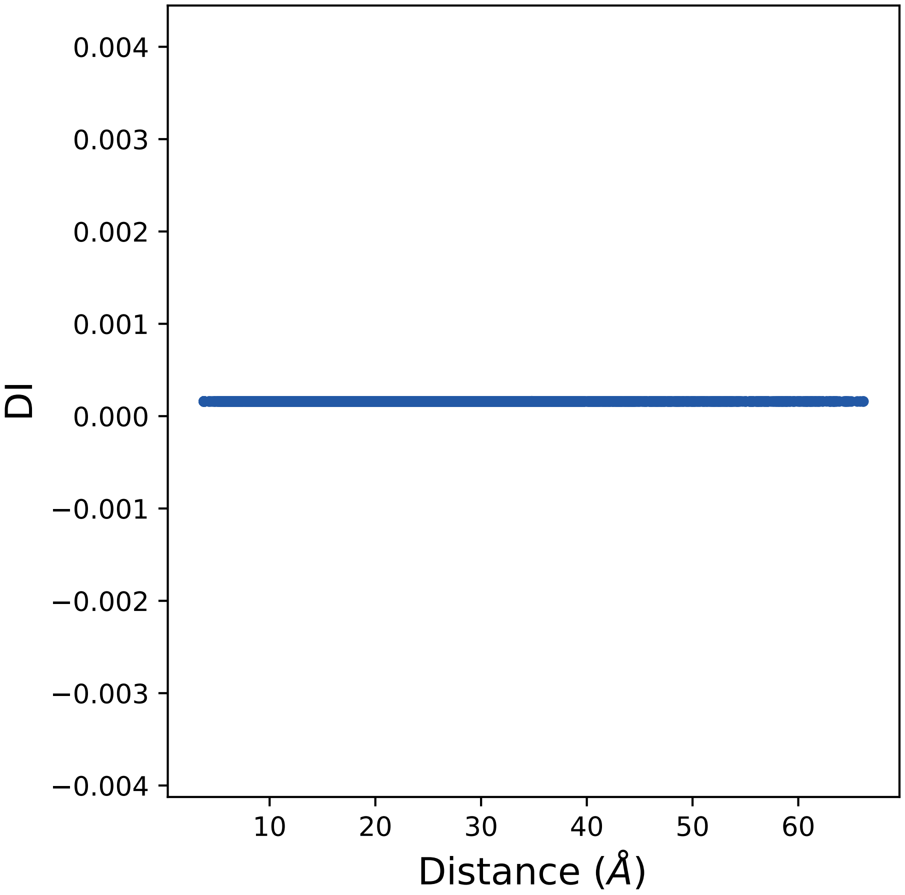

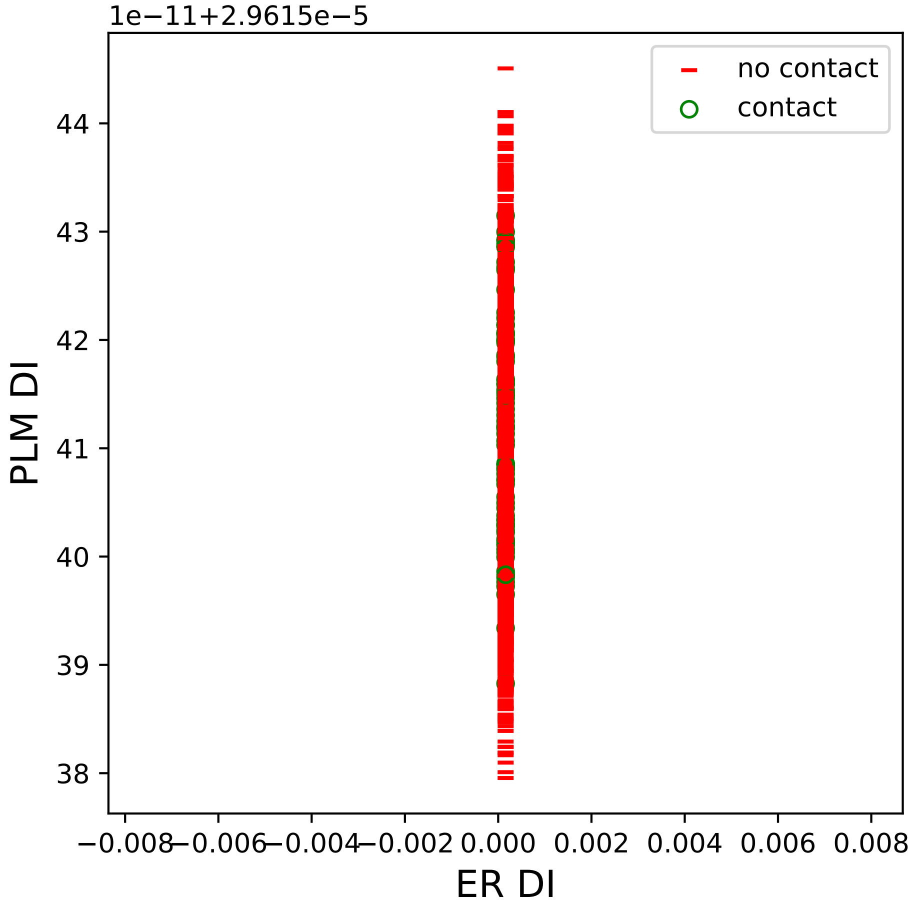

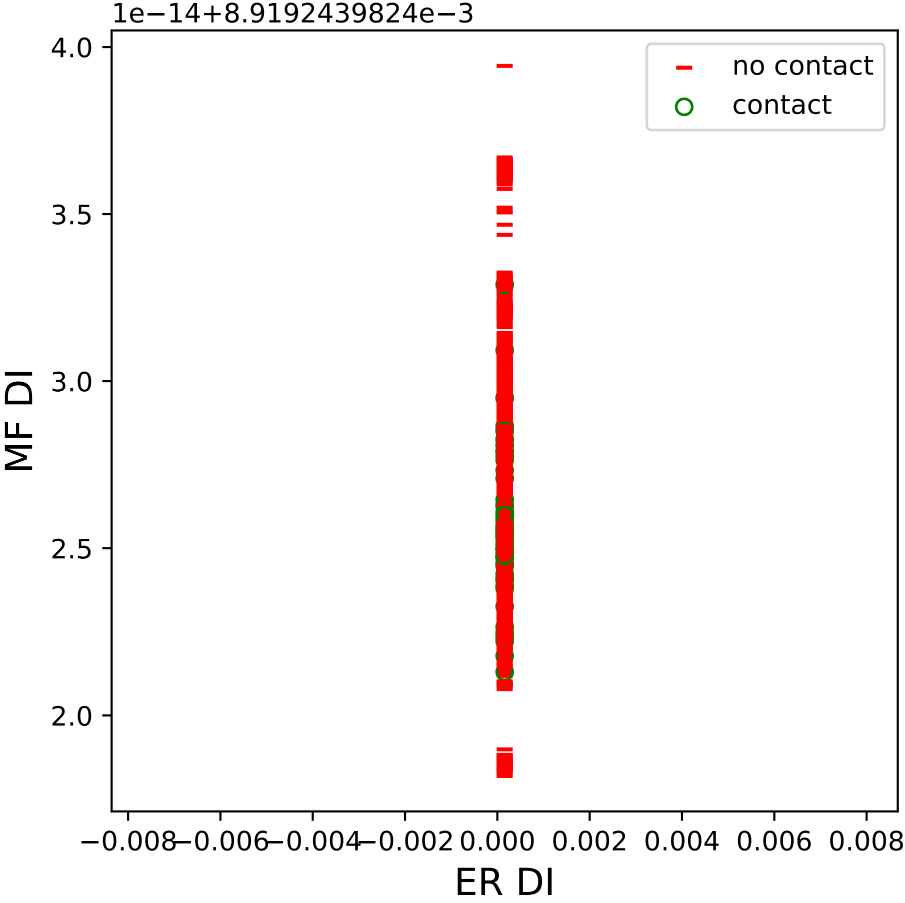

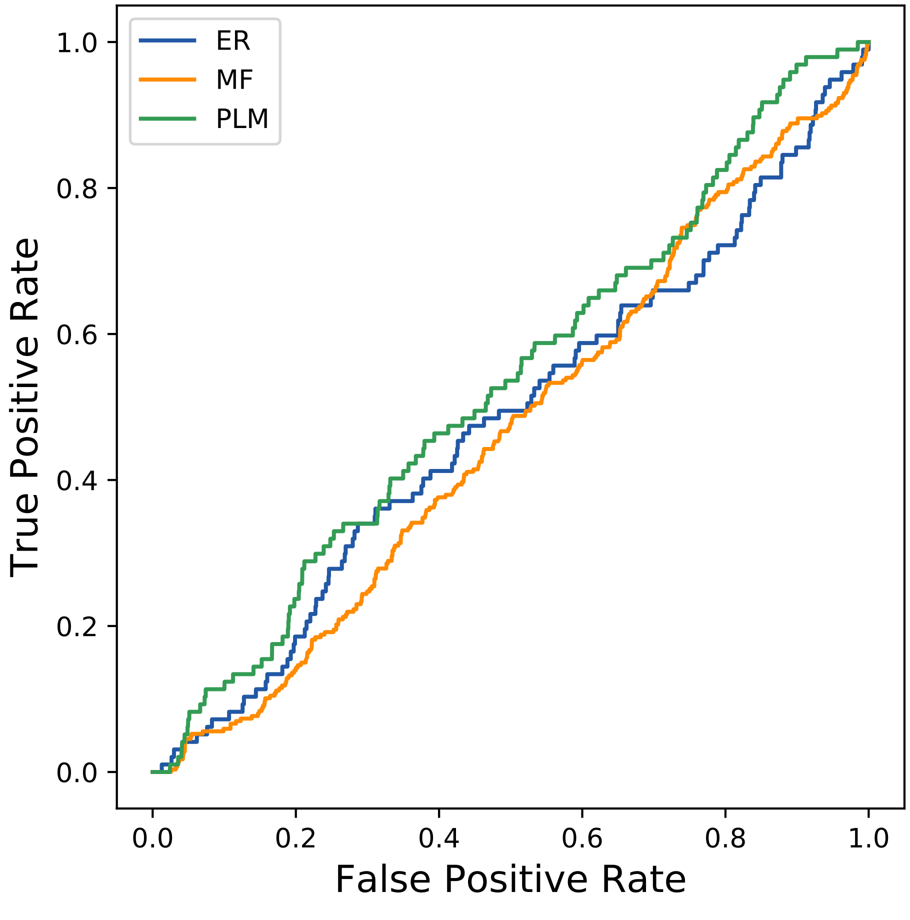

### 3seq_22col_5ii9_PF01303_plots.pdf

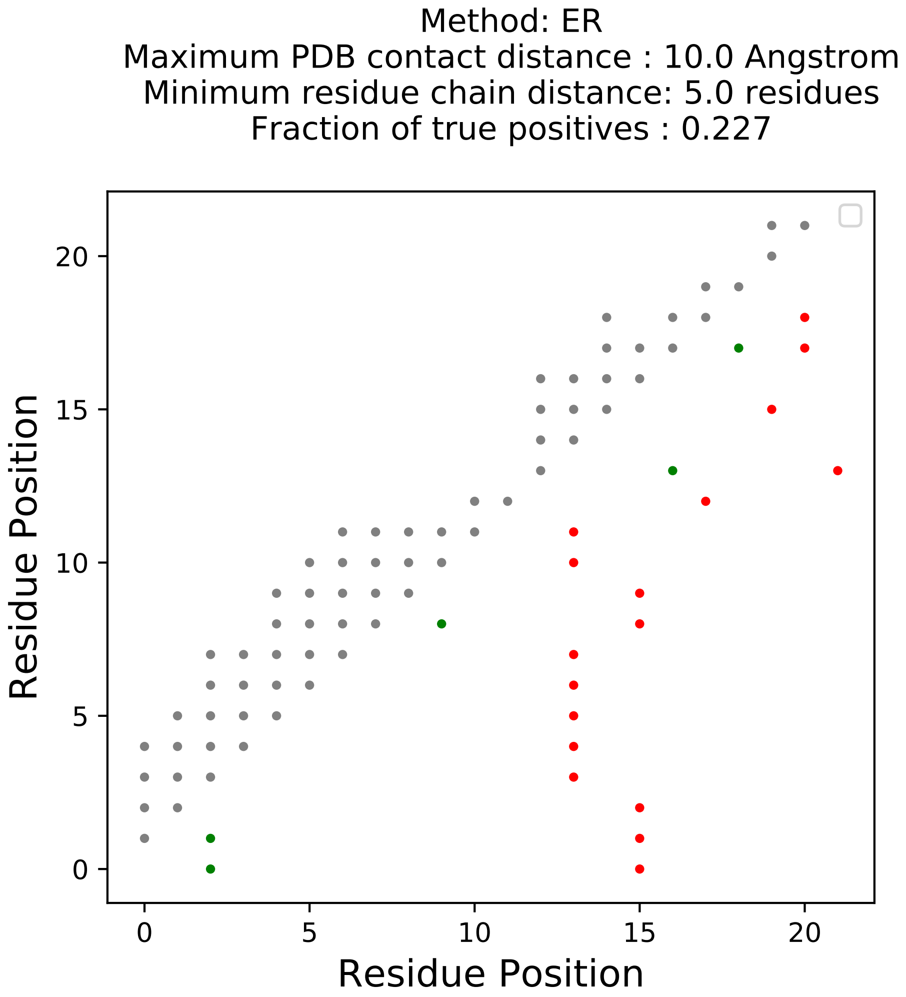

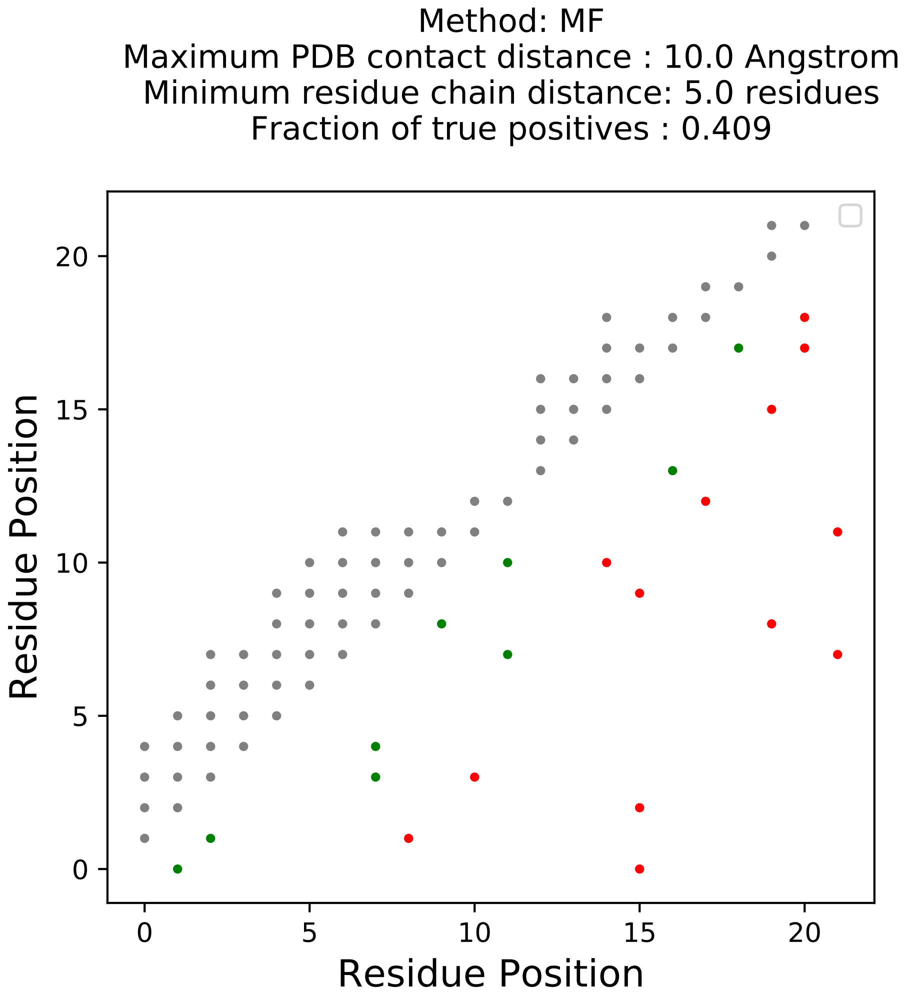

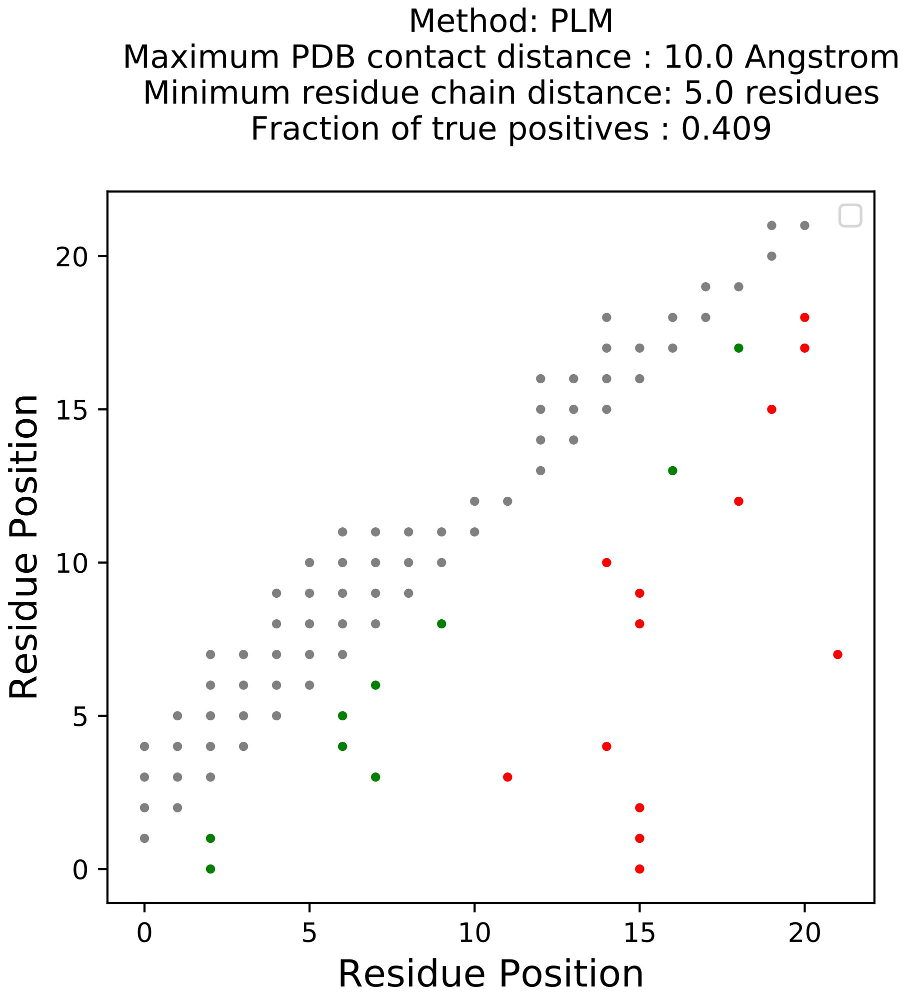

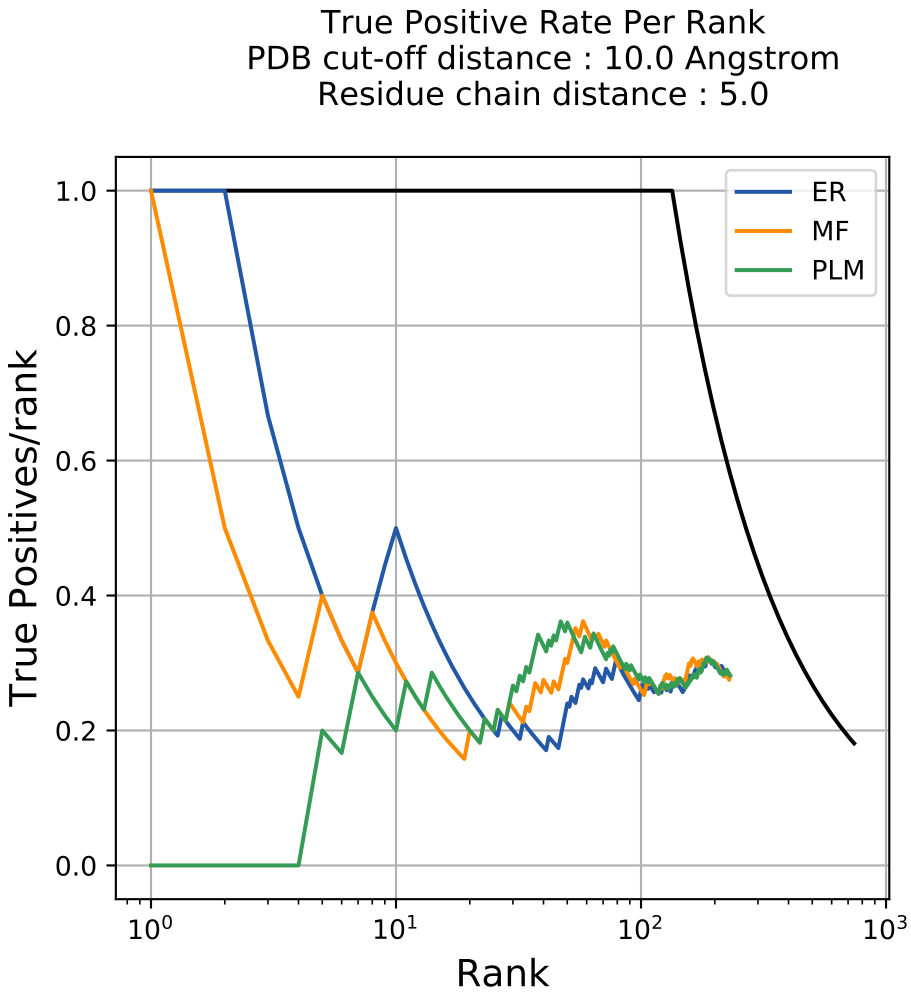

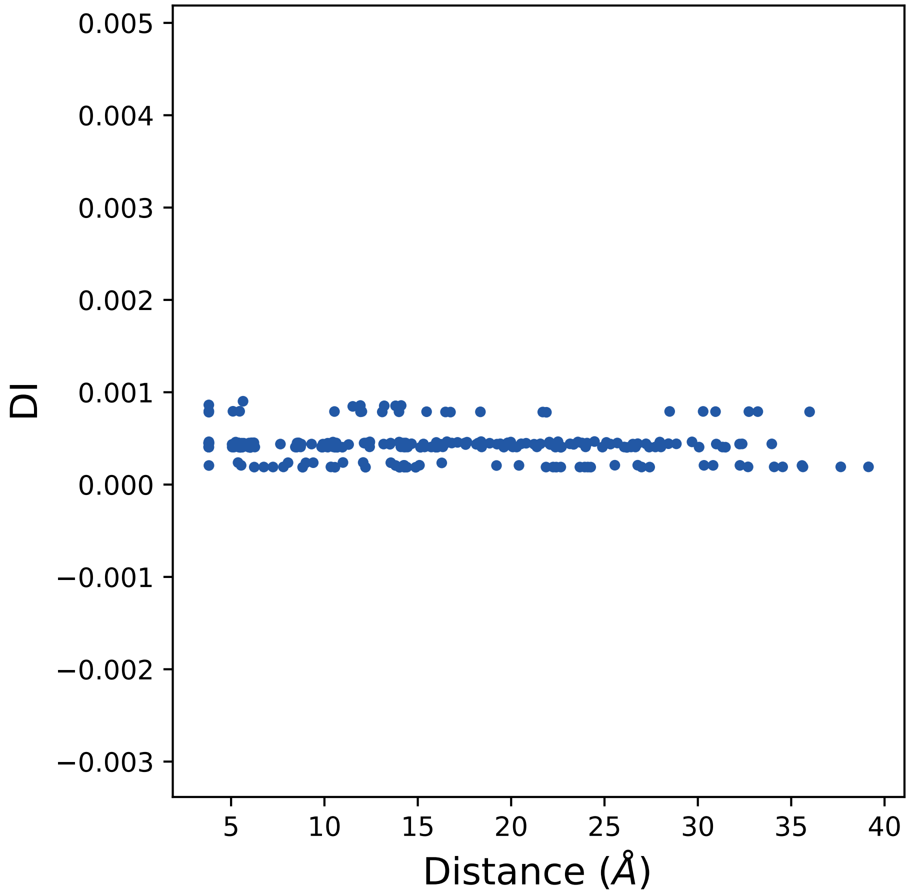

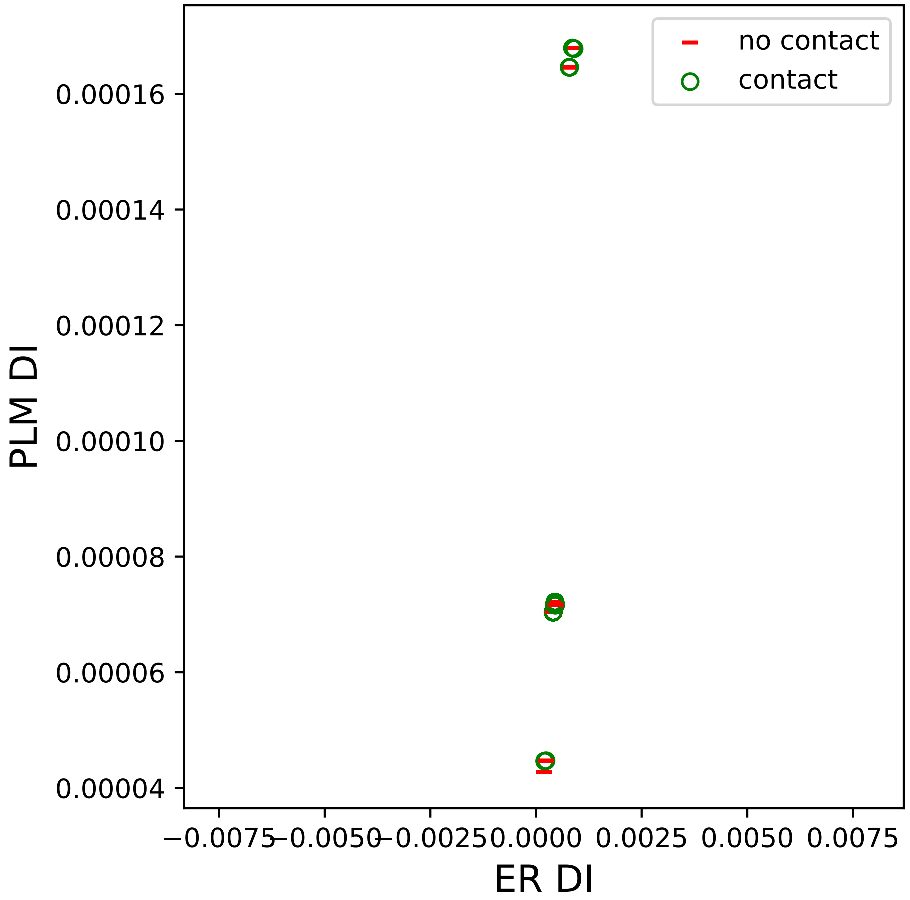

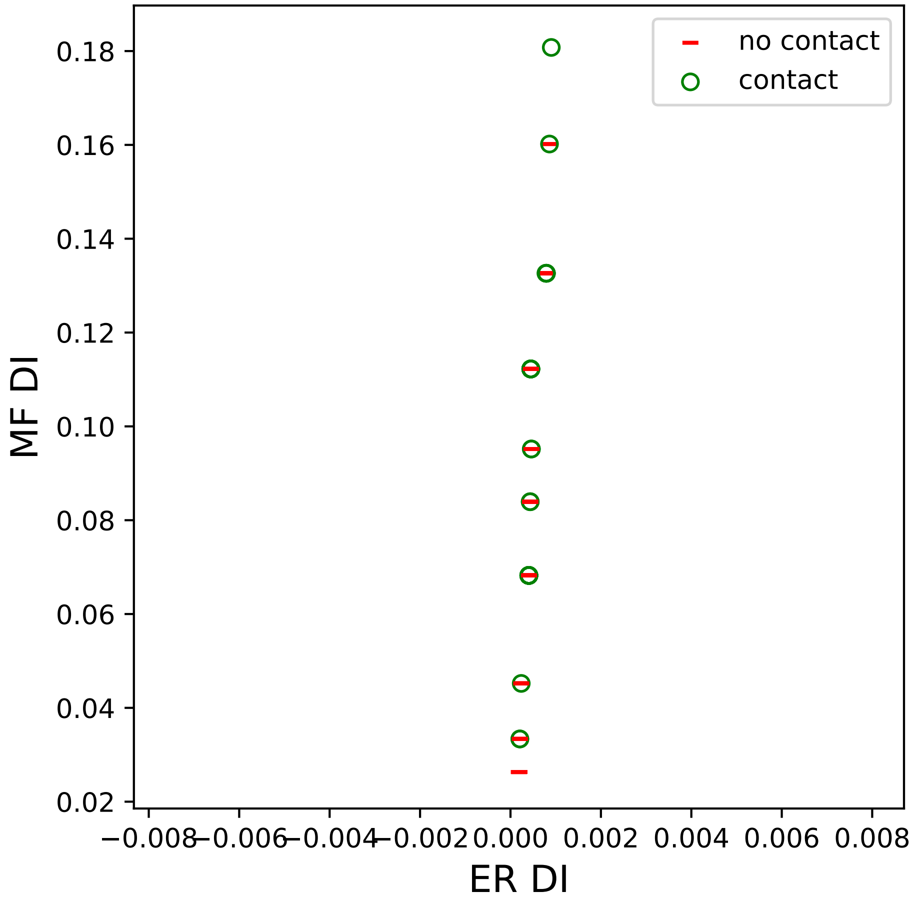

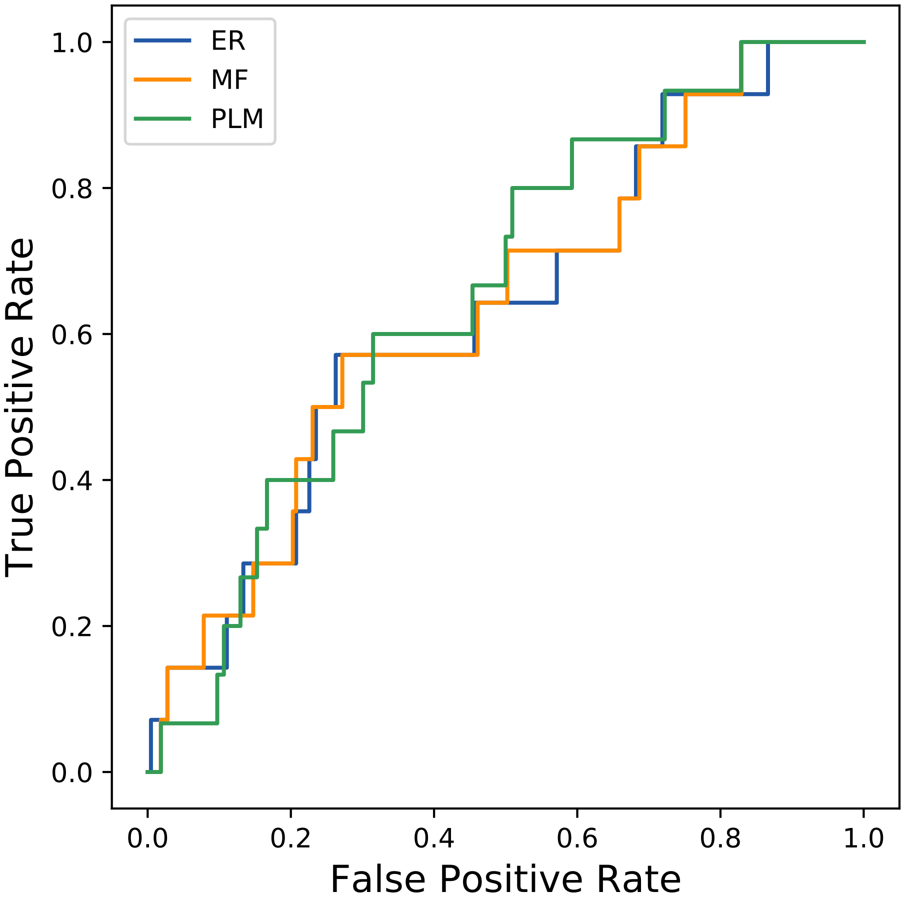

### 23seq_84col_2i88_PF01024_plots.pdf

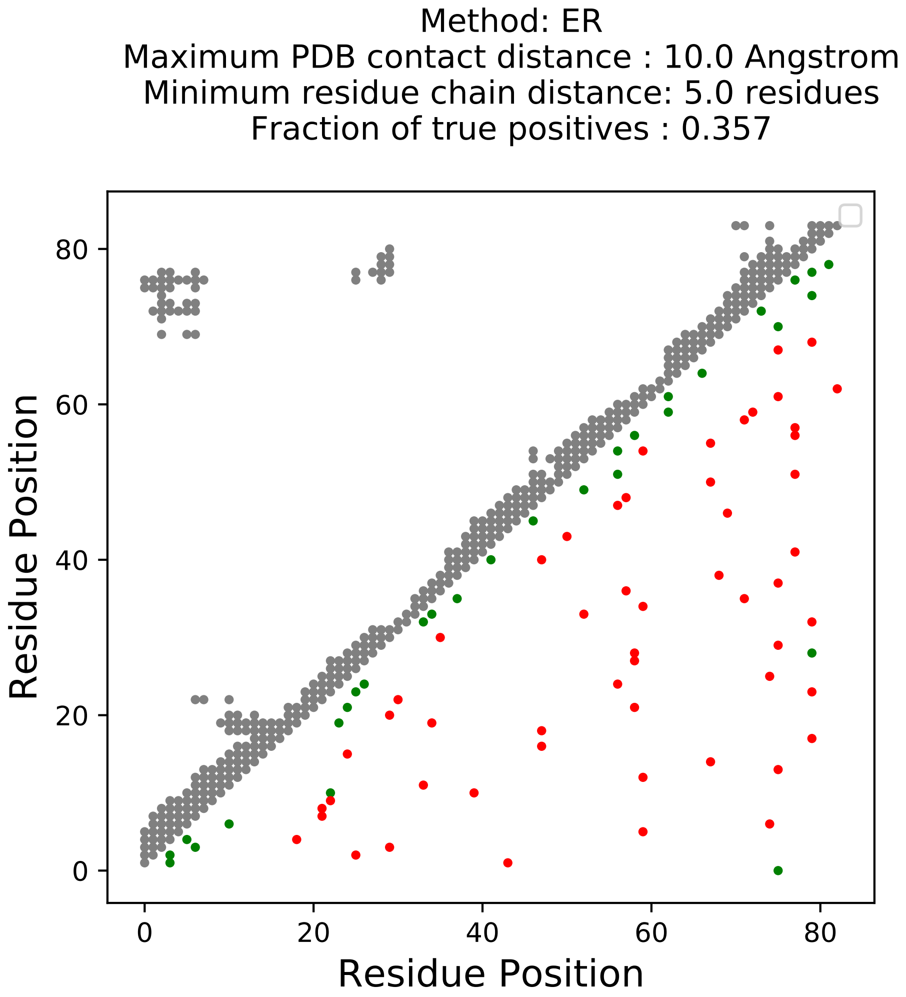

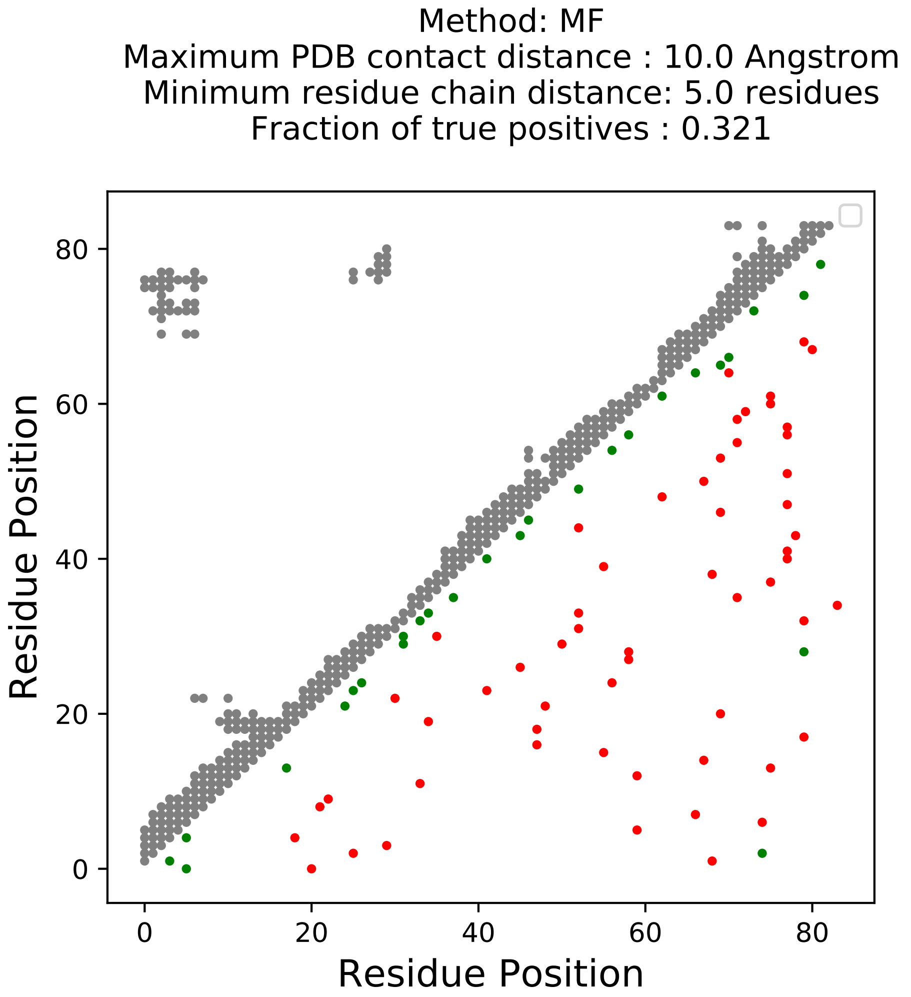

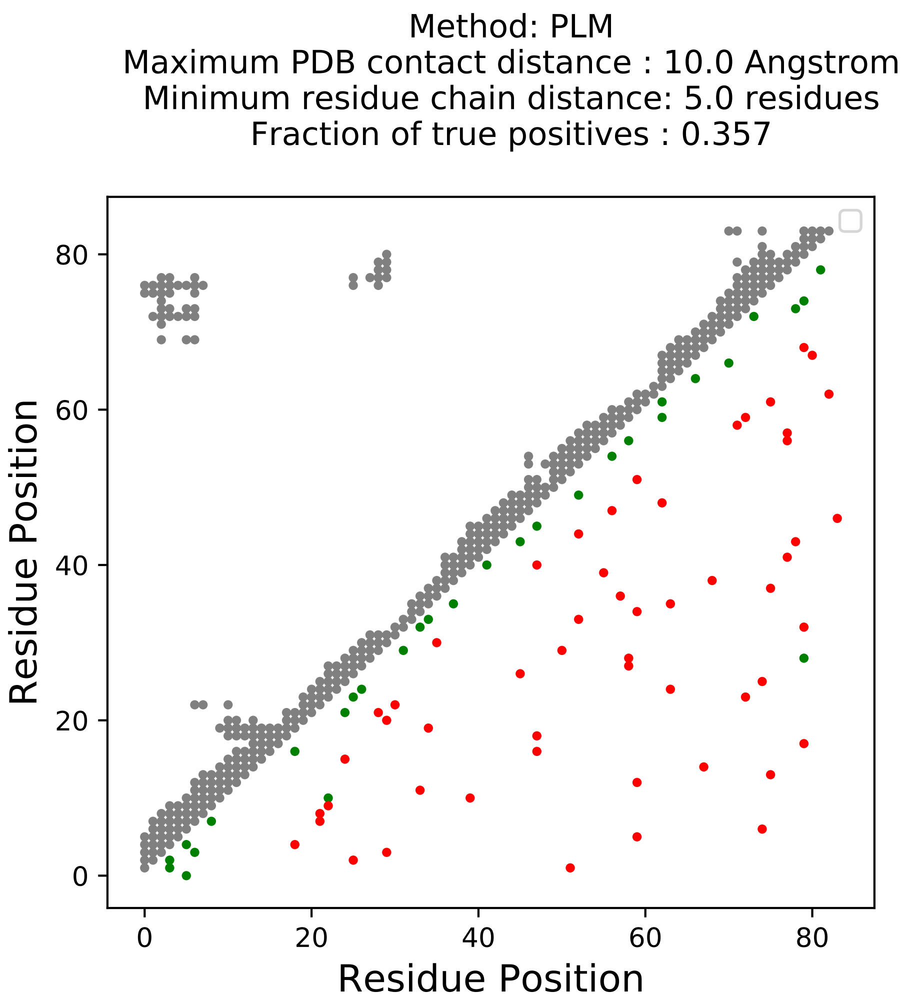

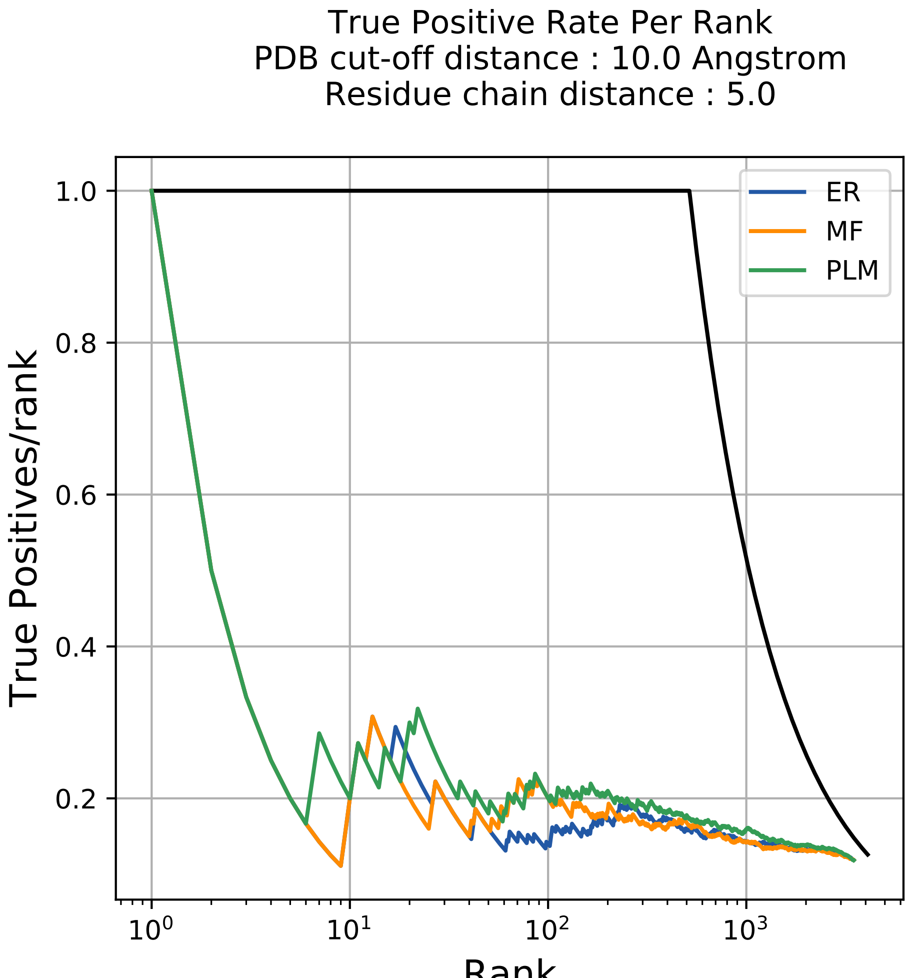

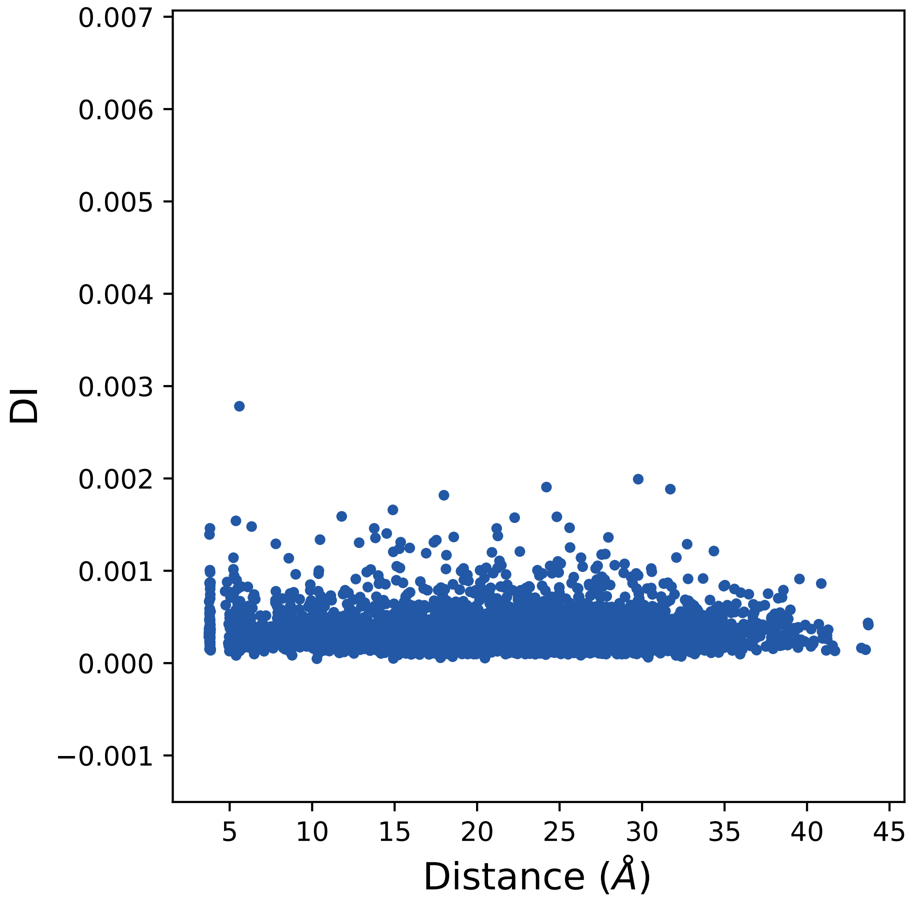

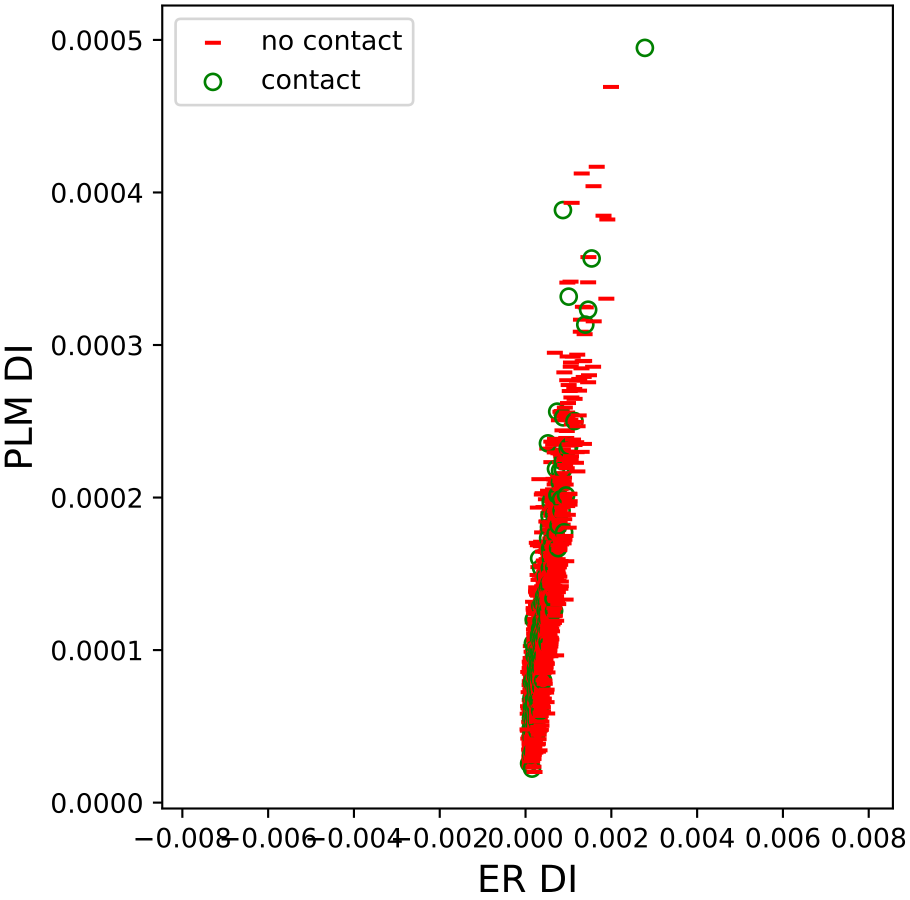

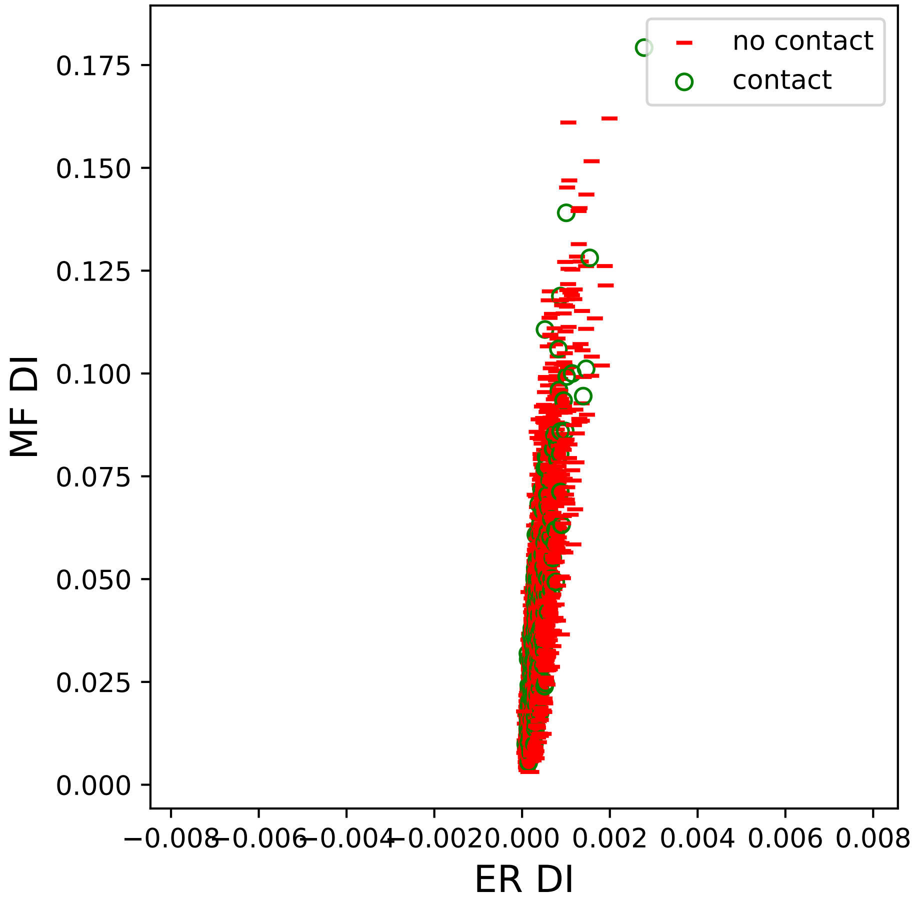

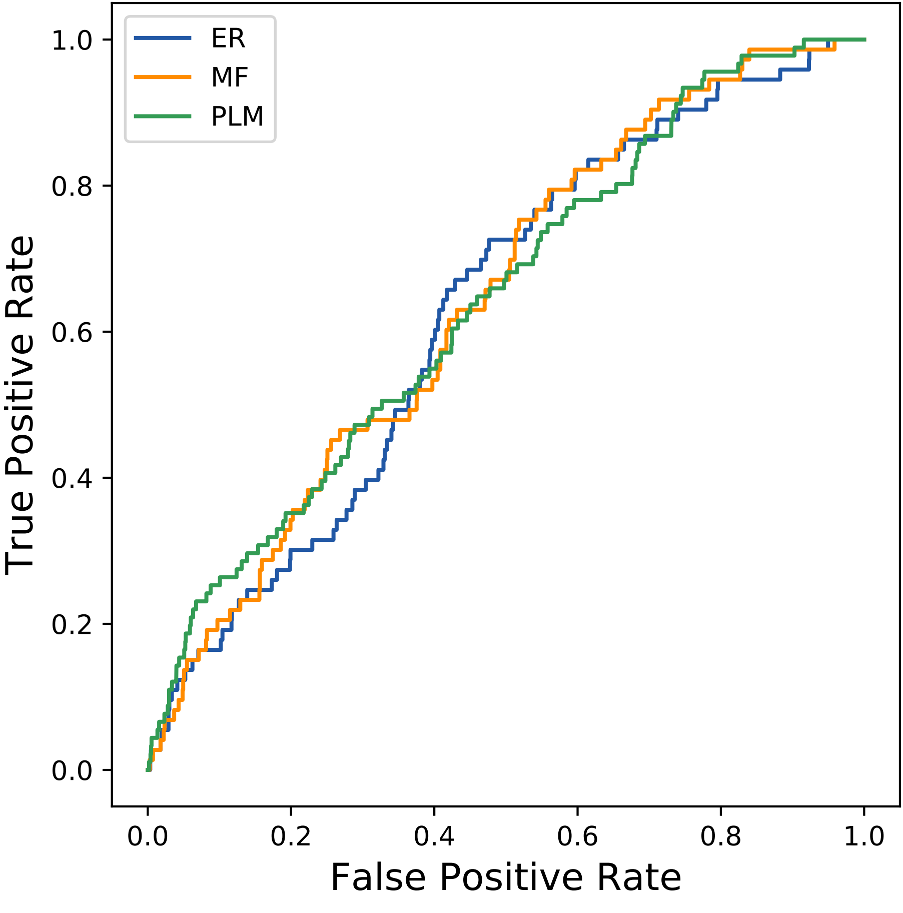

### 25seq_90col_7ps7_PF09408_plots.pdf

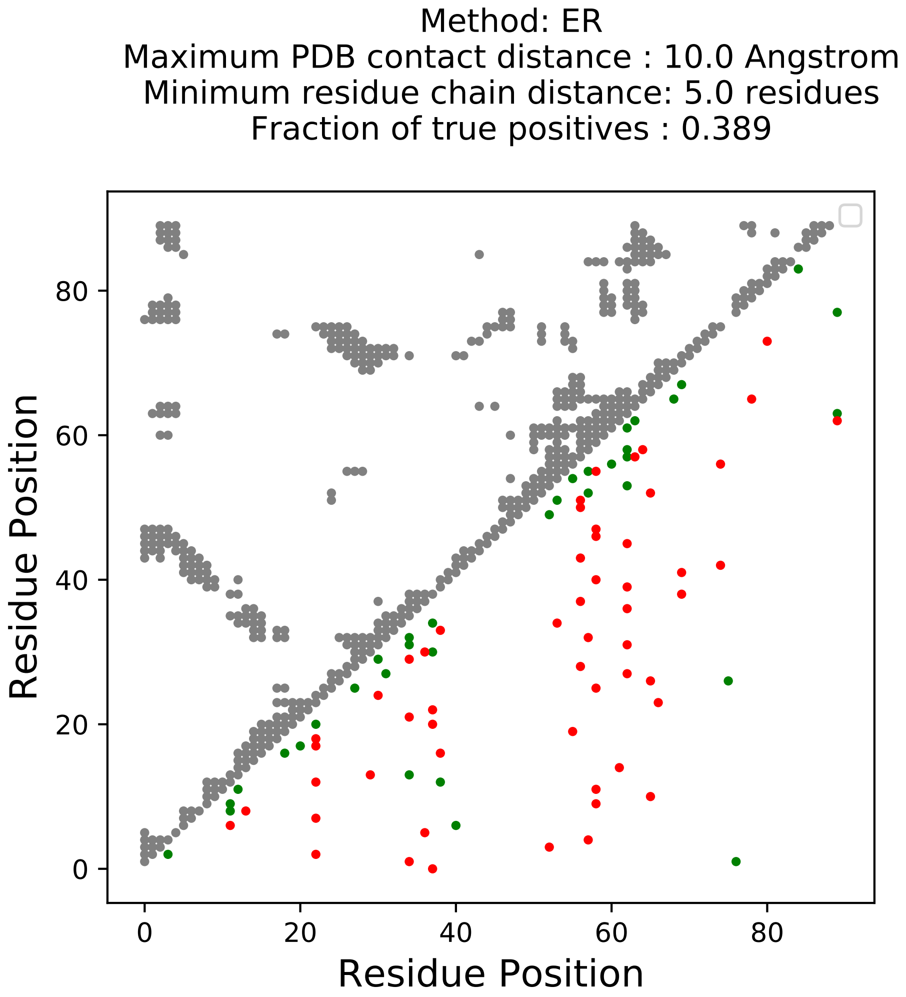

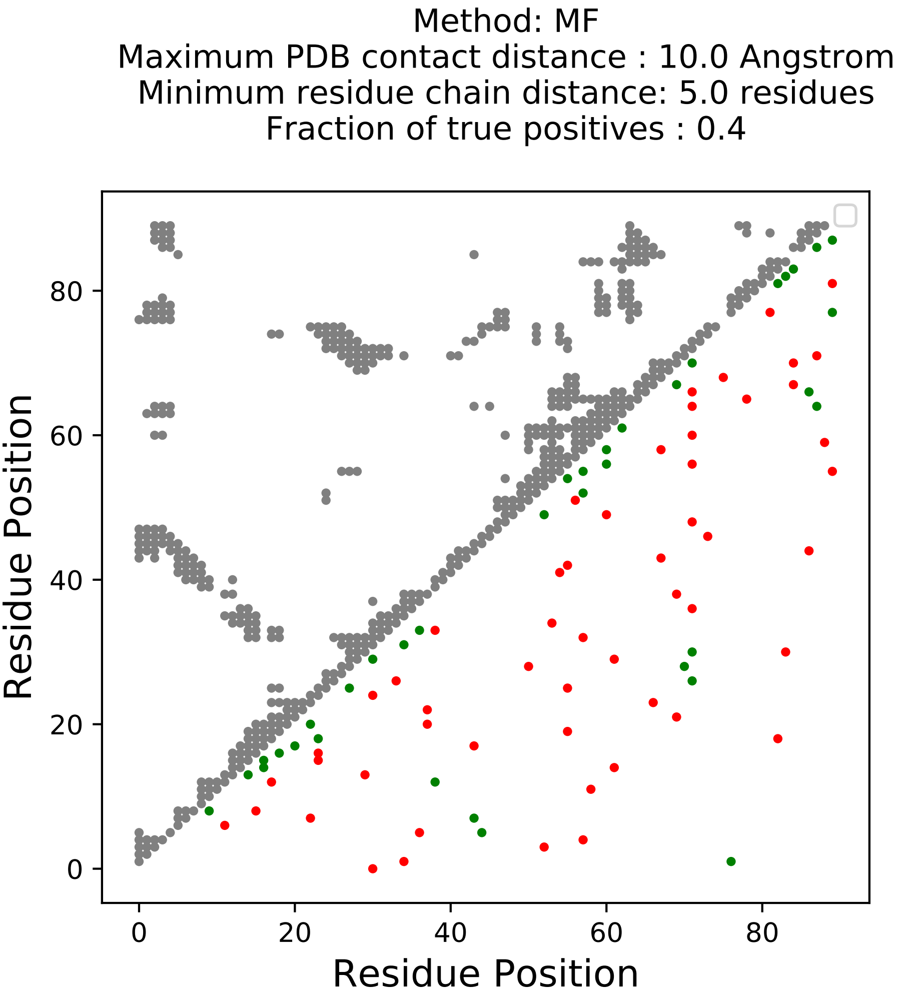

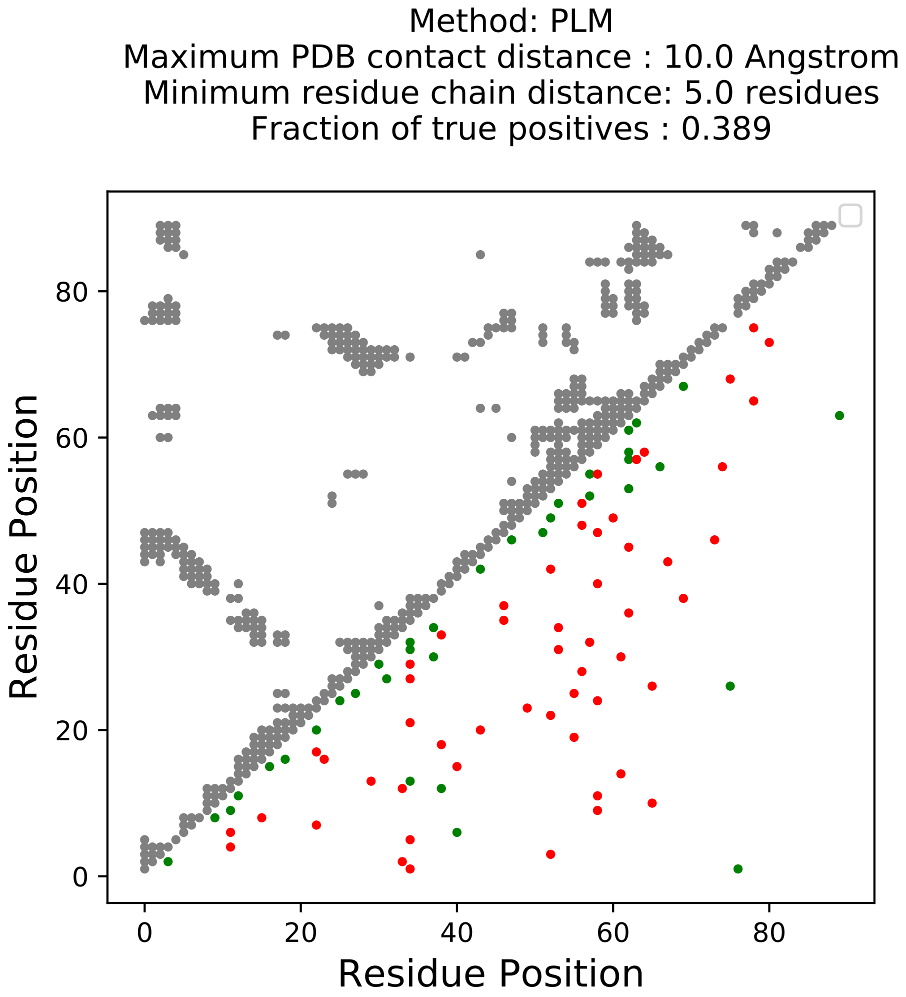

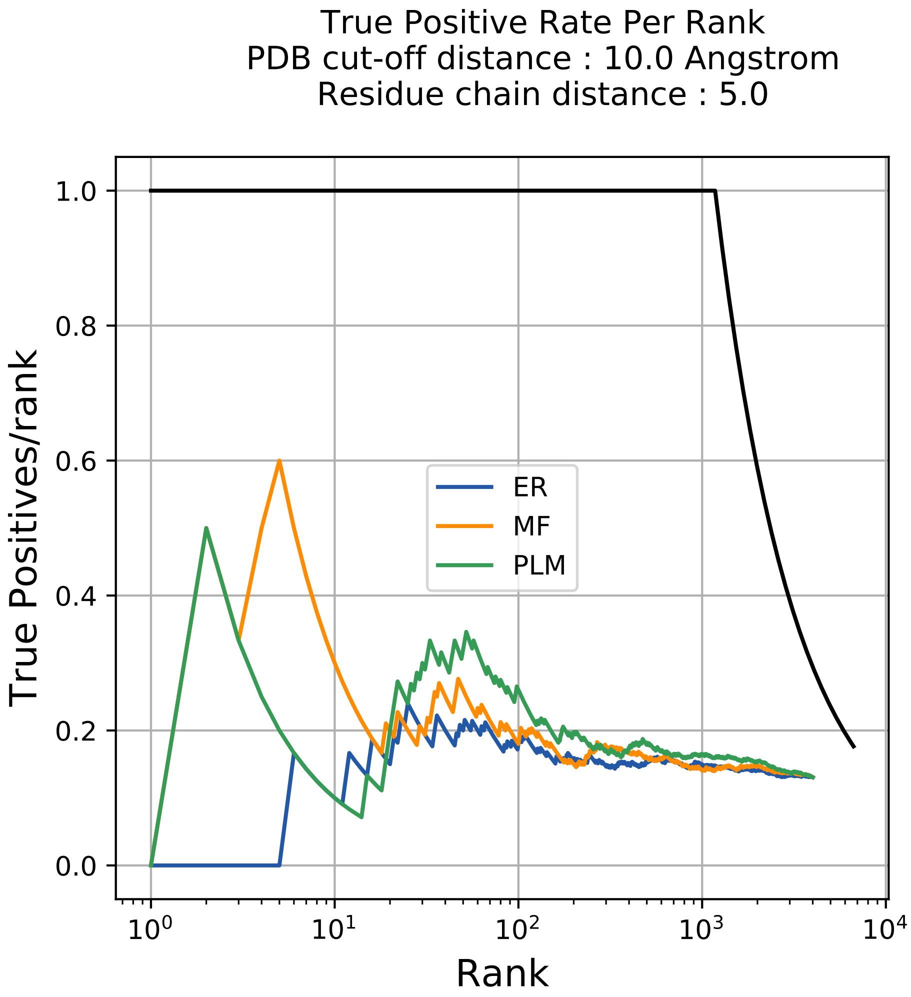

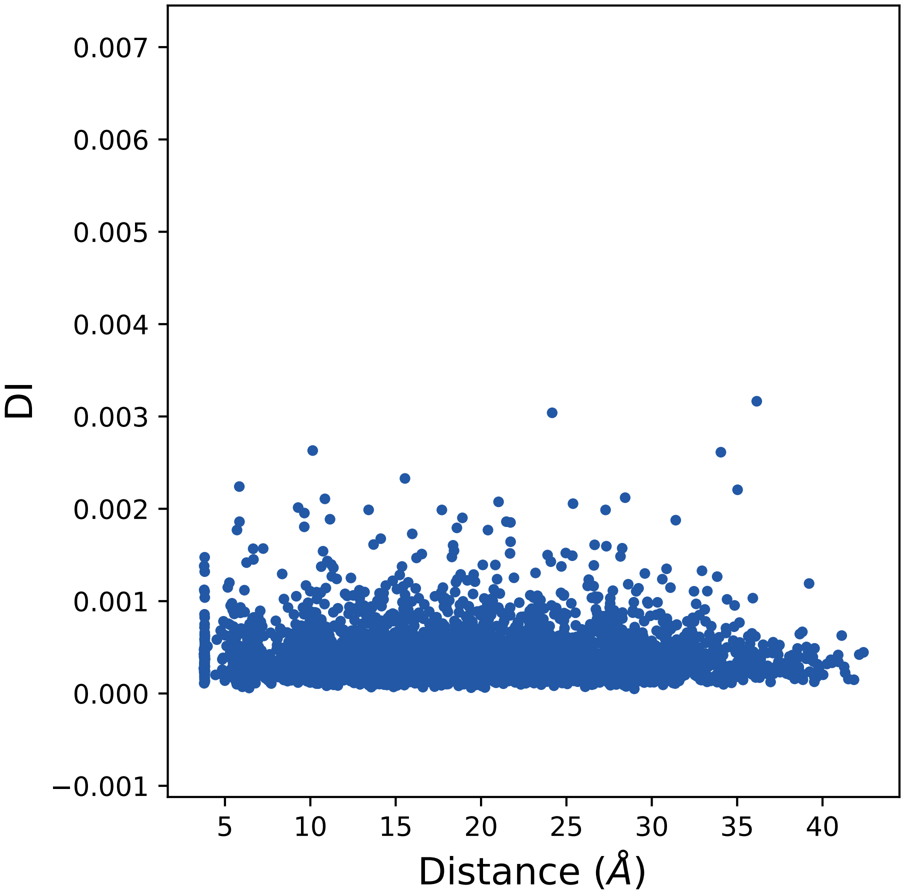

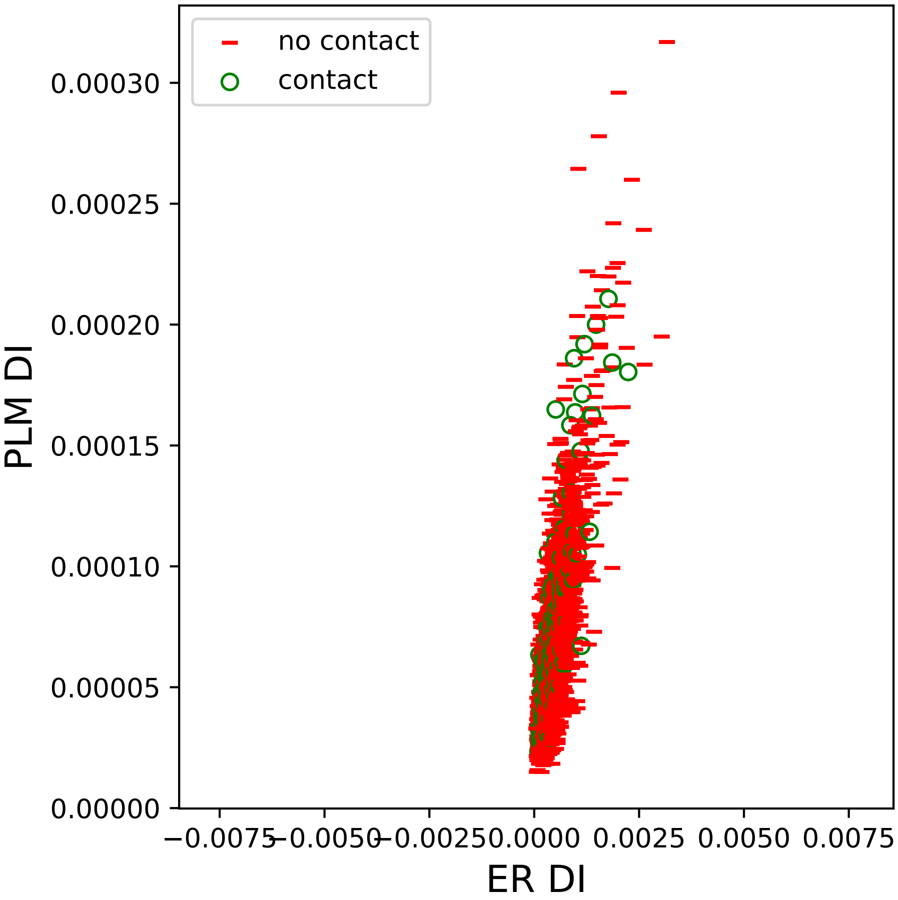

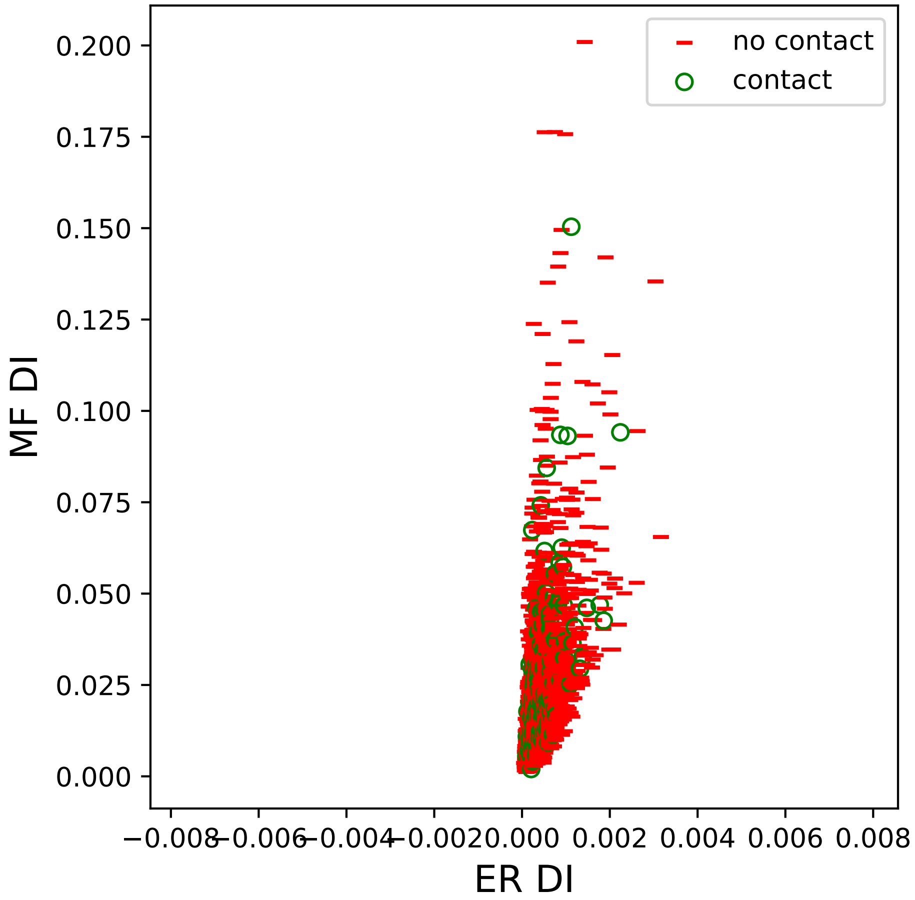

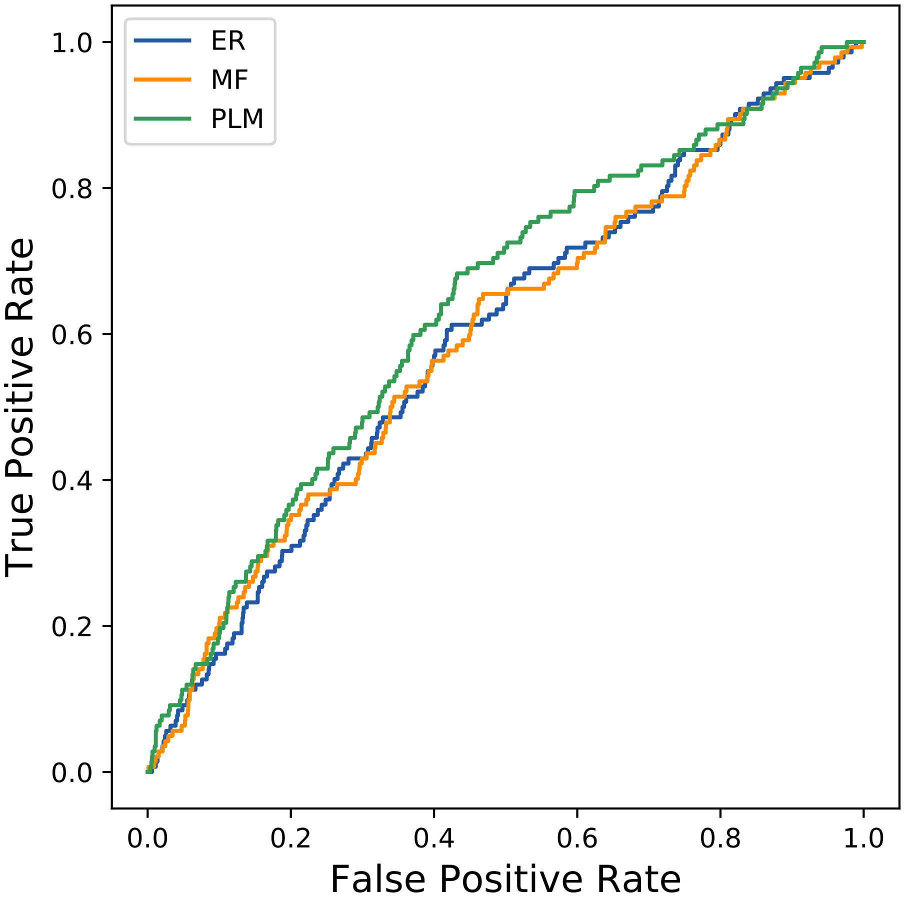

### 26seq_27col_4cfi_PF12613_plots.pdf

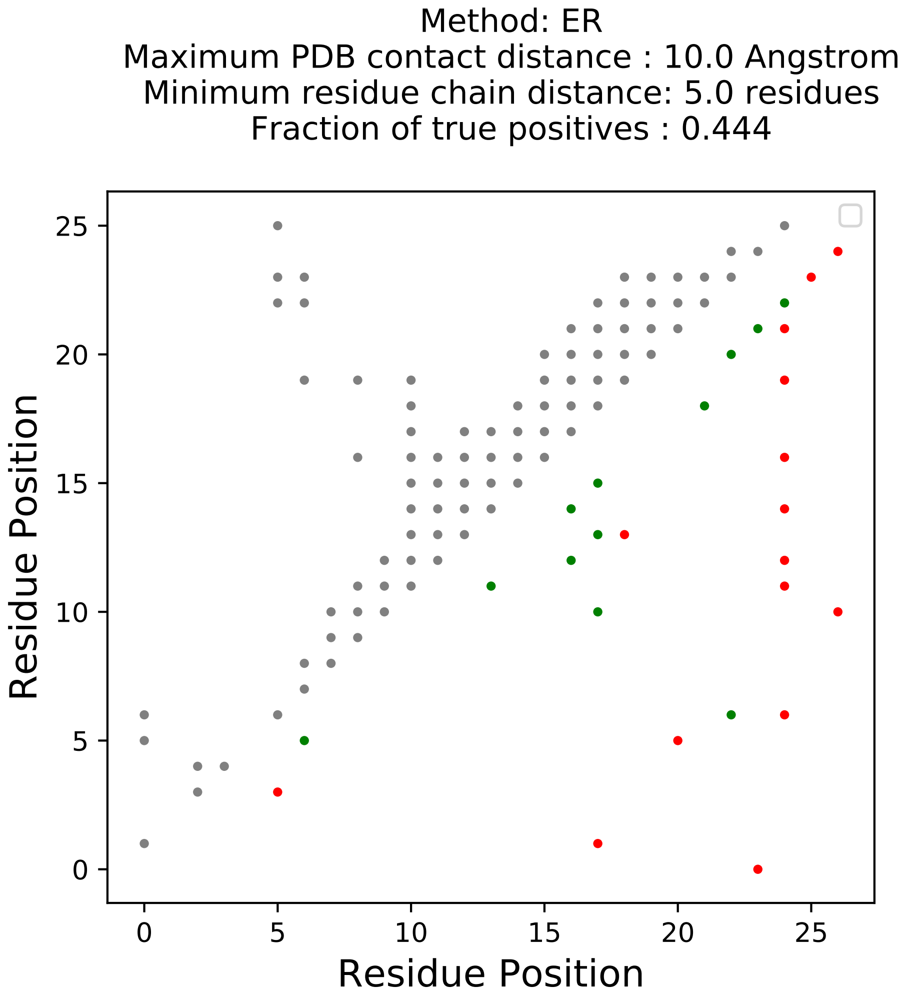

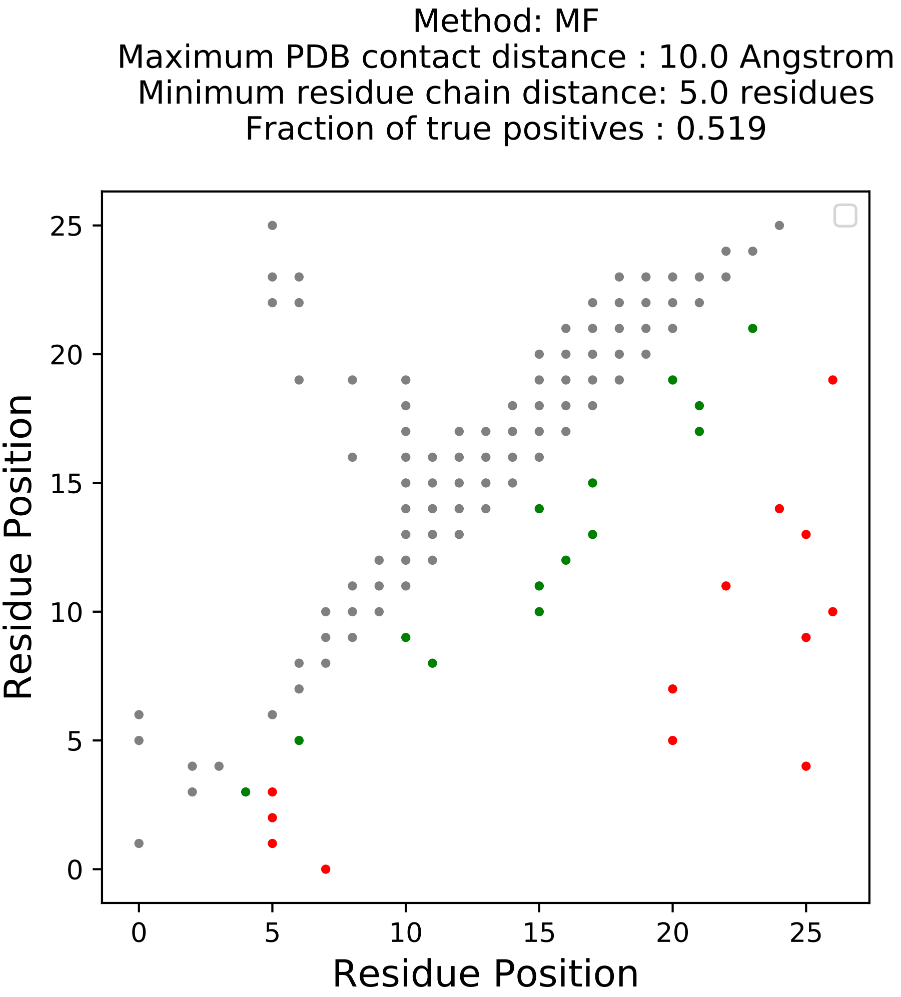

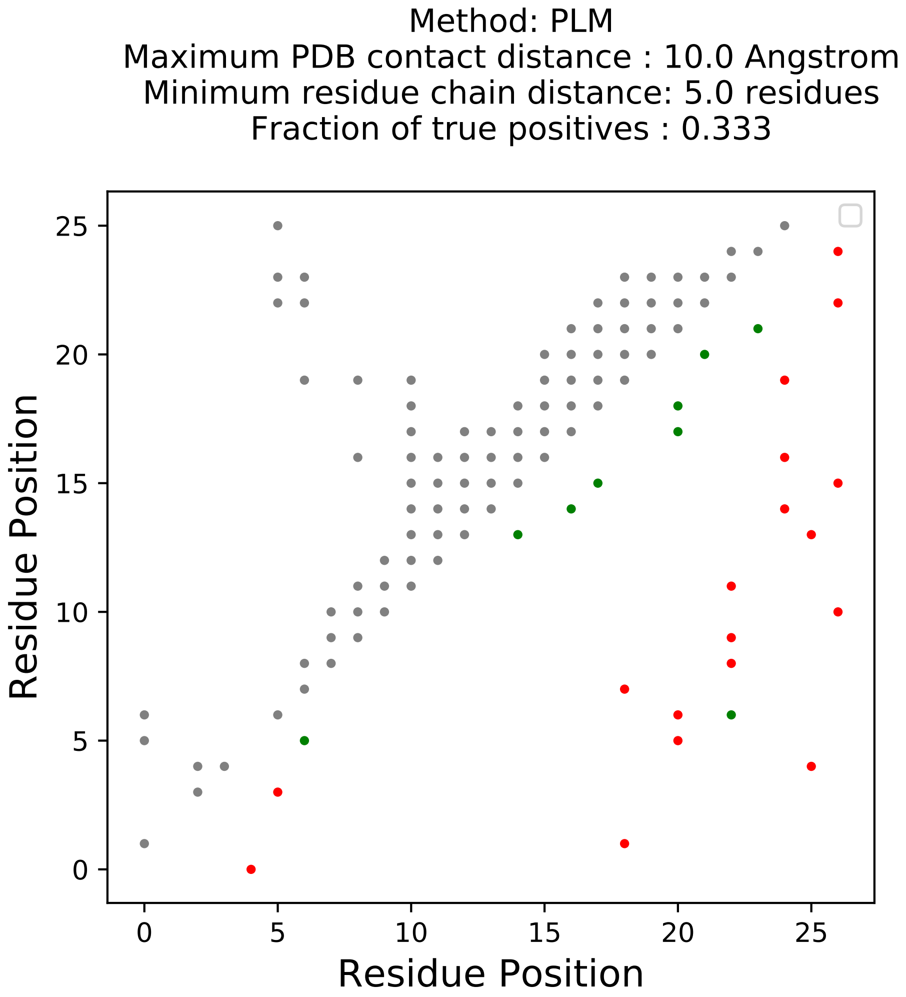

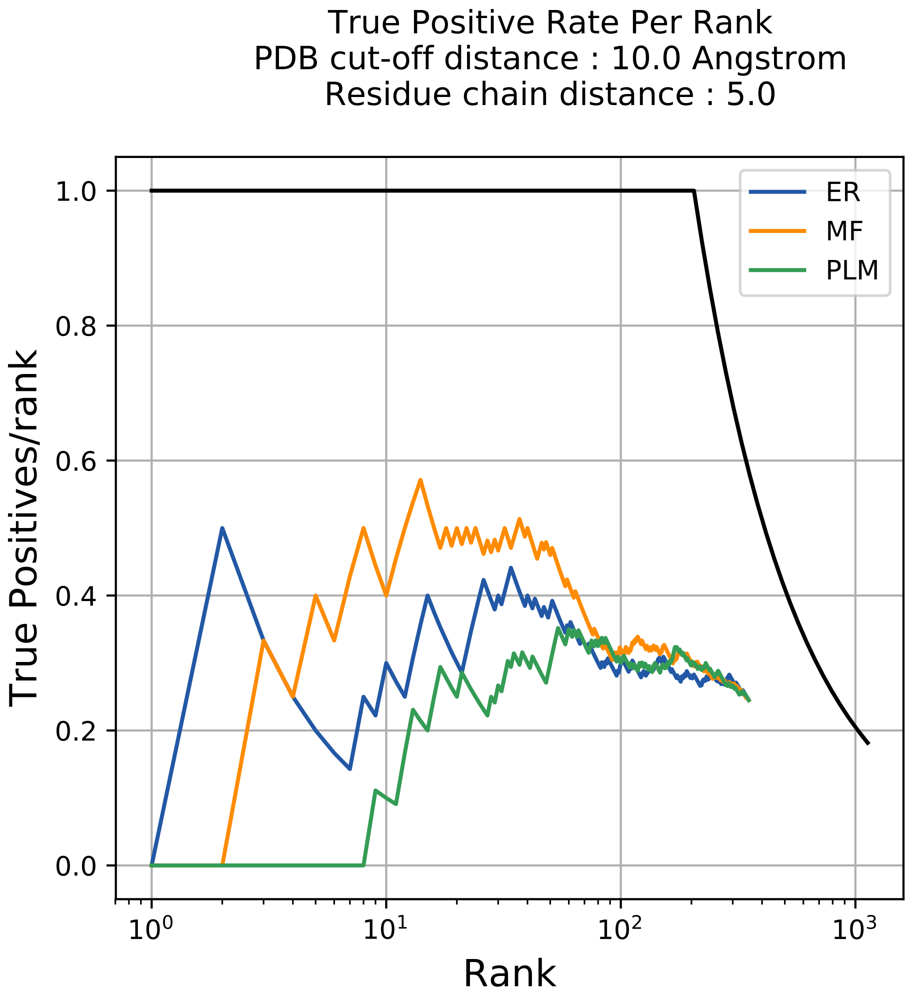

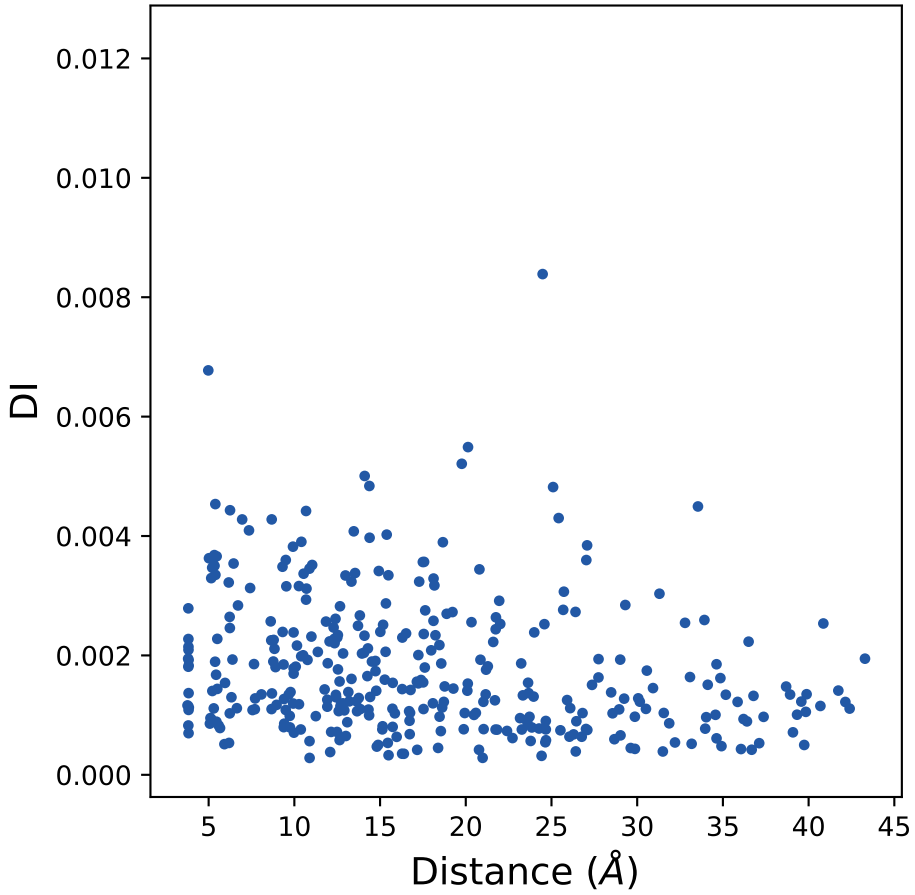

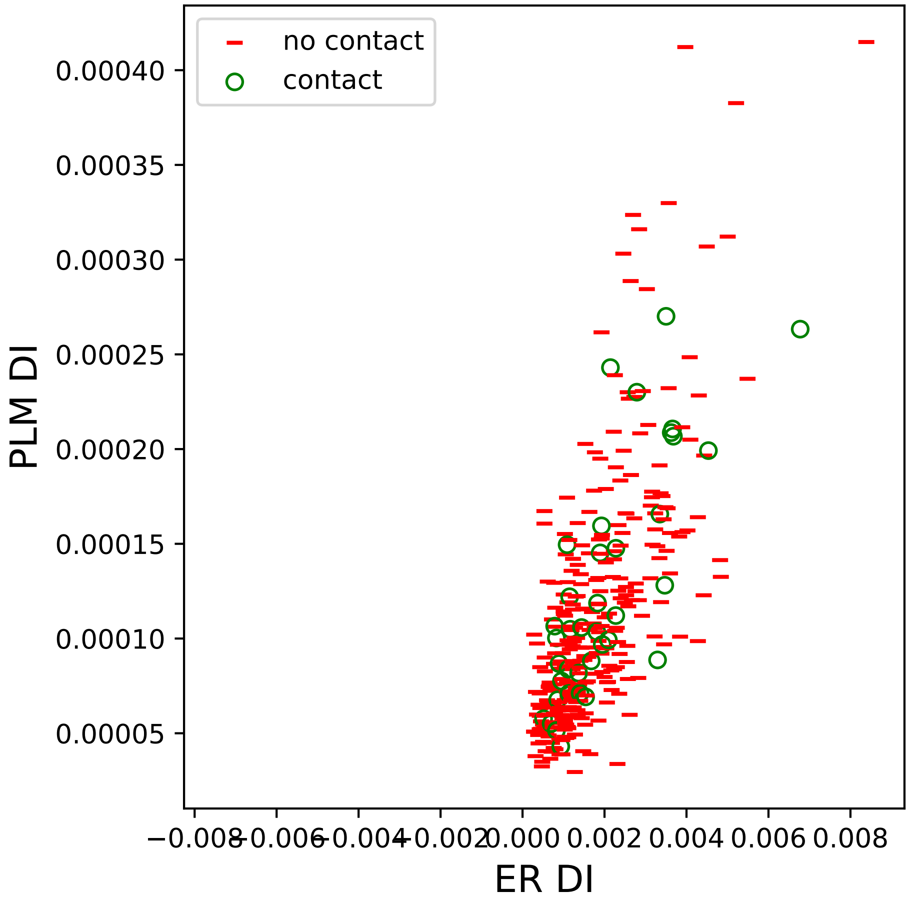

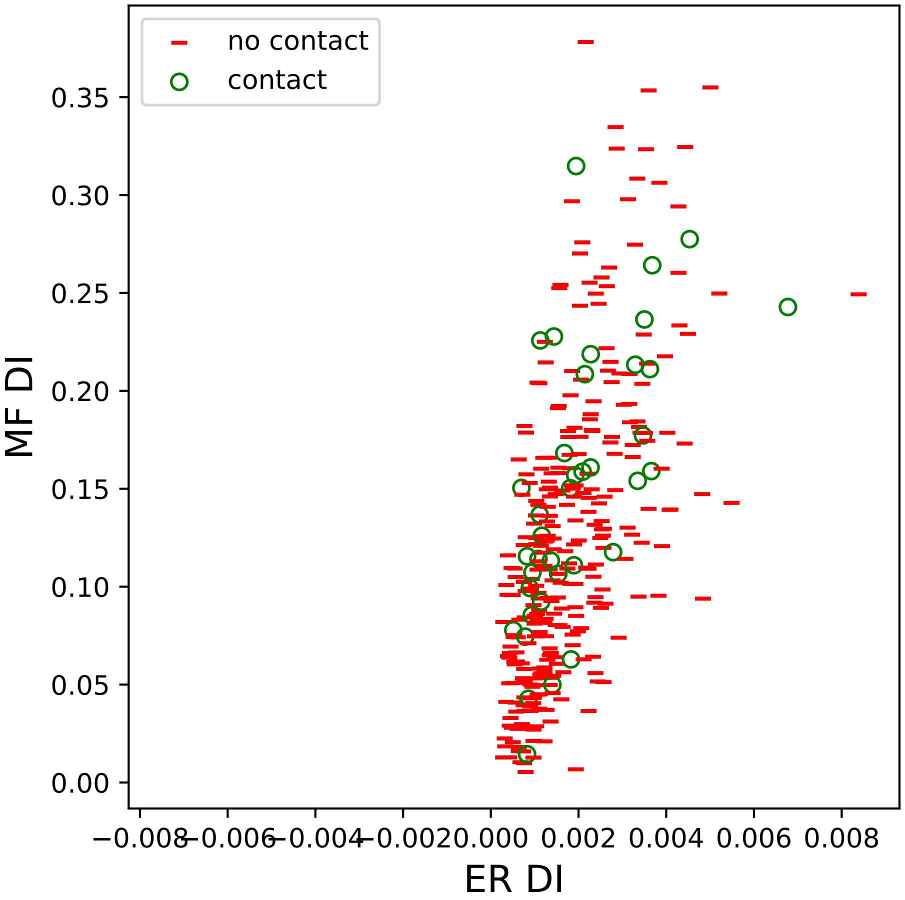

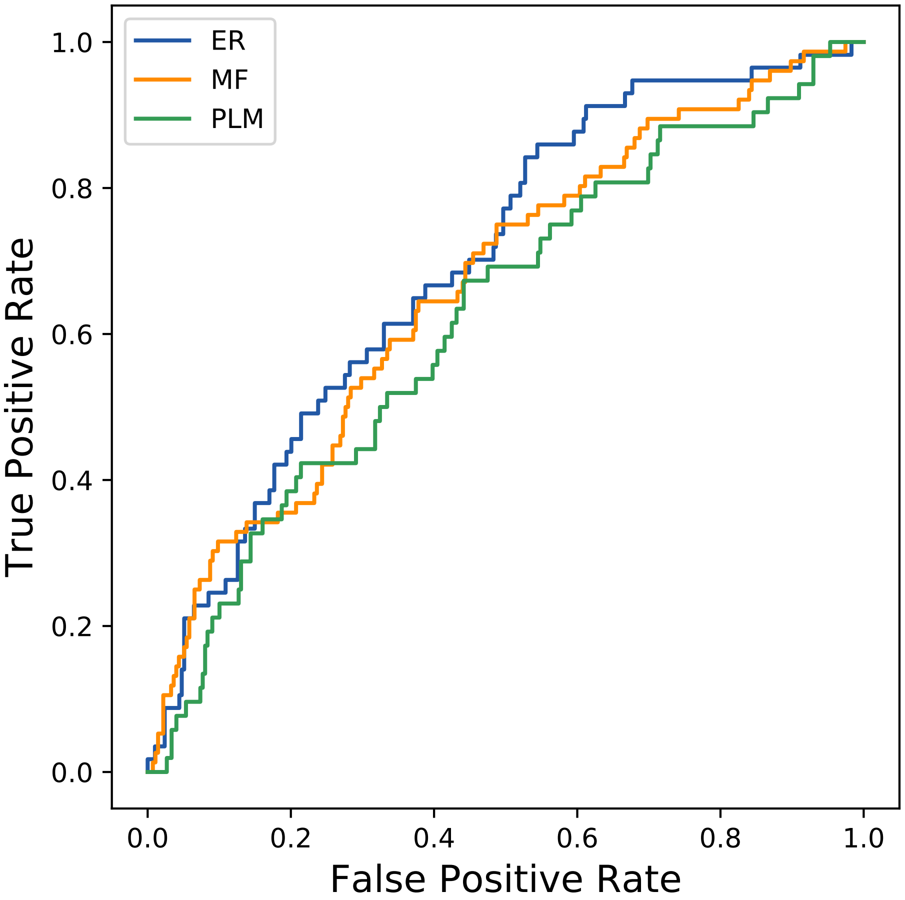

### 38seq_97col_6idd_PF00509_plots.pdf

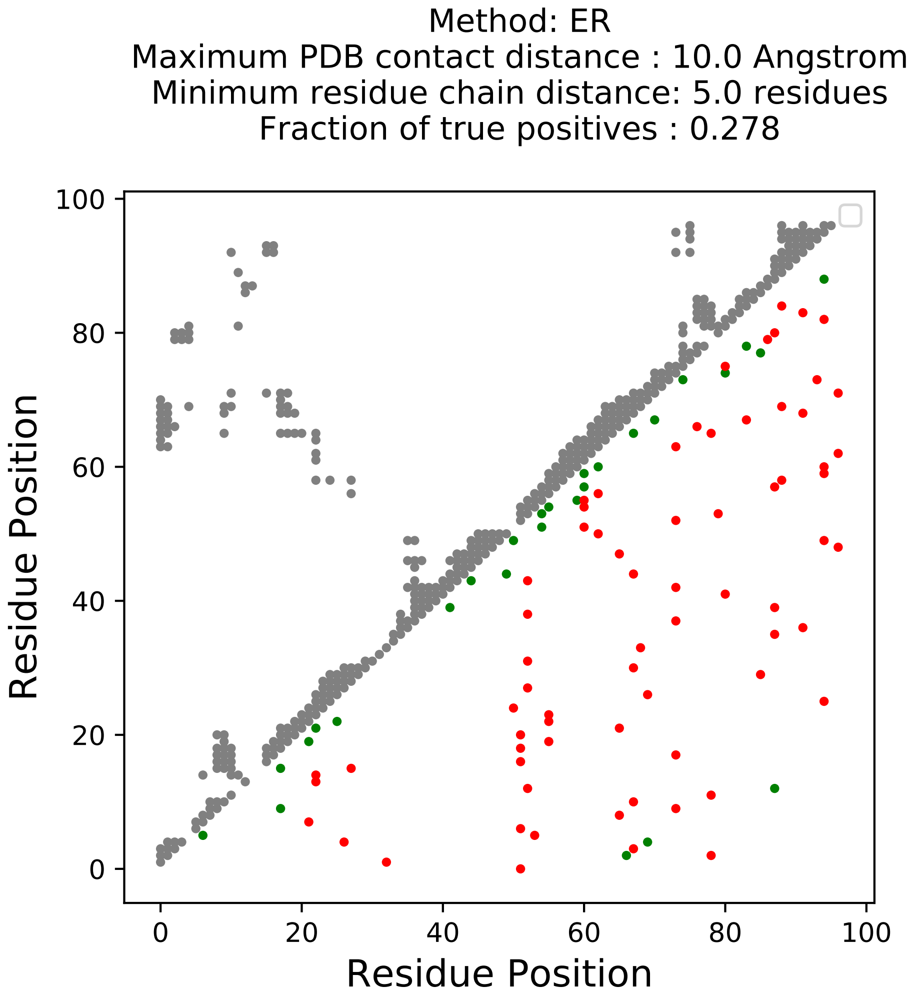

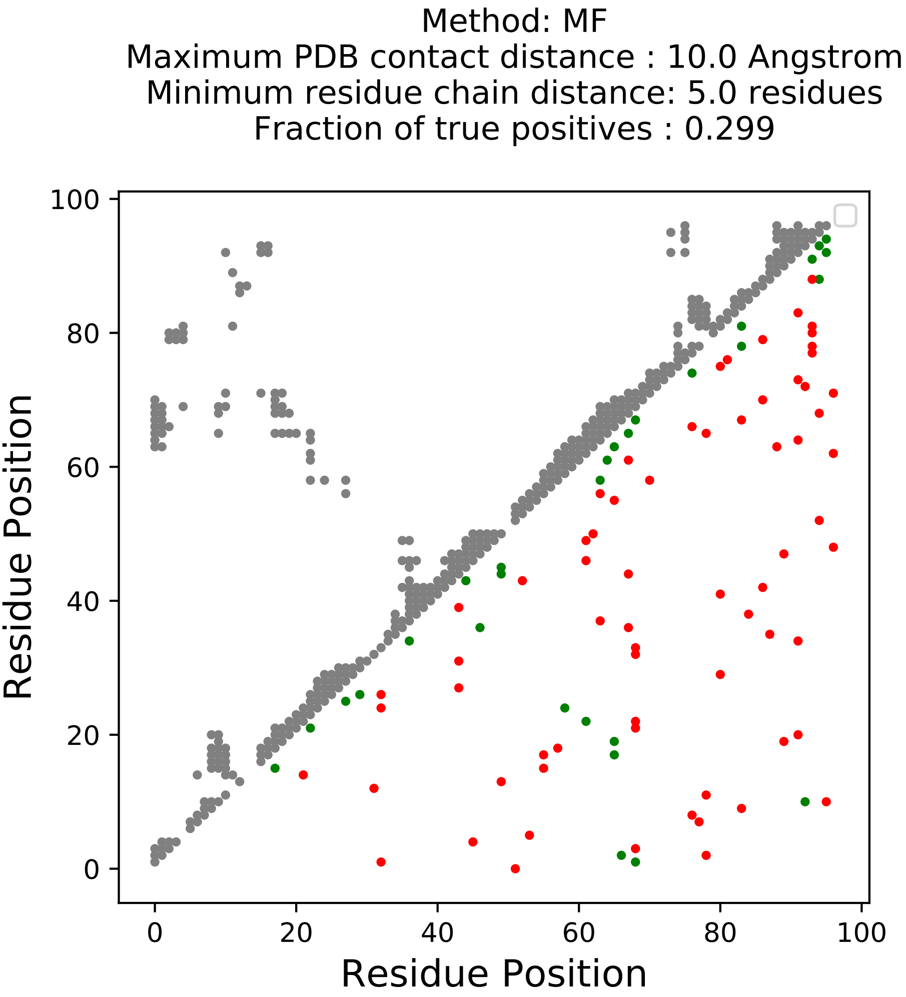

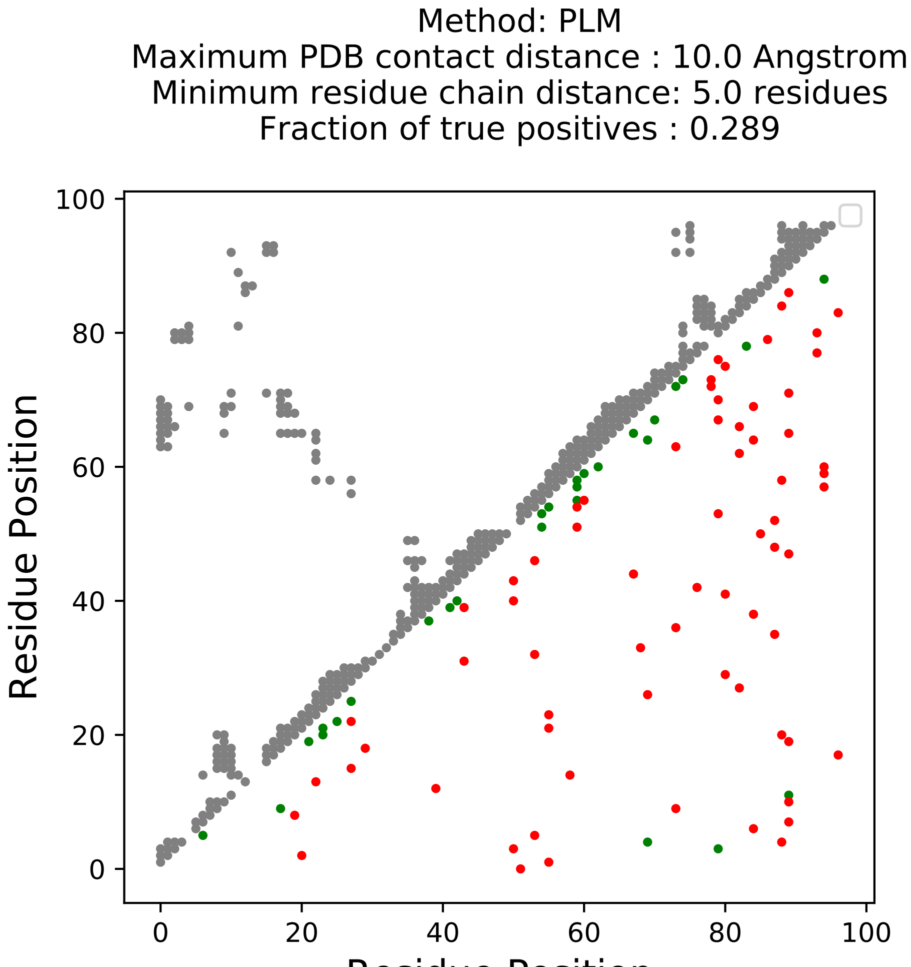

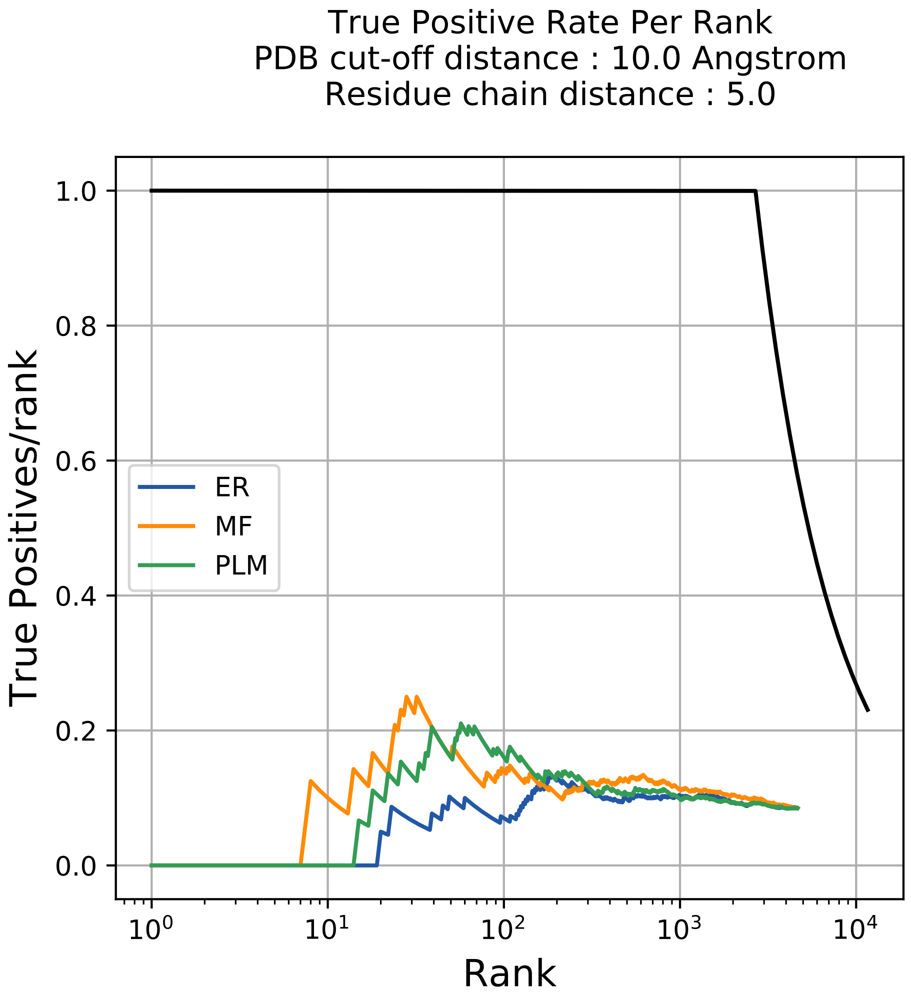

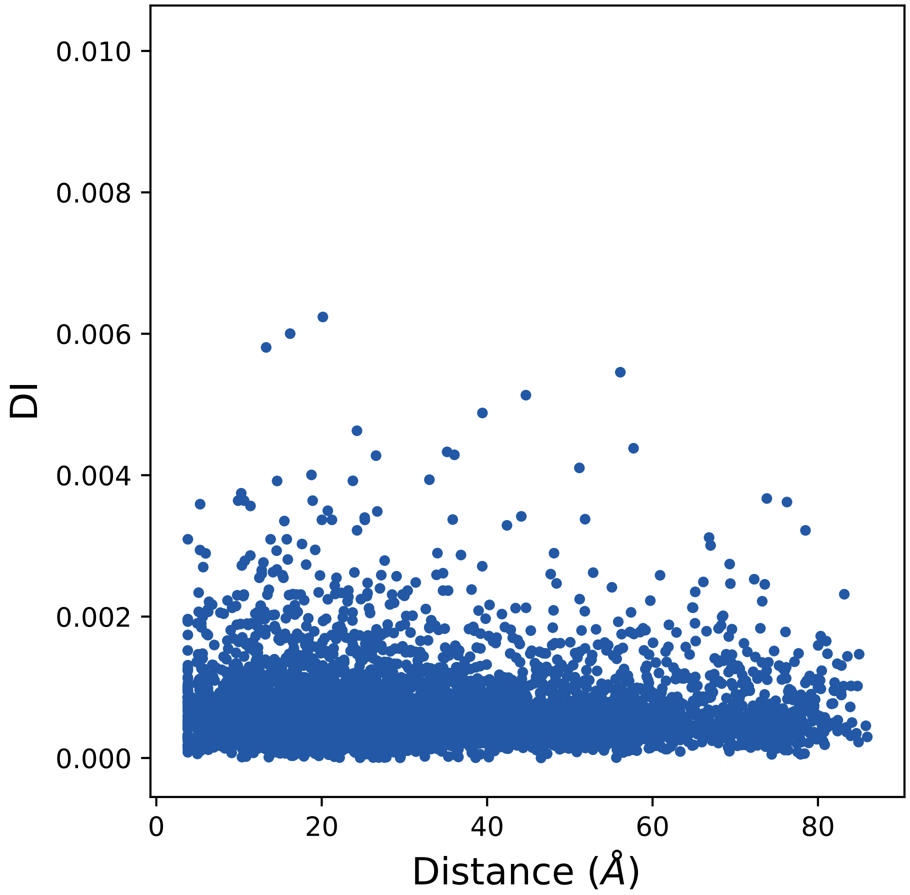

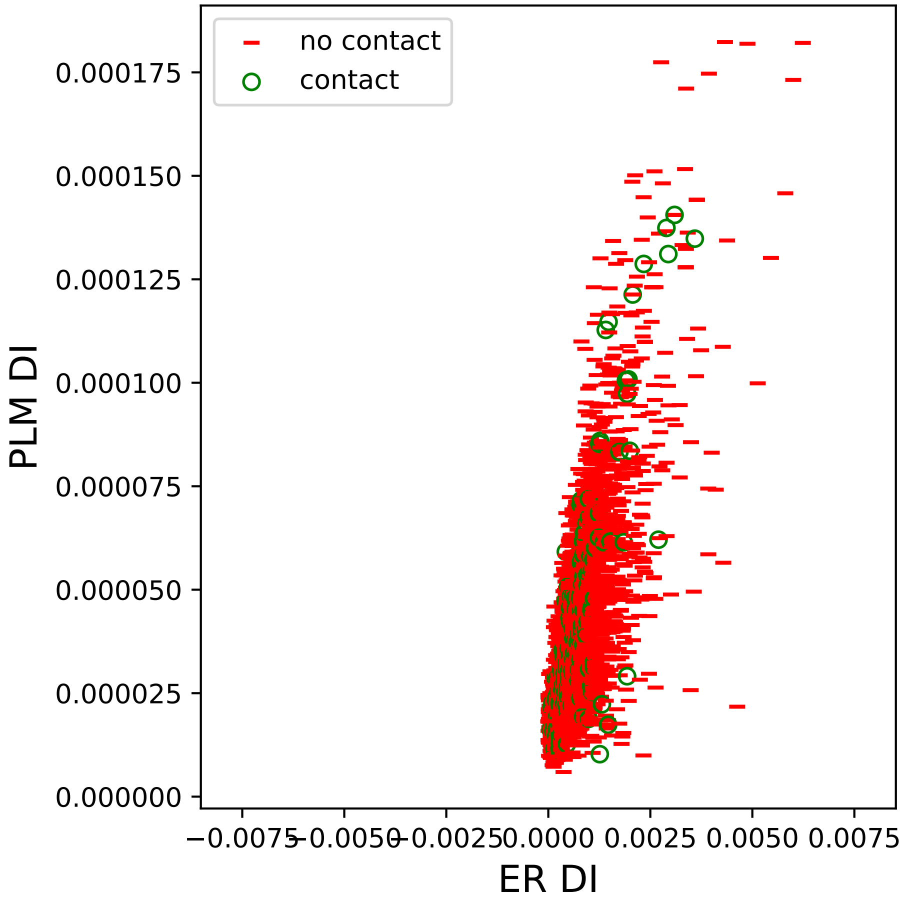

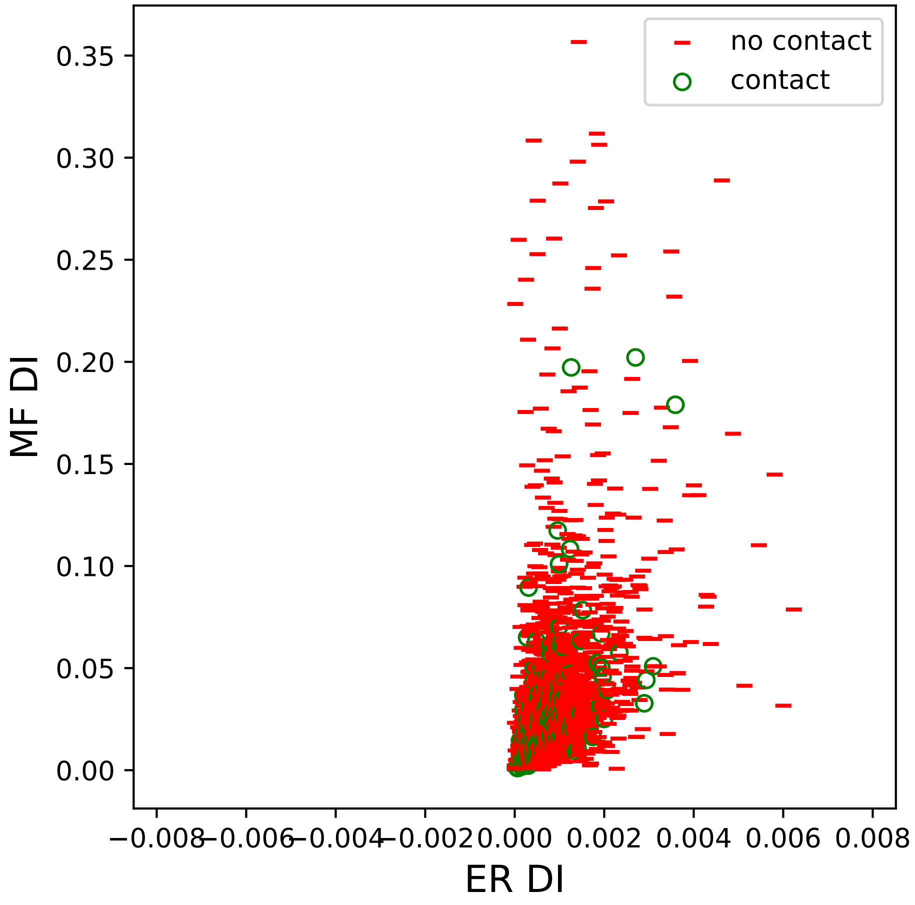

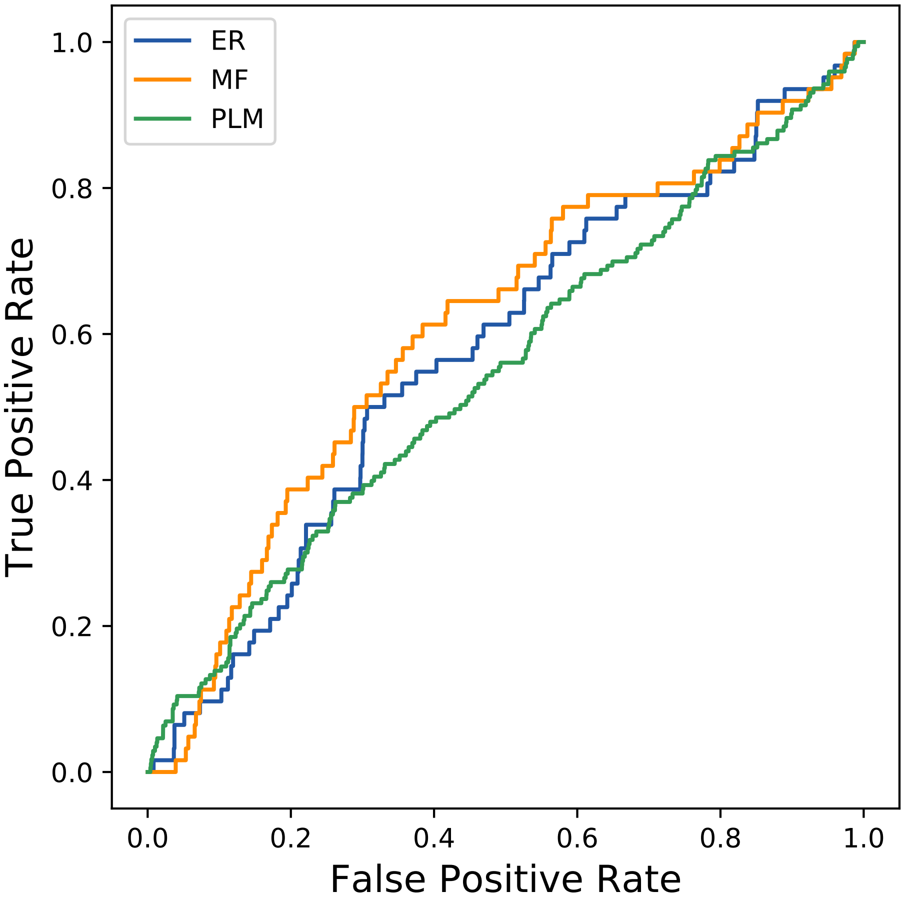

### 104seq_23col_1zwb_PF01279_plots.pdf

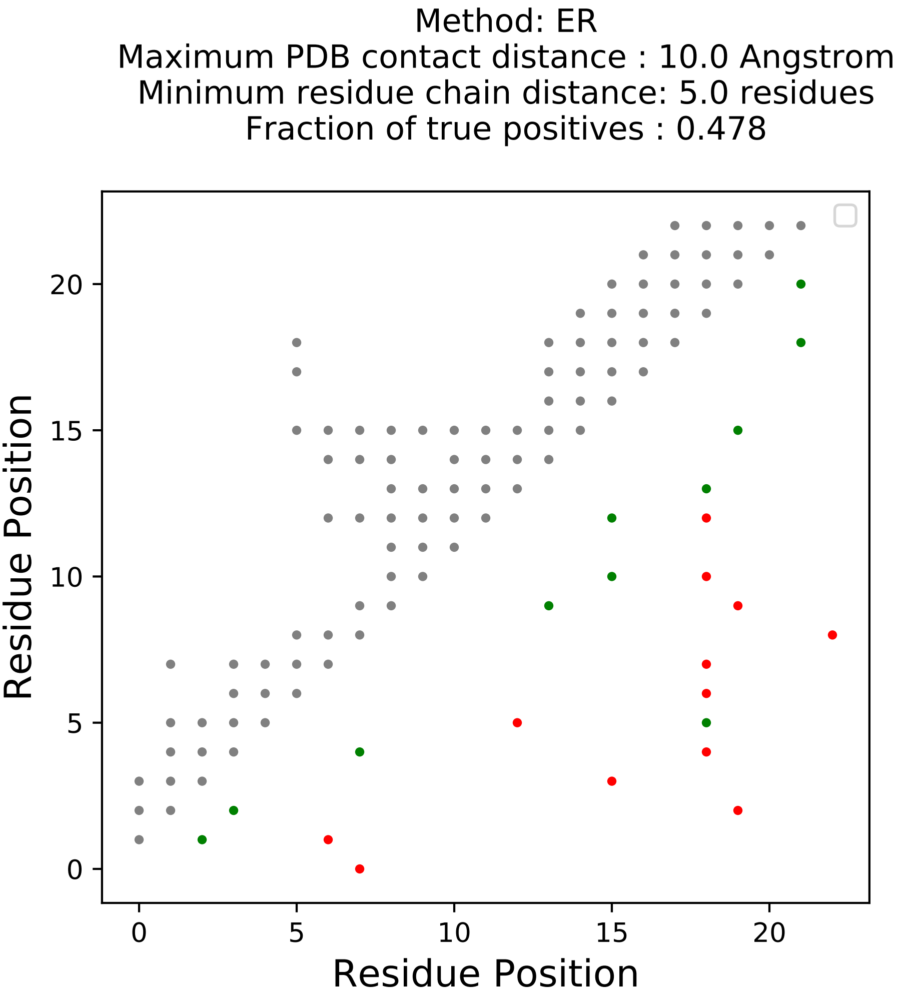

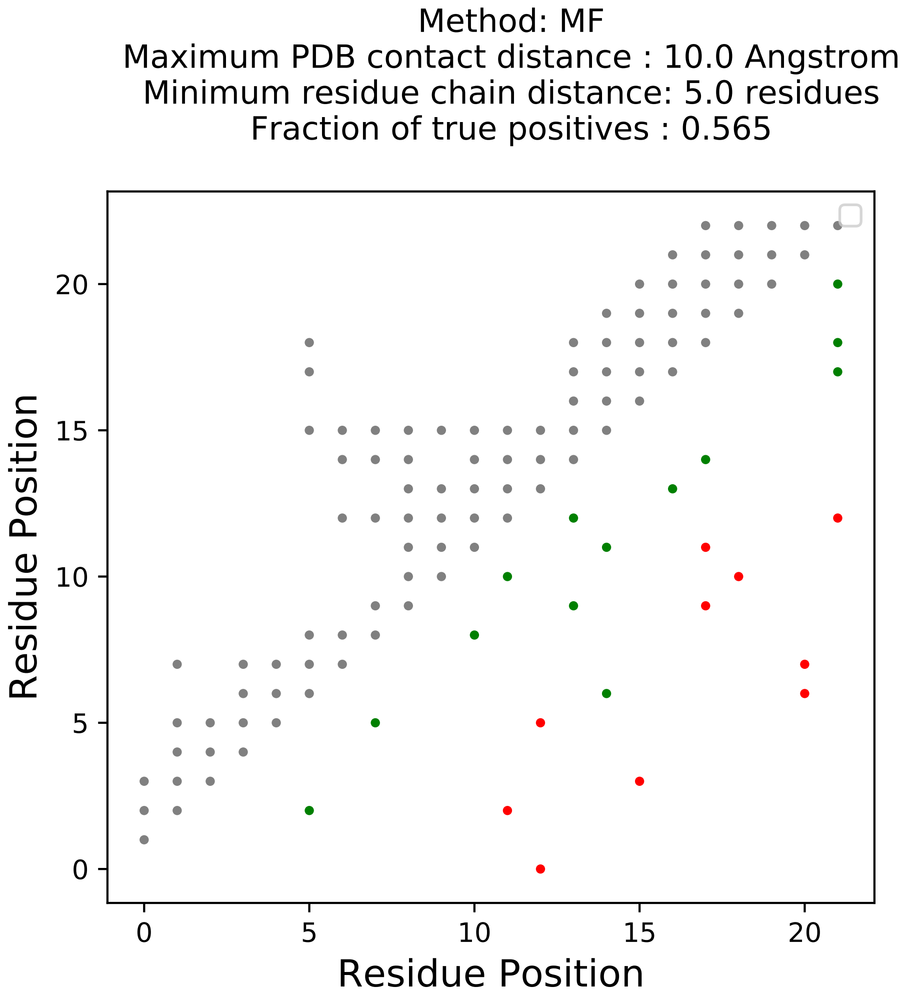

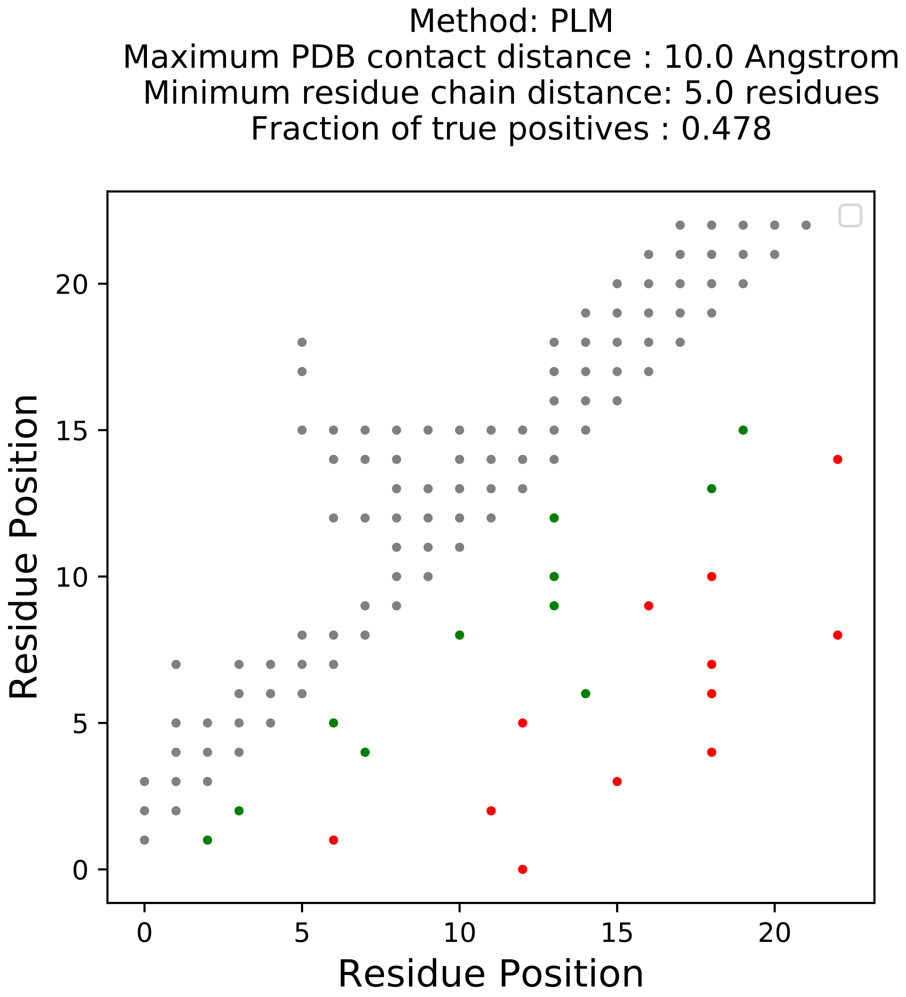

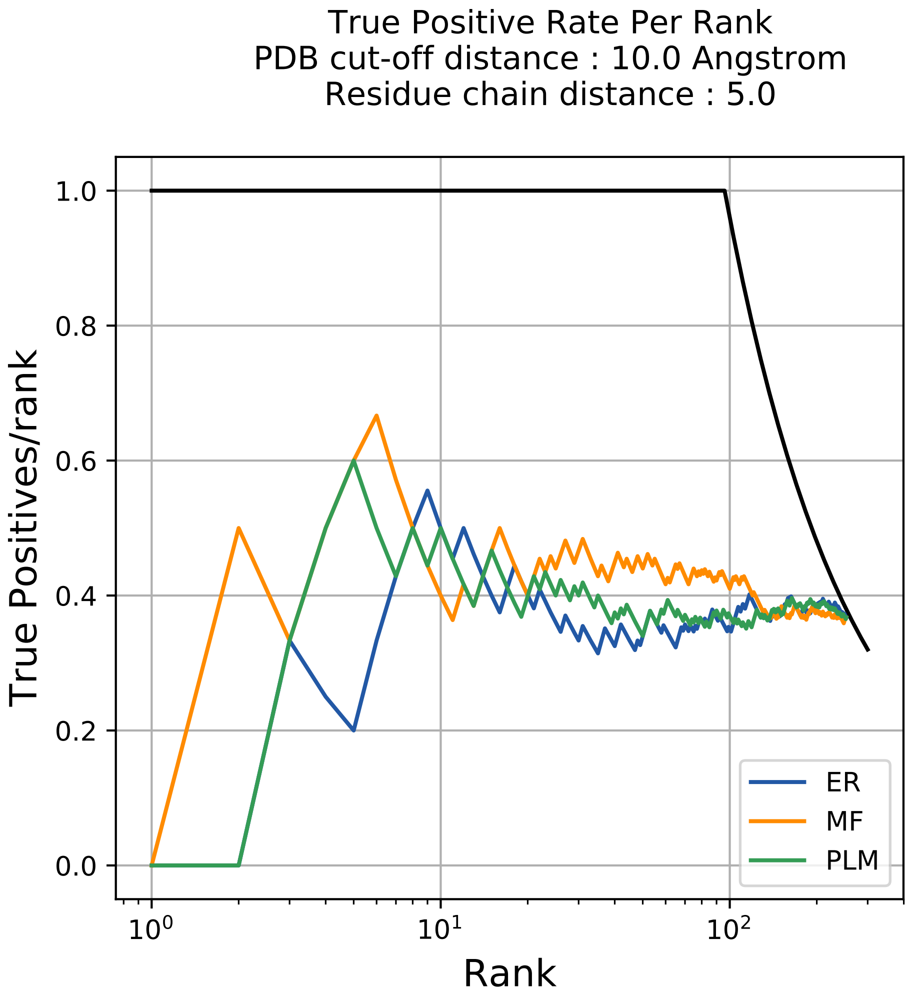

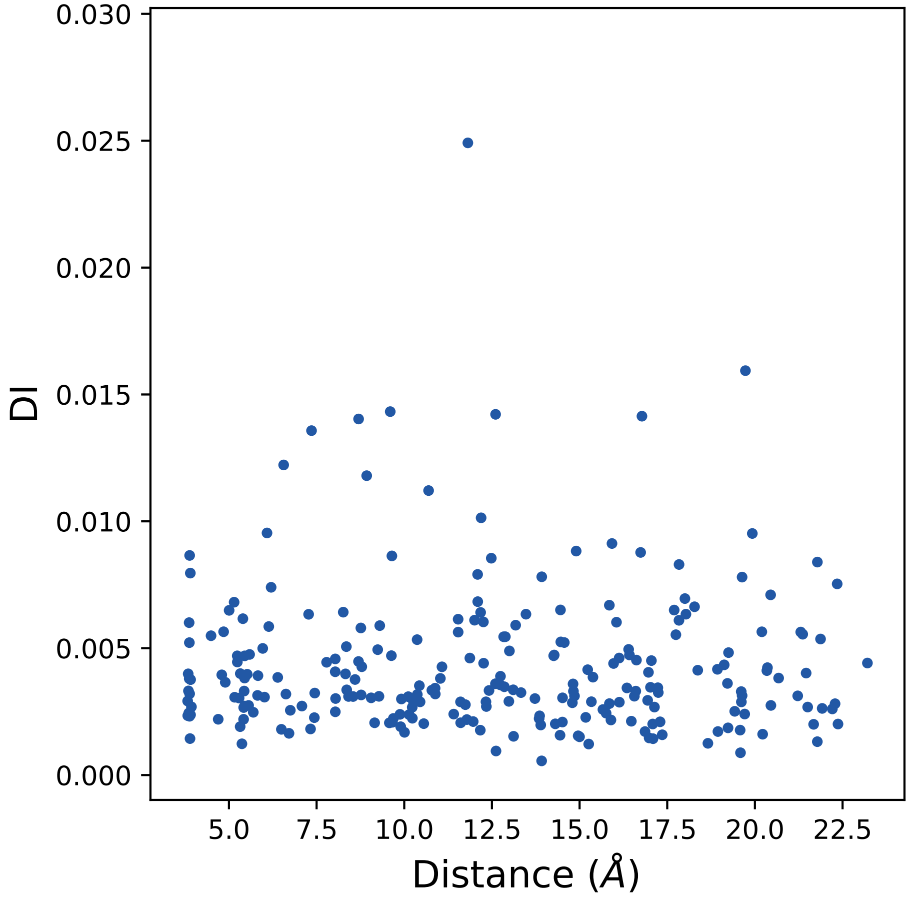

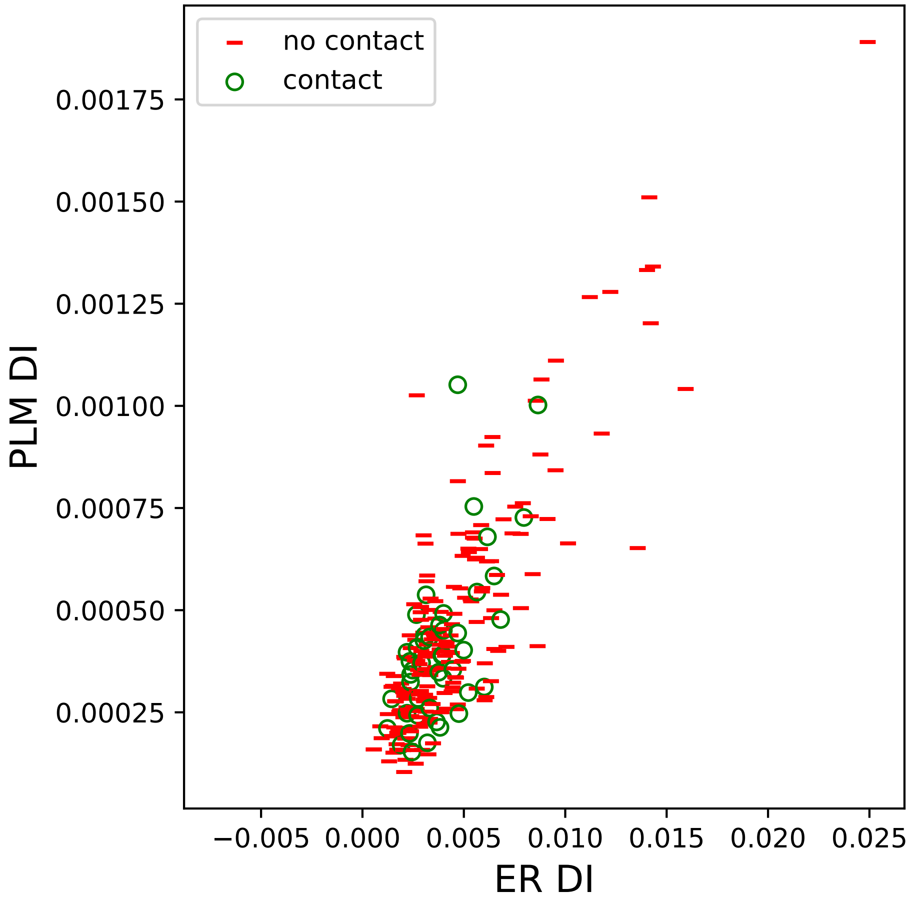

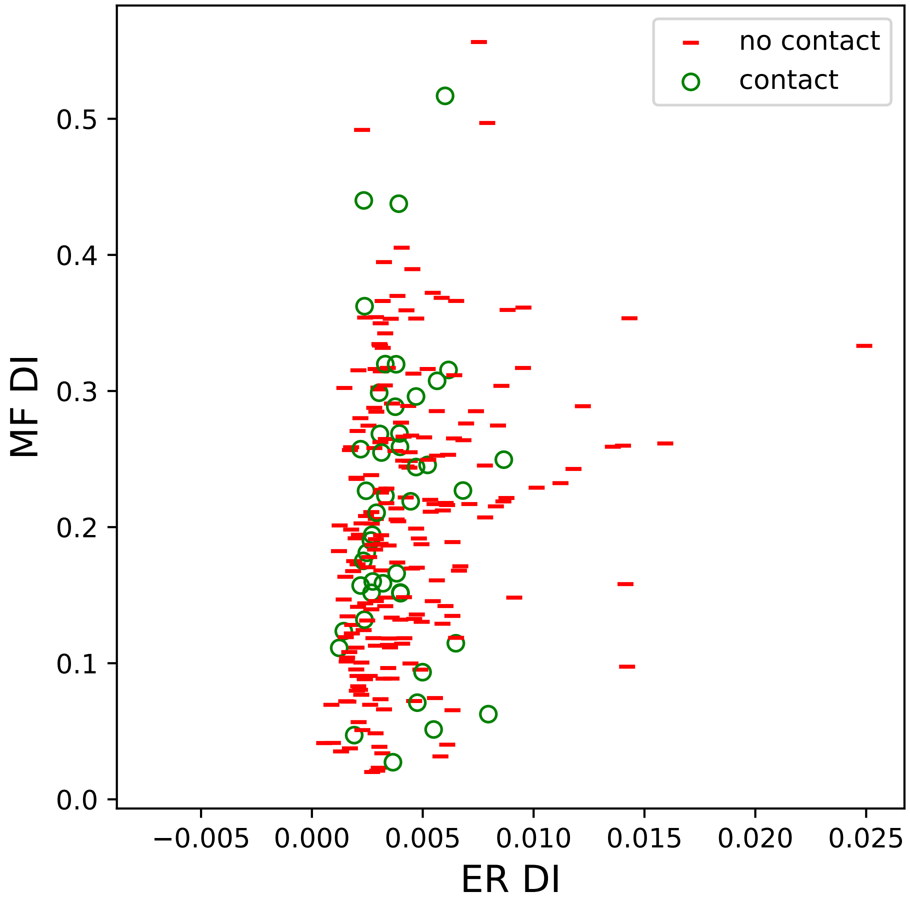

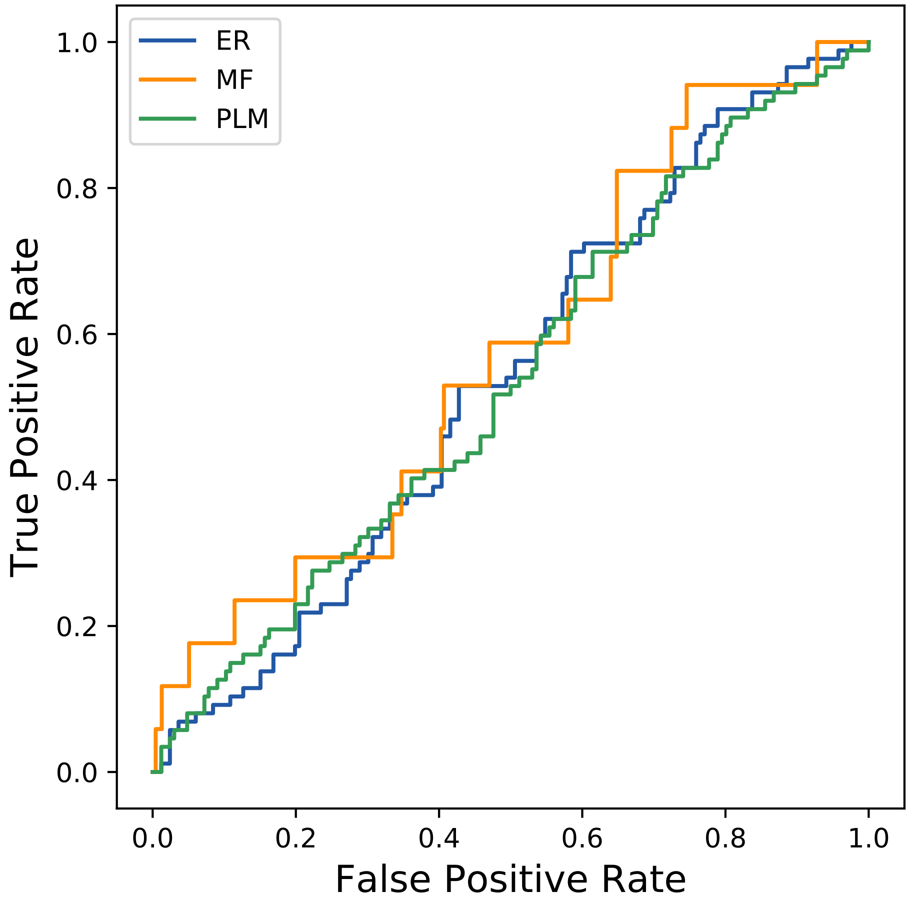

### 157seq_131col_5cj0_PF06632_plots.pdf

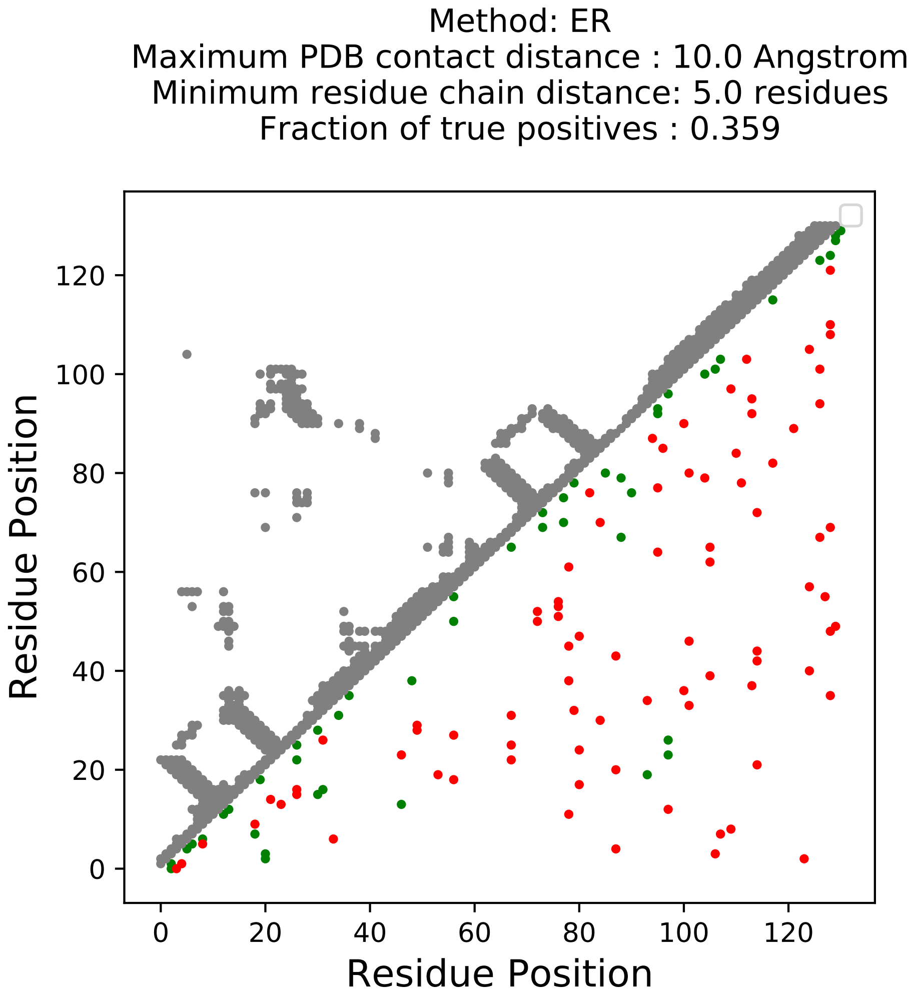

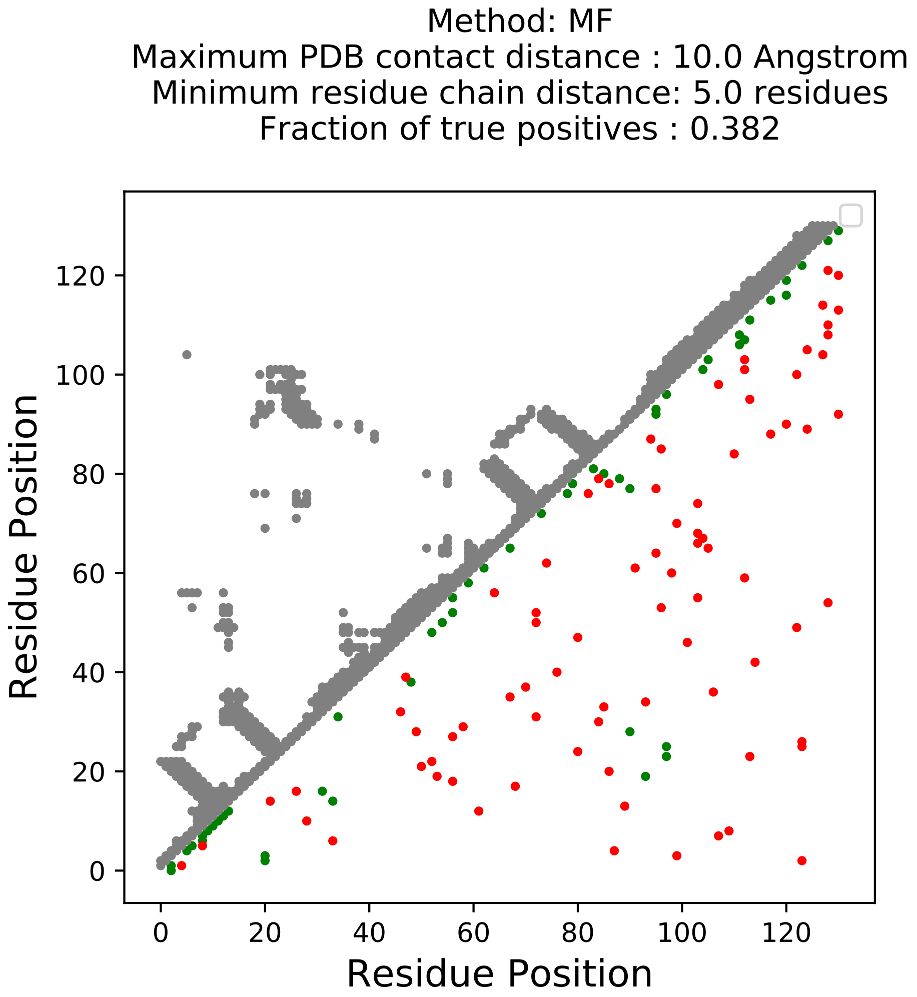

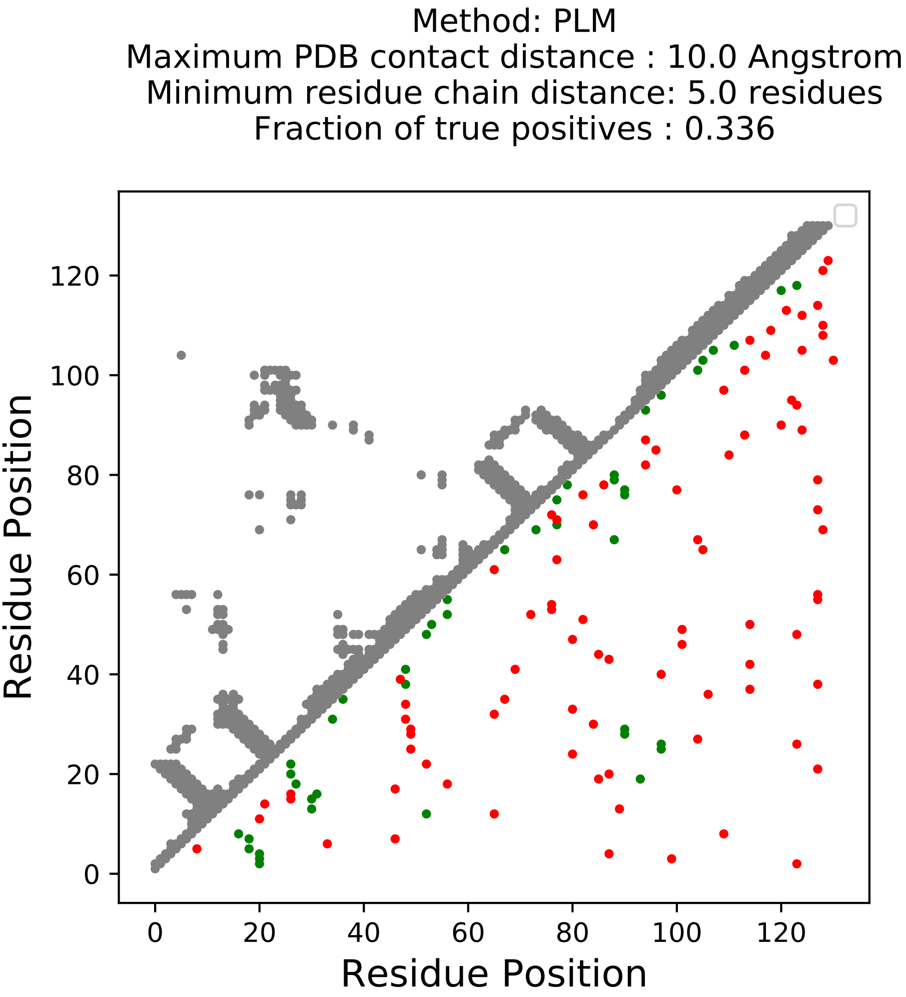

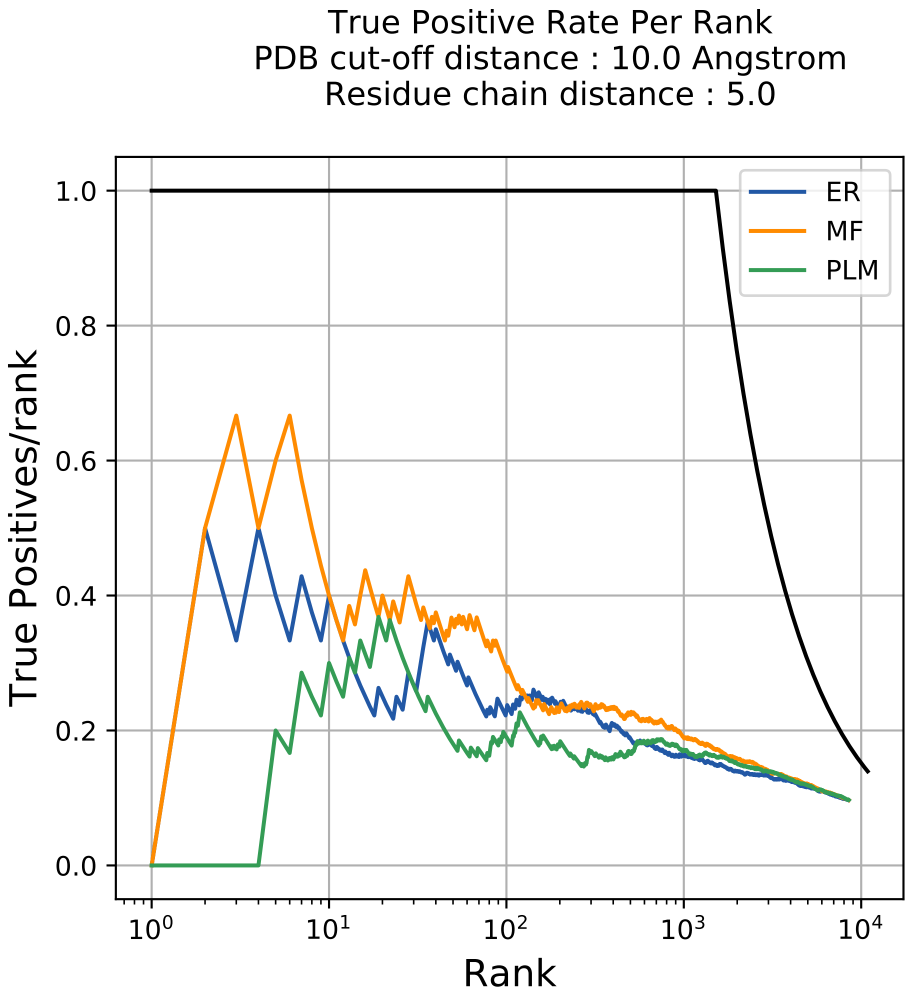

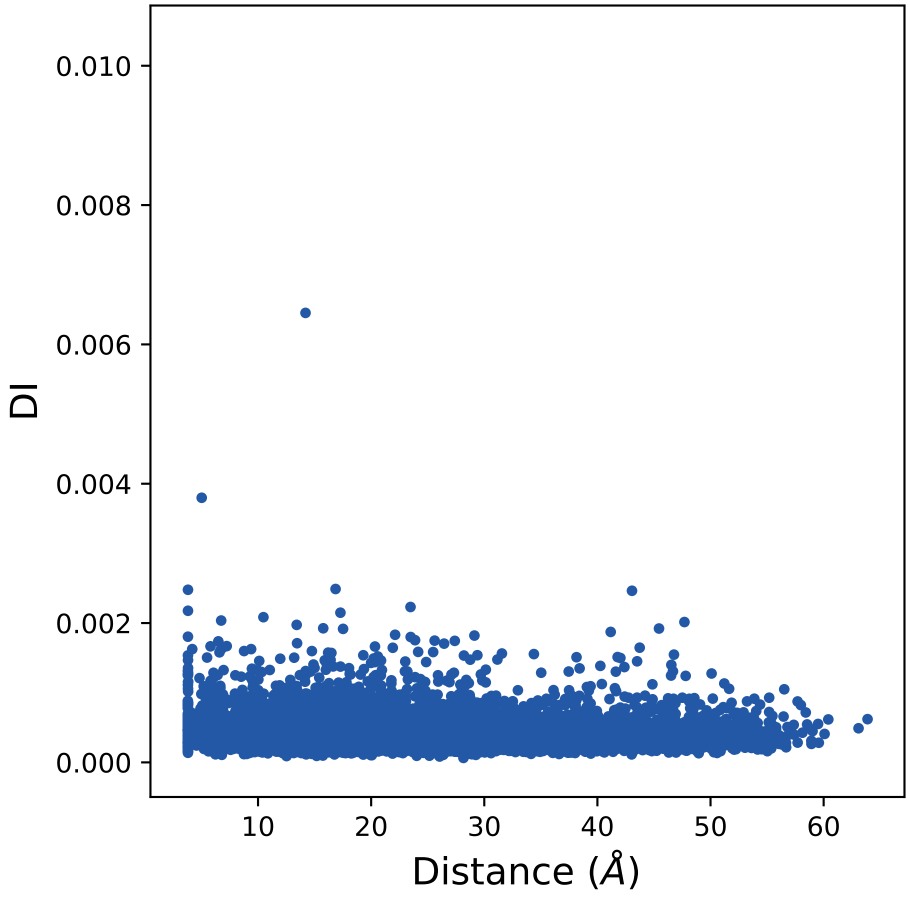

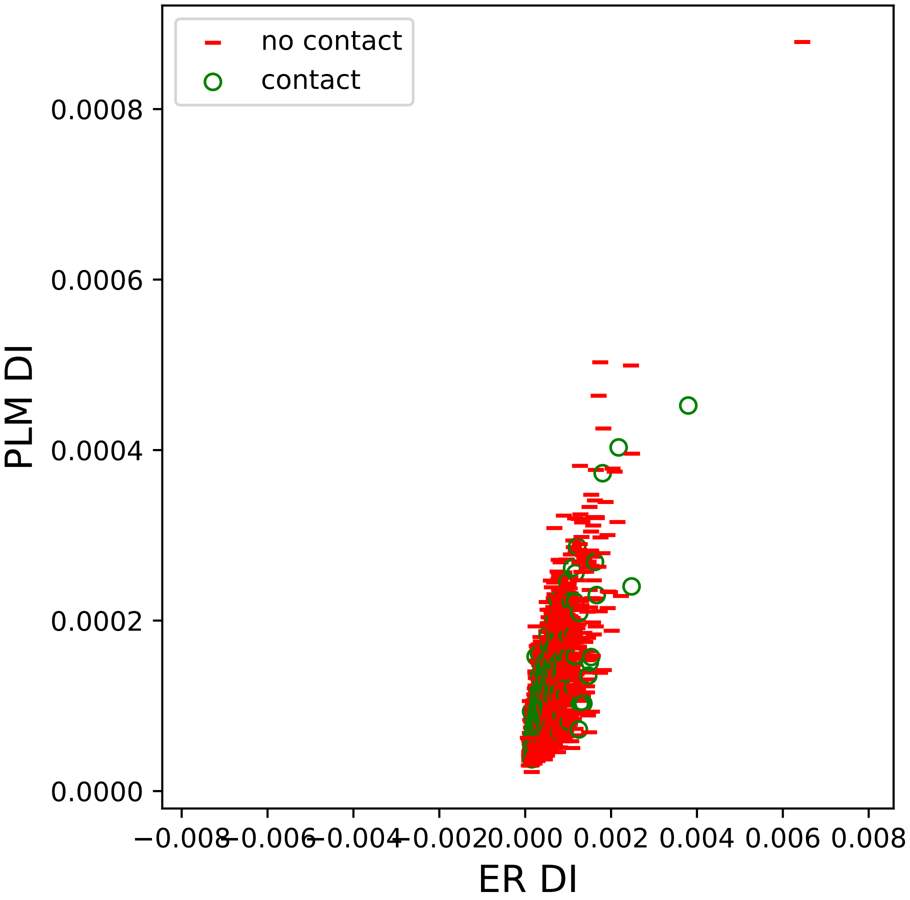

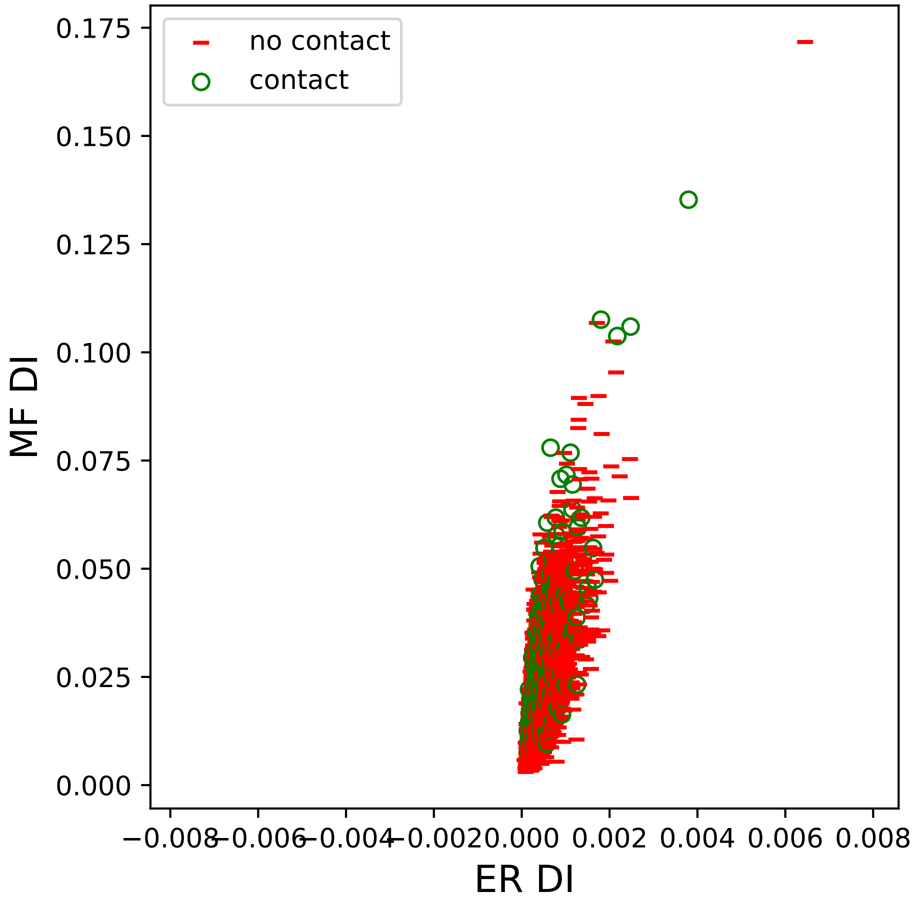

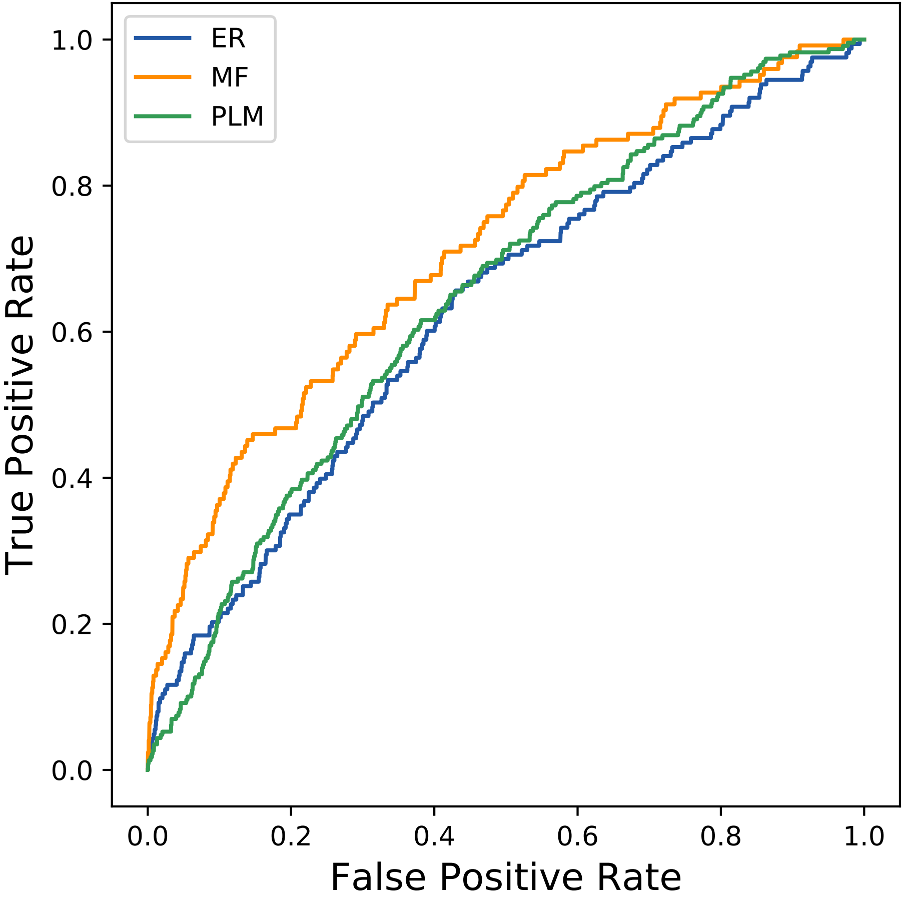

### 176seq_37col_6a6b_PF01387_plots.pdf

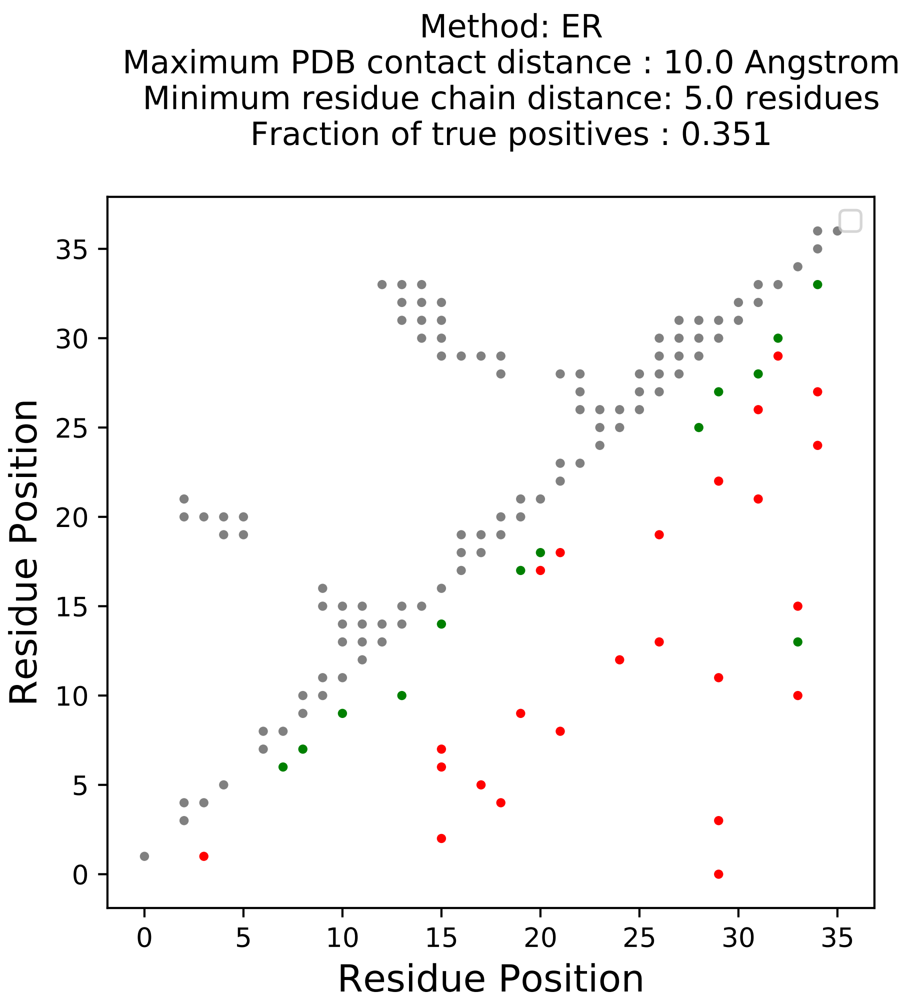

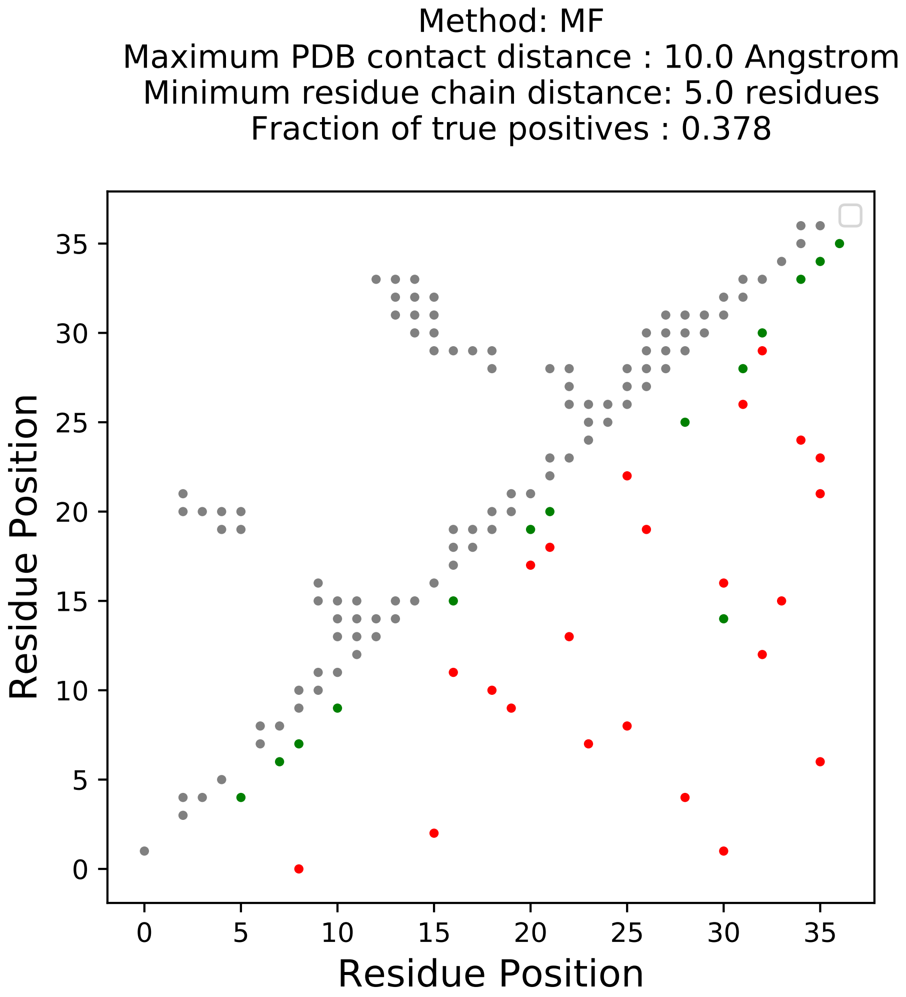

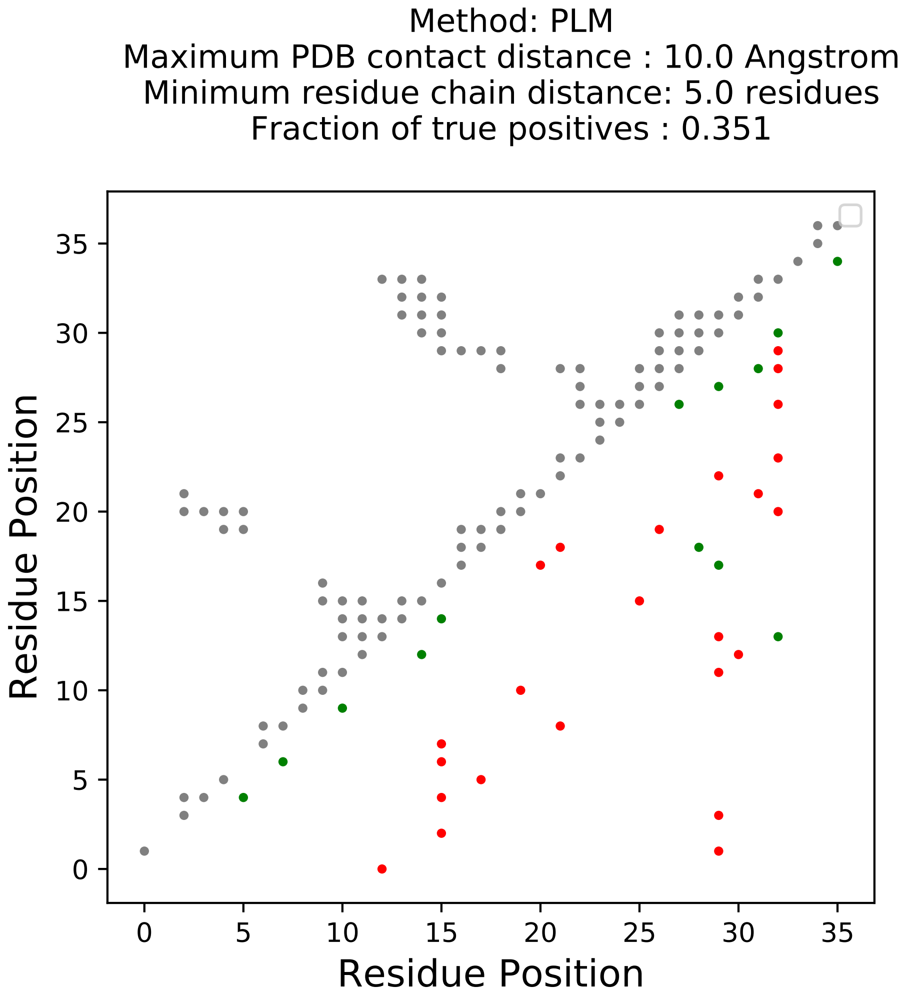

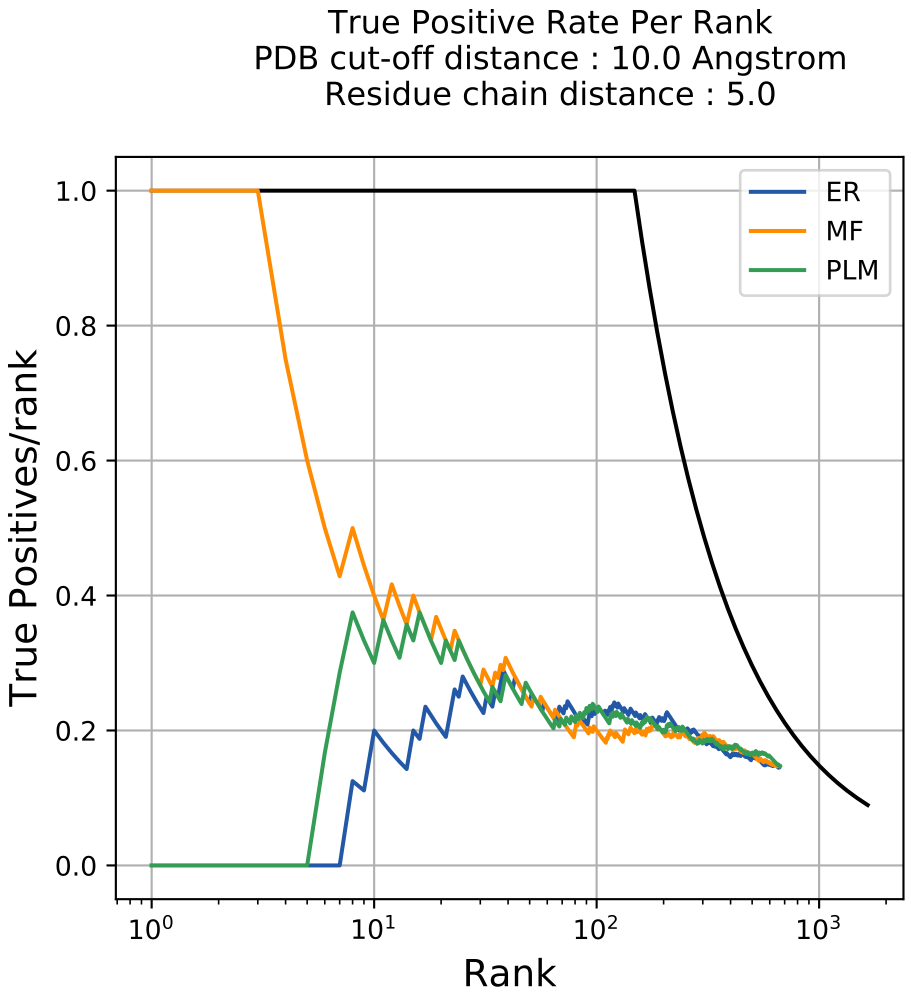

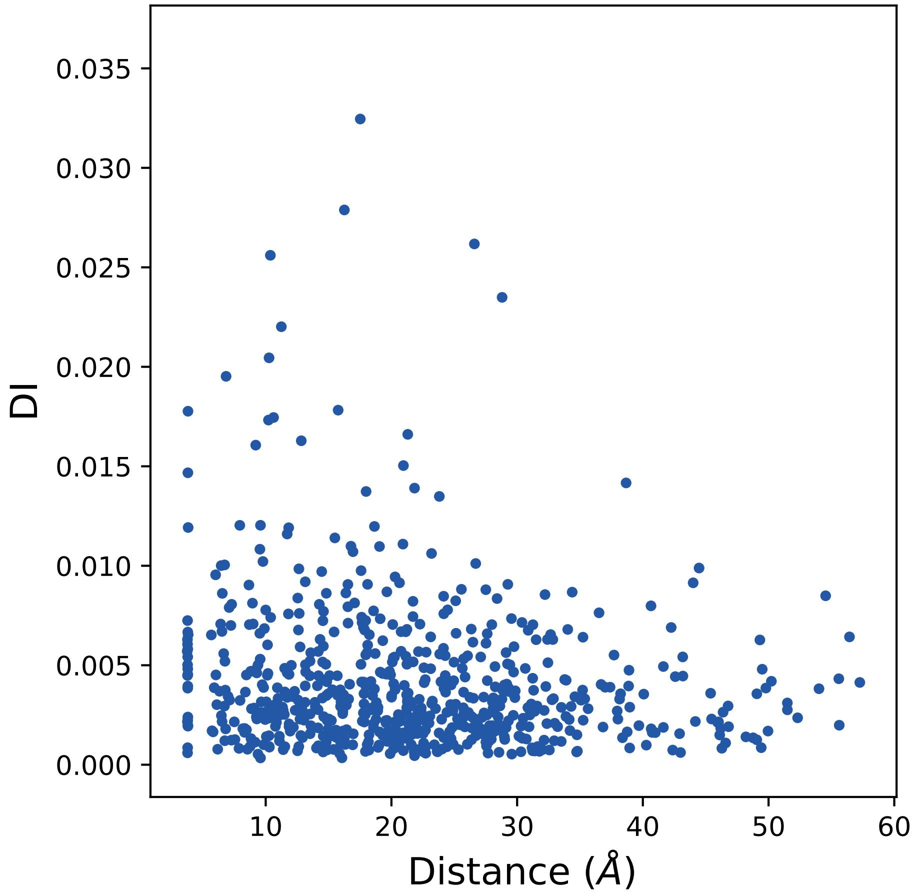

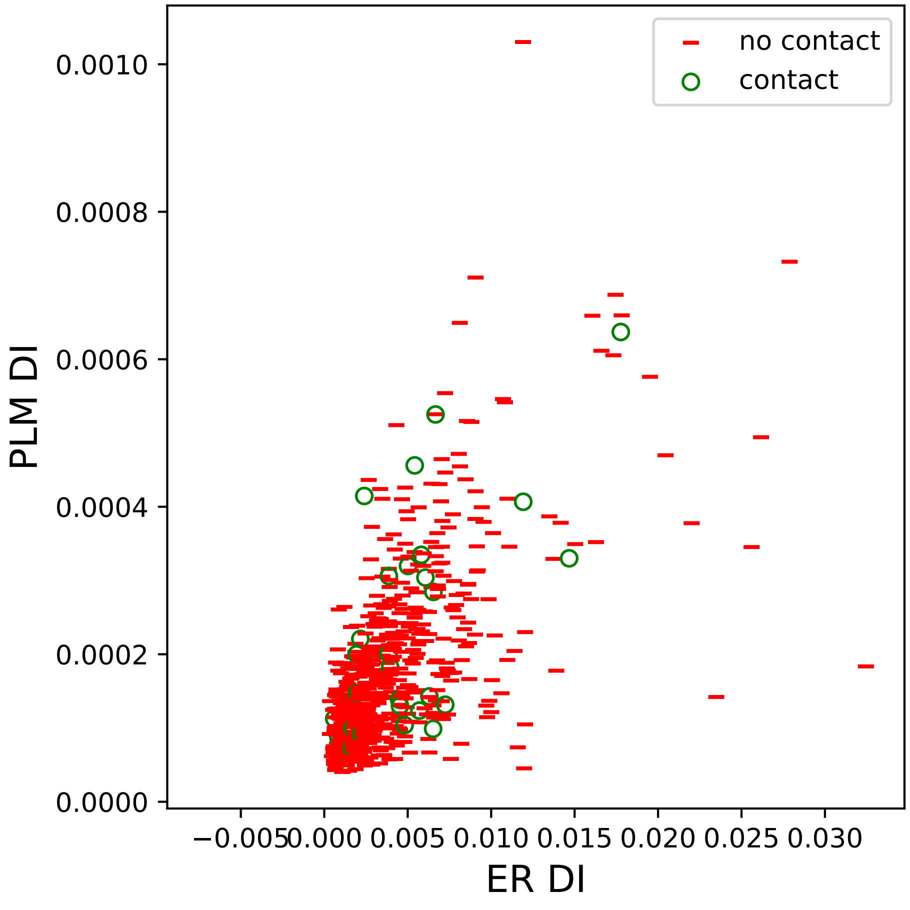

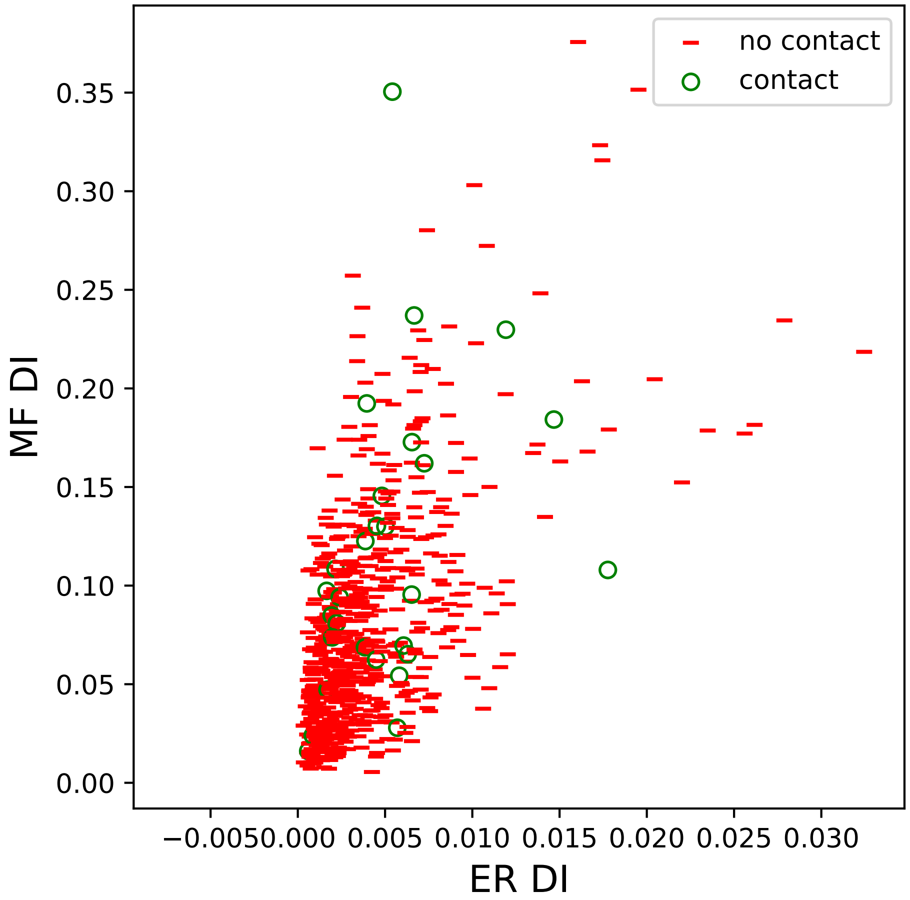

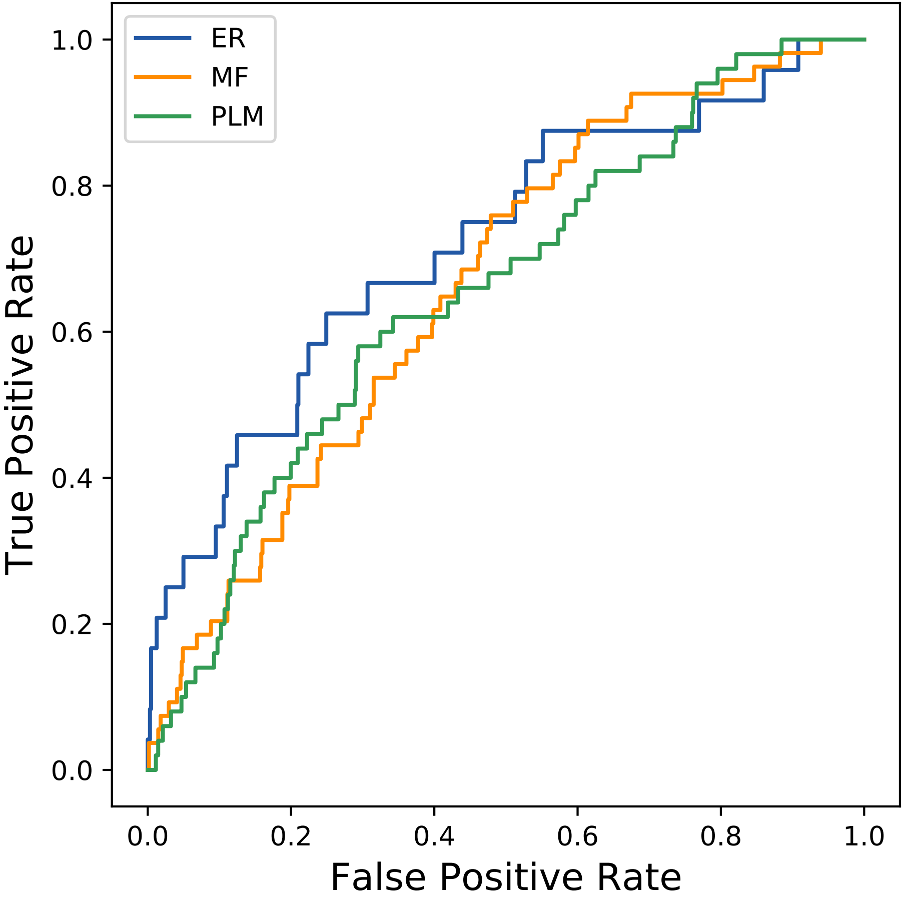

### 309seq_78col_6auo_PF01382_plots.pdf

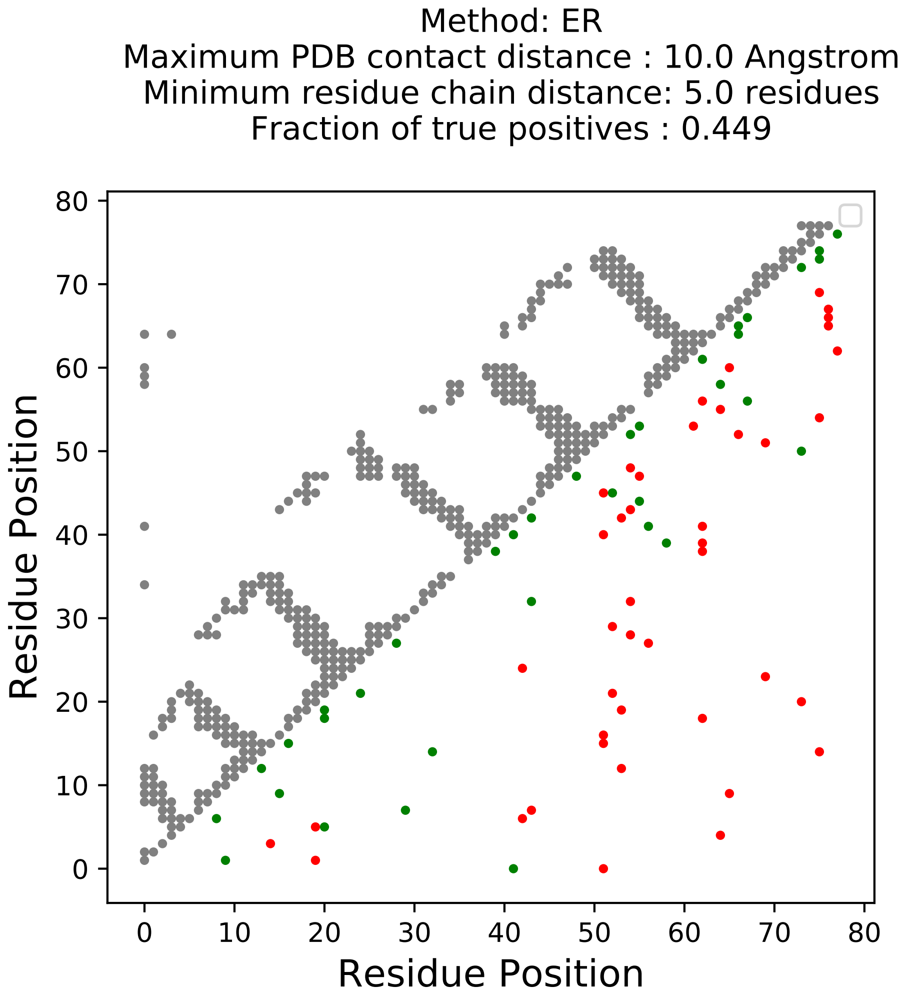

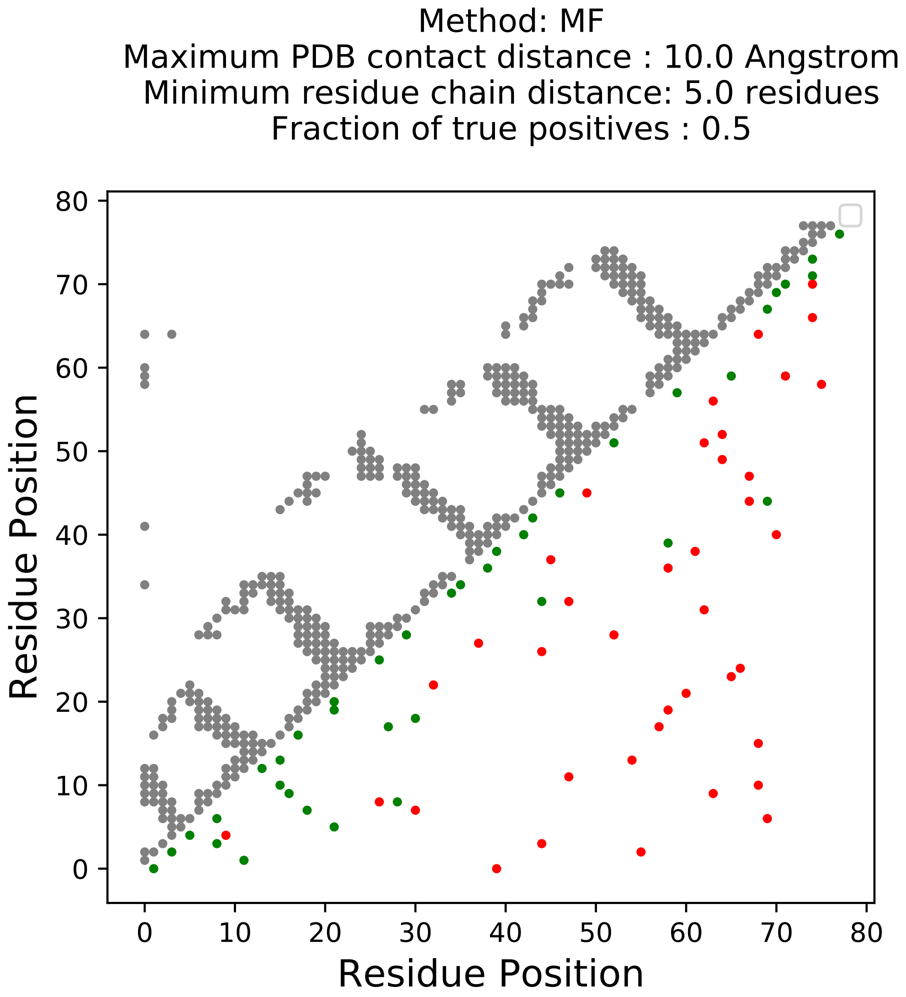

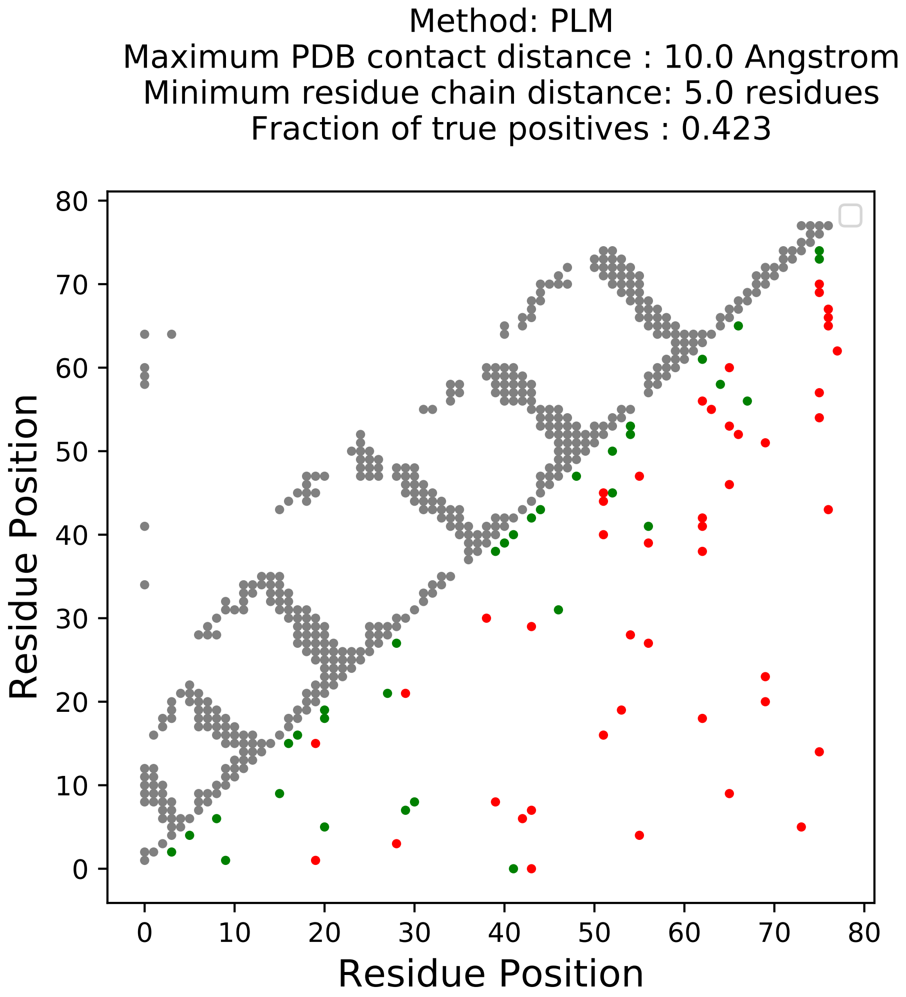

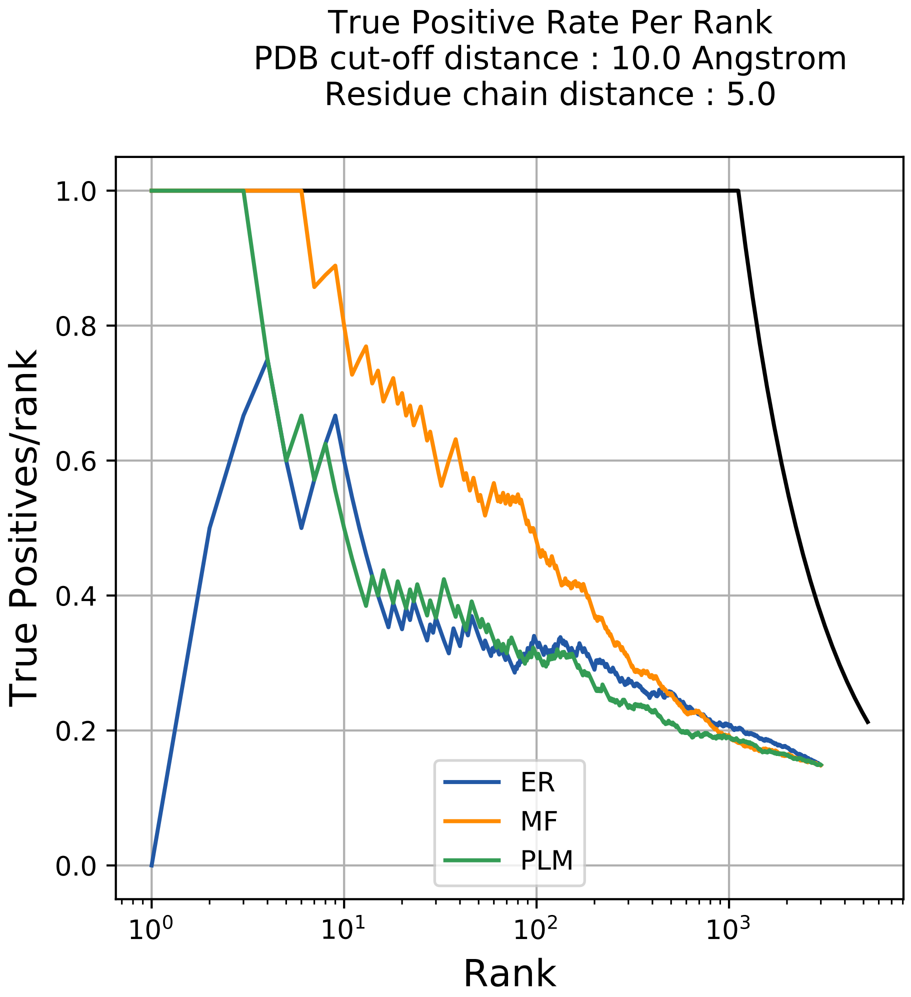

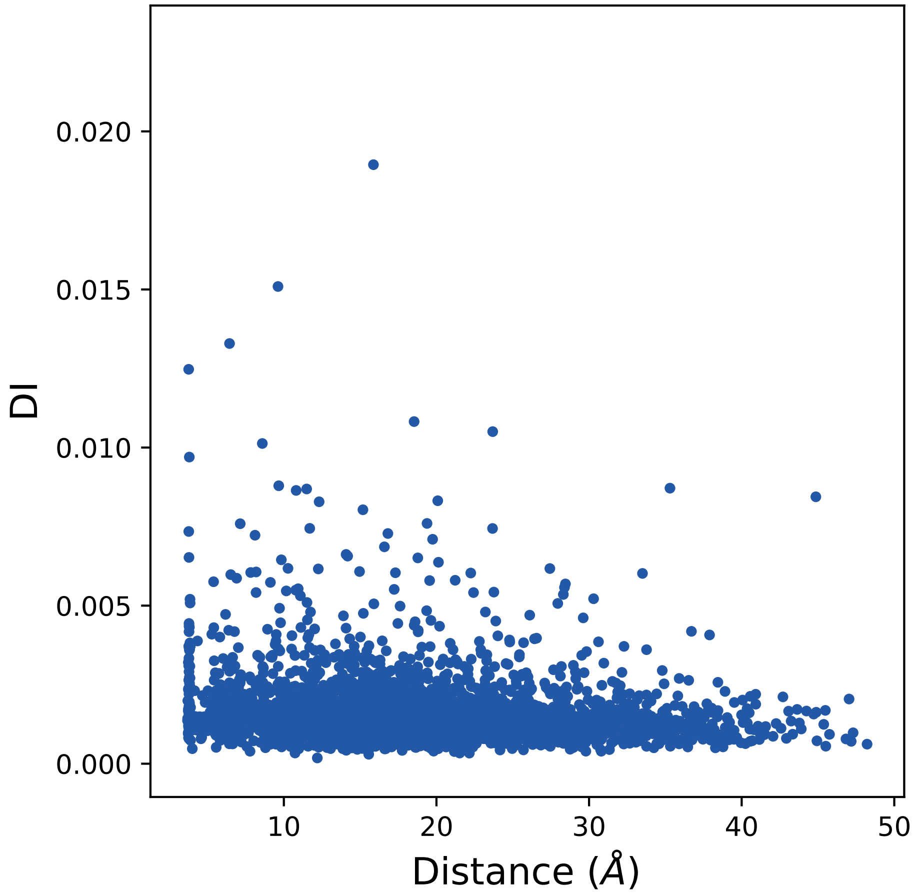

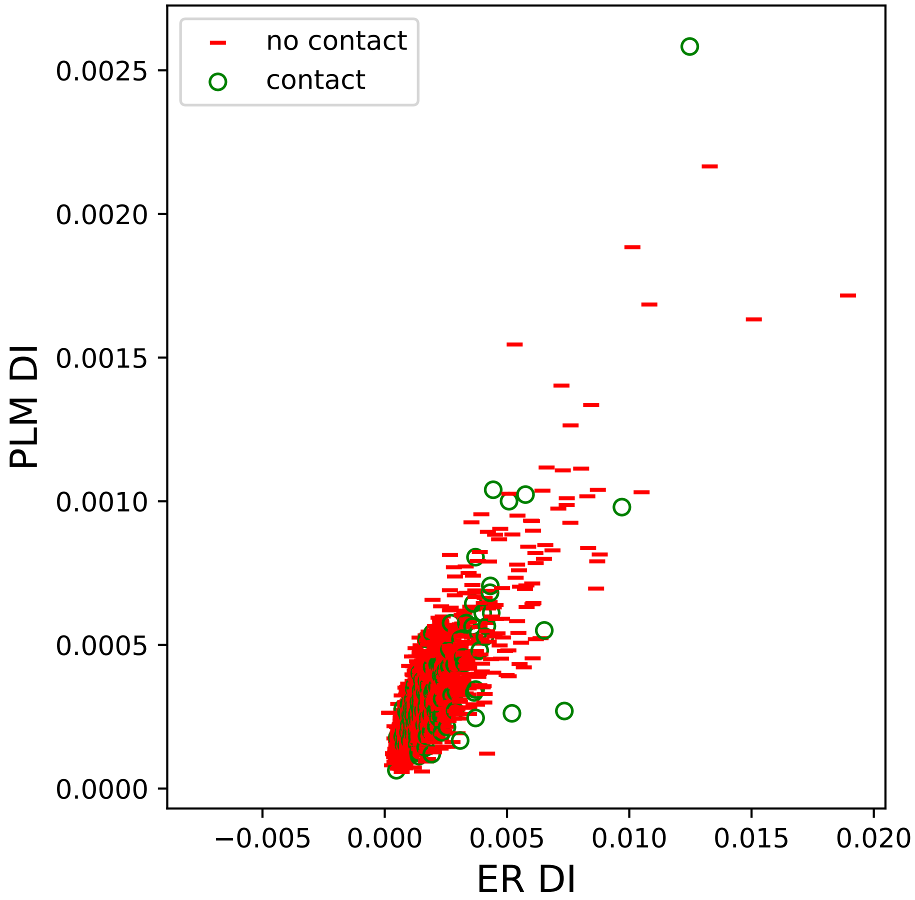

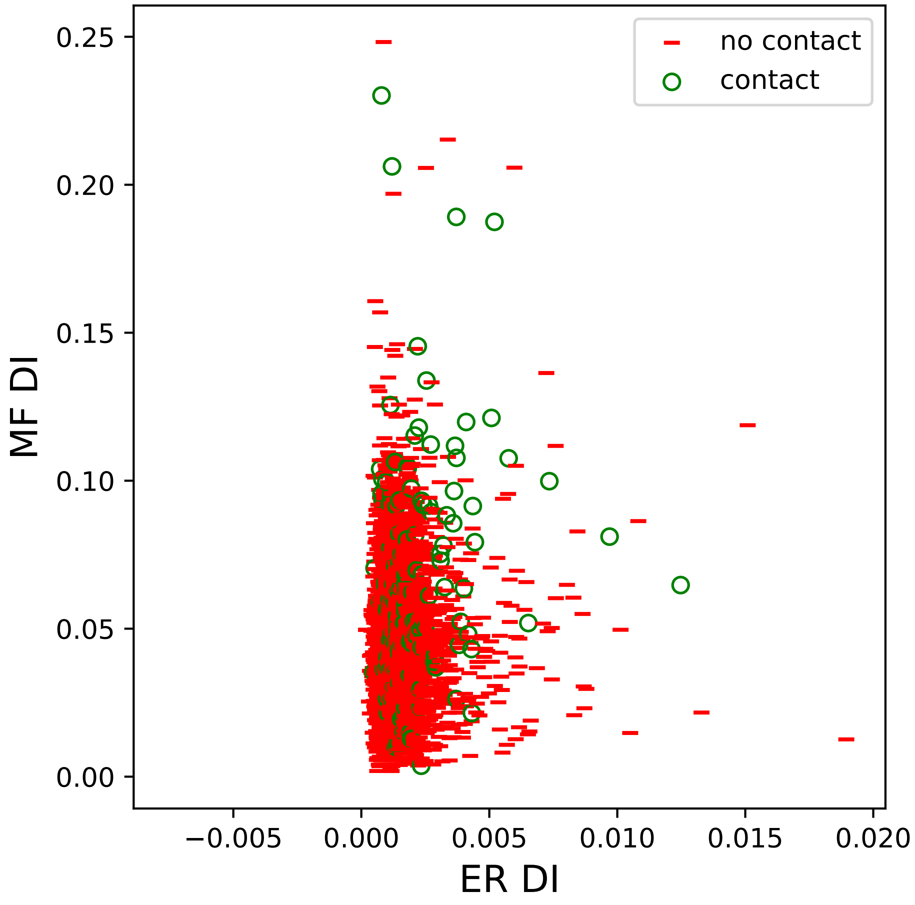

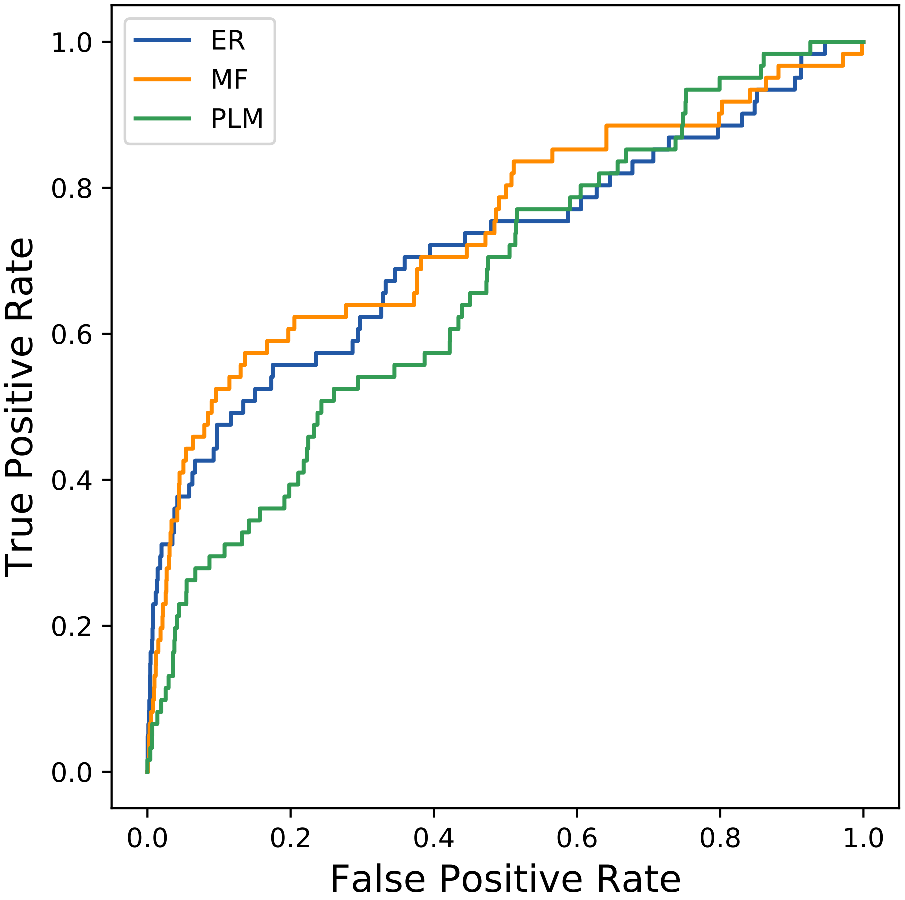

### 328seq_57col_6aeq_PF17948_plots.pdf

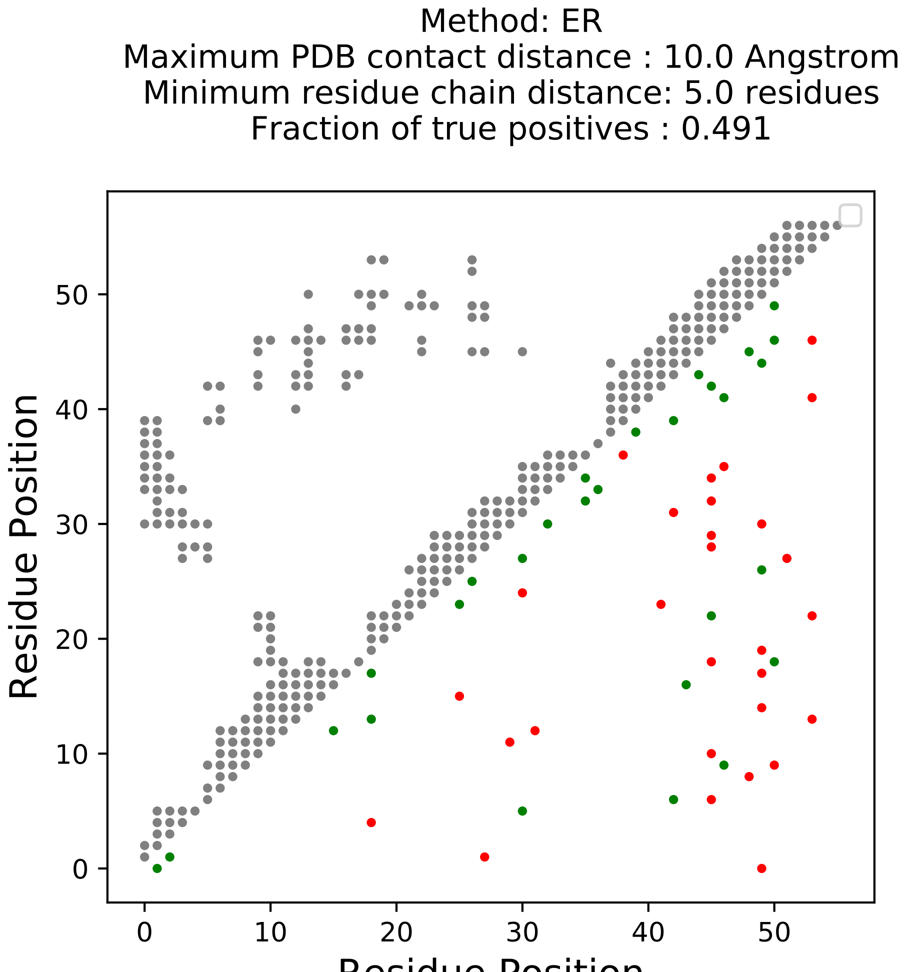

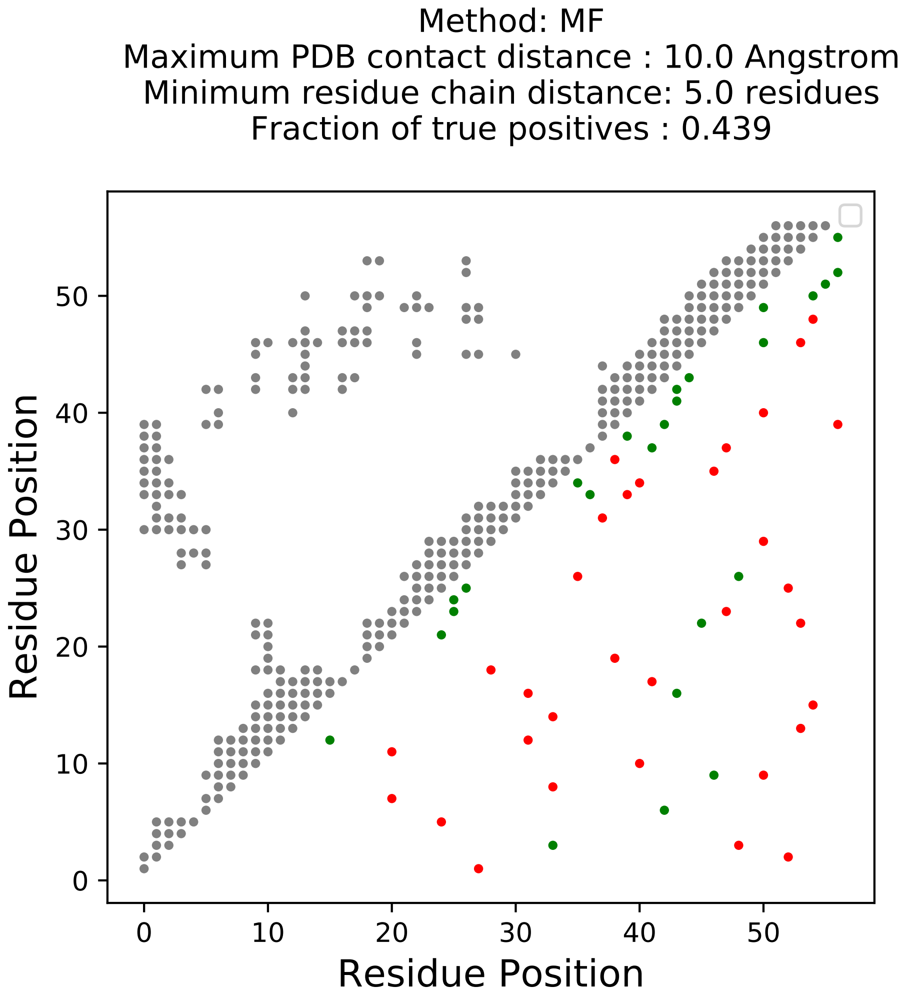

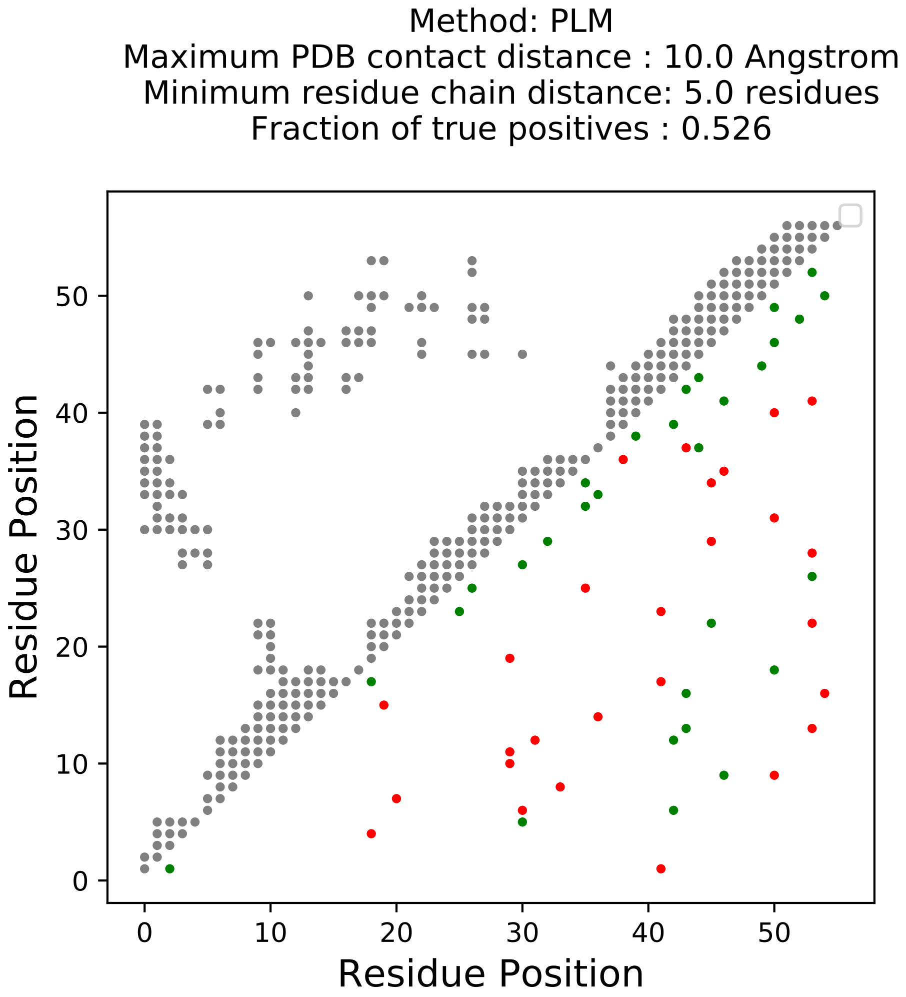

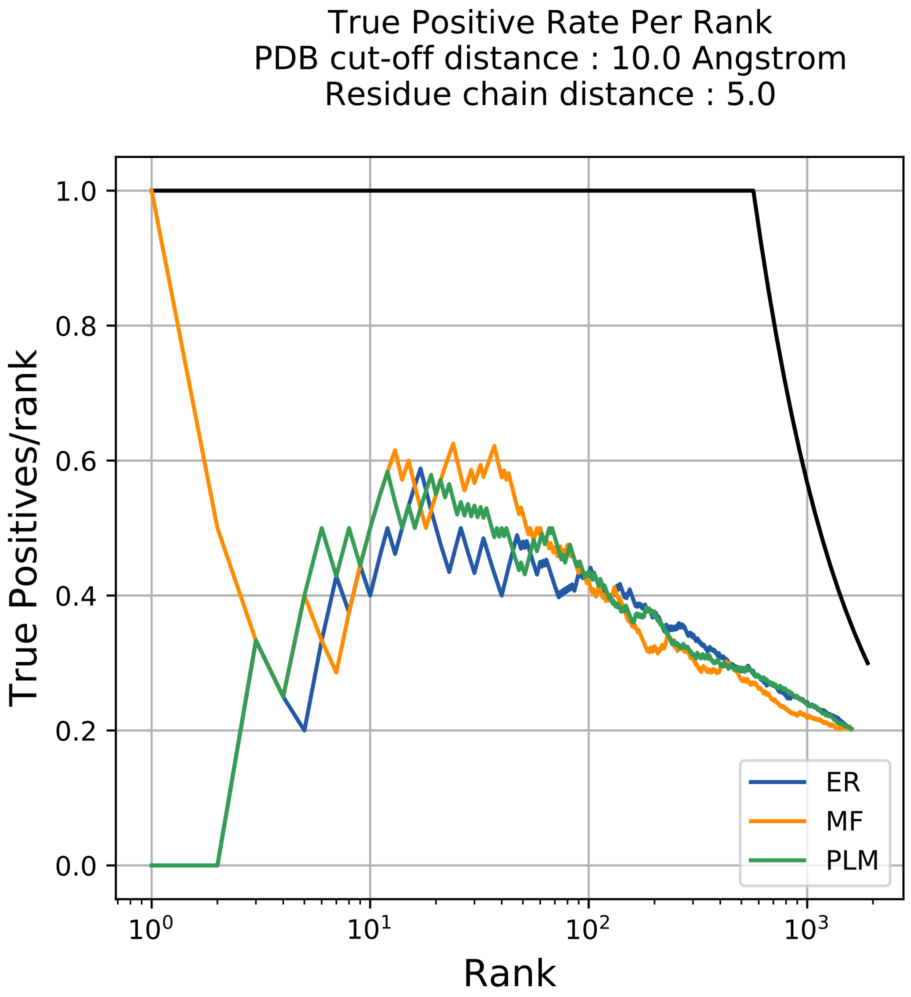

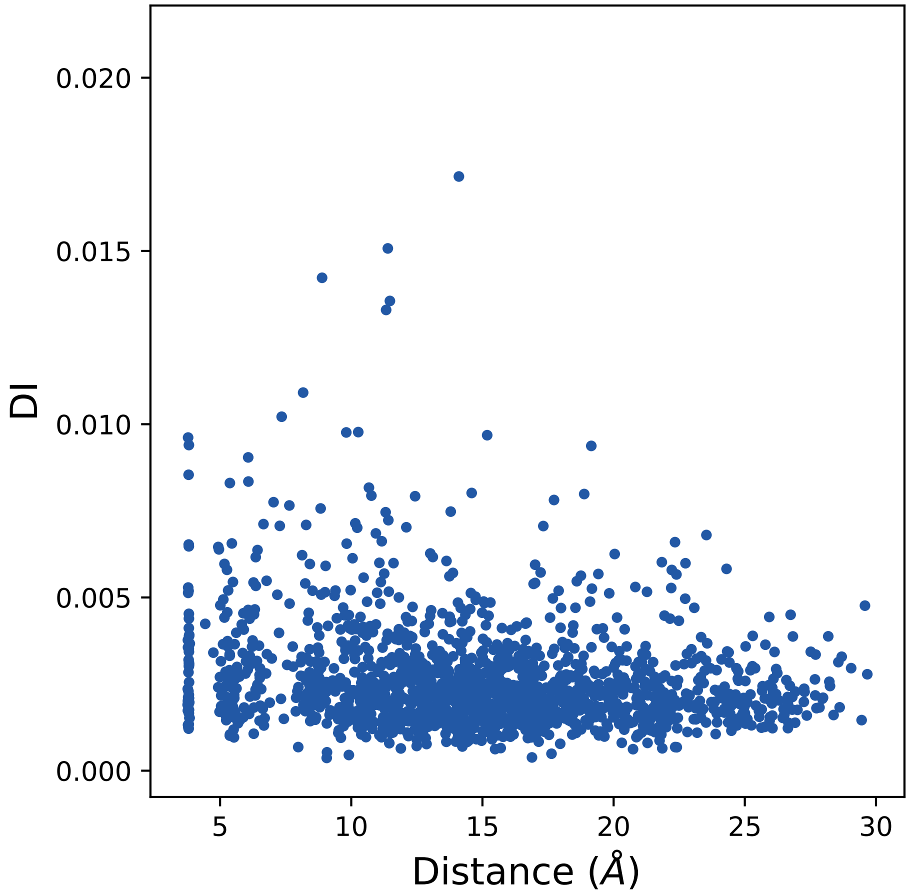

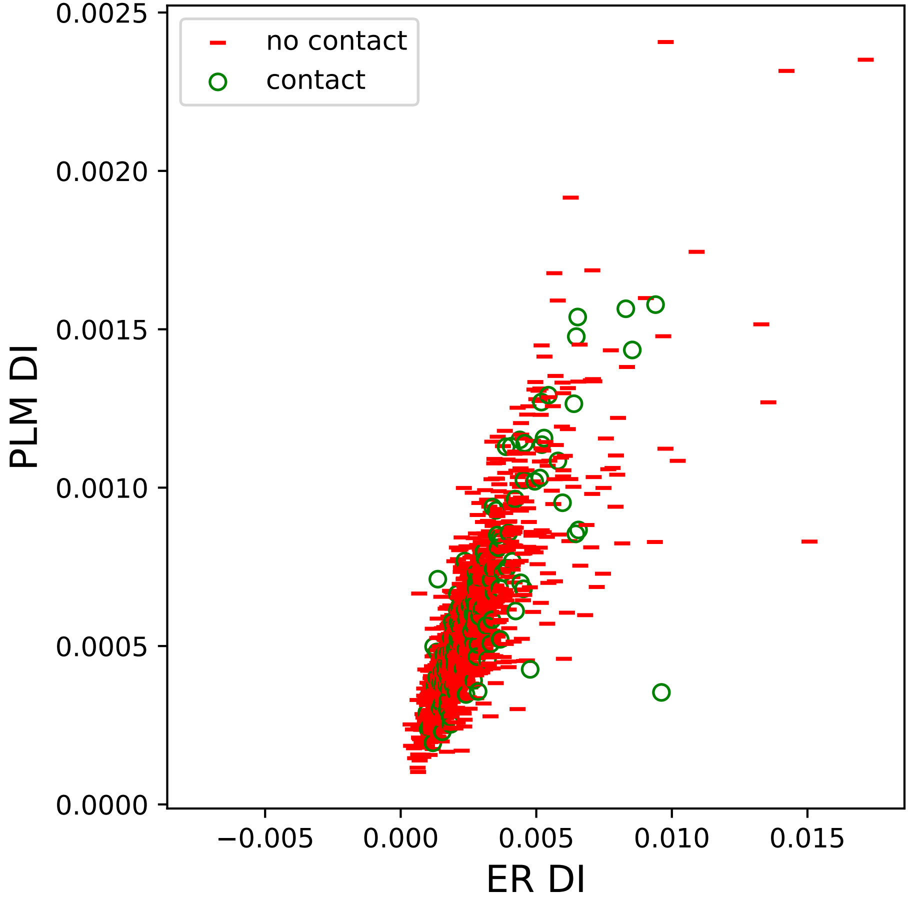

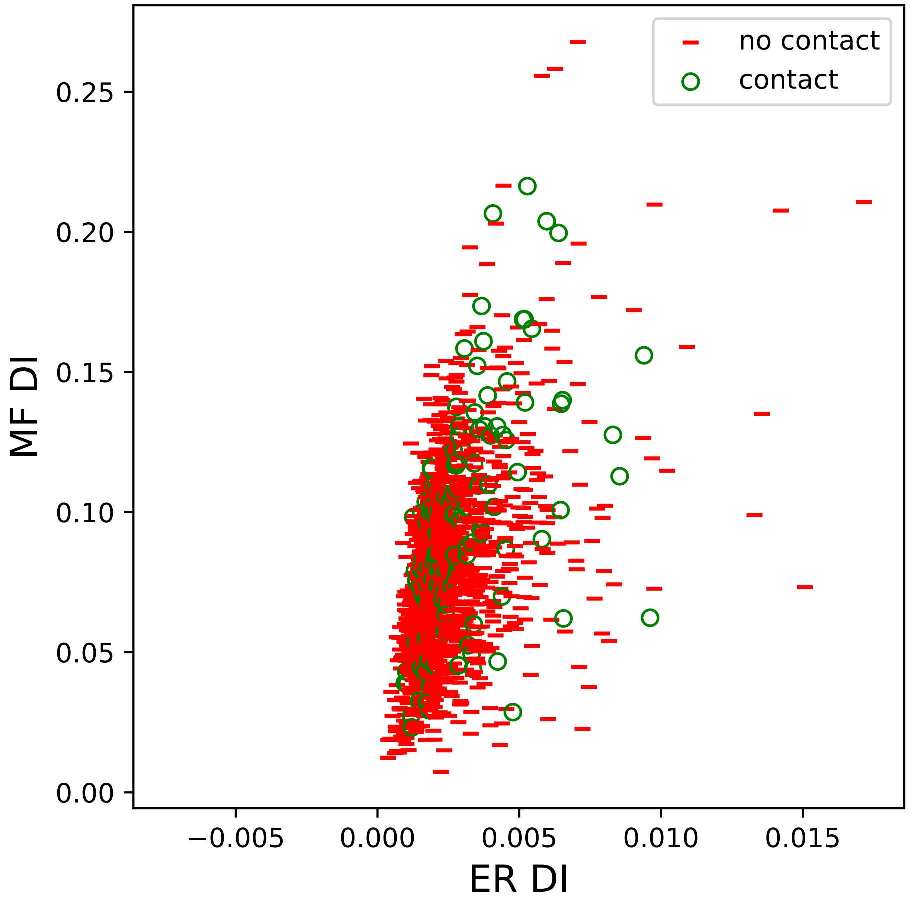

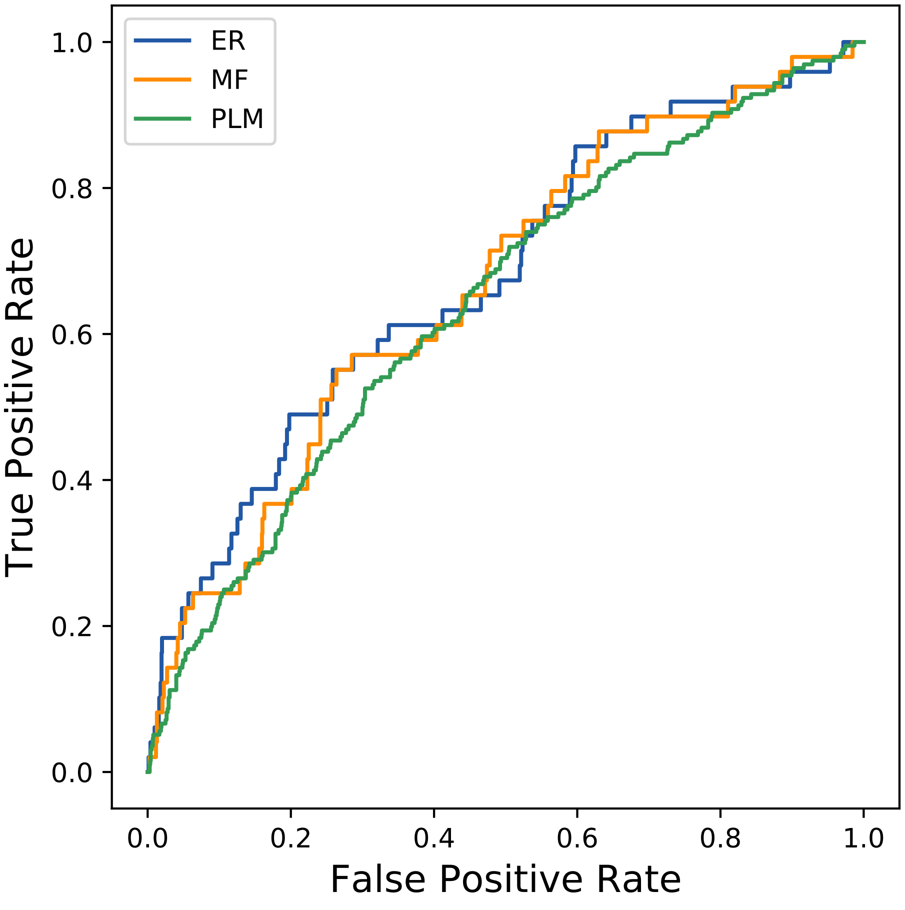

### 359seq_83col_2zh1_PF09249_plots.pdf

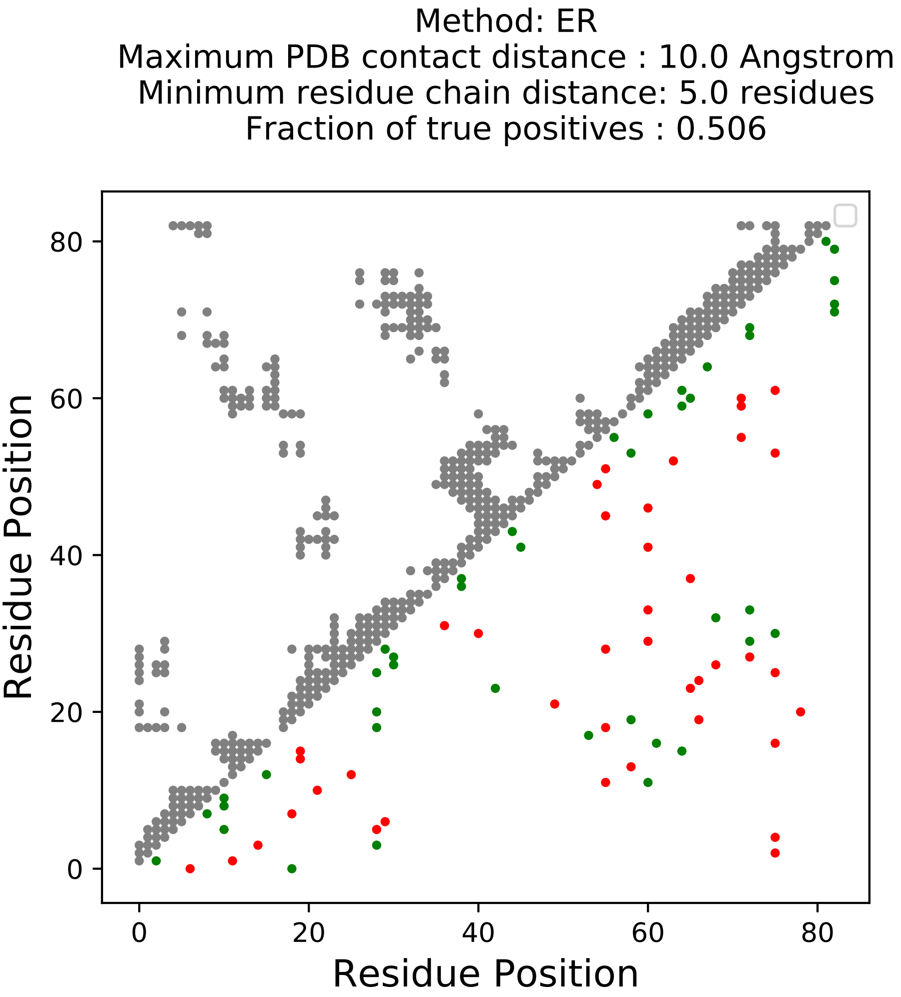

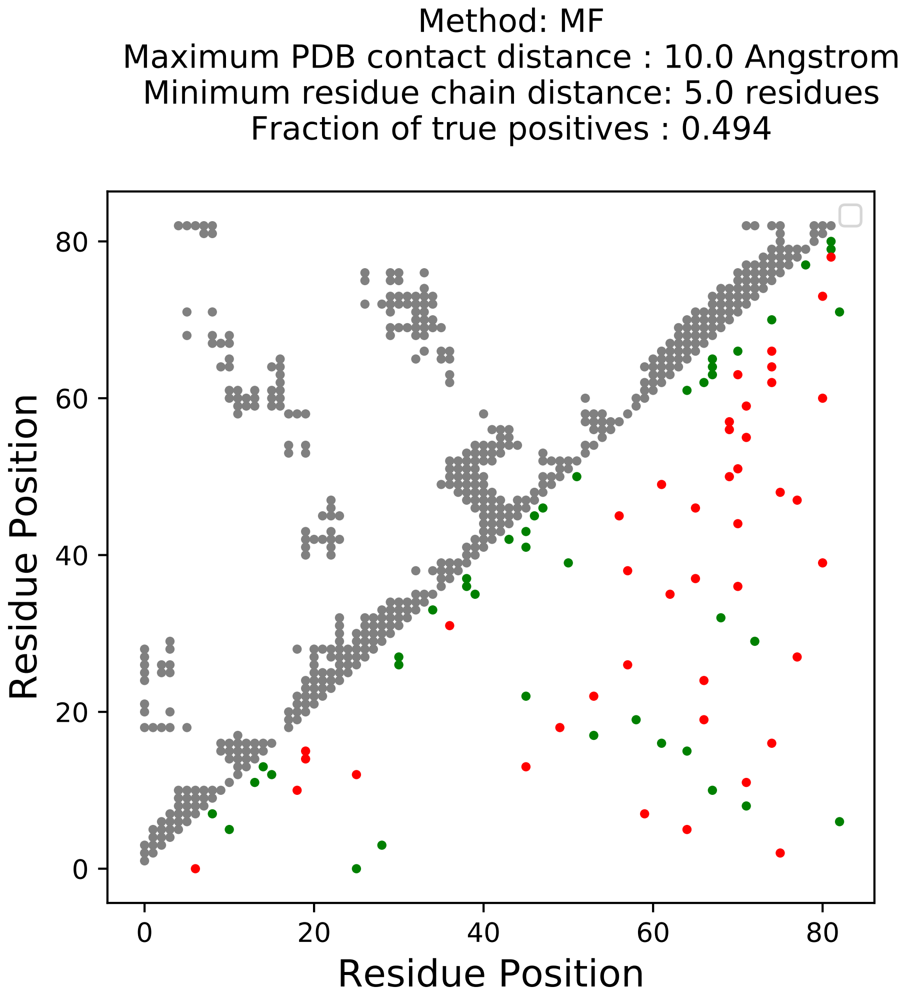

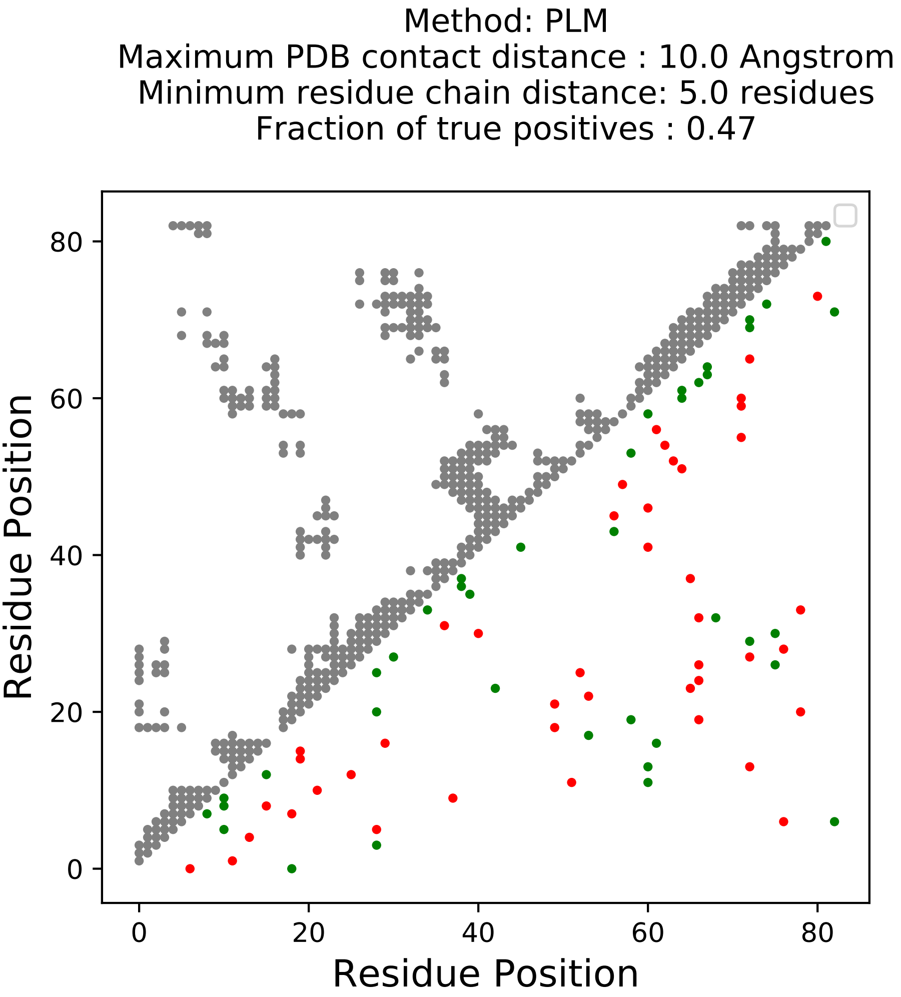

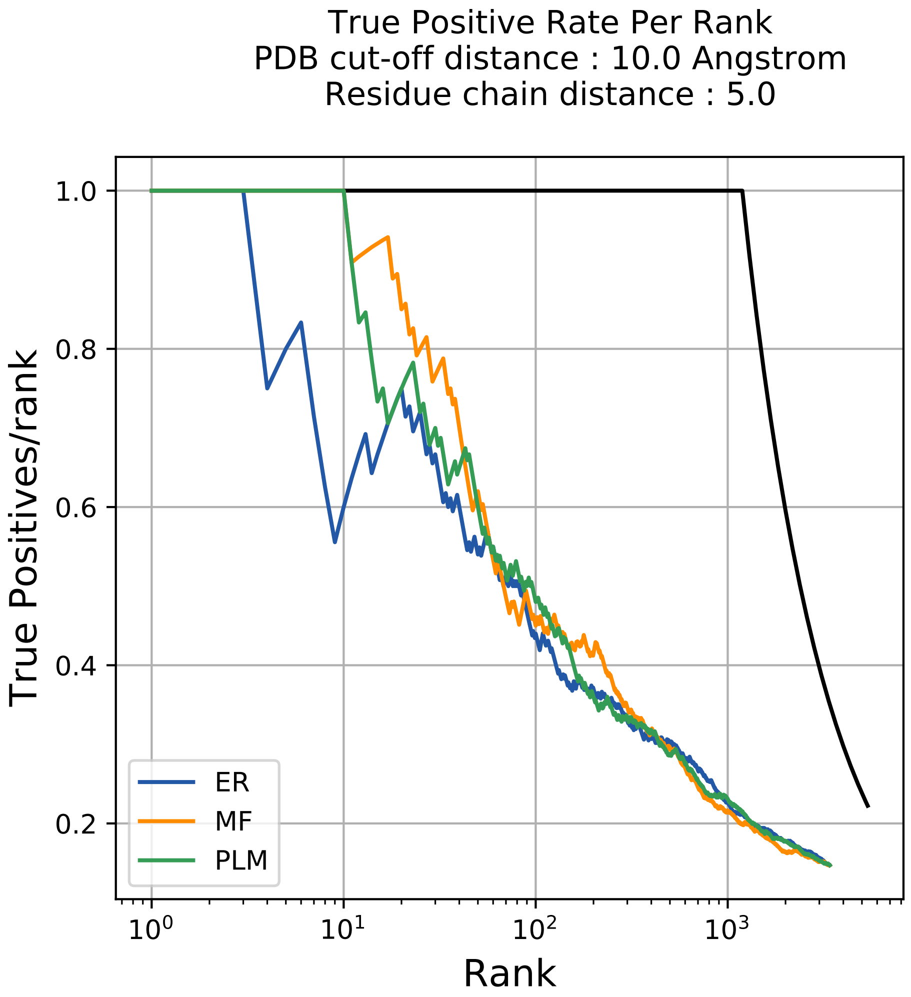

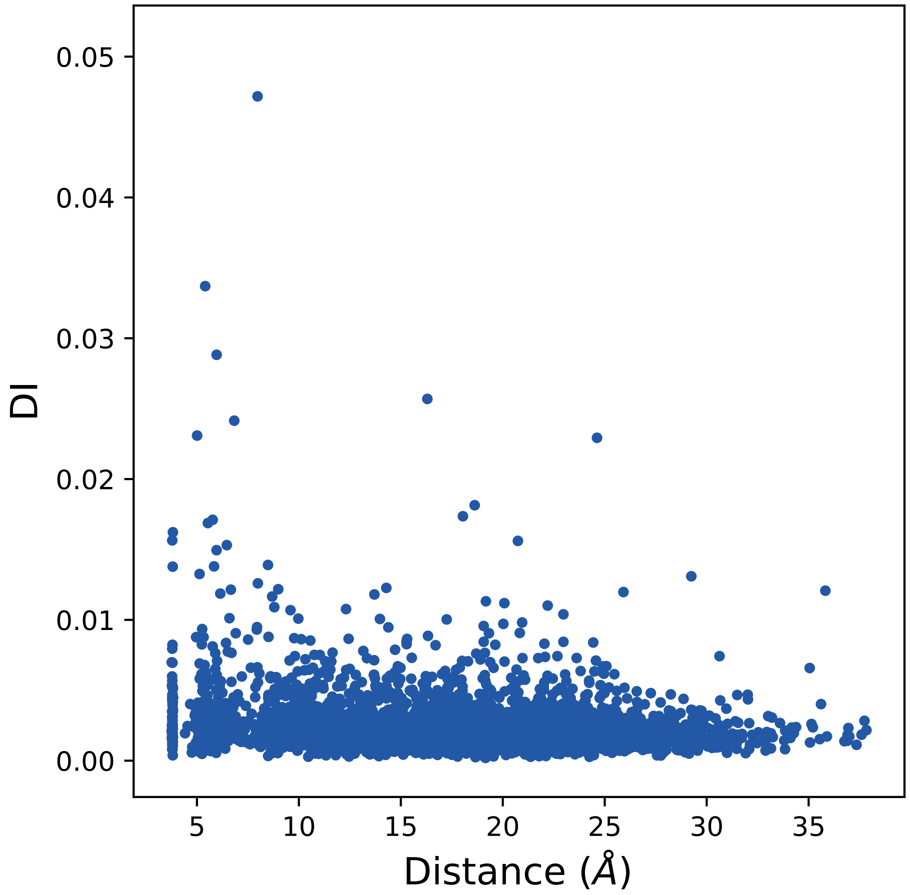

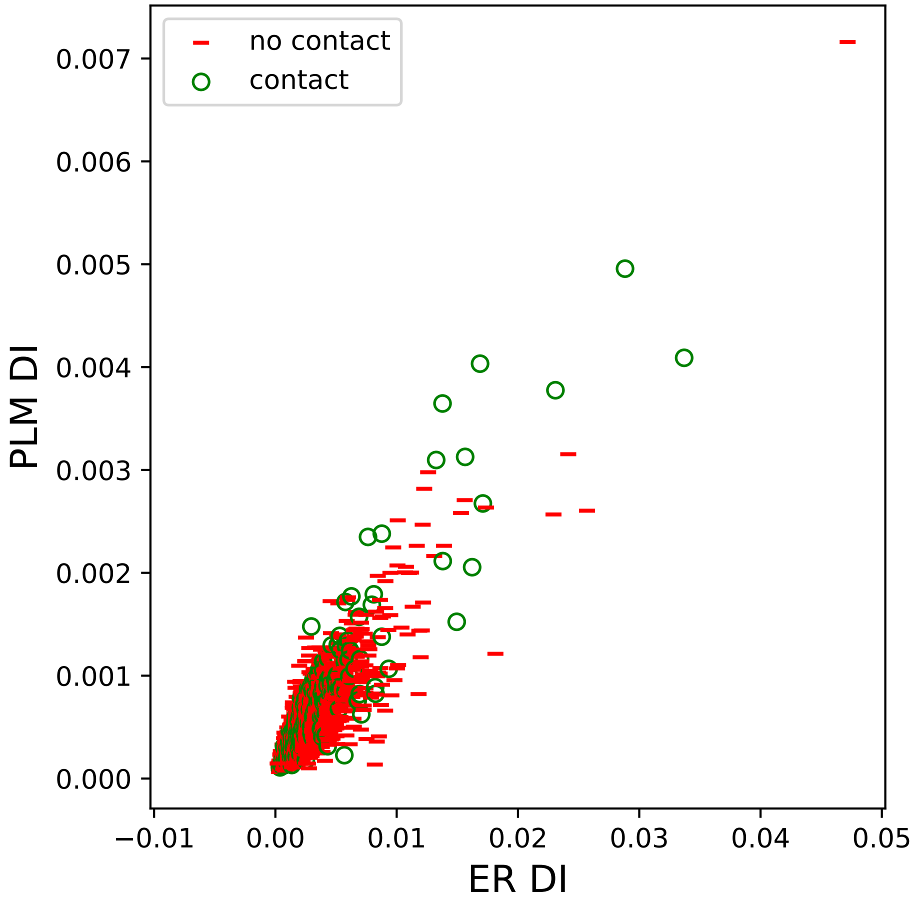

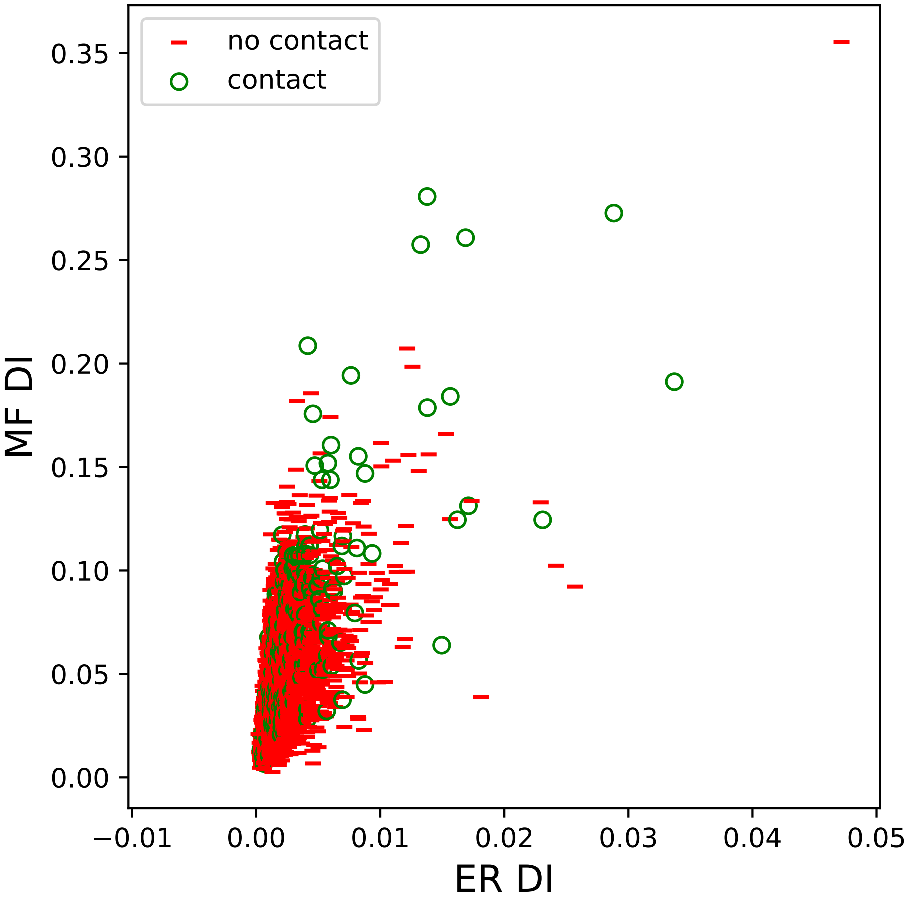

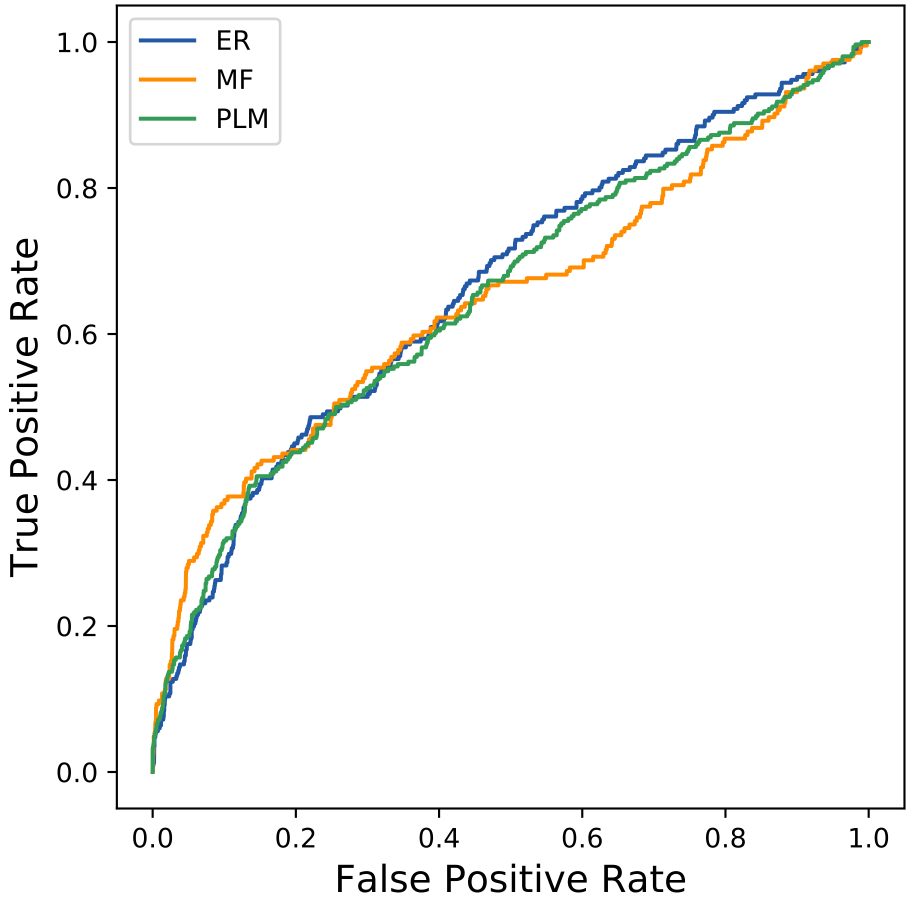

### 389seq_131col_4ilw_PF00965_plots.pdf

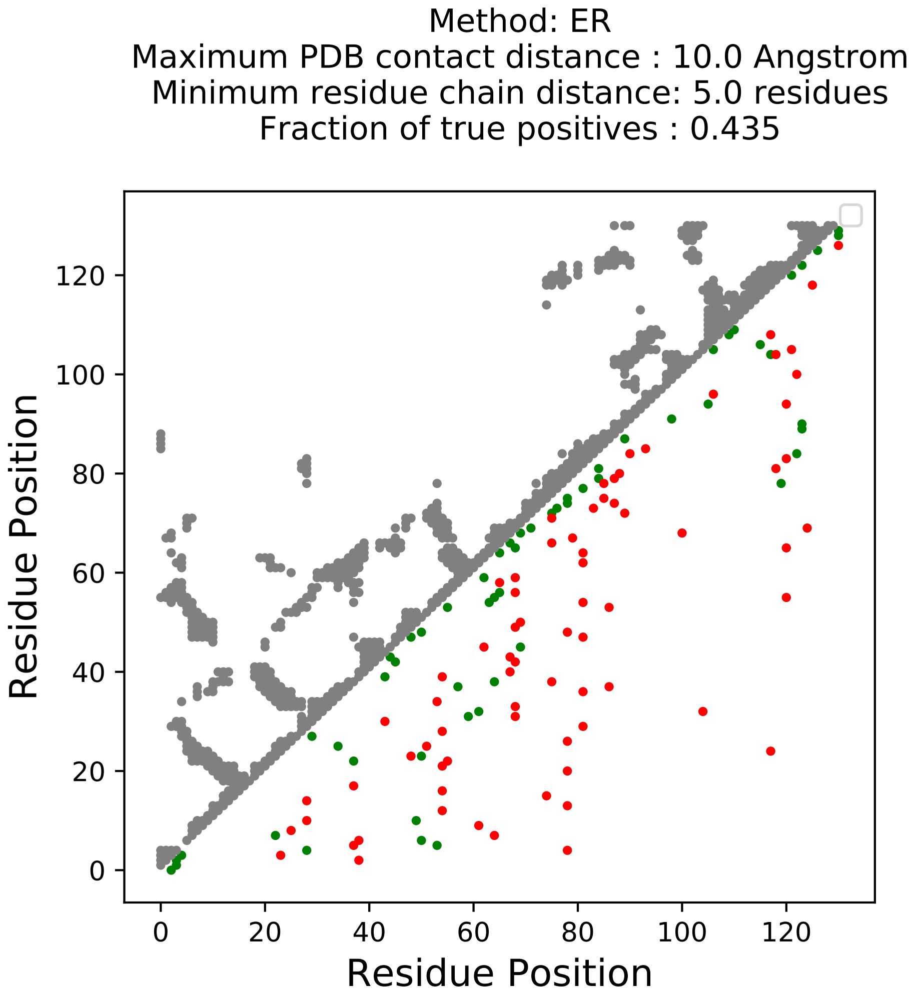

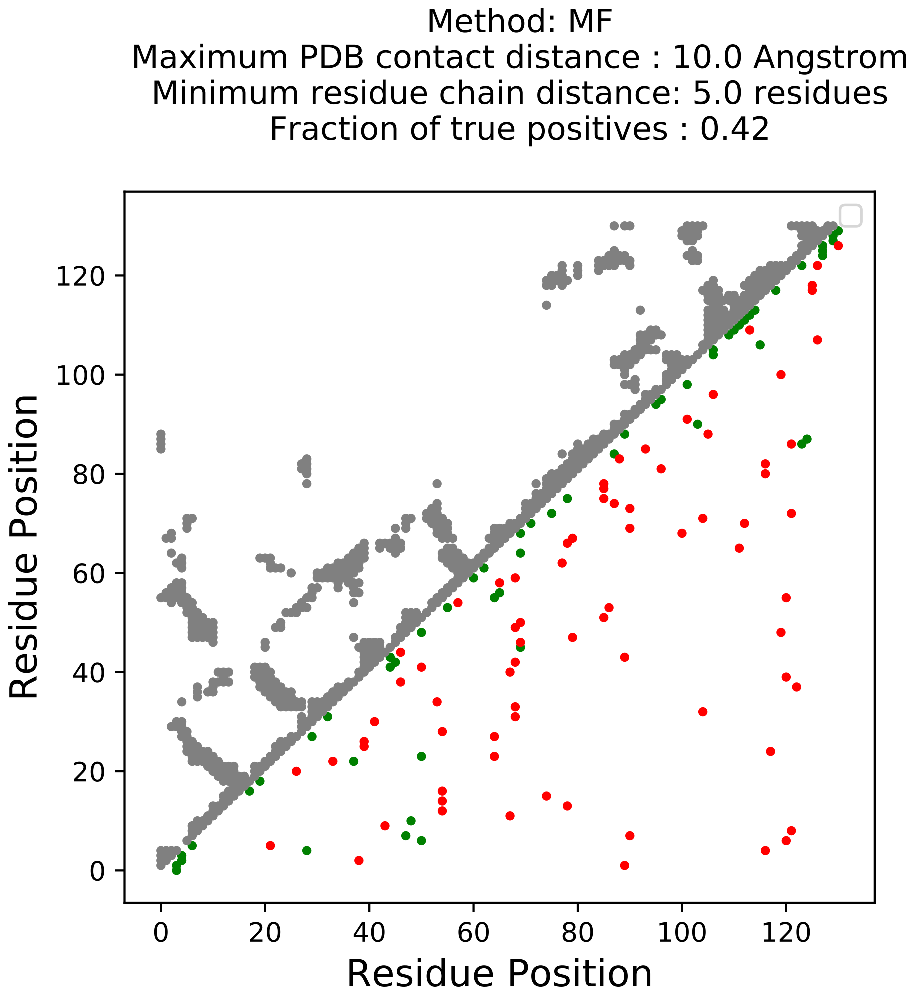

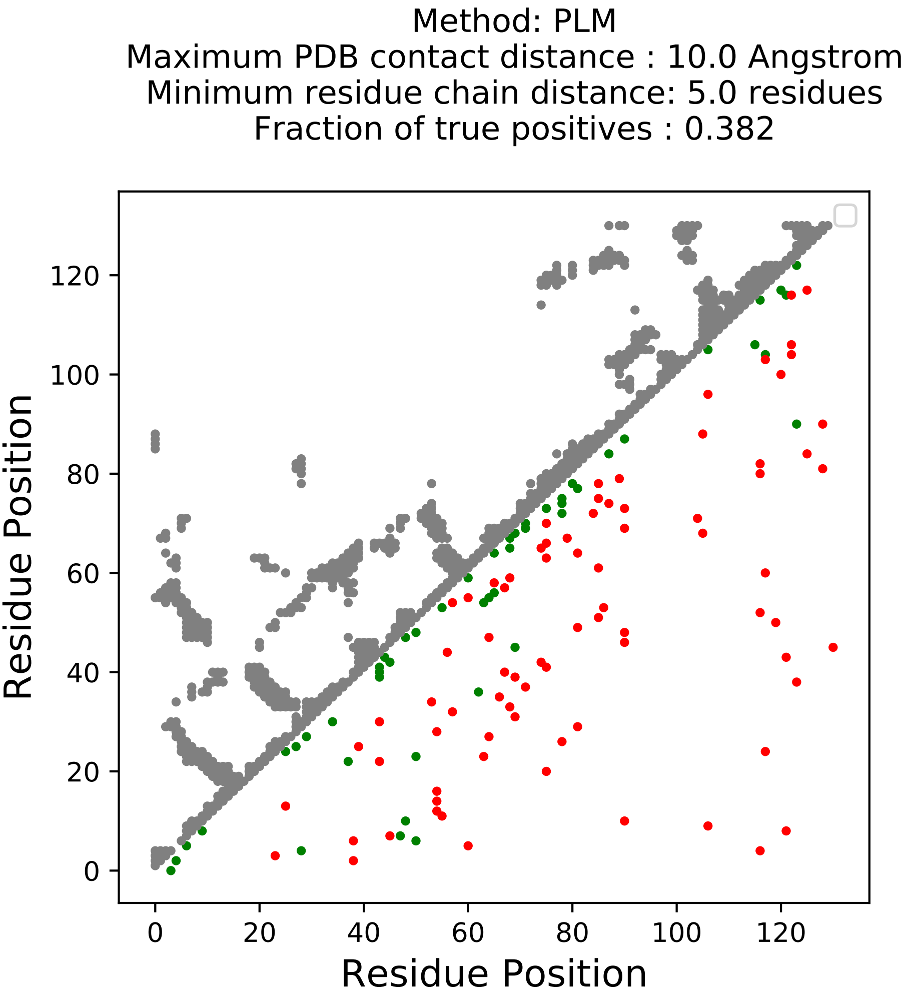

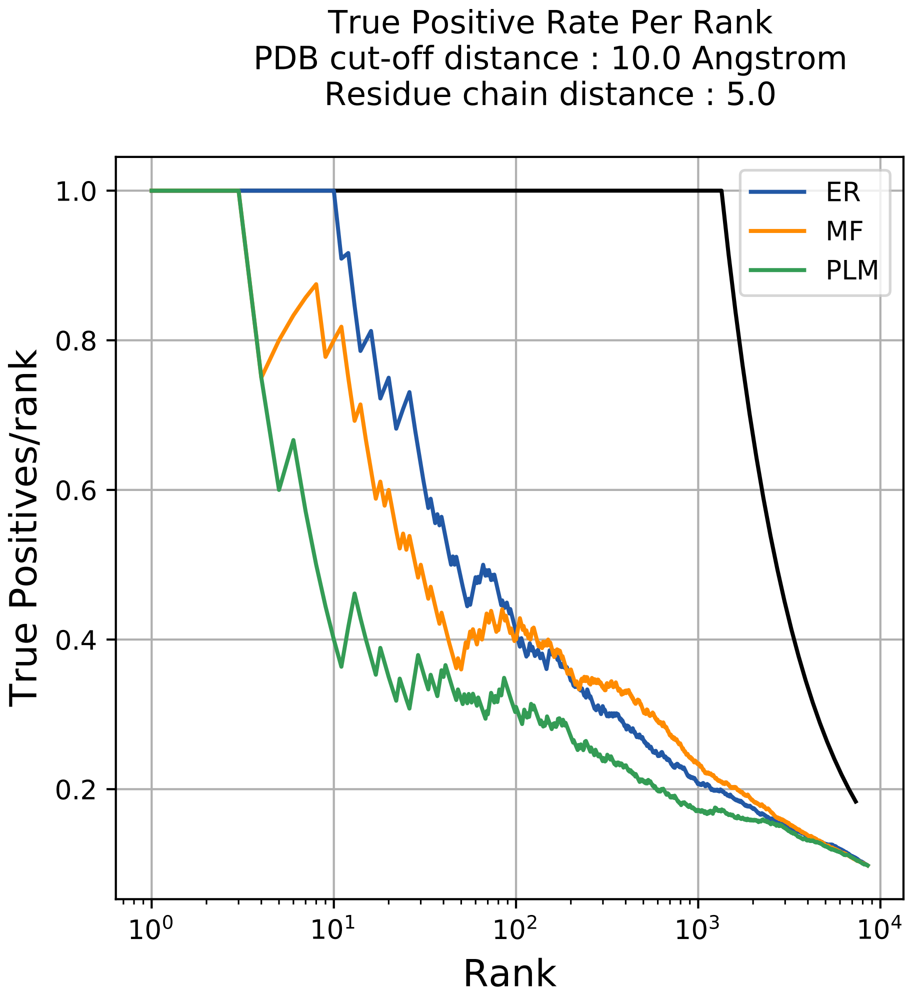

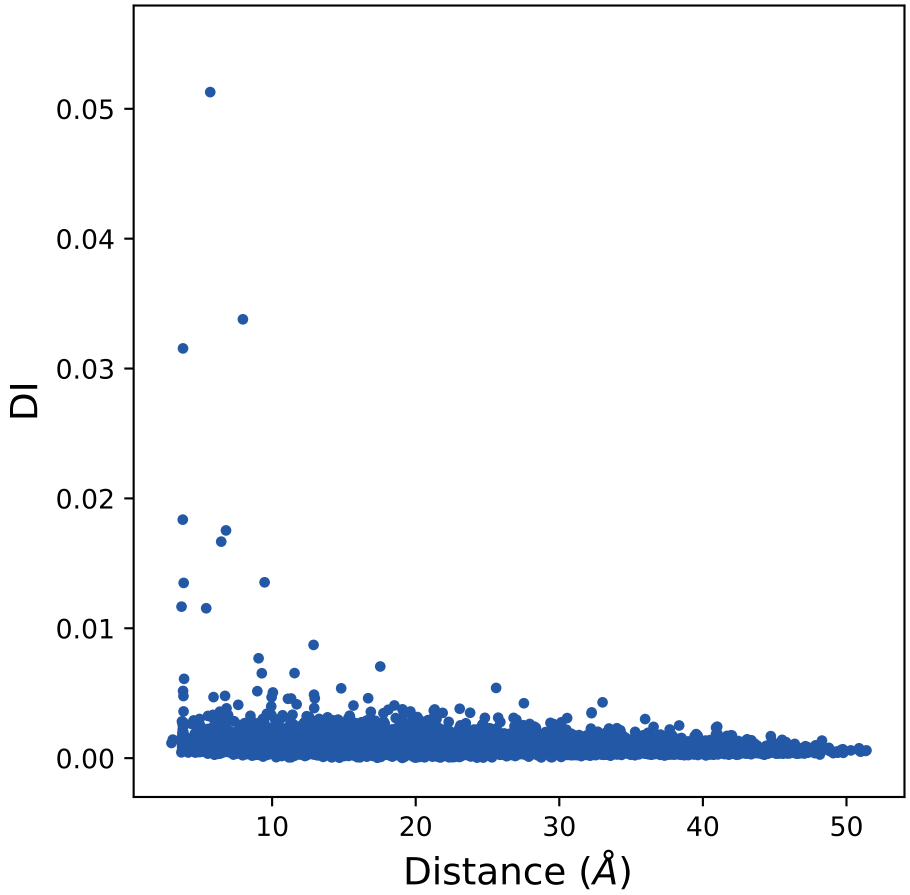

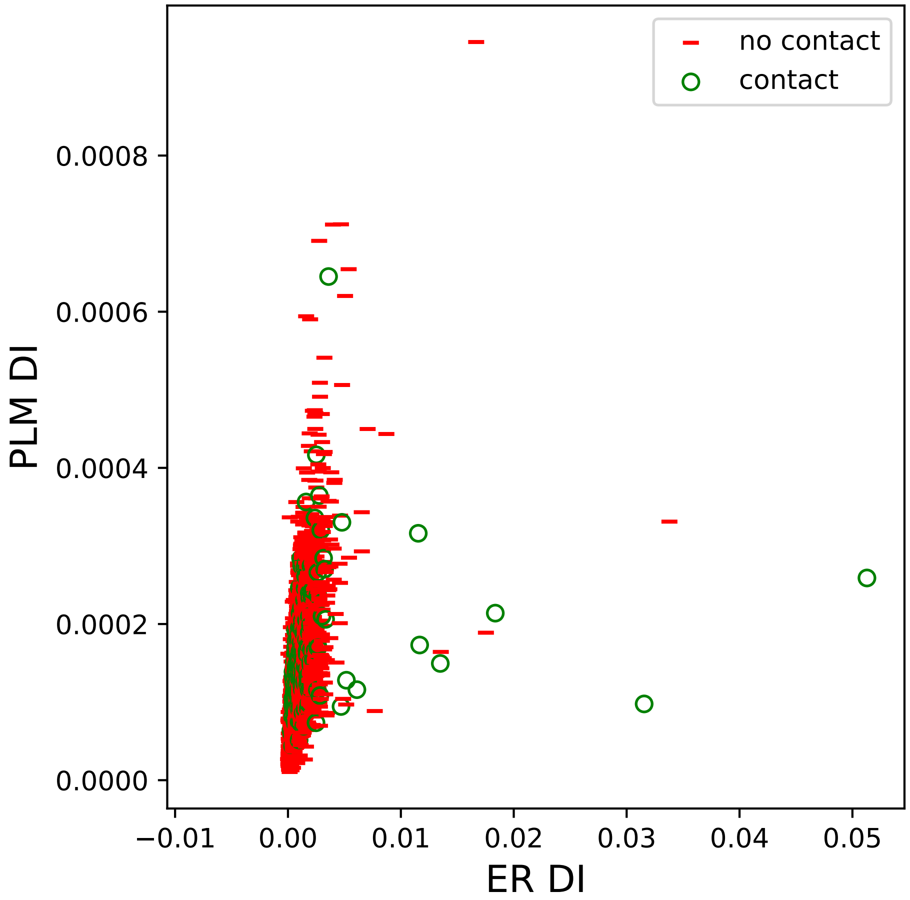

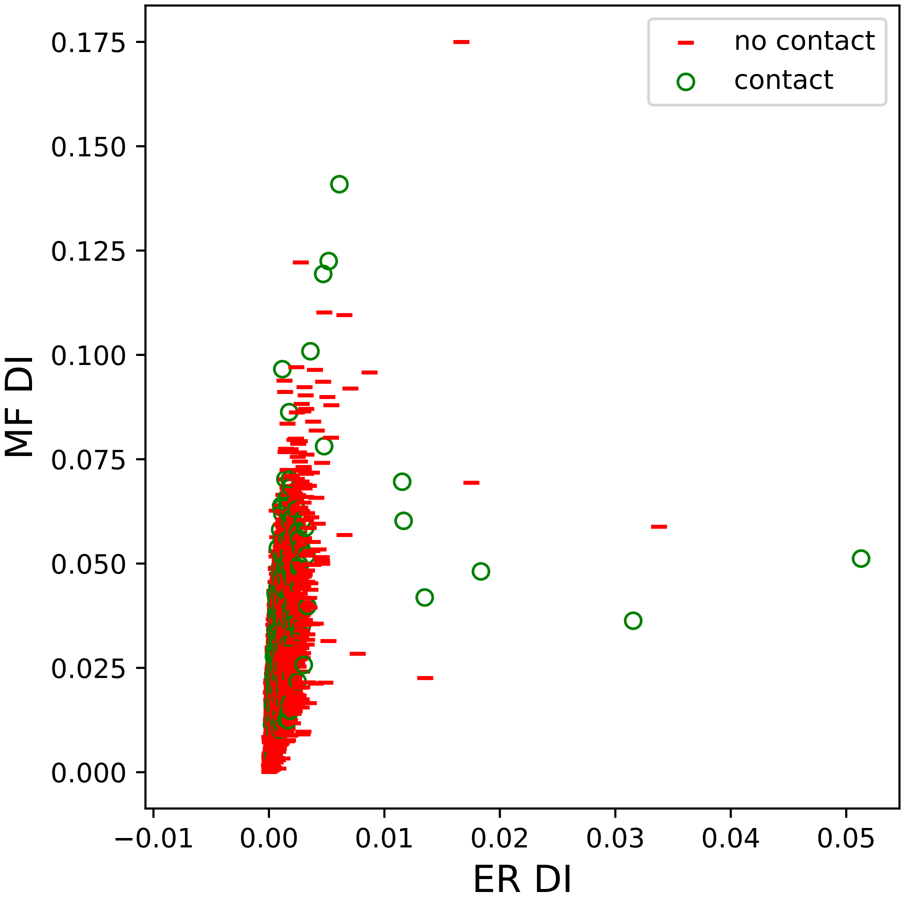

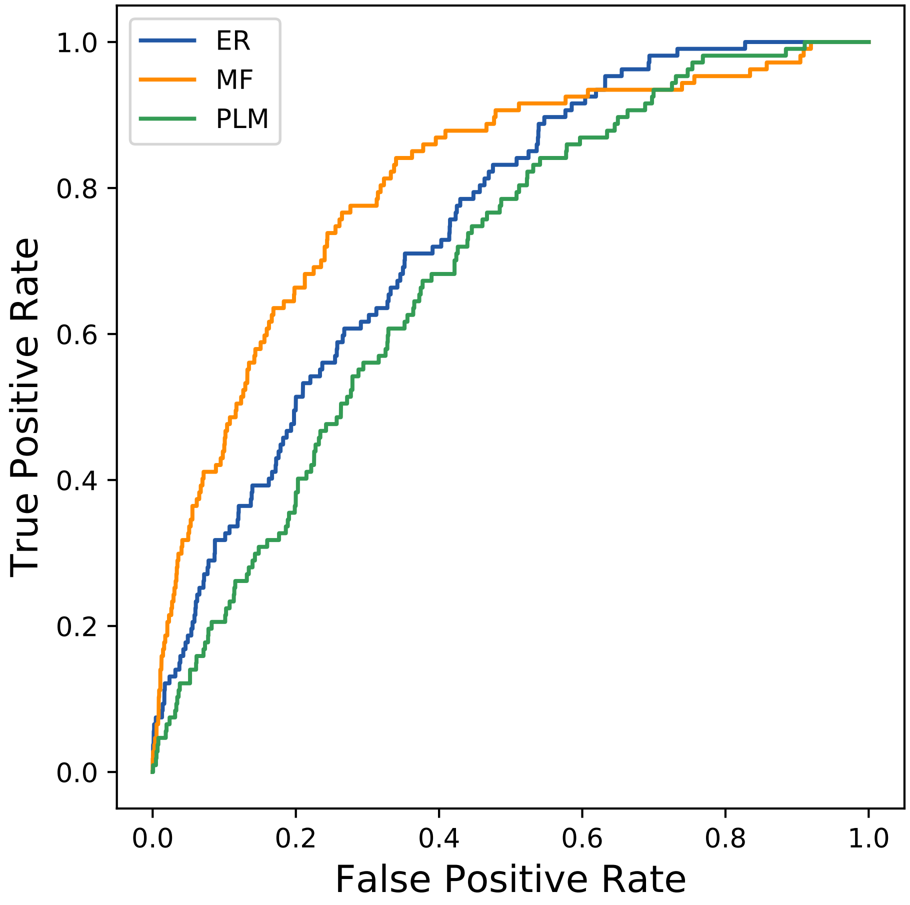

### 855seq_24col_2ikd_PF12032_plots.pdf

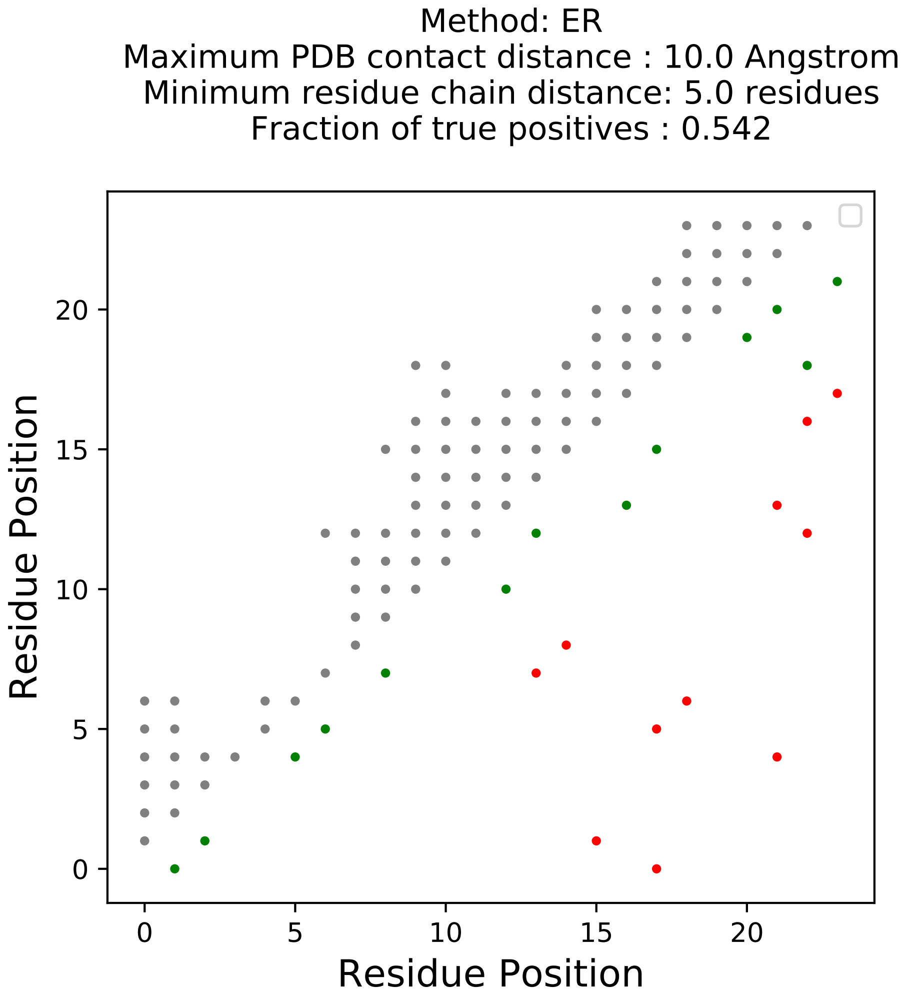

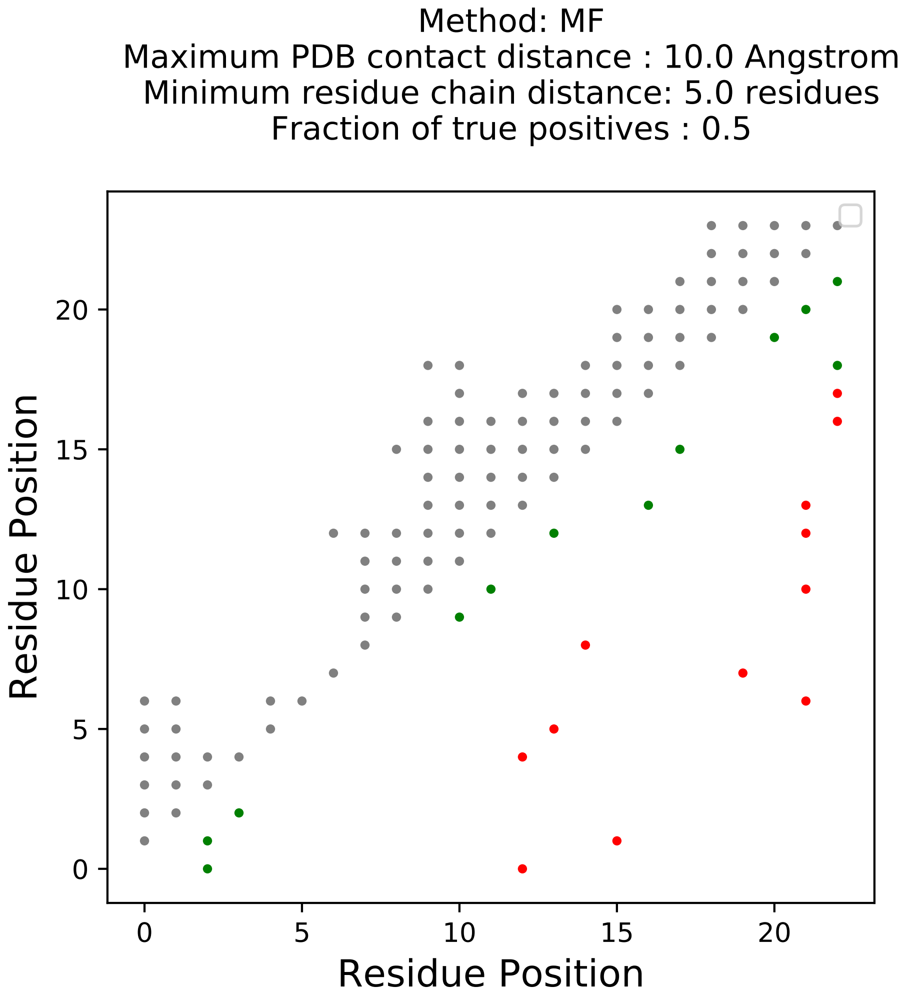

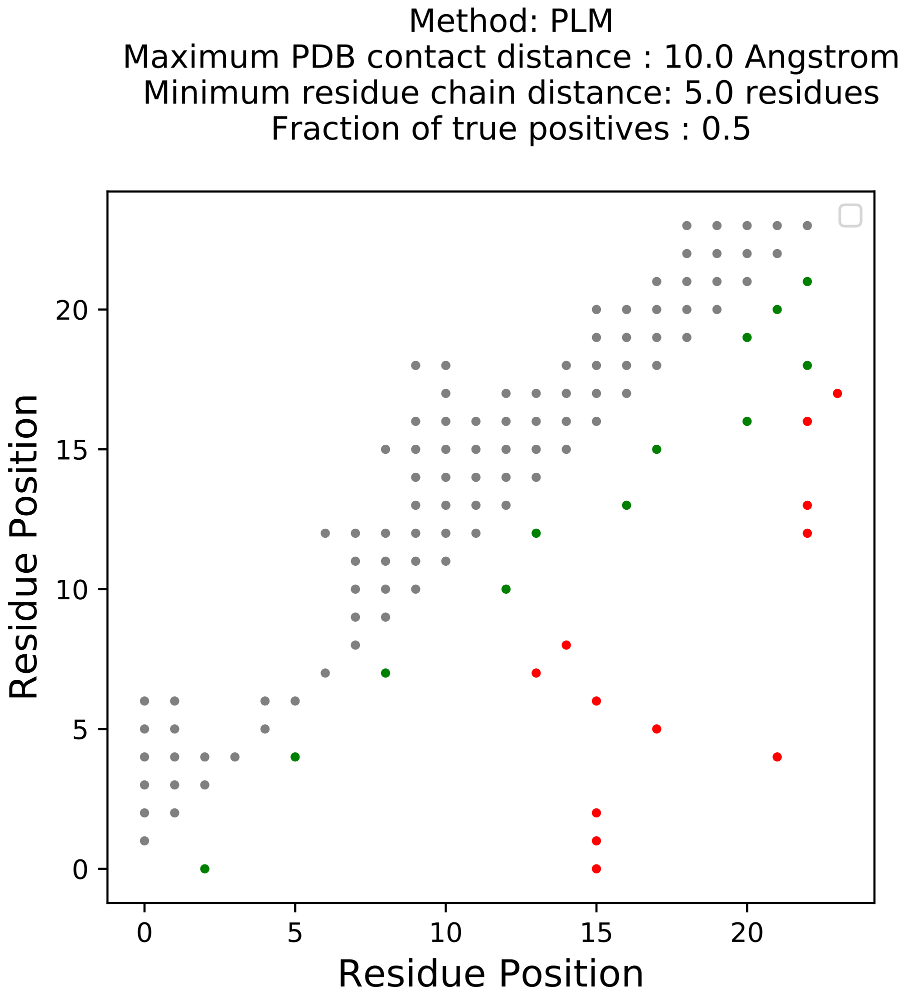

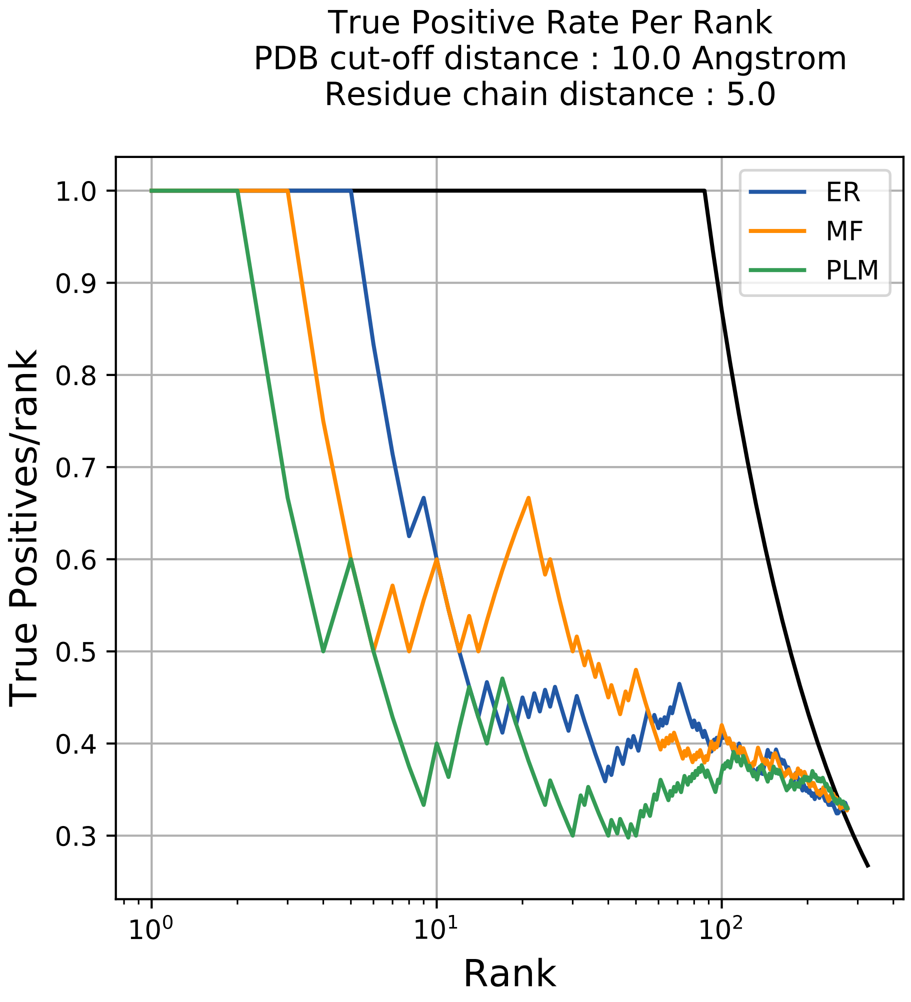

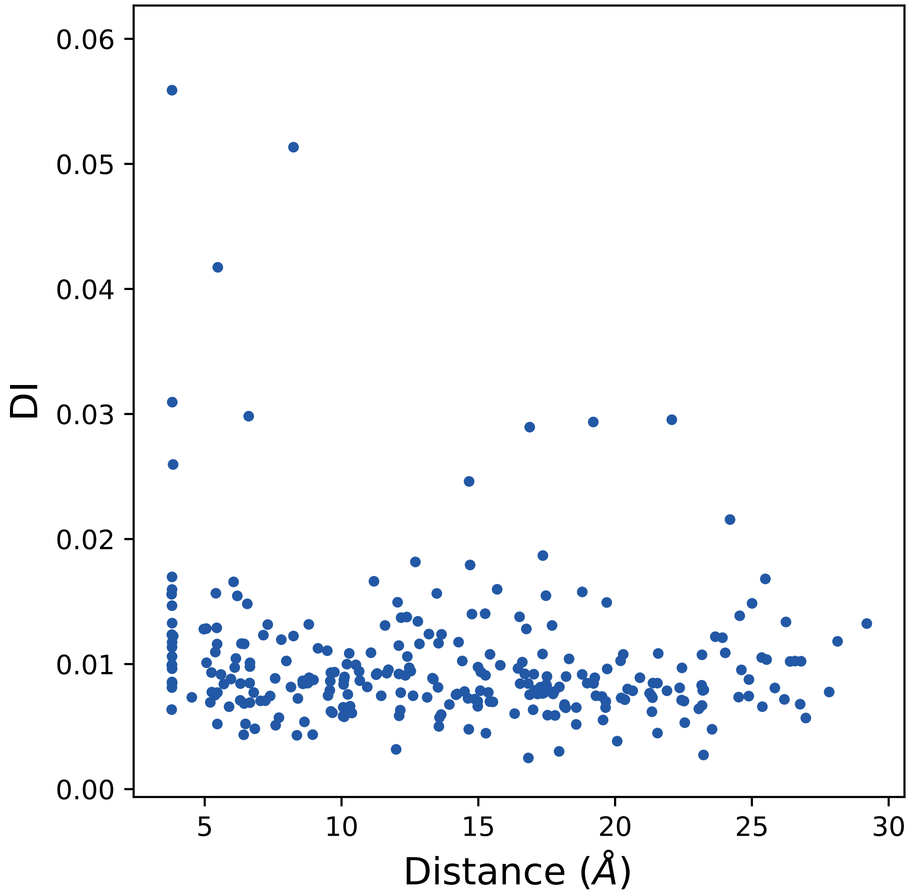

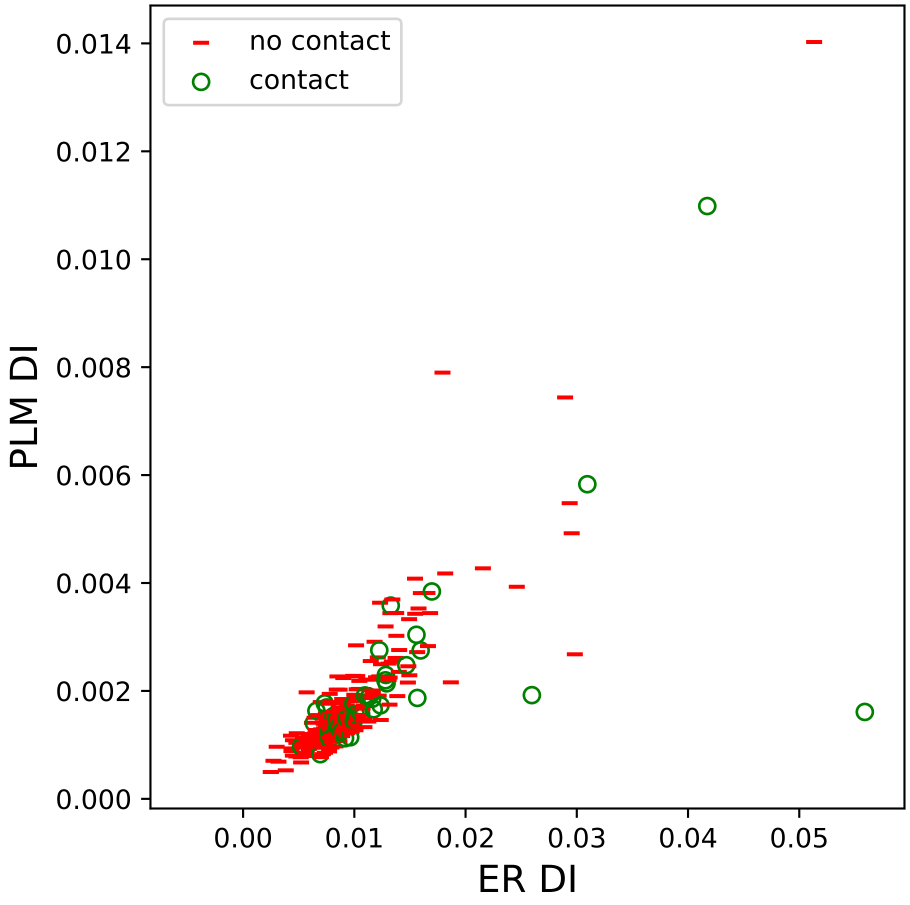

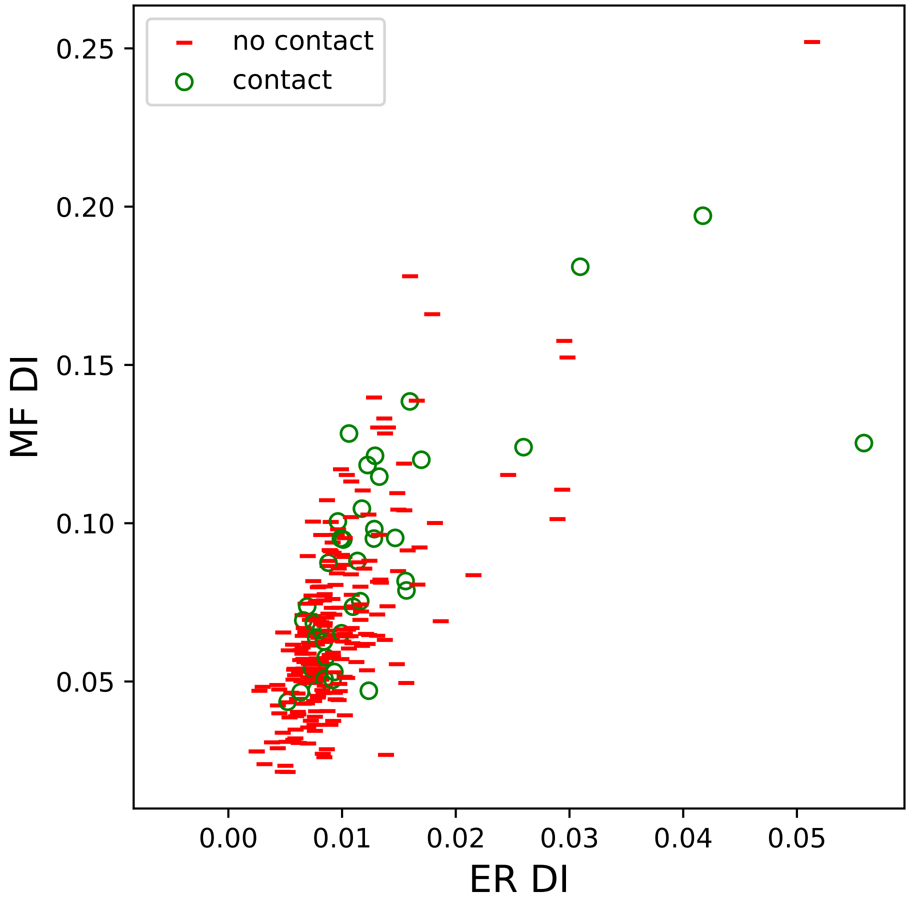

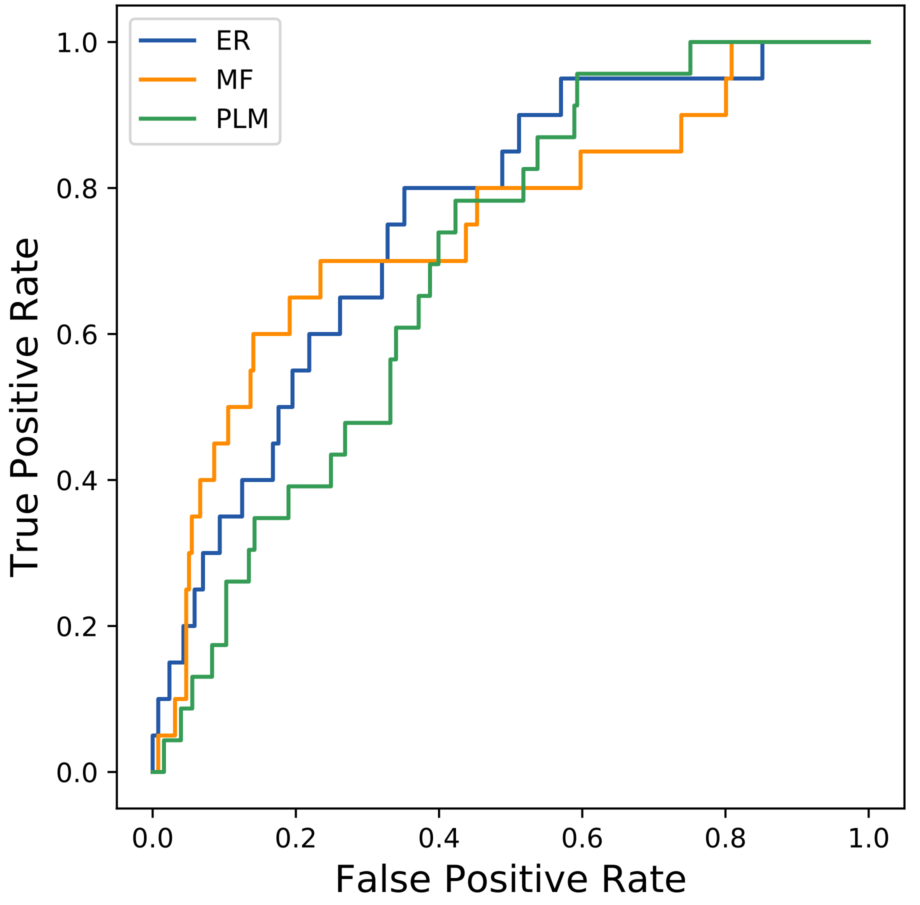

### 871seq_105col_4hwv_PF00544_plots.pdf

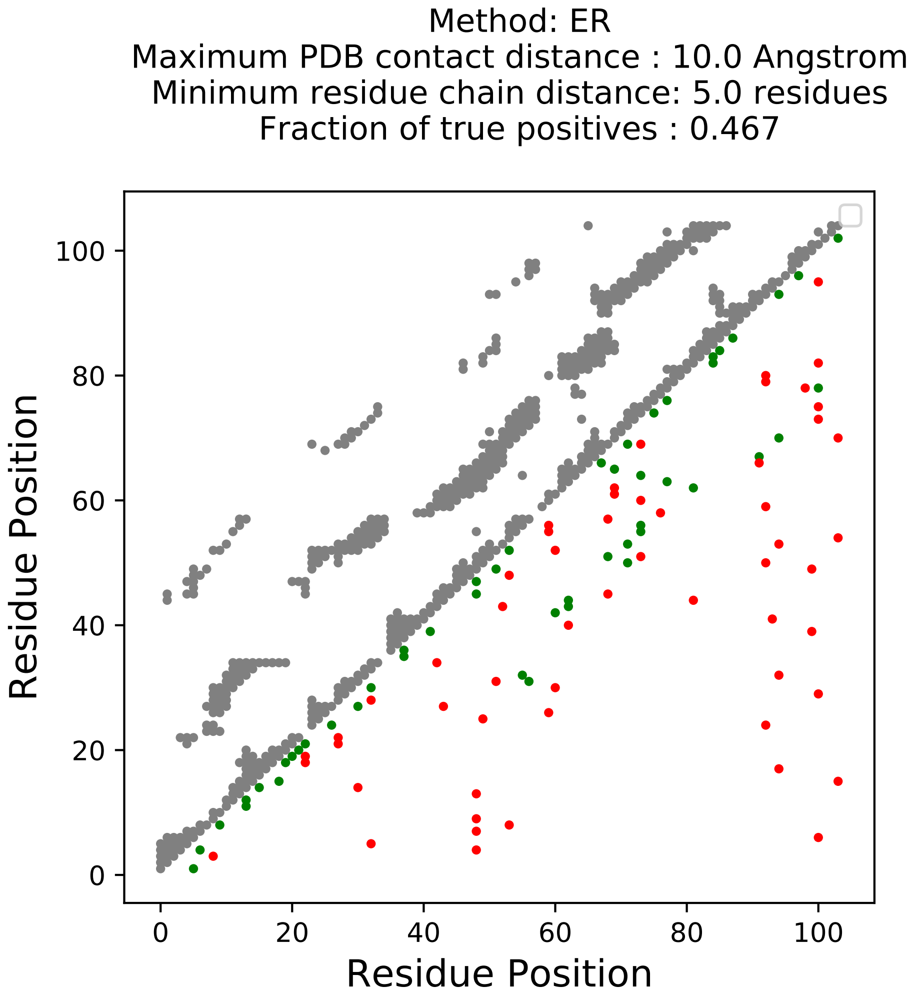

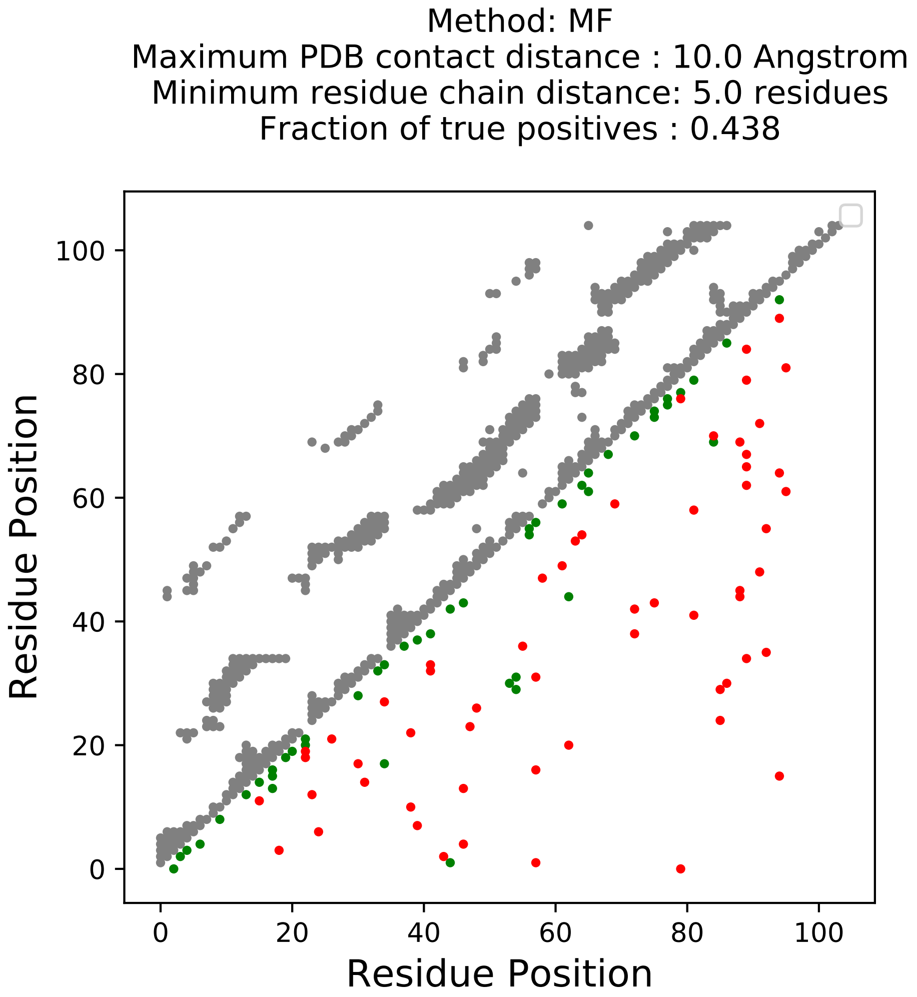

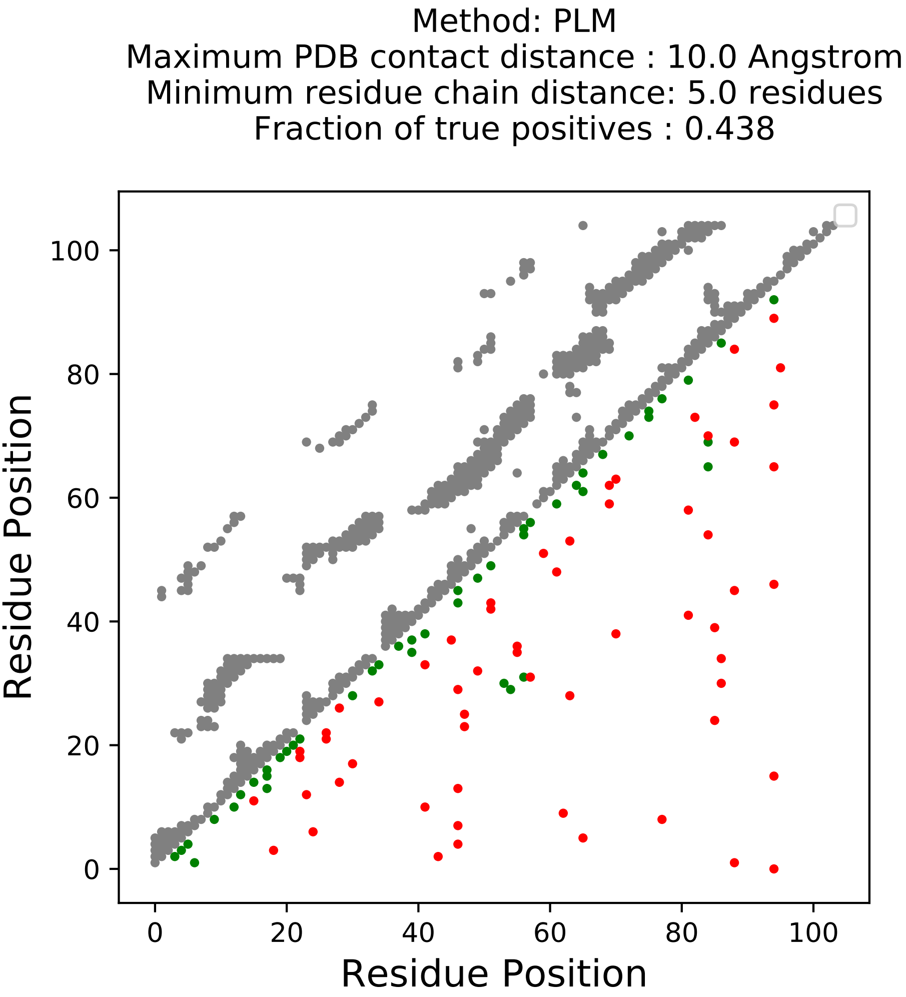

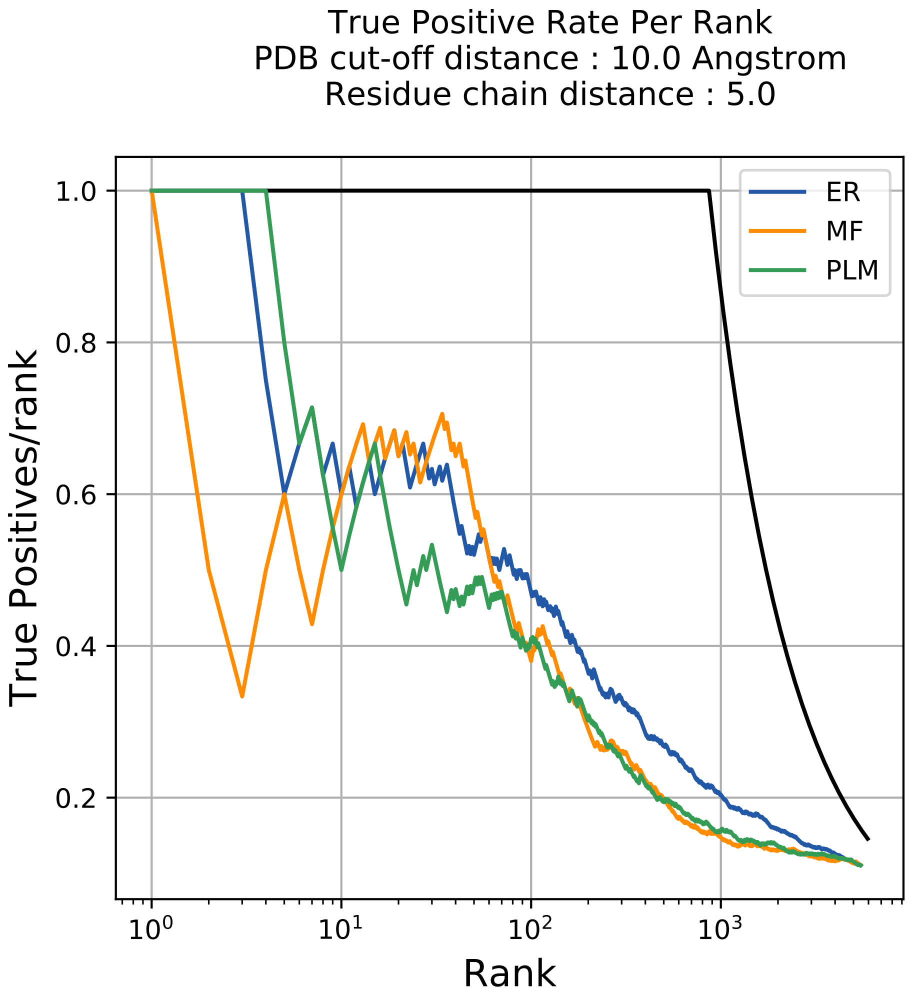

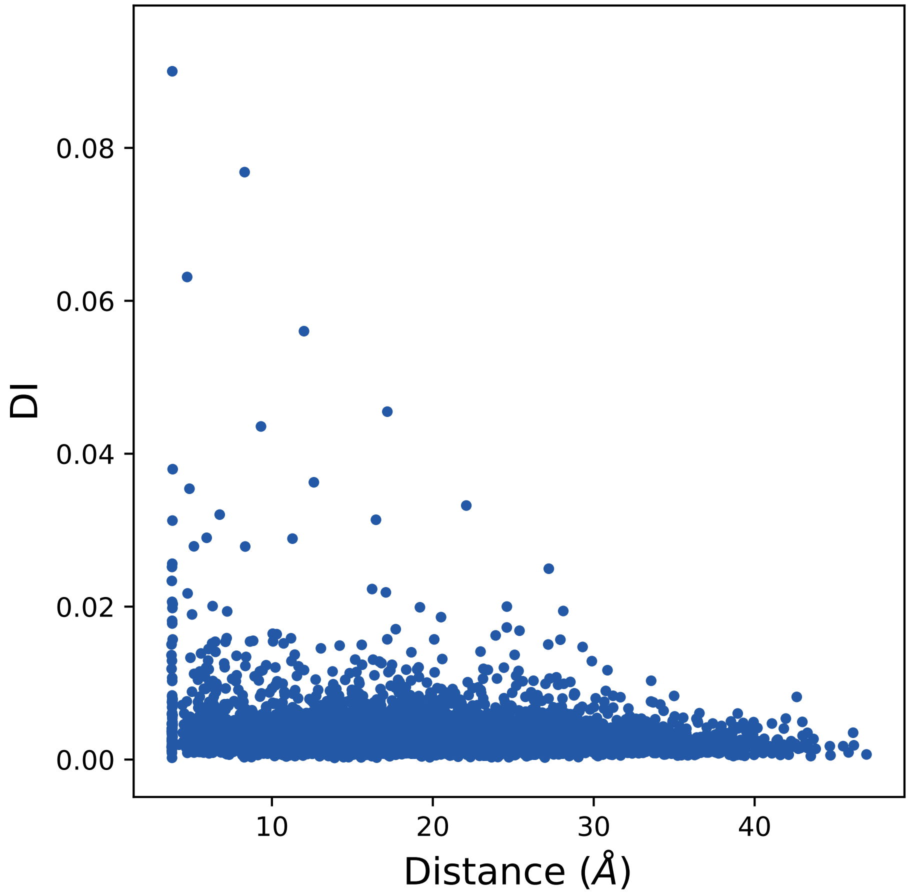

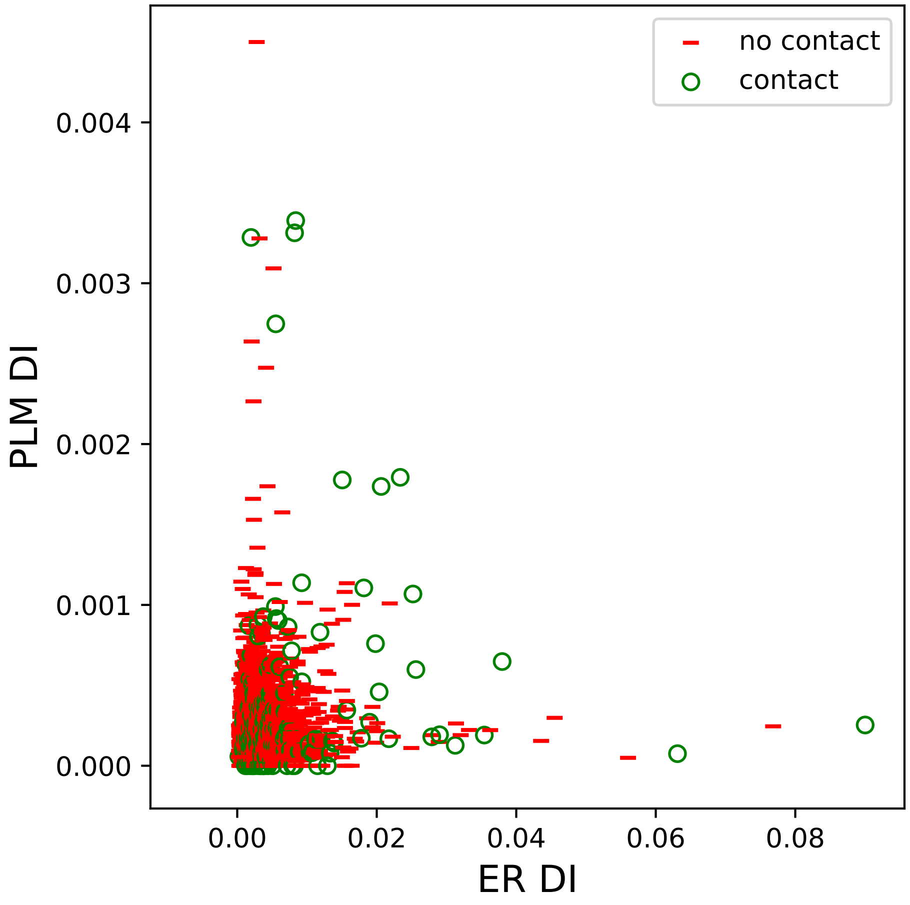

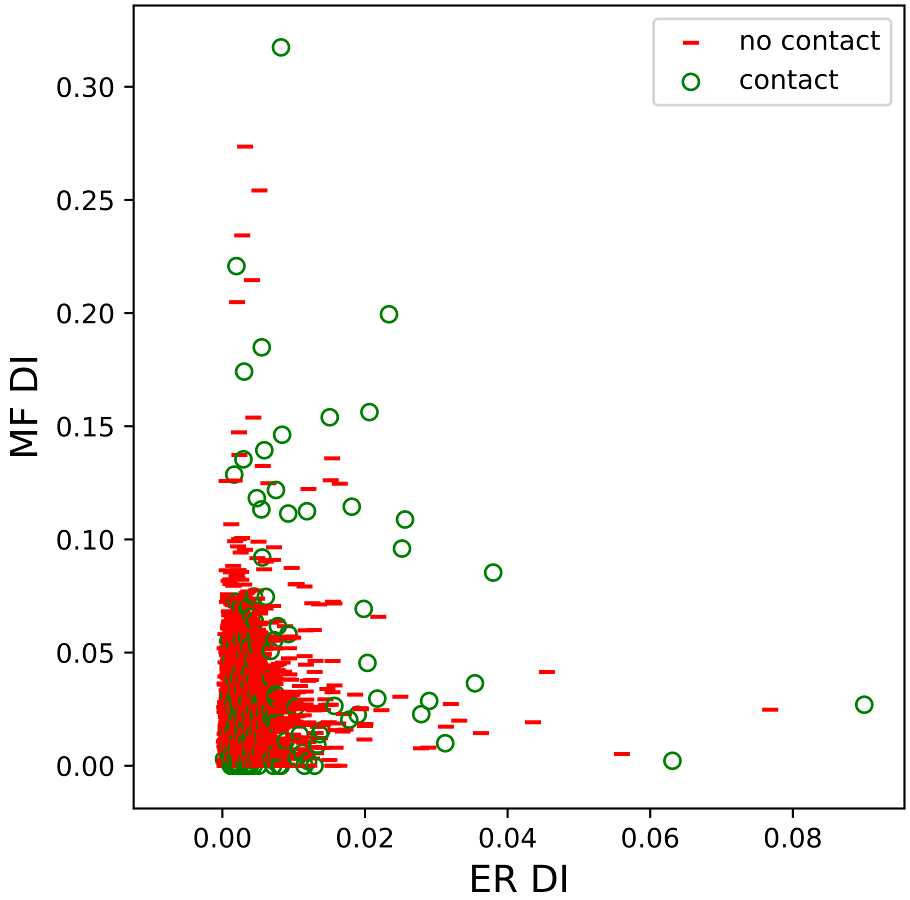

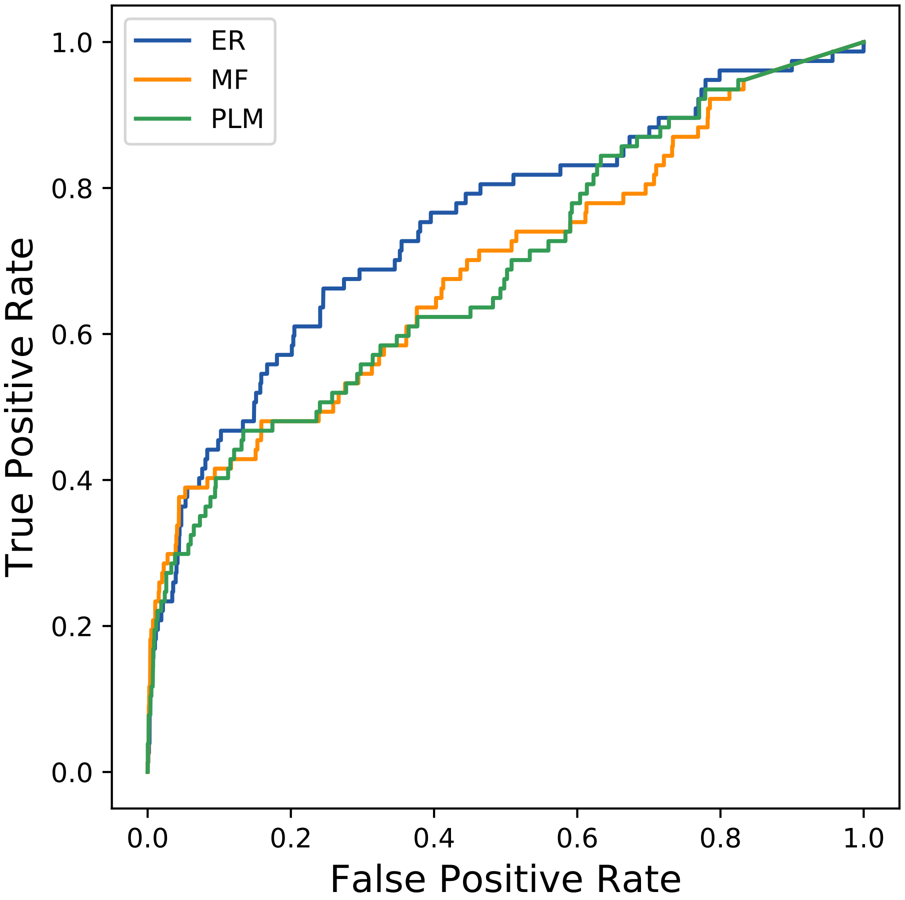

### 884seq_31col_3qy2_PF01111_plots.pdf

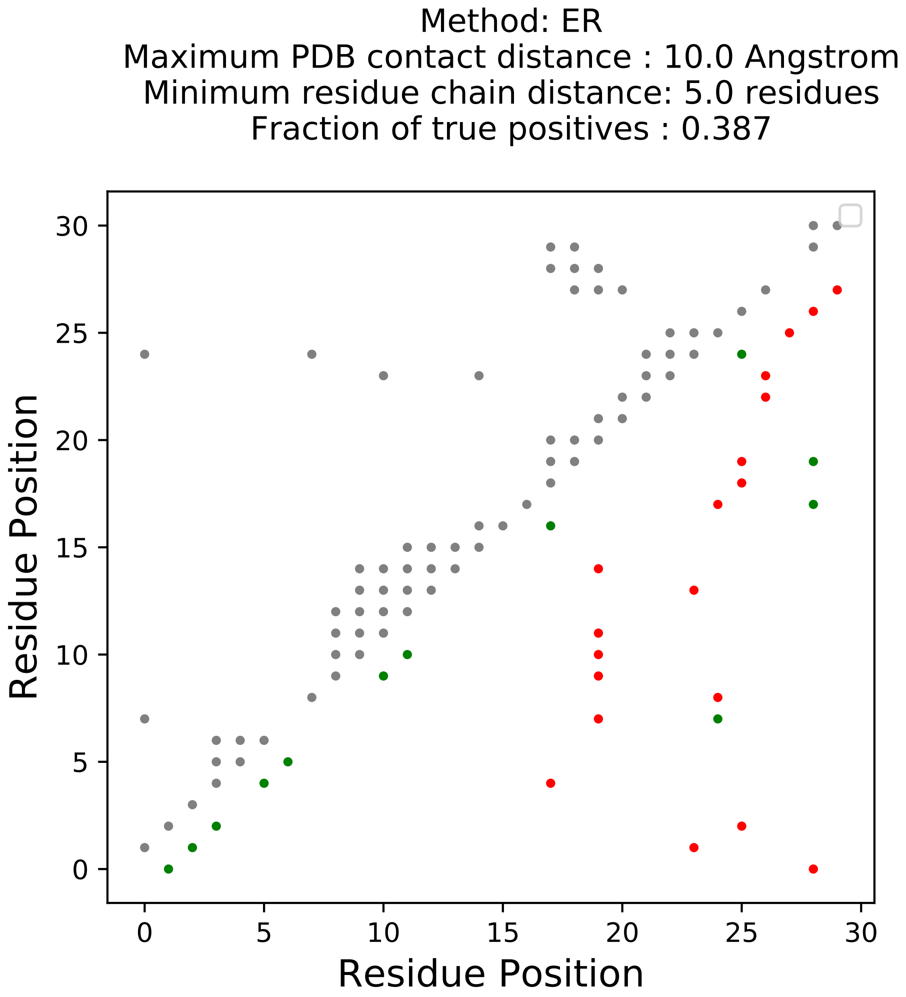

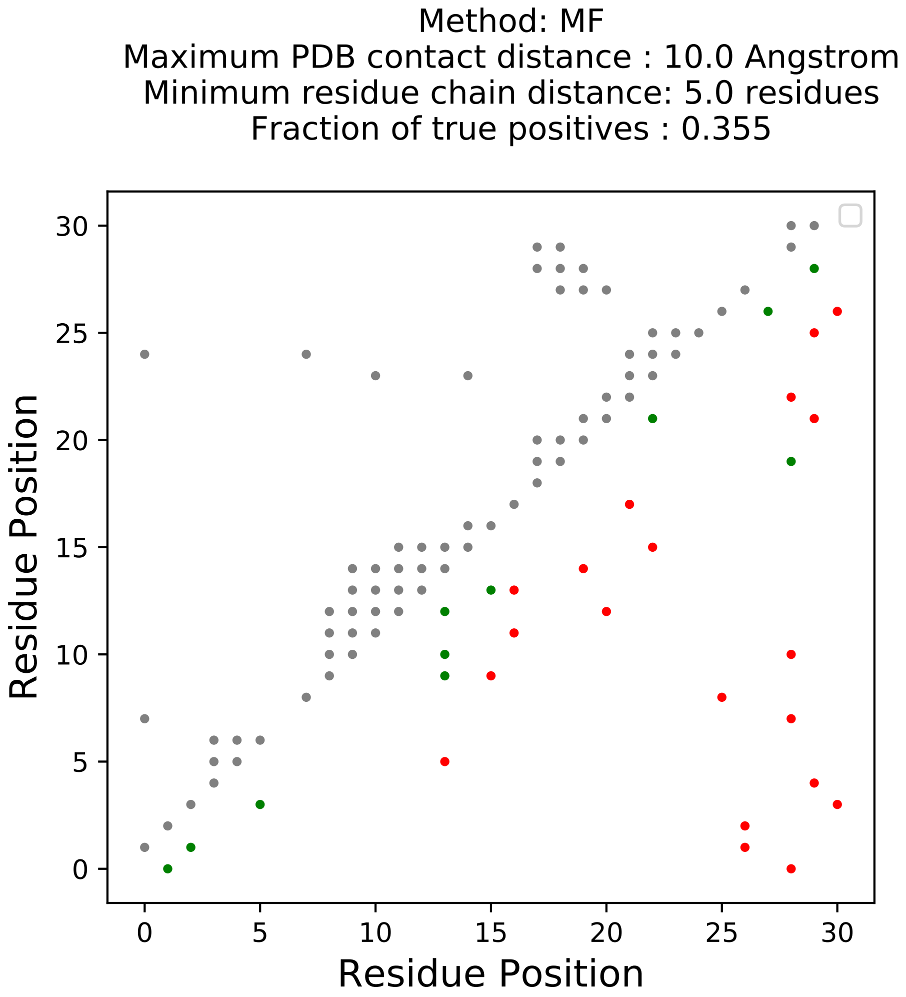

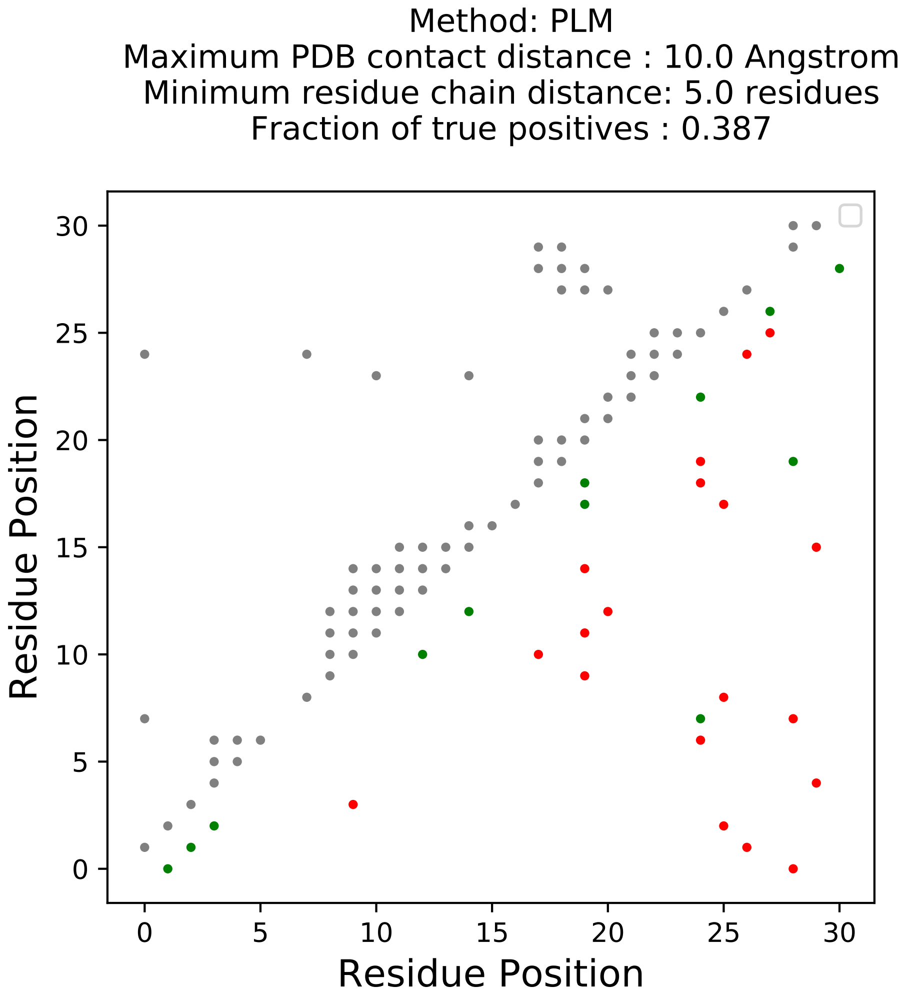

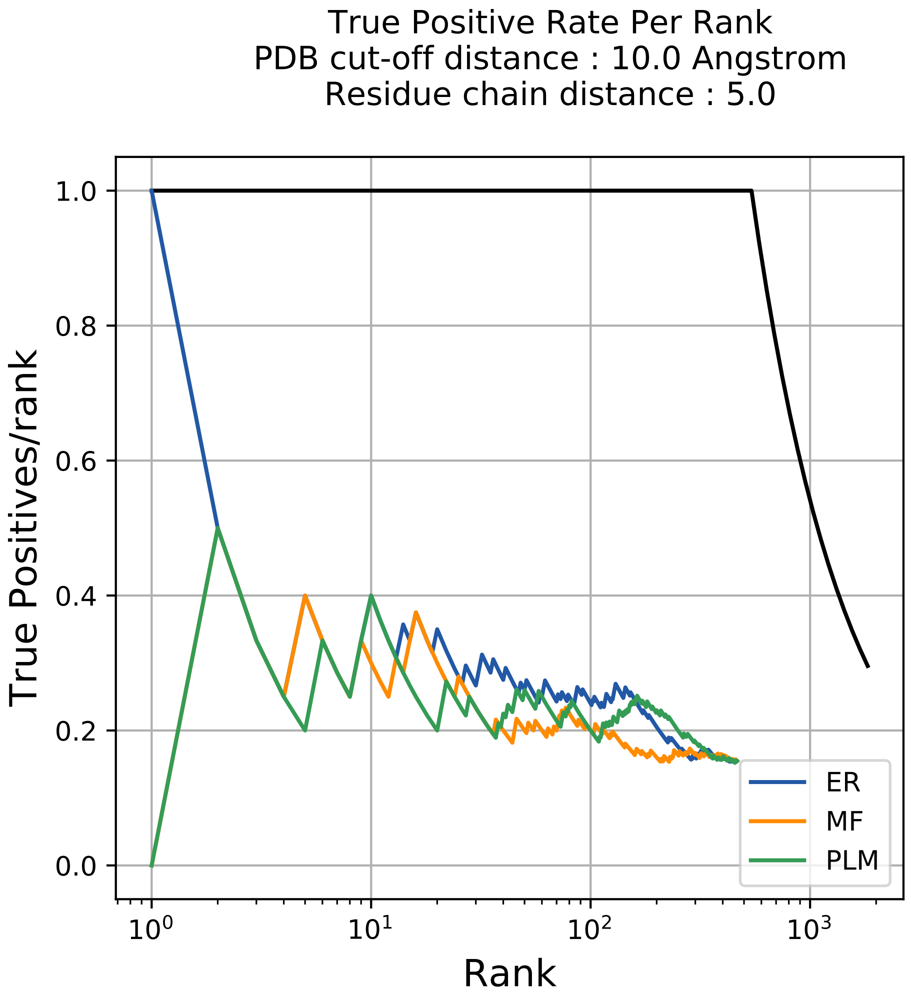

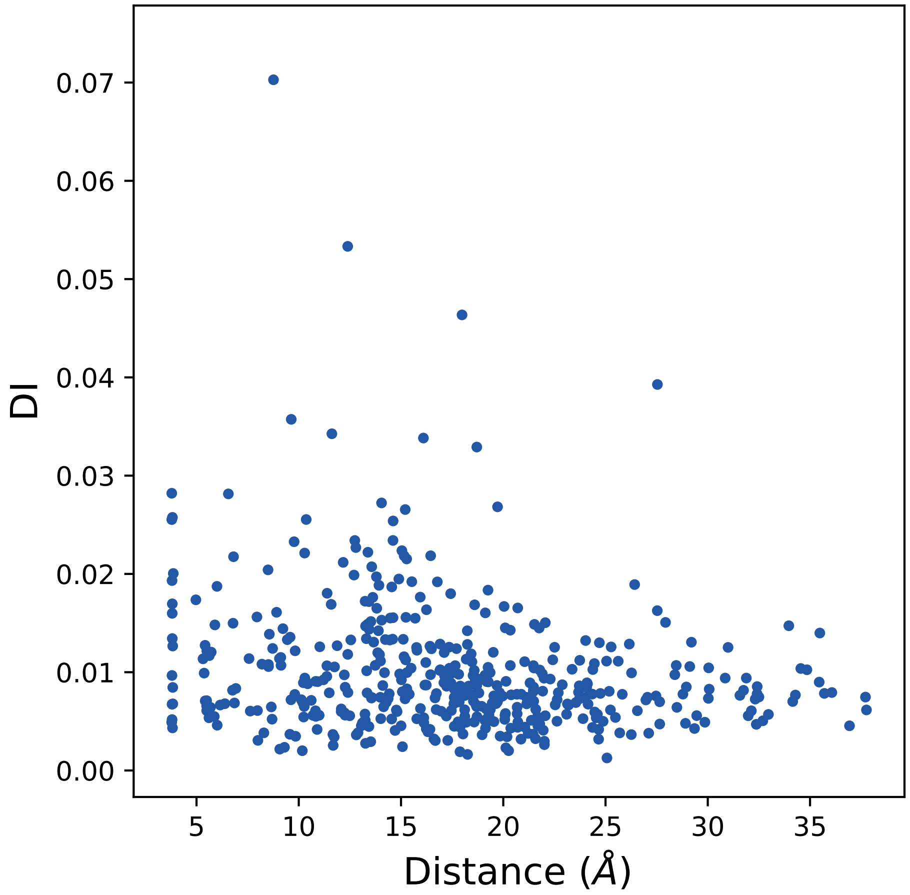

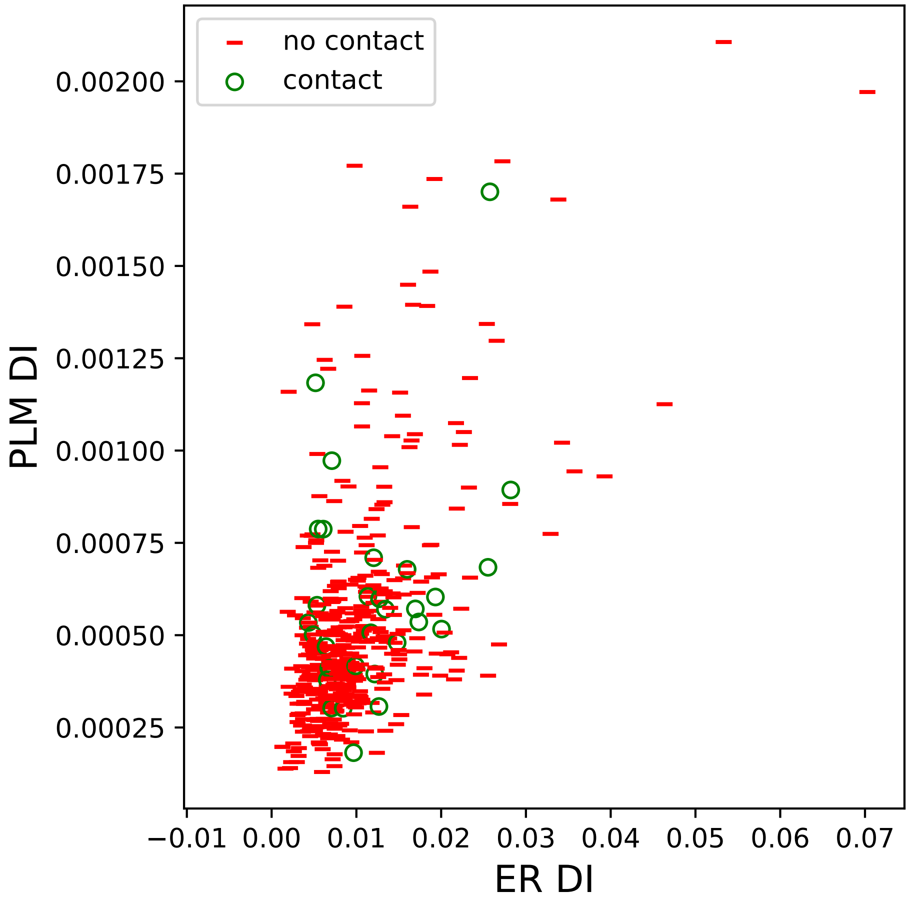

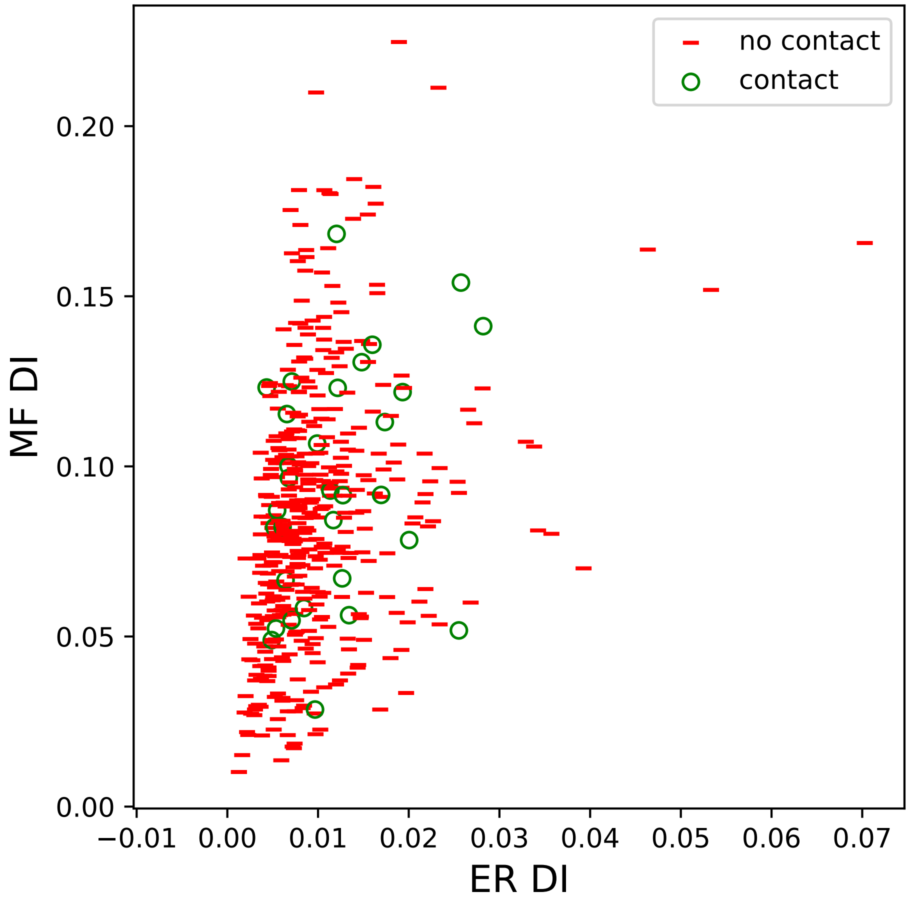

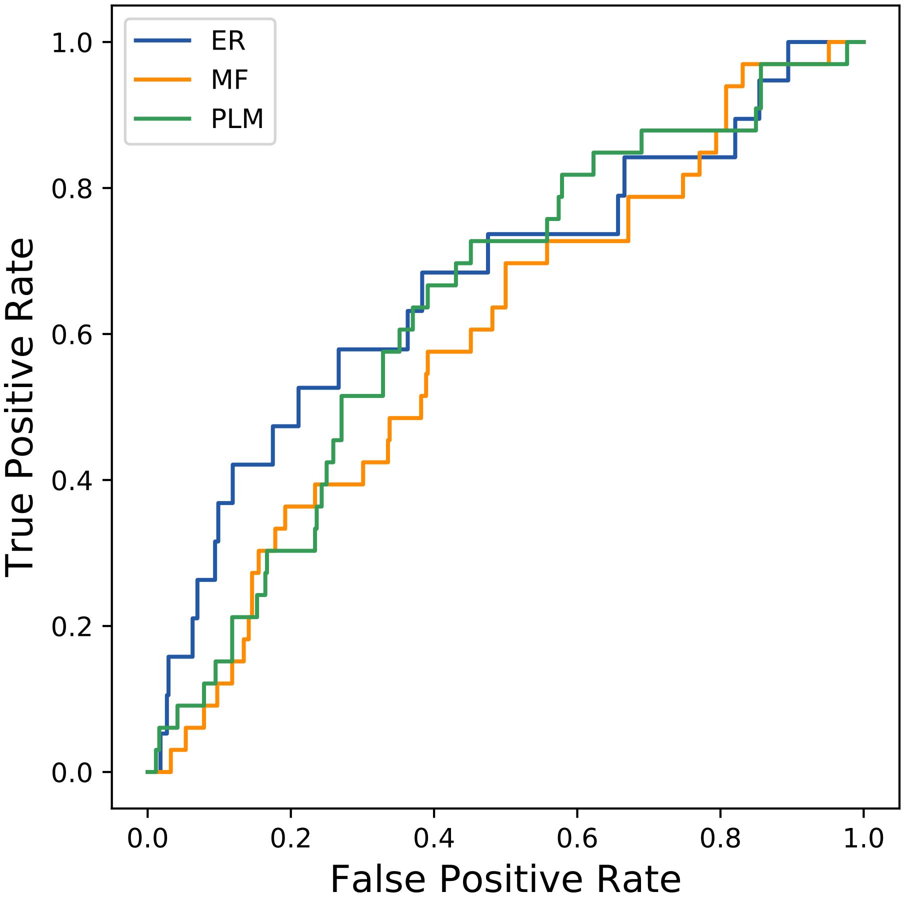

### 1125seq_148col_2czv_PF01876_plots.pdf

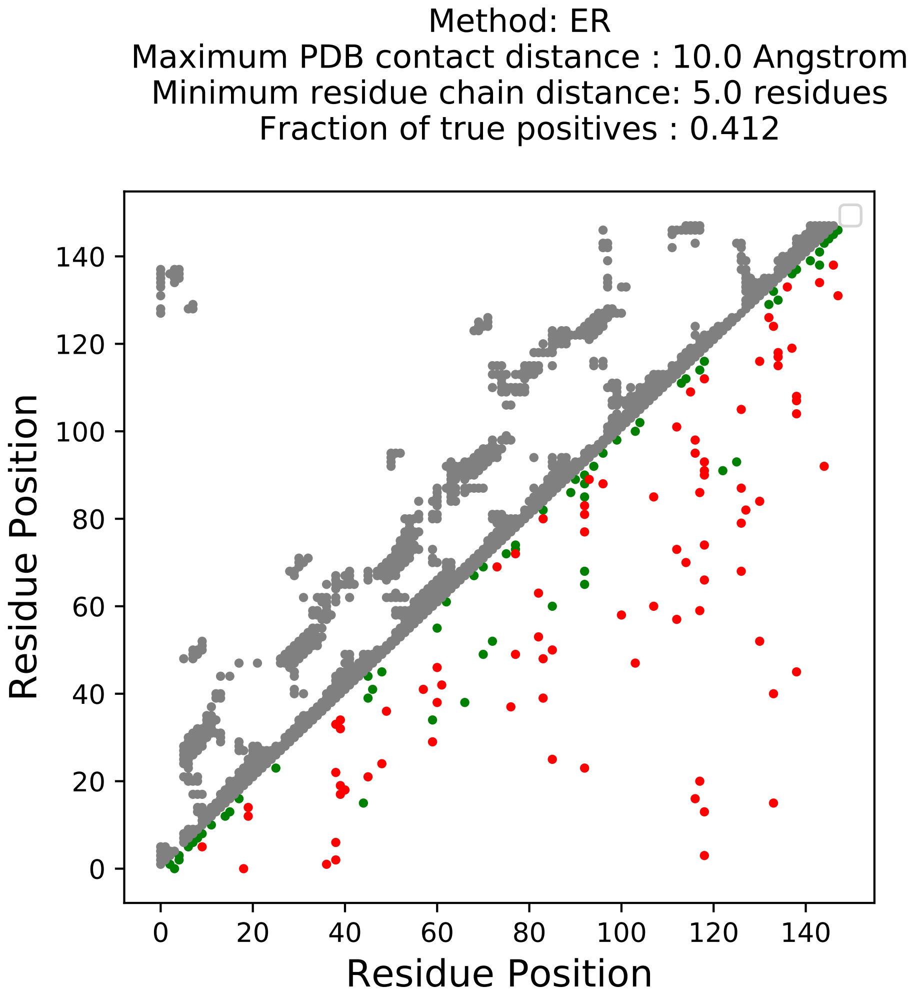

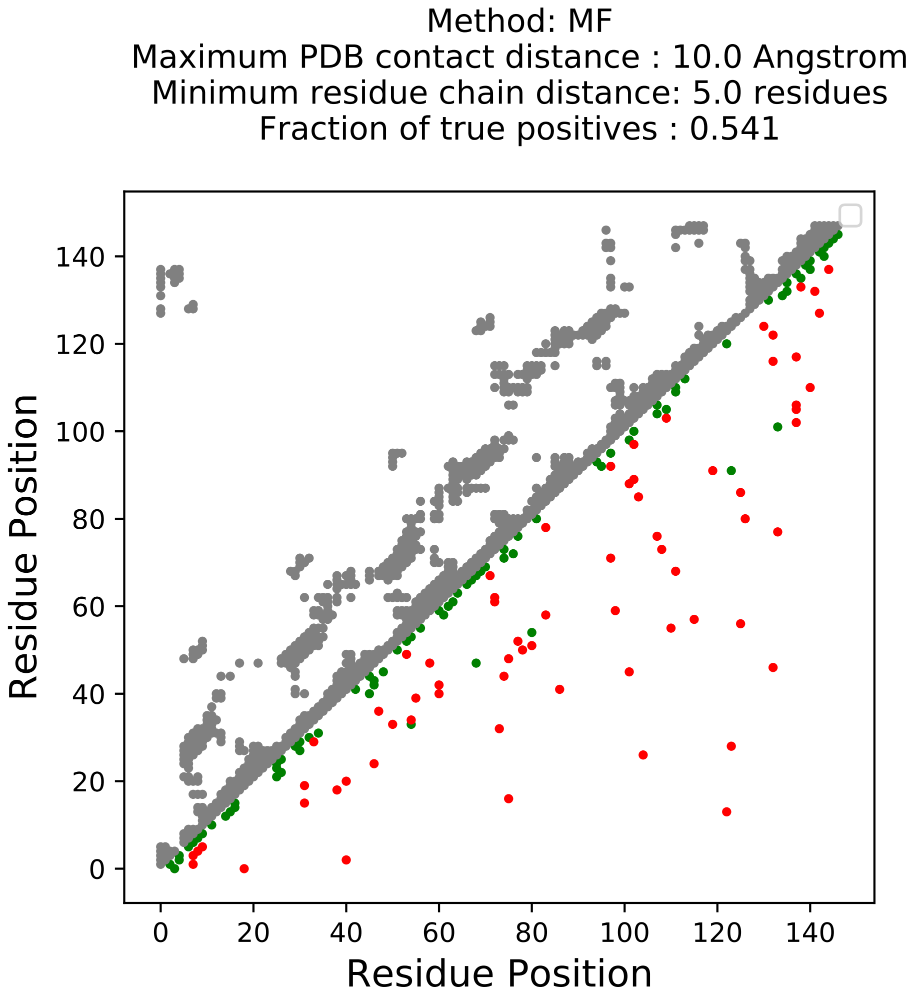

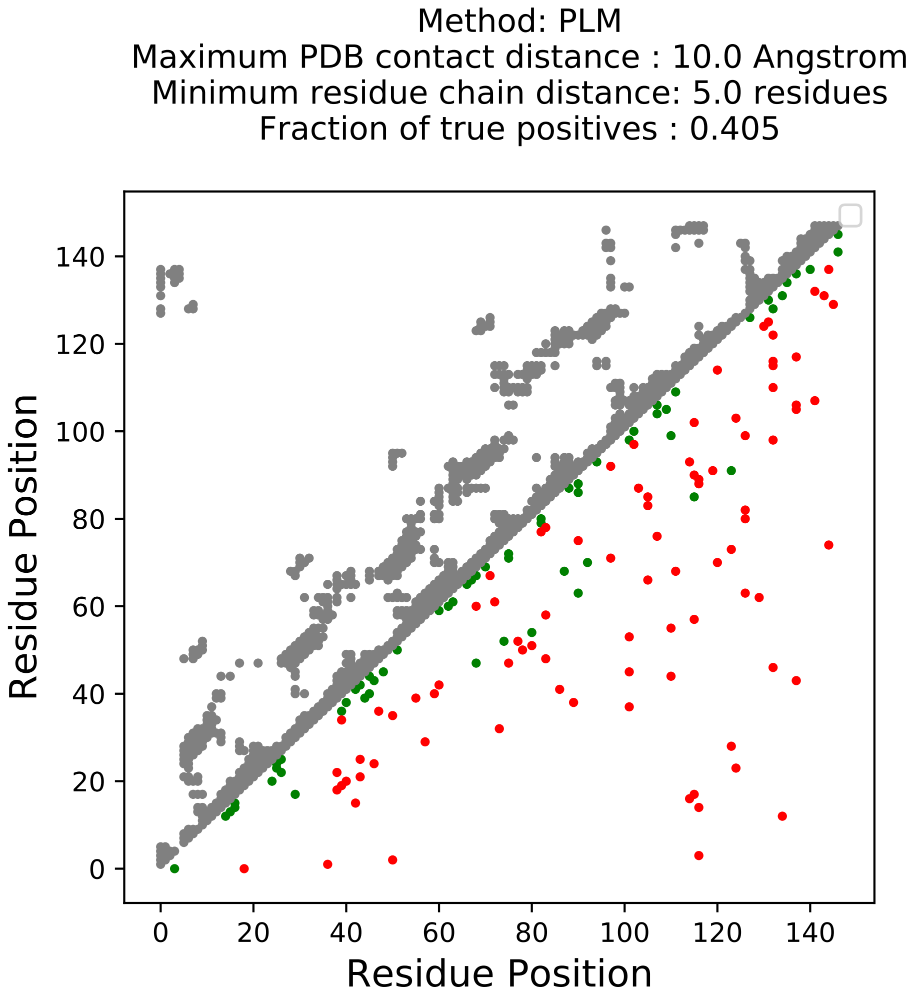

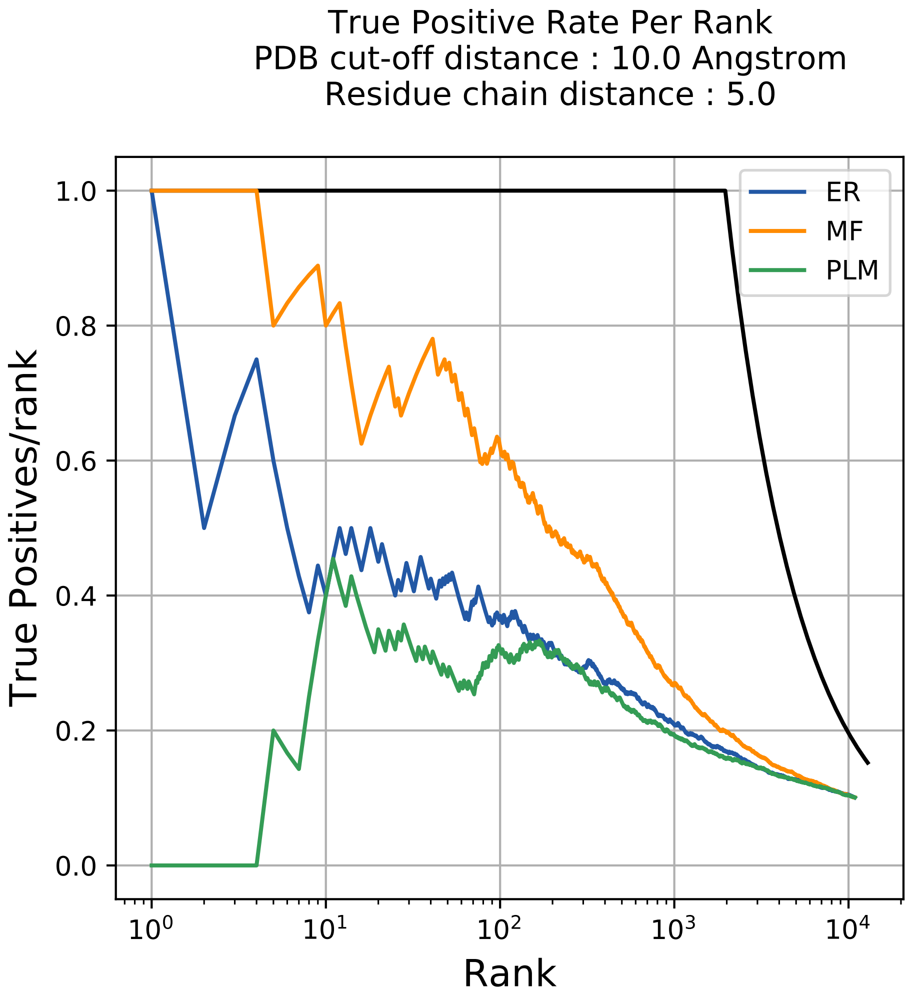

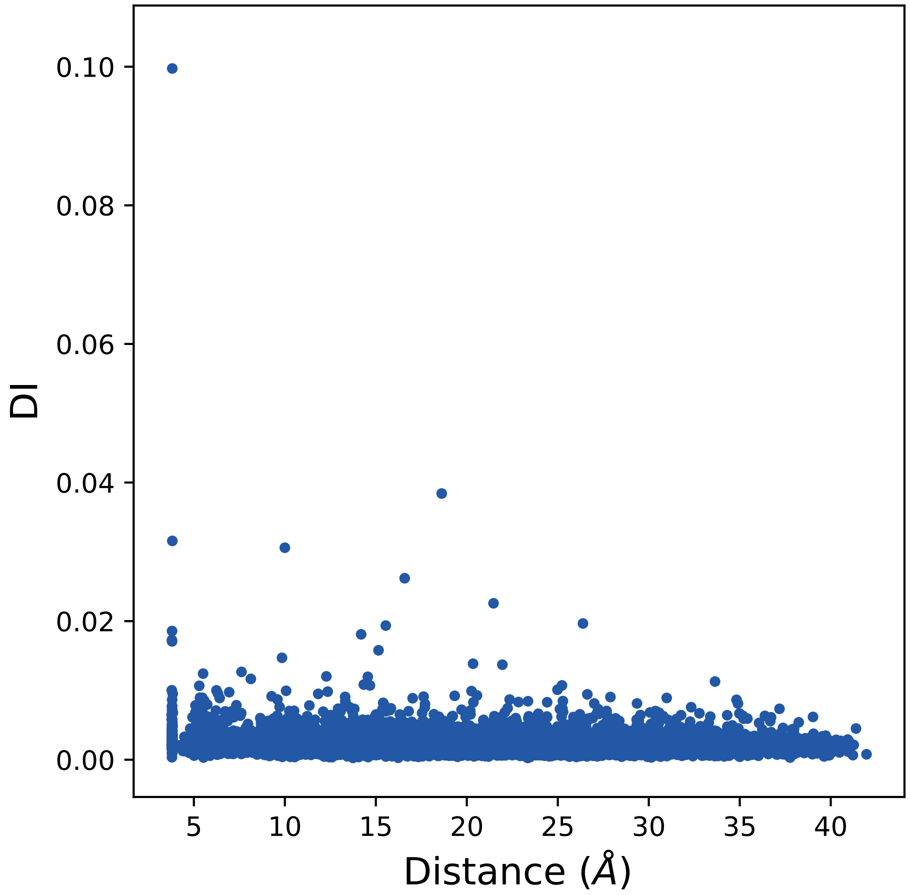

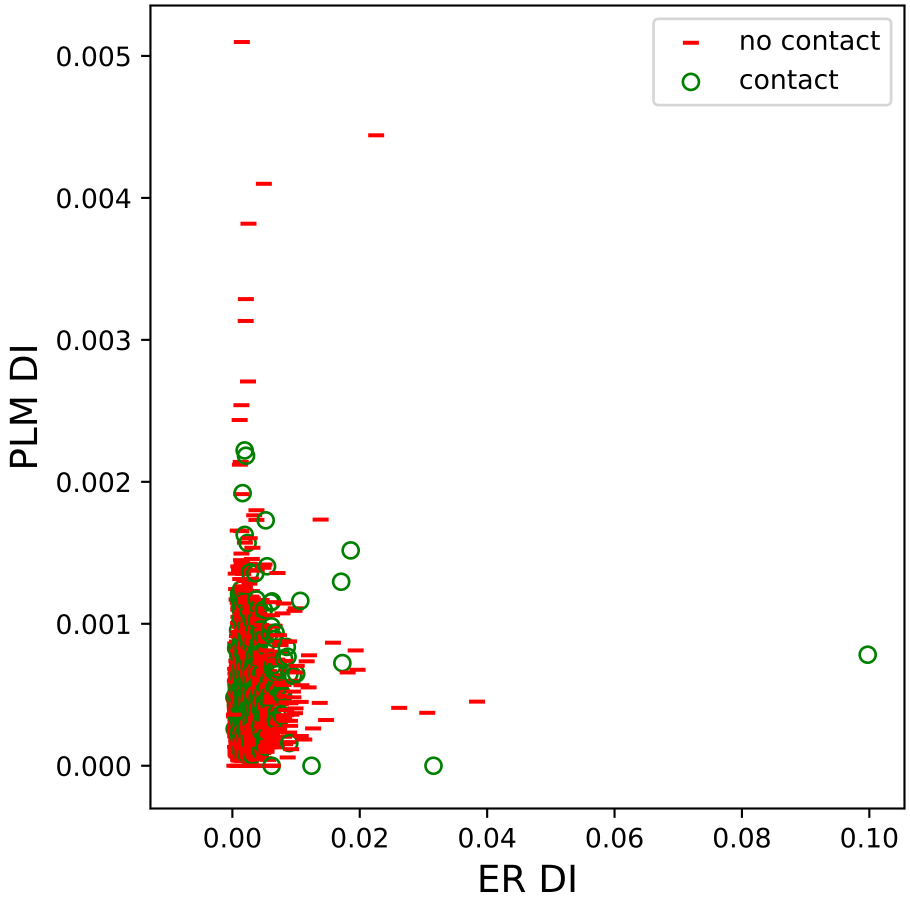

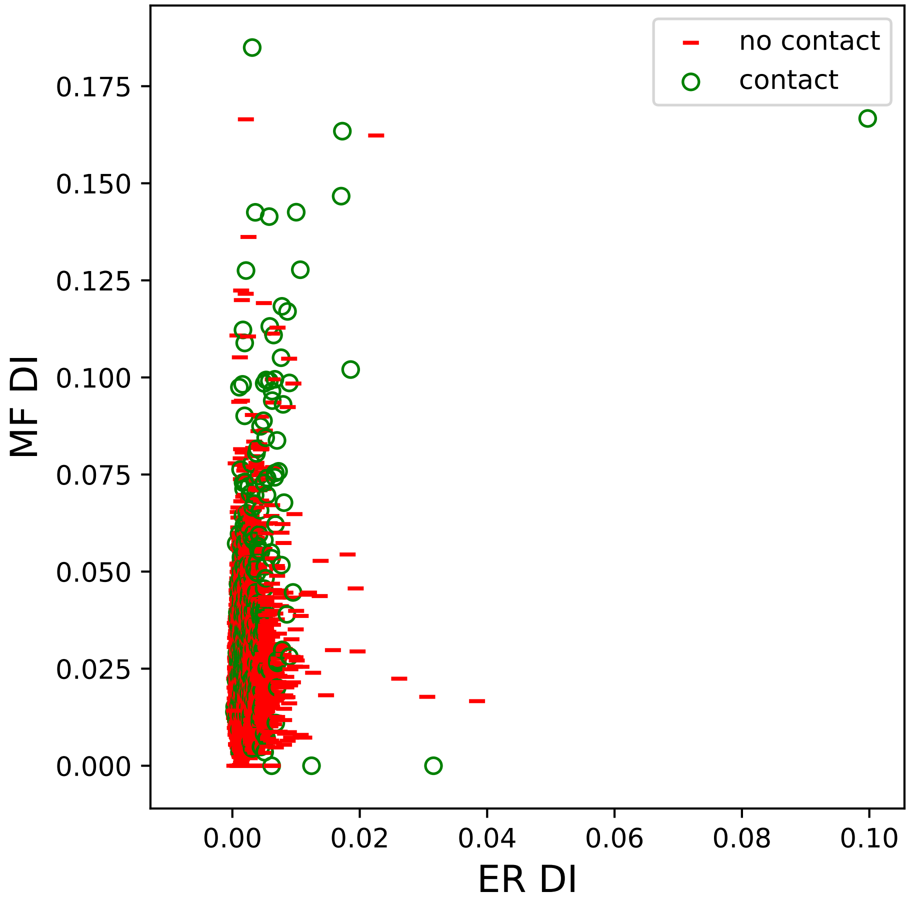

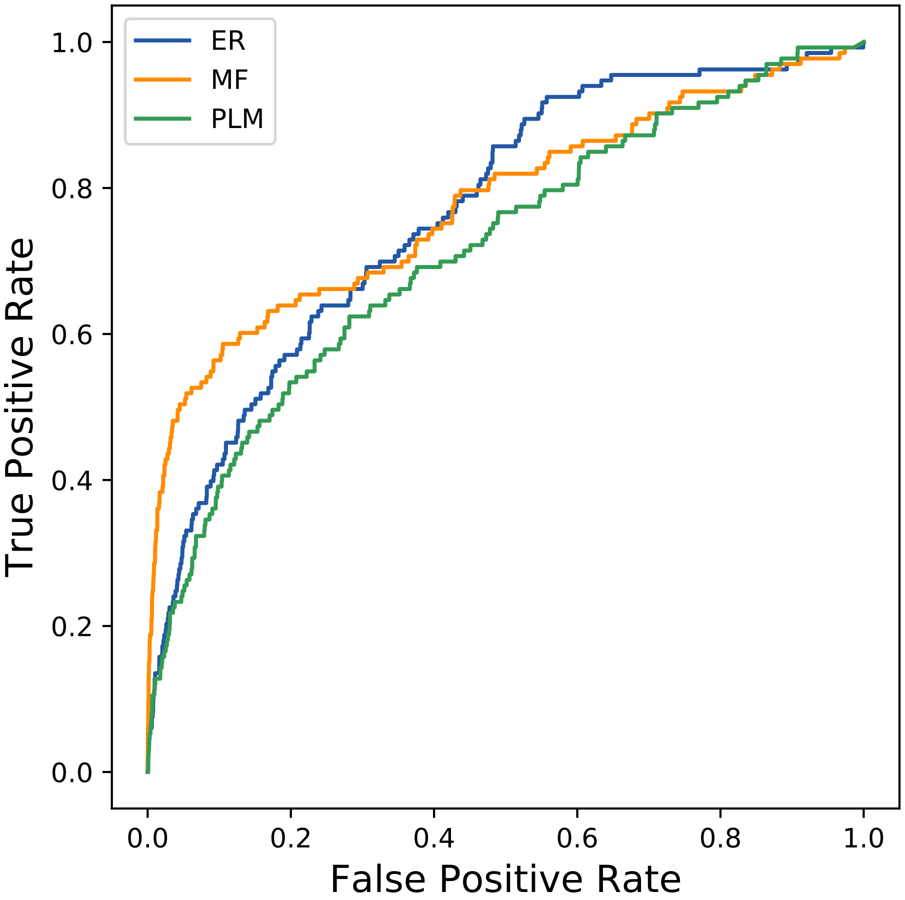

### 1184seq_104col_2jhw_PF02115_plots.pdf

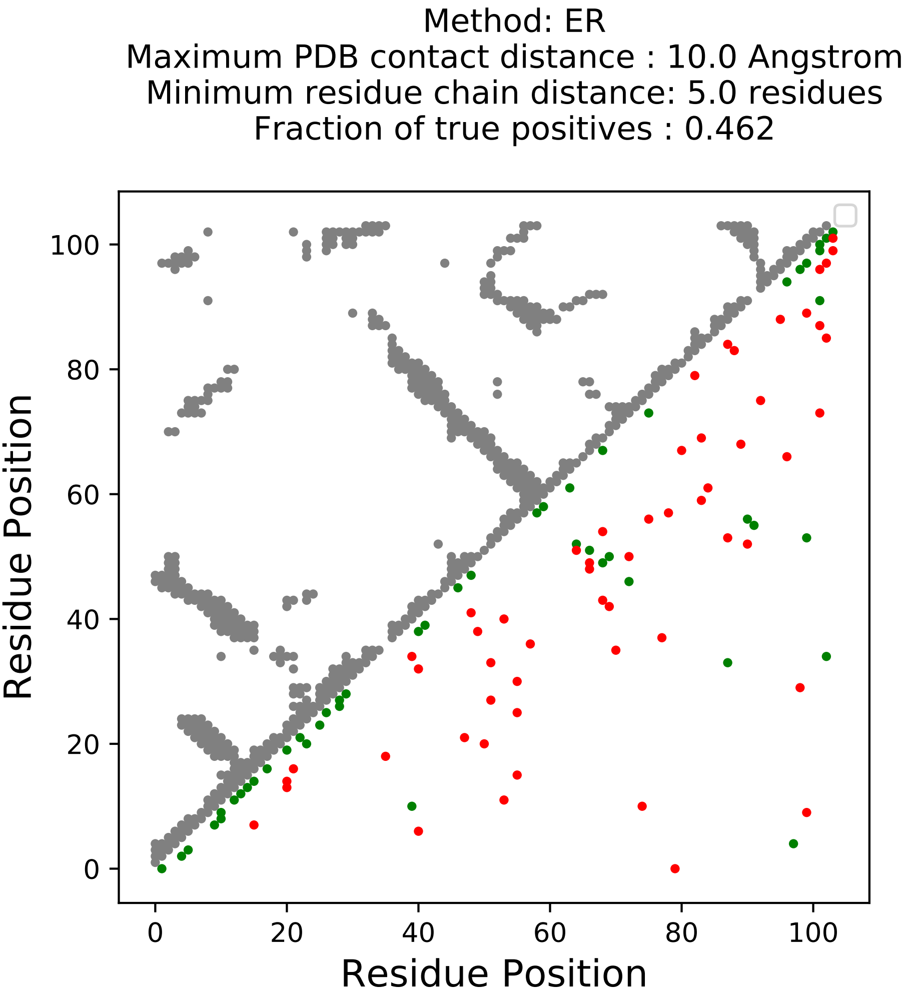

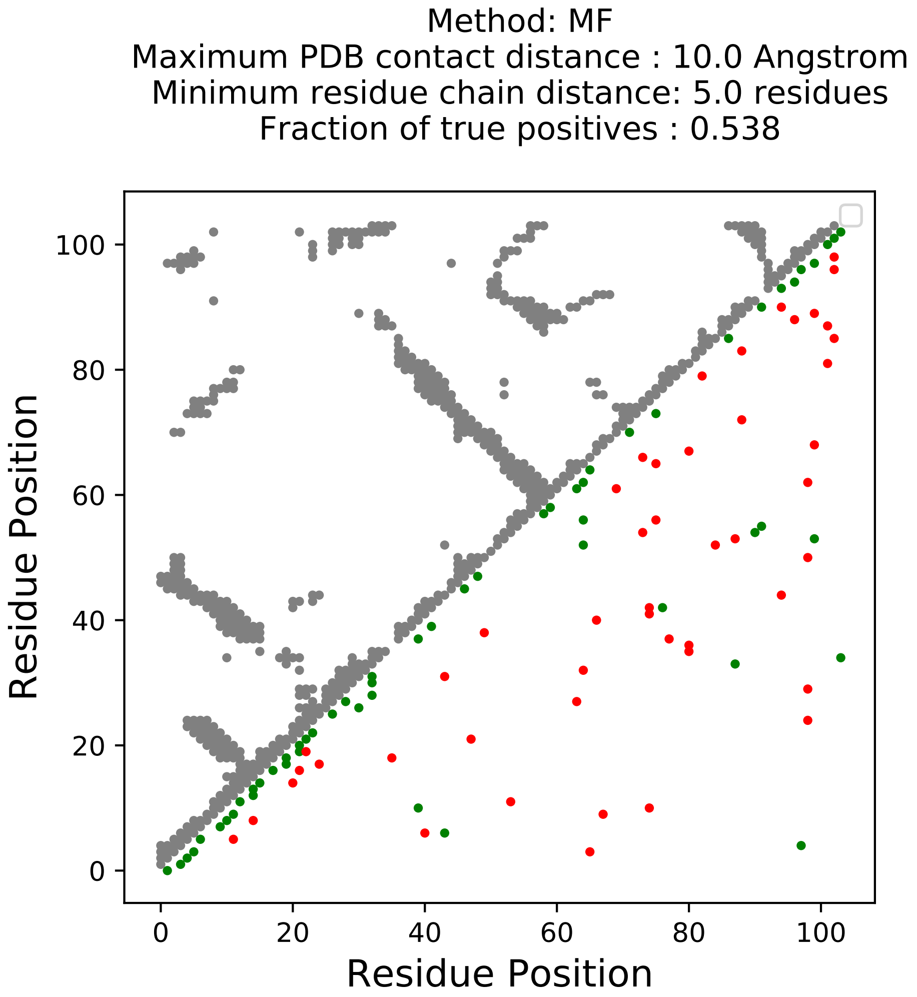

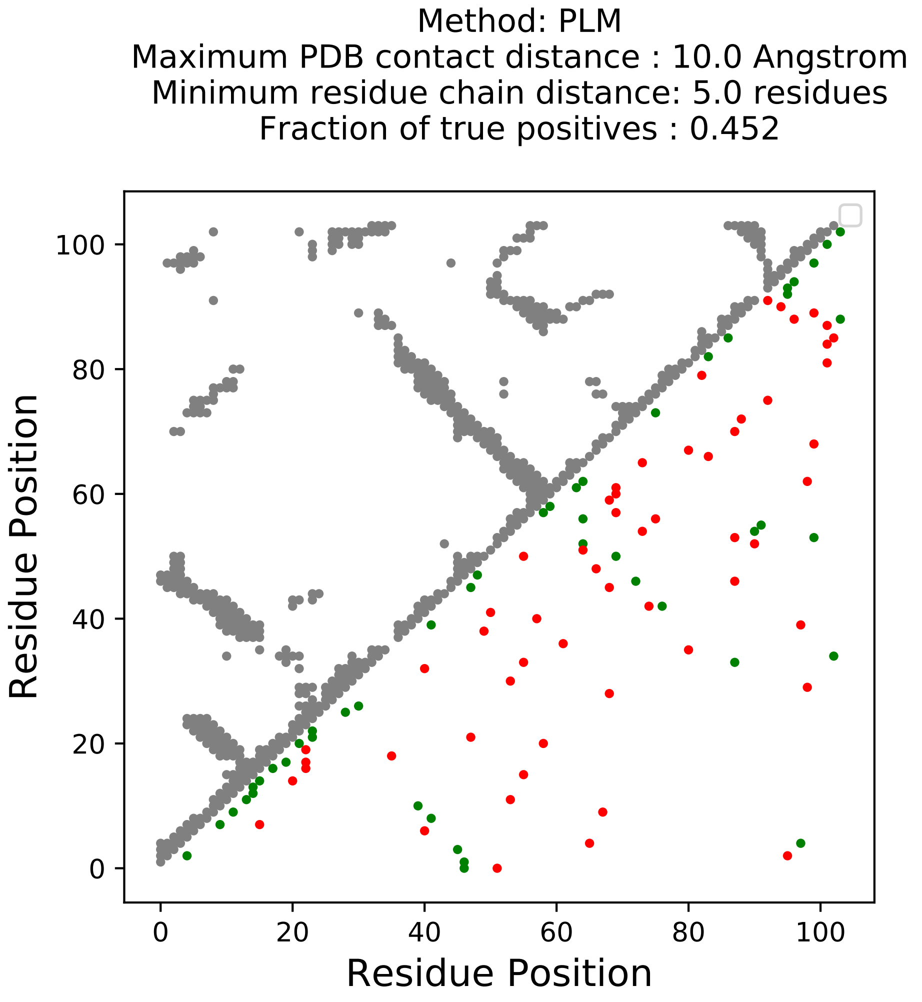

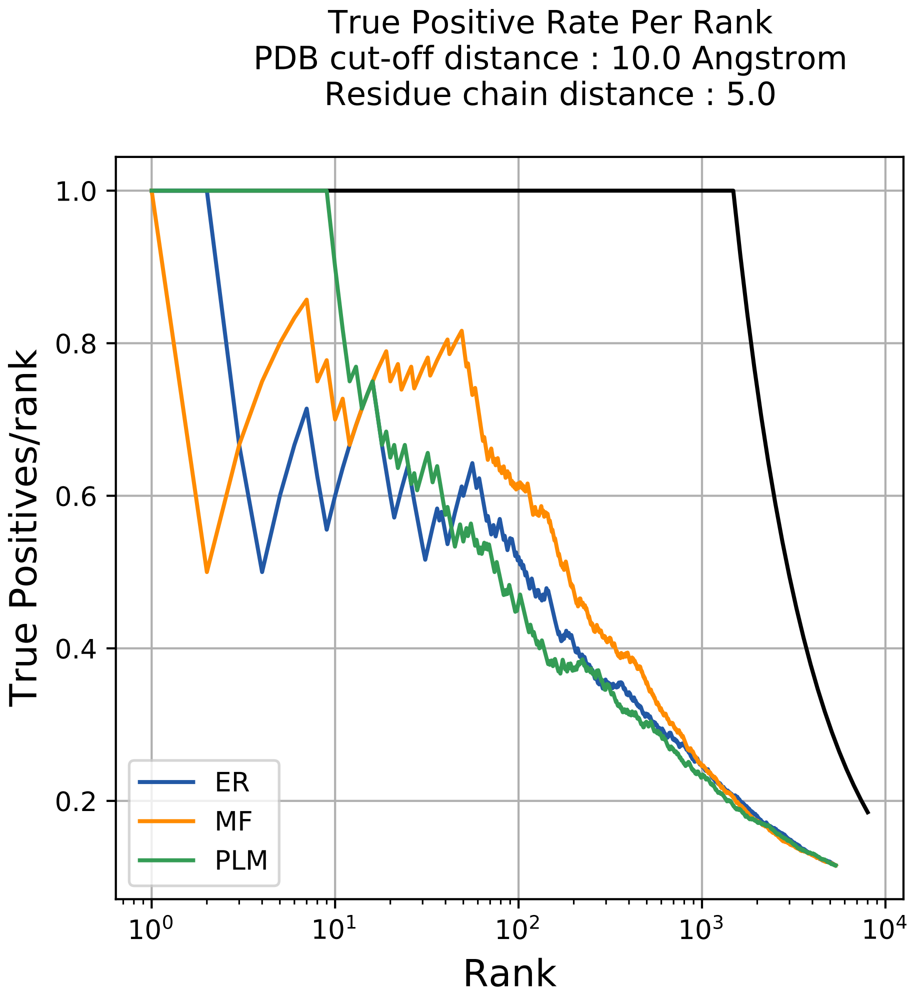

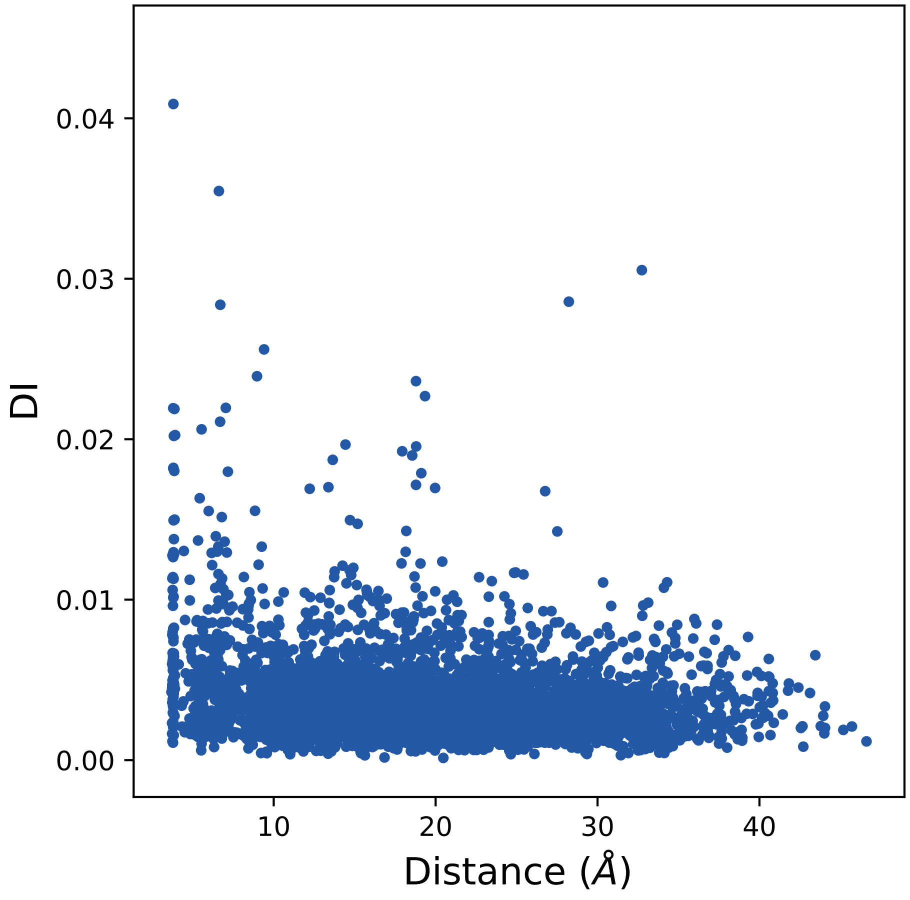

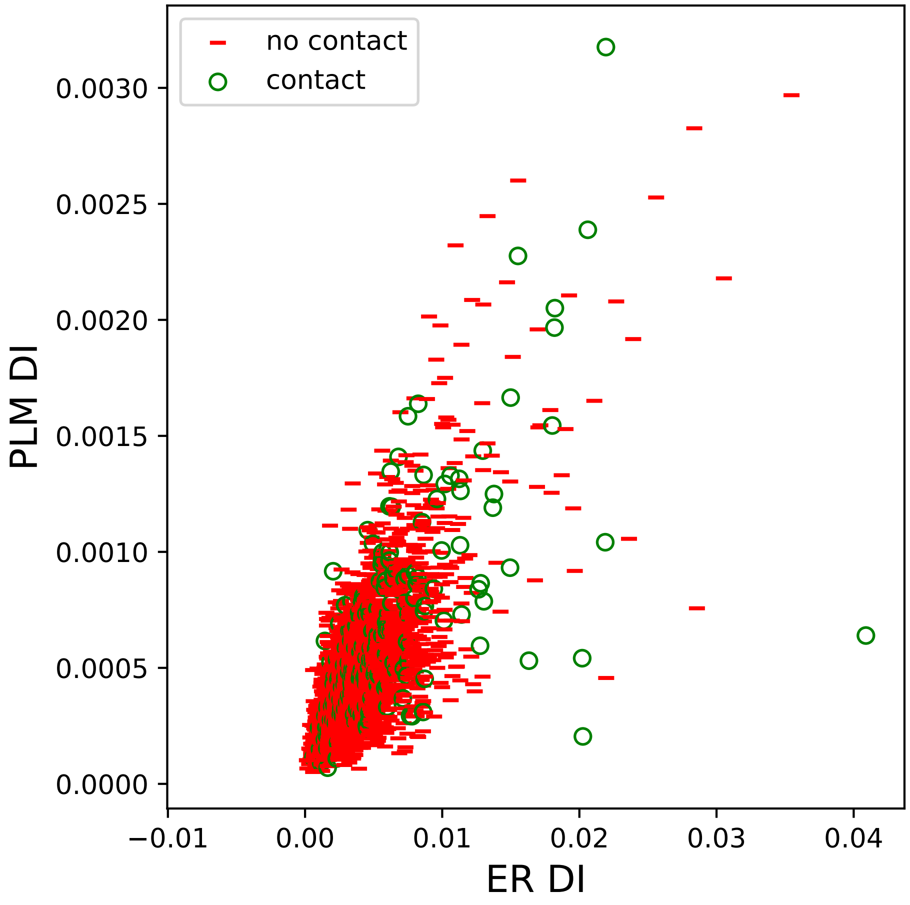

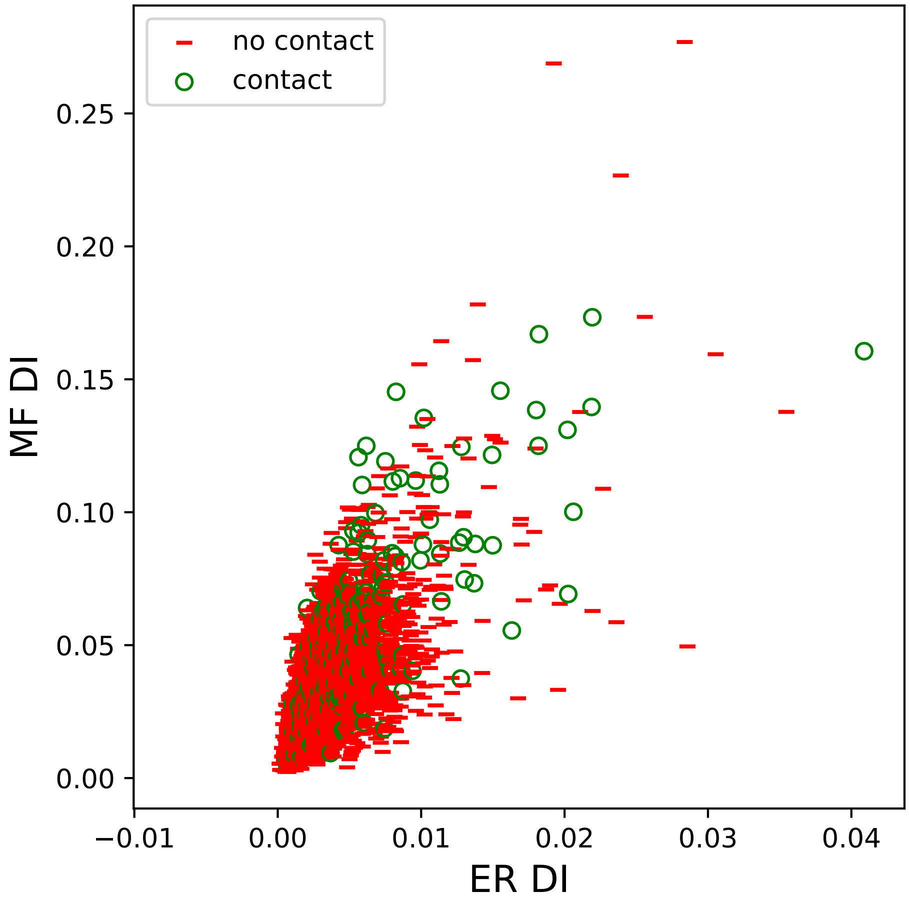

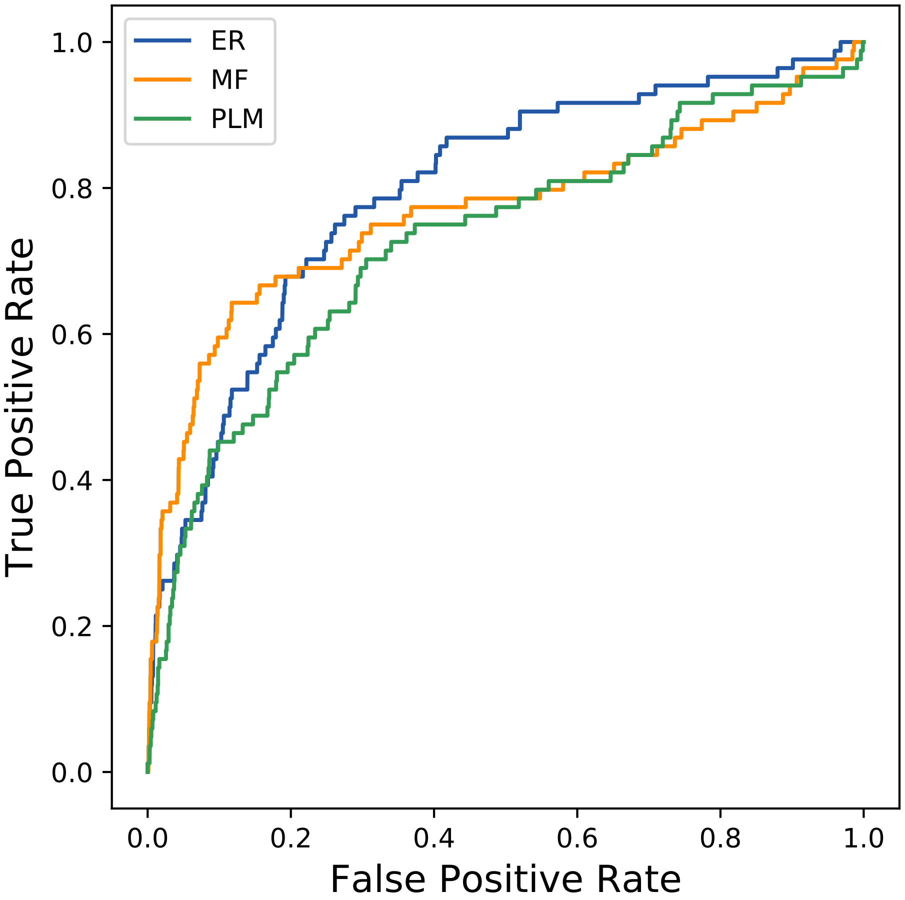

### 1257seq_84col_5q8a_PF07522_plots.pdf

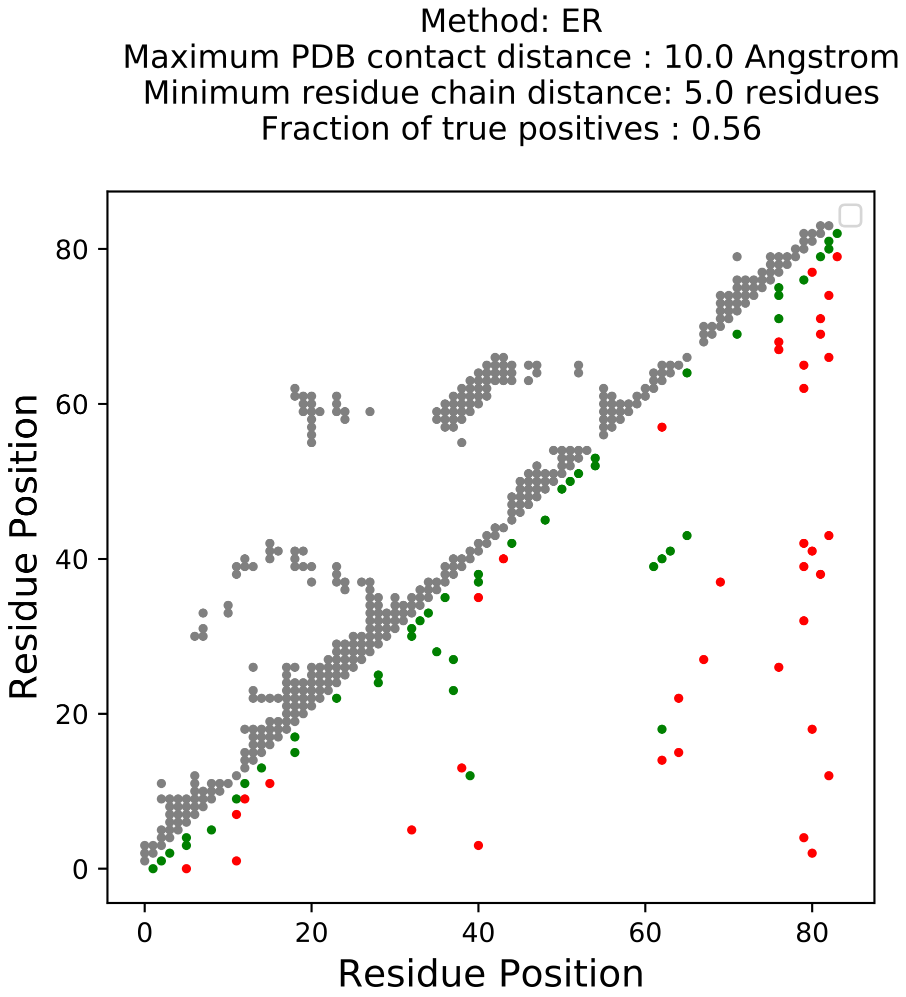

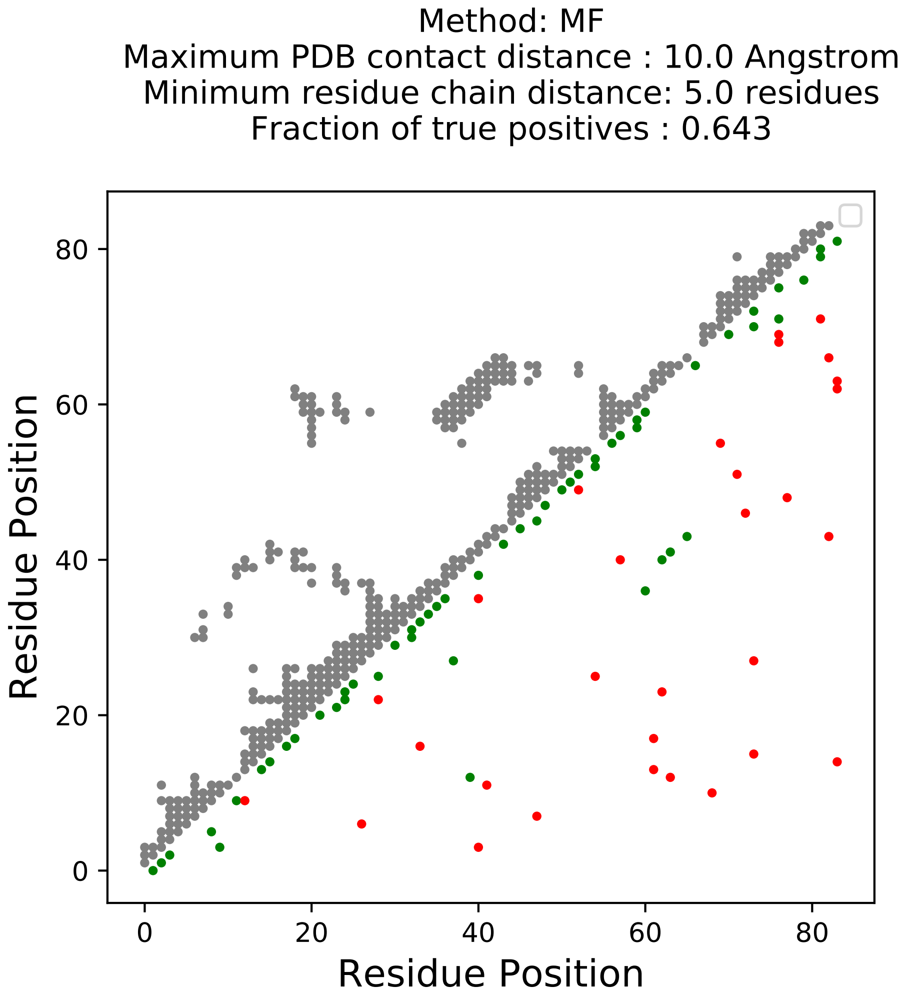

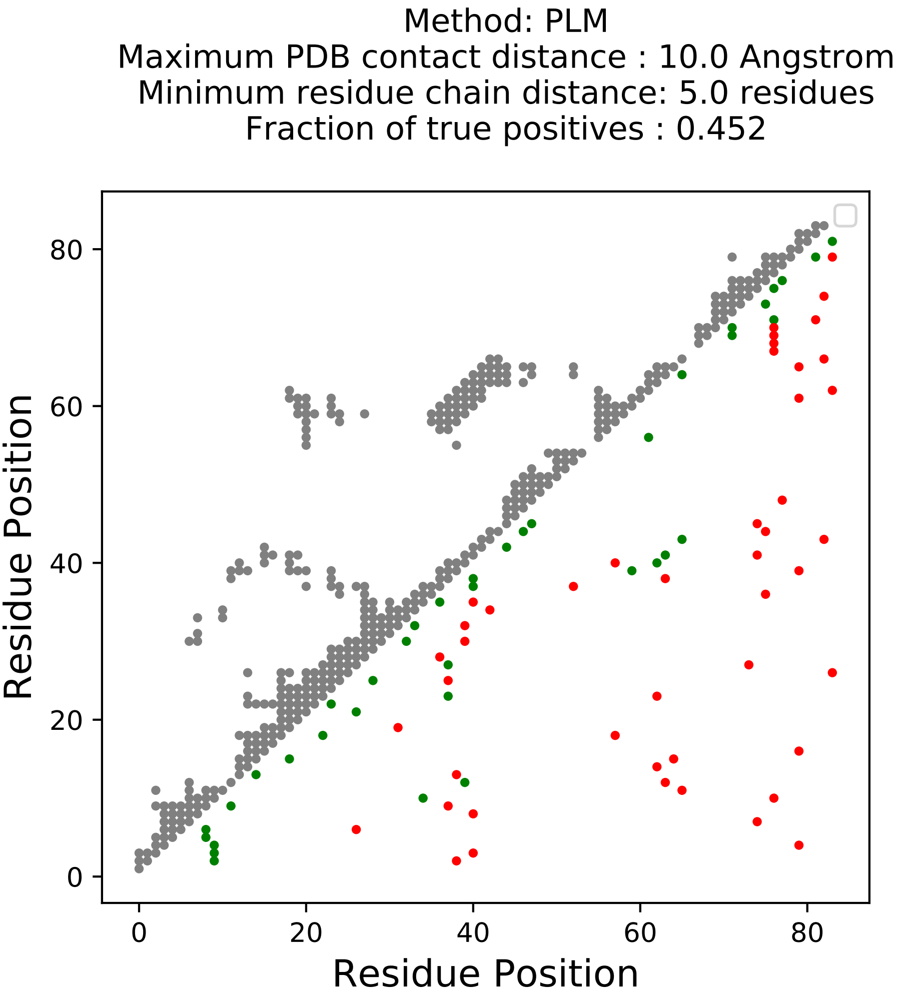

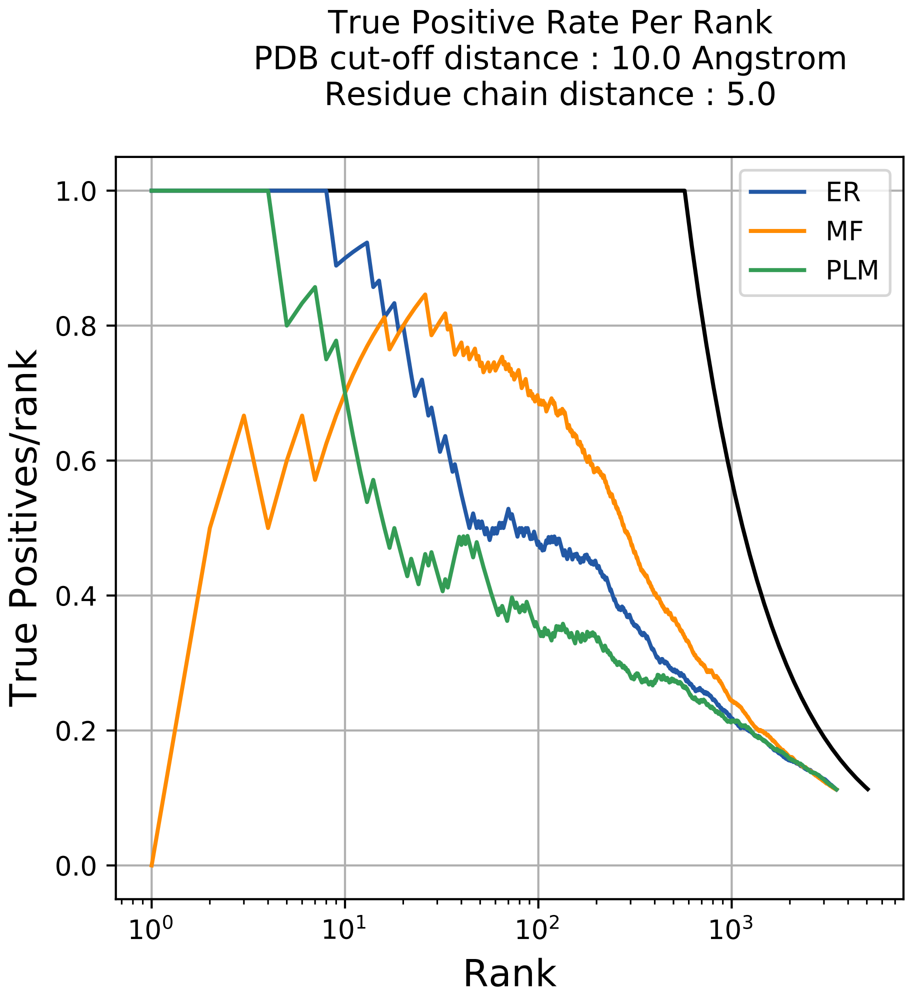

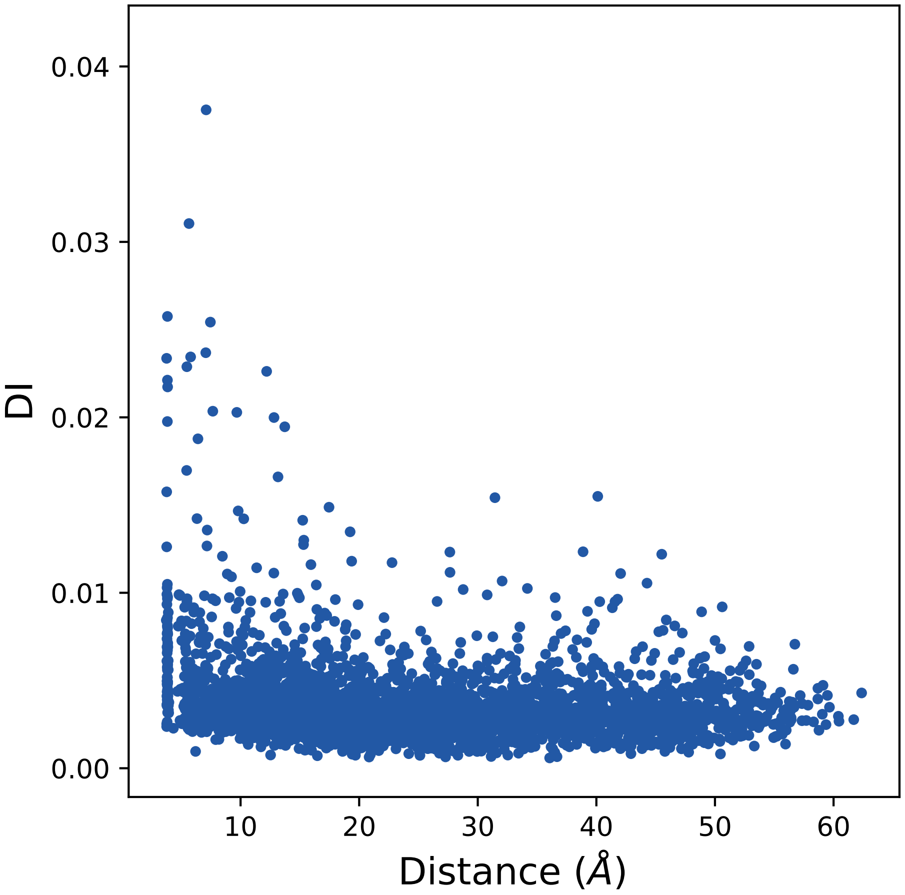

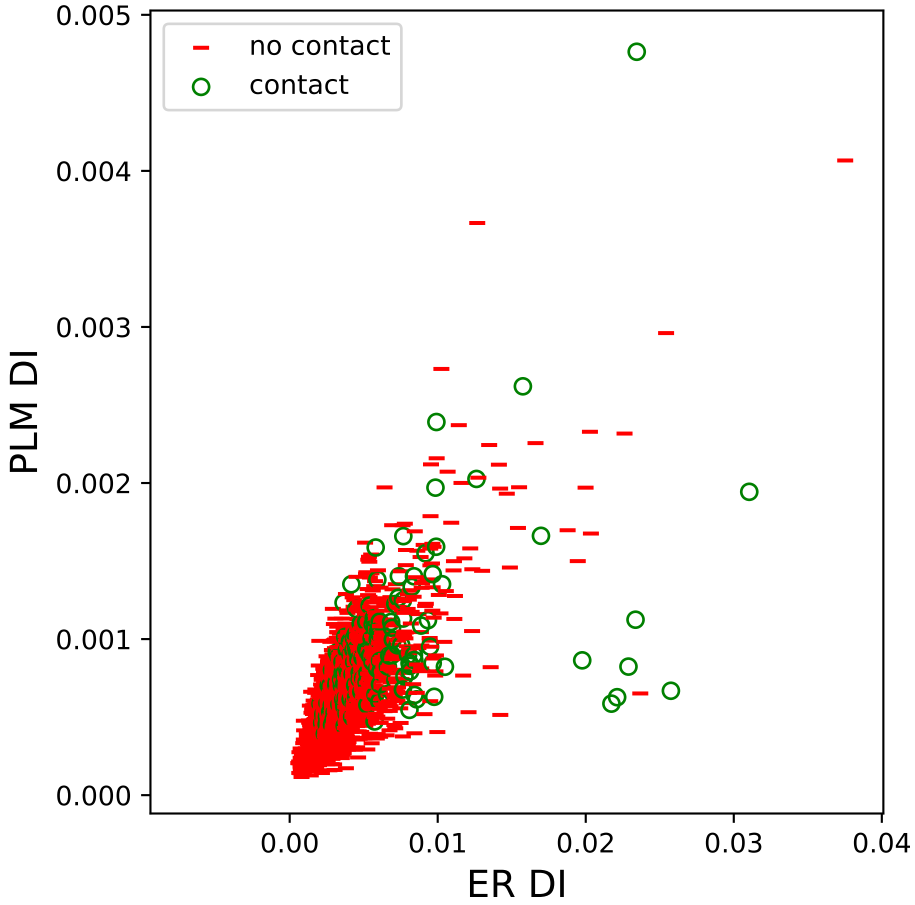

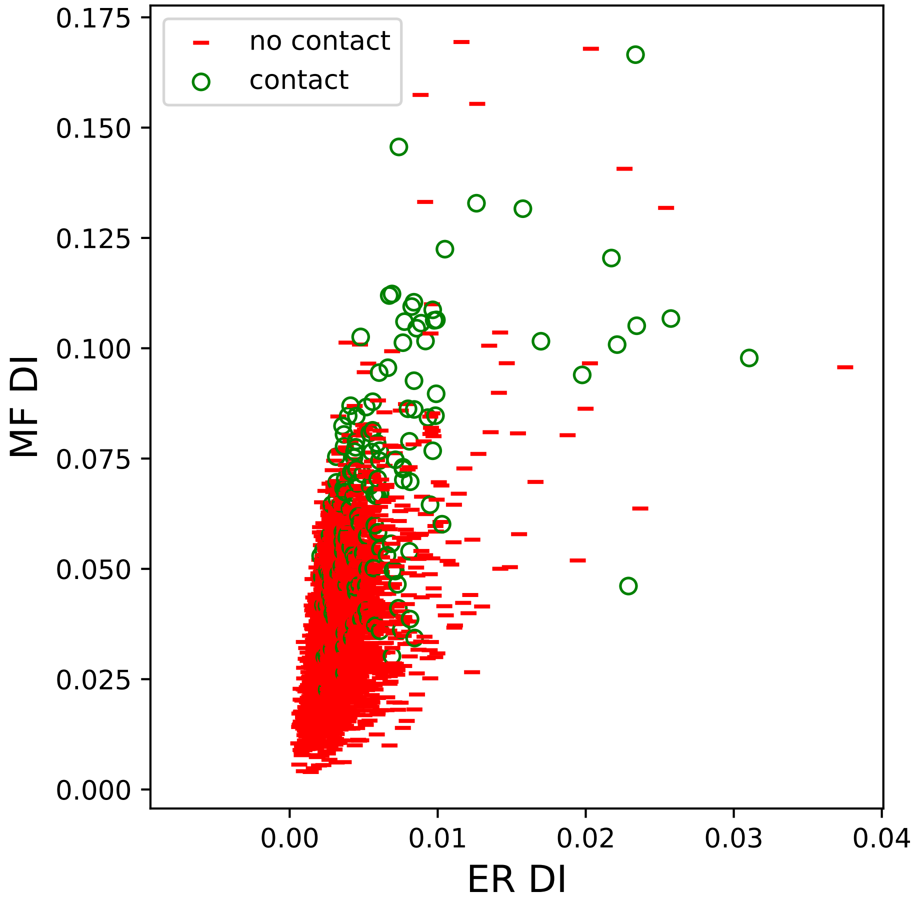

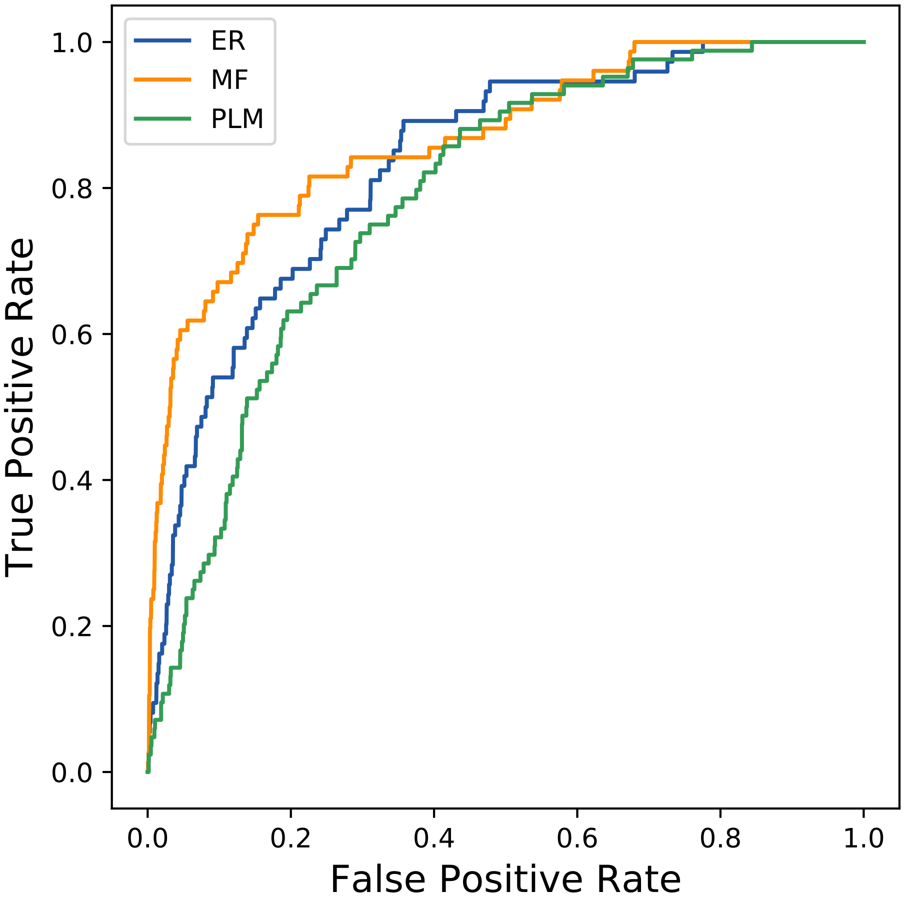

### 1257seq_84col_5q47_PF07522_plots.pdf

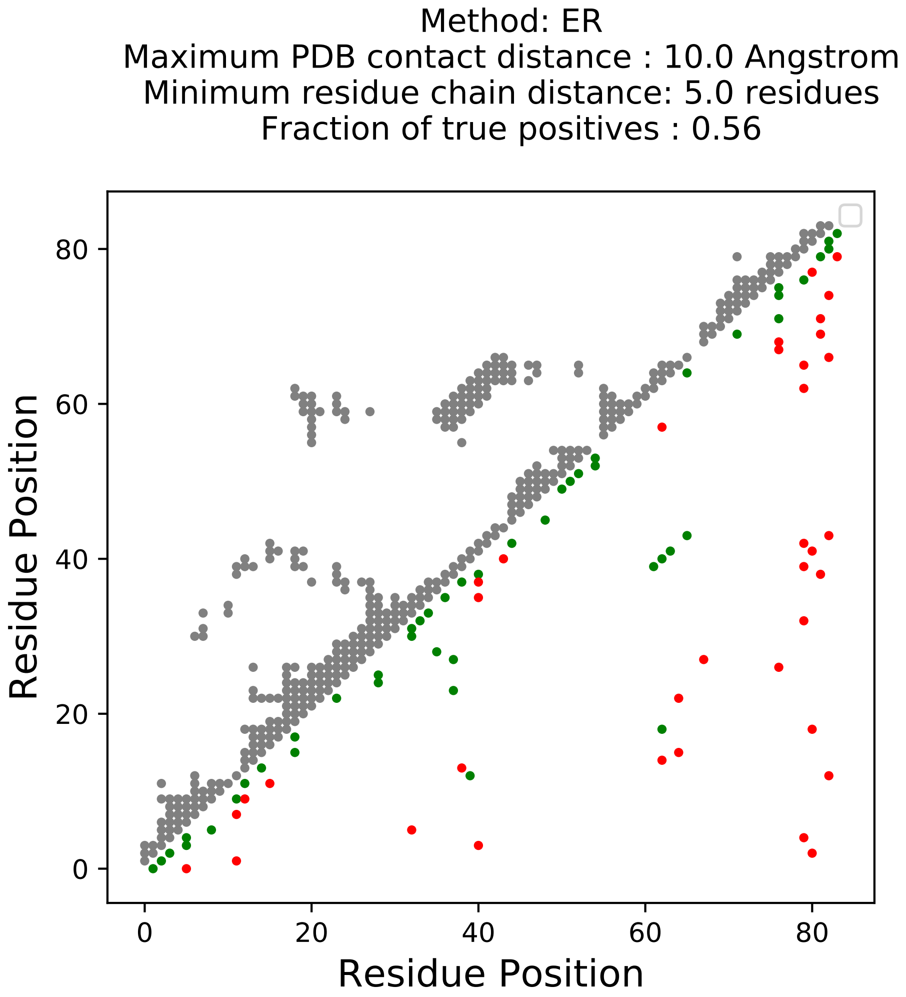

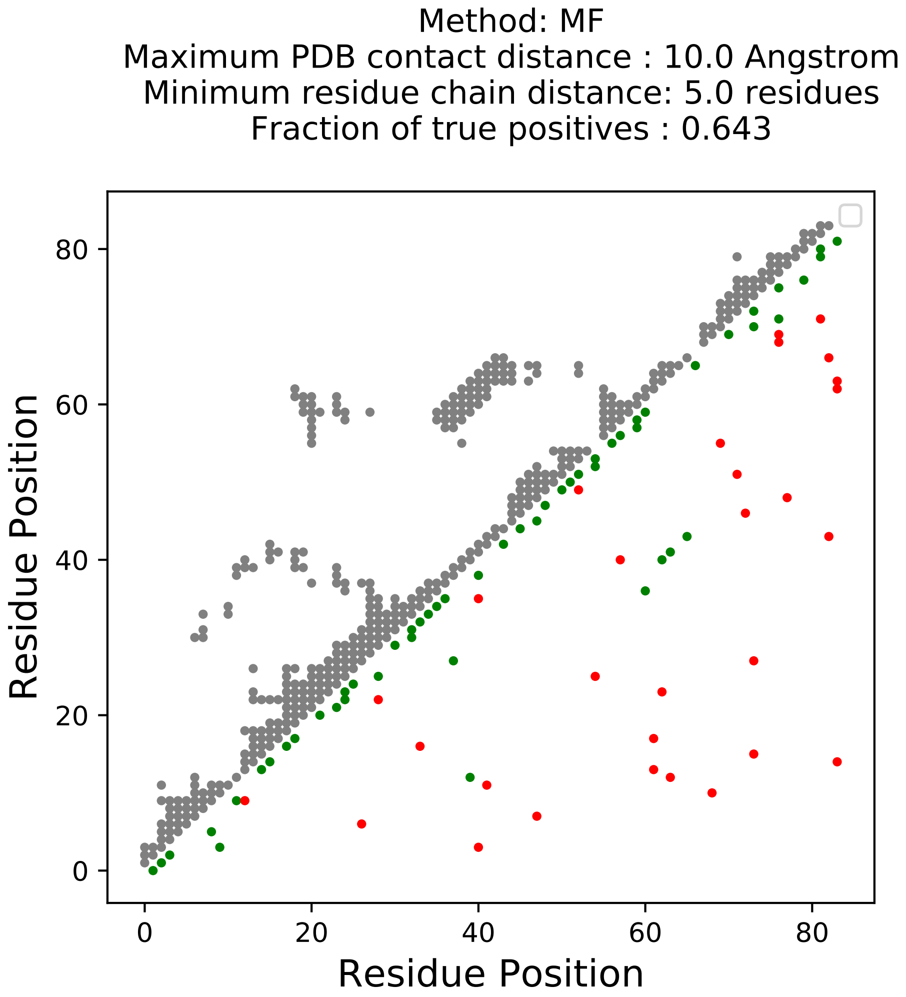

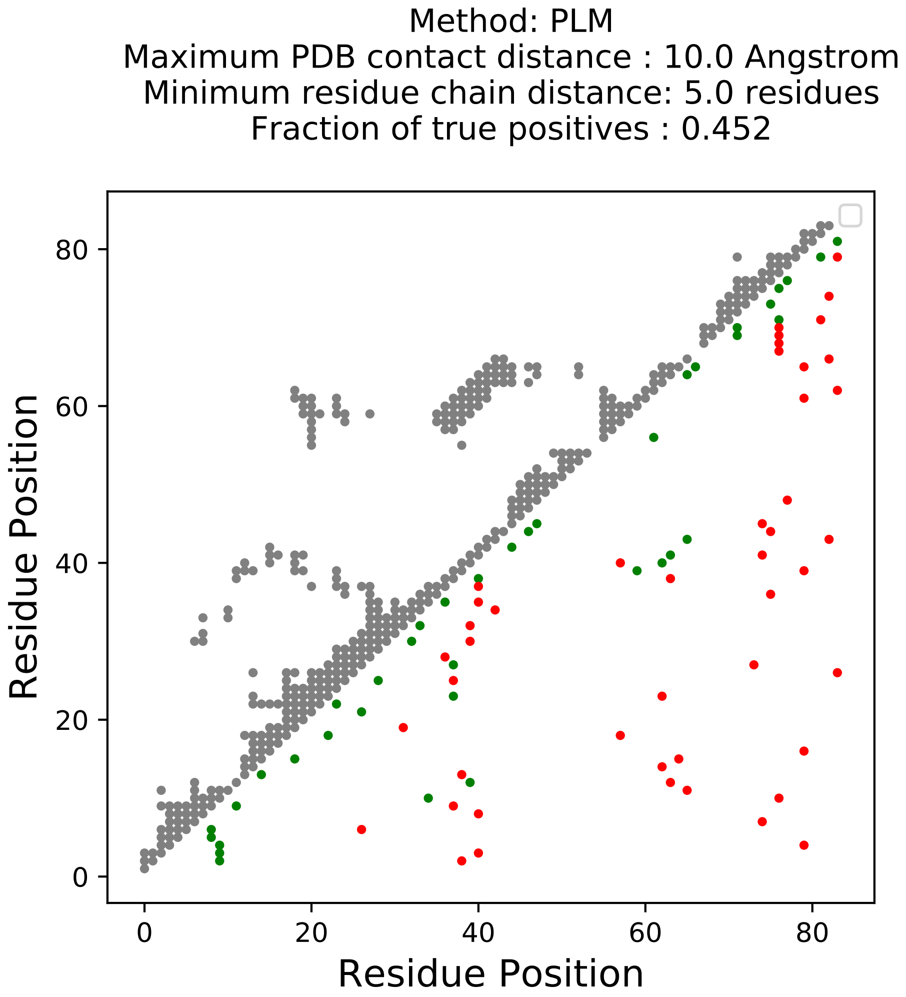

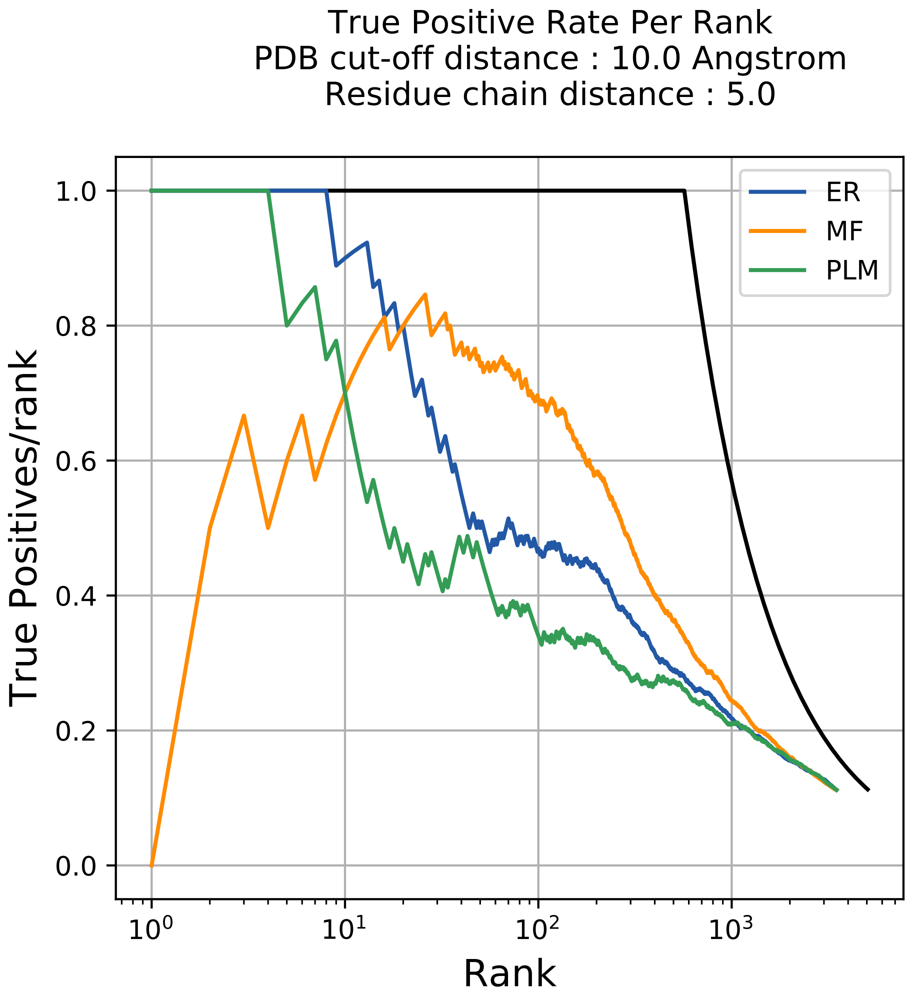

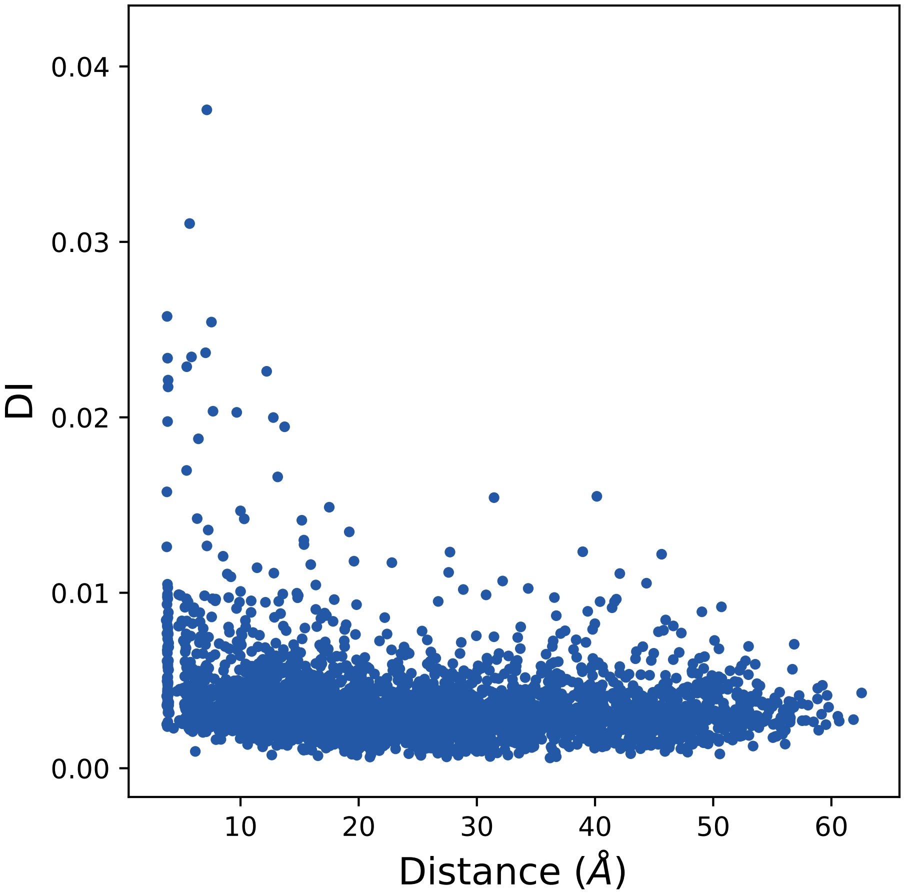

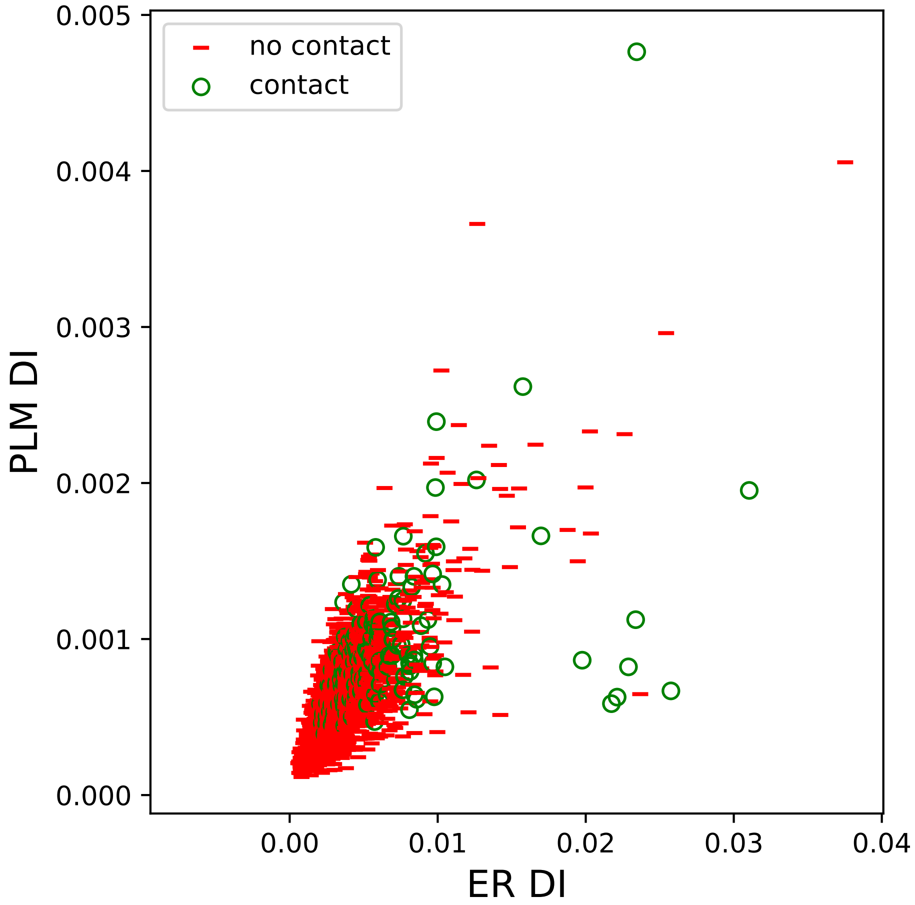

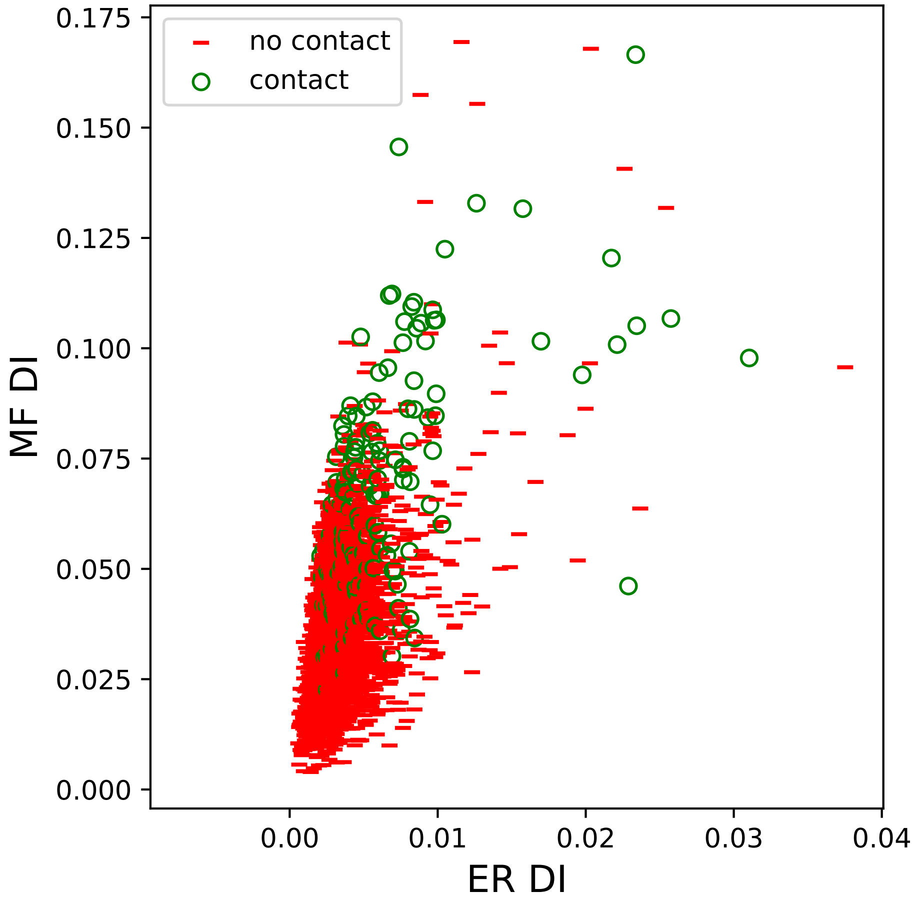

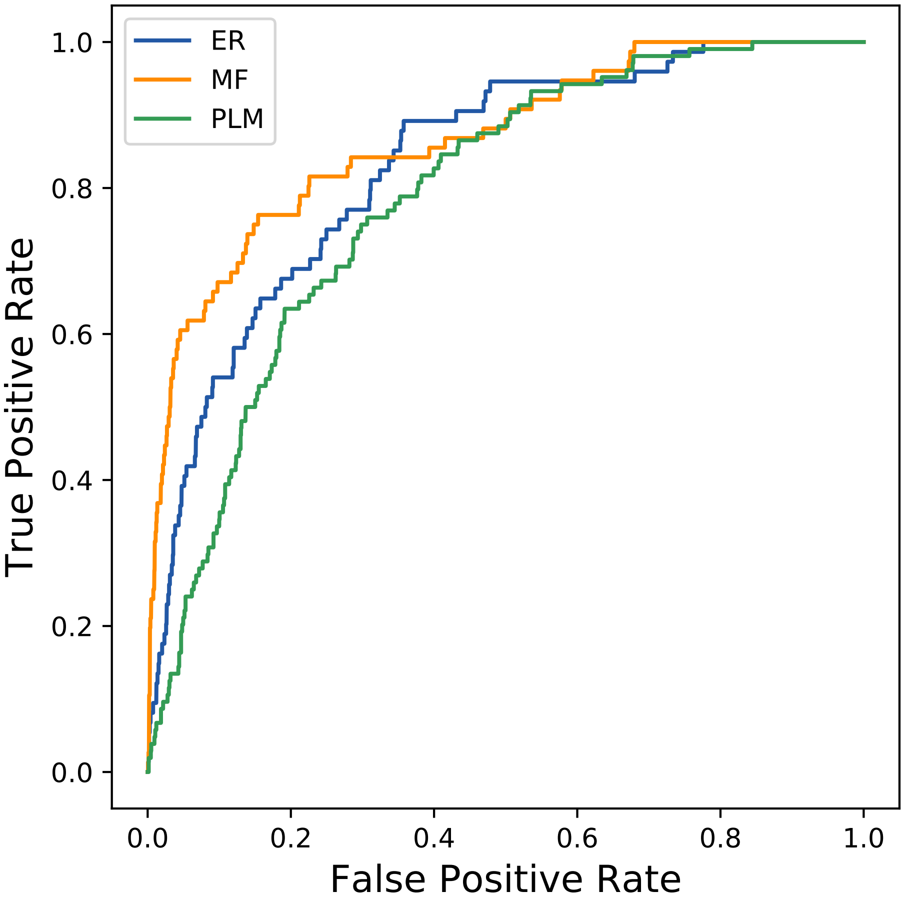

### 1400seq_49col_4rv9_PF08484_plots.pdf

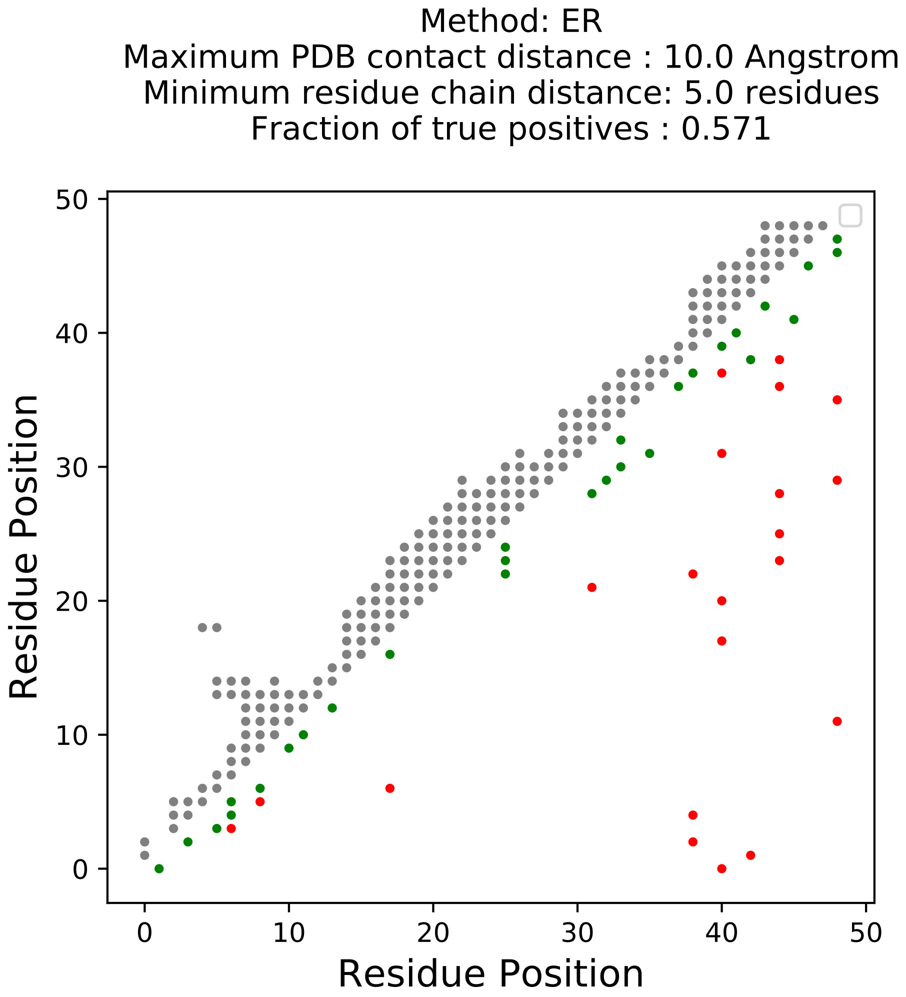

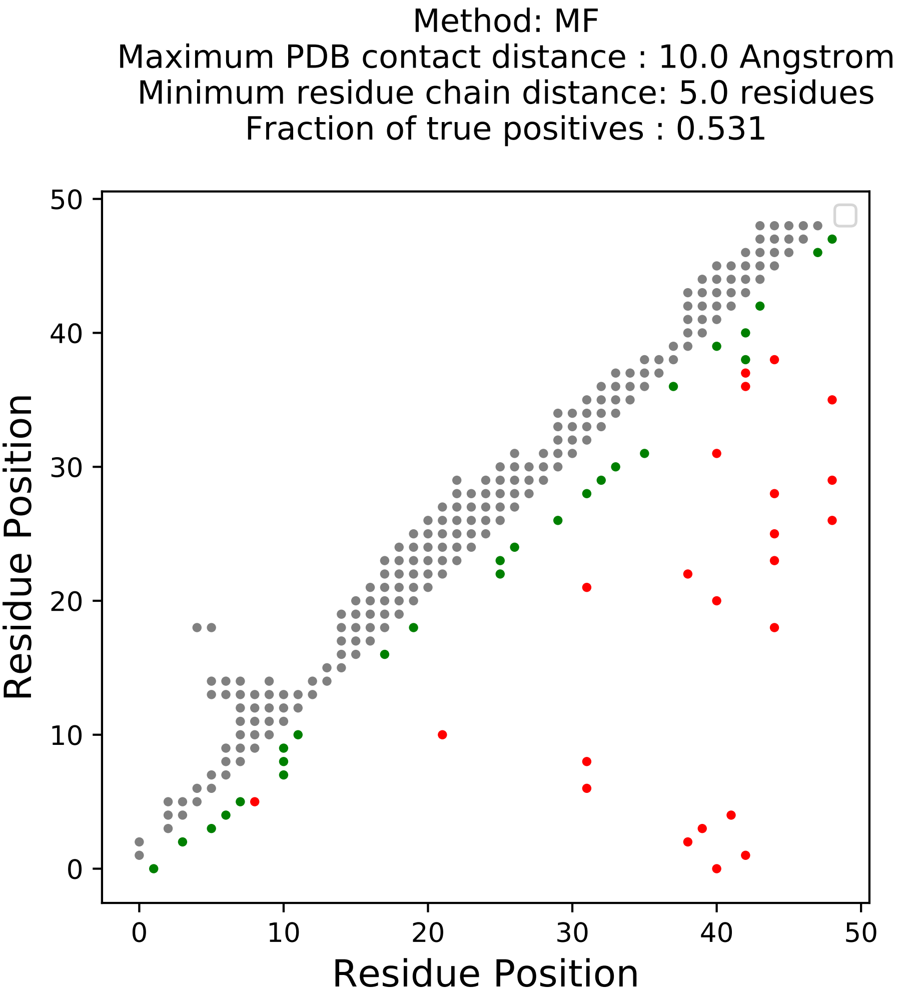

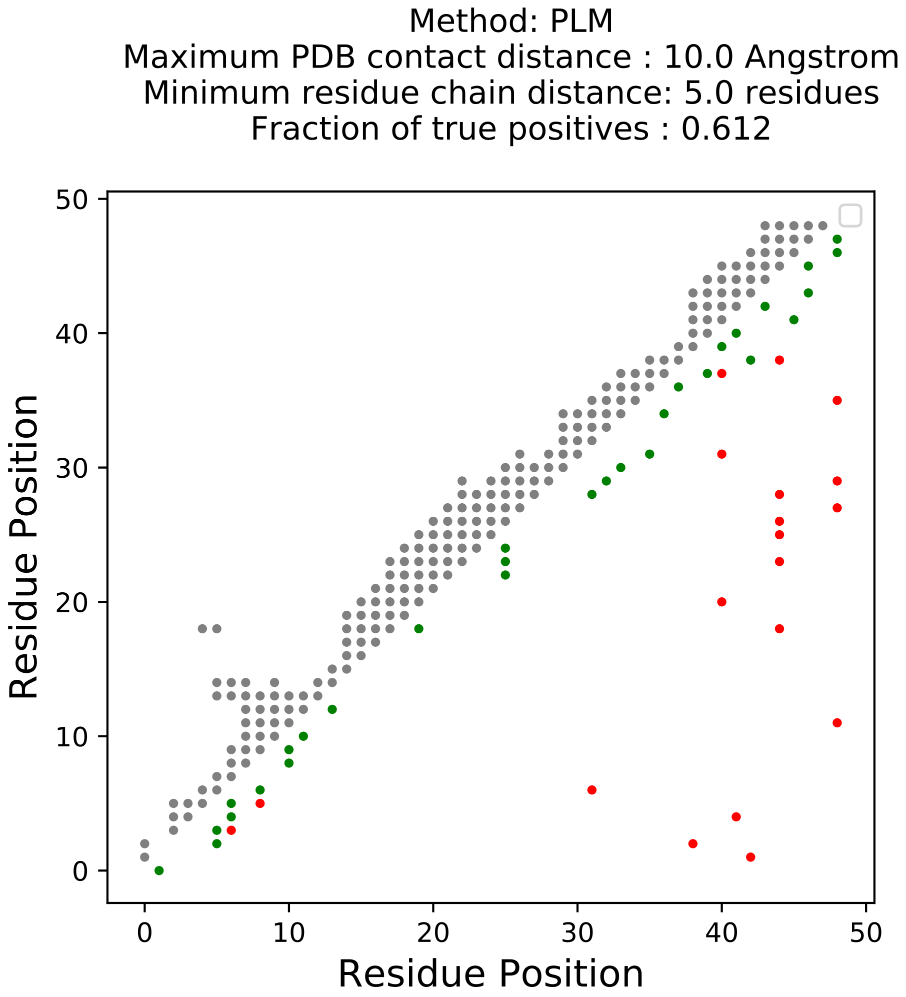

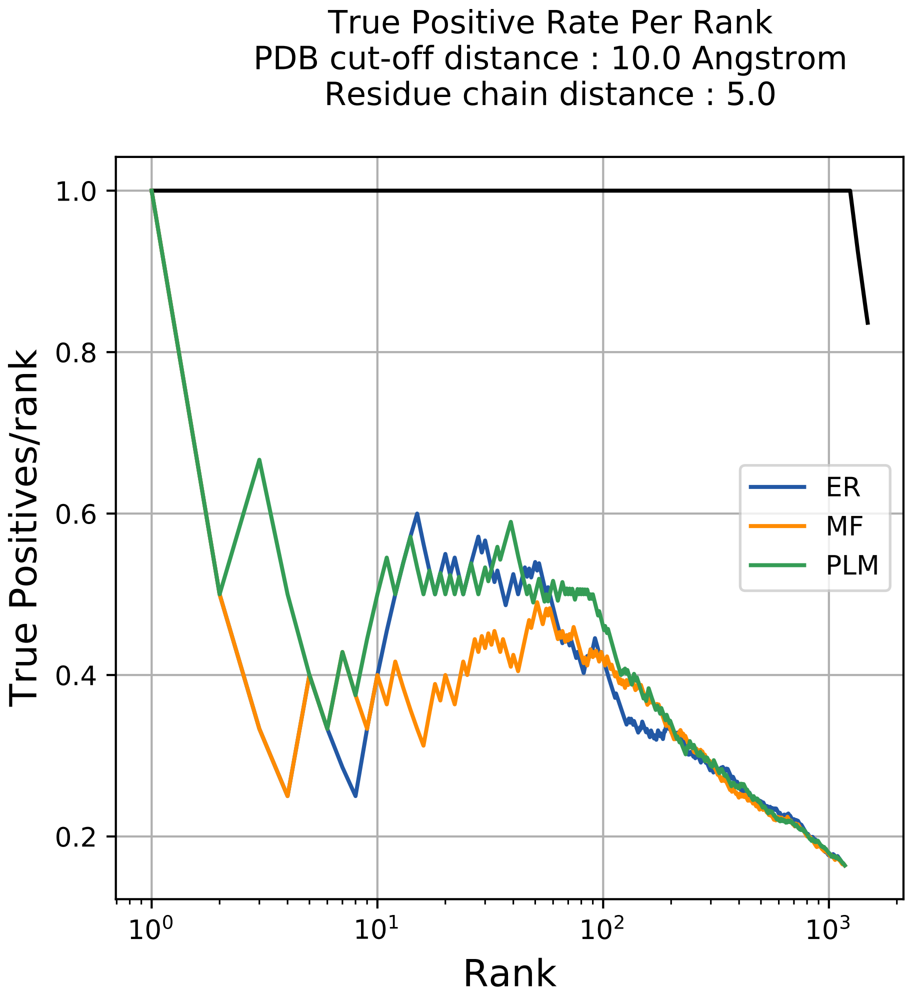

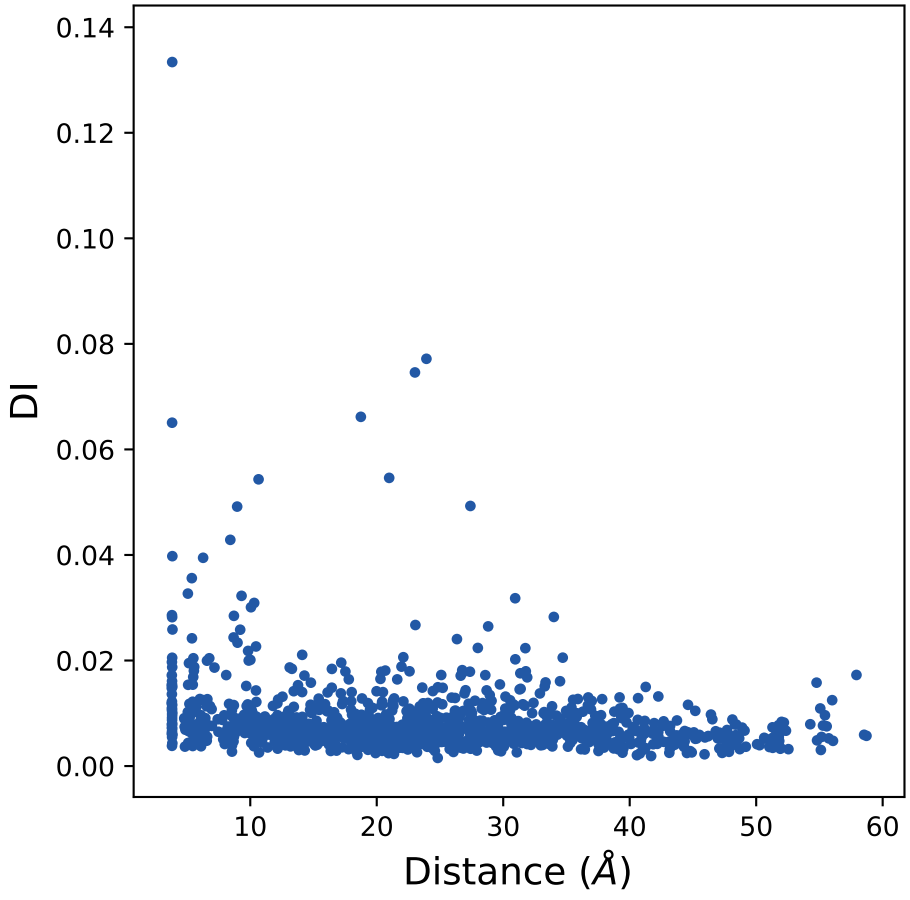

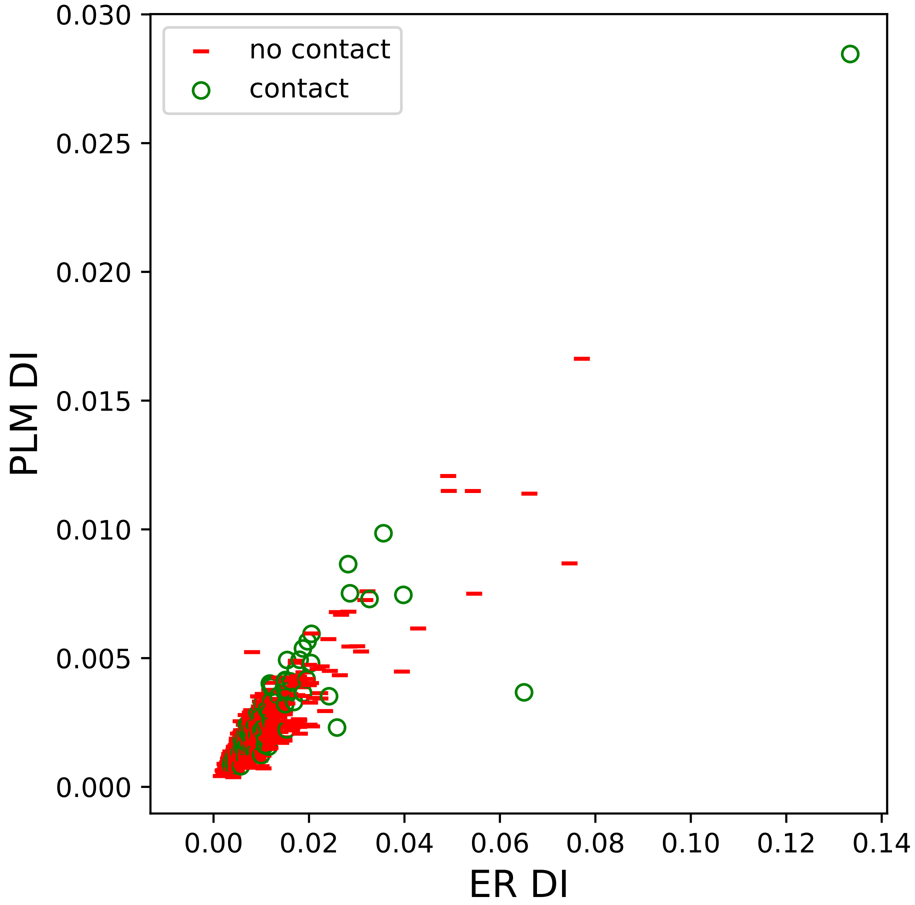

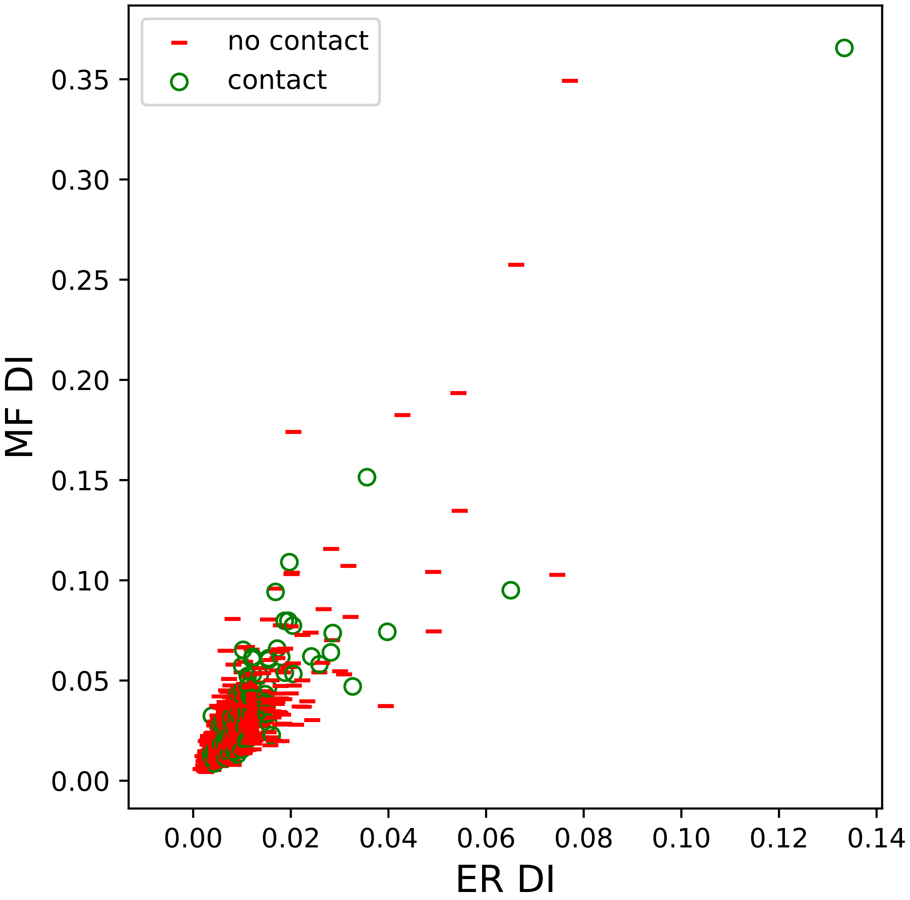

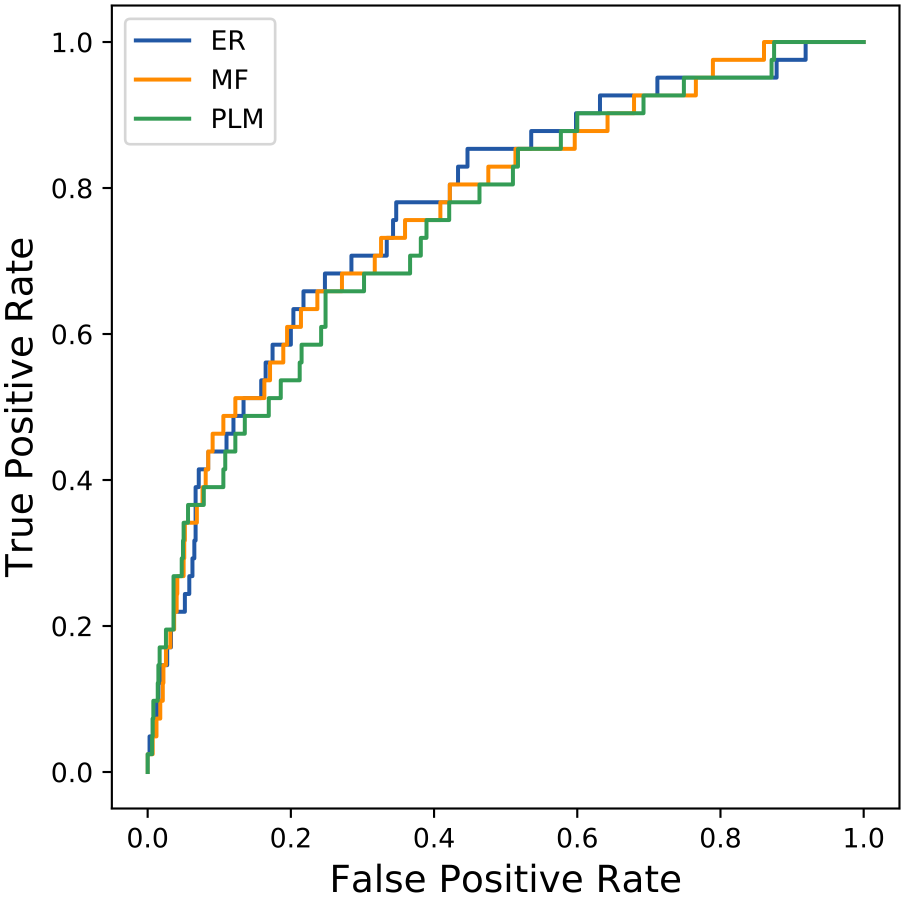

### 1423seq_156col_3pwl_PF00129_plots.pdf

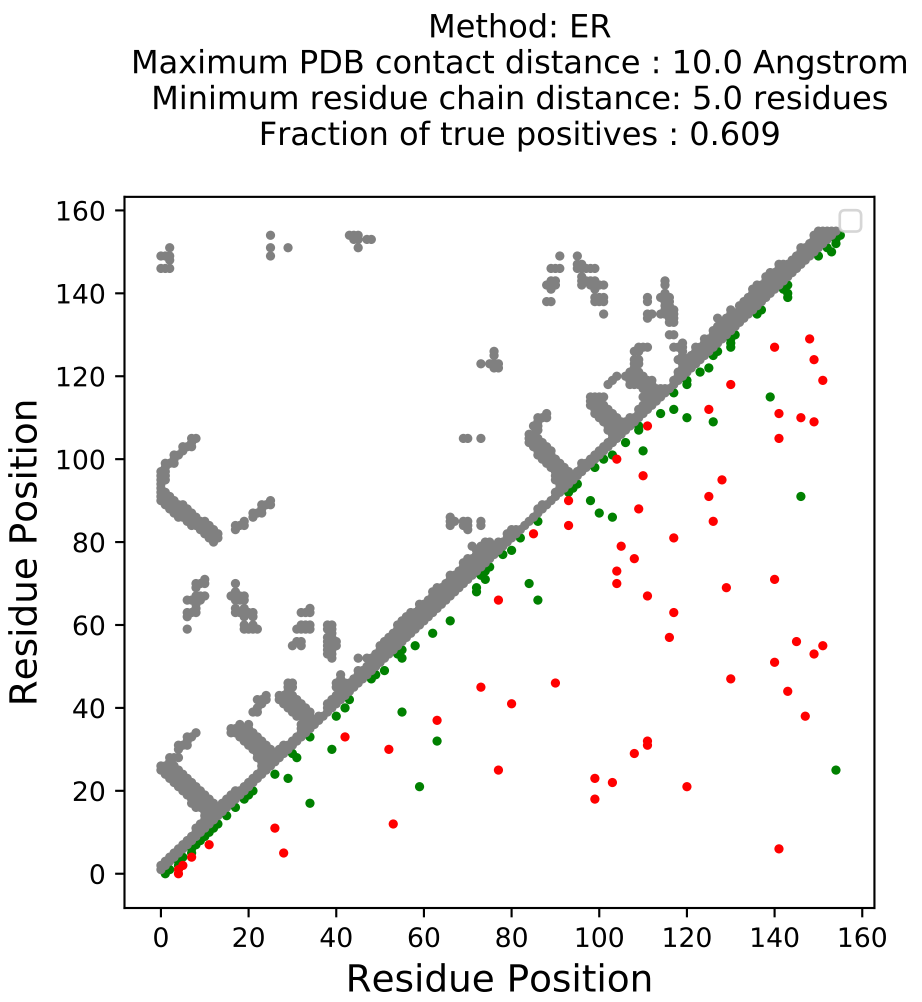

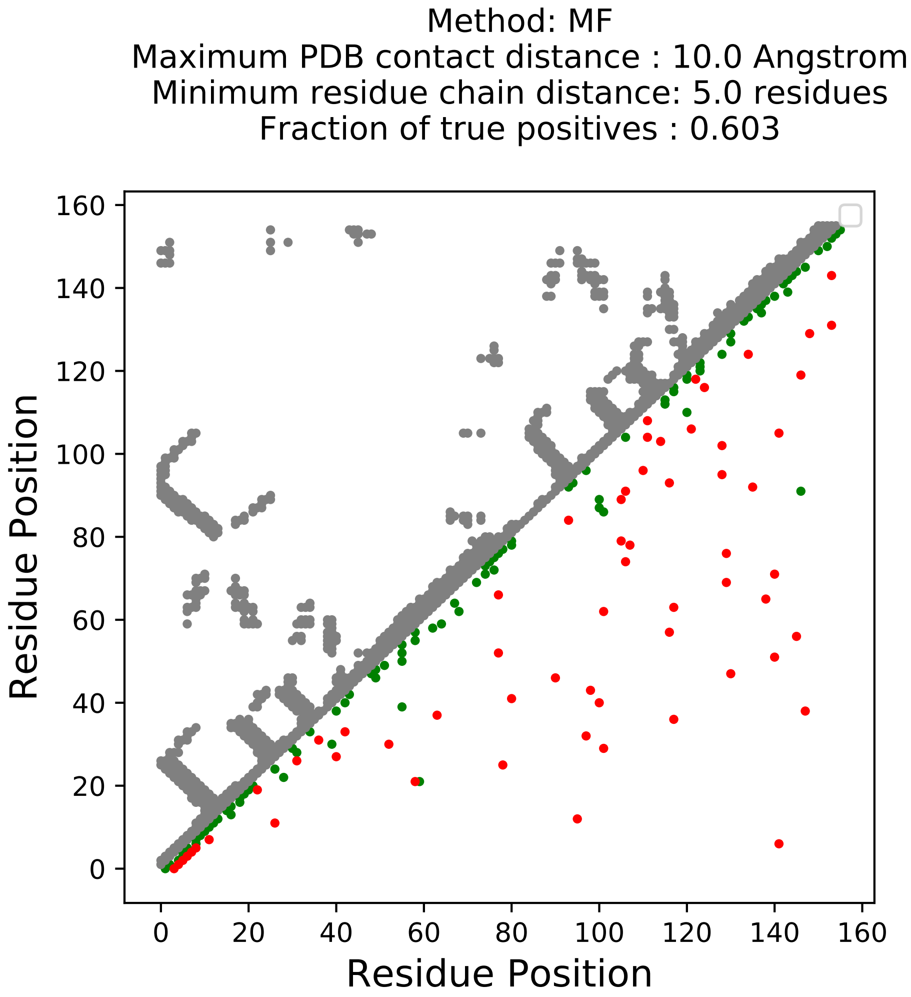

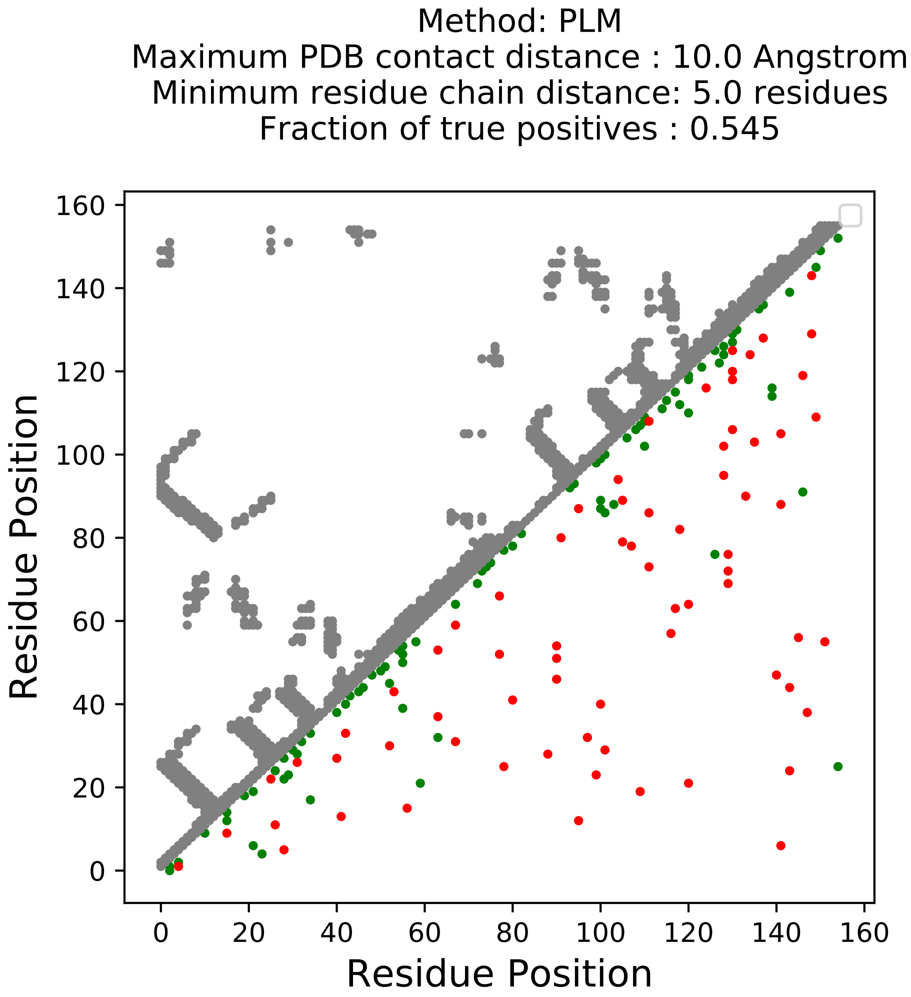

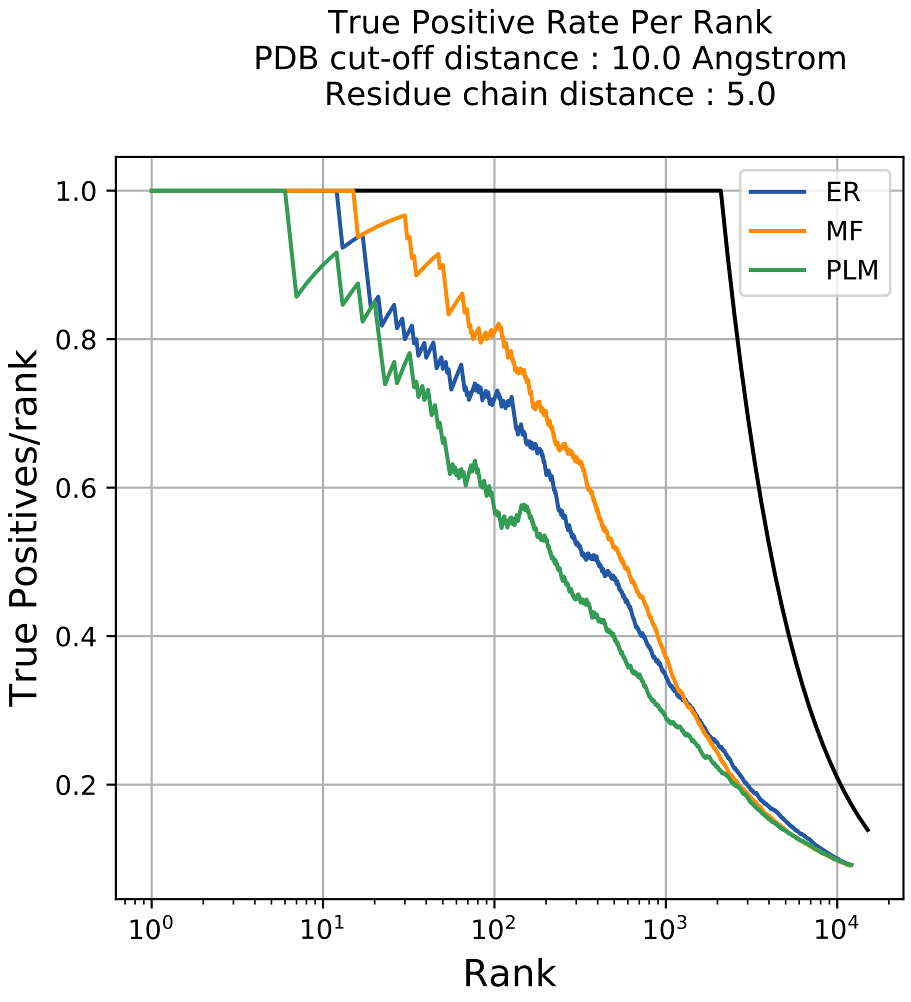

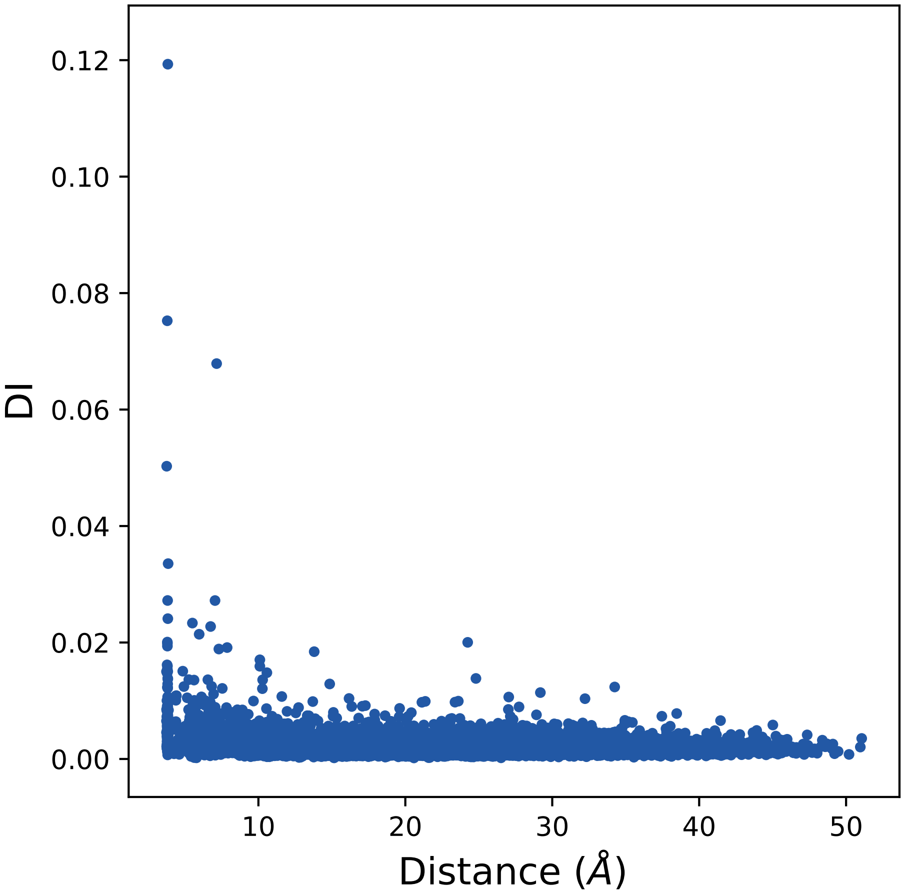

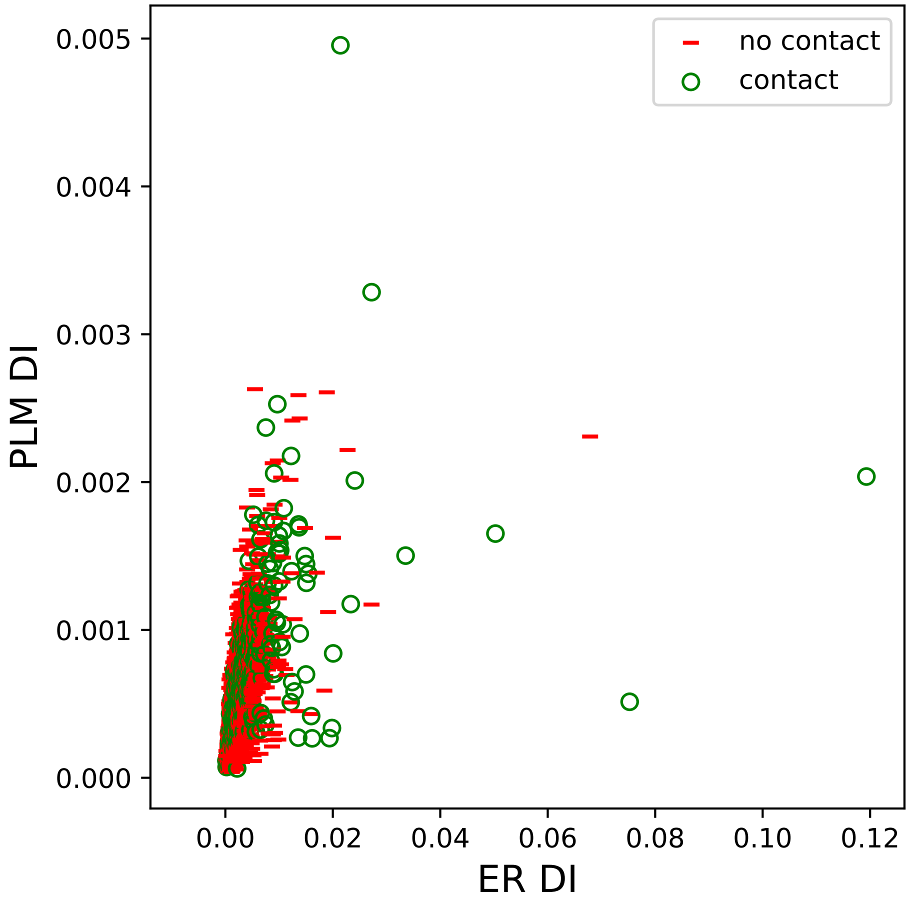

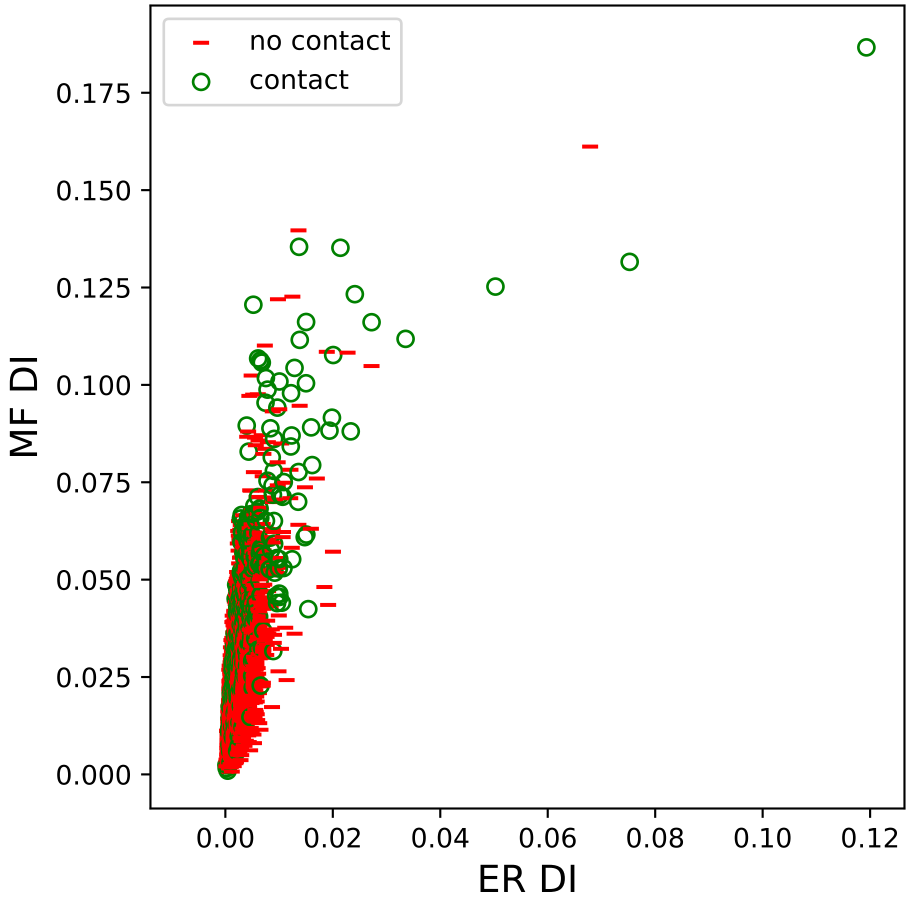

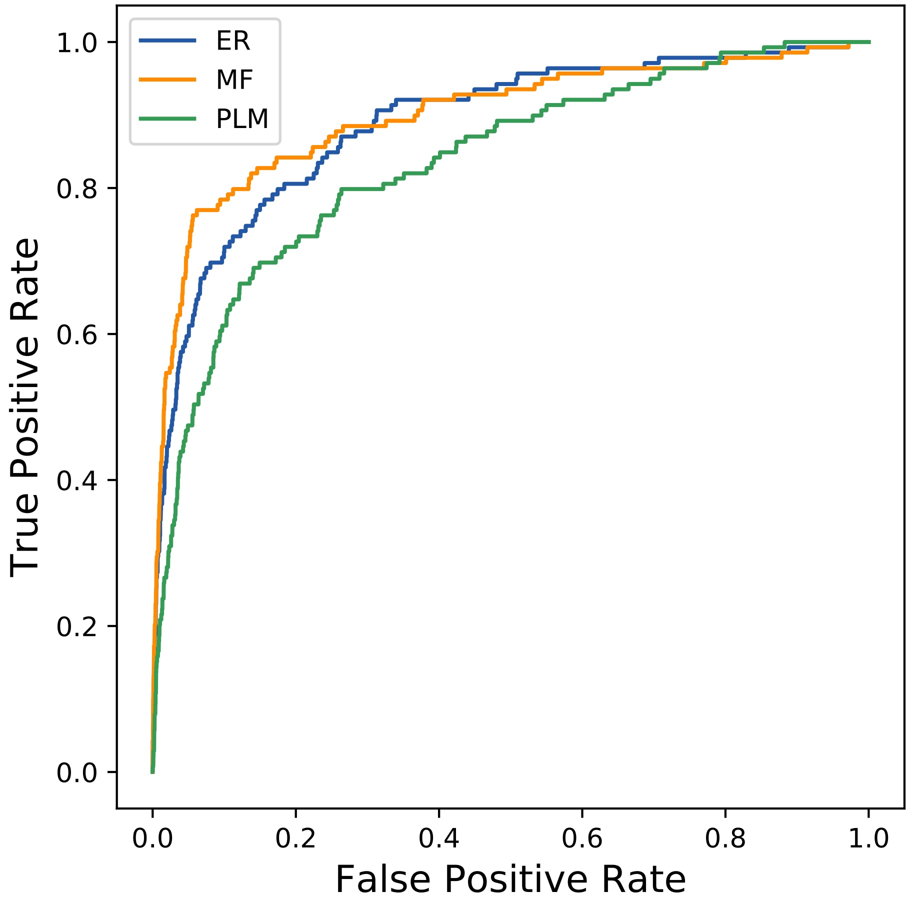

### 1432seq_151col_4s0u_PF00129_plots.pdf

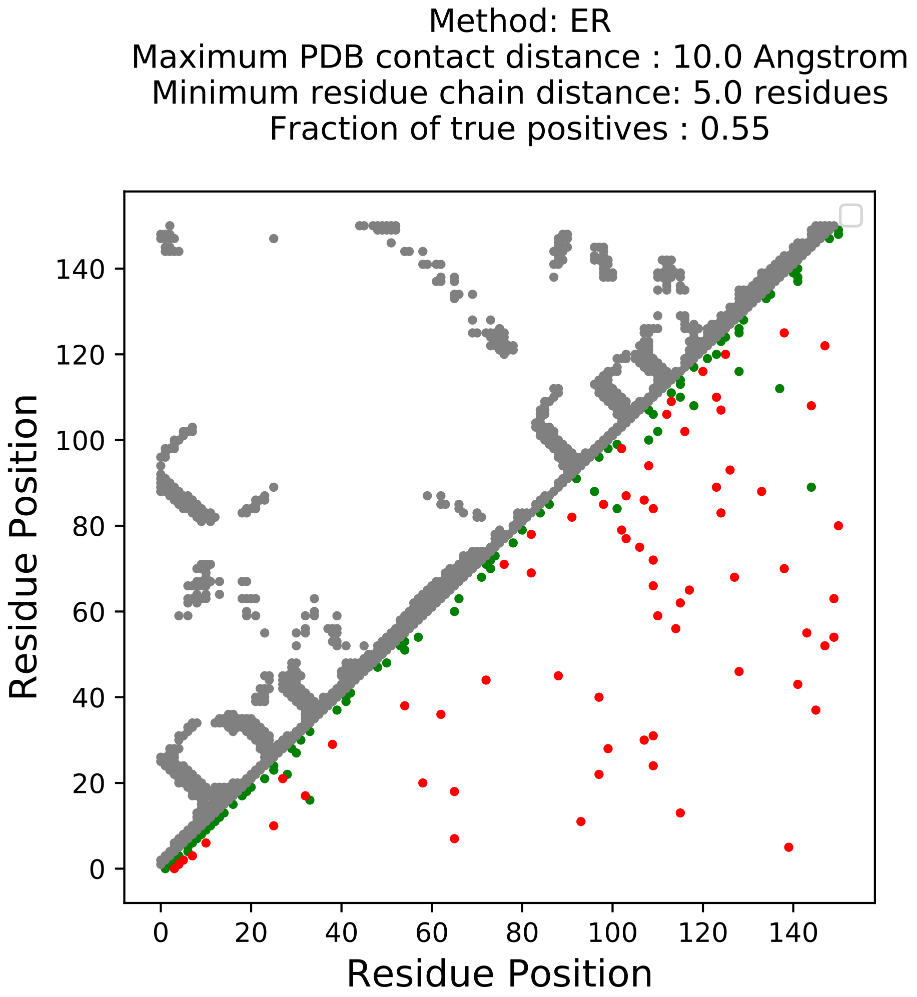

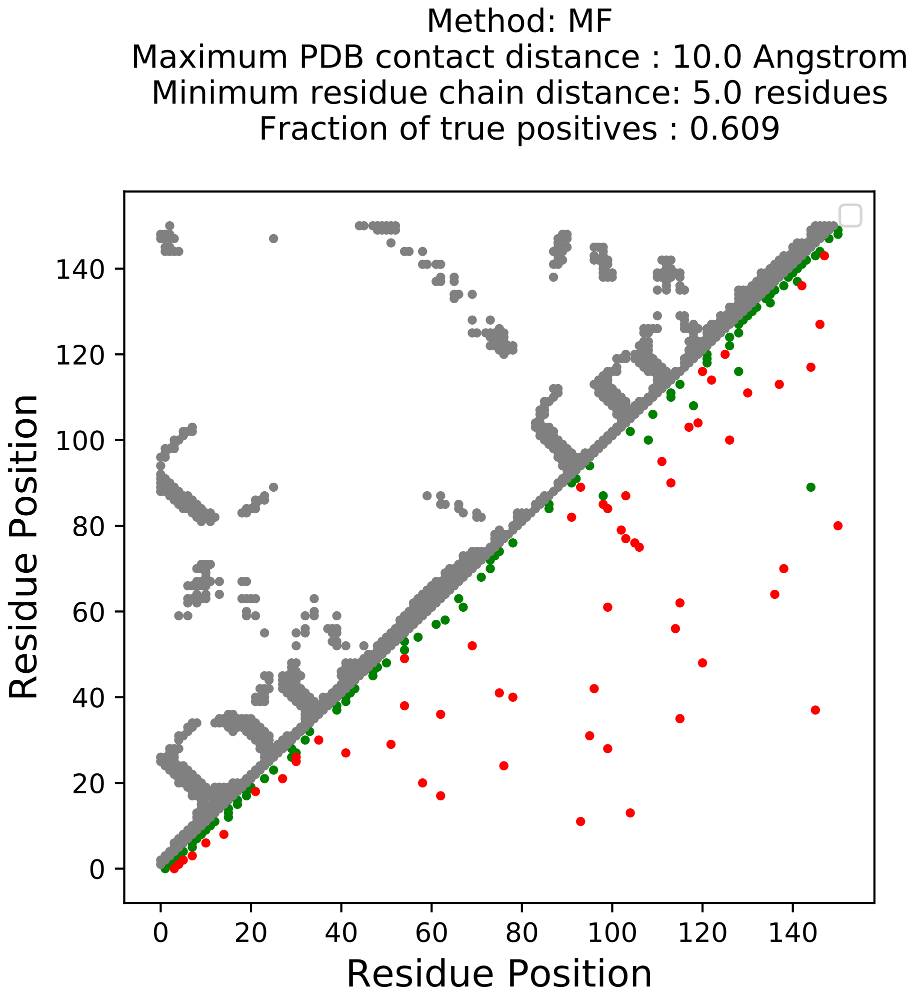

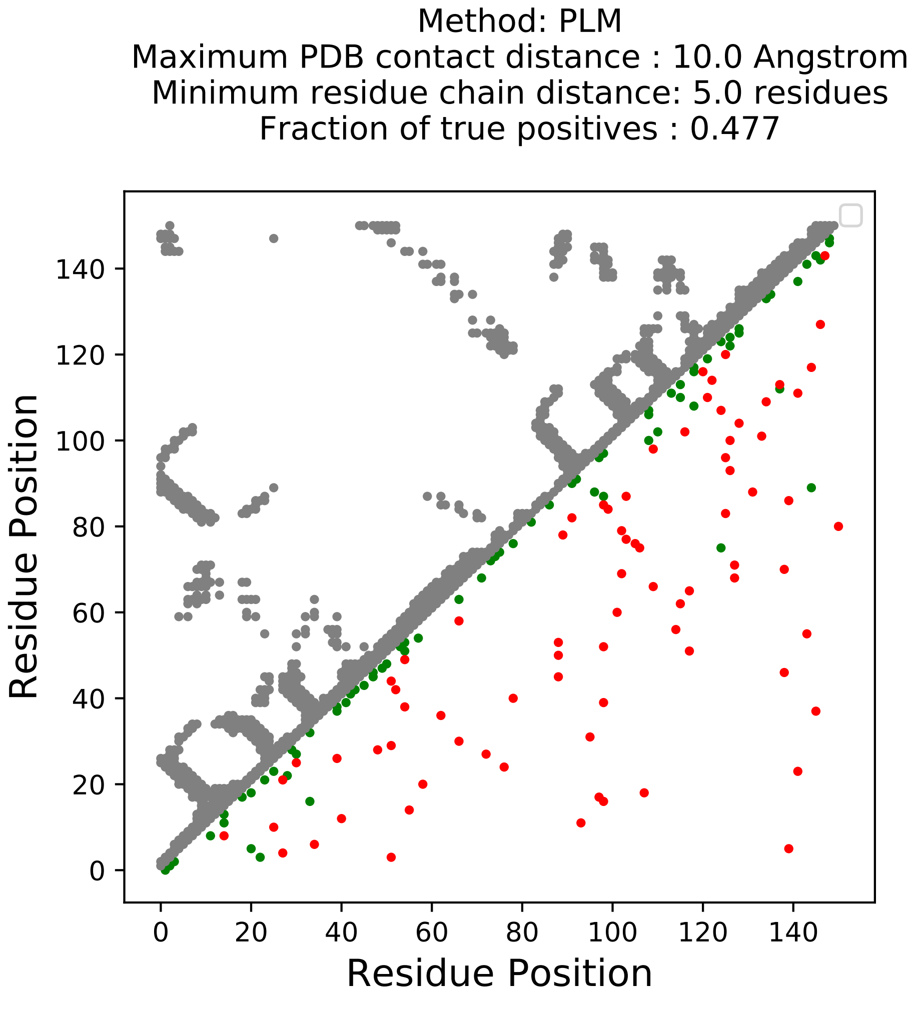

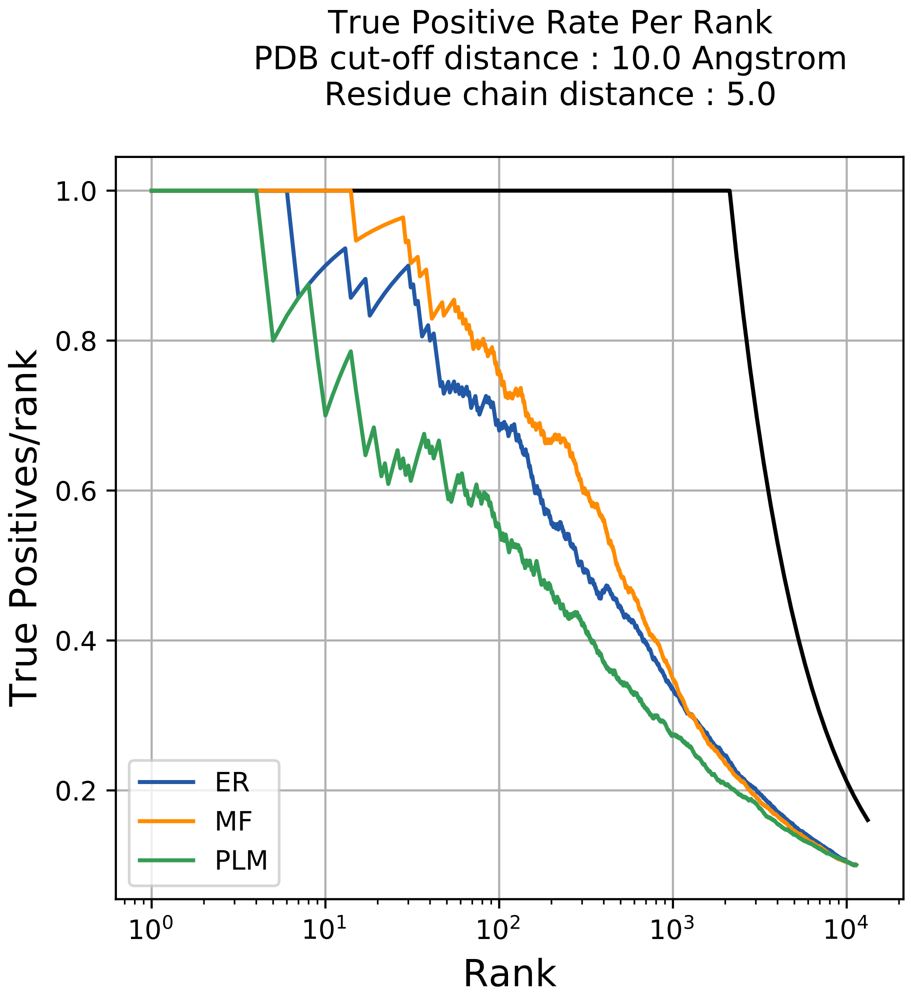

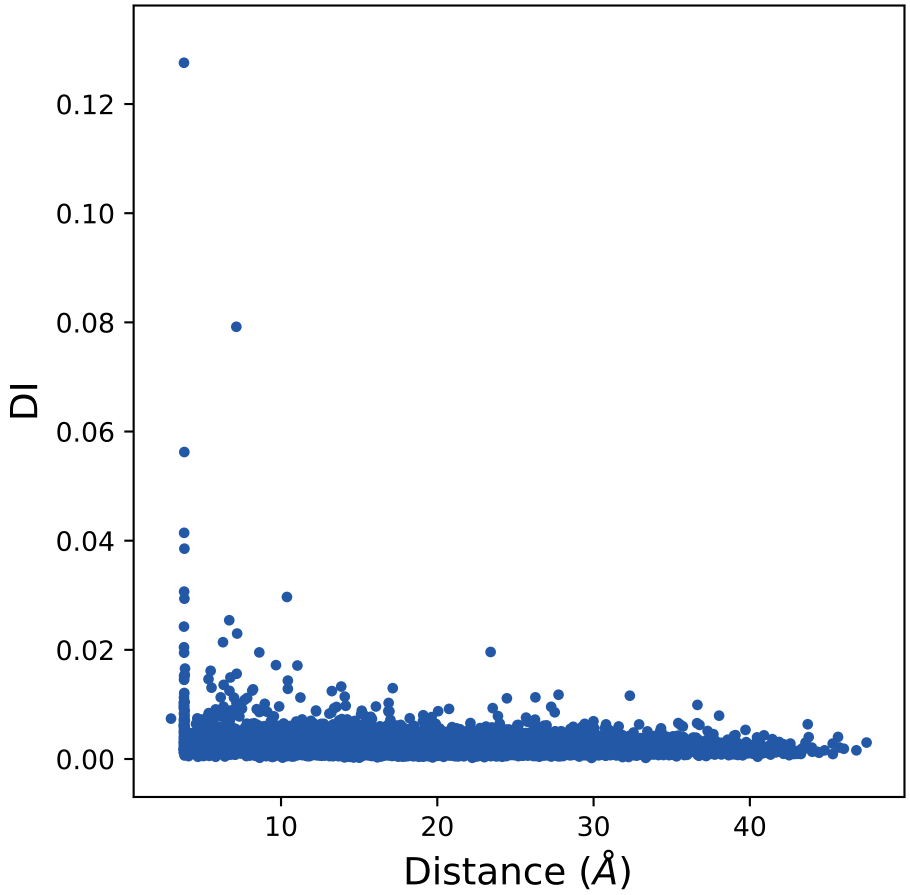

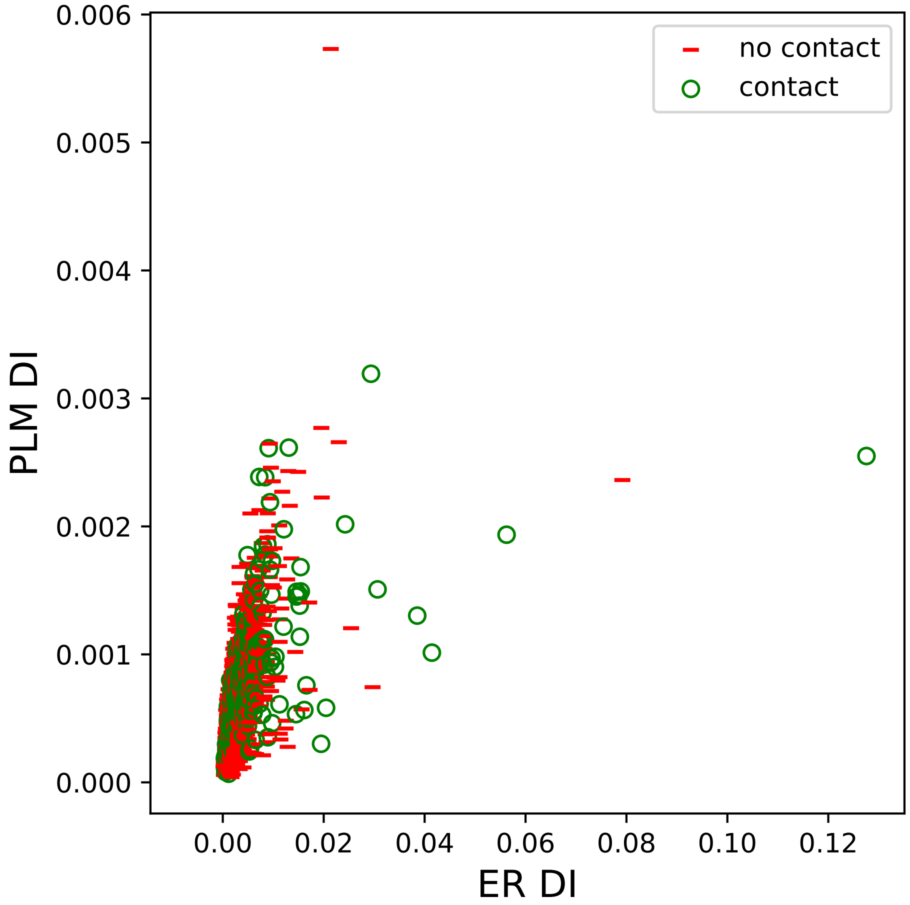

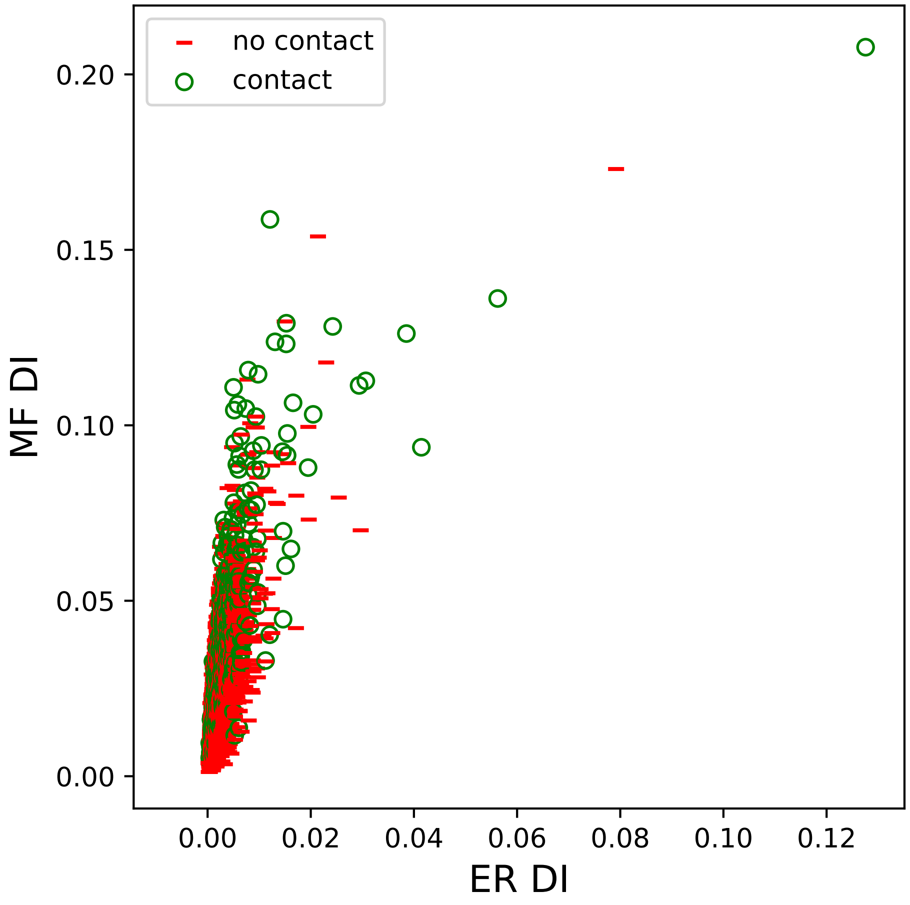

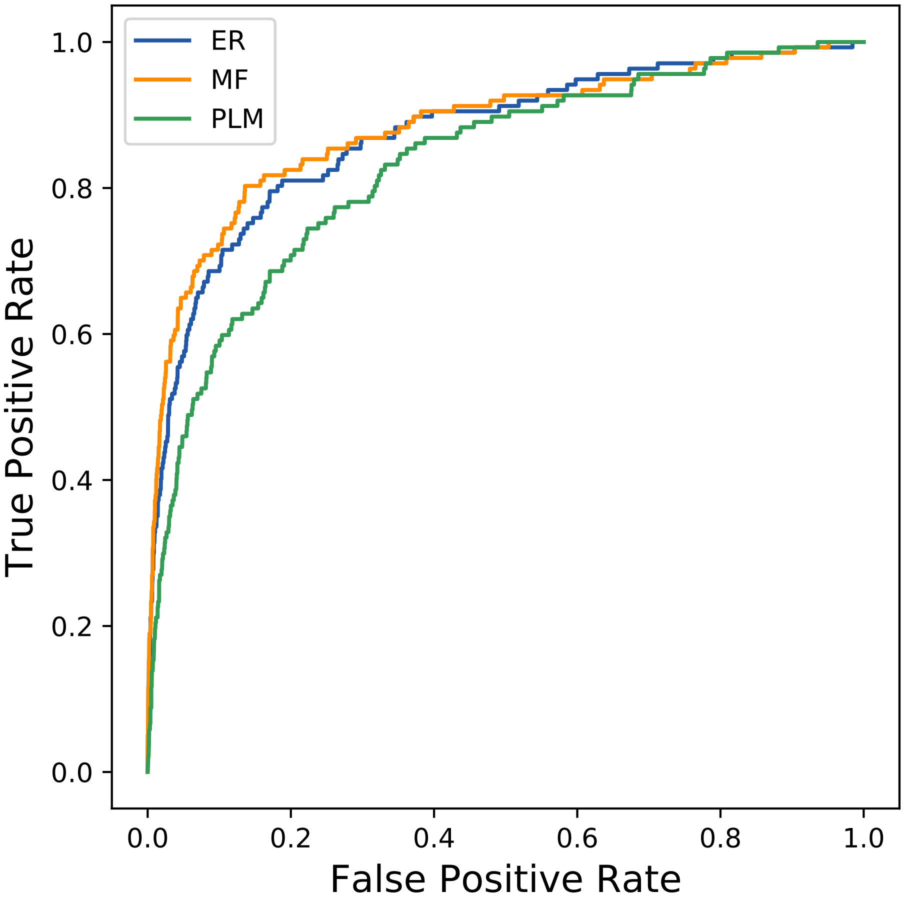

### 1480seq_221col_6ip1_PF14938_plots.pdf

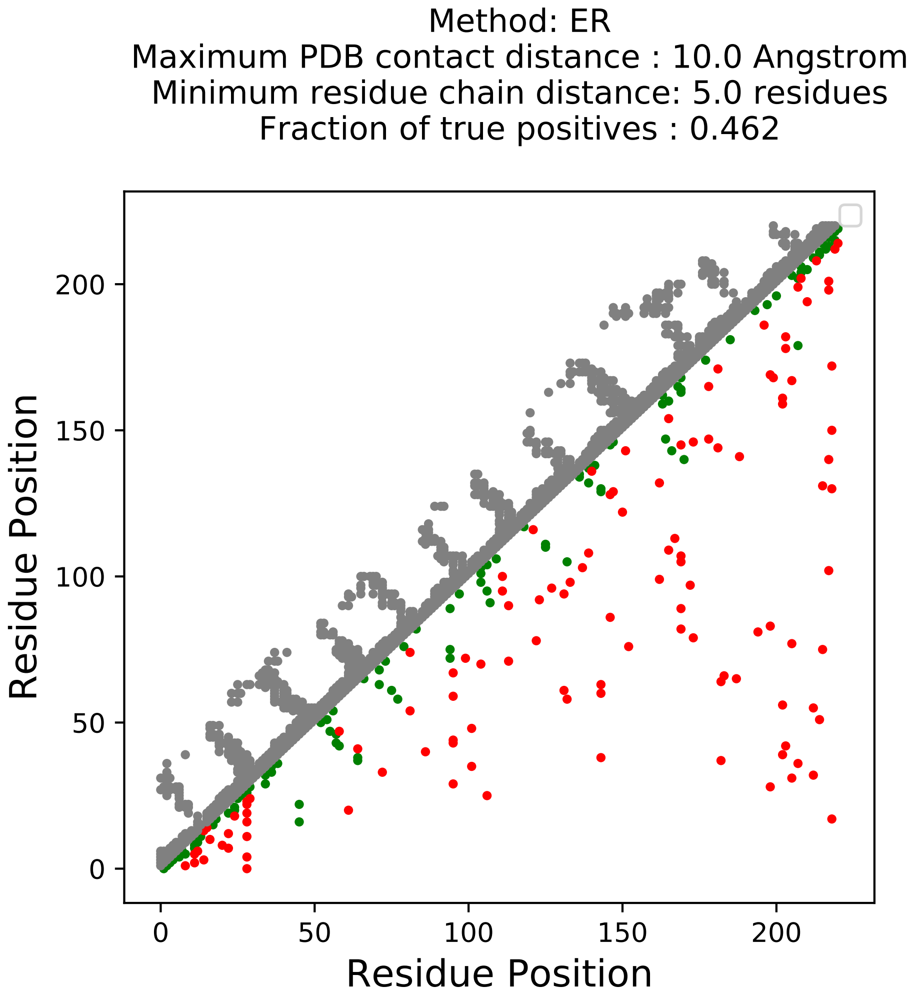

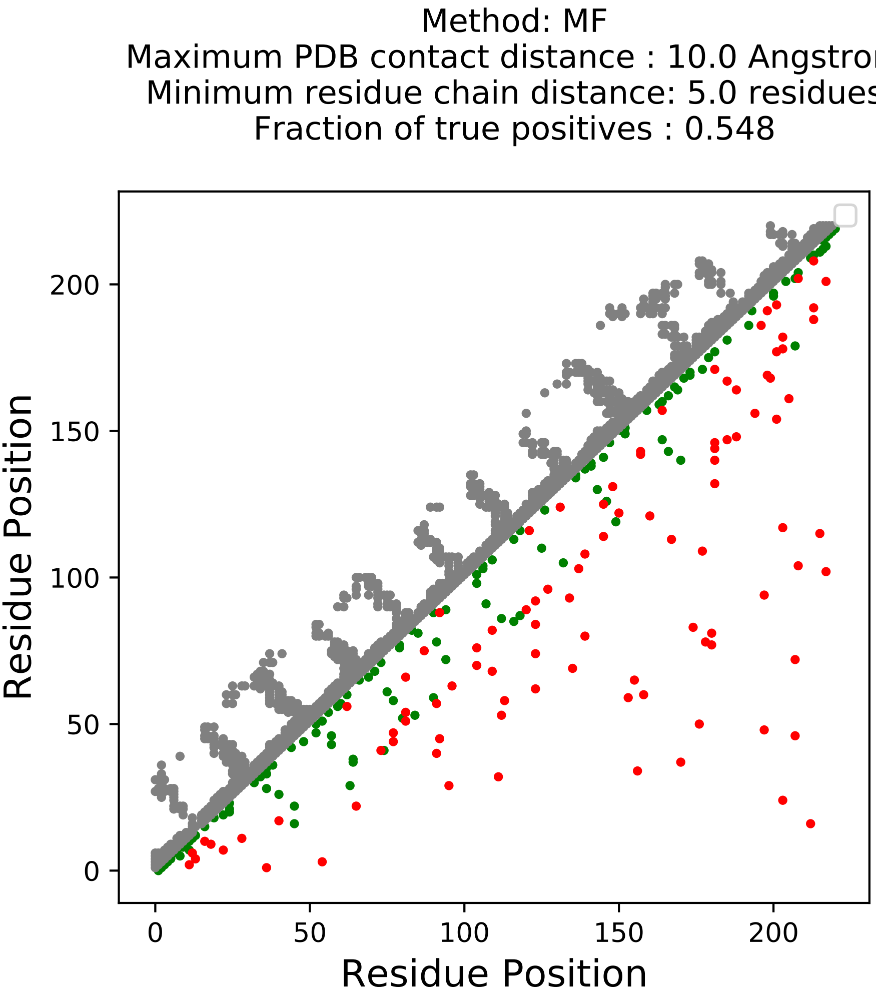

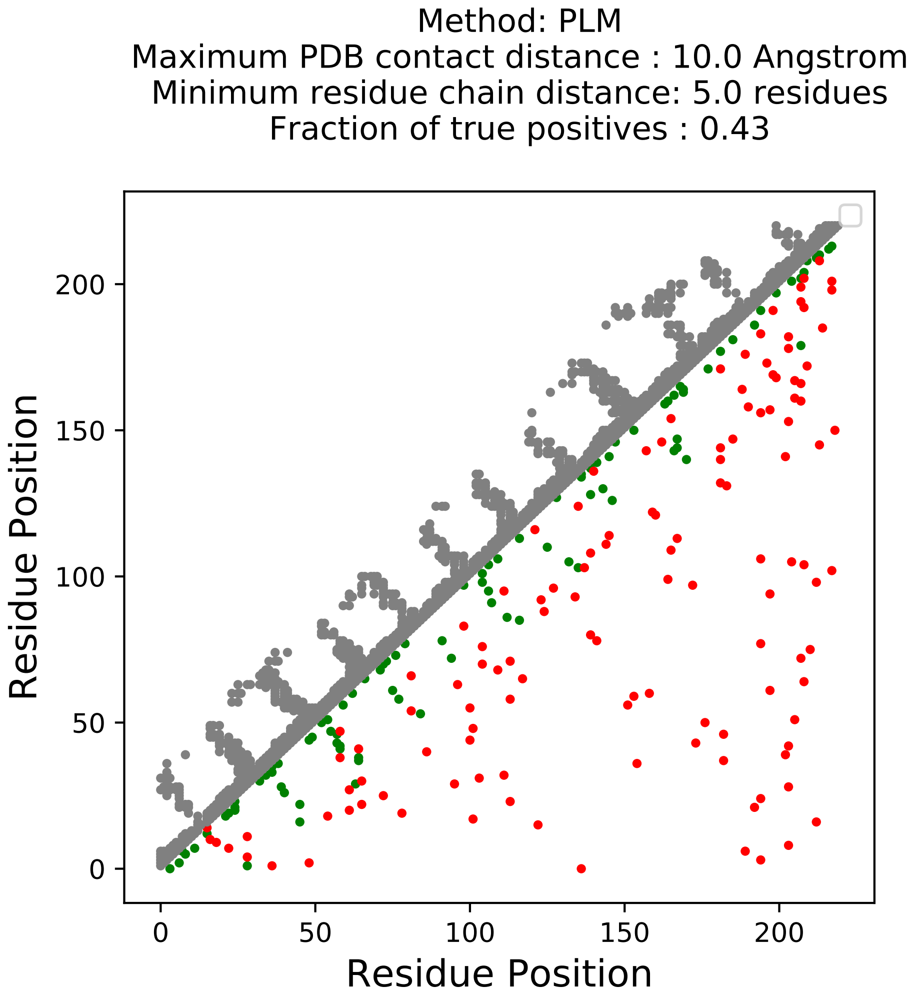

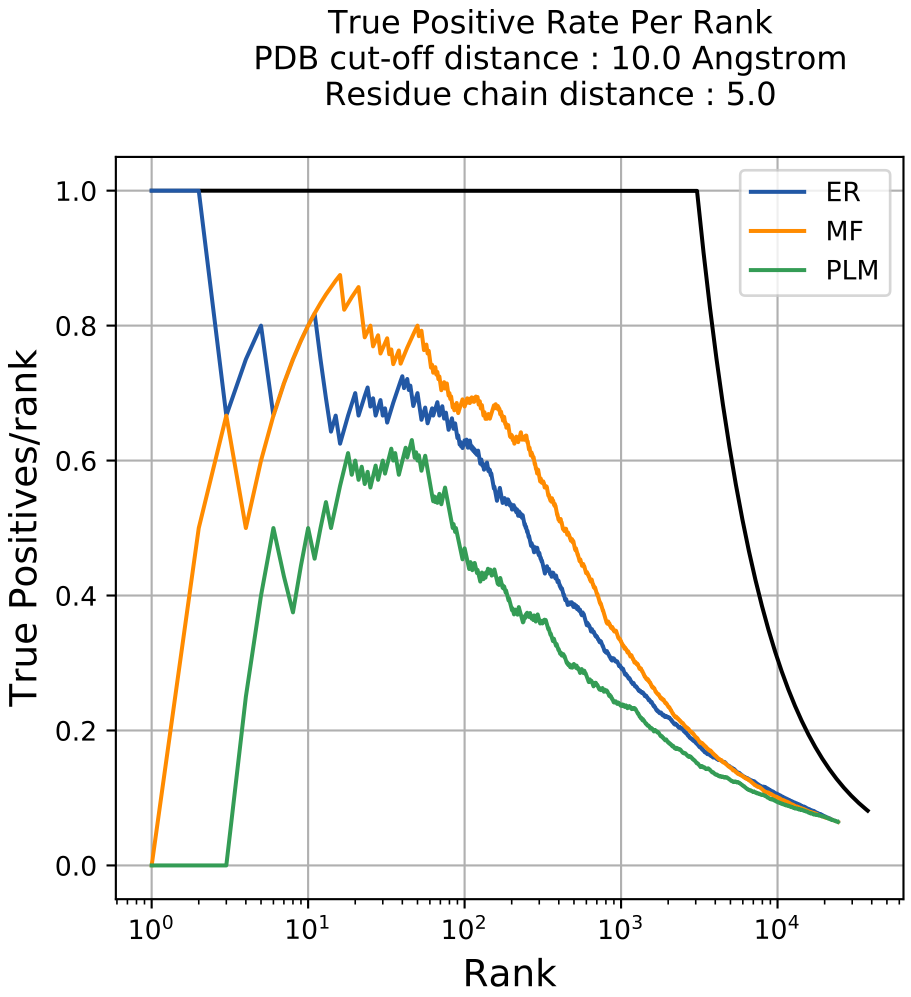

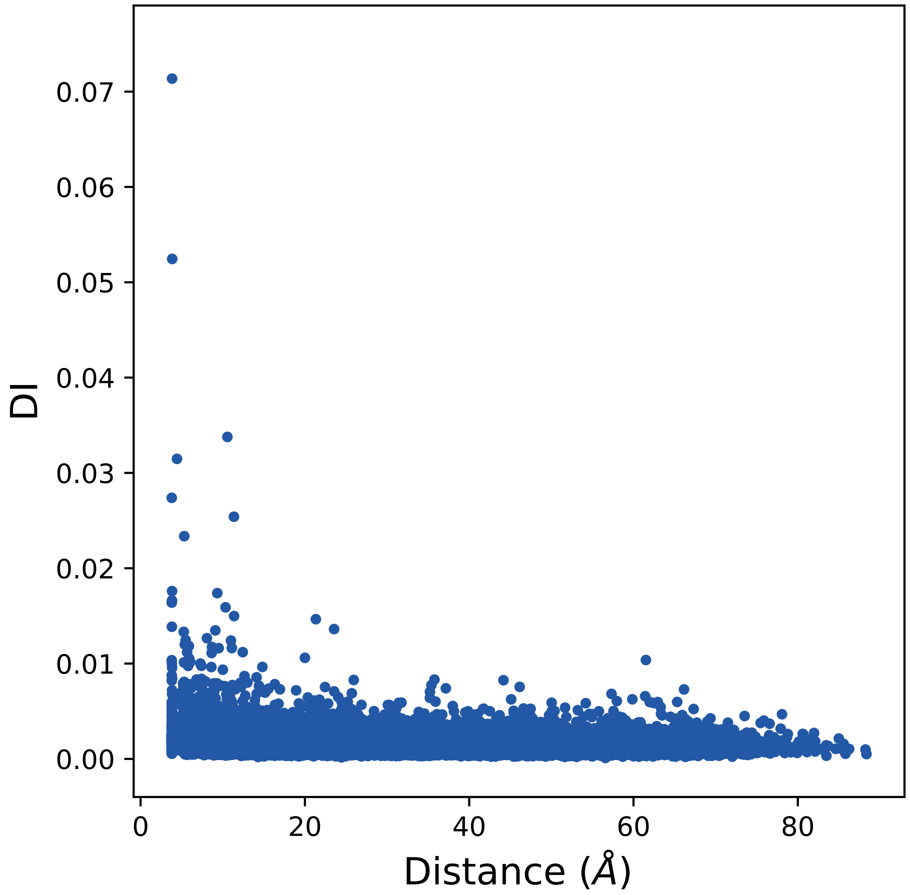

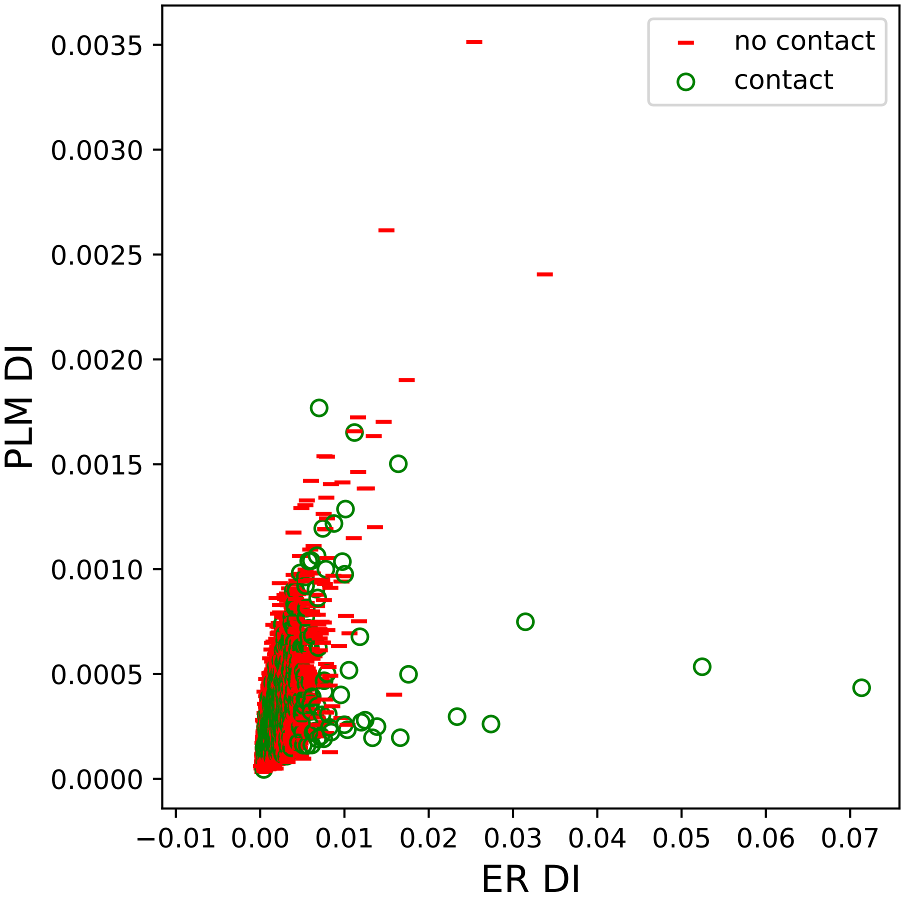

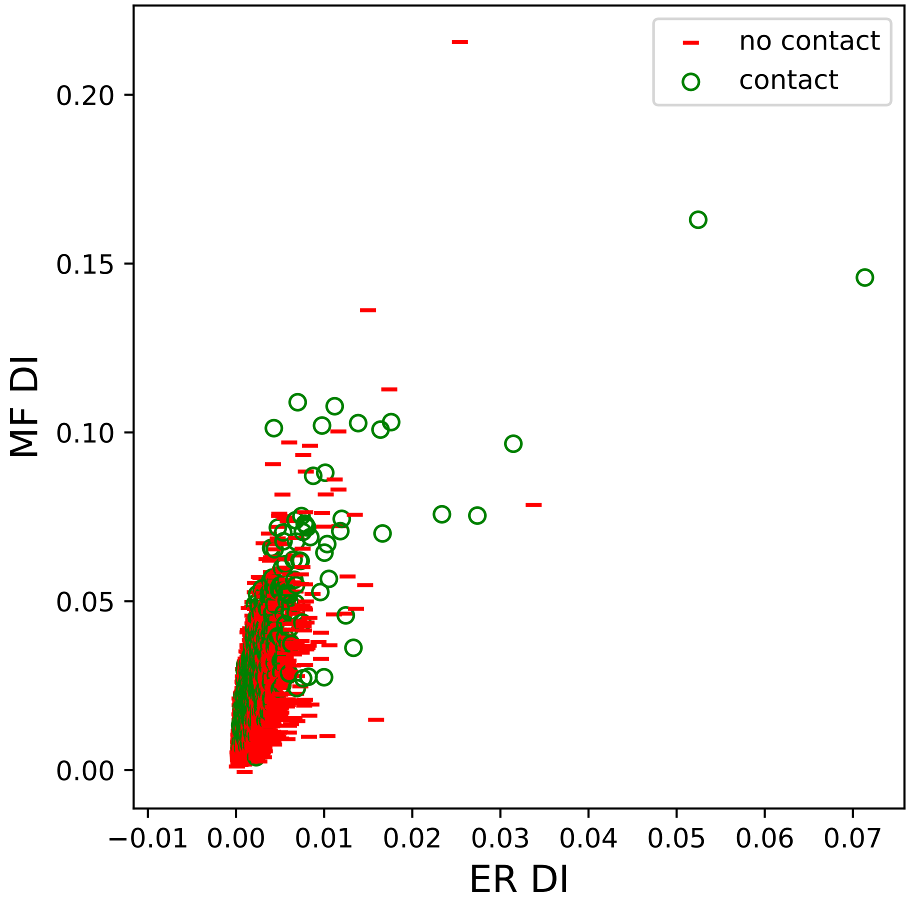

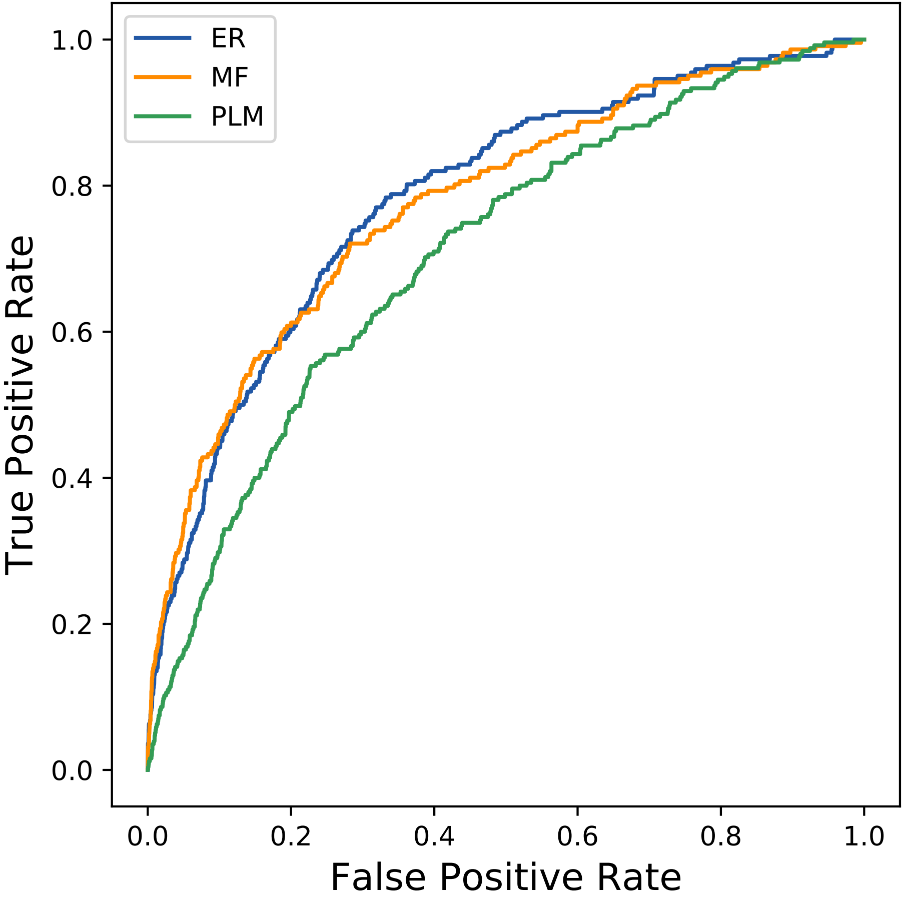

### 1533seq_165col_2a6z_PF03388_plots.pdf

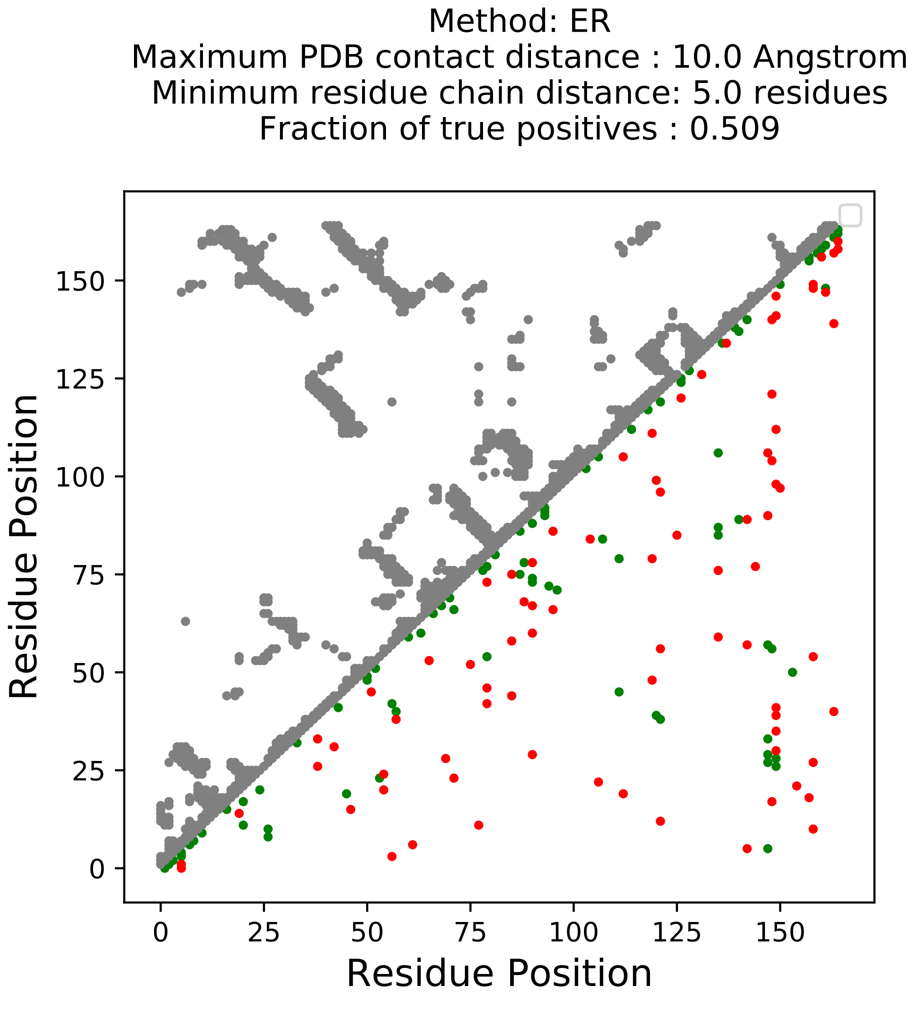

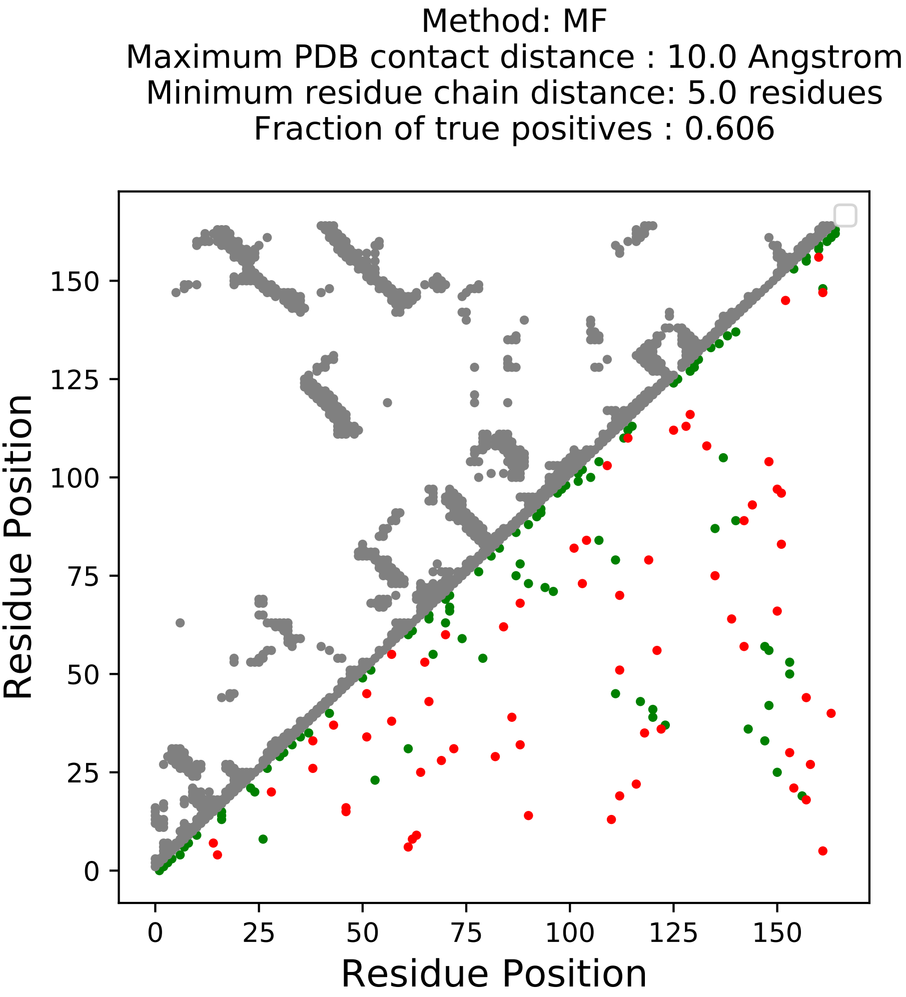

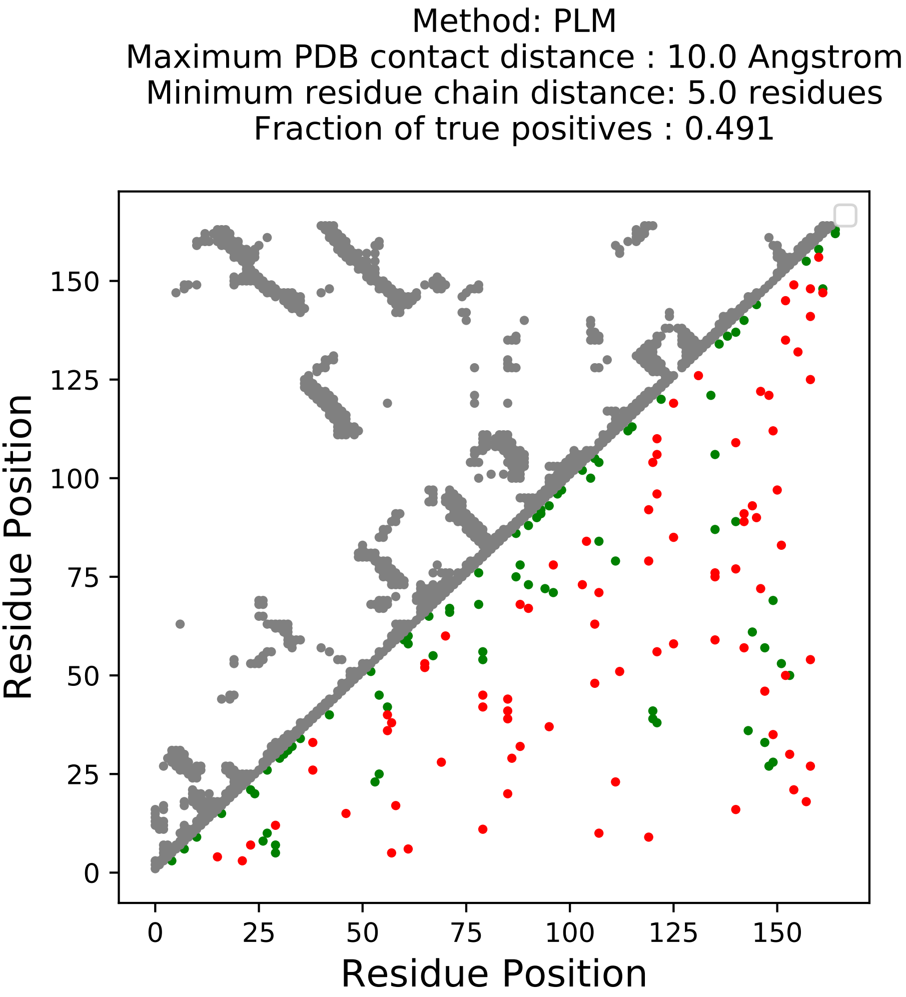

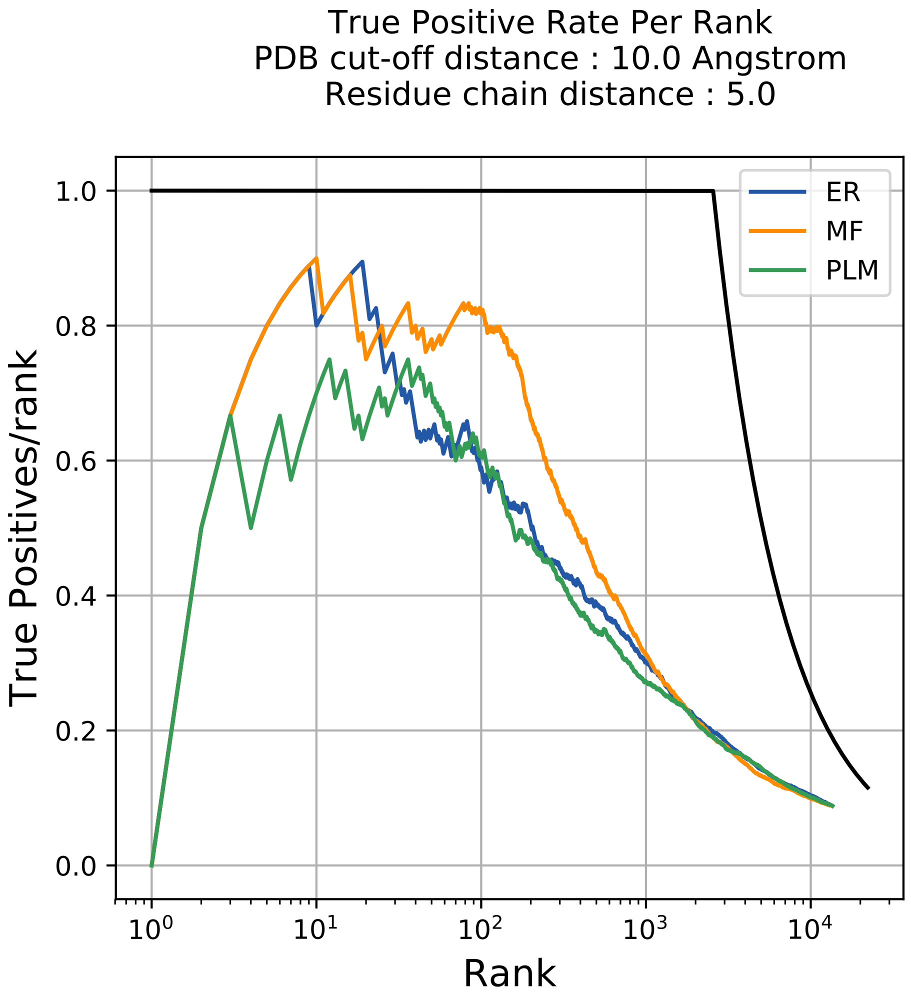

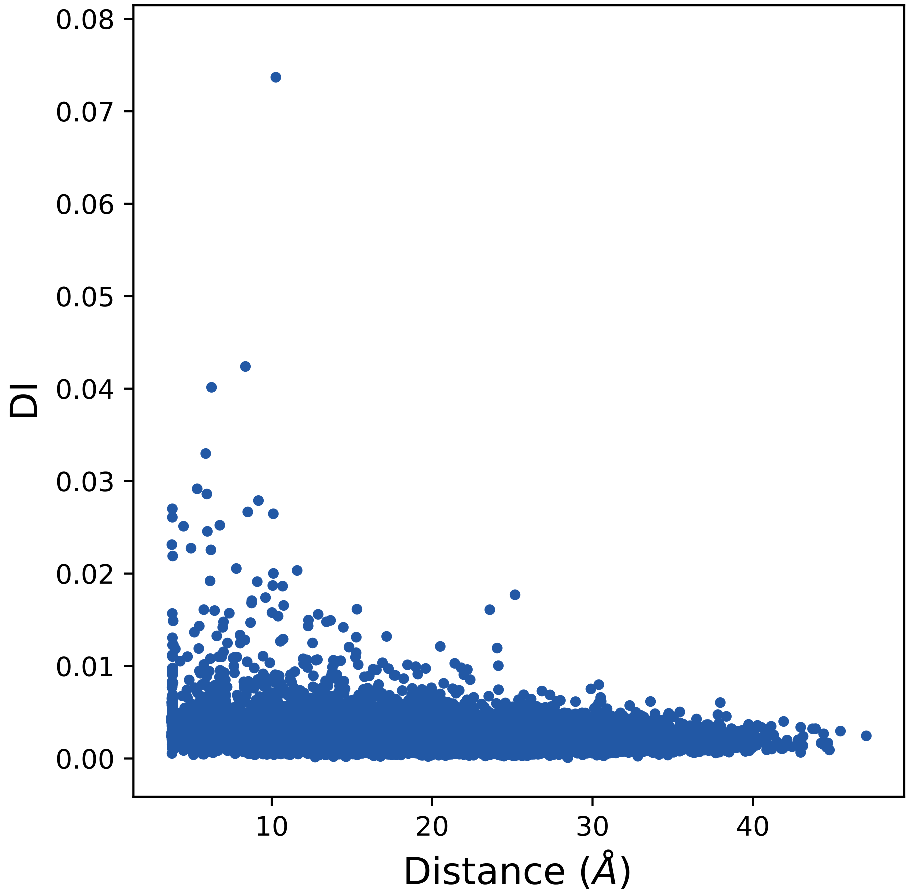

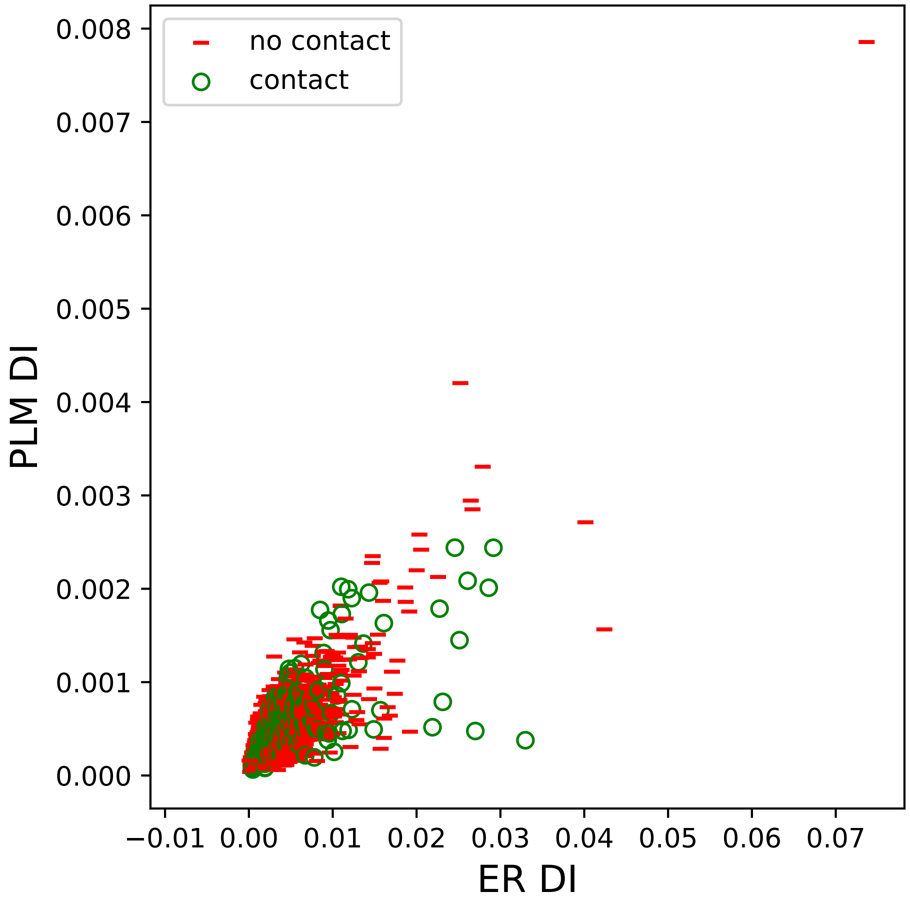

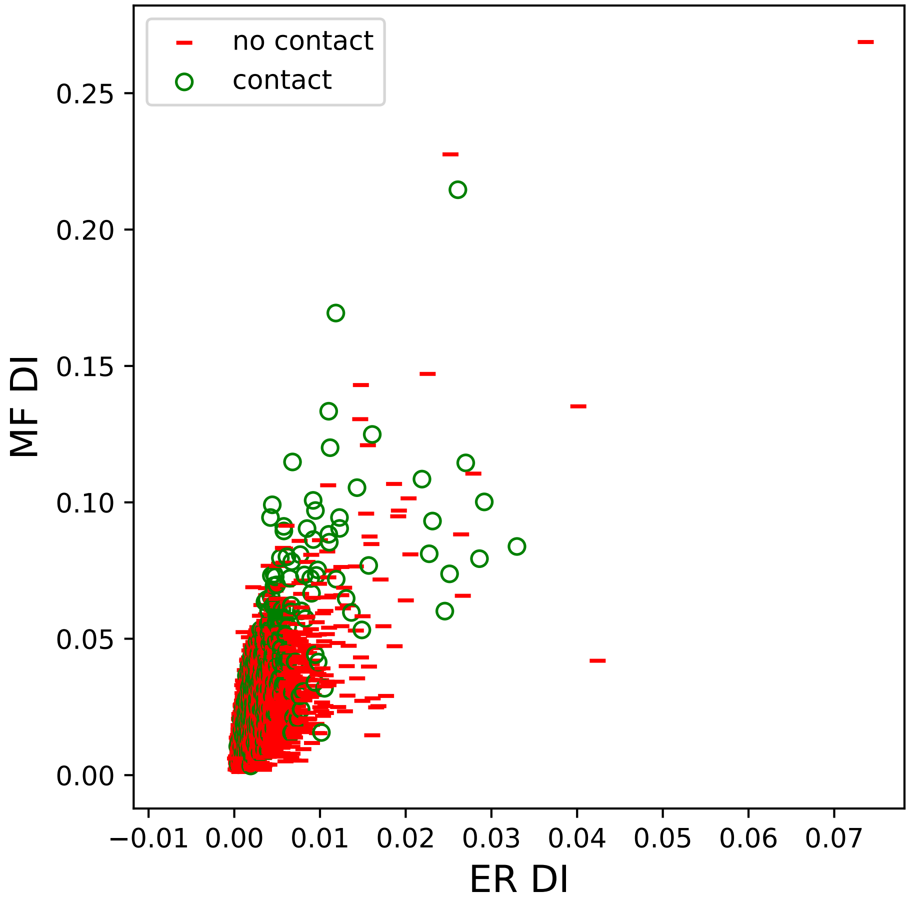

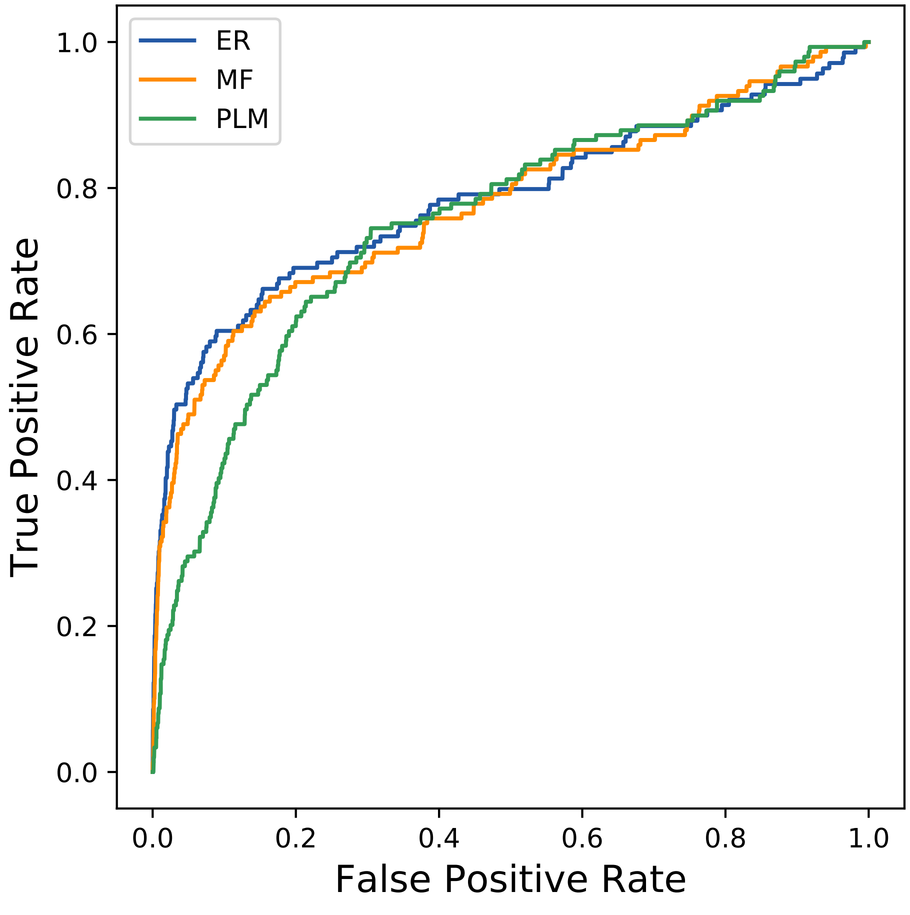

### 1574seq_112col_2acg_PF00235_plots.pdf

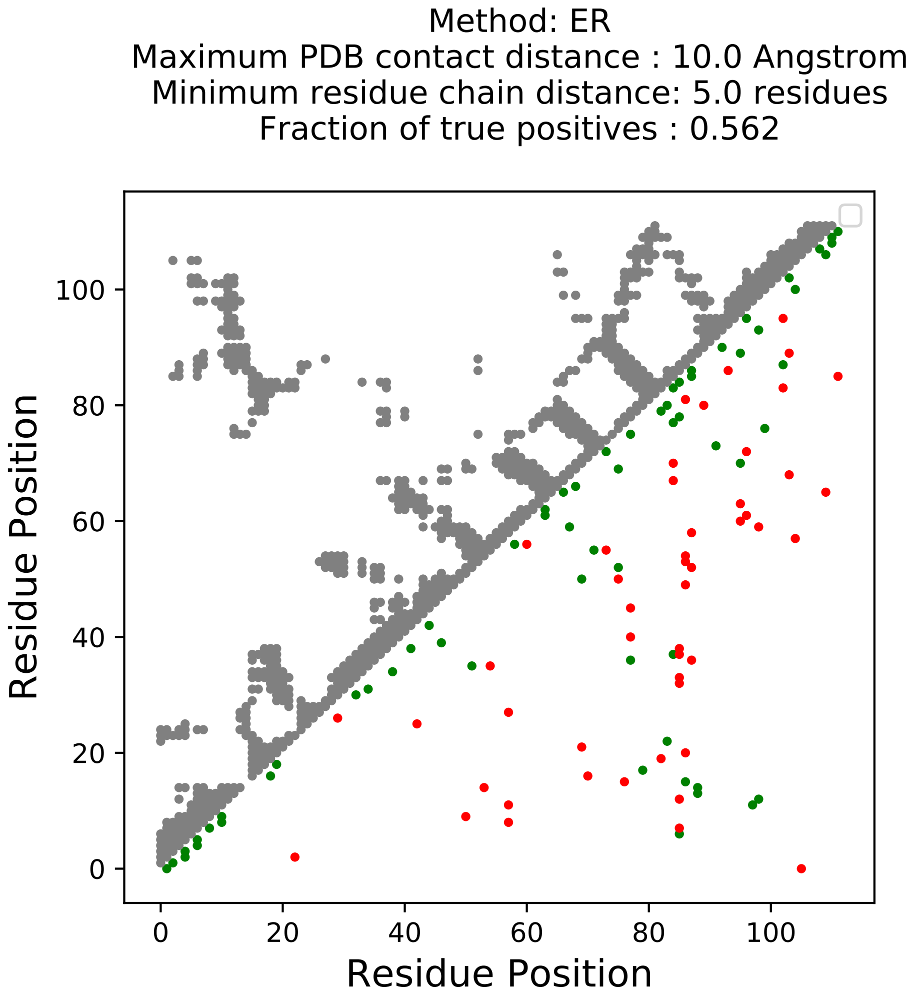

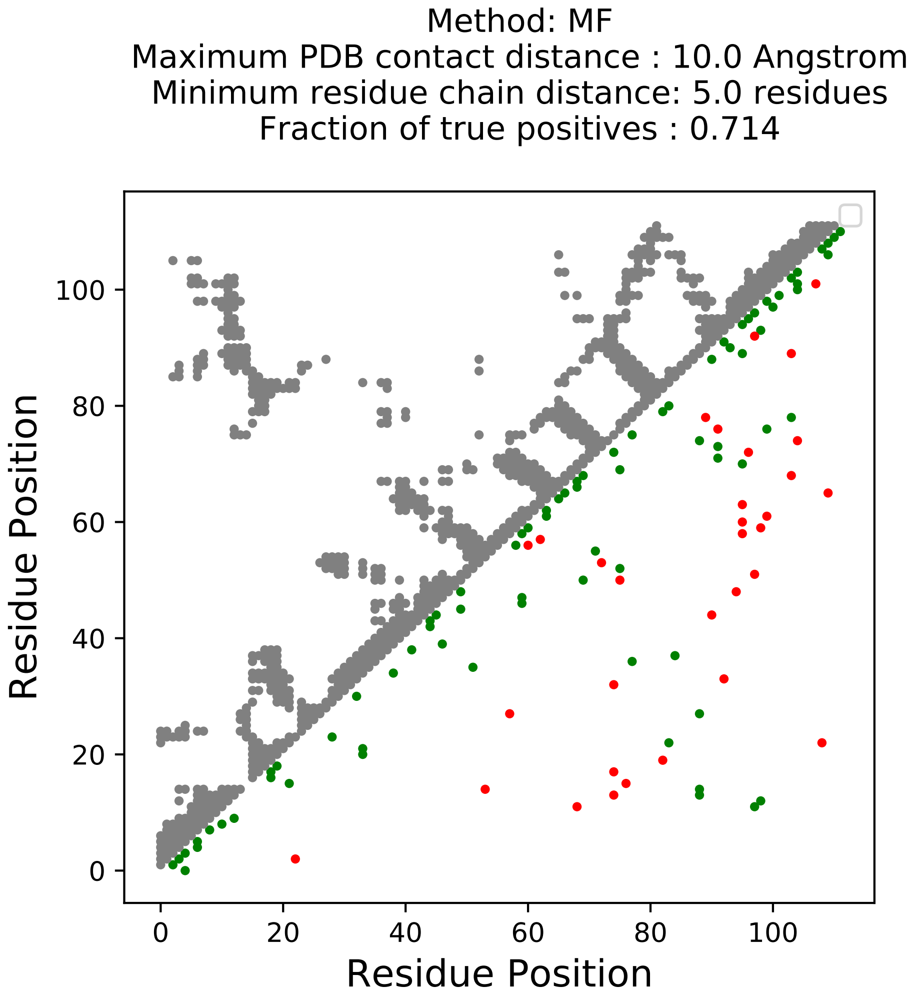

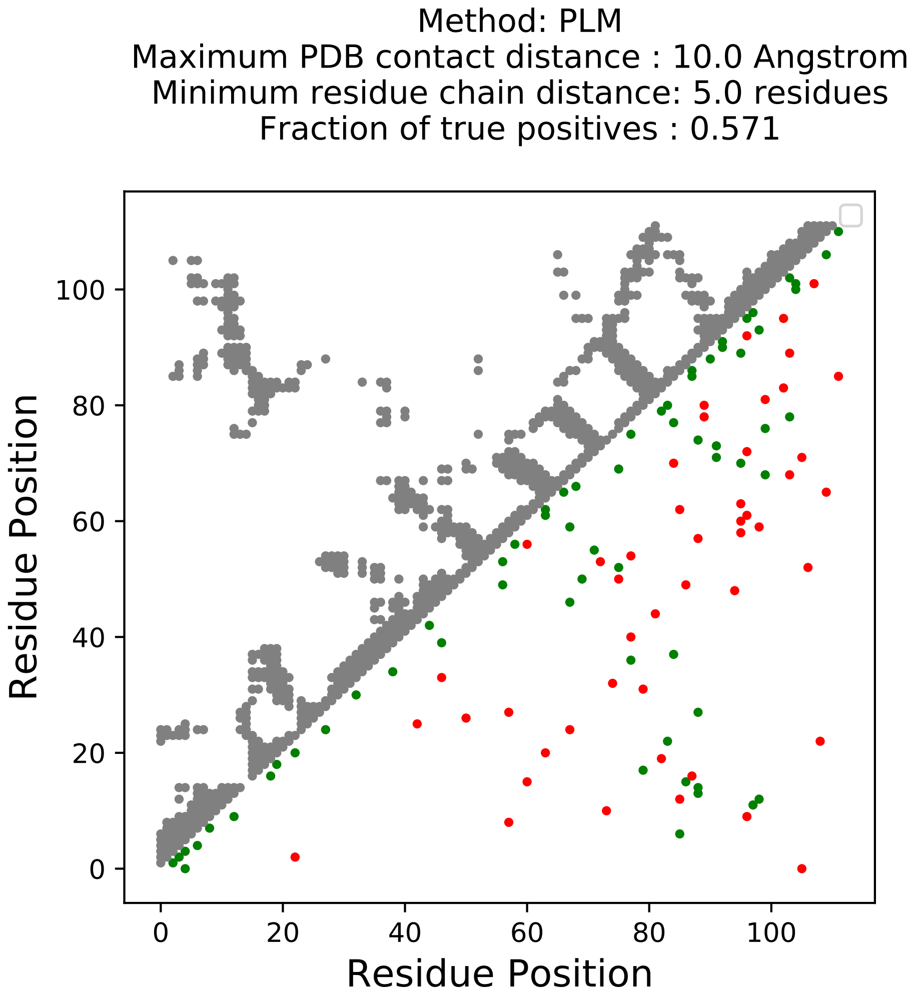

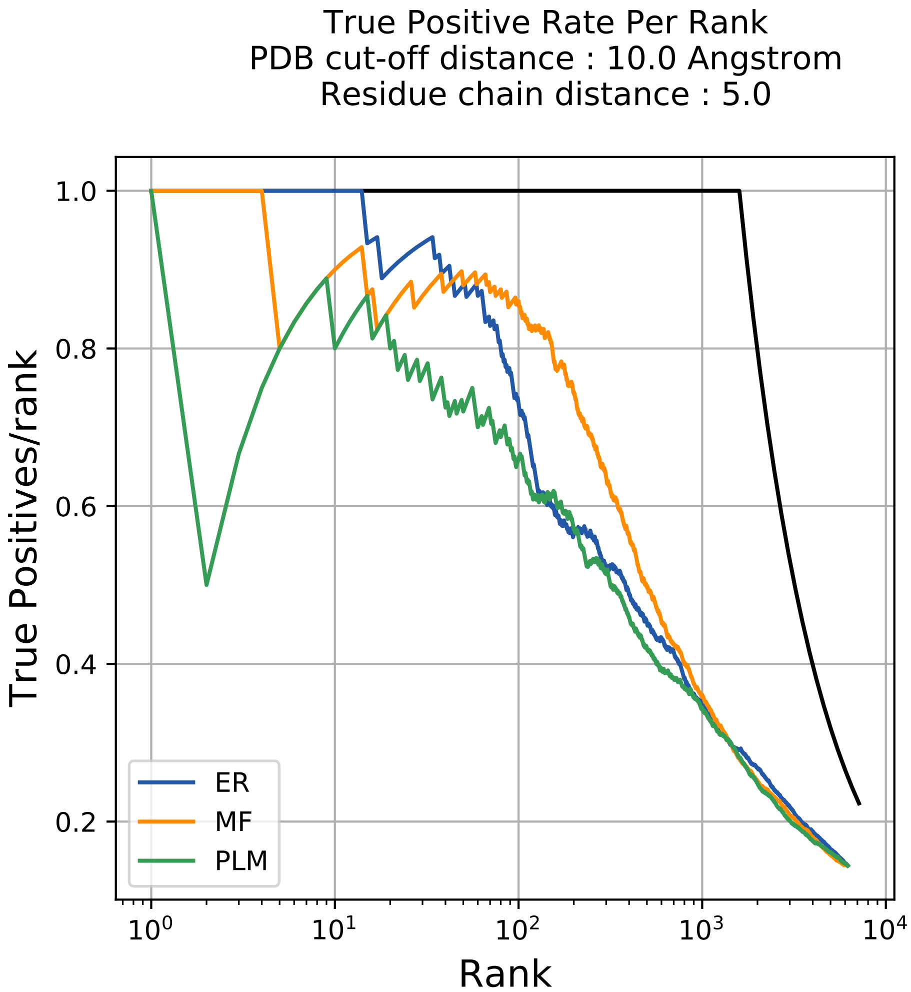

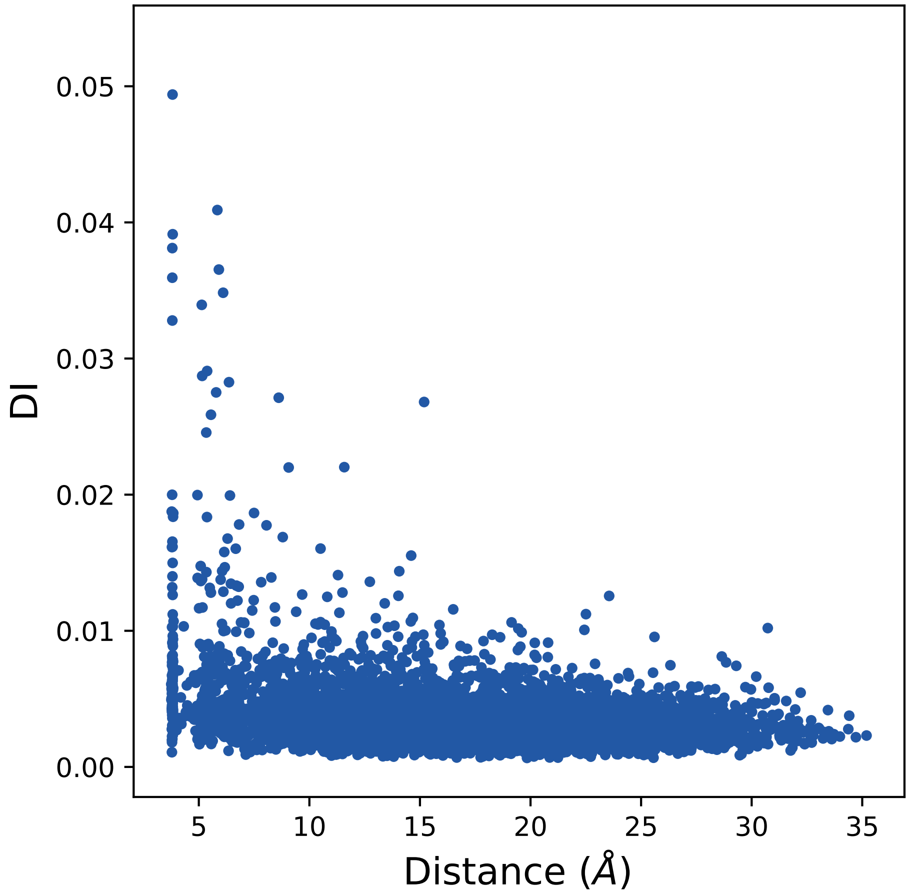

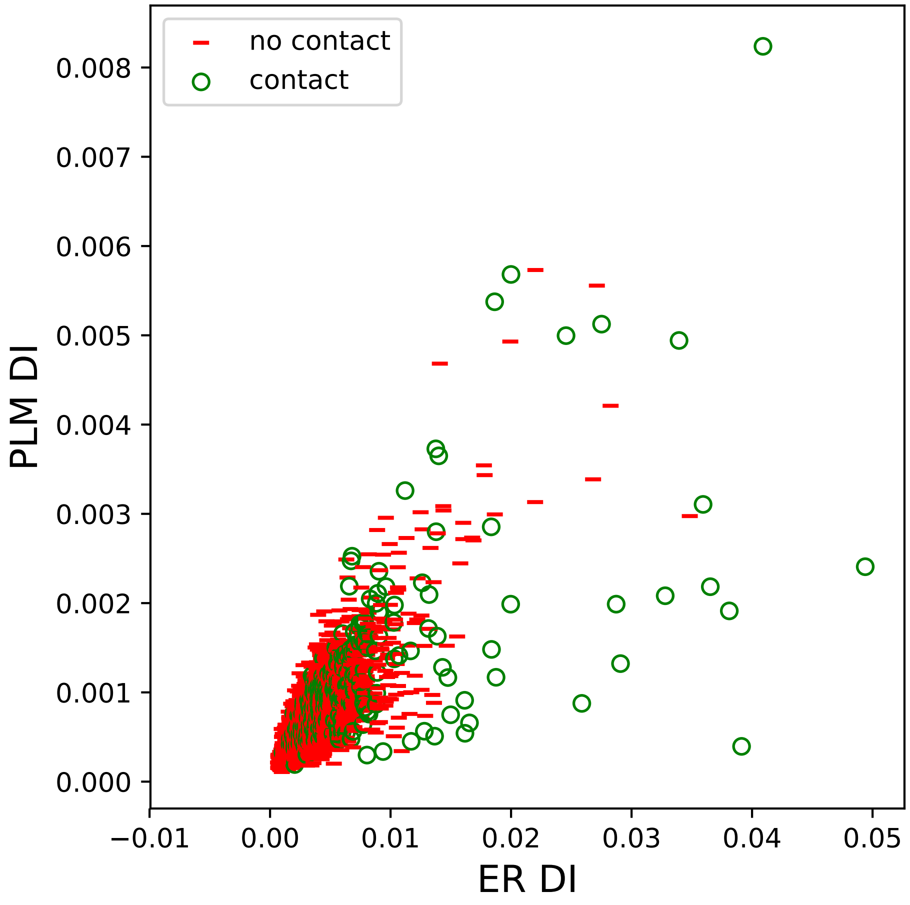

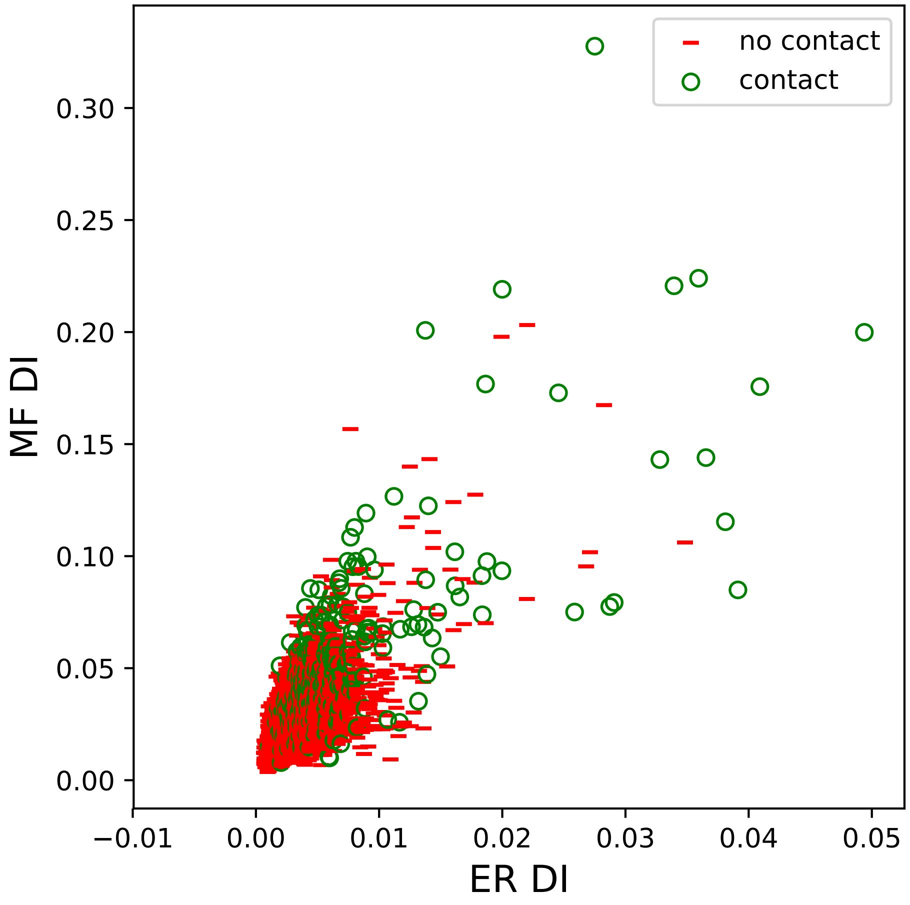

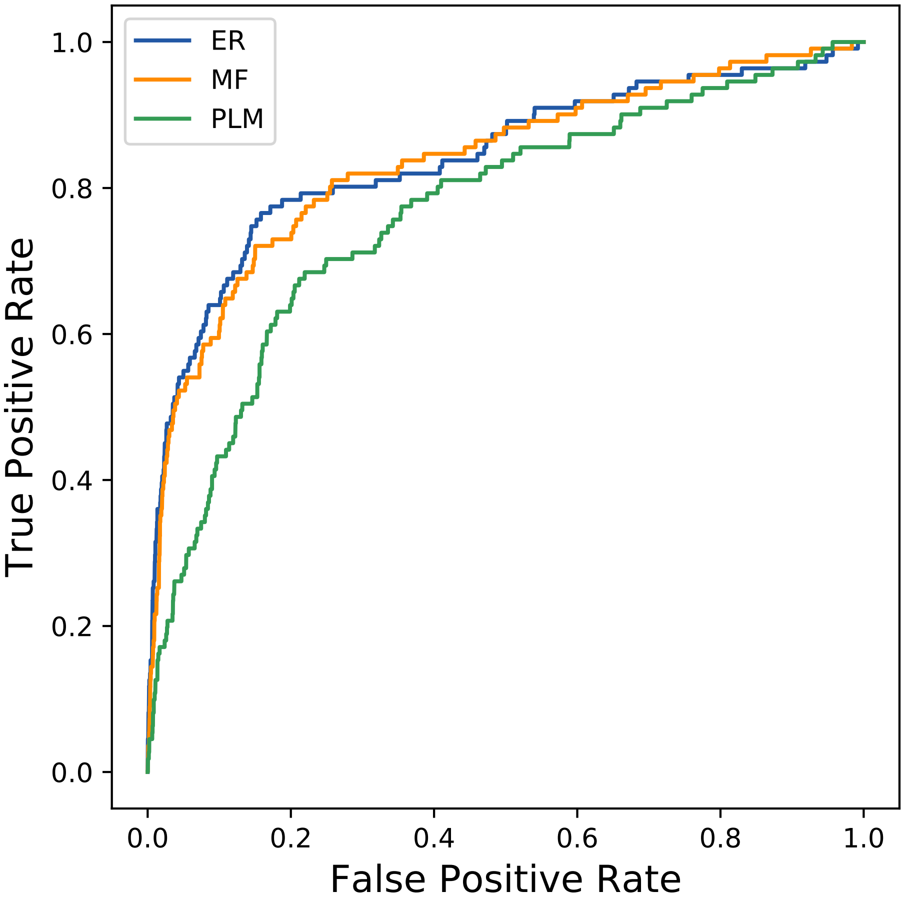

### 1739seq_139col_3ibe_PF00613_plots.pdf

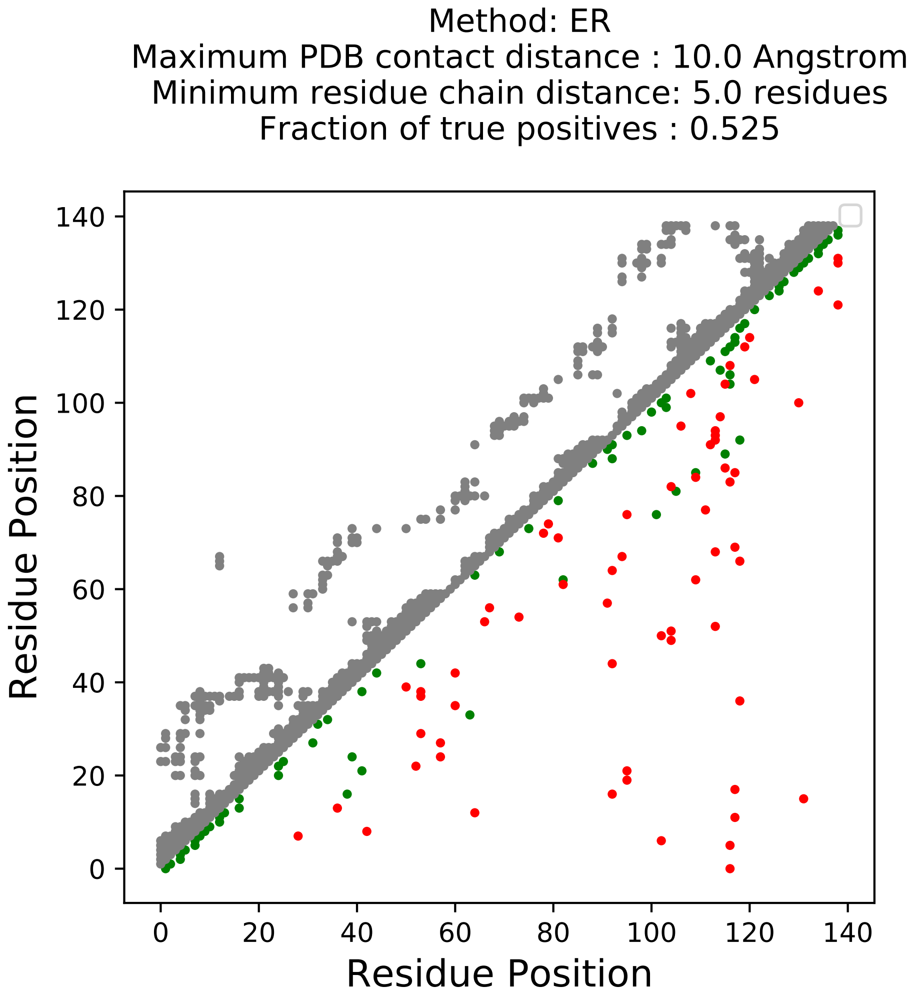

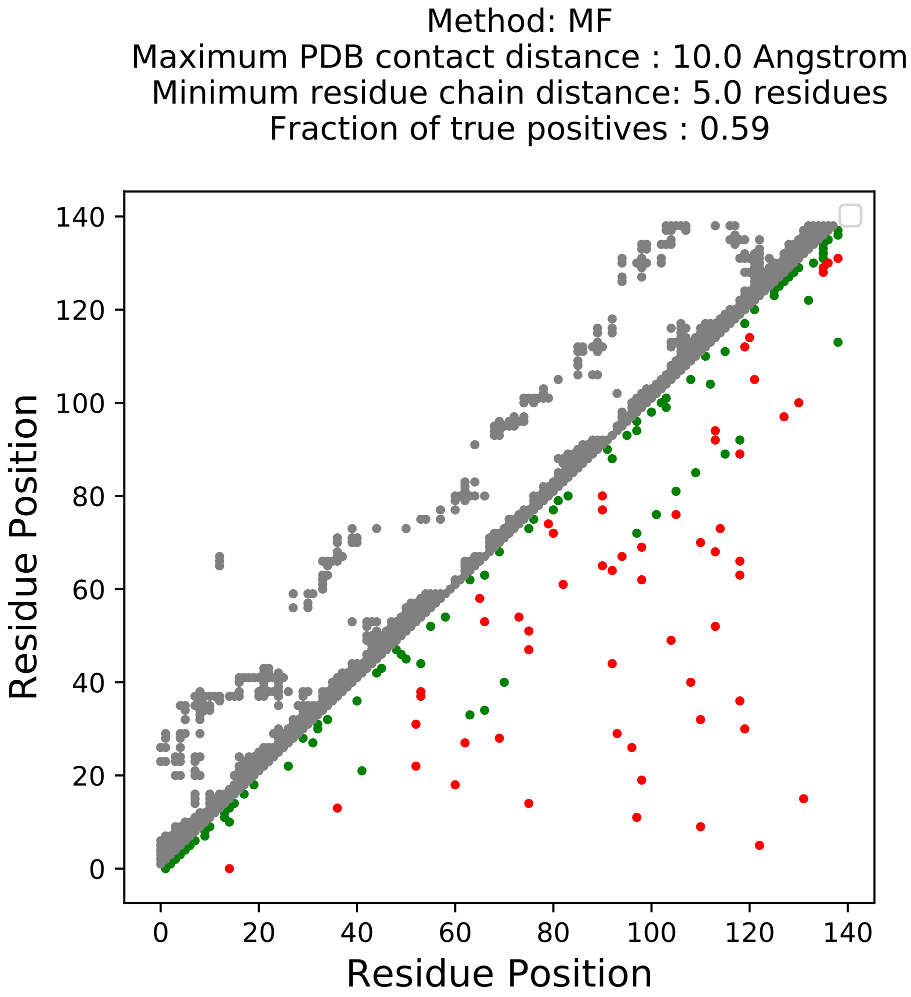

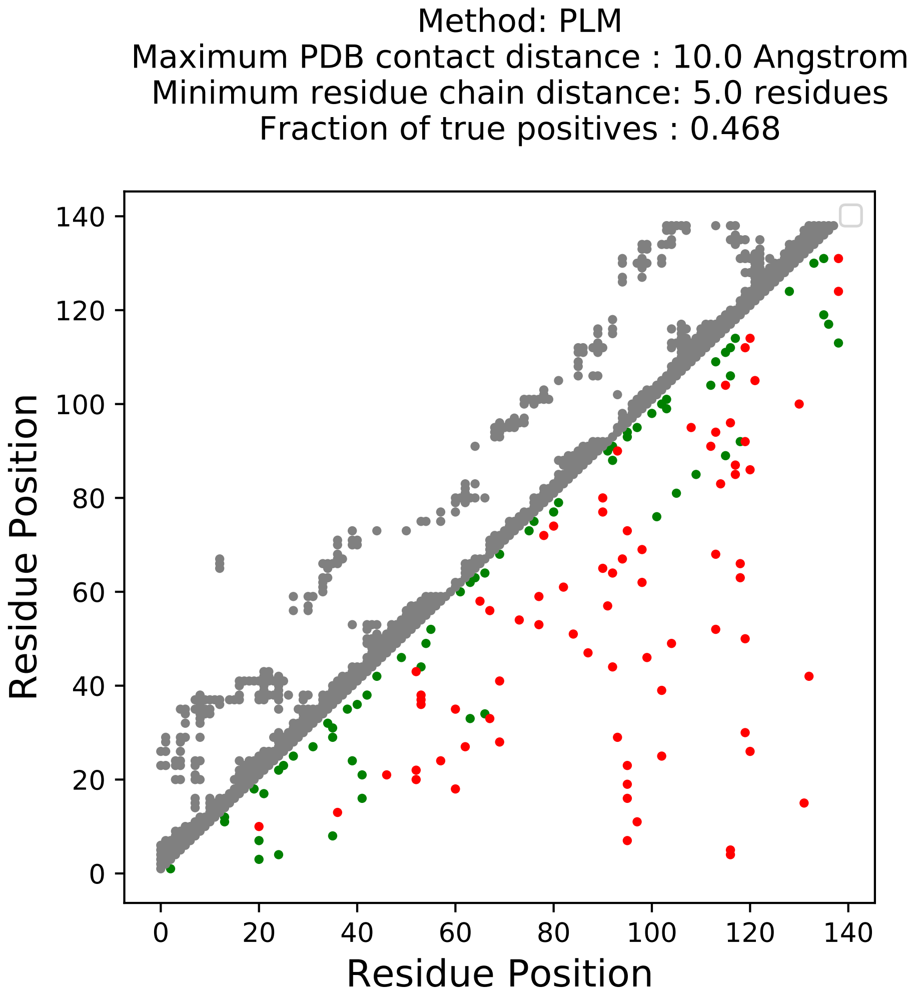

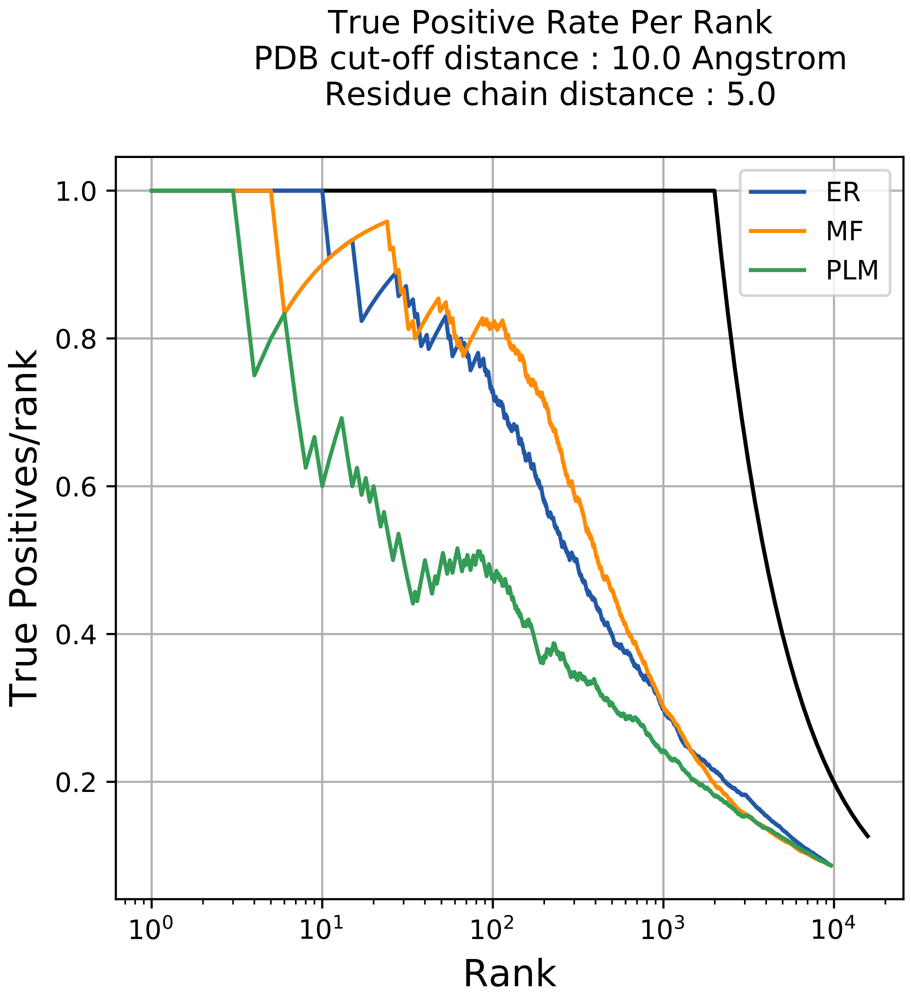

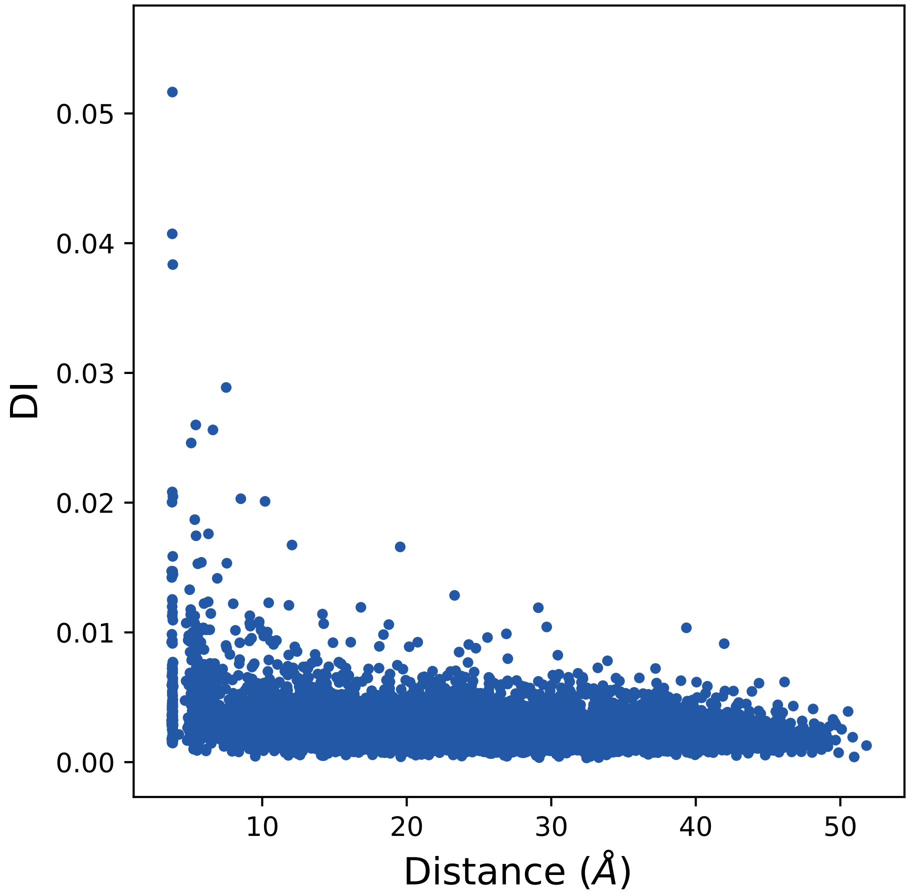

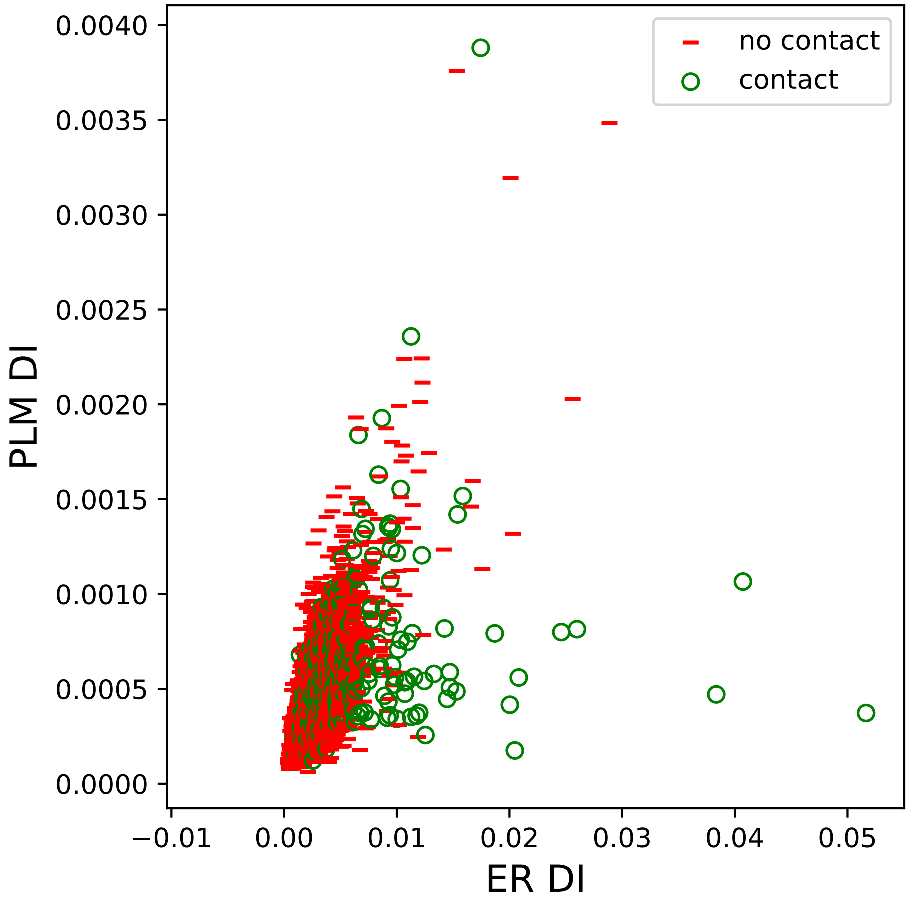

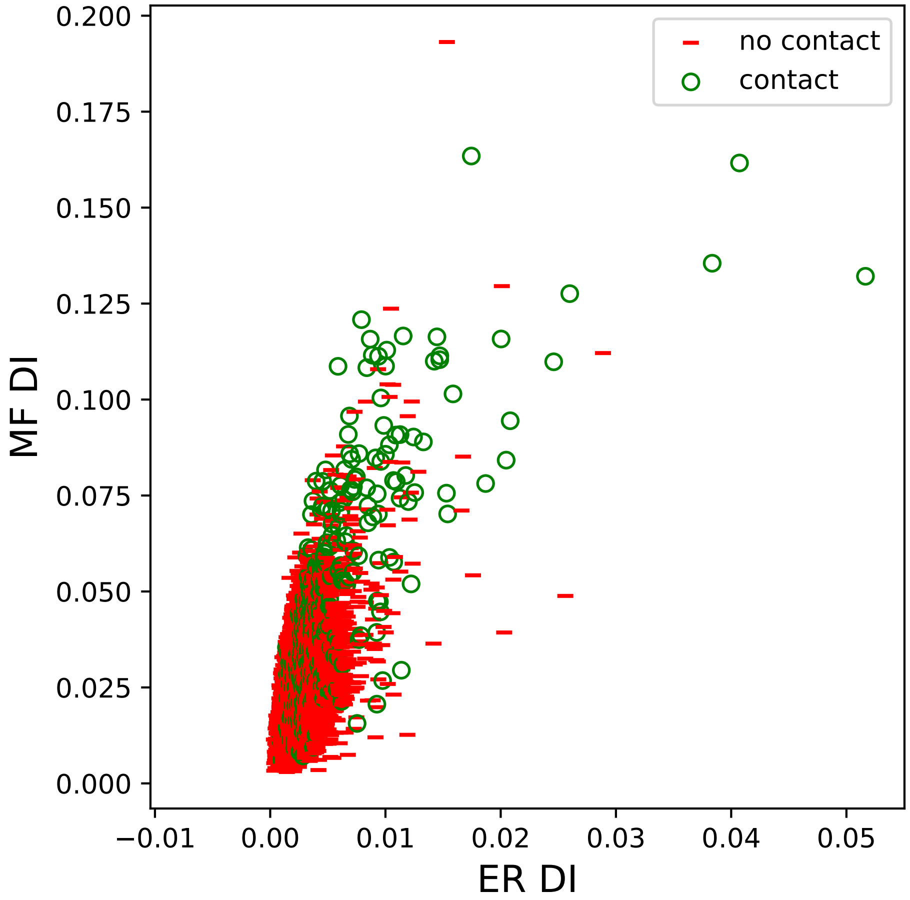

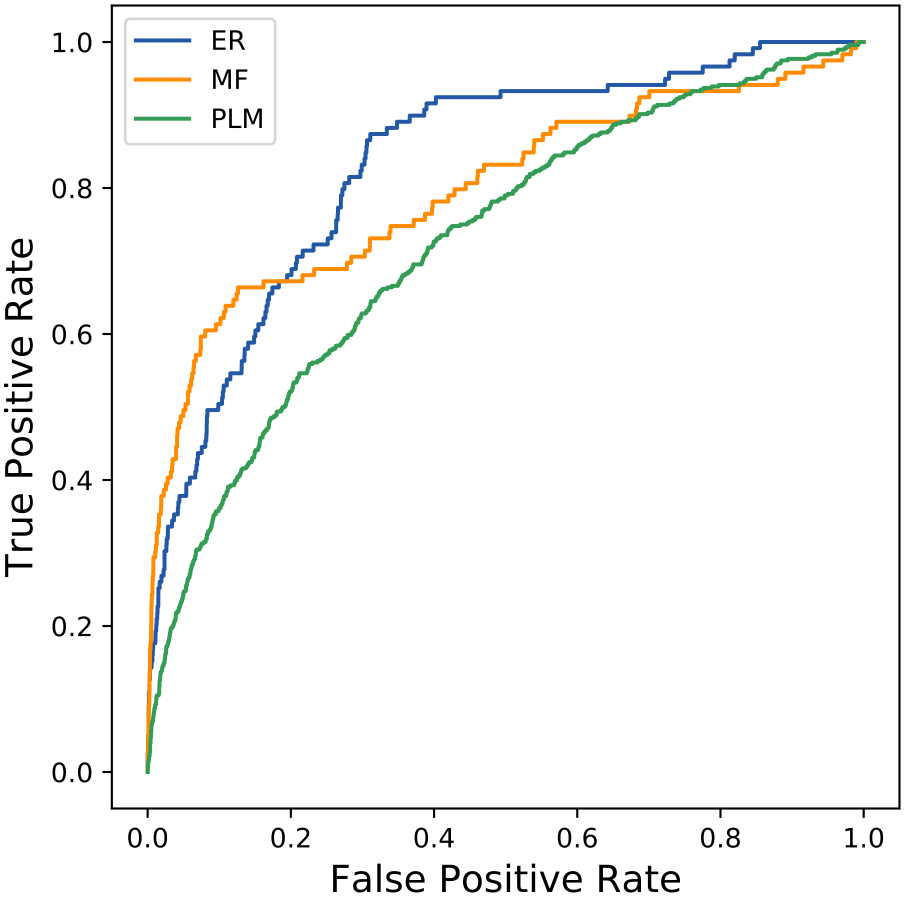

### 1833seq_37col_4ht9_PF17209_plots.pdf

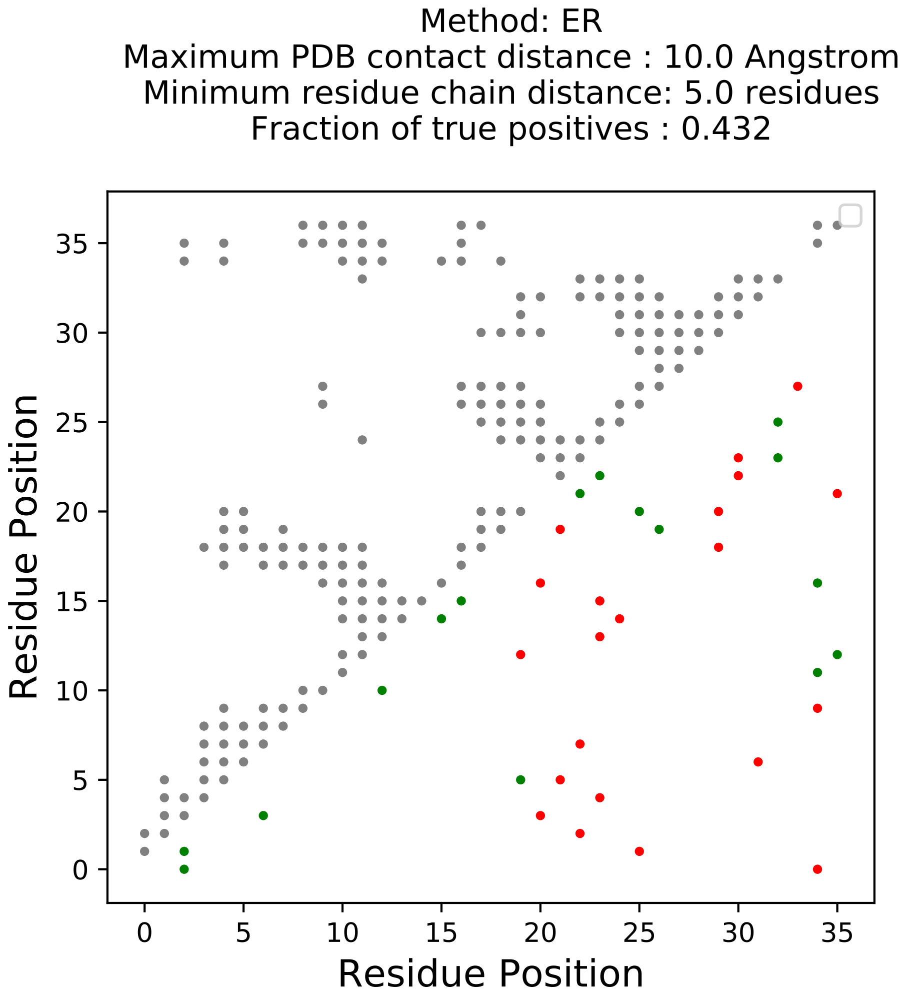

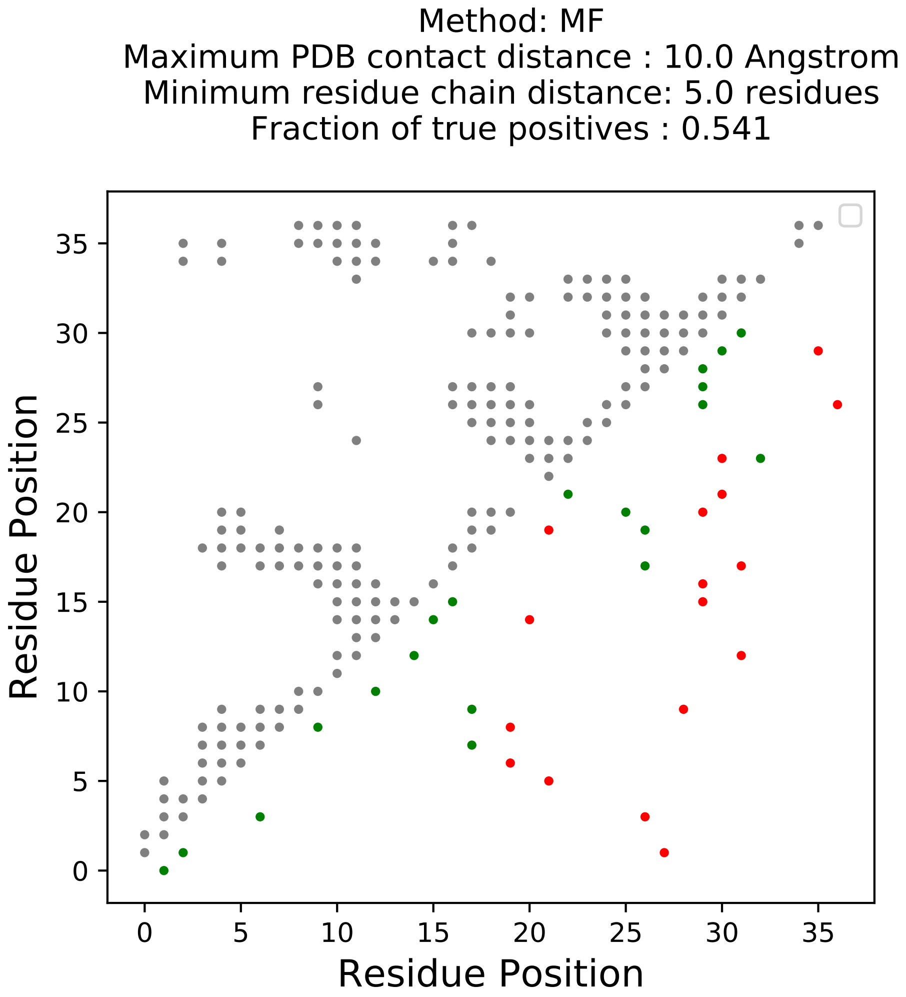

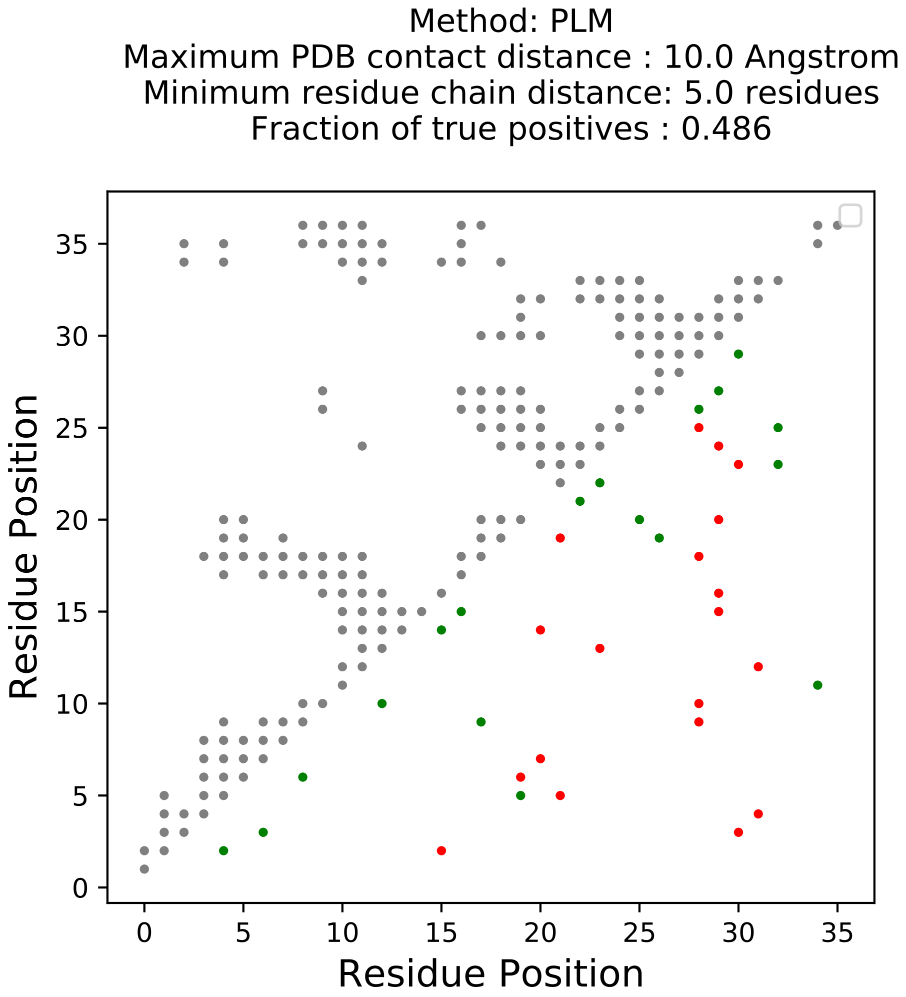

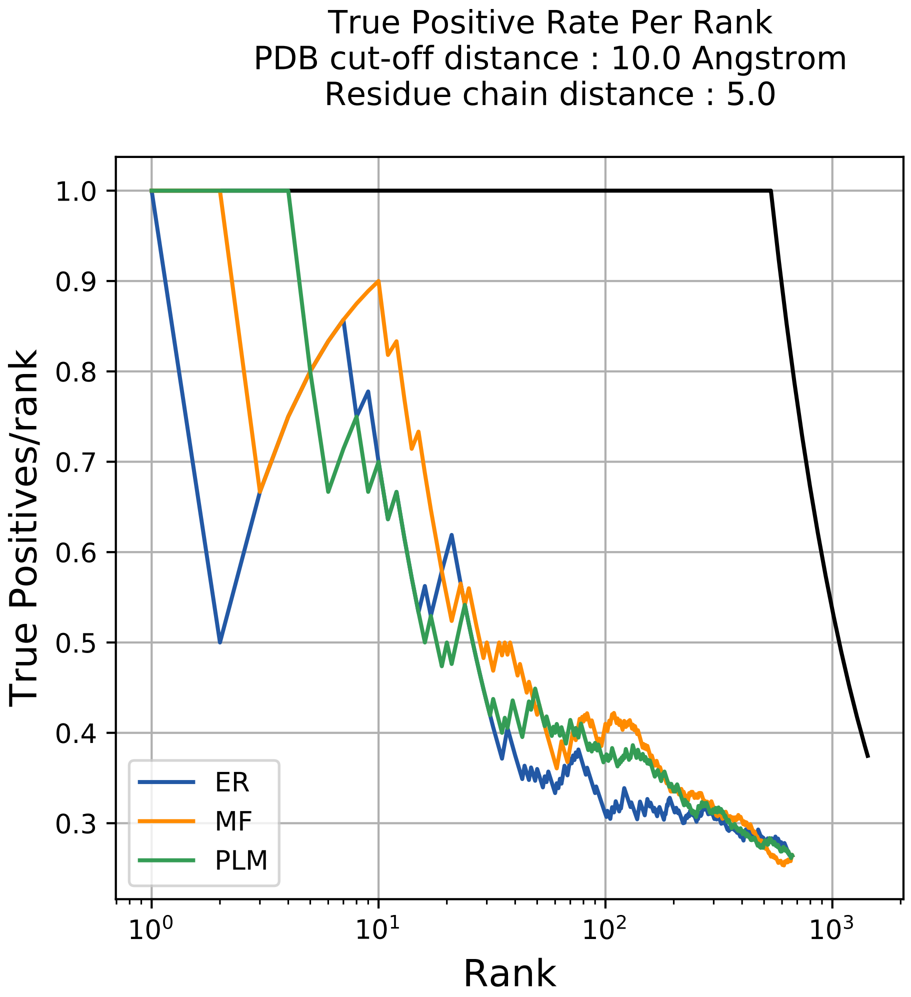

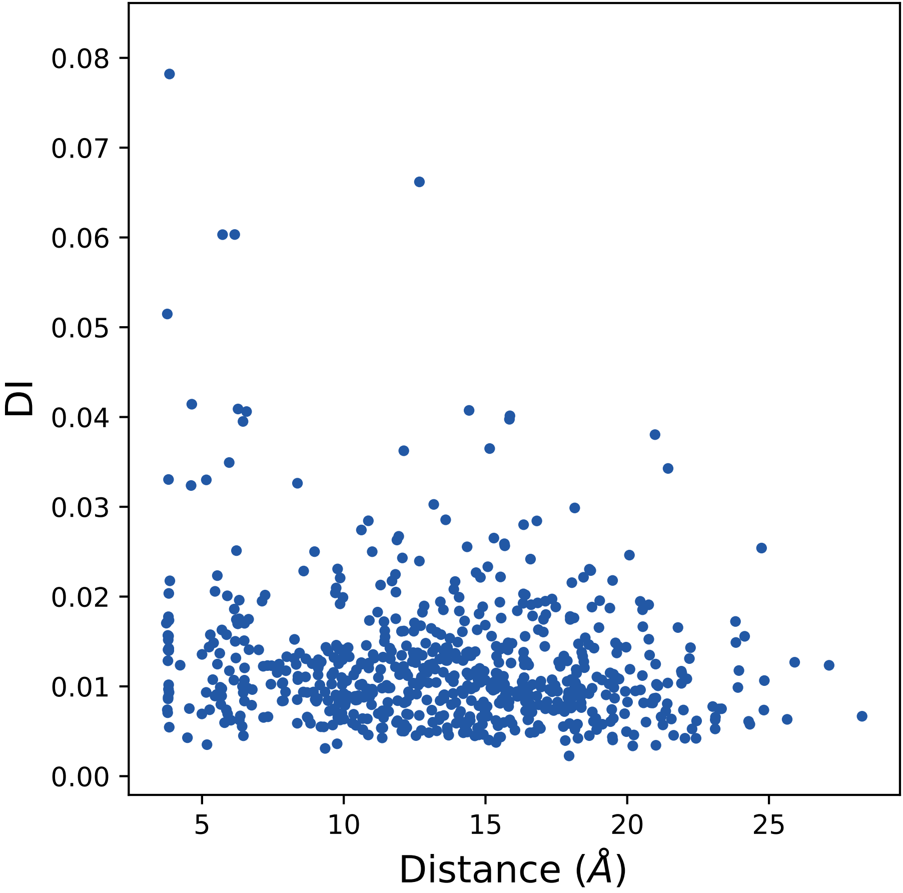

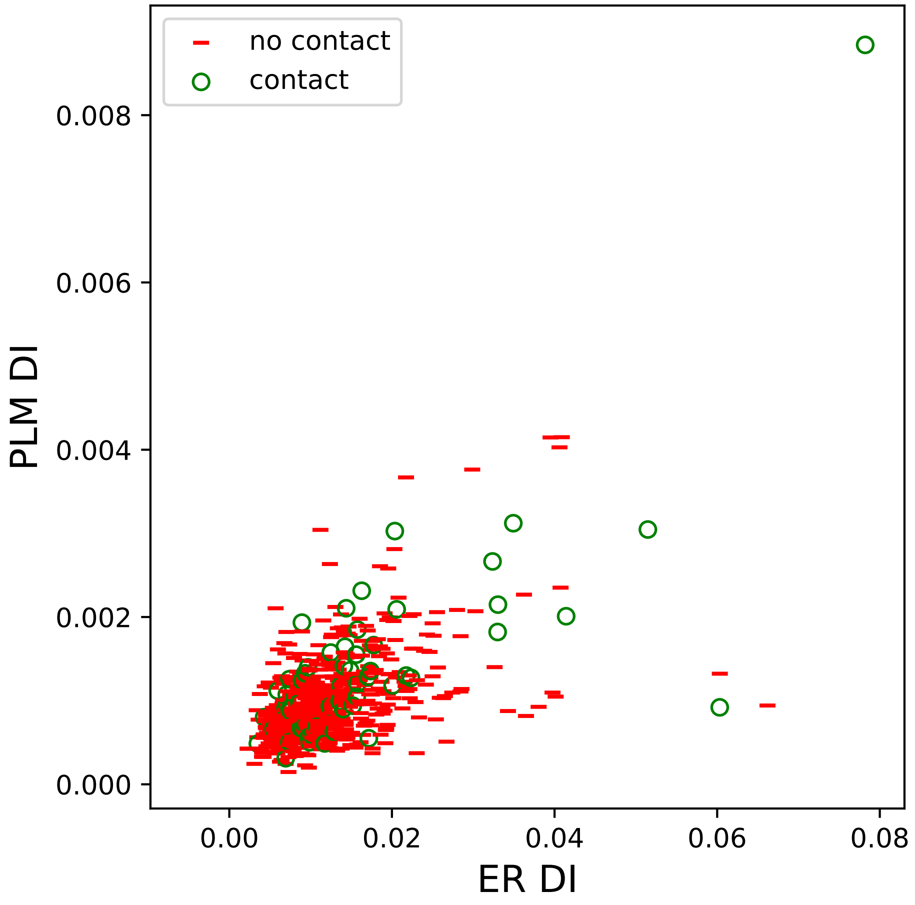

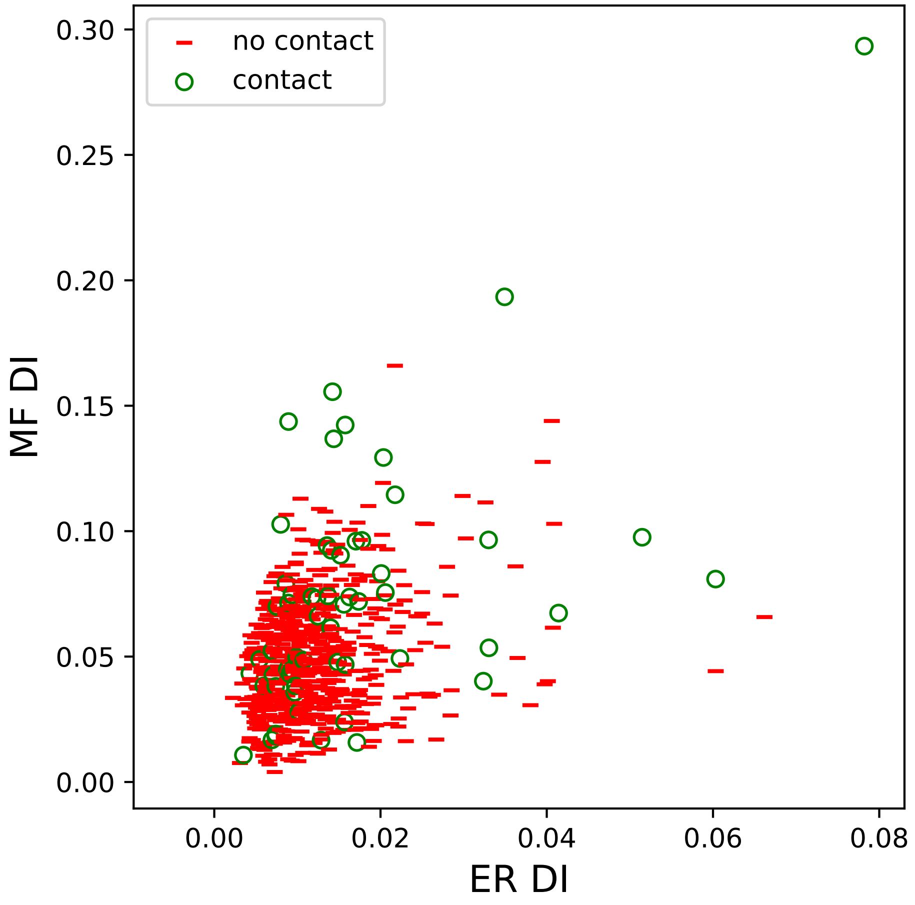

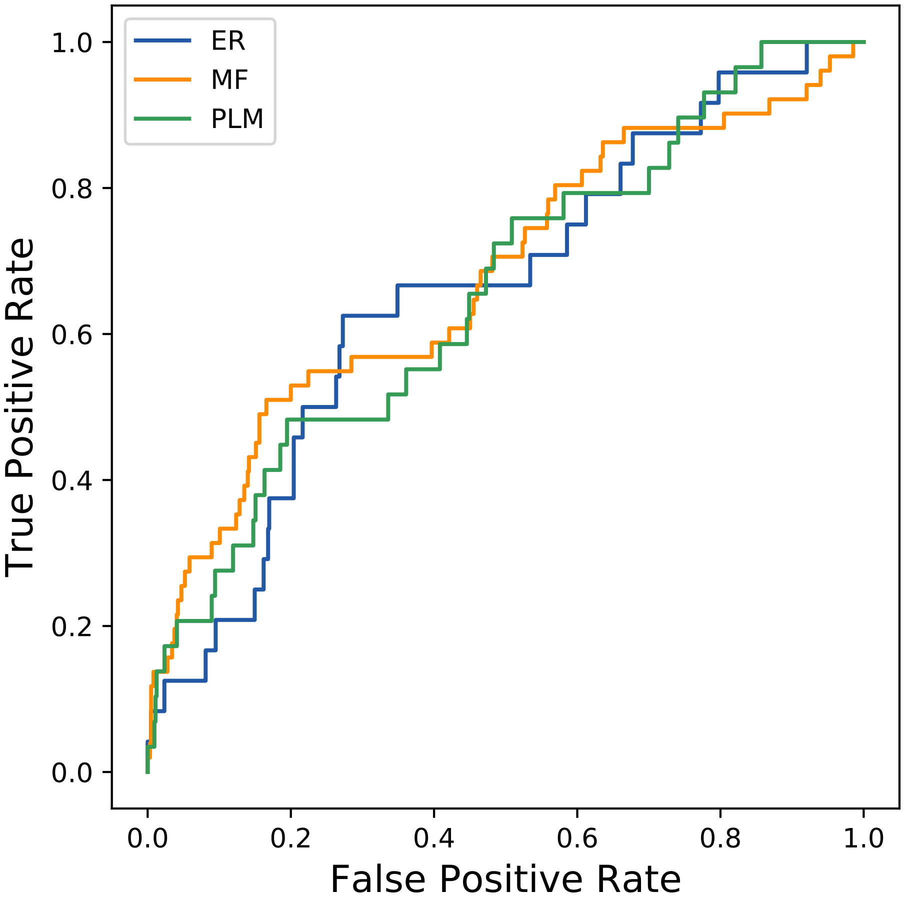

### 1894seq_100col_1ifv_PF00407_plots.pdf

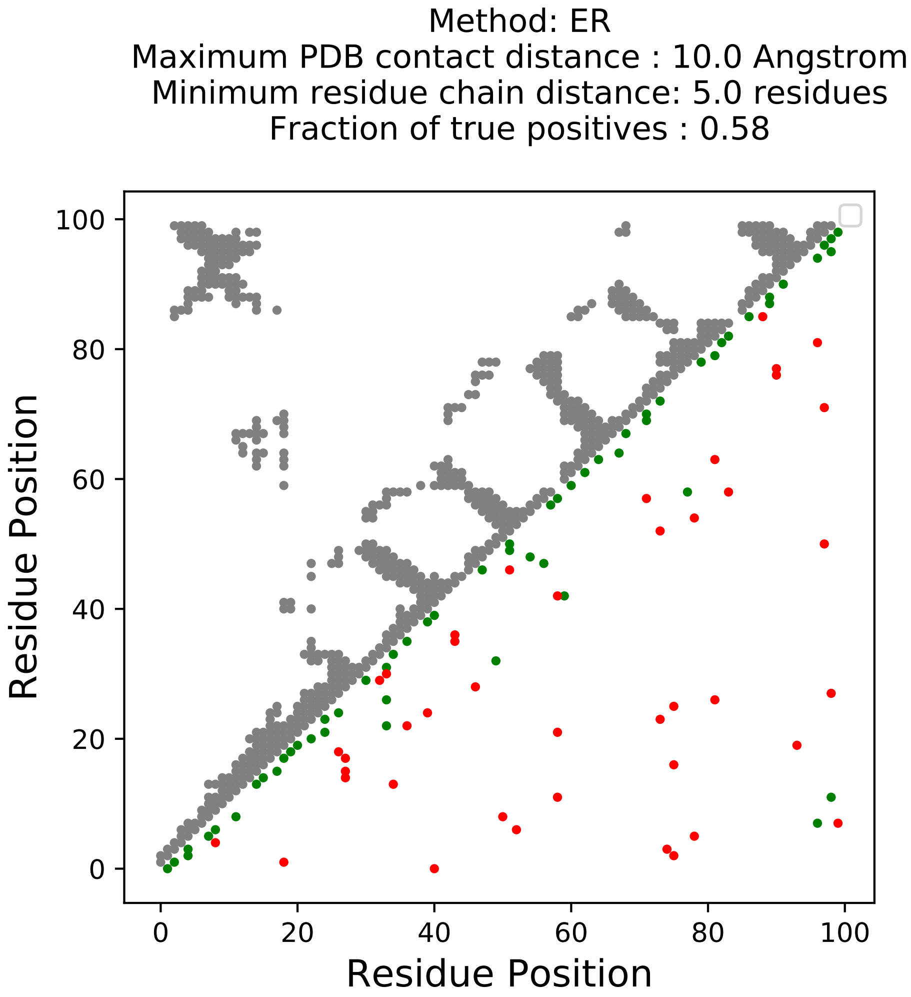

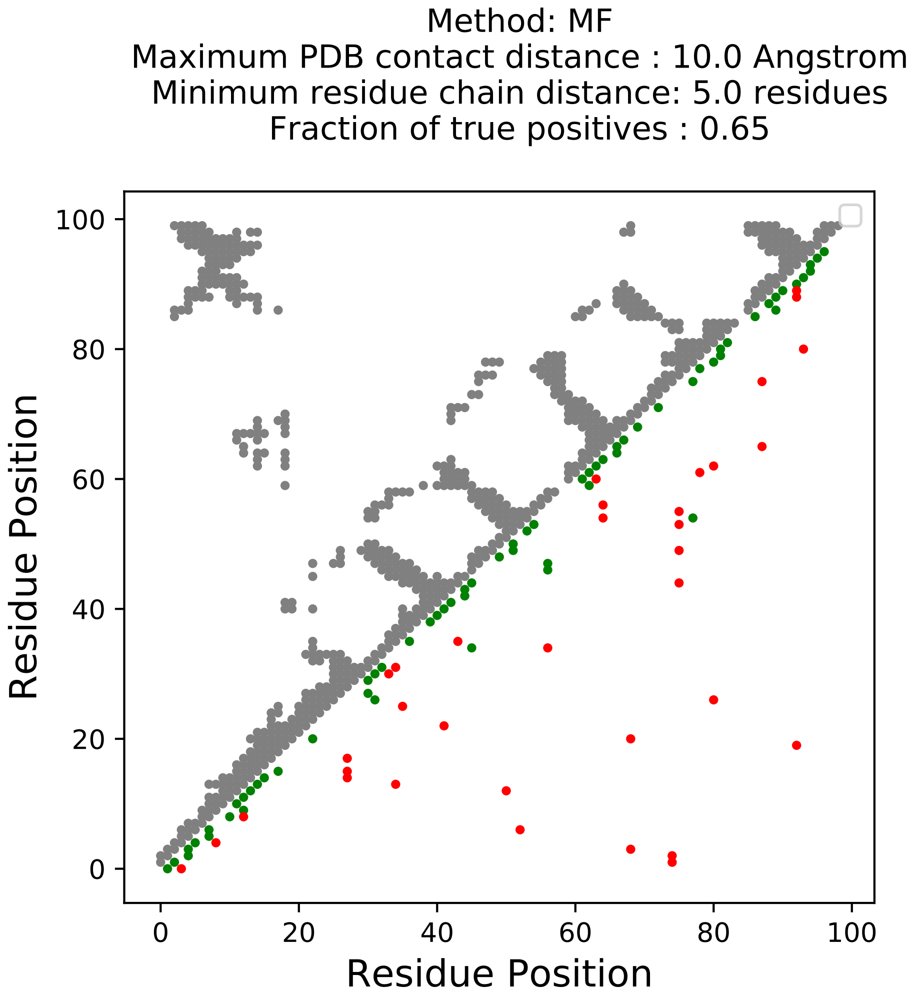

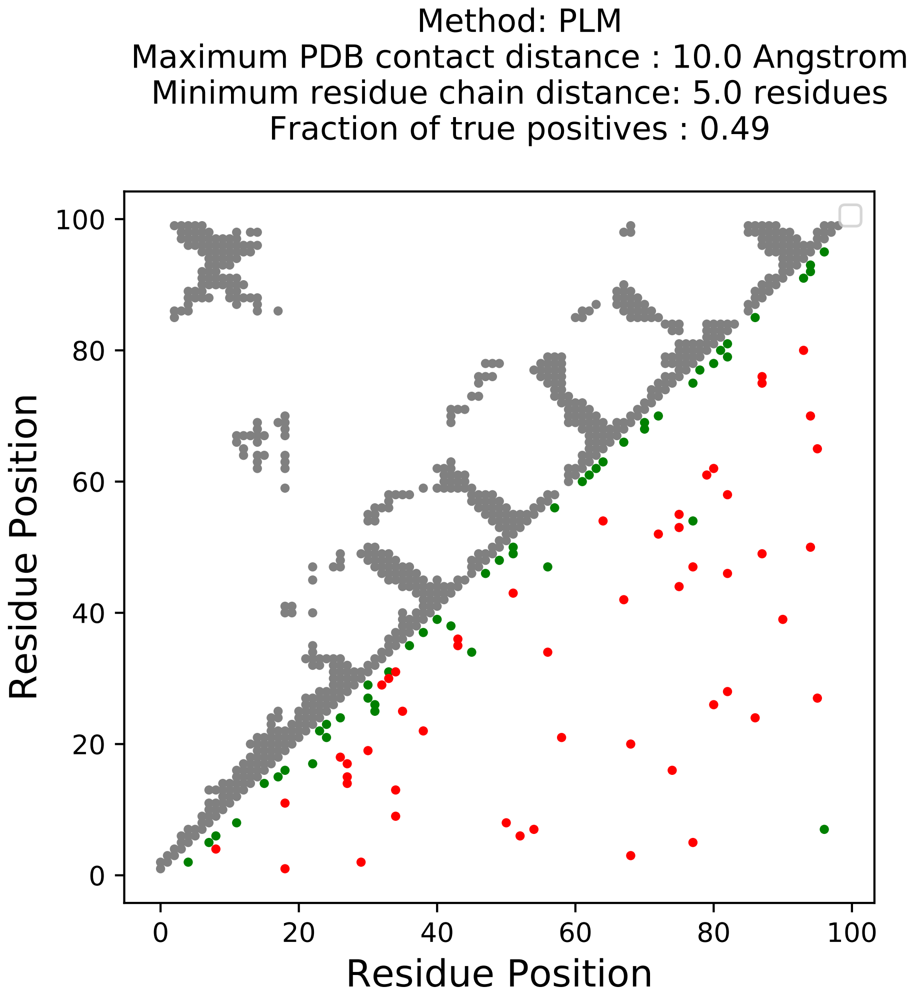

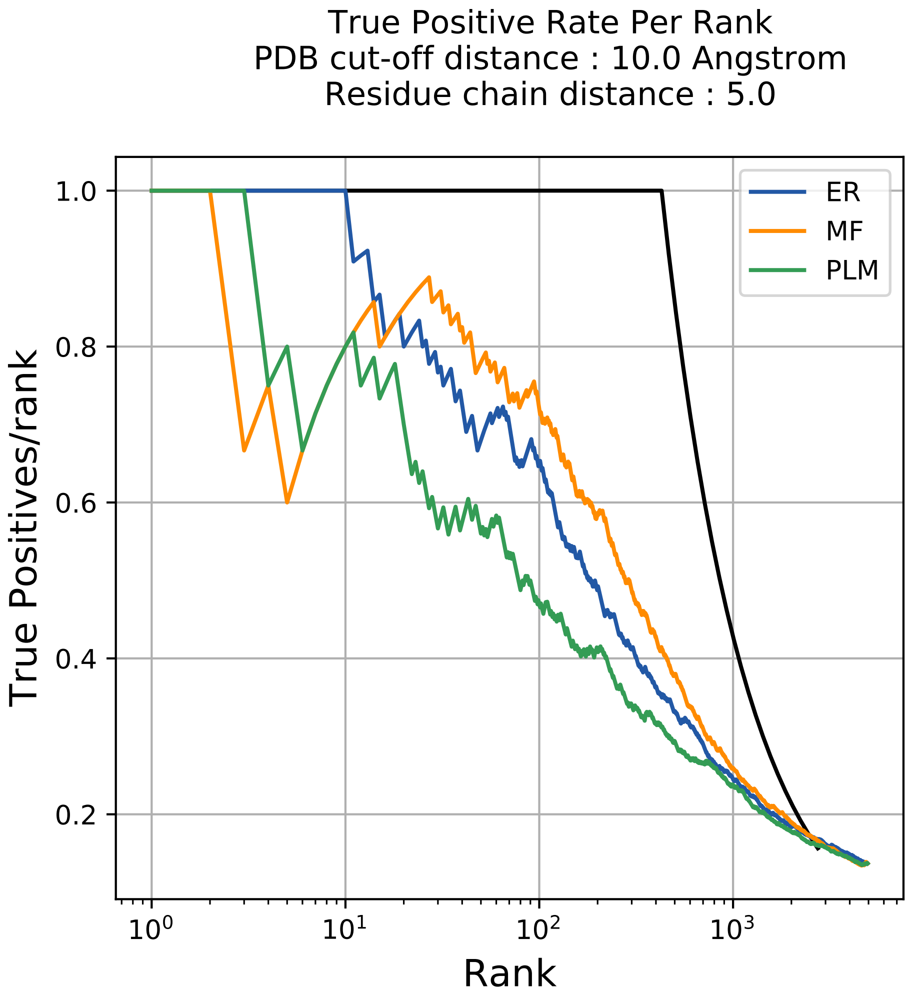

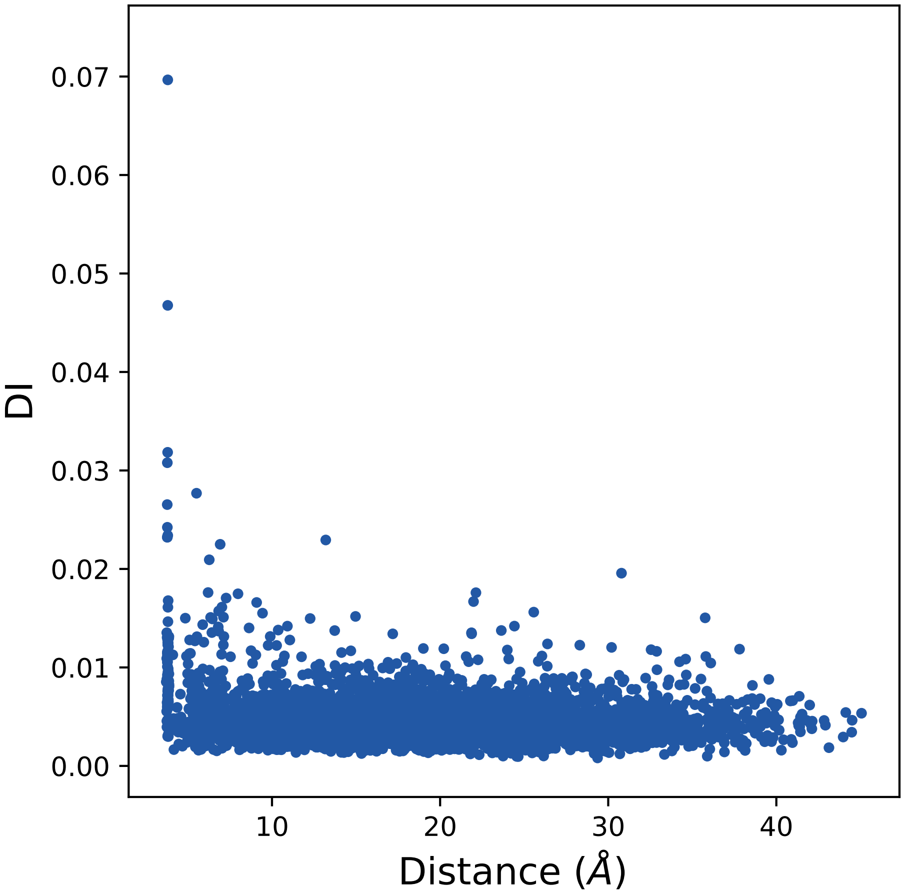

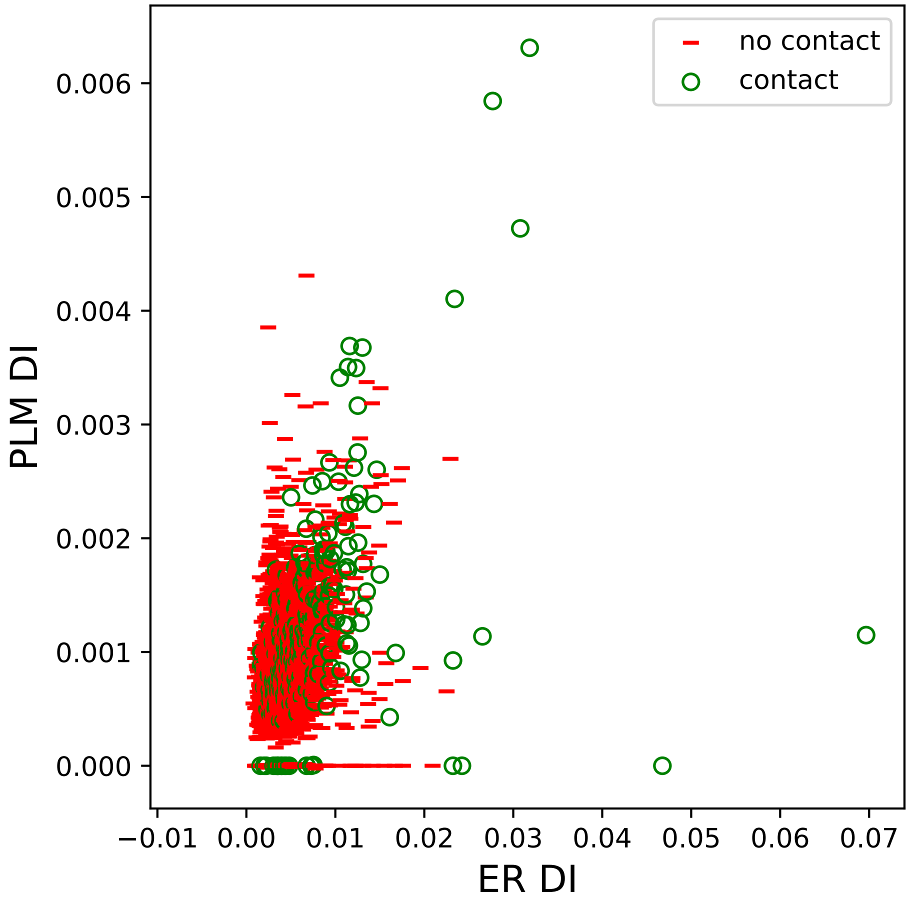

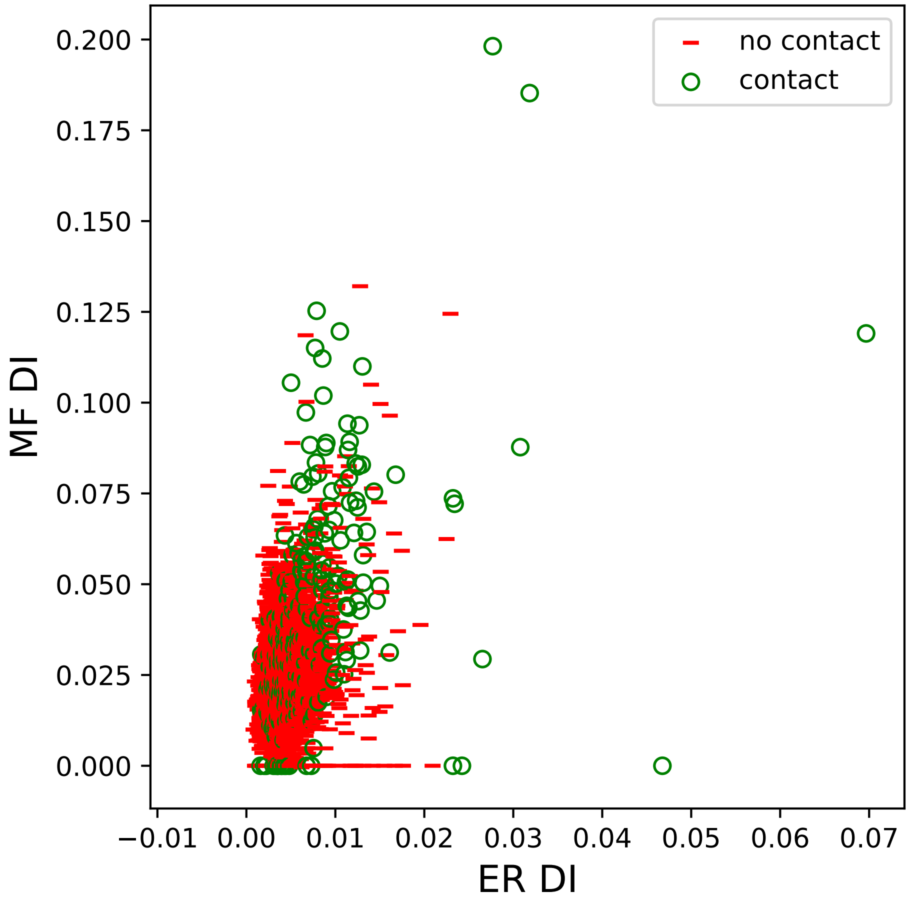

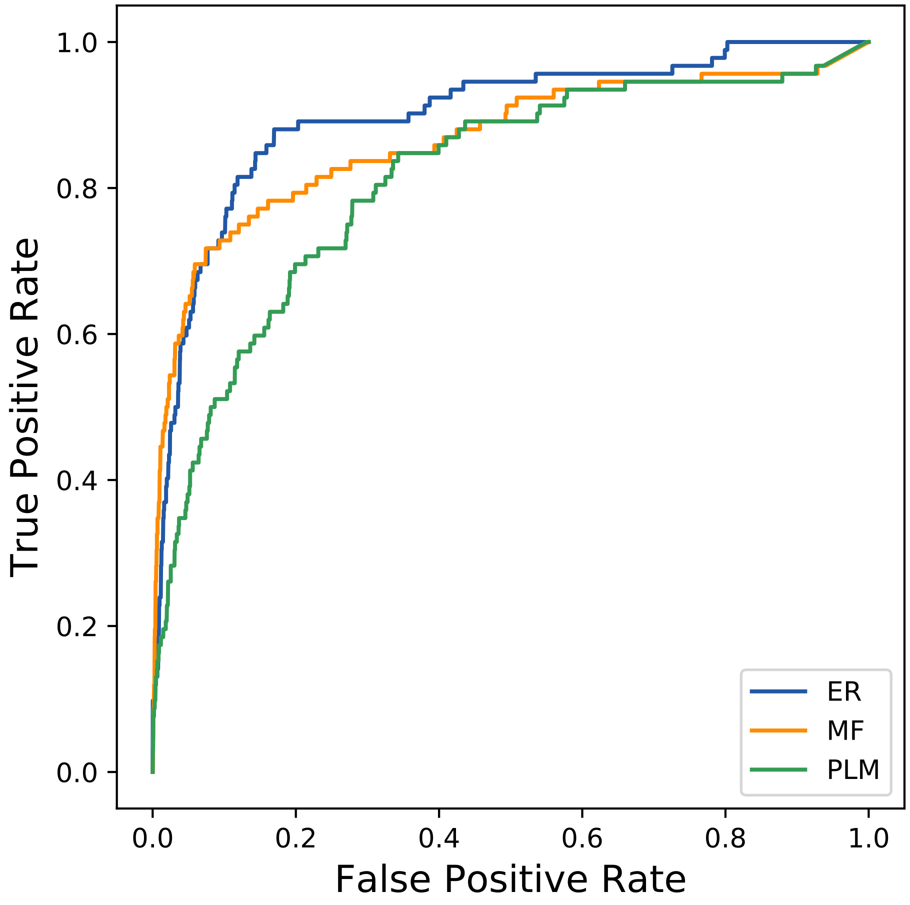

### 1938seq_159col_7ck1_PF03552_plots.pdf

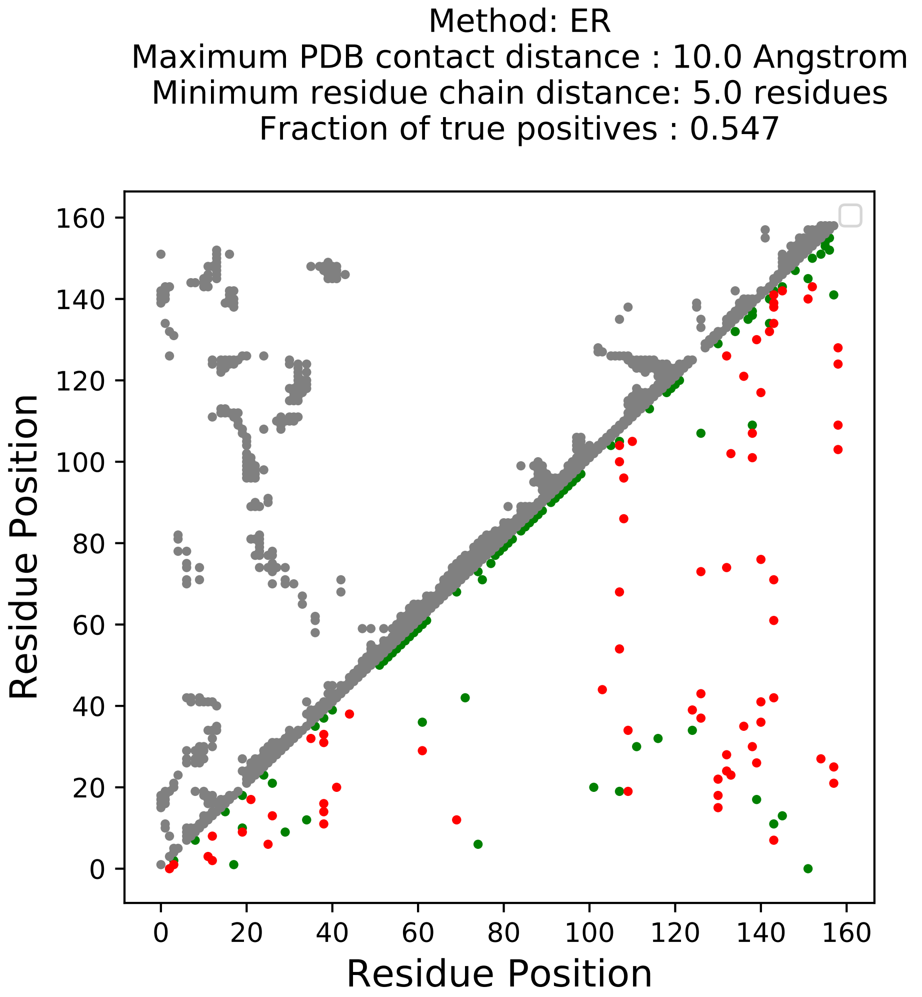

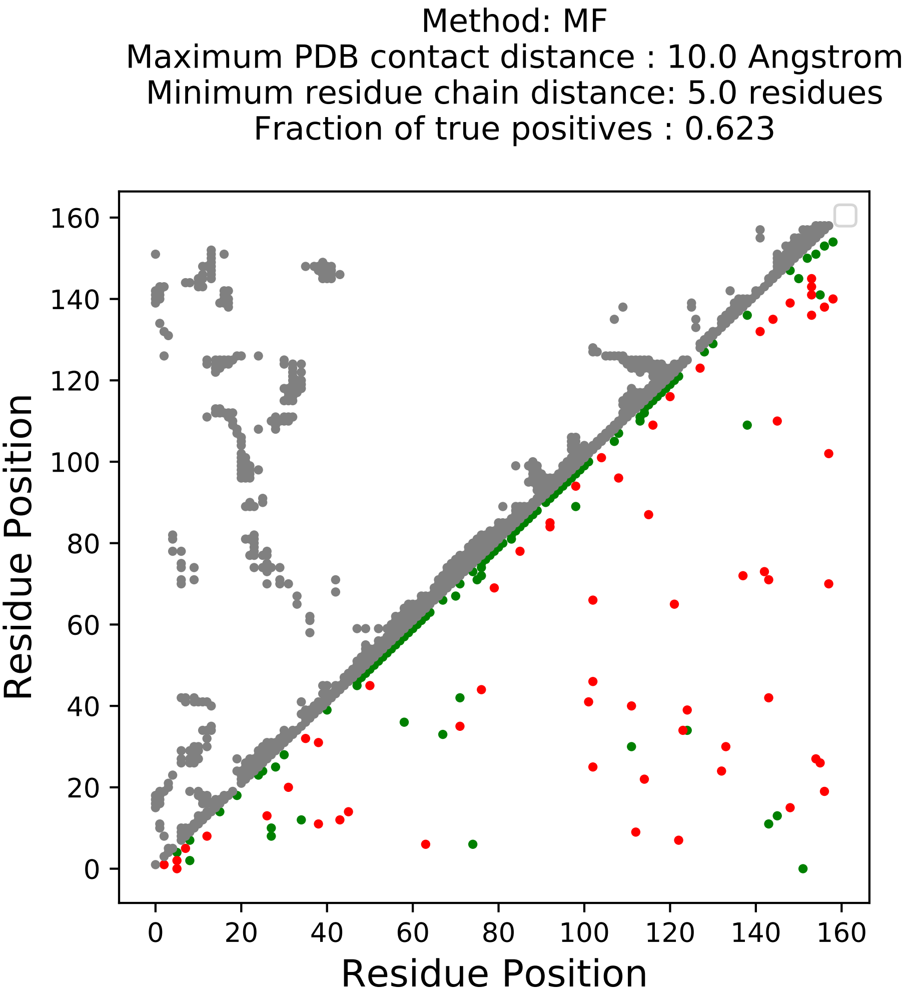

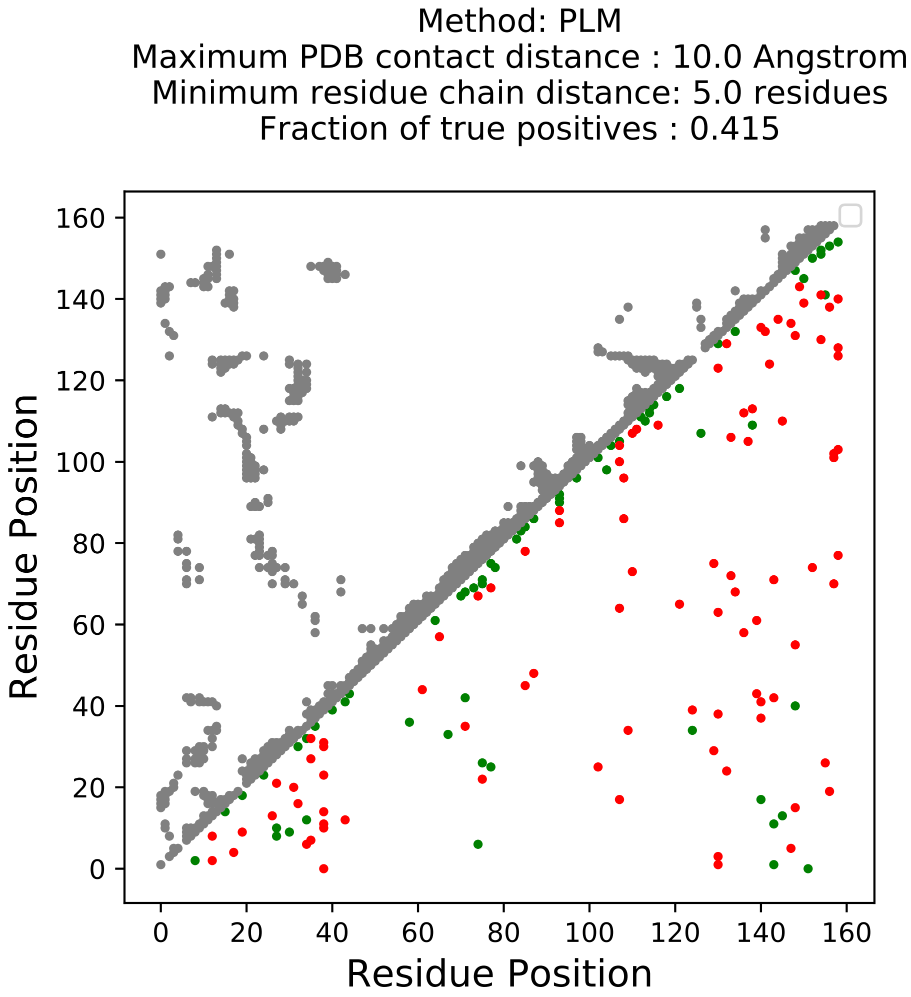

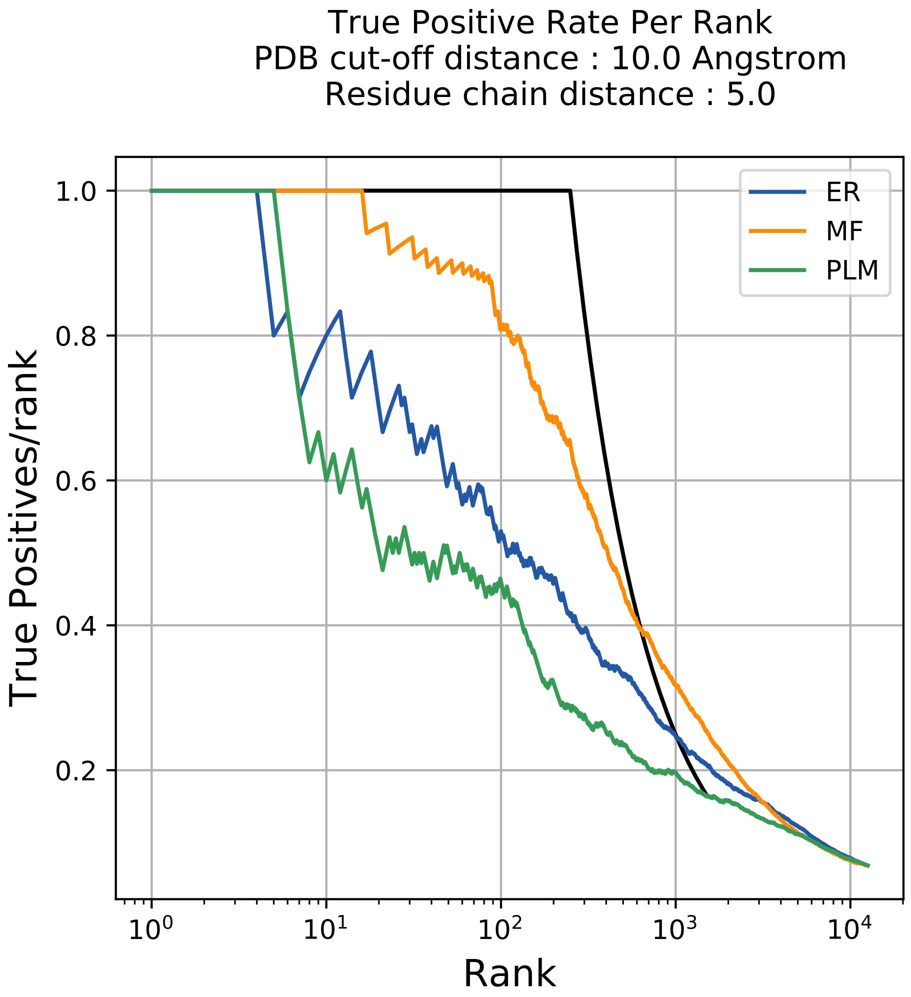

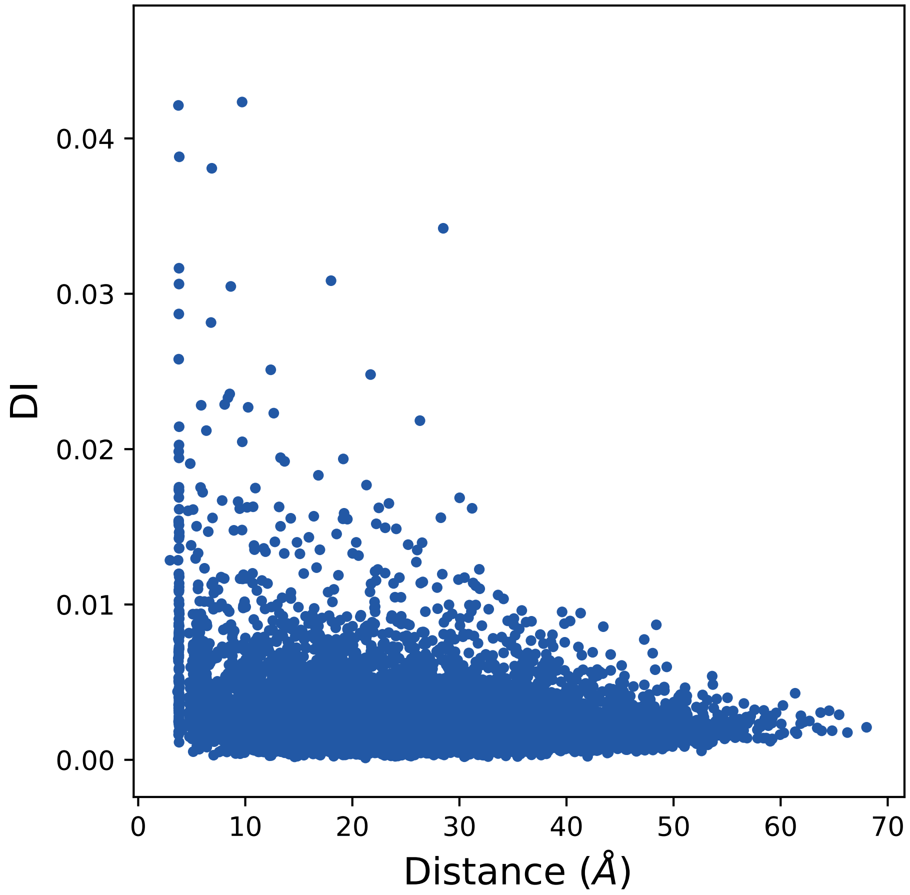

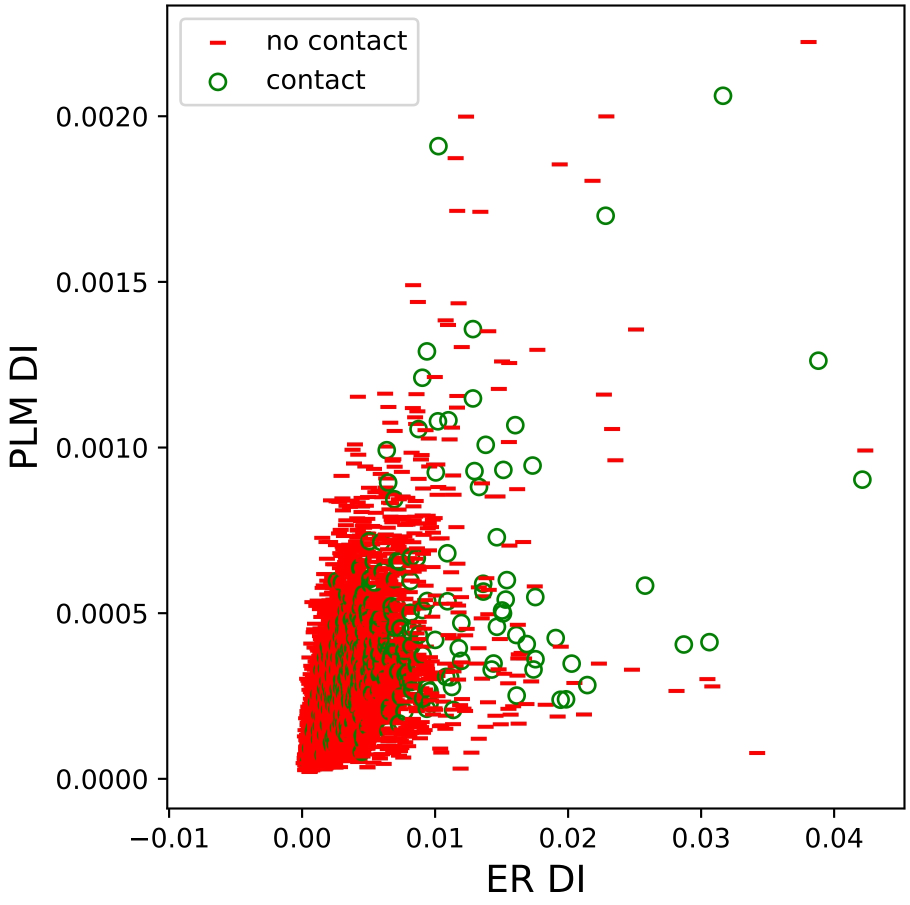

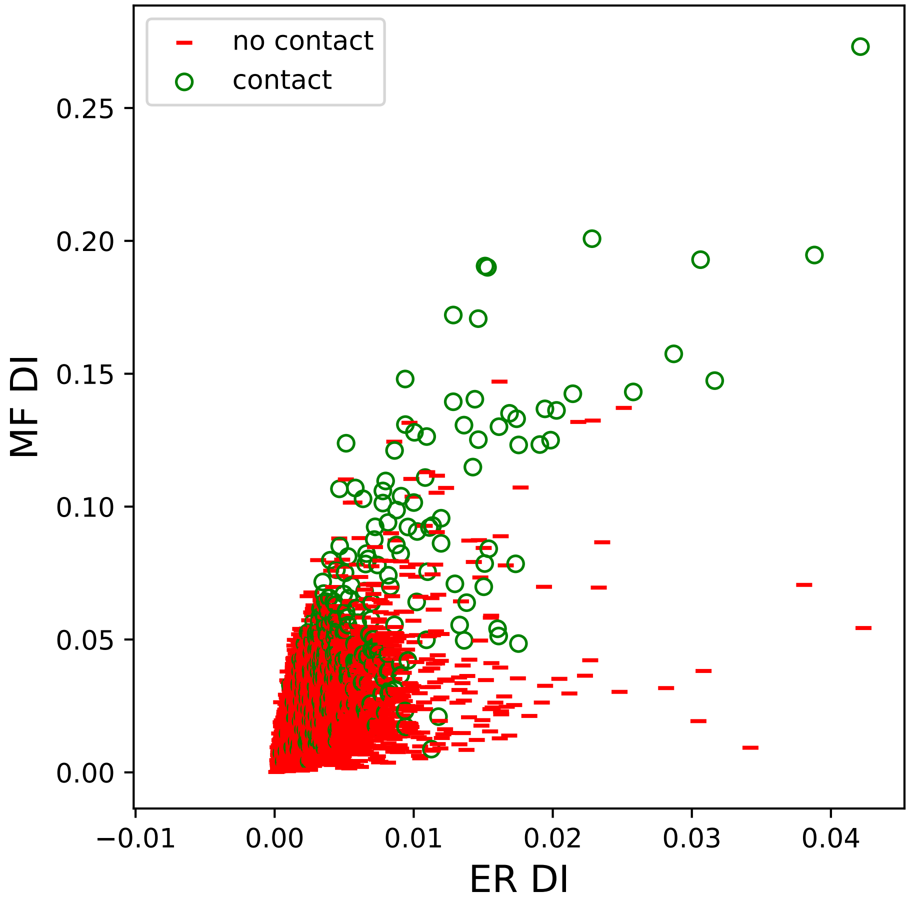

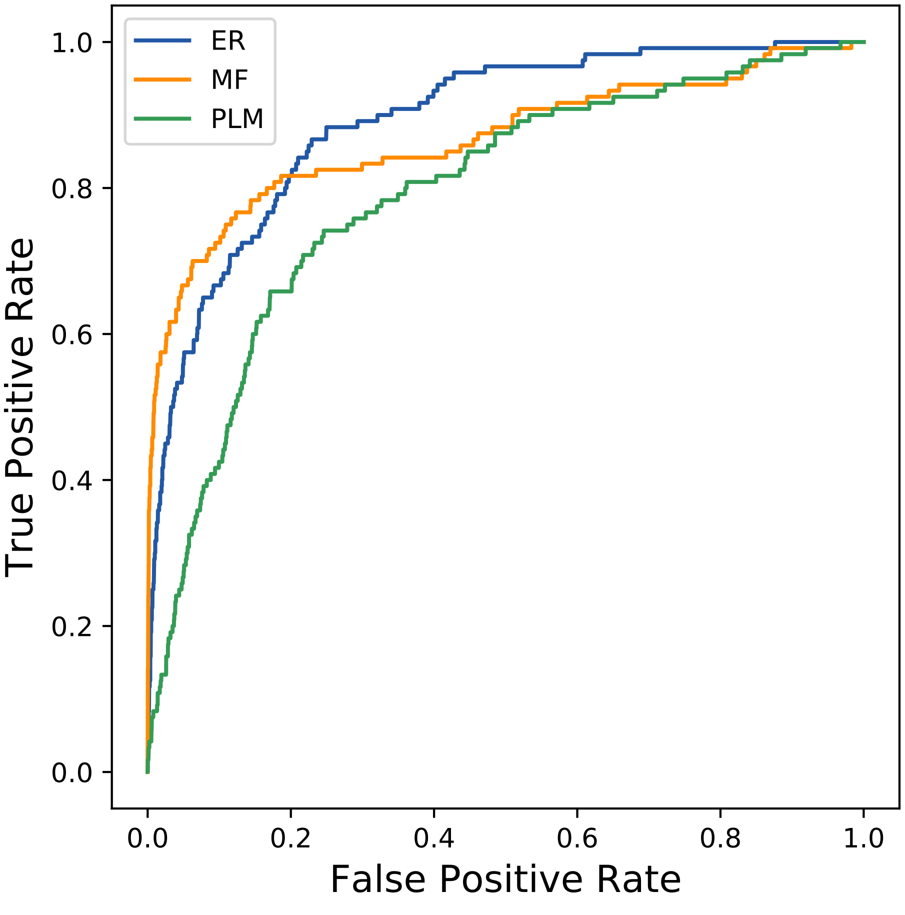

### 1955seq_166col_6hvh_PF01591_plots.pdf

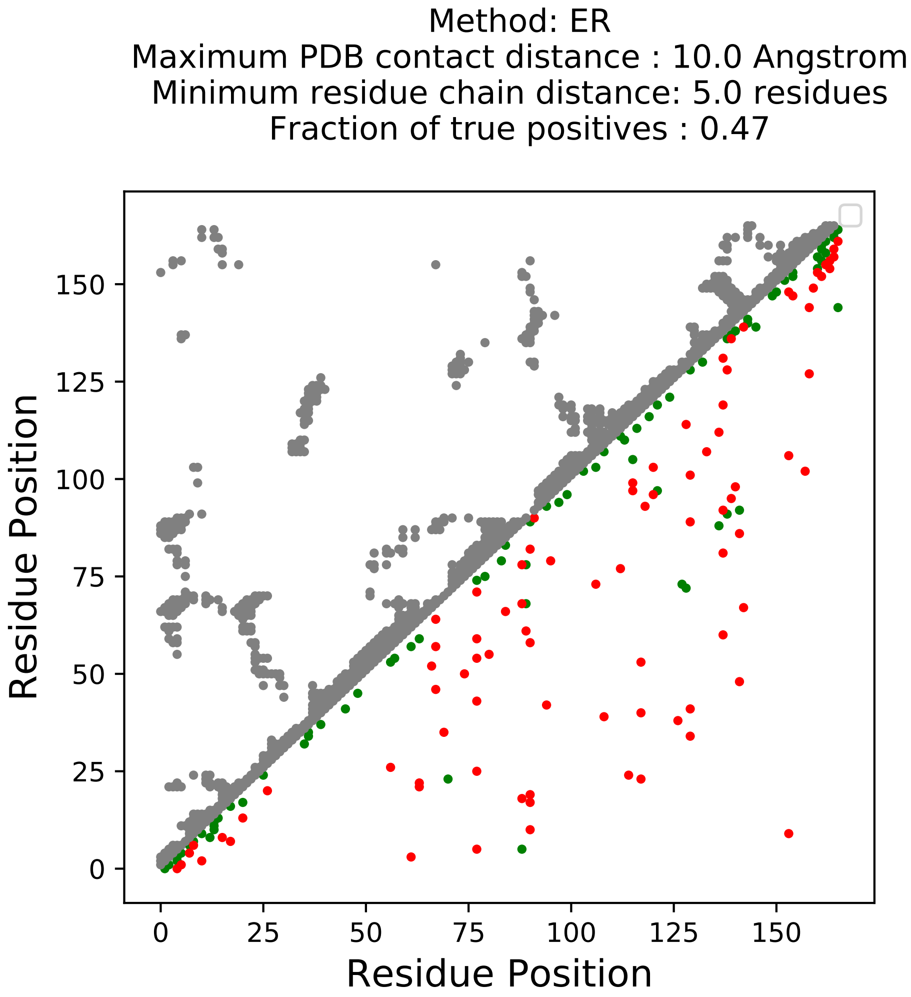

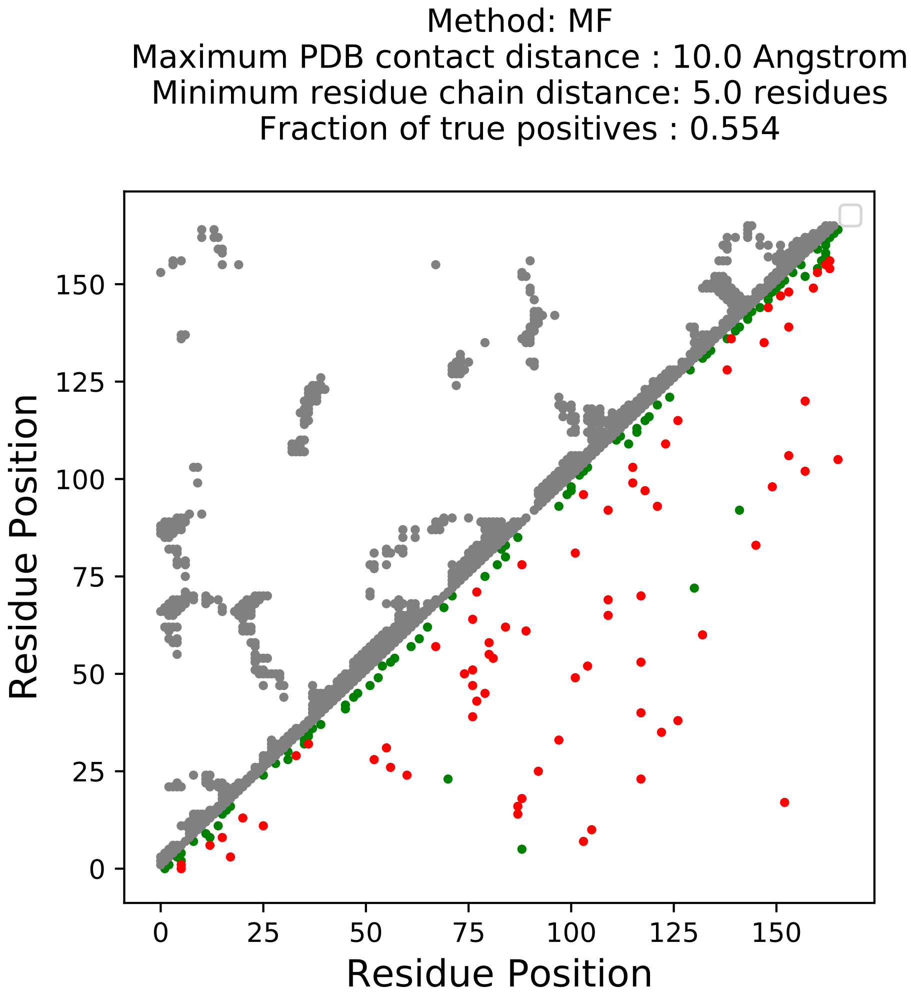

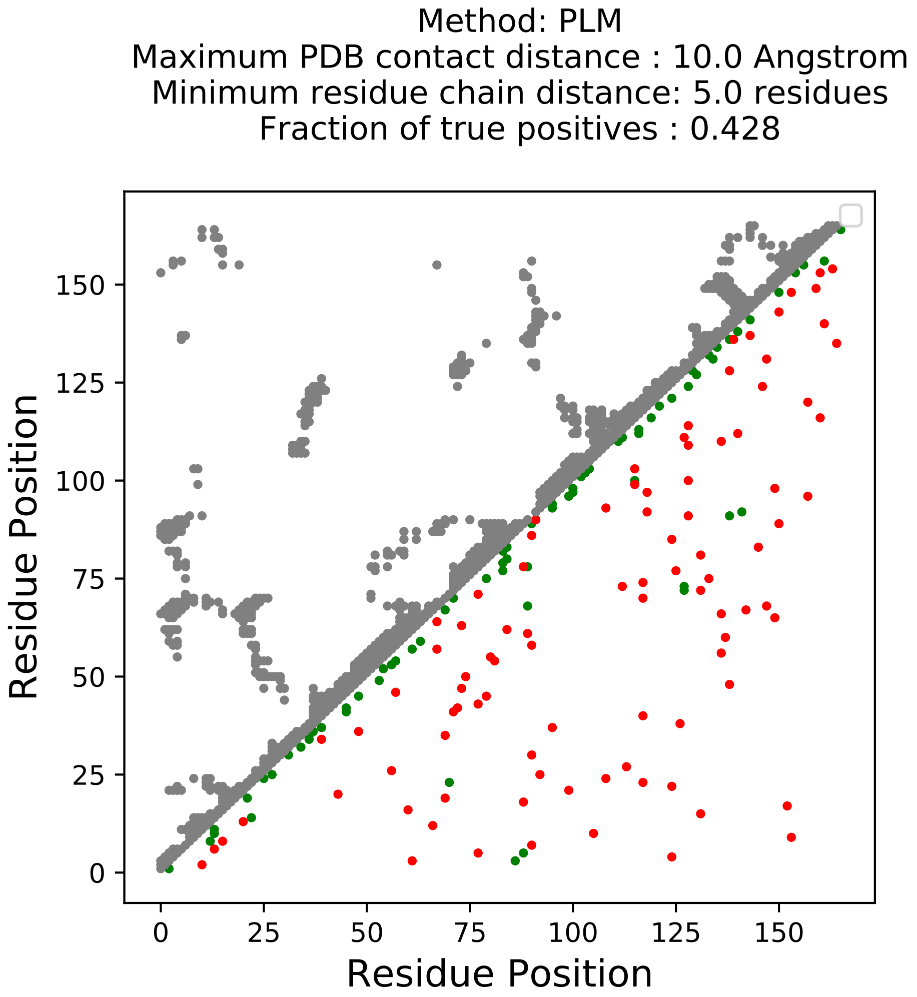

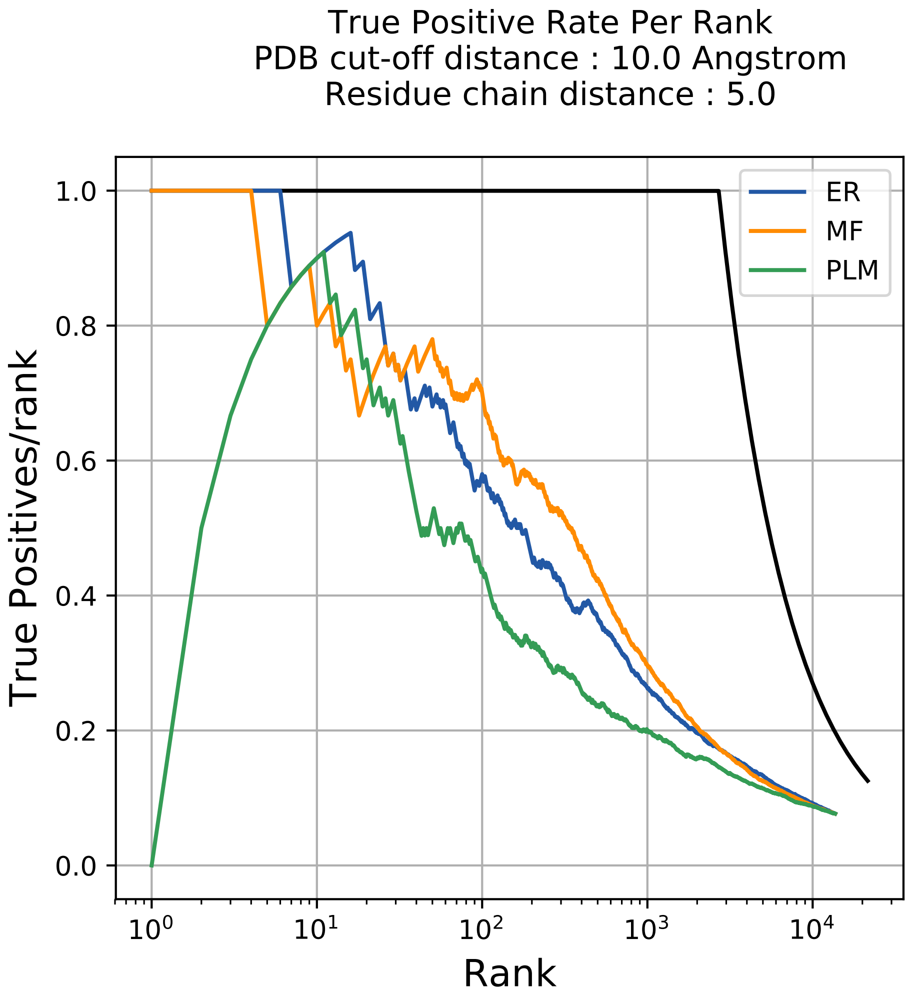

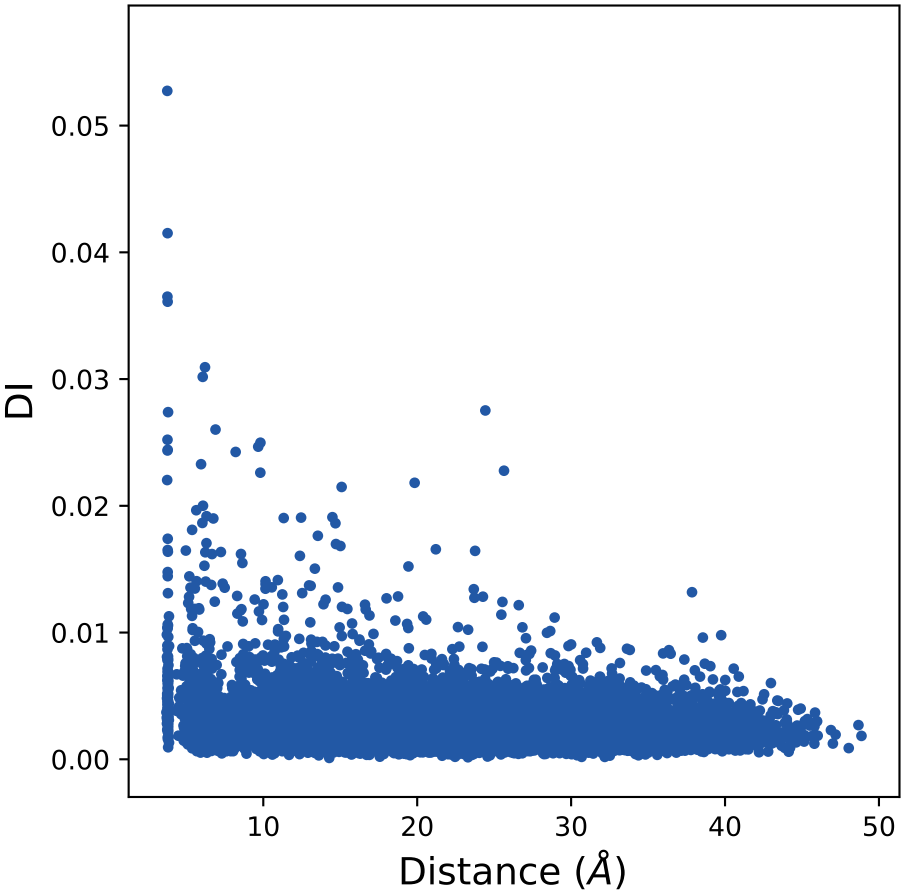

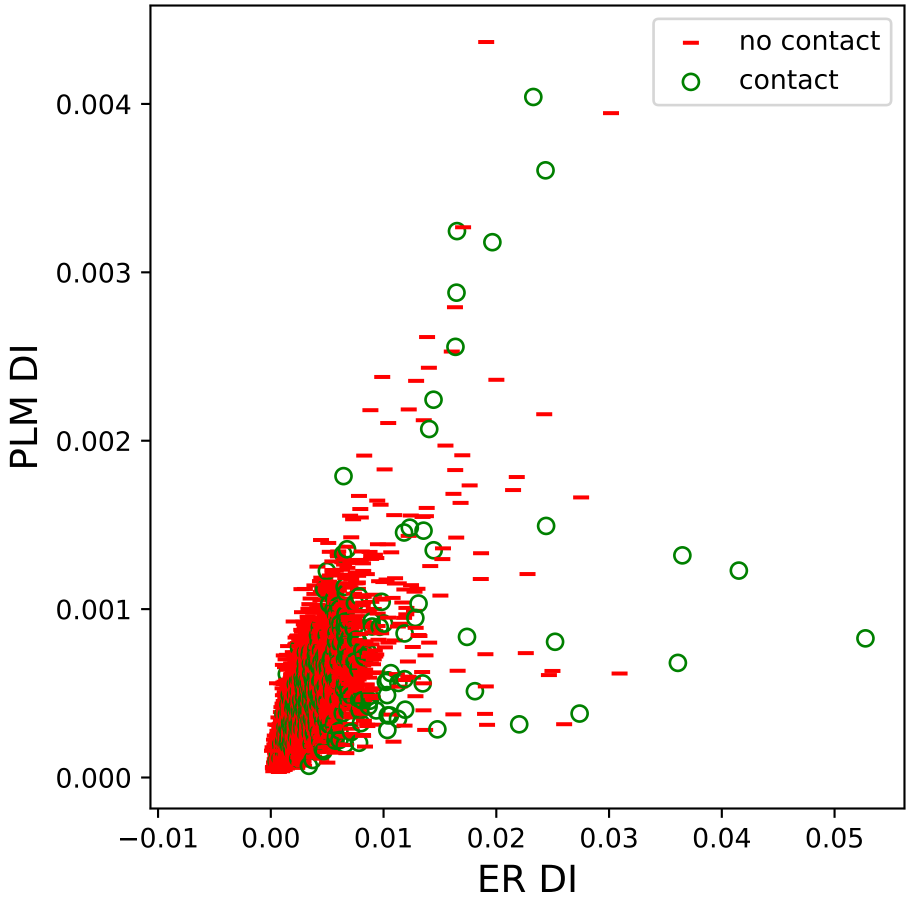

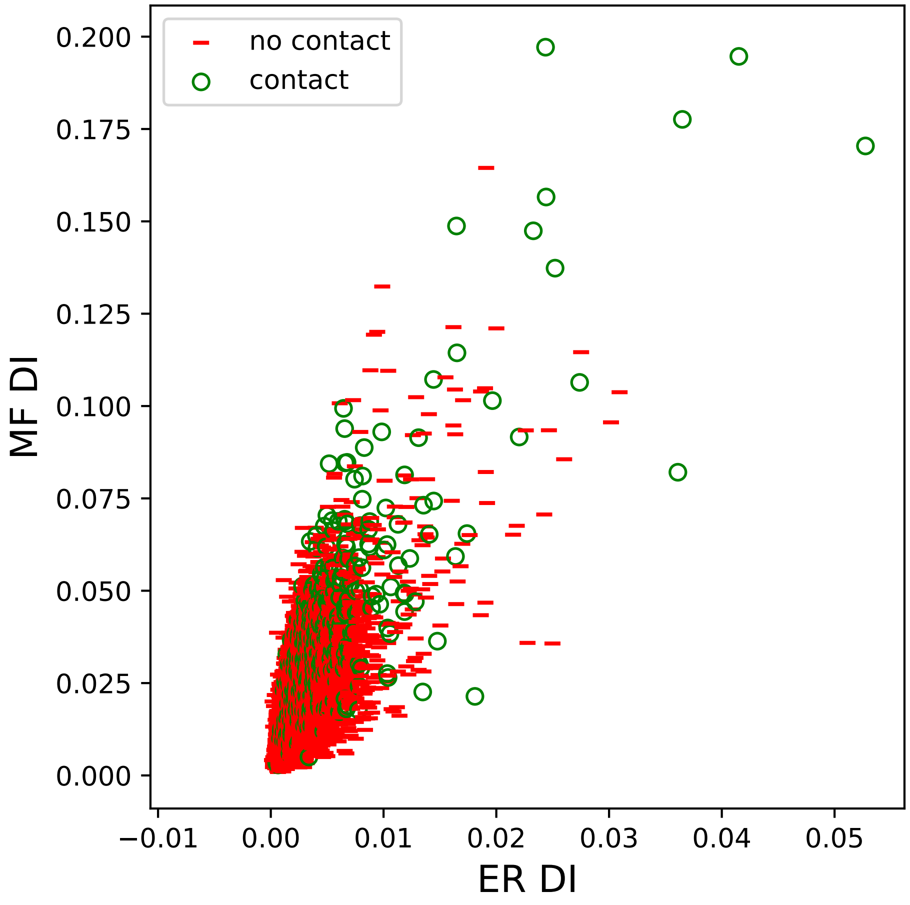

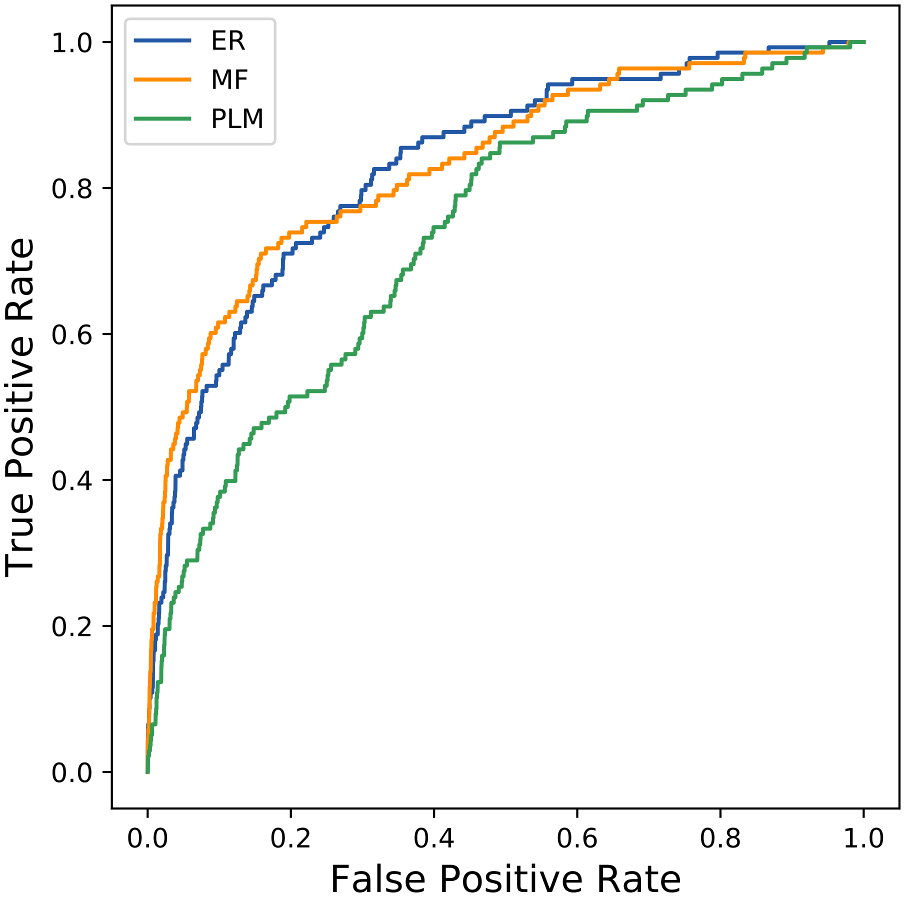

### 1968seq_132col_7s5t_PF17655_plots.pdf

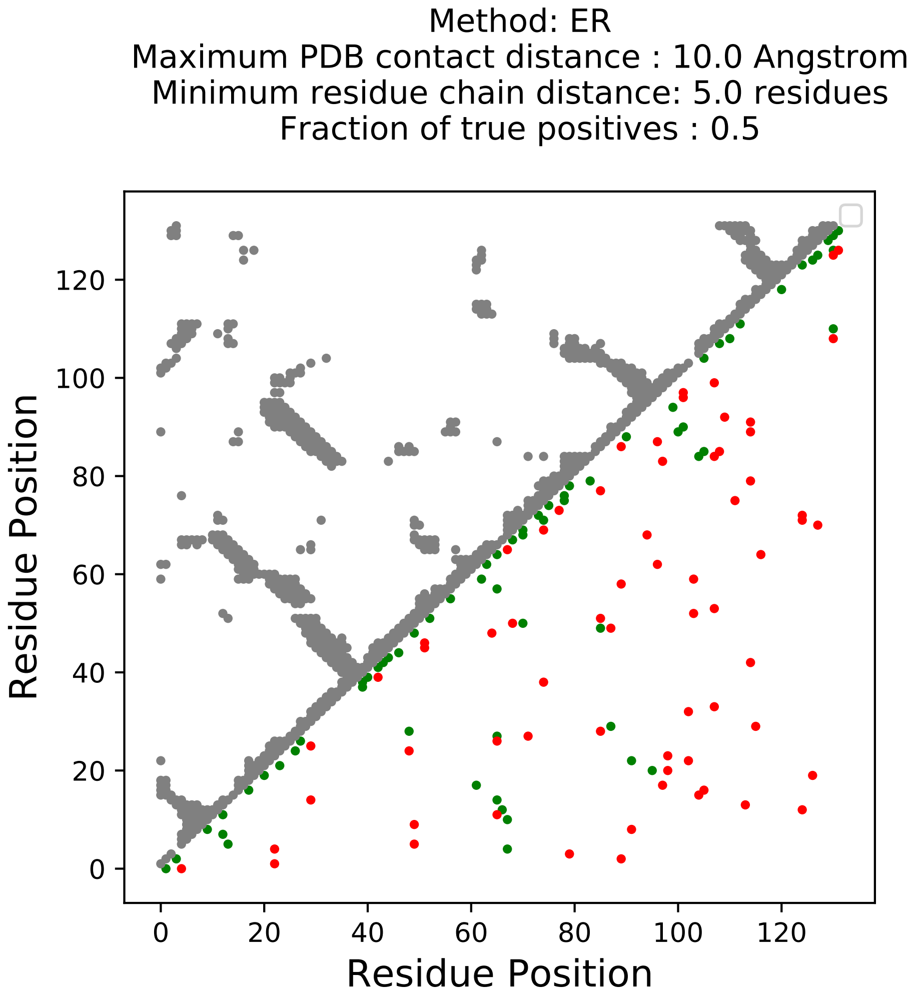

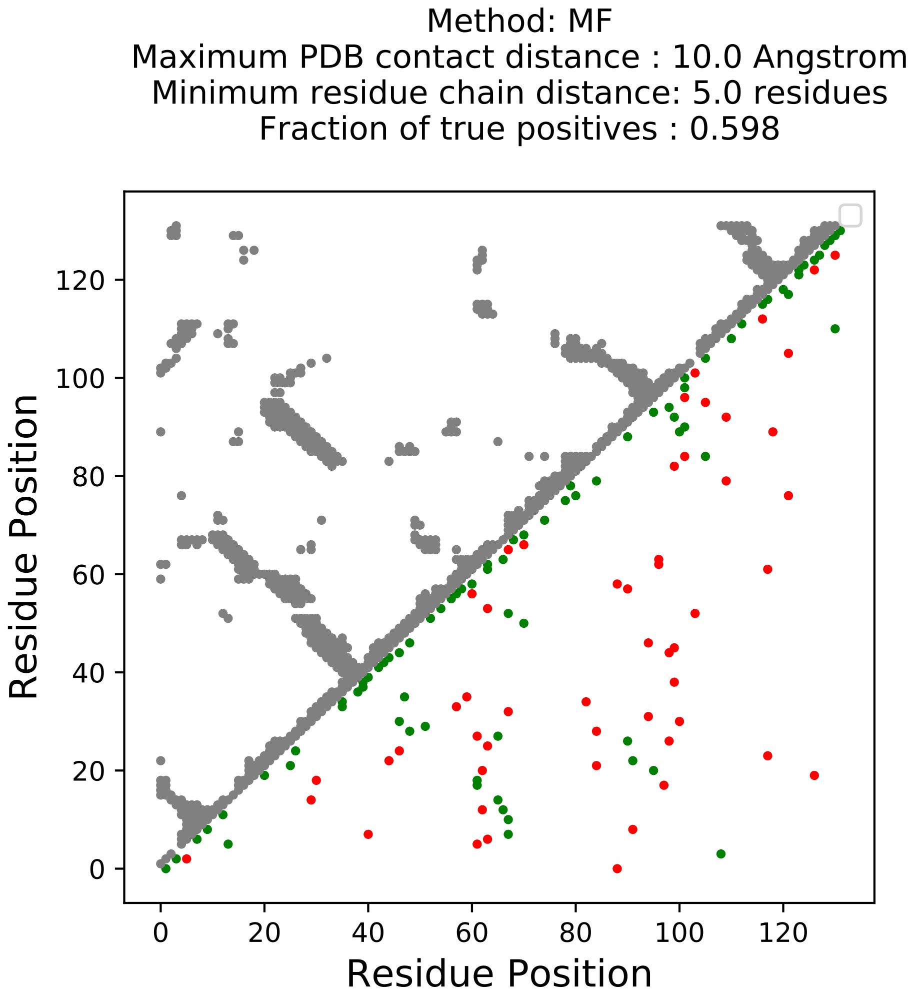

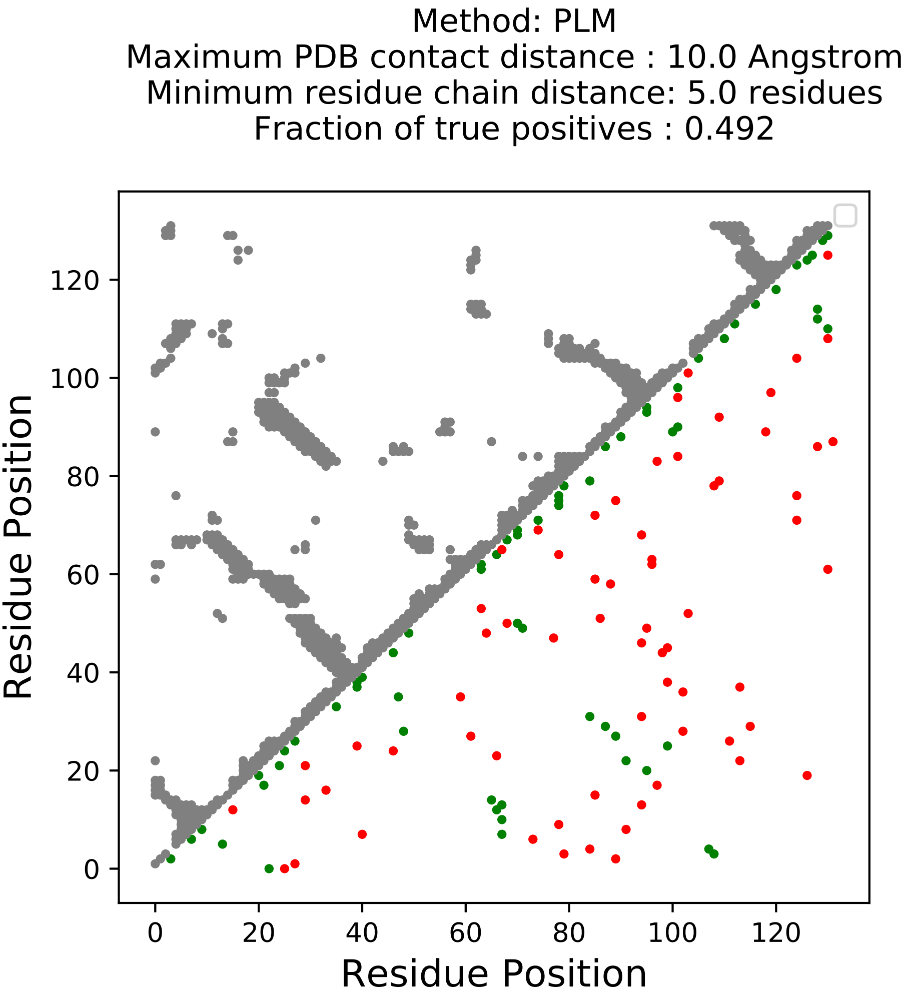

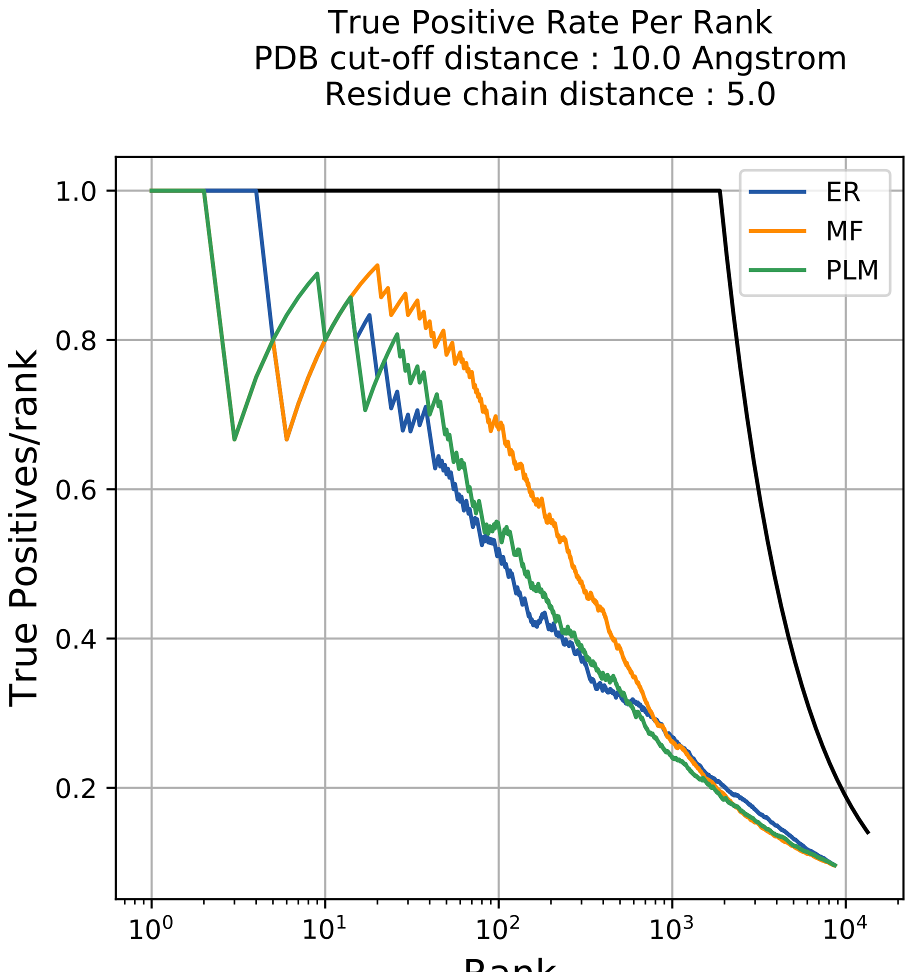

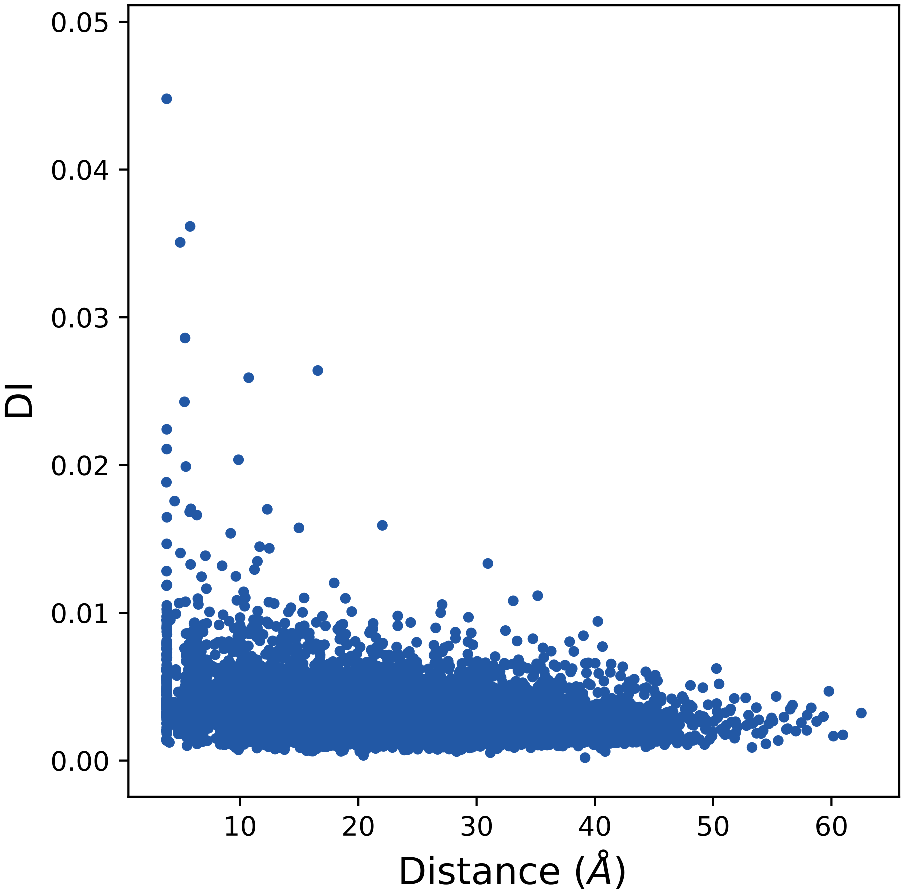

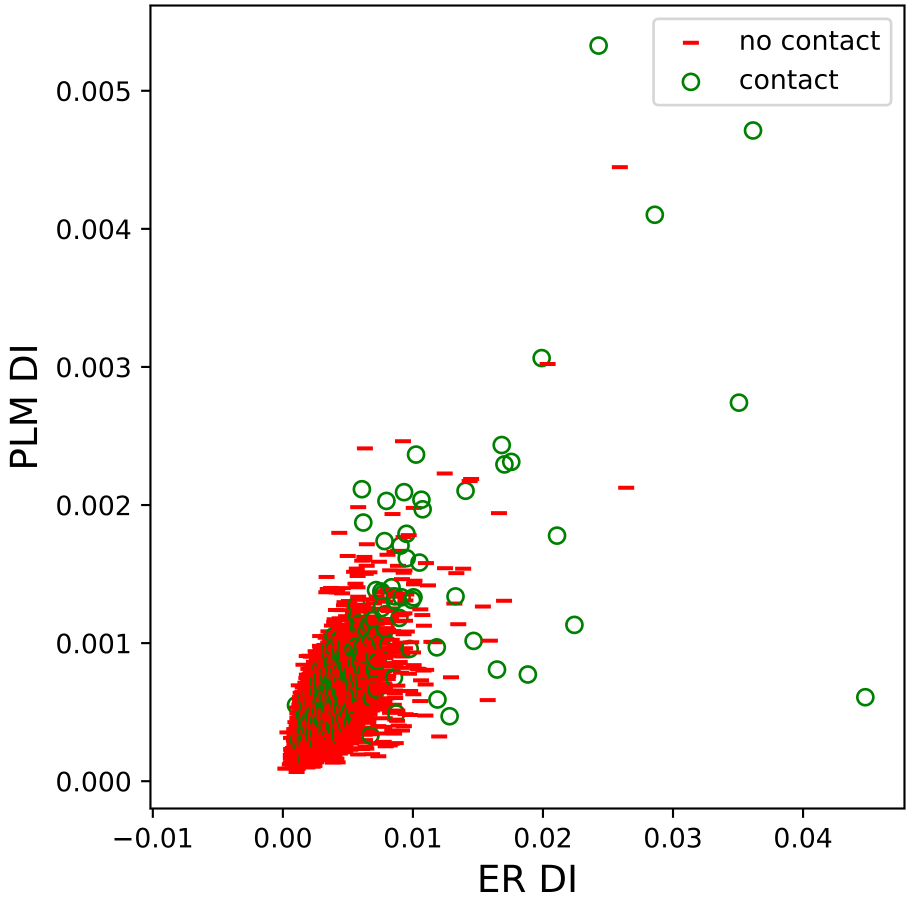

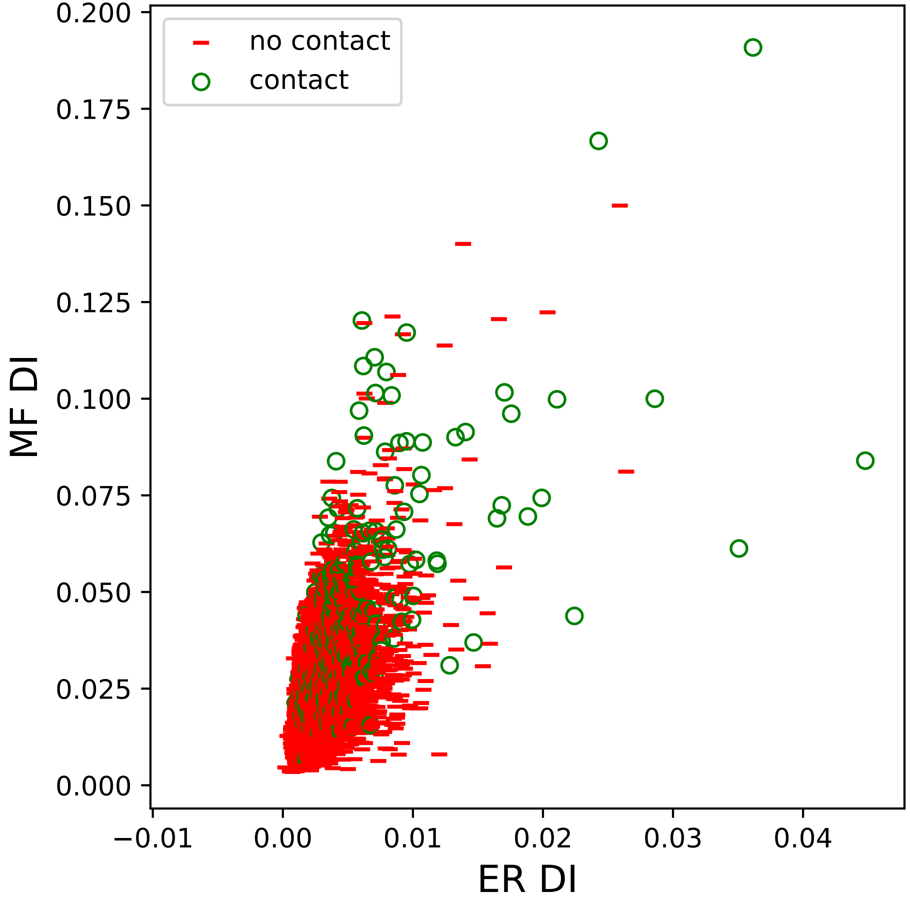

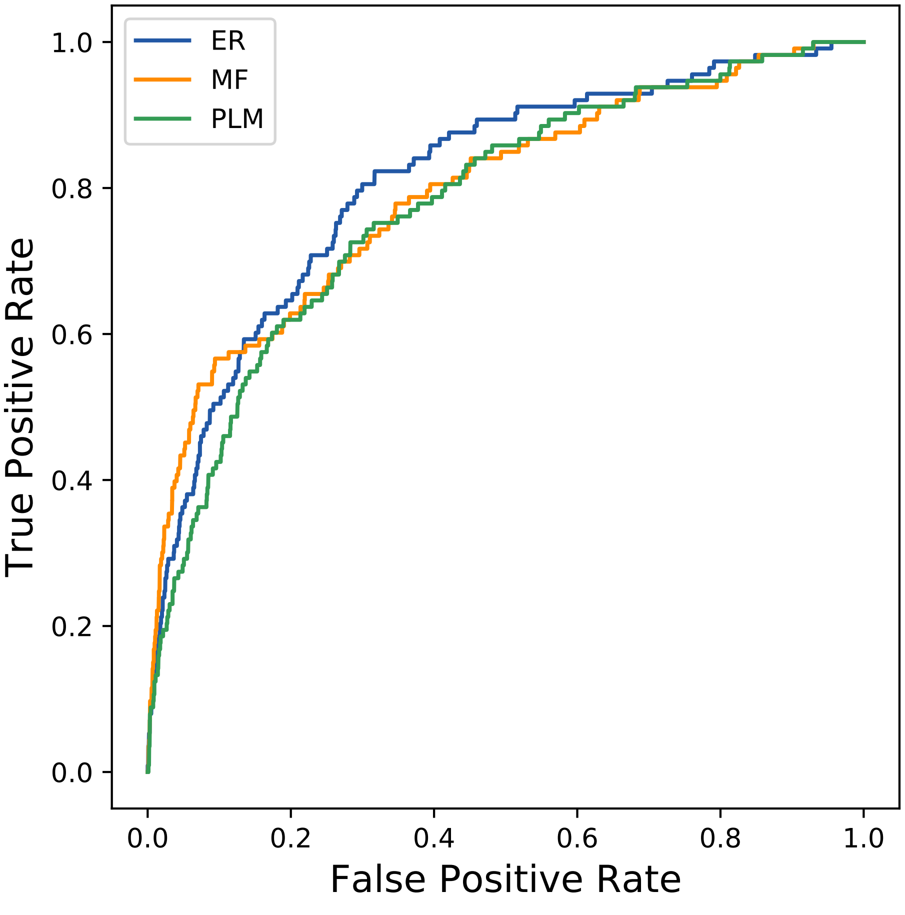

### 2169seq_26col_6zwx_PF11898_plots.pdf

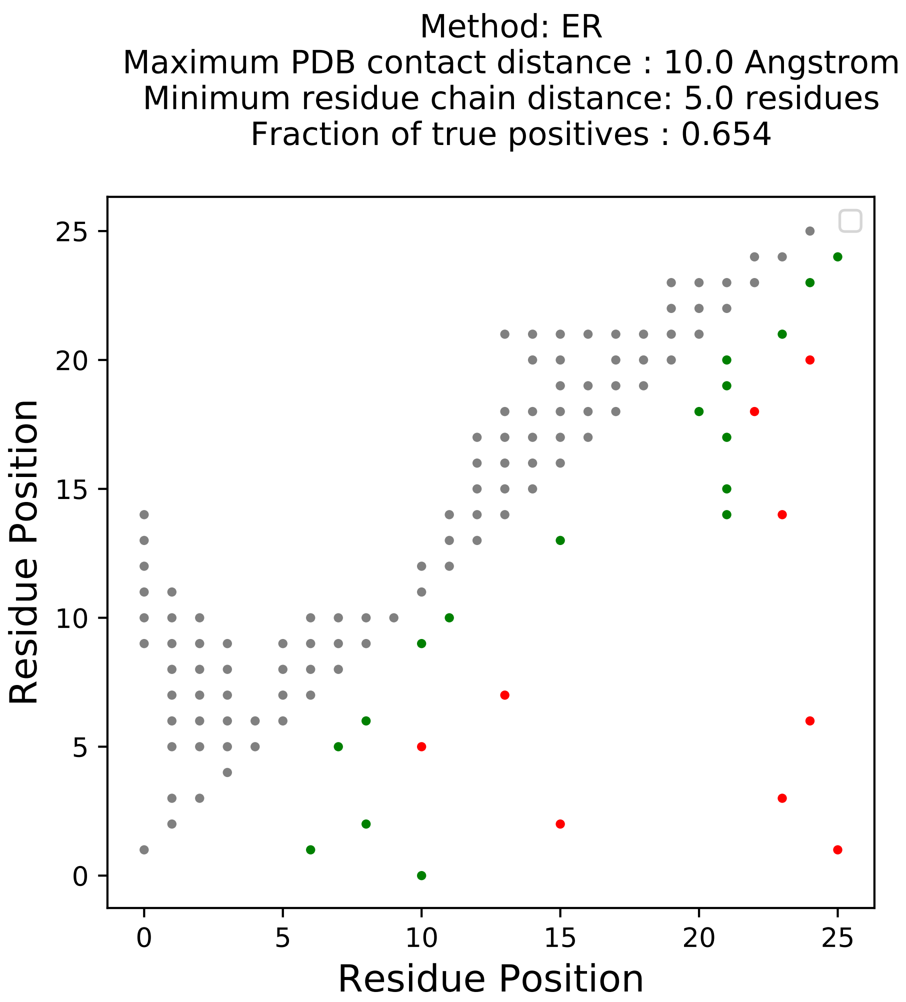

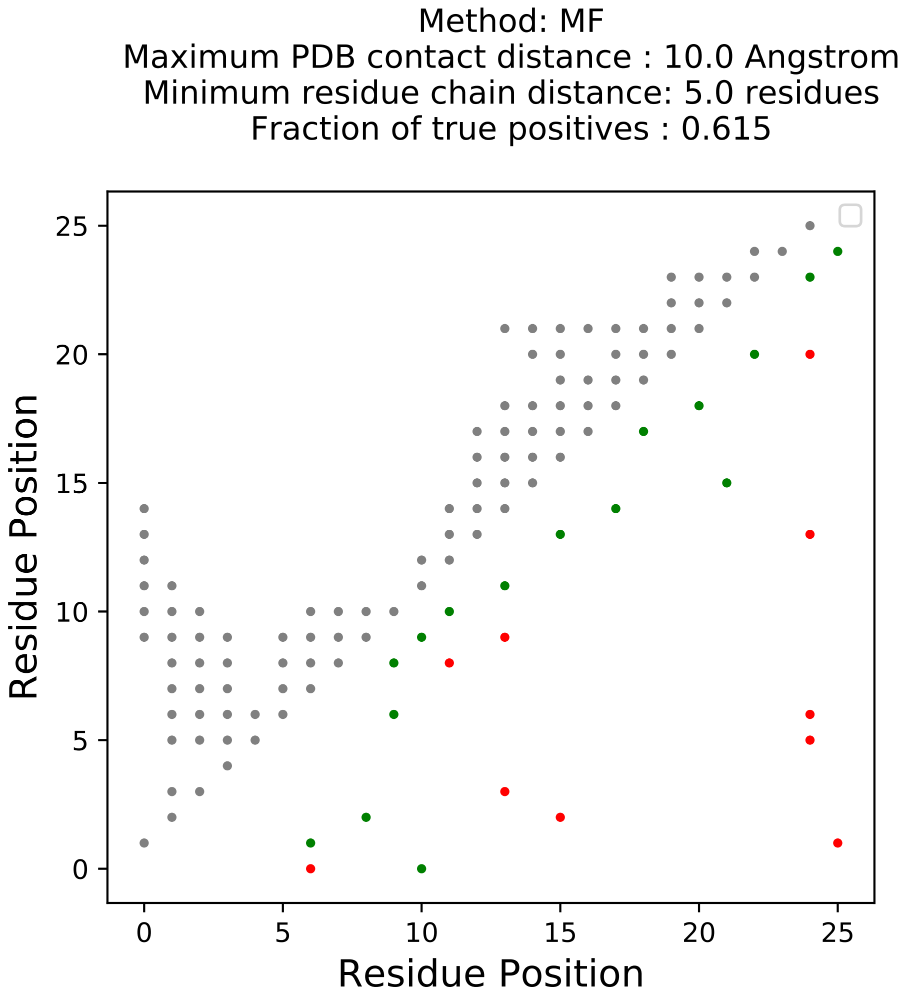

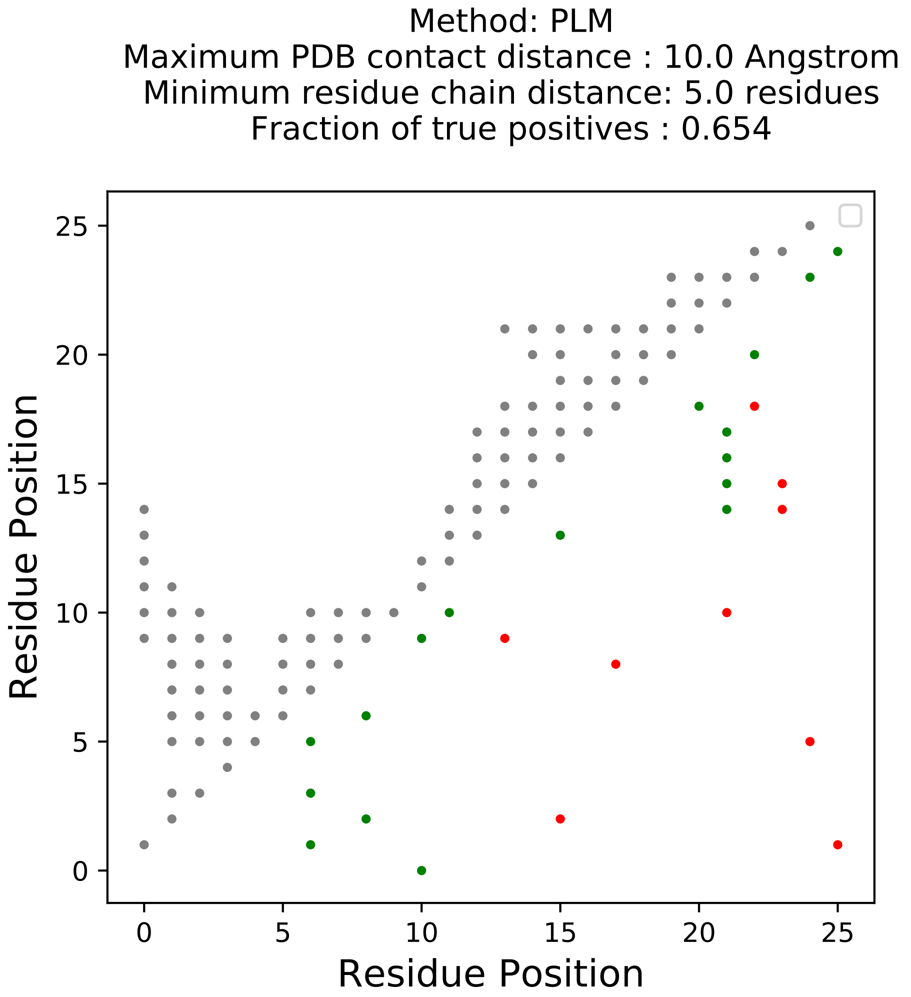

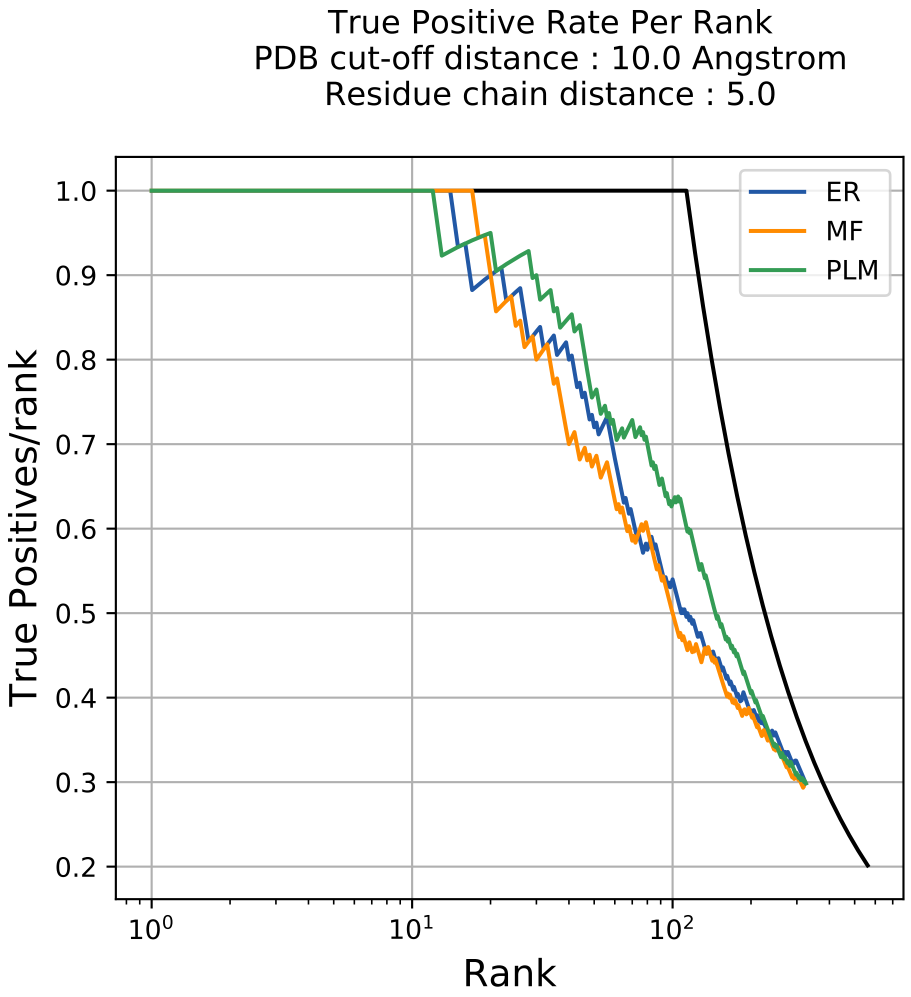

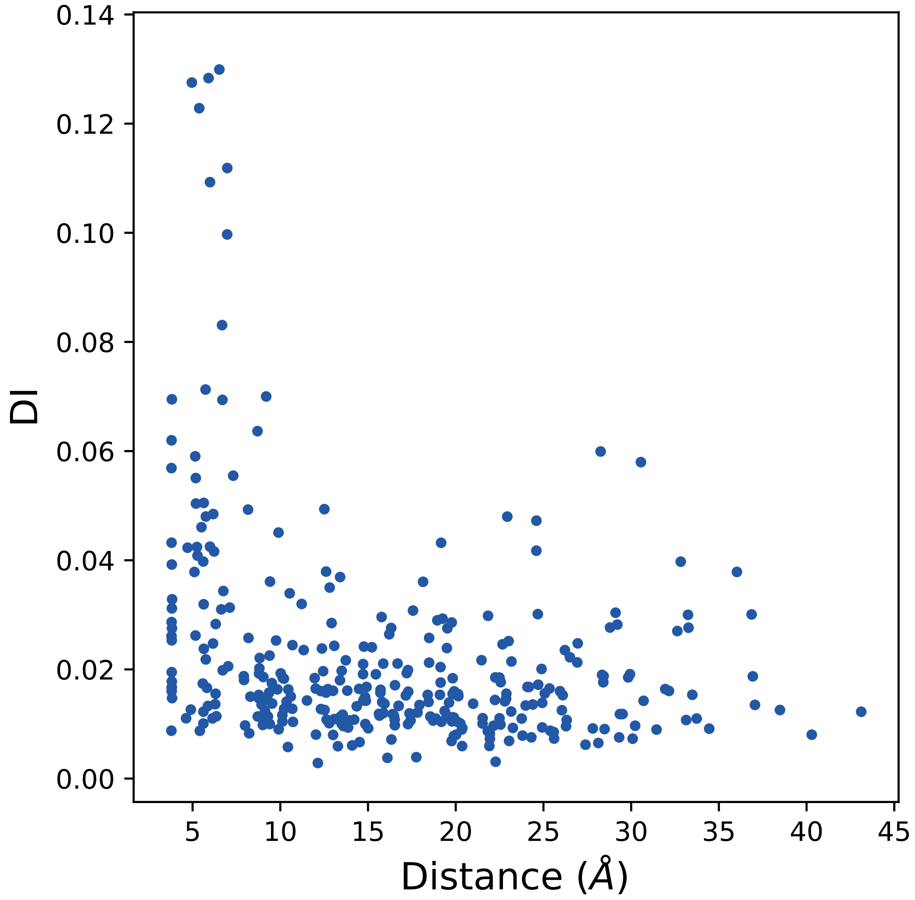

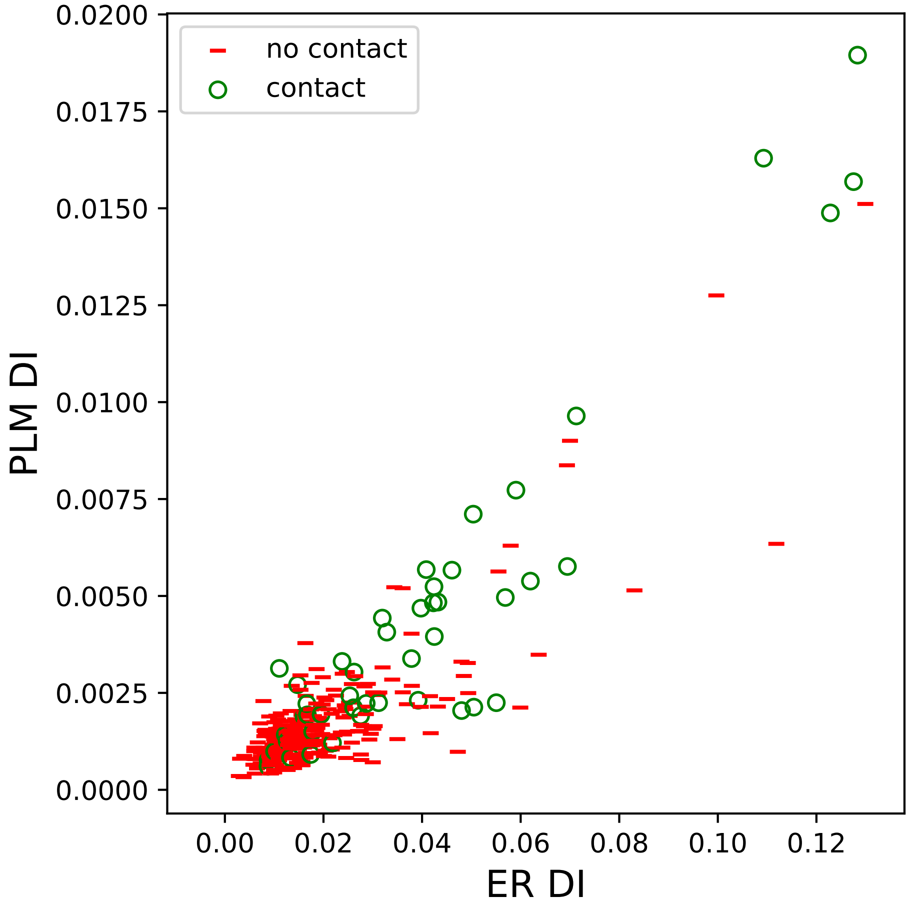

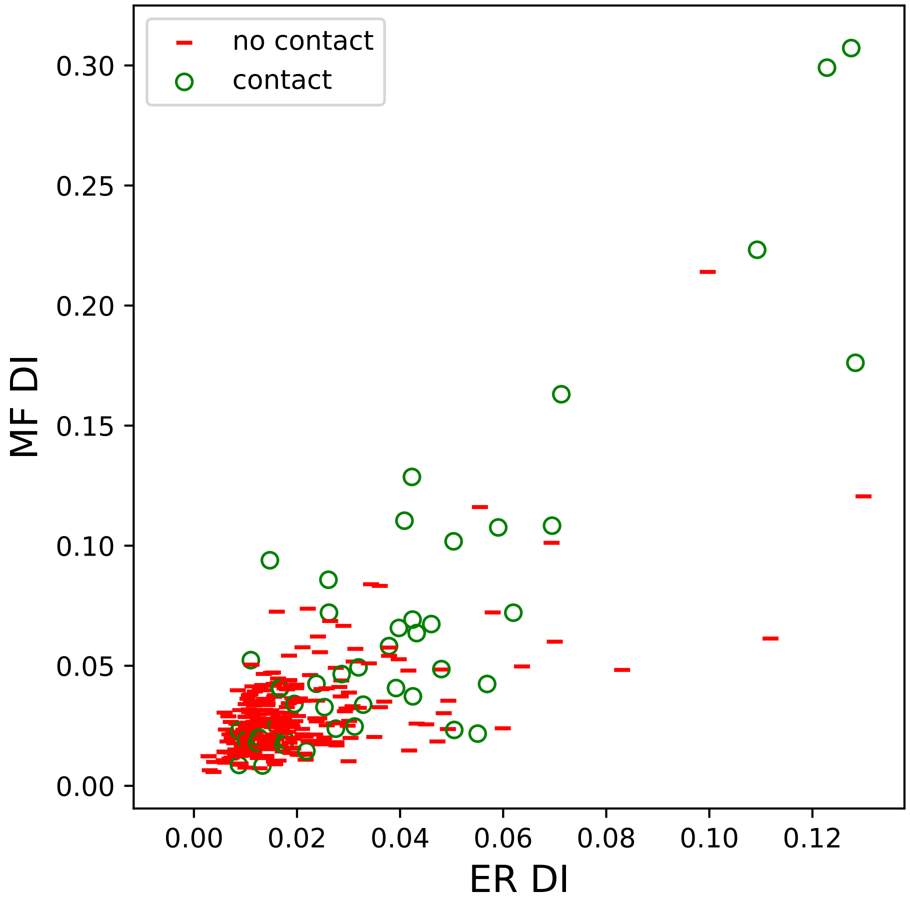

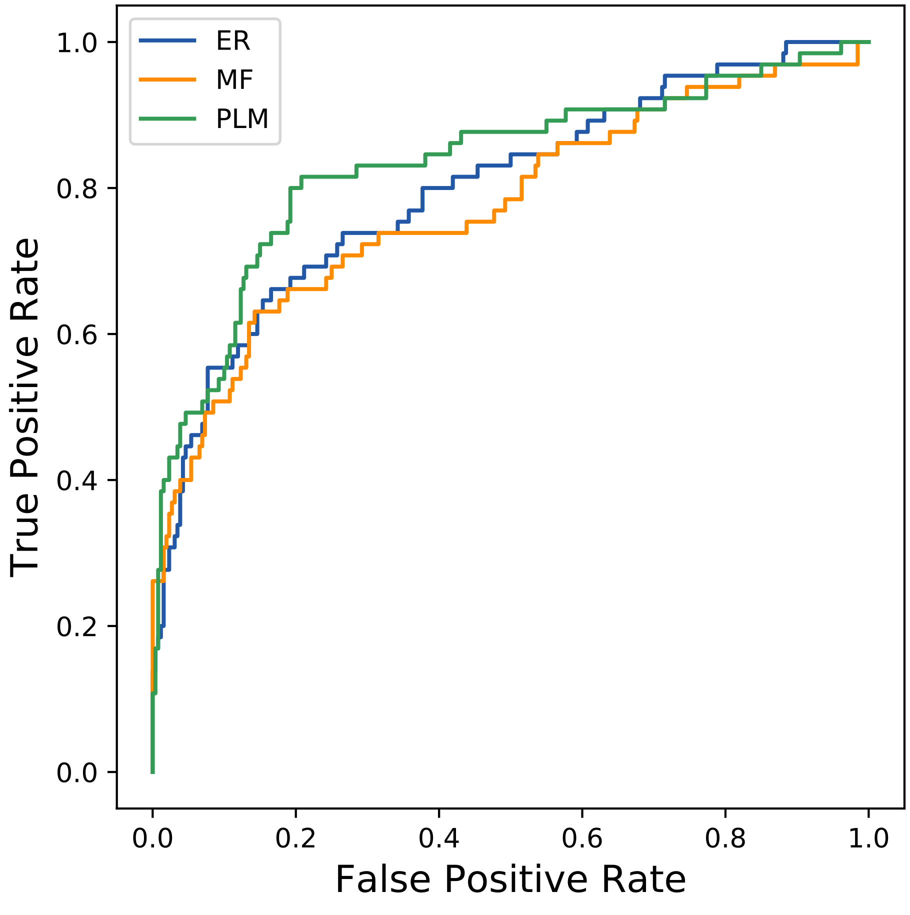

### 2255seq_184col_1ivj_PF01126_plots.pdf

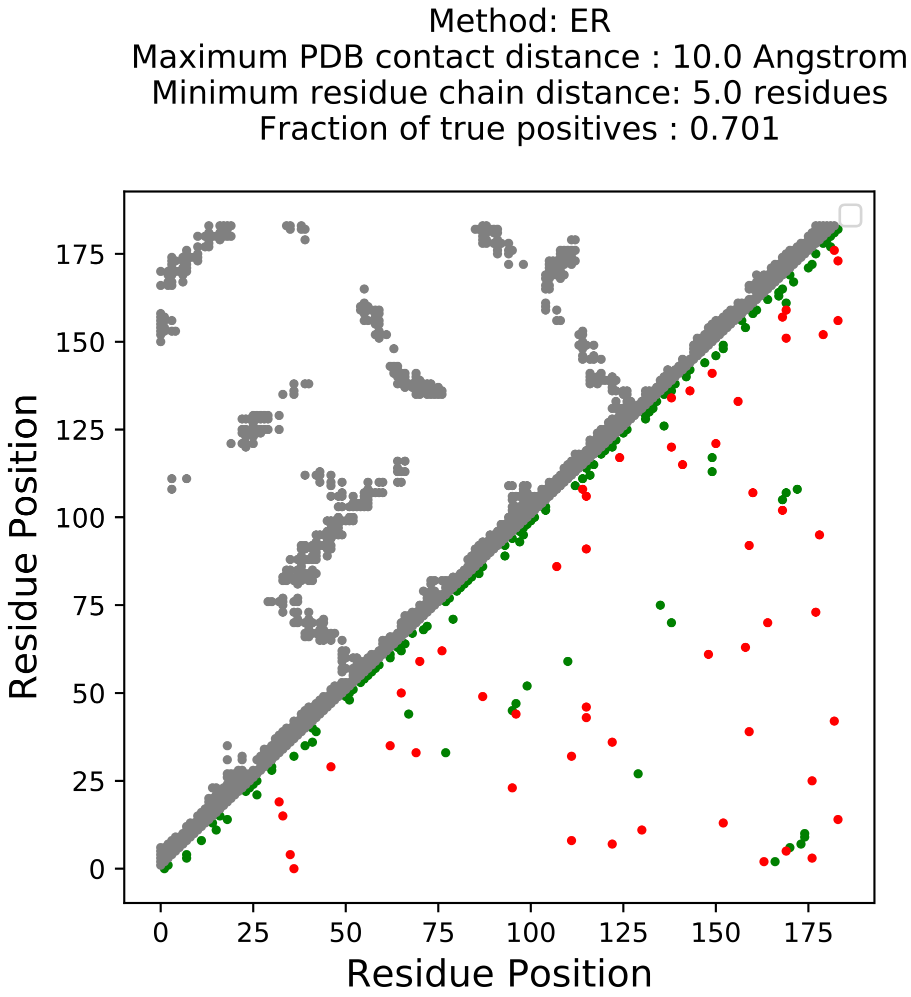

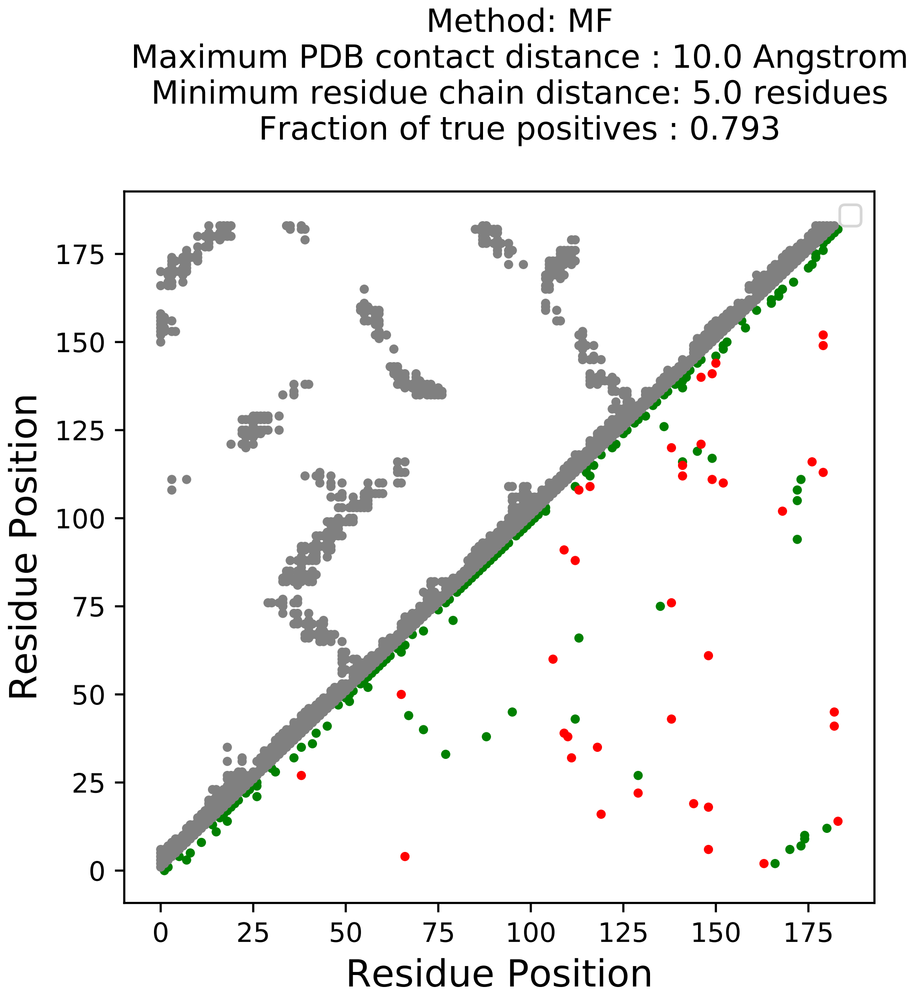

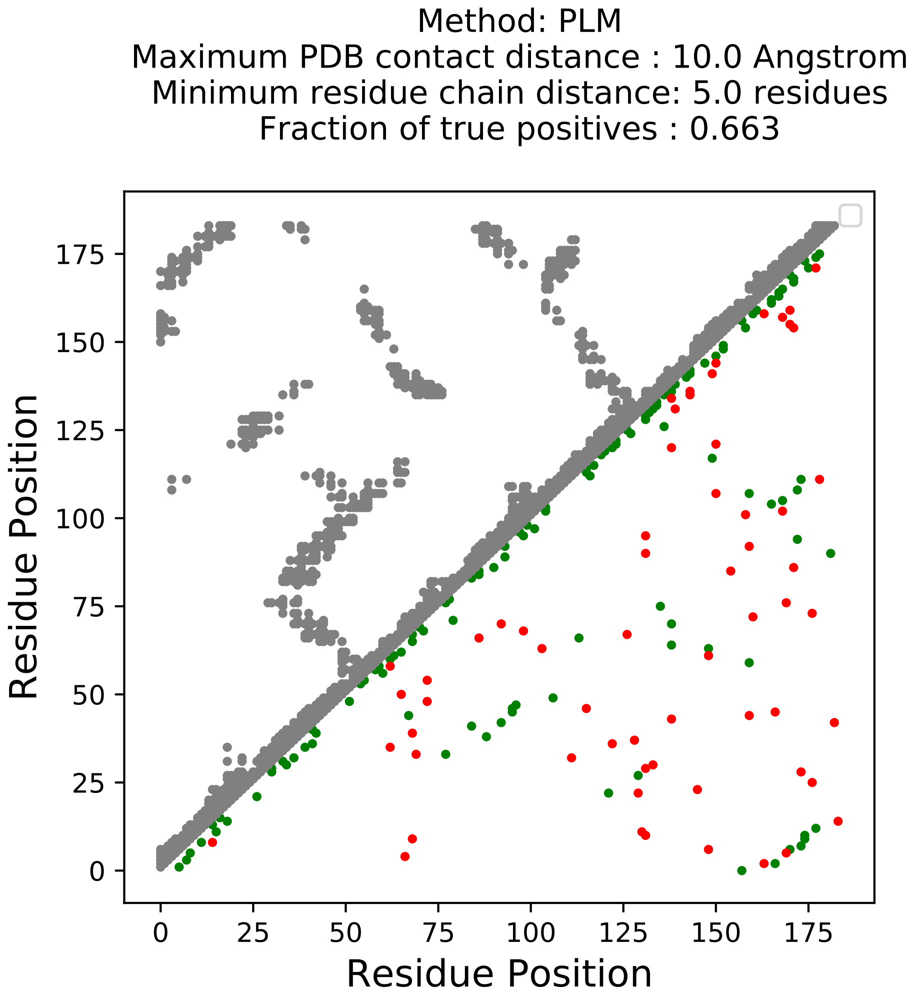

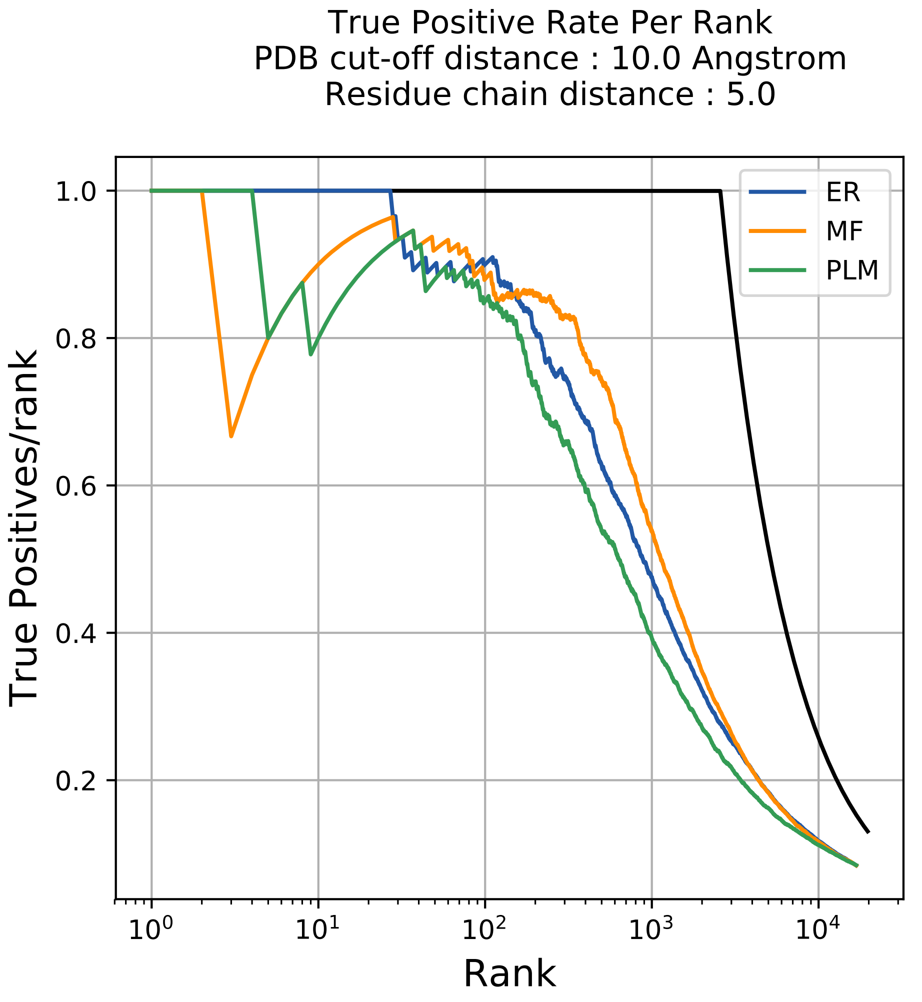

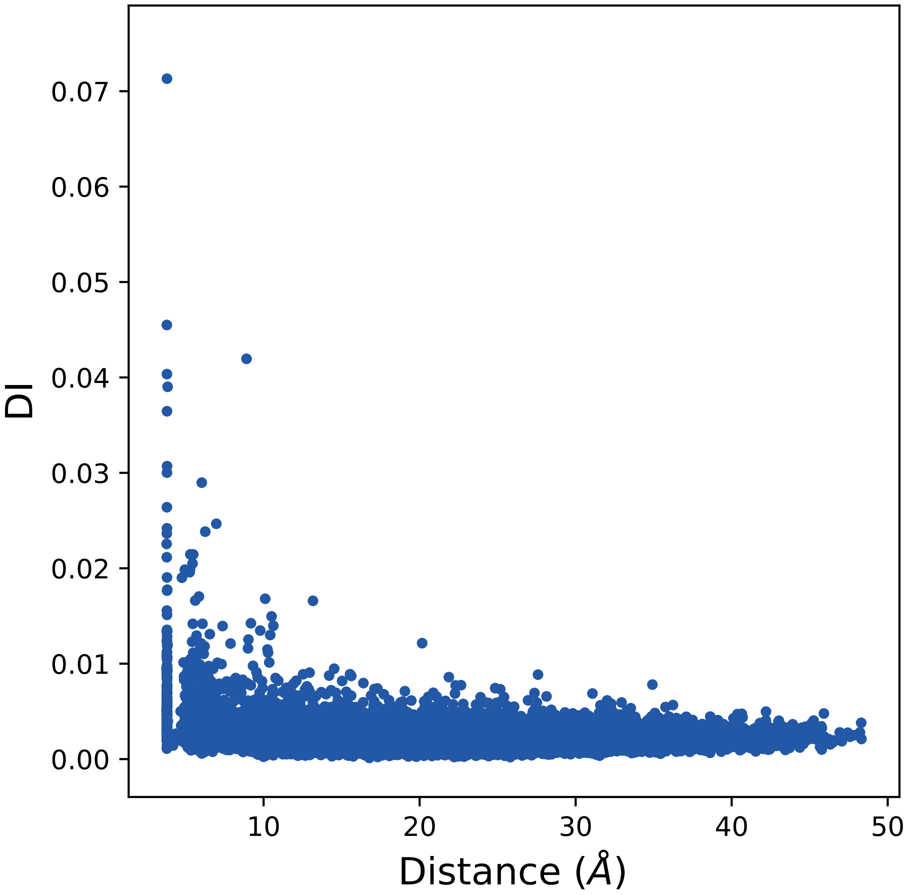

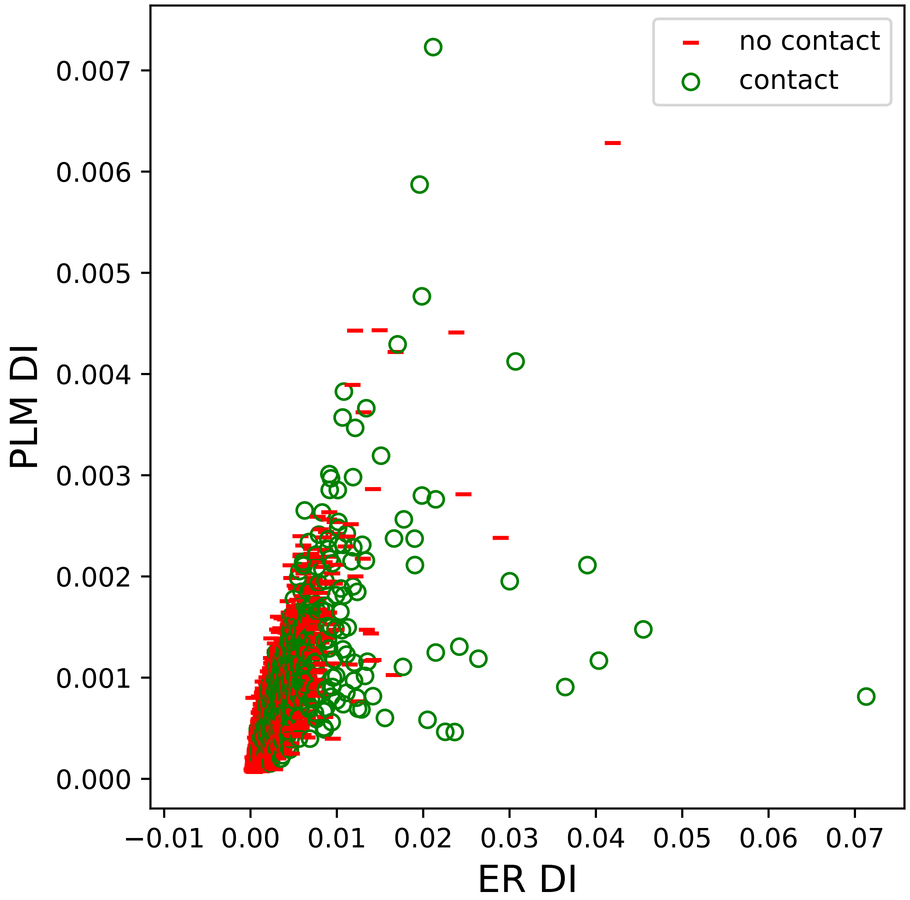

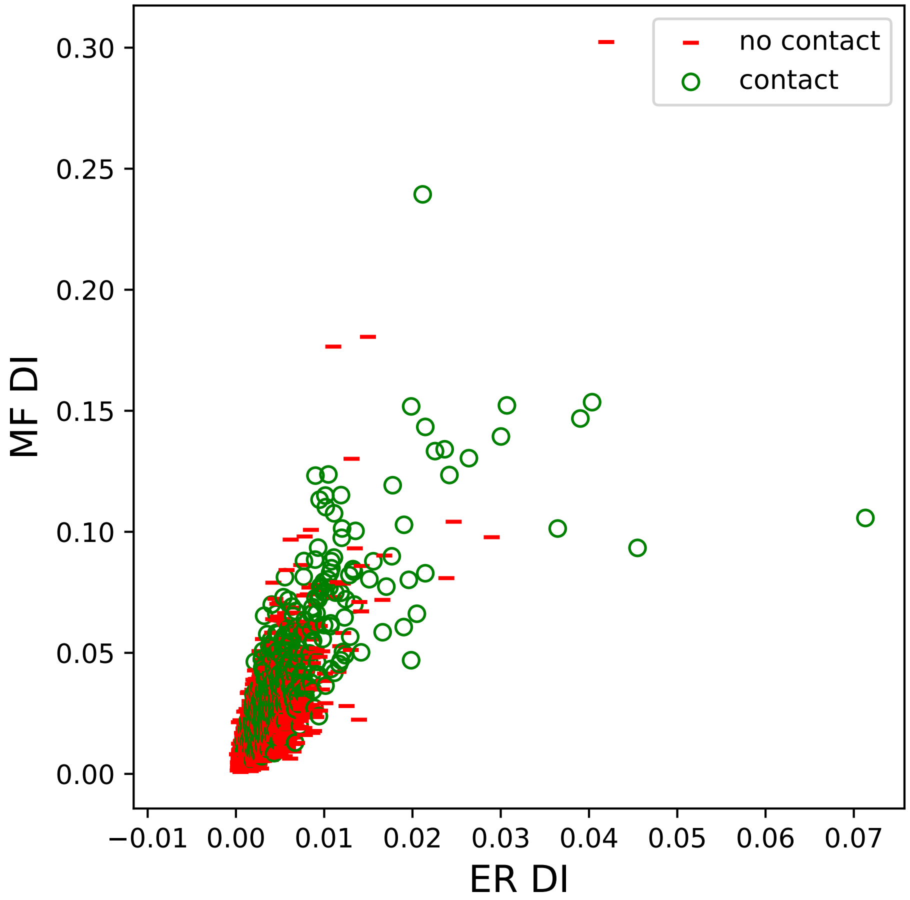

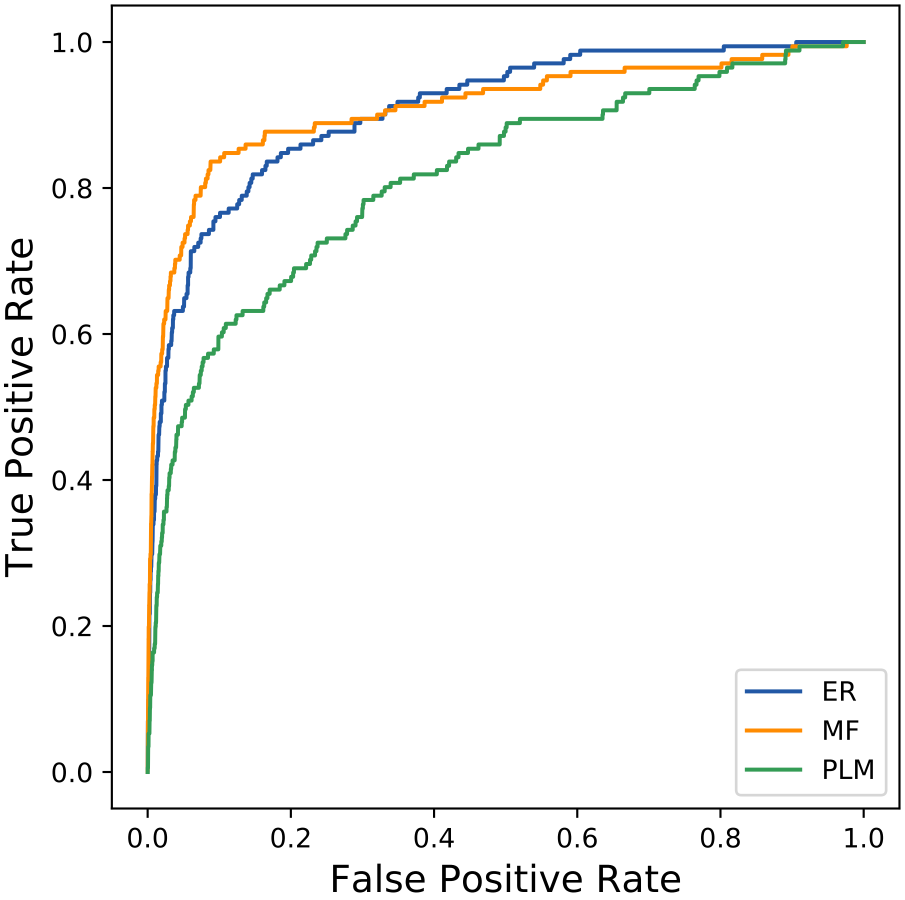

### 2334seq_60col_2cr9_PF05406_plots.pdf

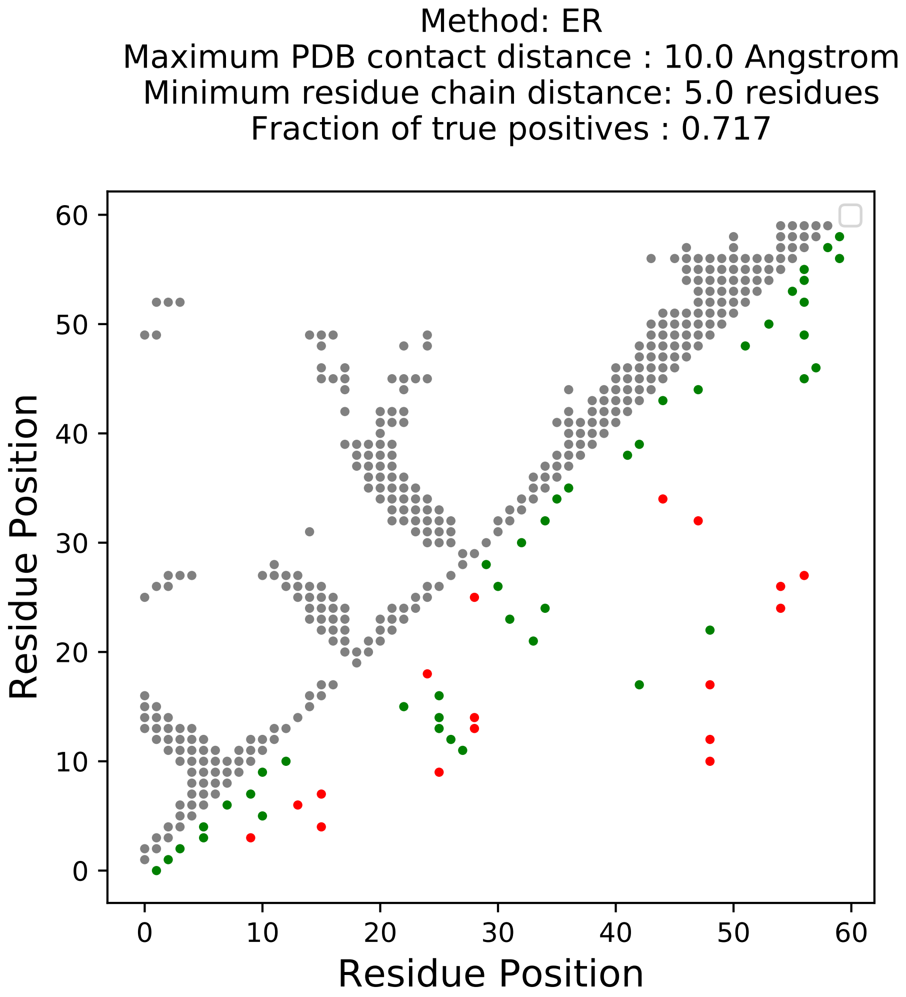

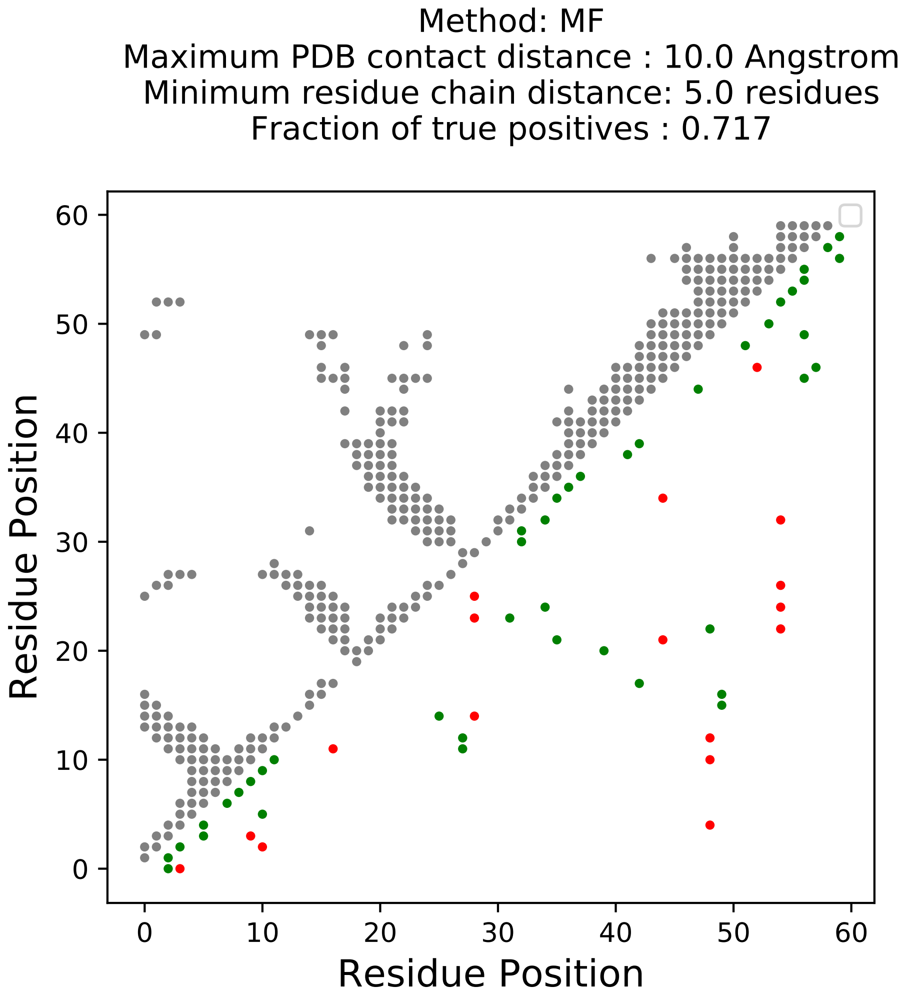

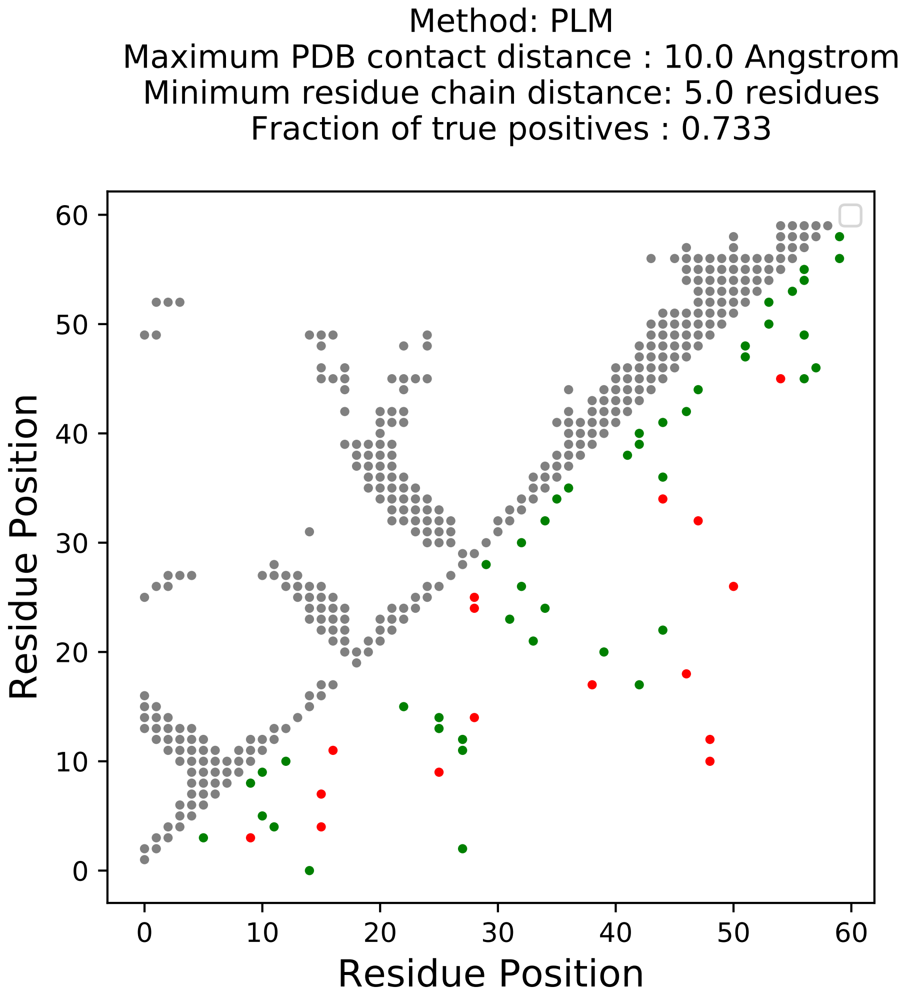

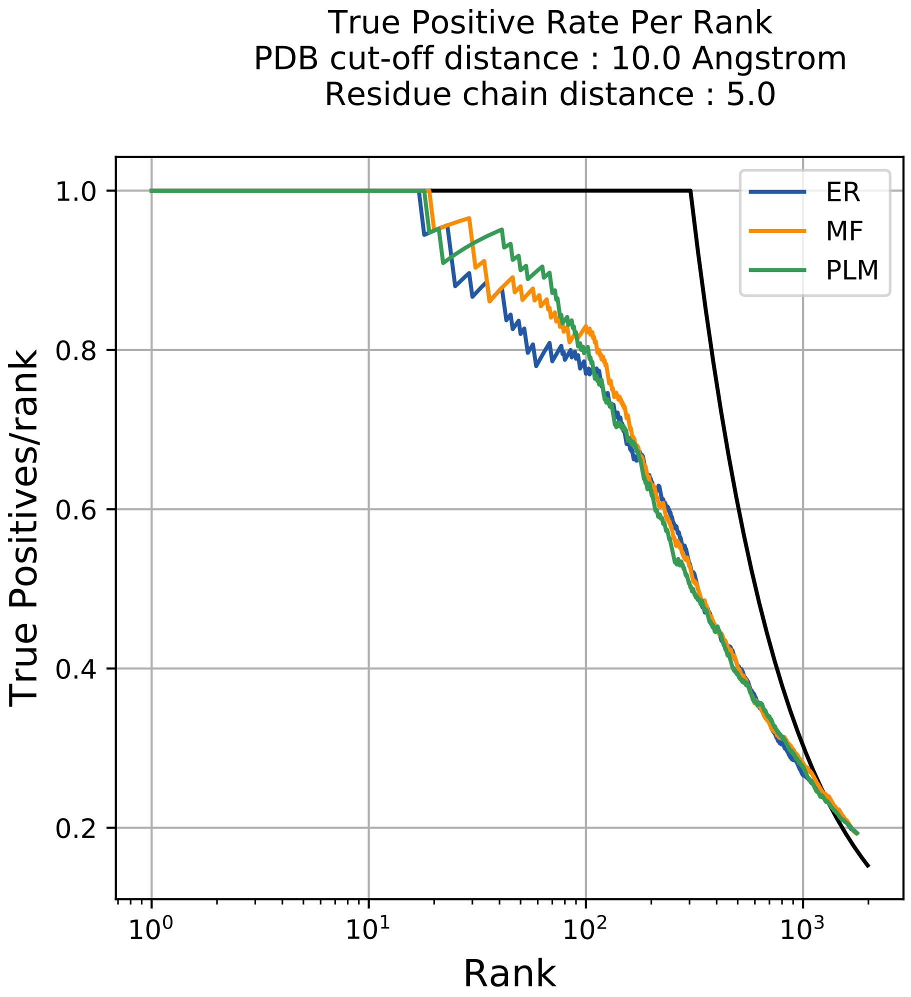

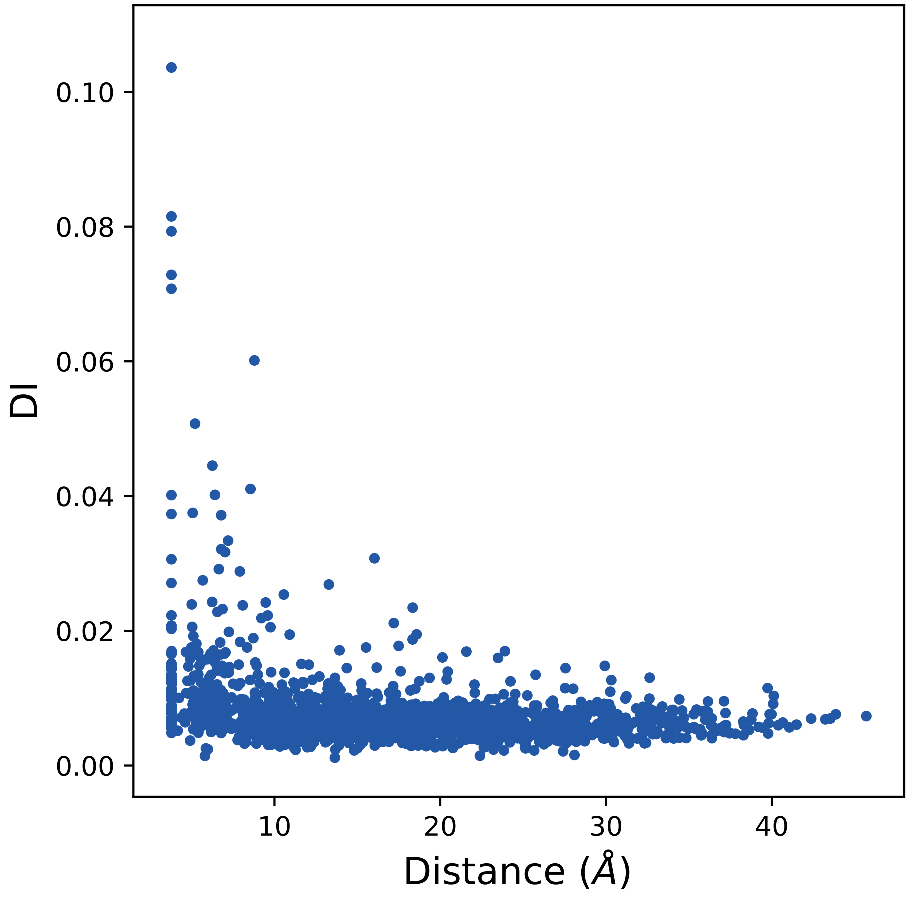

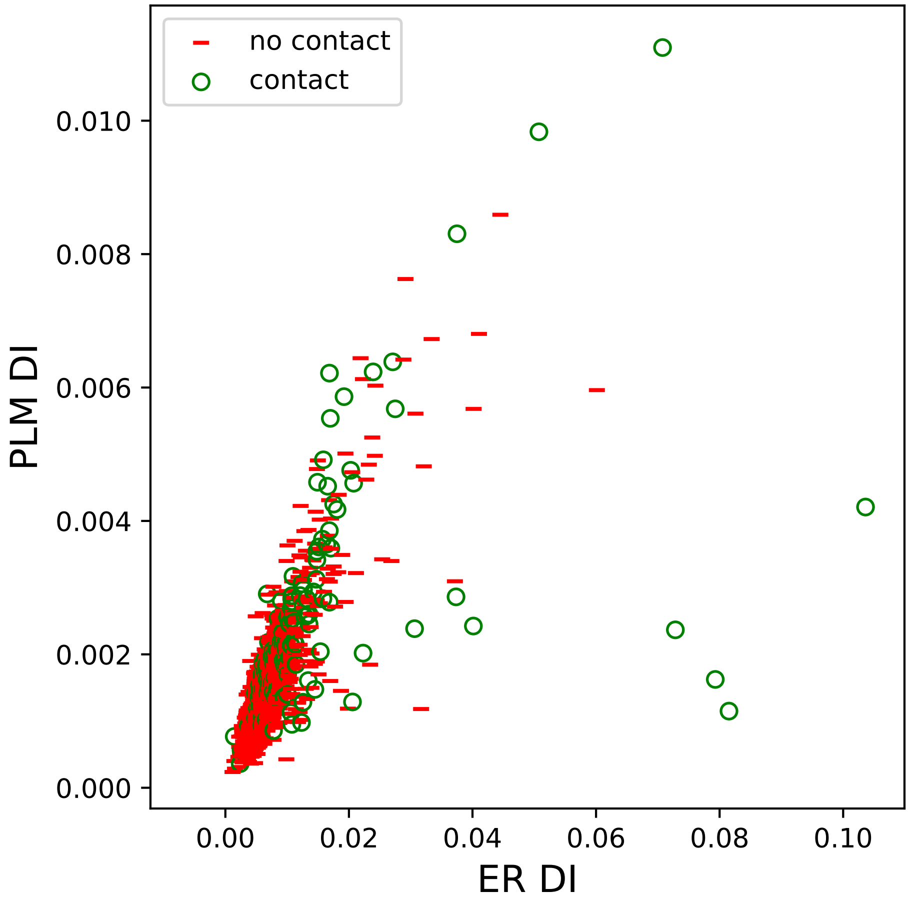

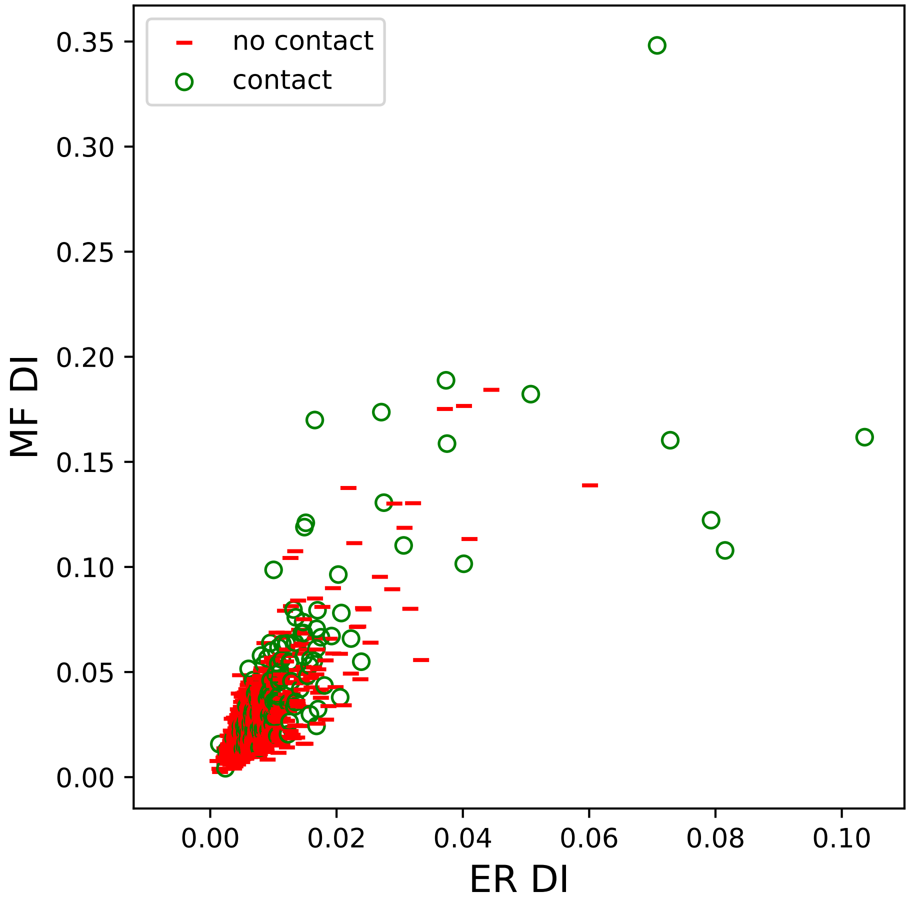

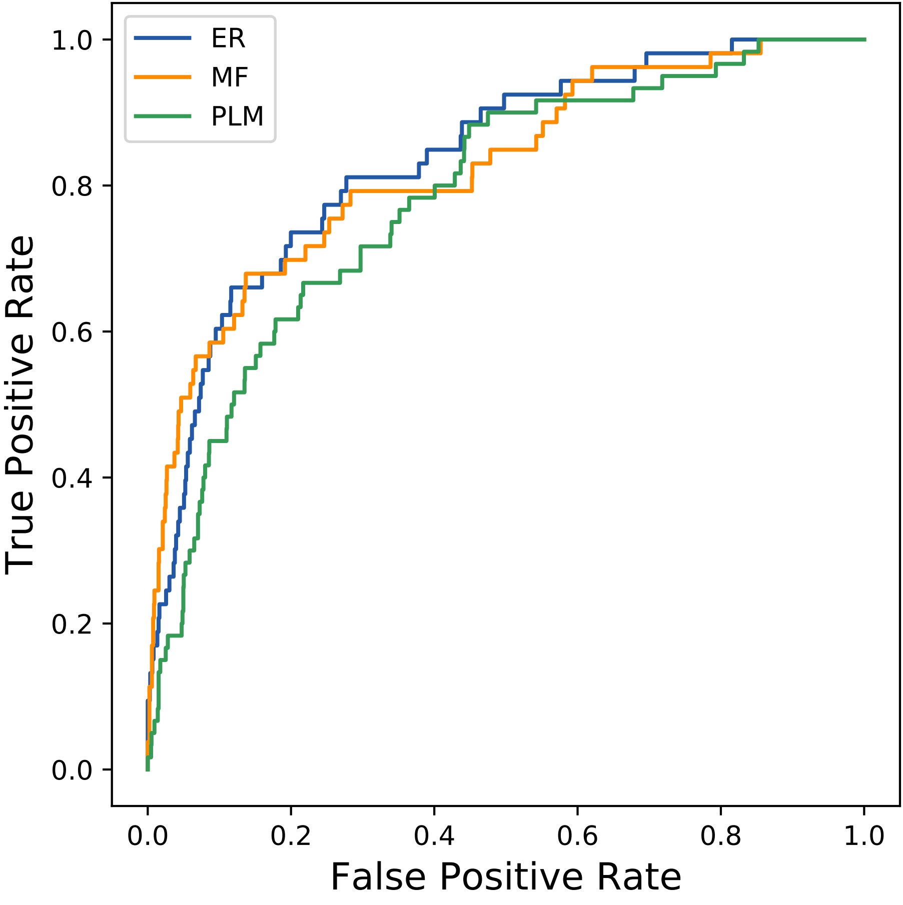

### 2645seq_185col_5jm8_PF04183_plots.pdf

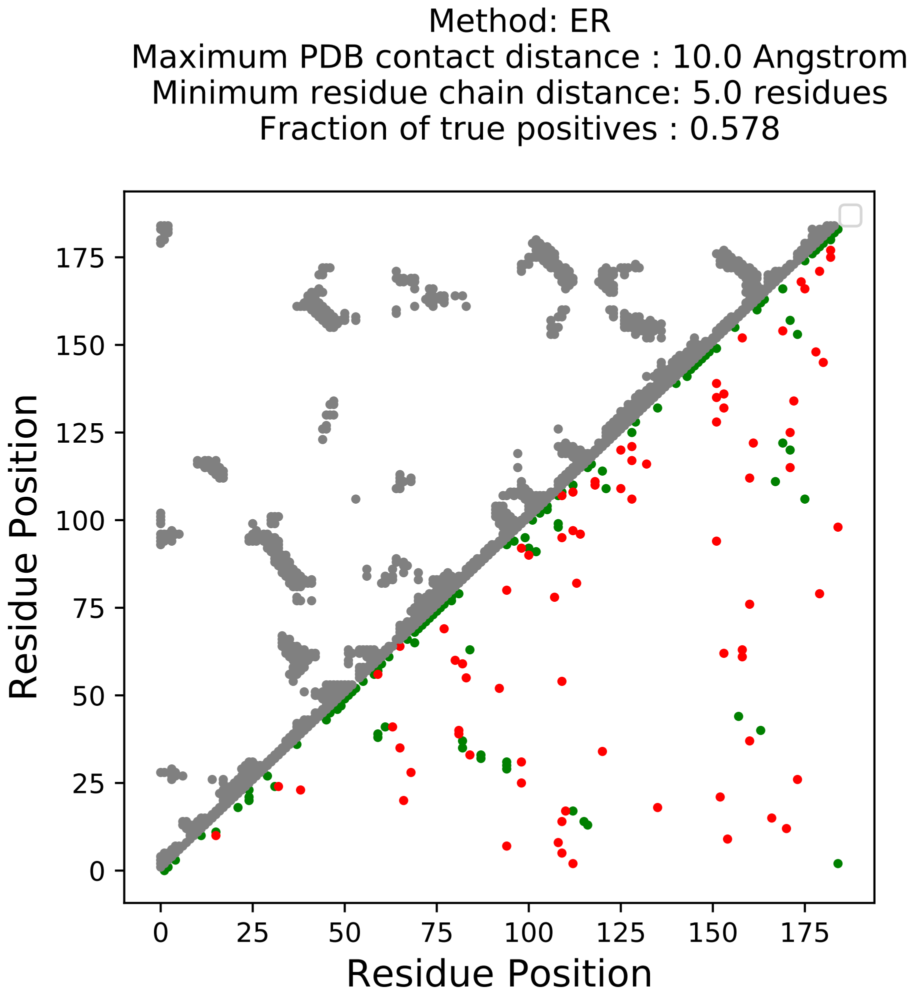

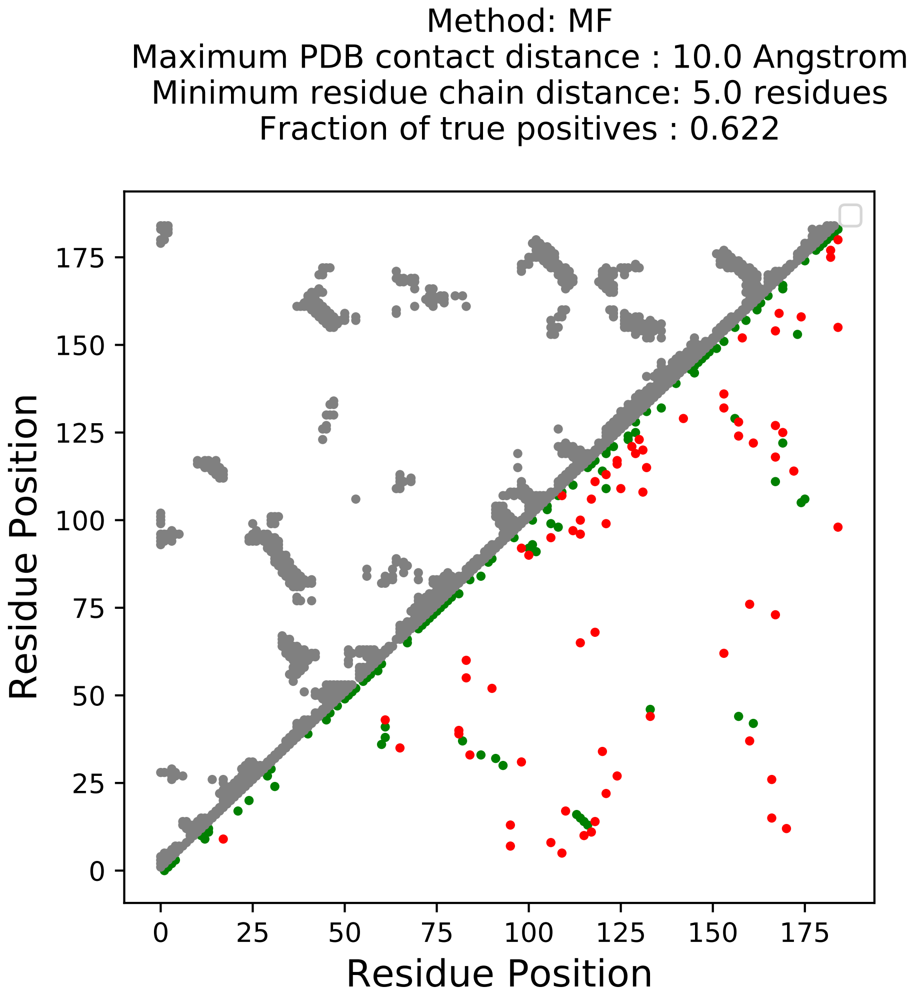

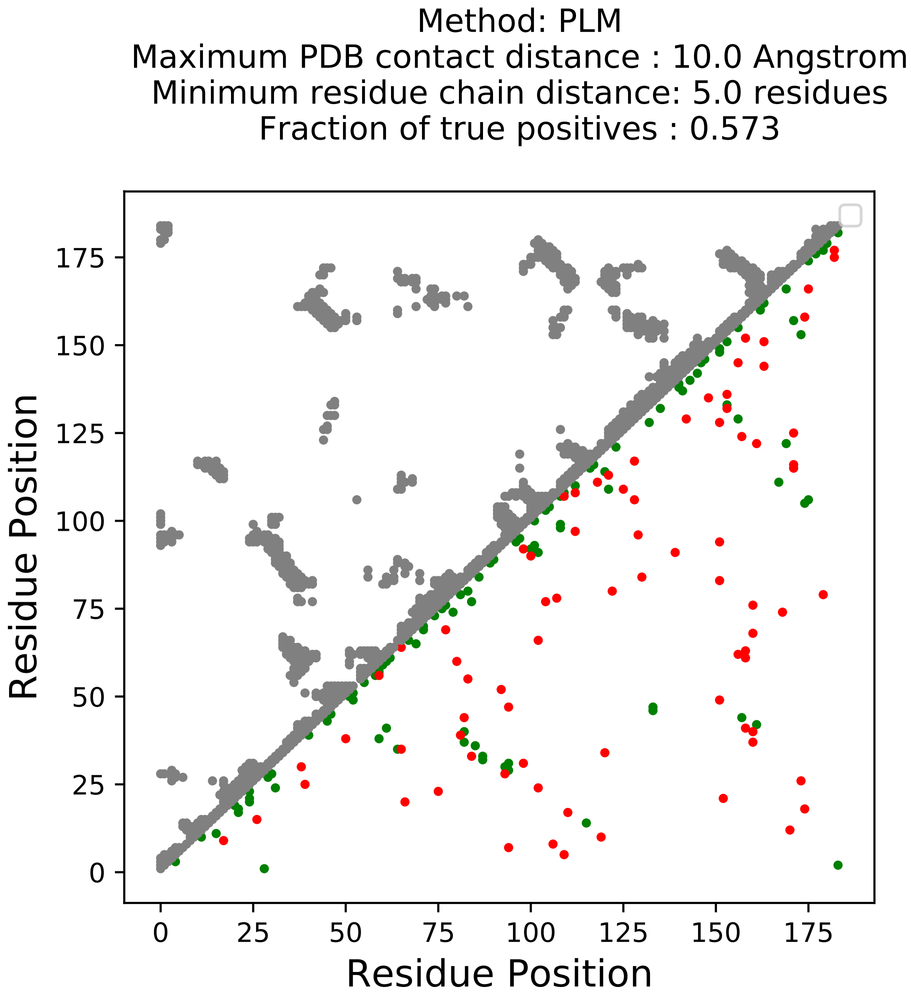

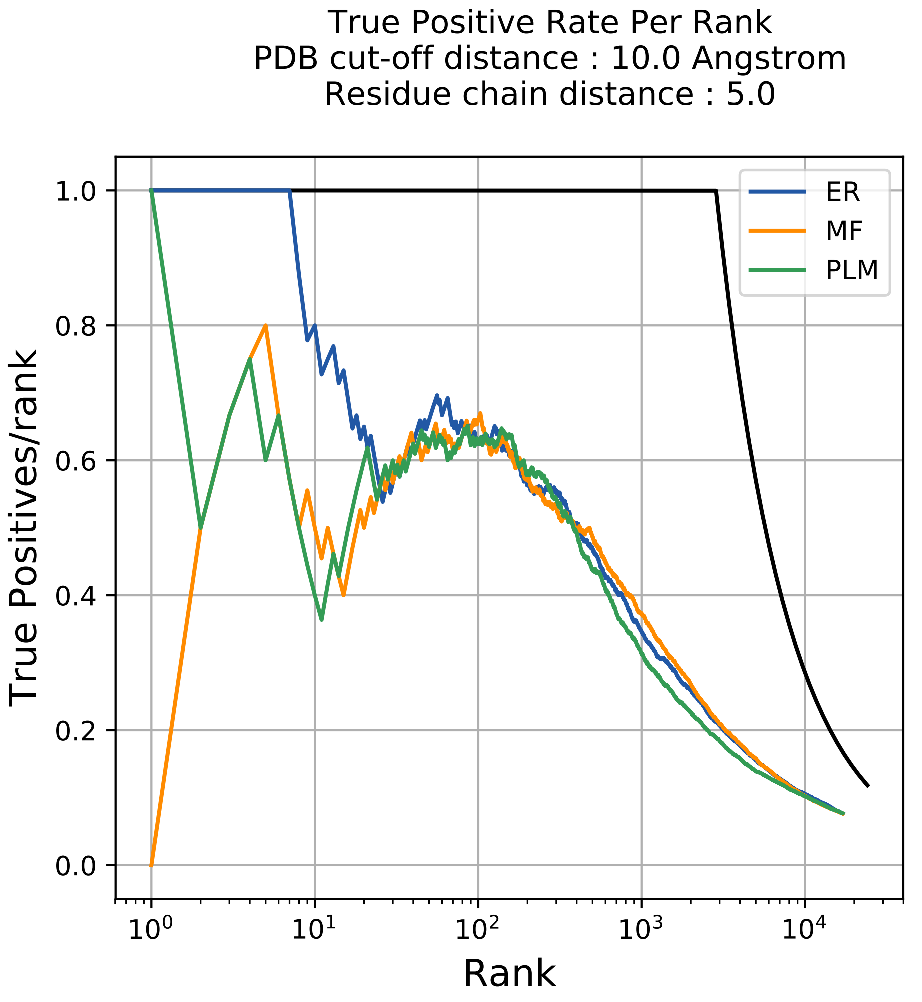

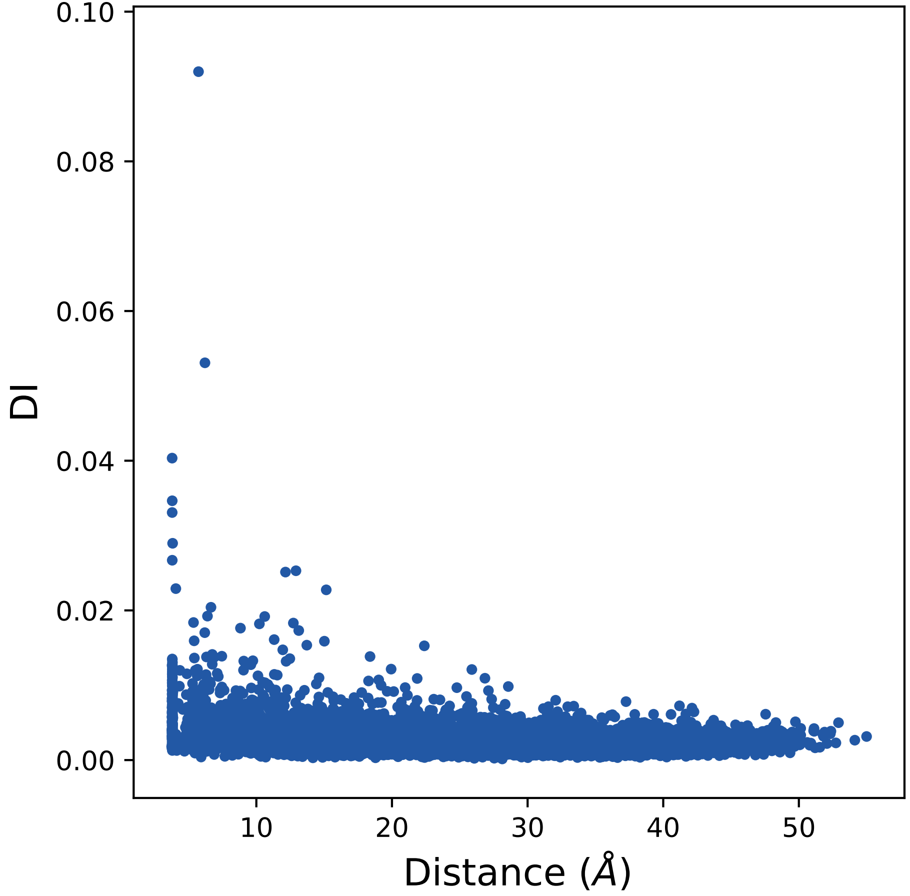

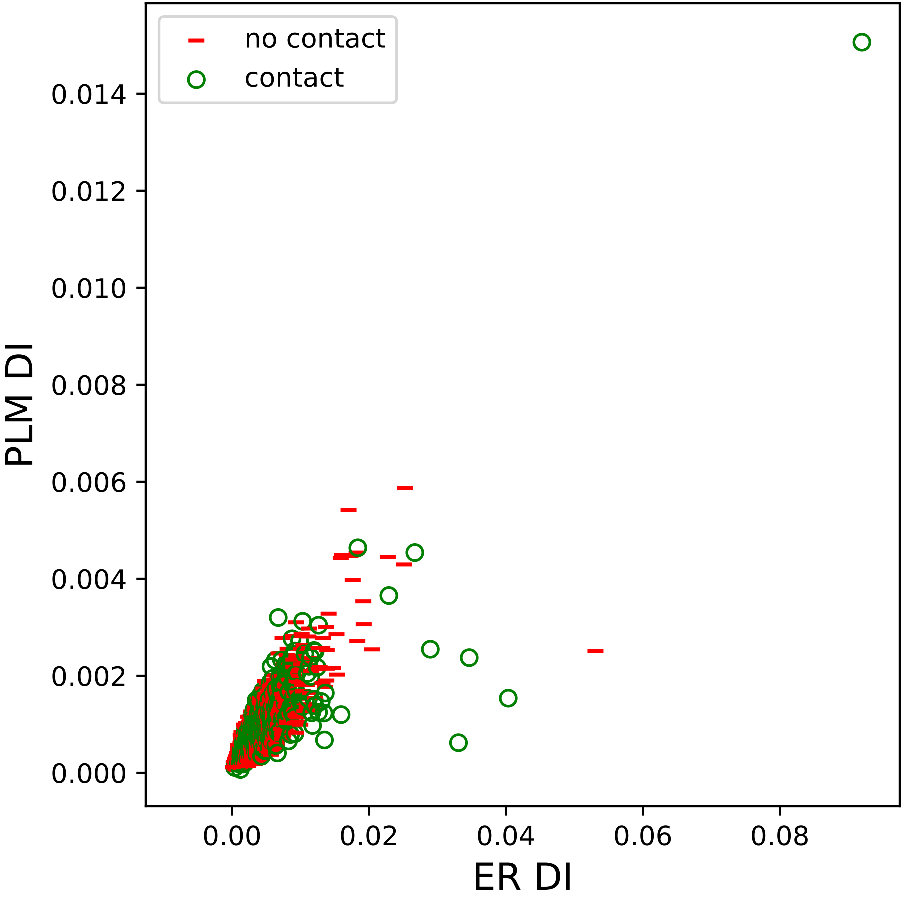

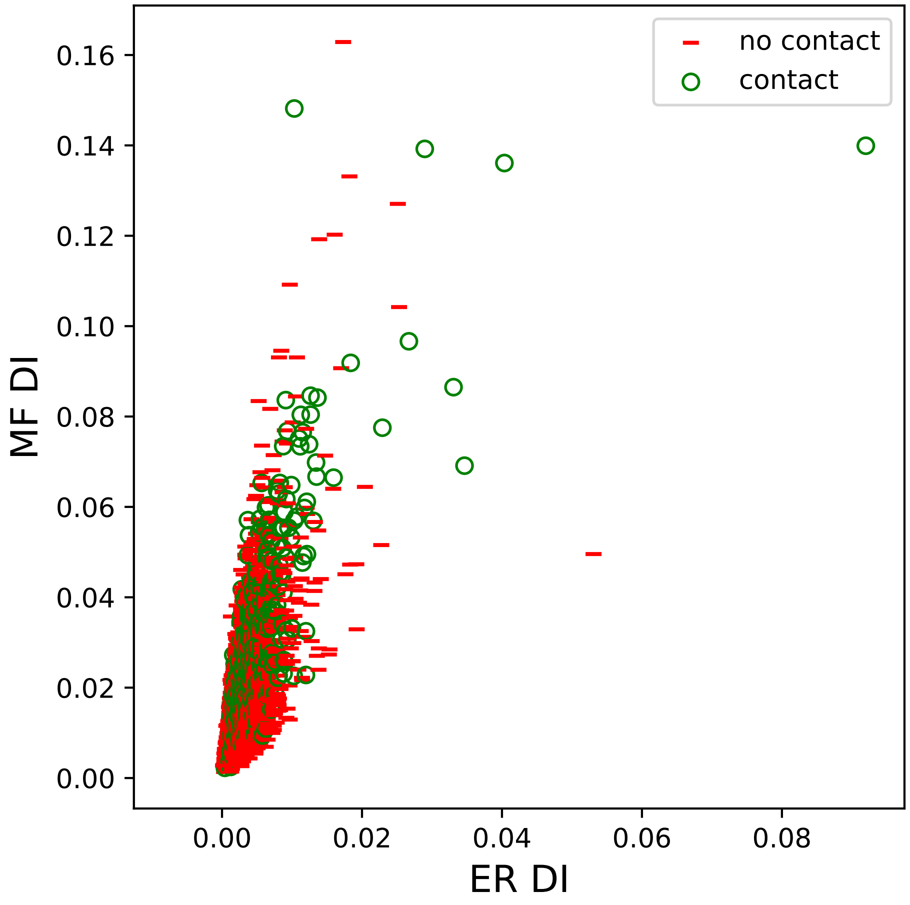

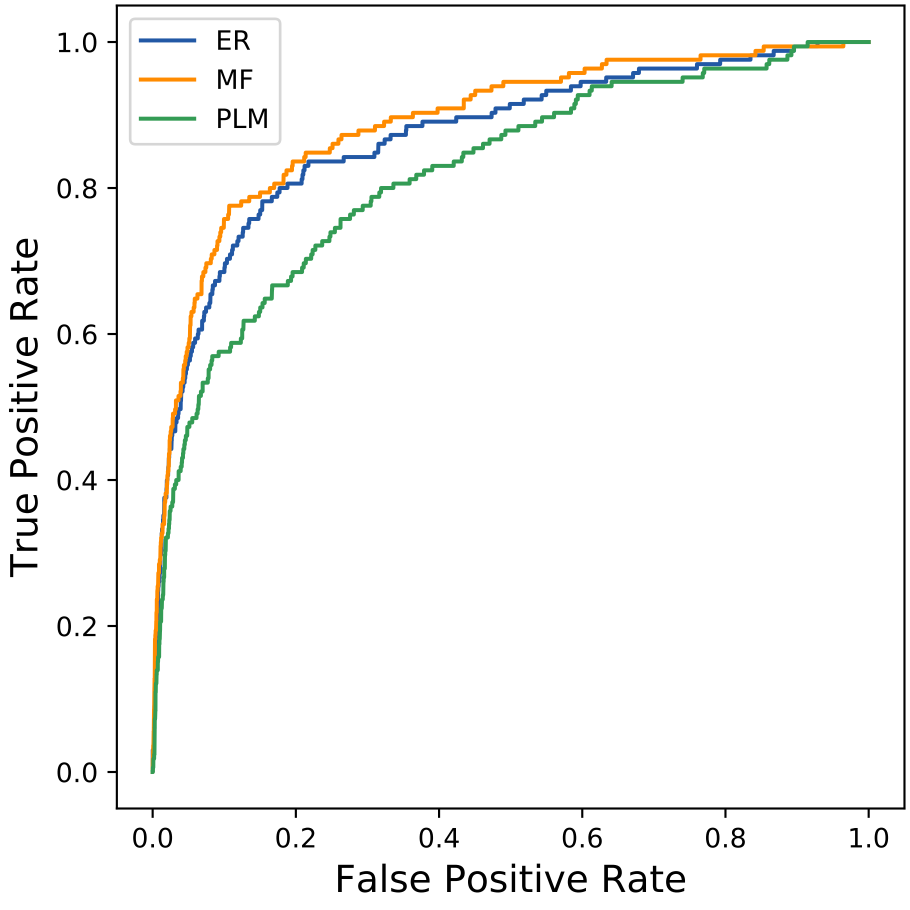

### 2861seq_54col_7cmx_PF00463_plots.pdf

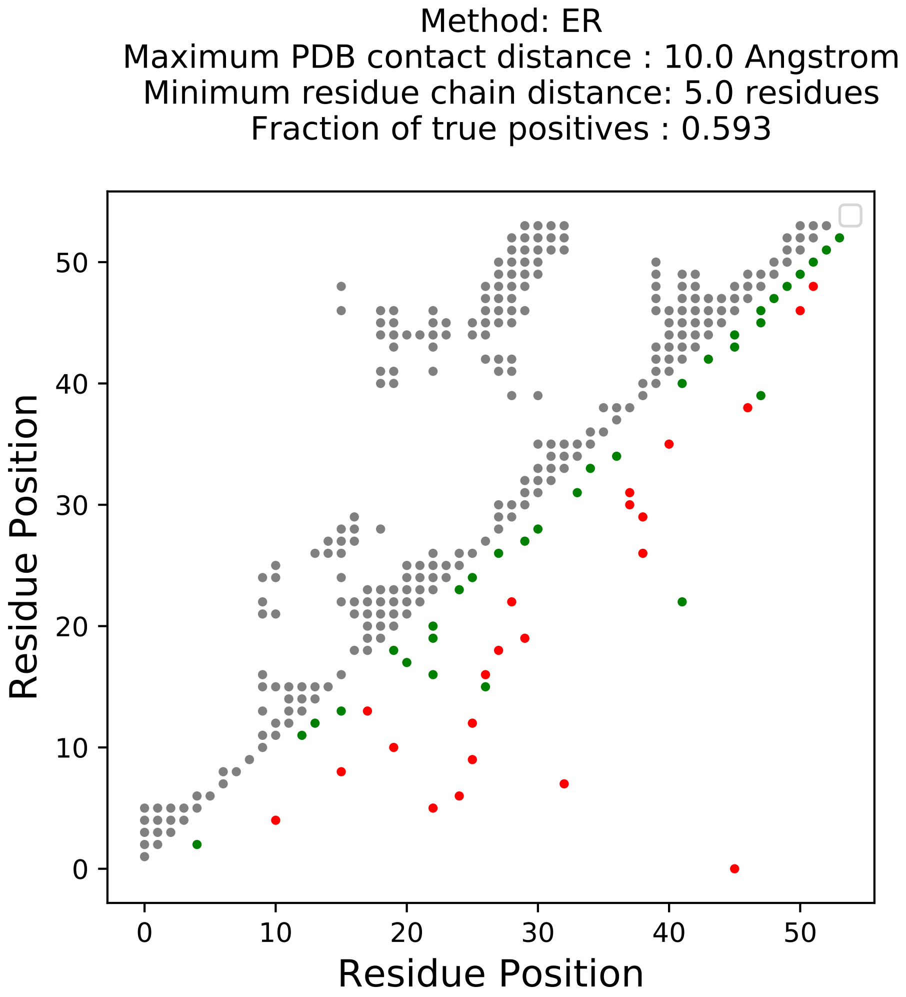

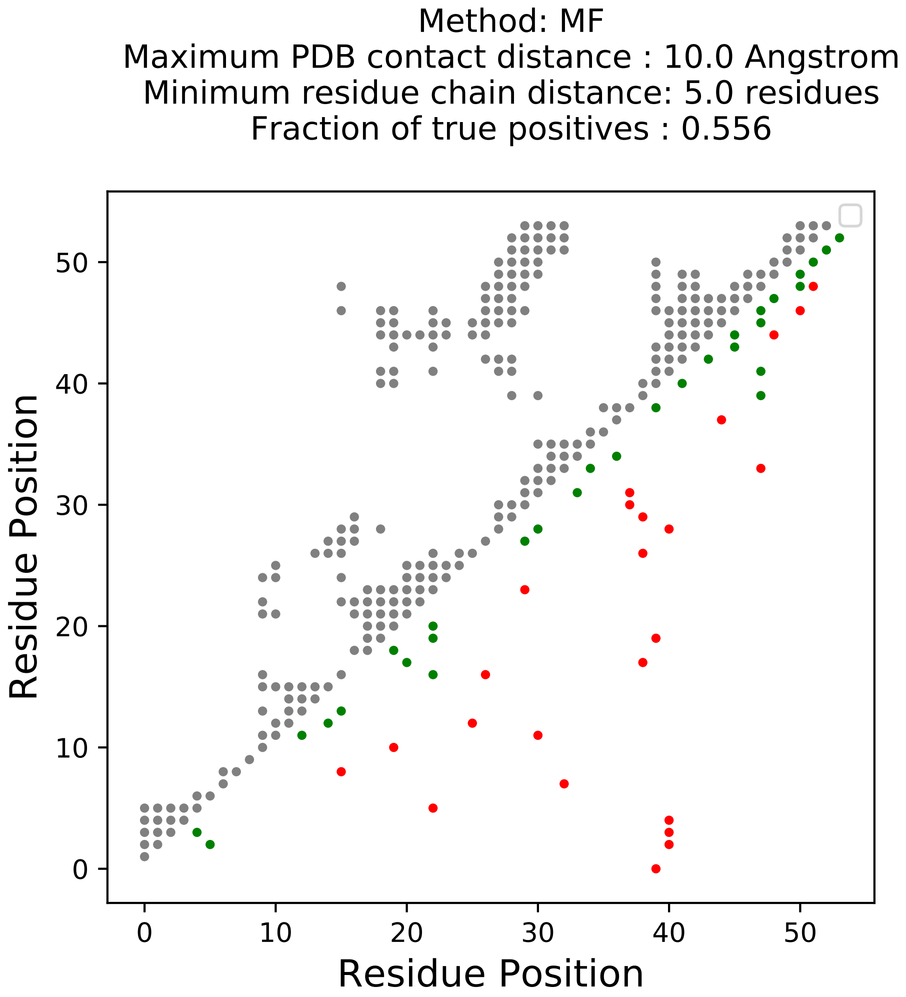

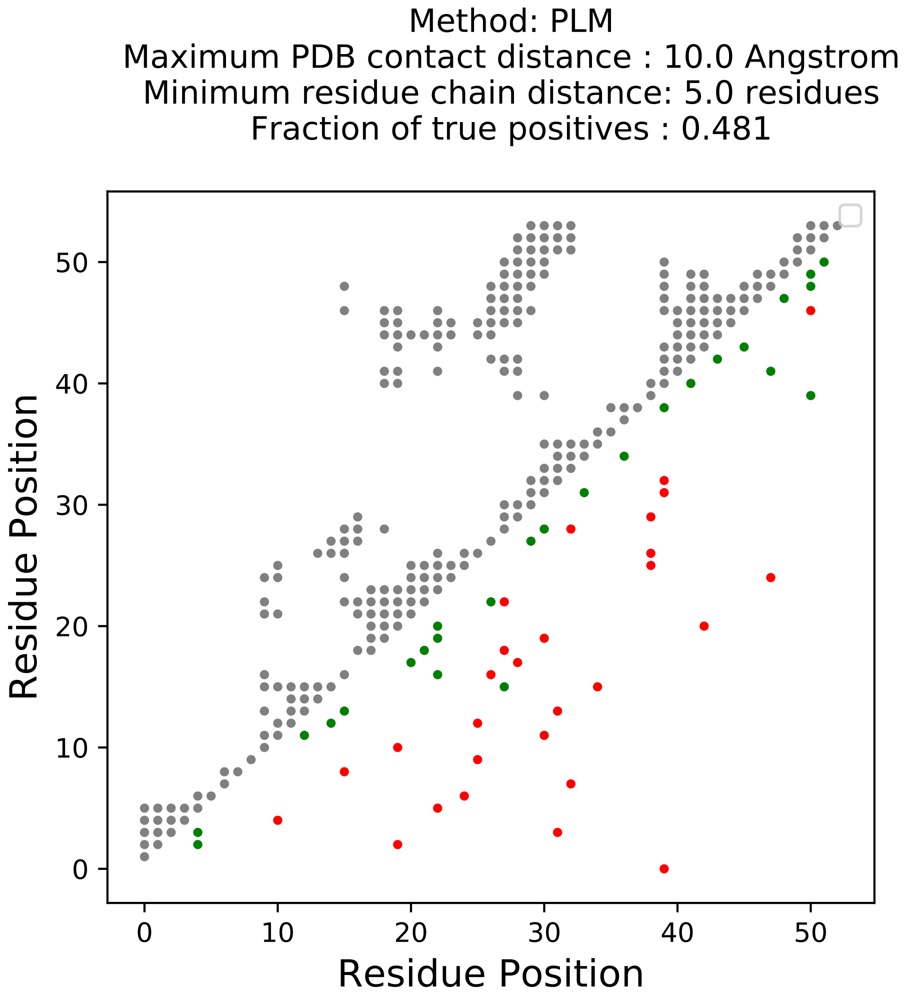

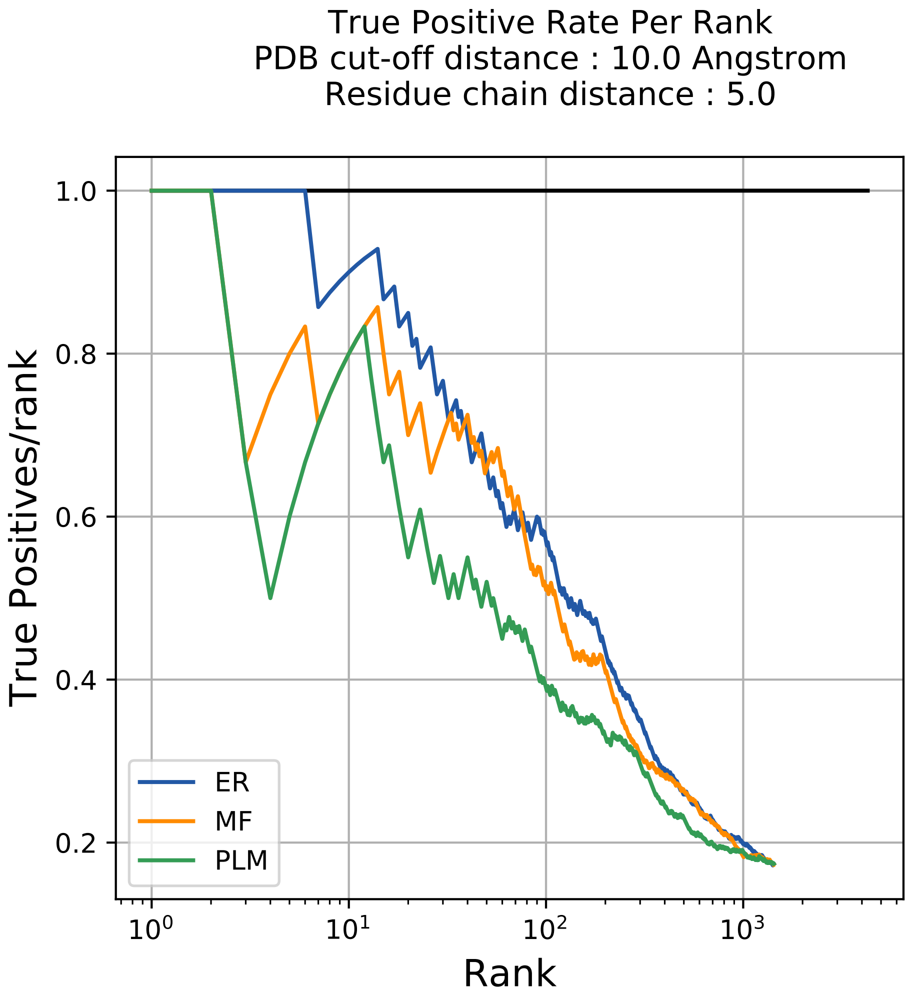

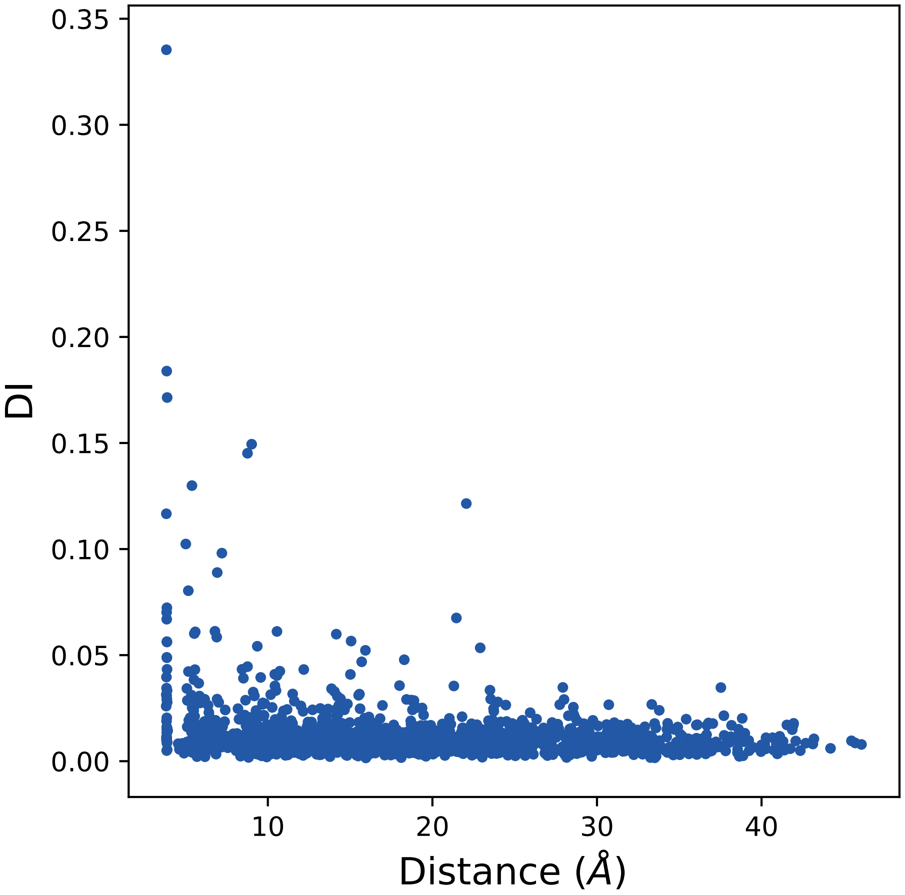

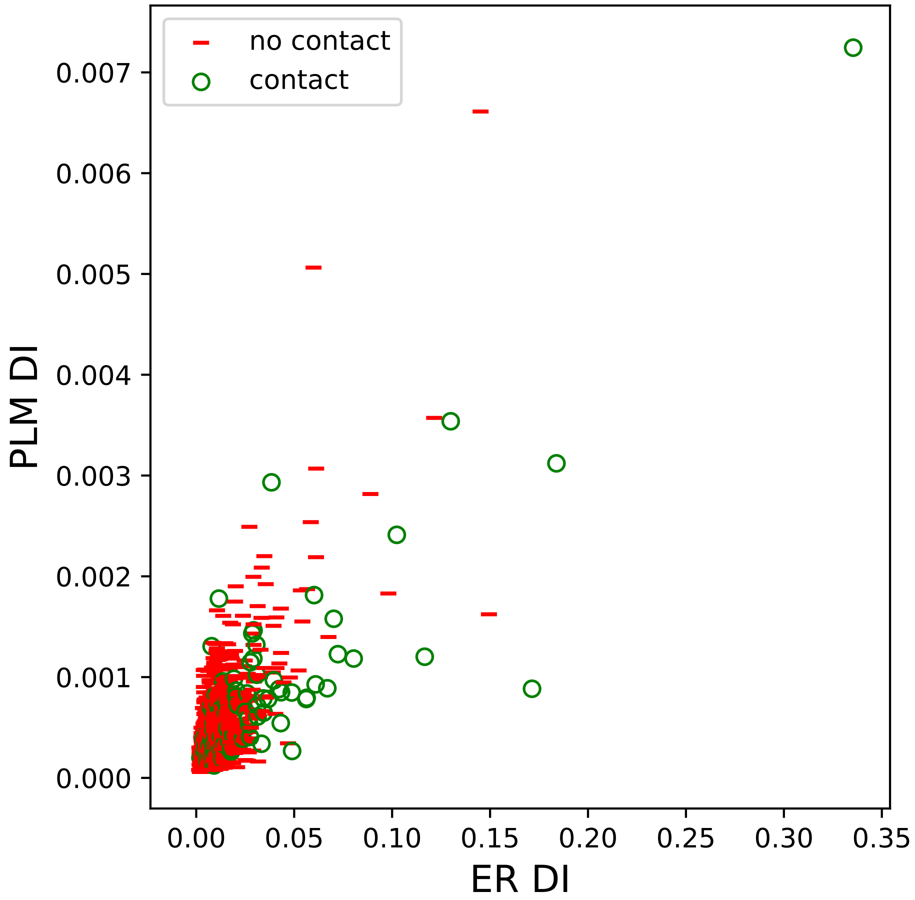

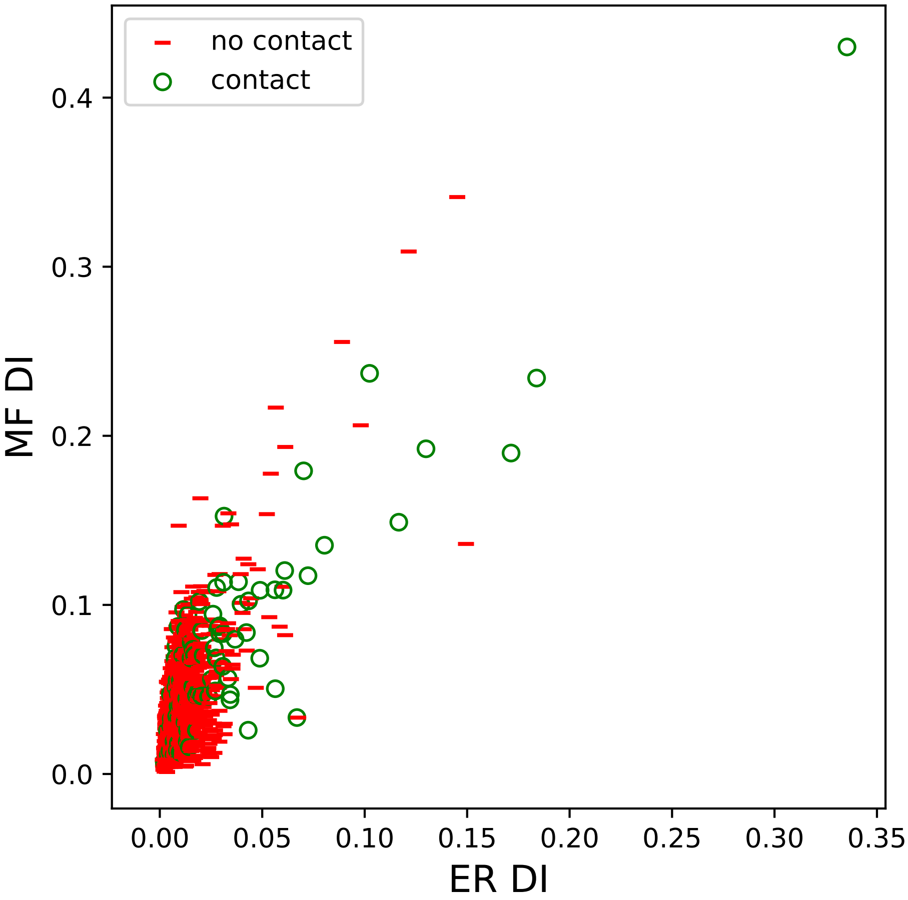

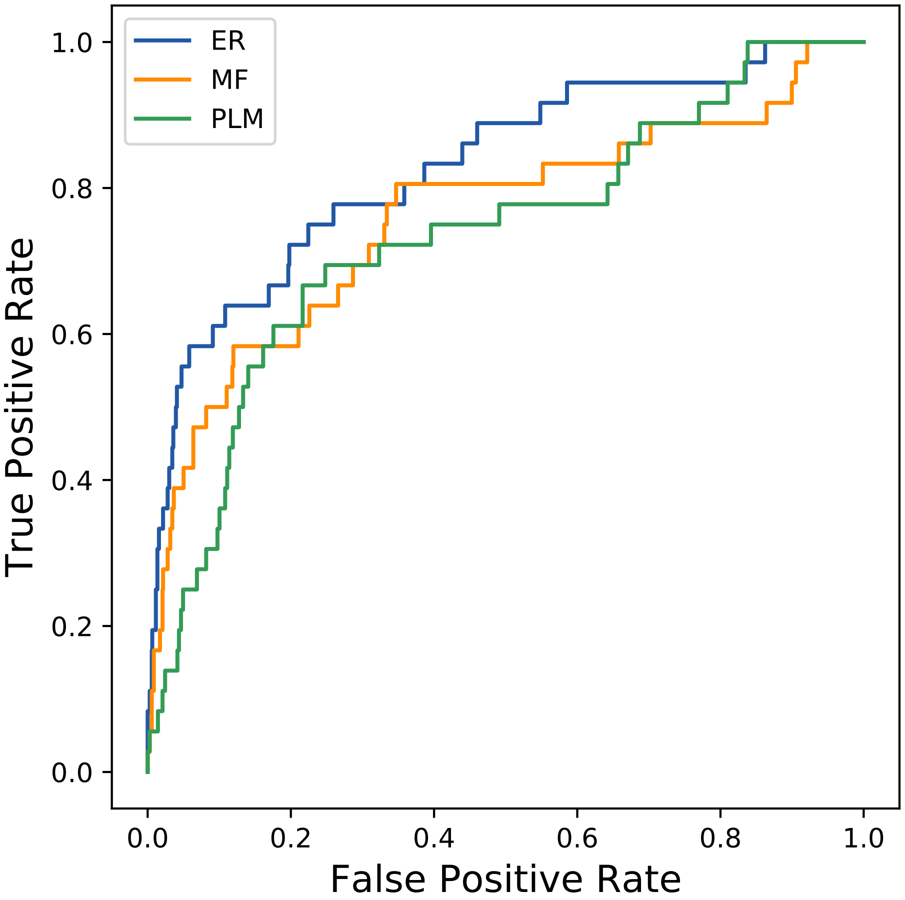

### 3125seq_138col_4psd_PF01083_plots.pdf

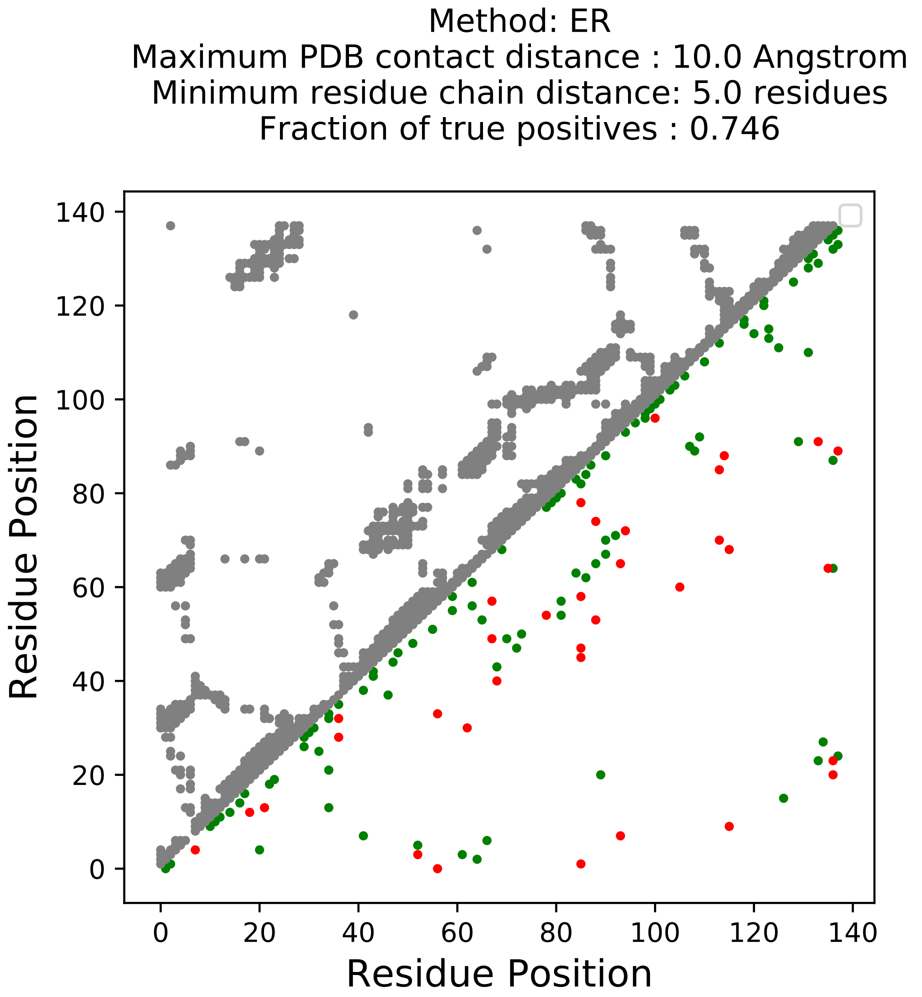

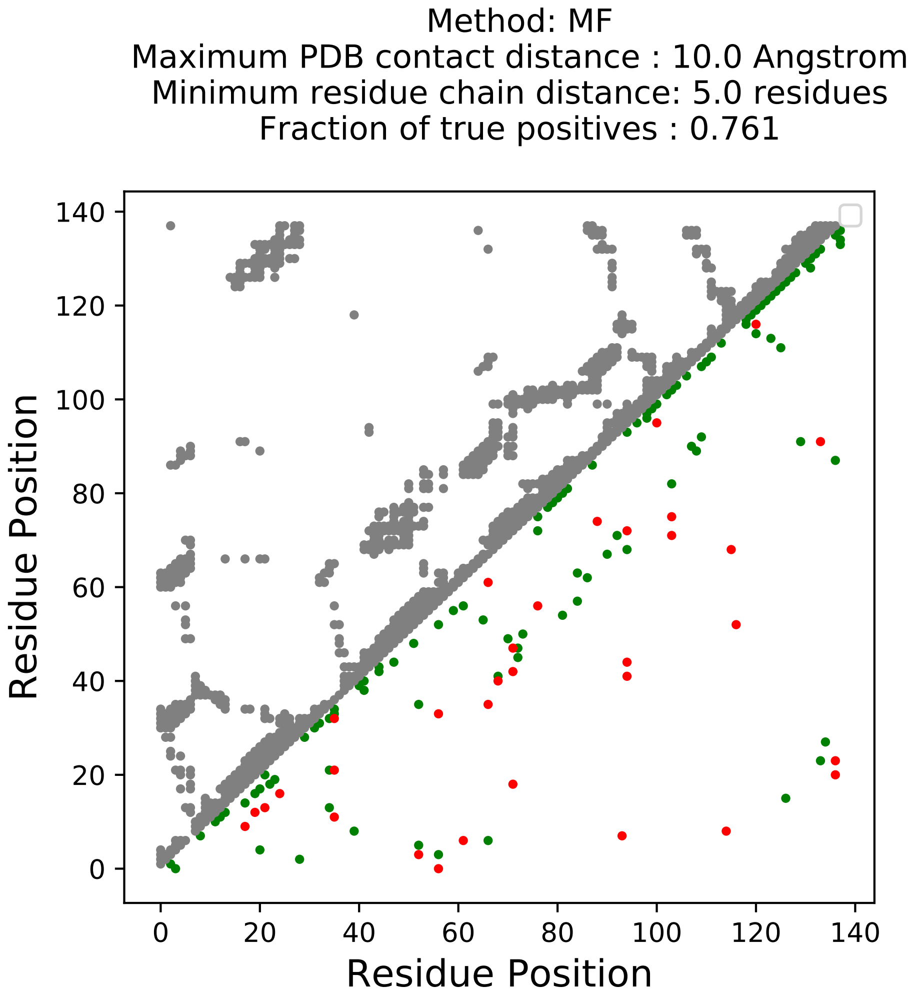

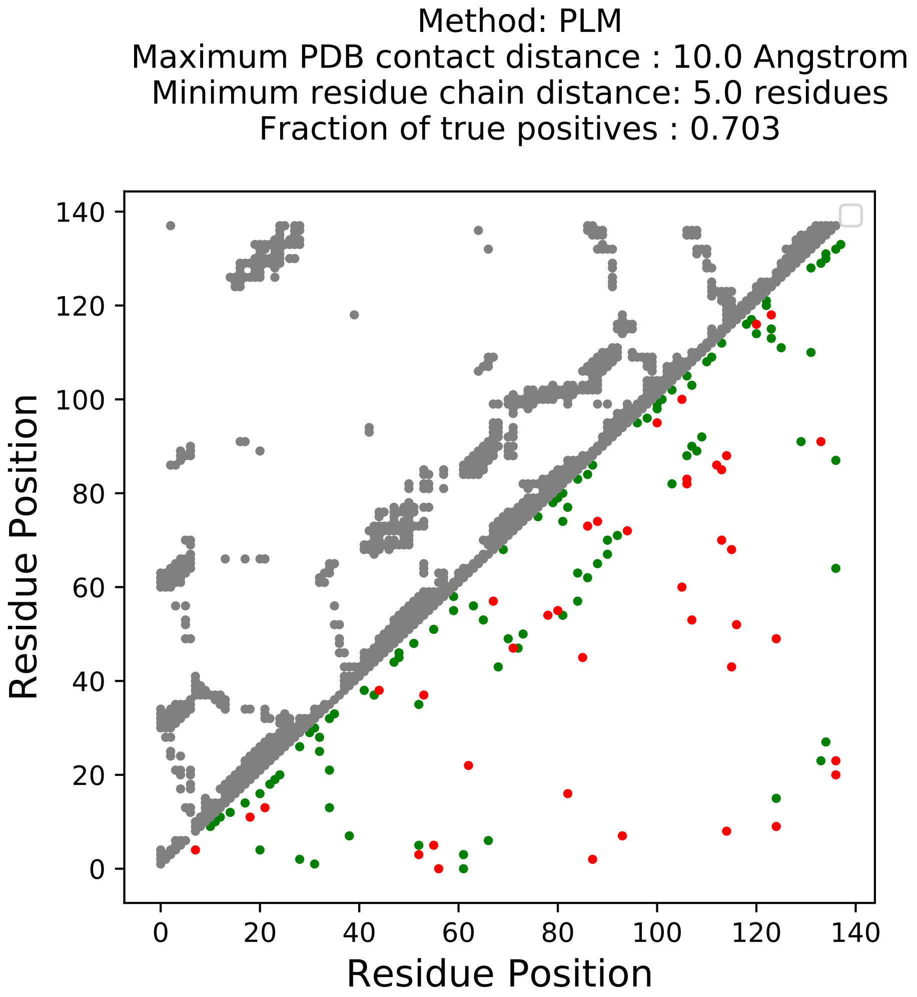

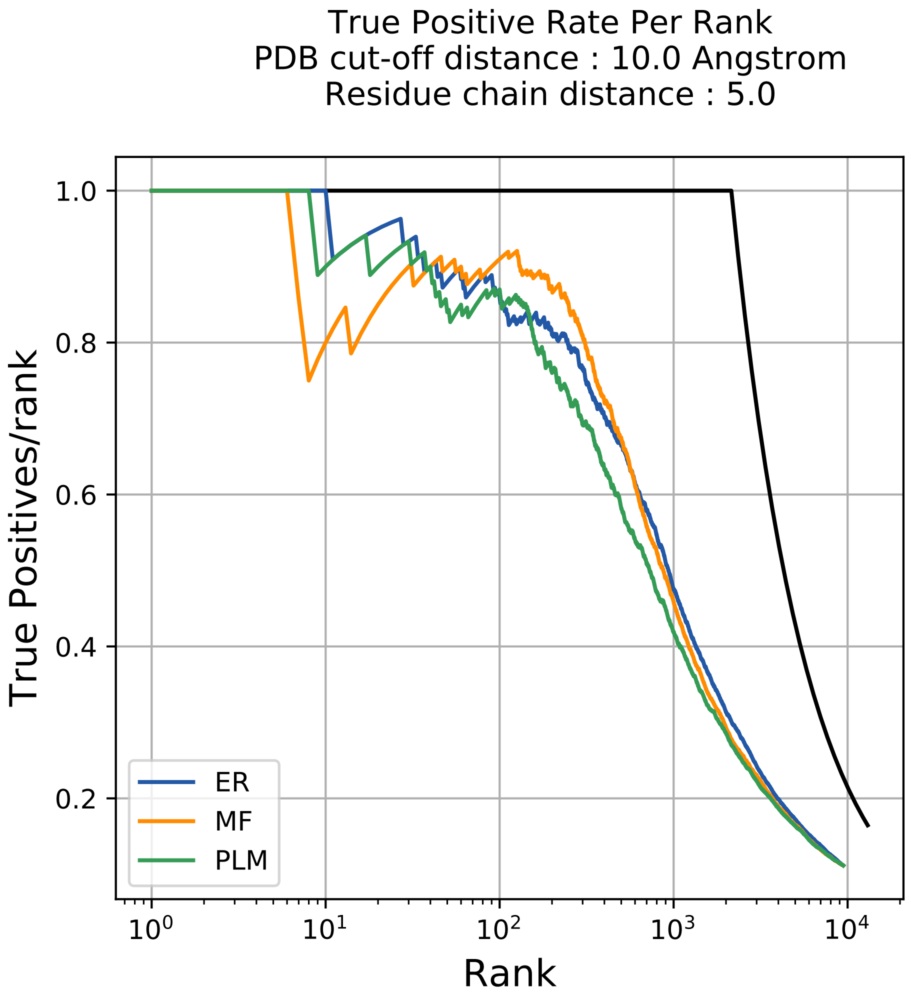

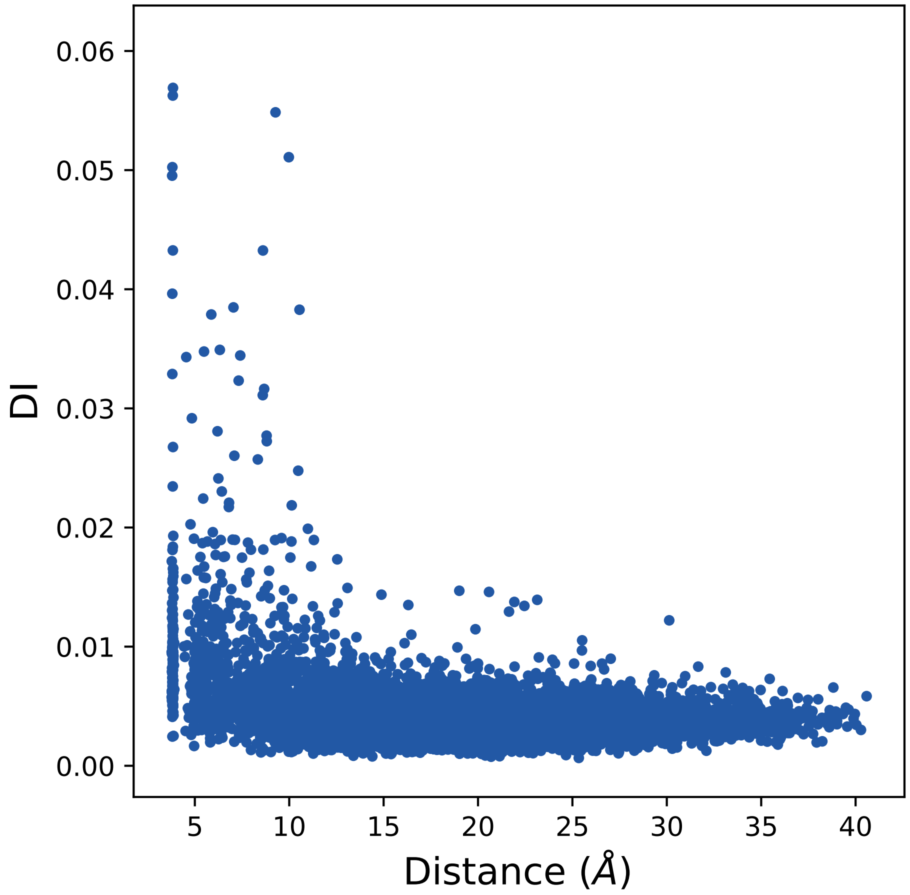

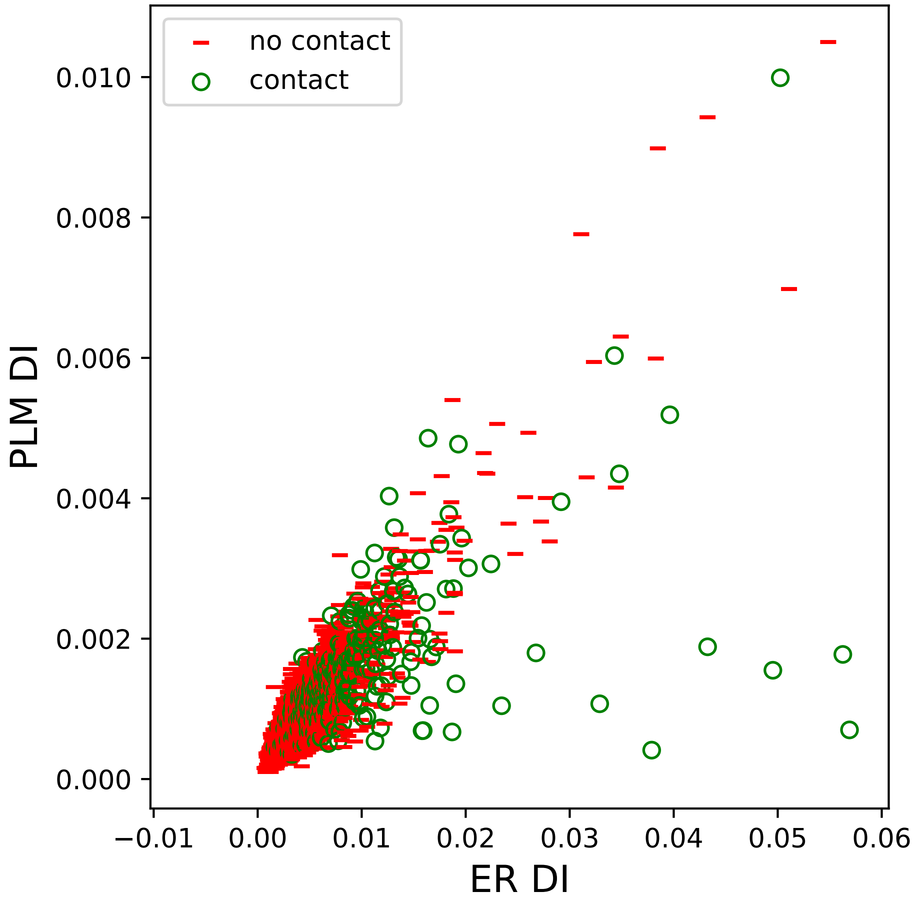

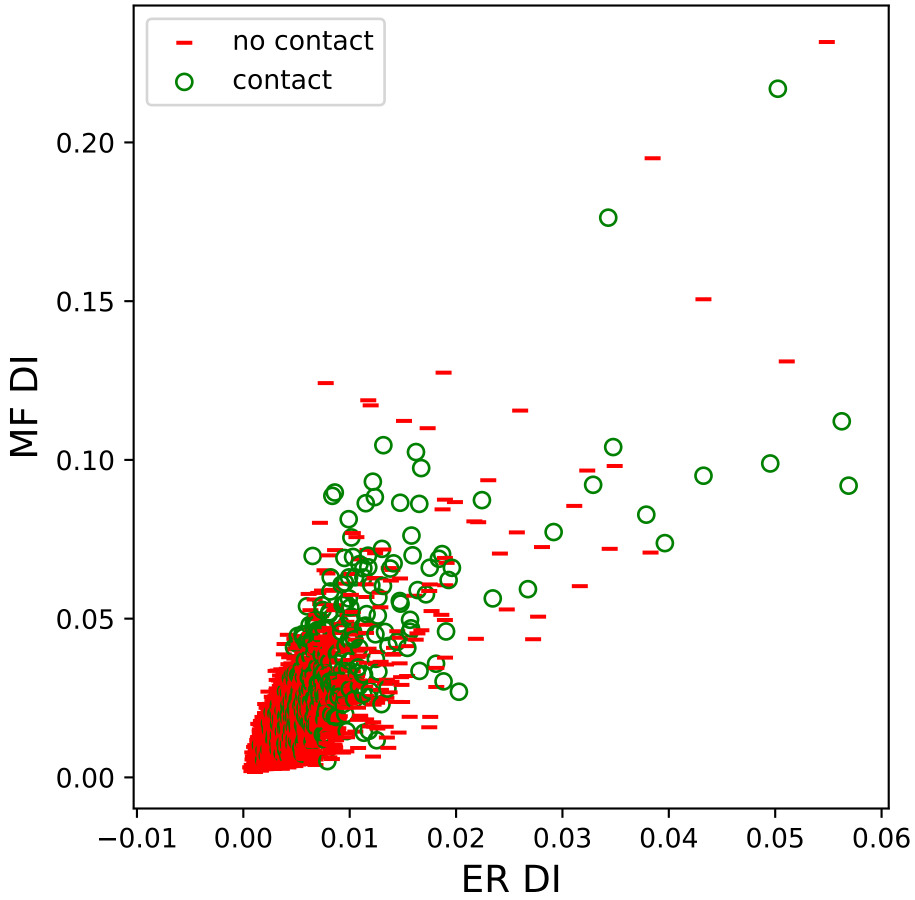

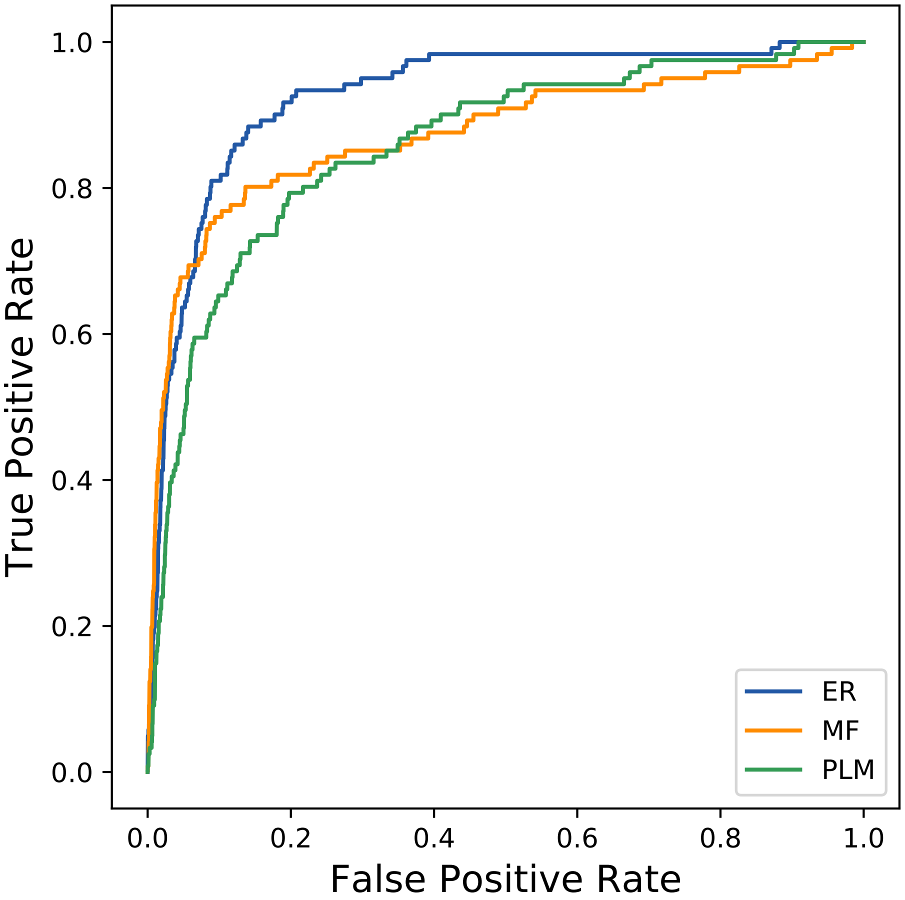

### 3251seq_65col_4zaz_PF02441_plots.pdf

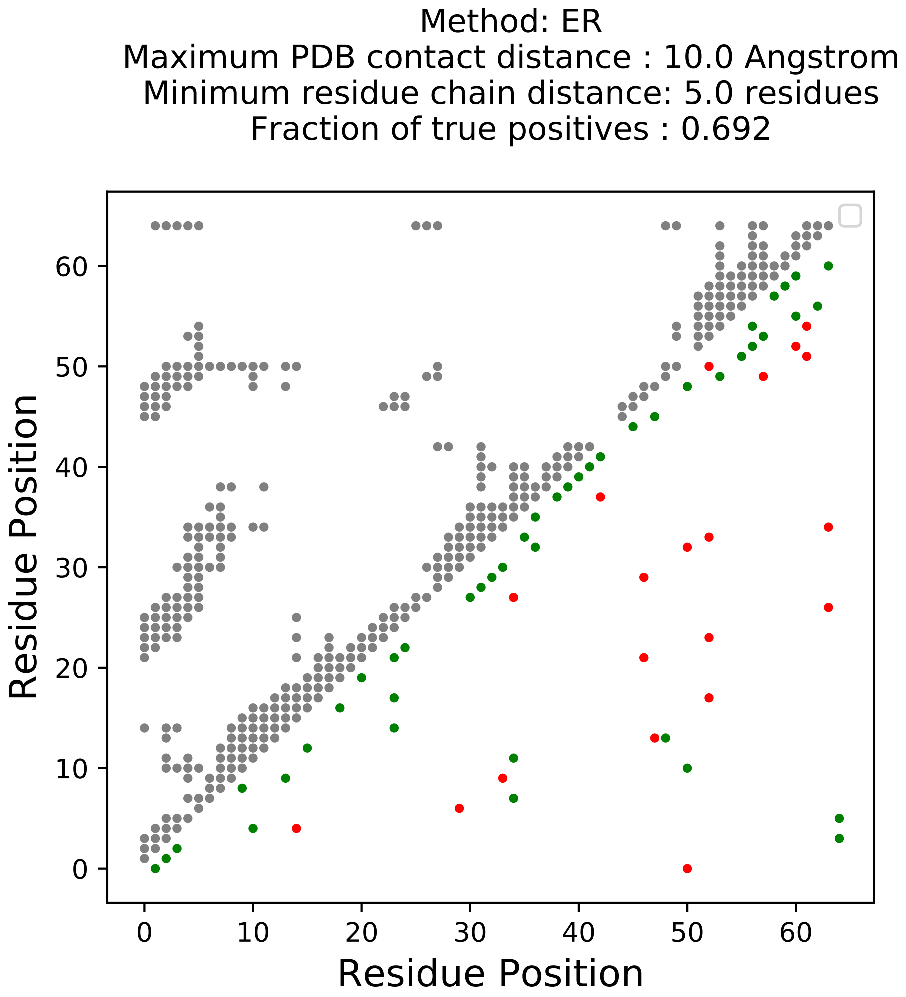

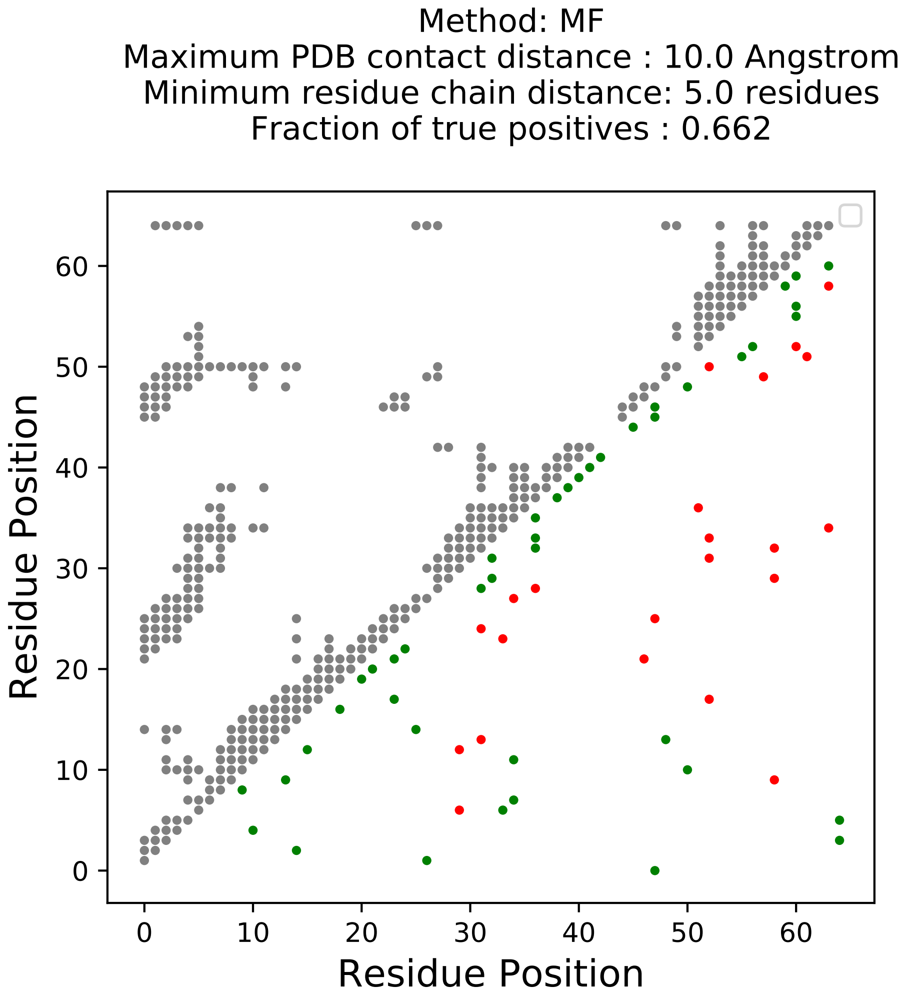

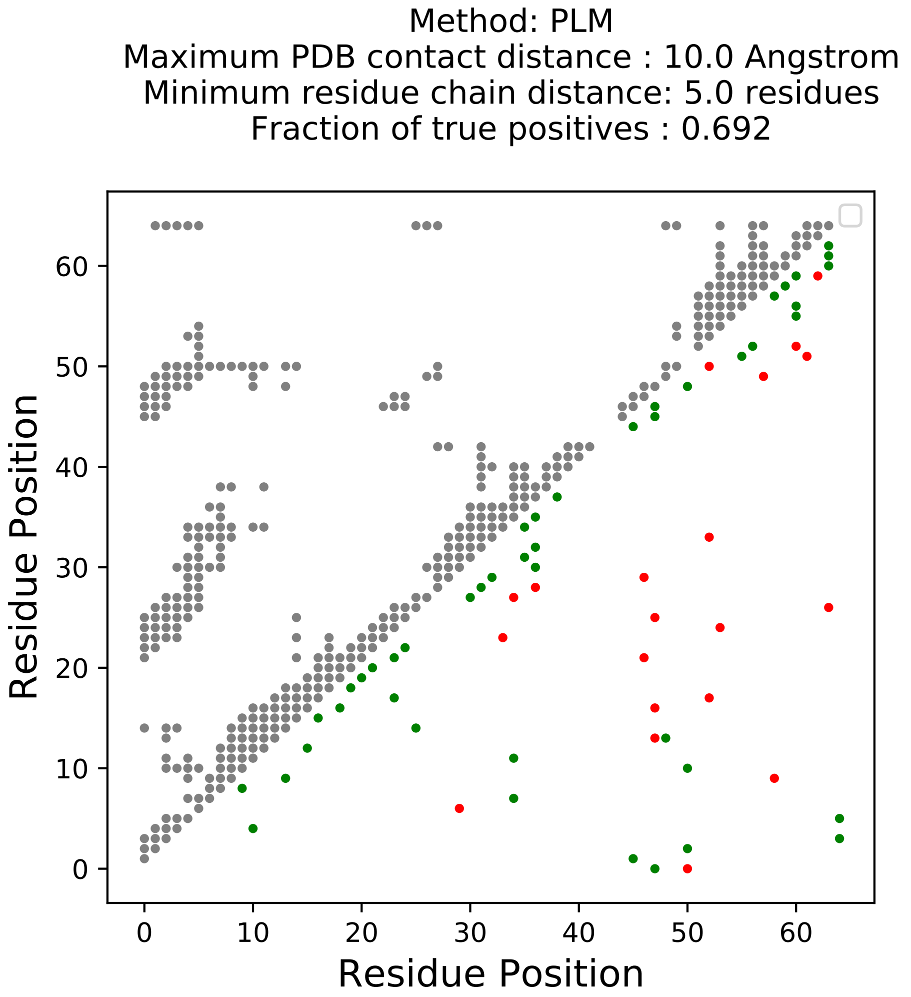

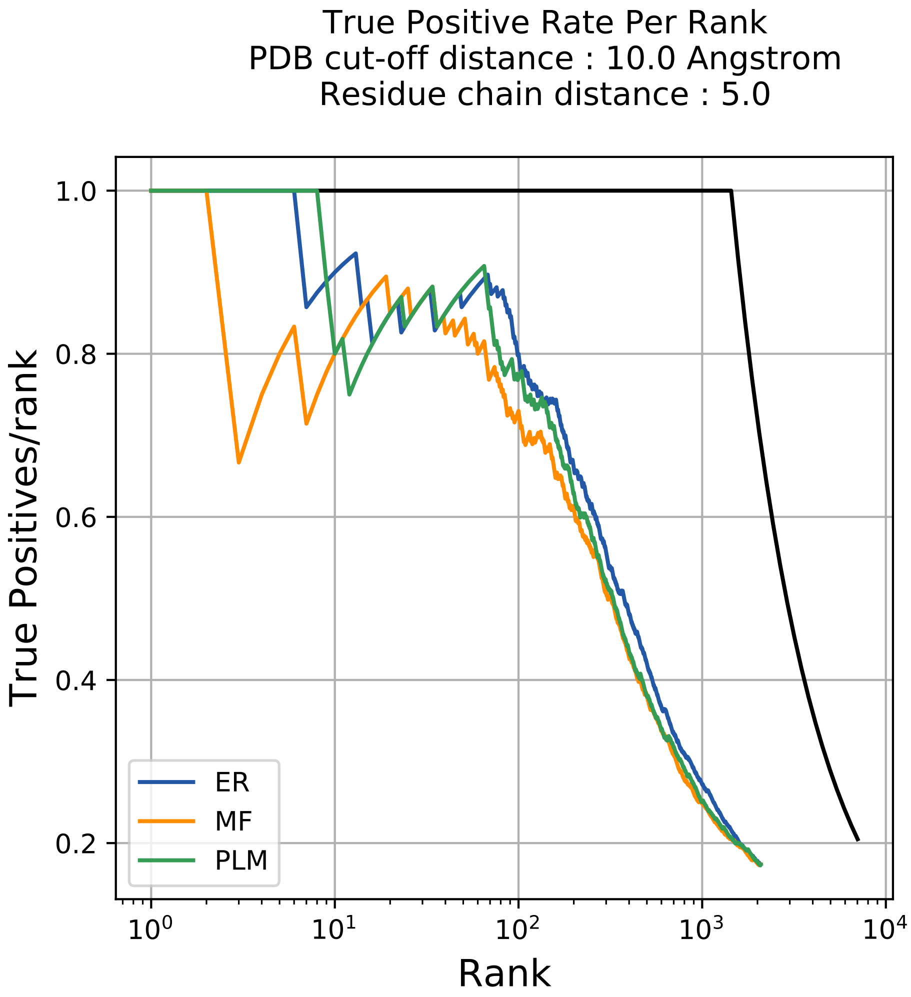

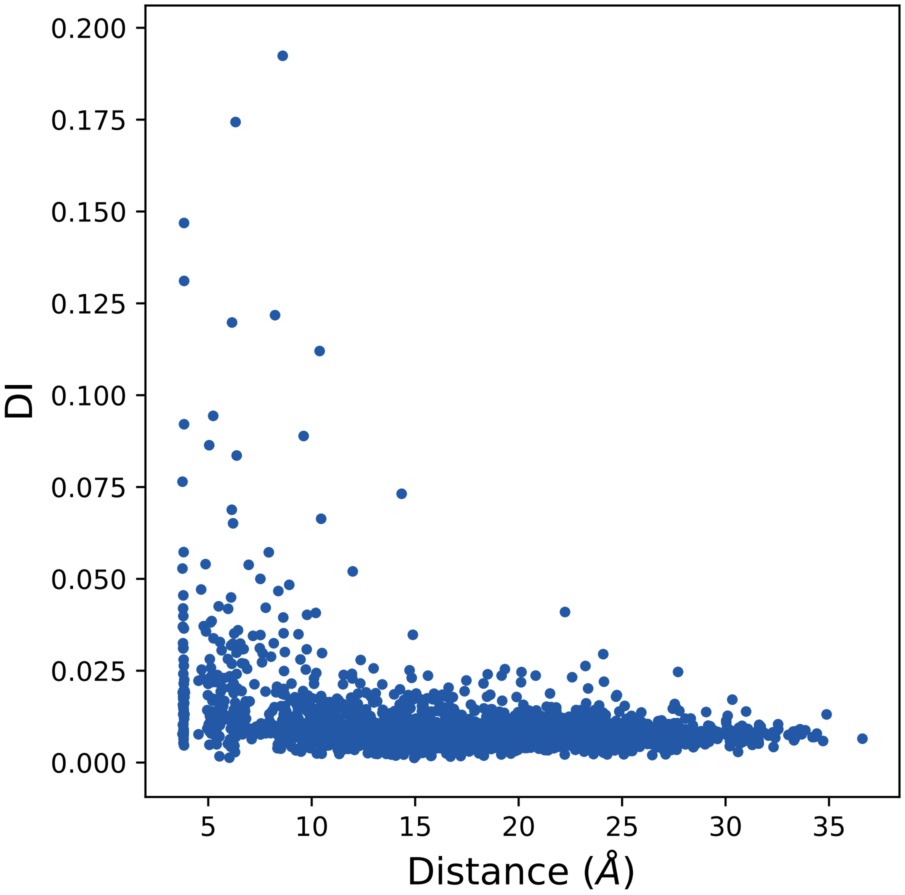

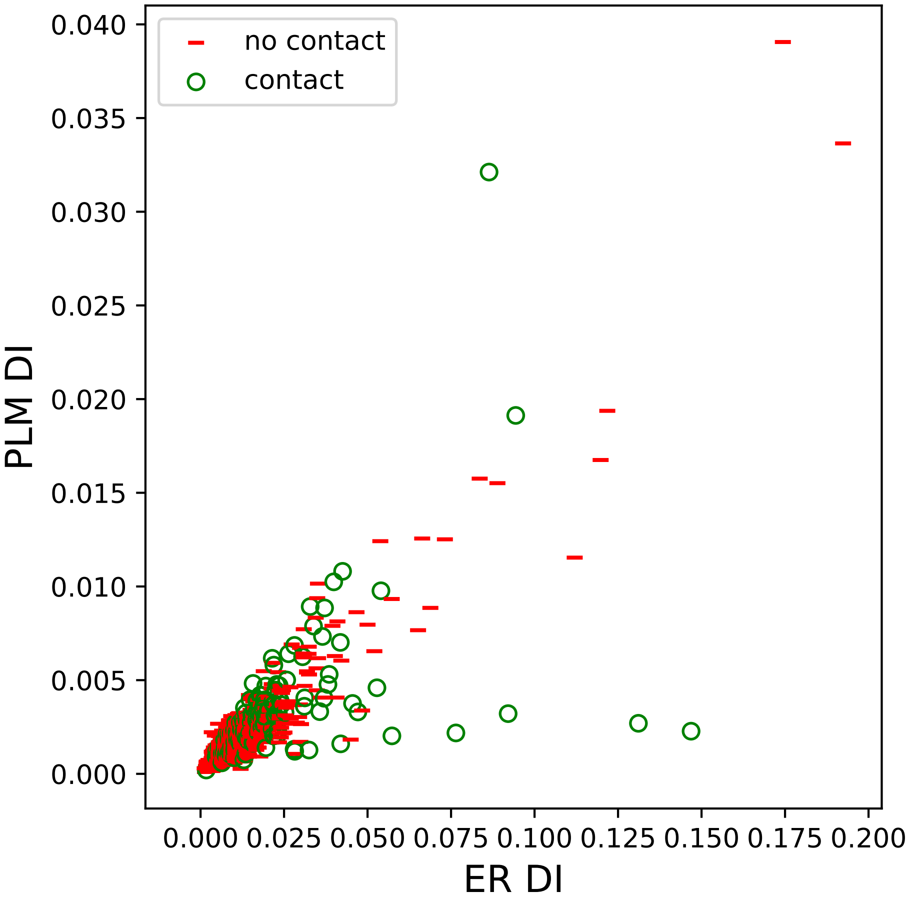

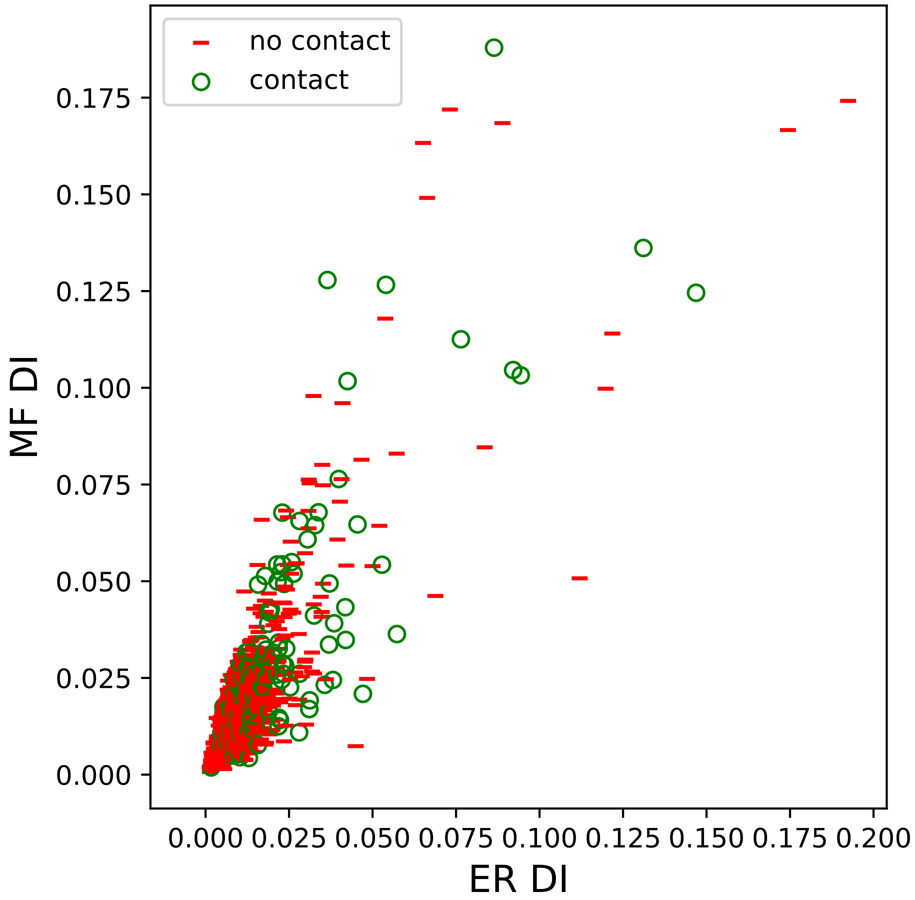

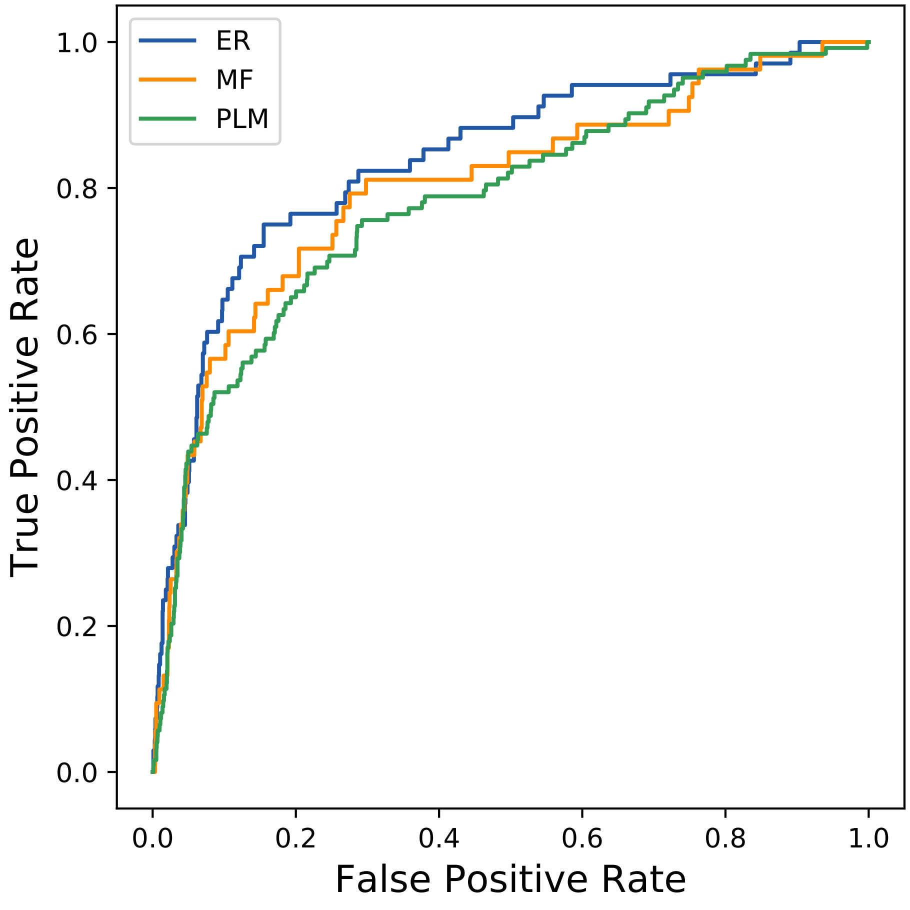

### 3408seq_111col_1d8f_PF00413_plots.pdf

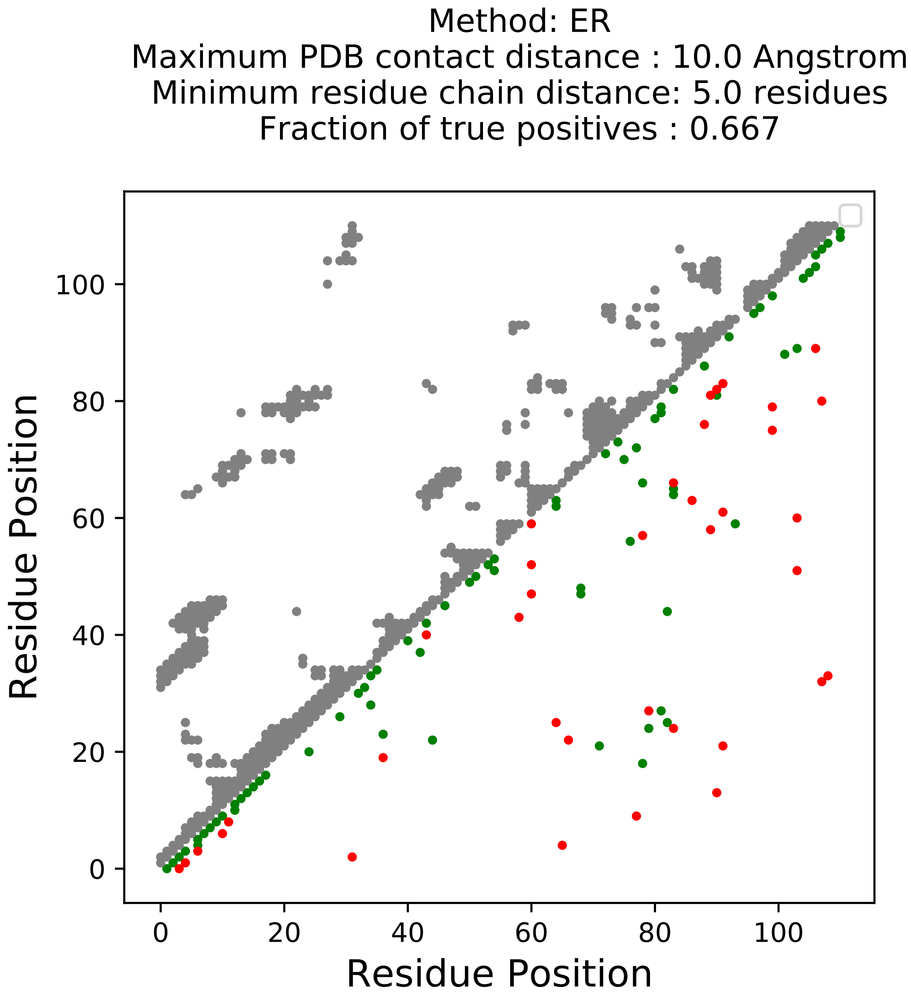

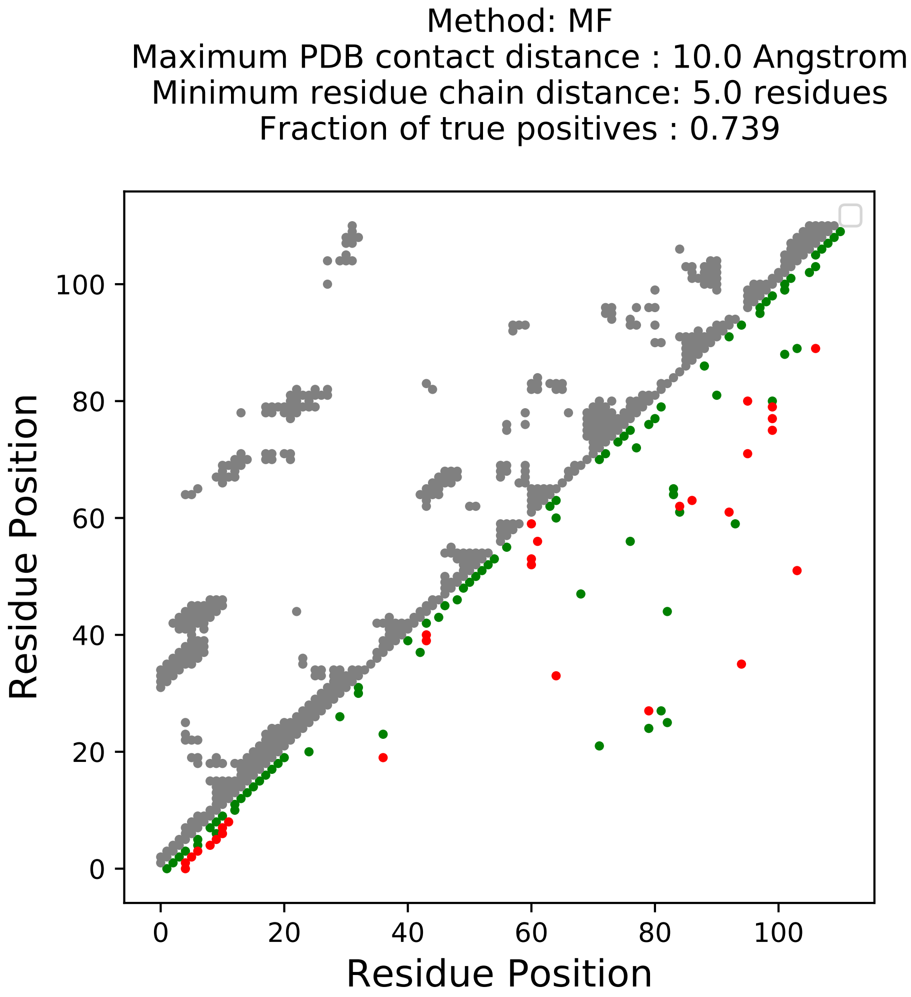

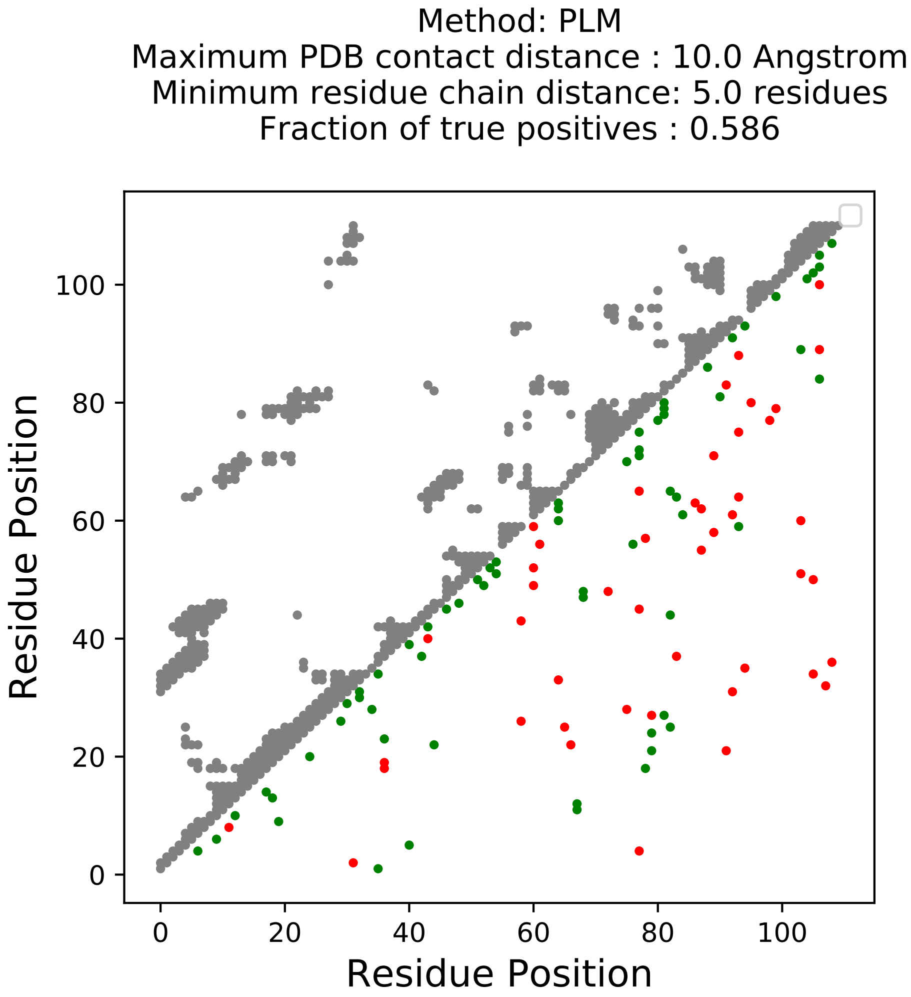

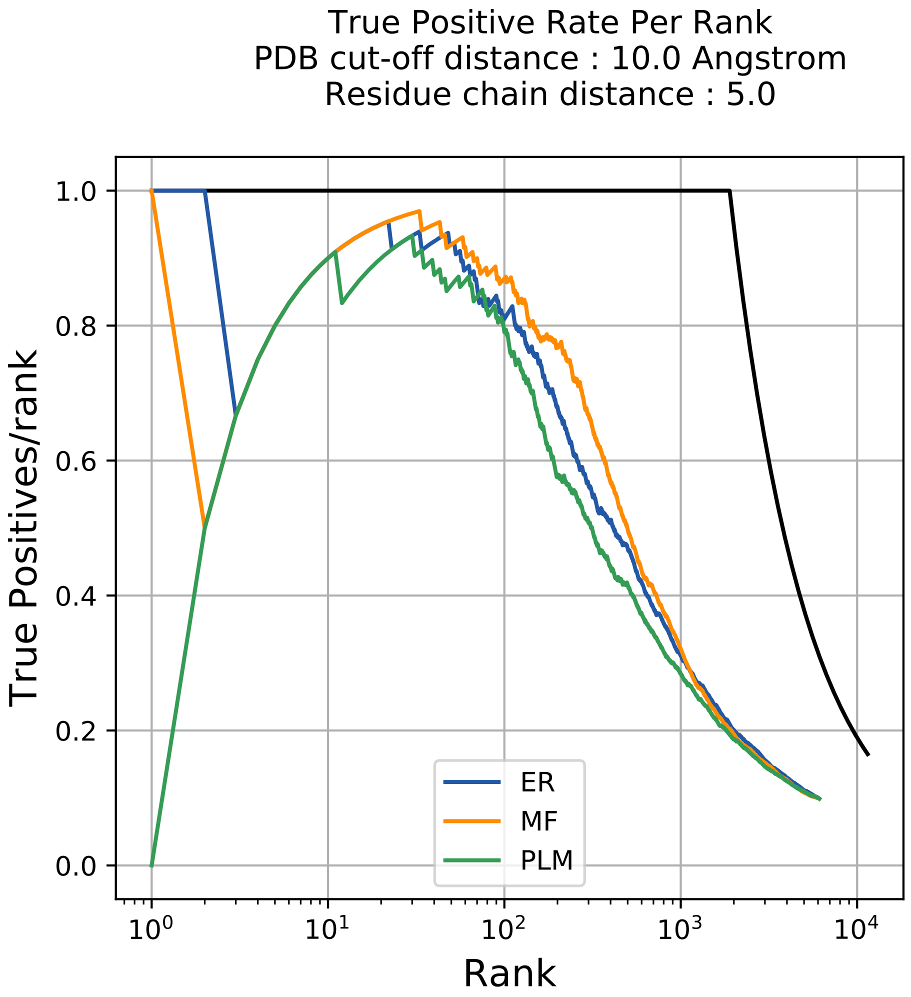

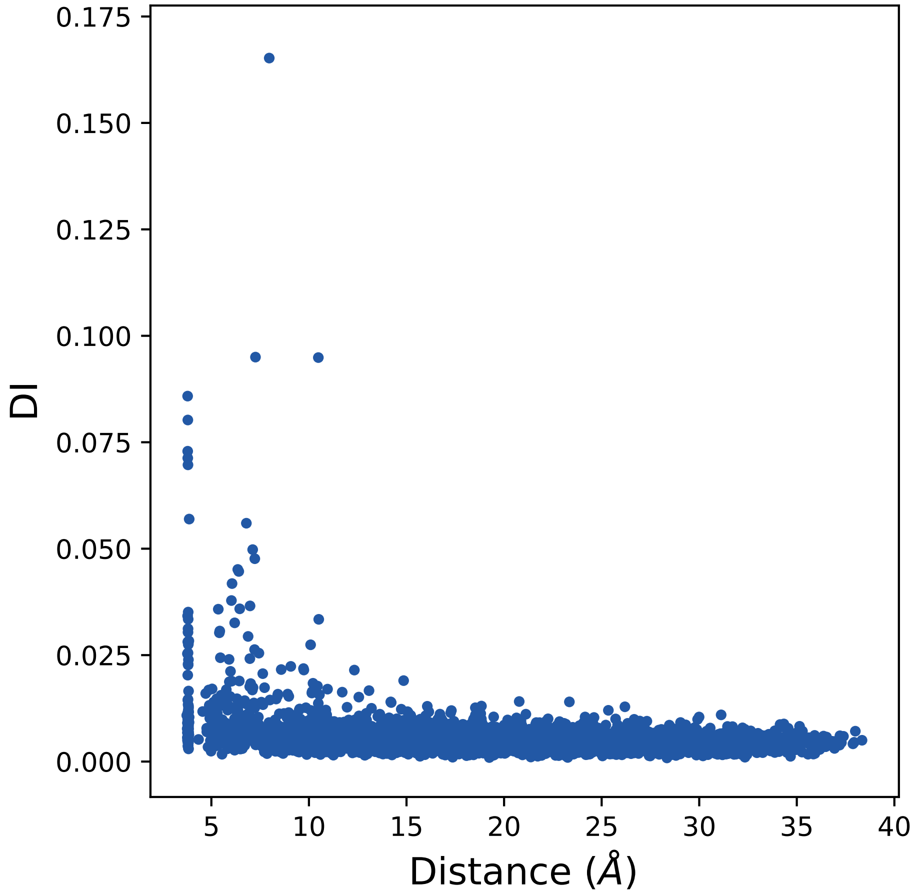

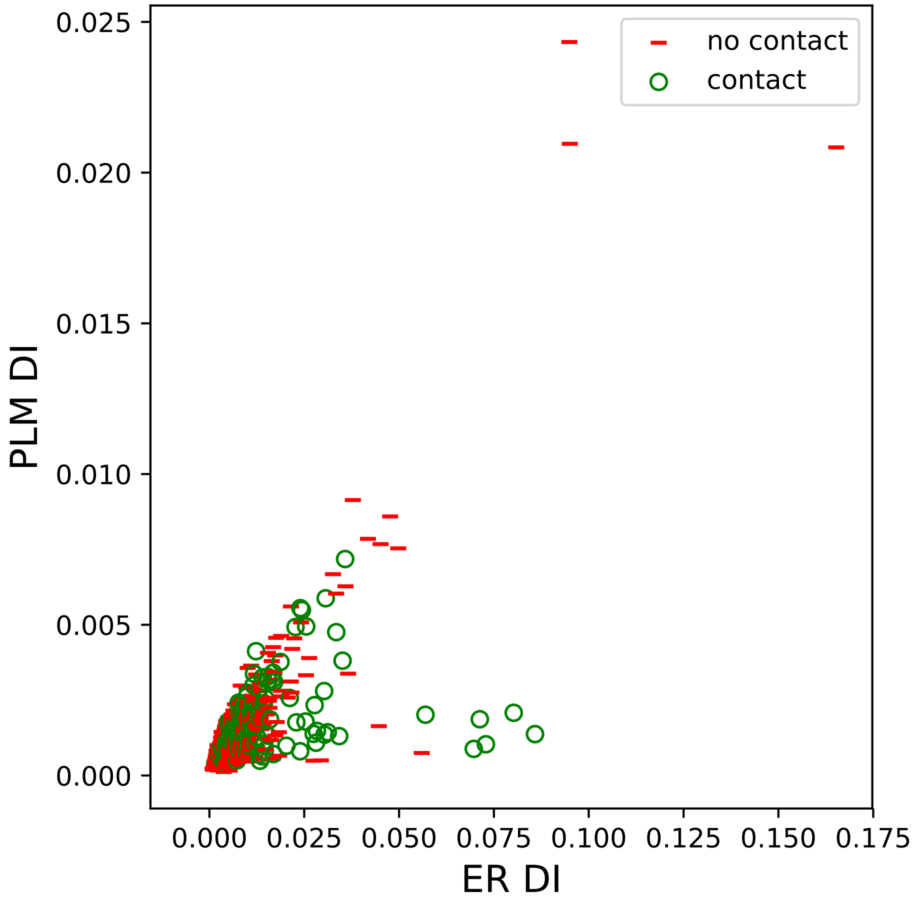

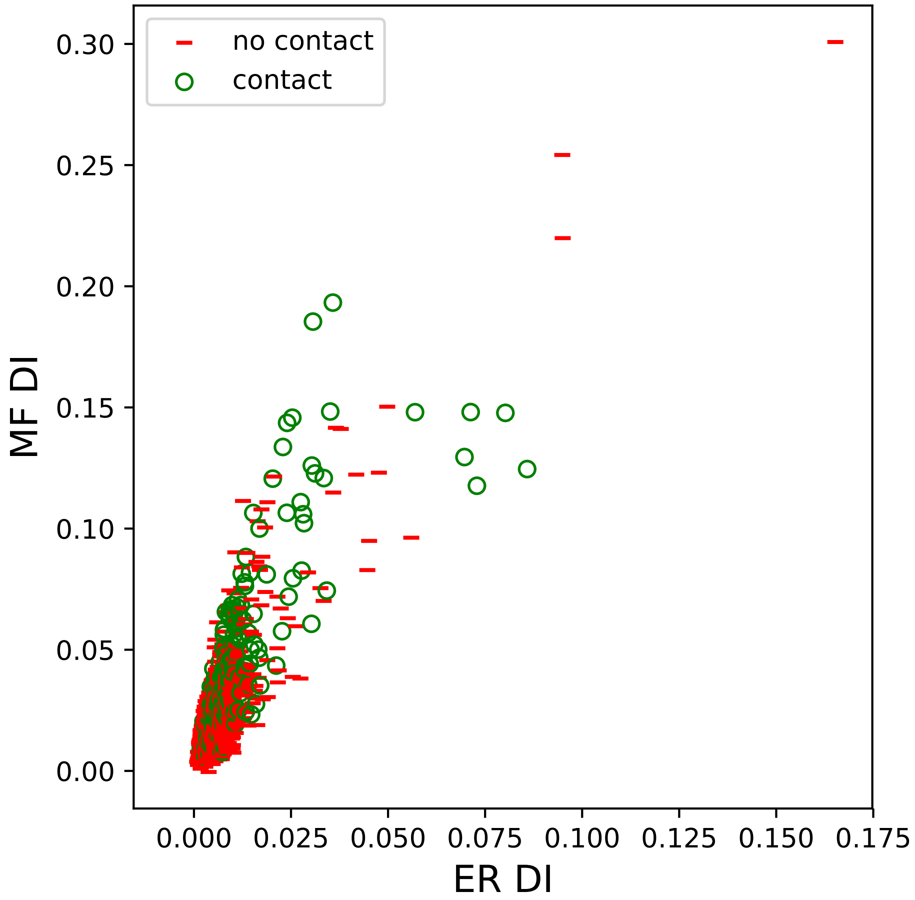

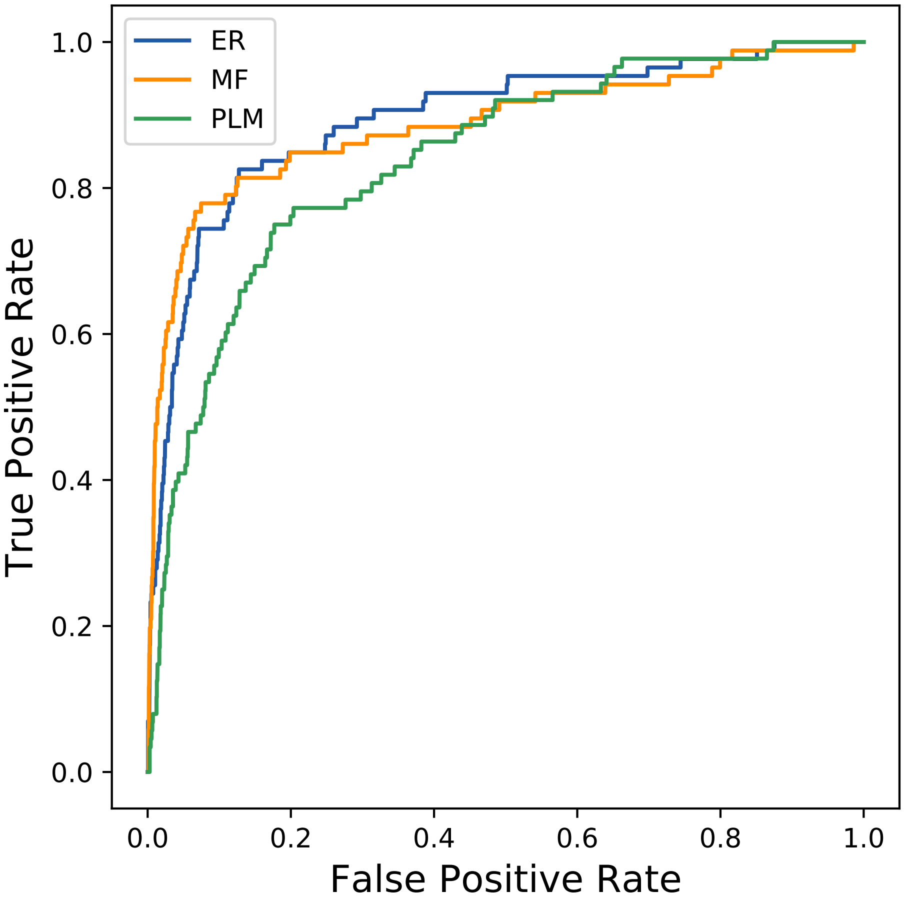

### 3492seq_206col_5zpq_PF01179_plots.pdf

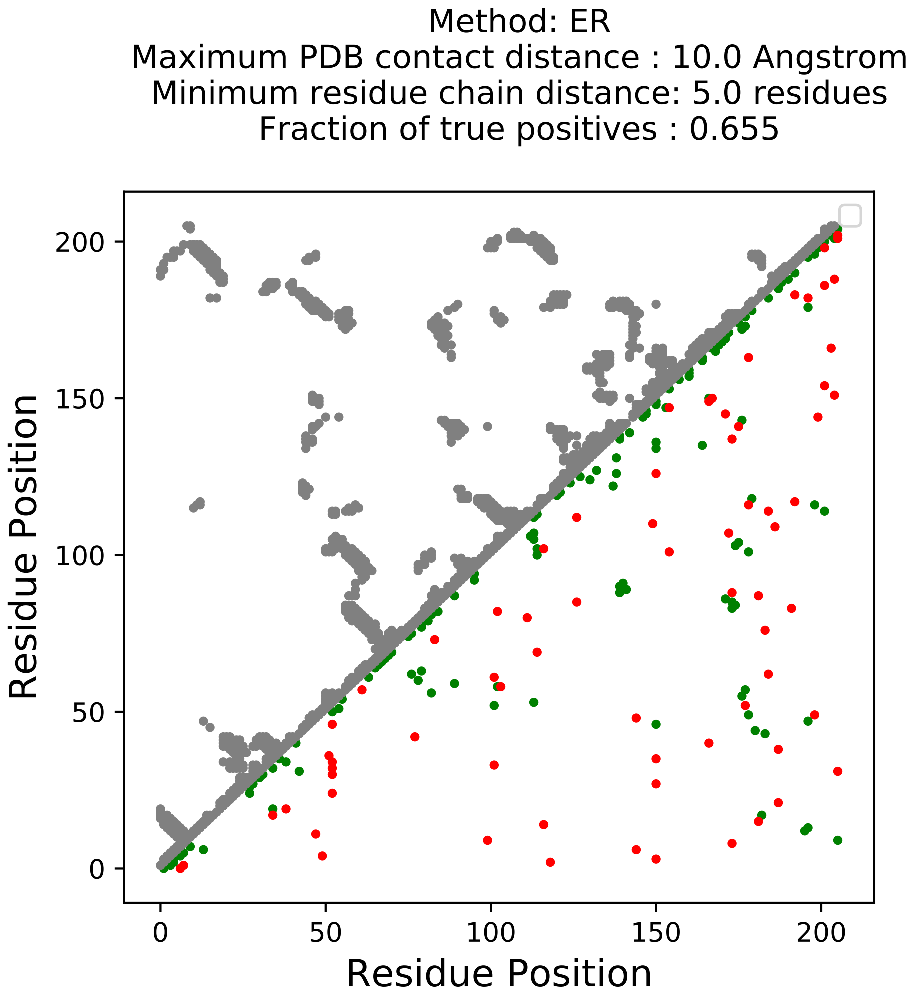

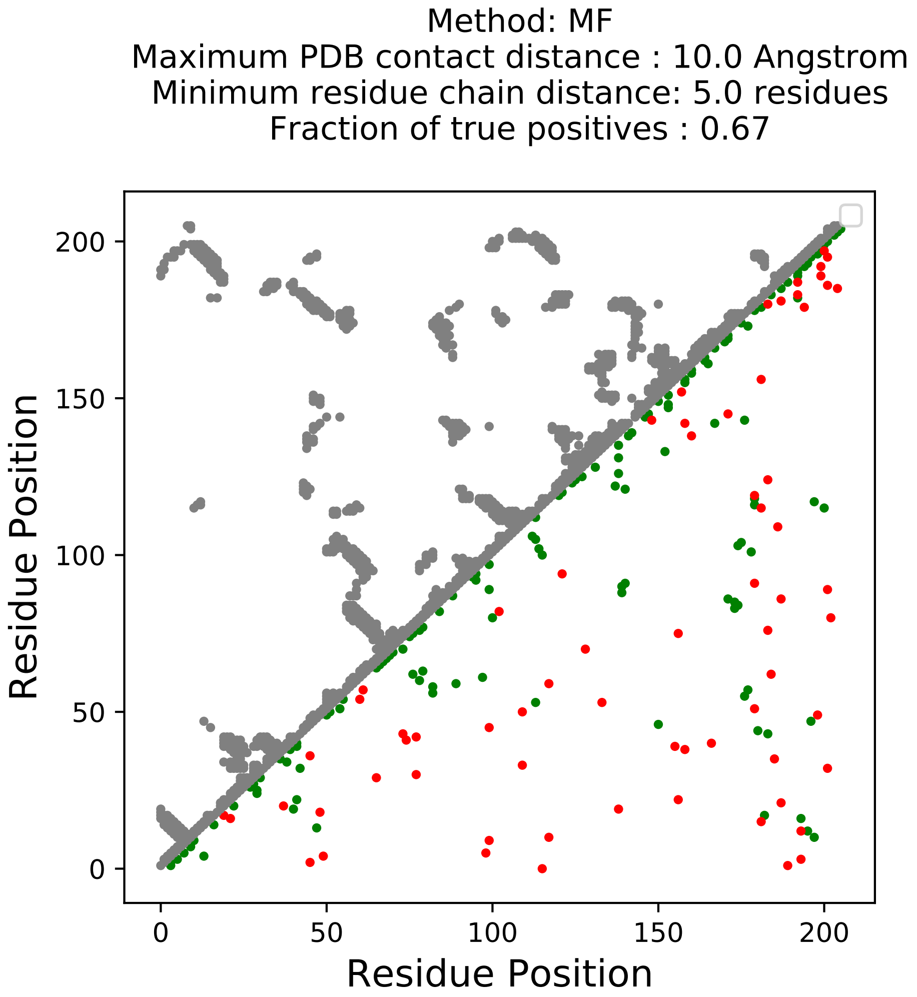

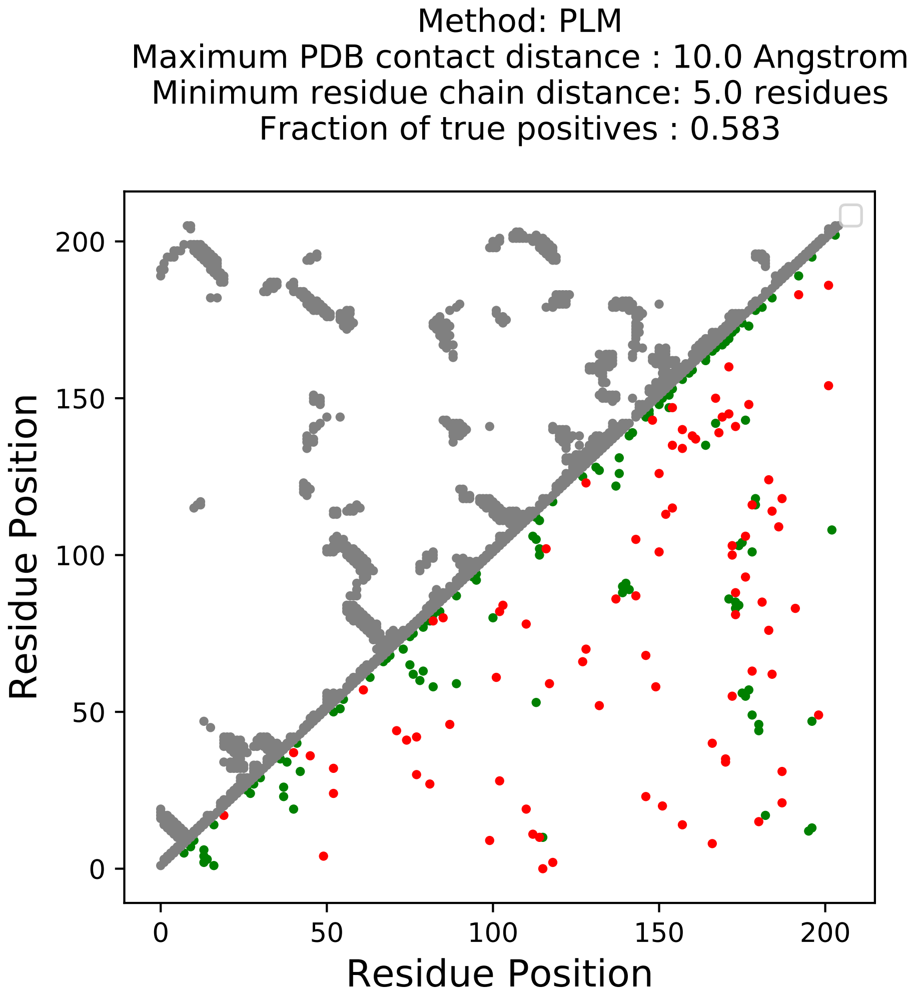

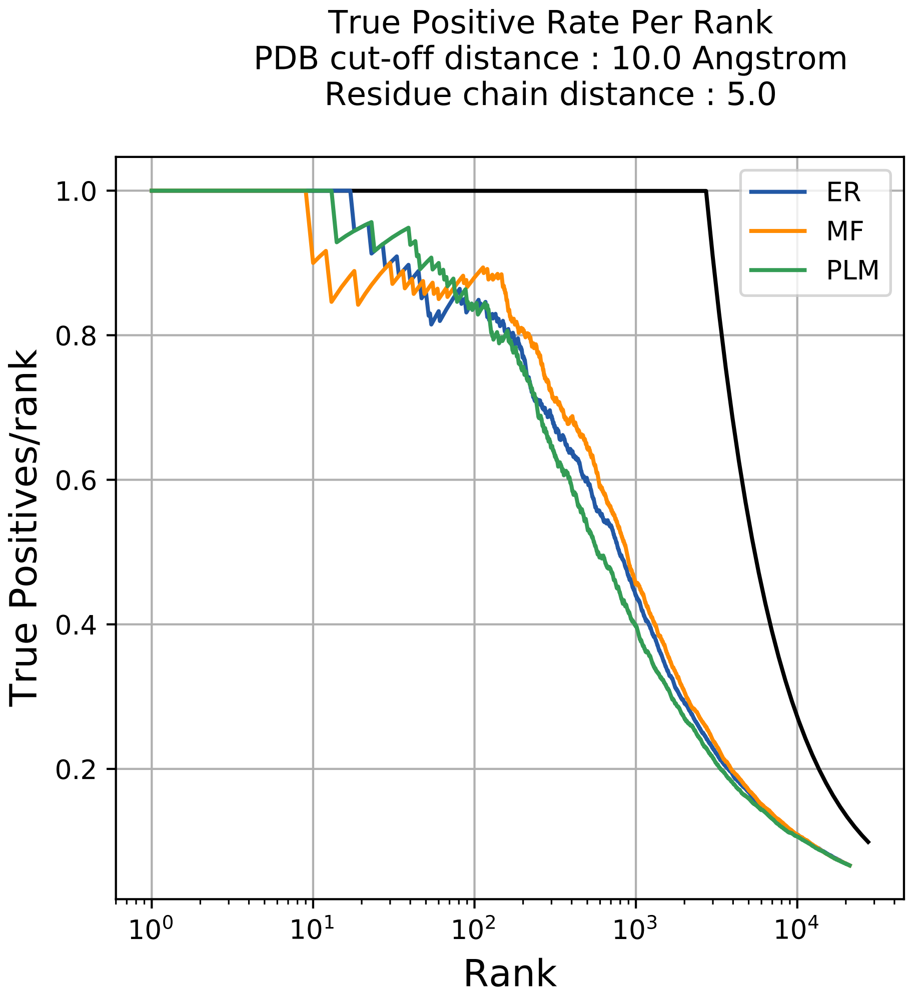

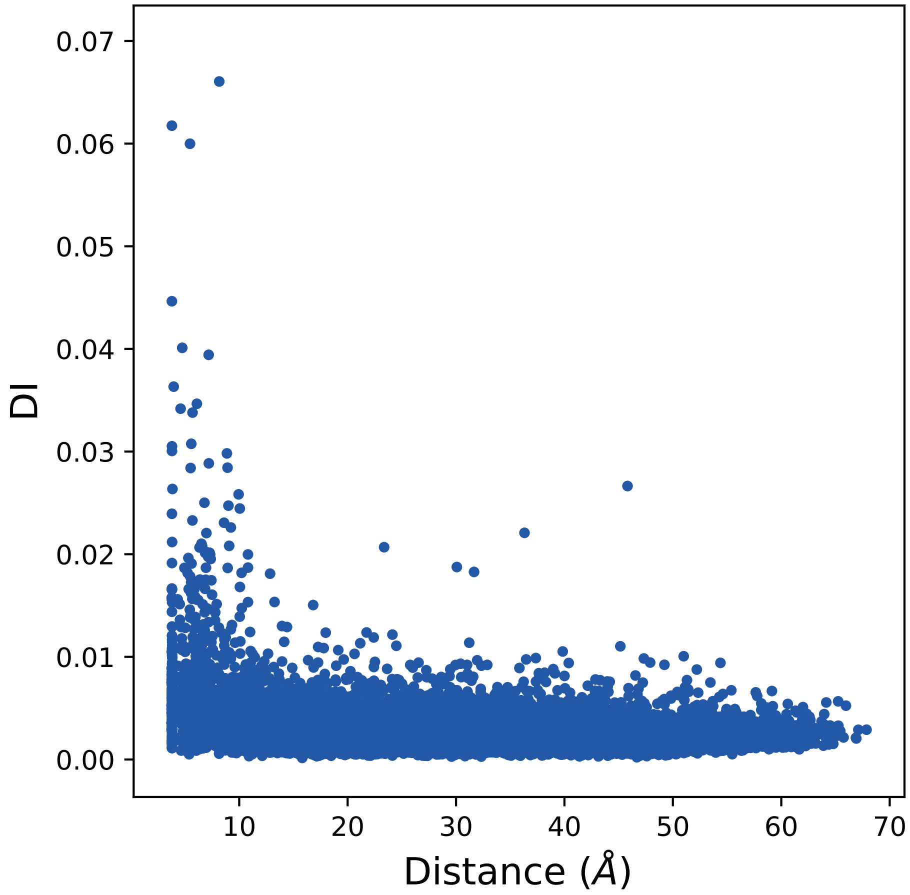

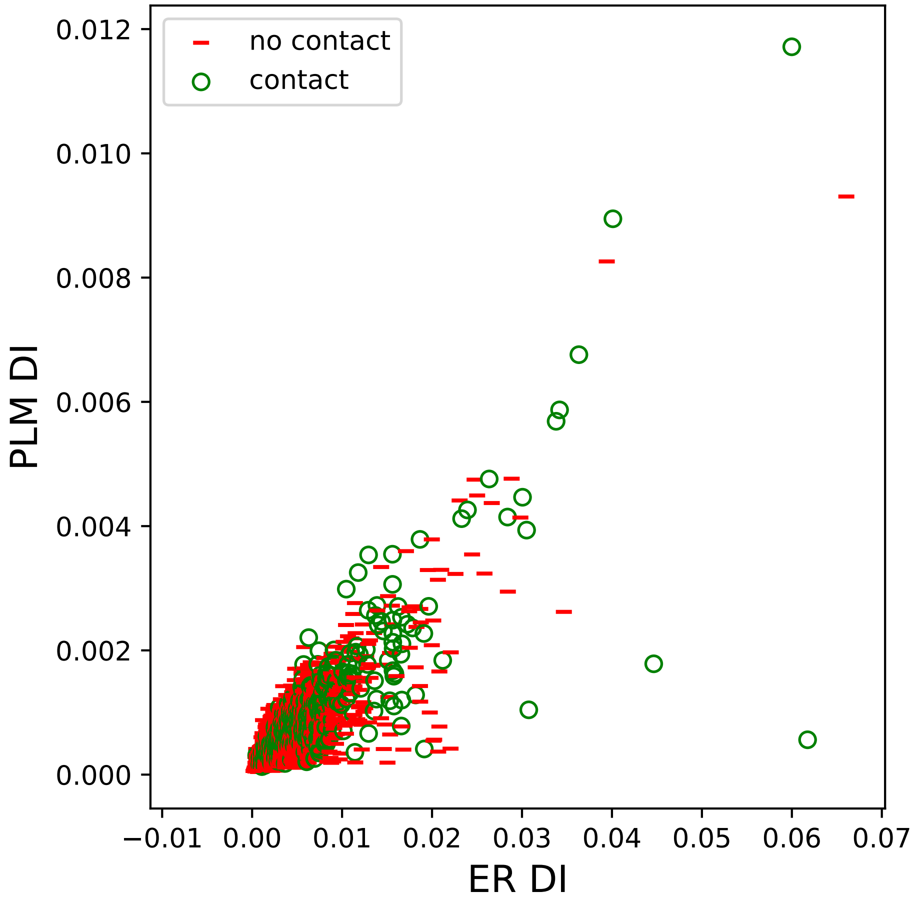

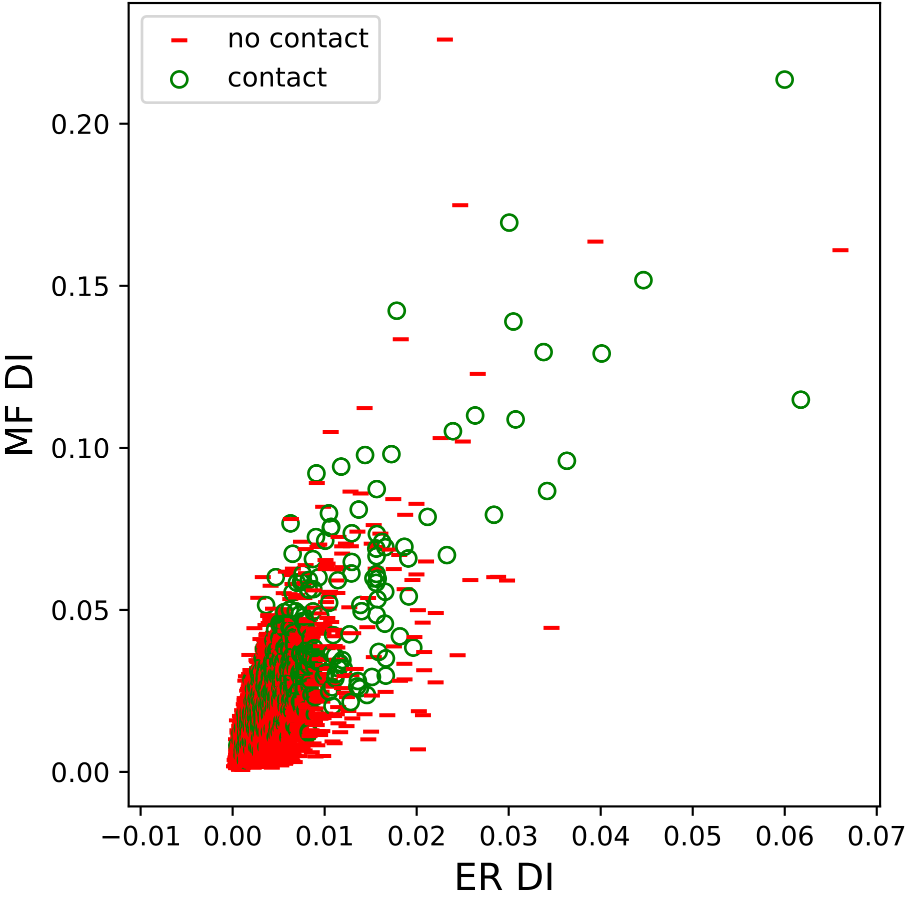

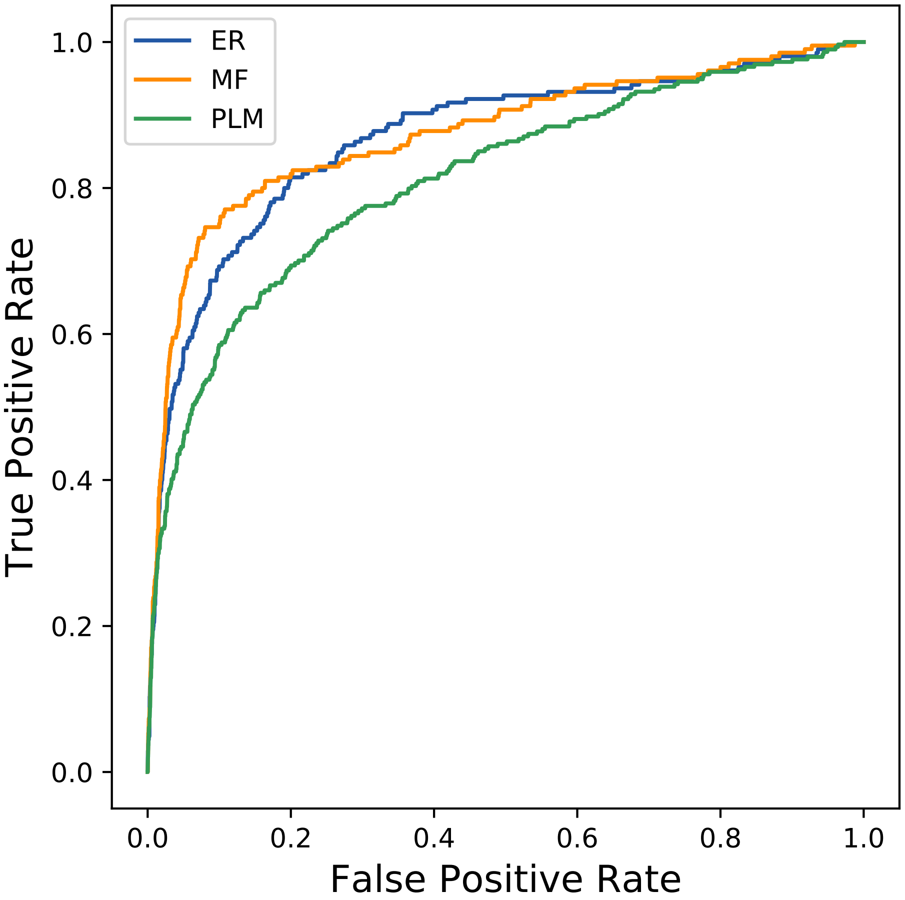

### 3619seq_84col_3zxf_PF00337_plots.pdf

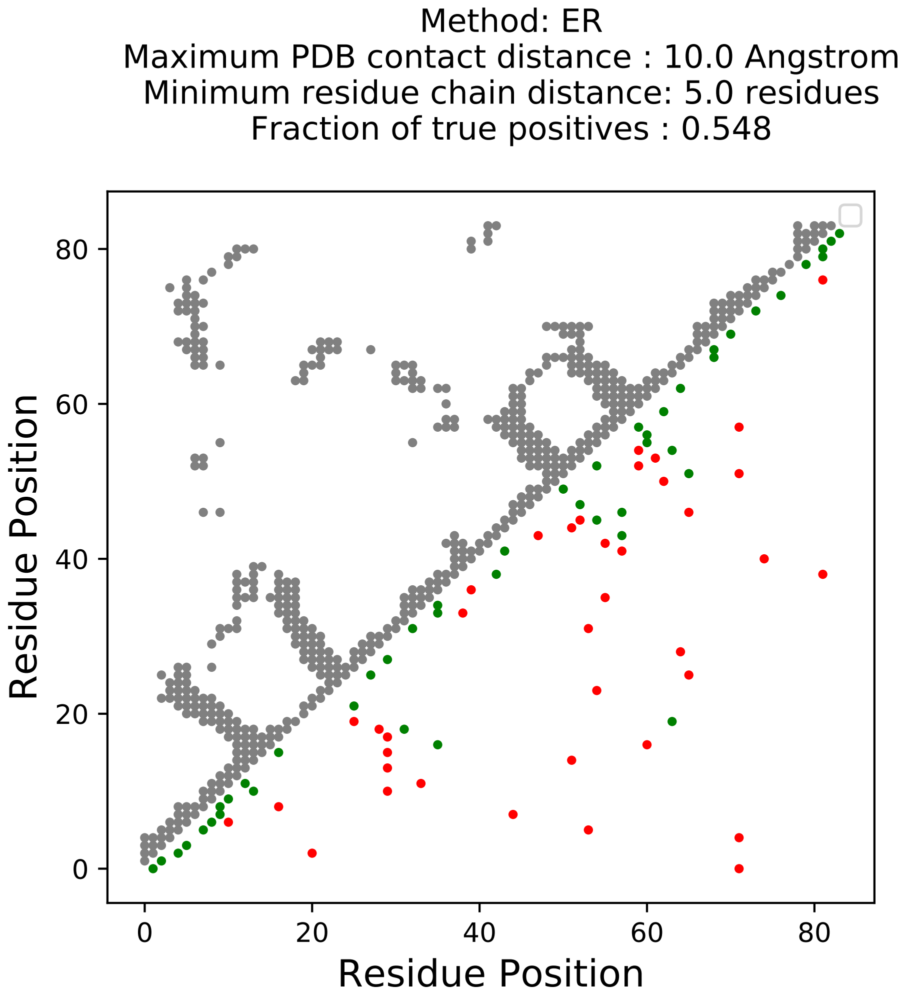

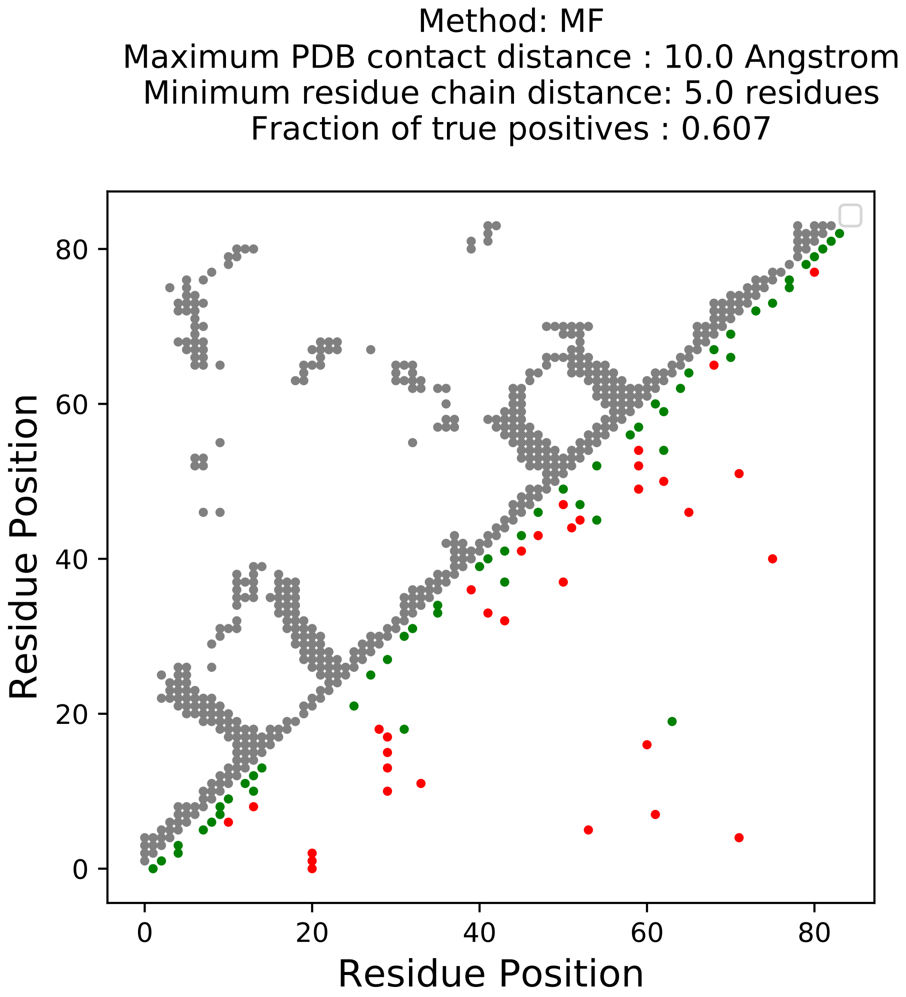

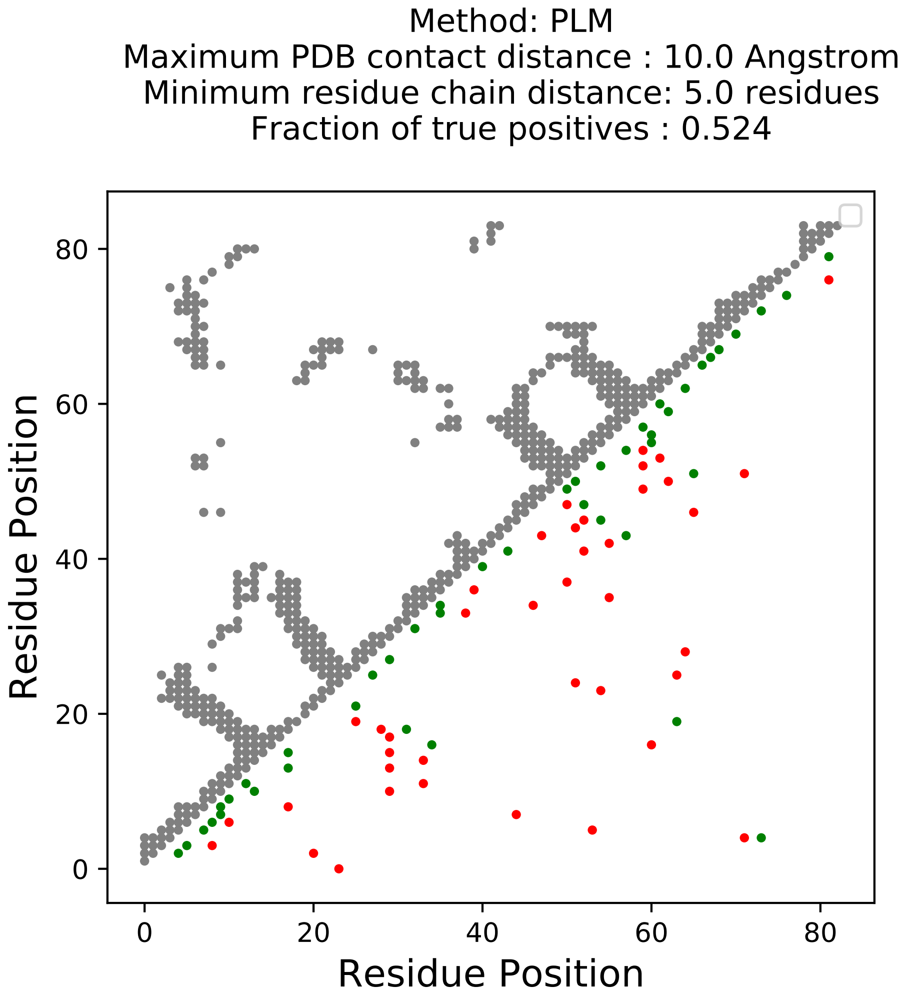

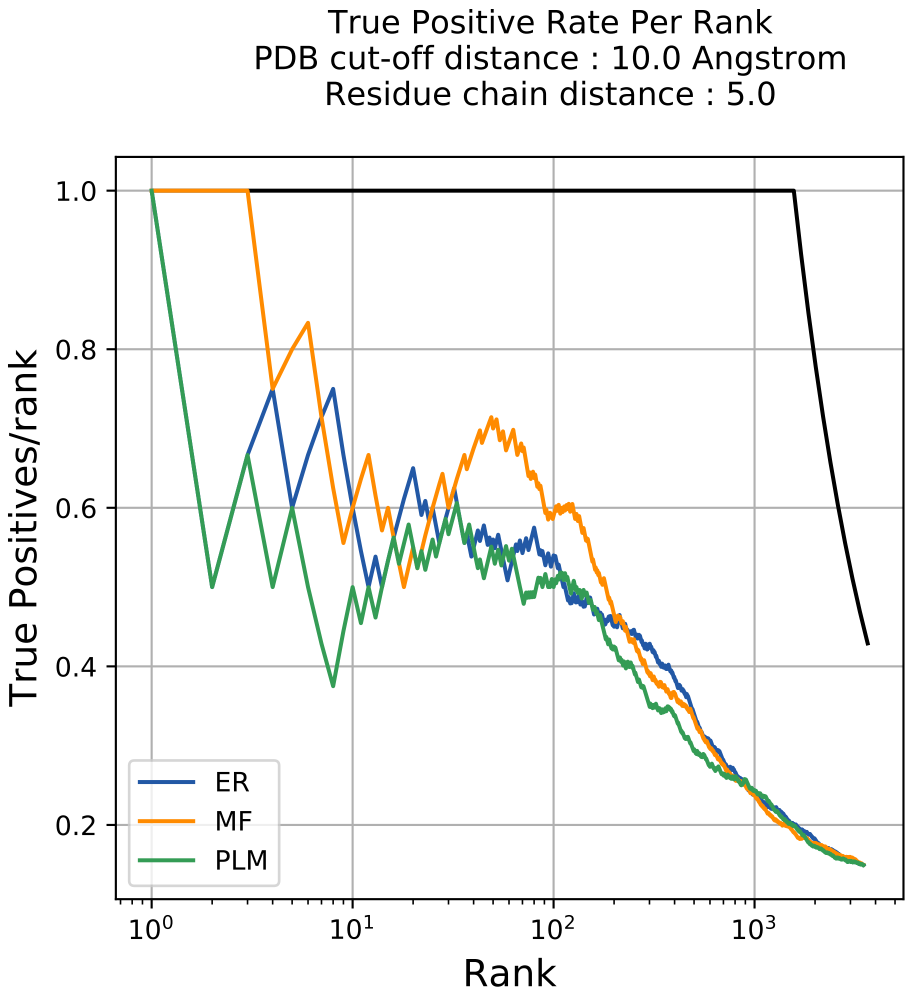

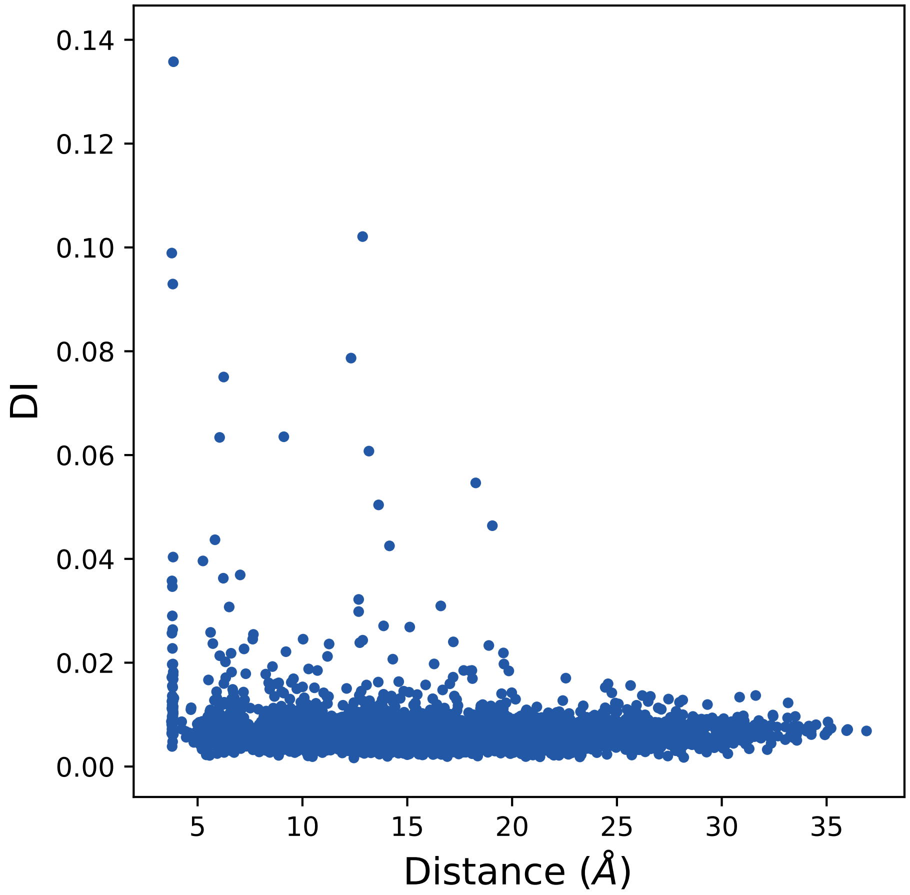

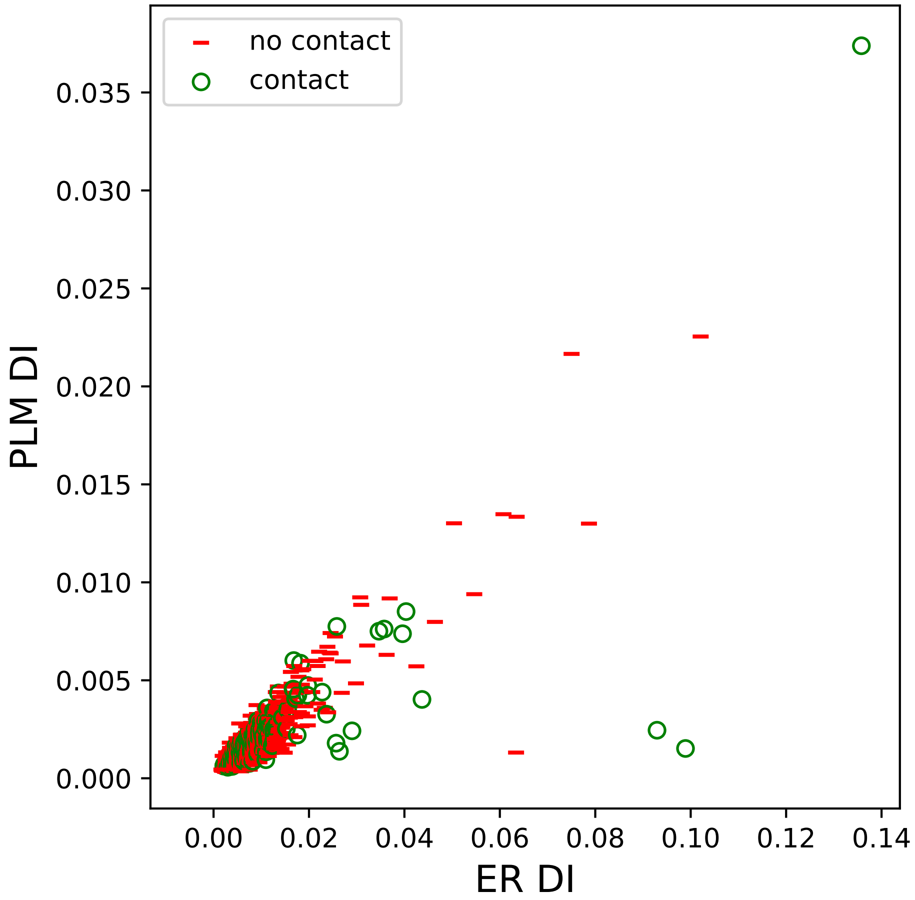

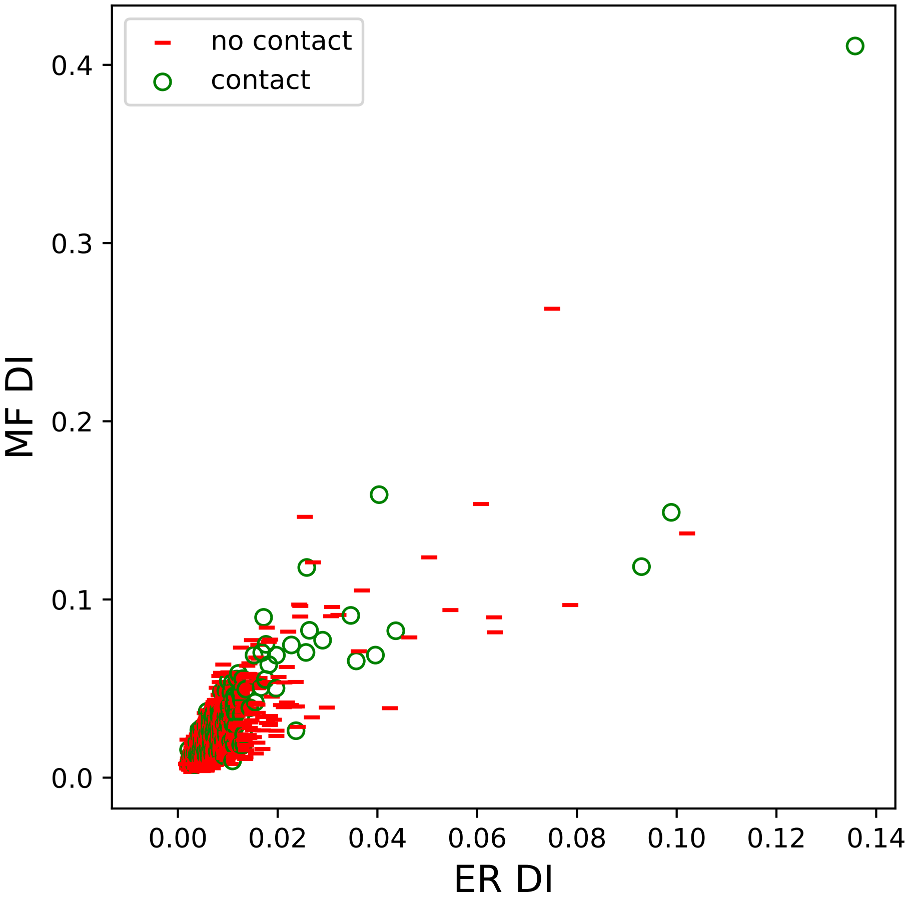

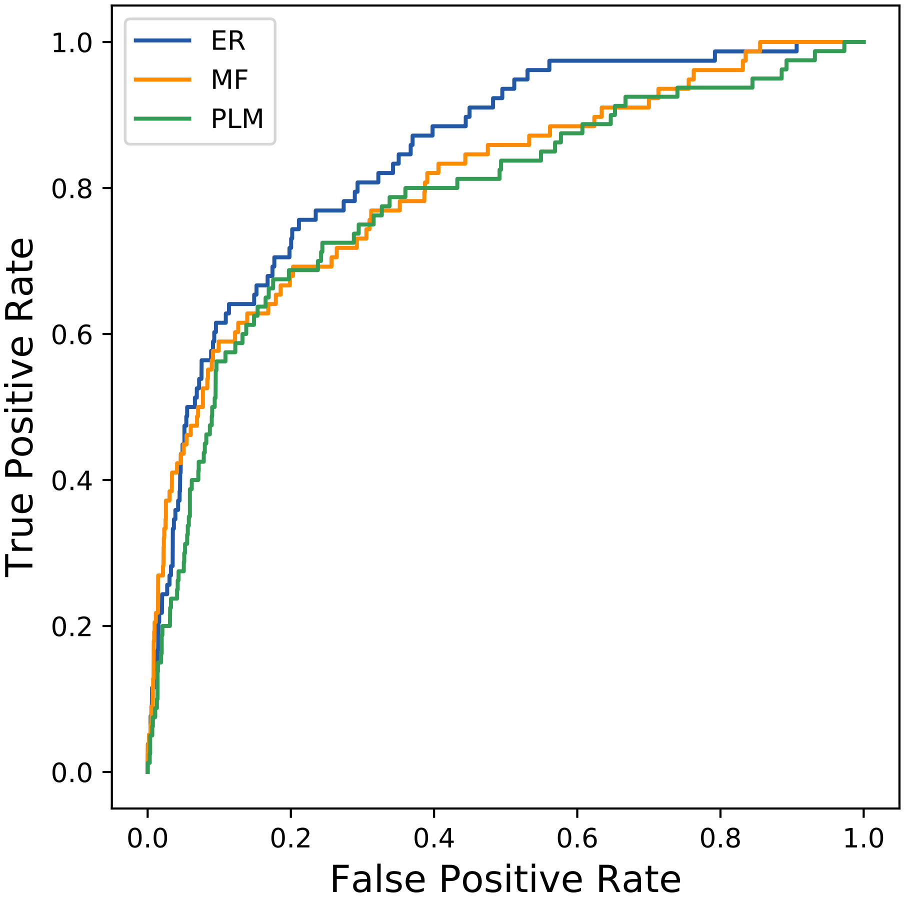

### 3873seq_61col_1jck_PF07654_plots.pdf

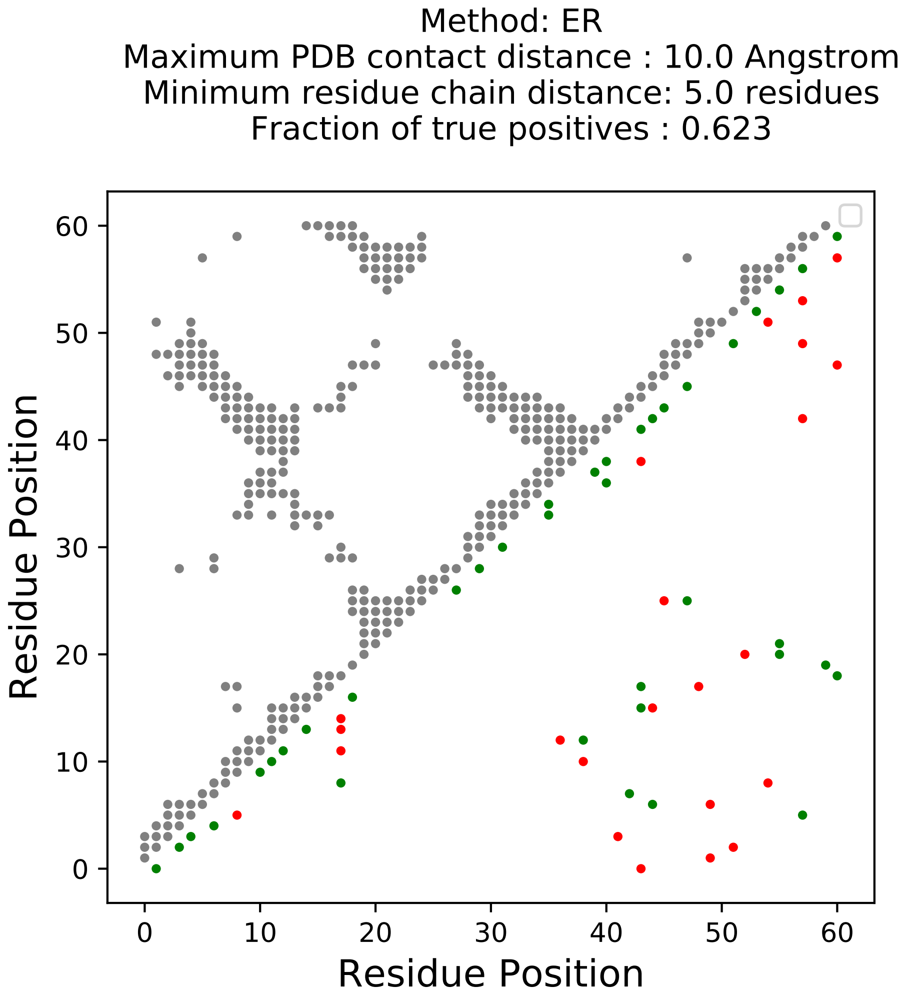

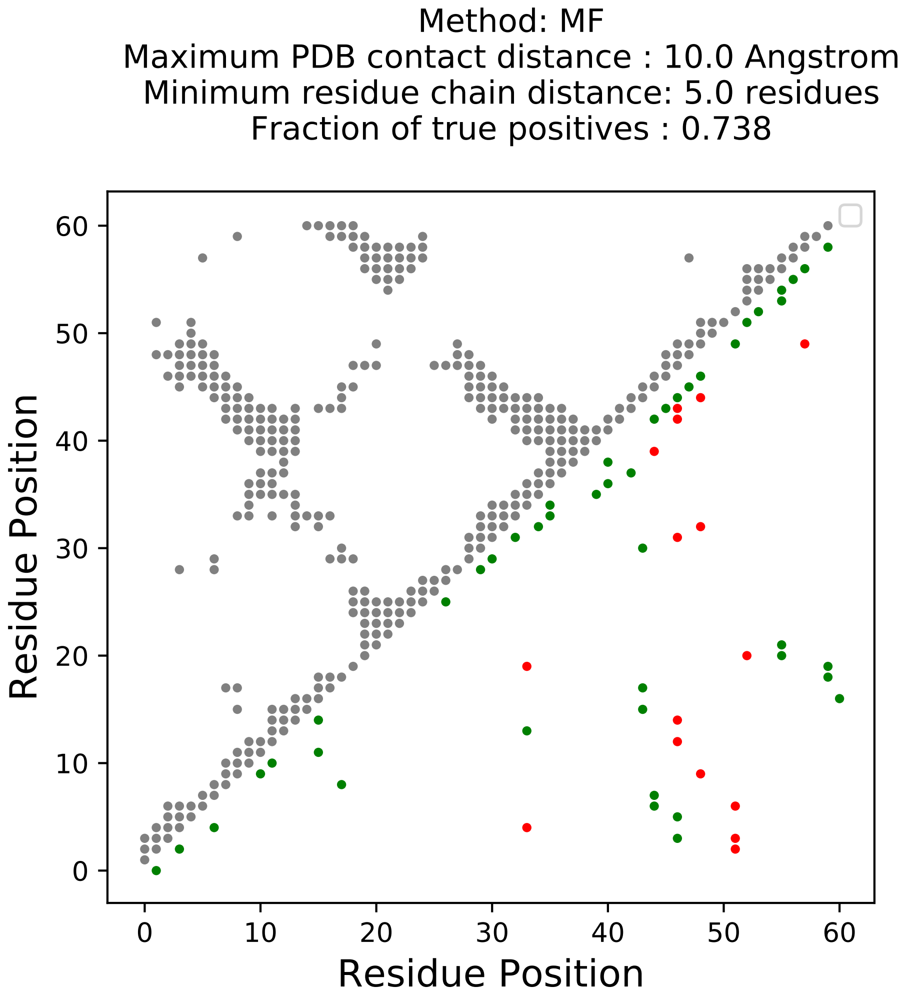

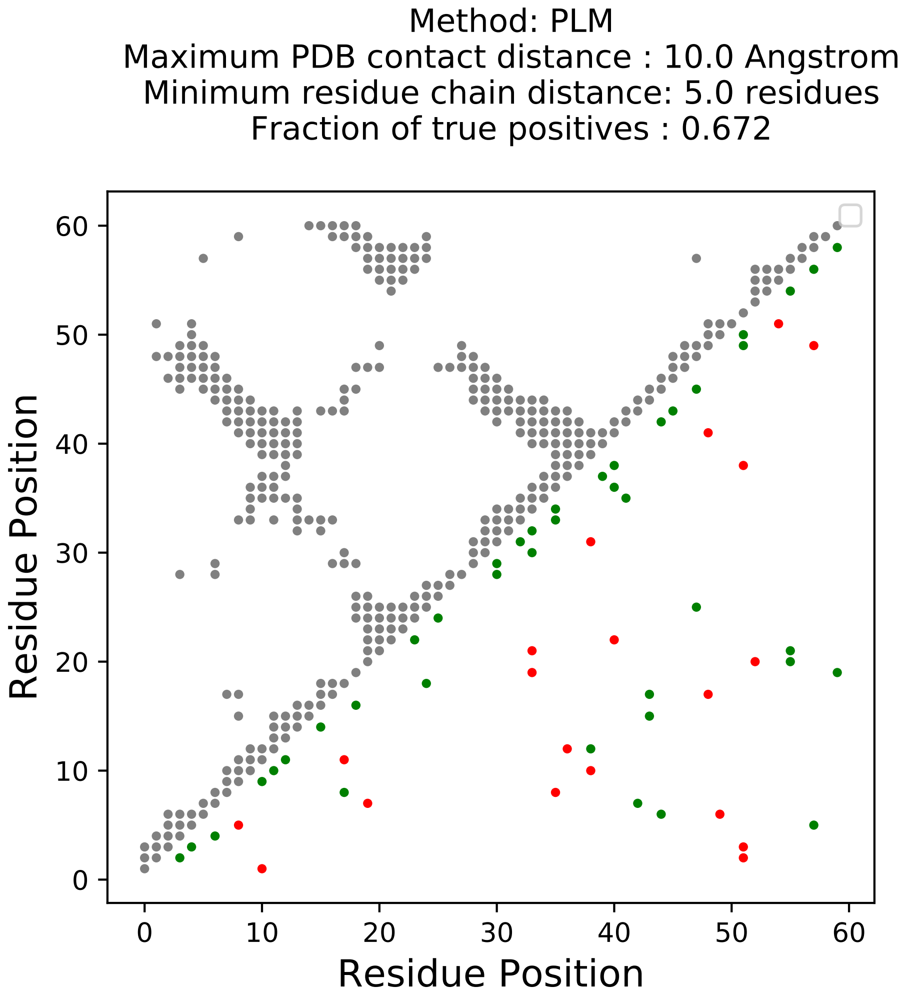

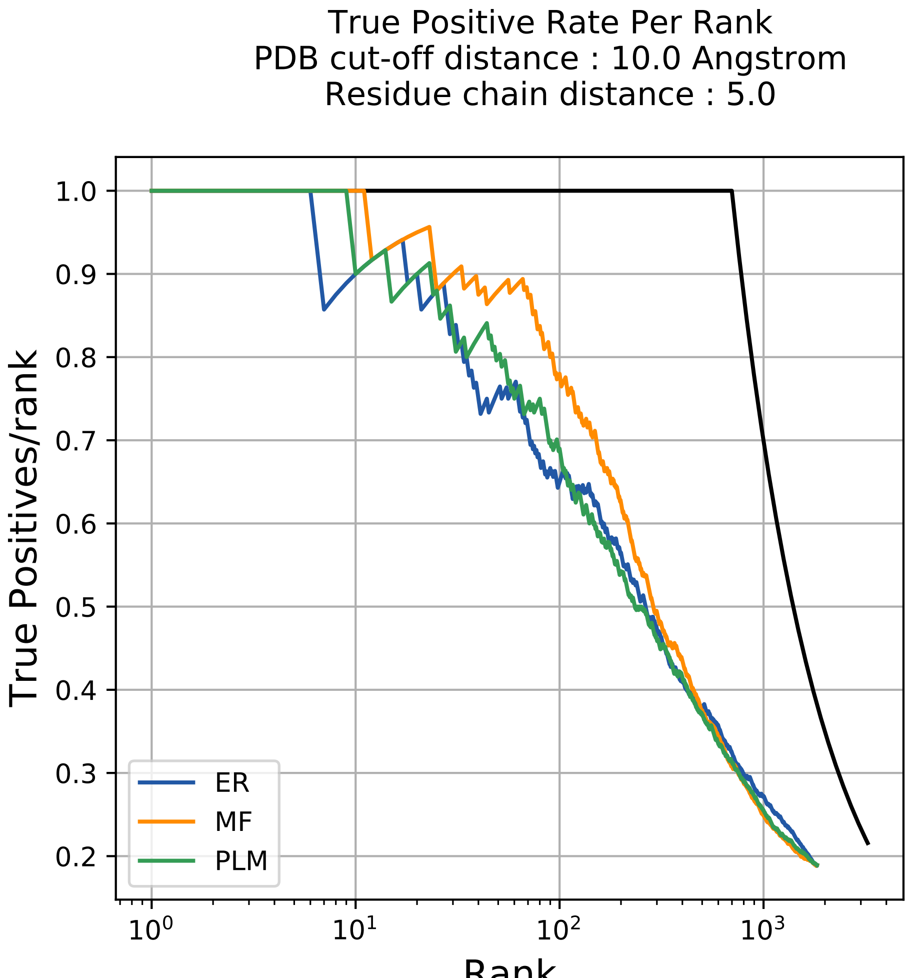

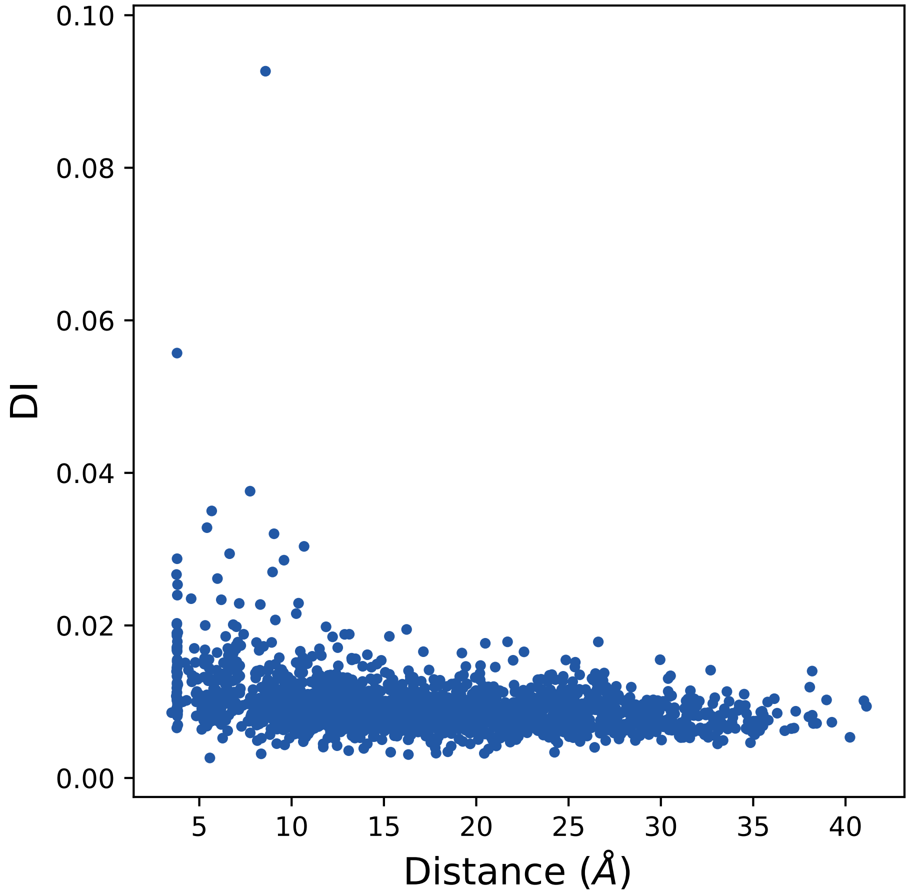

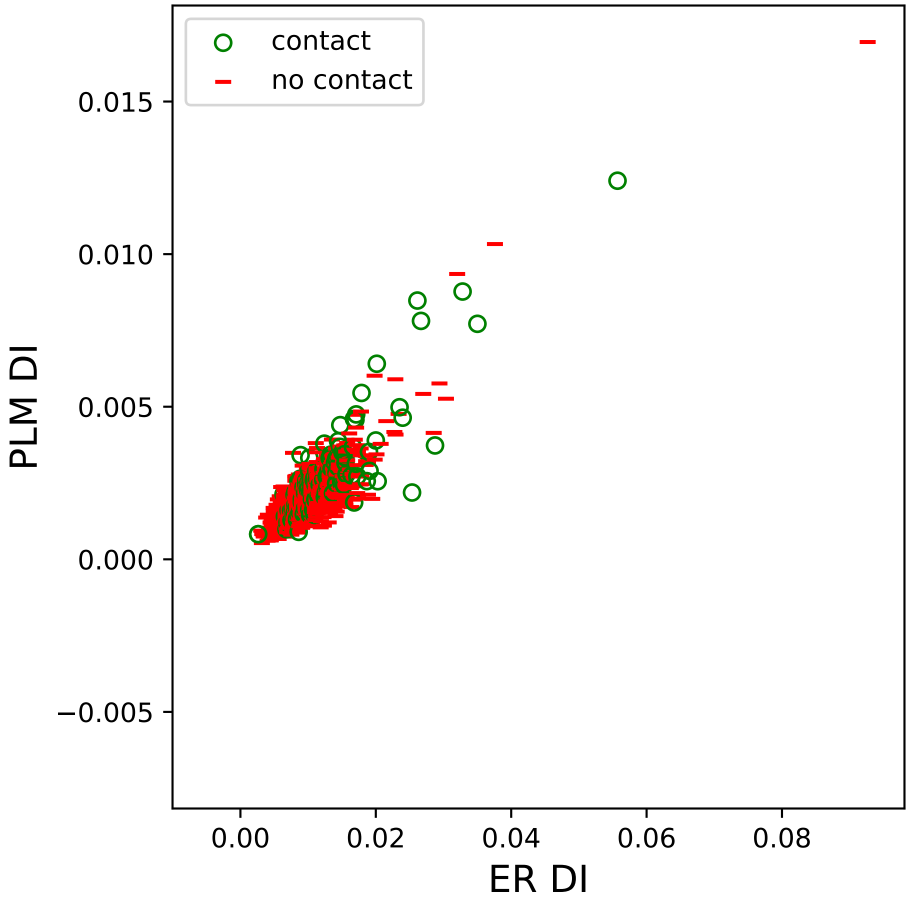

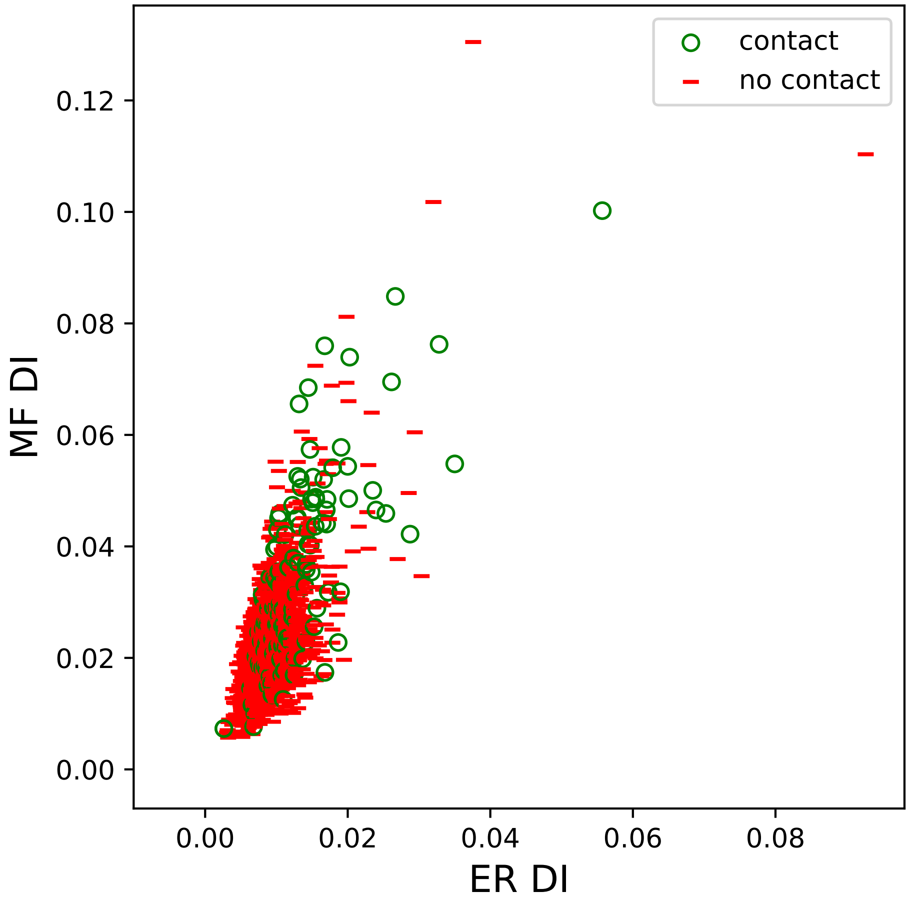

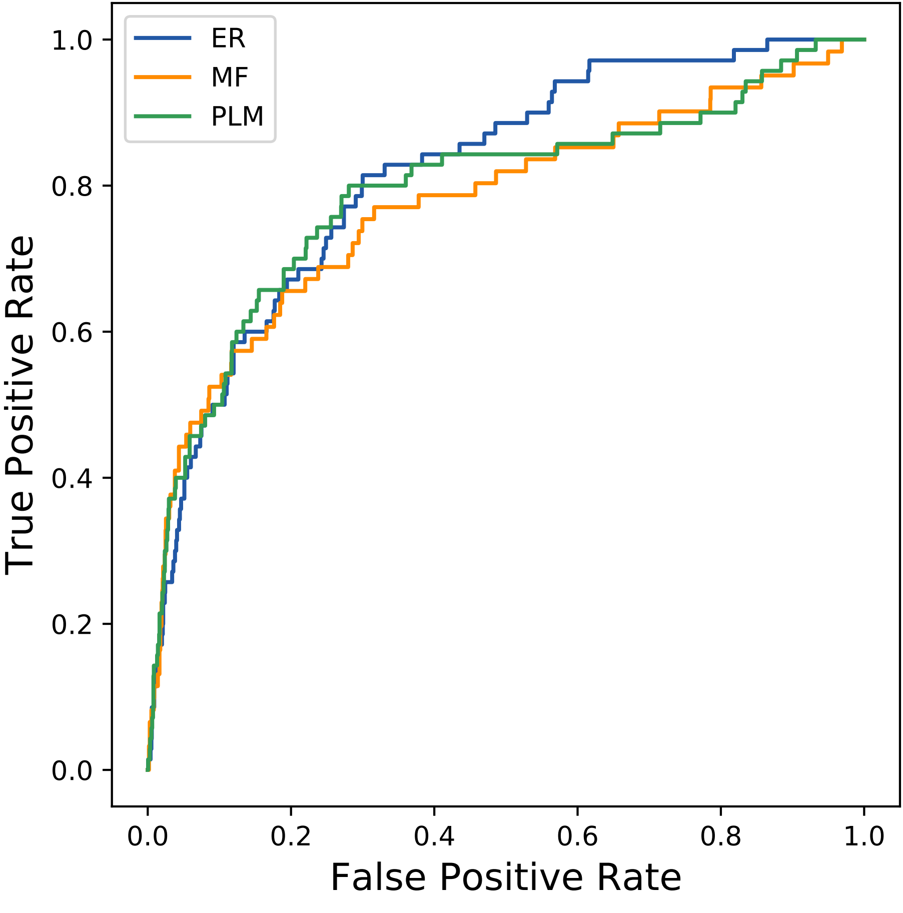

### 4038seq_55col_6bcs_PF13895_plots.pdf

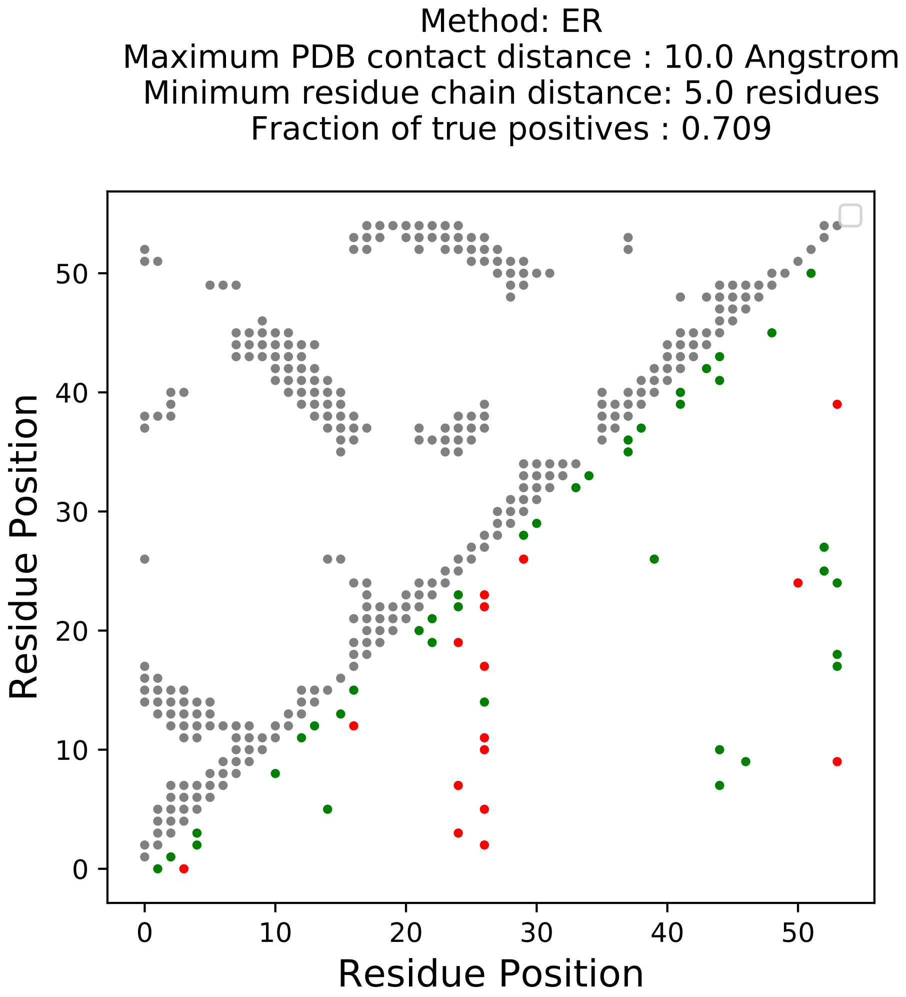

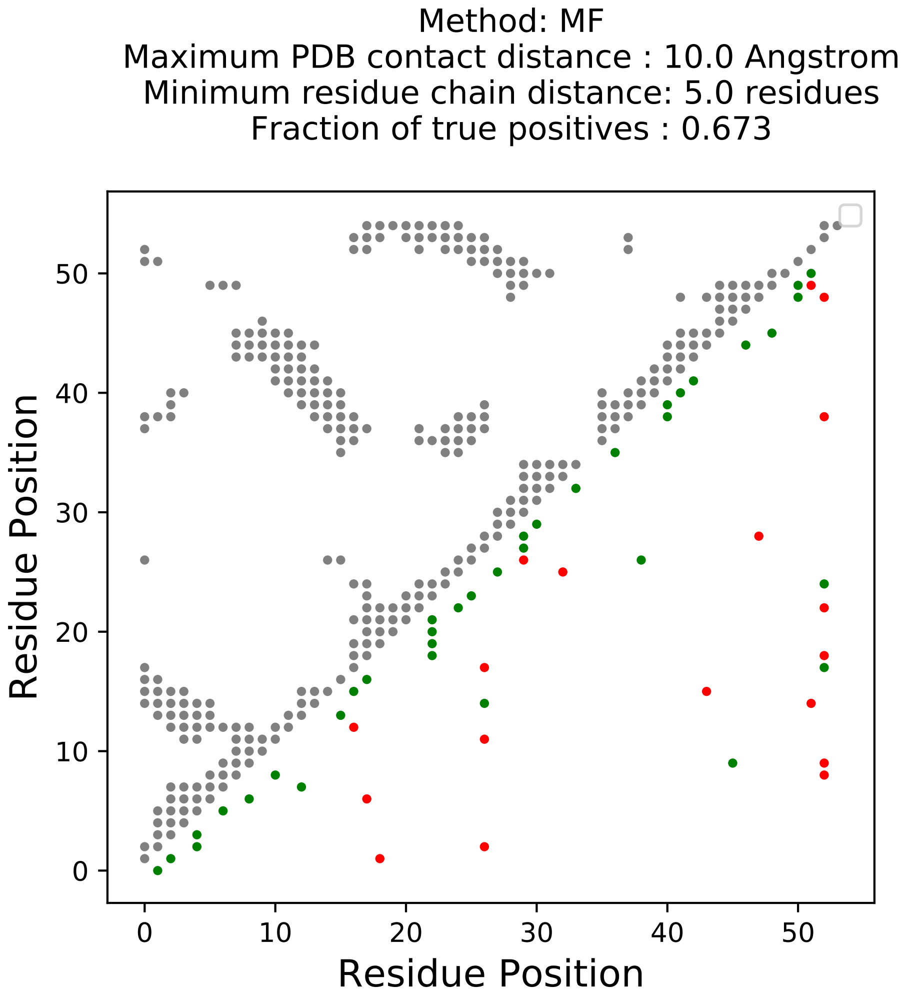

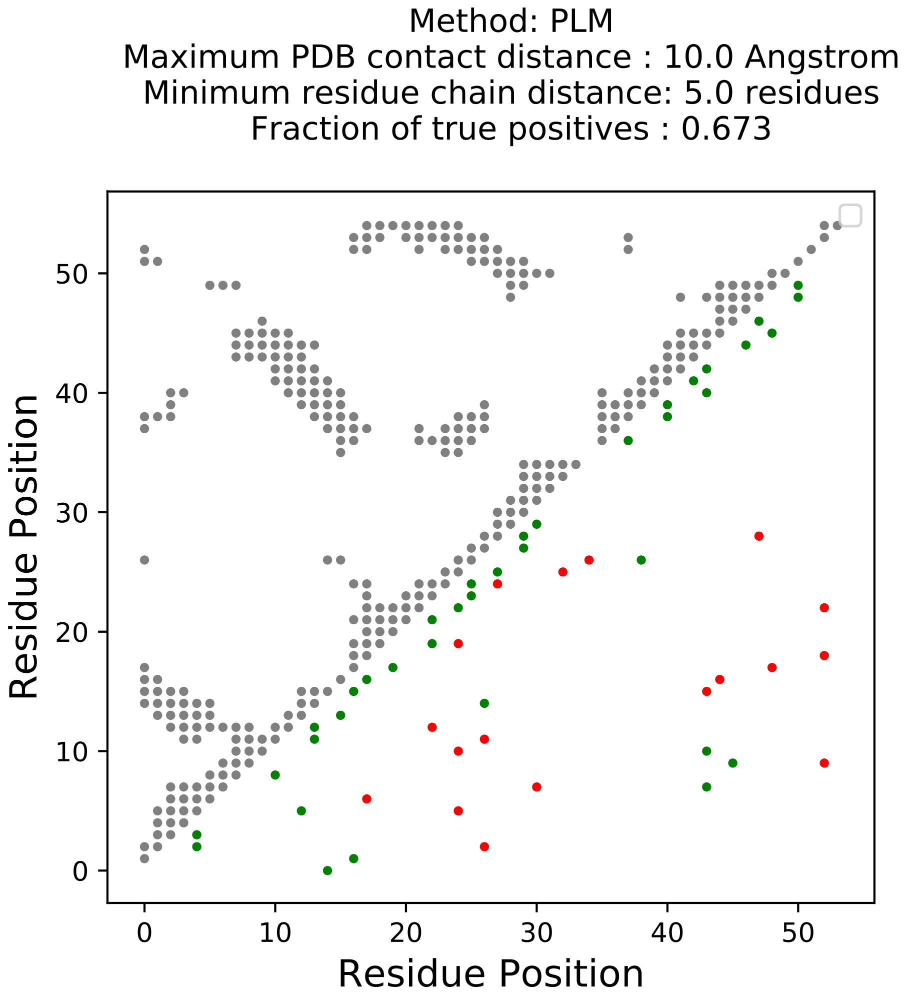

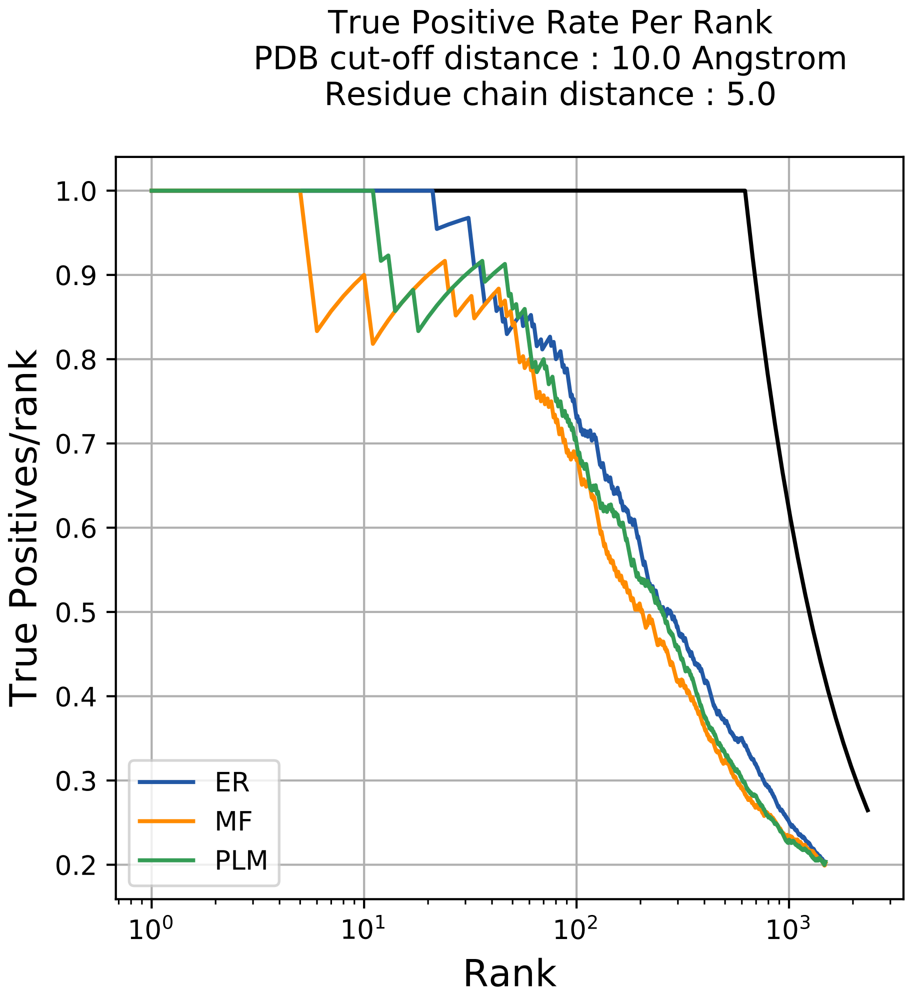

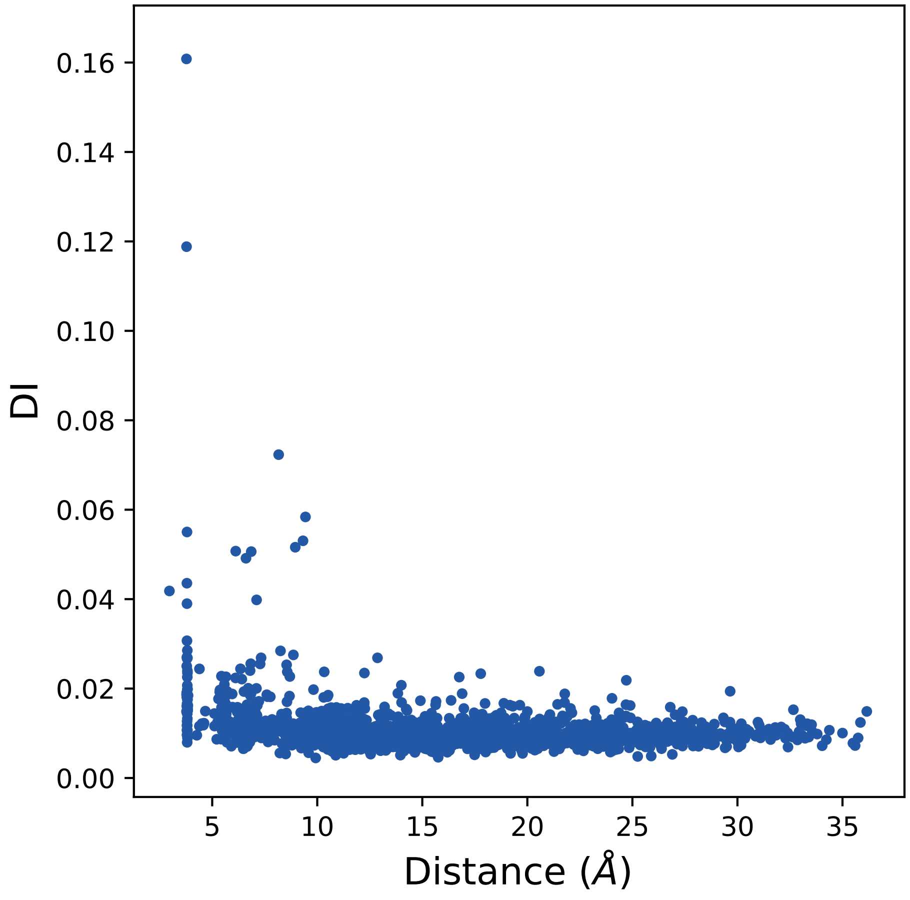

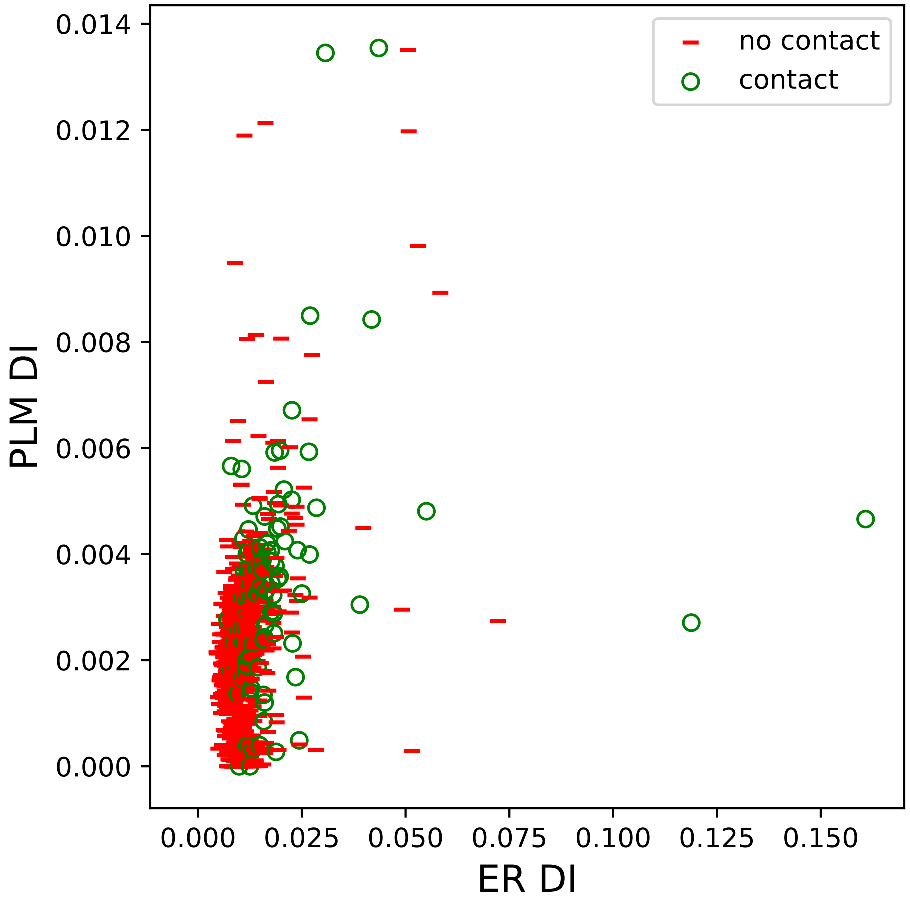

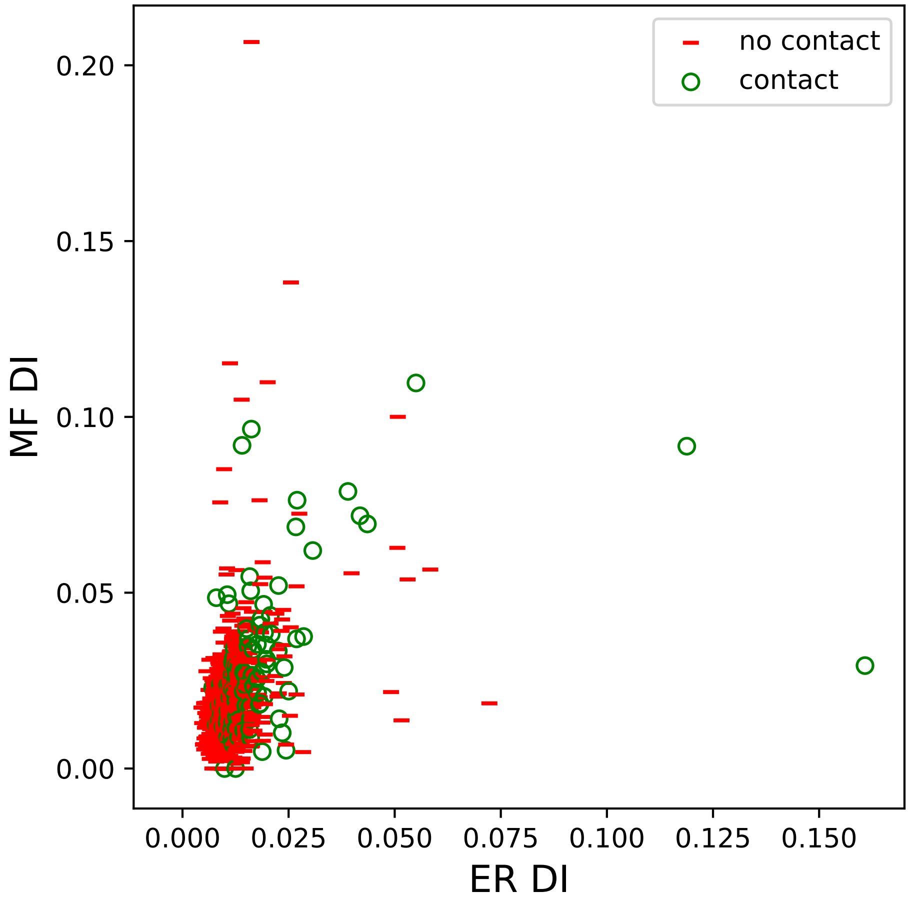

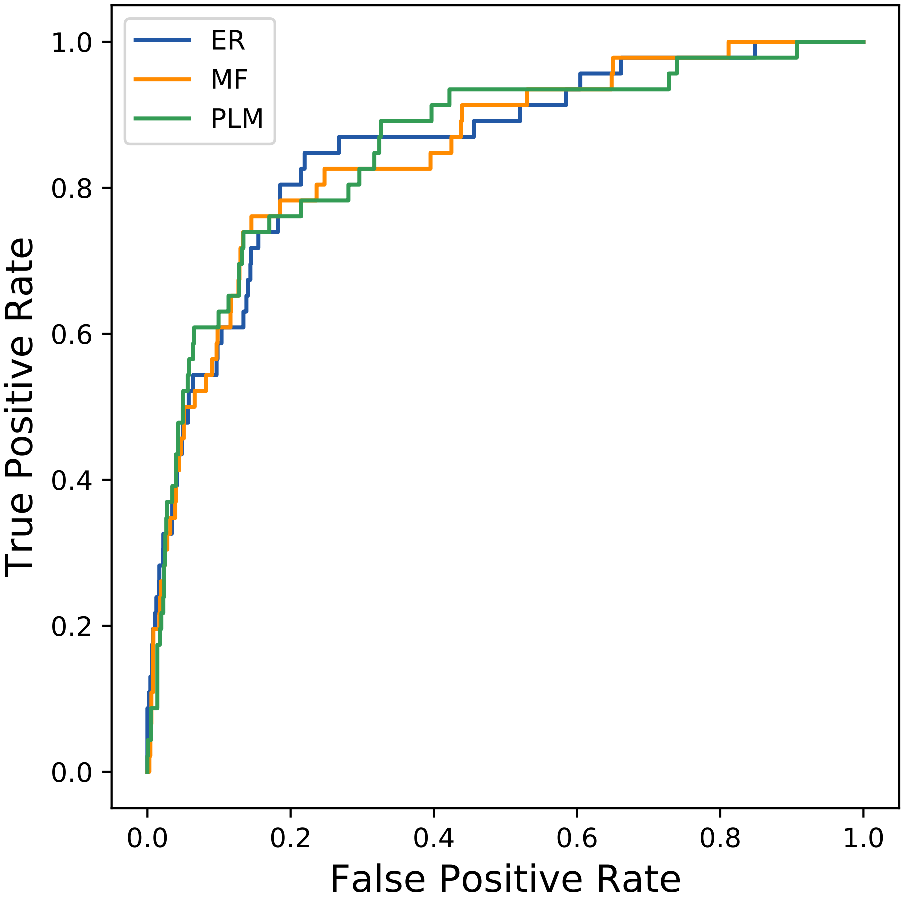

### 4168seq_31col_1b2j_PF00301_plots.pdf

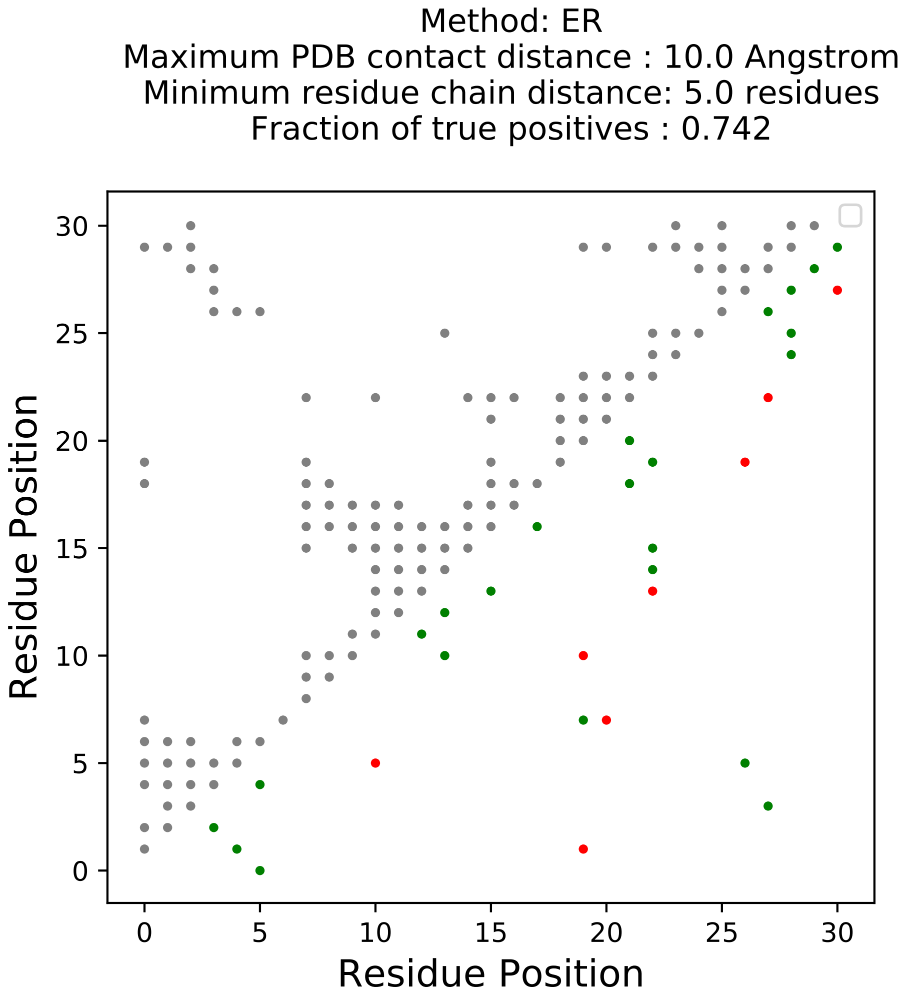

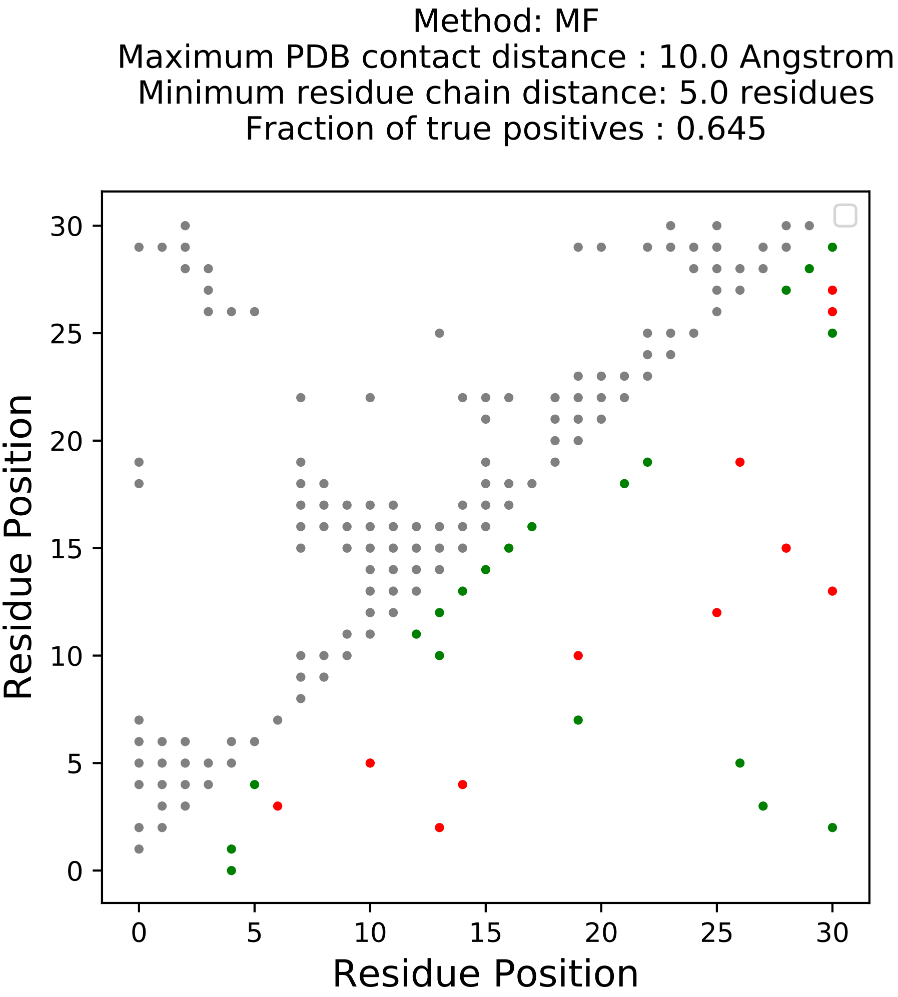

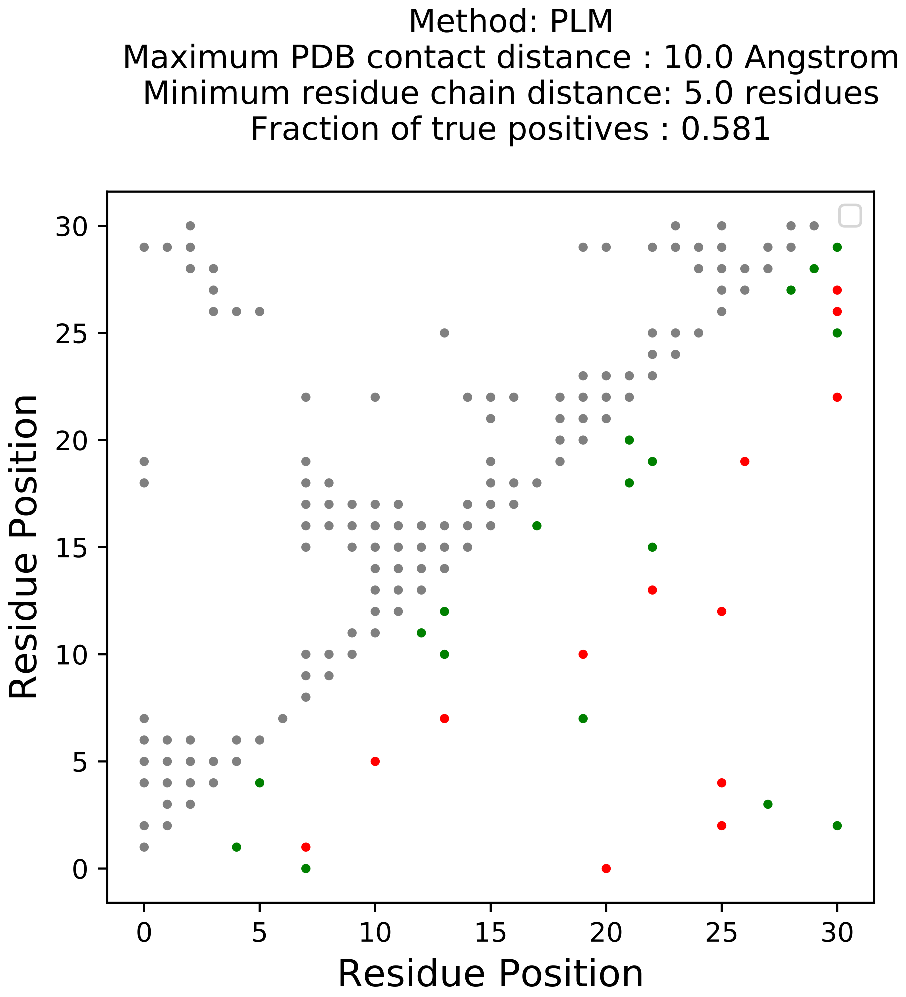

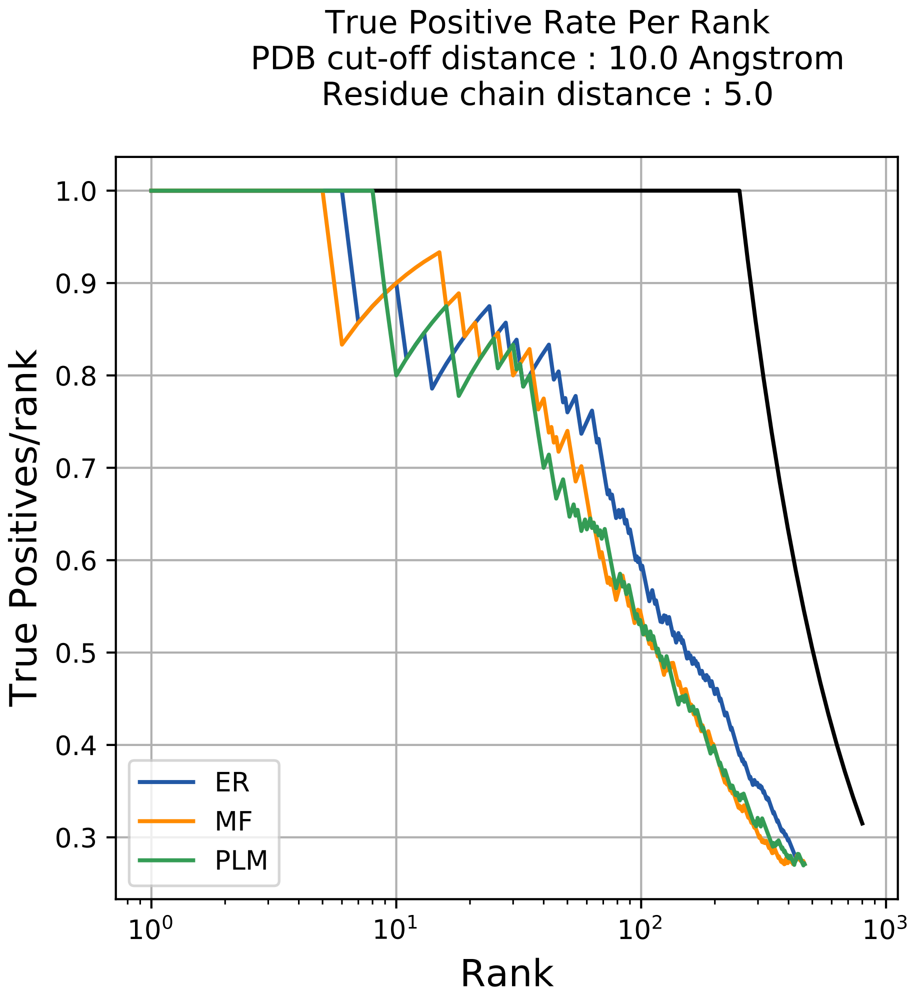

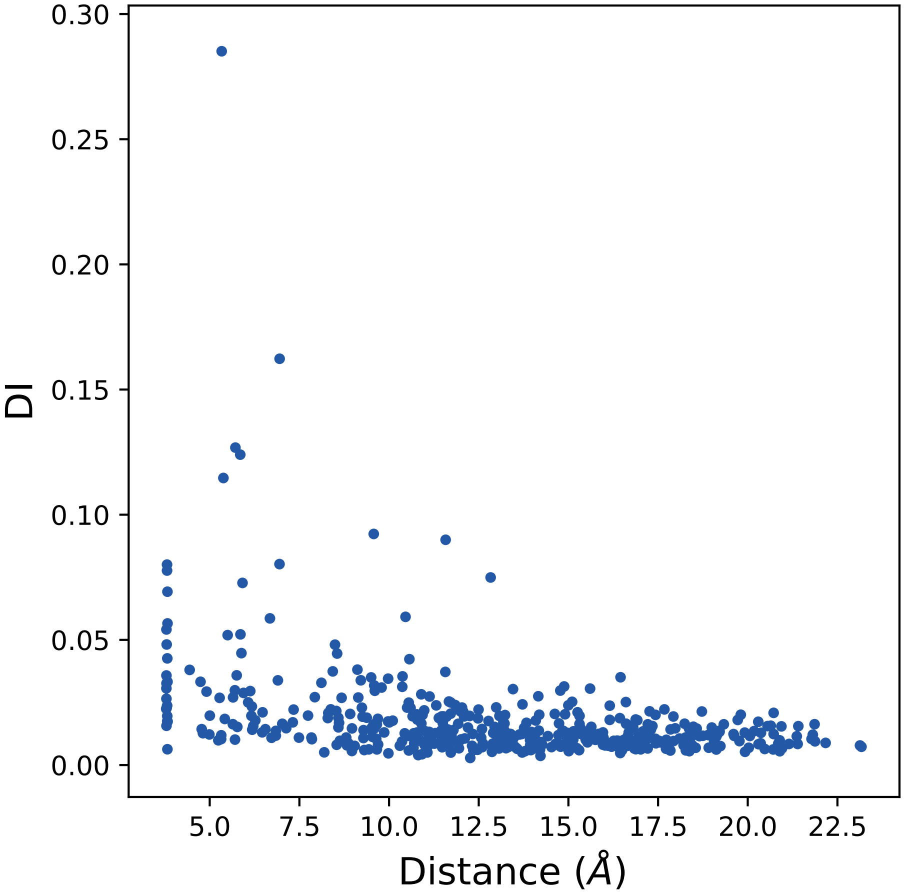

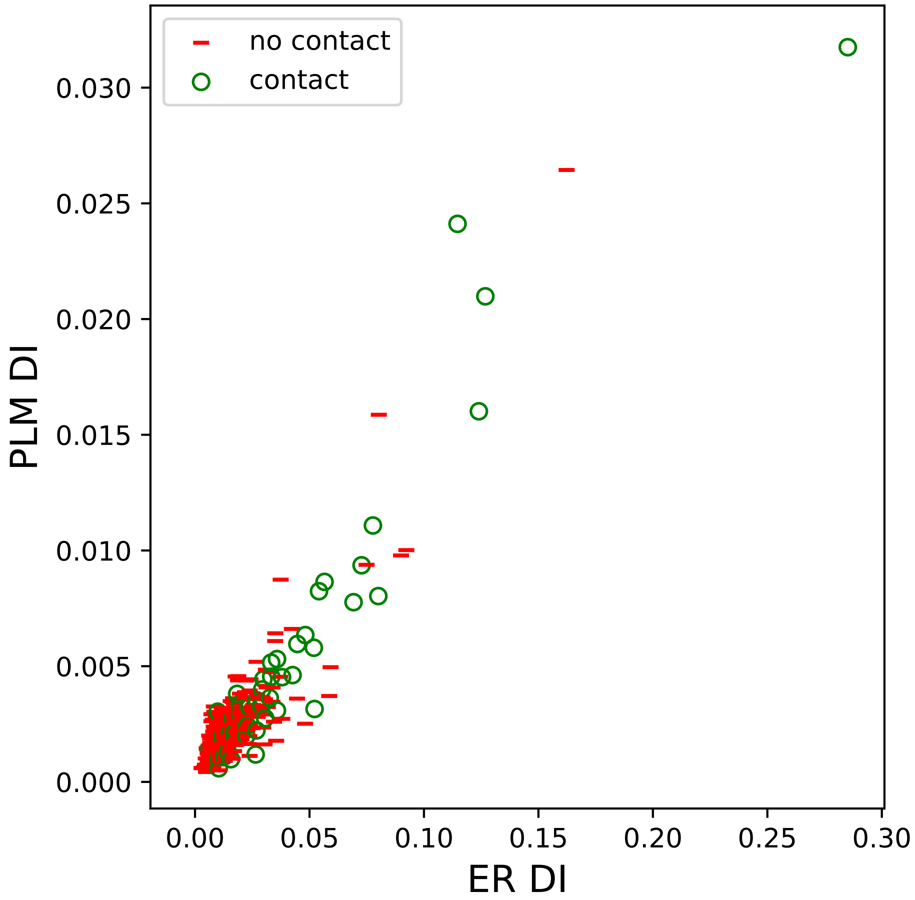

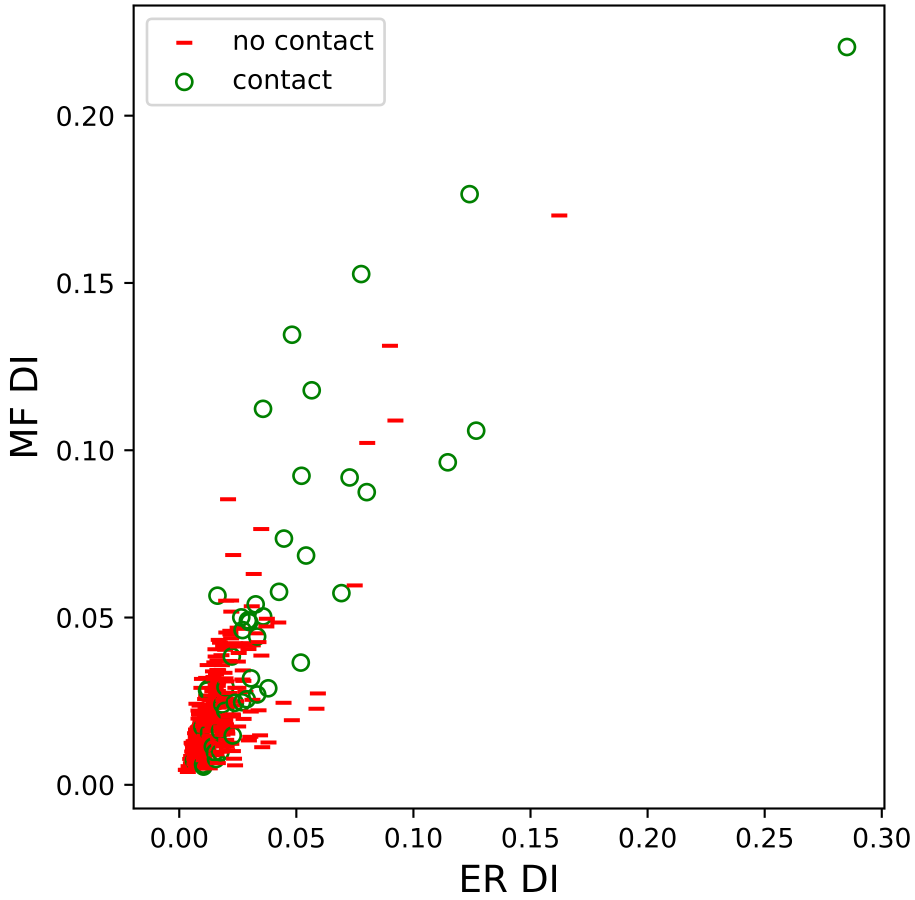

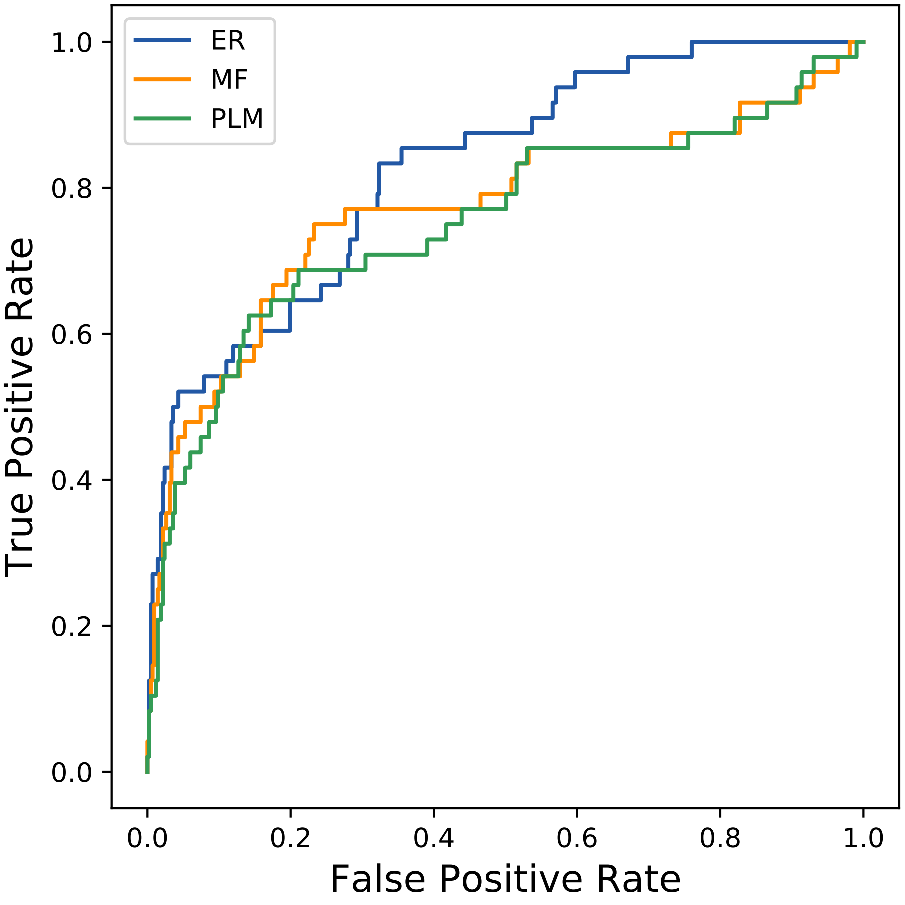

### 4348seq_100col_5pmf_PF02373_plots.pdf

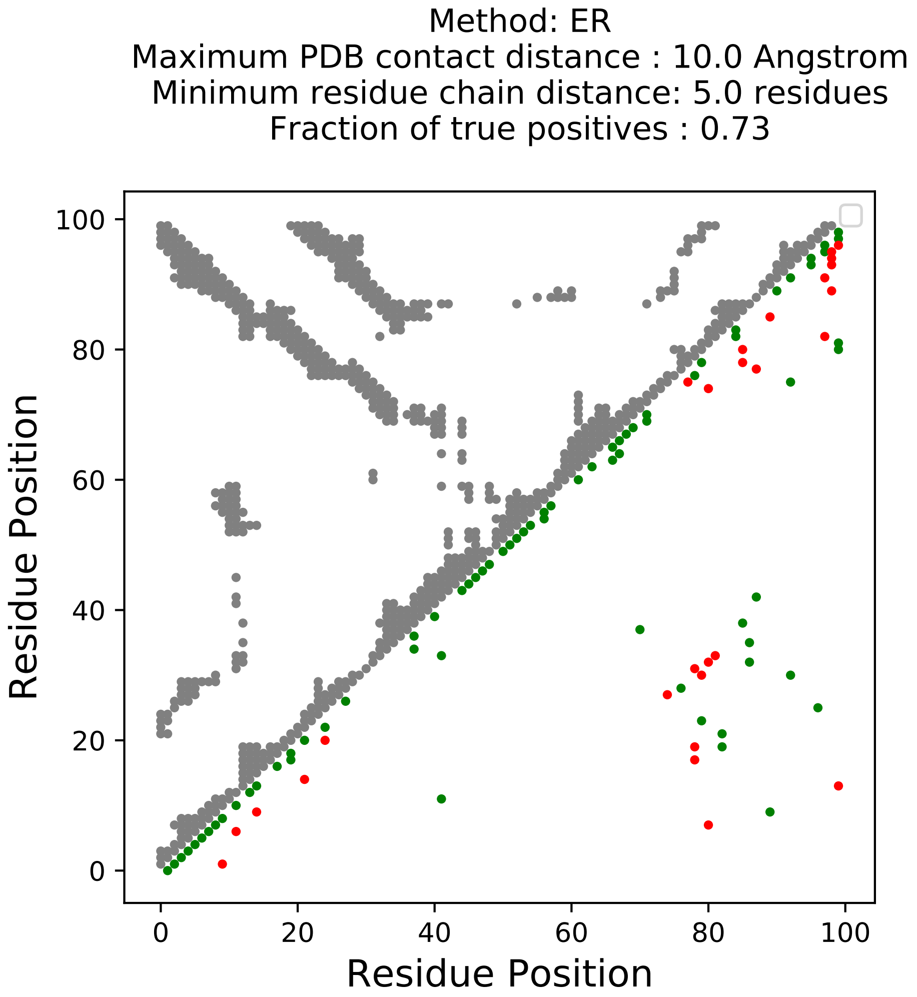

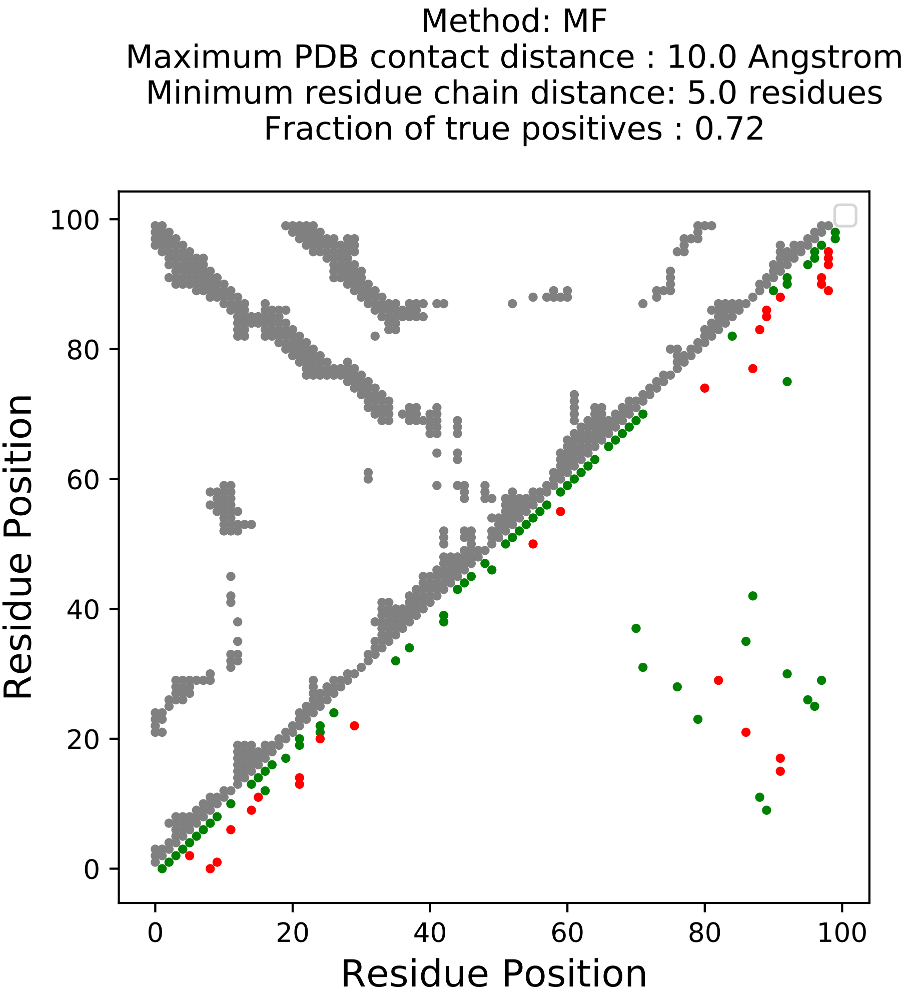

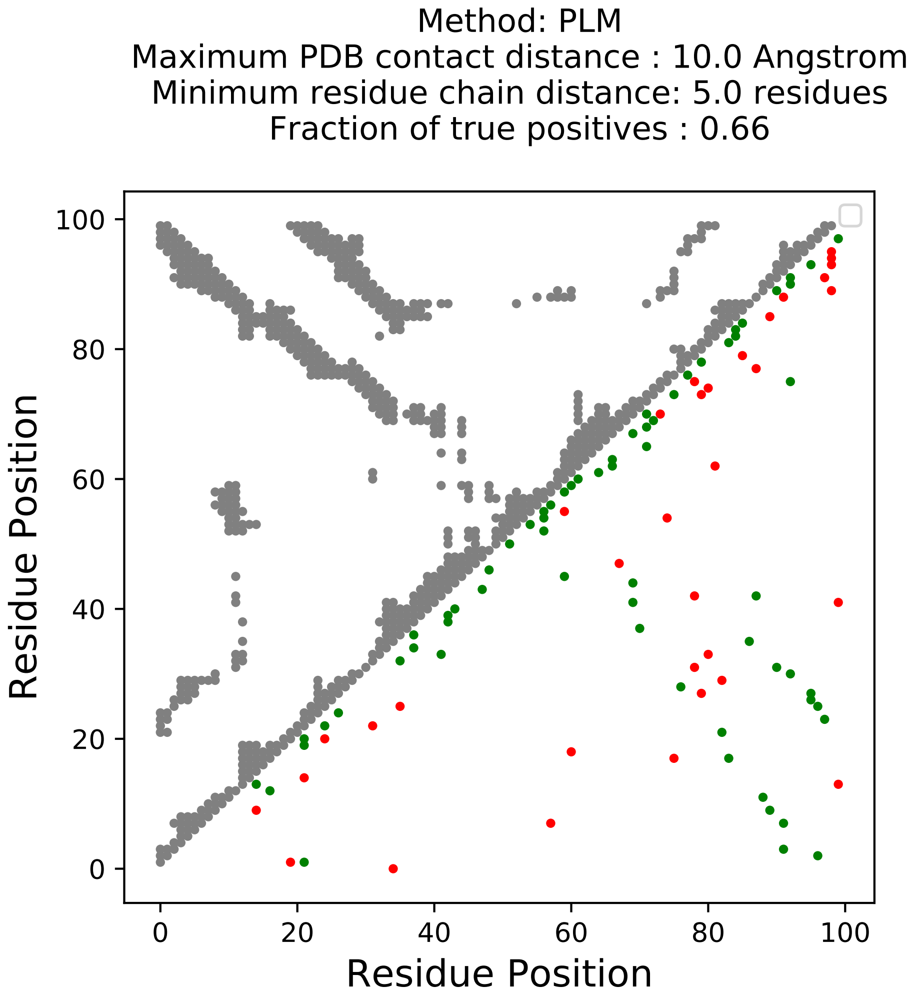

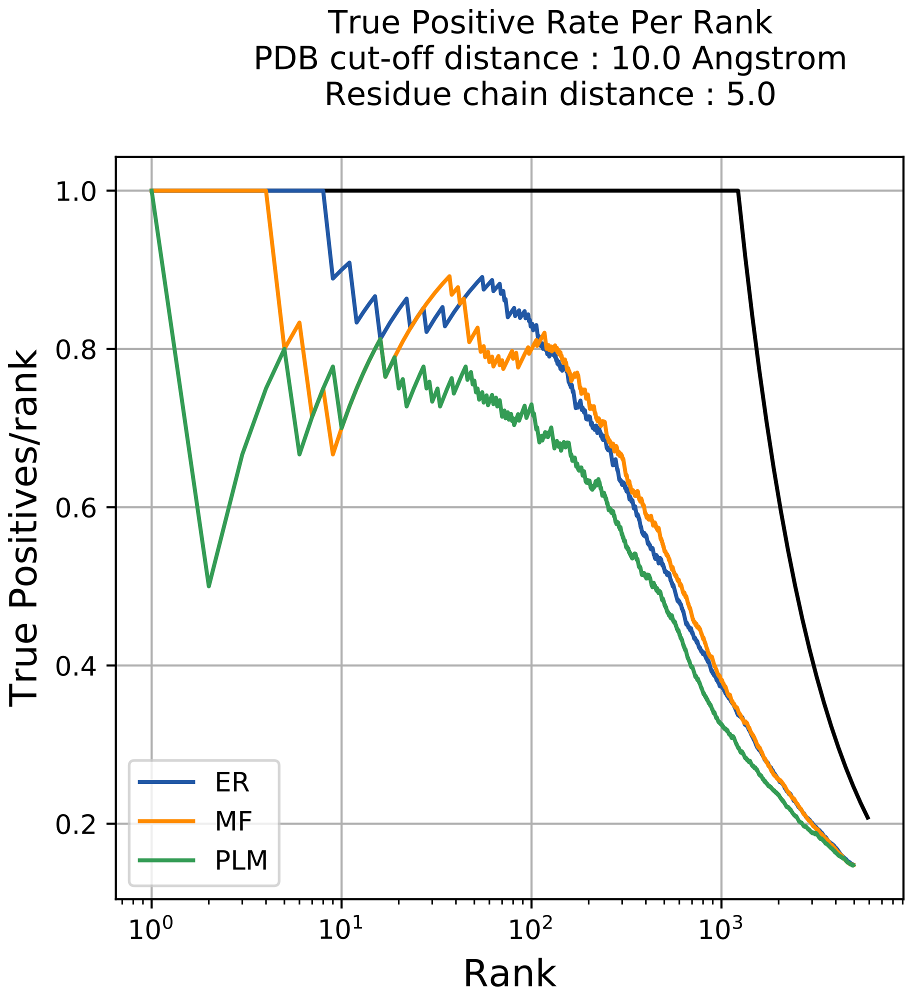

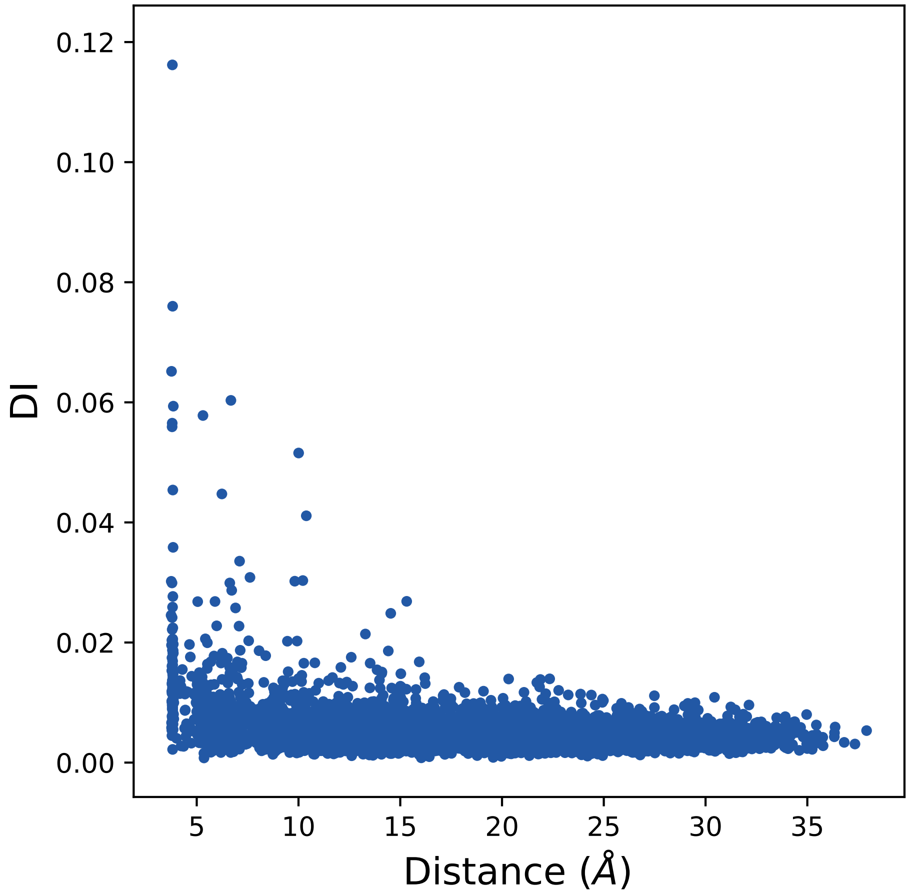

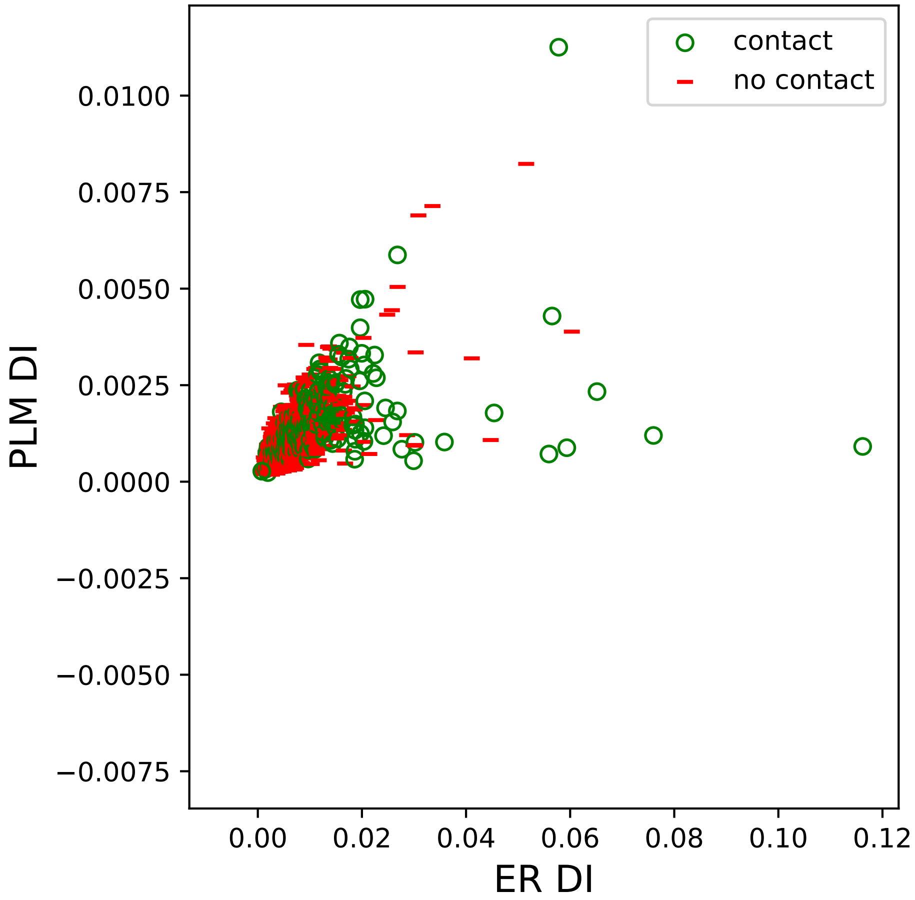

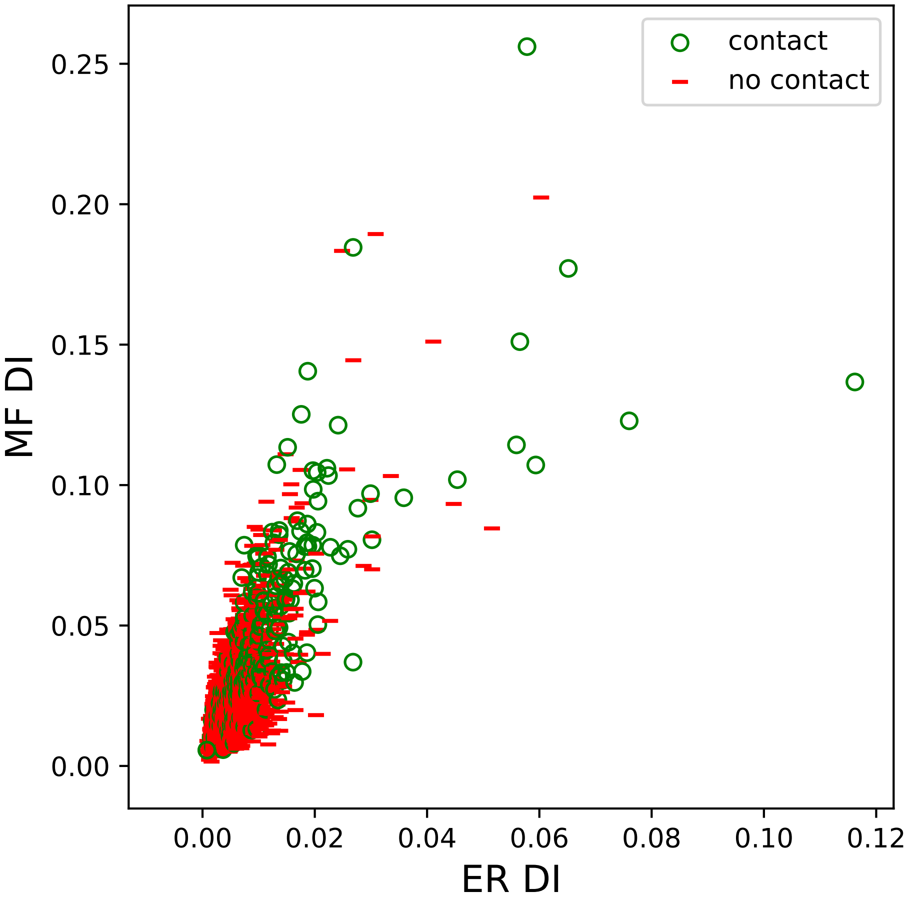

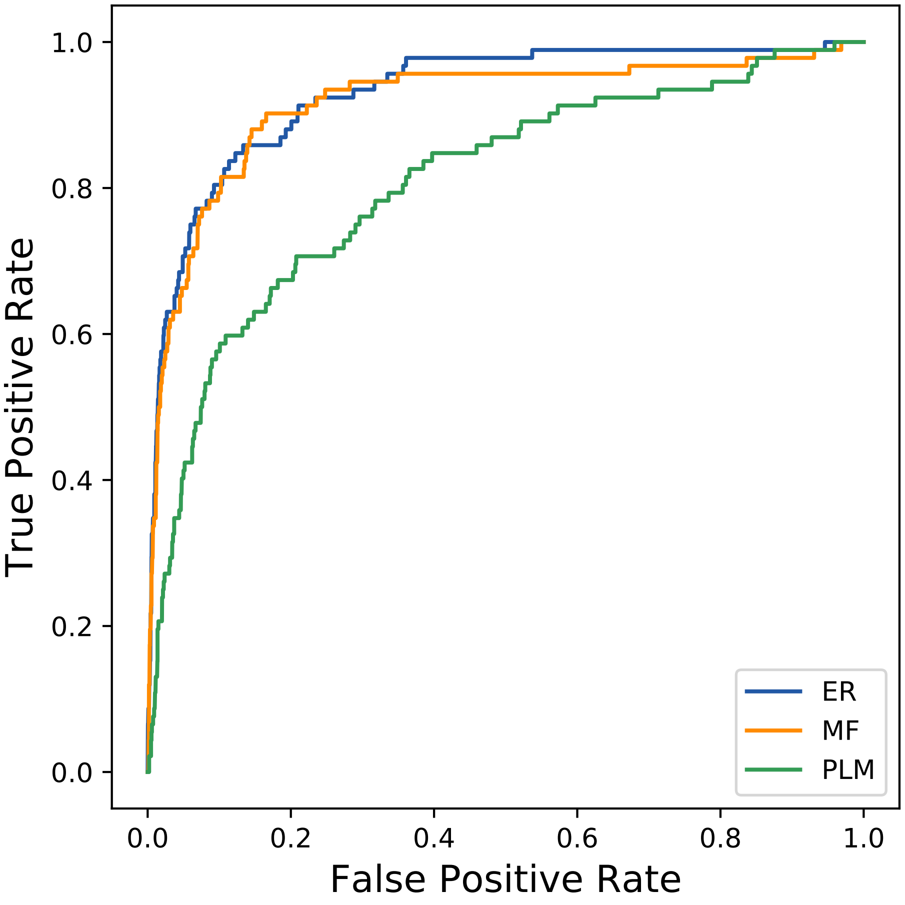

### 4389seq_62col_2ak1_PF07654_plots.pdf

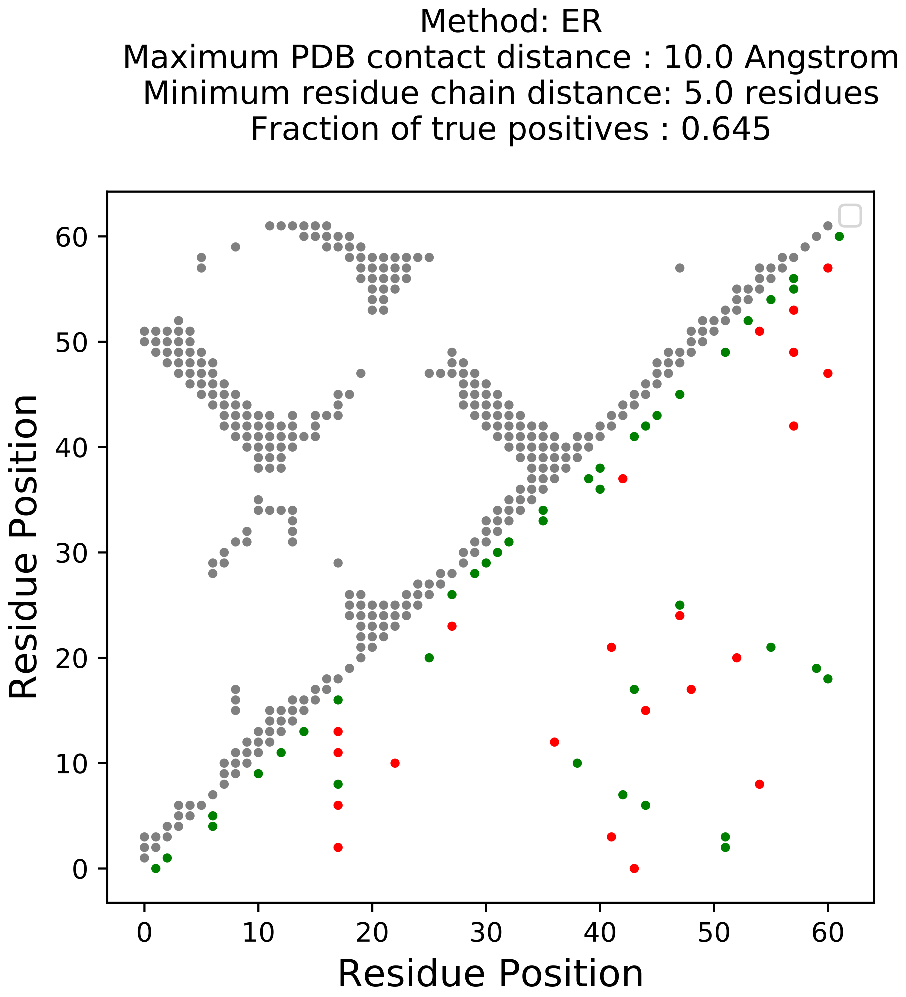

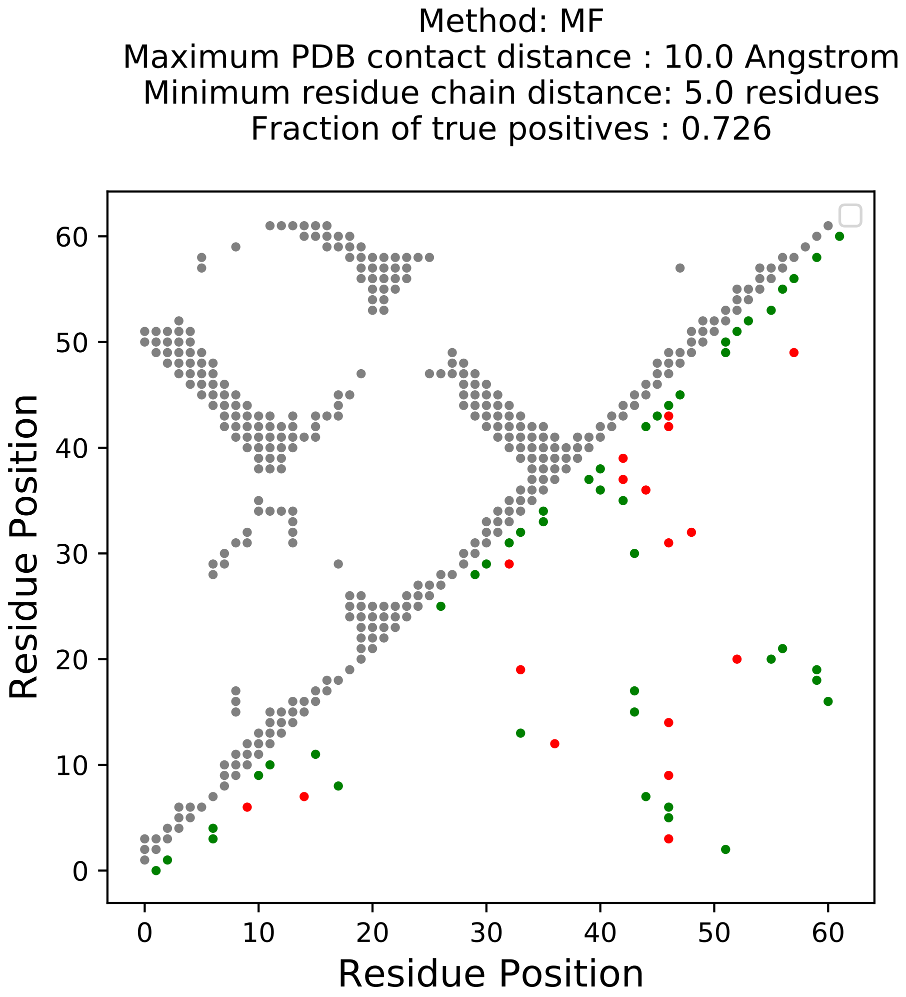

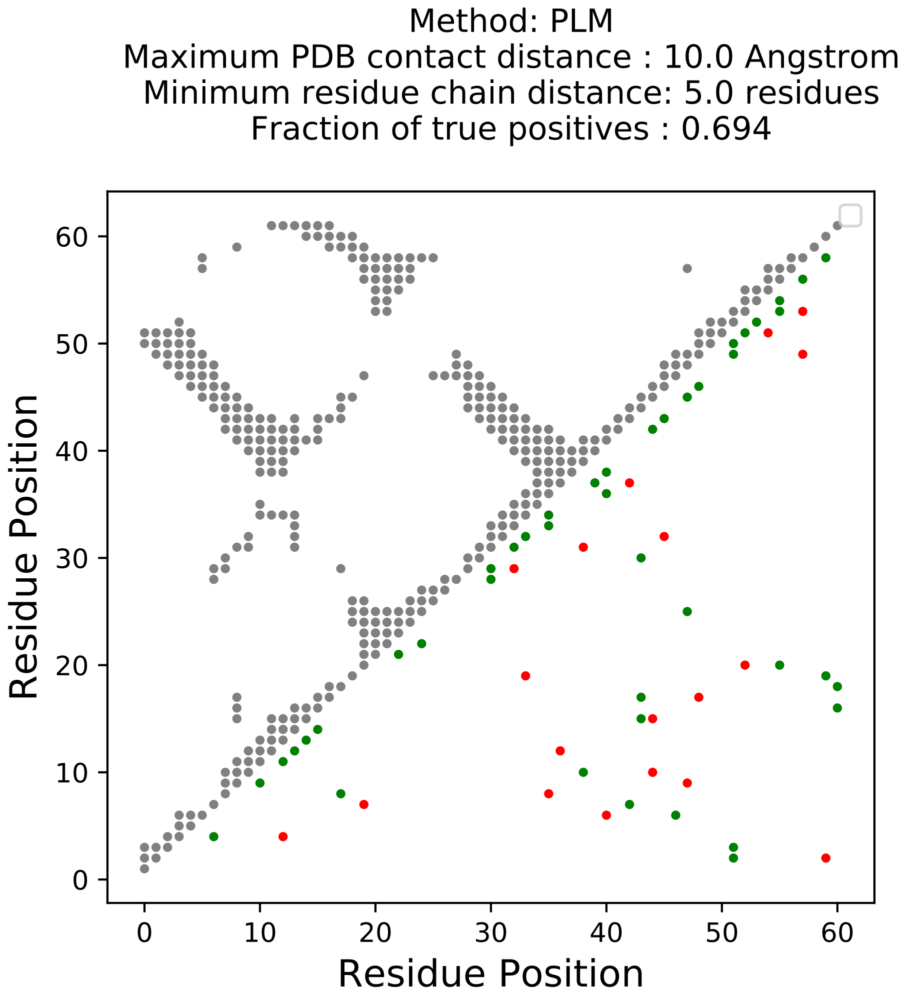

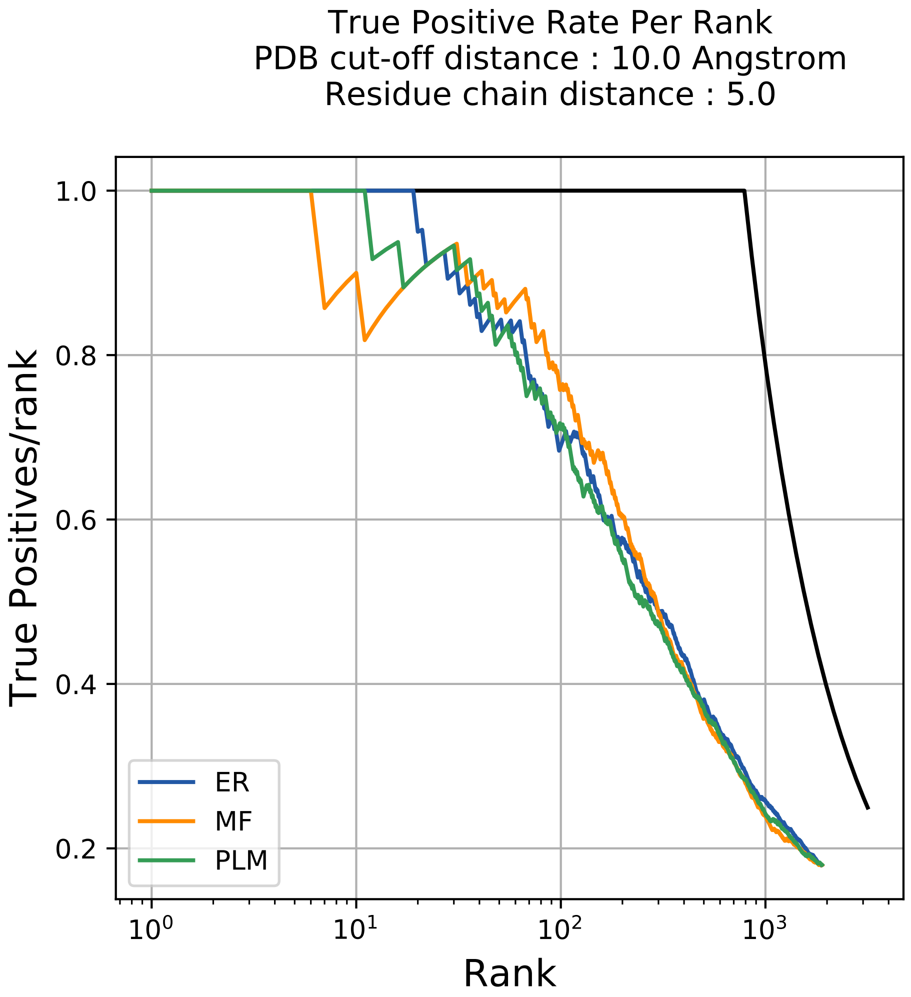

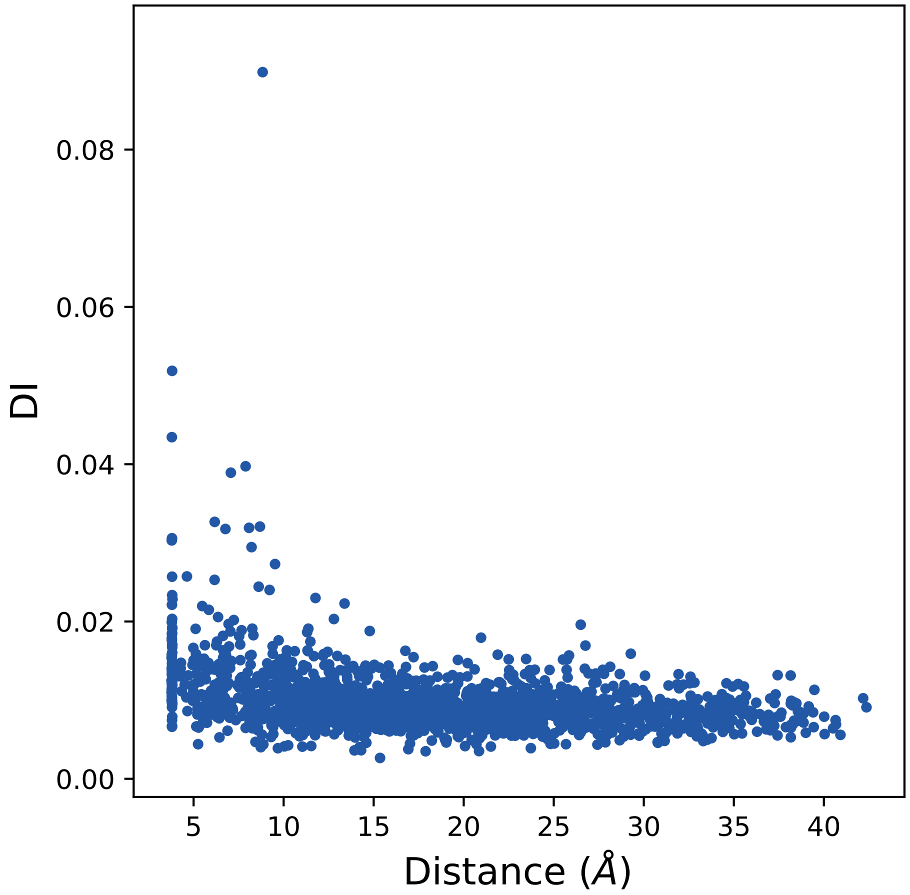

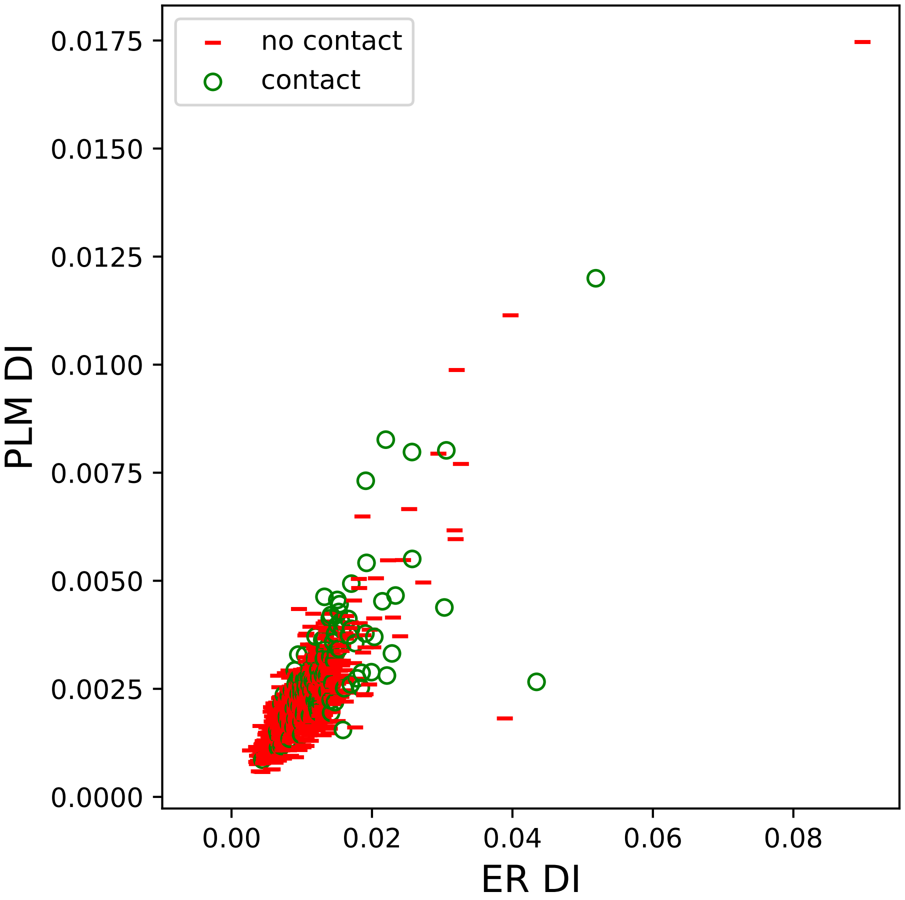

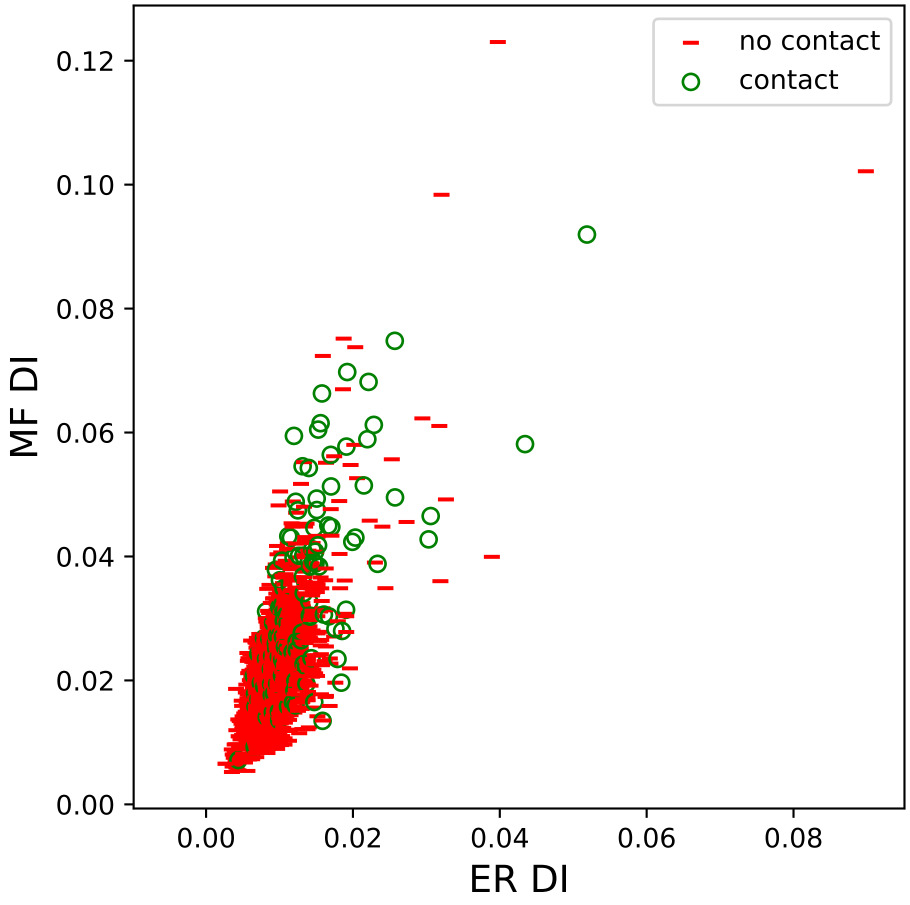

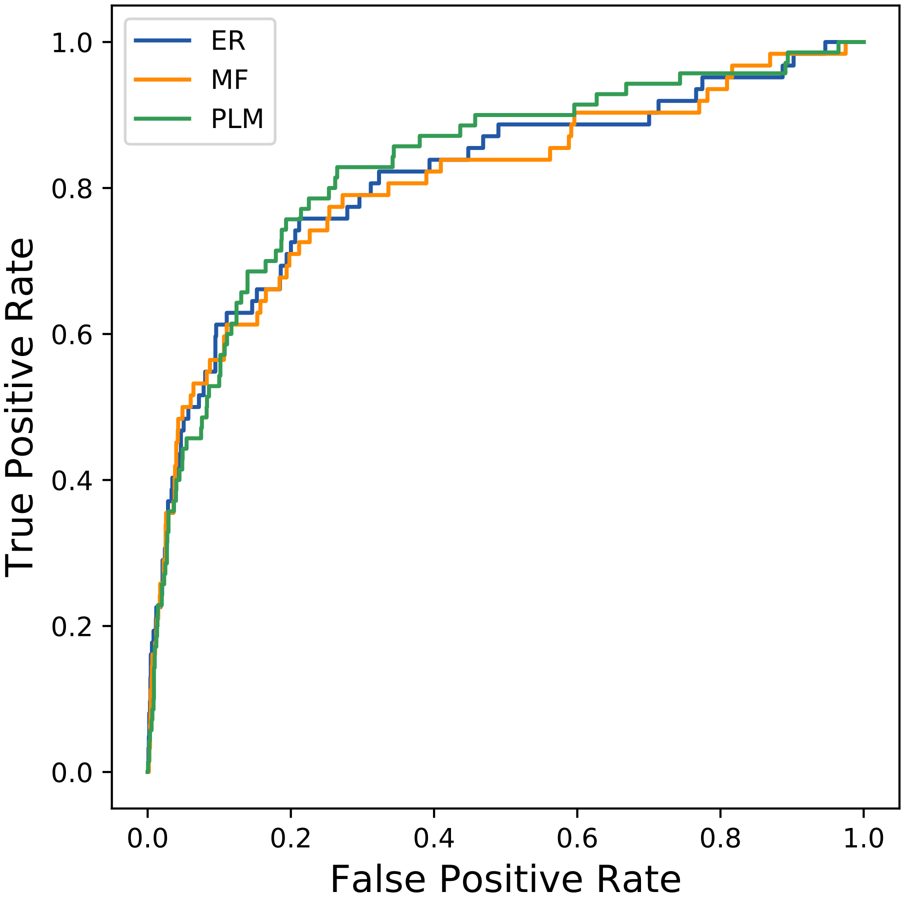

### 4766seq_182col_3ibl_PF00194_plots.pdf

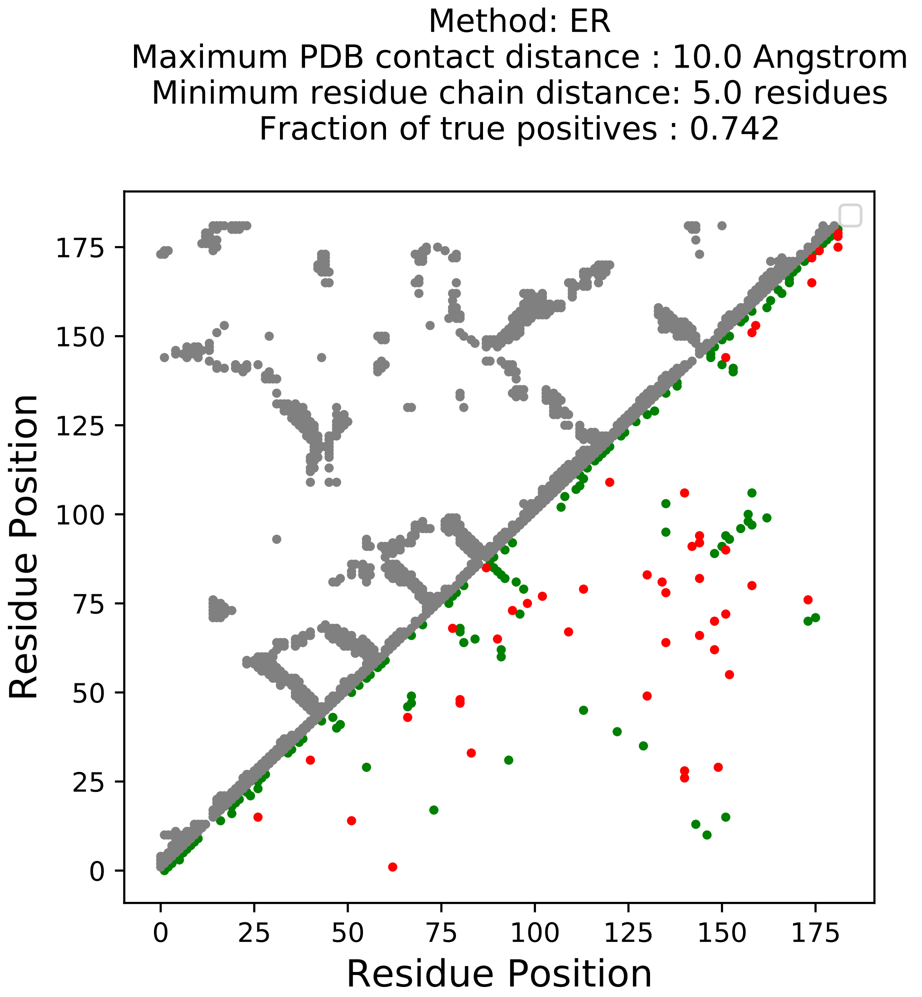

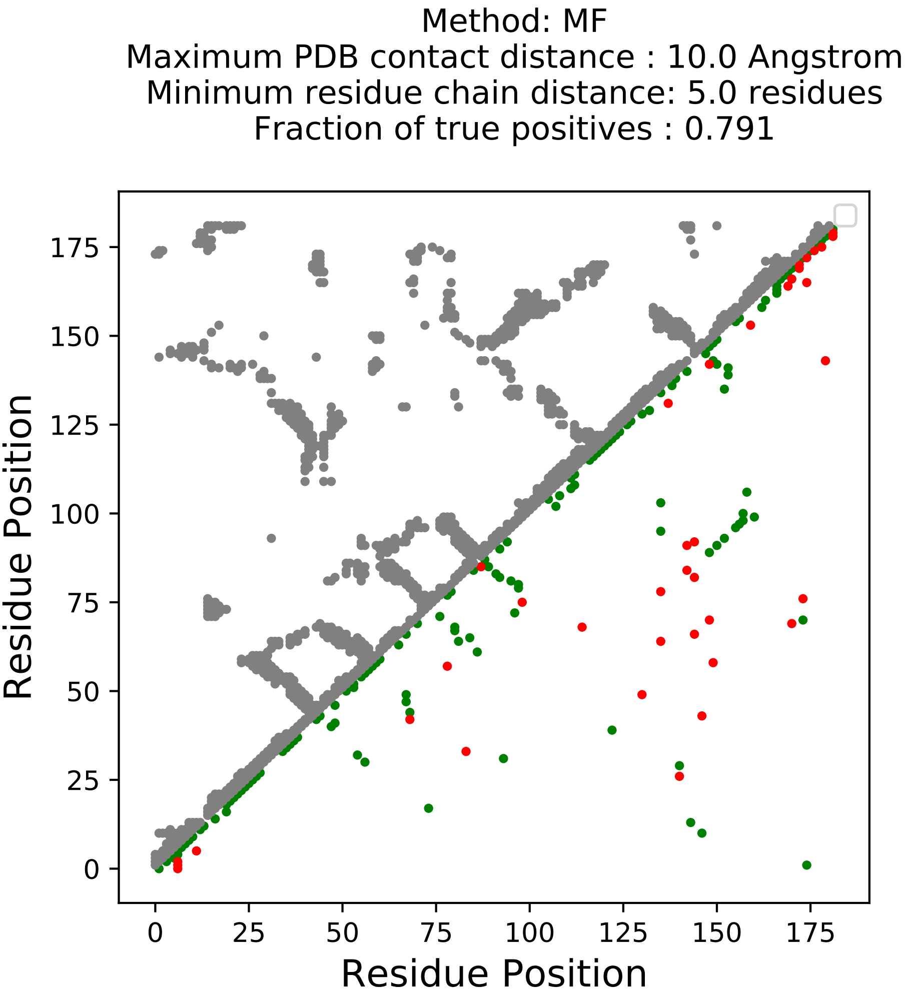

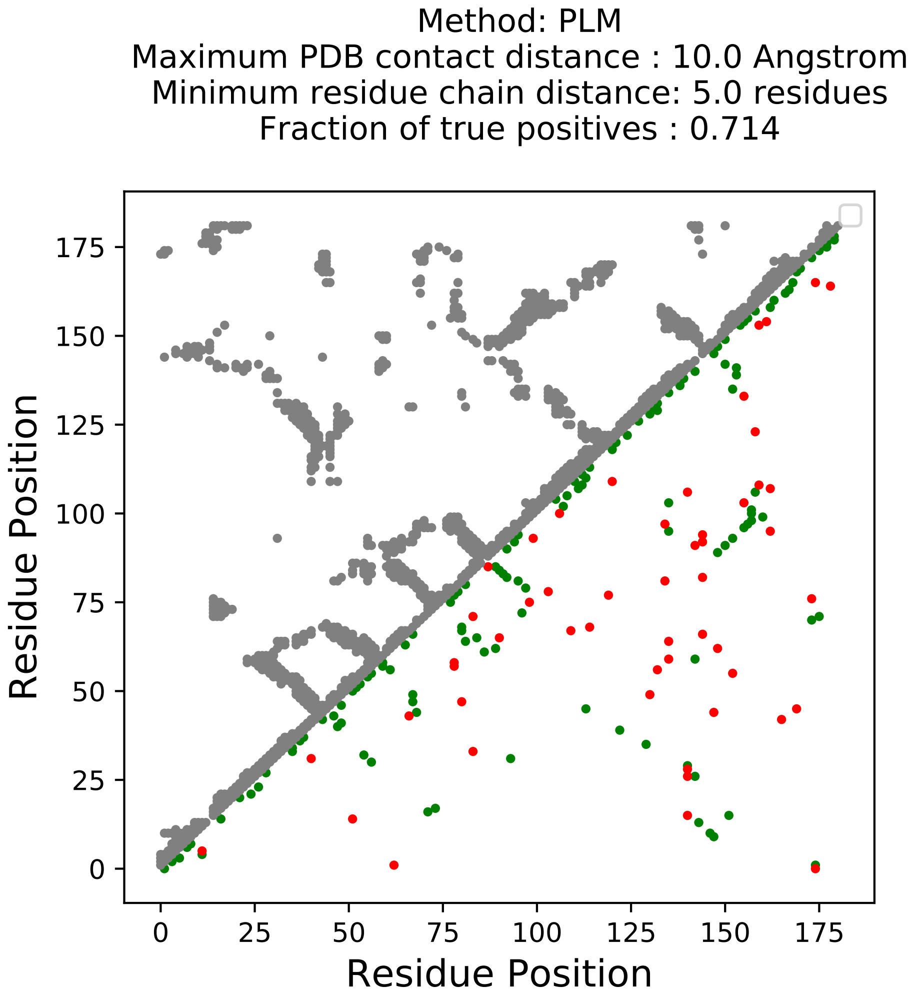

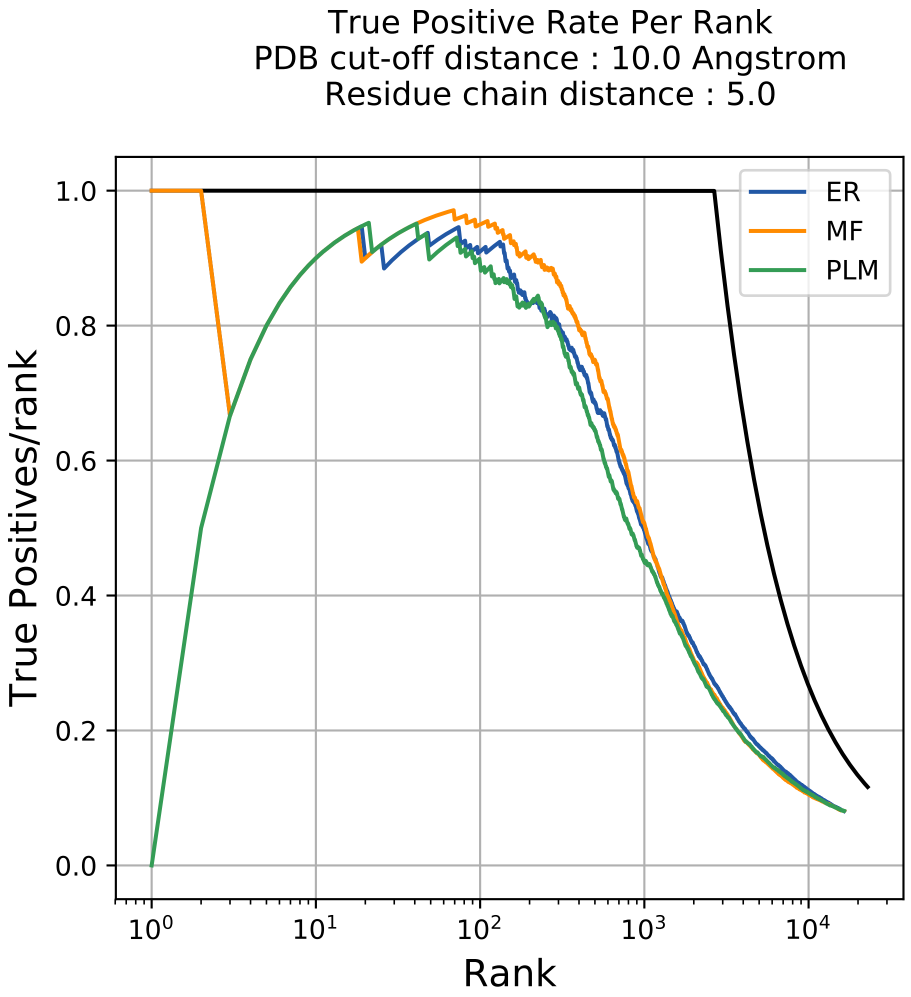

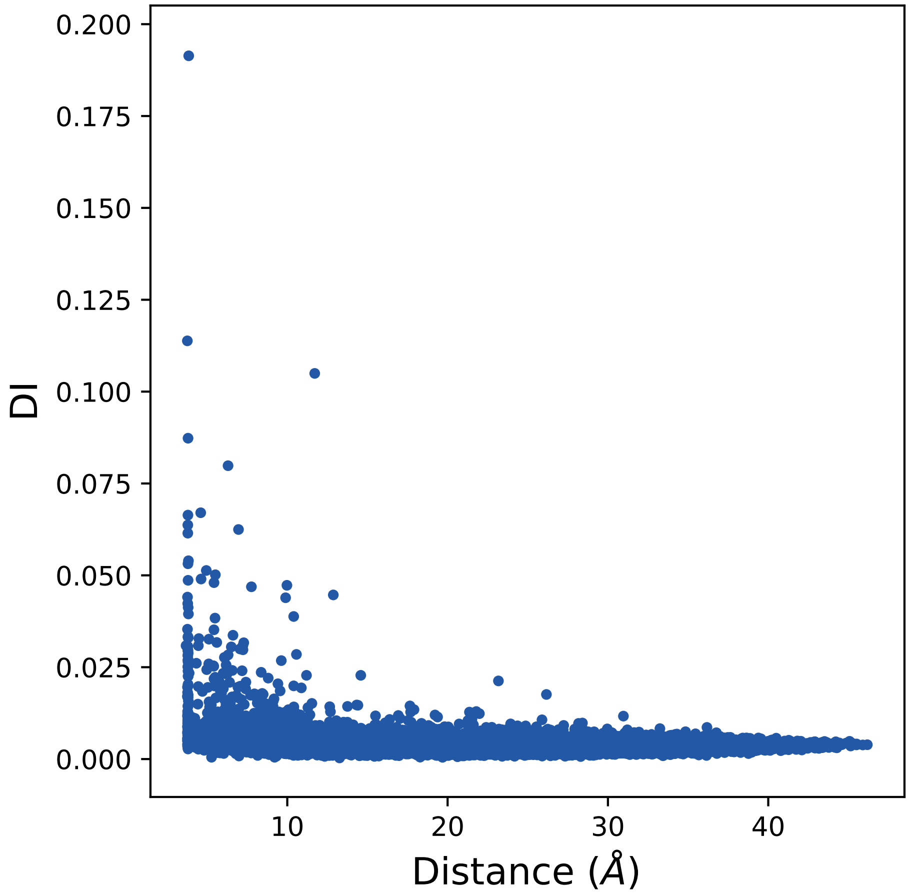

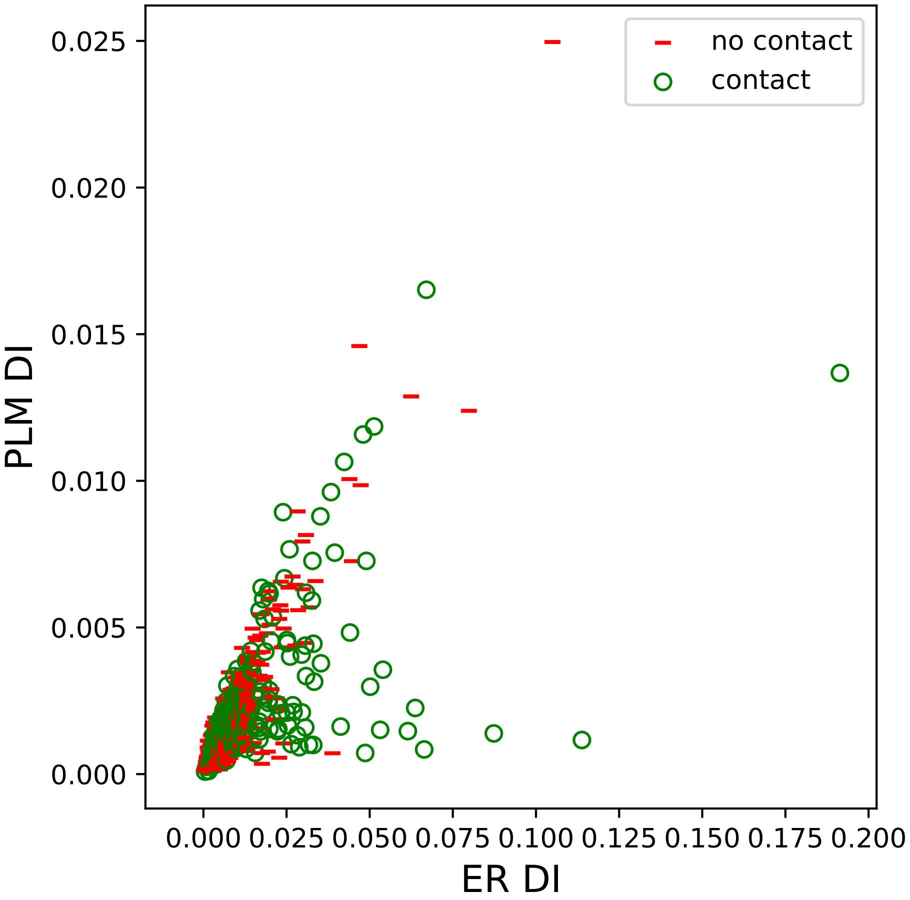

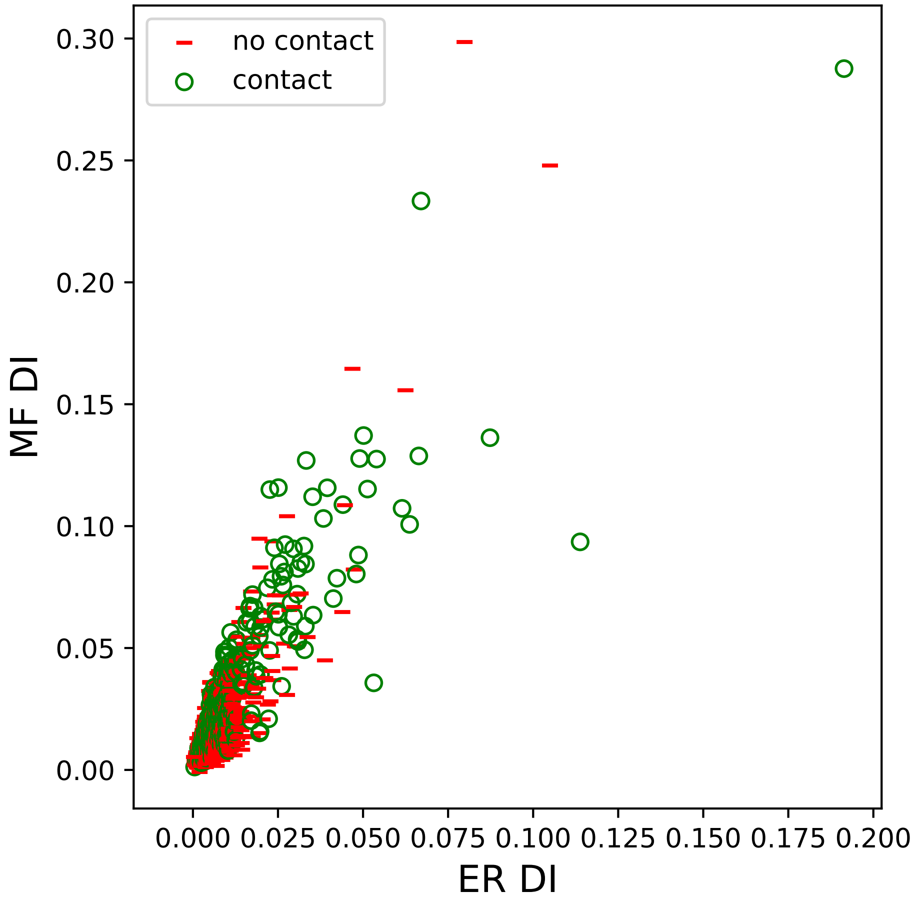

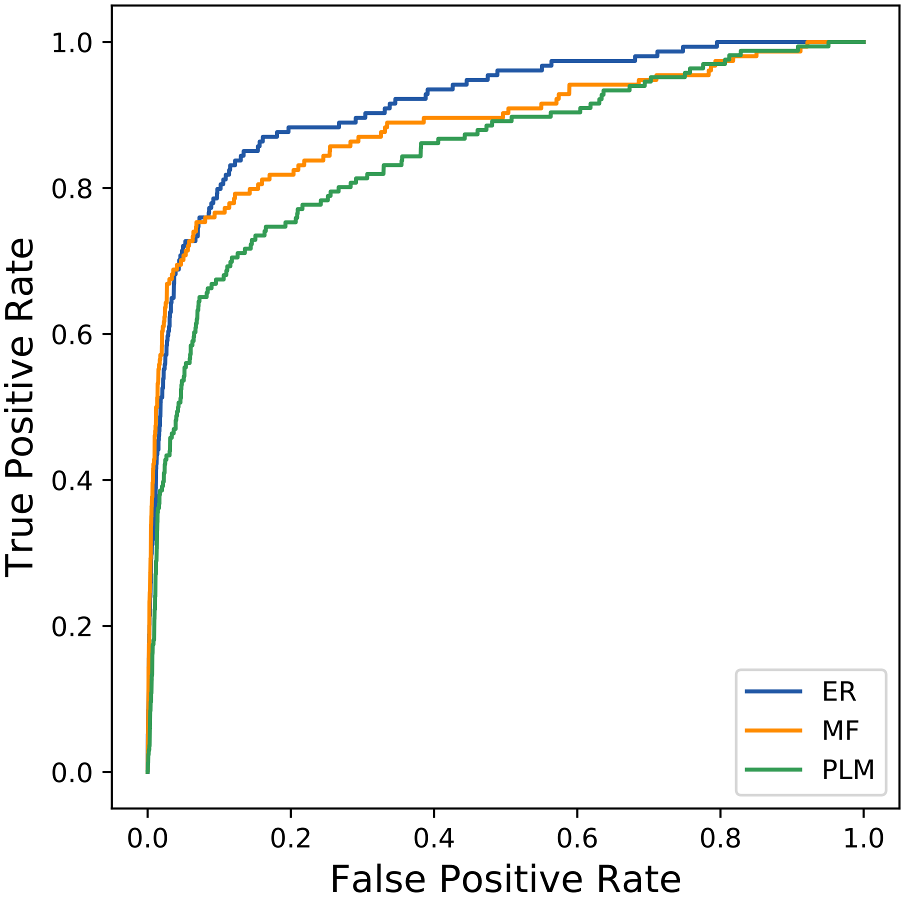

### 4806seq_63col_5cs7_PF07654_plots.pdf

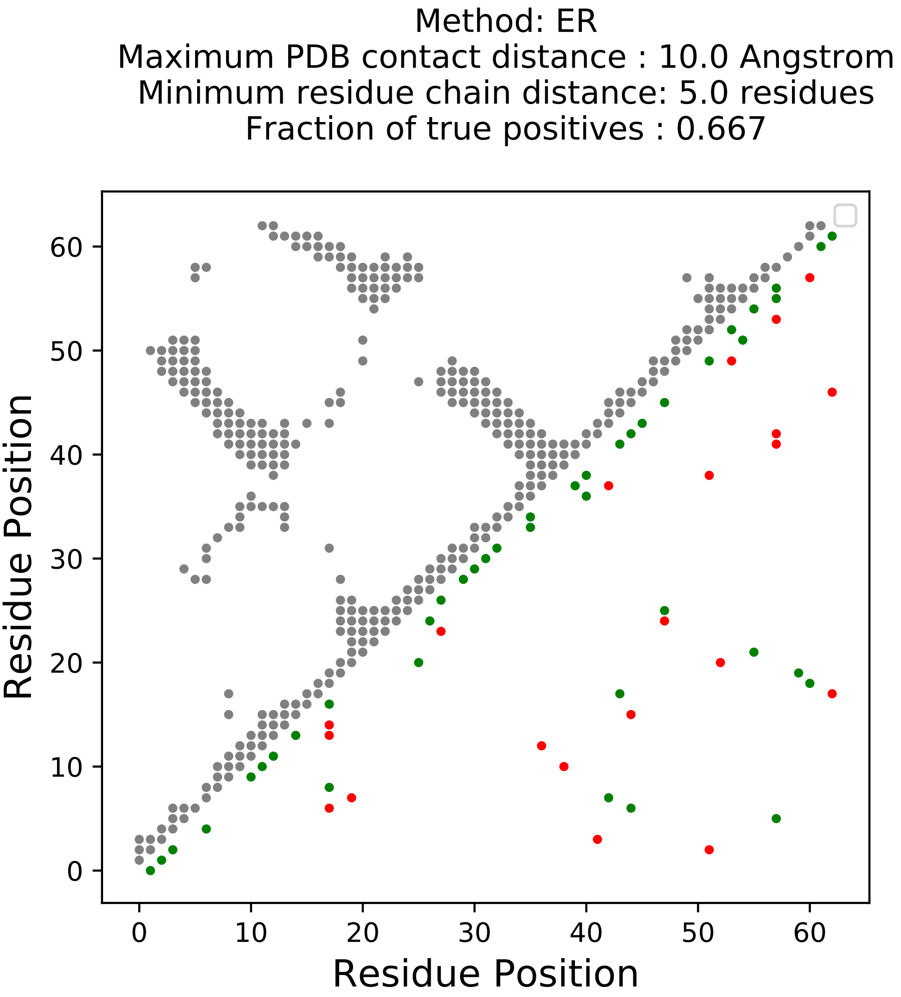

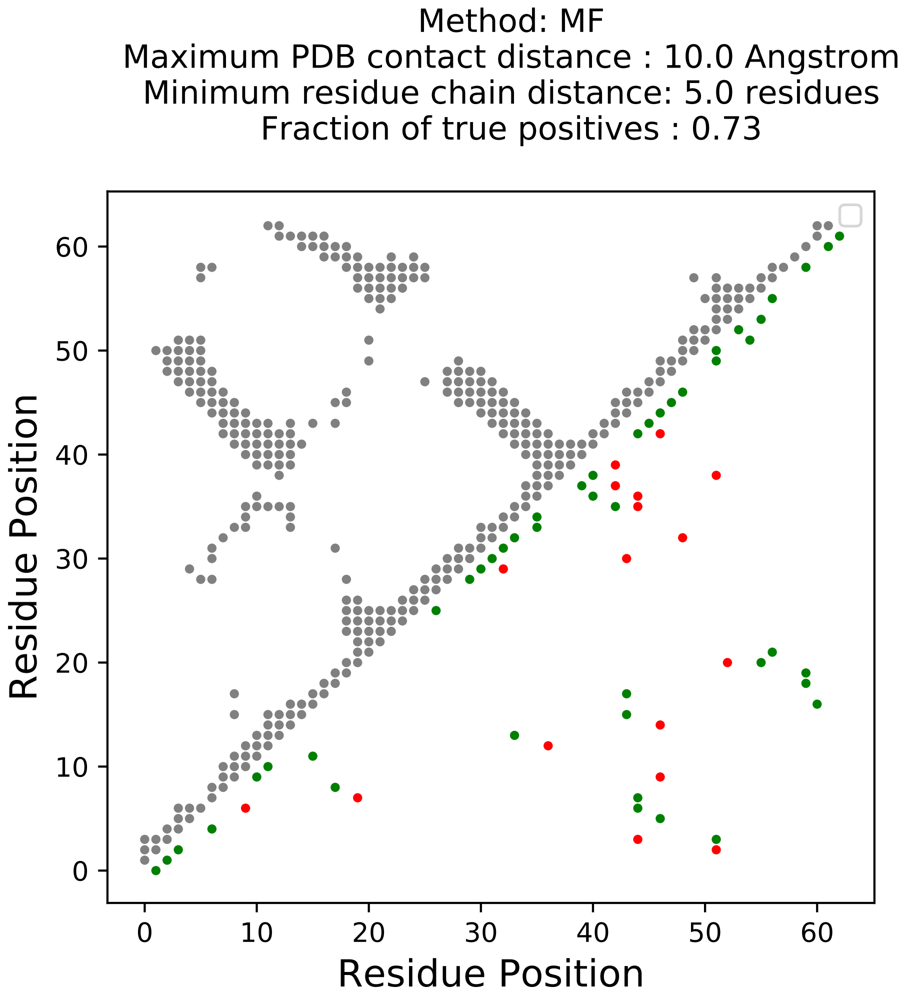

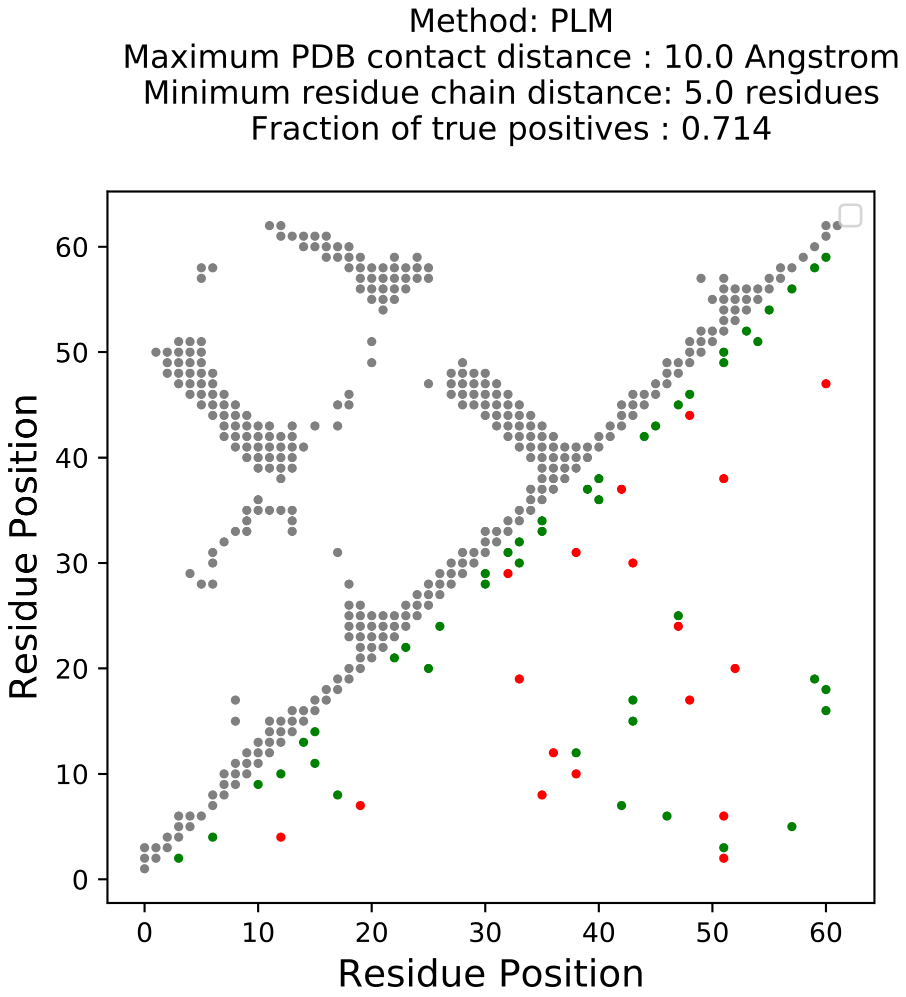

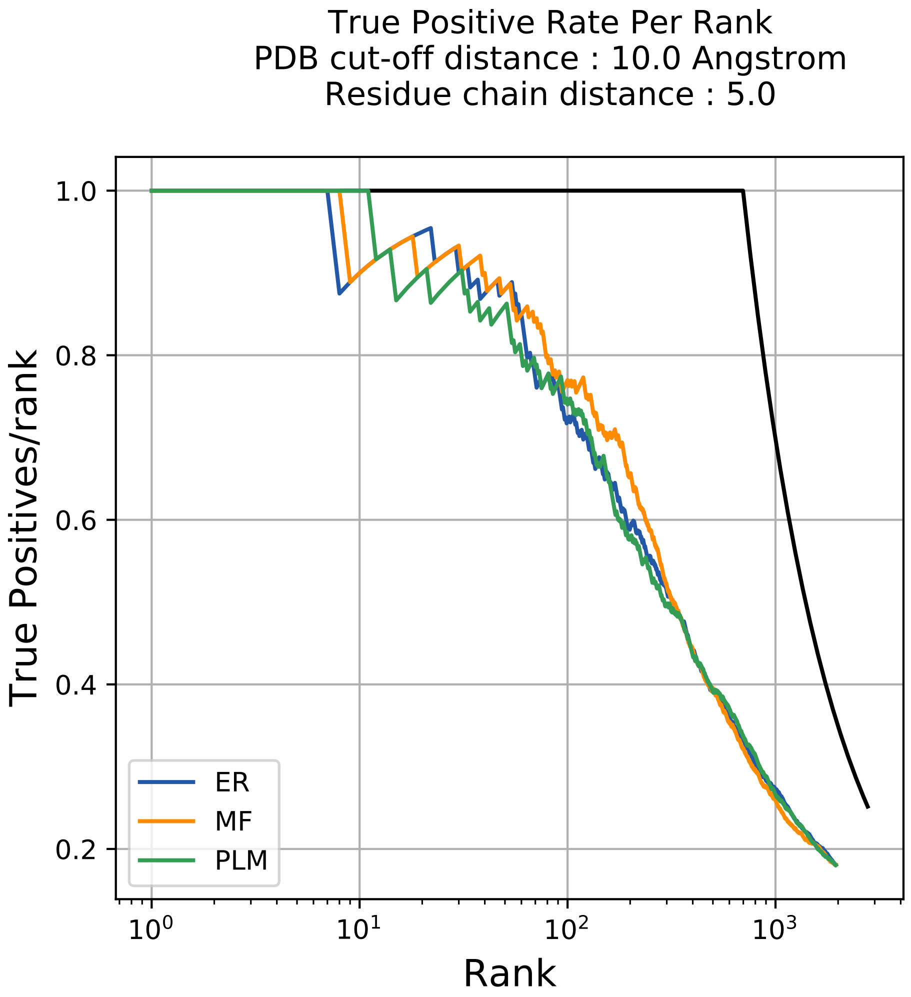

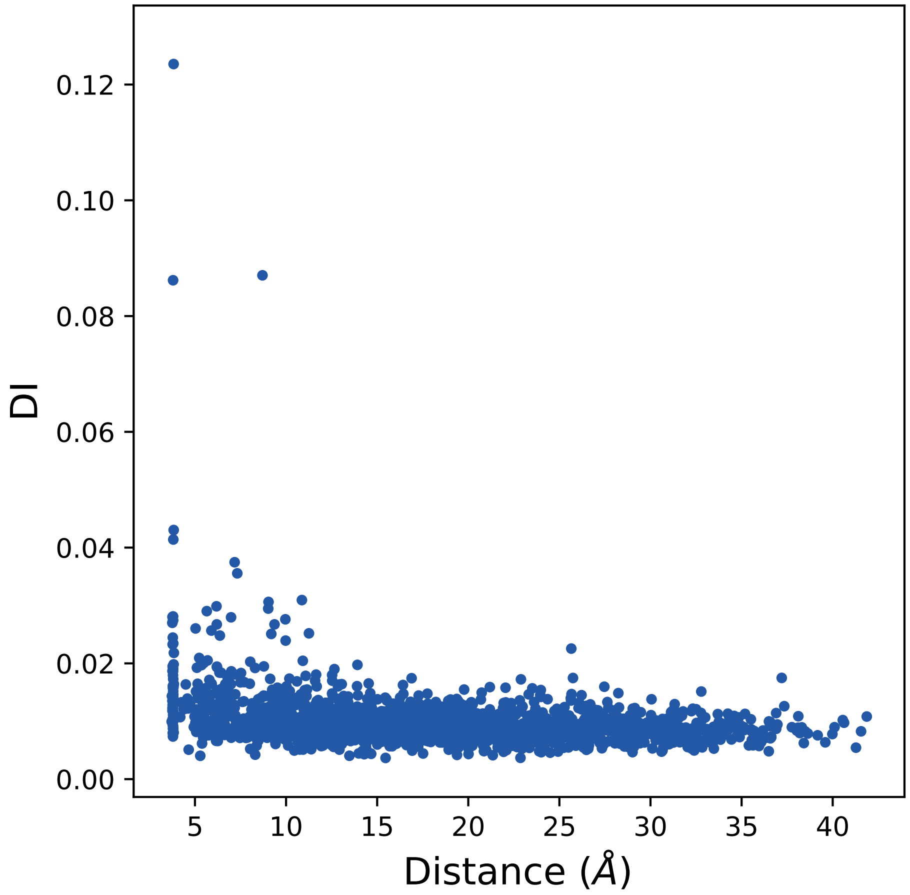

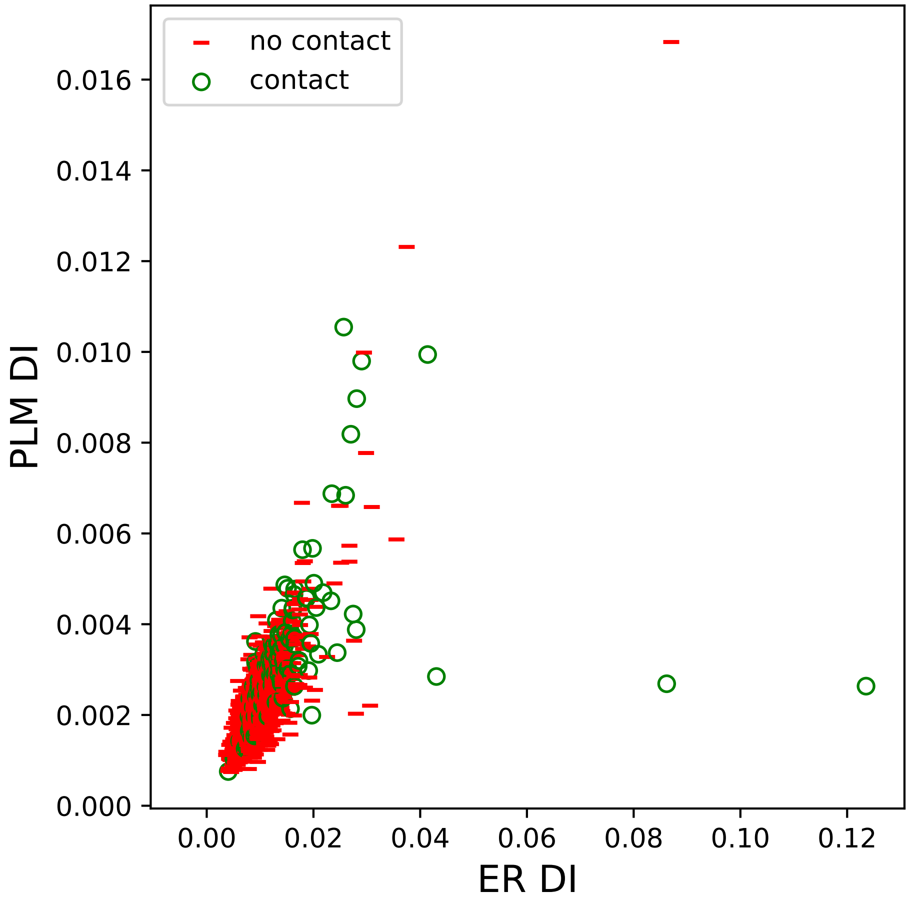

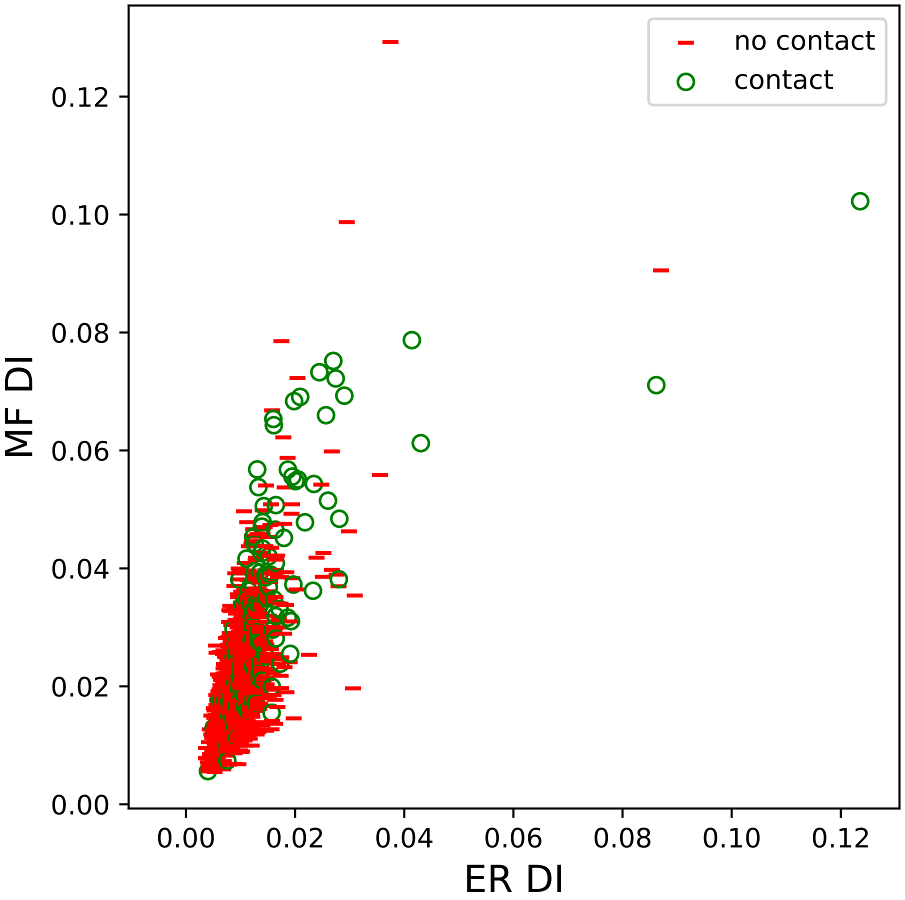

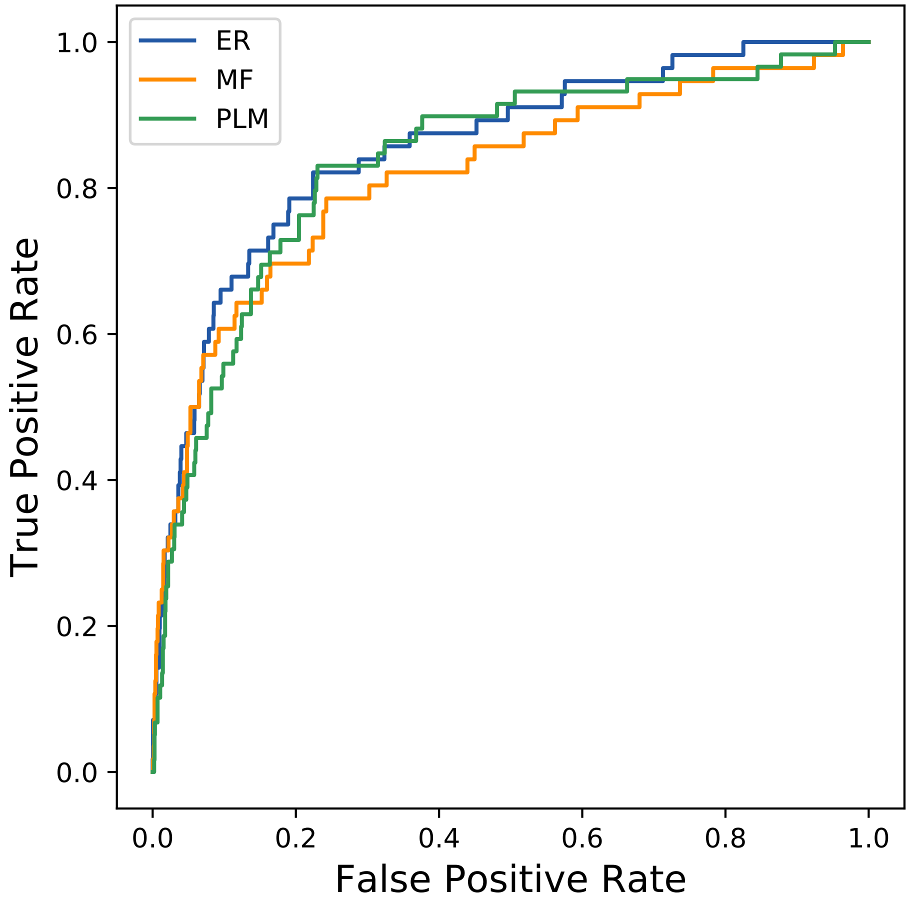

### 8143seq_98col_4izo_PF02222_plots.pdf

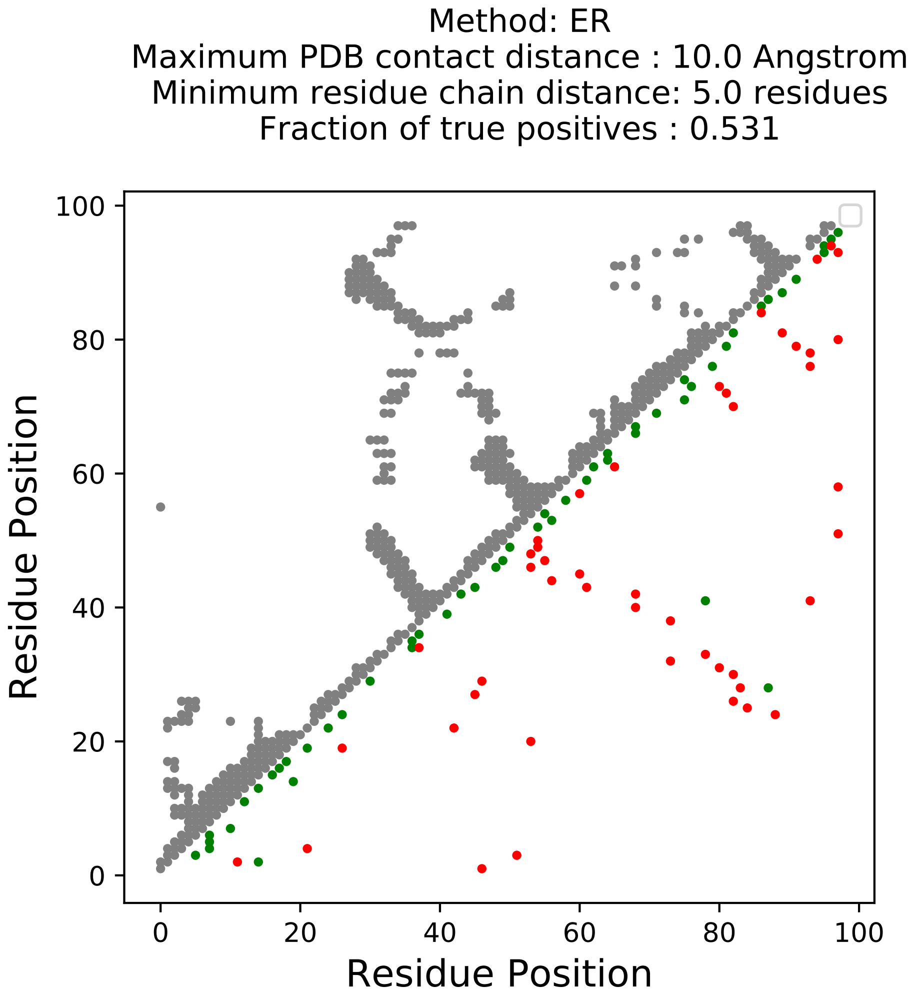

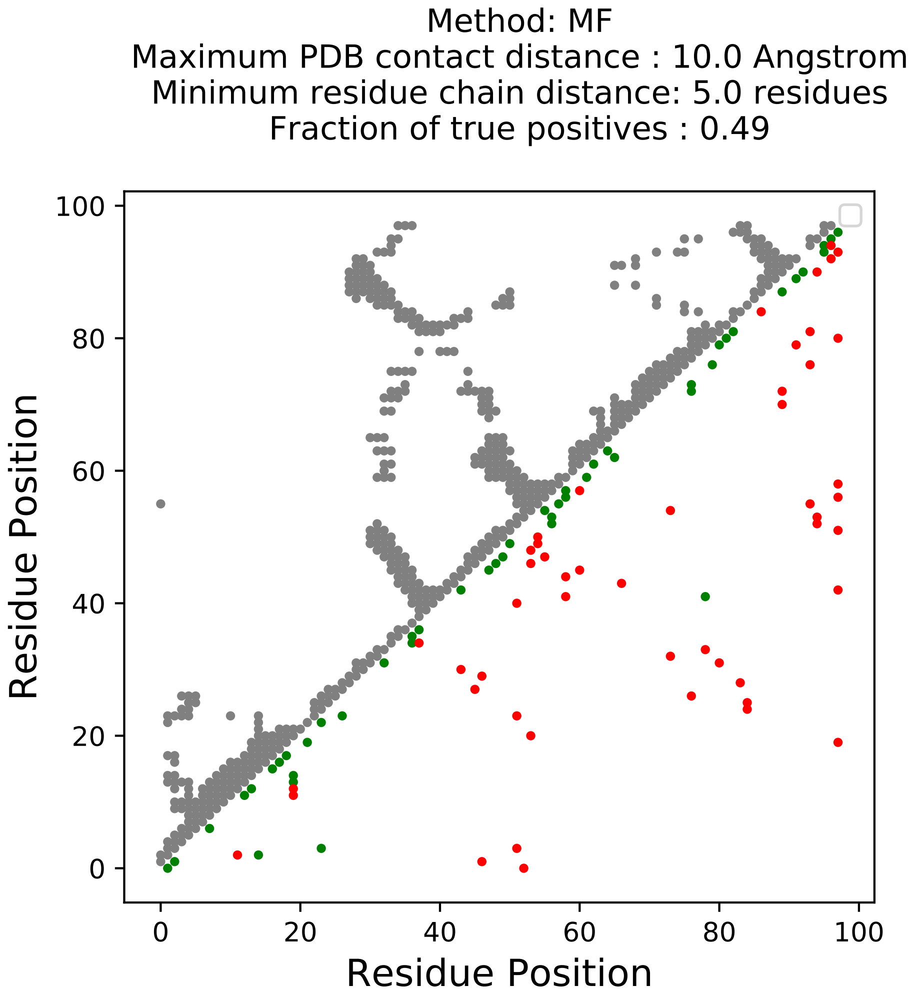

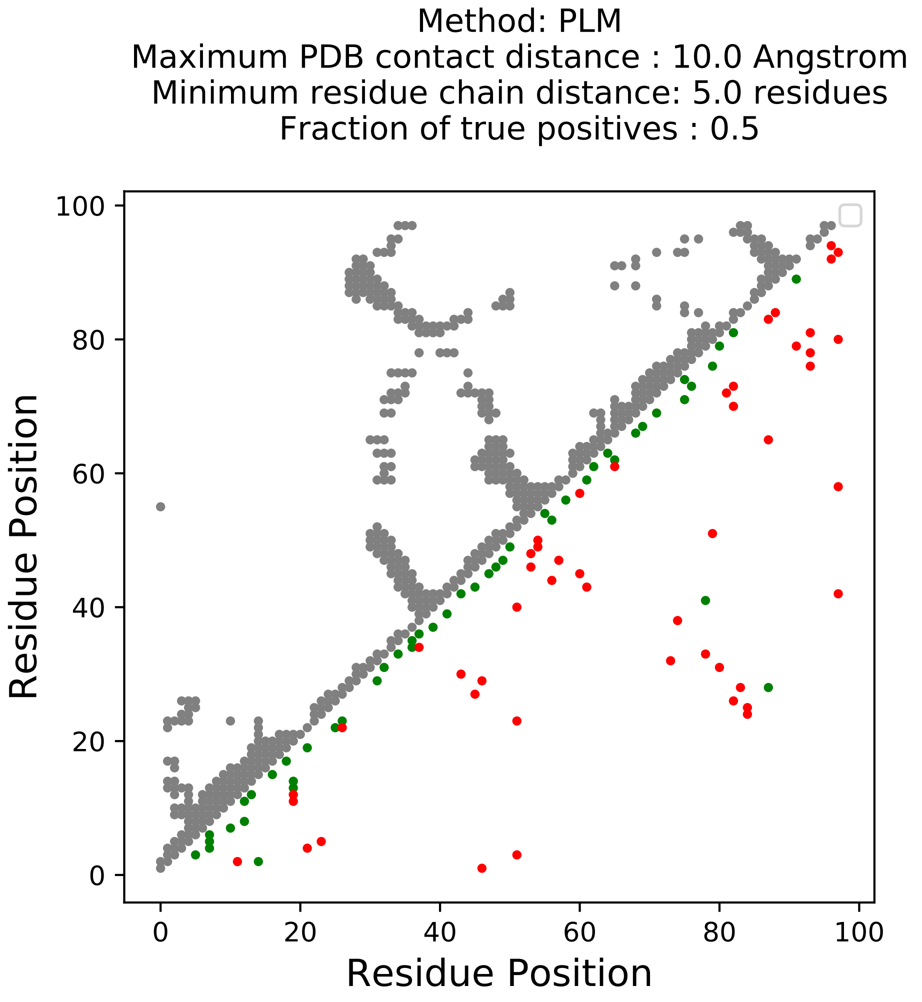

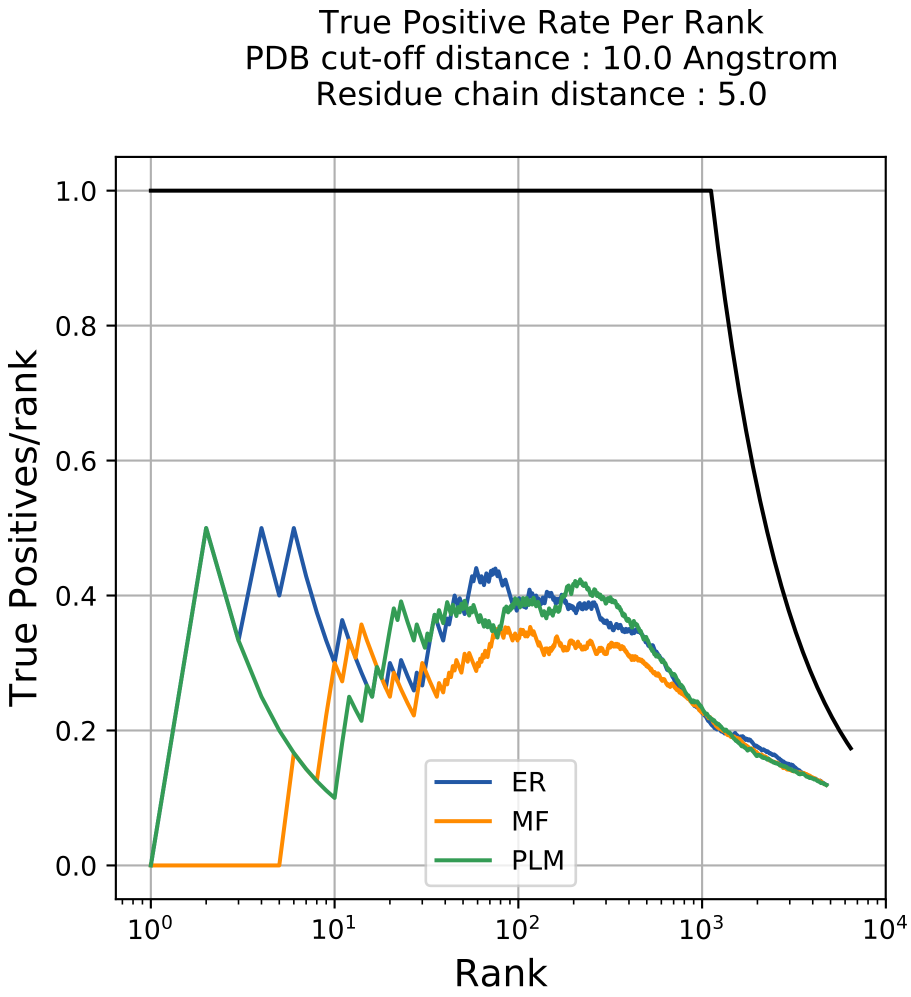

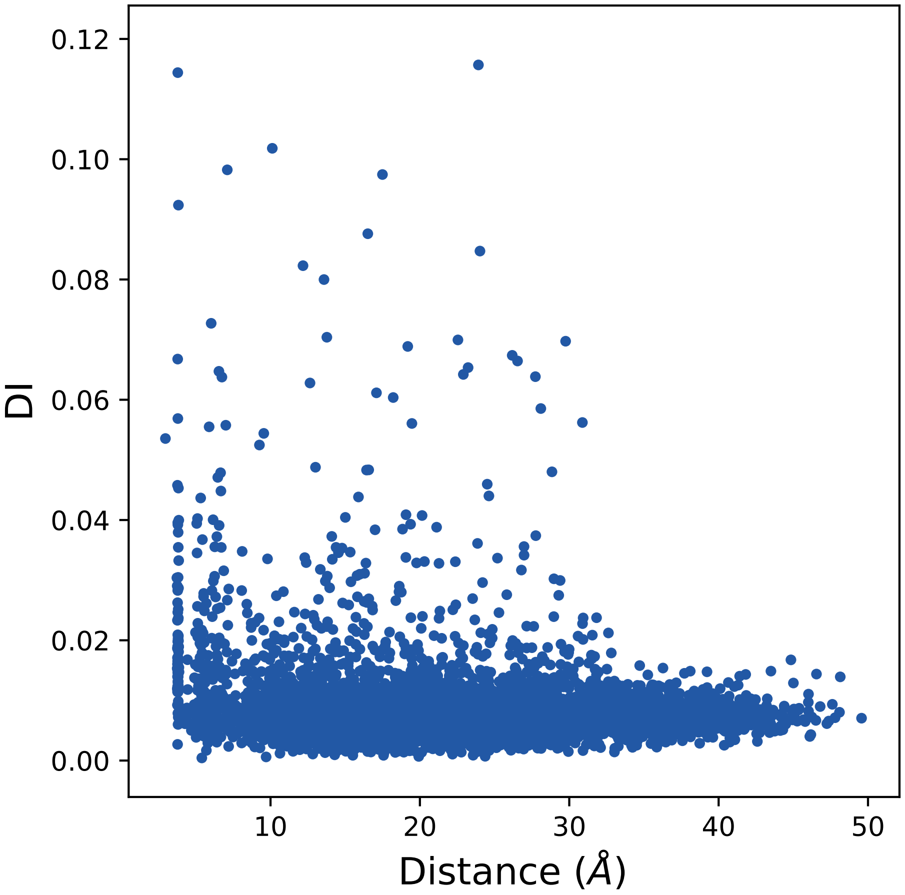

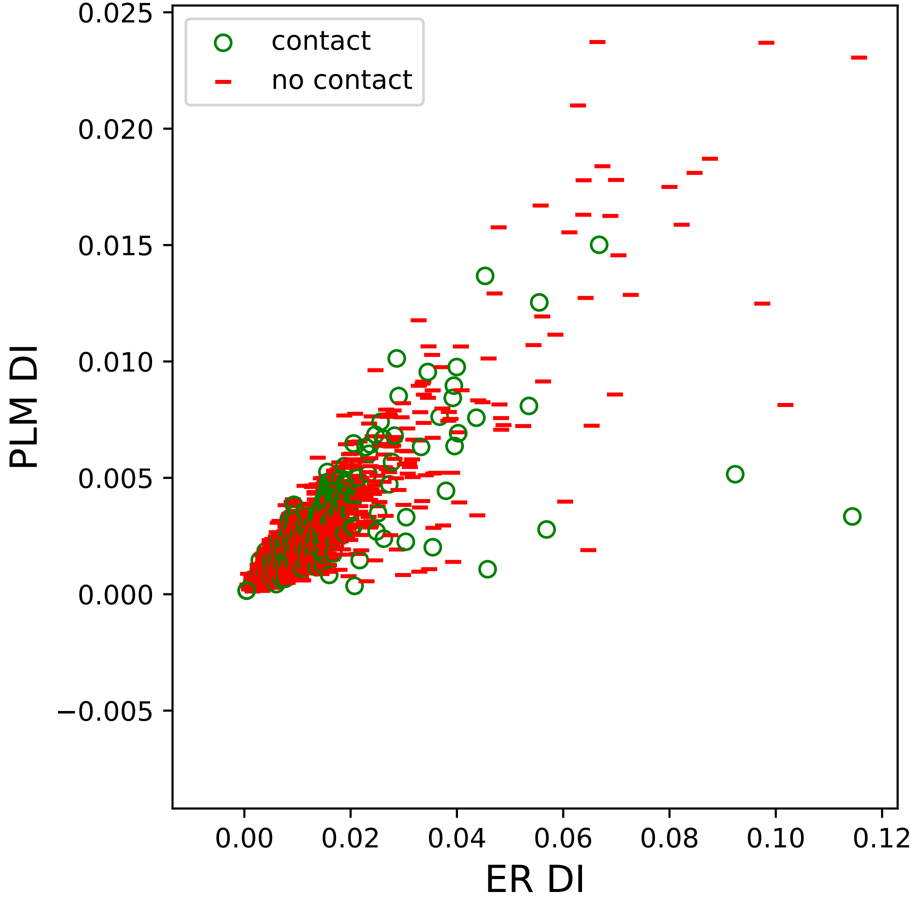

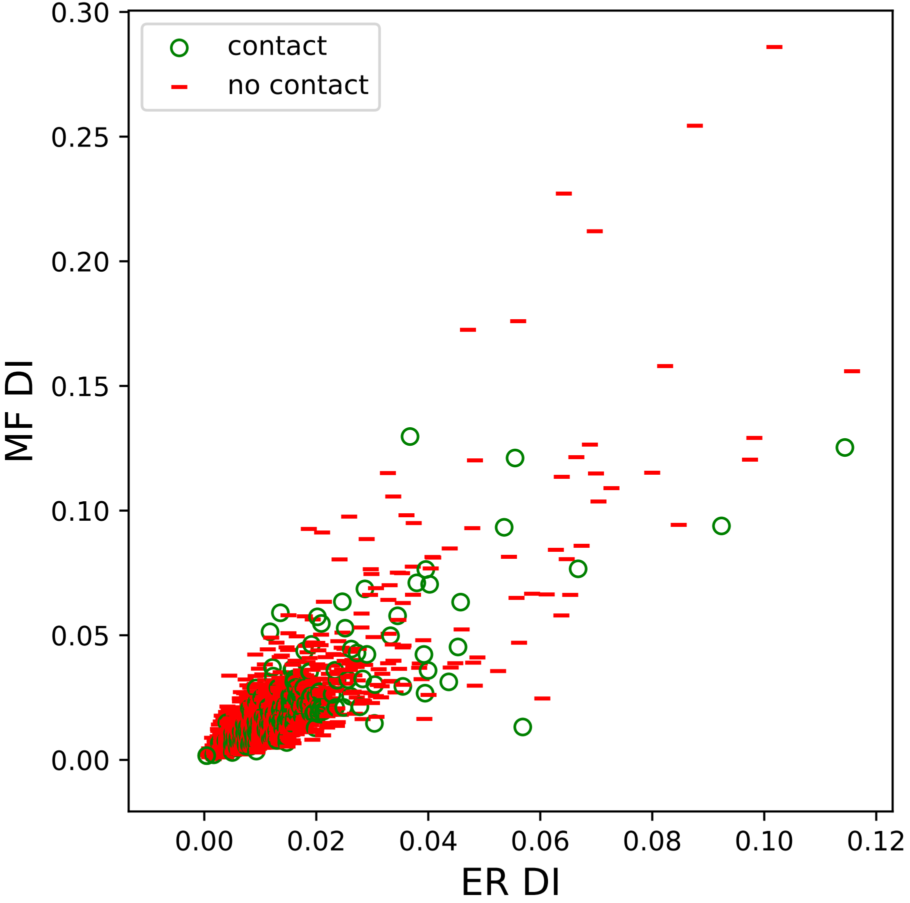

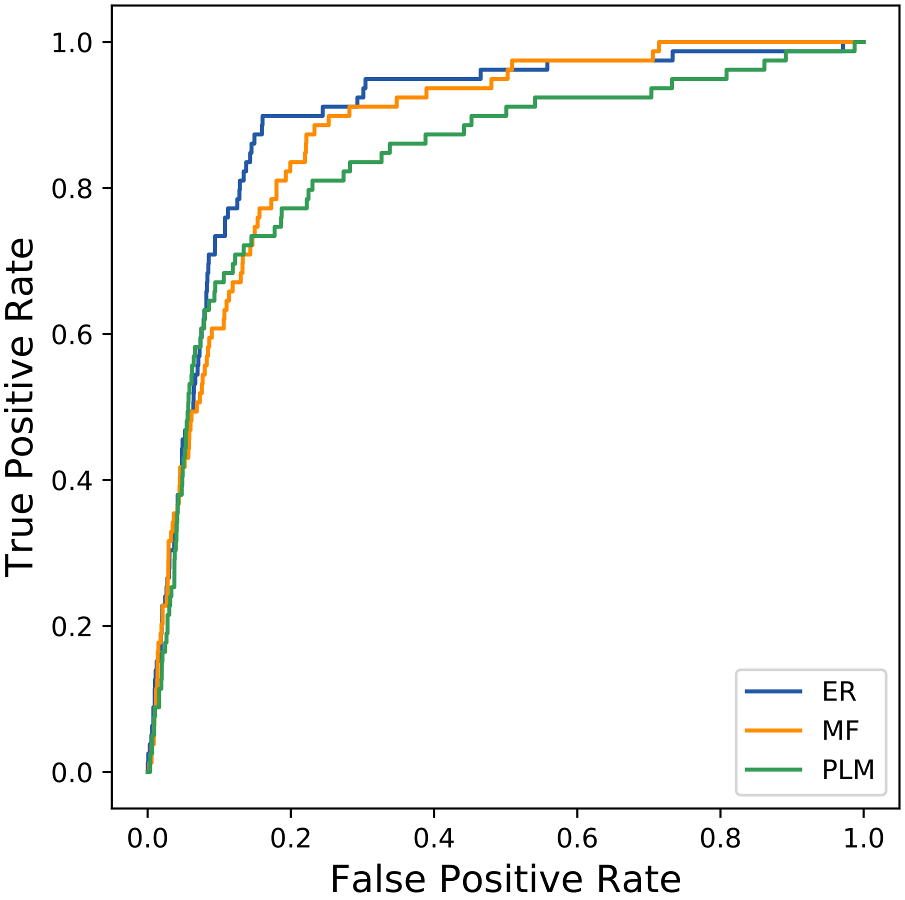

### 8352seq_60col_4qao_PF00034_plots.pdf

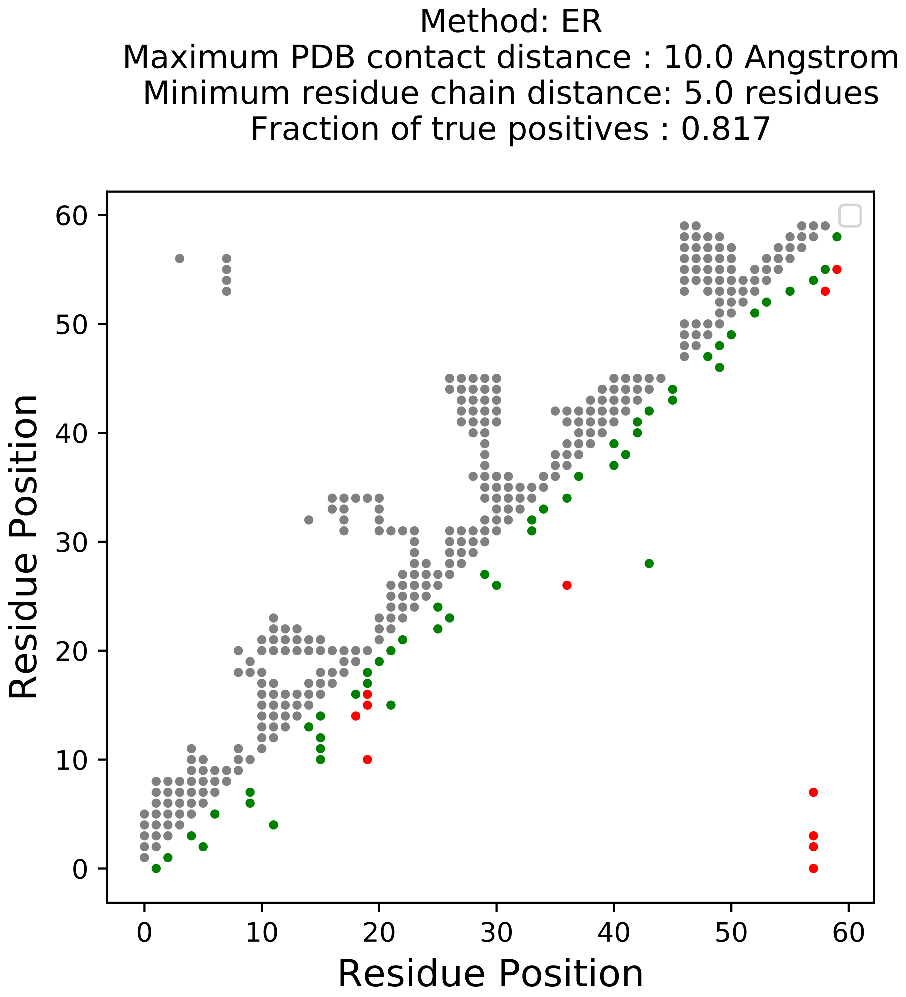

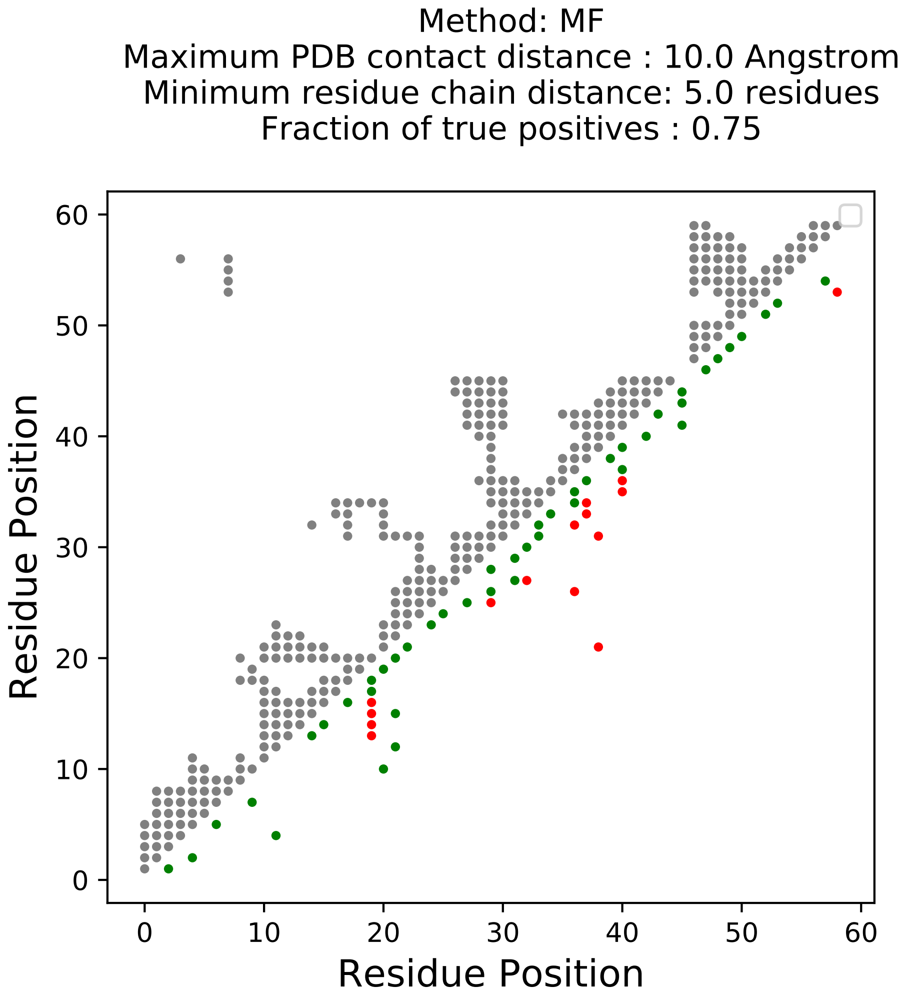

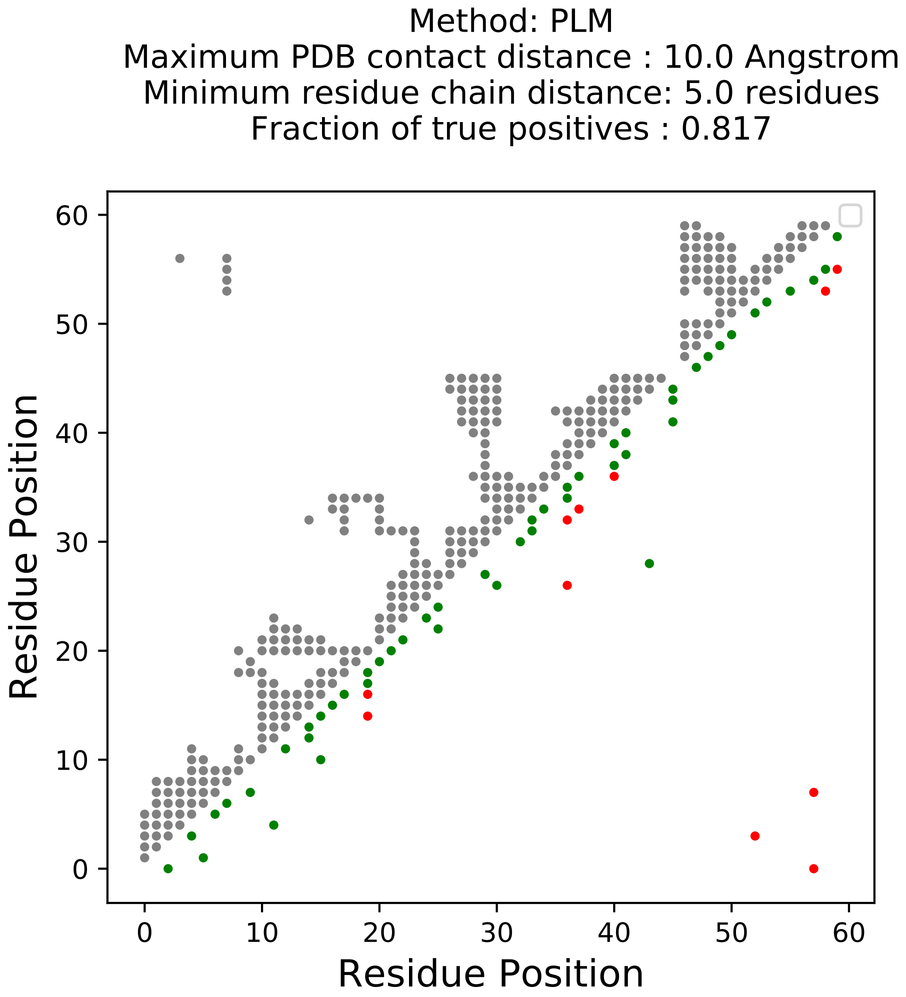

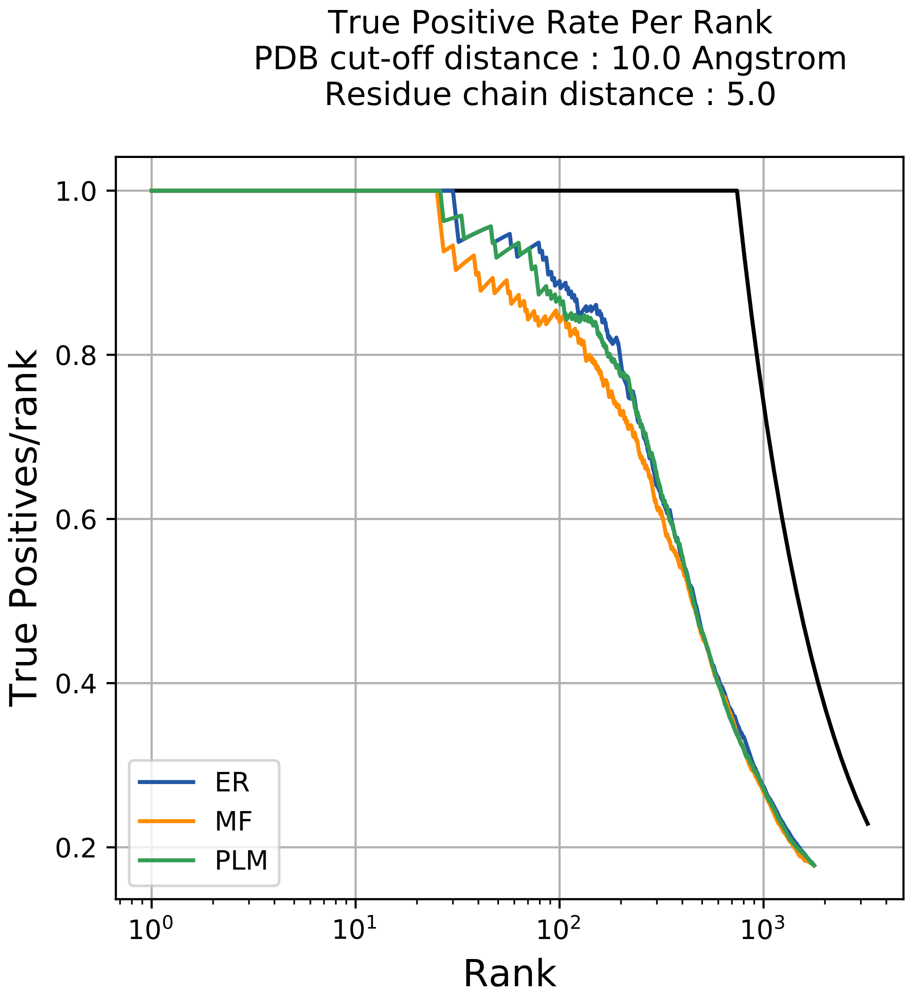

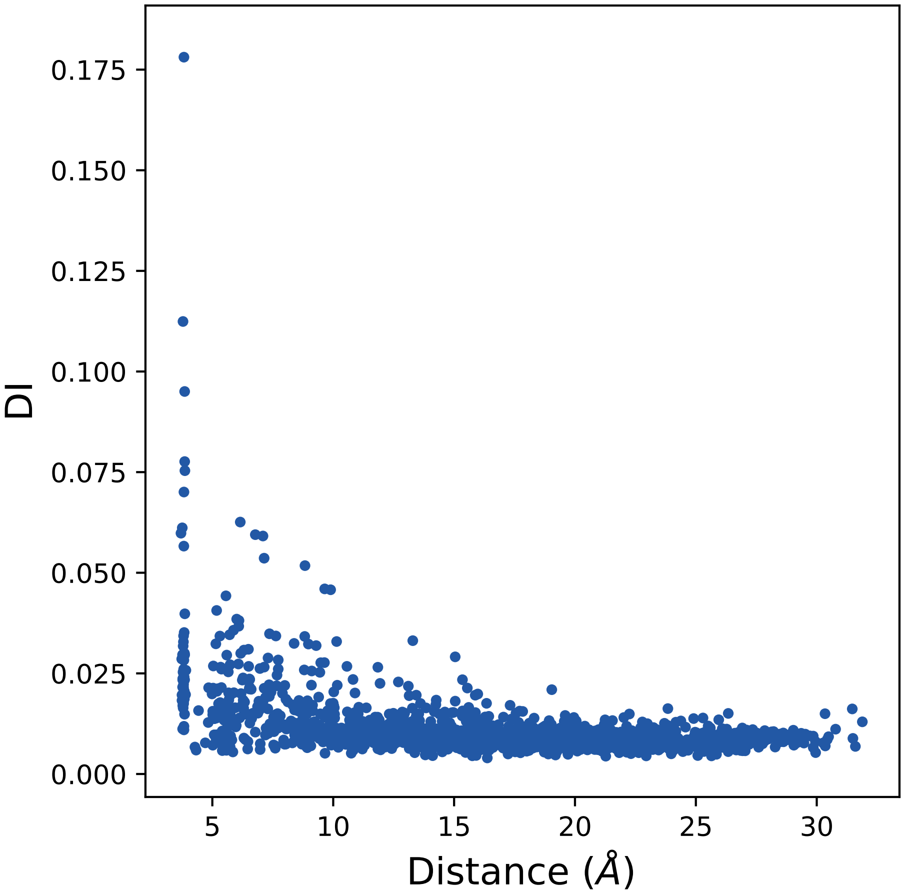

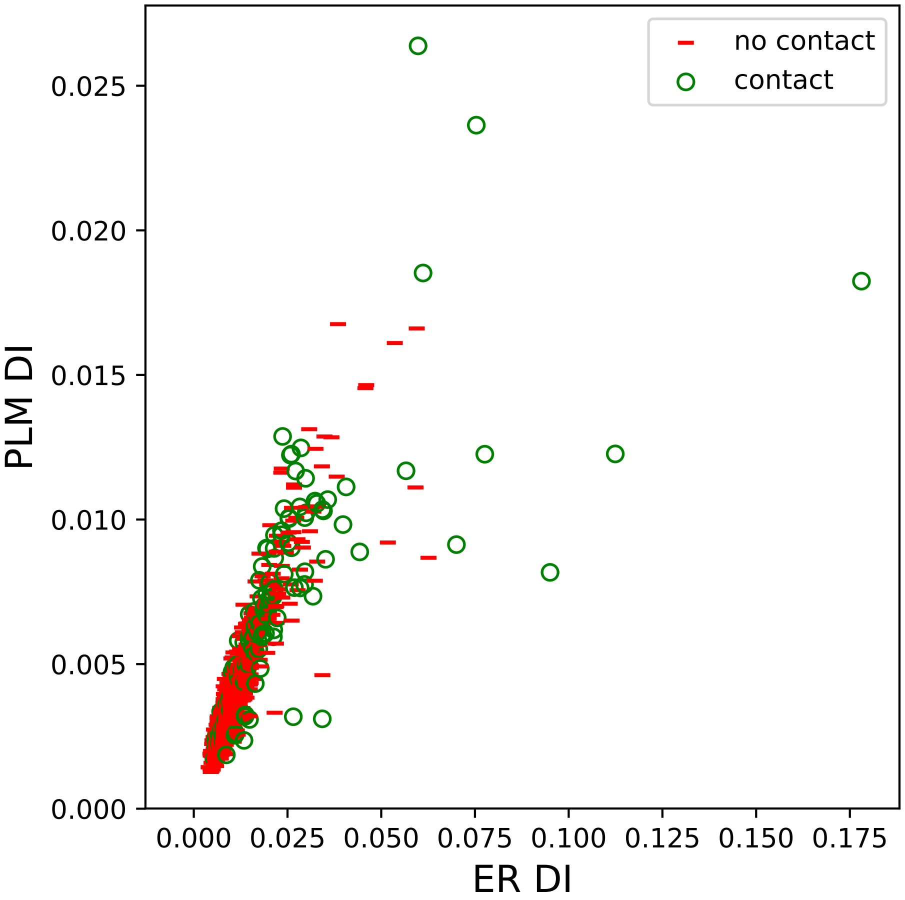

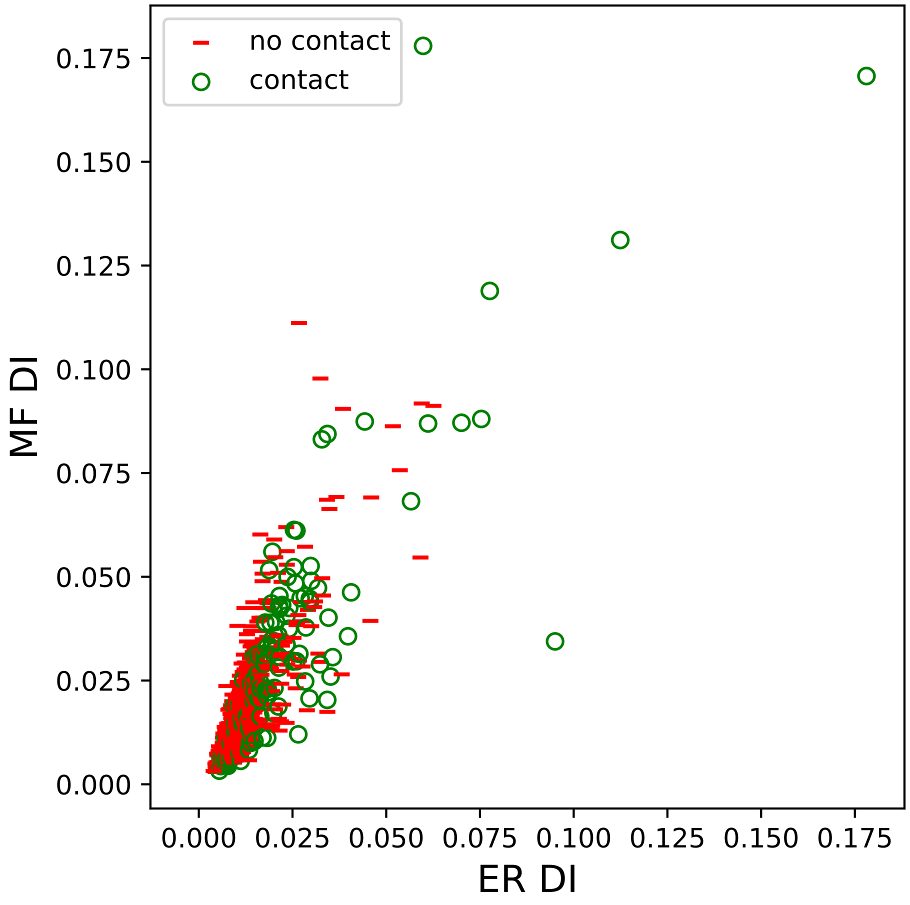

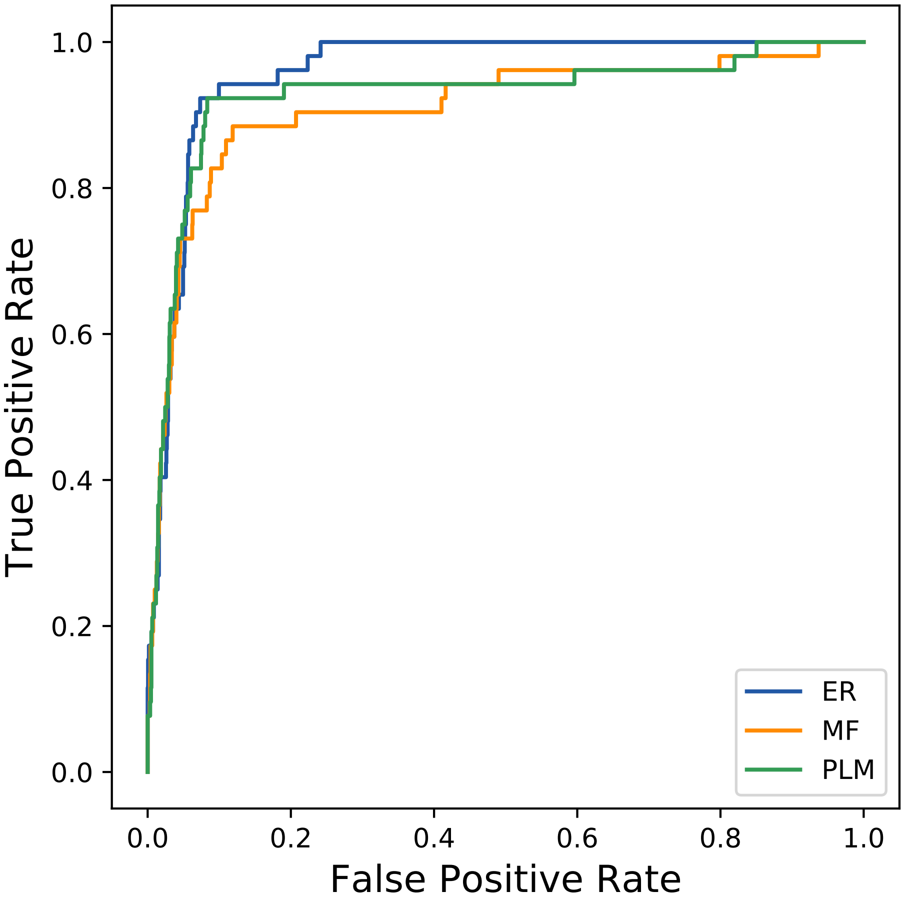

### 8416seq_25col_5jb5_PF00014_plots.pdf

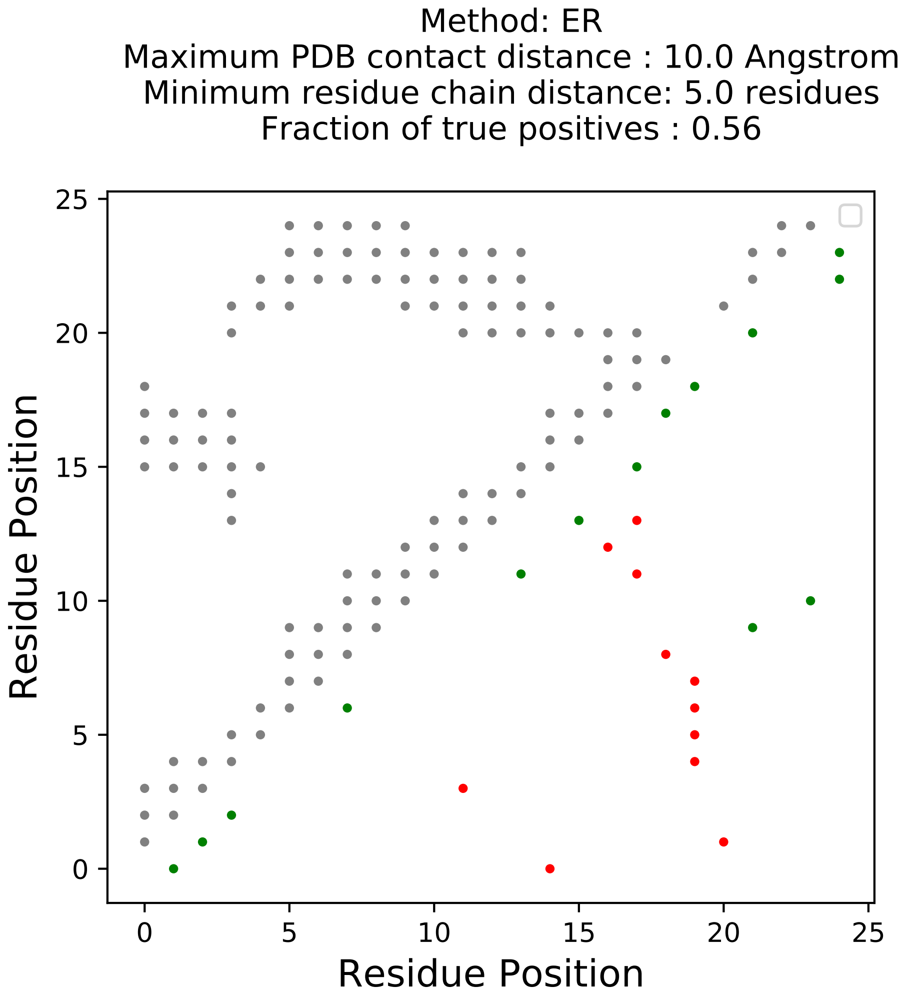

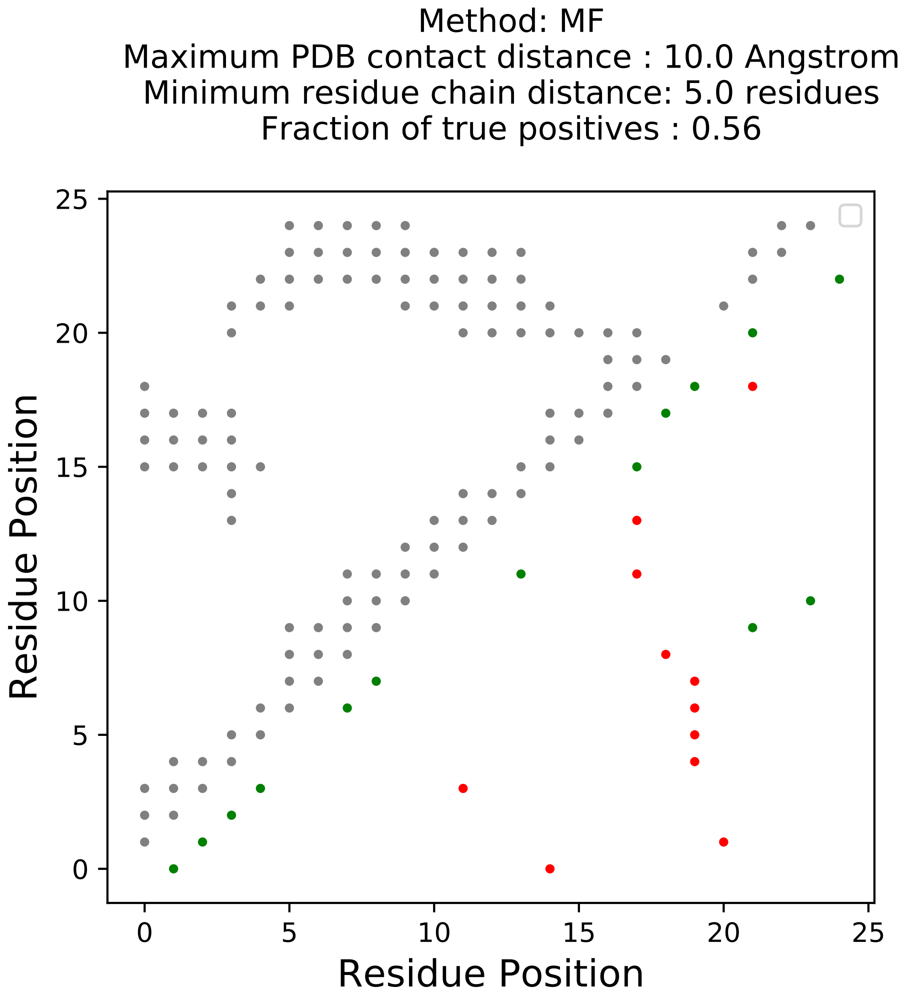

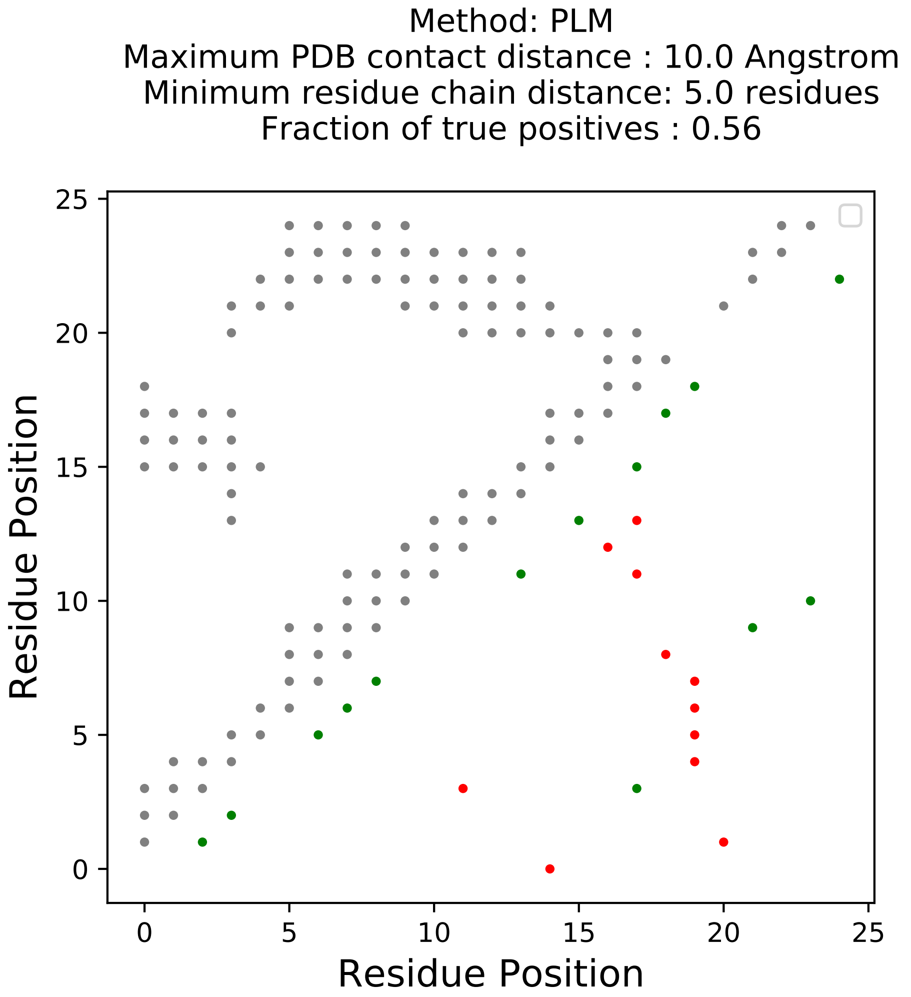

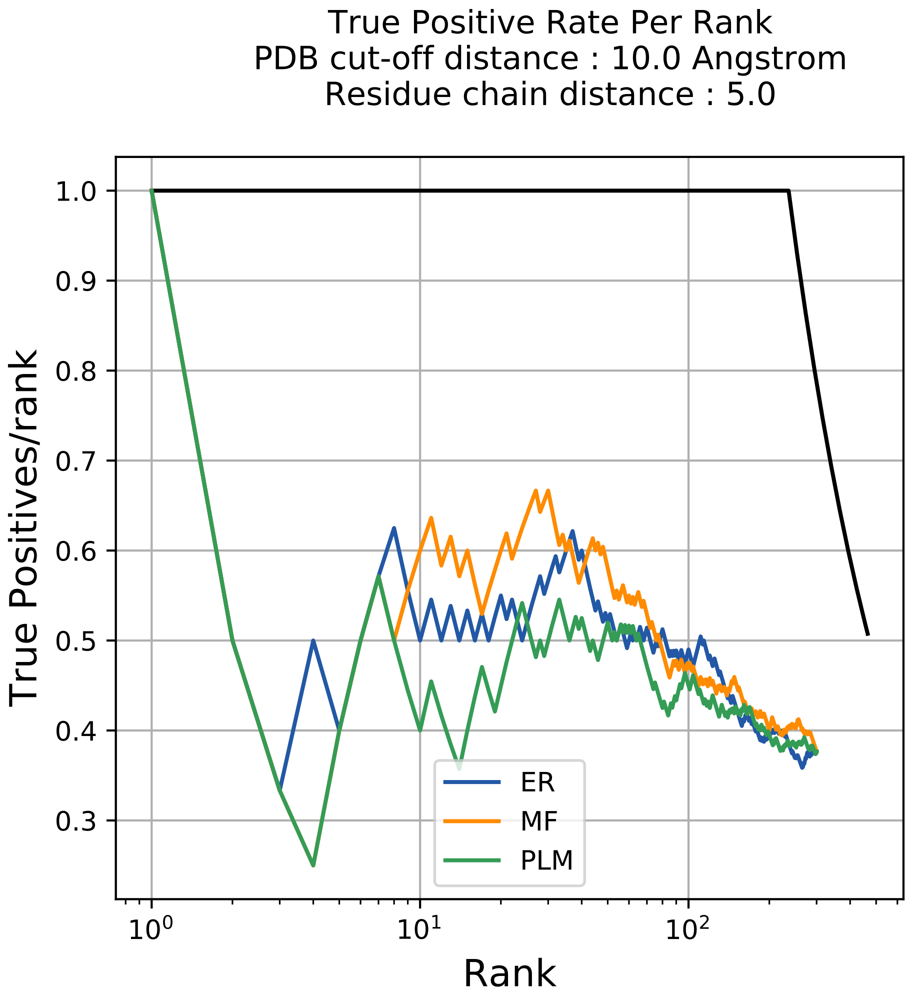

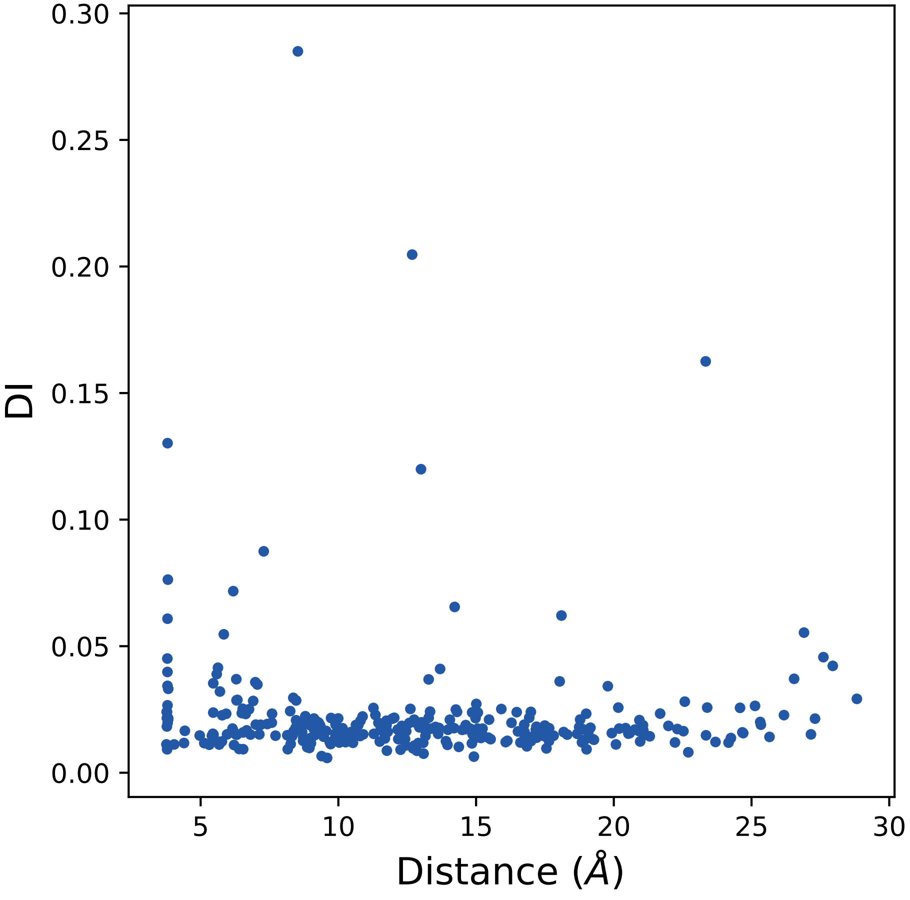

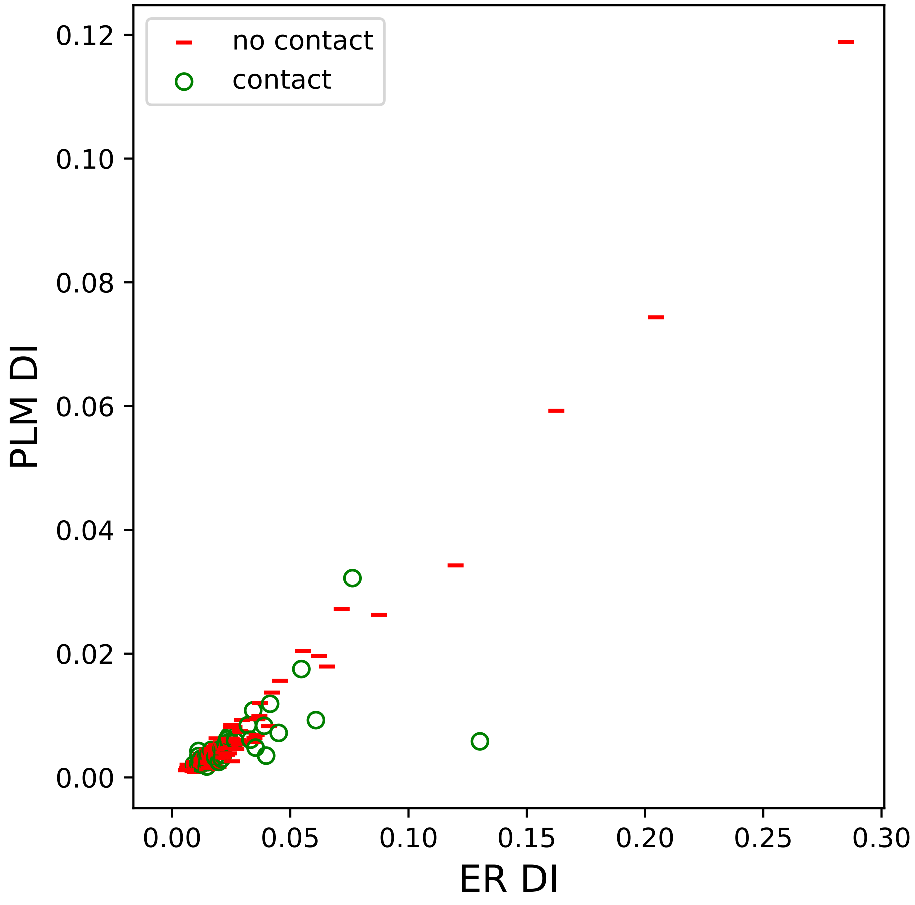

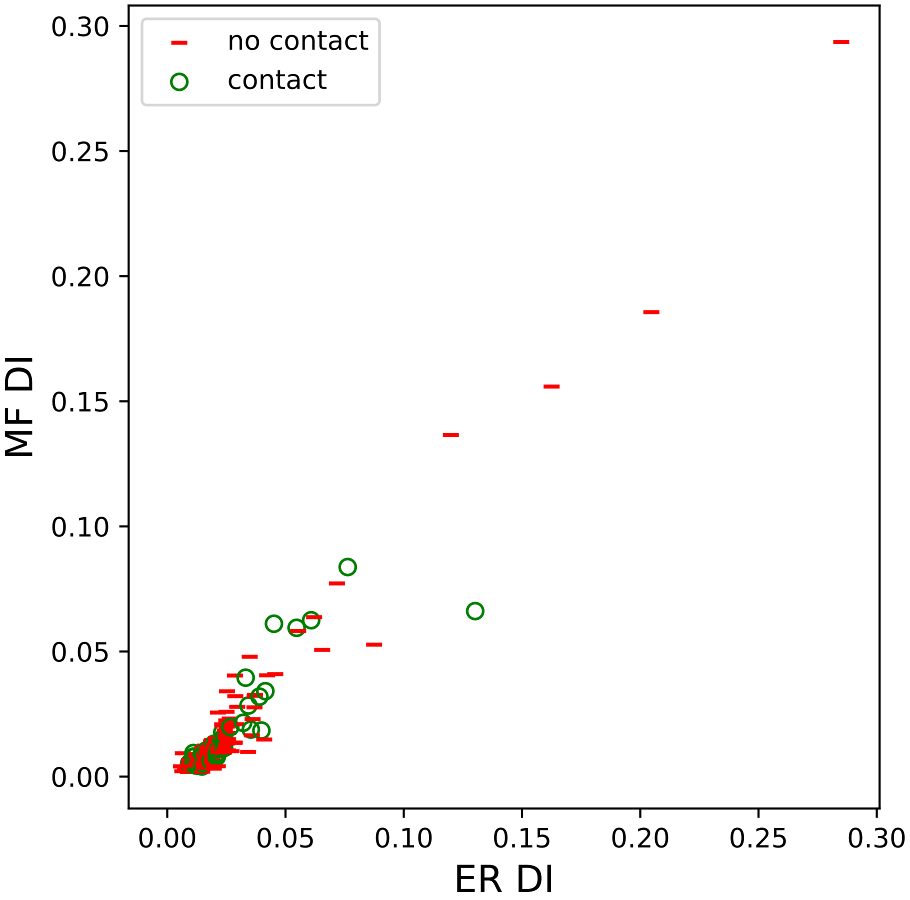

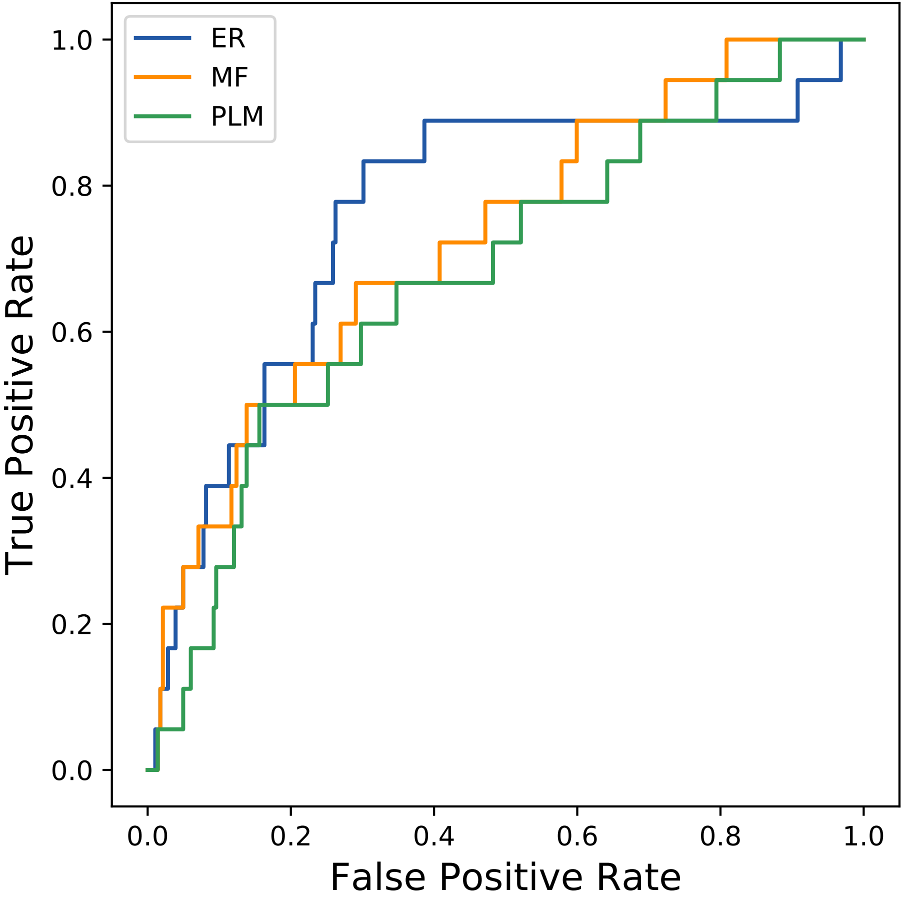

### 10059seq_92col_3rz2_PF00782_plots.pdf

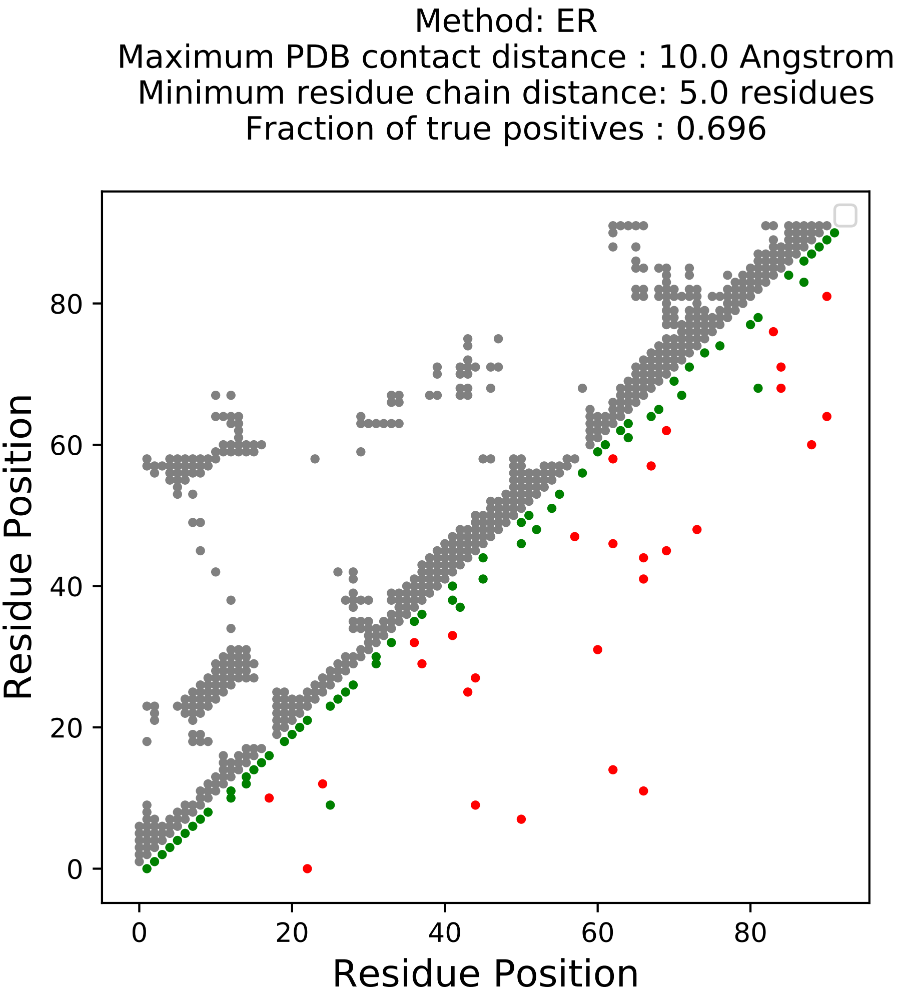

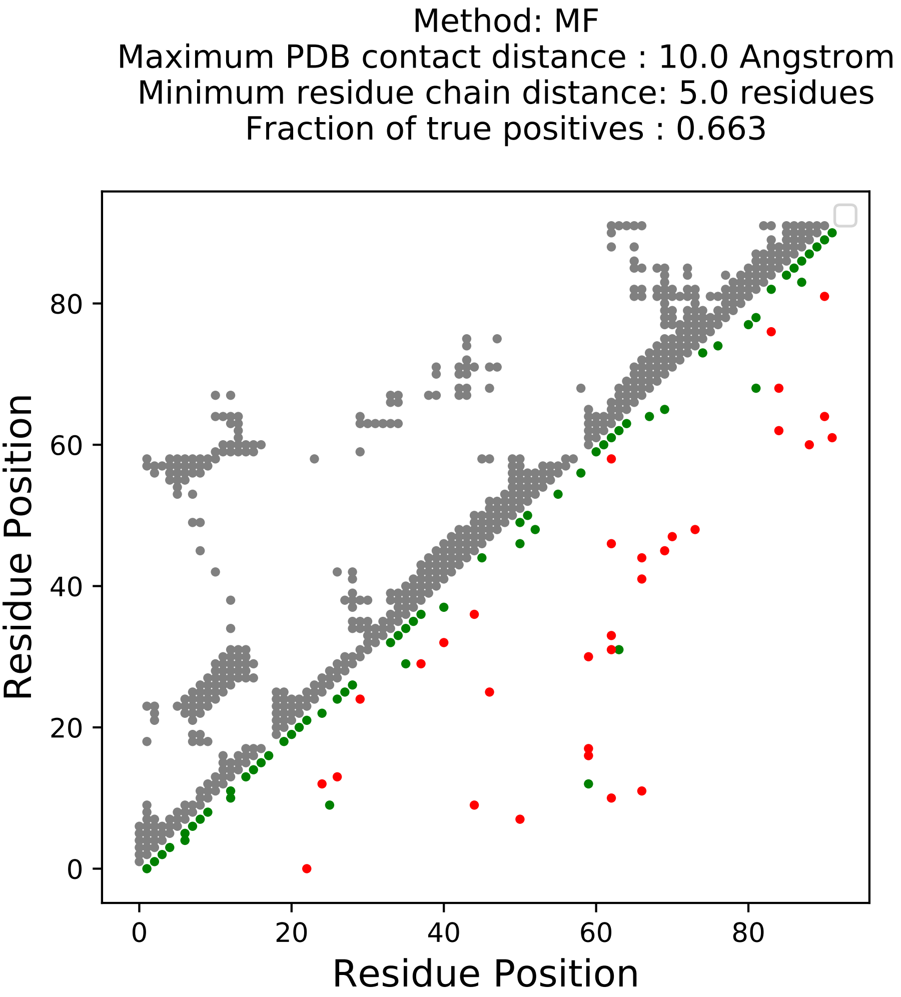

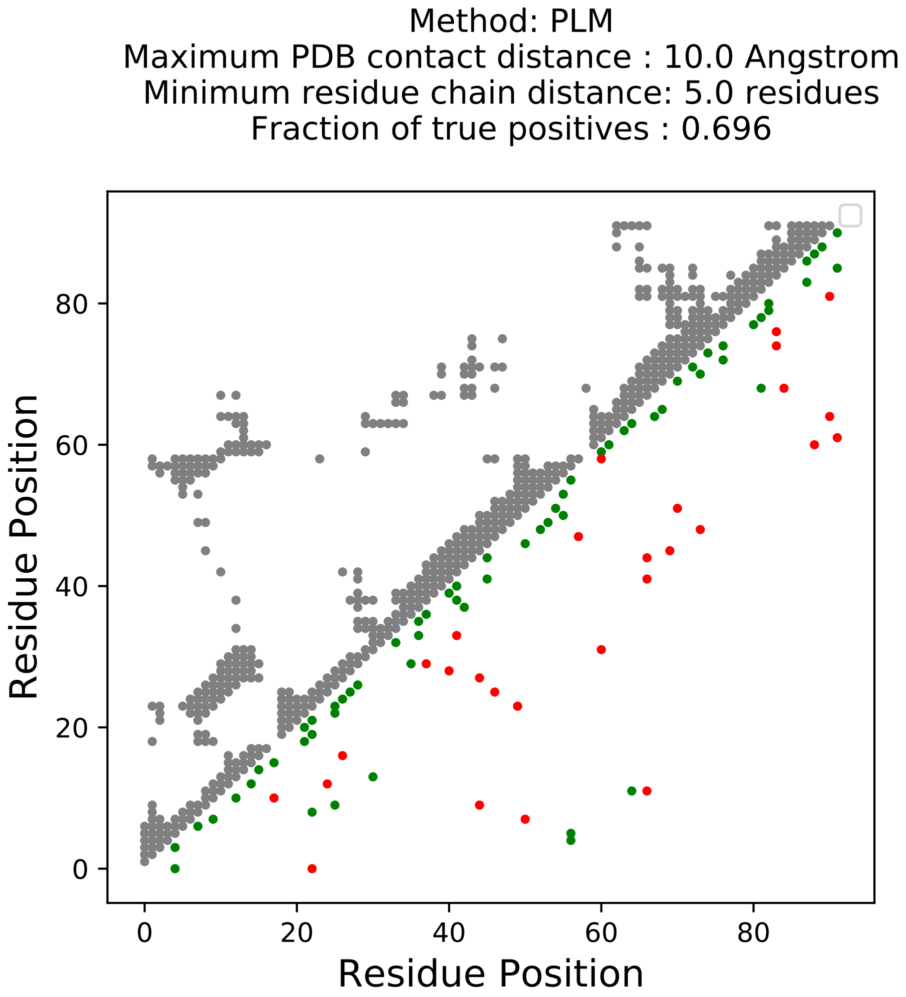

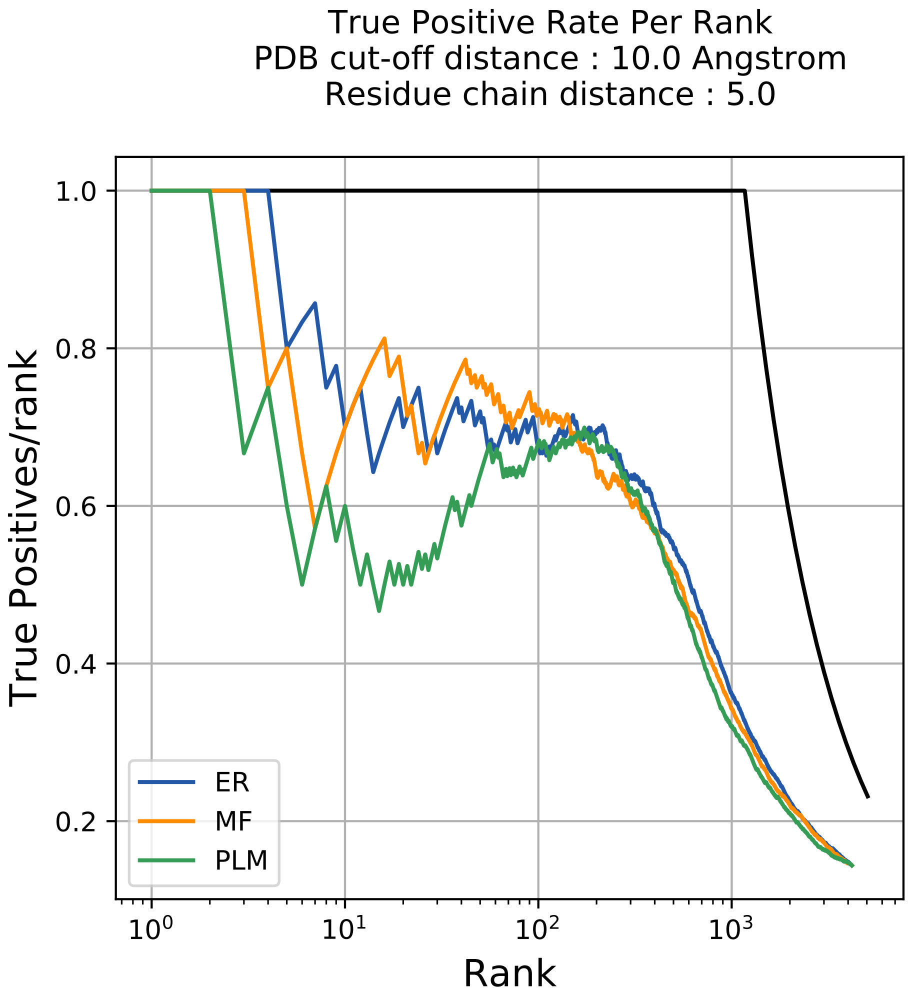

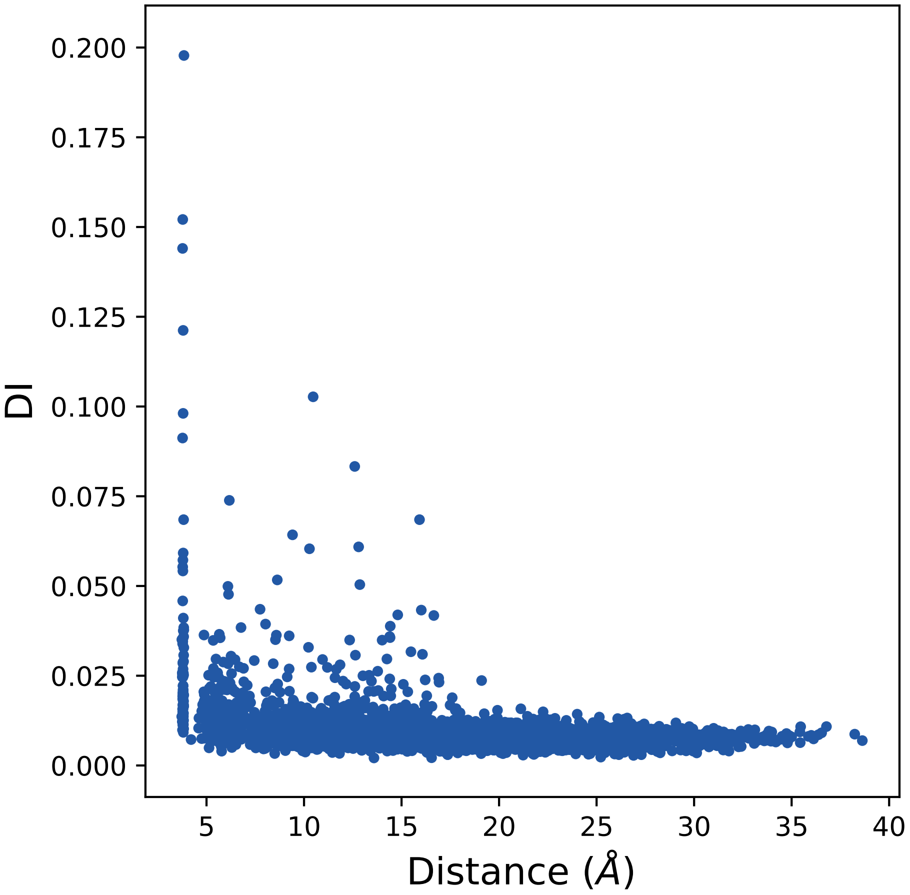

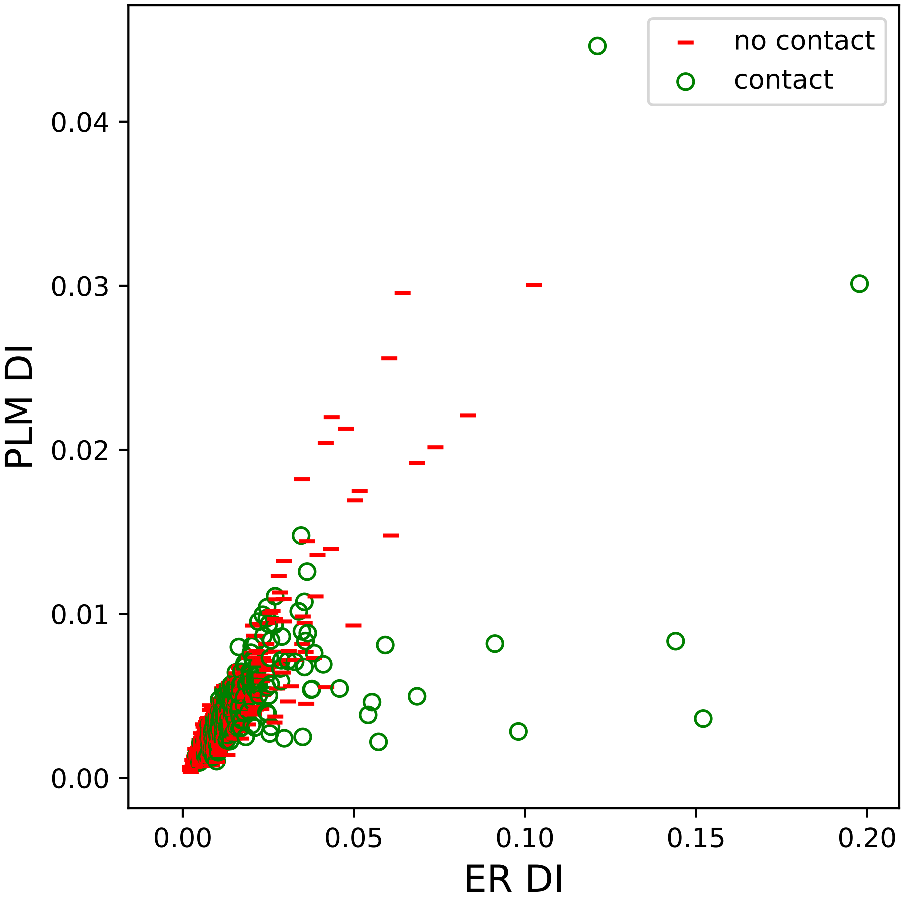

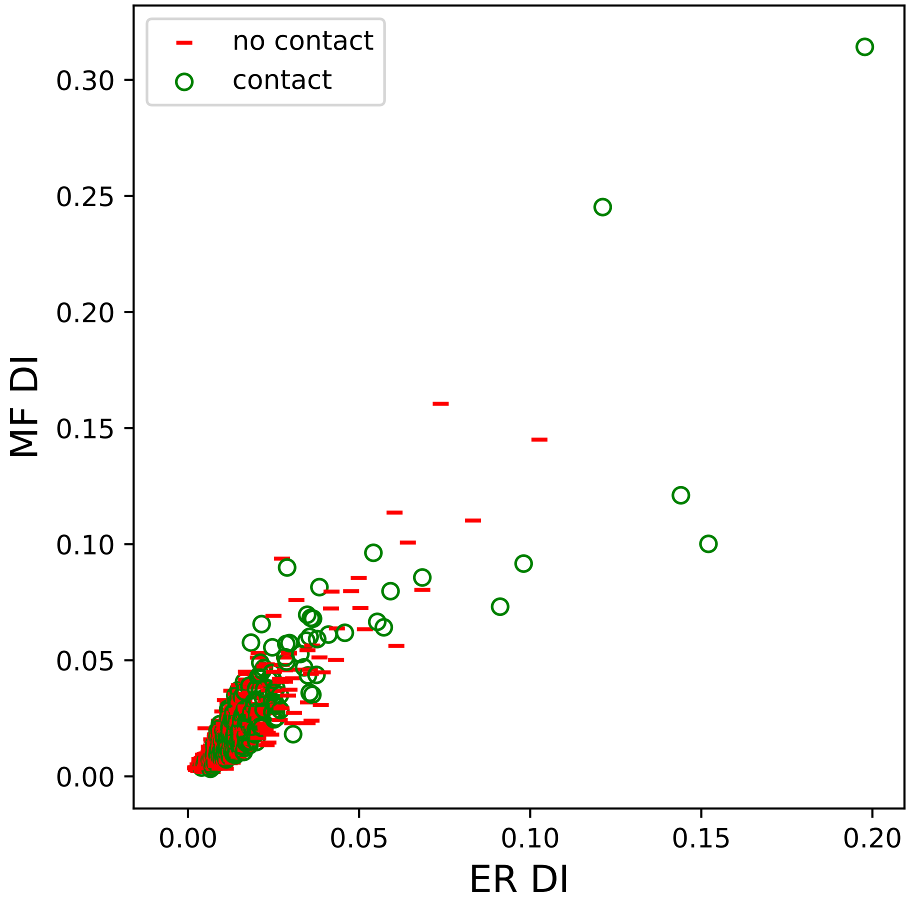

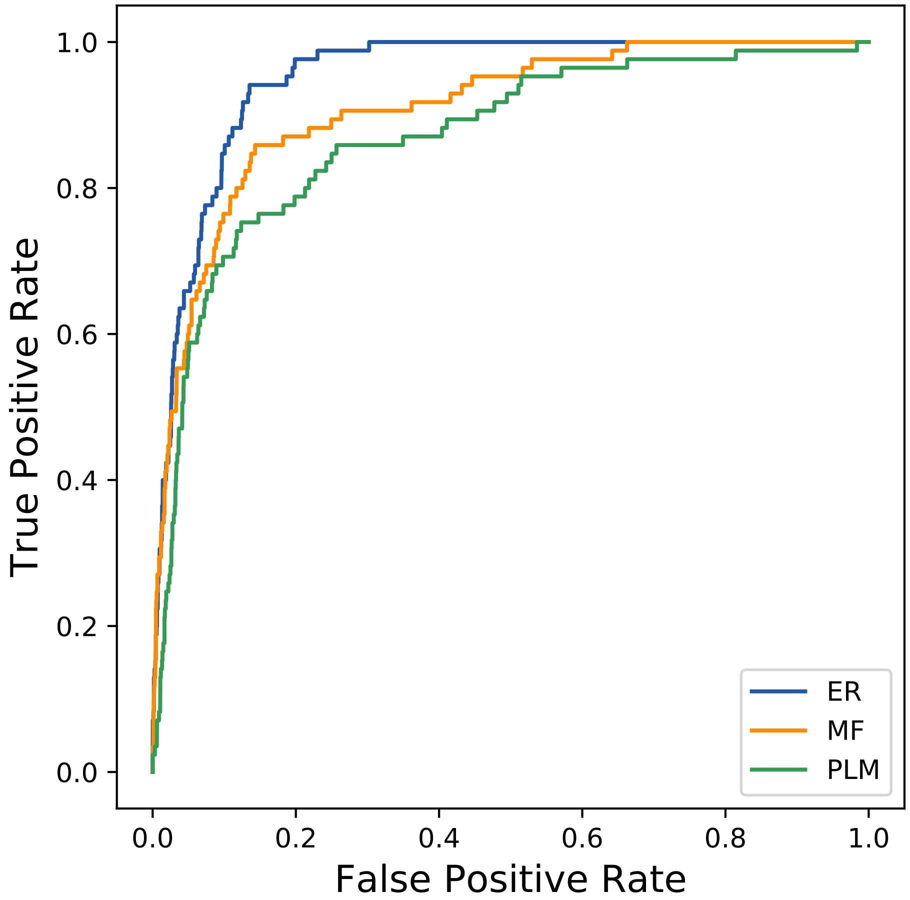

### 10112seq_110col_6deb_PF02882_plots.pdf

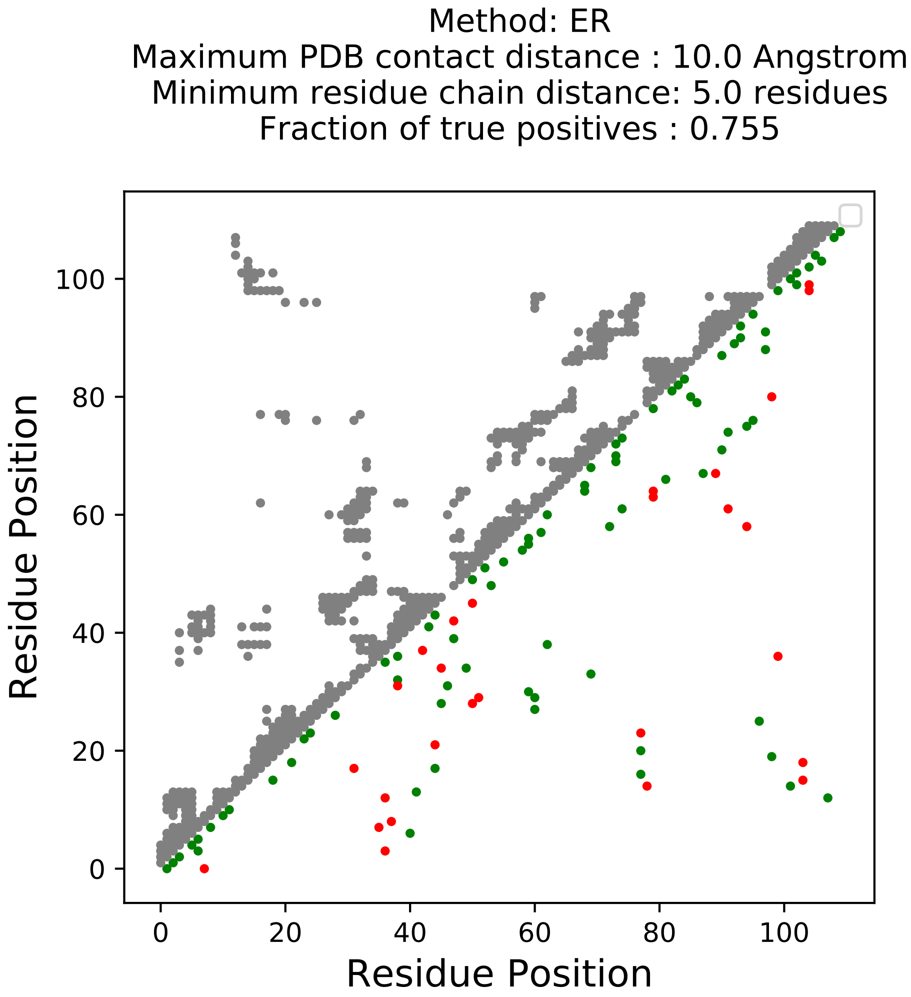

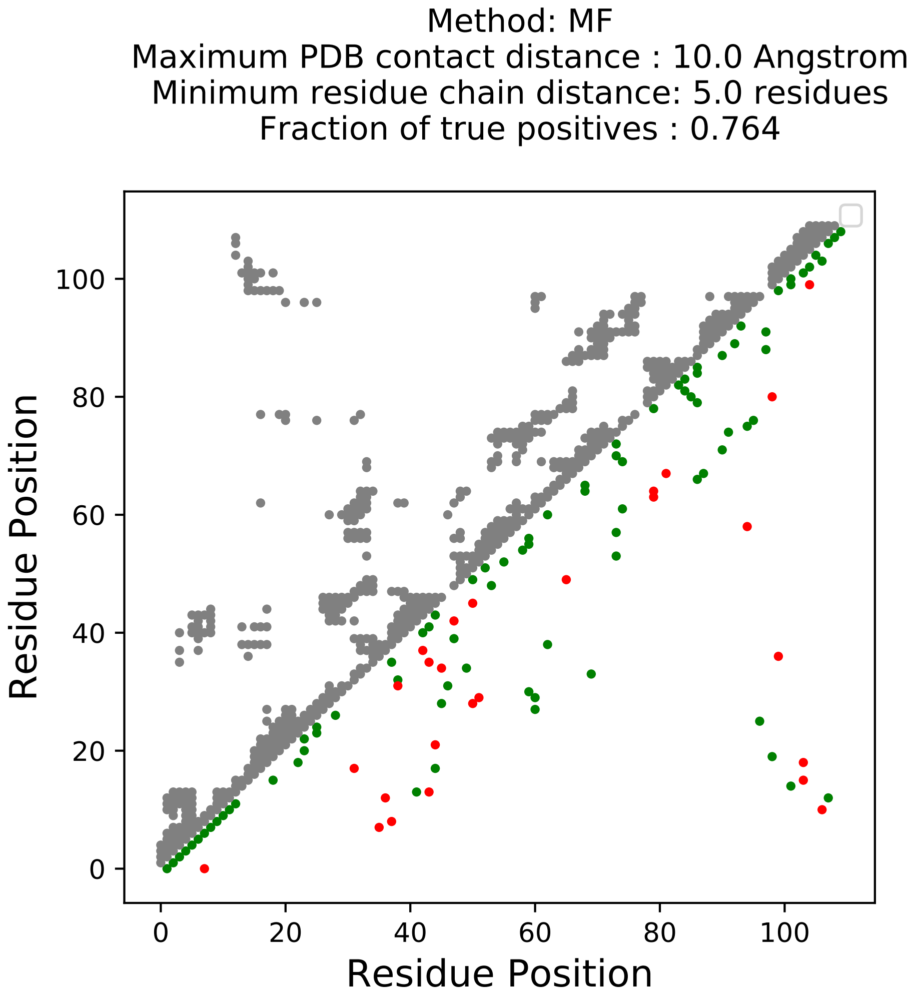

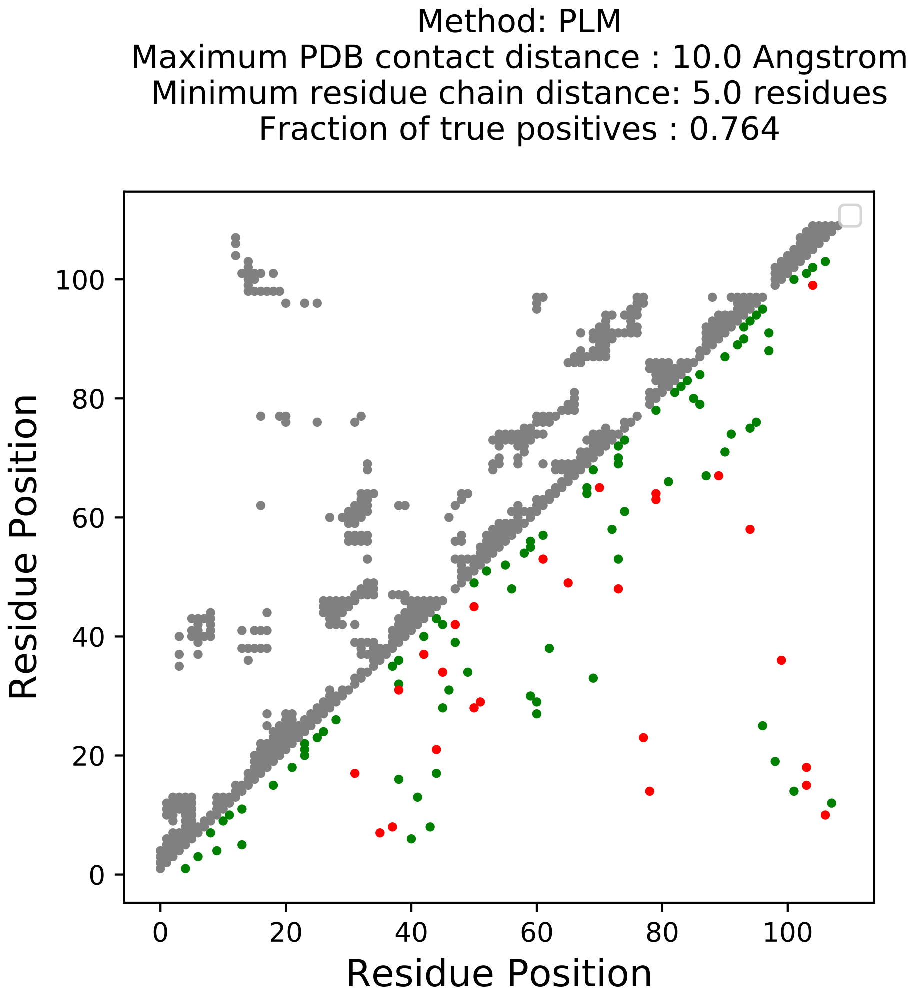

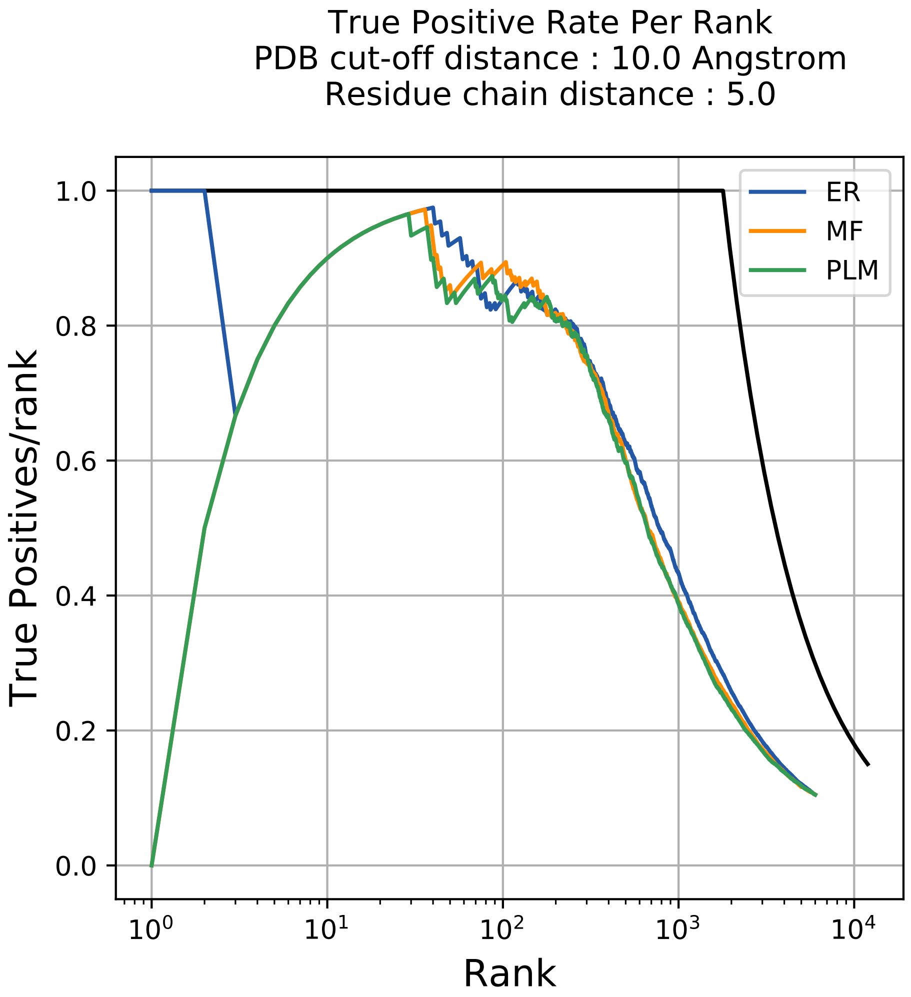

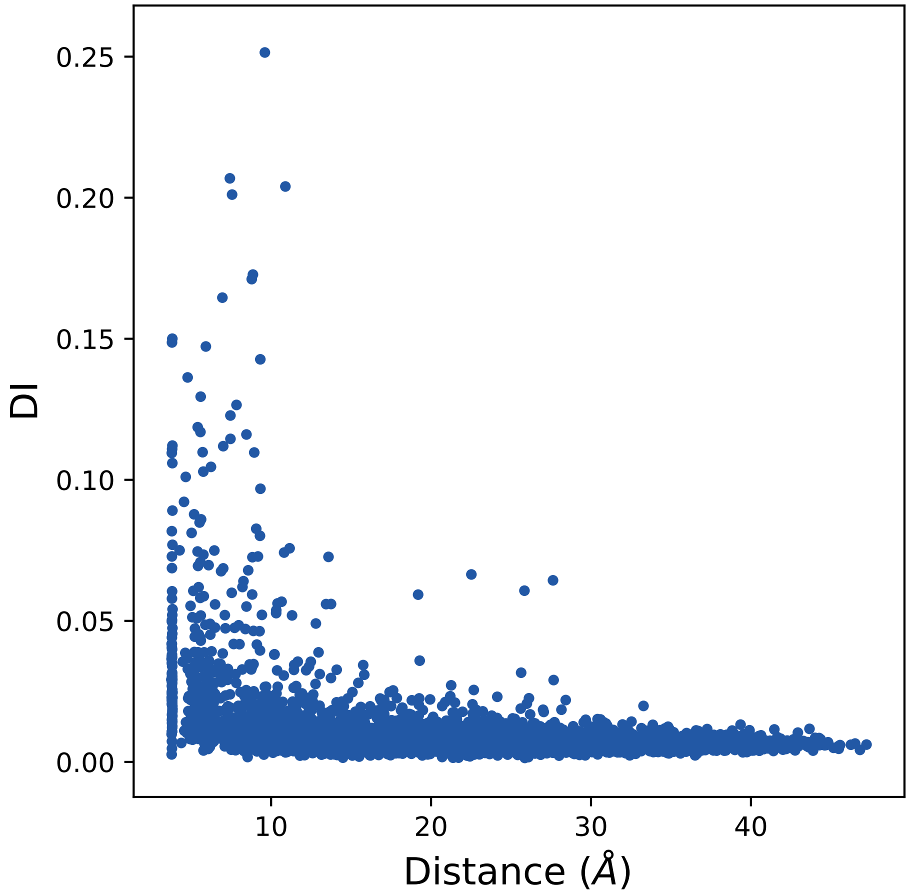

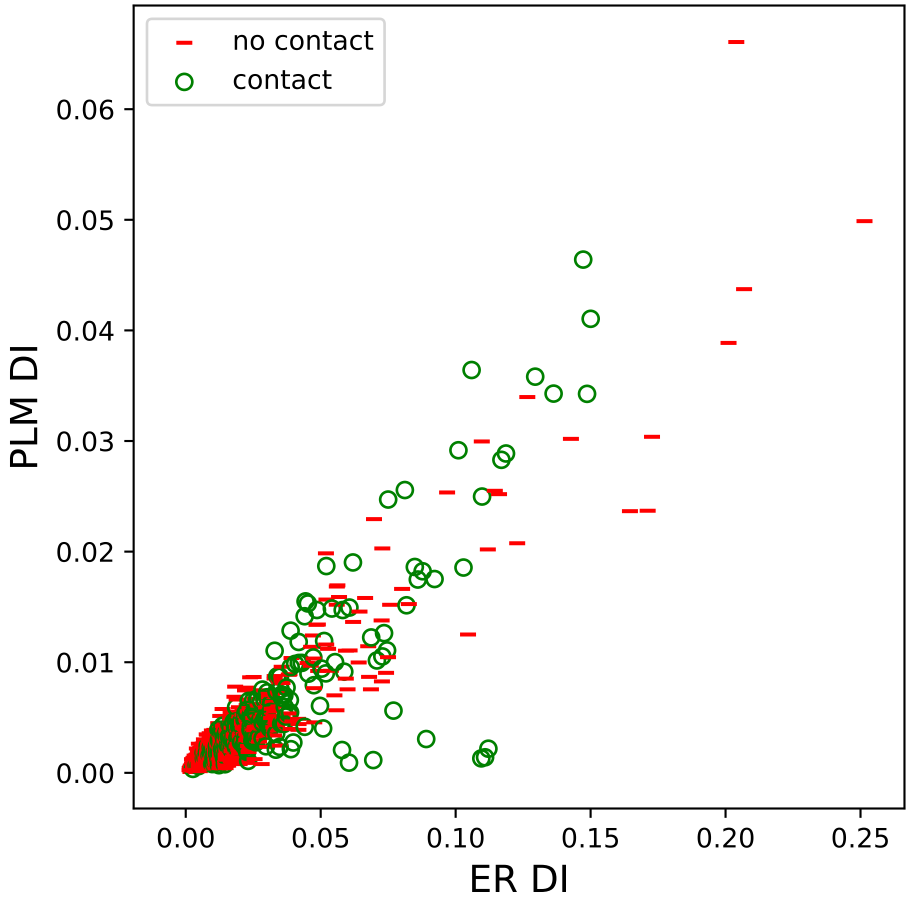

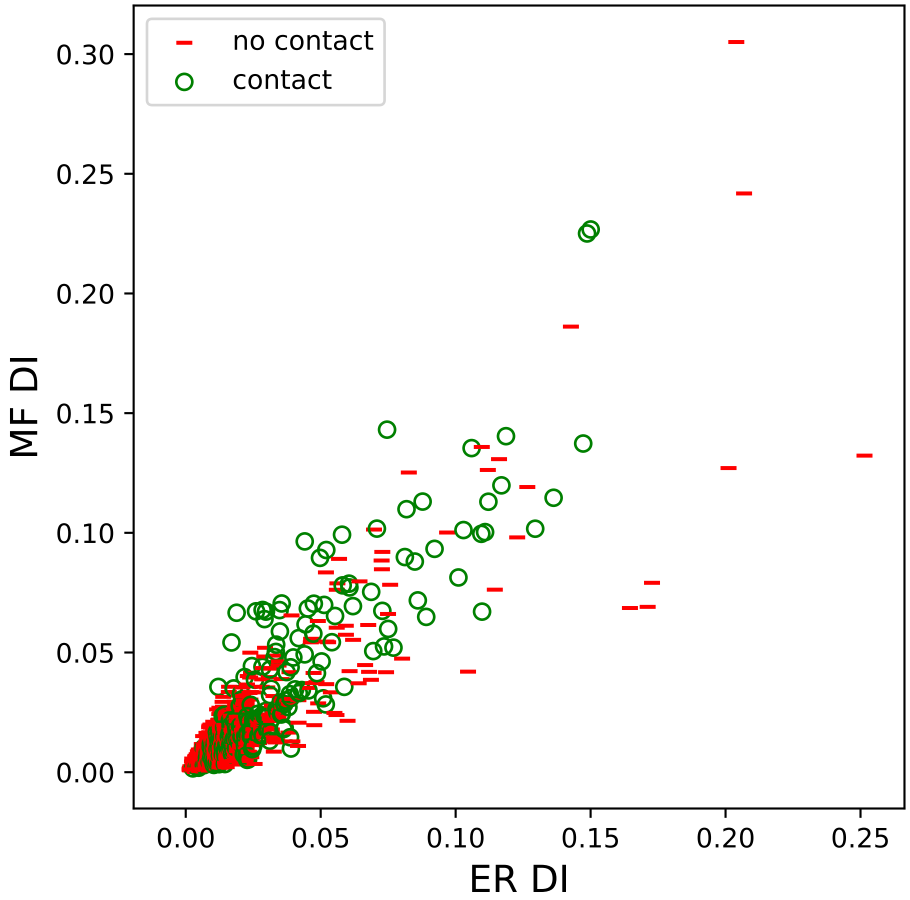

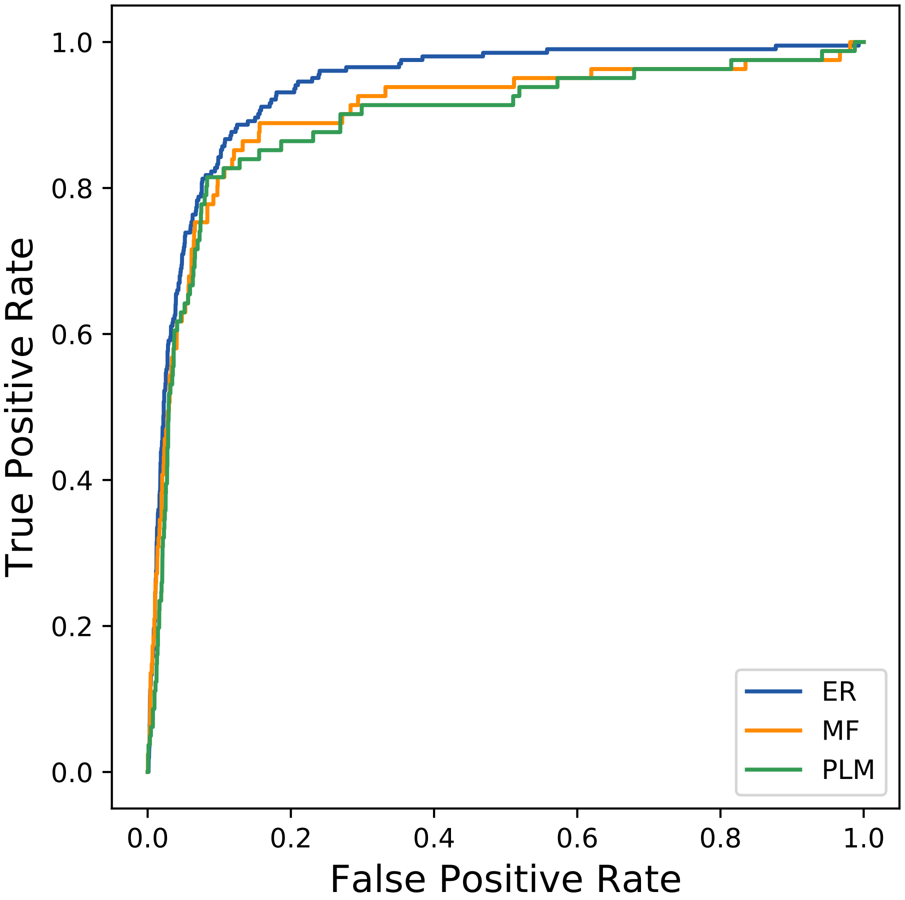

### 10175seq_225col_5rd3_PF00026_plots.pdf

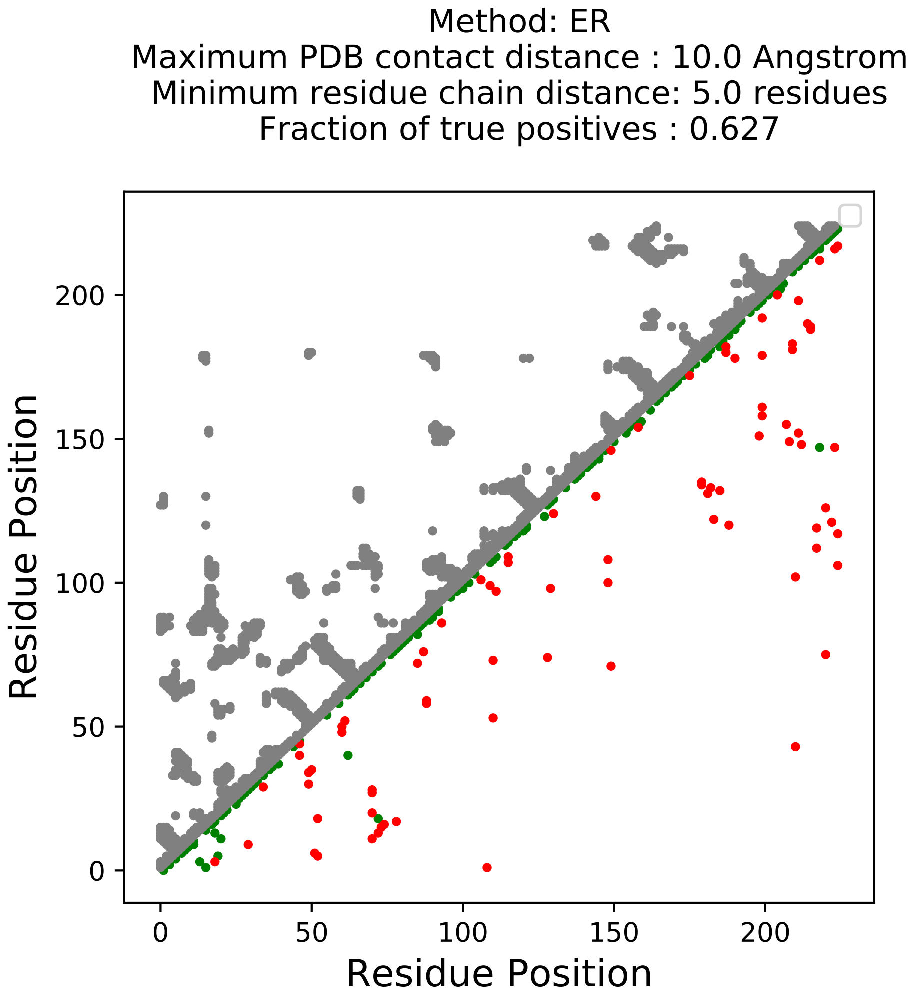

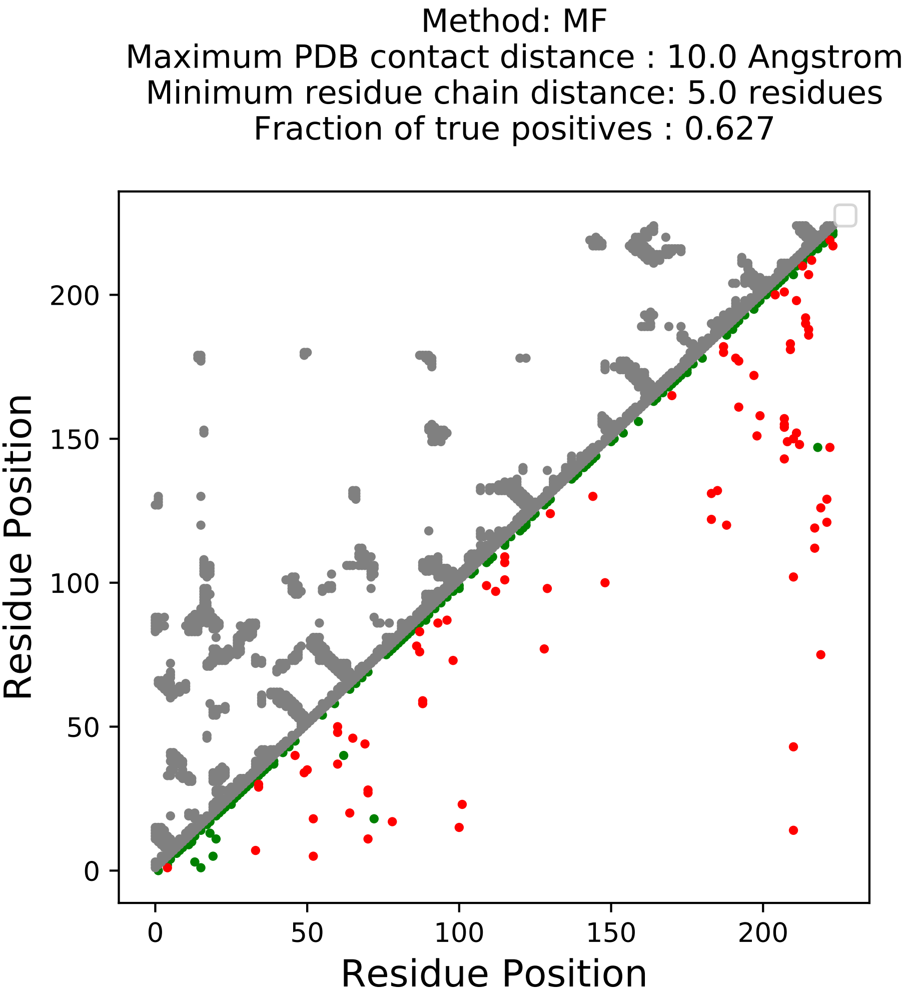

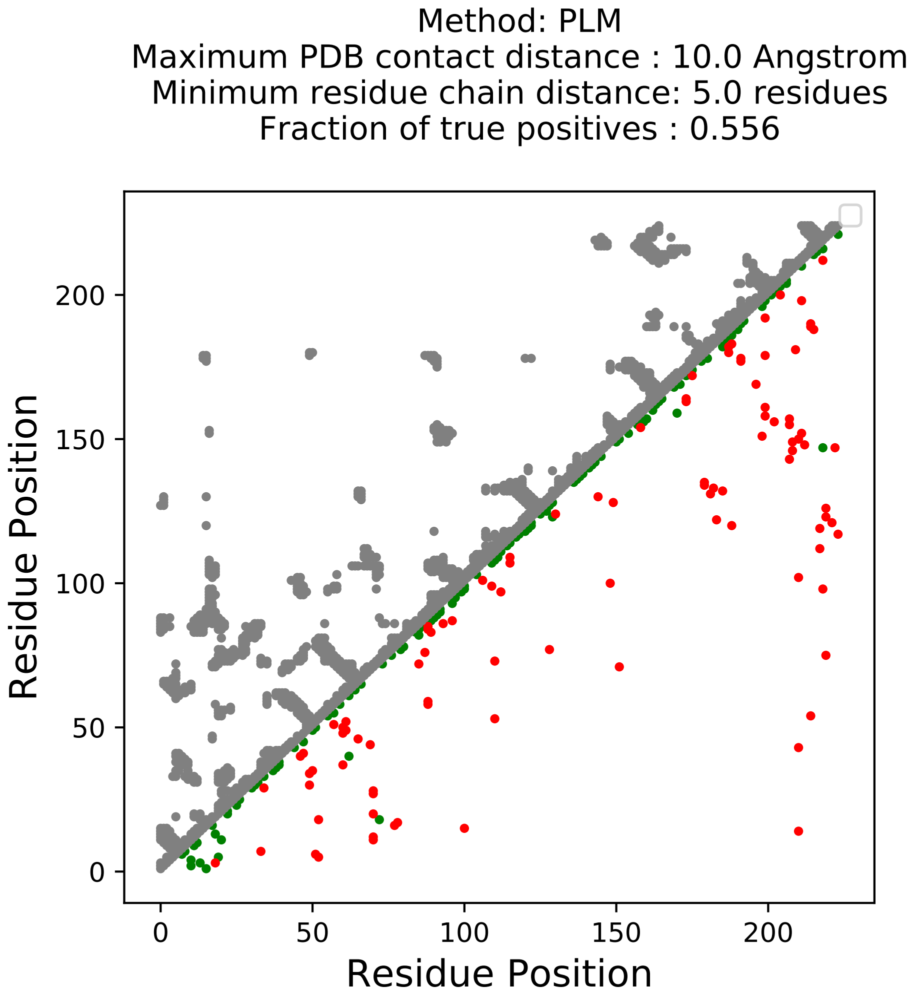

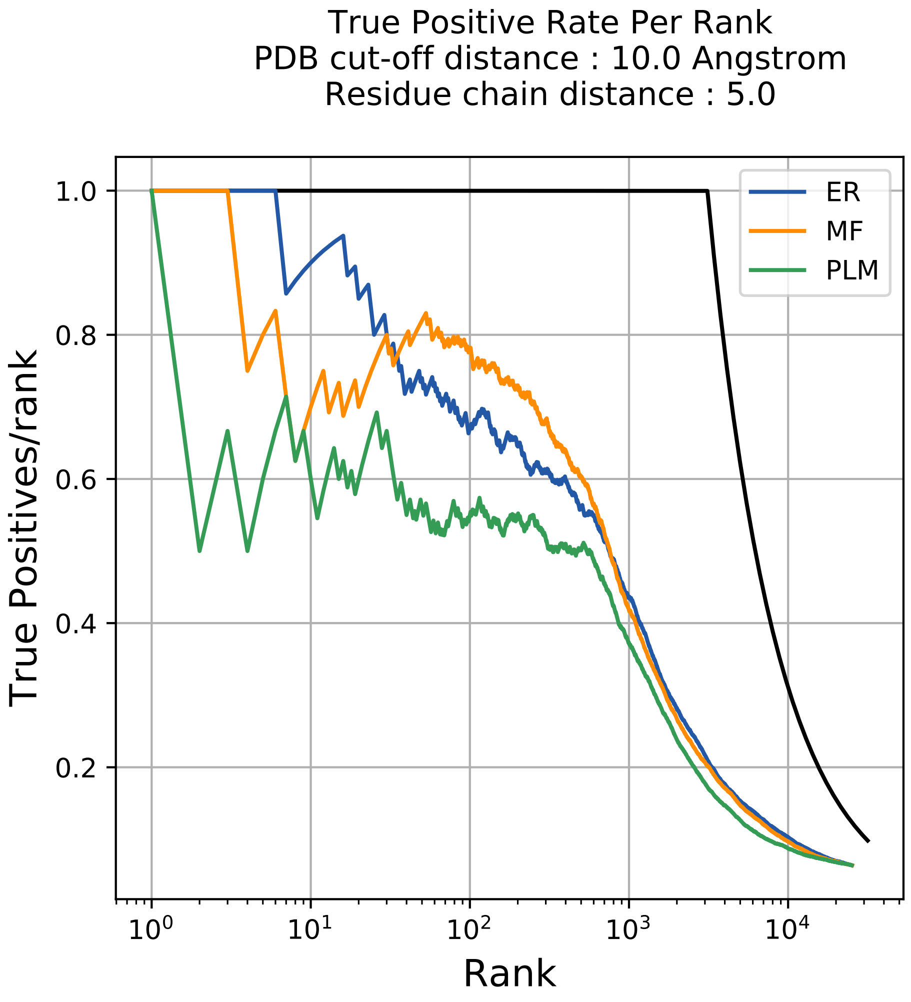

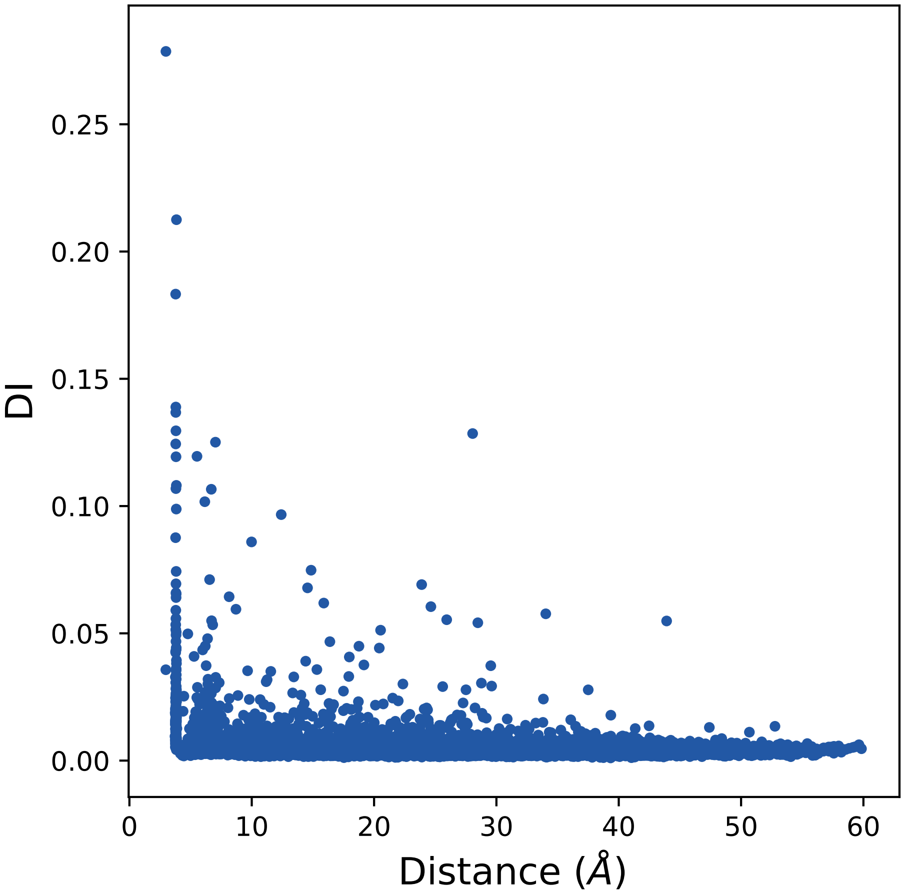

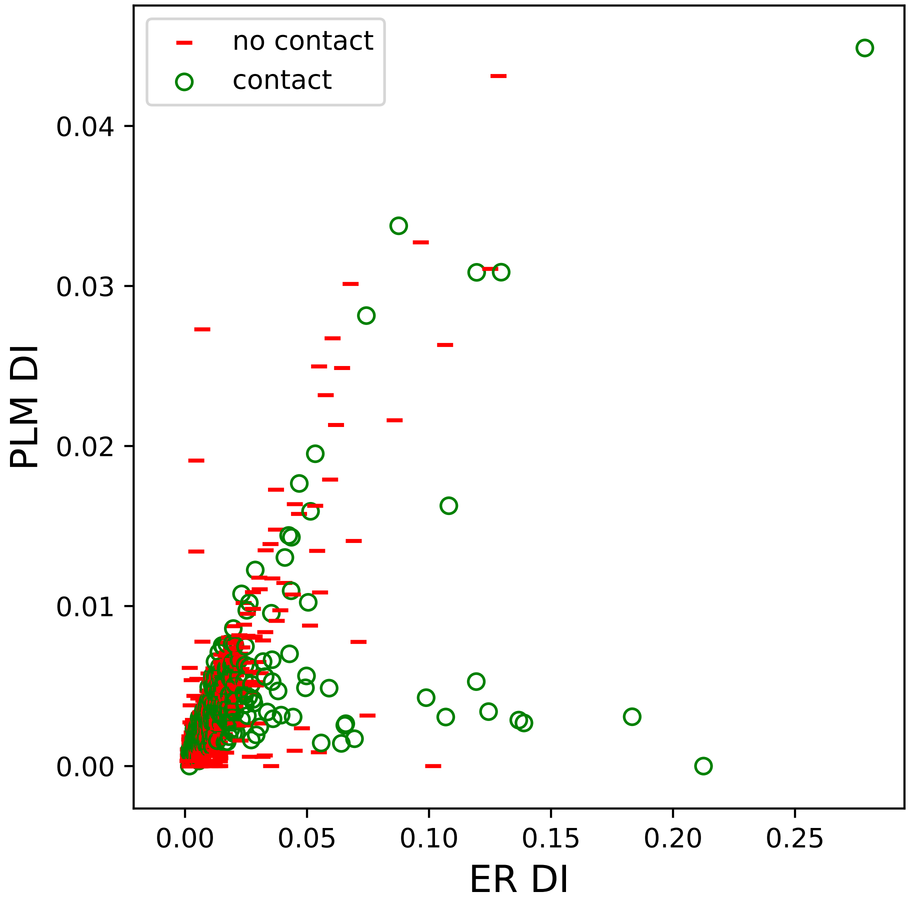

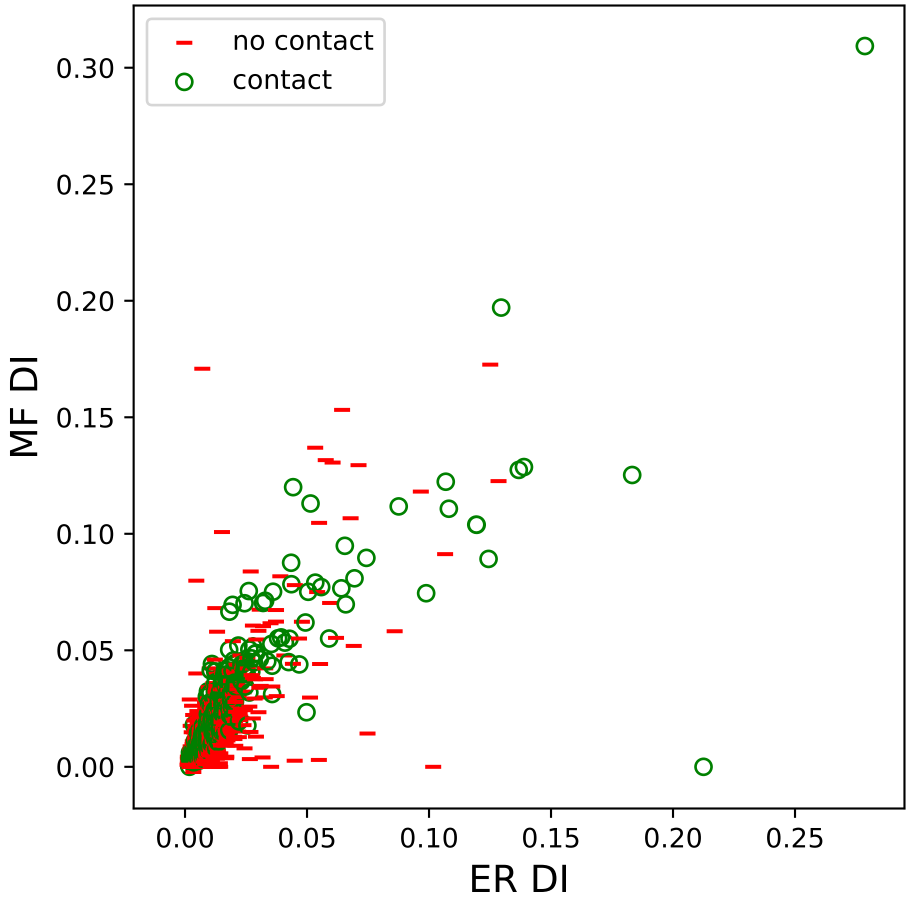

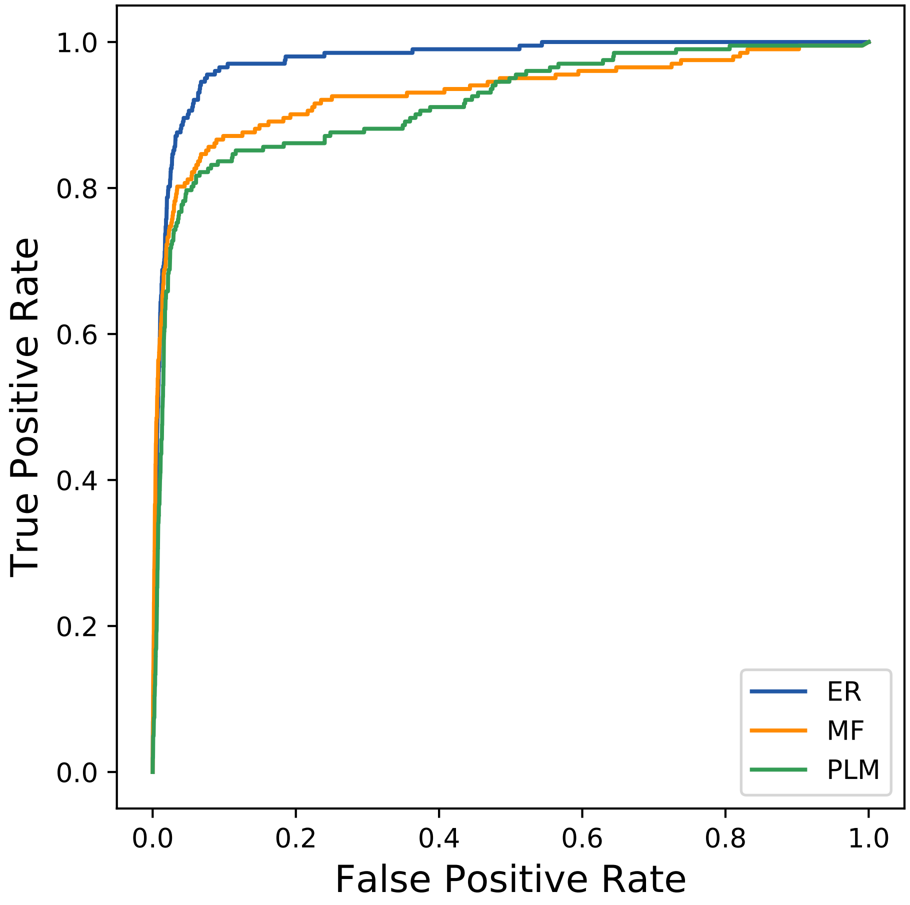

### 10249seq_118col_2irz_PF00026_plots.pdf

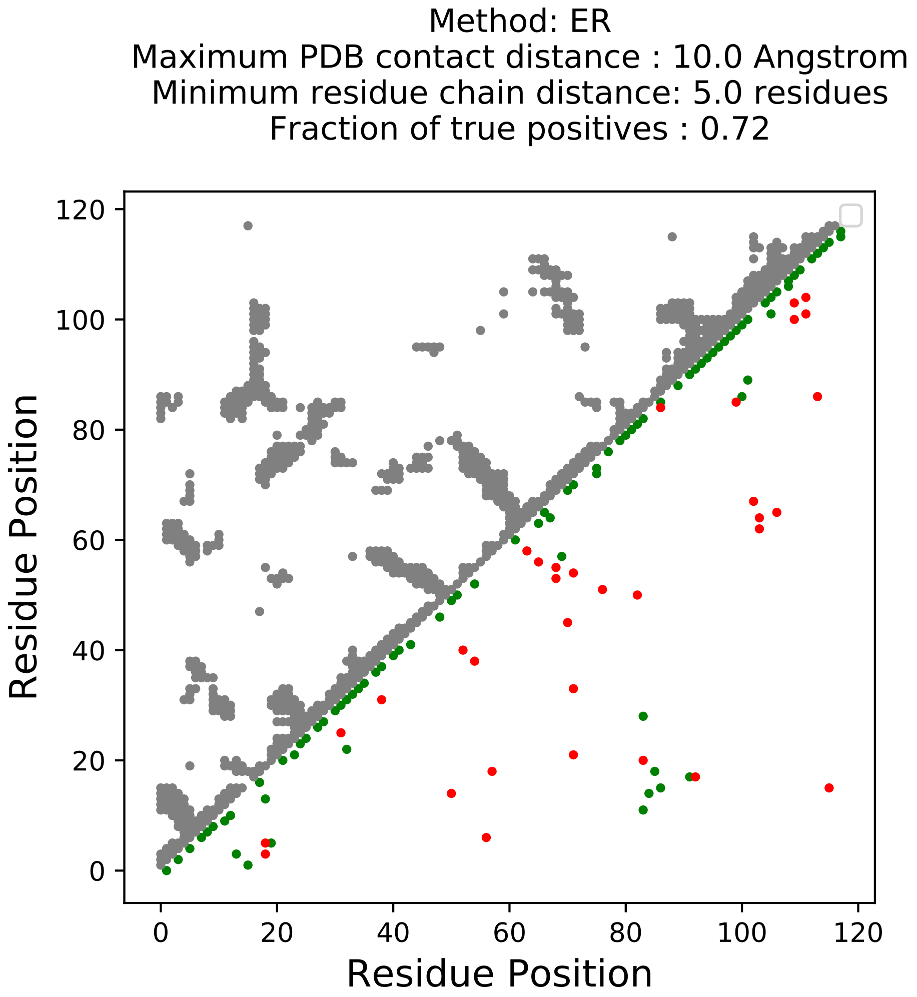

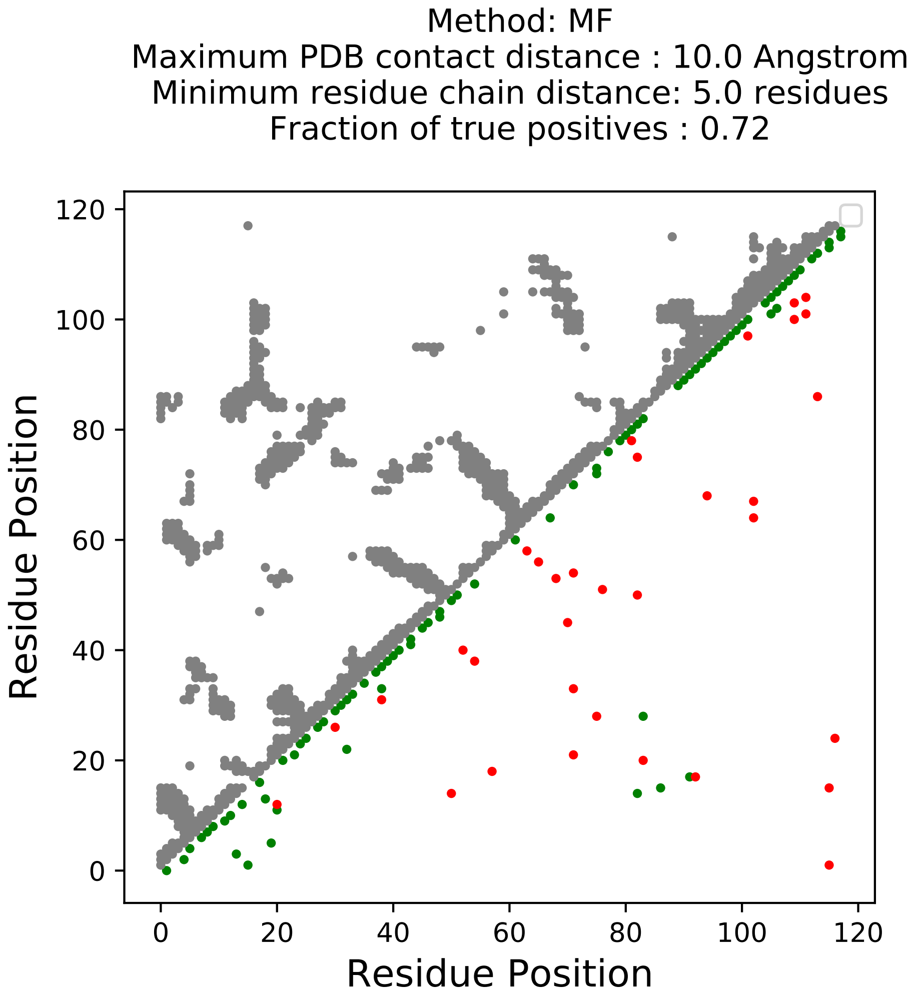

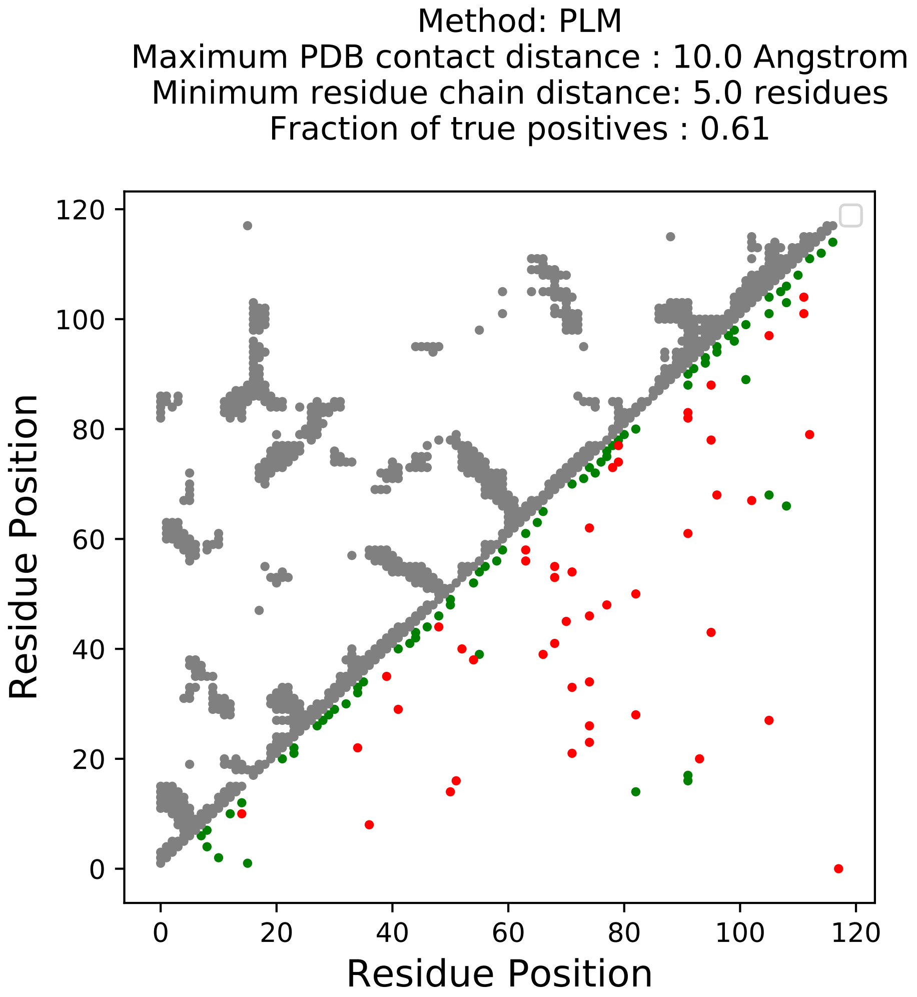

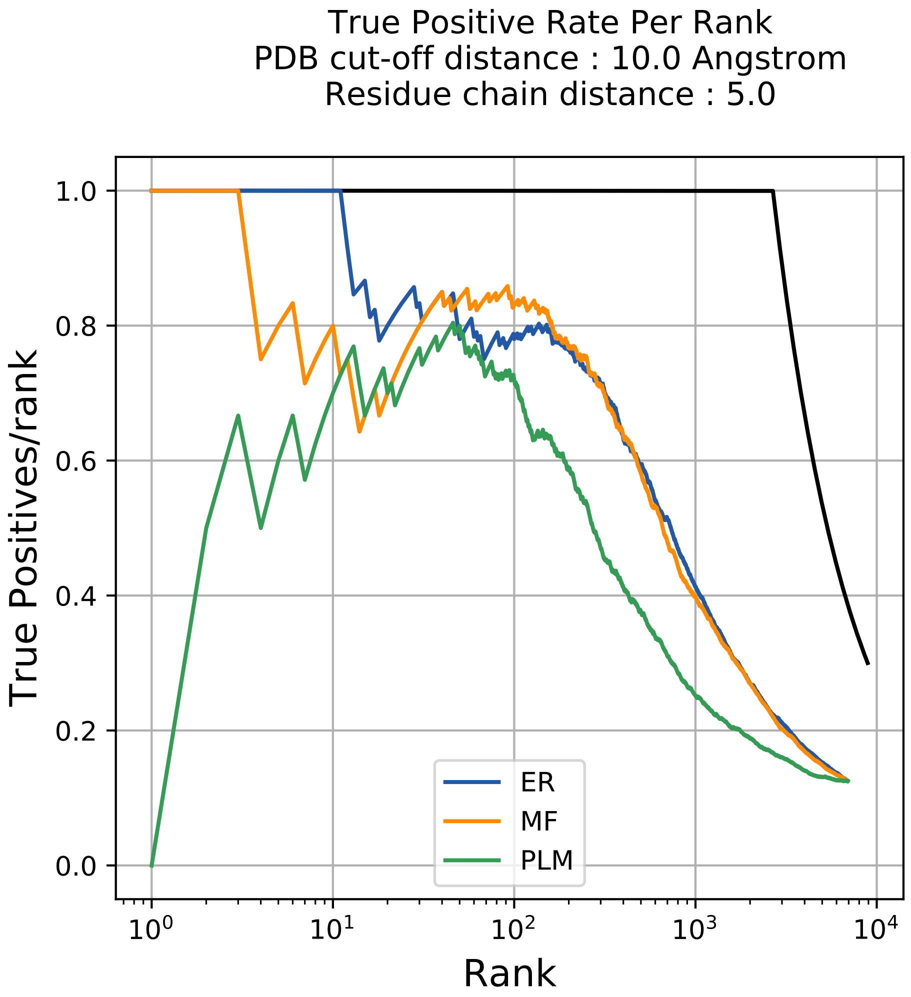

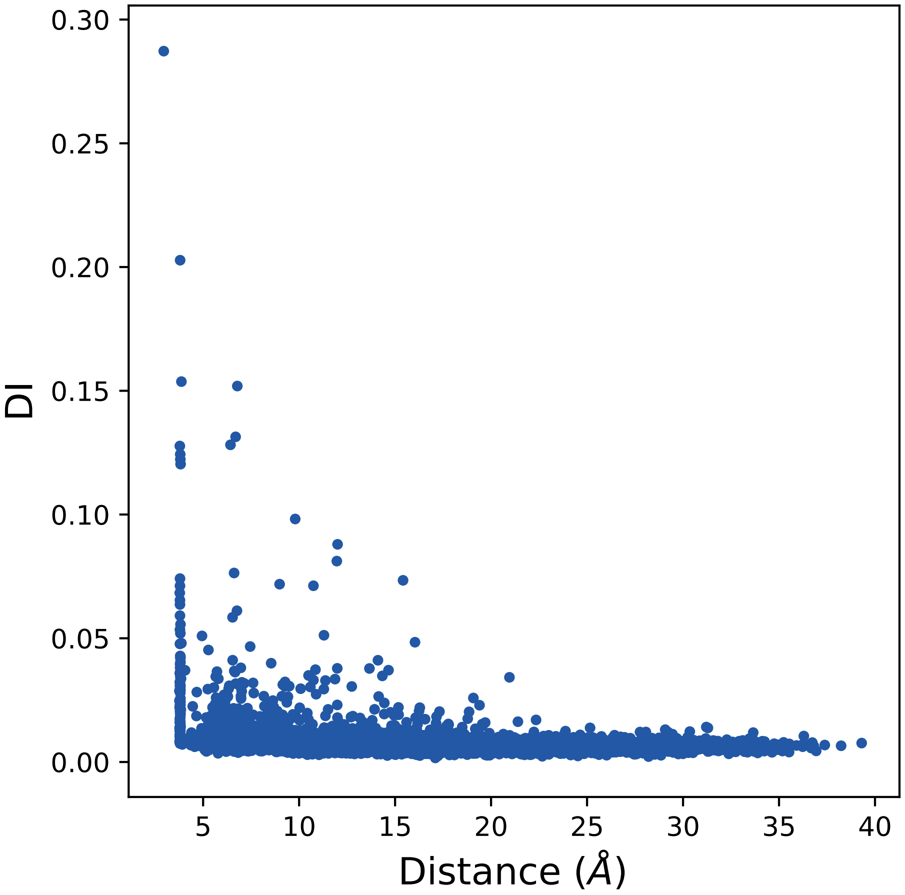

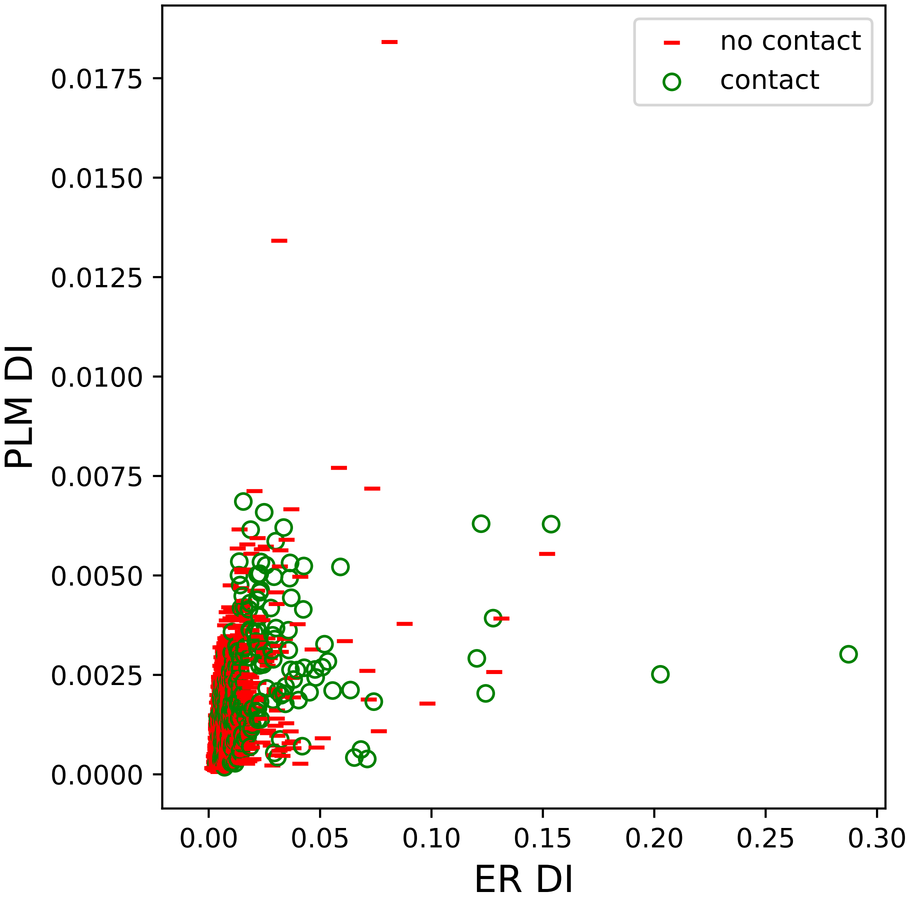

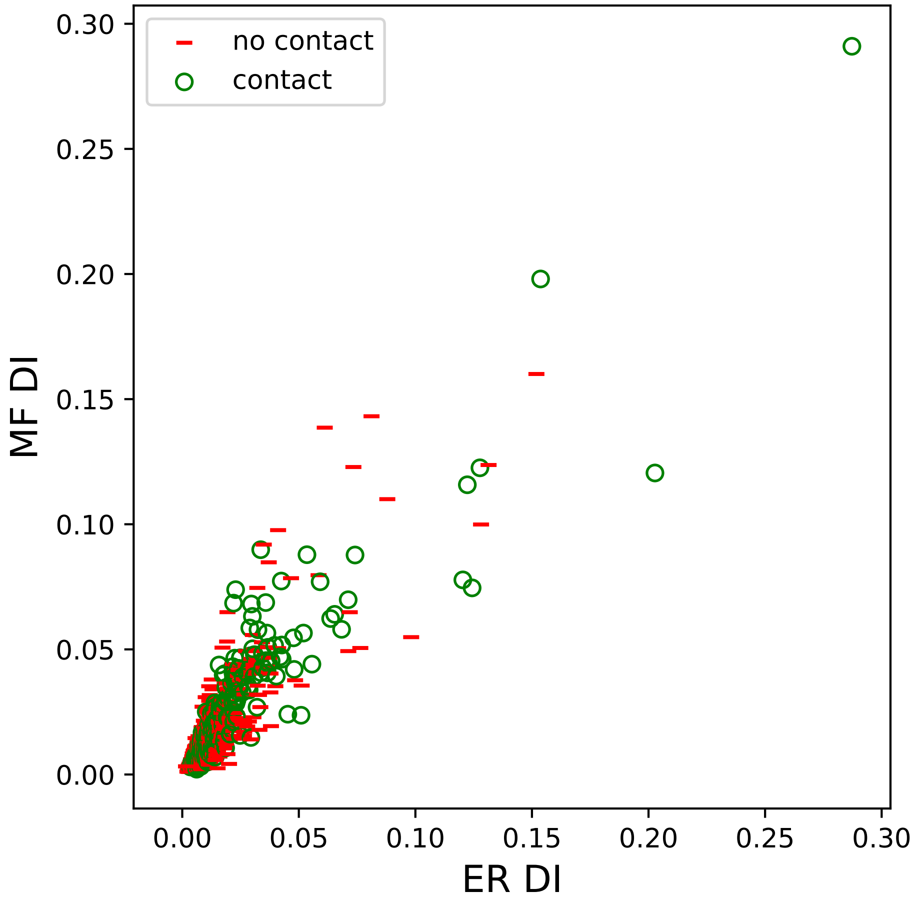

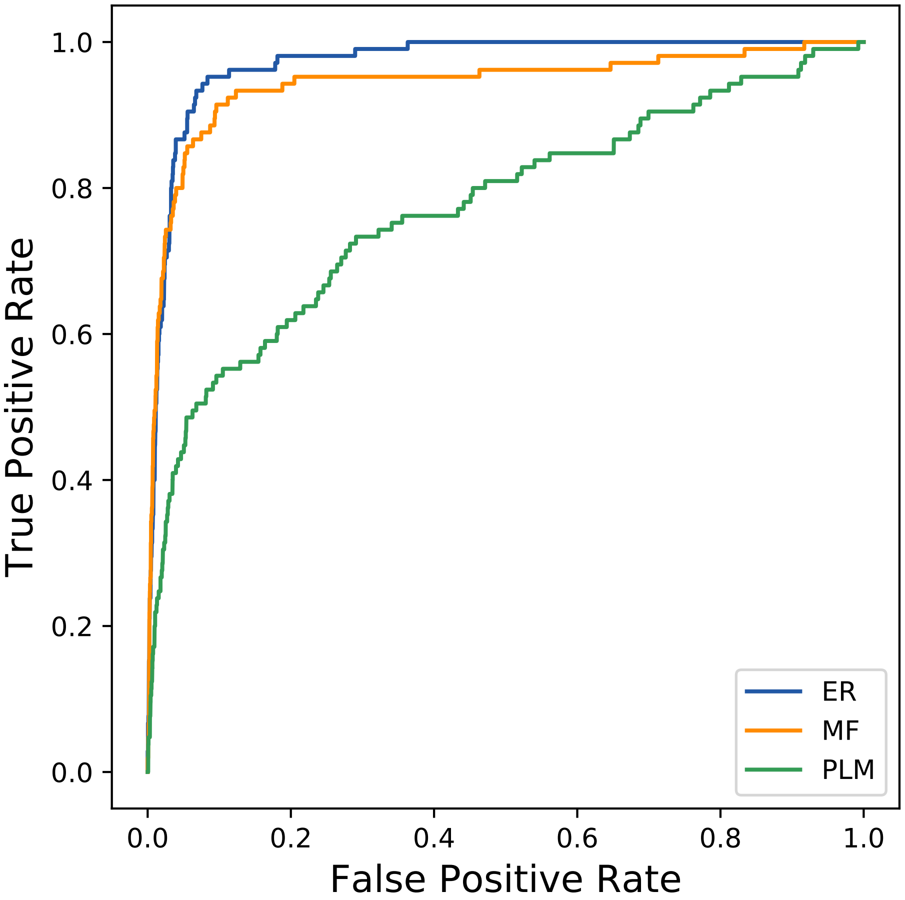

### 10251seq_68col_6ji1_PF00854_plots.pdf

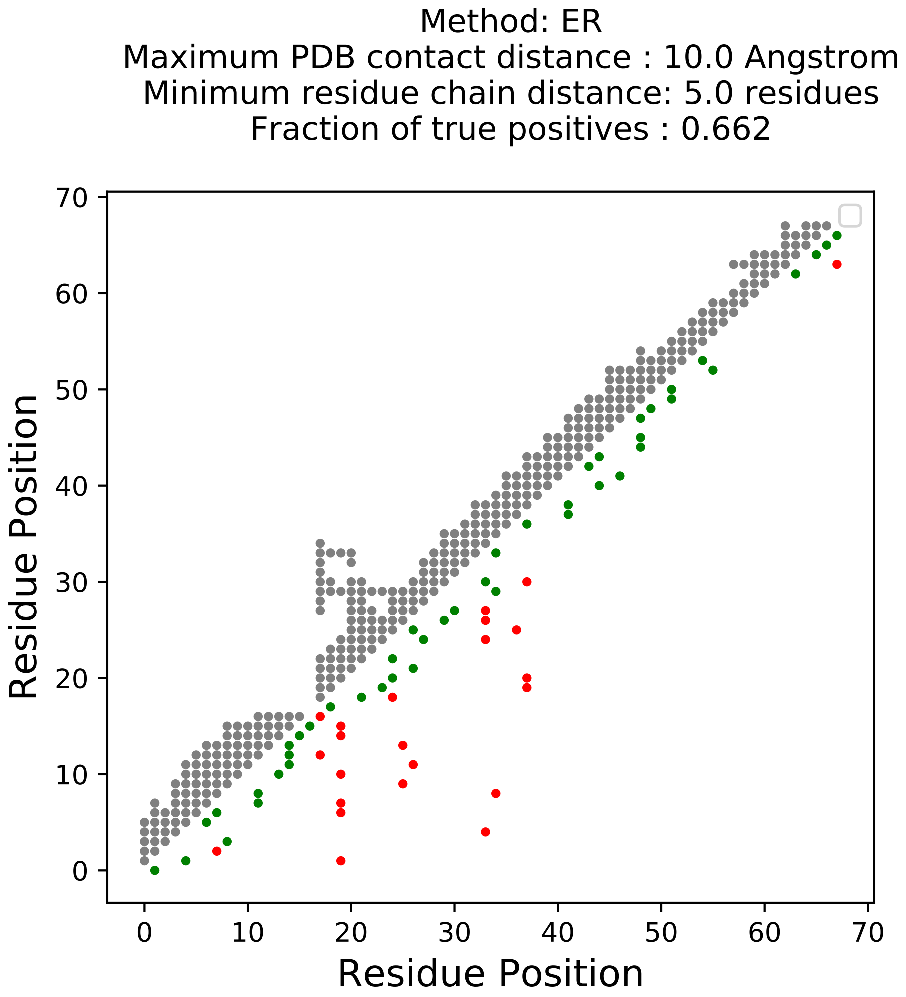

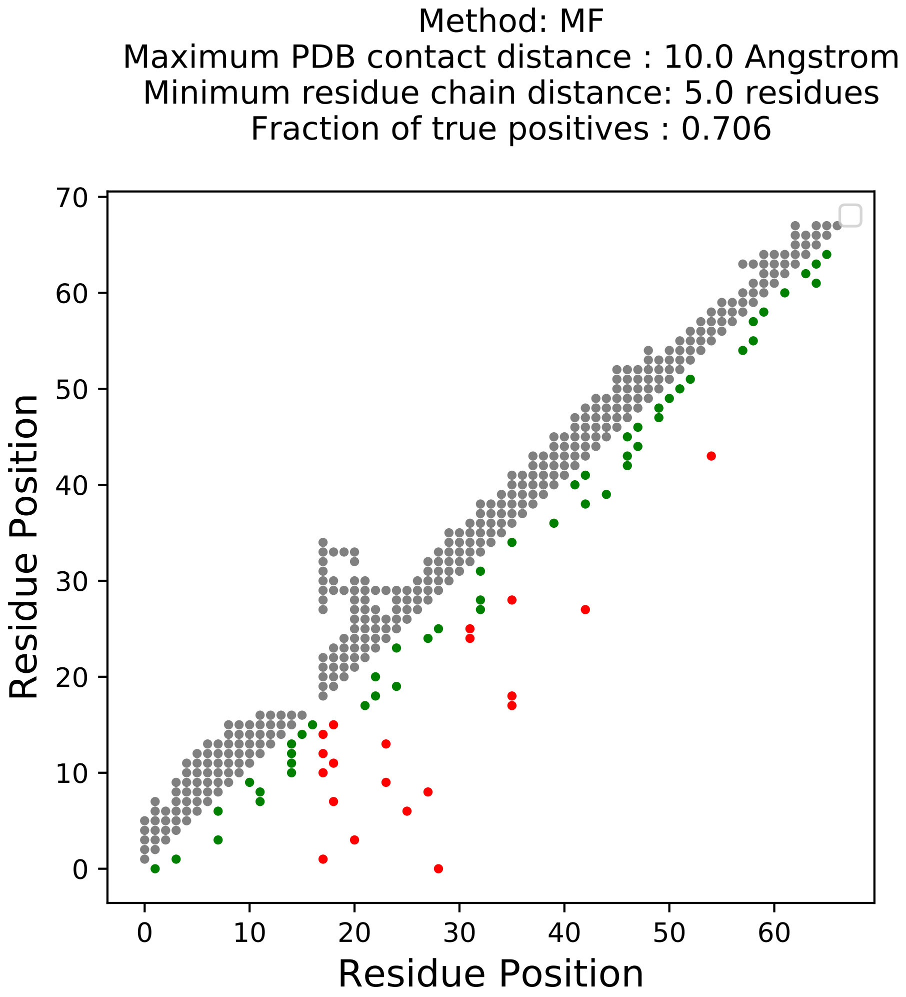

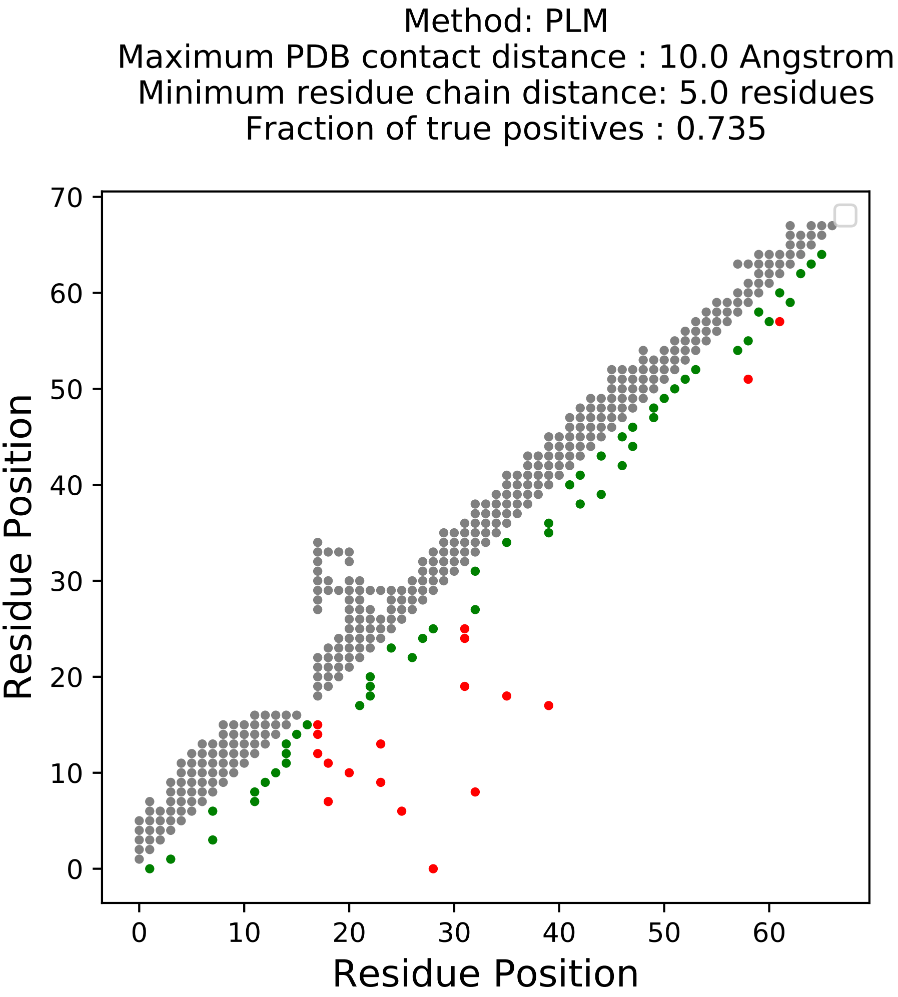

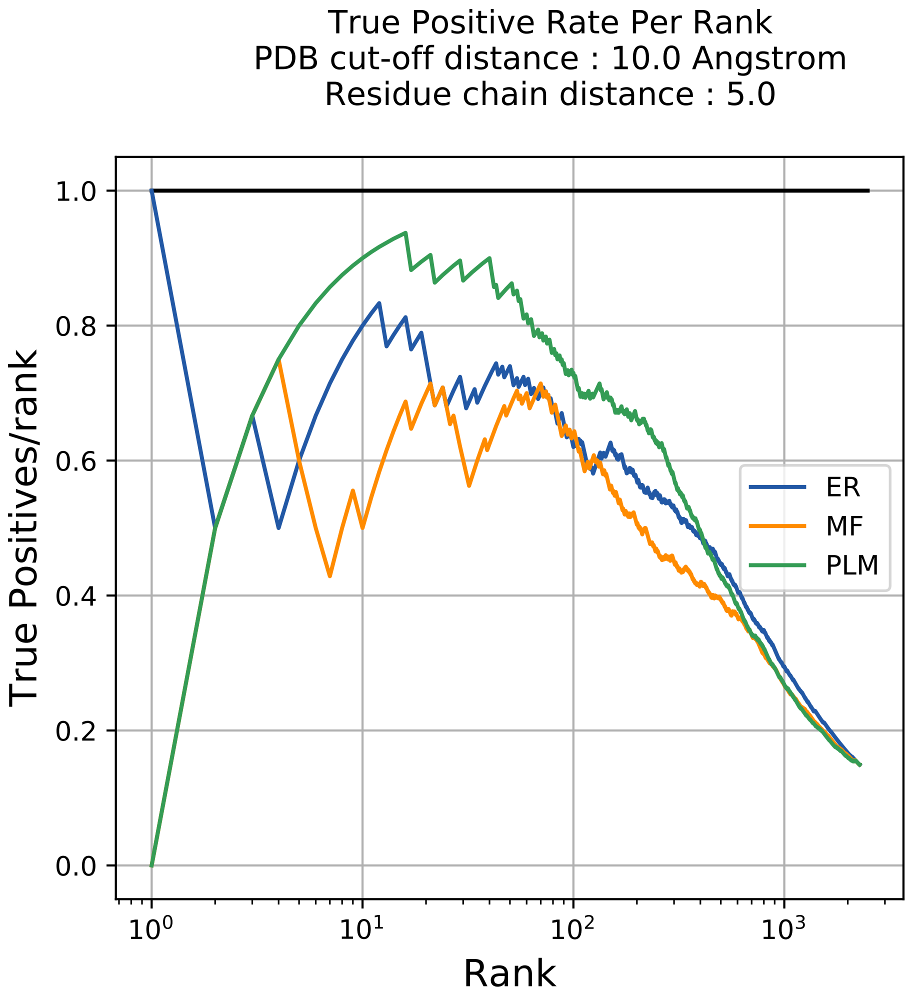

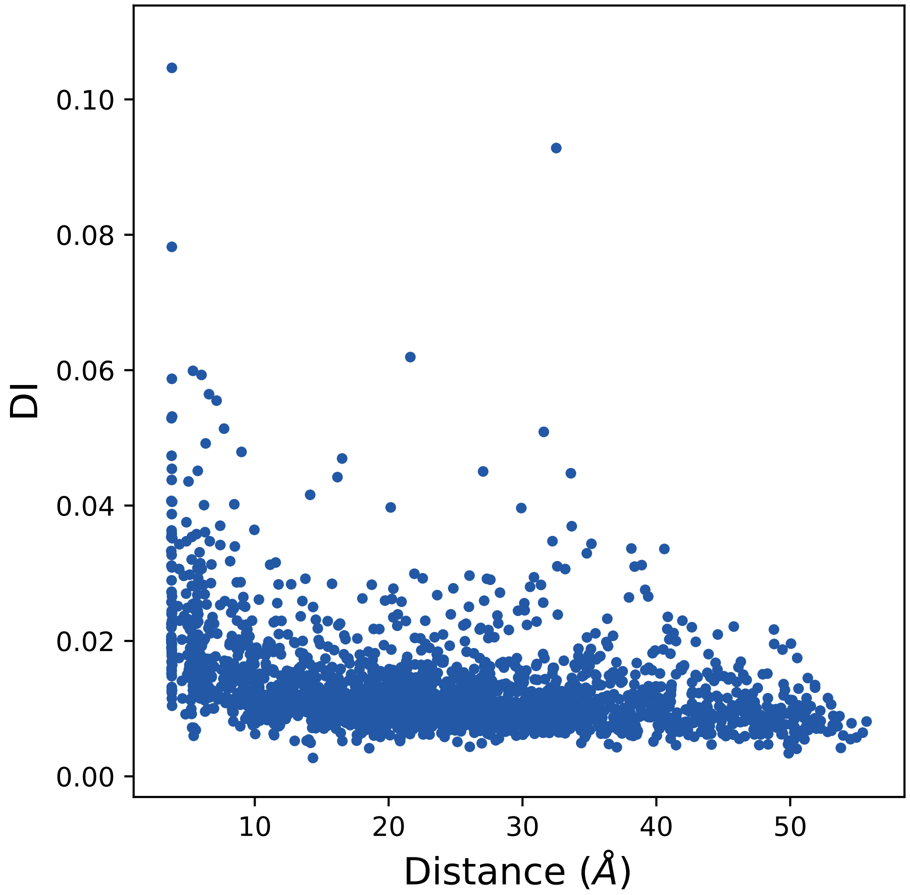

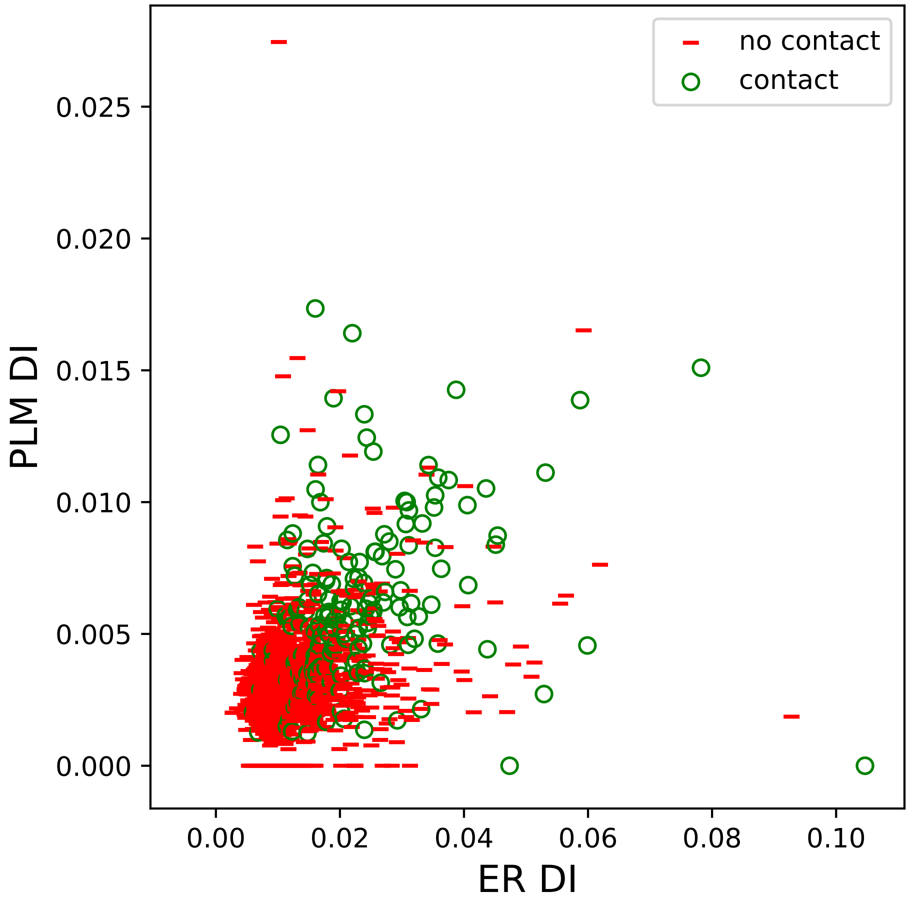

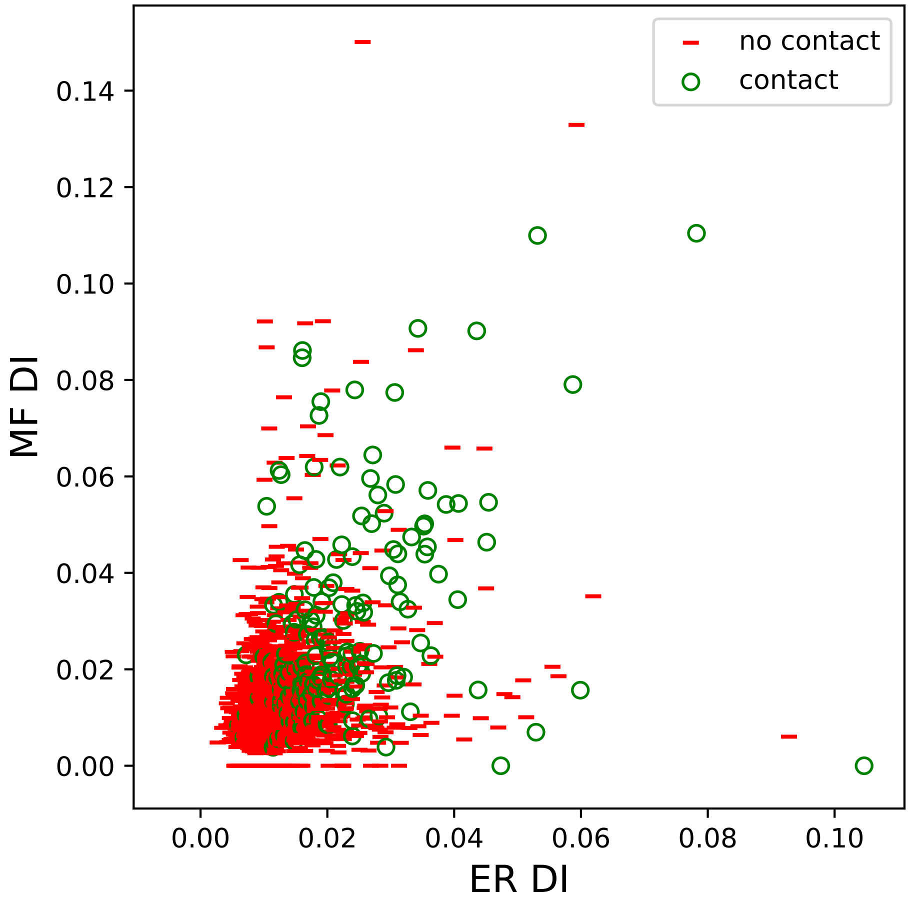

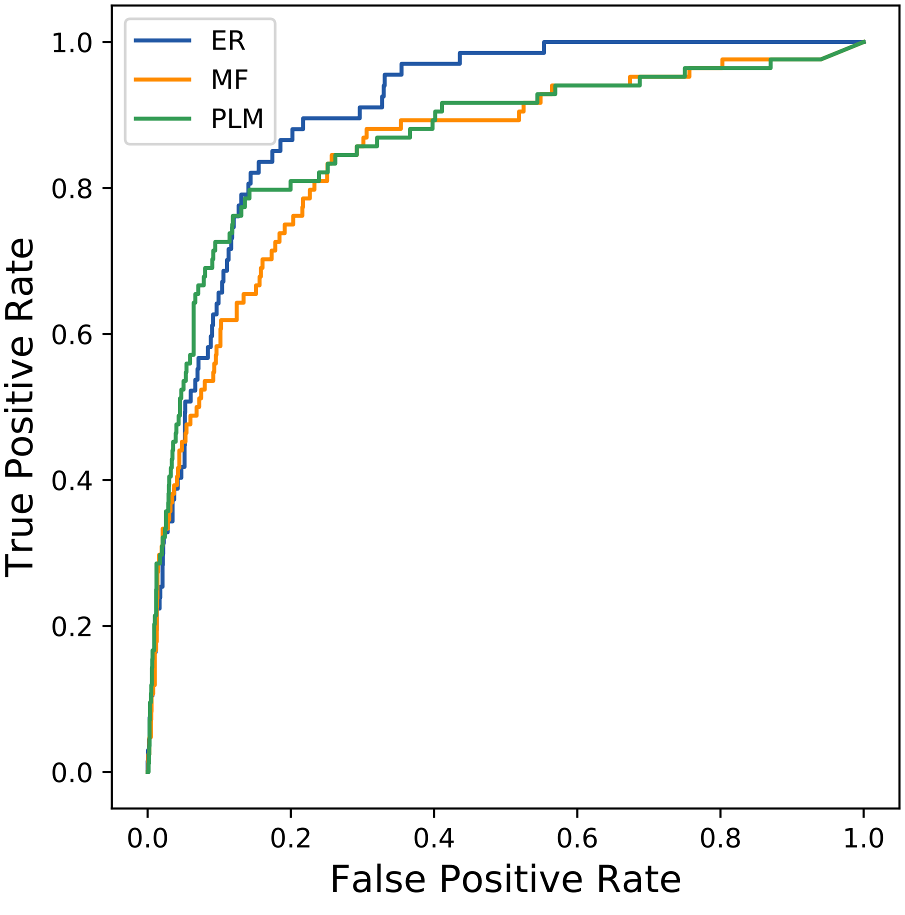

### 10286seq_120col_4j1i_PF00026_plots.pdf

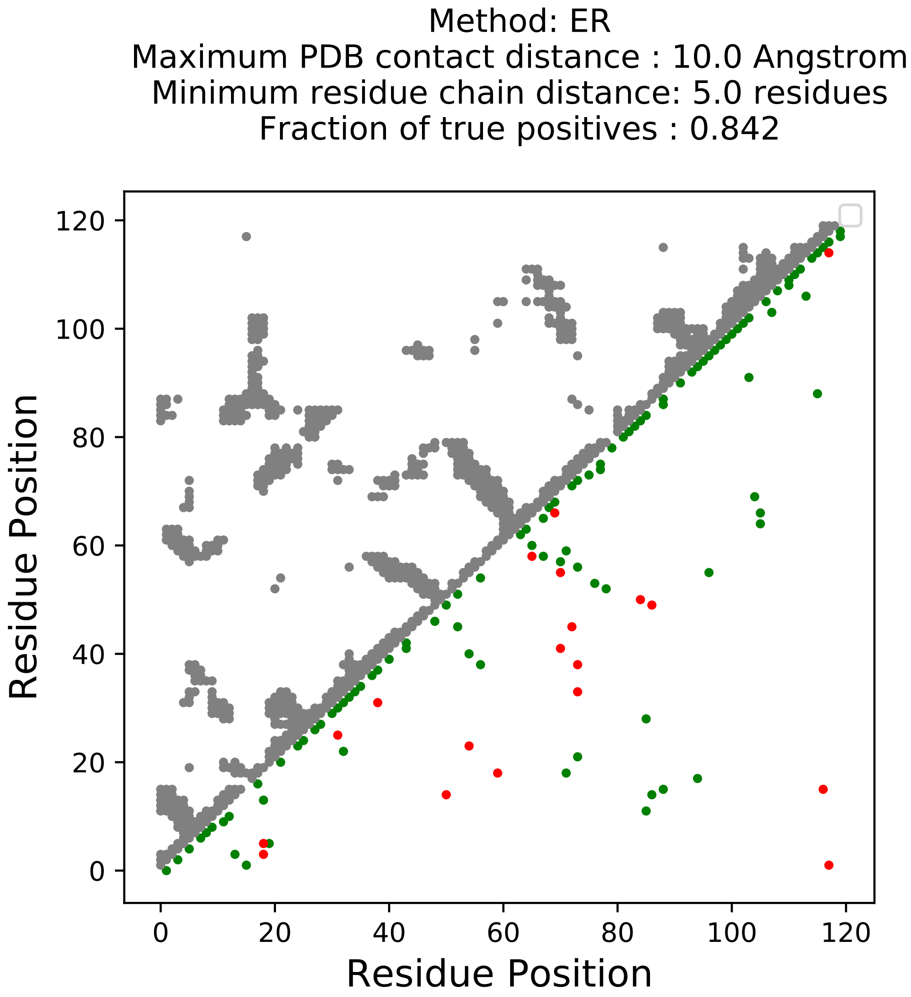

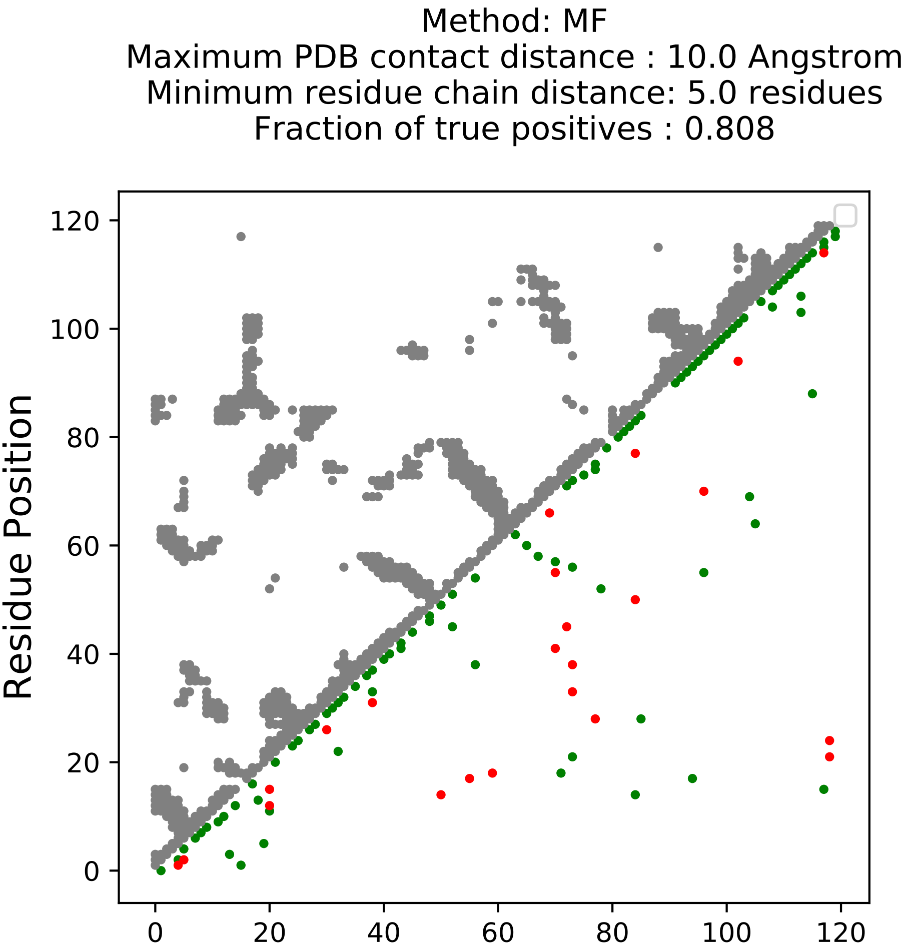

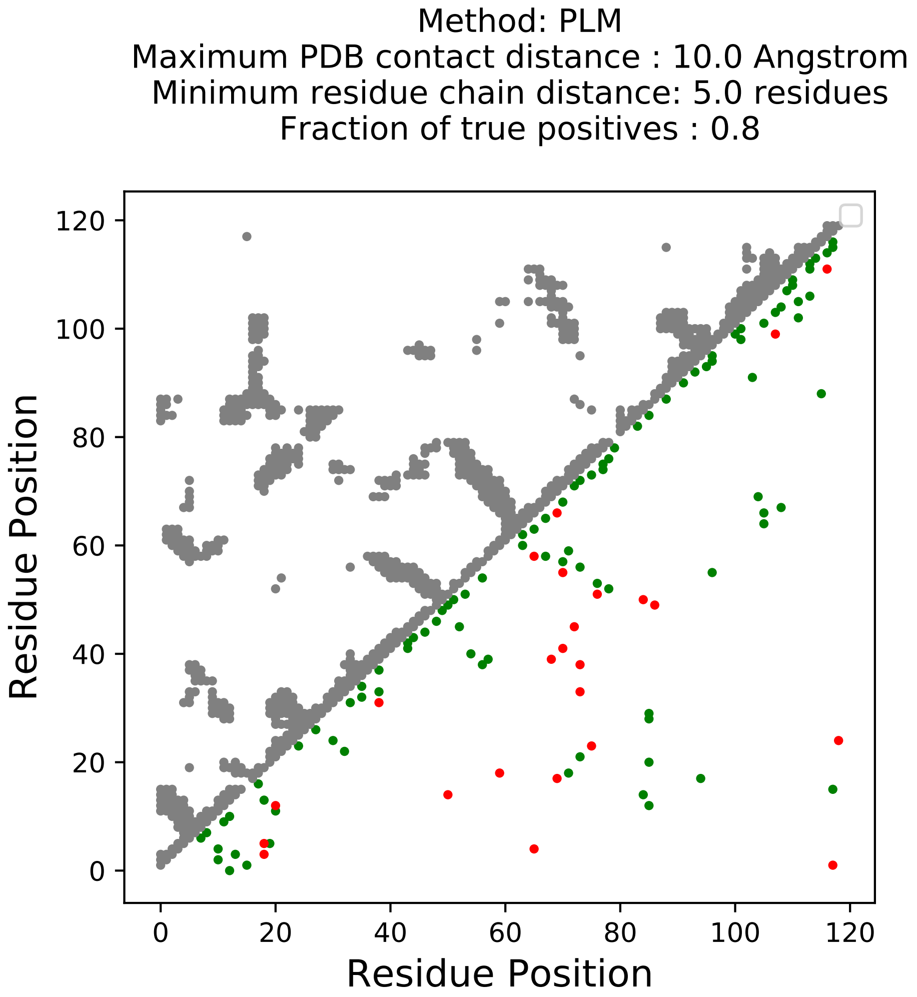

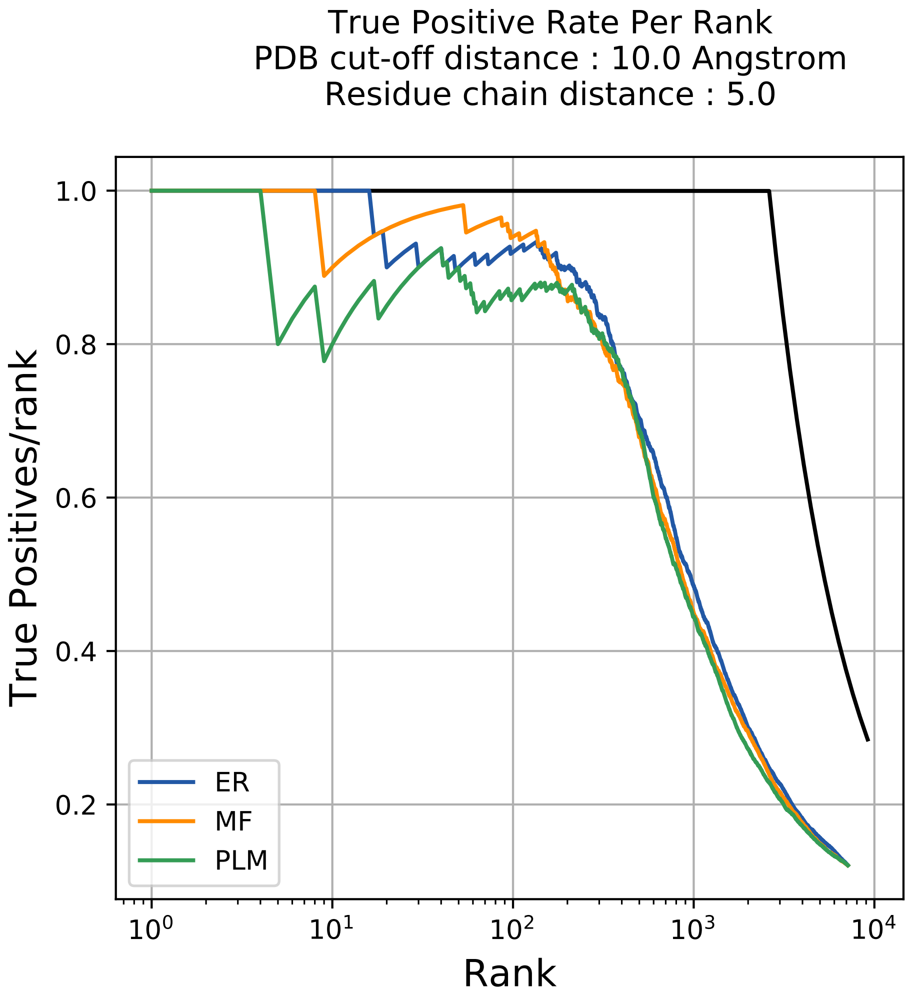

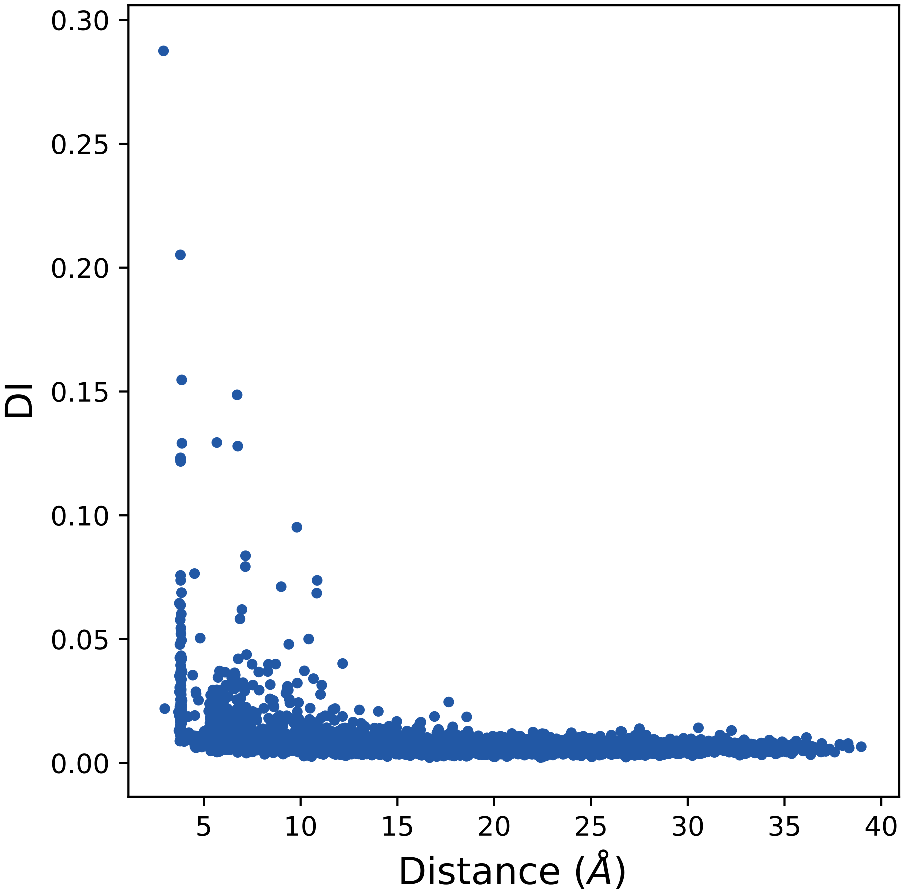

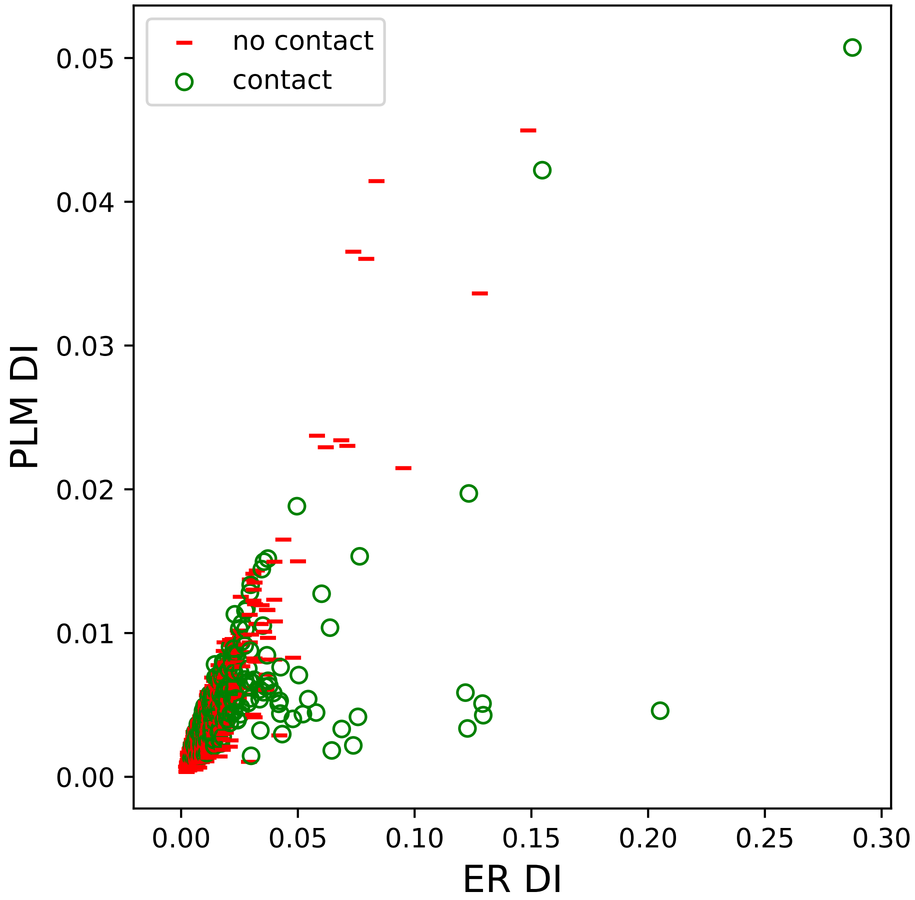

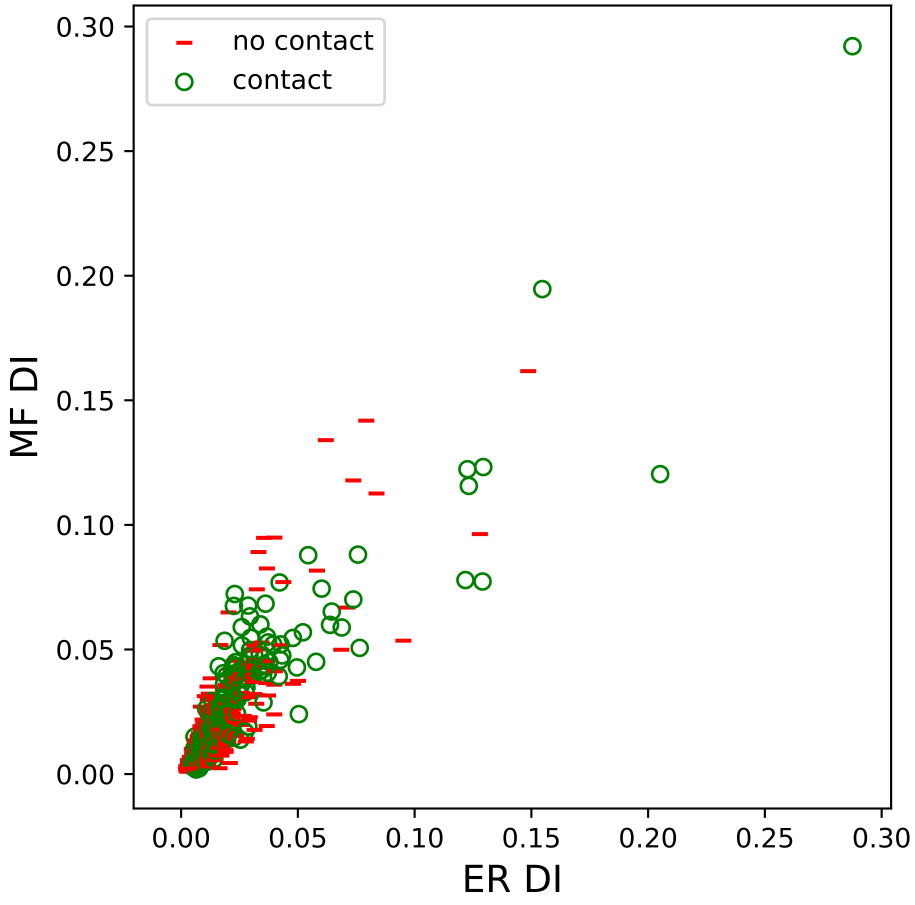

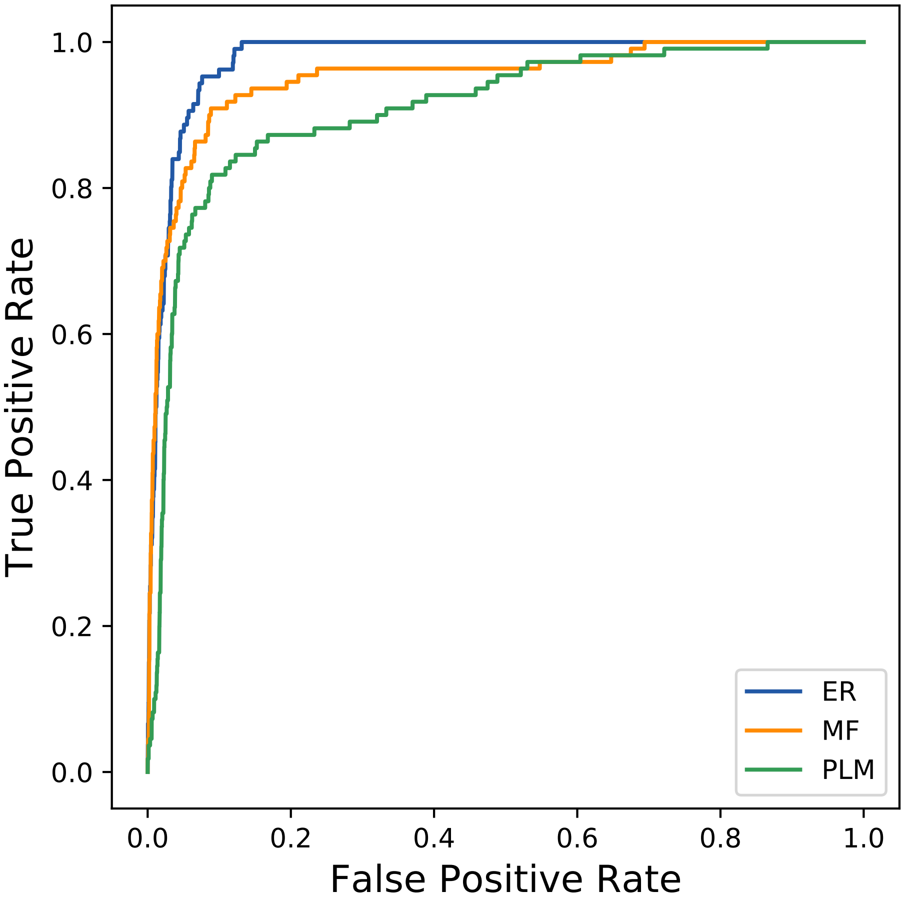

### 10286seq_120col_6pz4_PF00026_plots.pdf

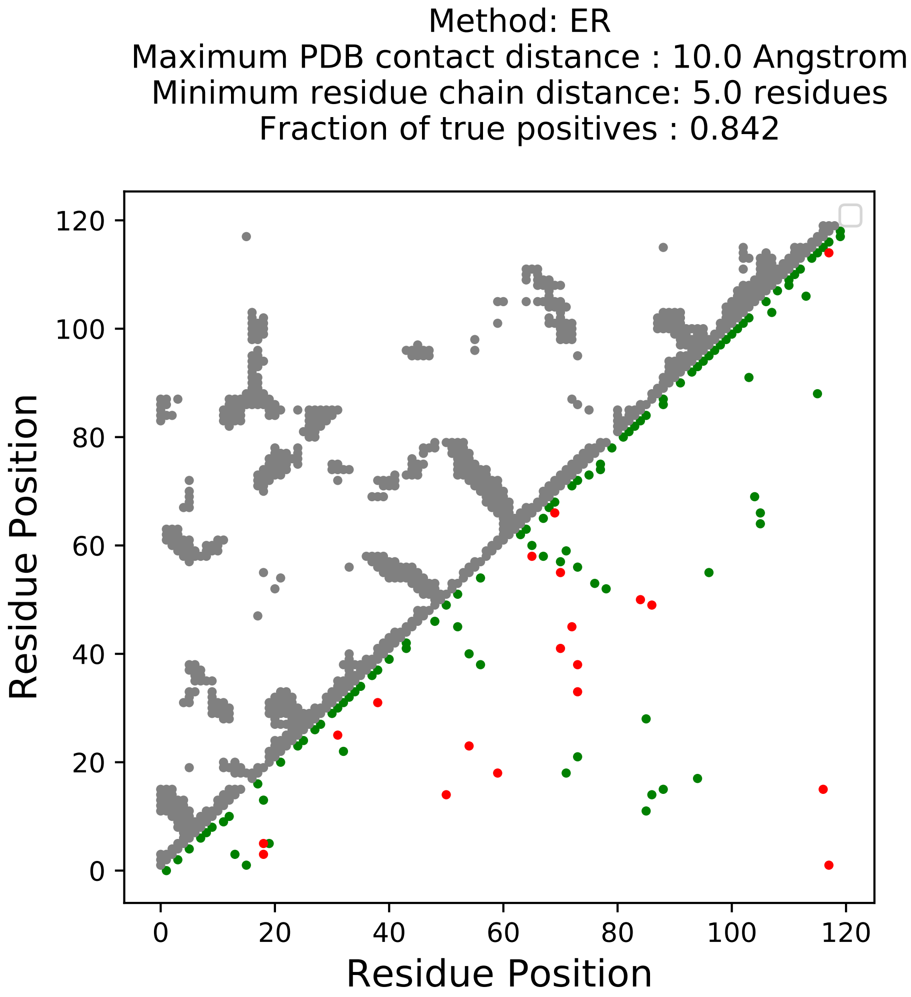

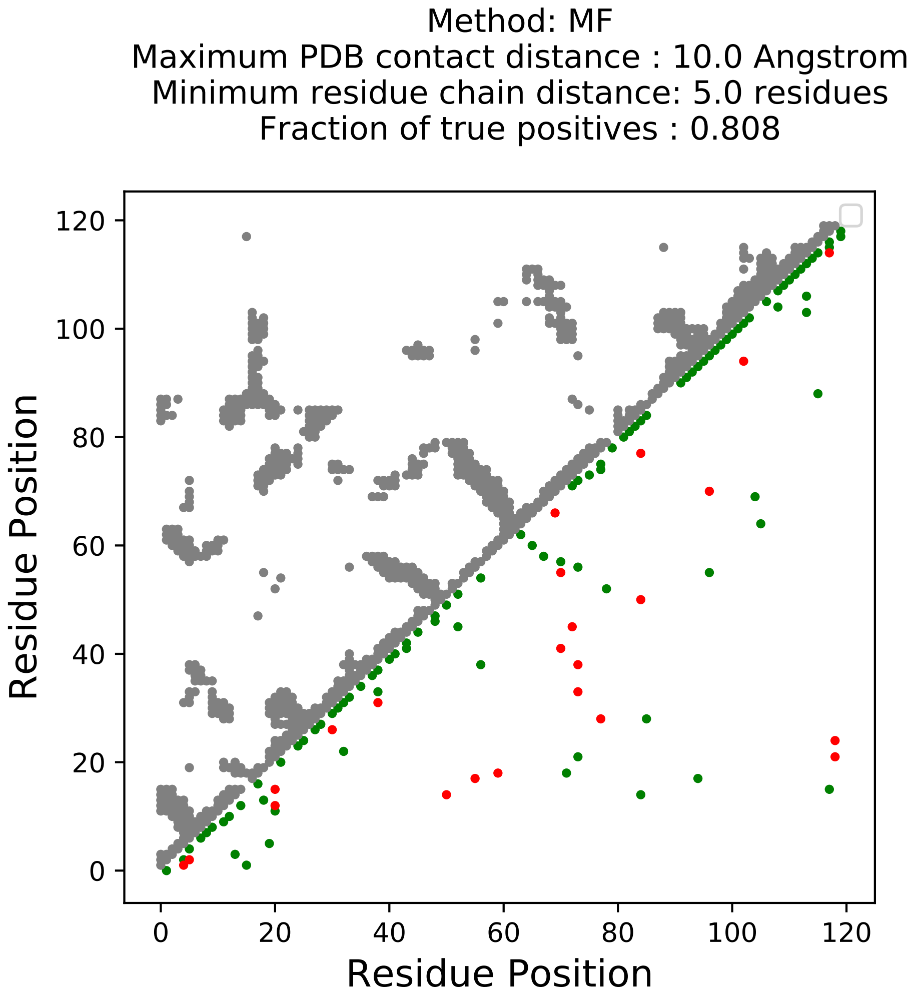

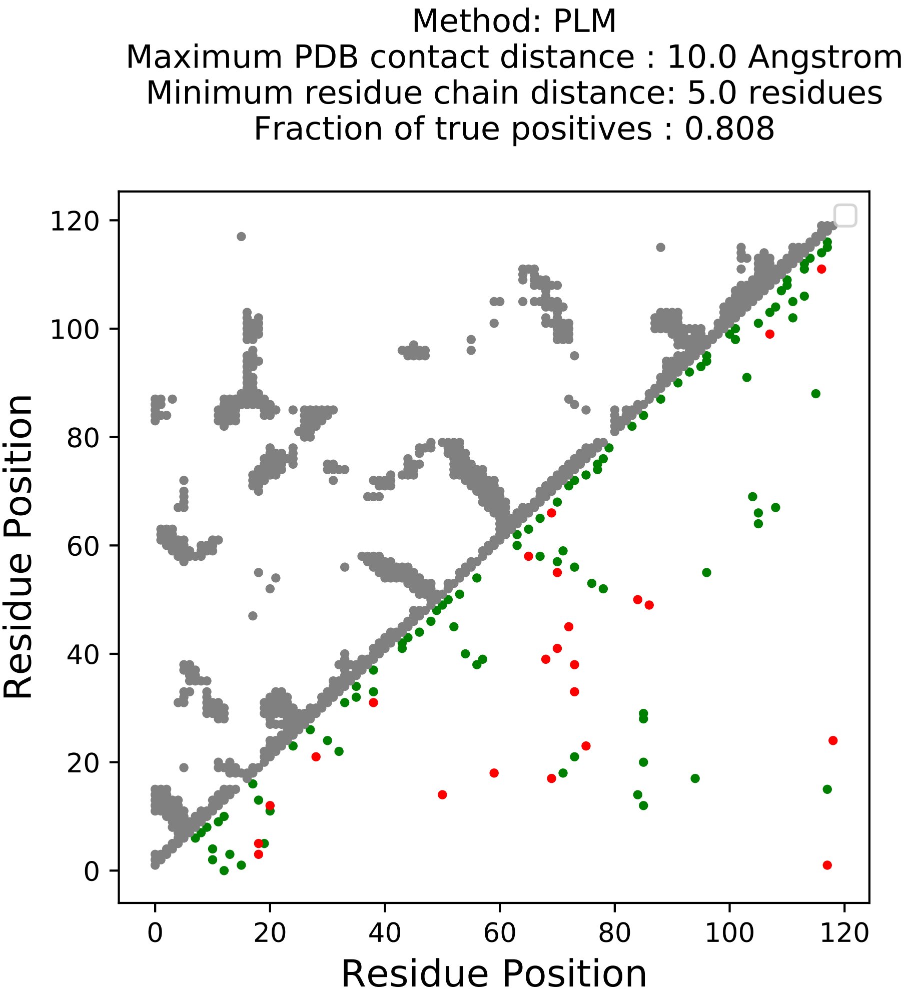

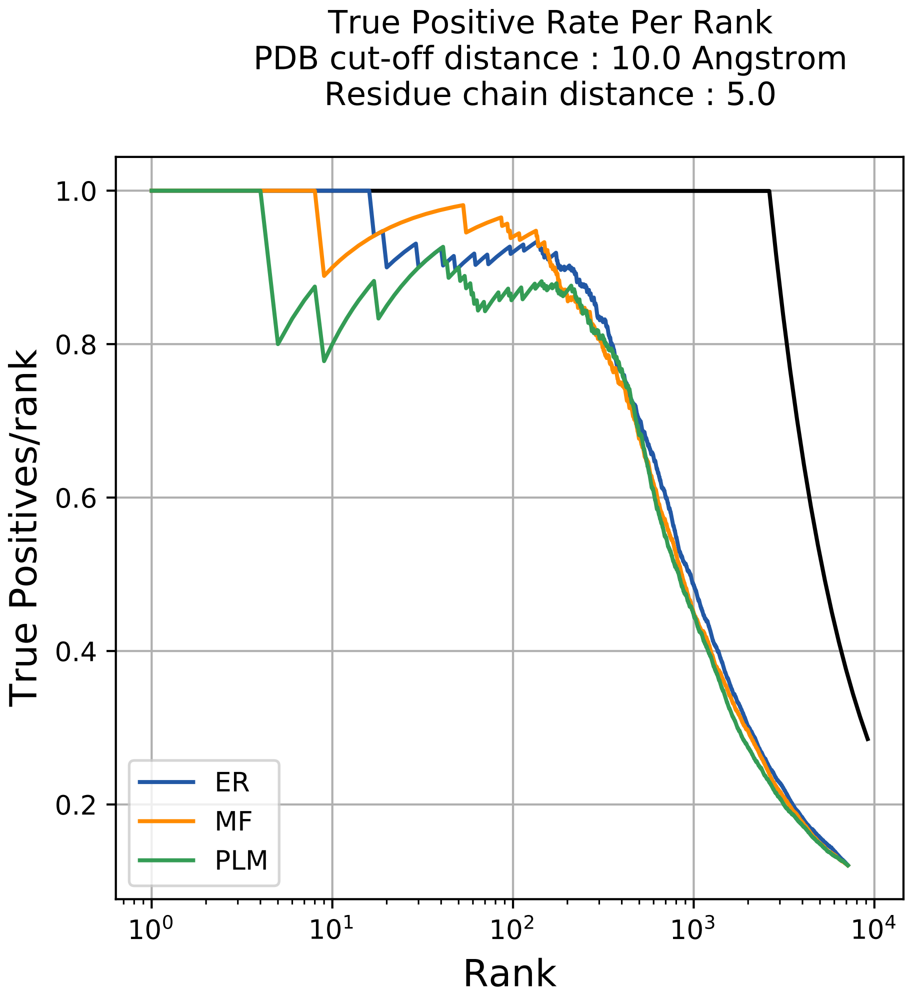

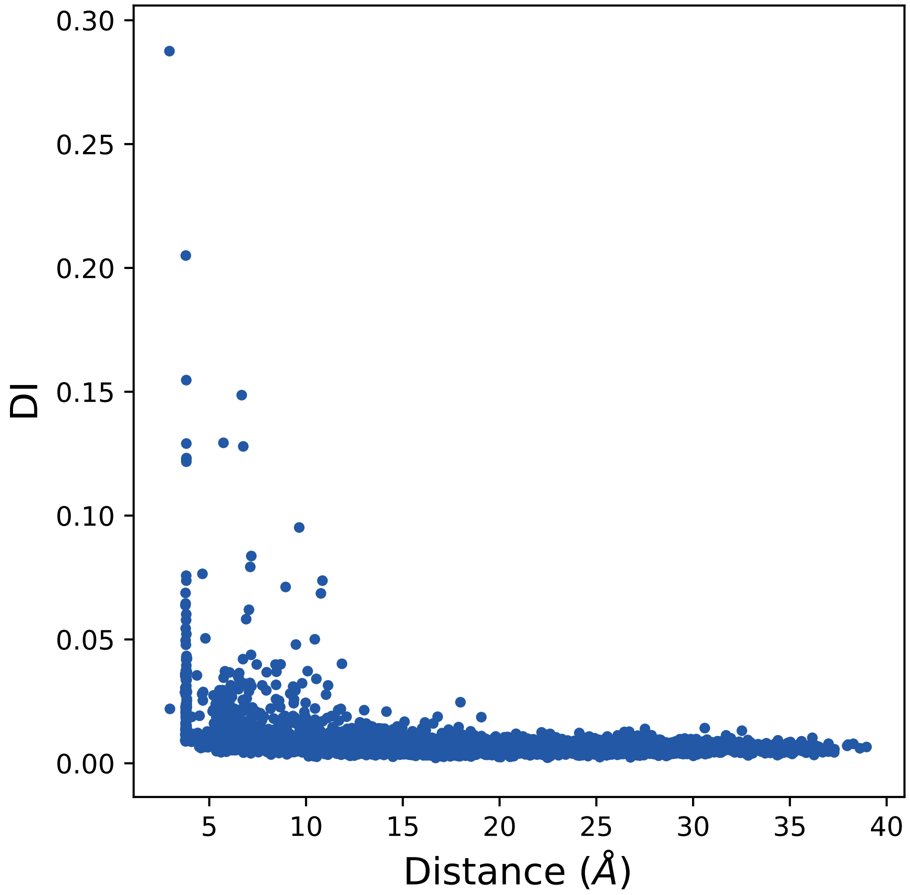

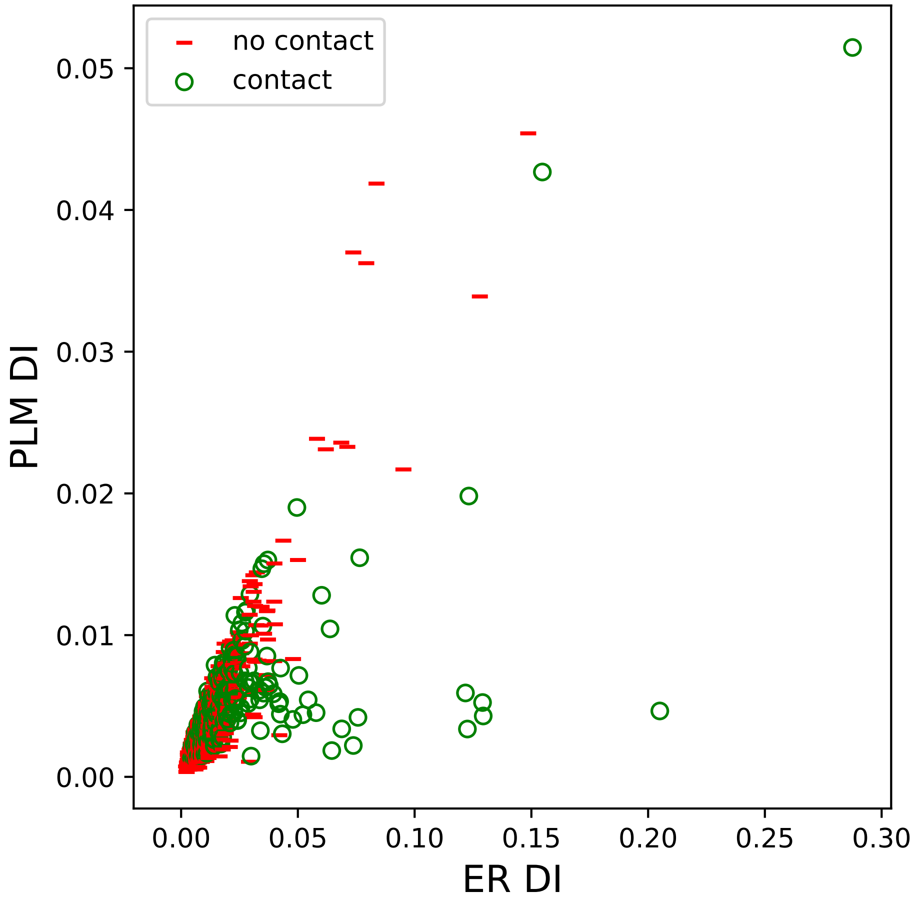

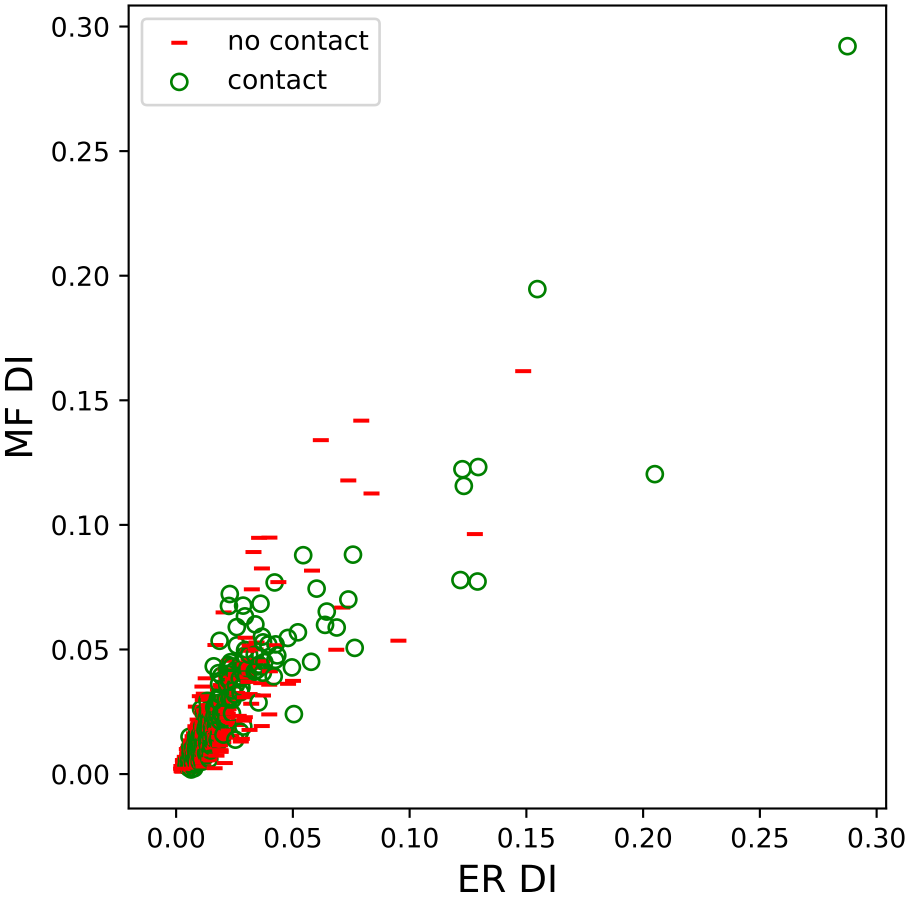

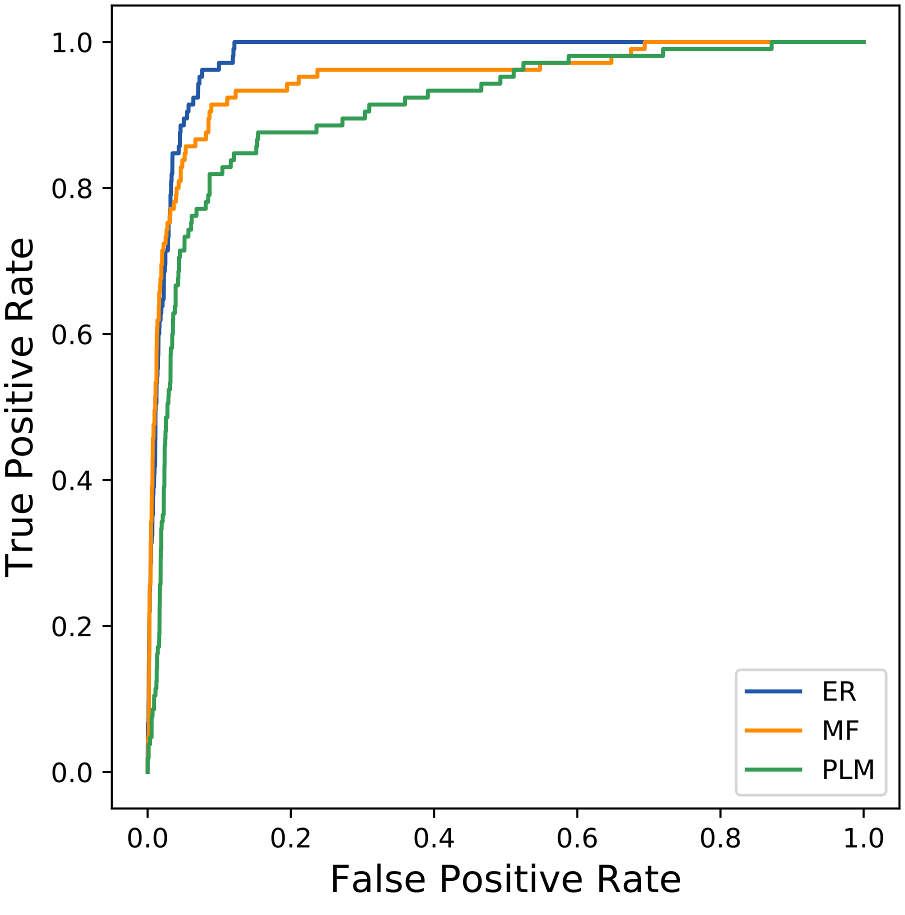

### 11510seq_126col_4qt0_PF00056_plots.pdf

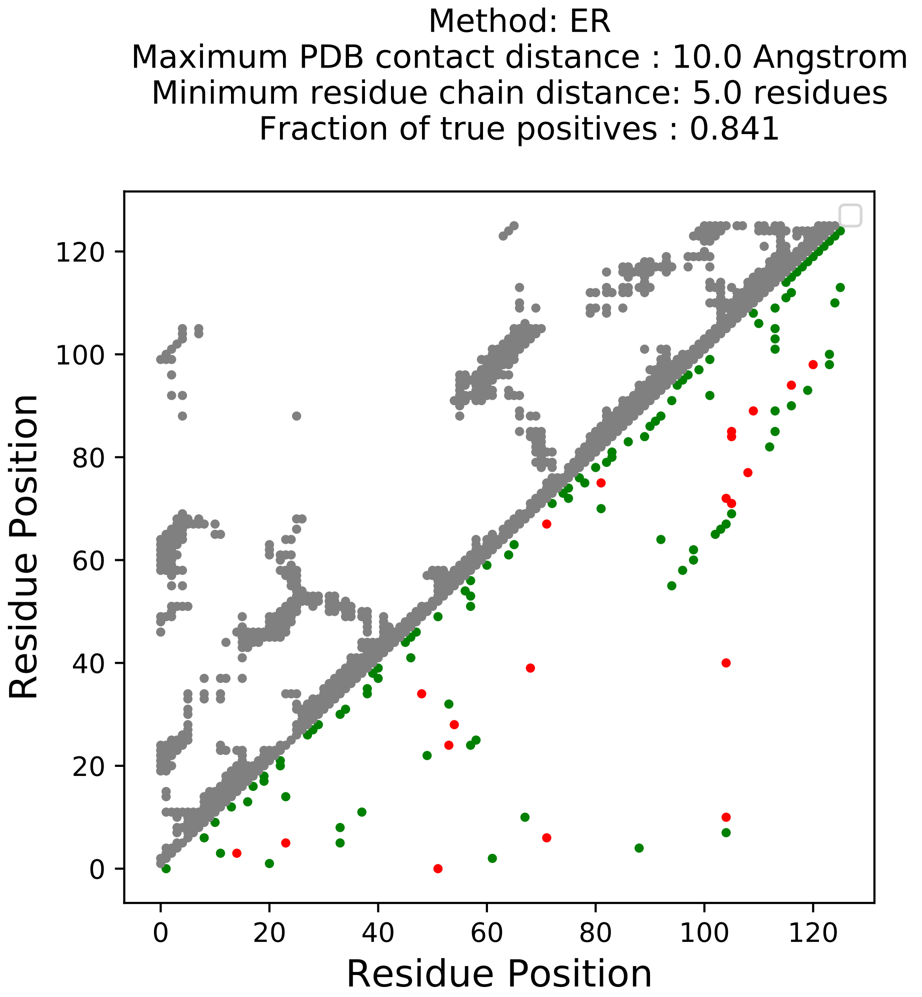

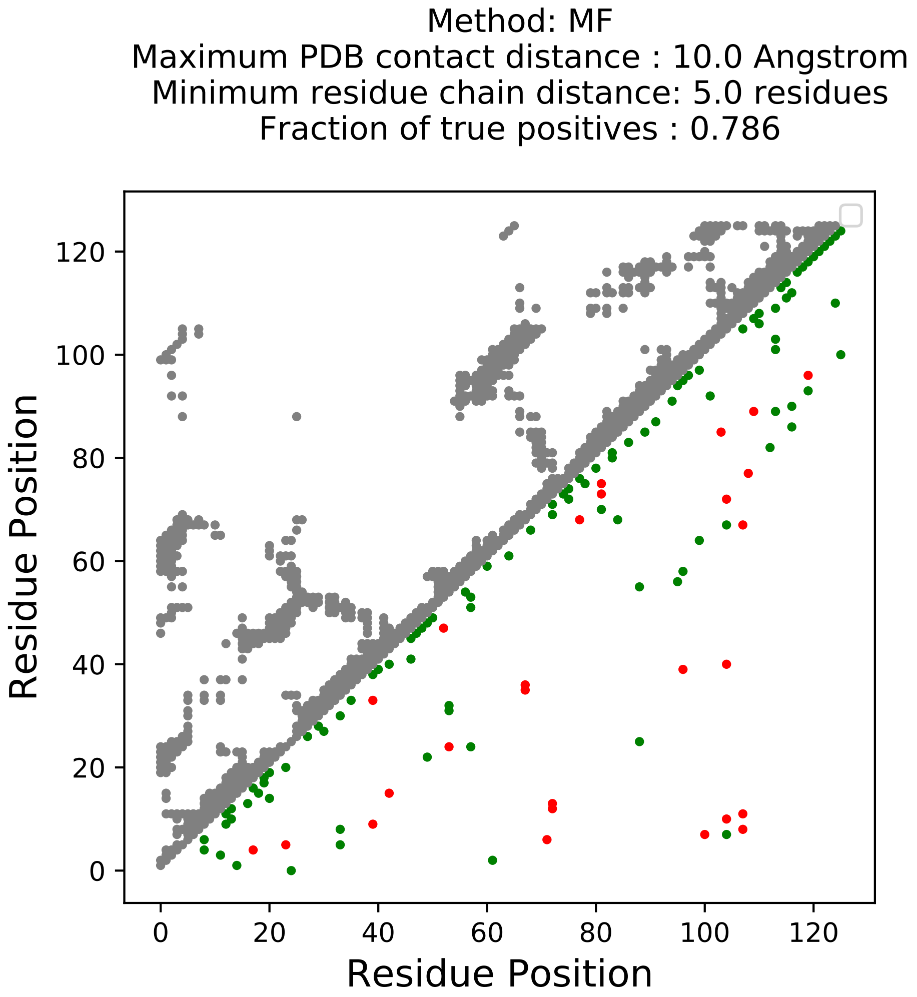

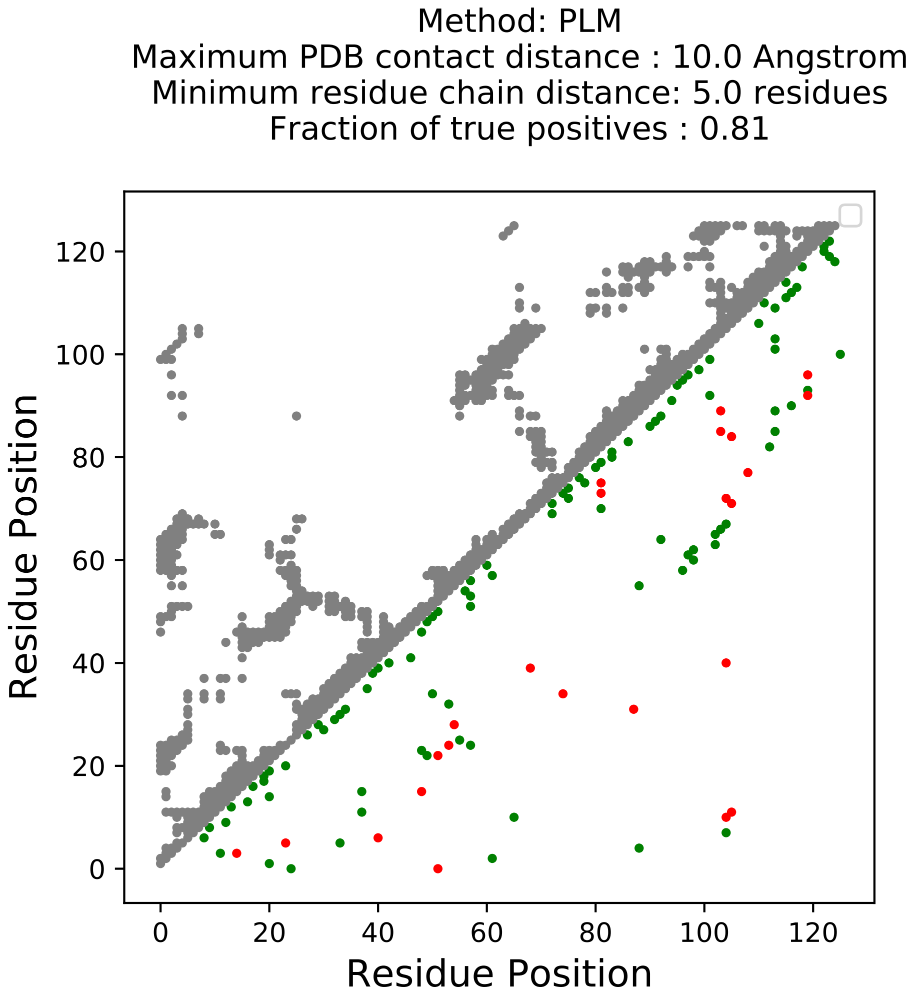

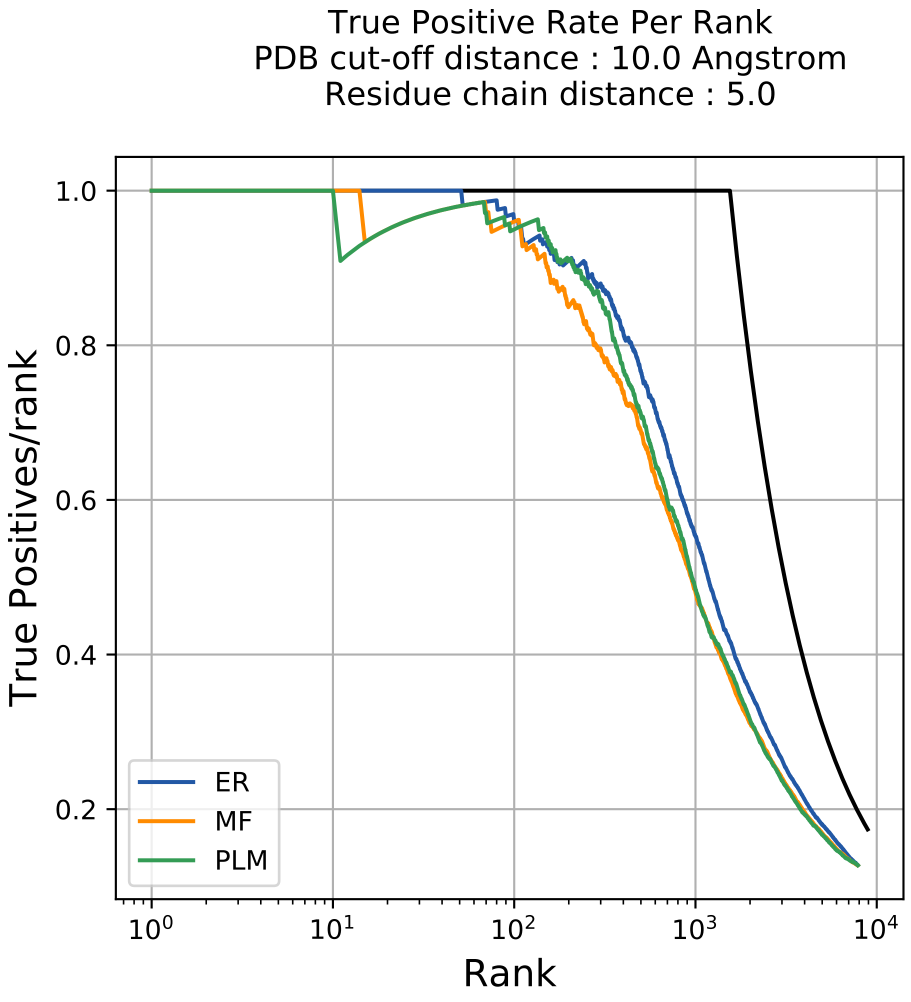

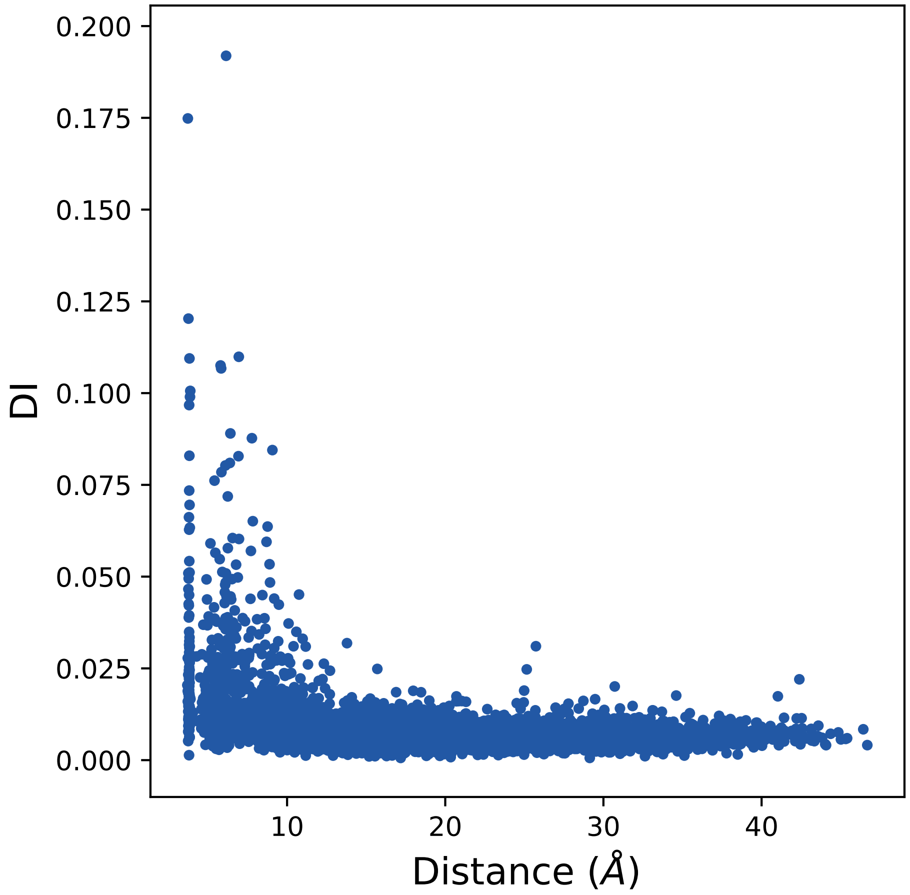

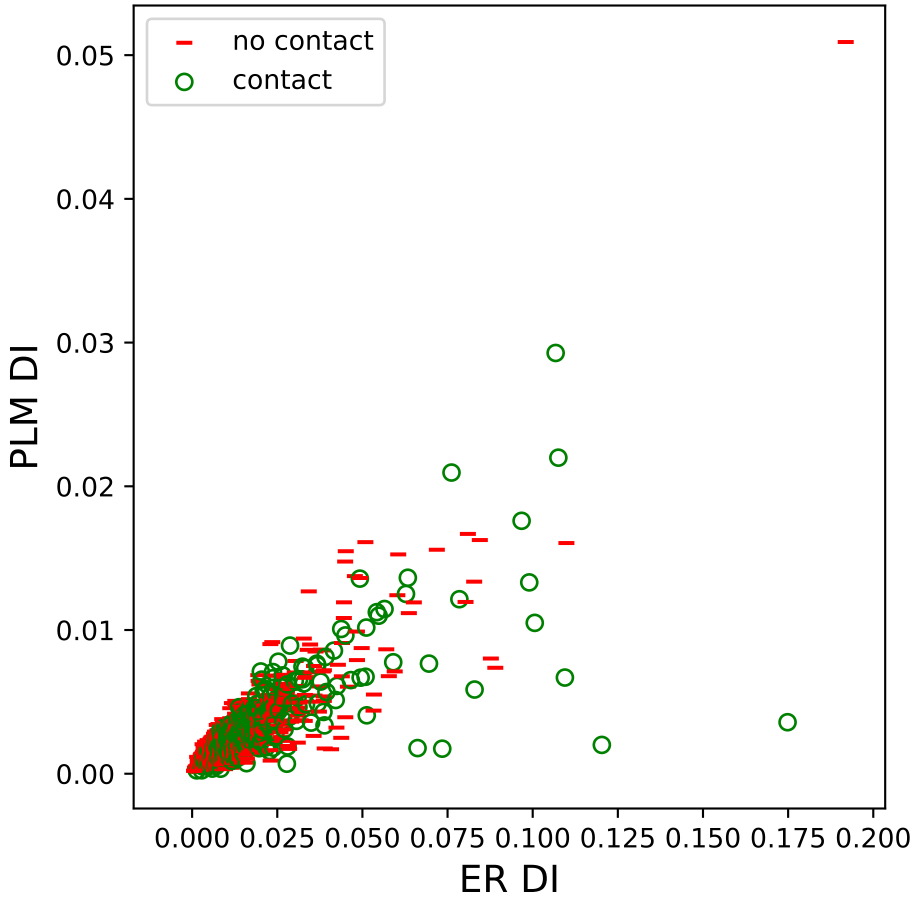

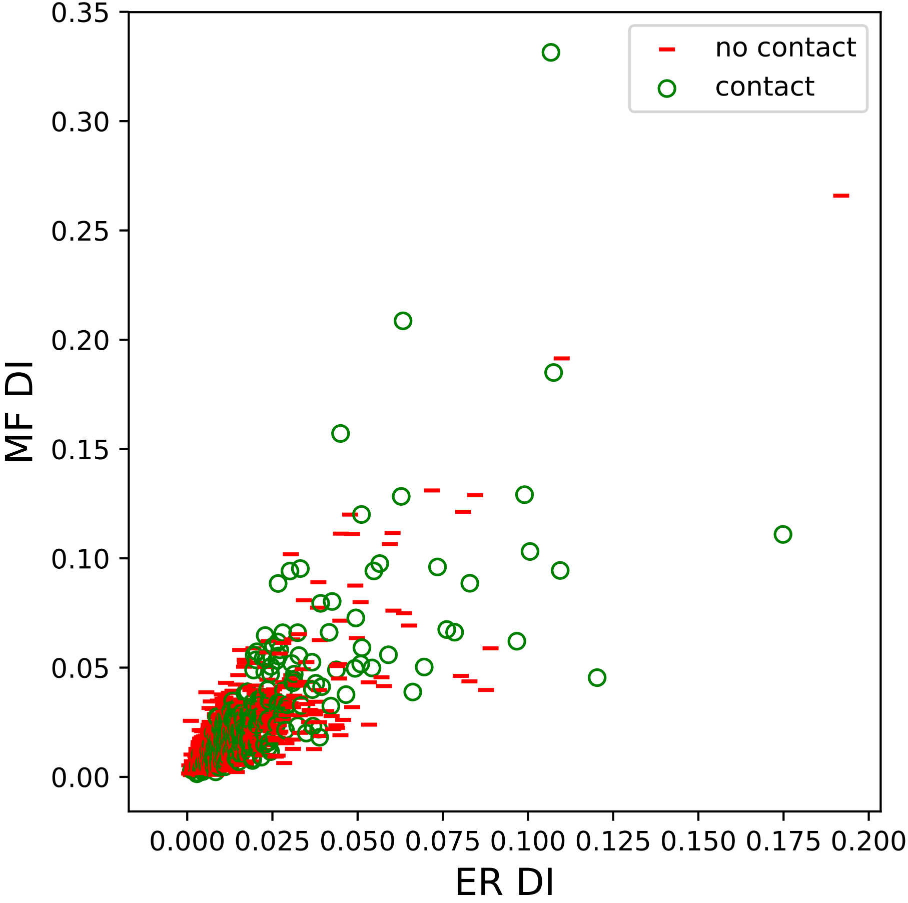

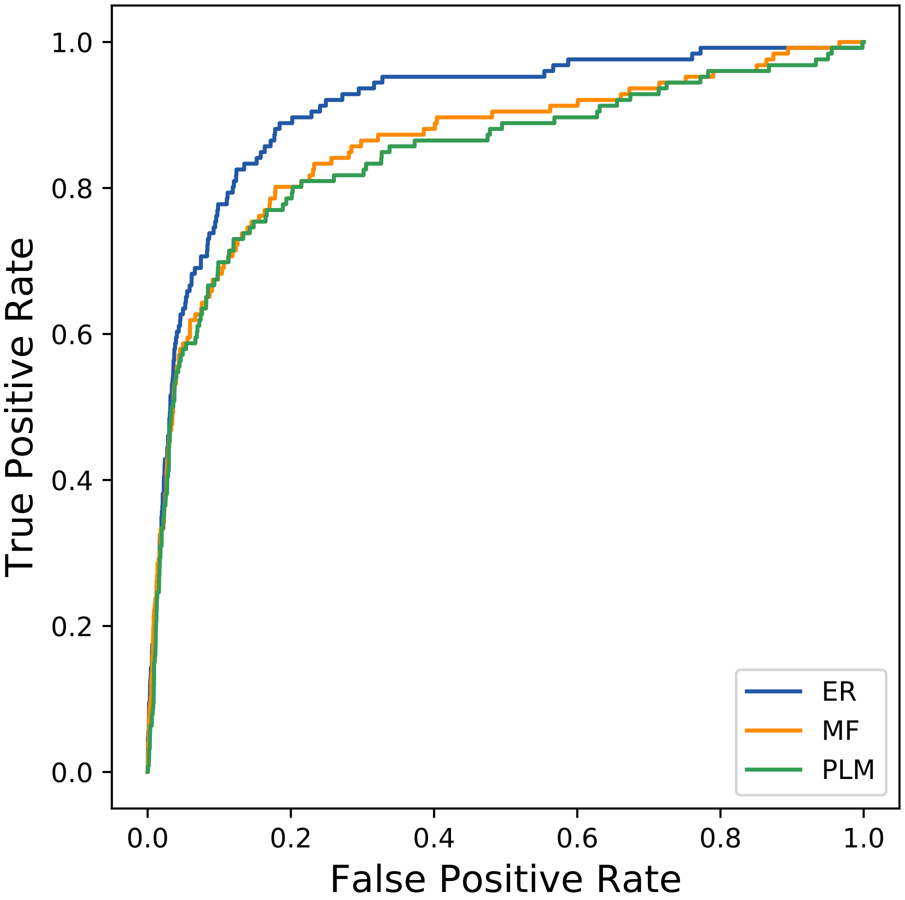

### 12172seq_232col_3ahz_PF00232_plots.pdf

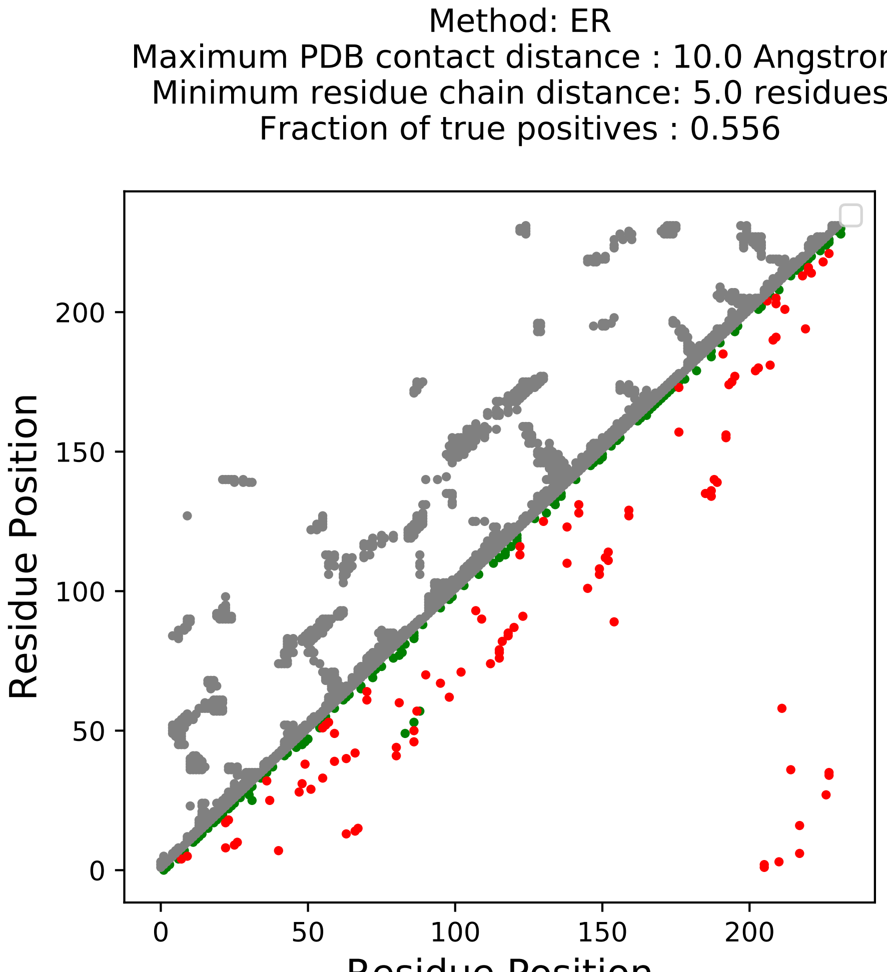

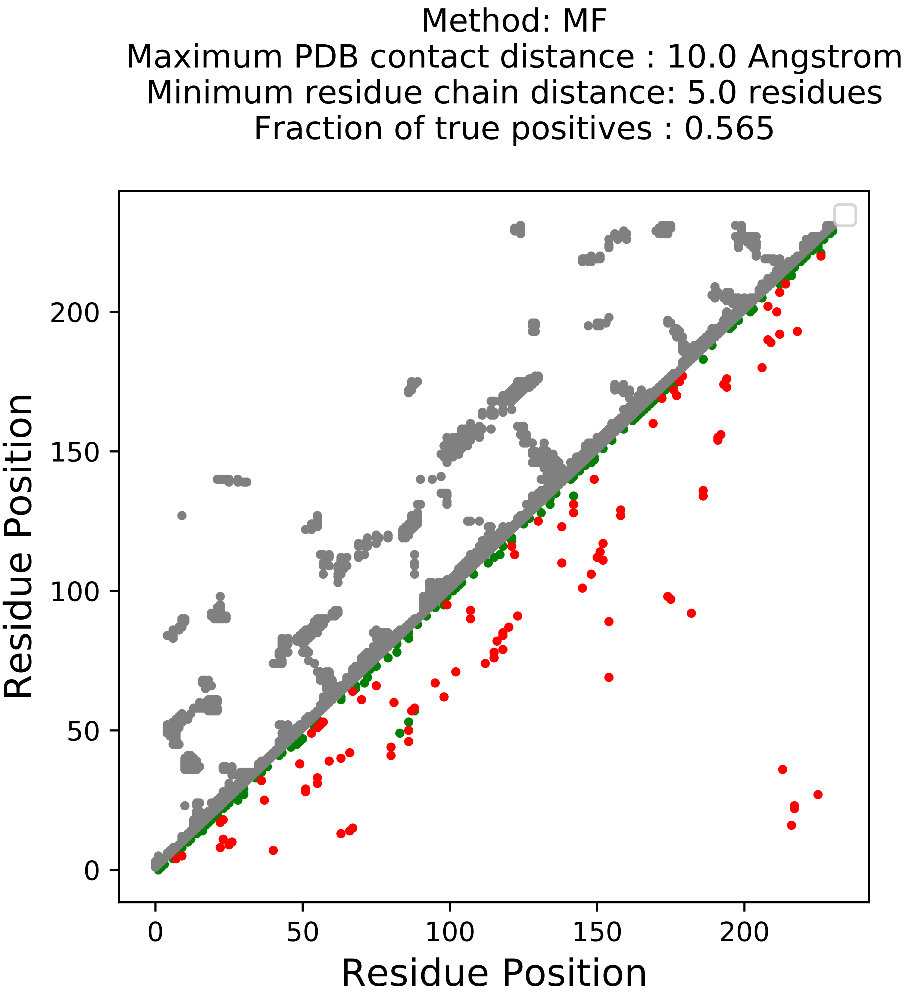

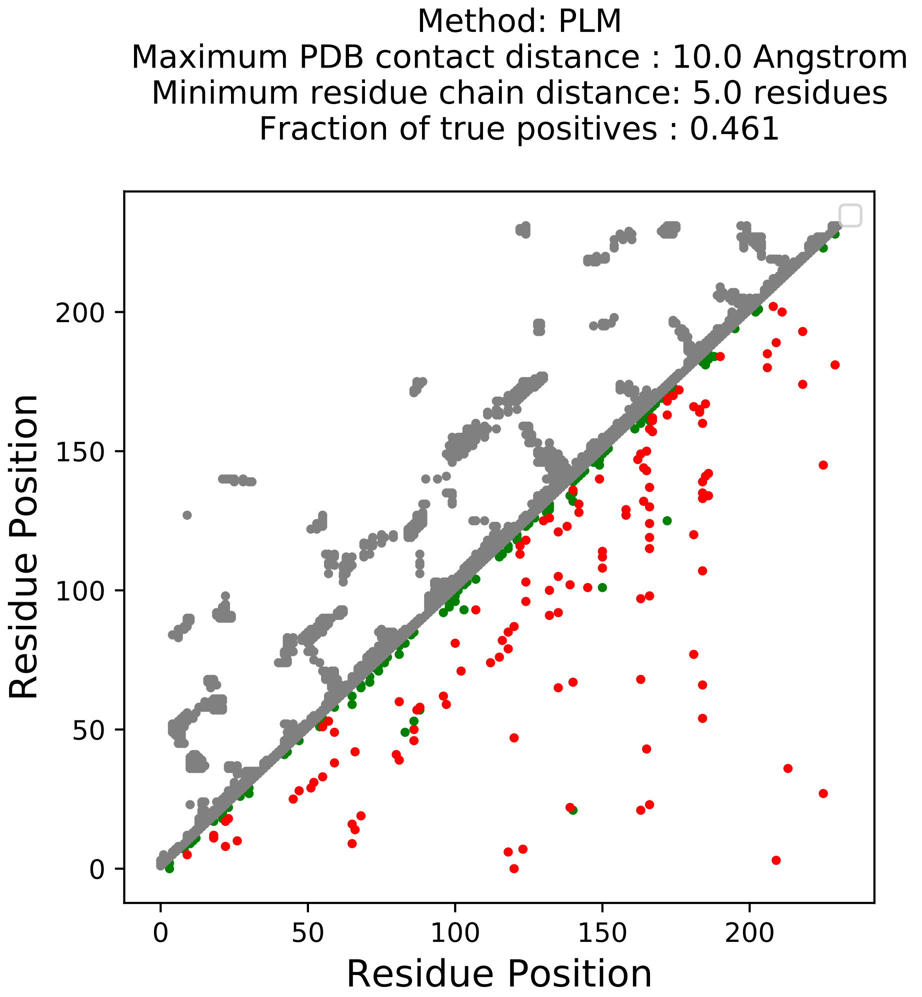

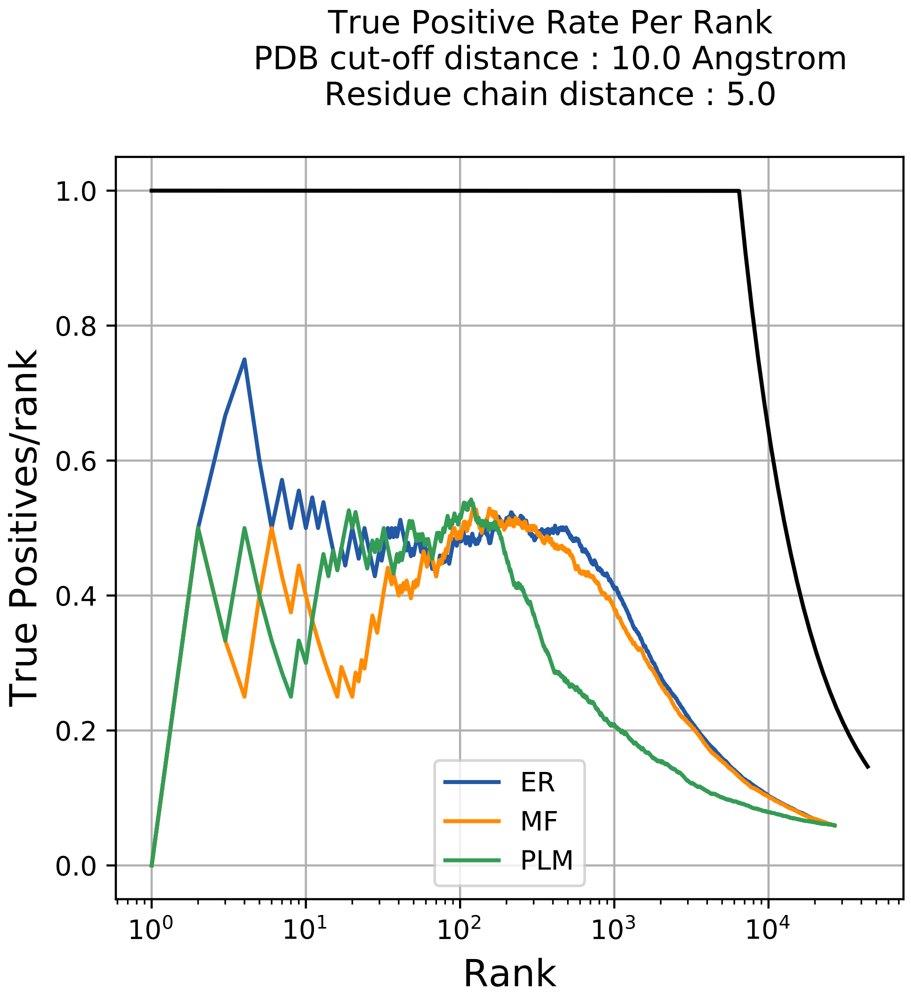

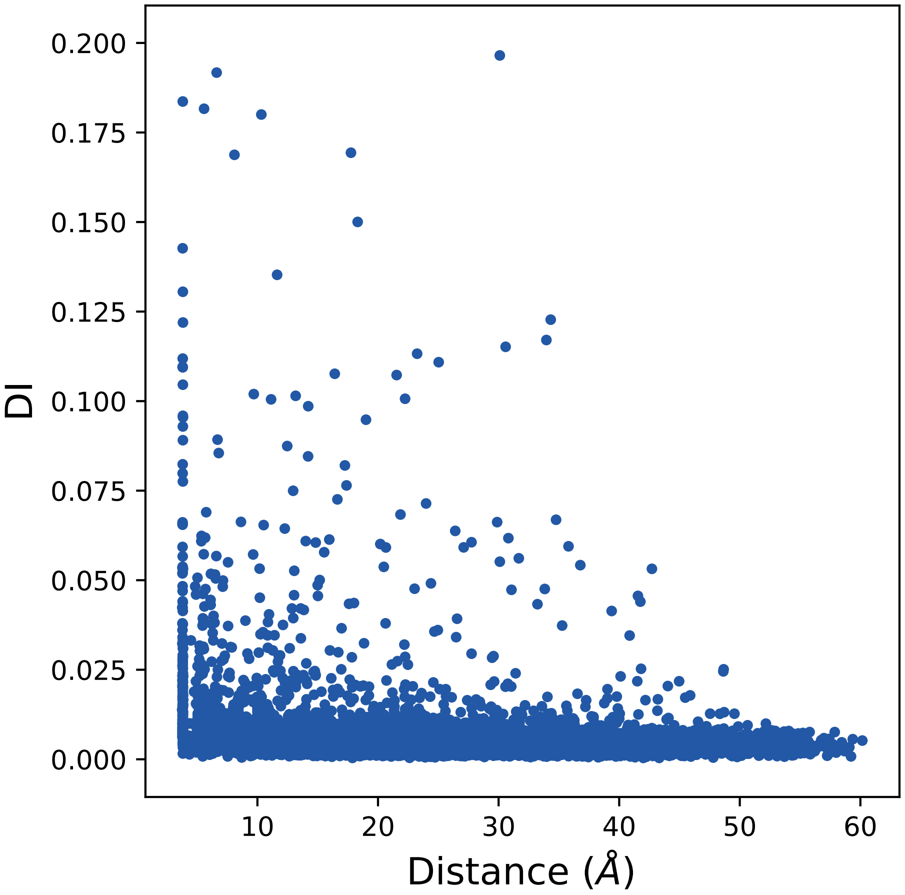

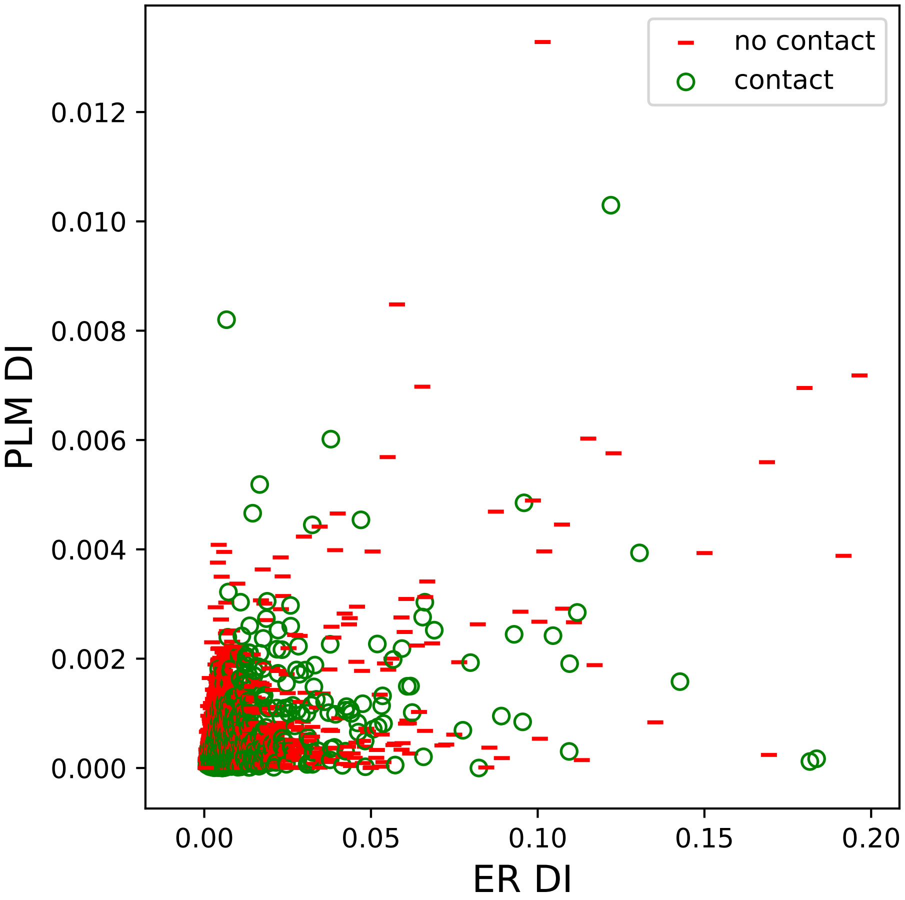

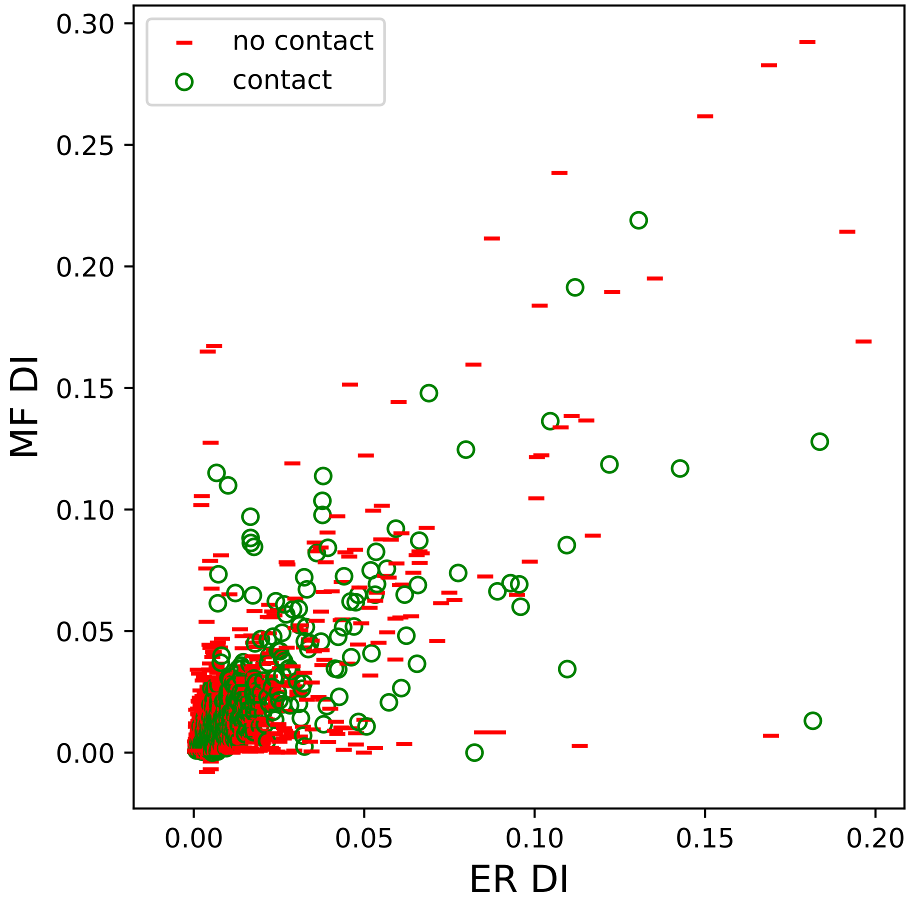

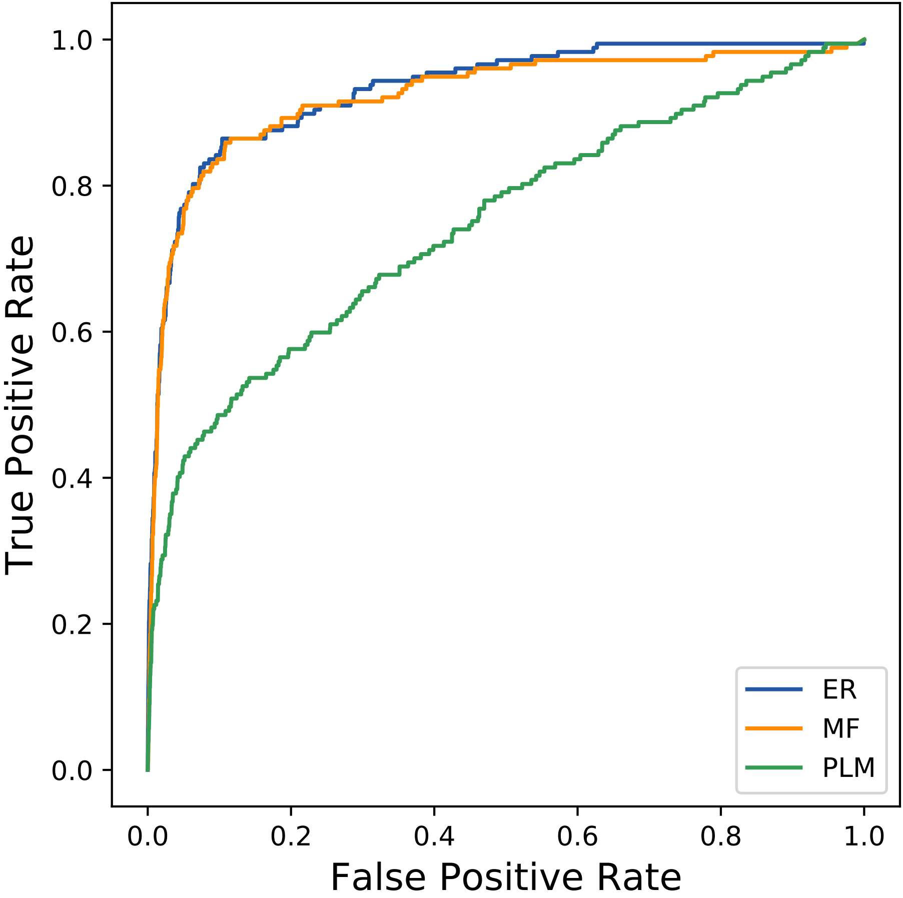

### 12823seq_22col_4zfh_PF13857_plots.pdf

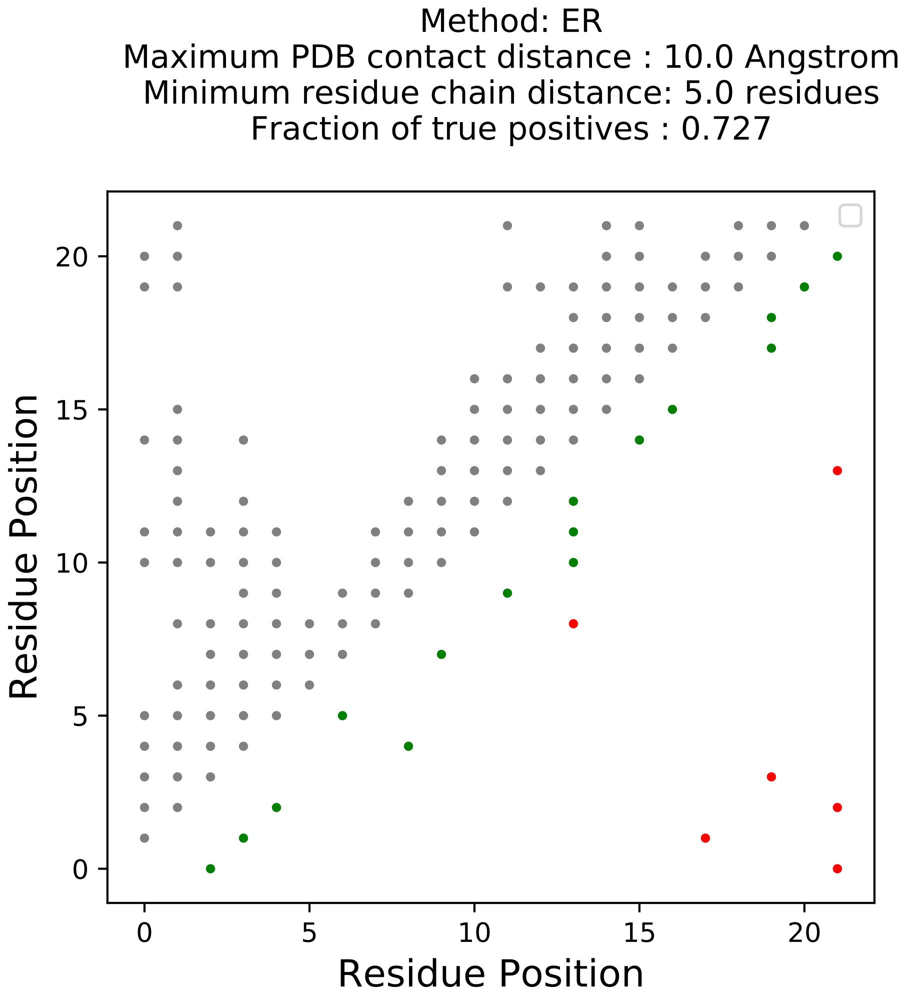

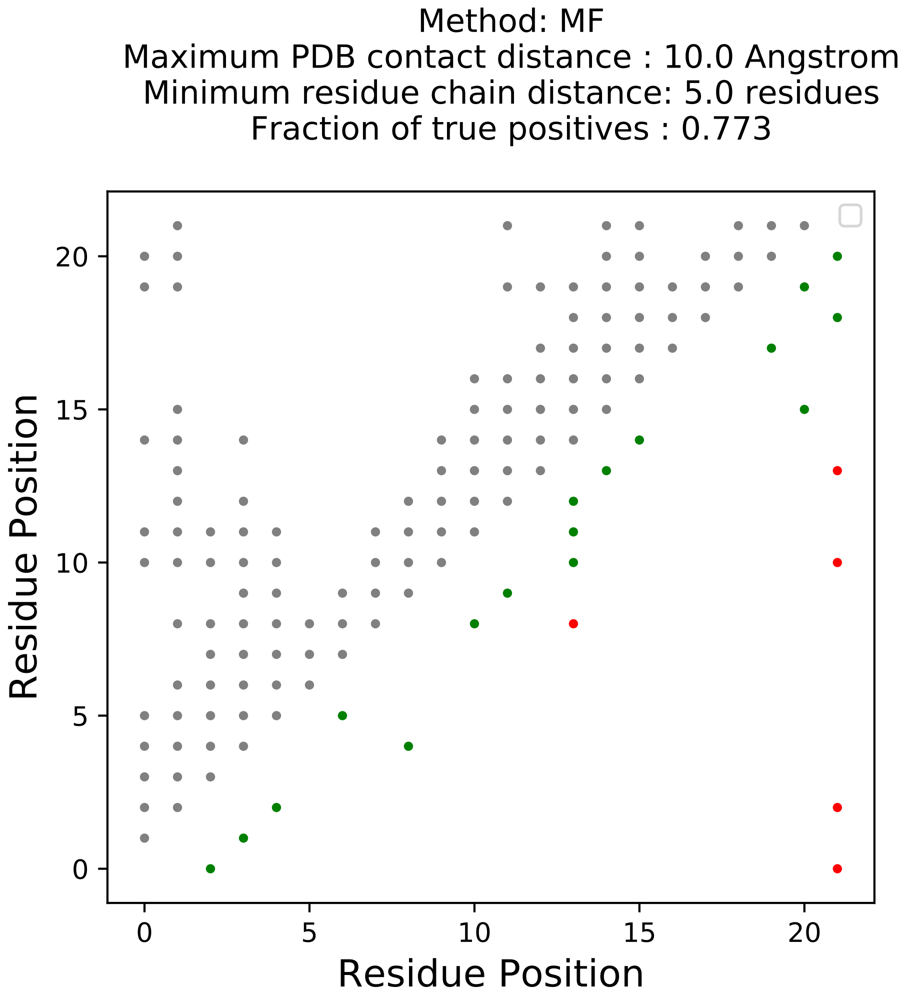

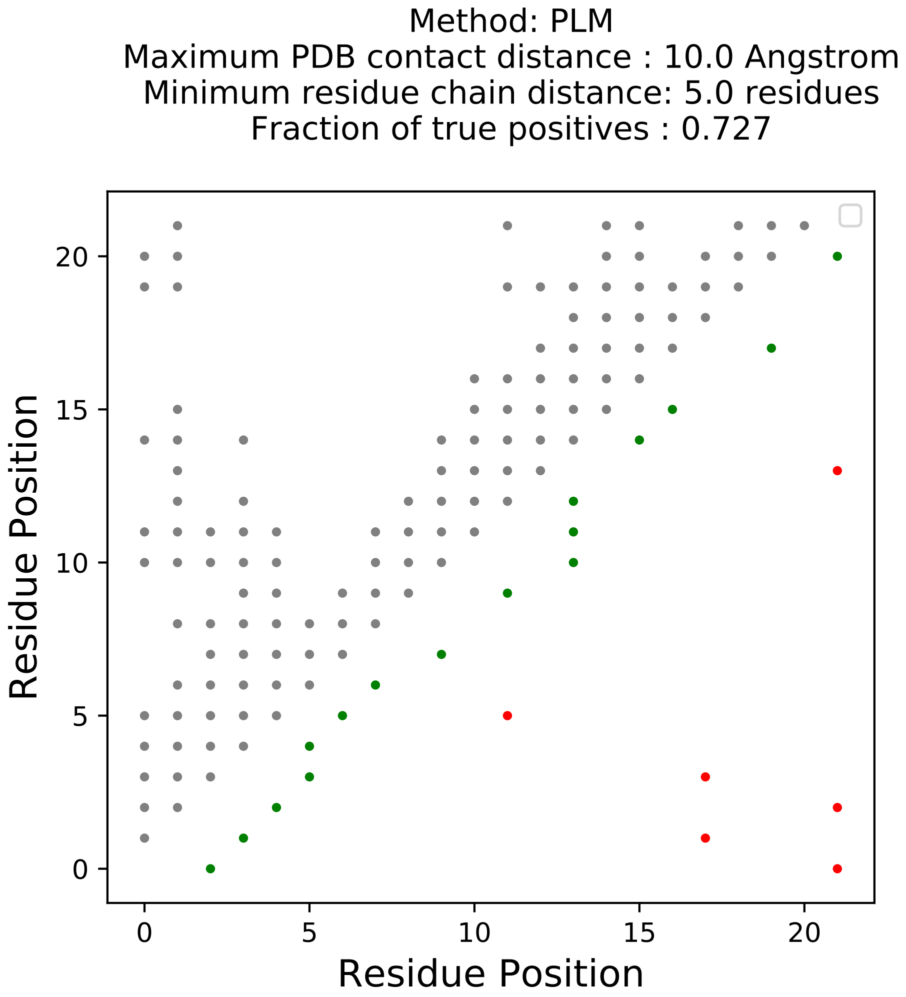

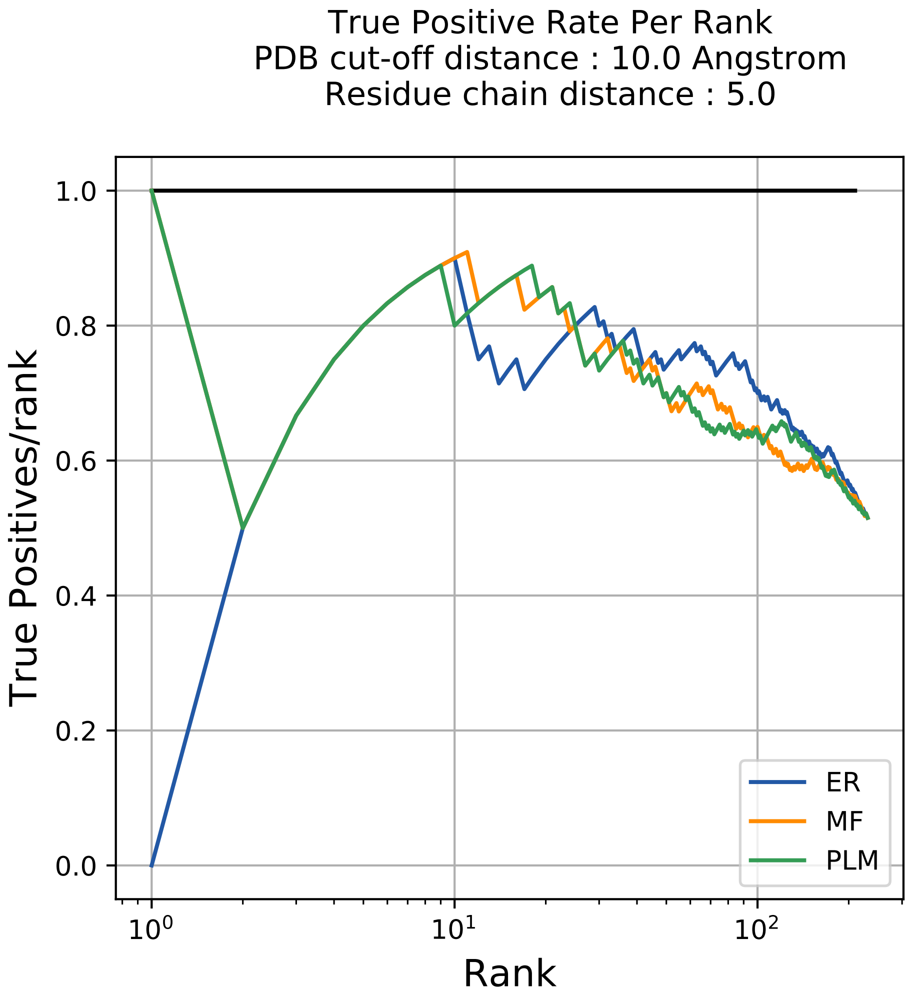

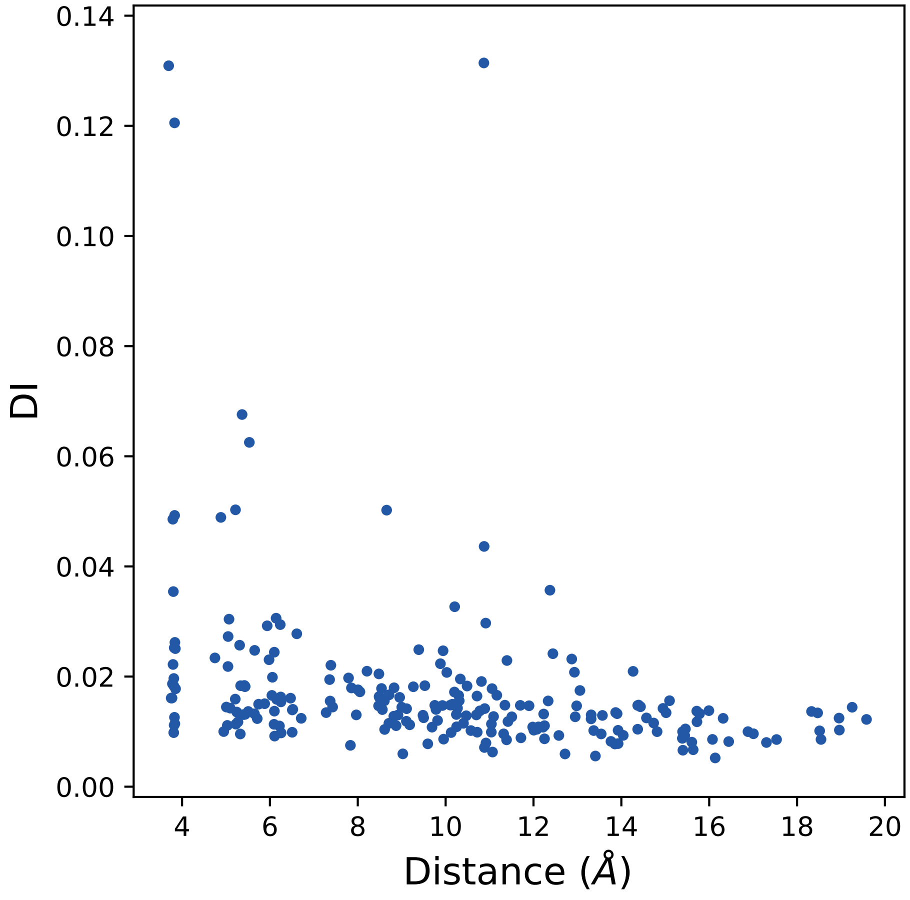

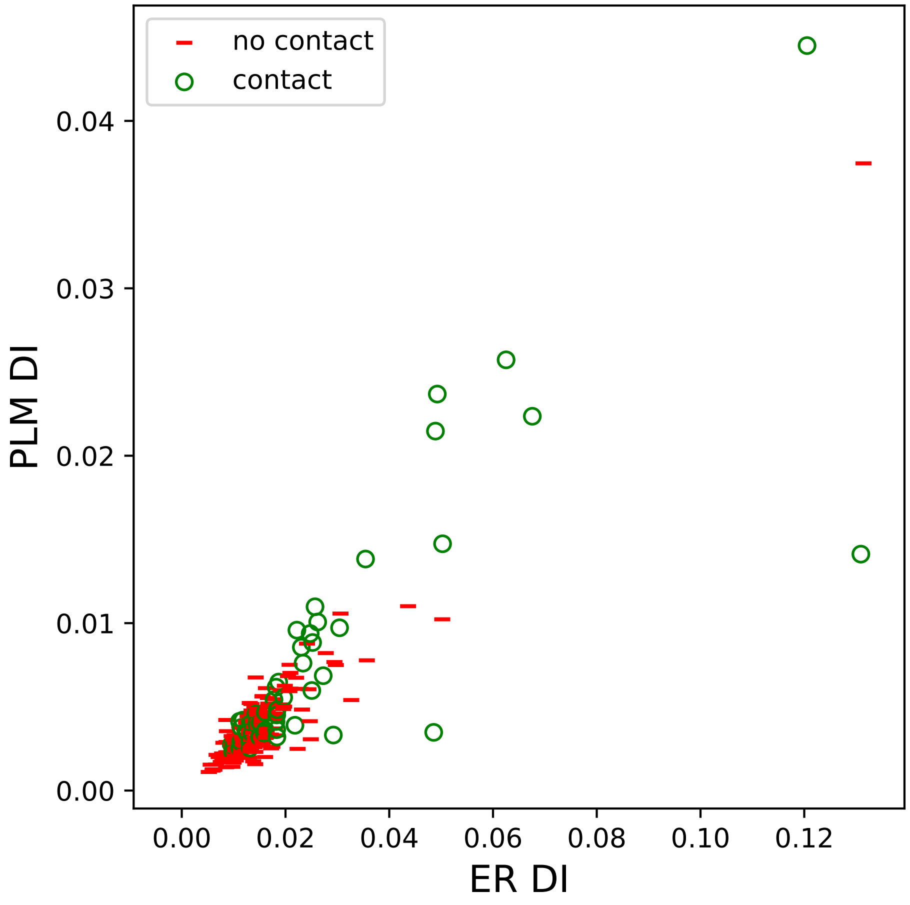

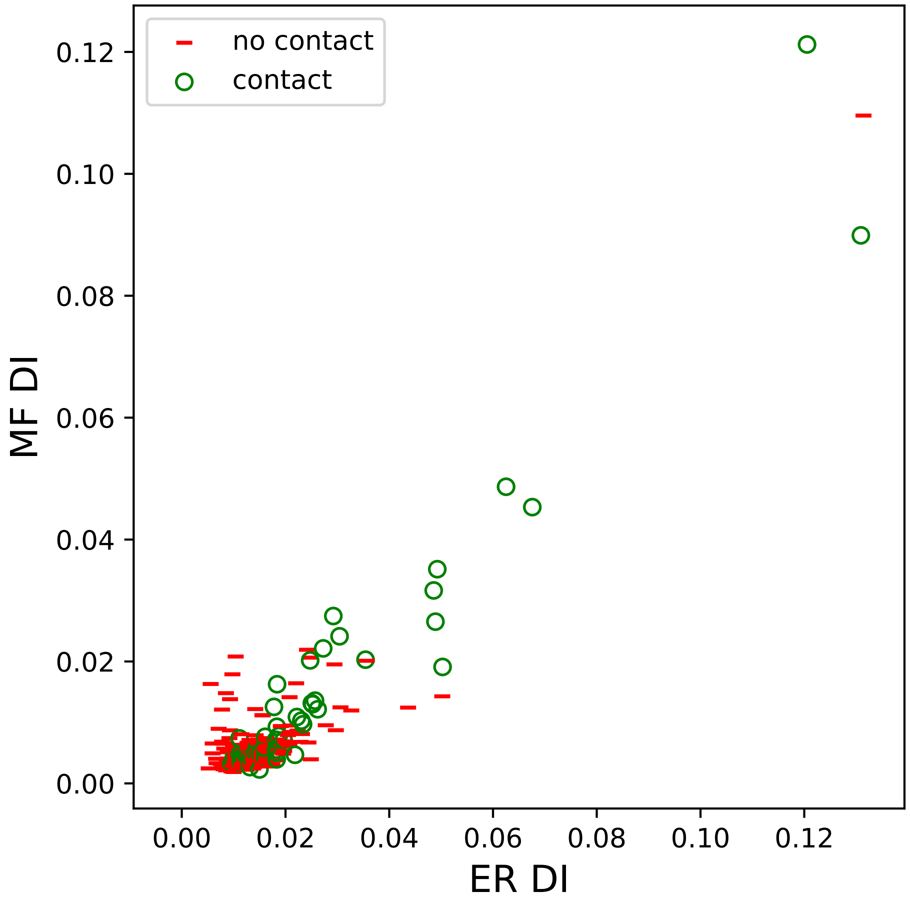

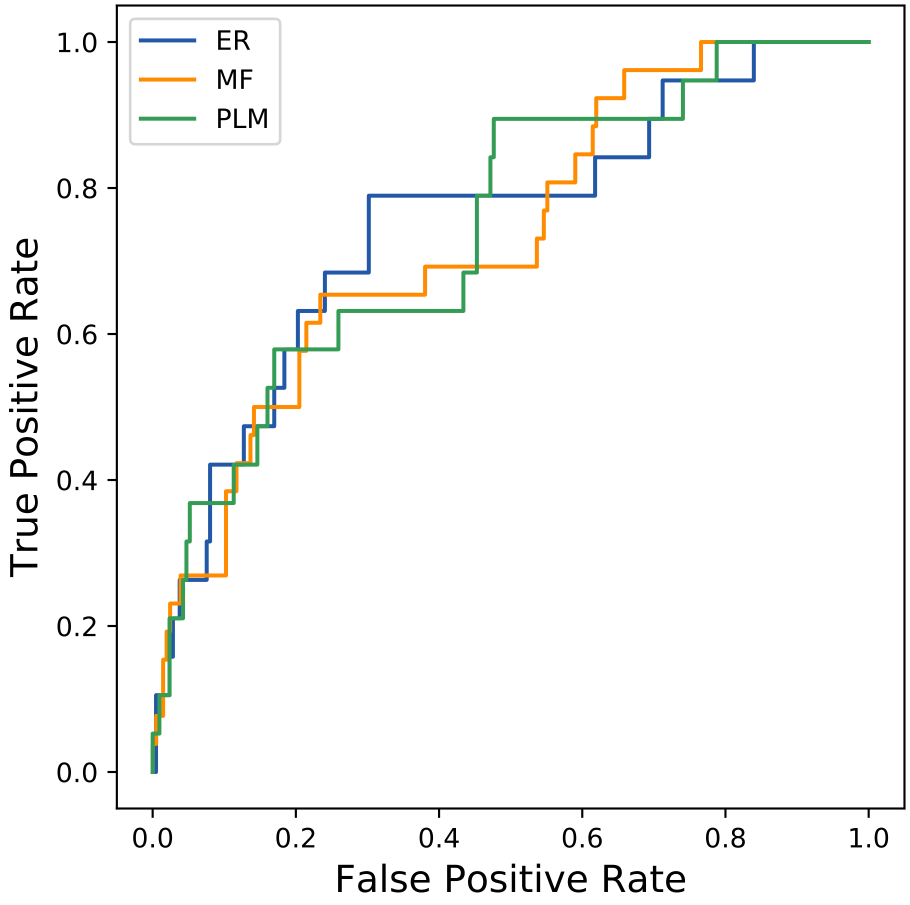

### 12971seq_32col_5pzh_PF00439_plots.pdf

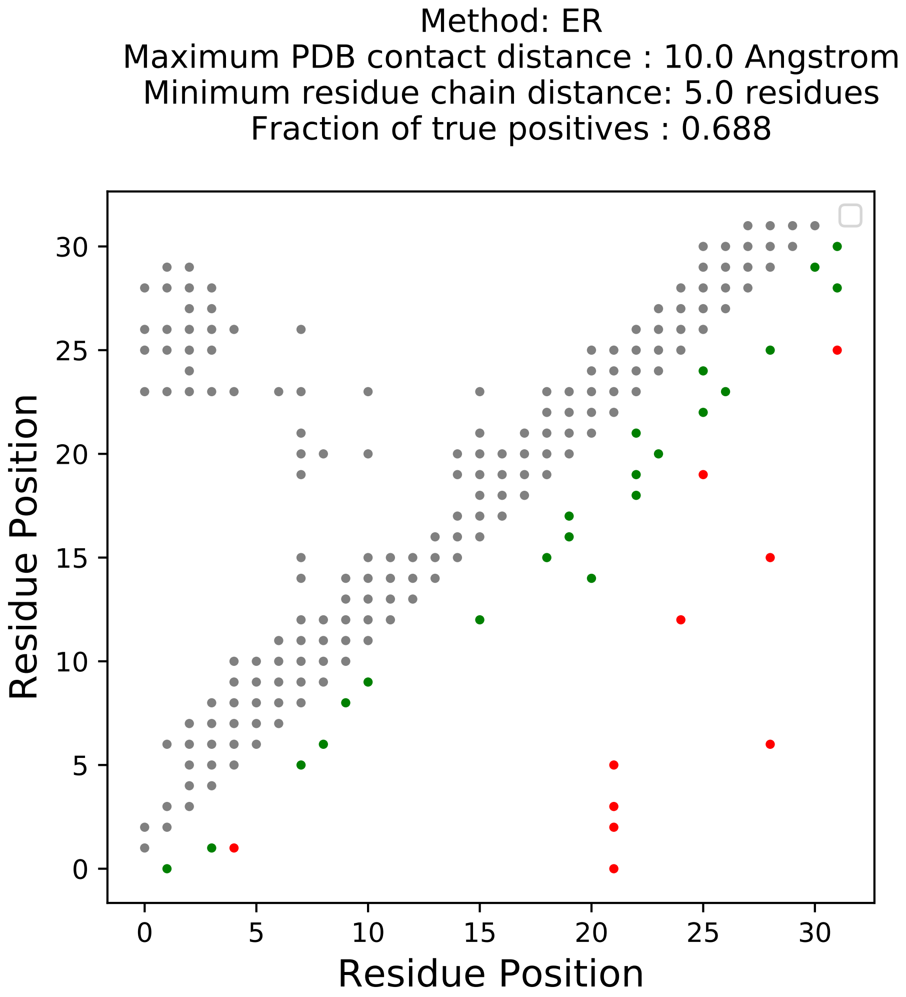

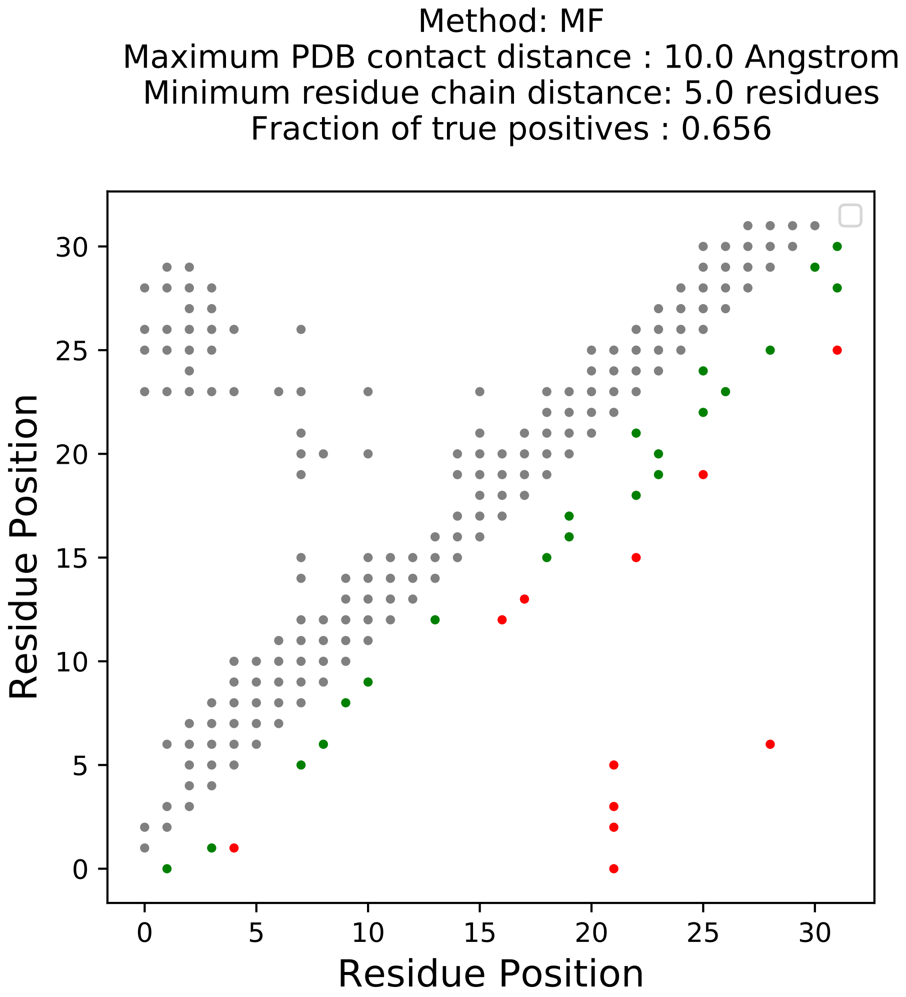

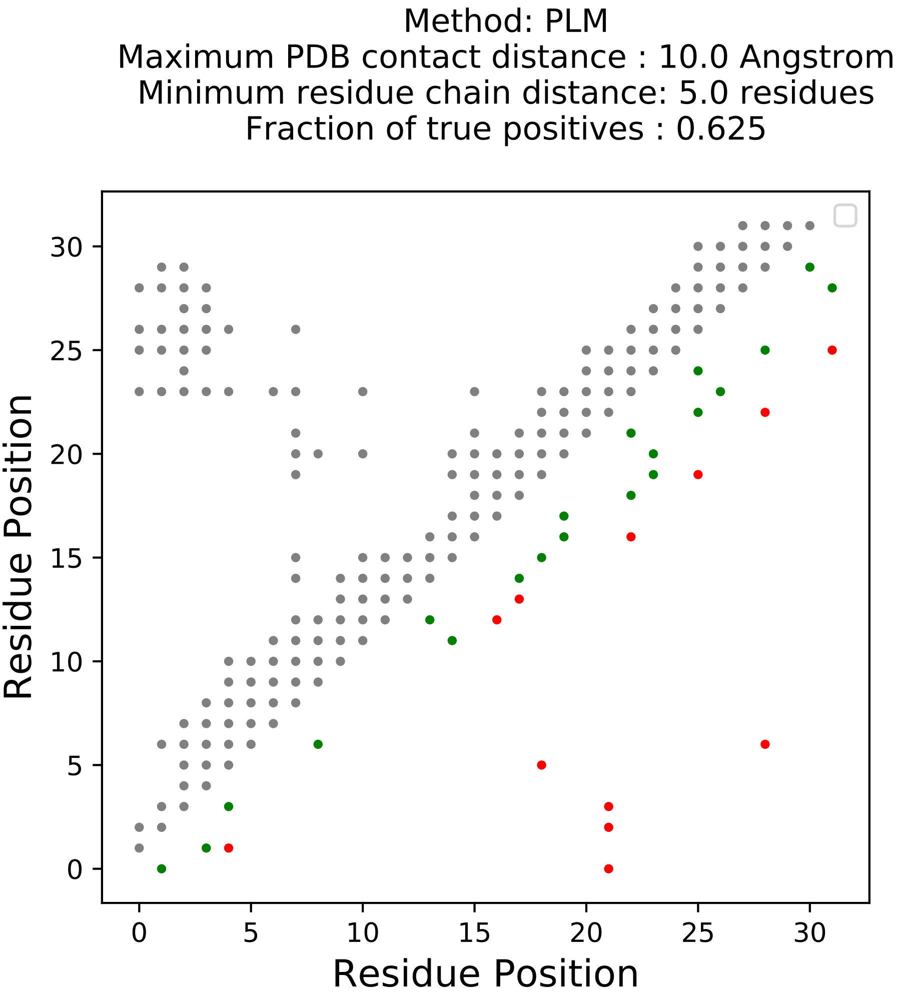

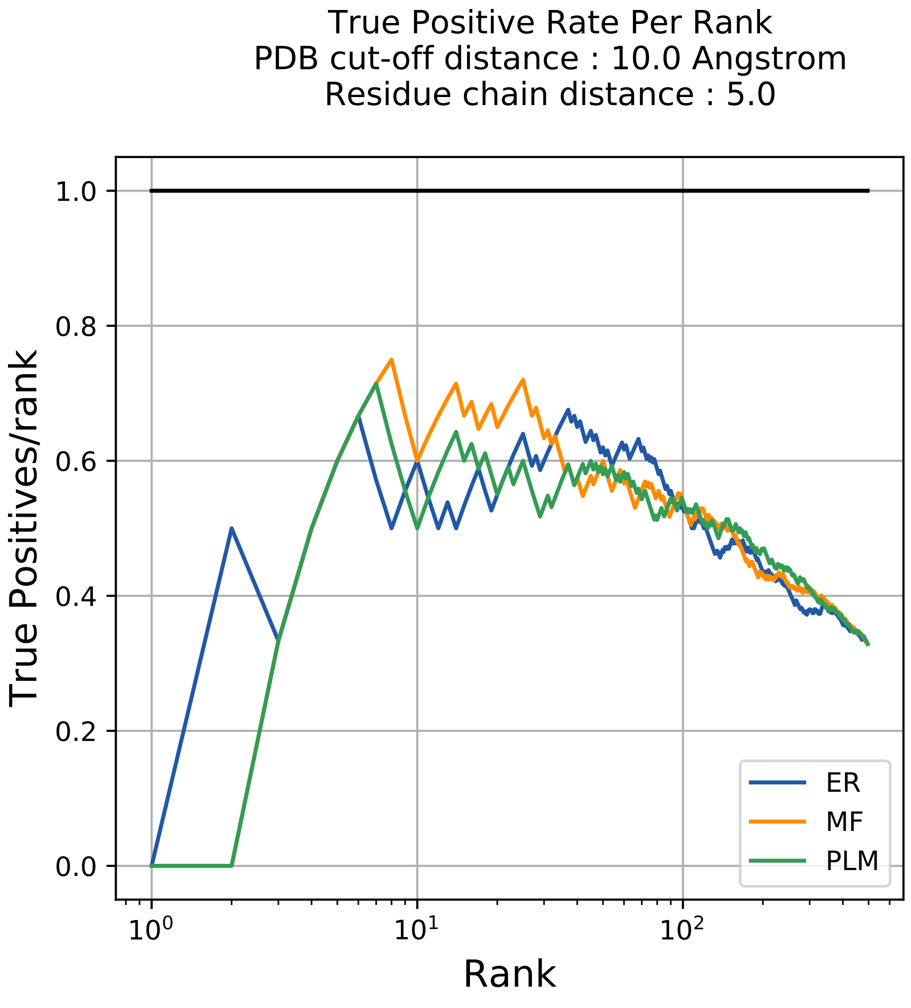

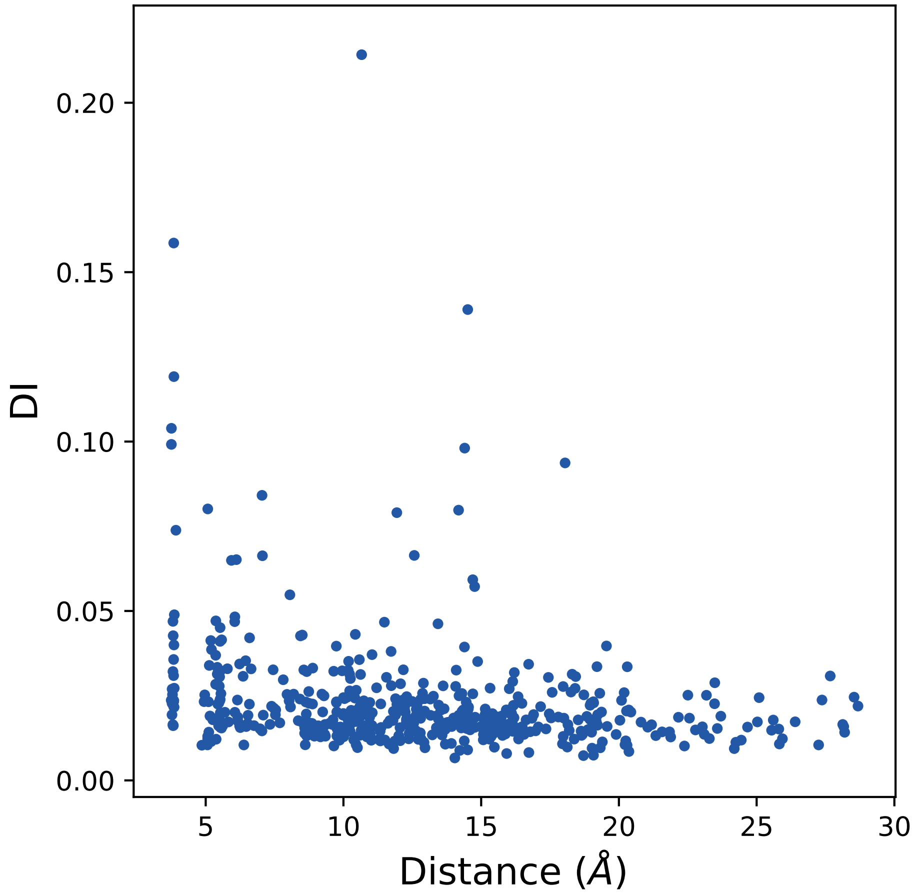

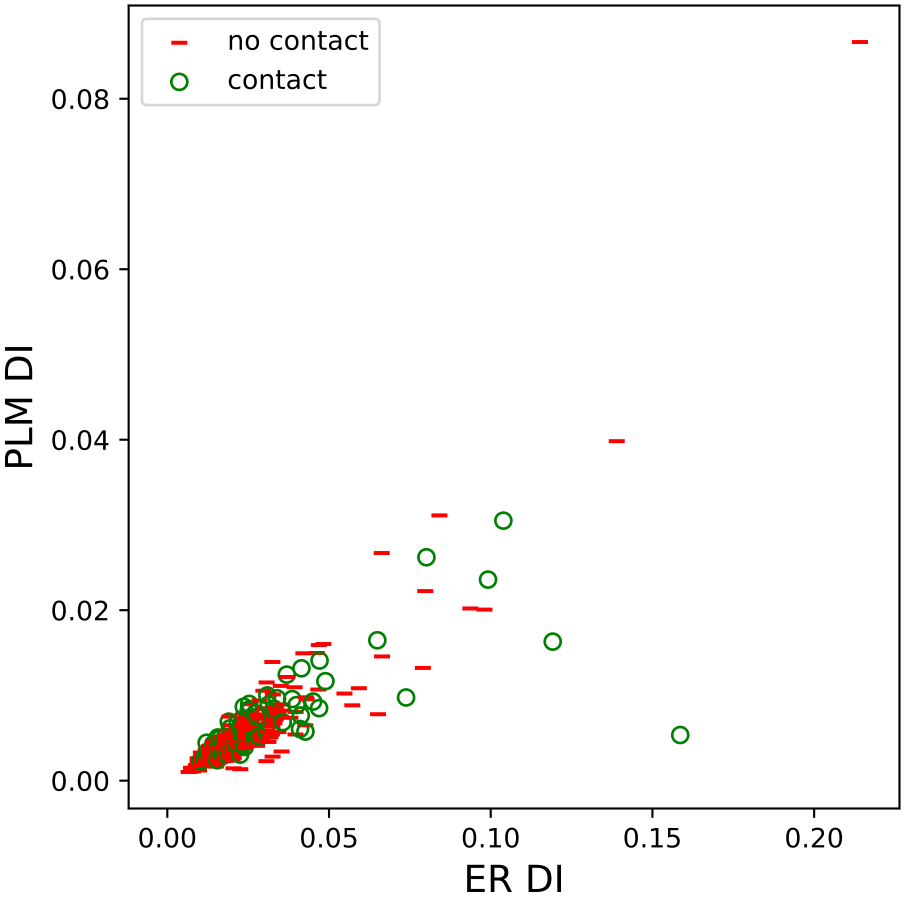

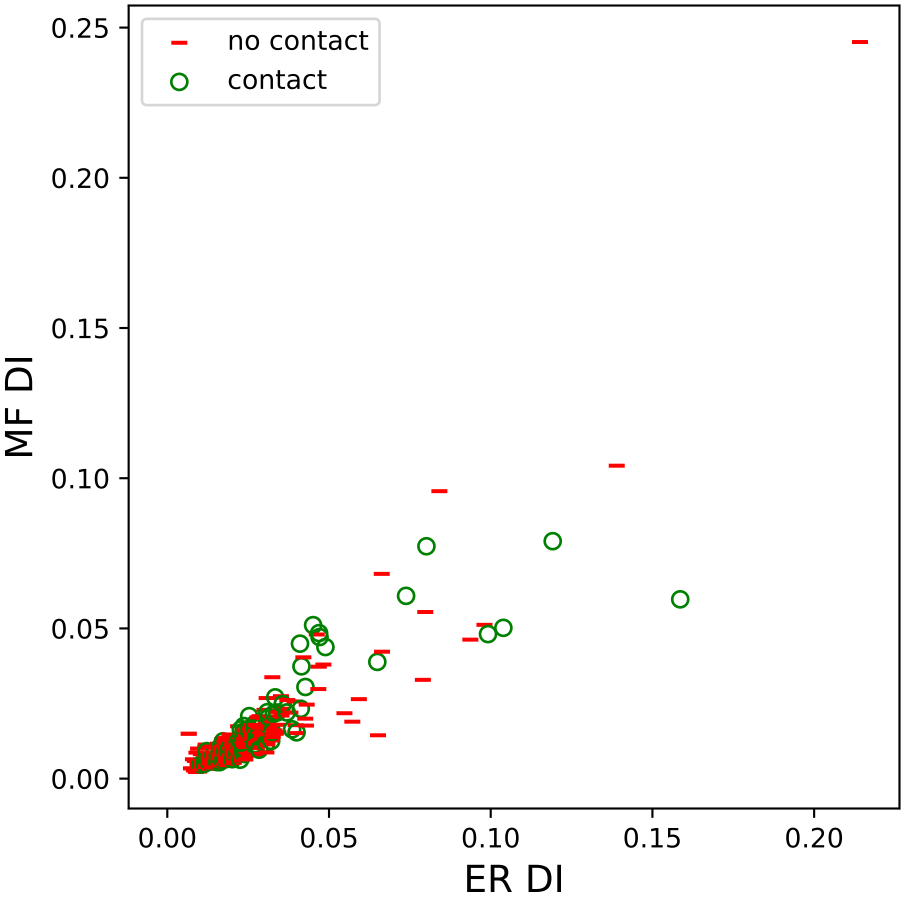

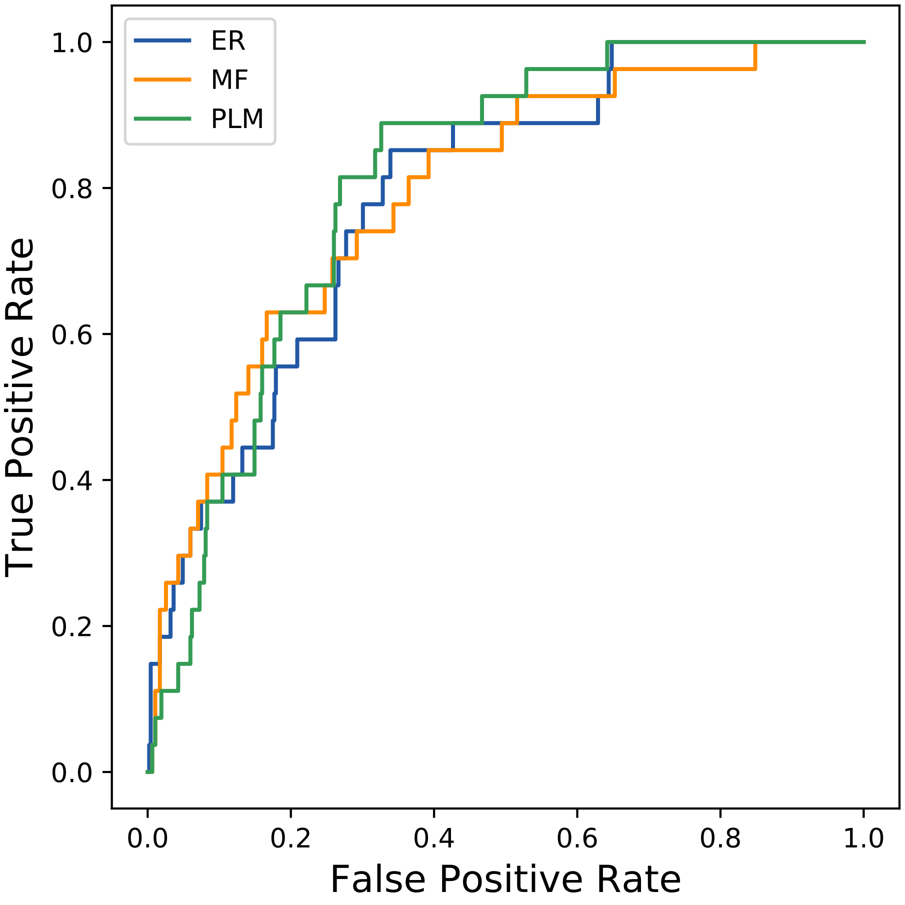

### 13244seq_61col_2bgf_PF00240_plots.pdf

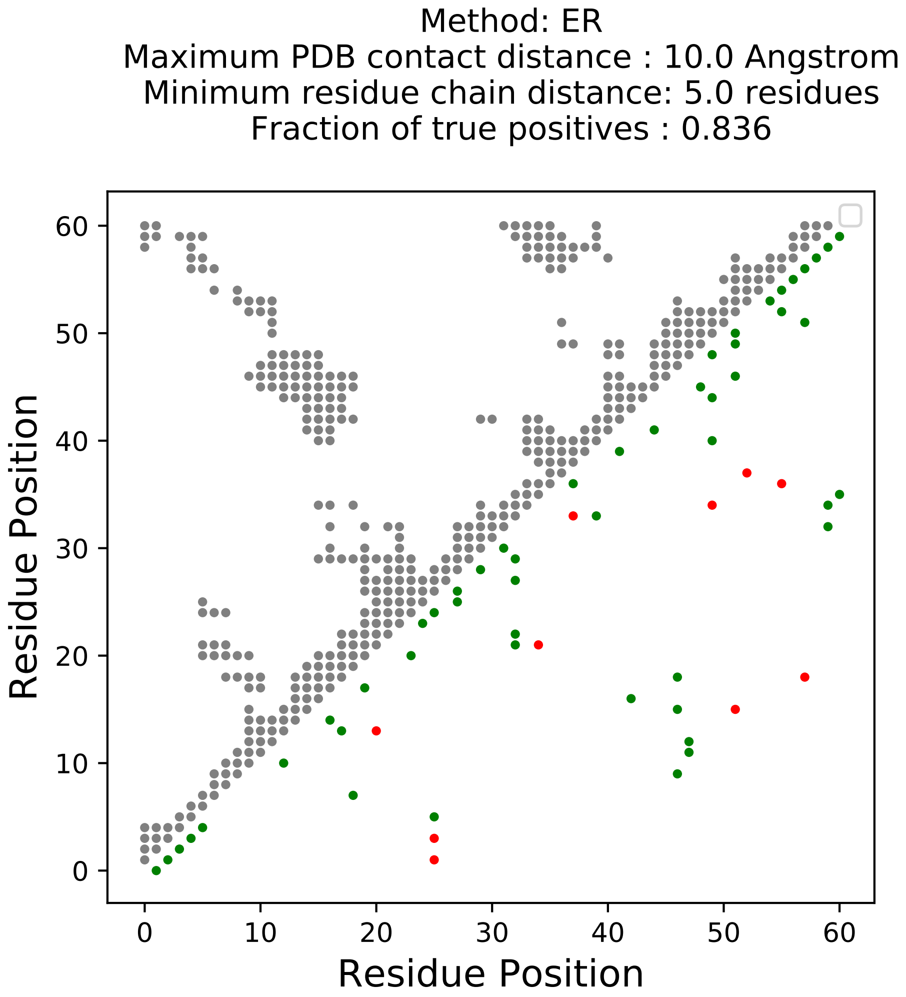

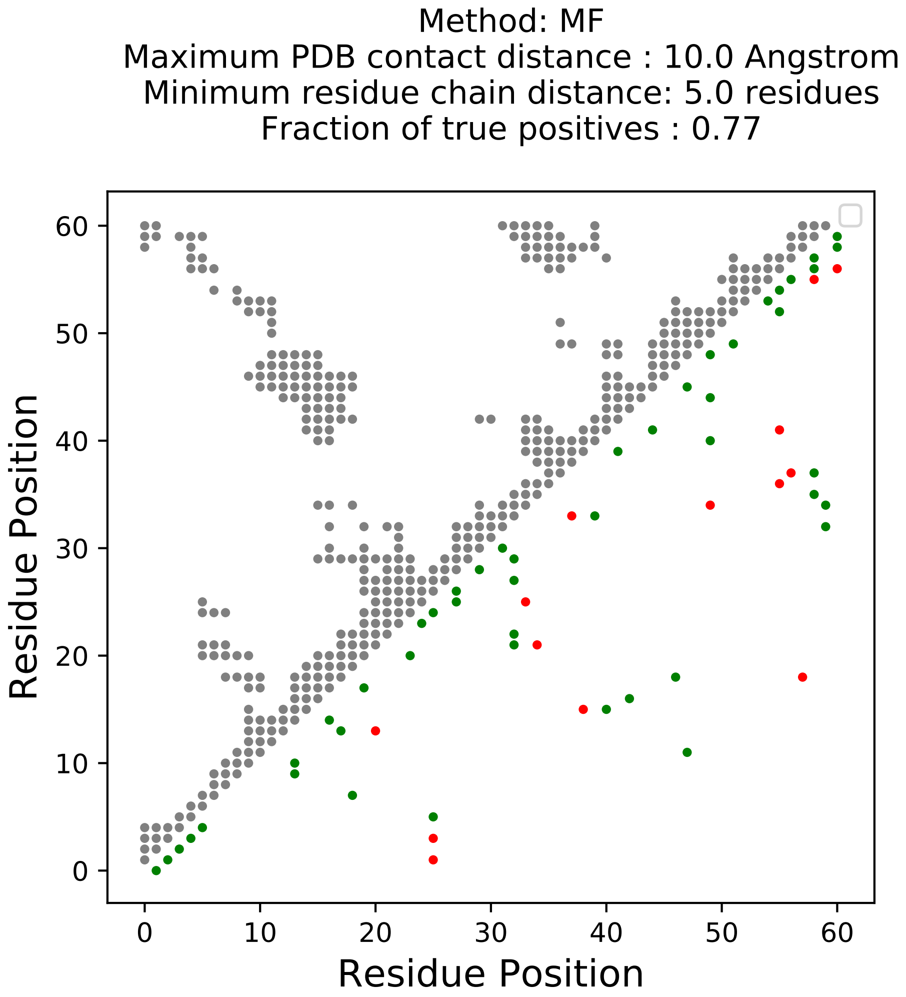

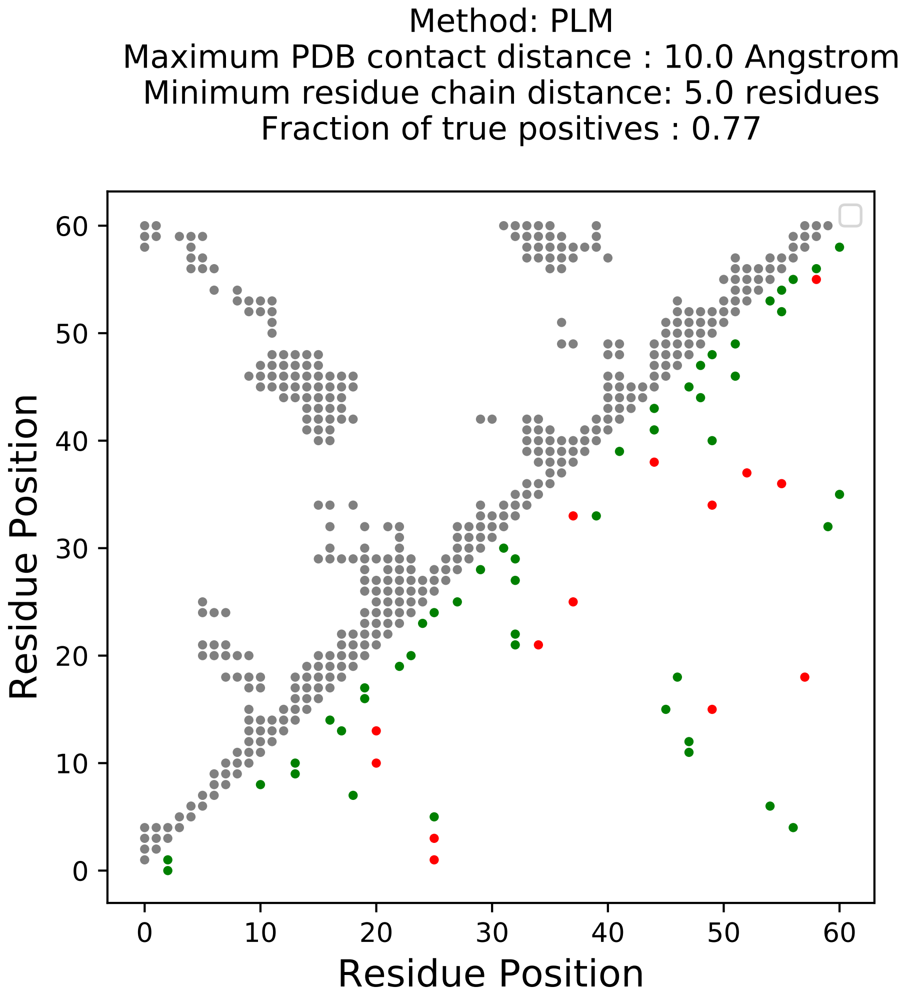

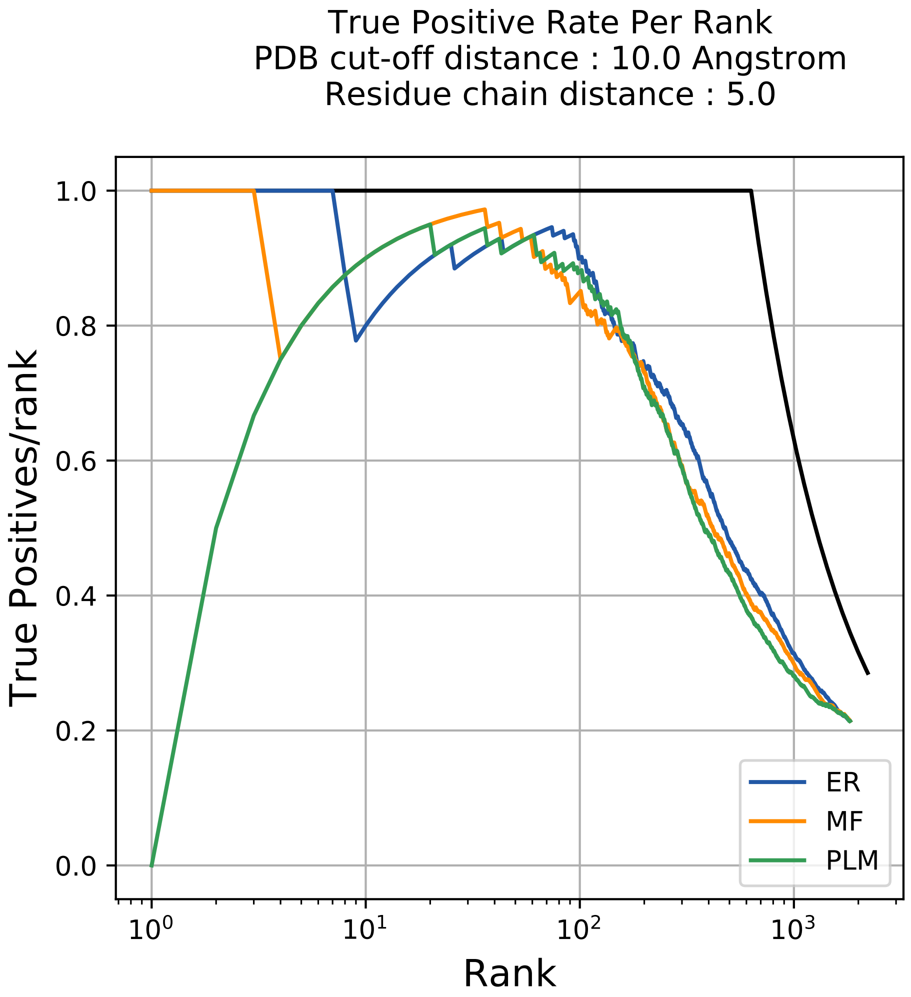

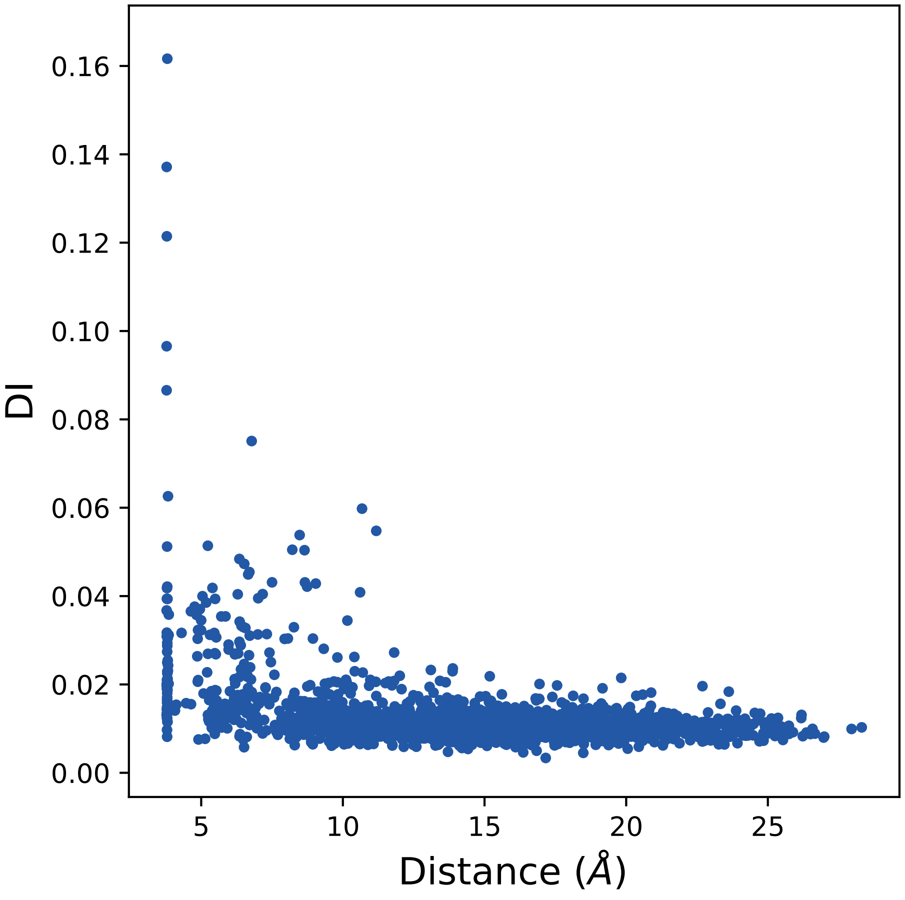

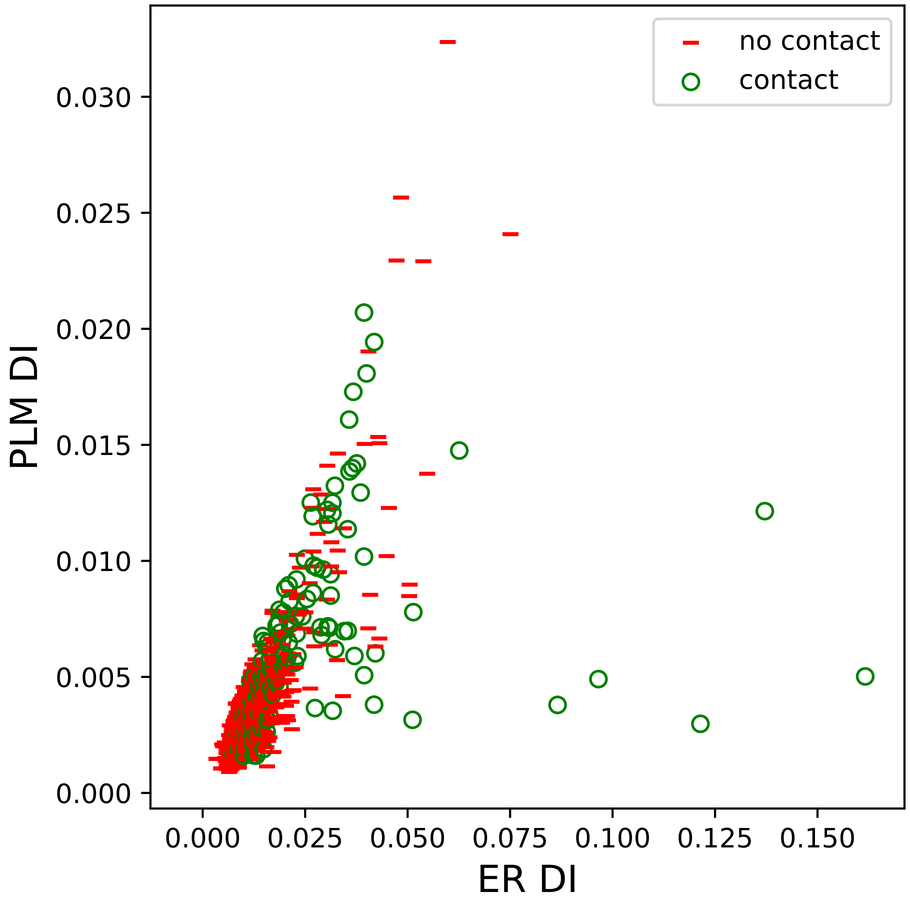

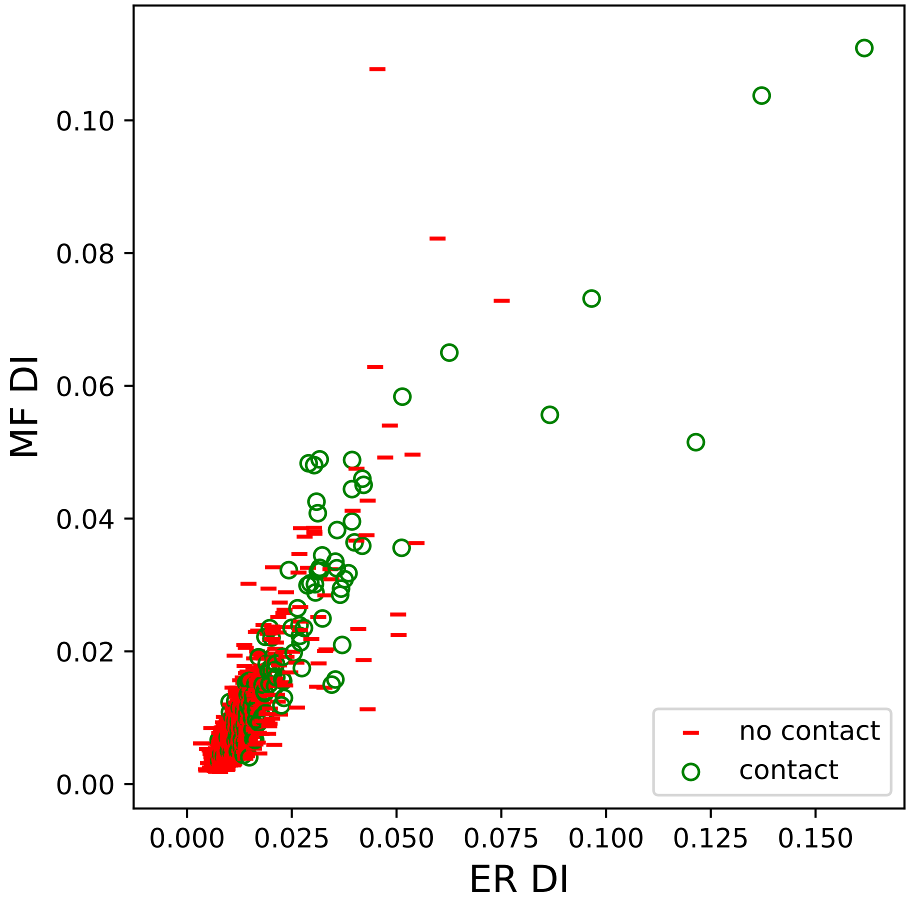

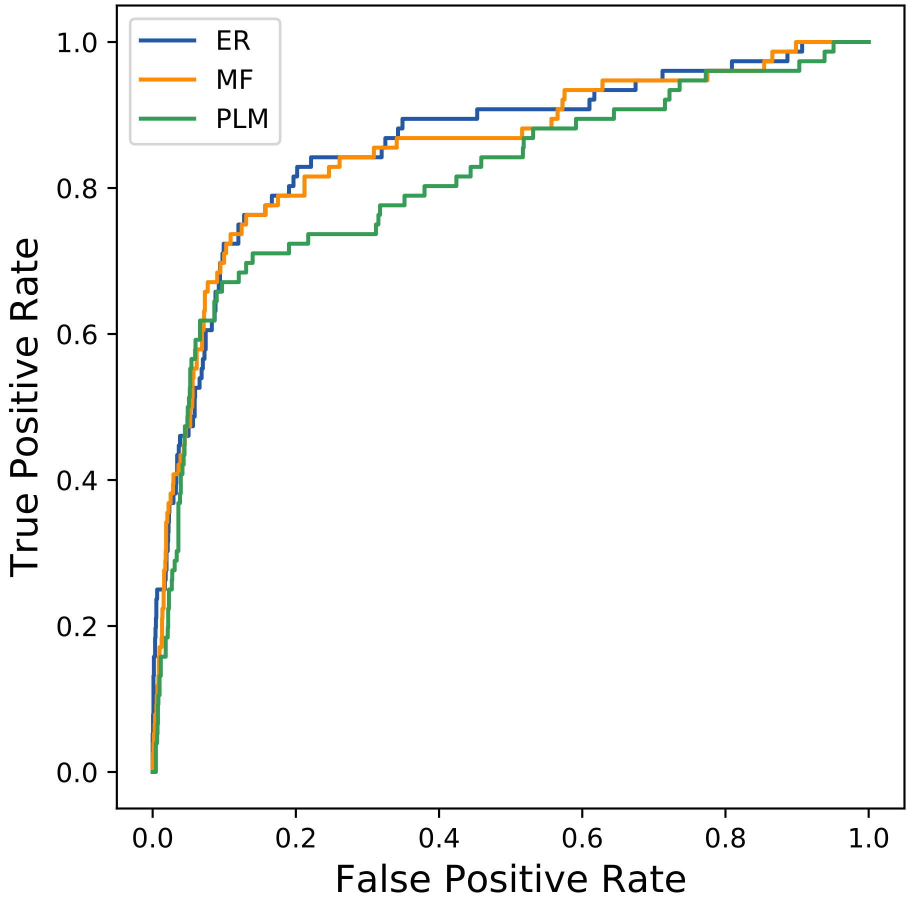

### 13566seq_62col_4ir3_PF00439_plots.pdf

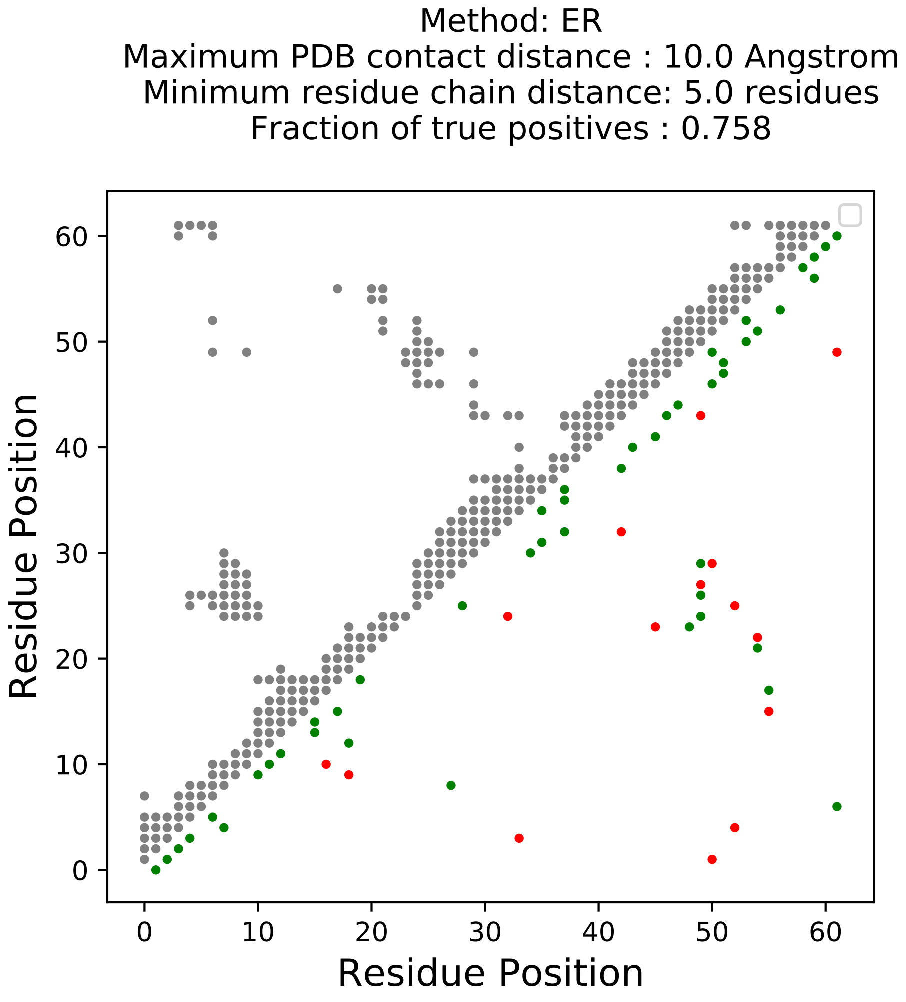

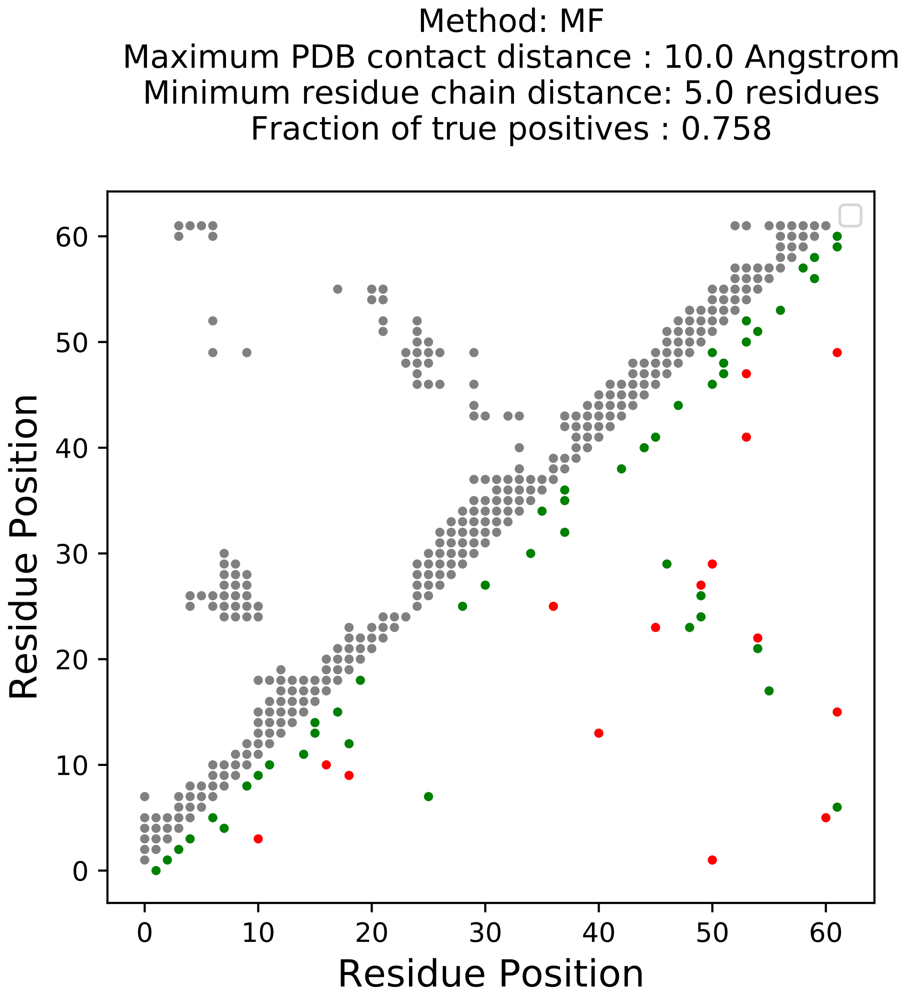

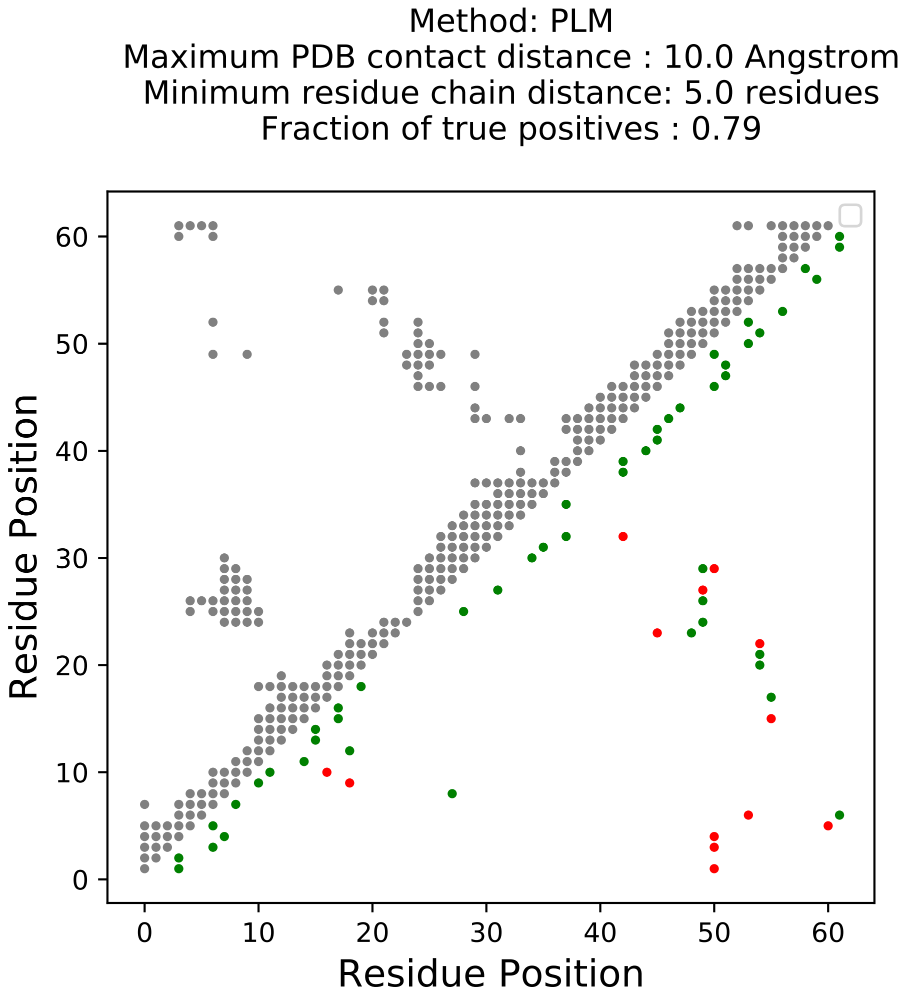

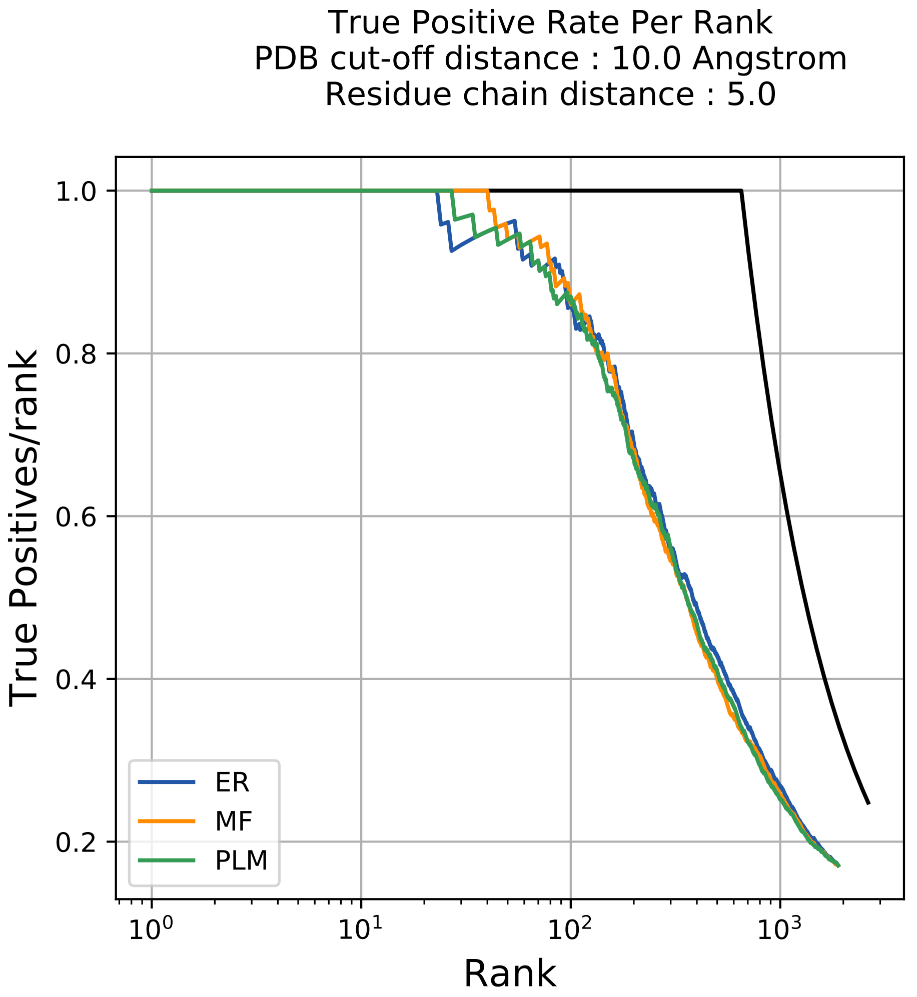

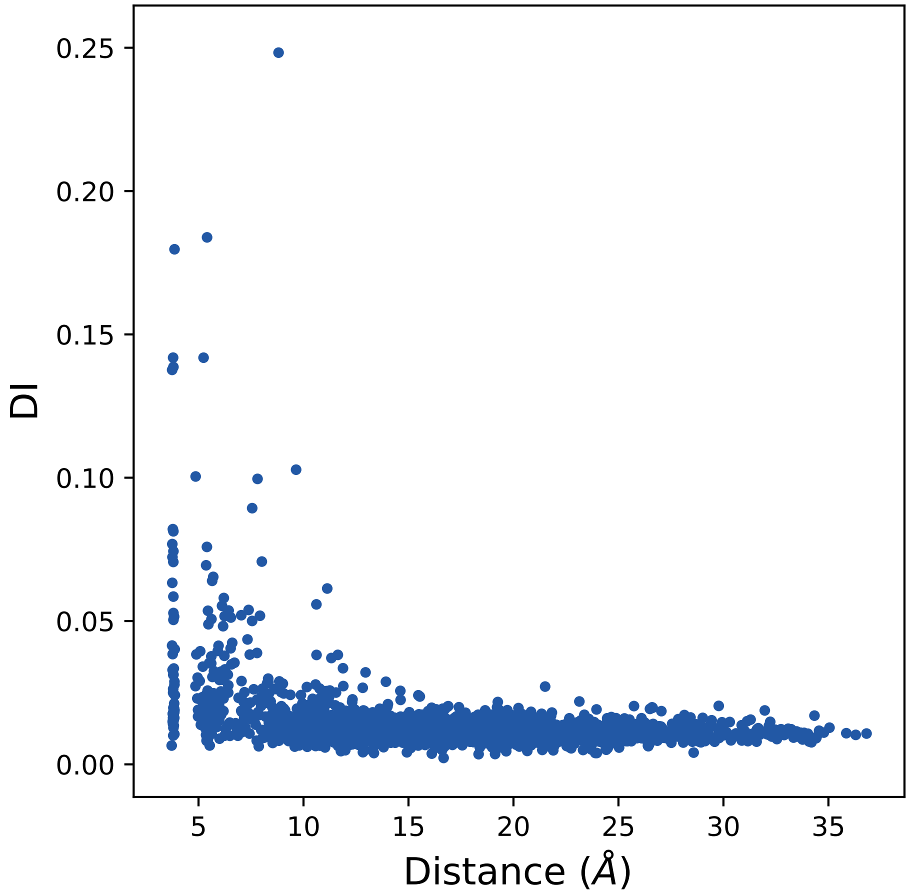

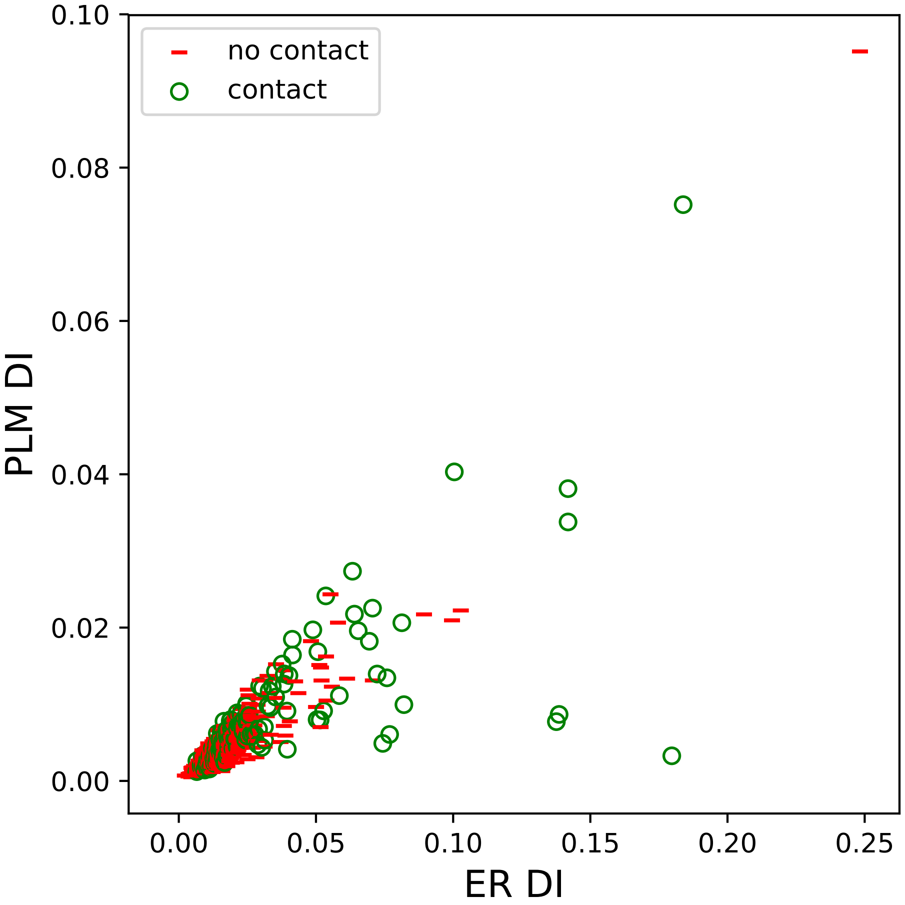

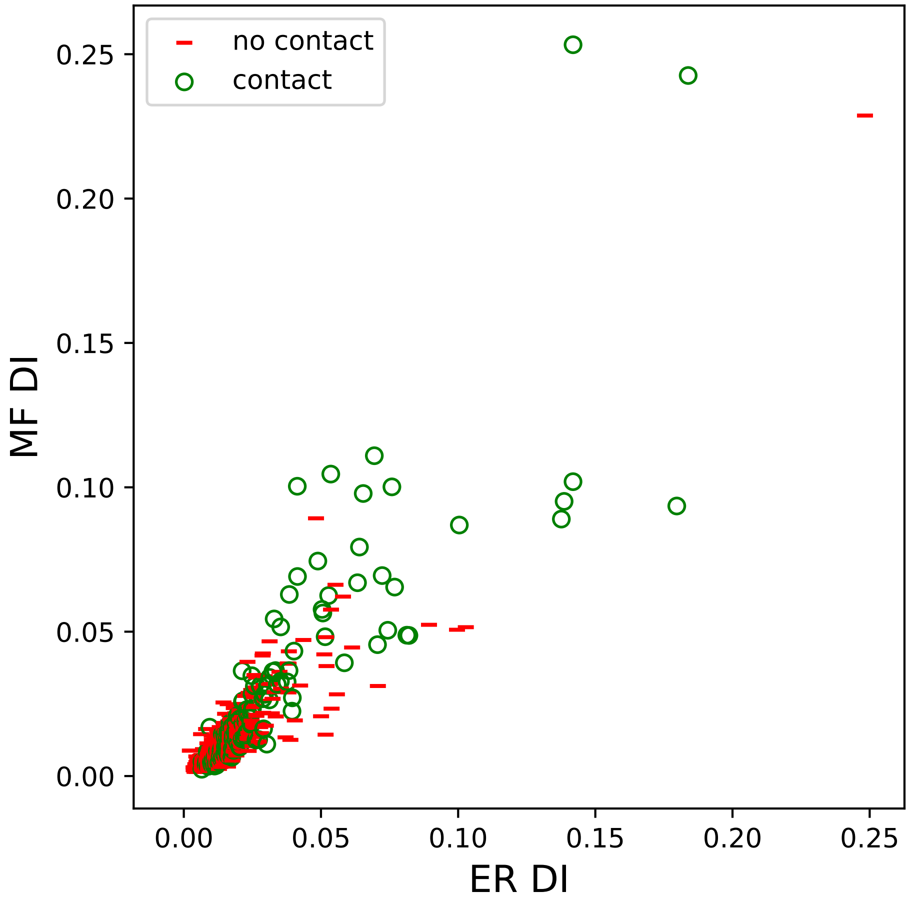

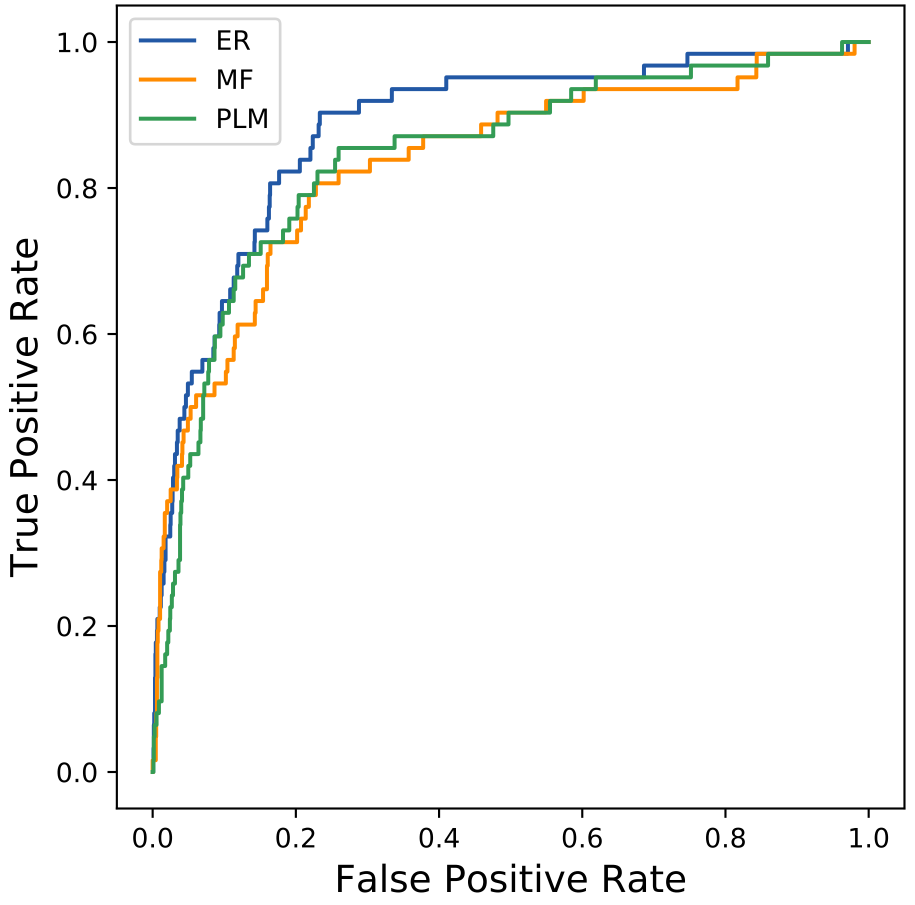

### 13816seq_58col_6eps_PF00439_plots.pdf

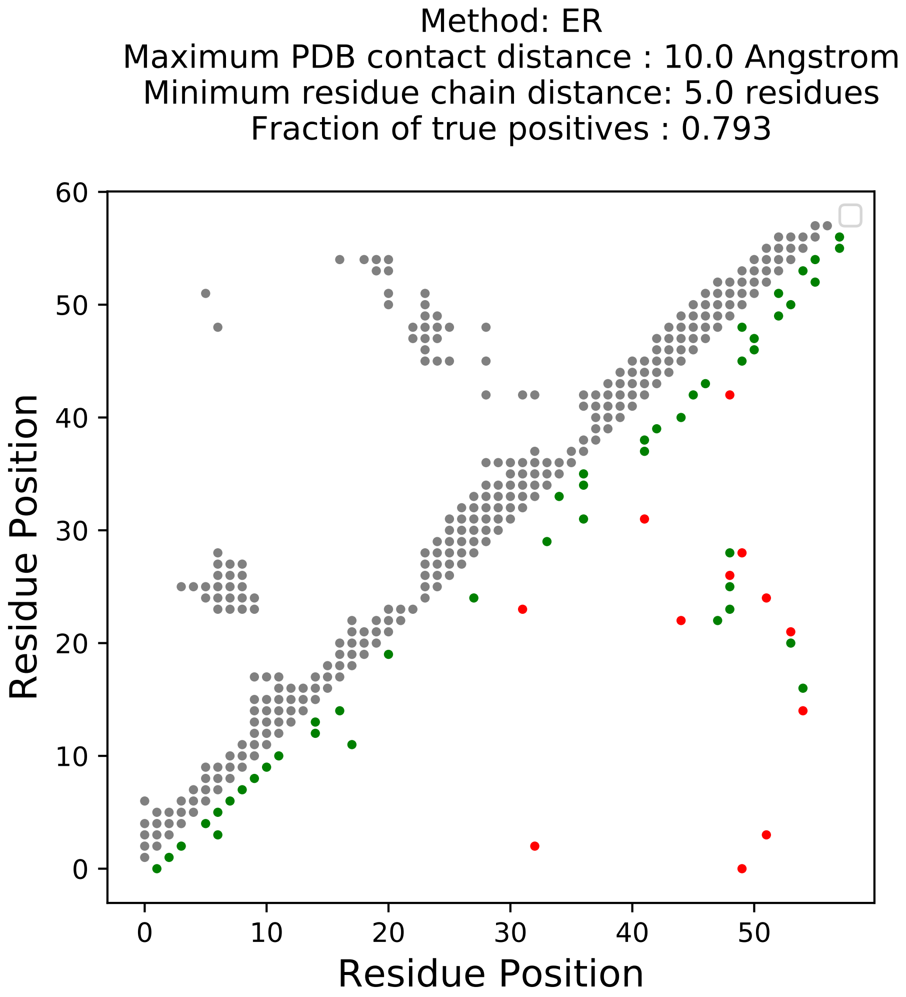

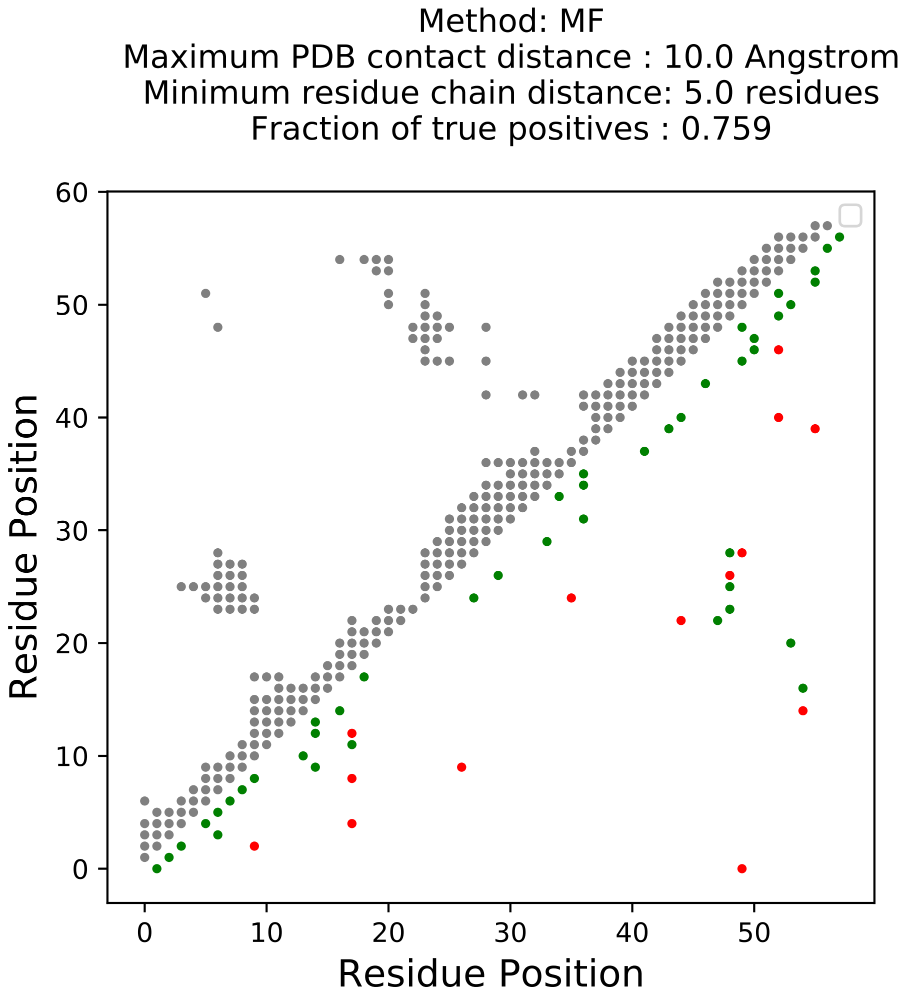

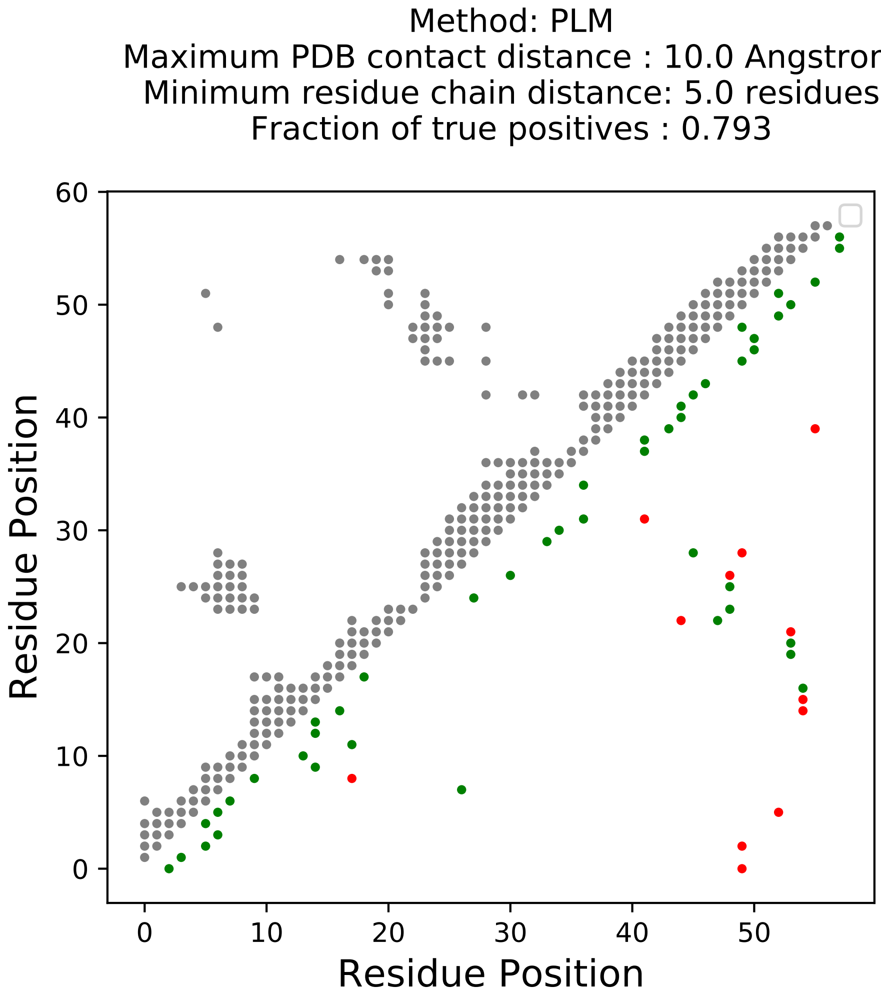

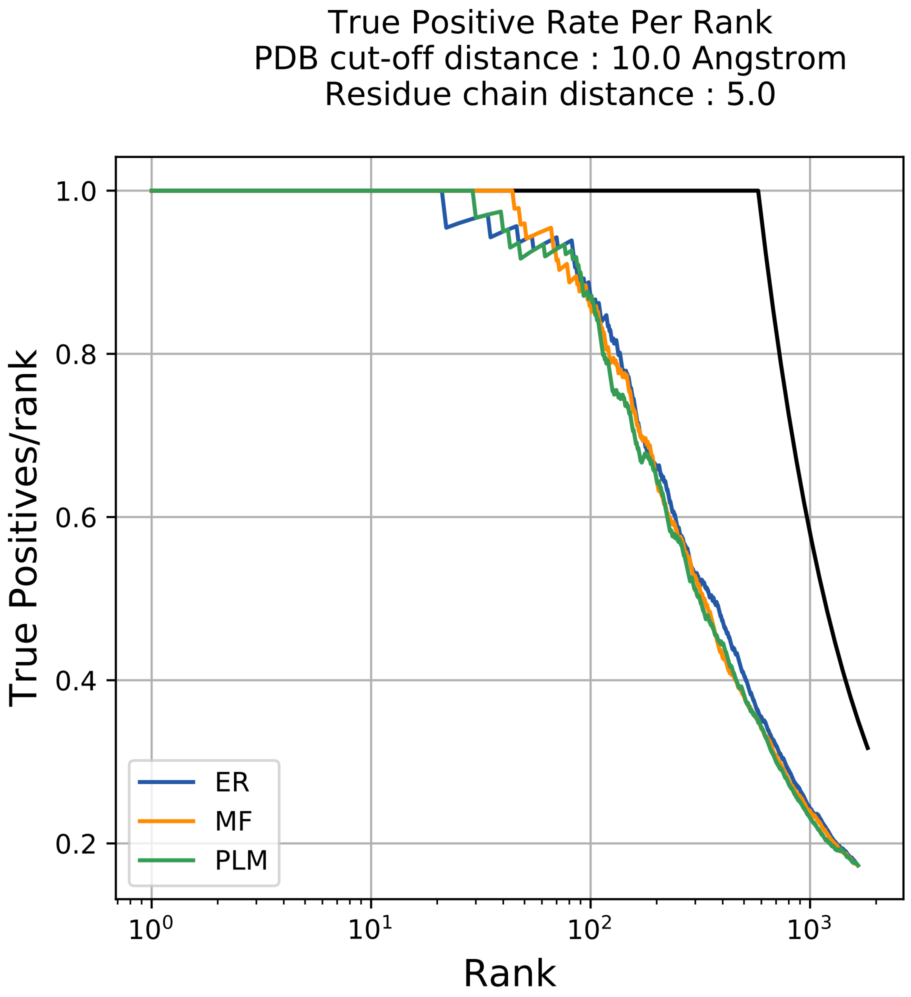

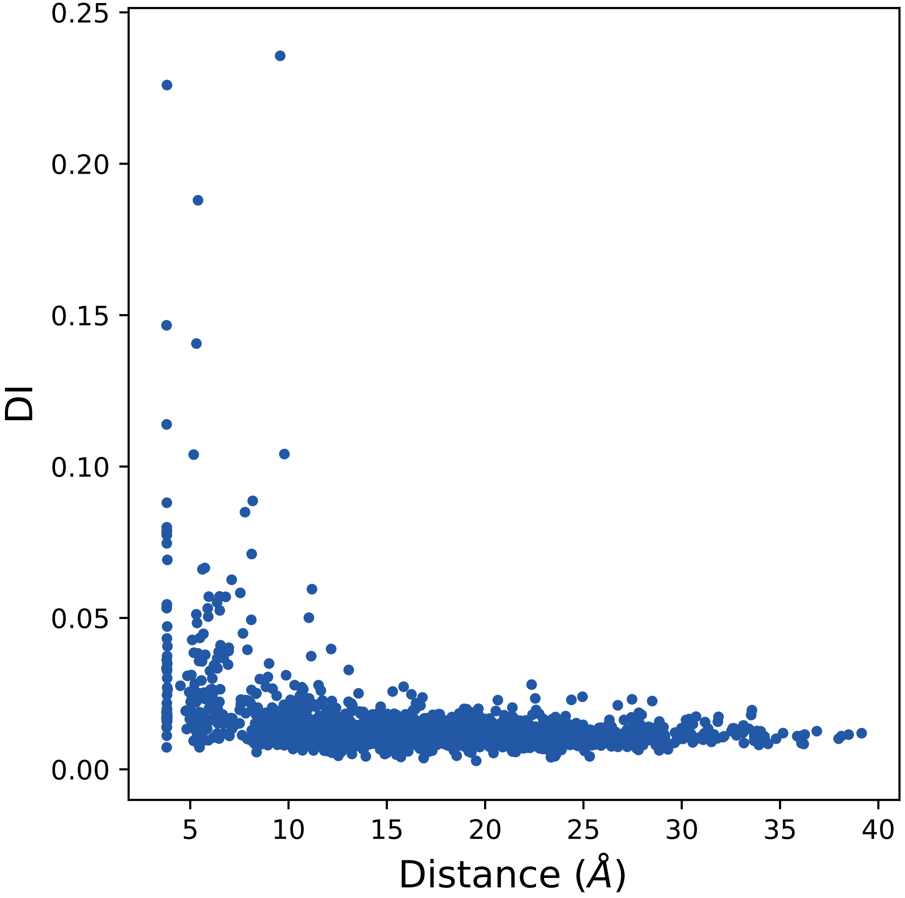

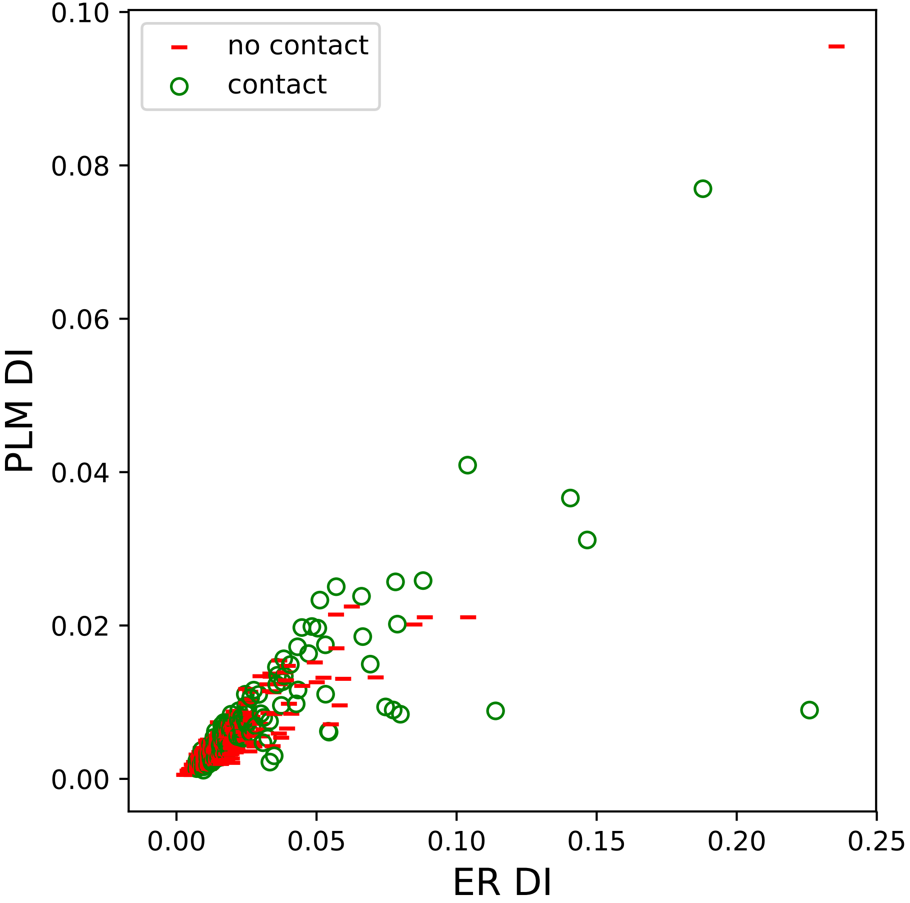

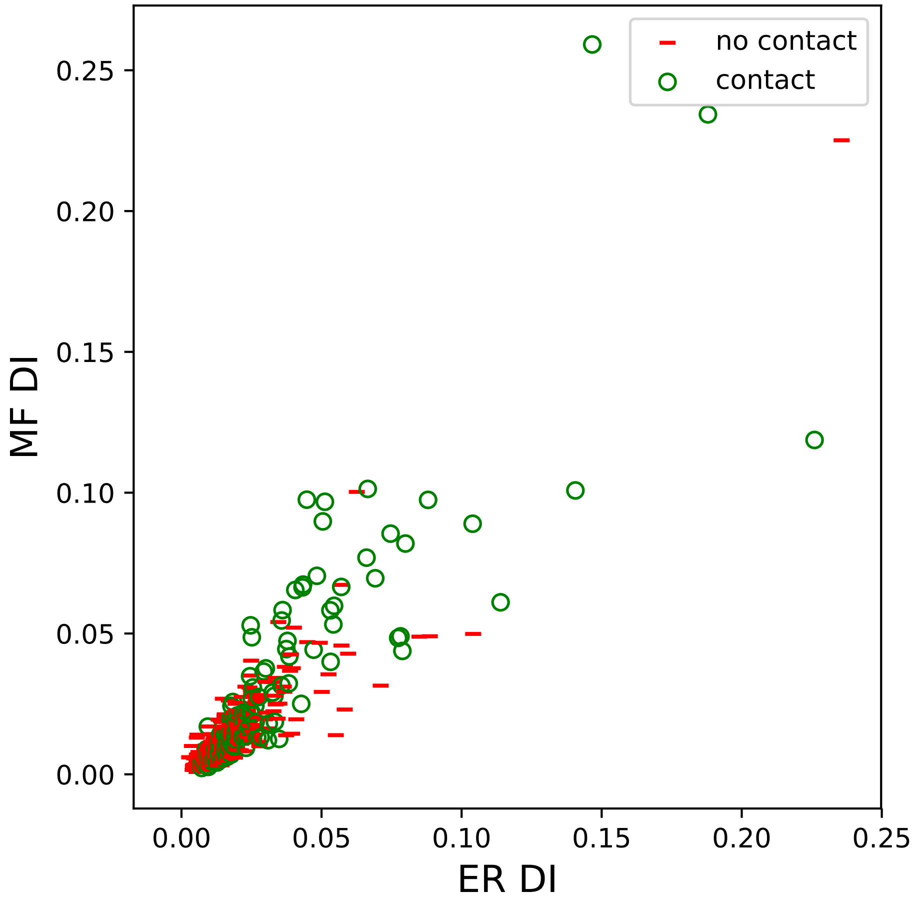

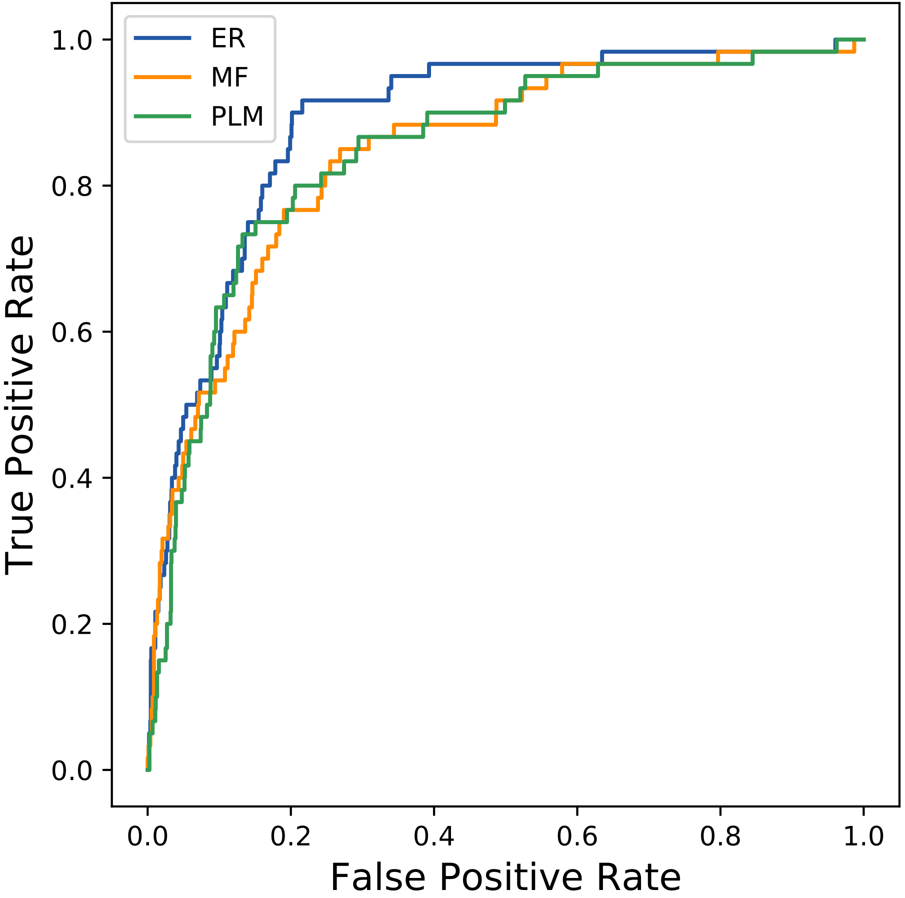

### 14121seq_207col_1aj0_PF00809_plots.pdf

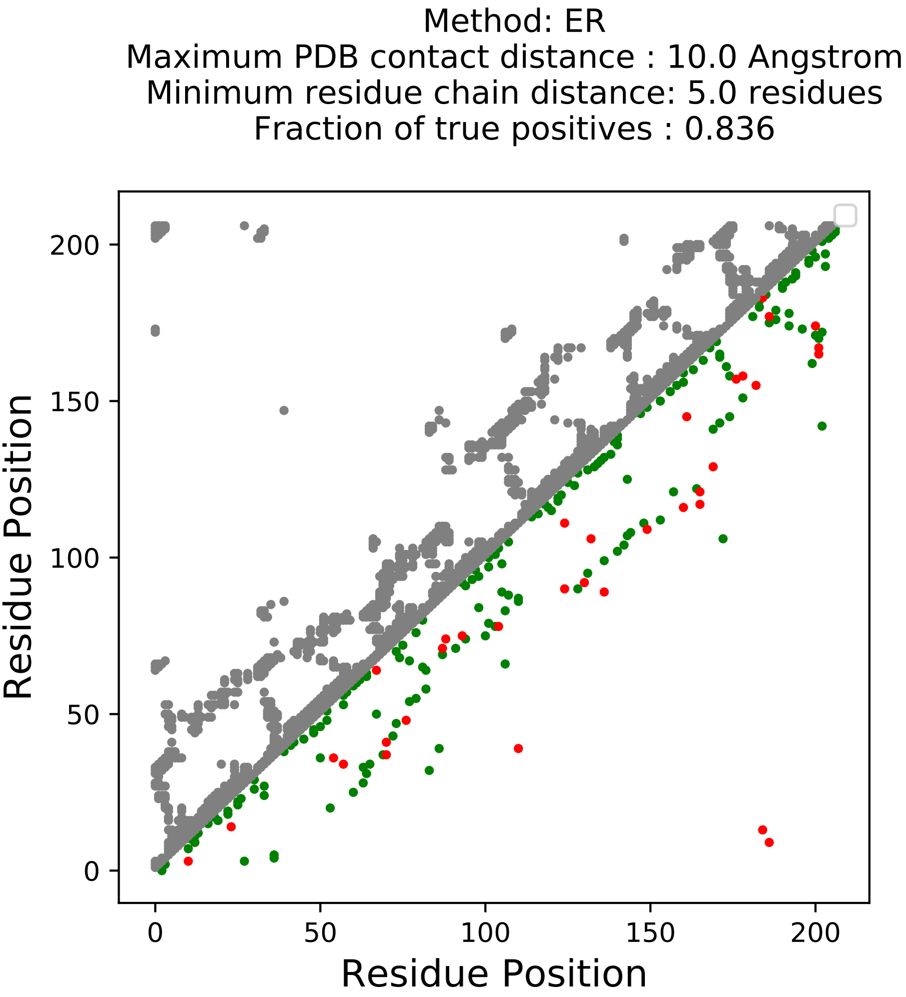

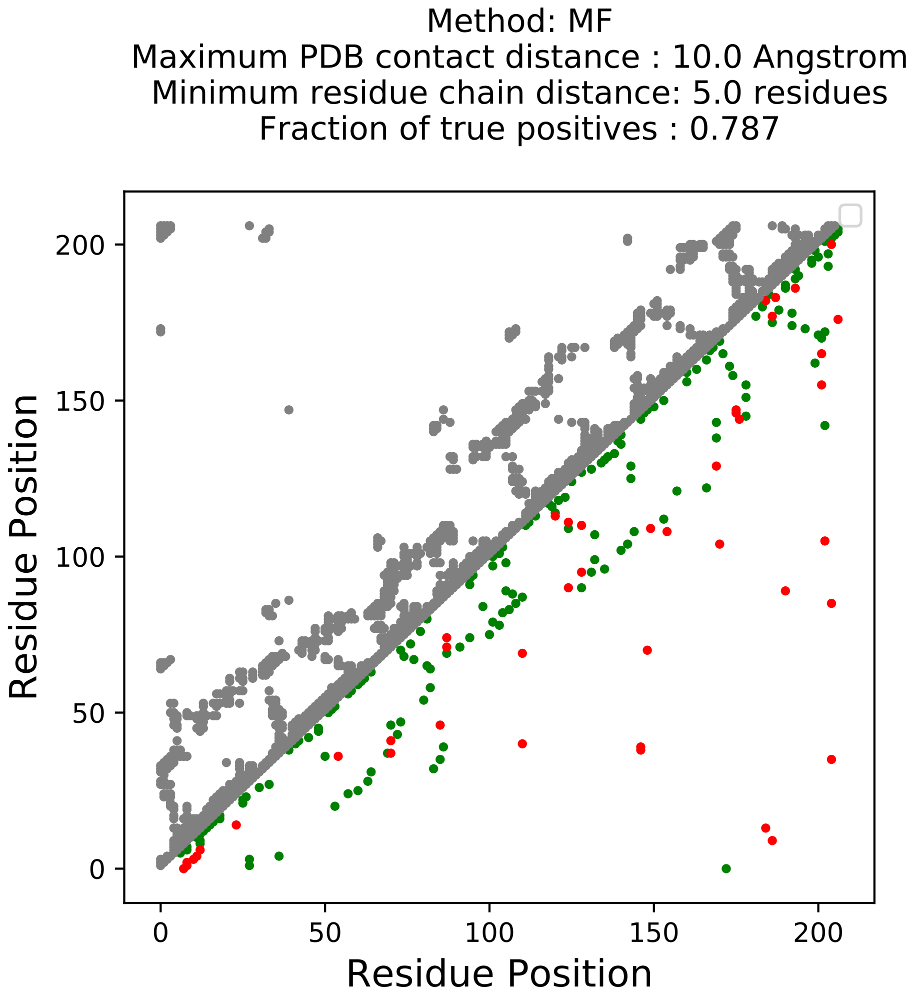

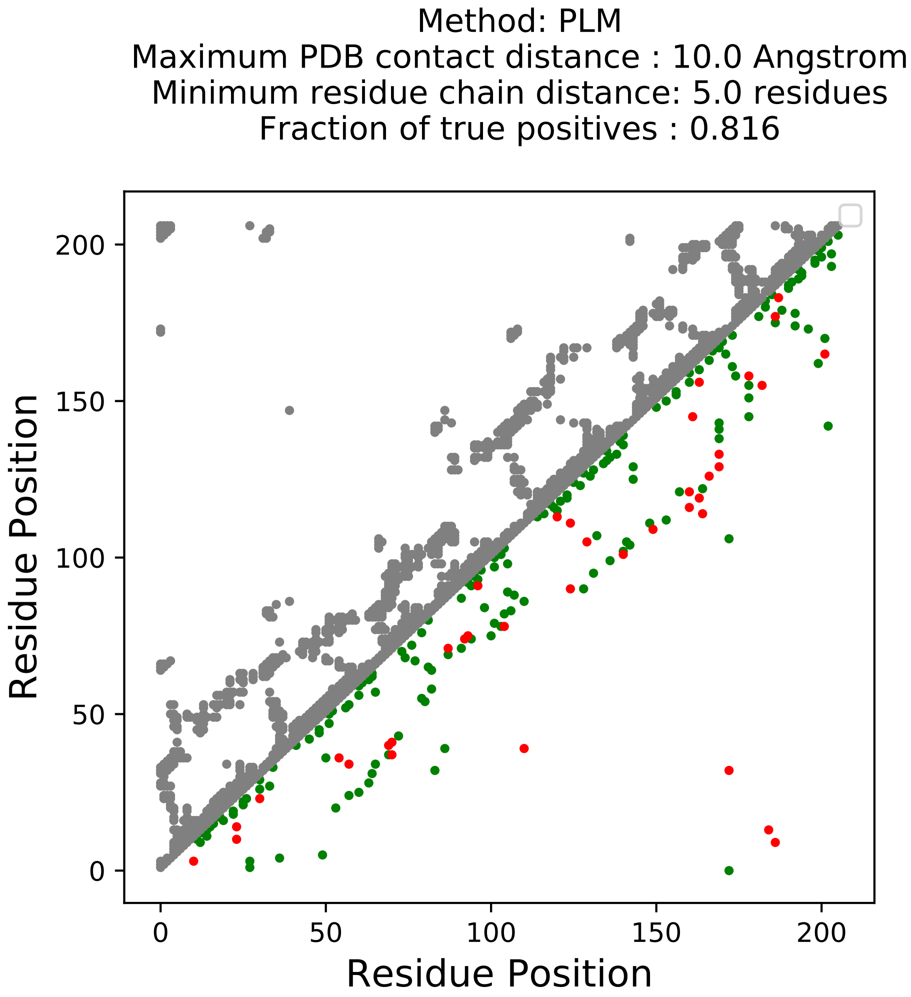

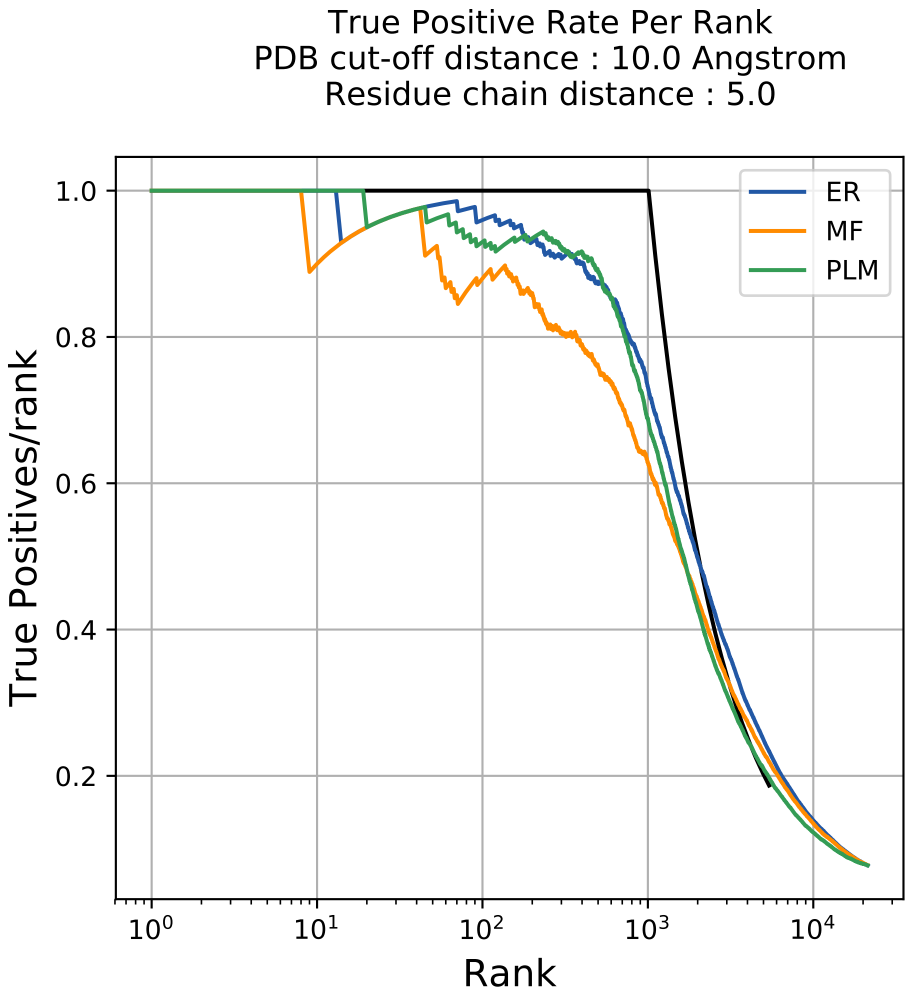

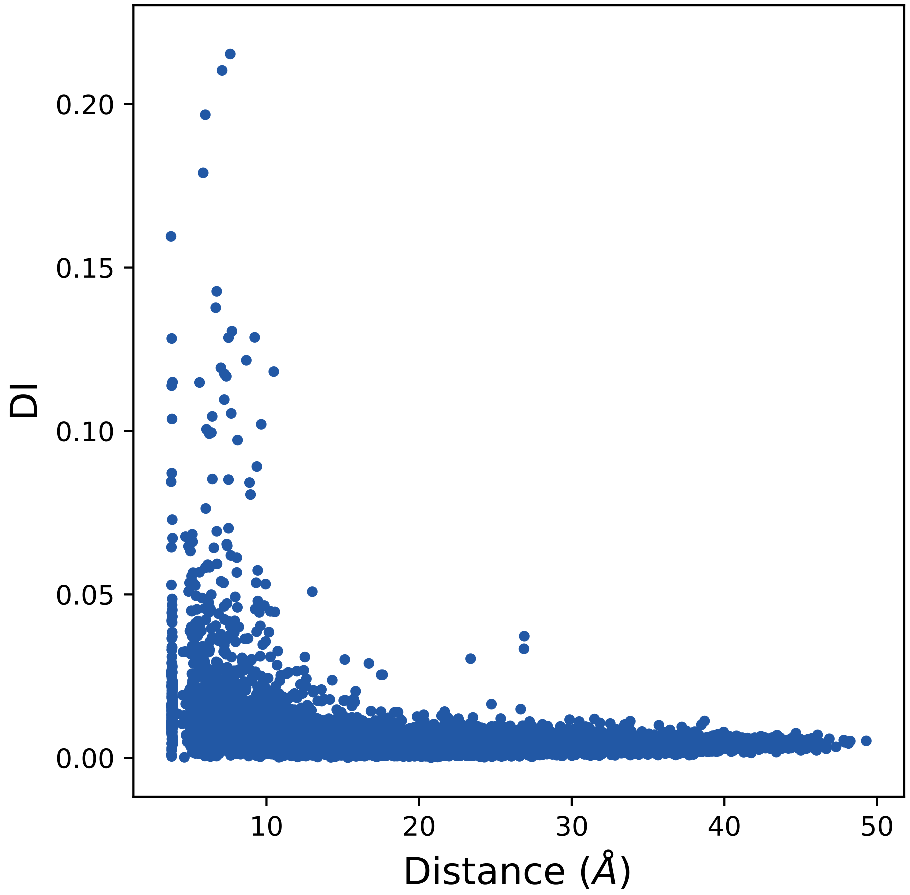

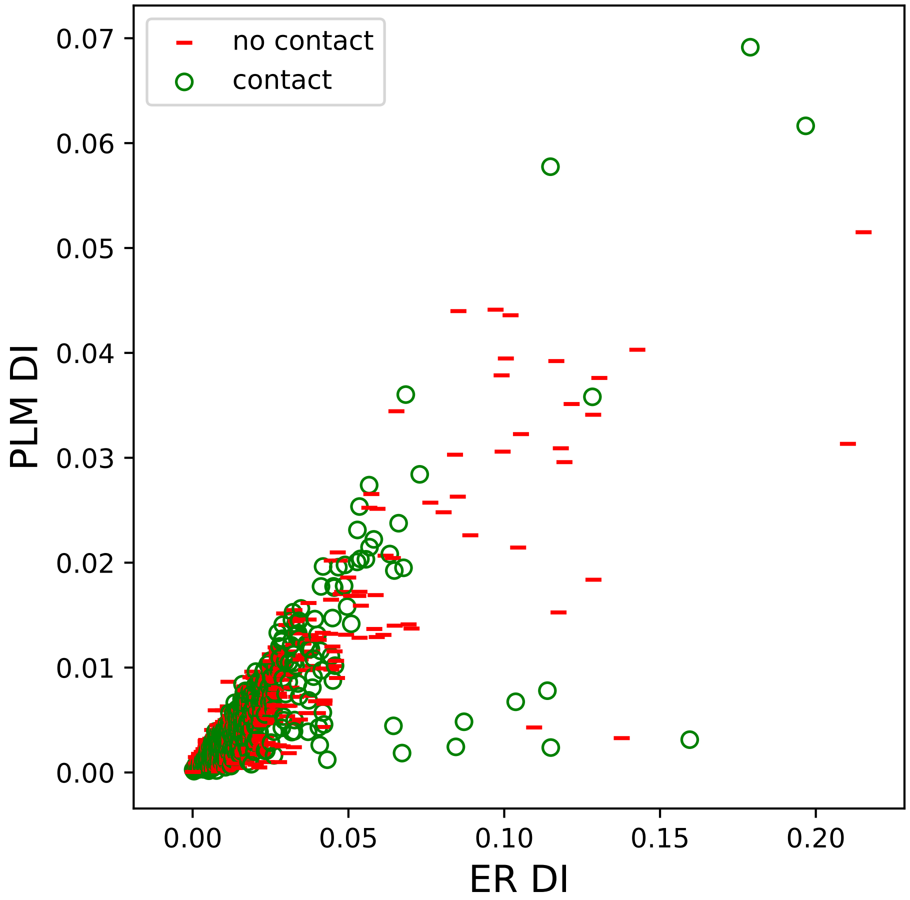

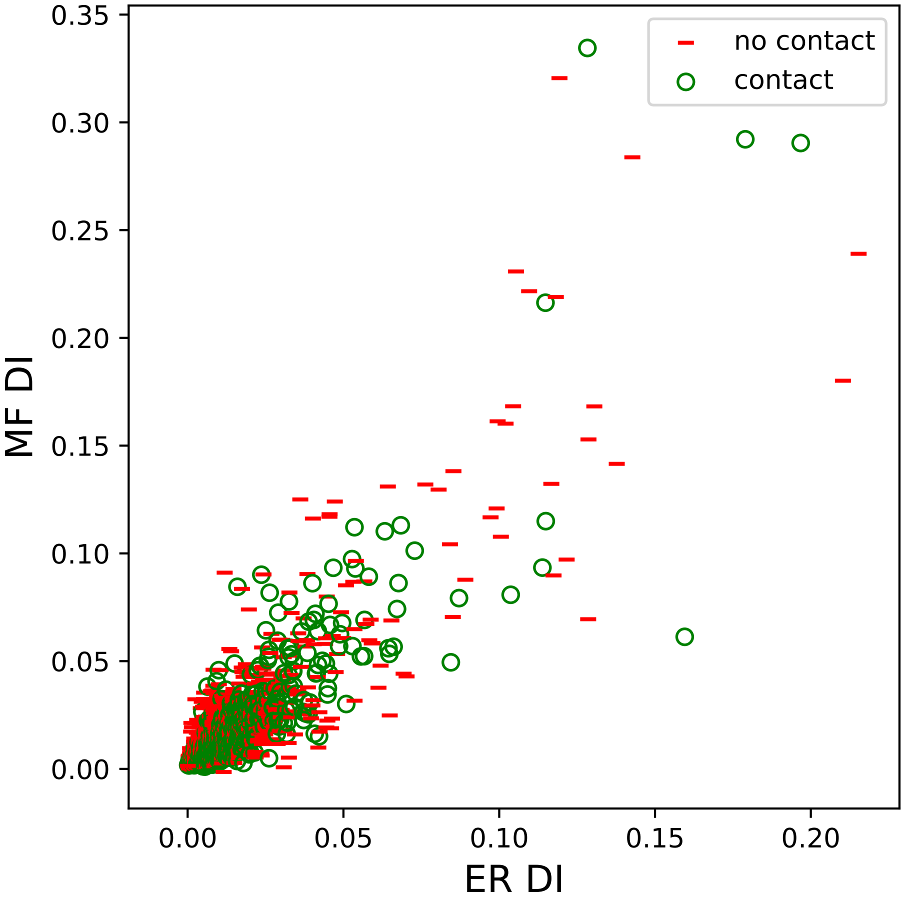

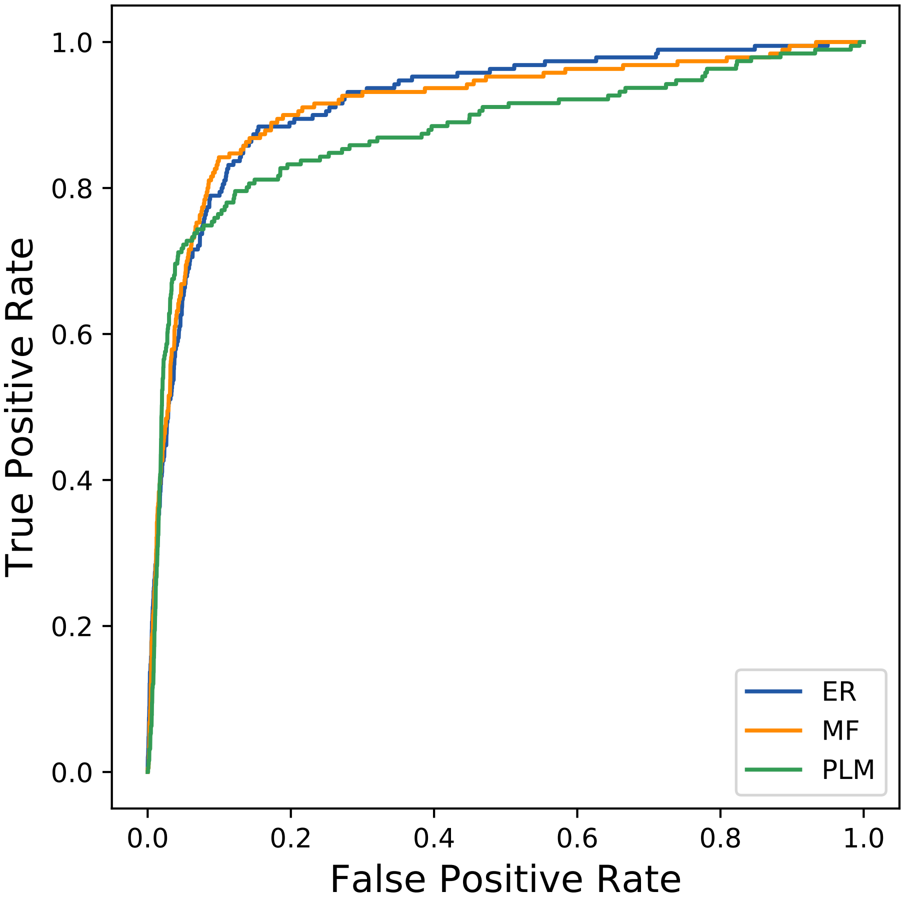

### 14767seq_186col_5d2w_PF00977_plots.pdf

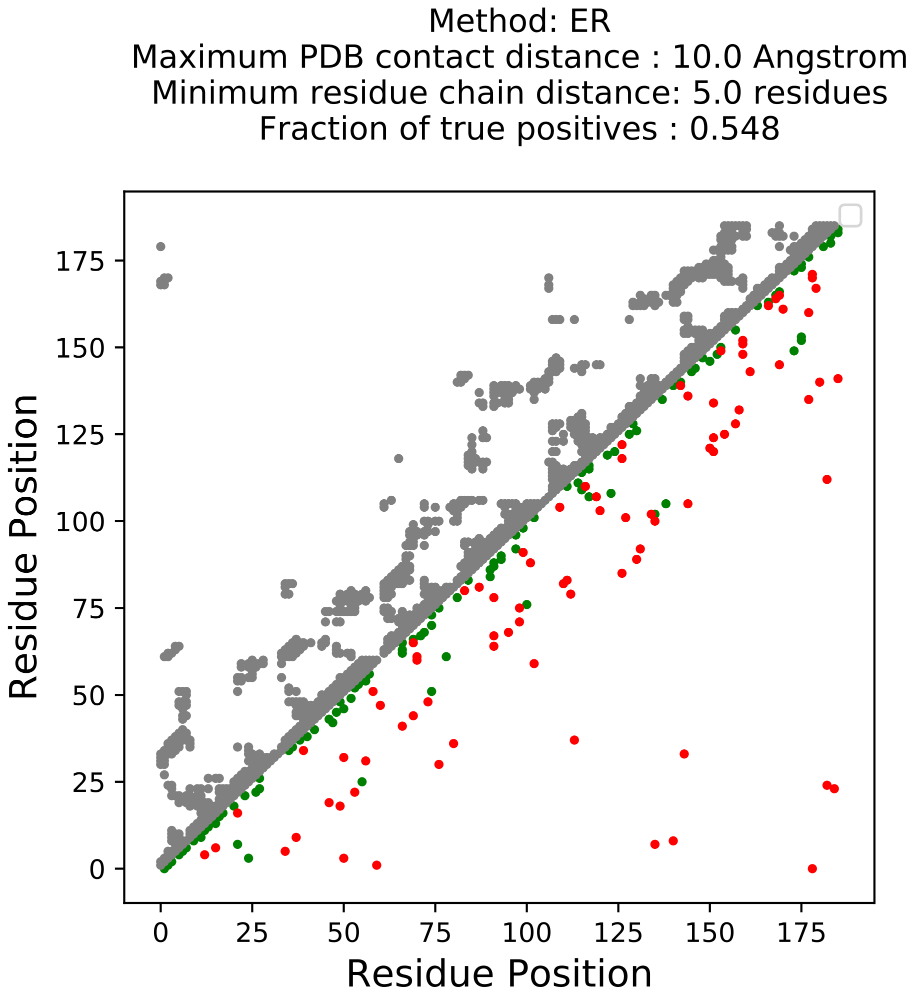

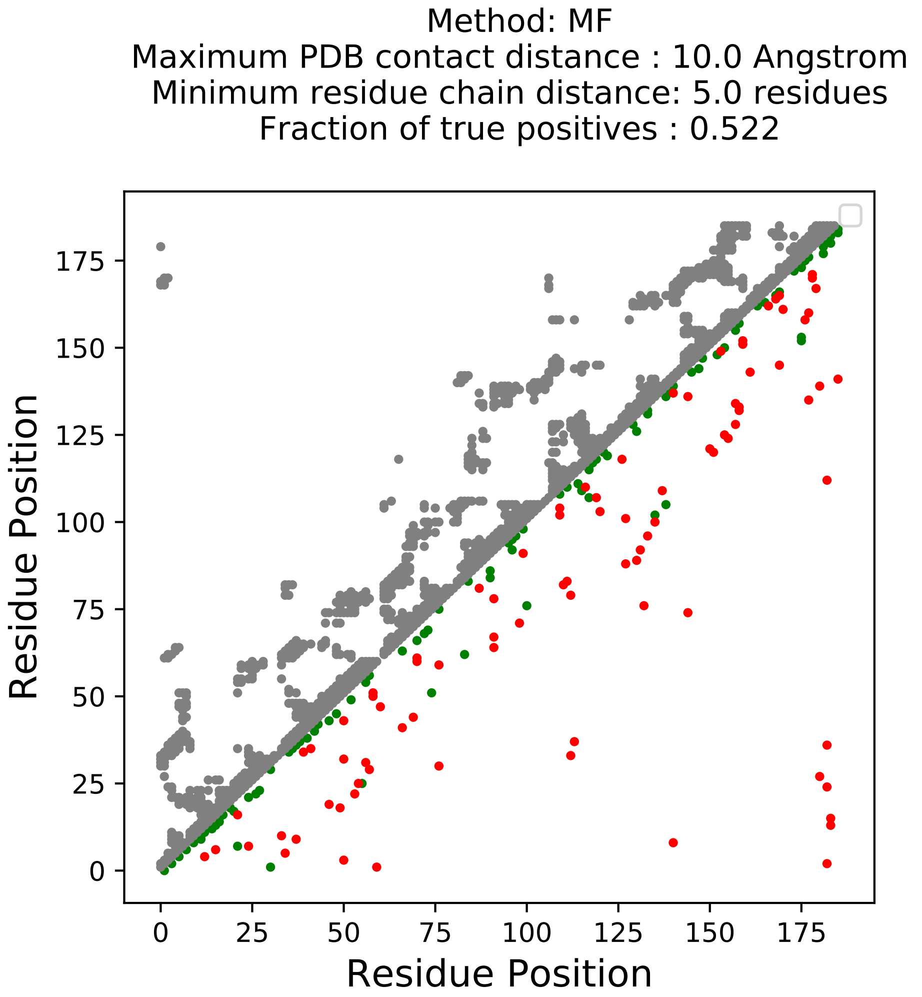

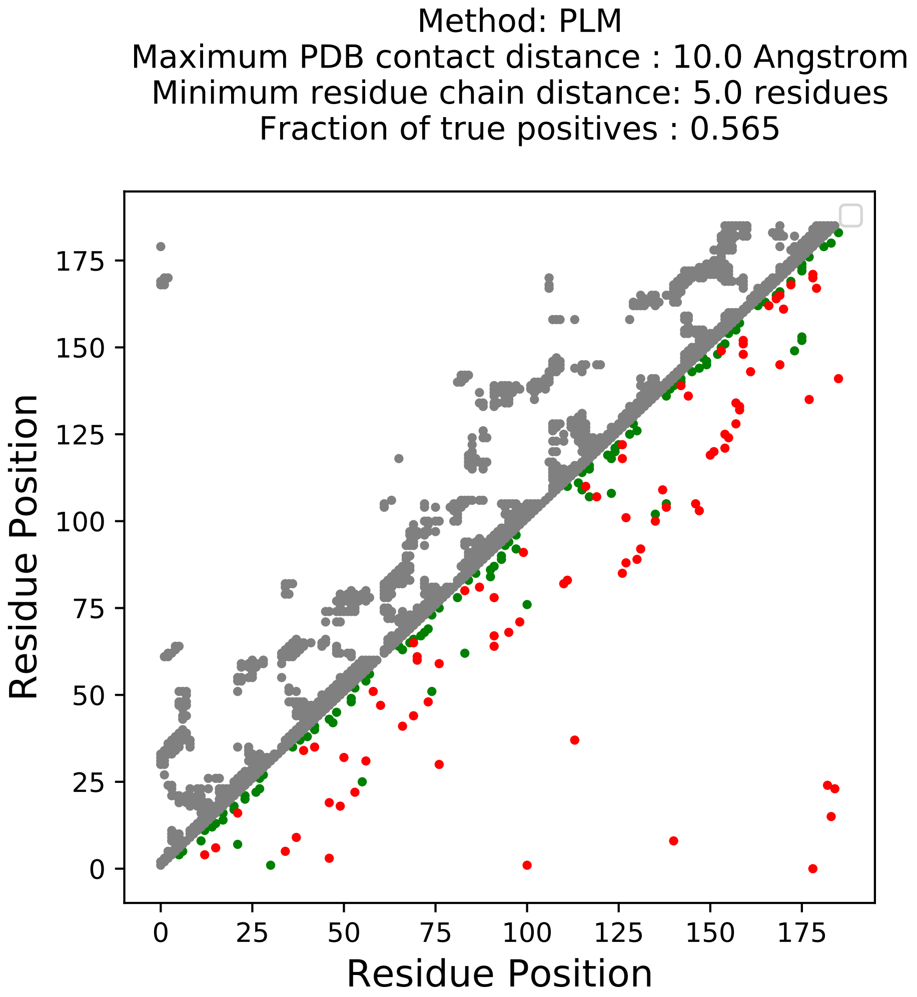

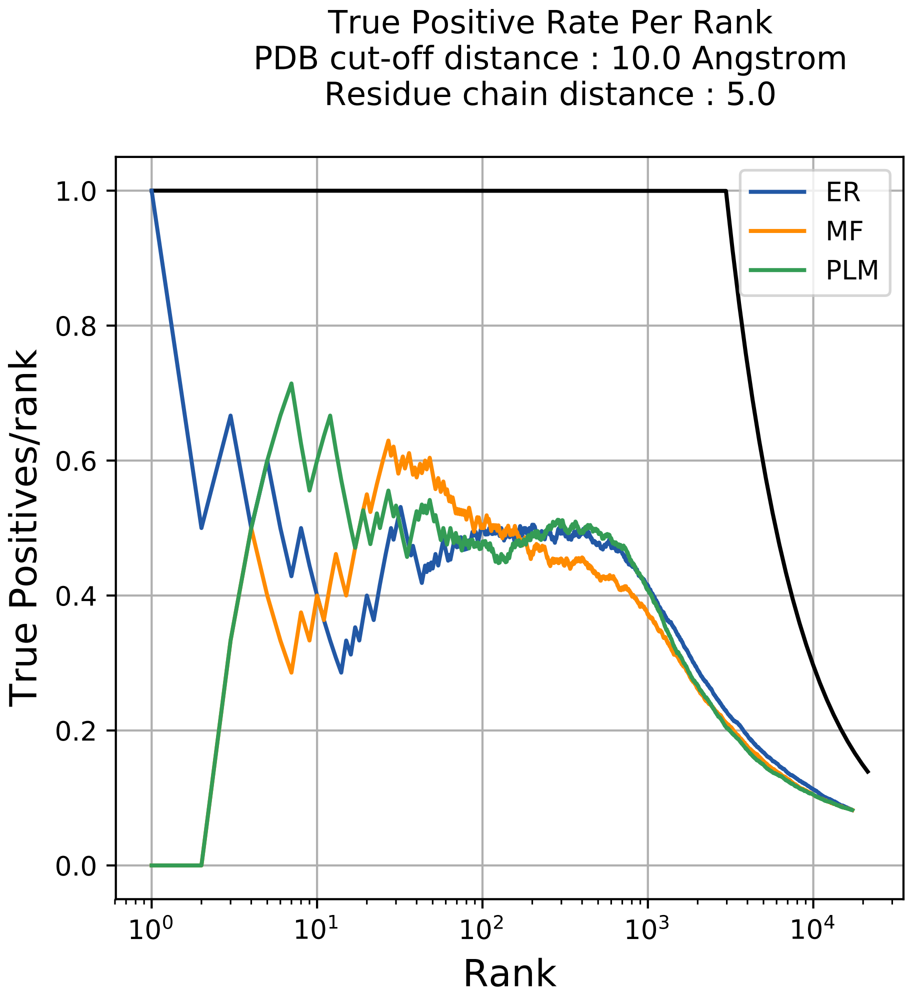

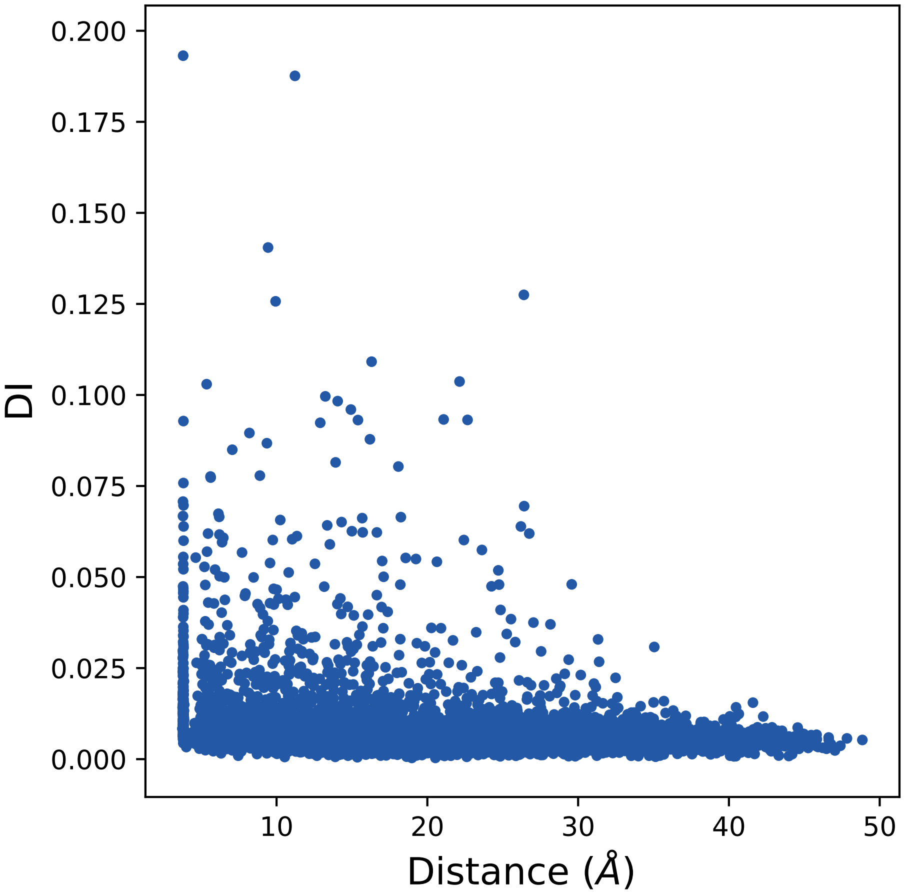

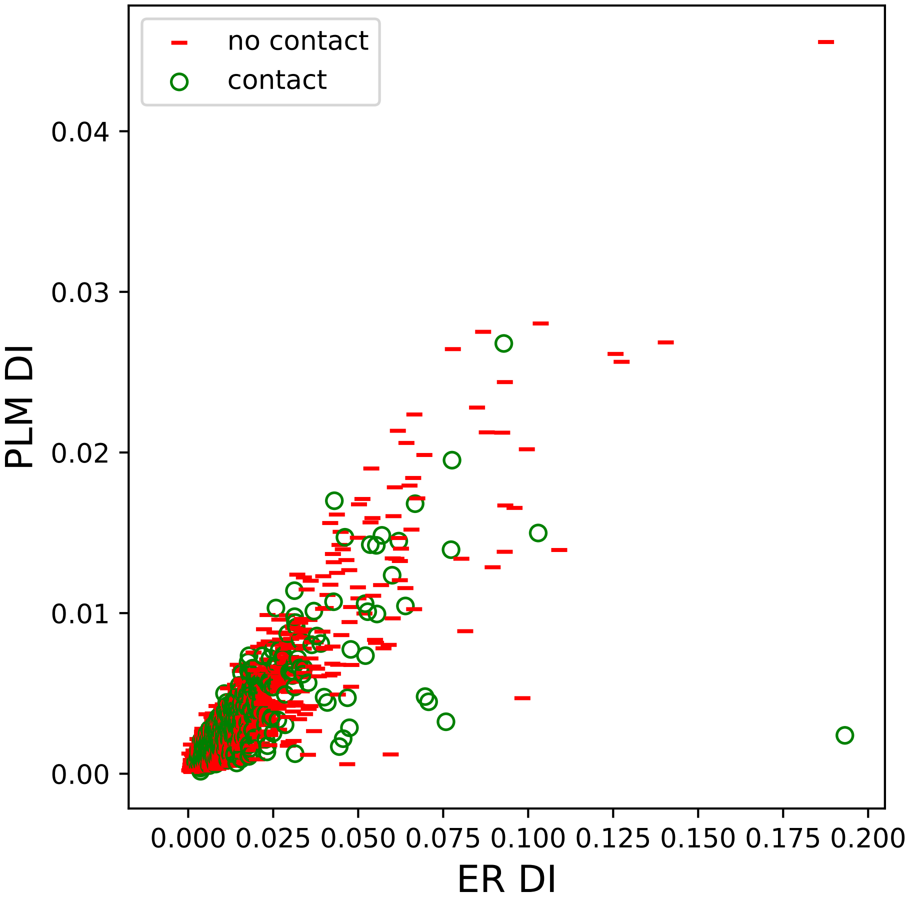

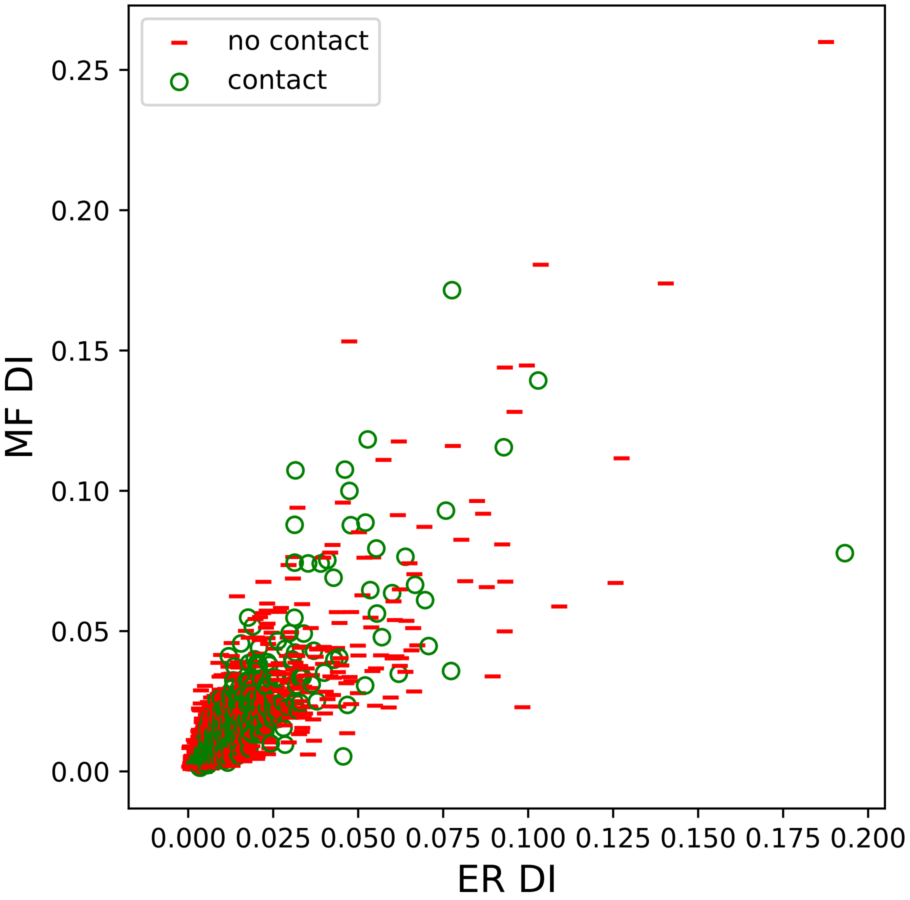

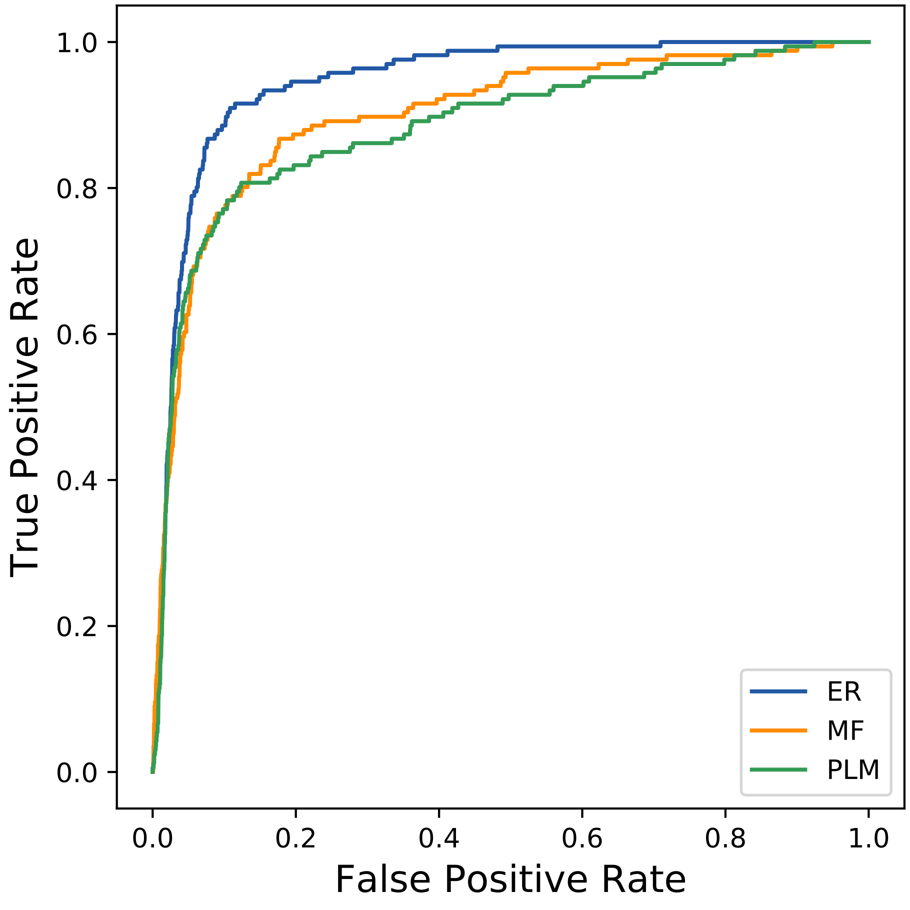

### 14806seq_117col_6b0d_PF00210_plots.pdf

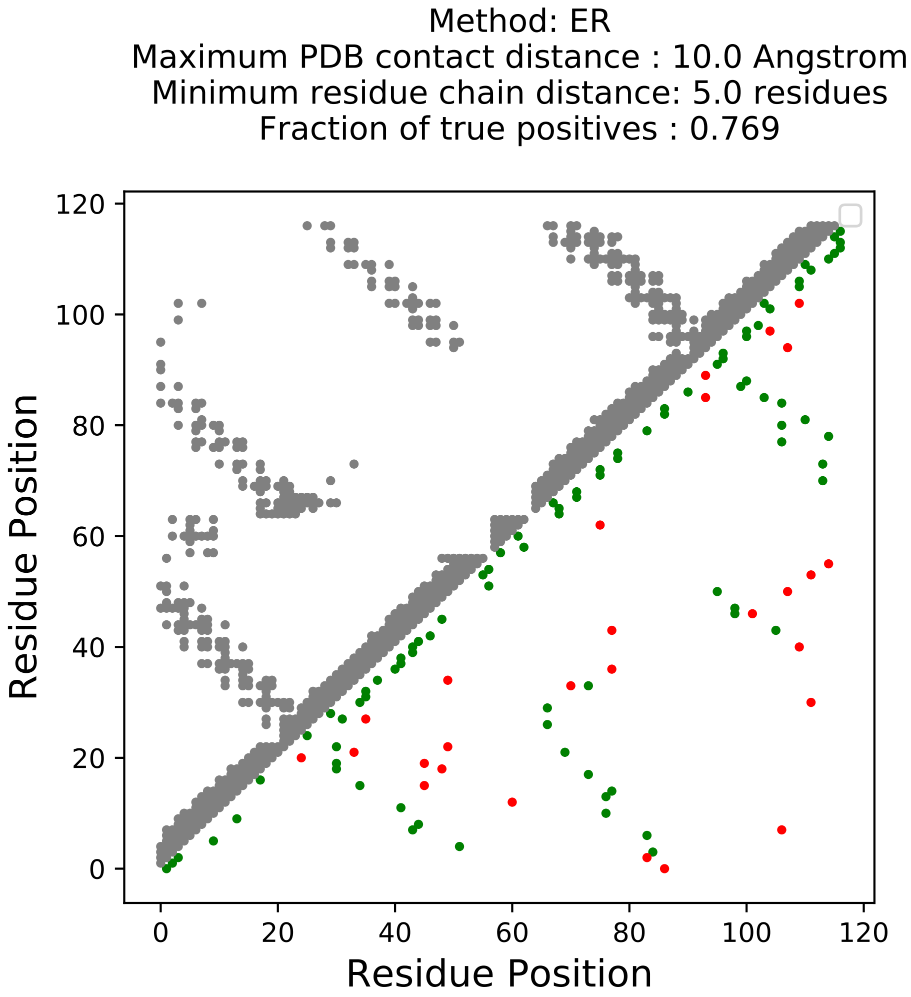

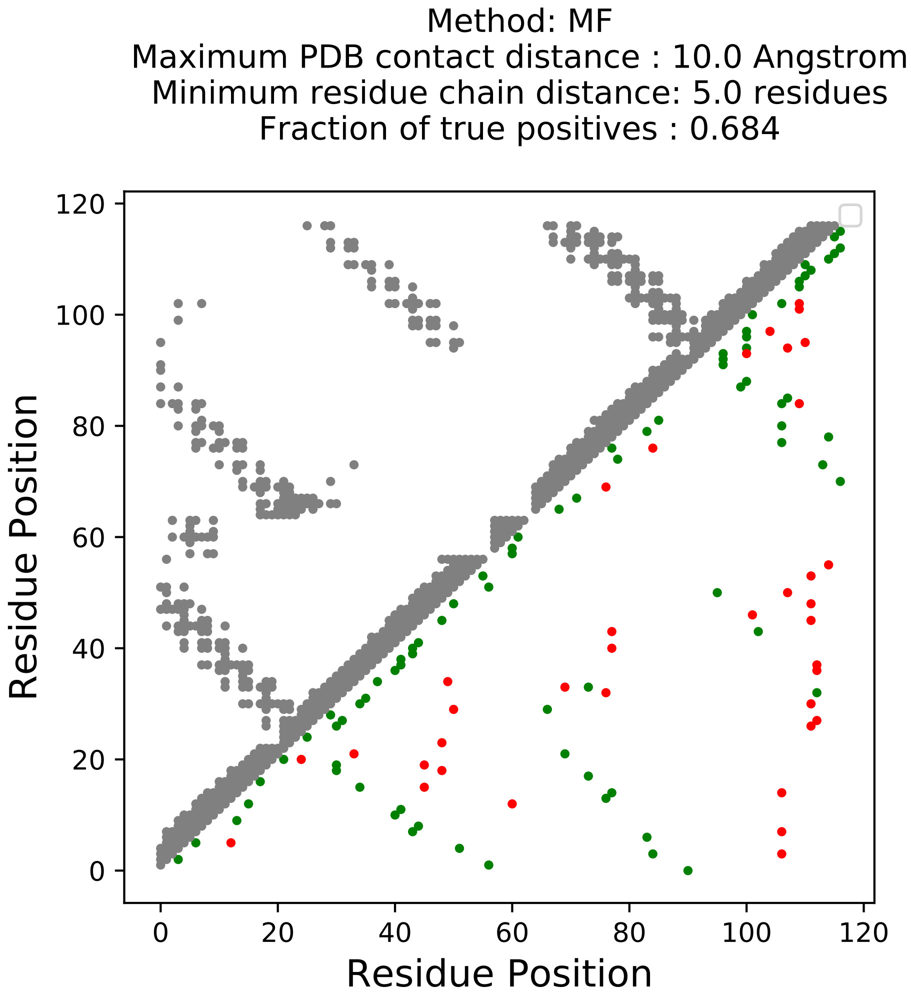

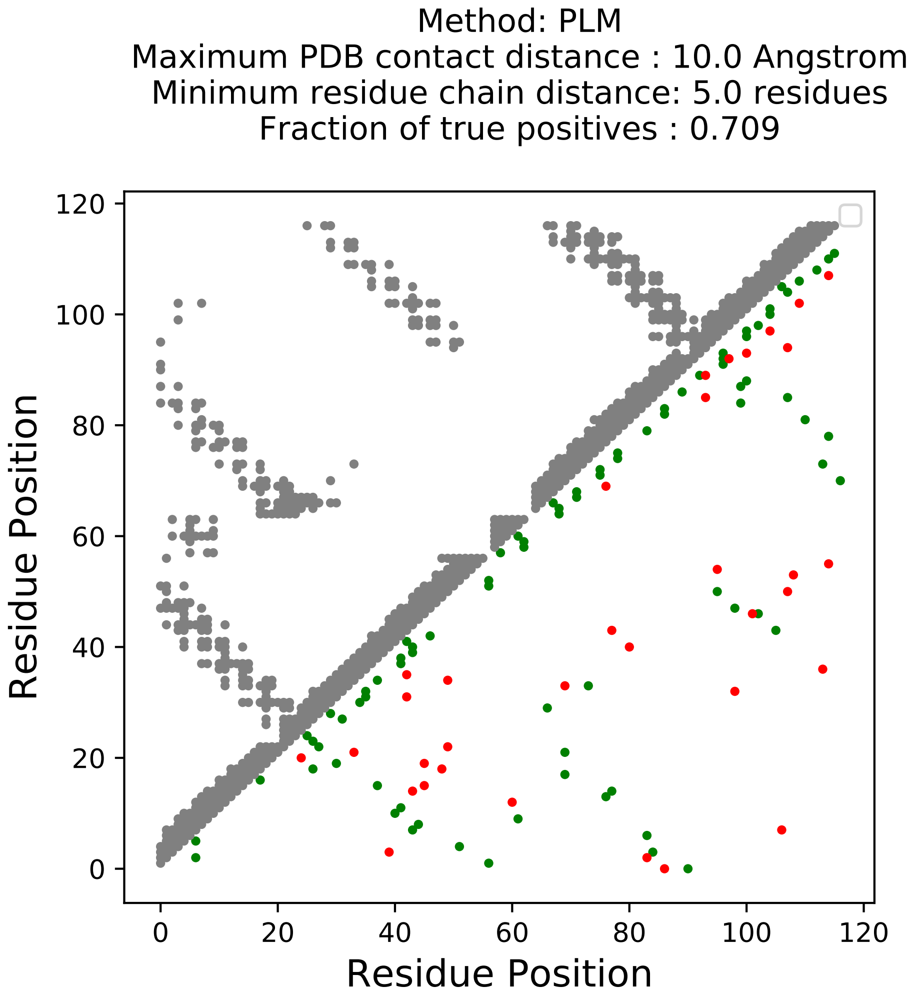

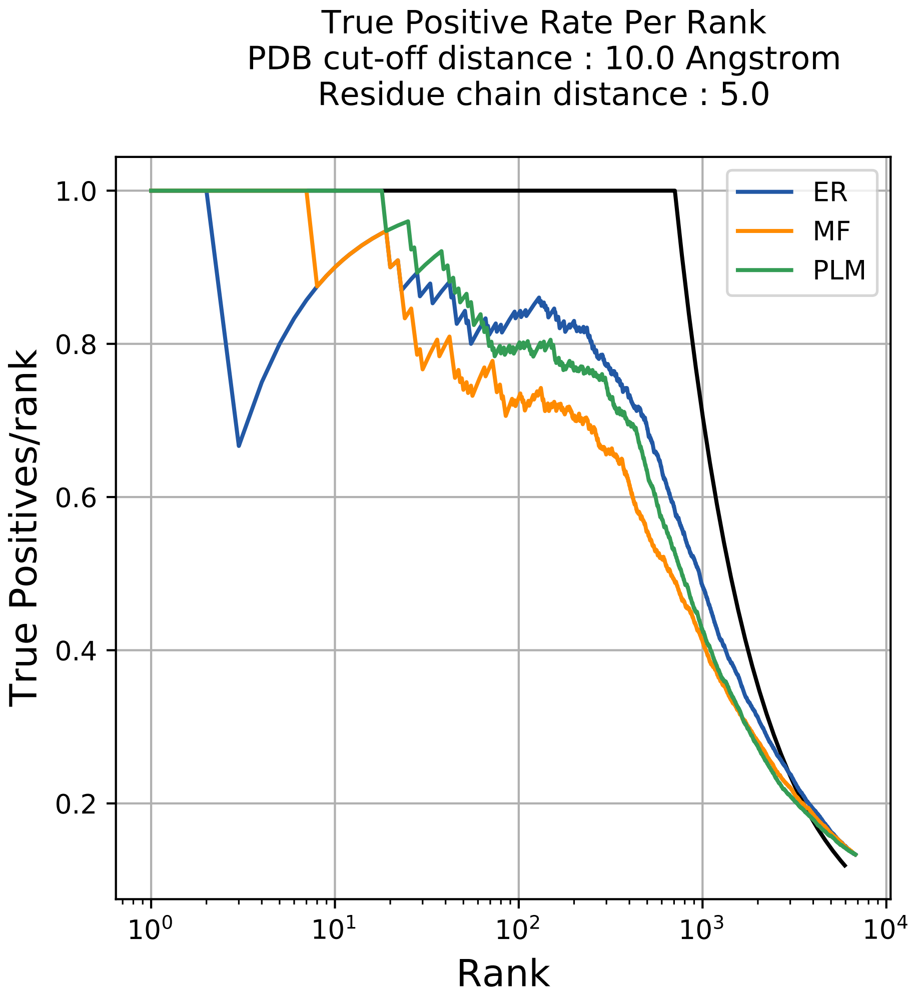

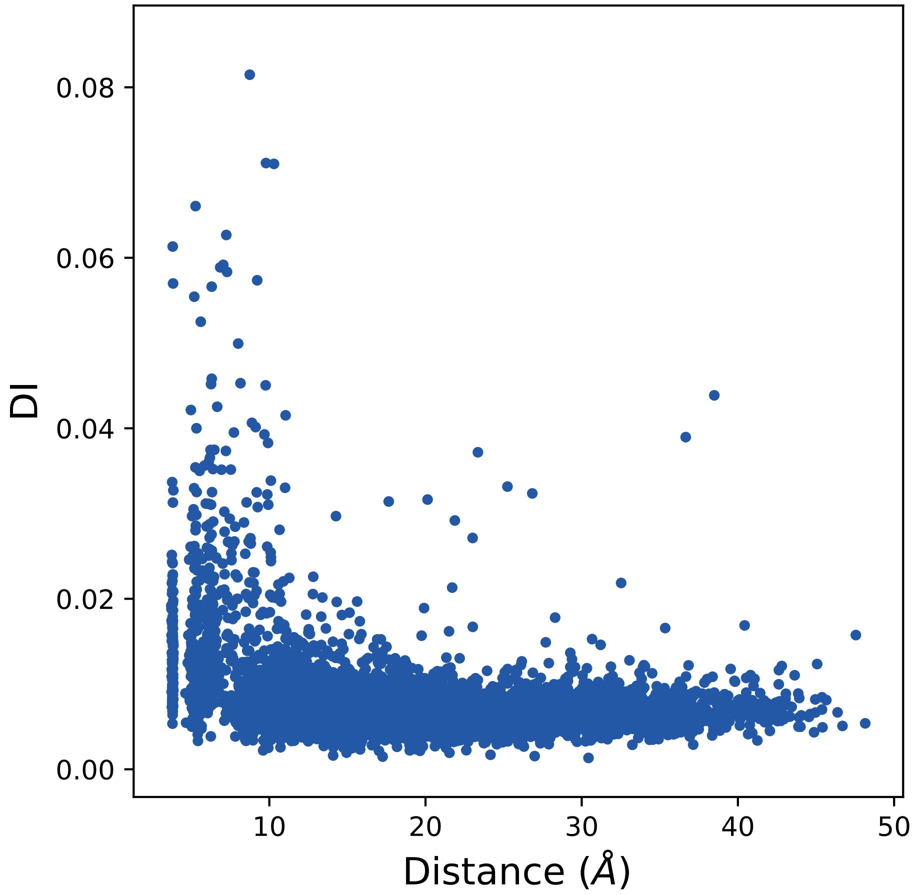

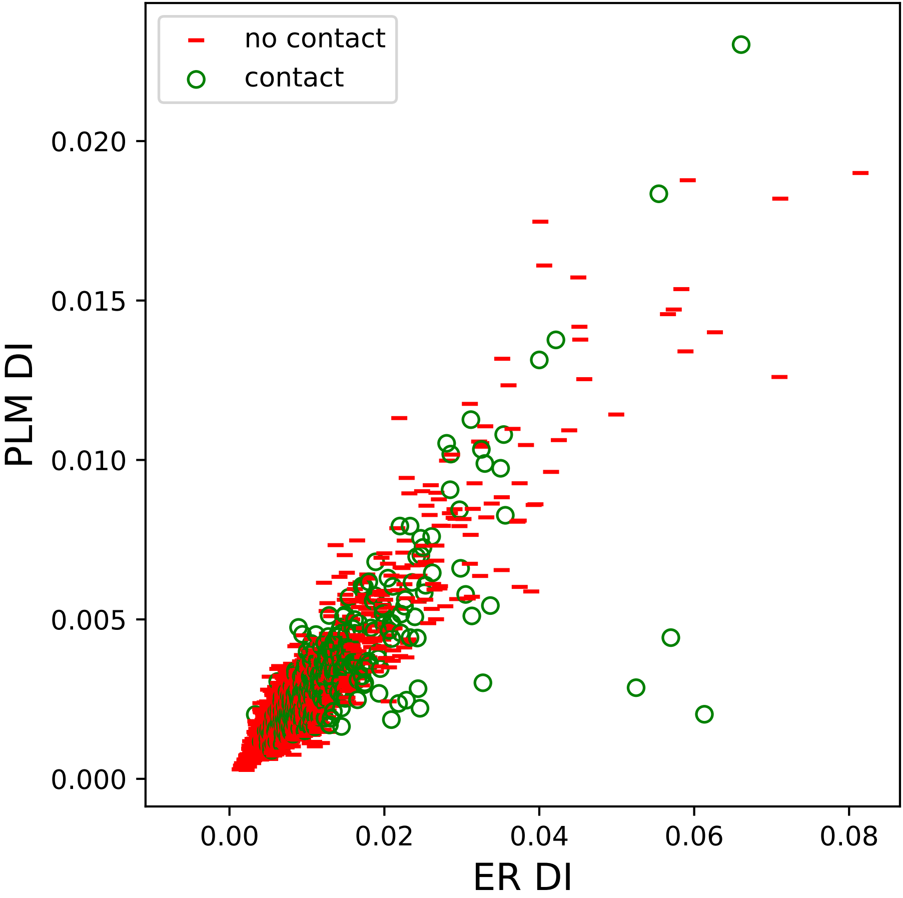

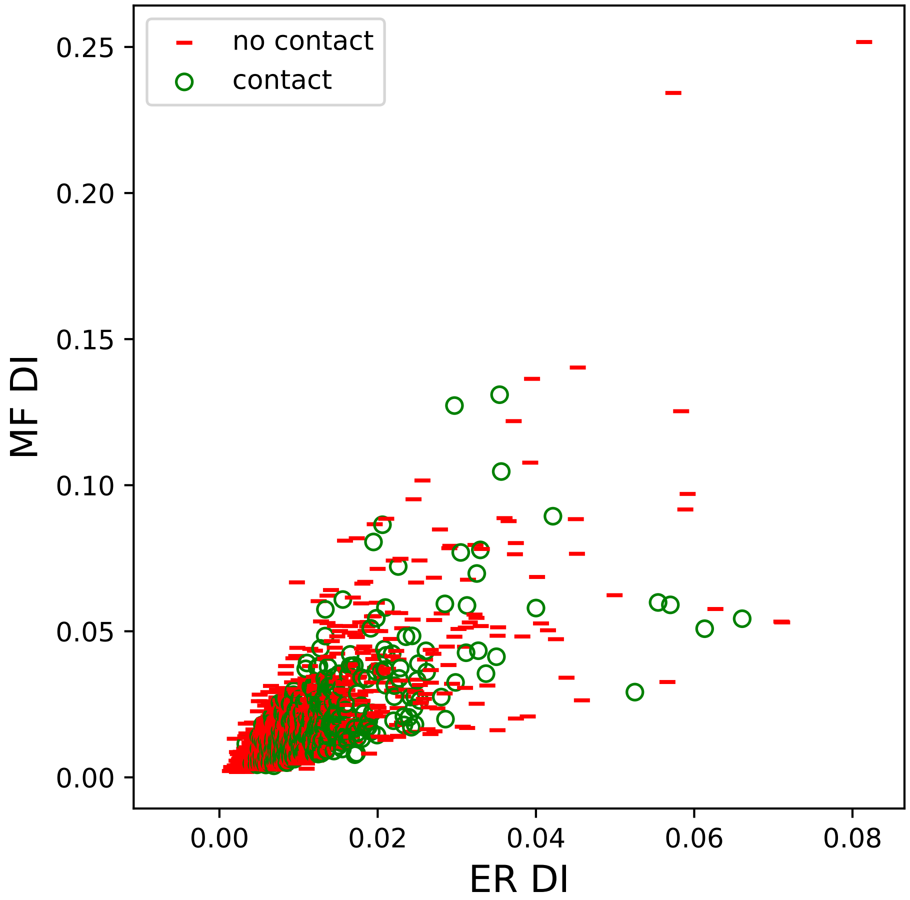

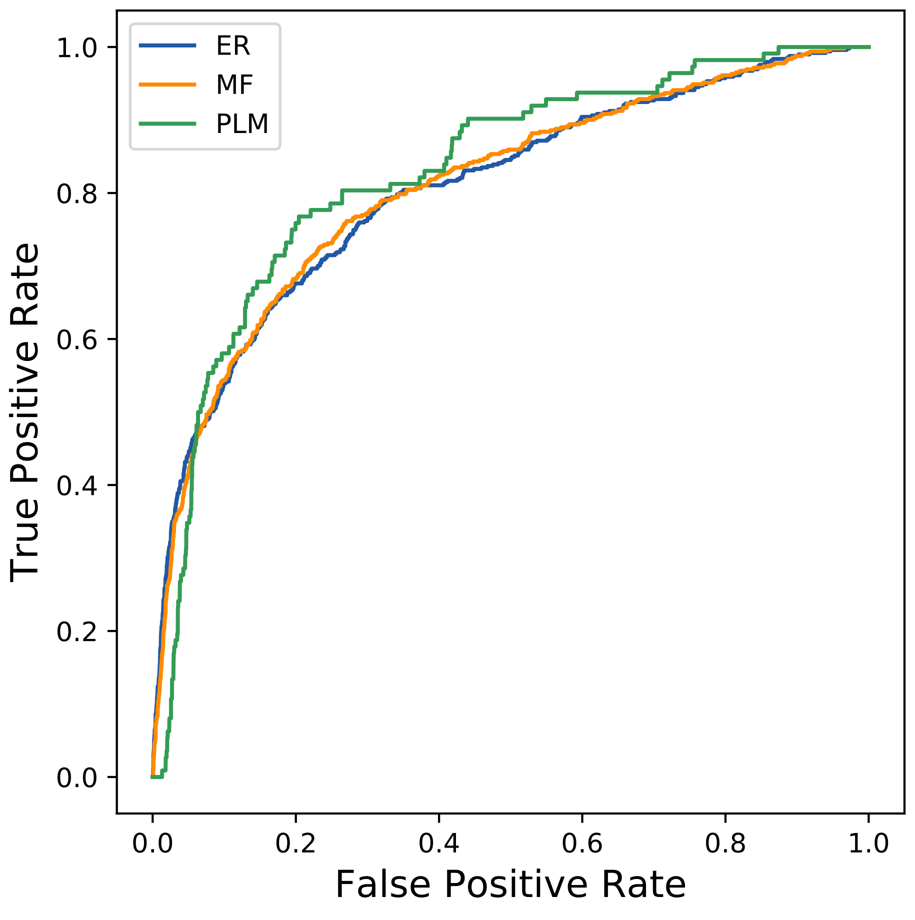

### 15694seq_98col_4dpl_PF01118_plots.pdf

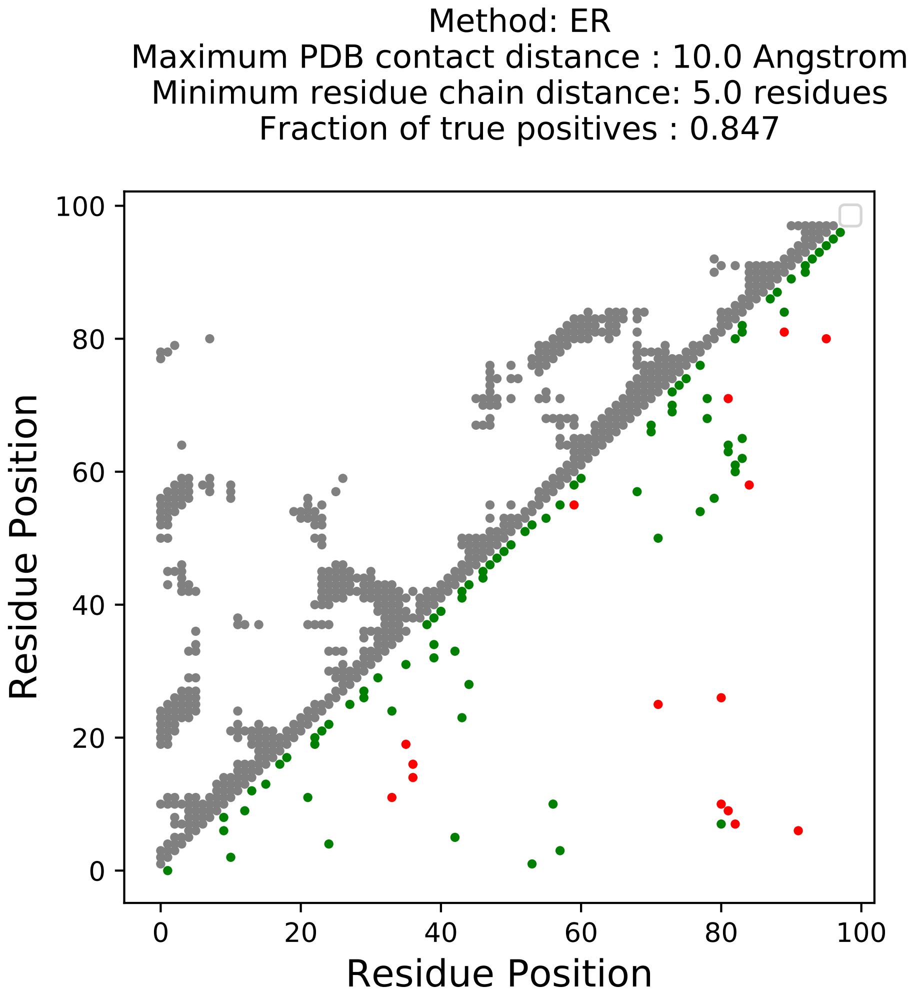

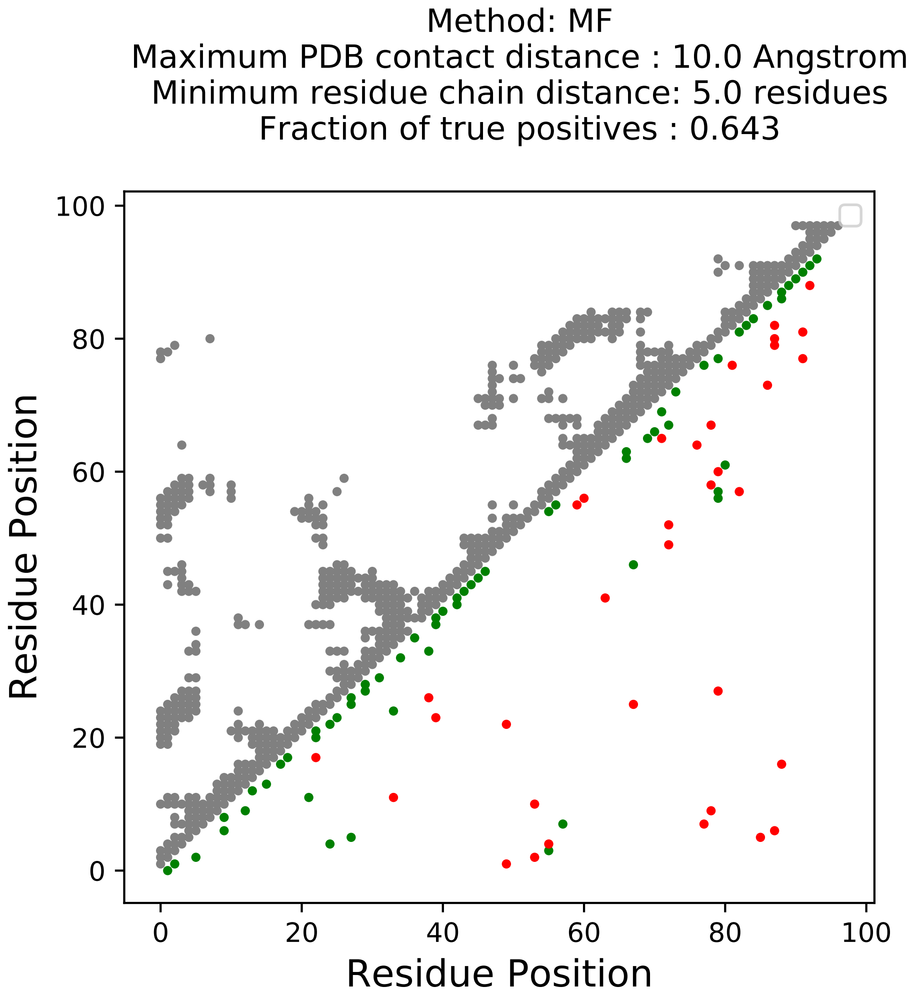

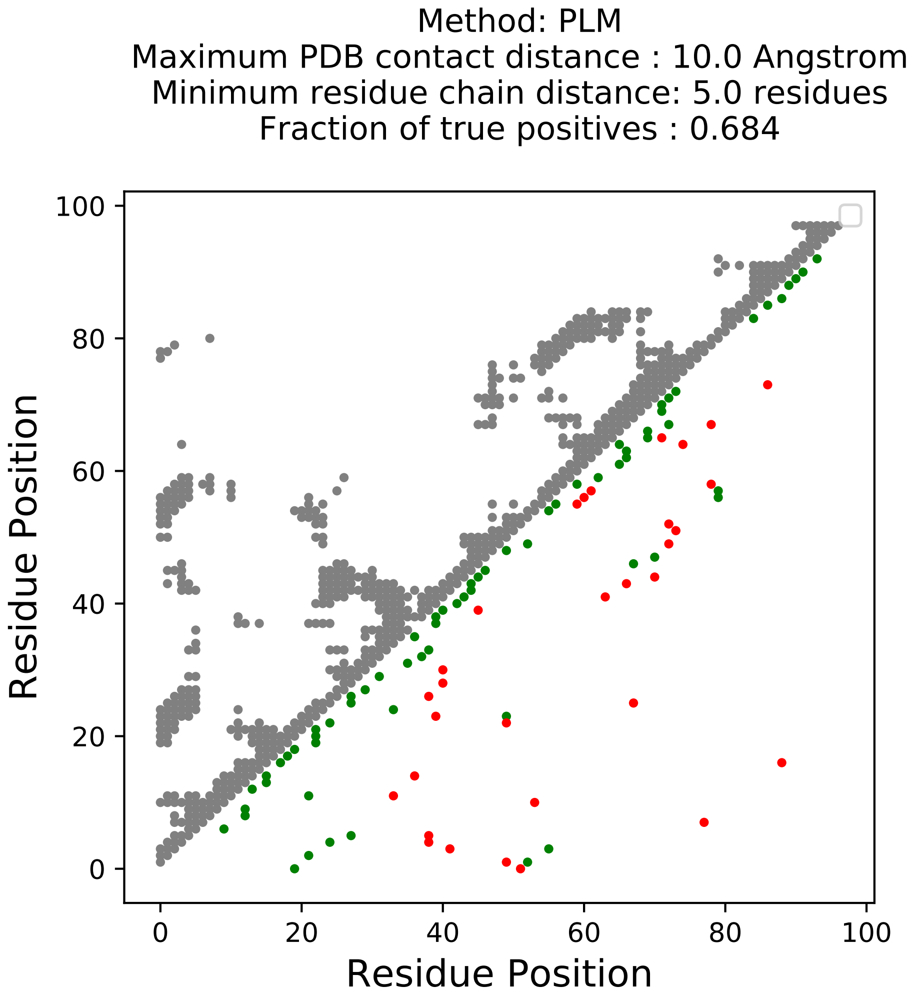

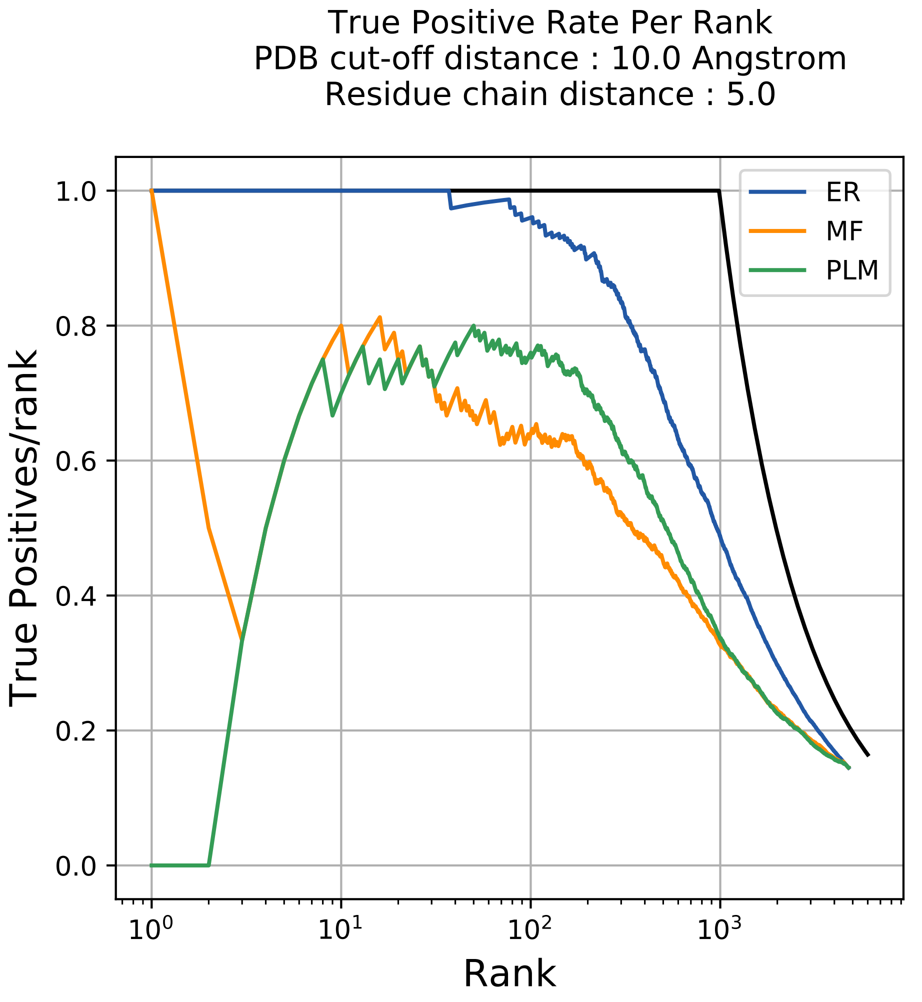

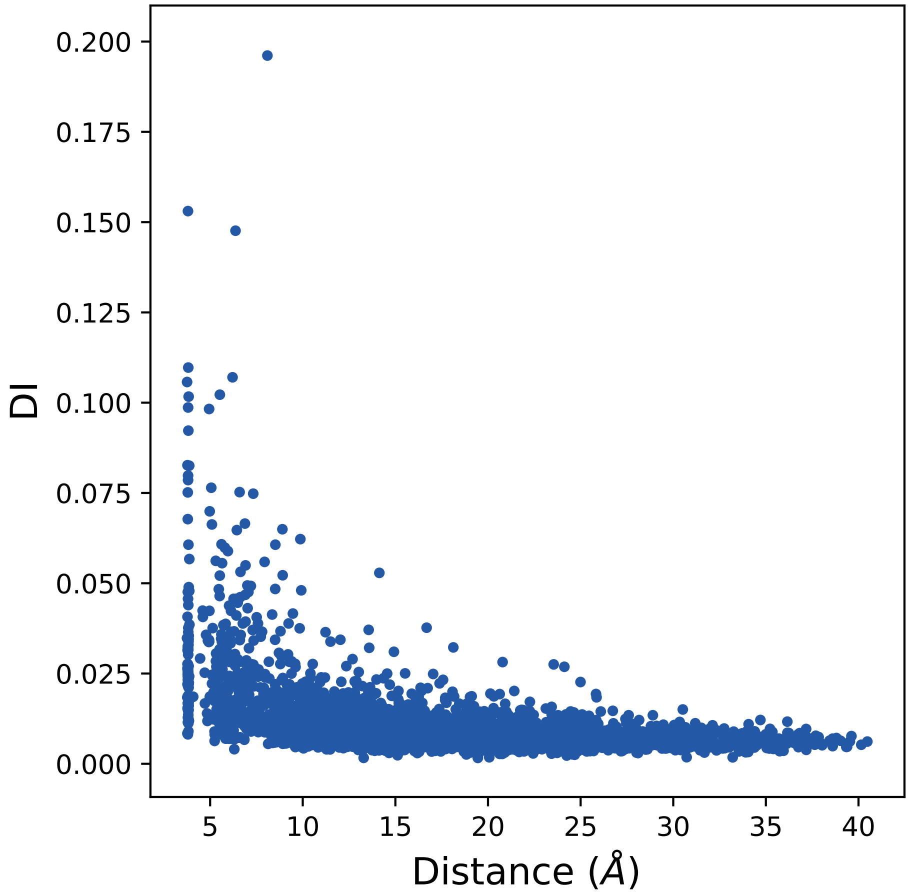

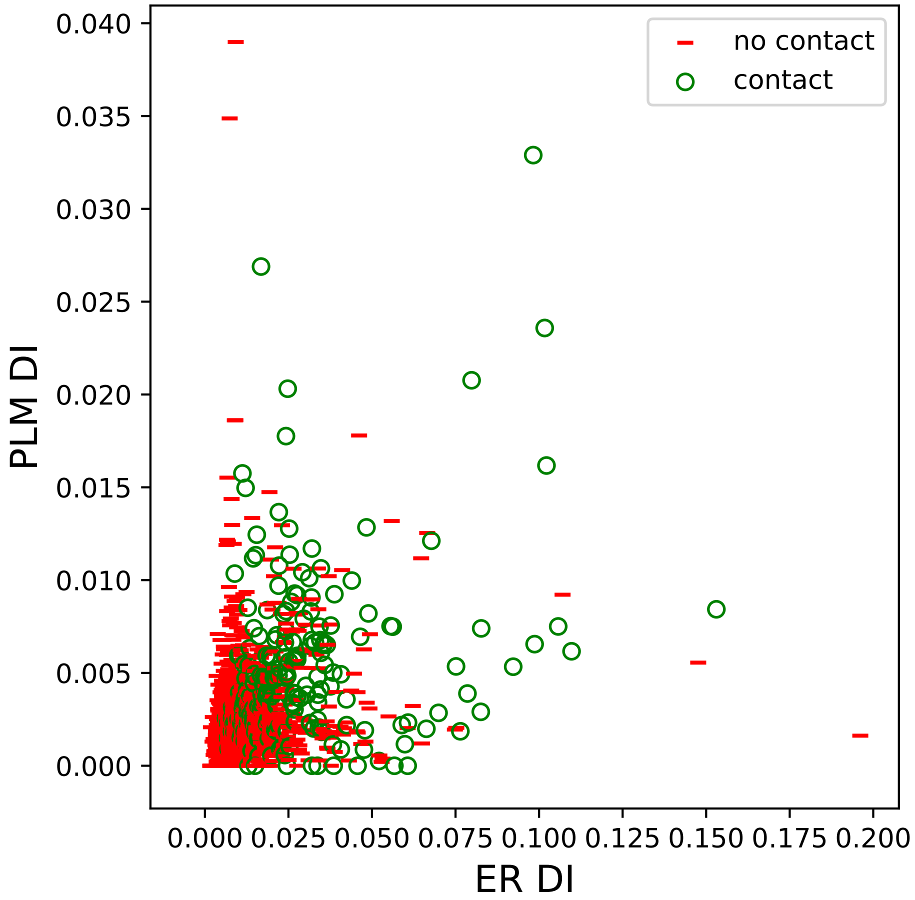

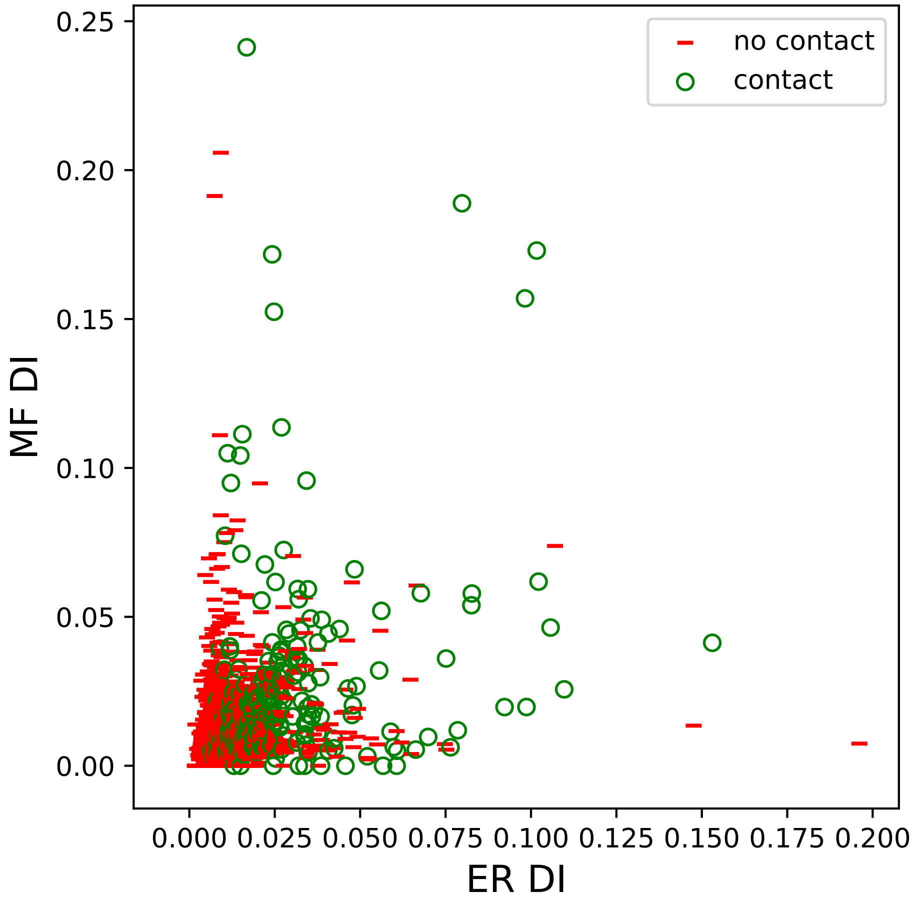

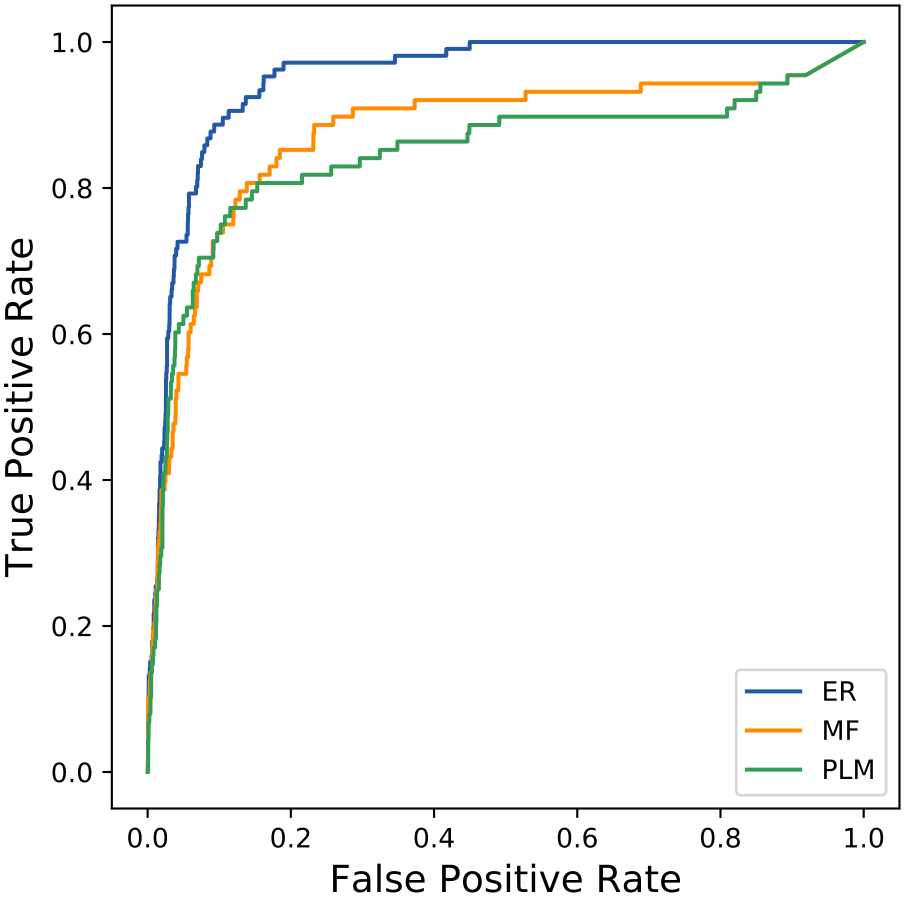

### 18213seq_88col_5ckm_PF00431_plots.pdf

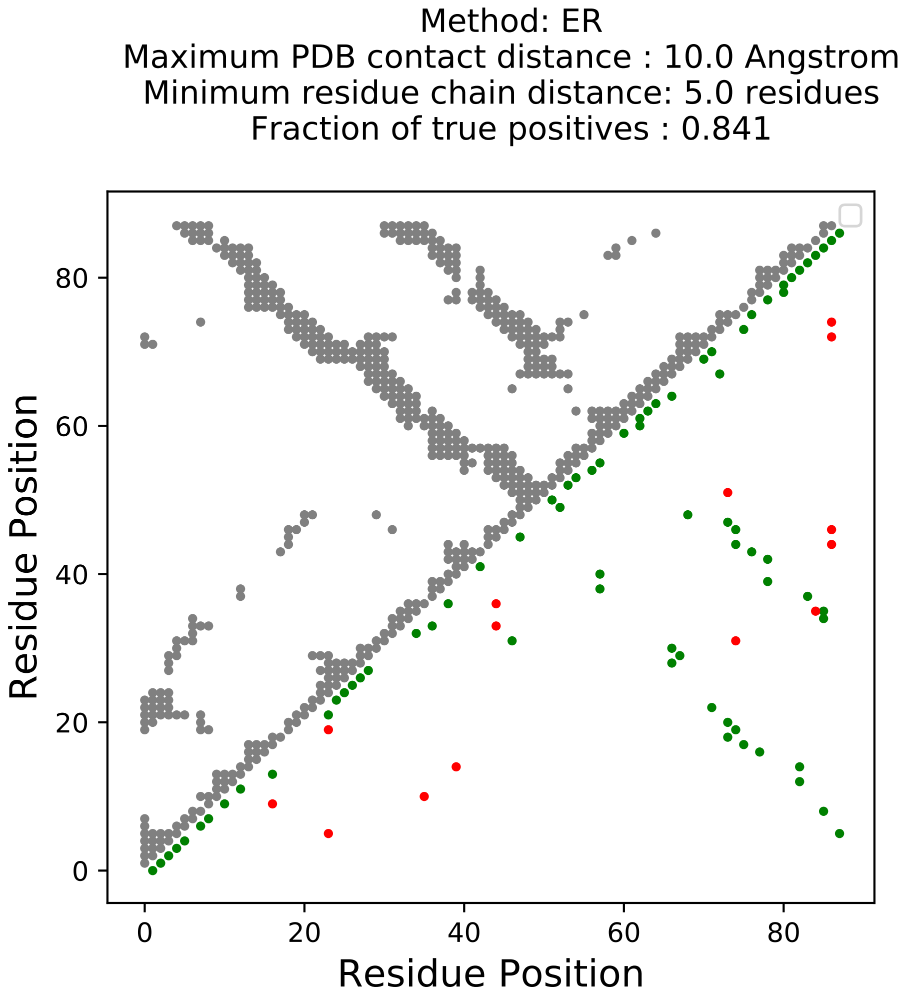

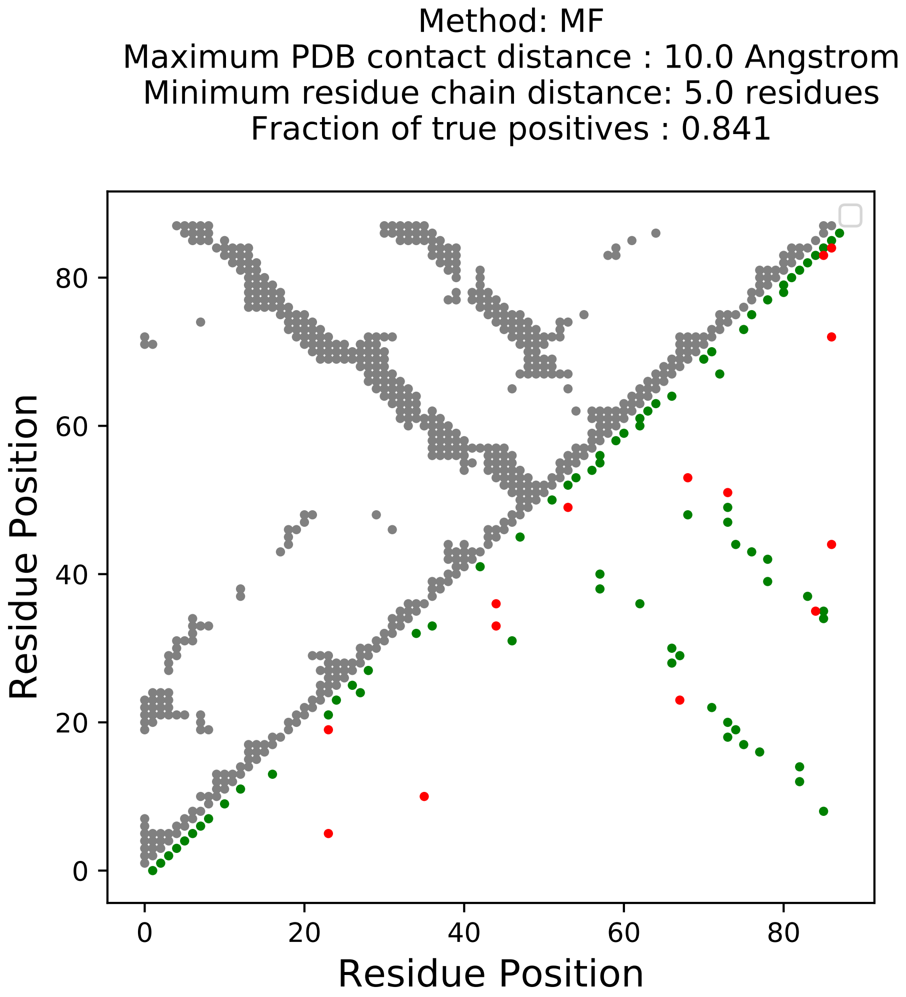

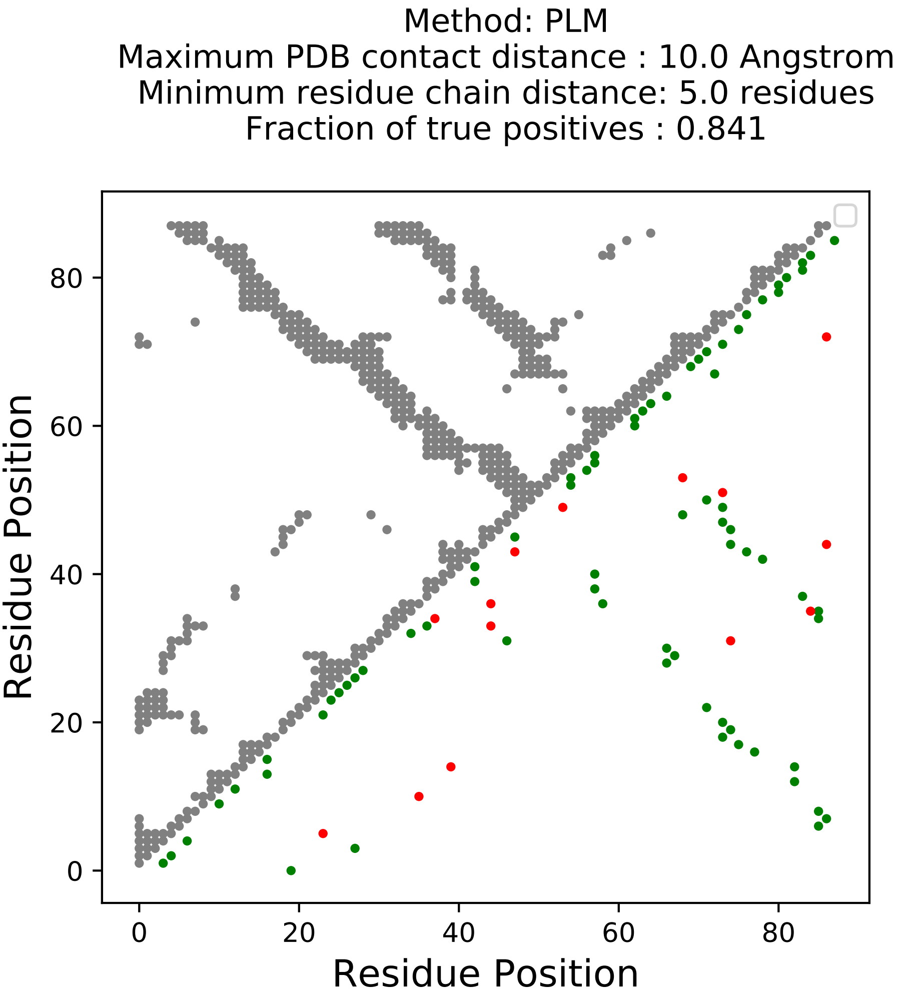

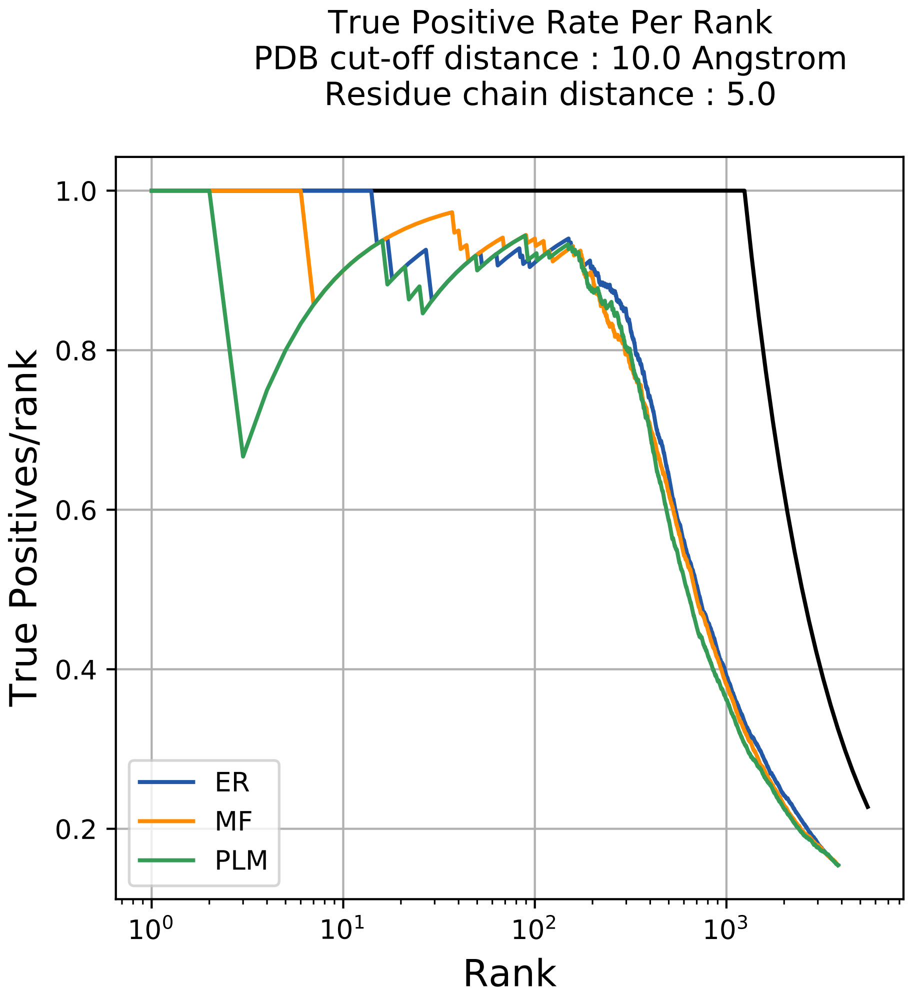

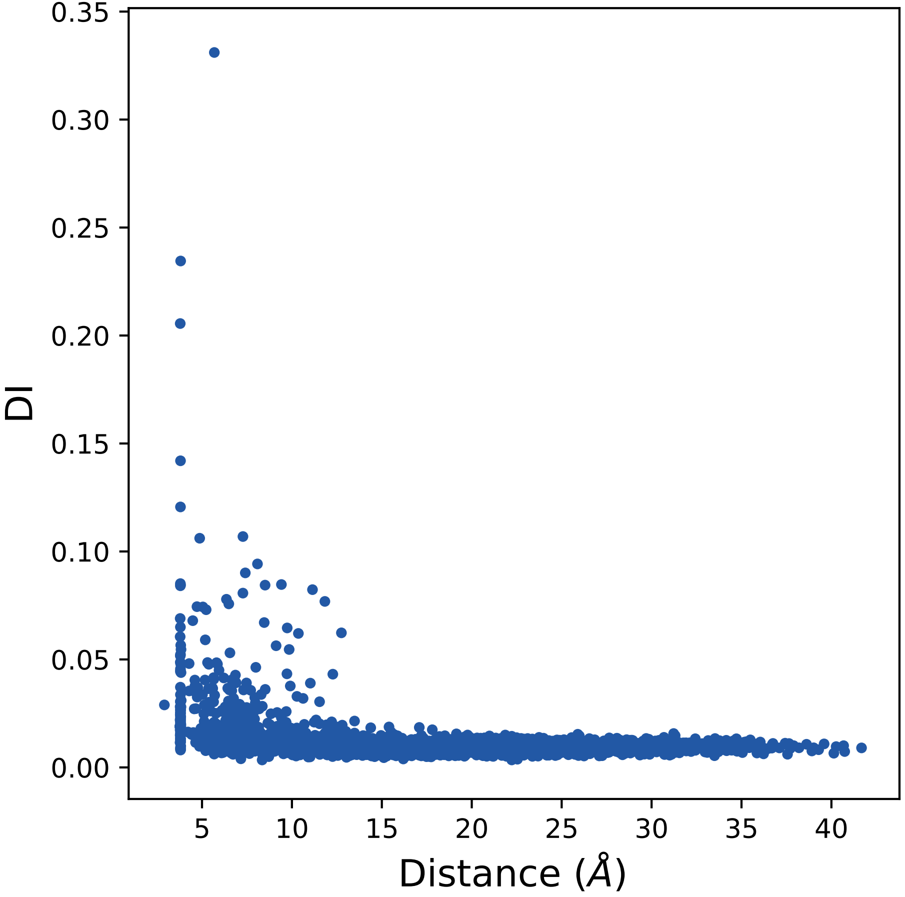

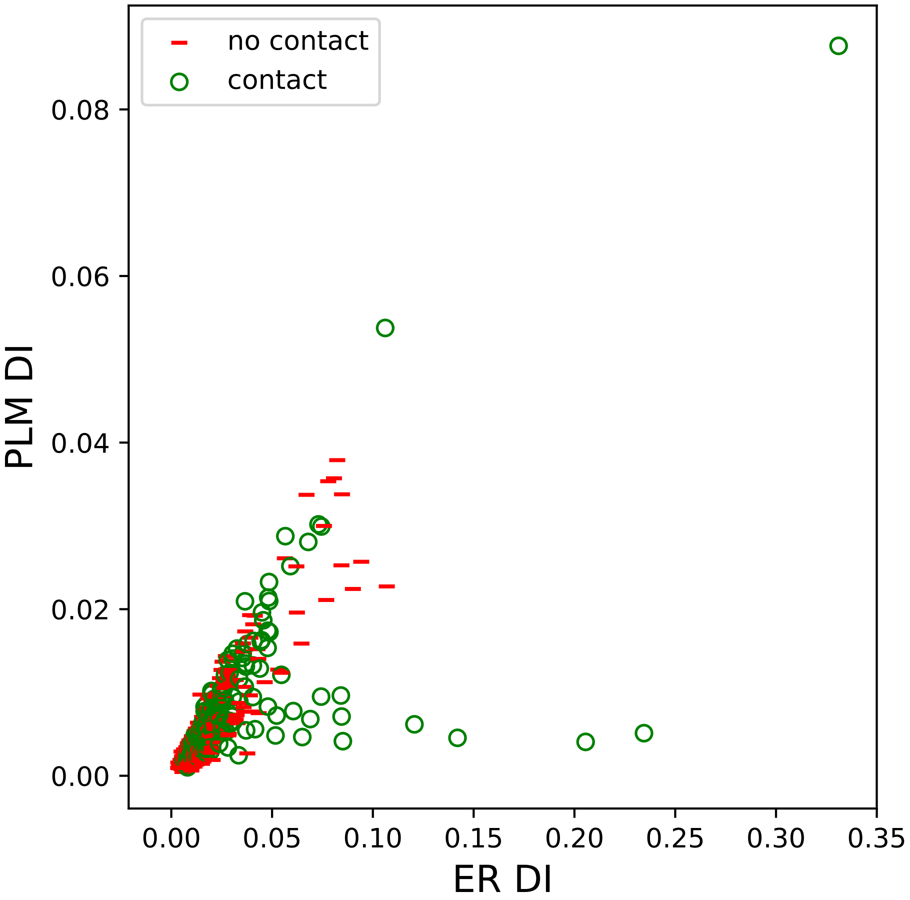

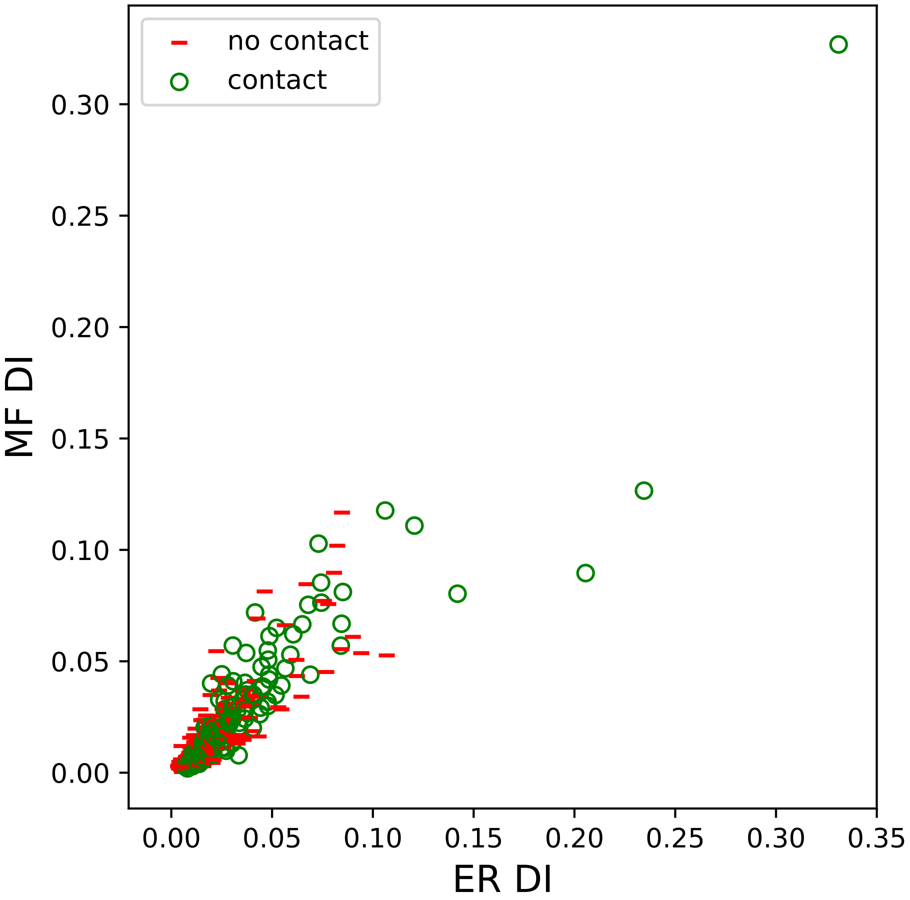

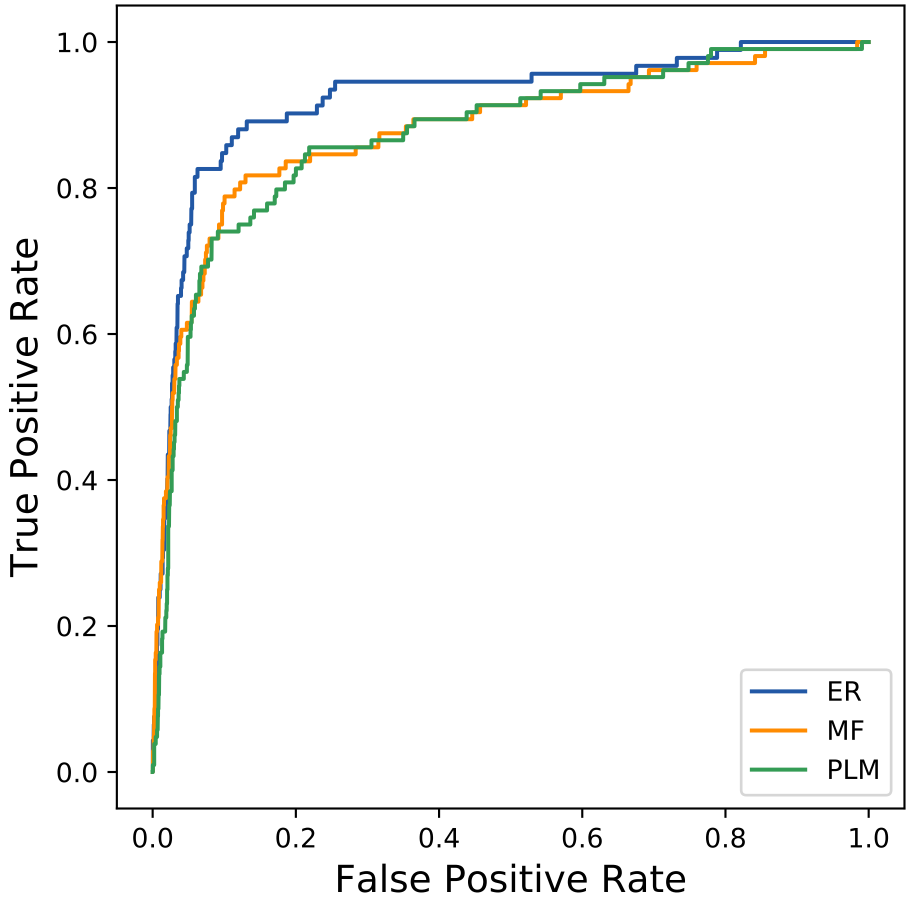

### 18873seq_258col_6cfo_PF00676_plots.pdf

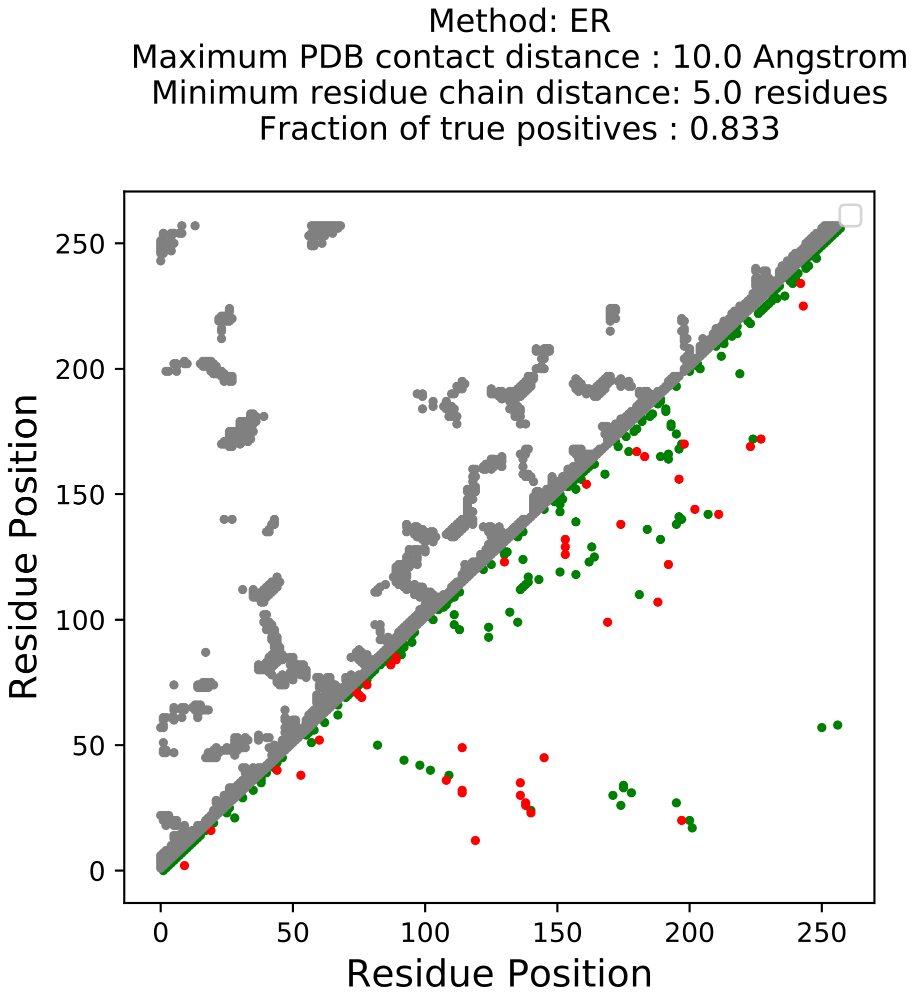

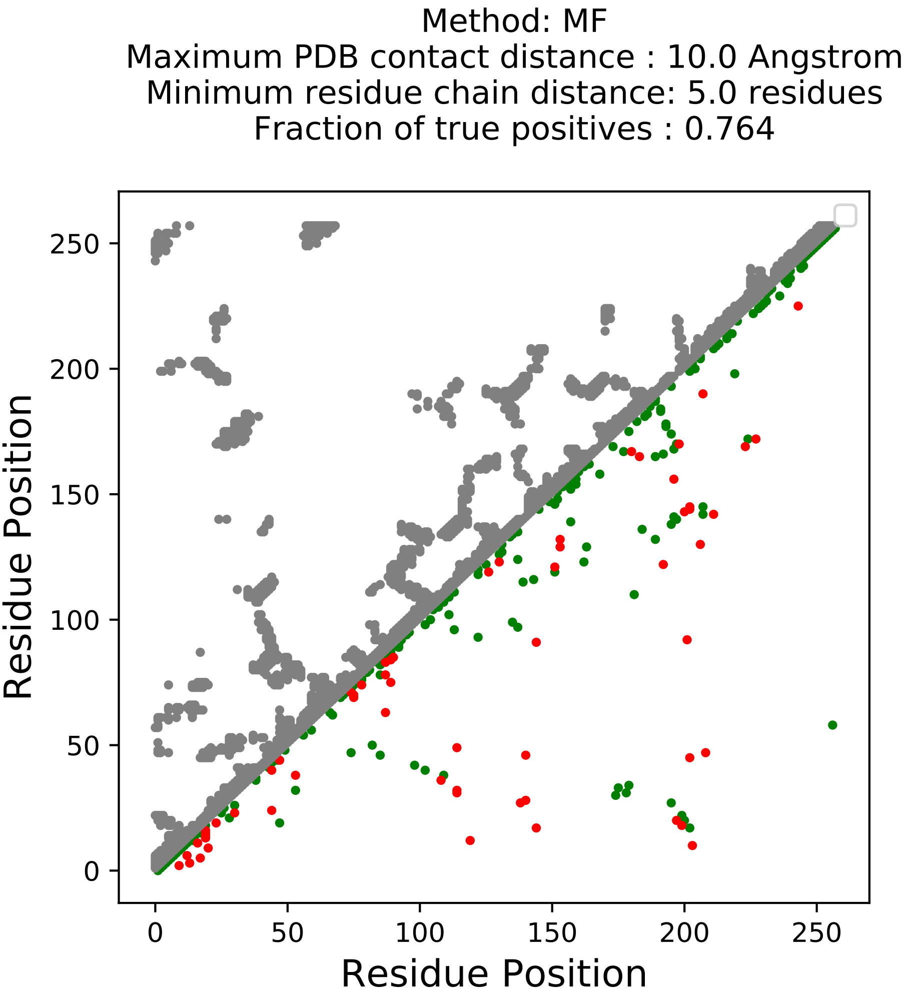

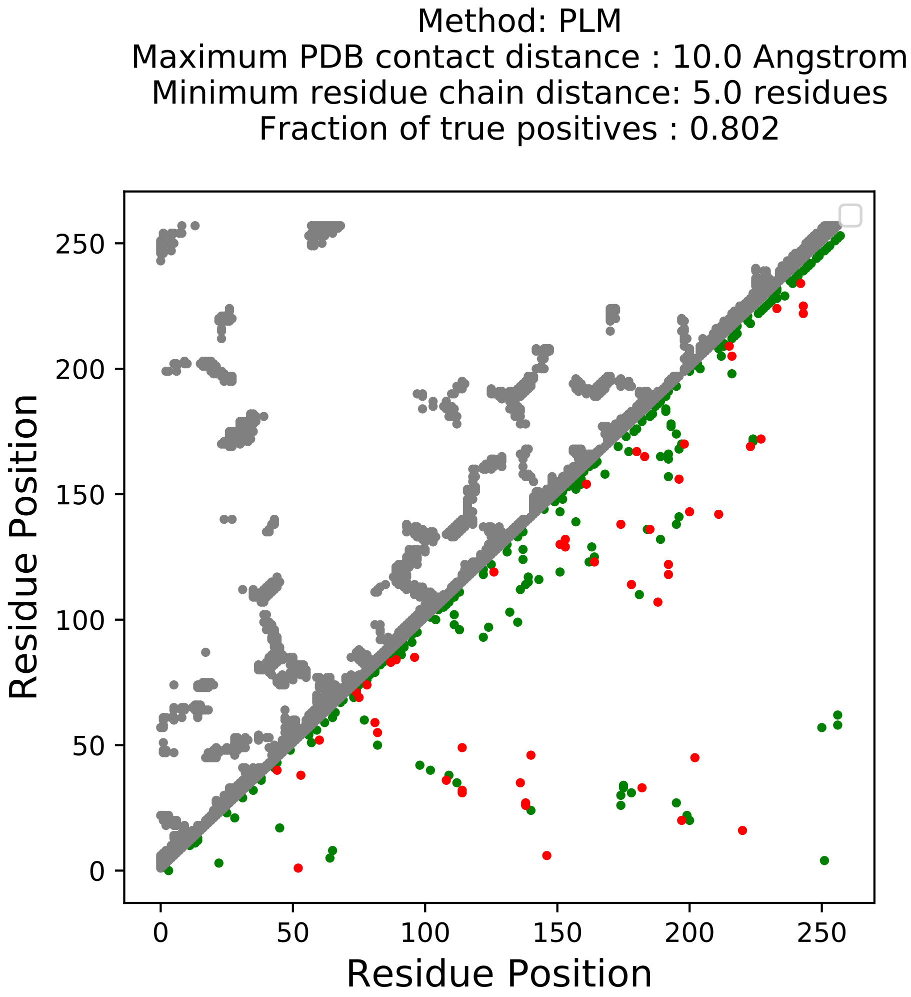

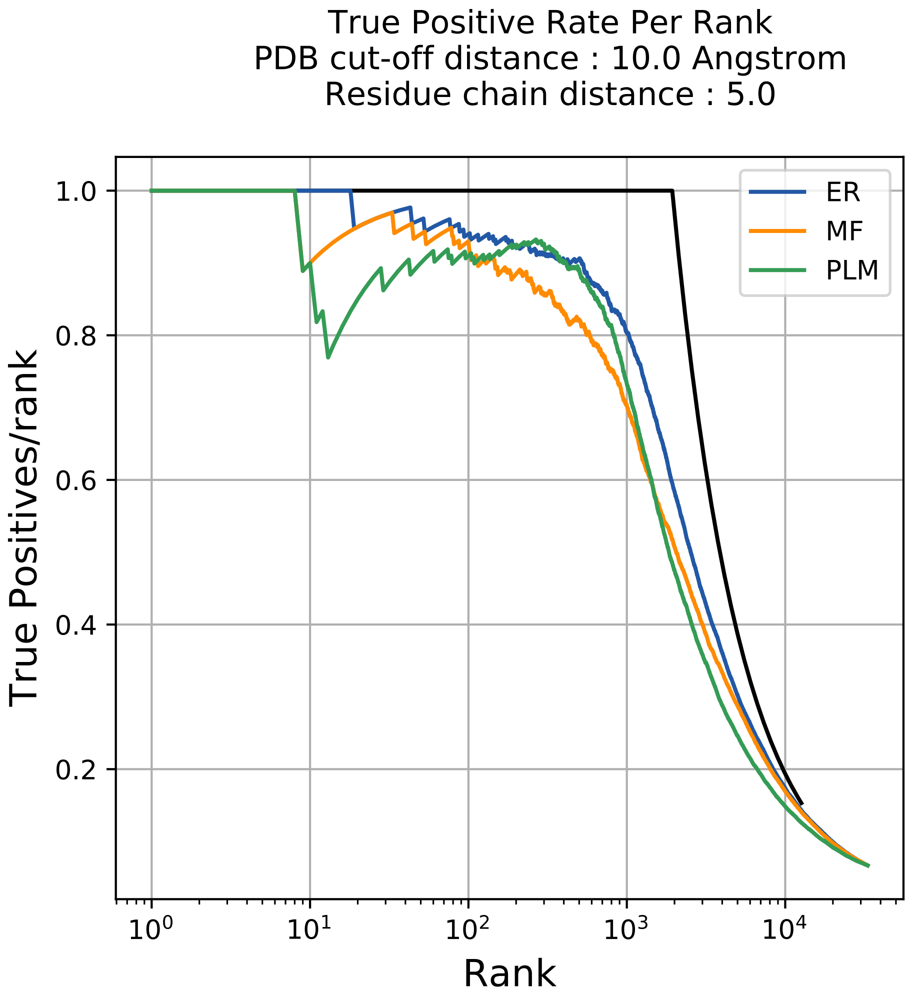

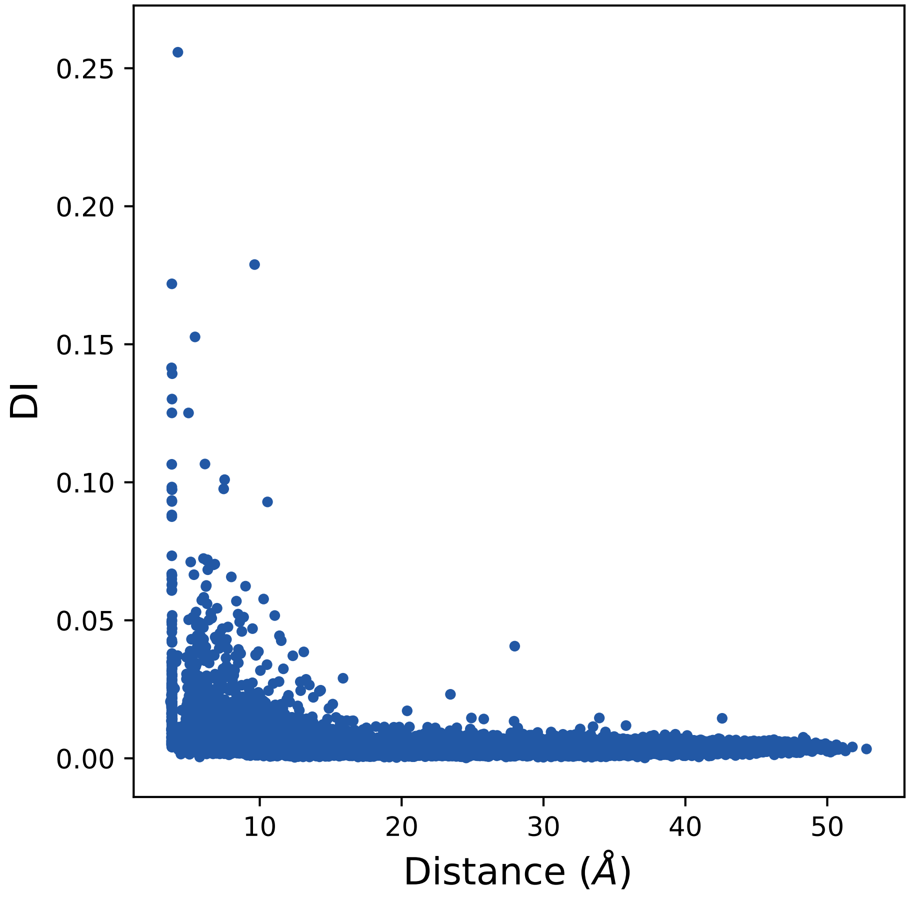

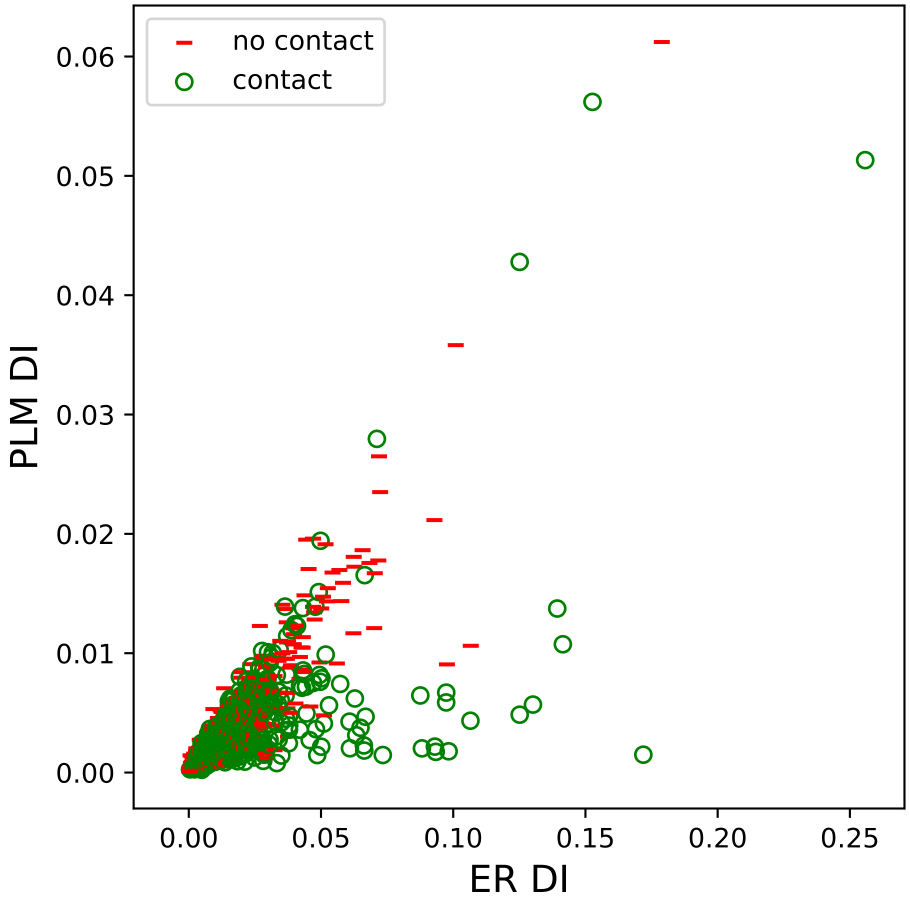

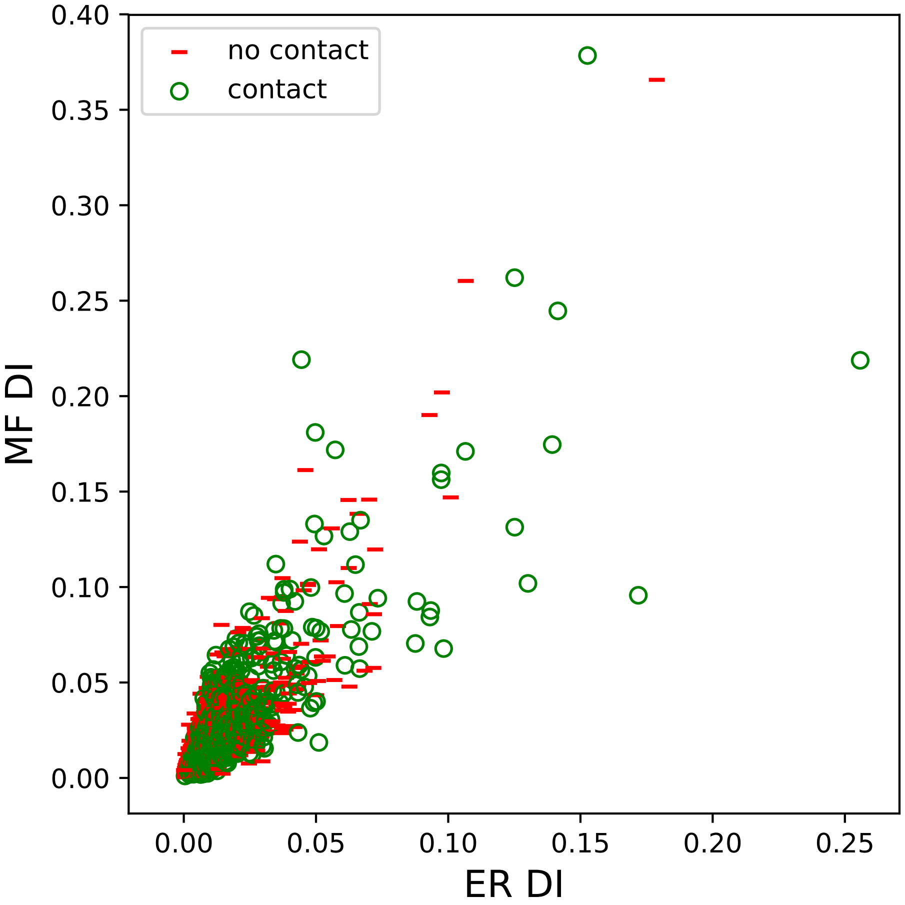

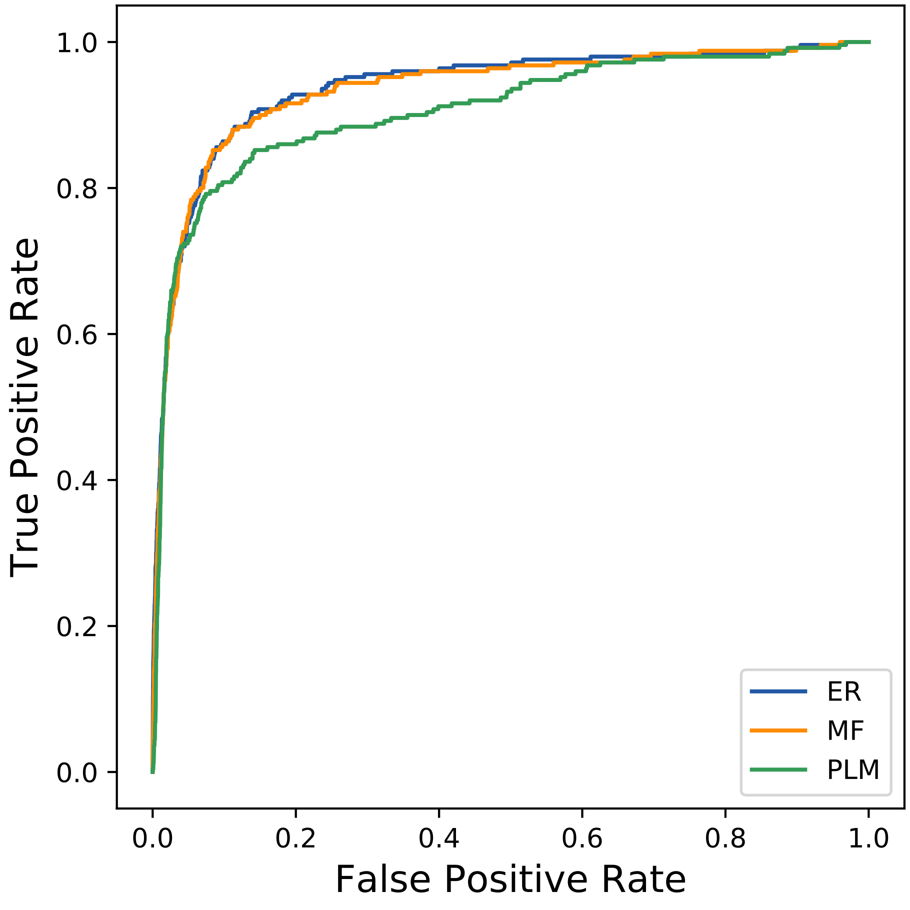

### 18980seq_63col_1bfv_PF07686_plots.pdf

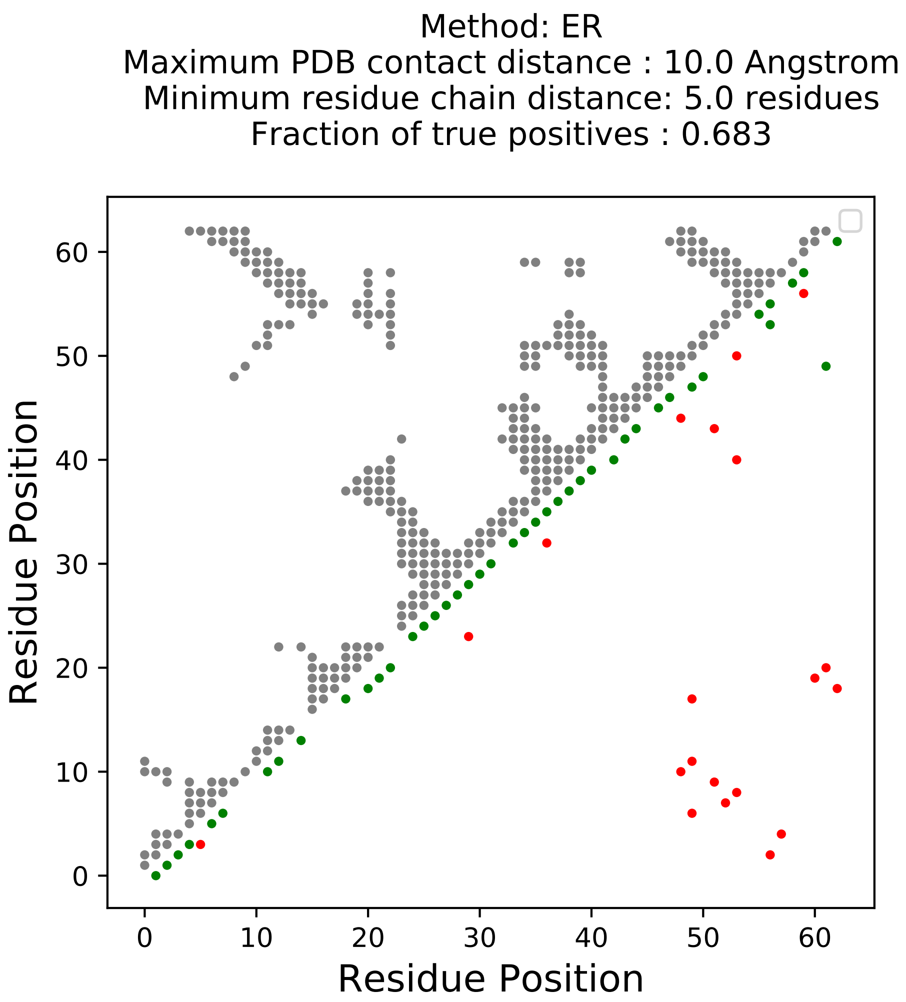

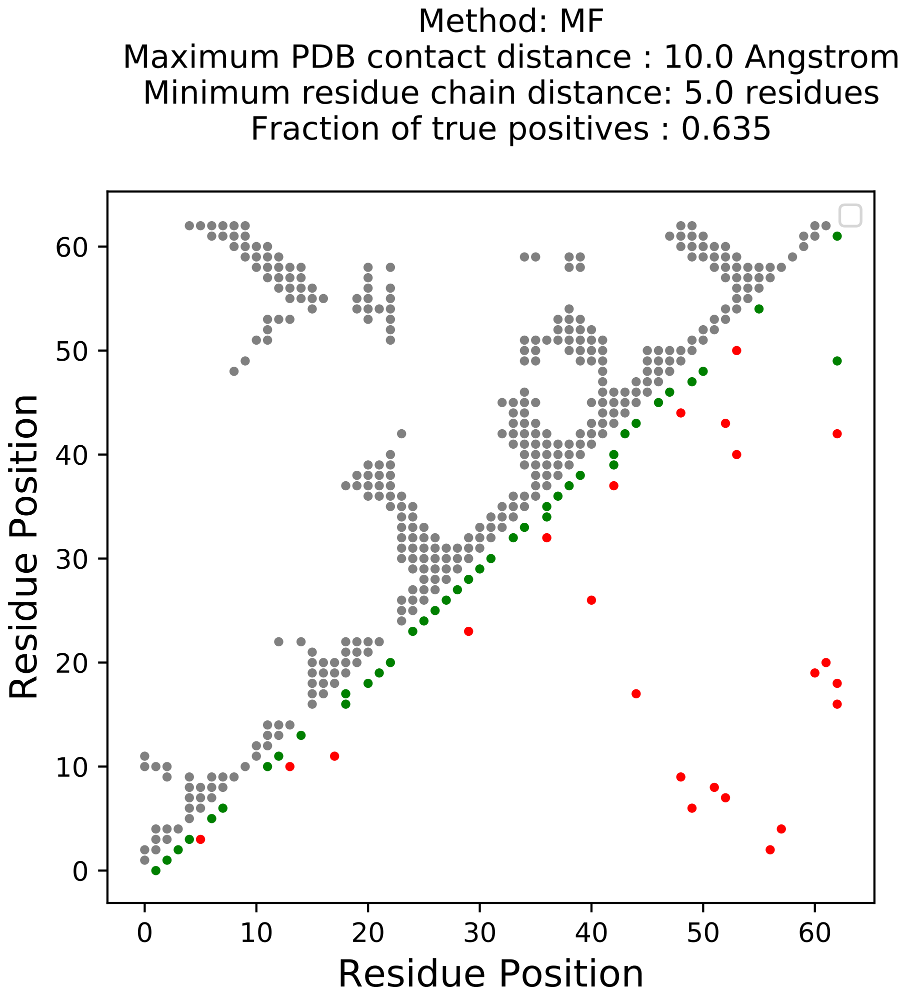

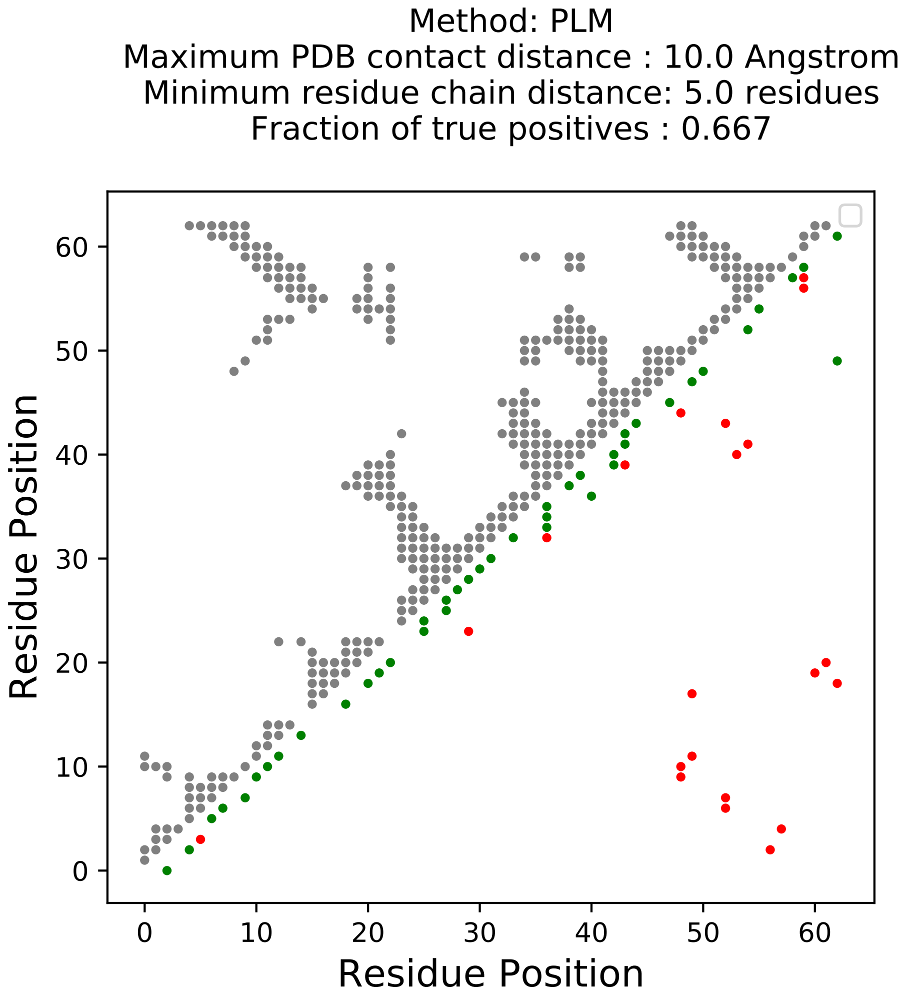

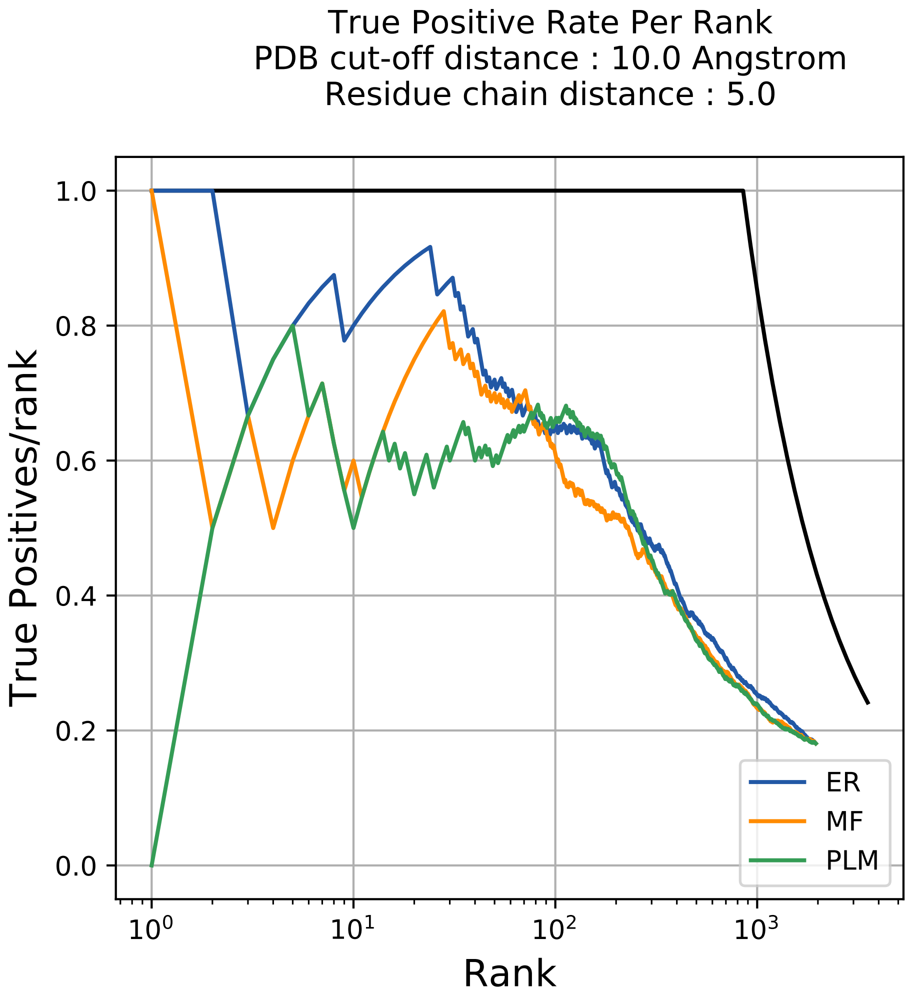

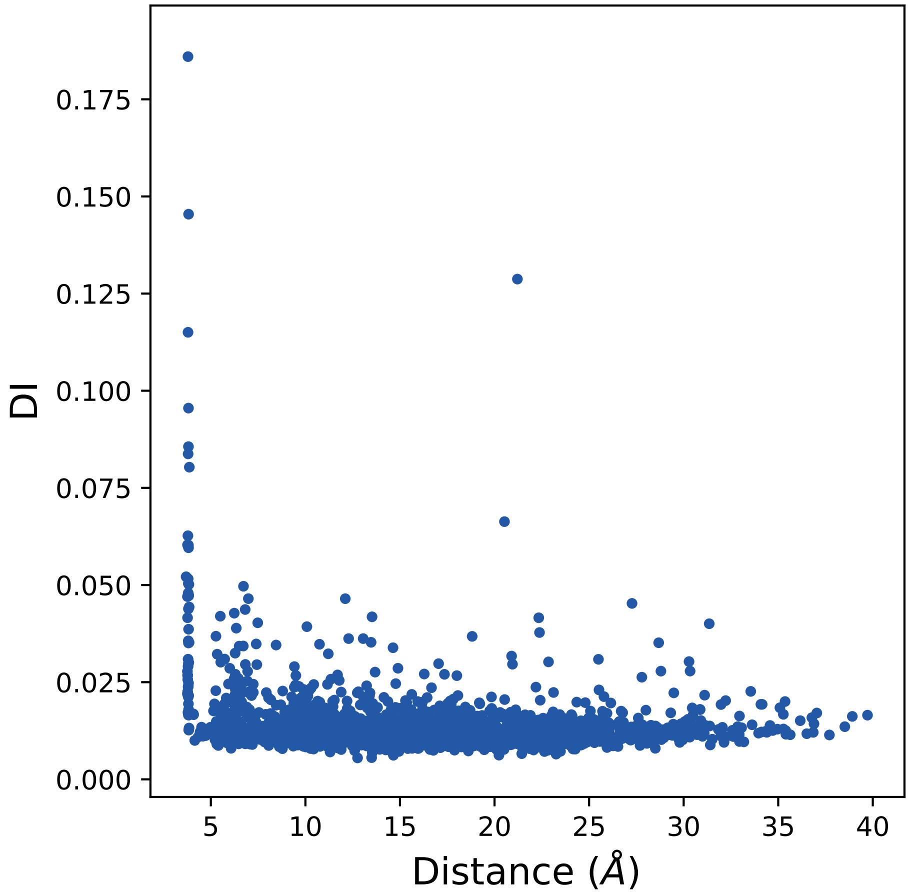

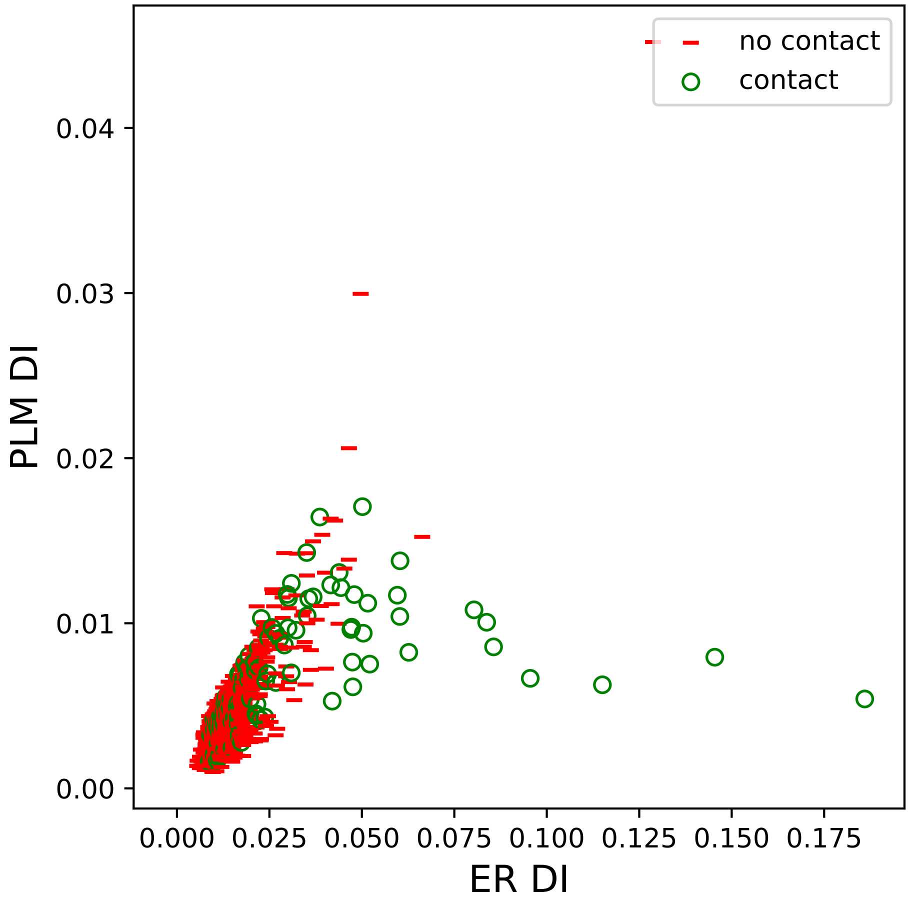

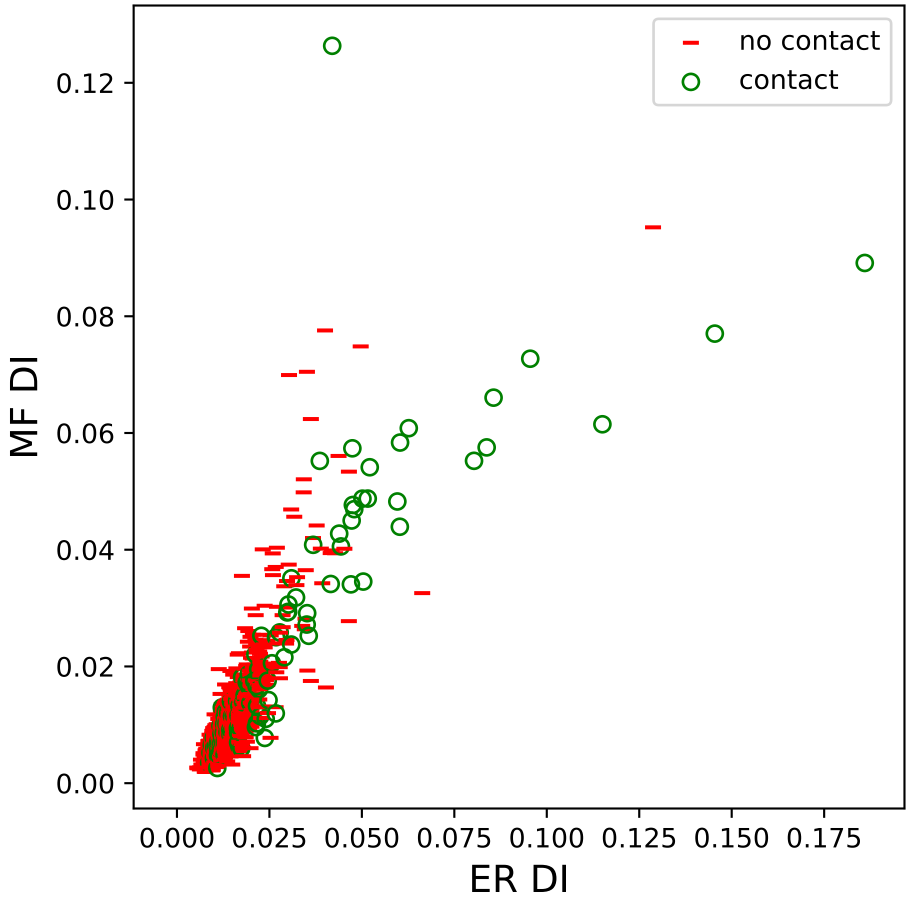

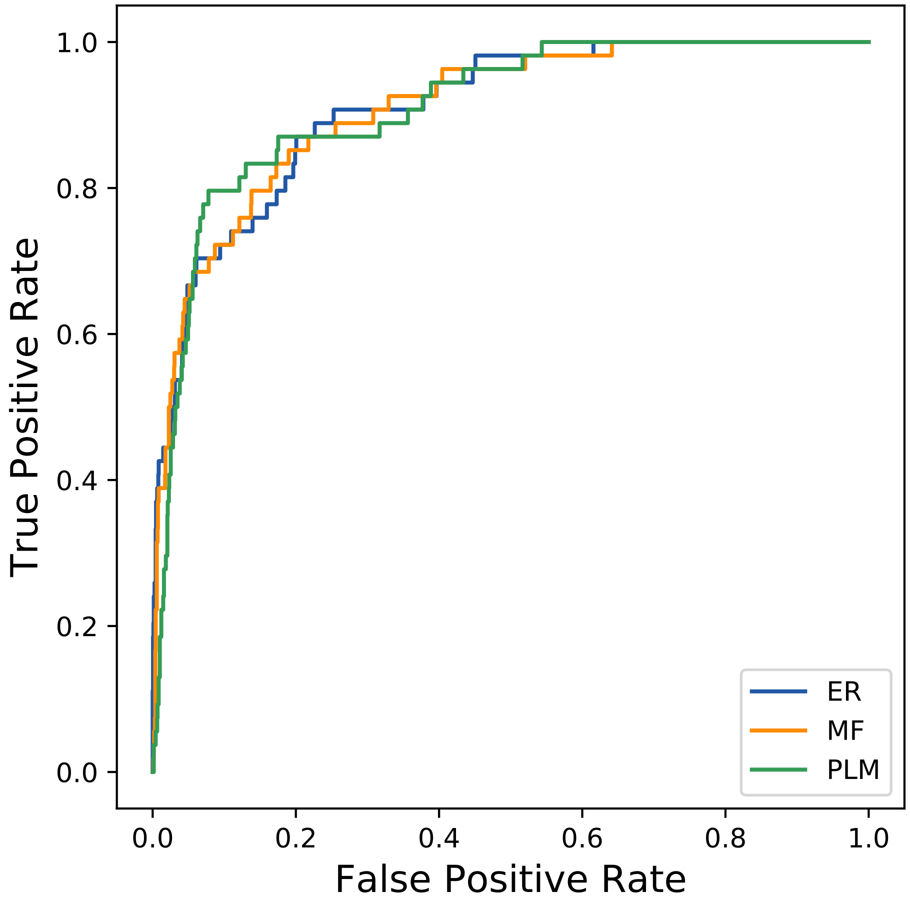

### 19175seq_70col_3aac_PF00011_plots.pdf

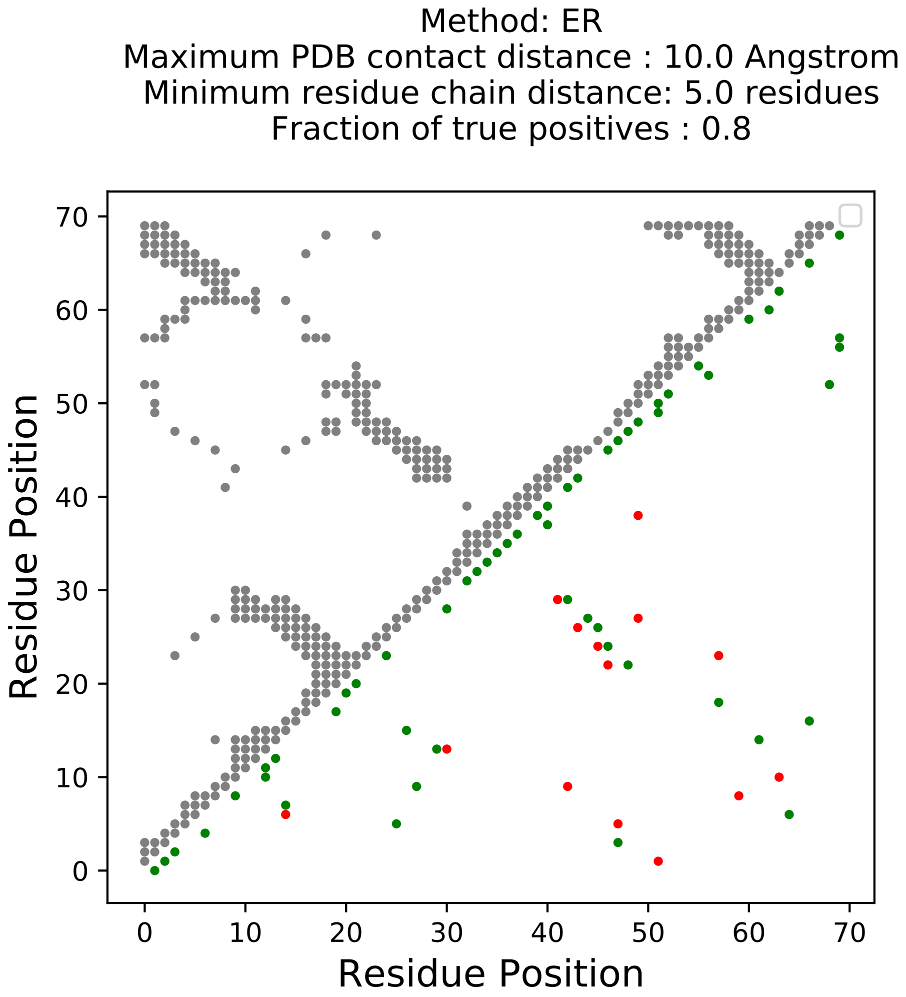

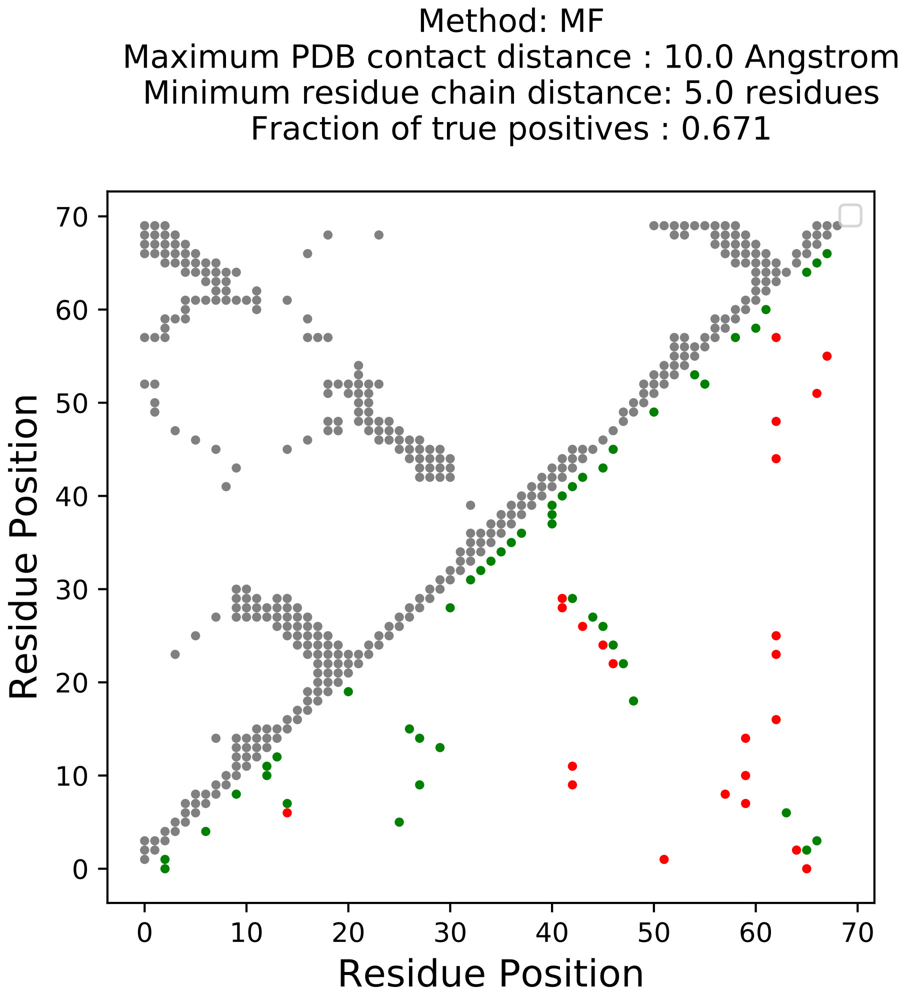

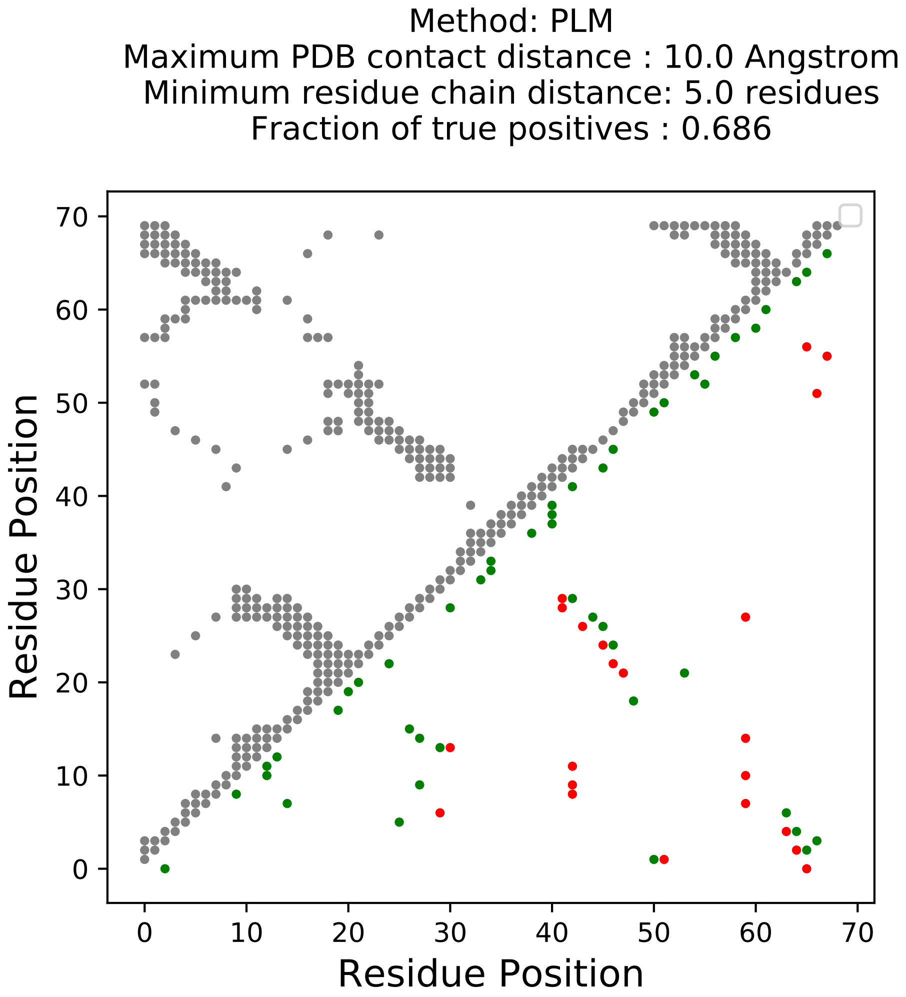

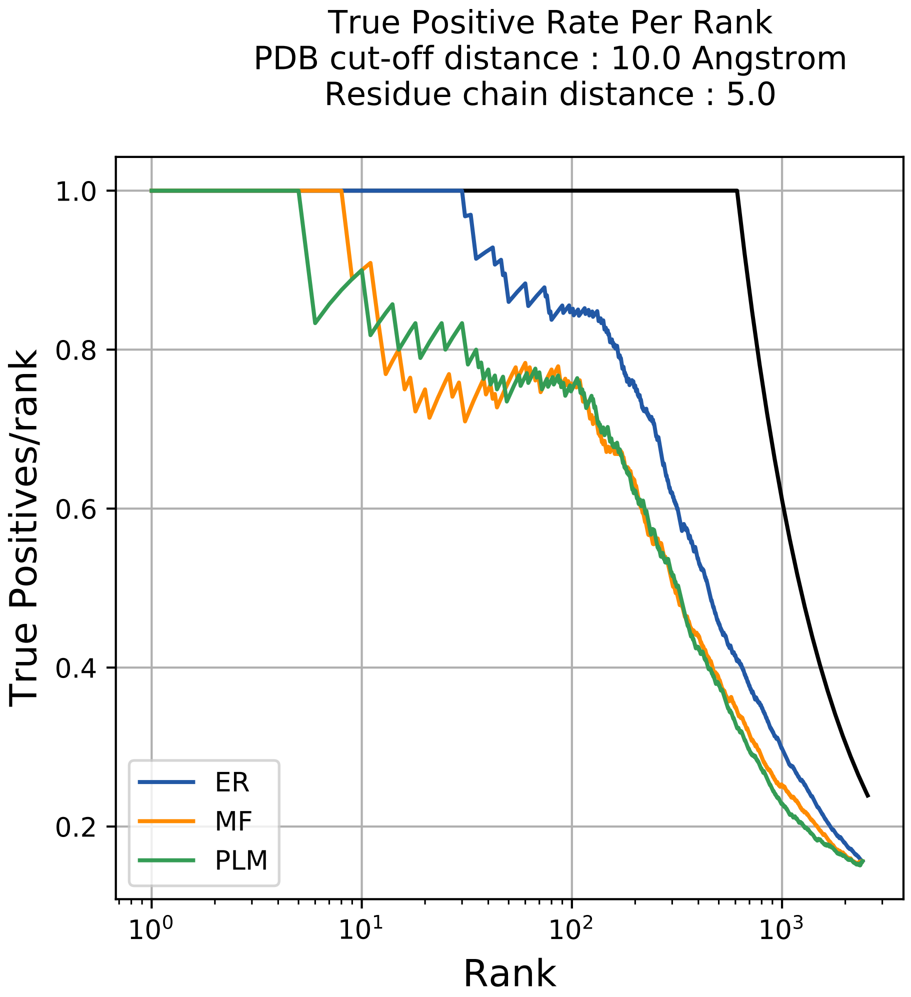

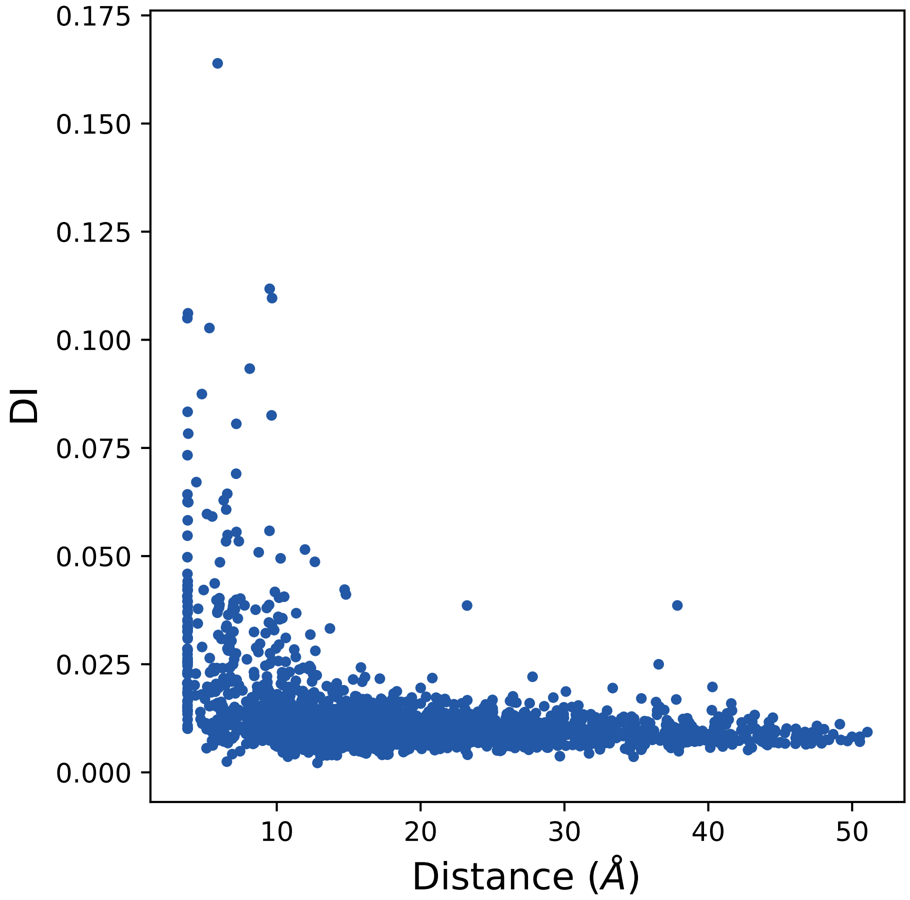

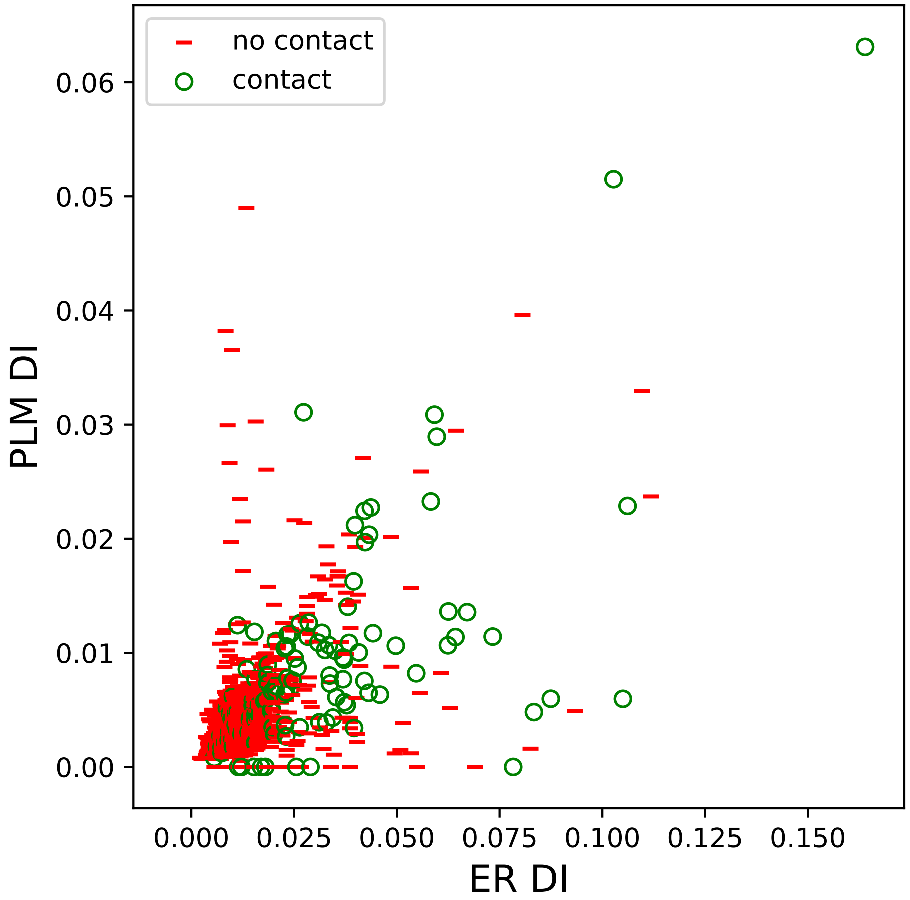

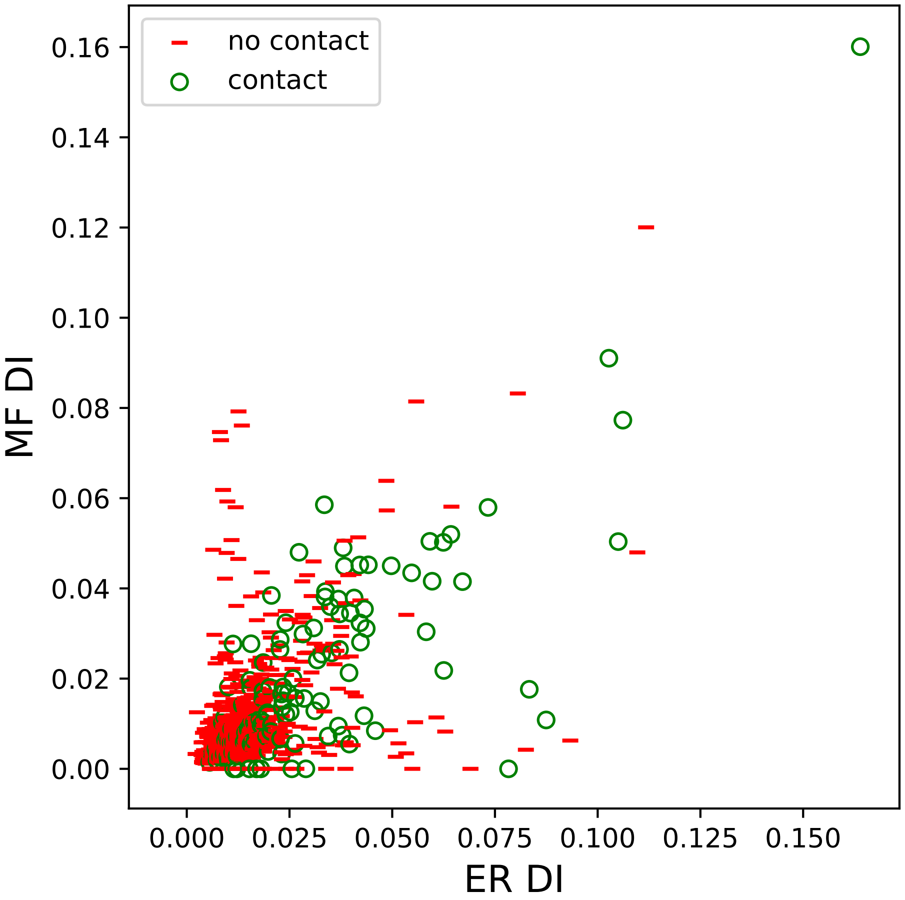

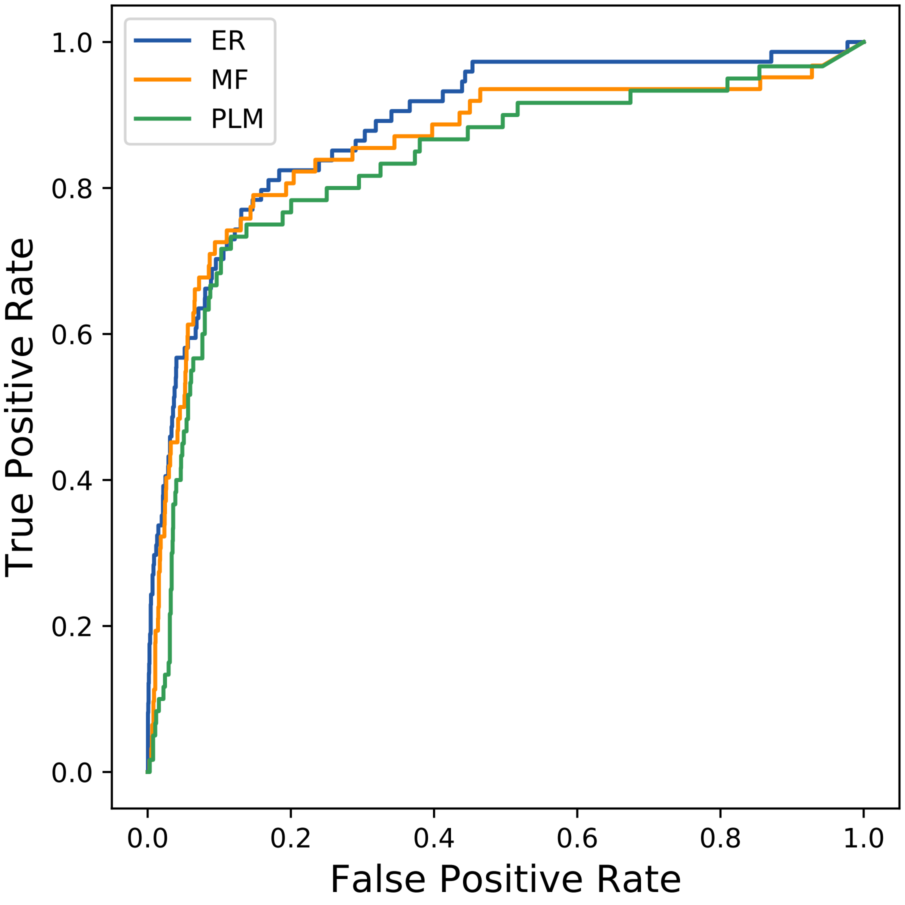

### 19970seq_63col_5j4u_PF02798_plots.pdf

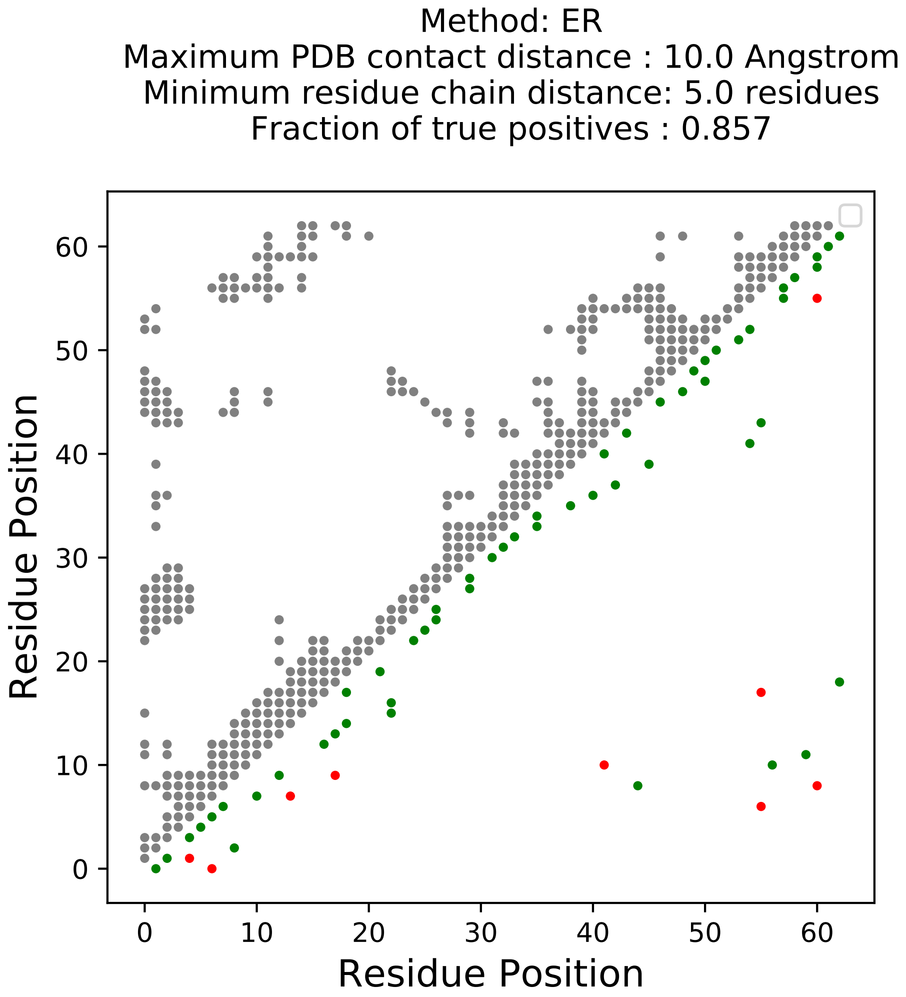

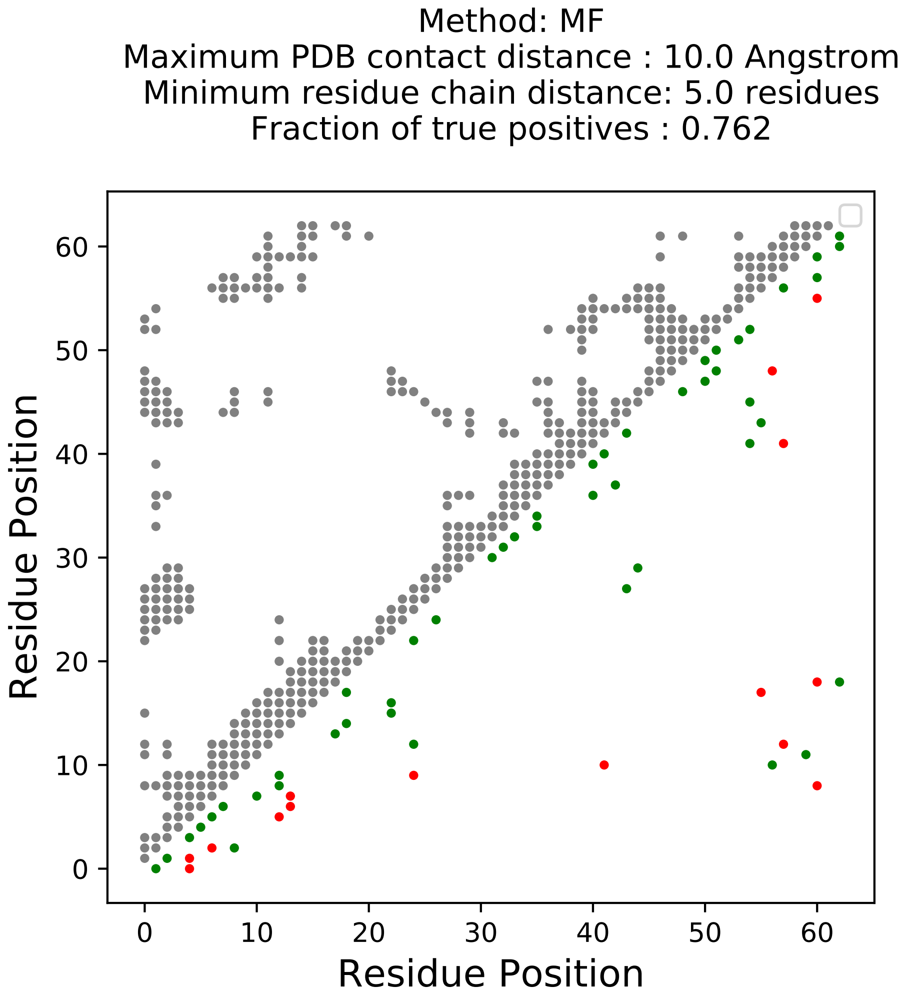

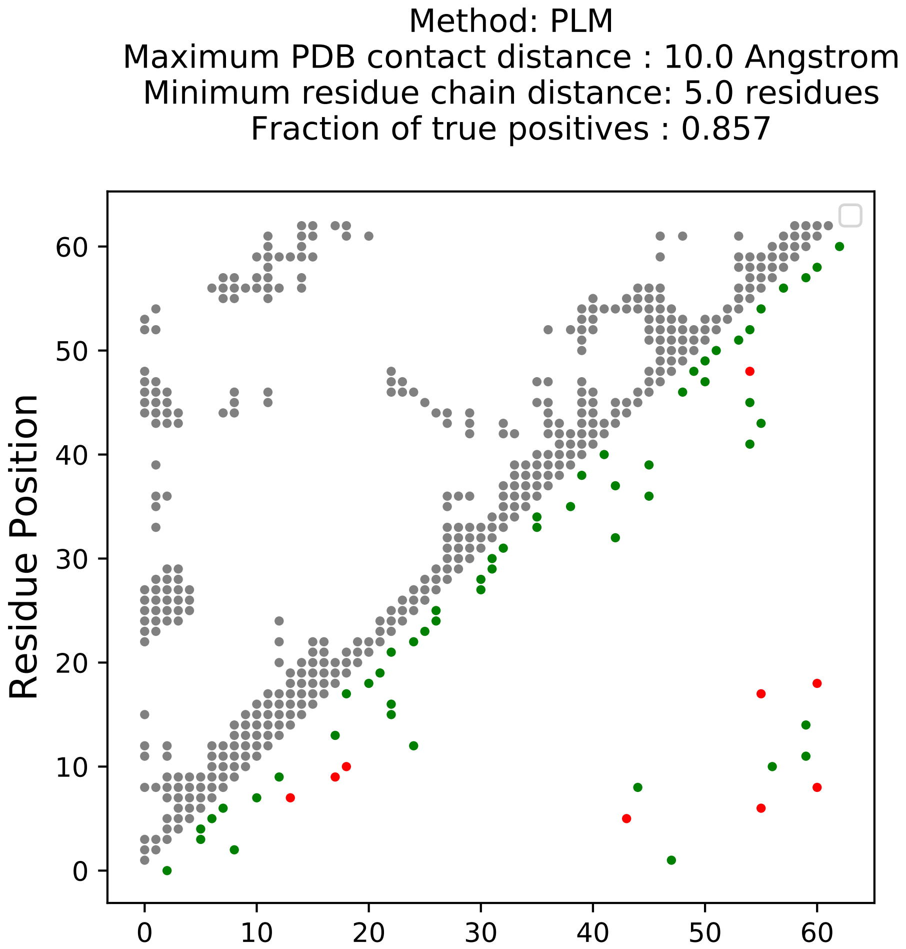

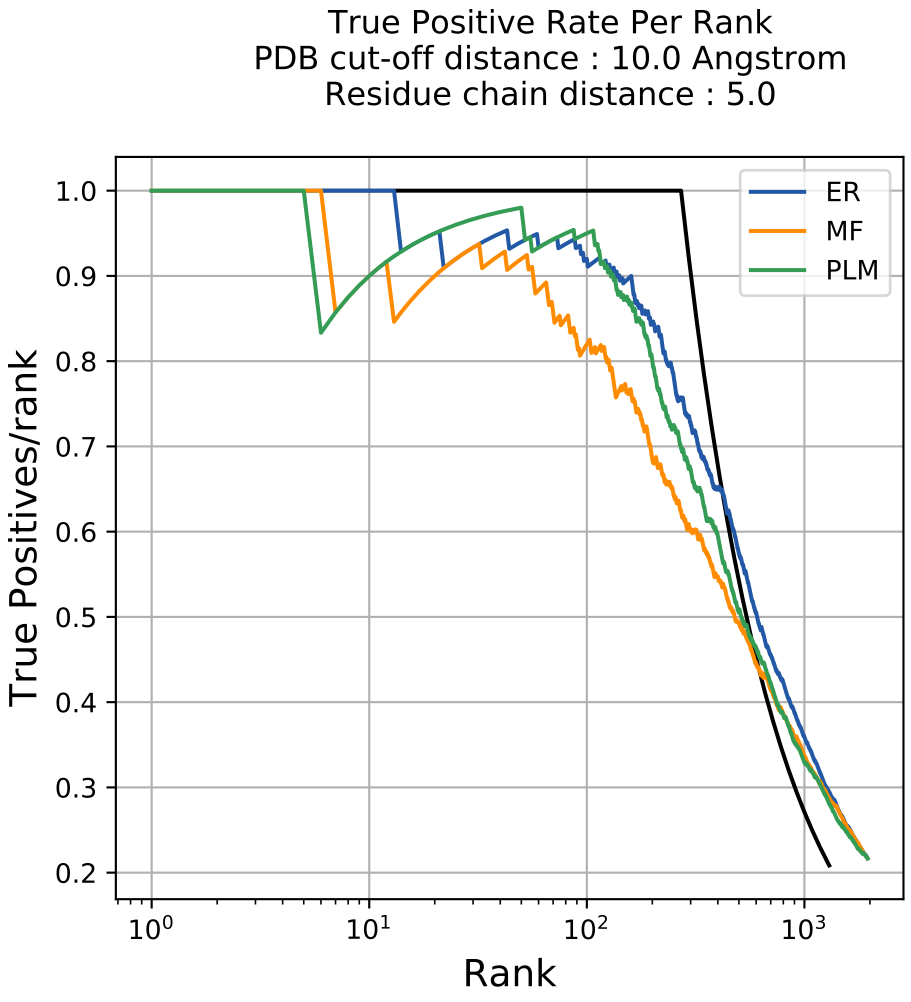

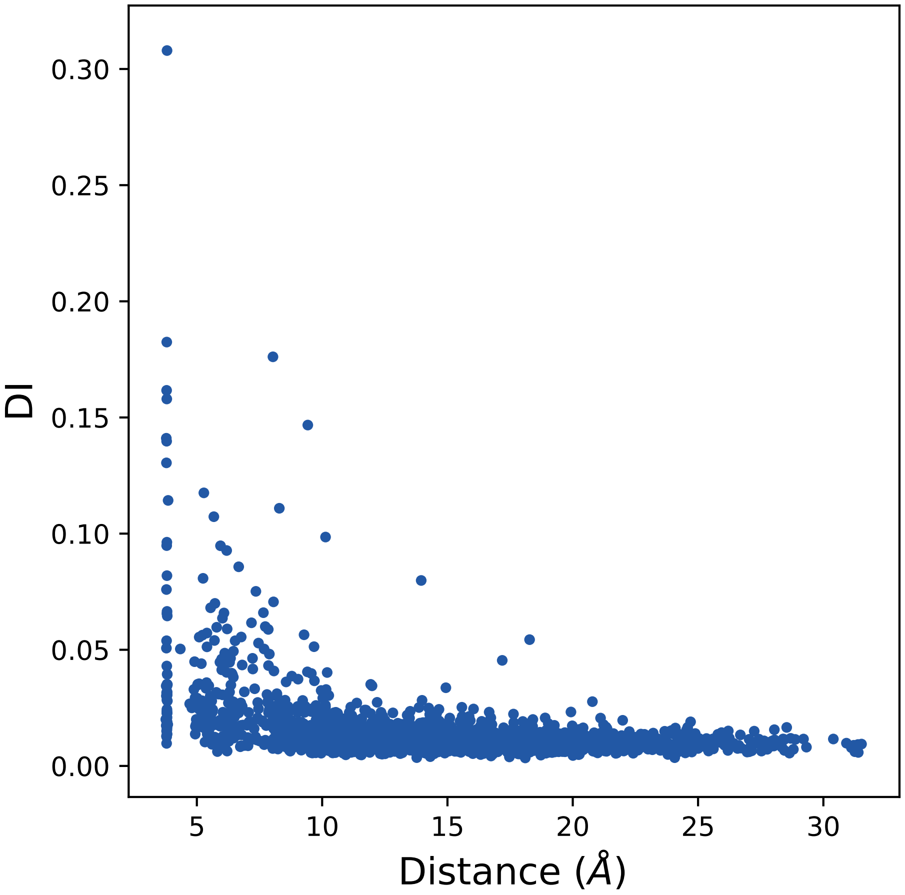

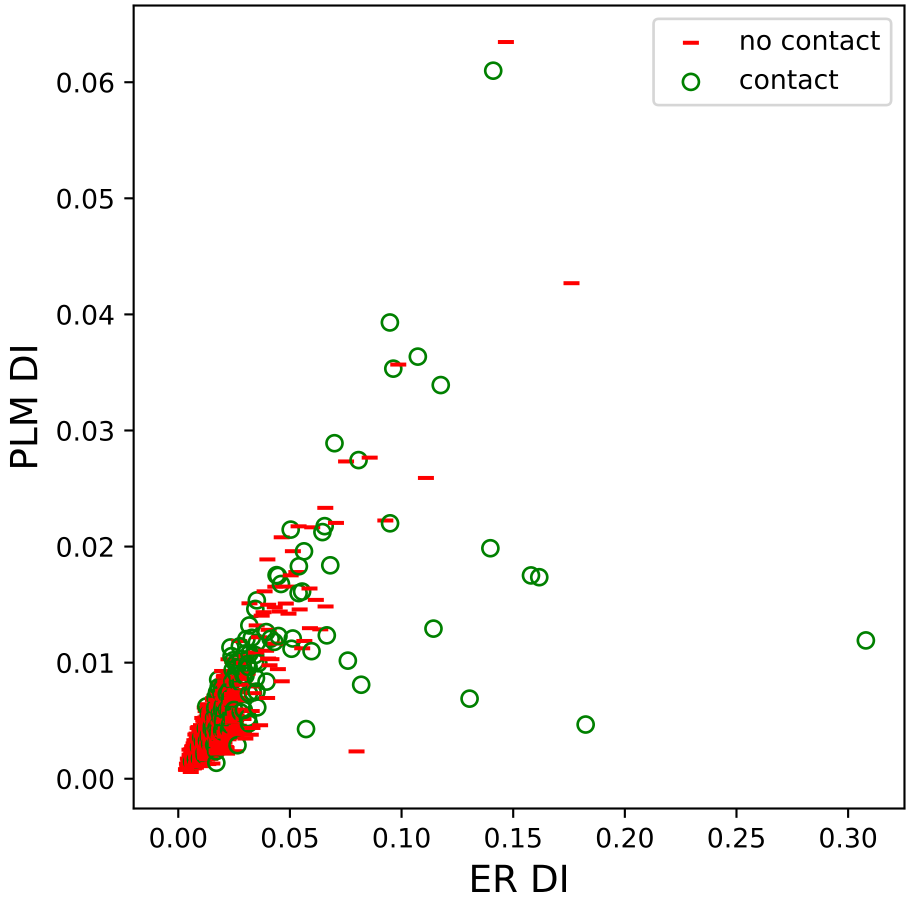

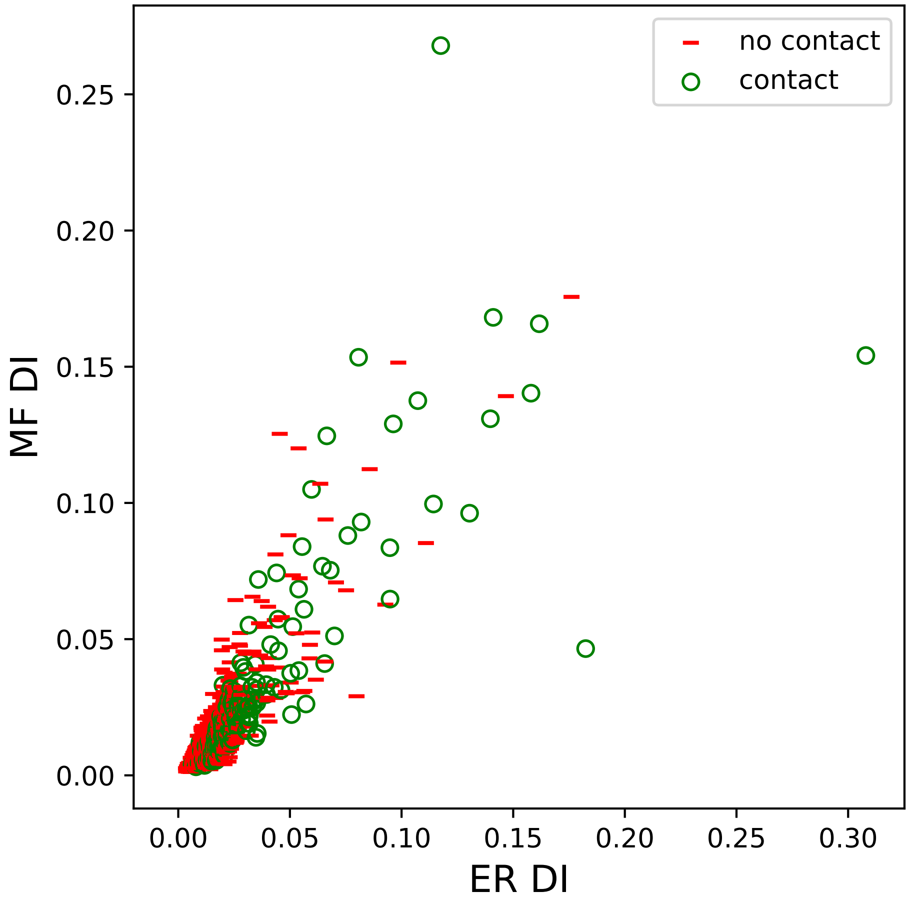

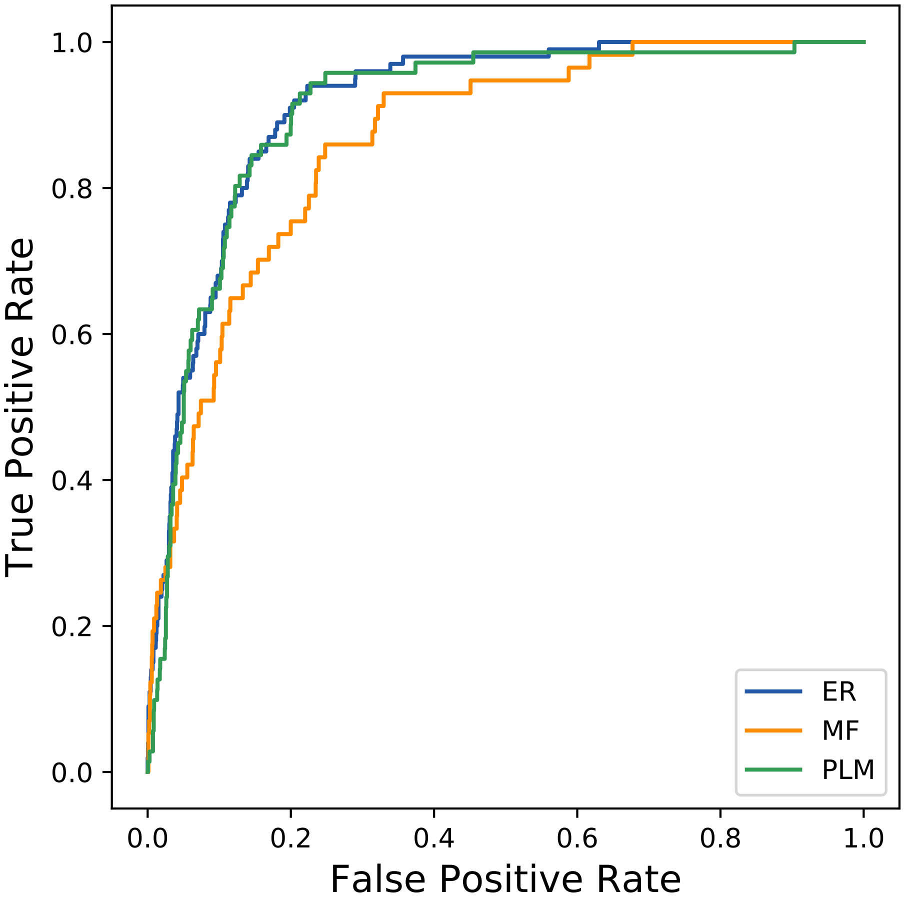

### 20118seq_145col_5a8g_PF00701_plots.pdf

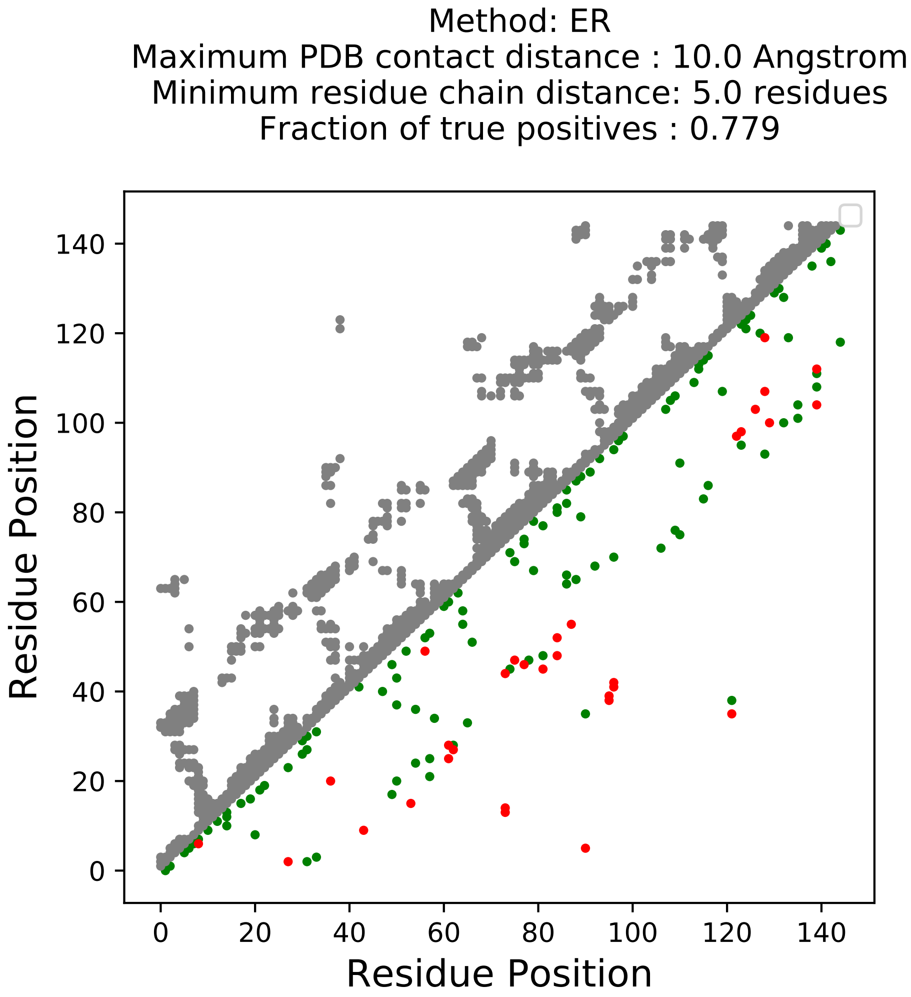

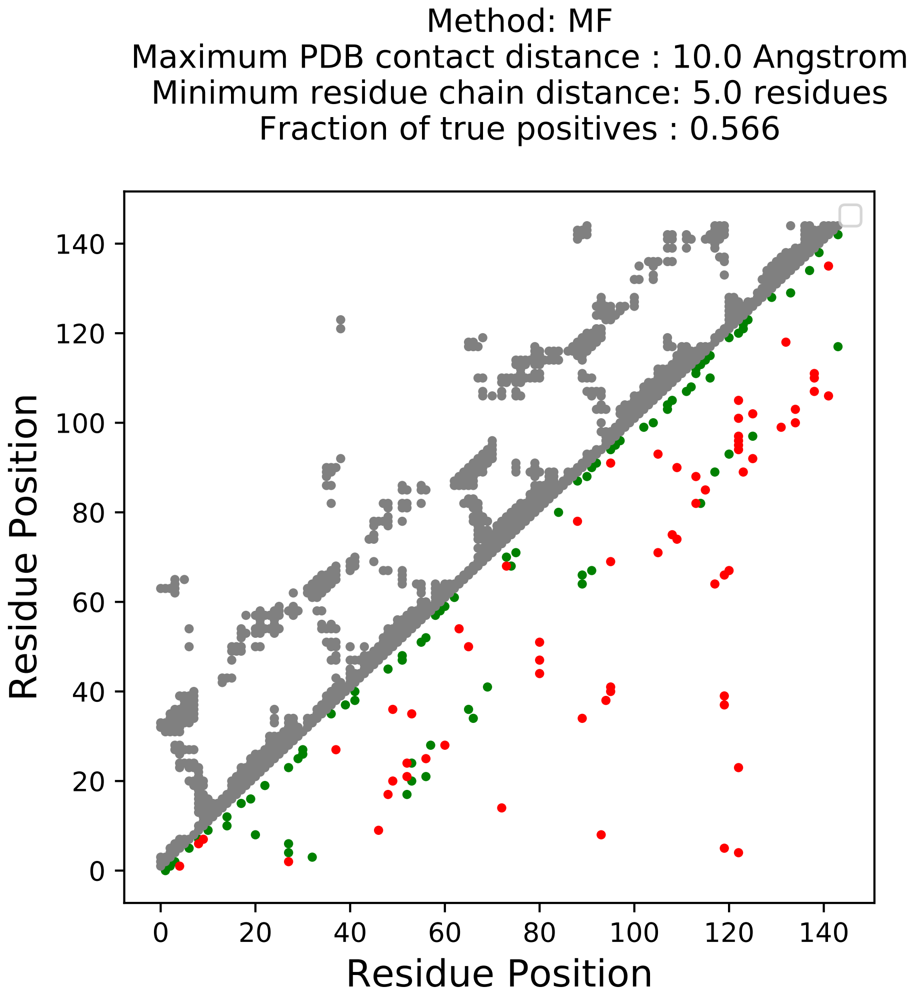

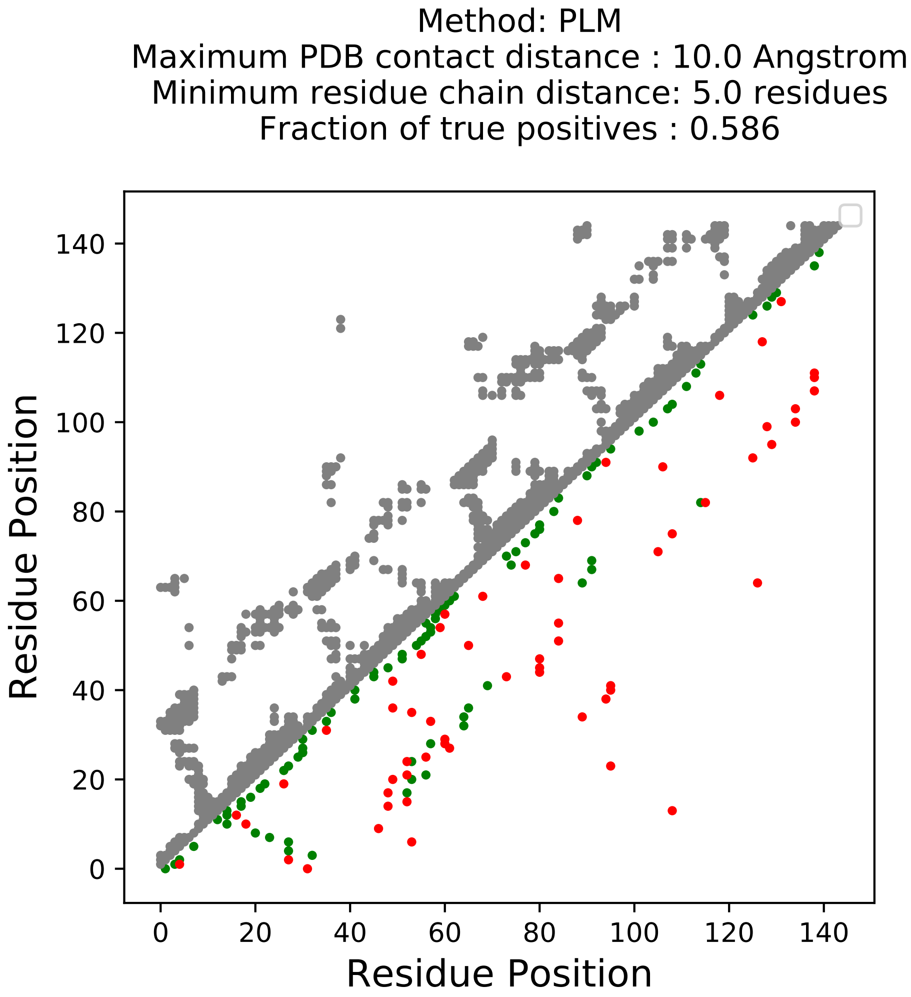

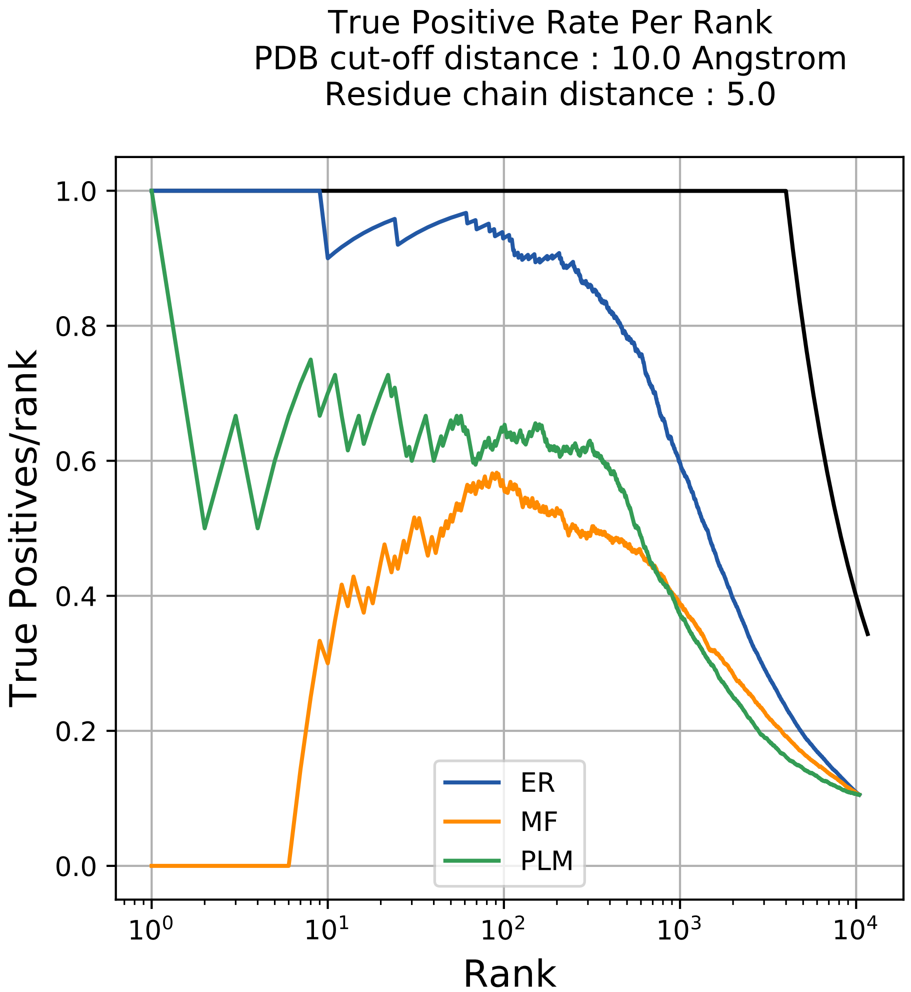

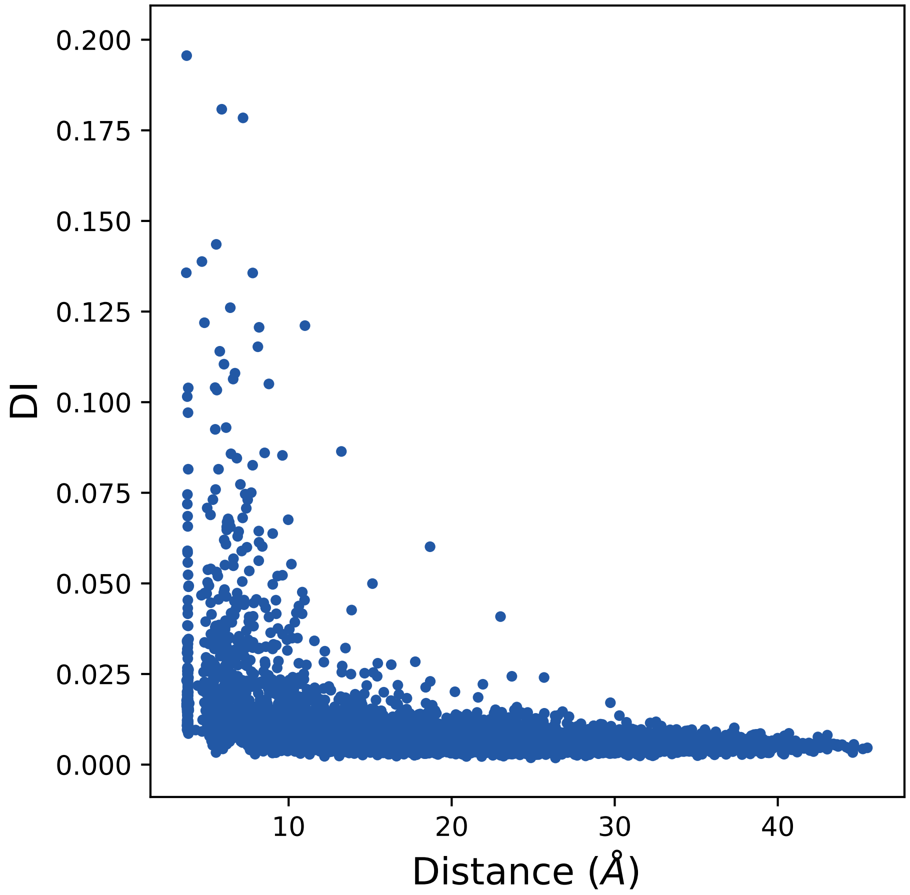

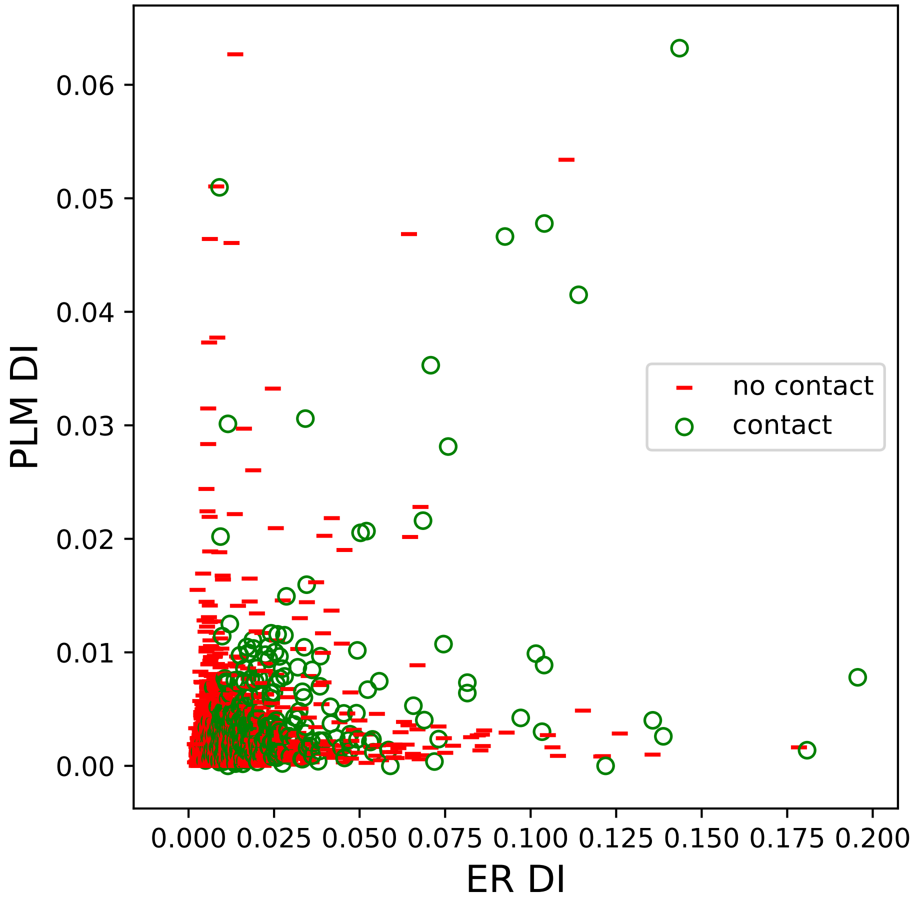

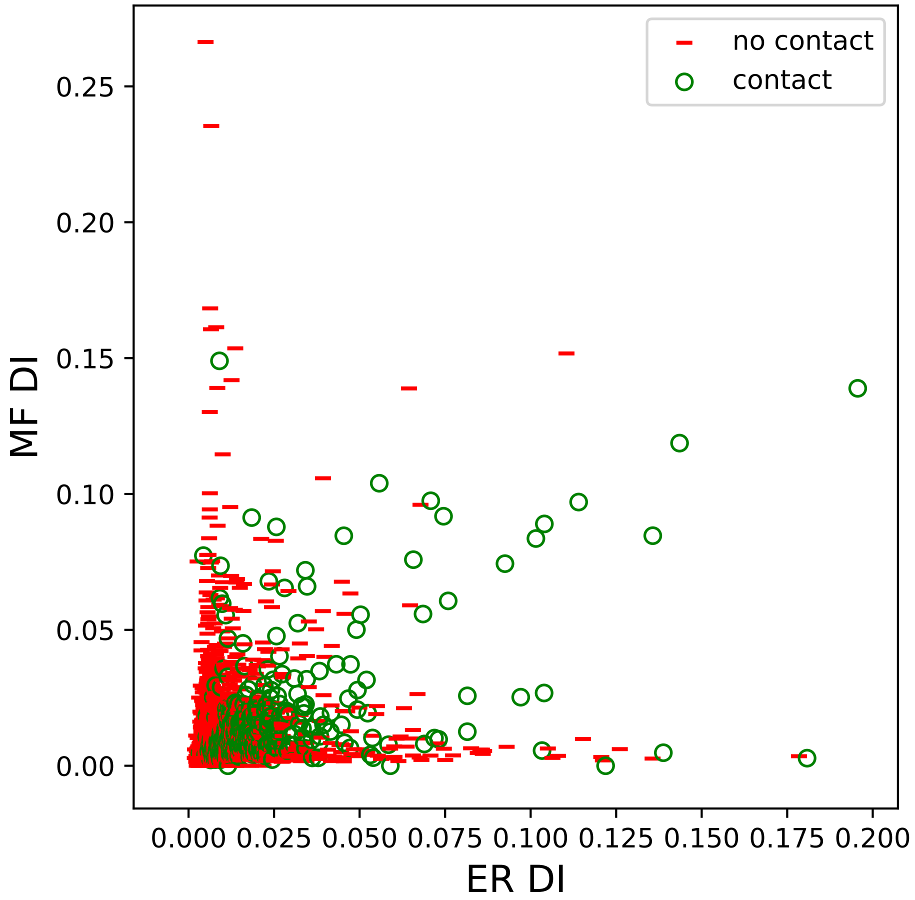

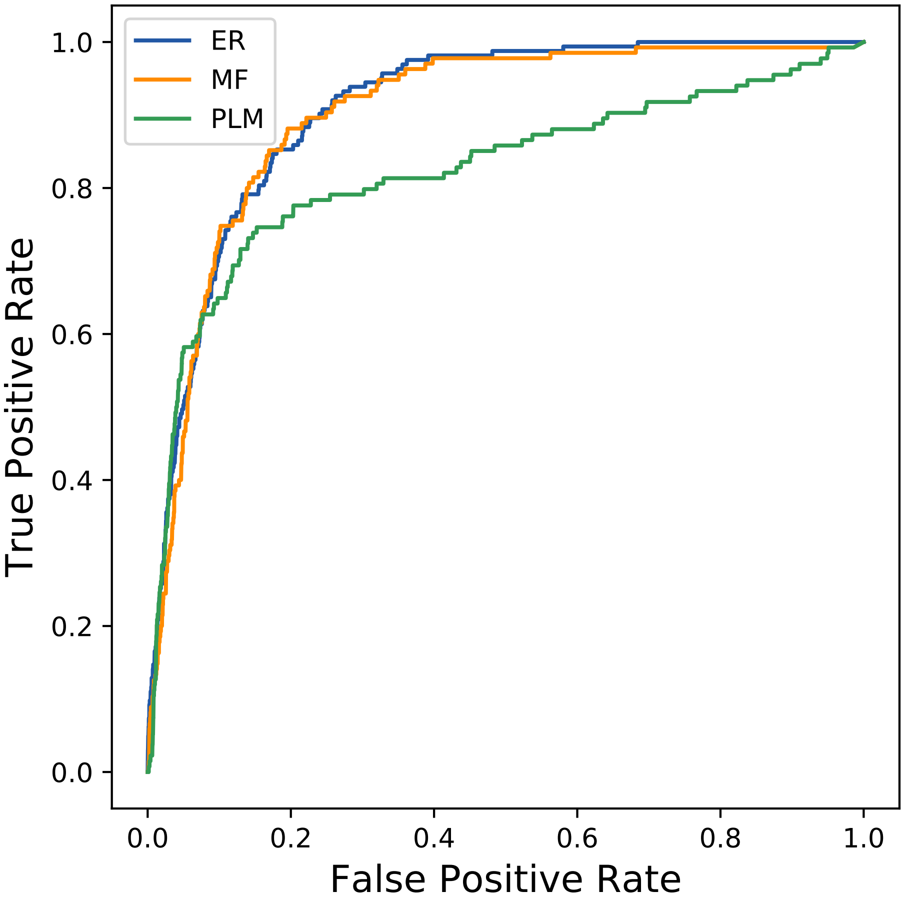

### 20391seq_140col_3e2m_PF00092_plots.pdf

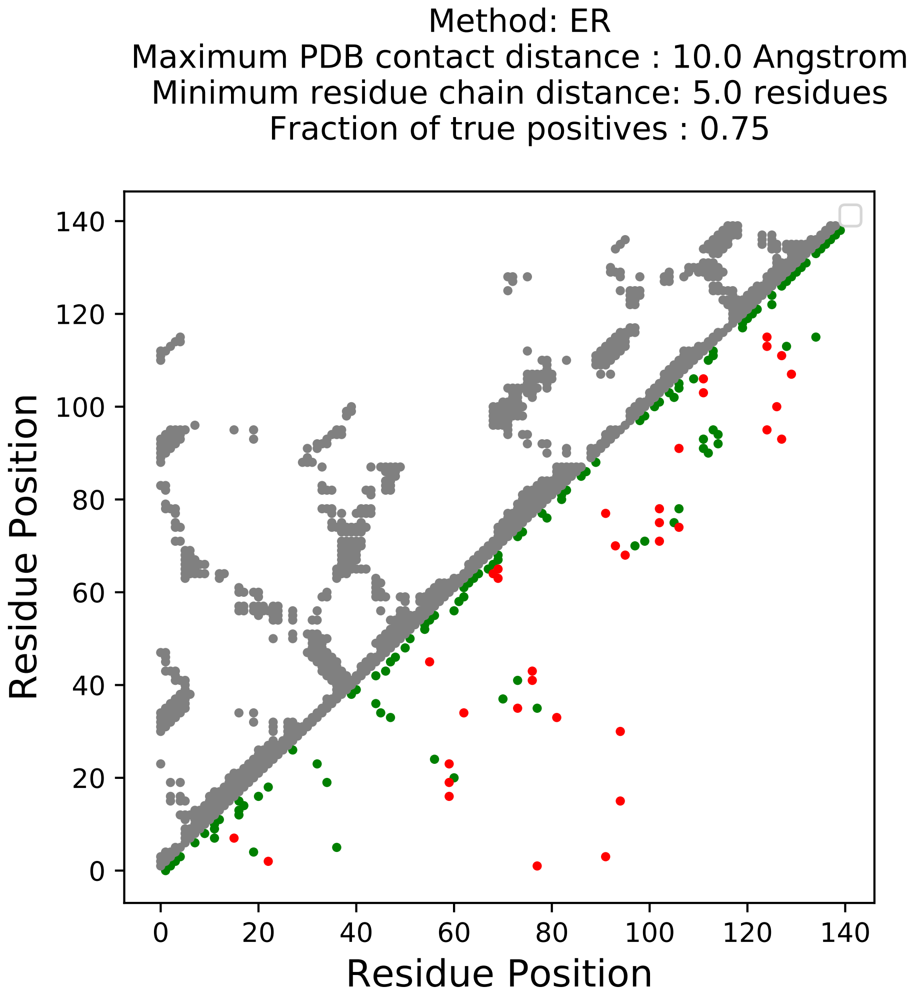

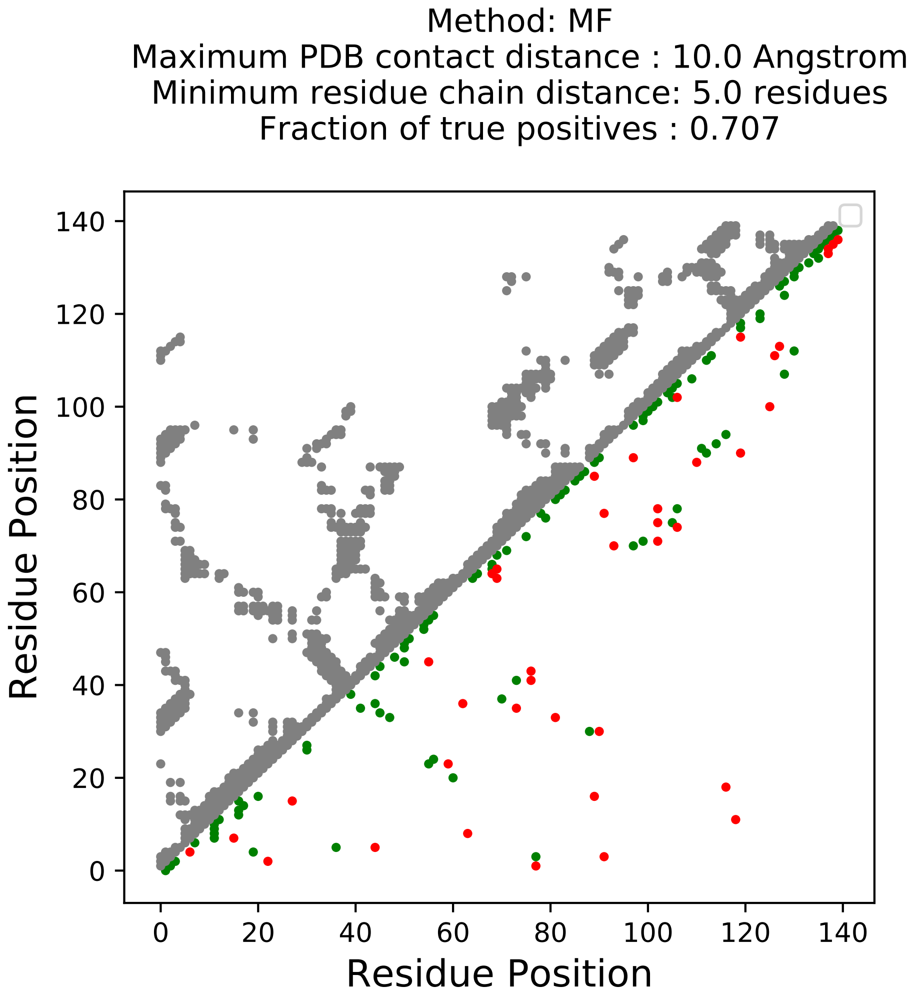

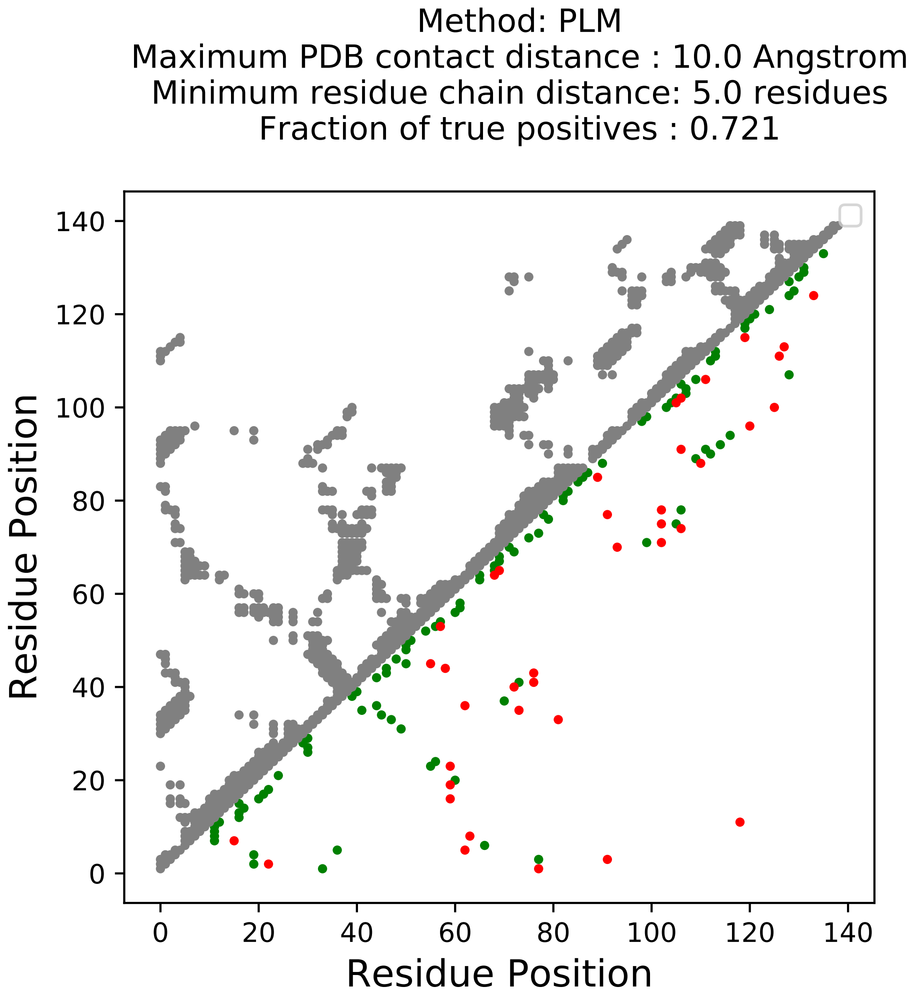

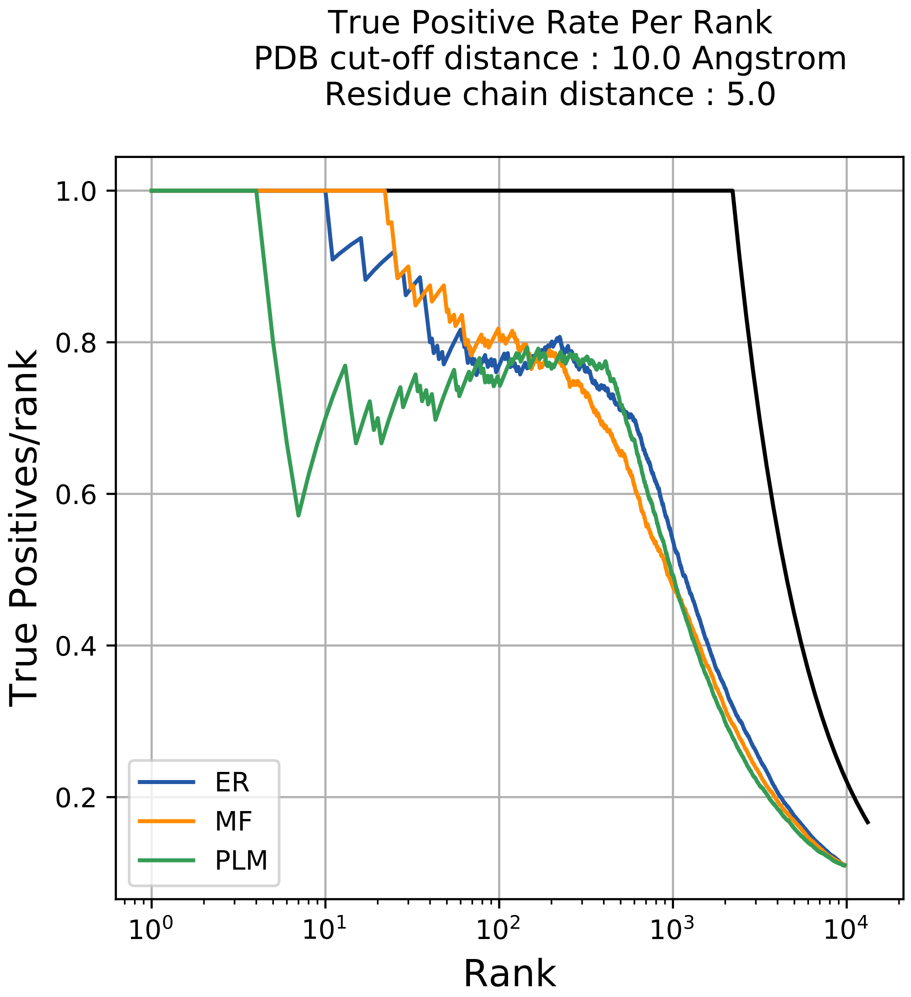

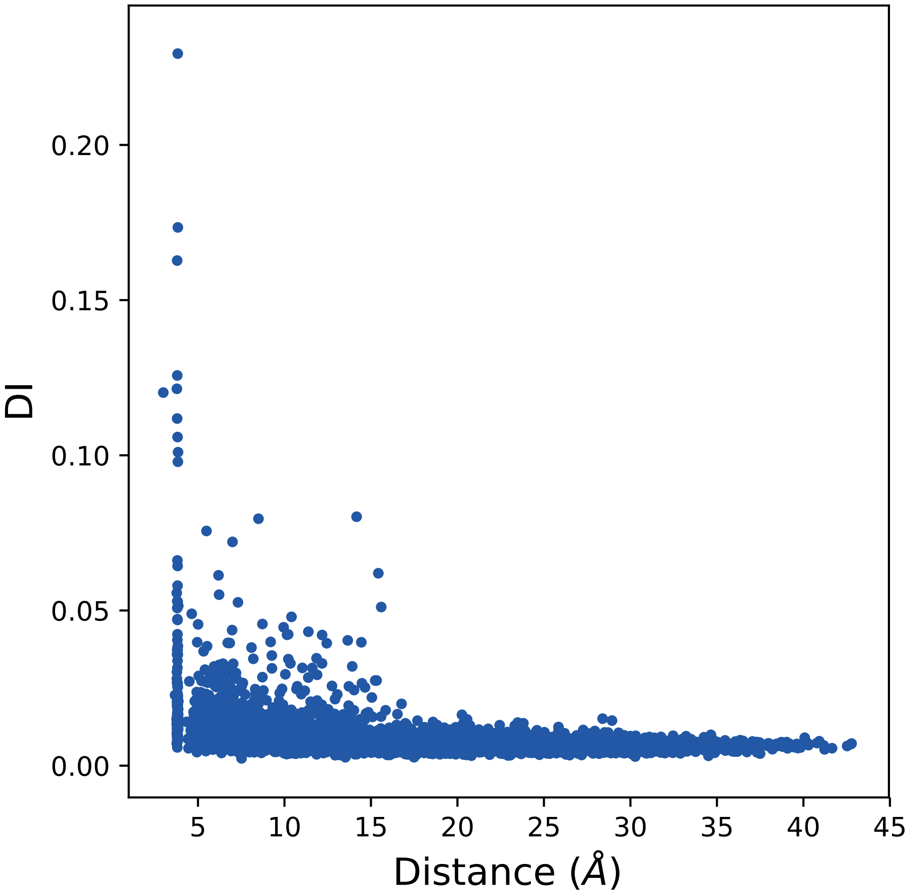

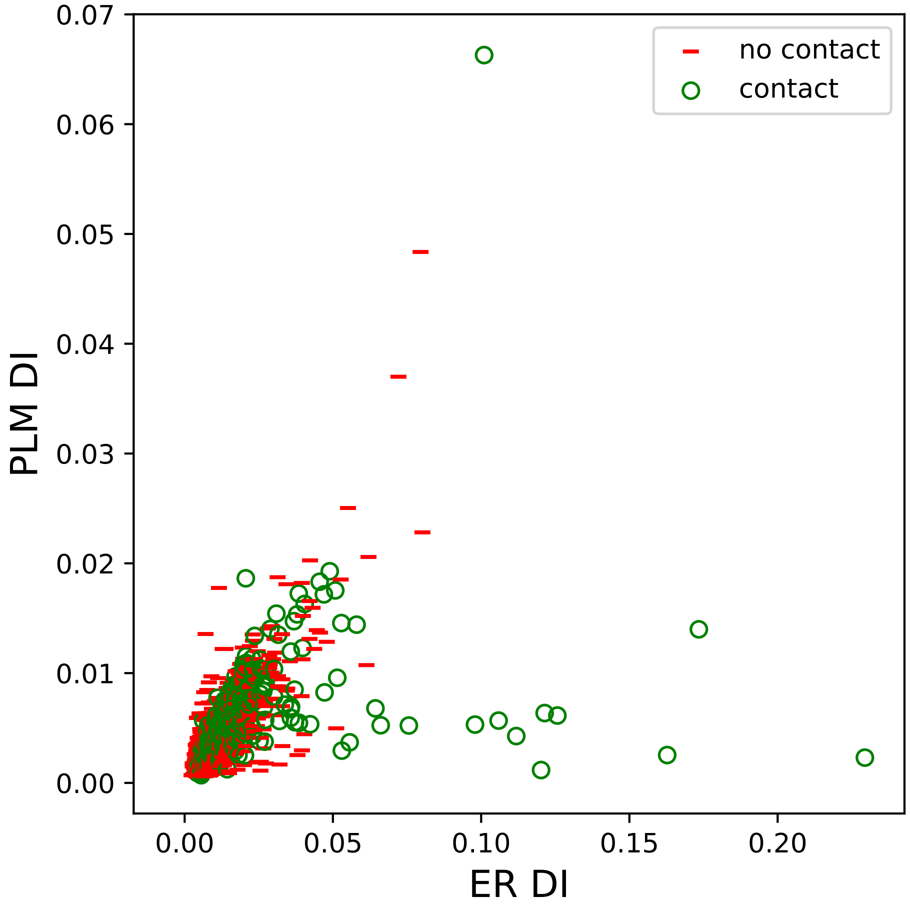

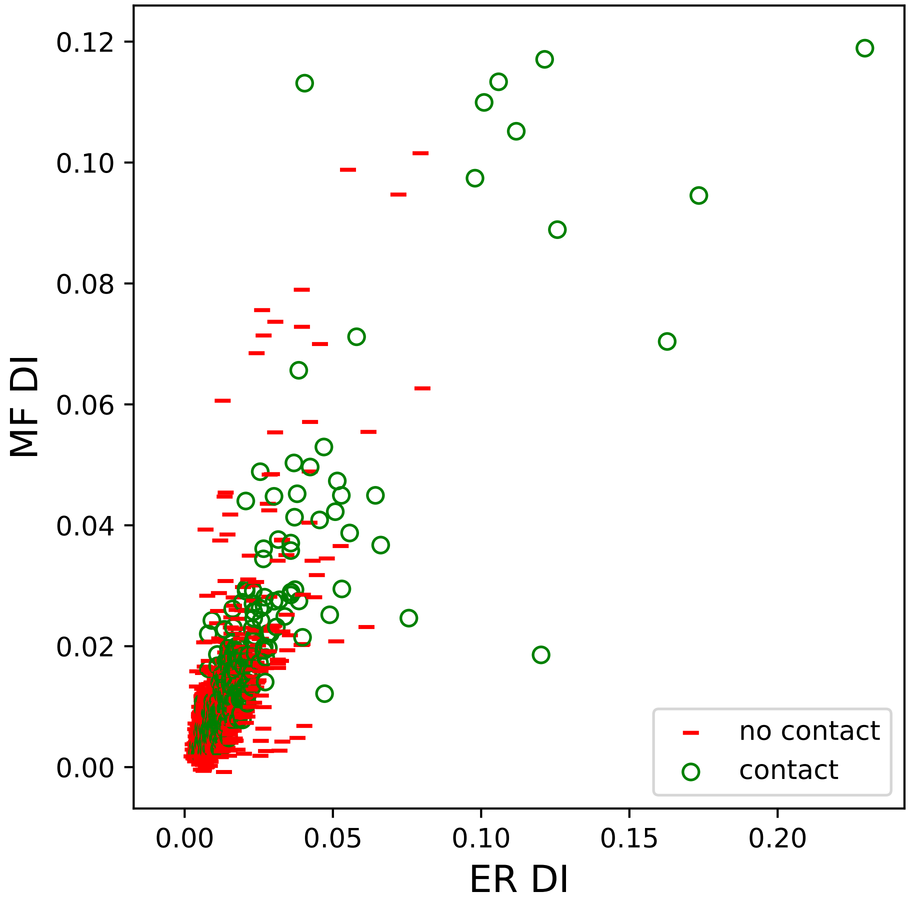

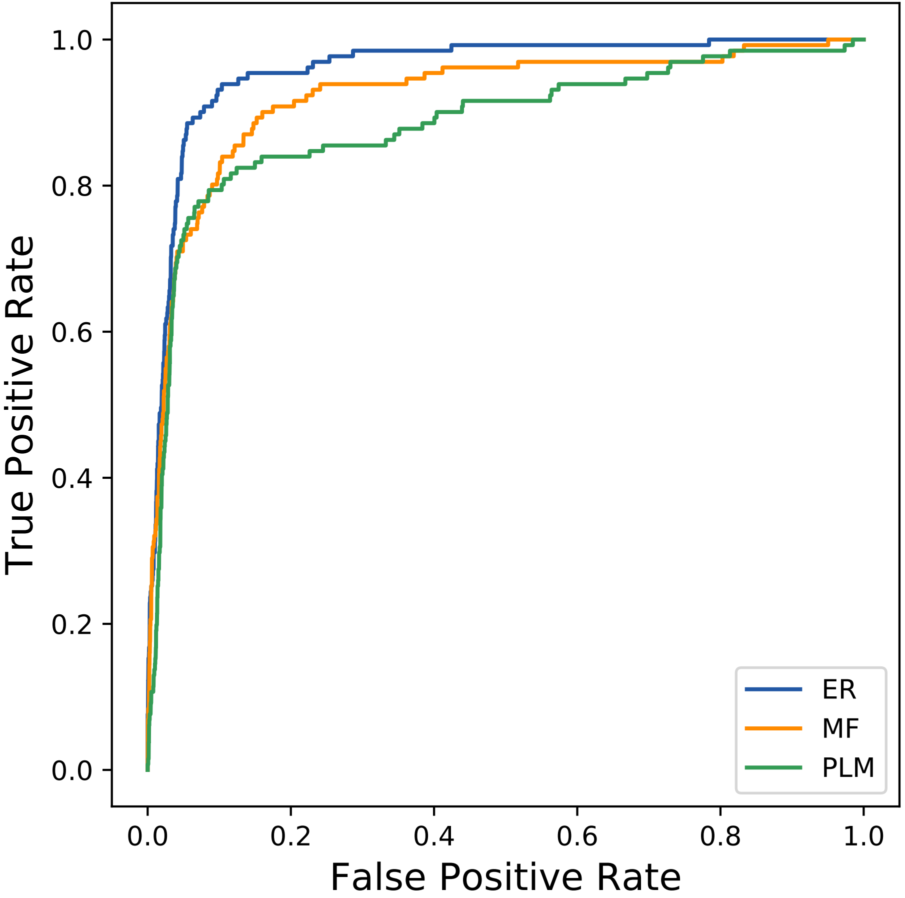

### 20421seq_34col_1k0t_PF12838_plots.pdf

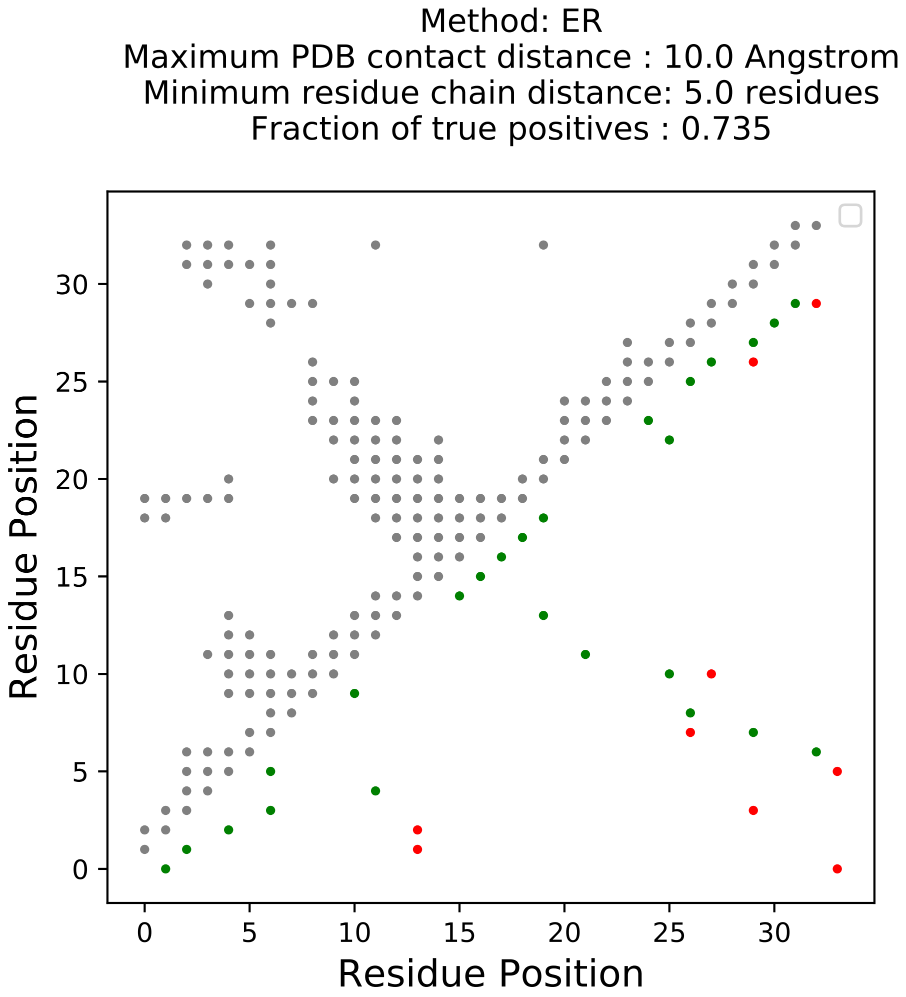

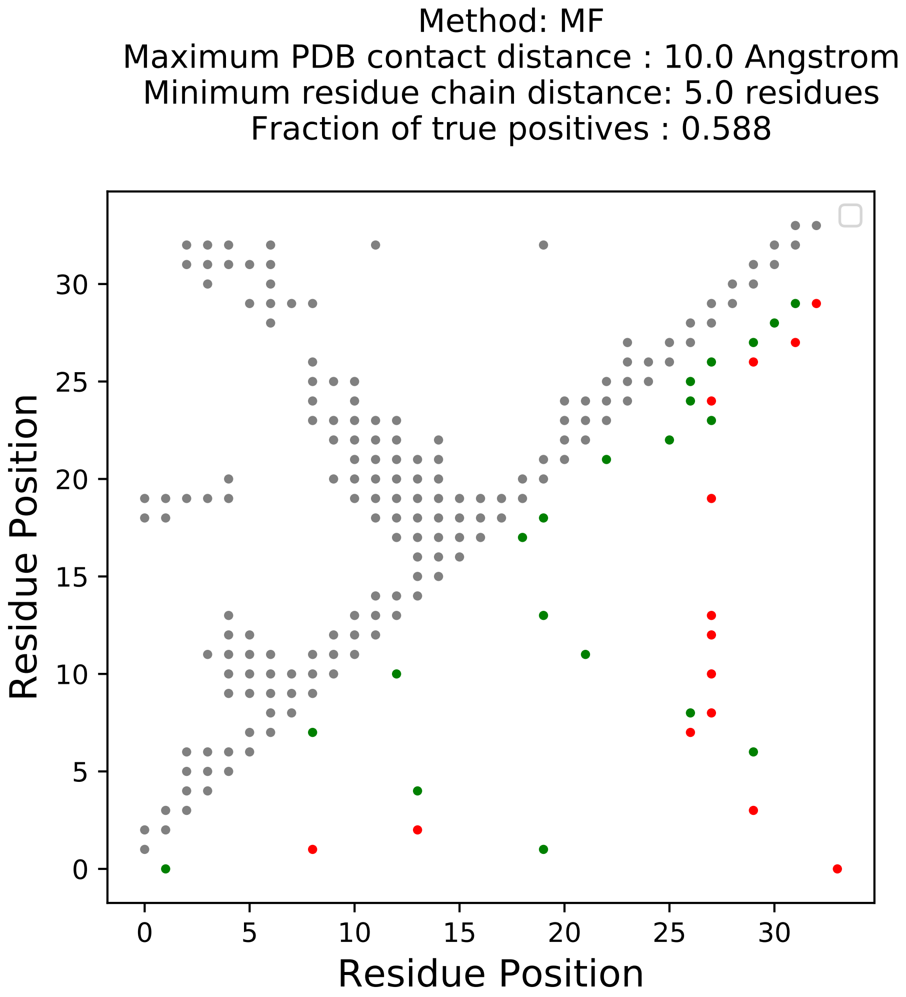

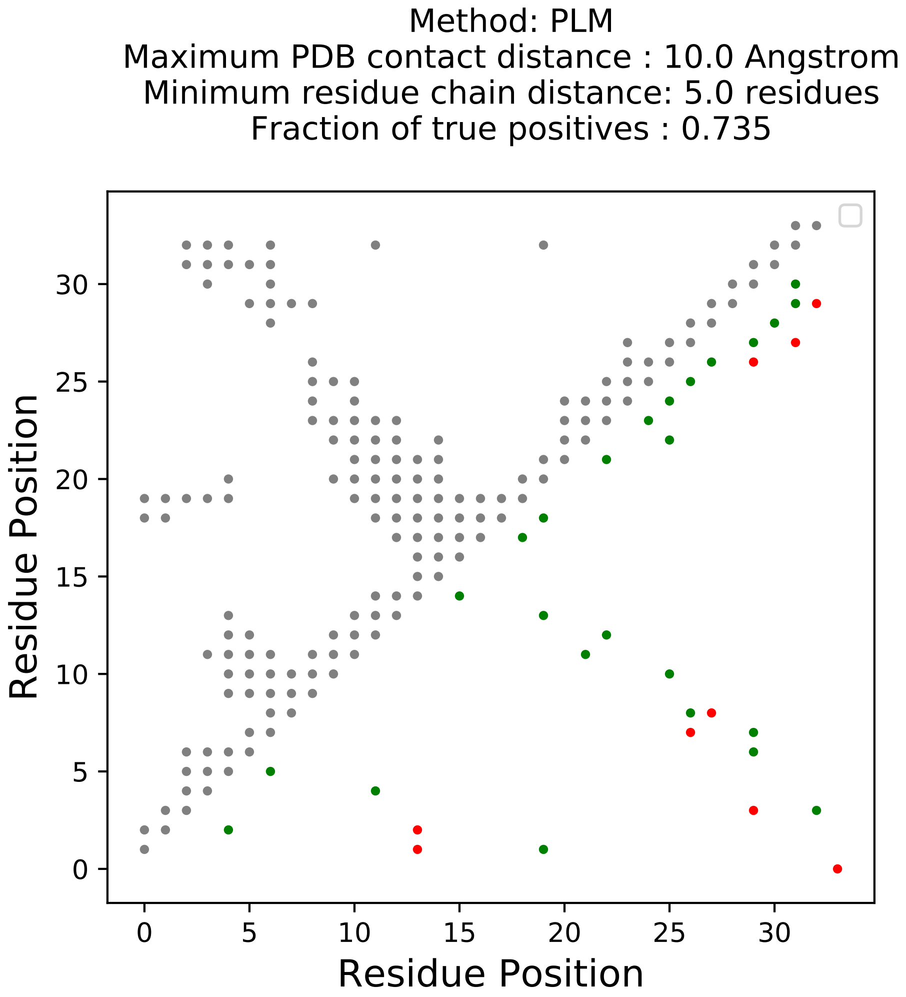

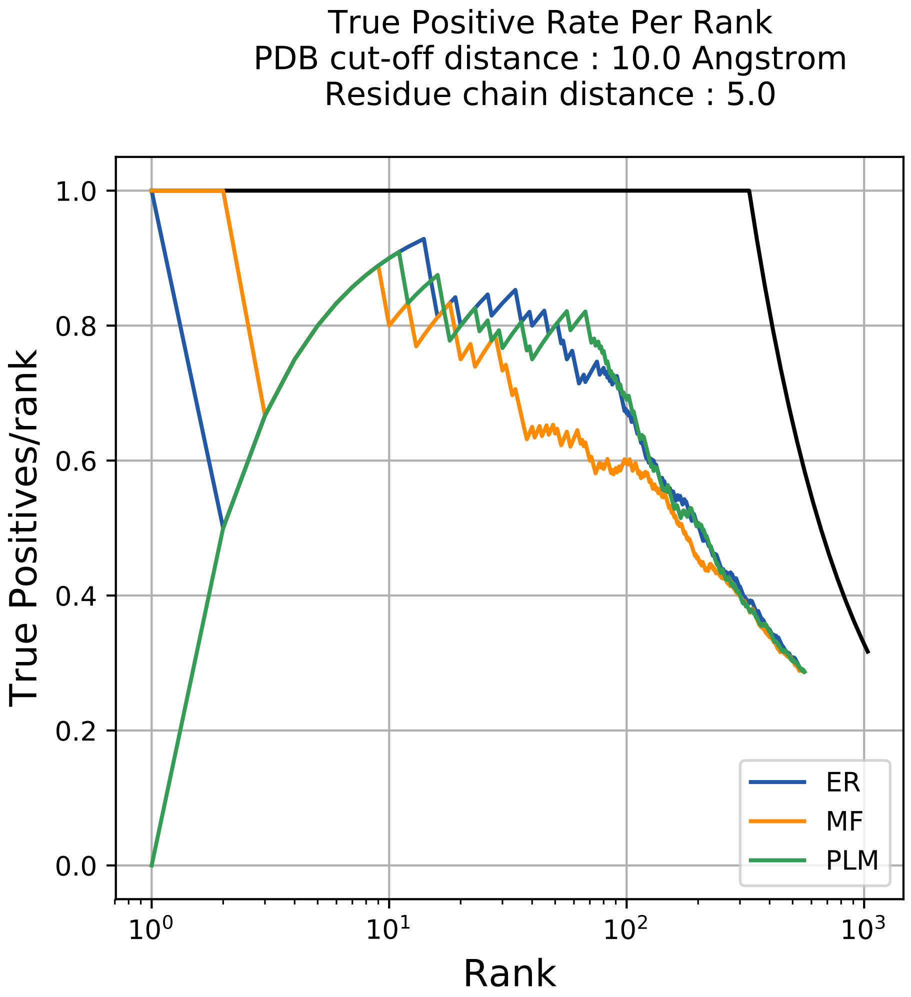

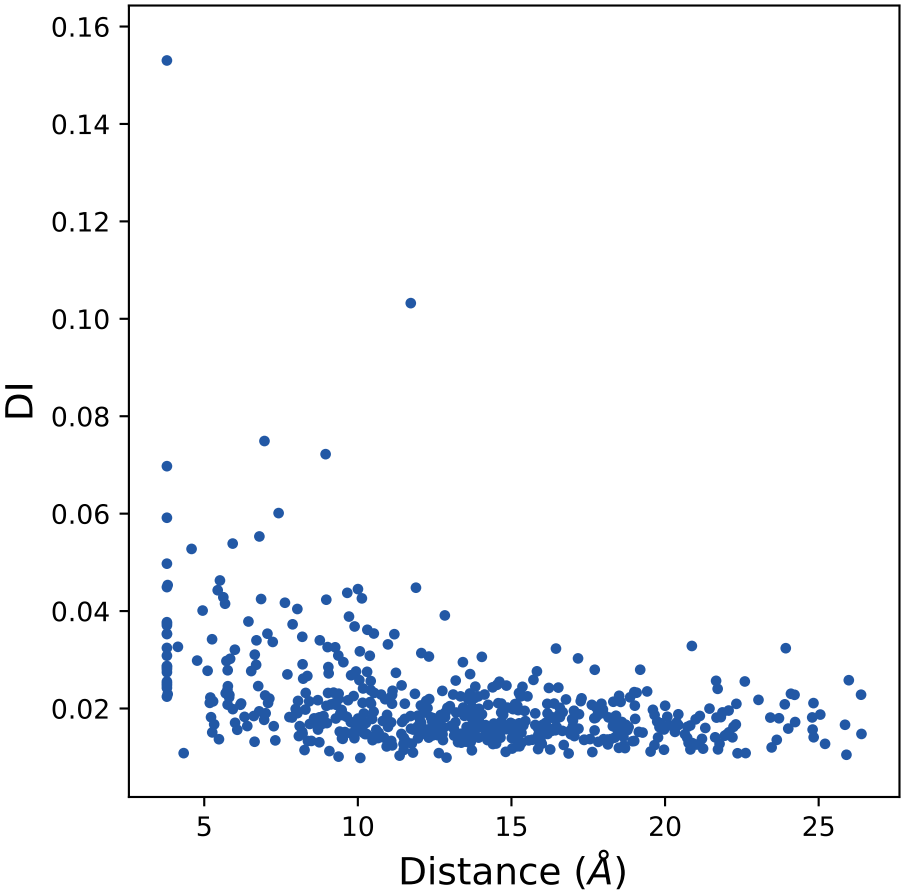

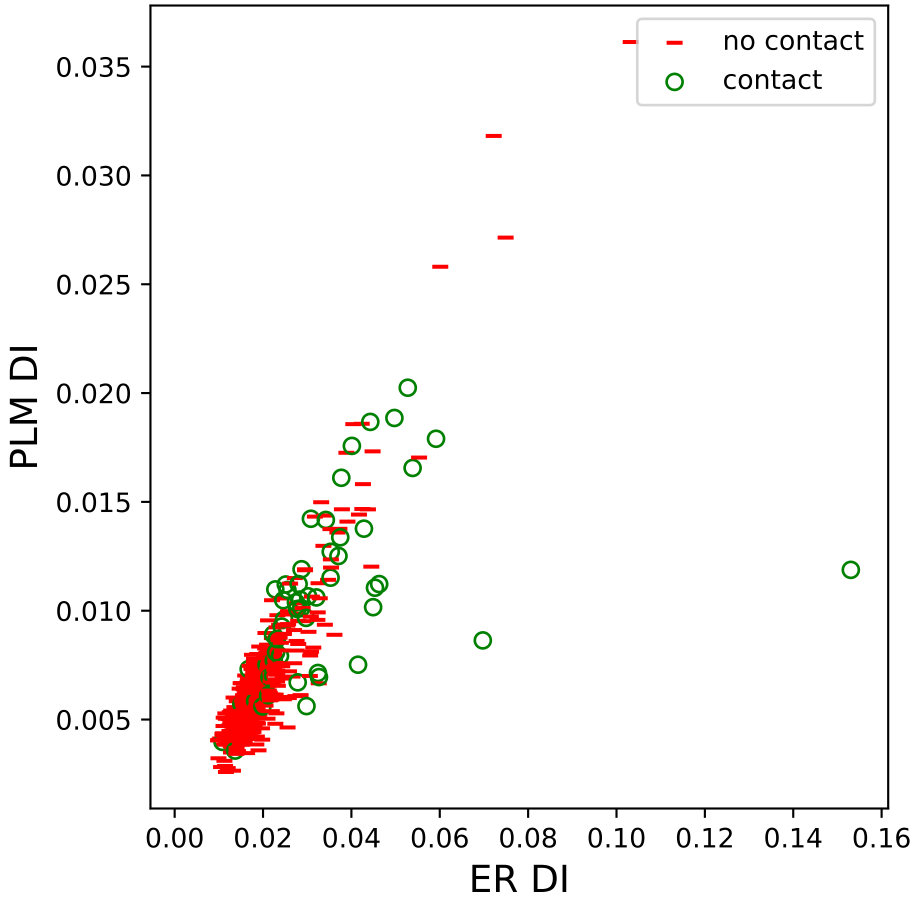

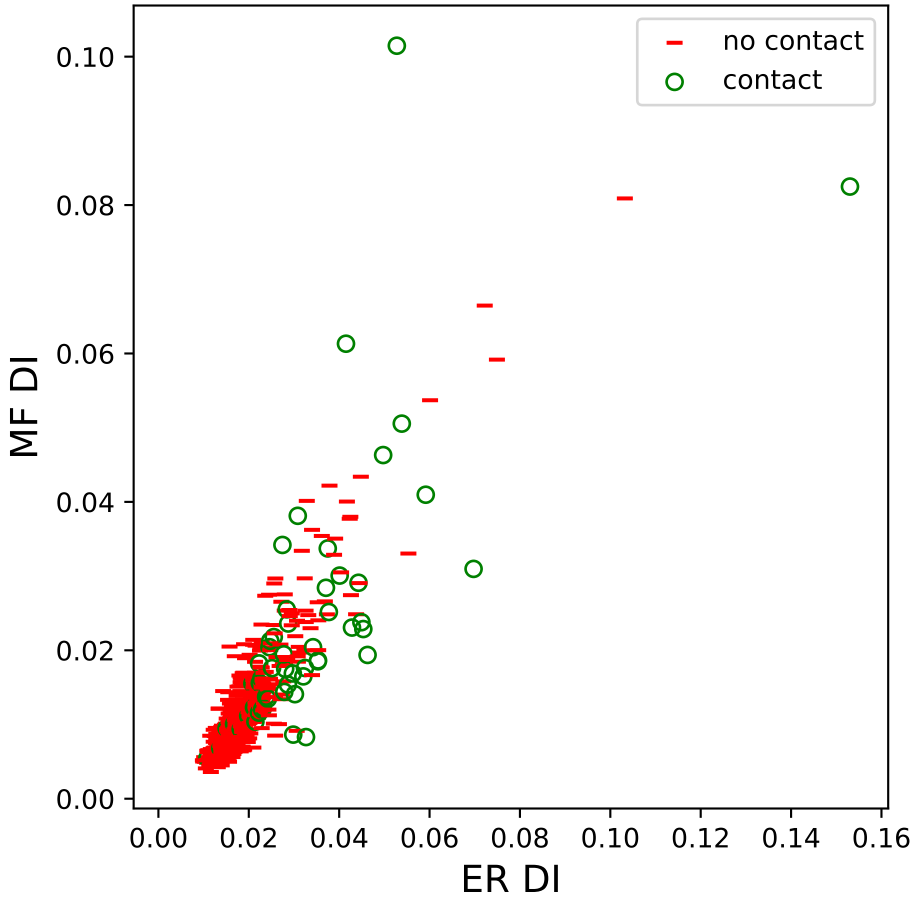

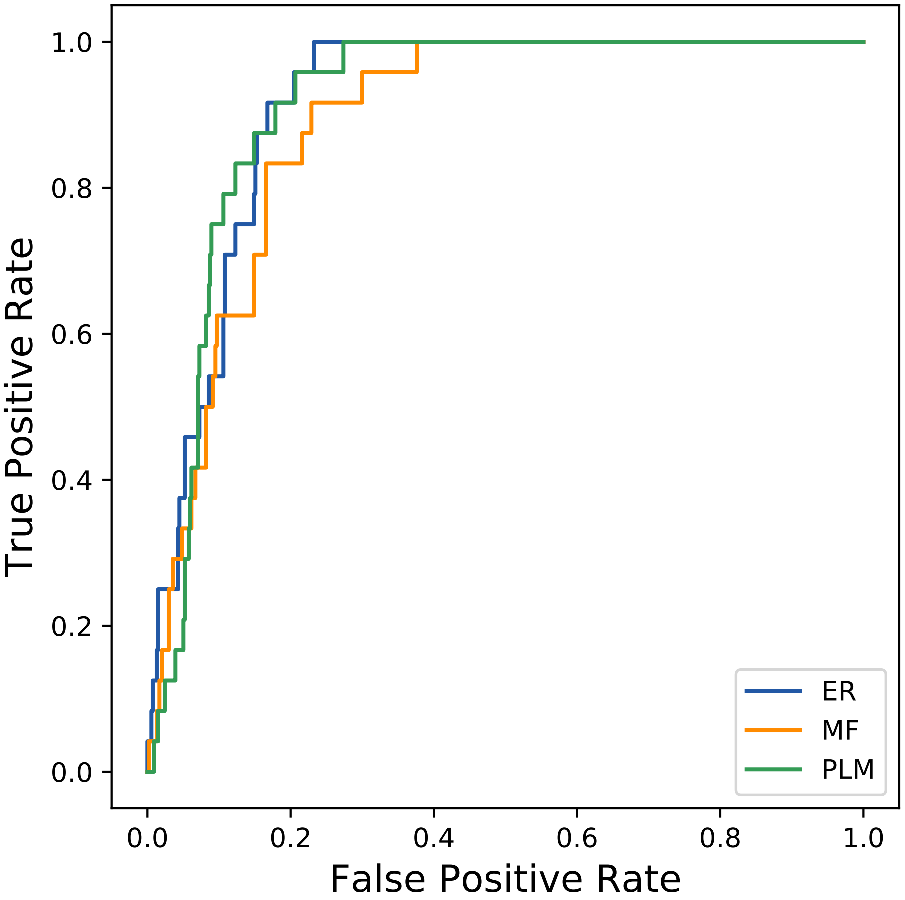

### 20473seq_183col_1jp7_PF01048_plots.pdf

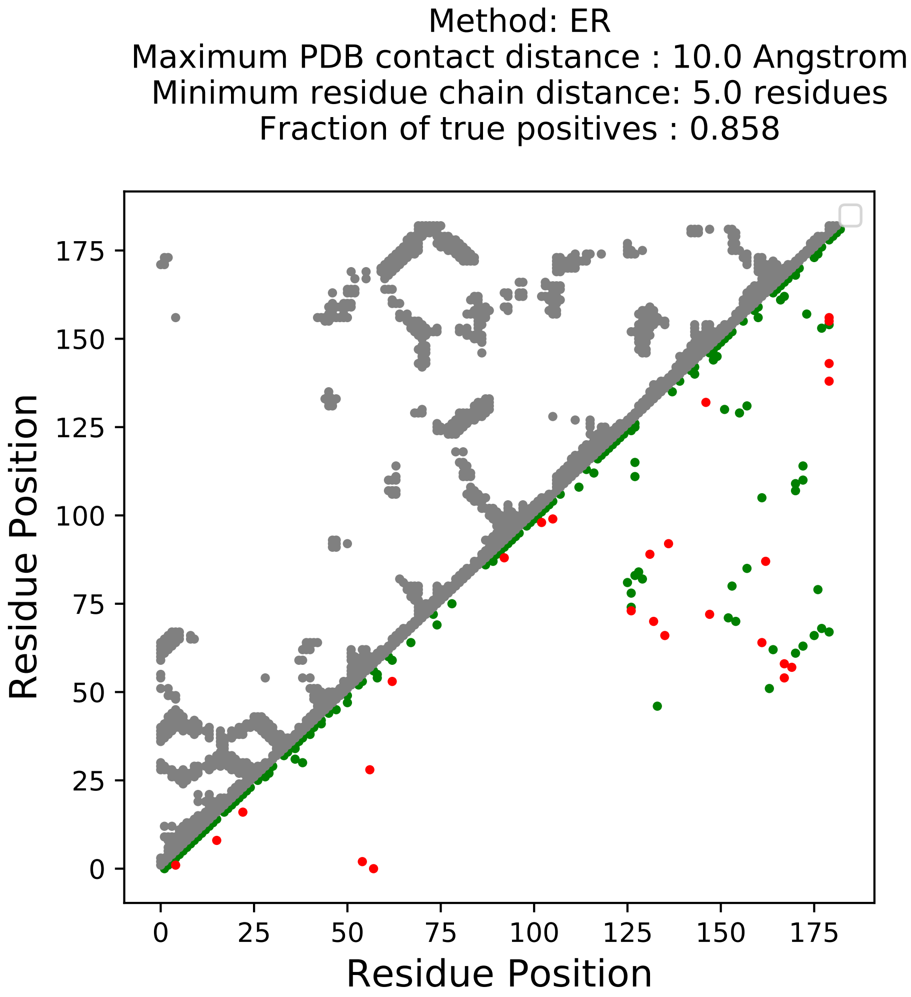

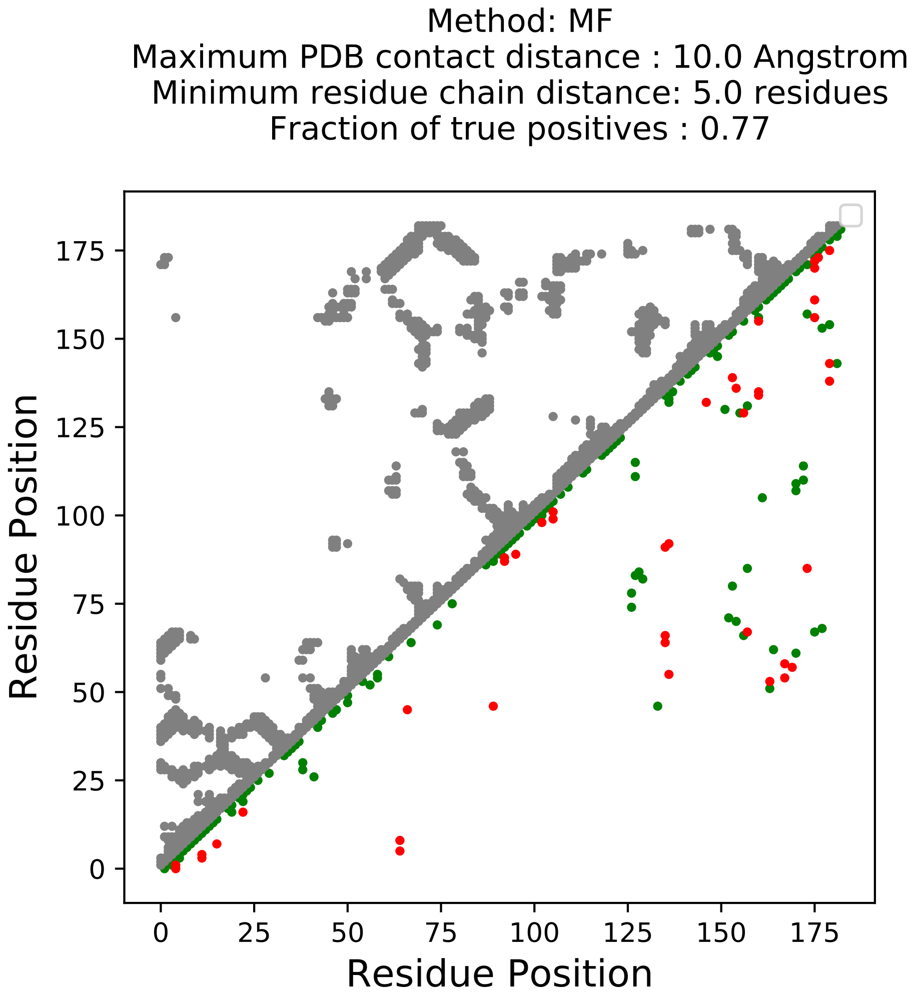

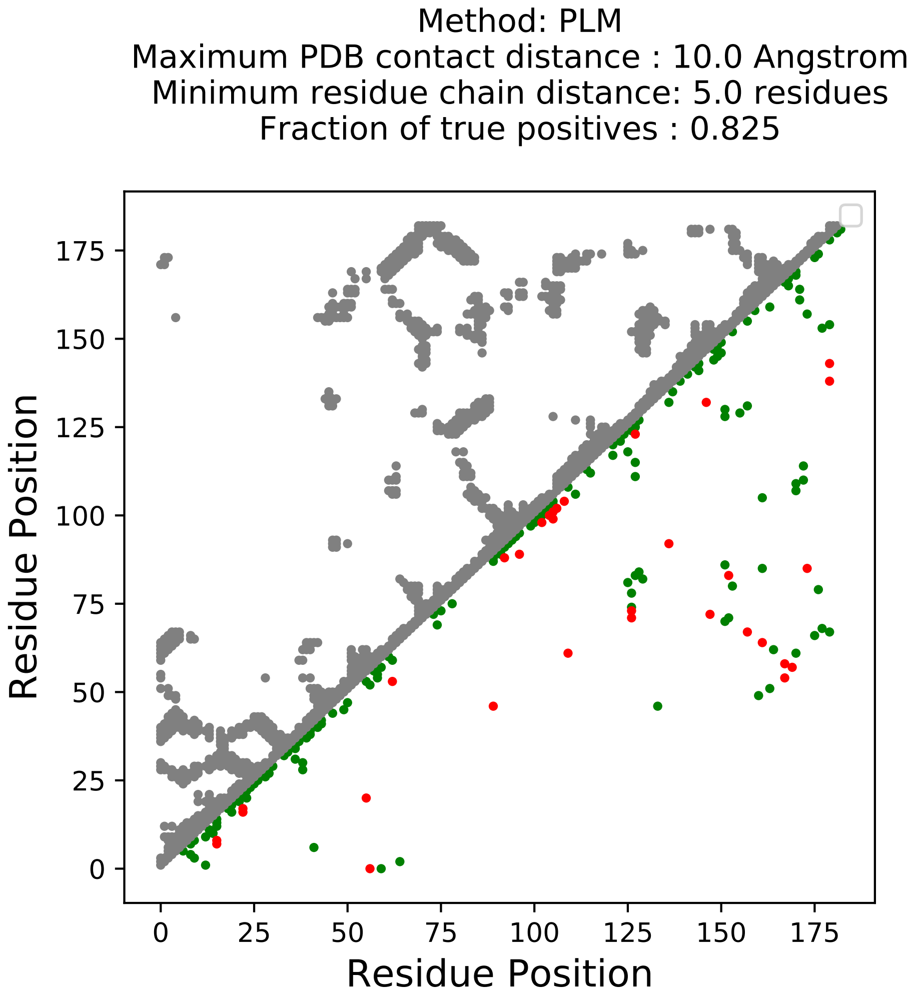

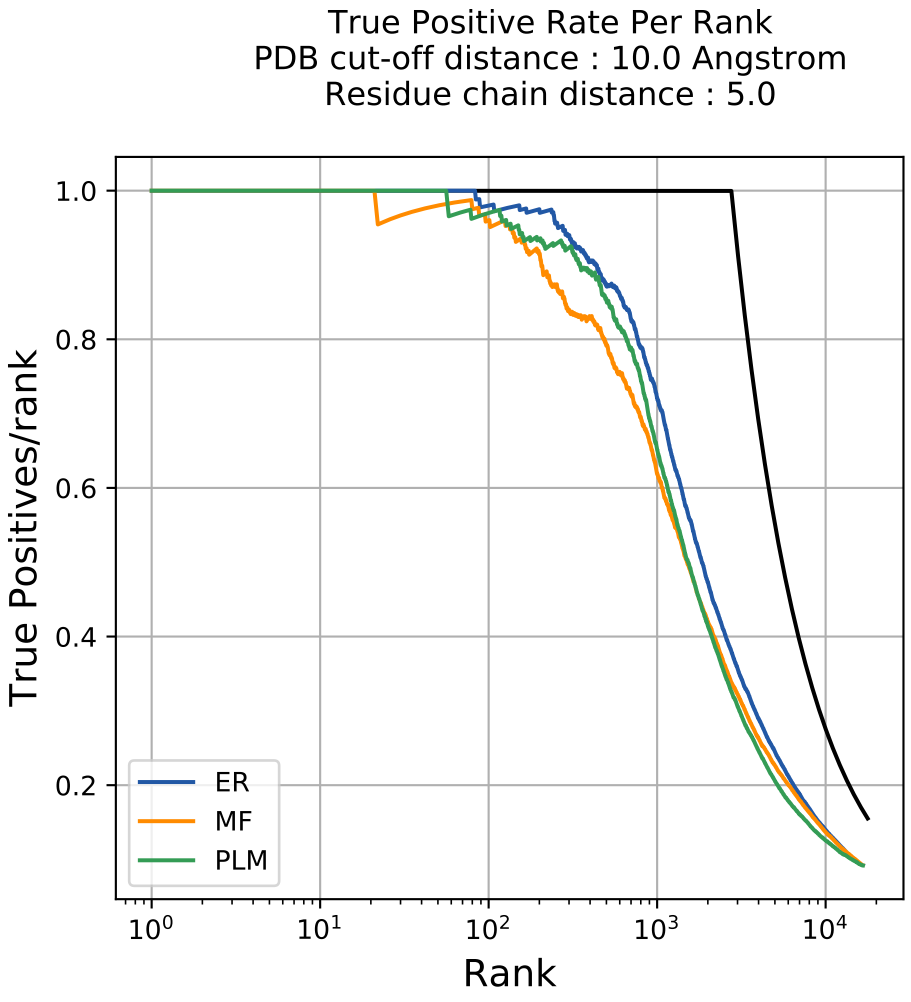

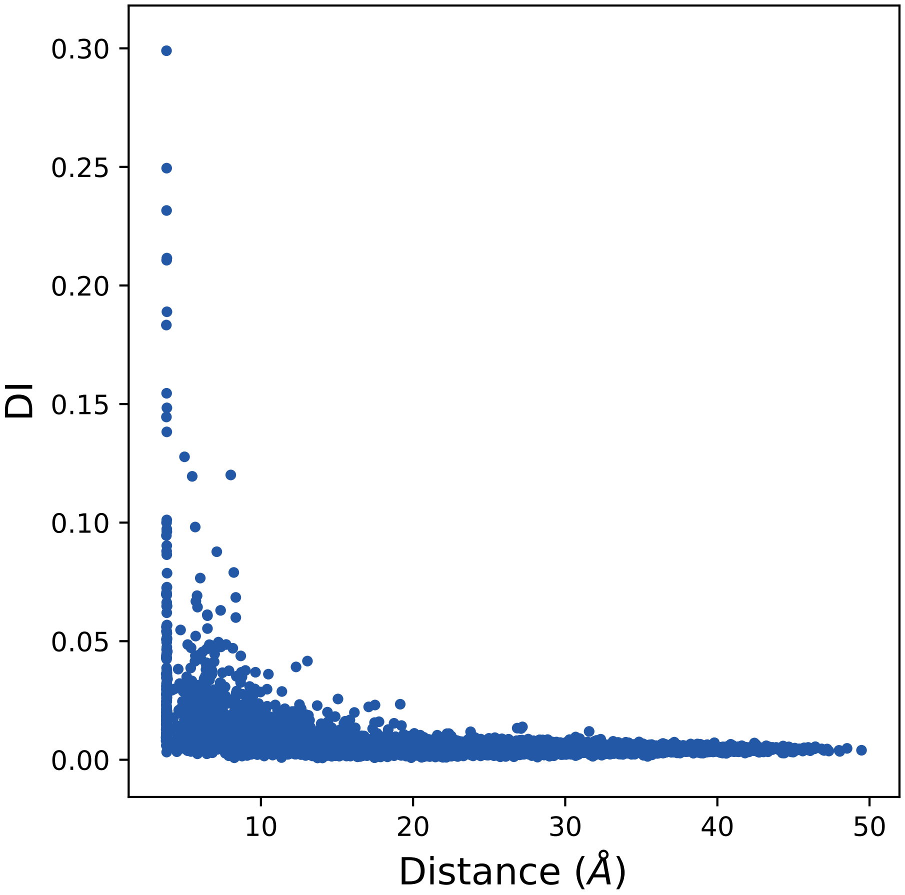

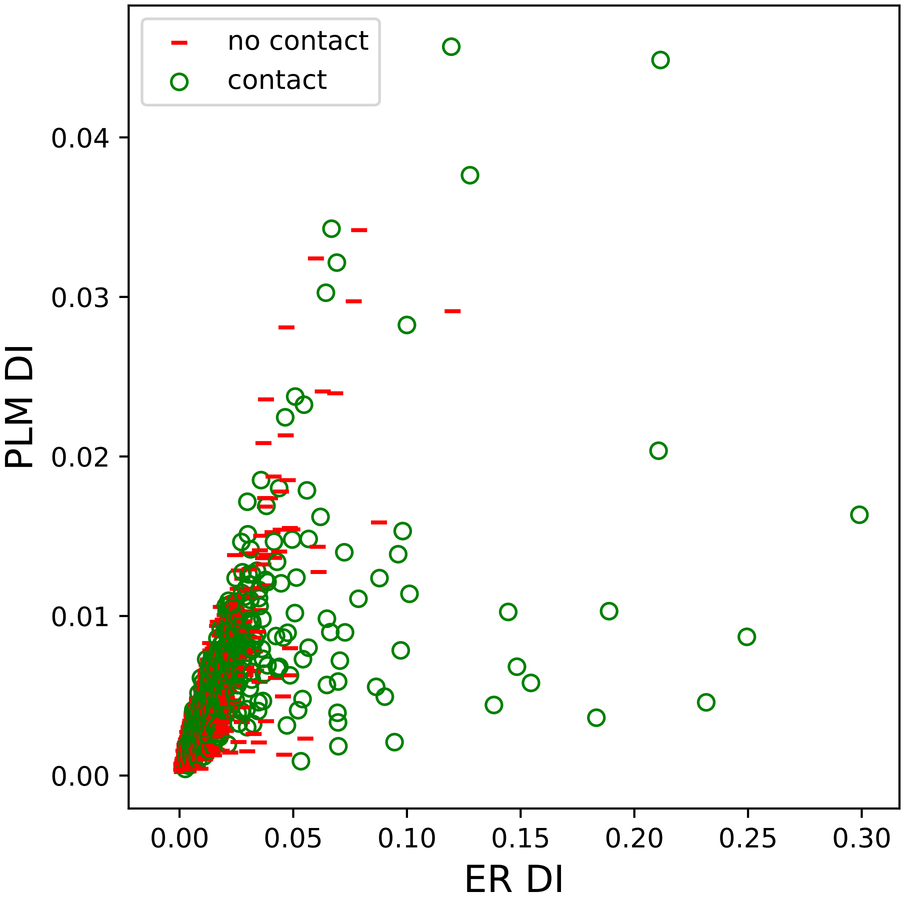

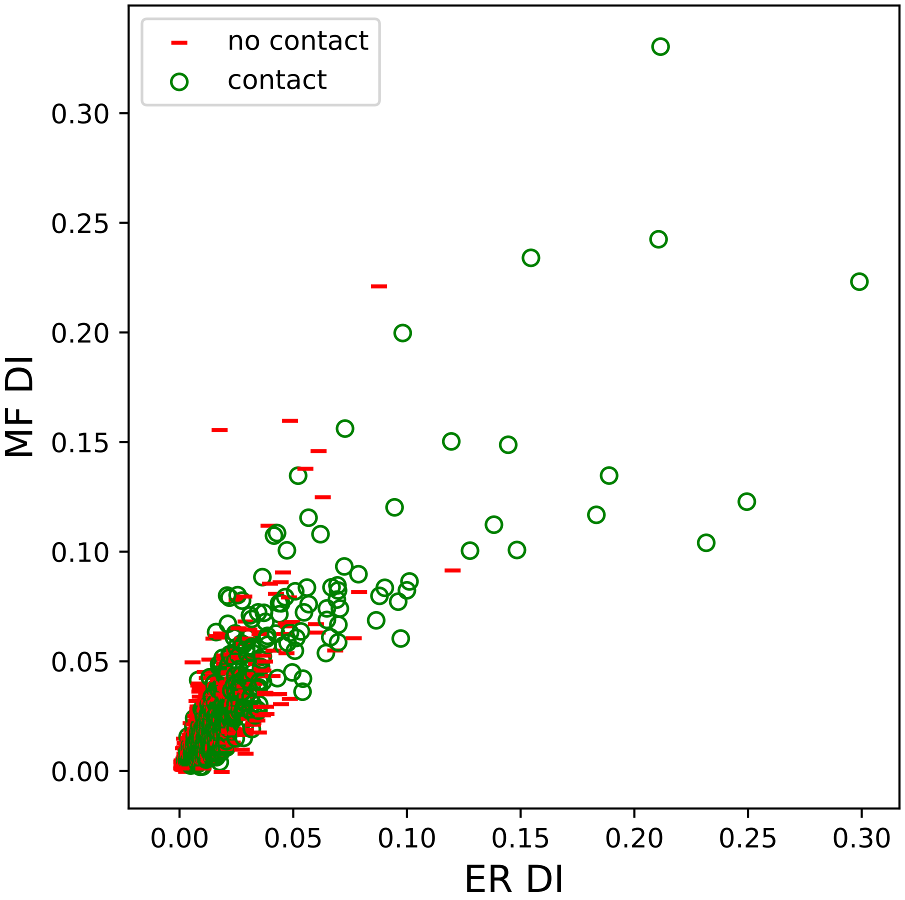

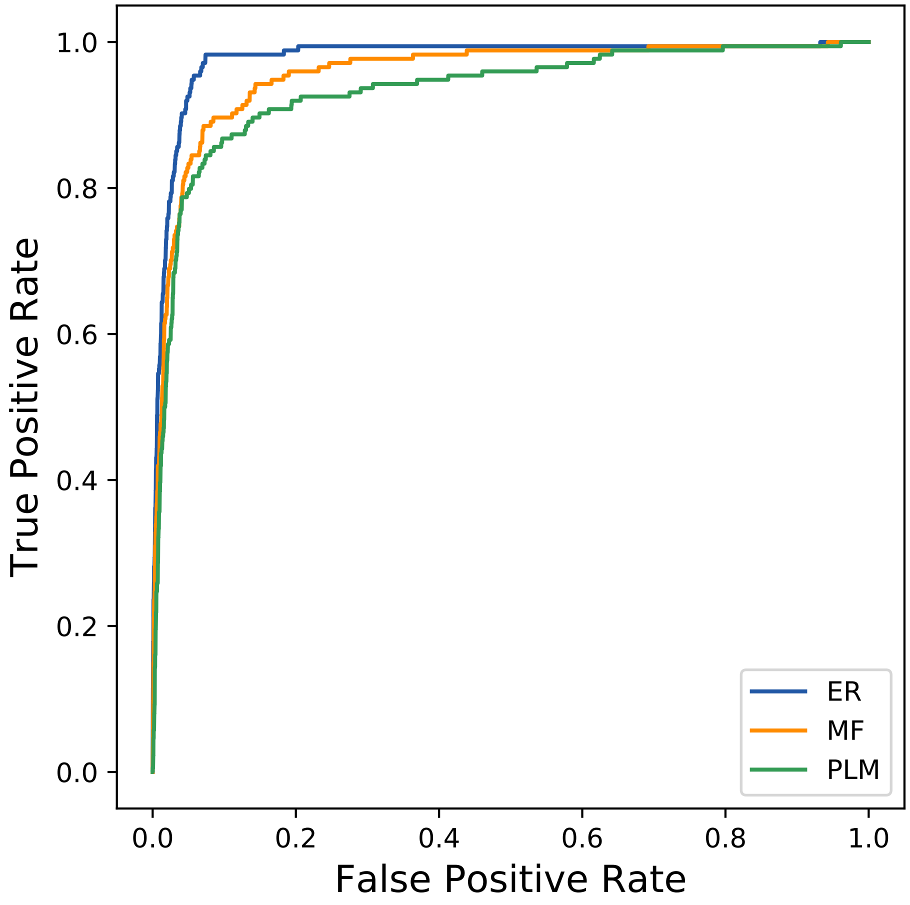

### 21586seq_59col_2i5r_PF01751_plots.pdf

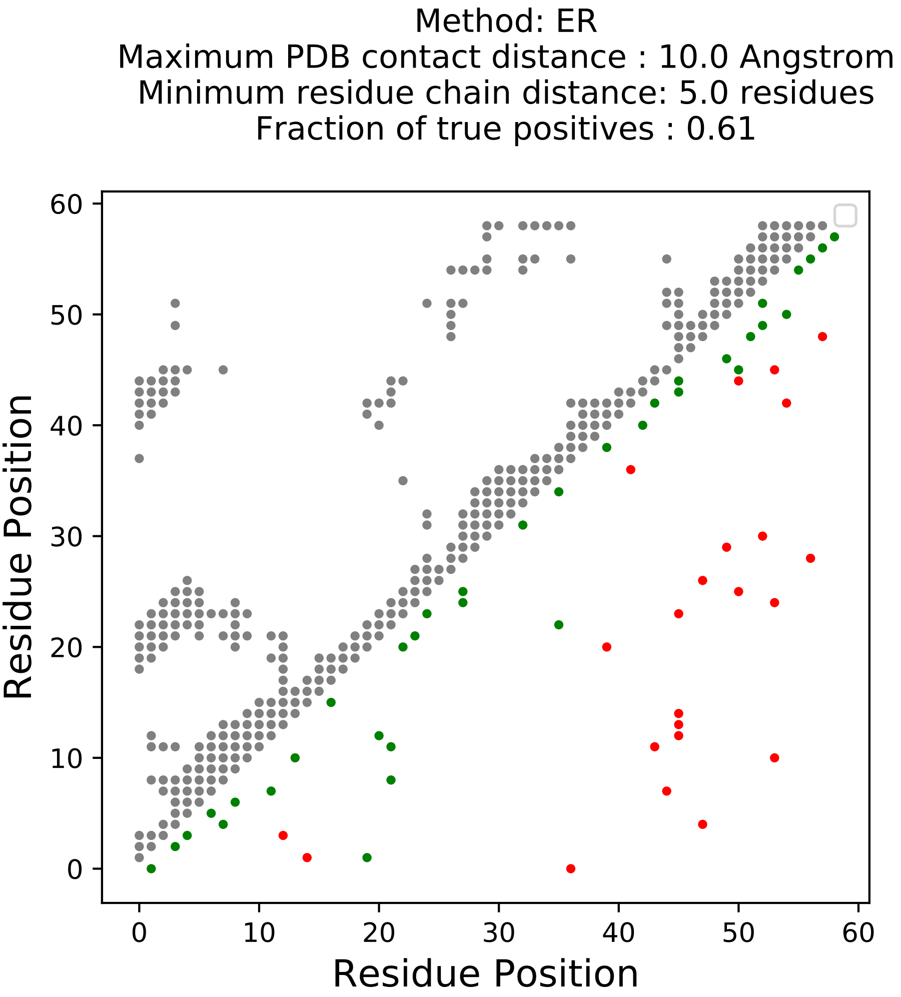

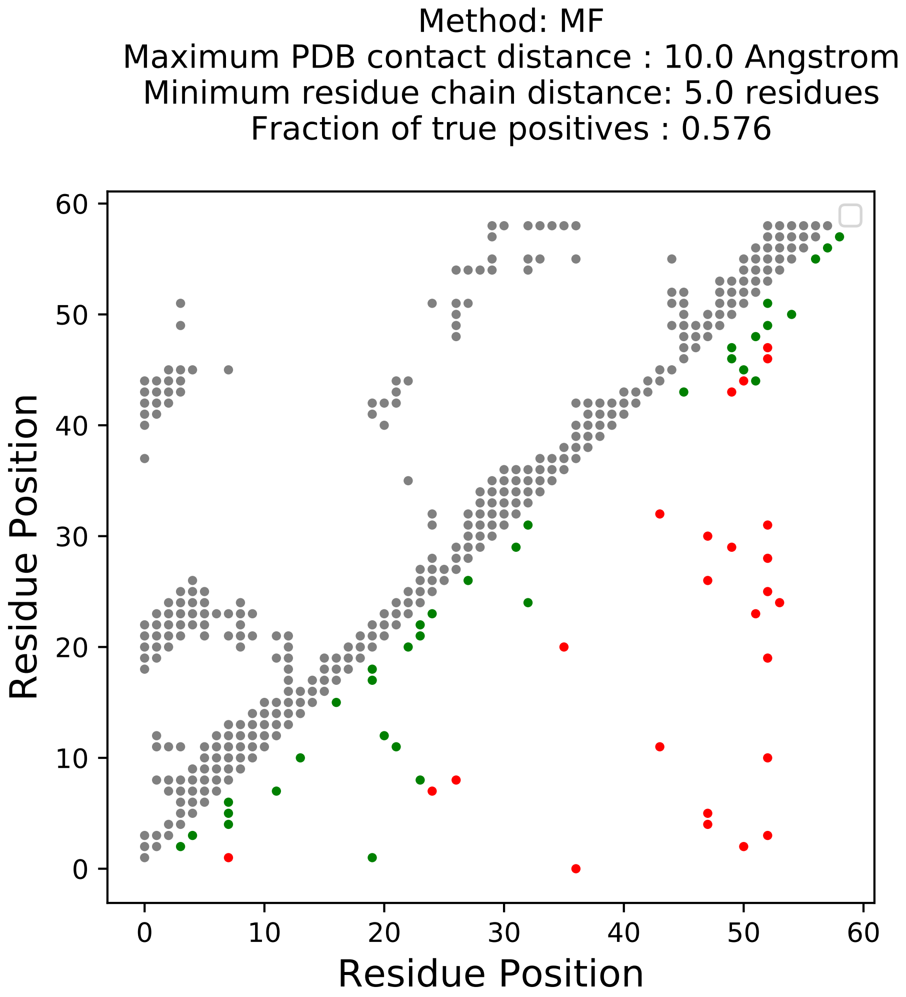

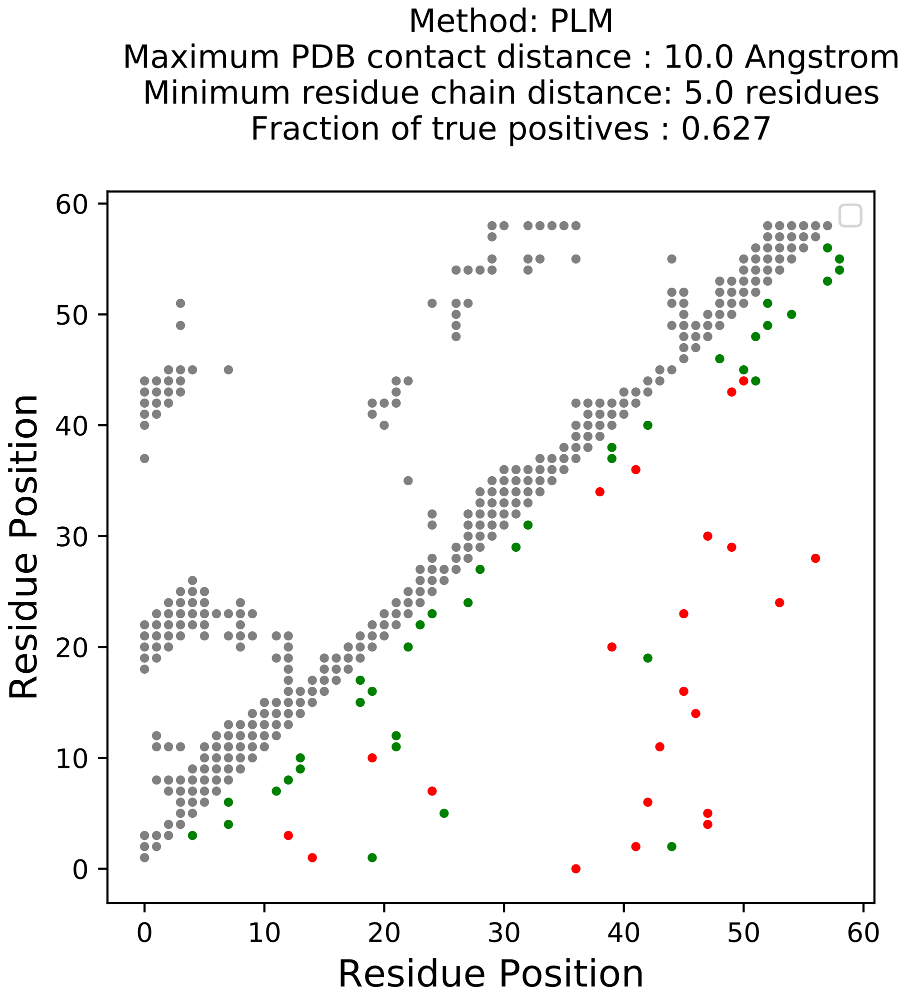

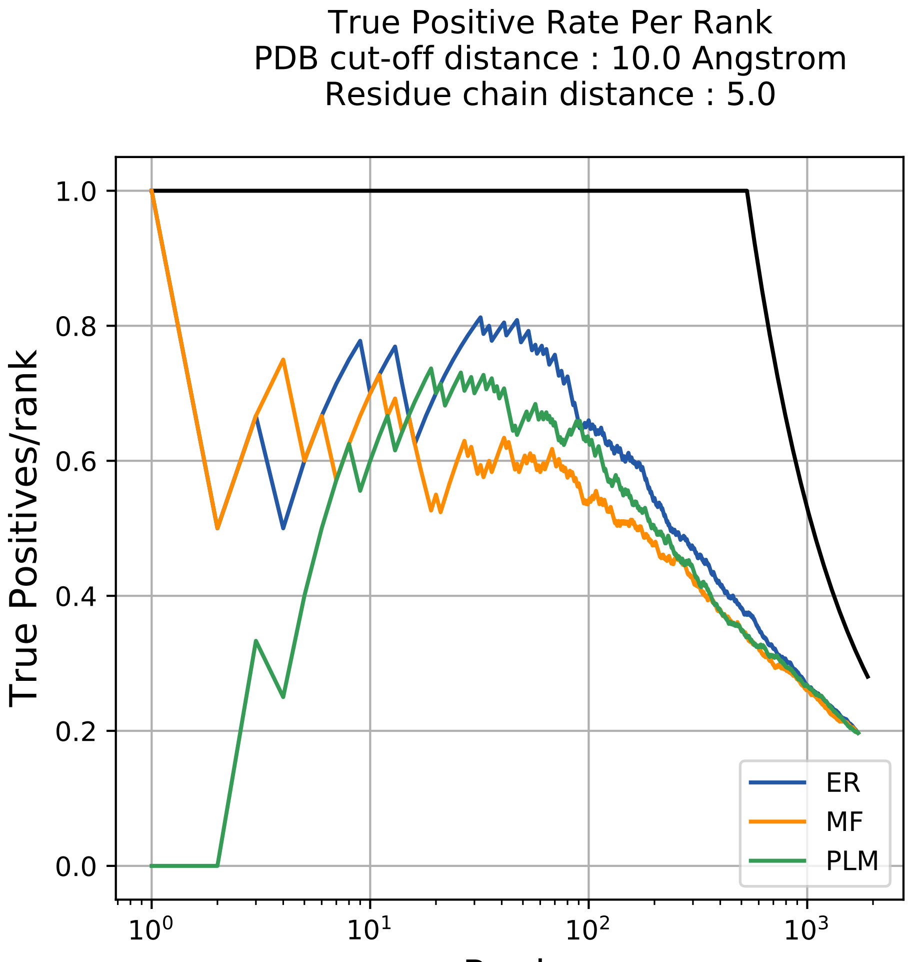

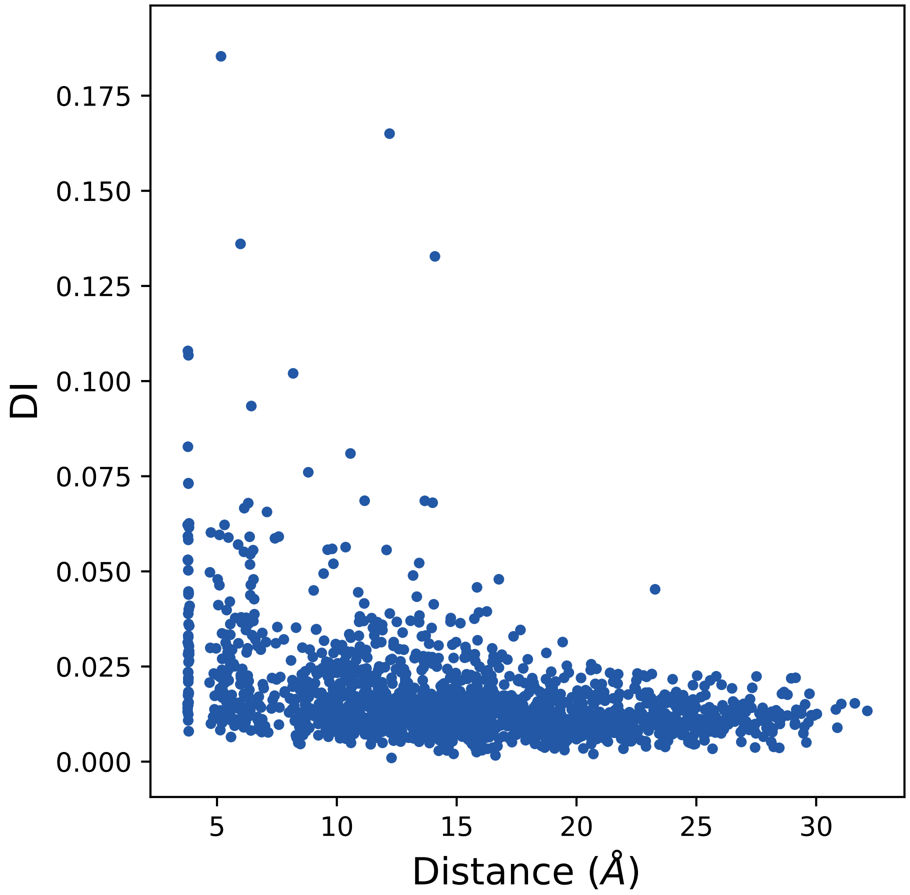

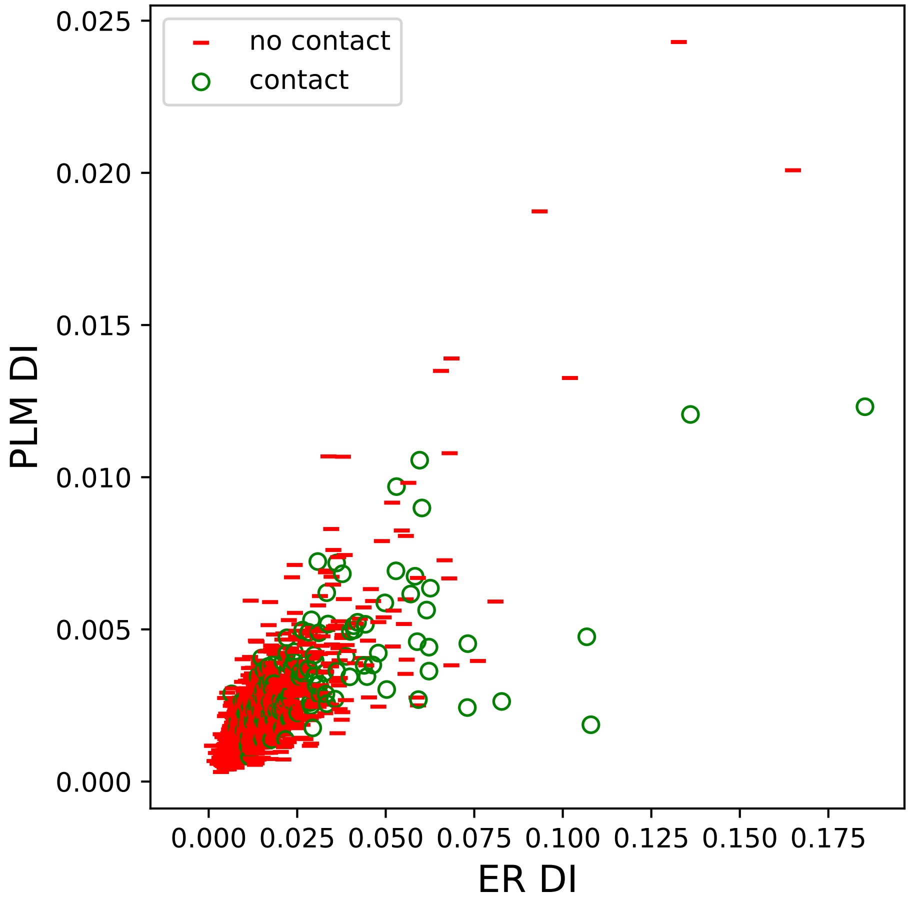

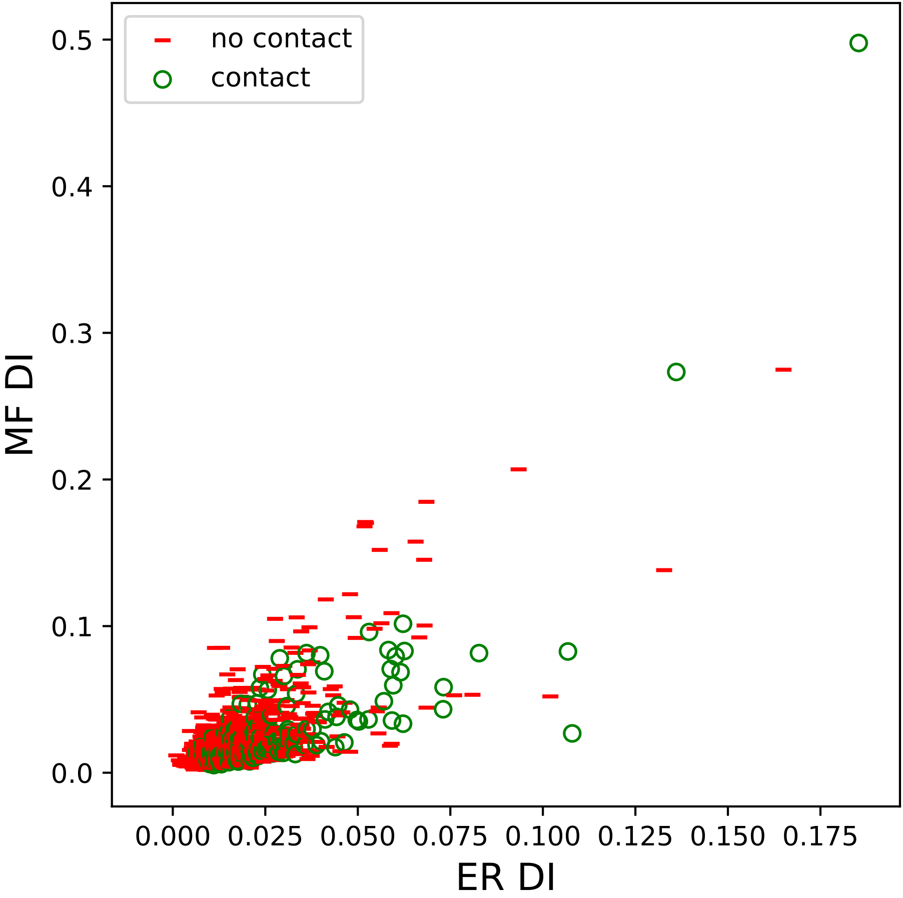

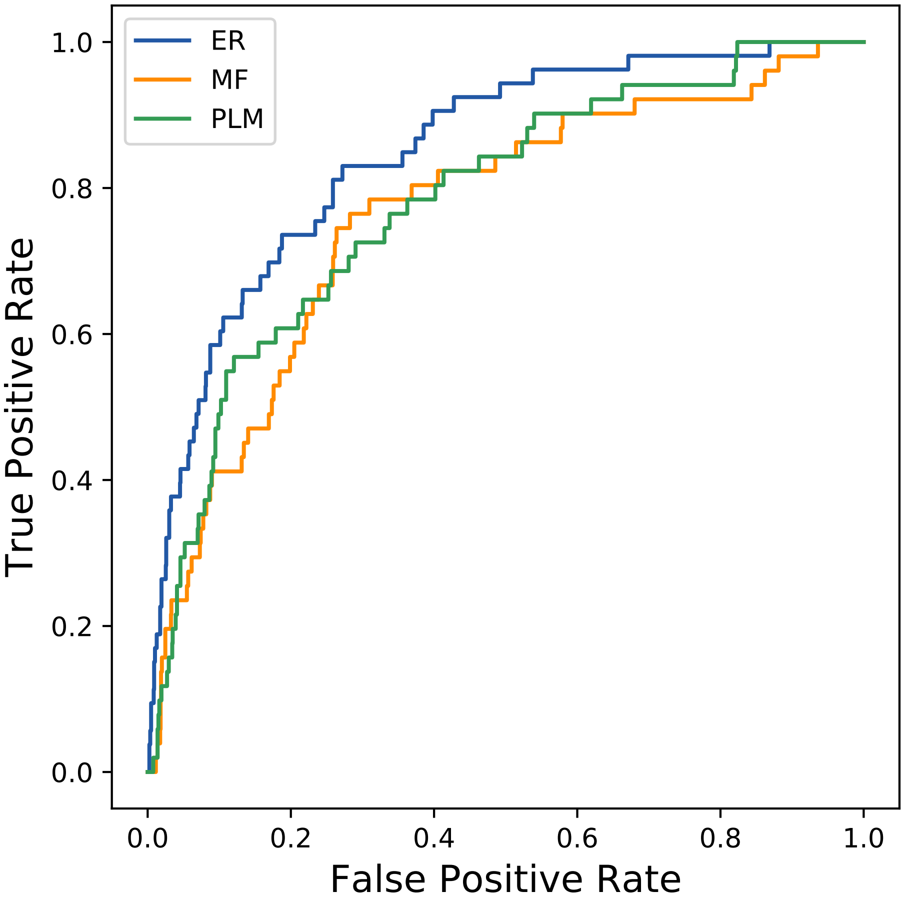

### 22601seq_69col_2imm_PF07686_plots.pdf

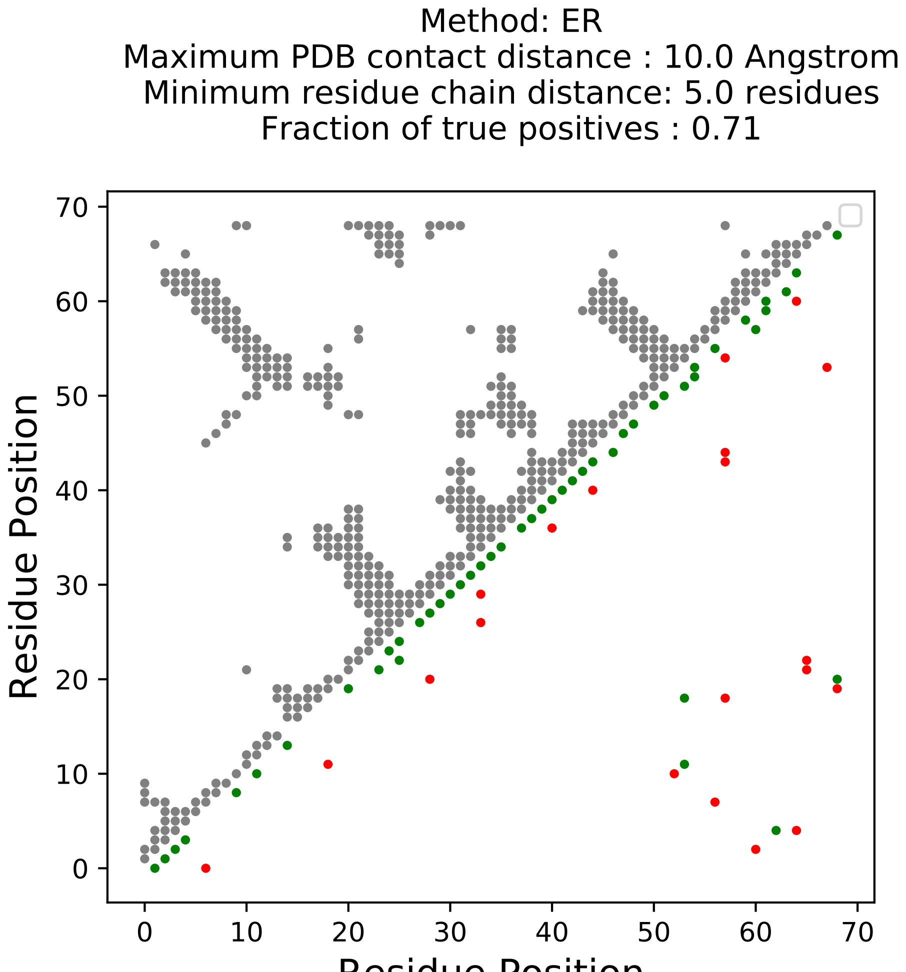

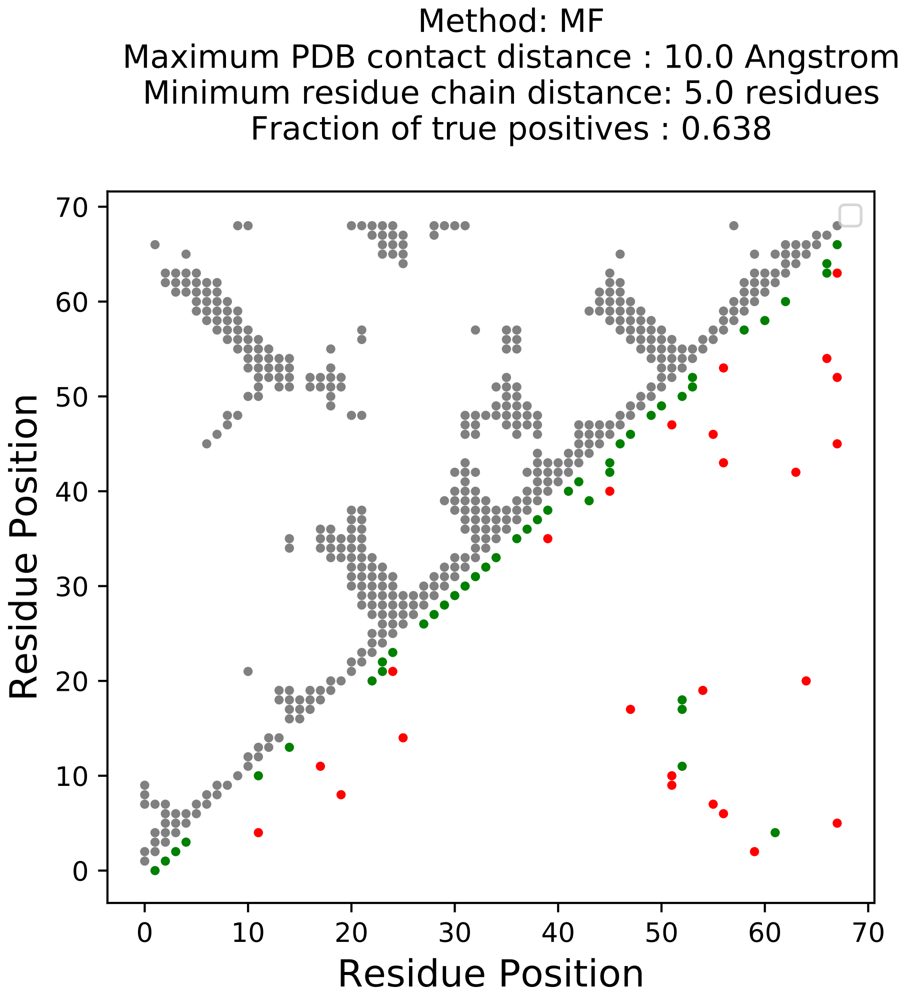

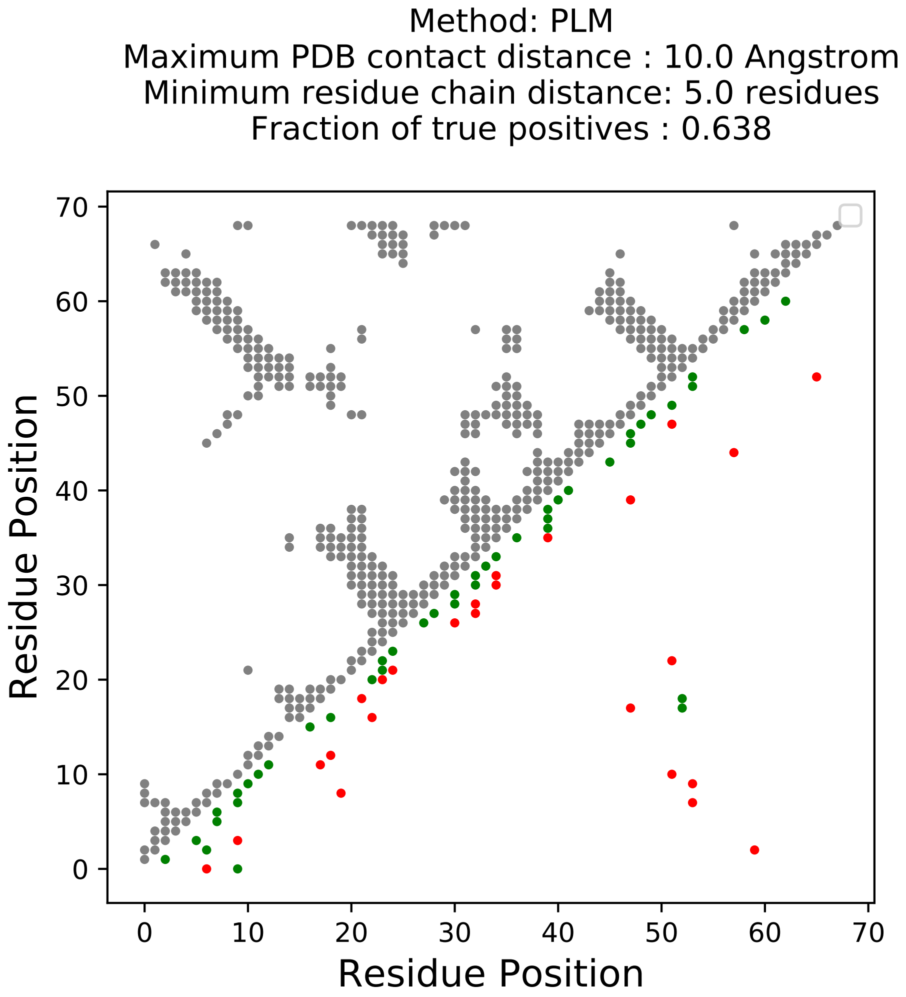

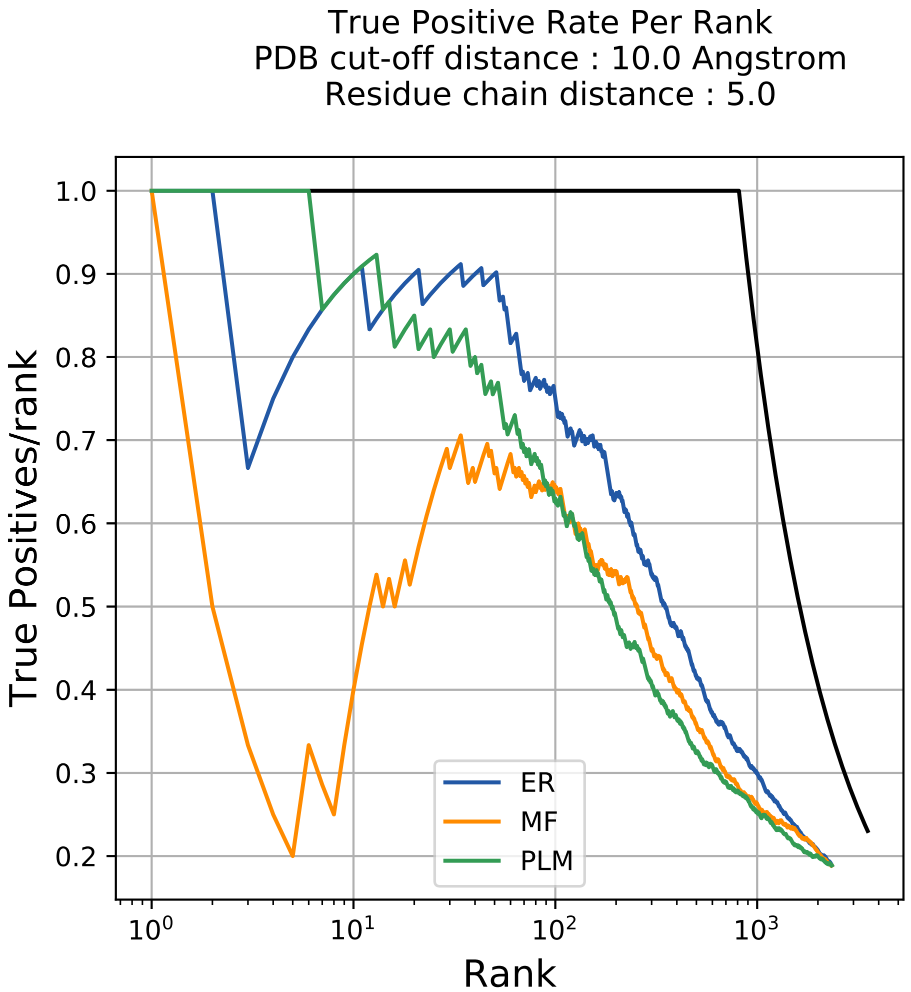

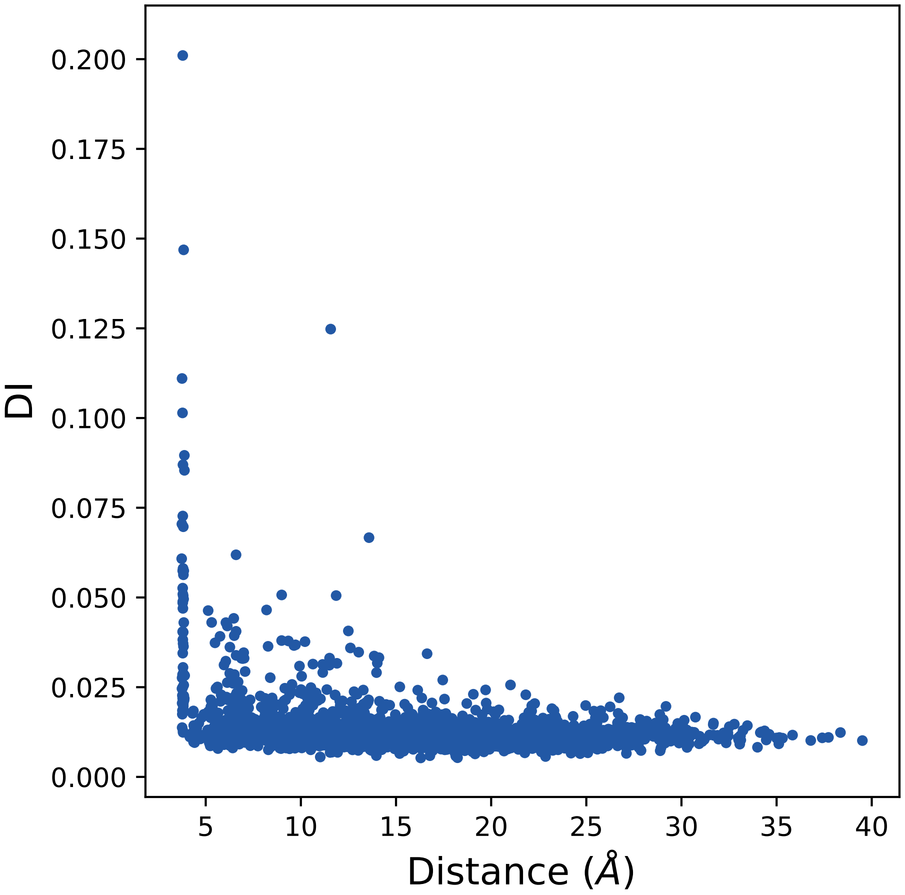

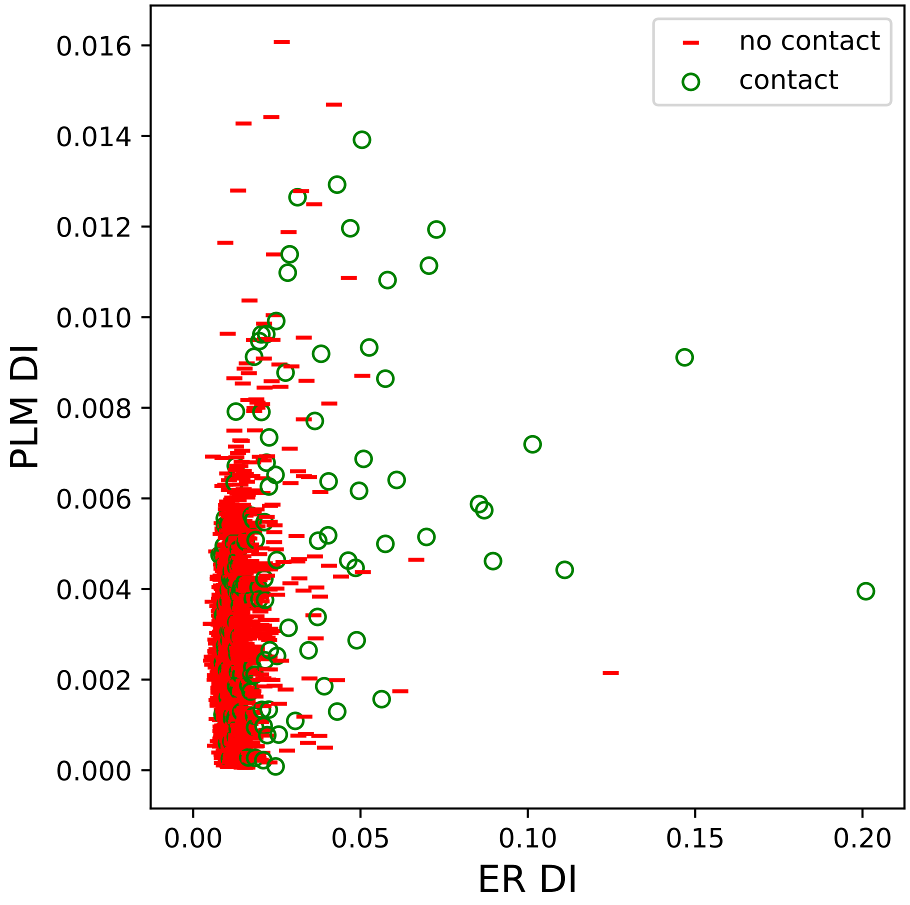

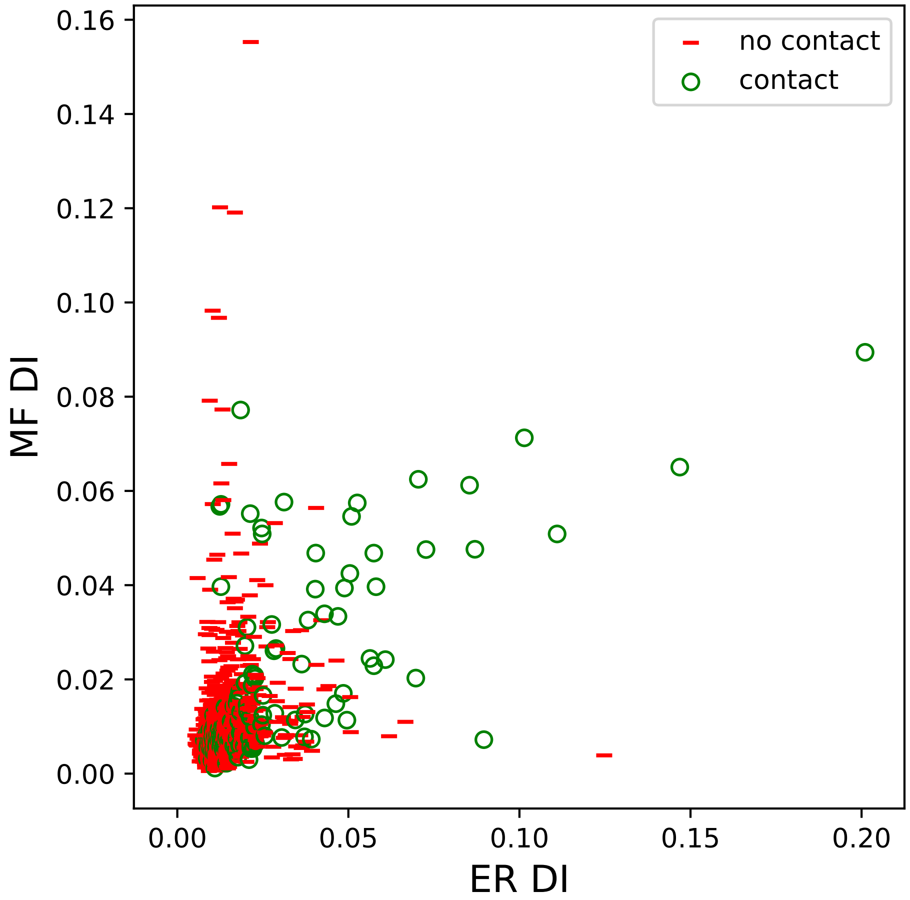

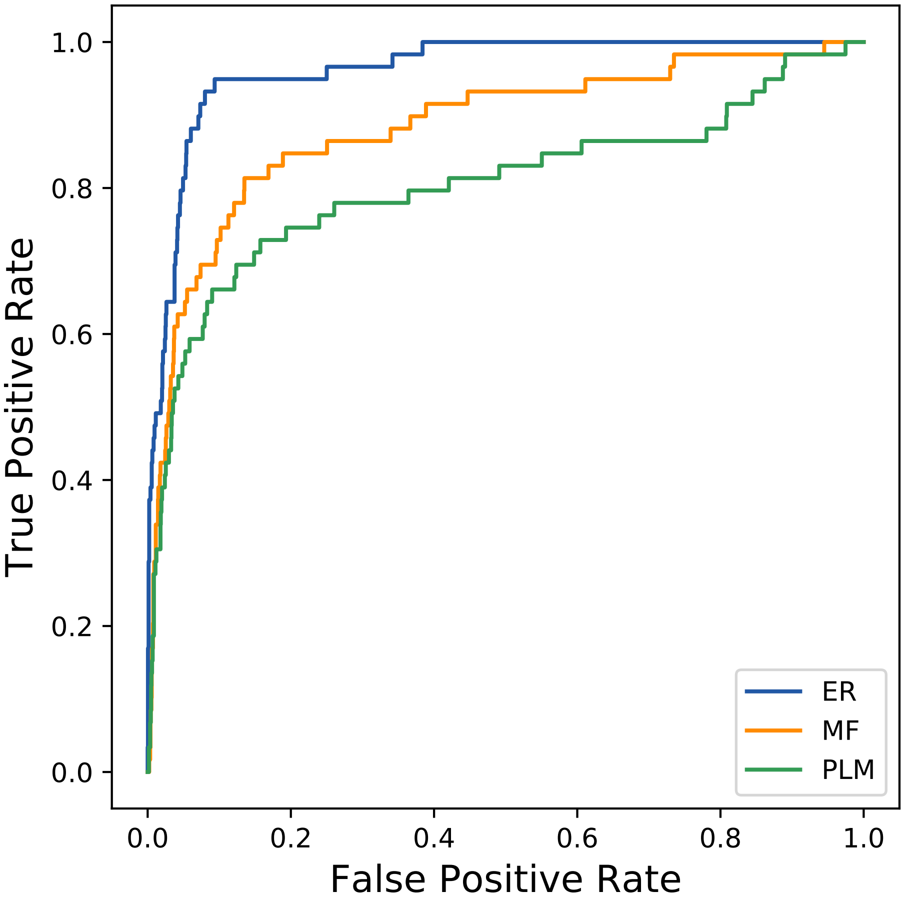

### 24293seq_183col_3aim_PF07859_plots.pdf

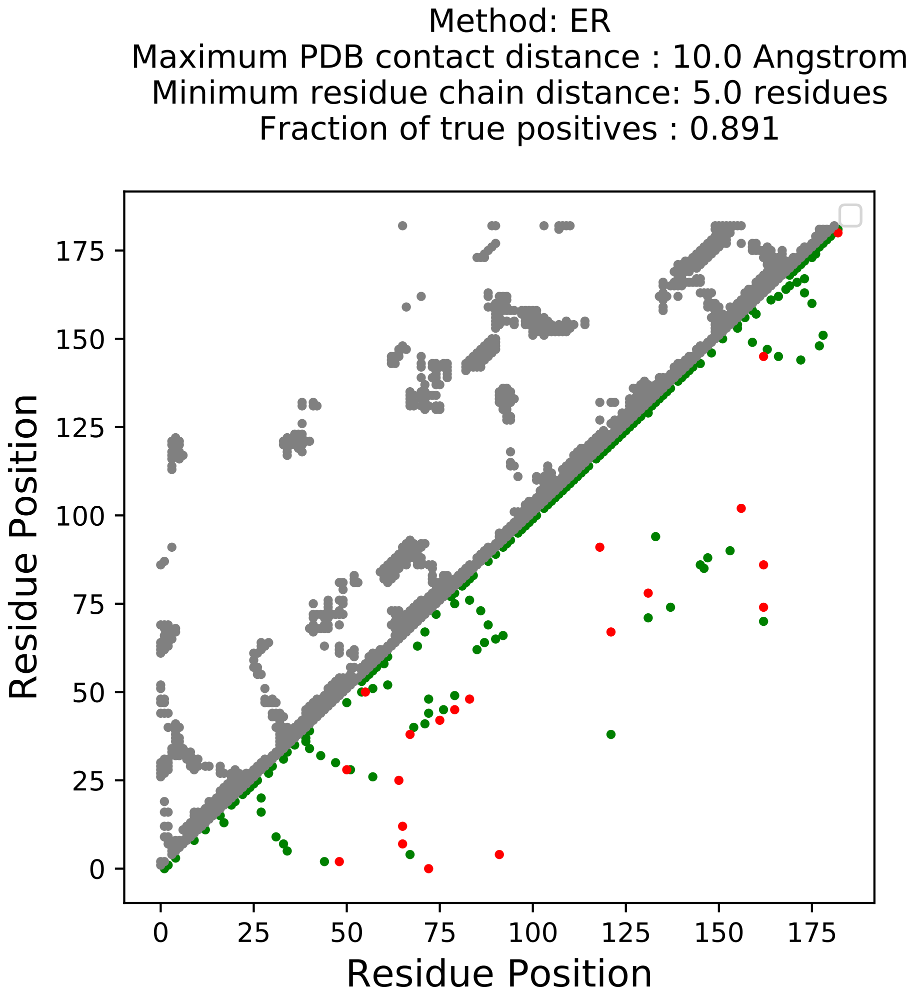

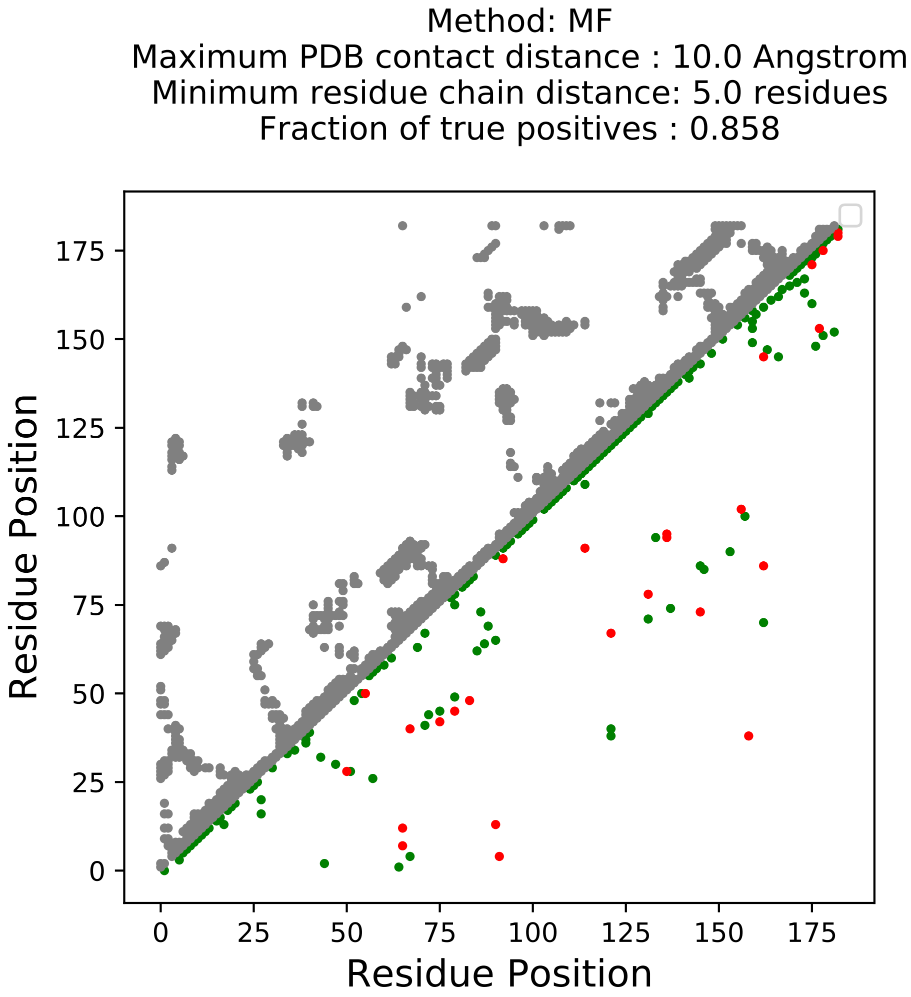

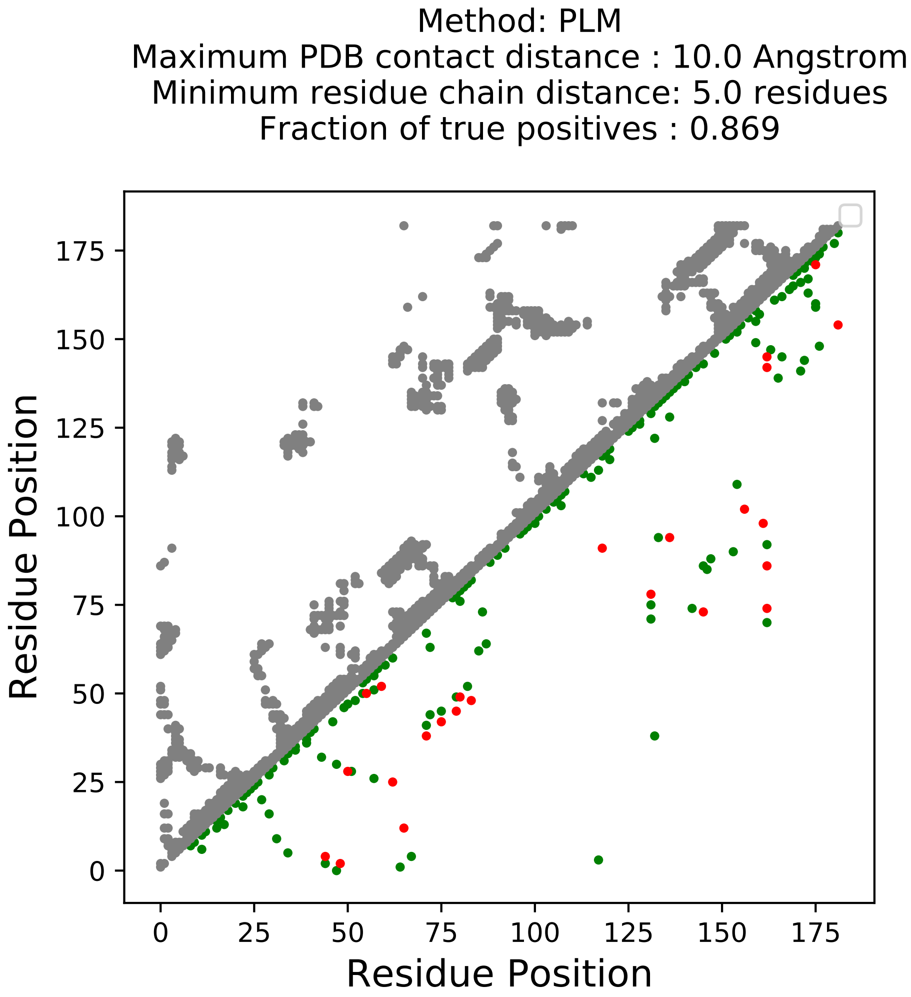

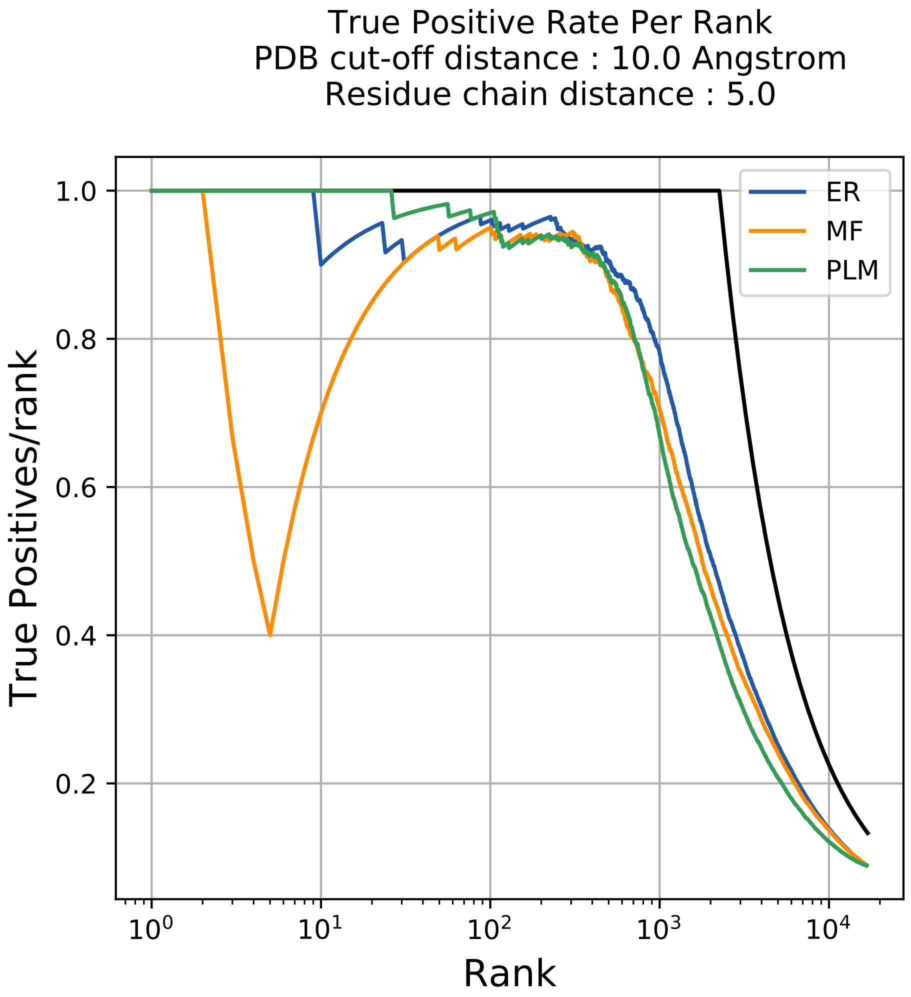

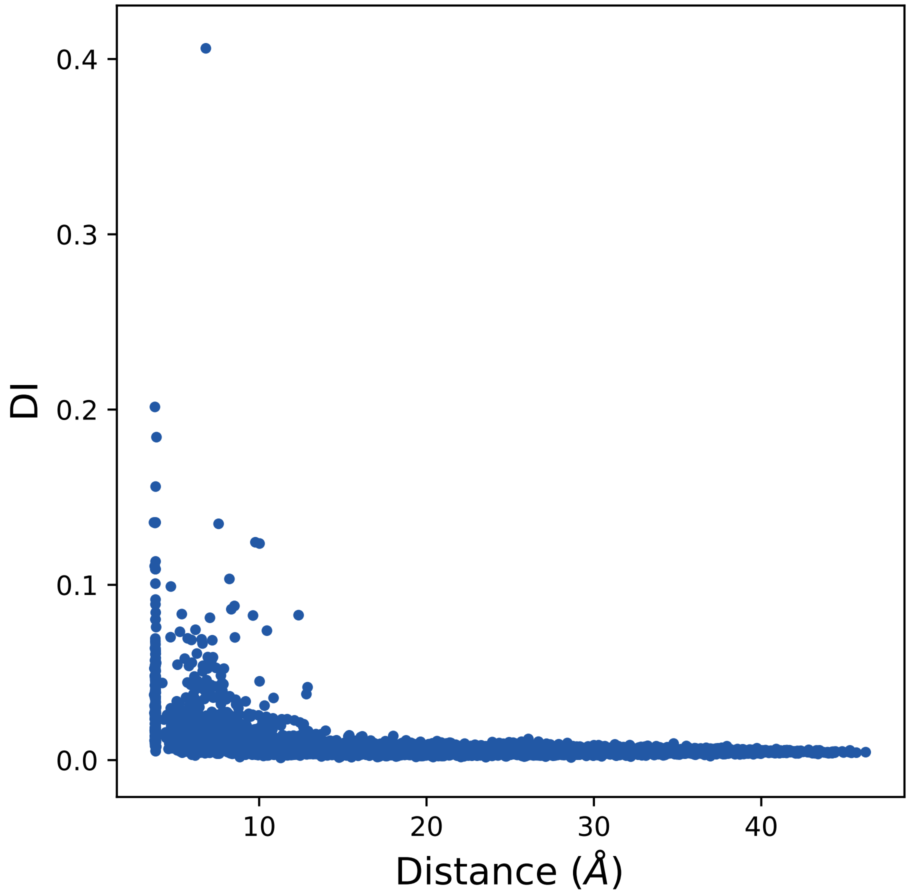

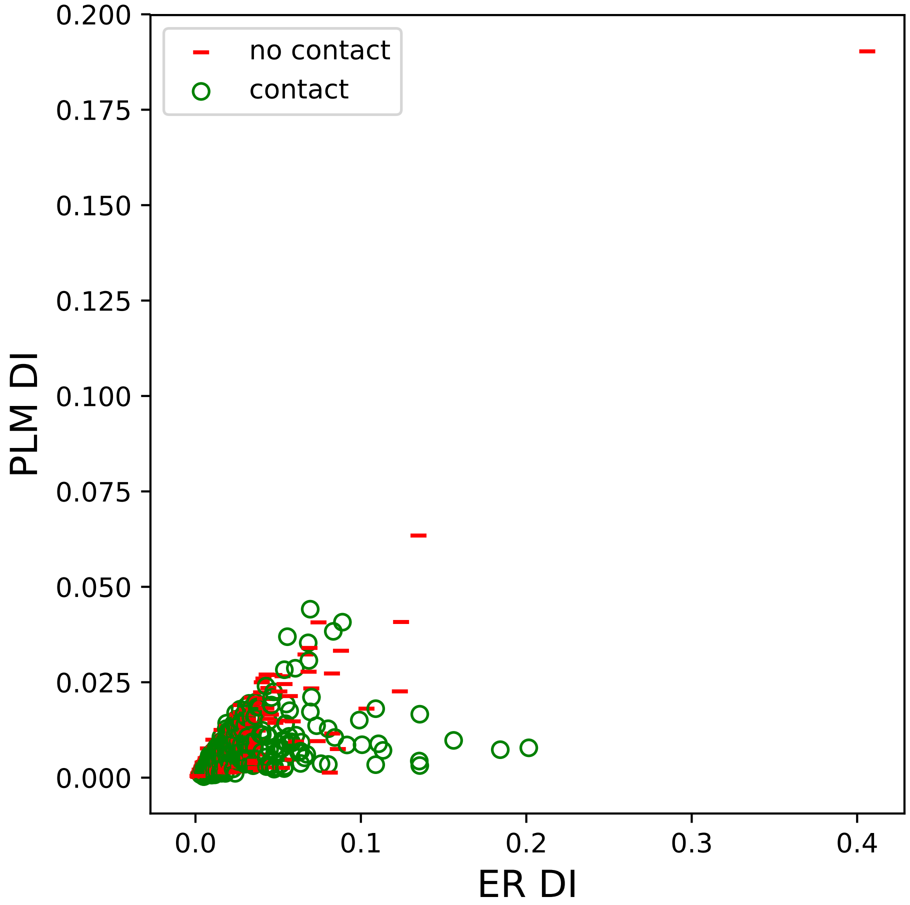

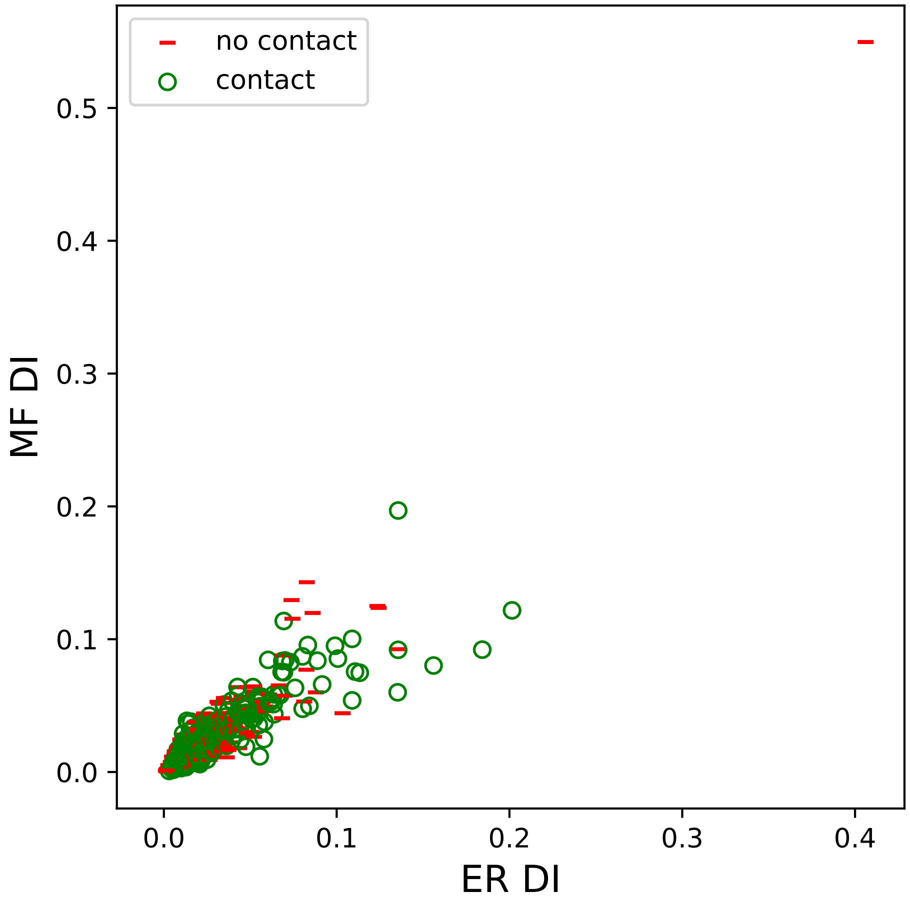

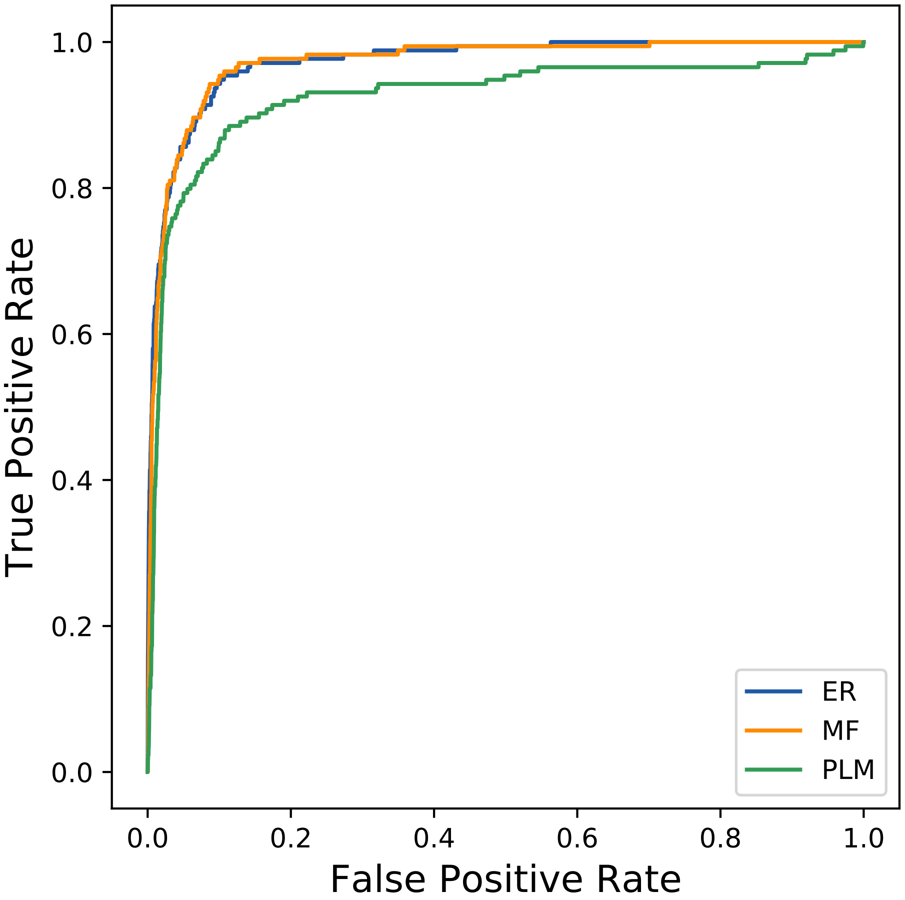

### 25204seq_37col_1ity_PF00249_plots.pdf

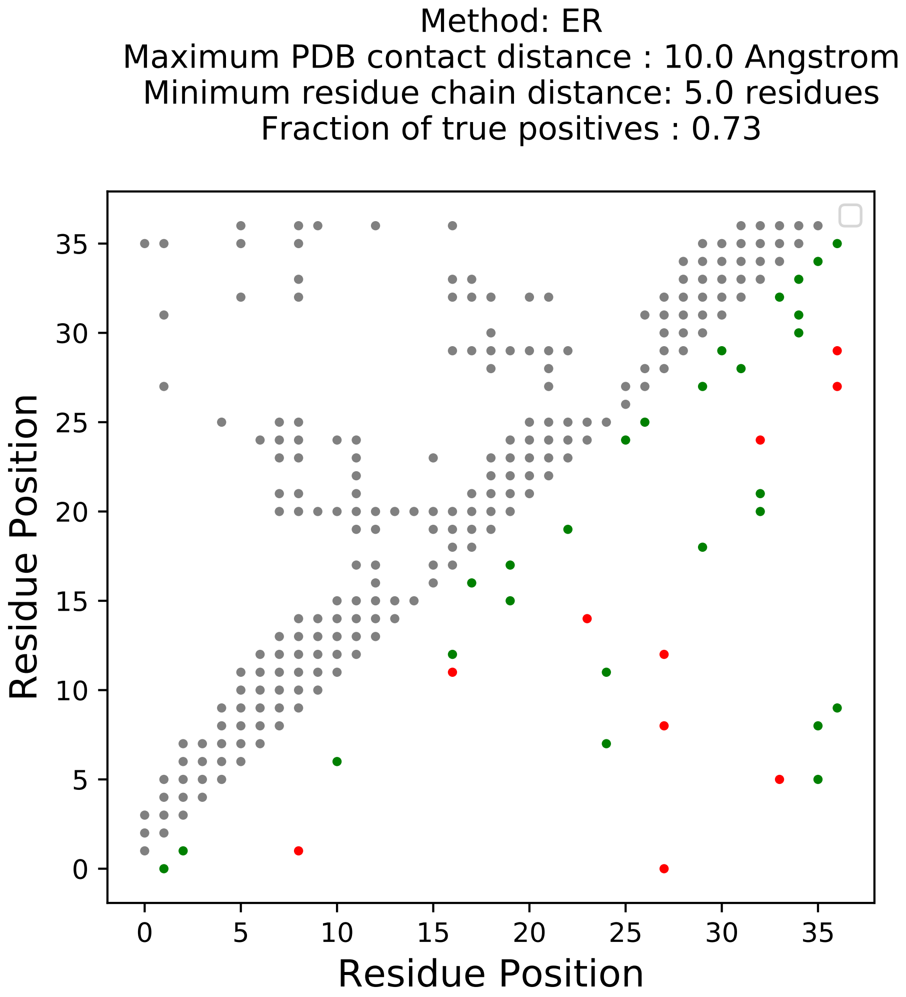

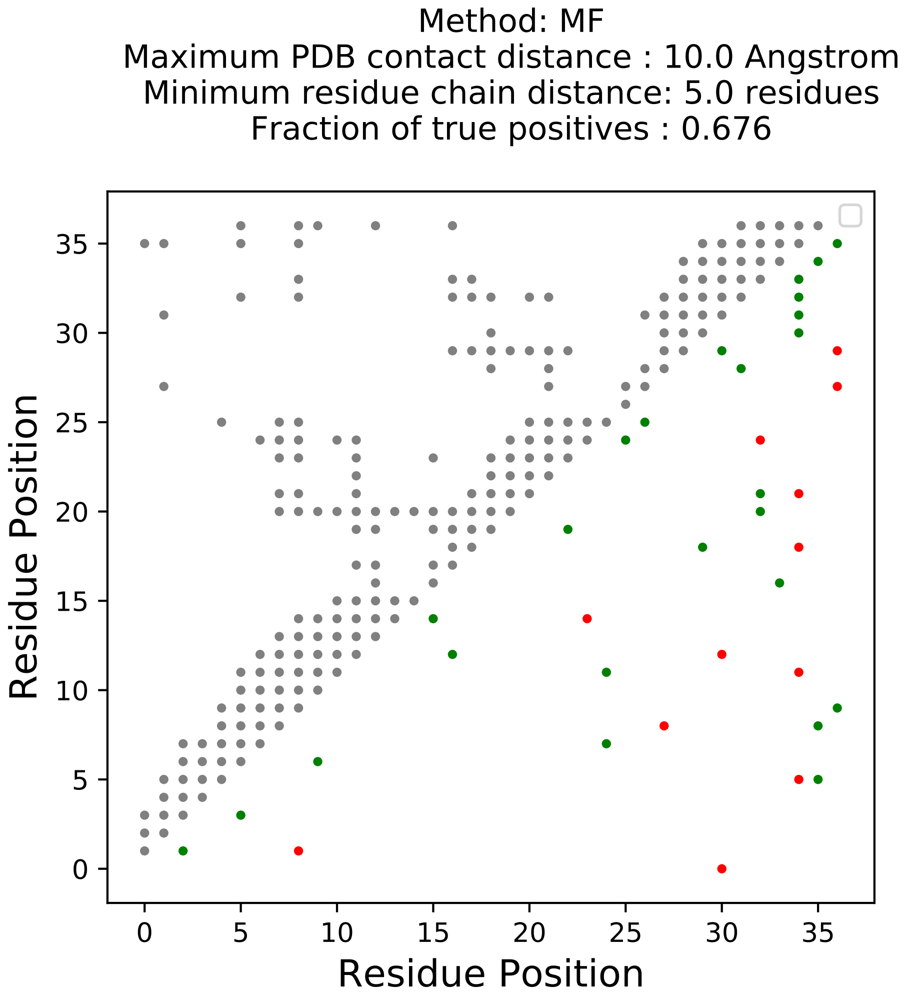

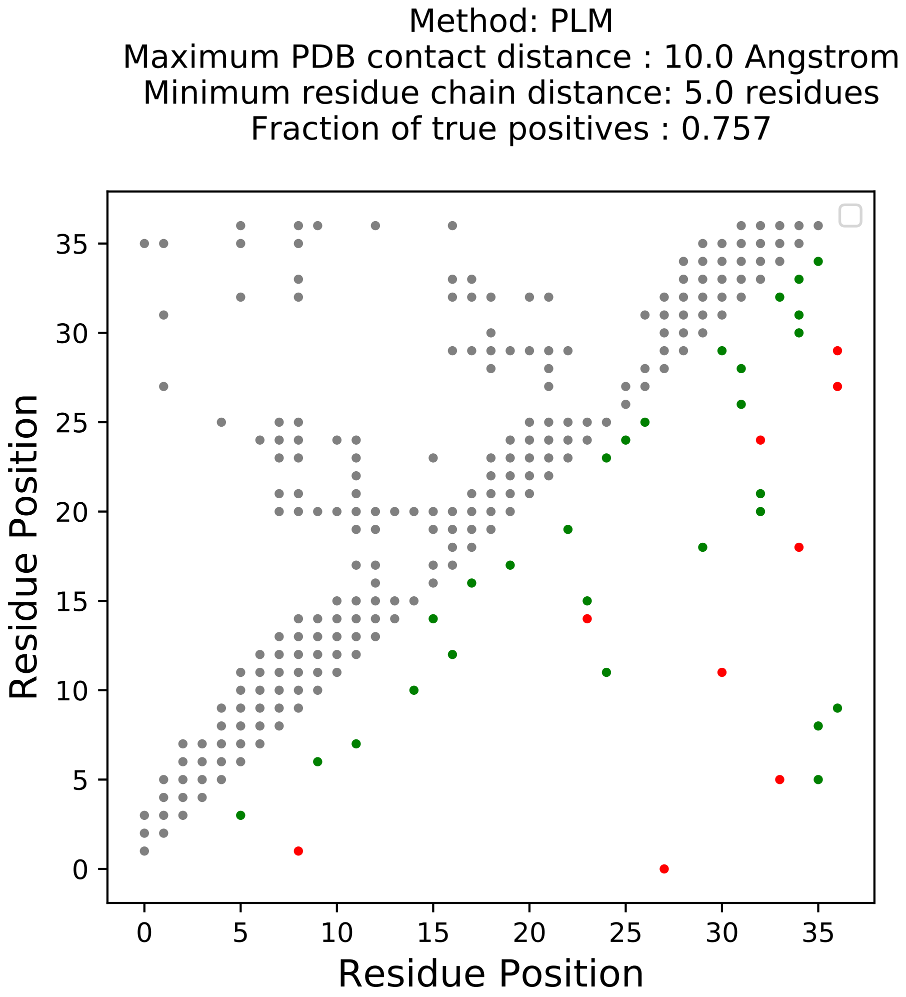

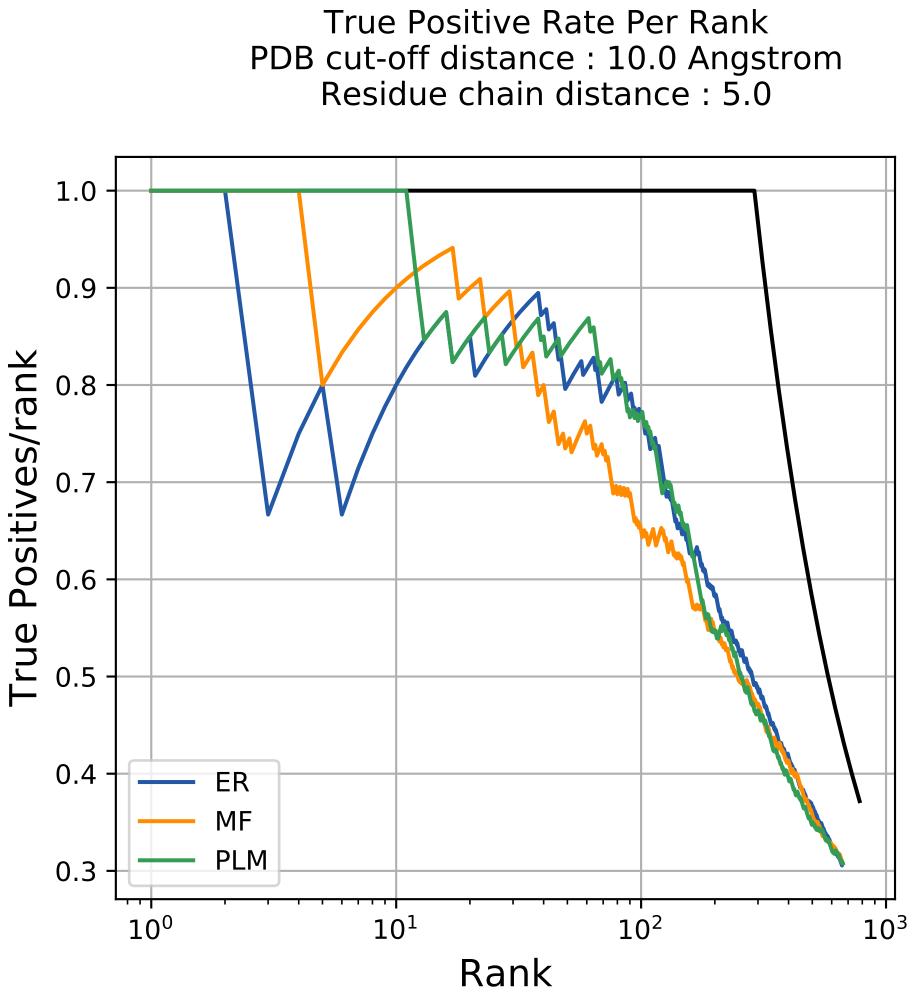

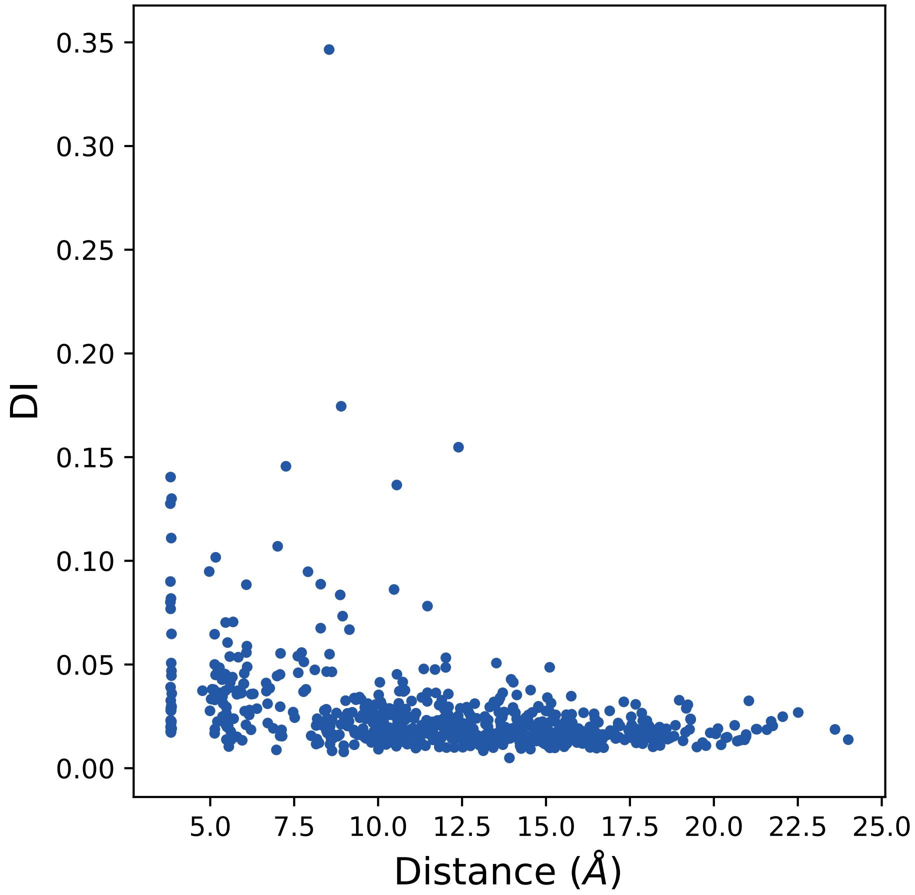

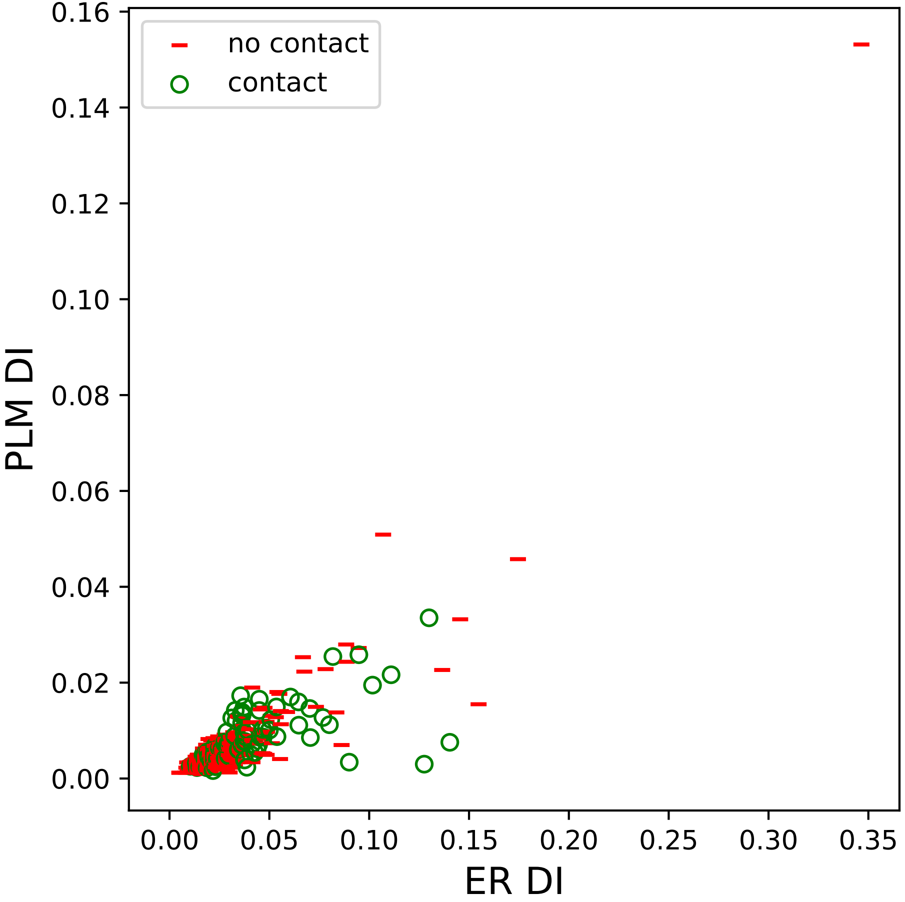

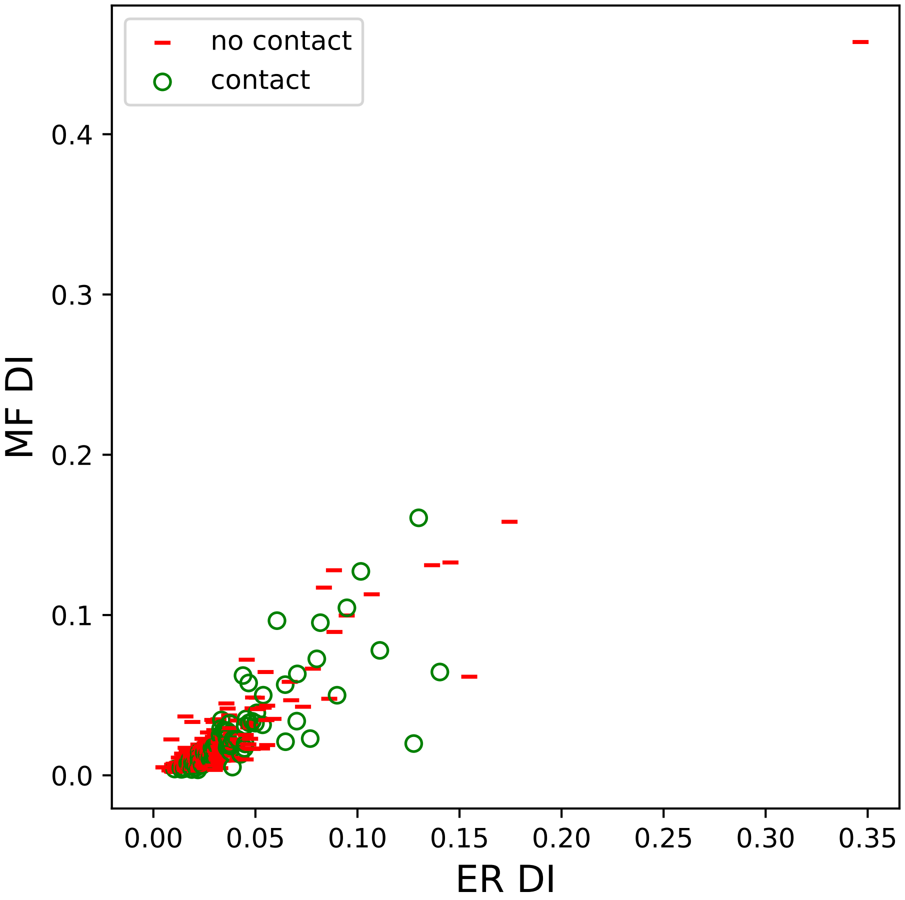

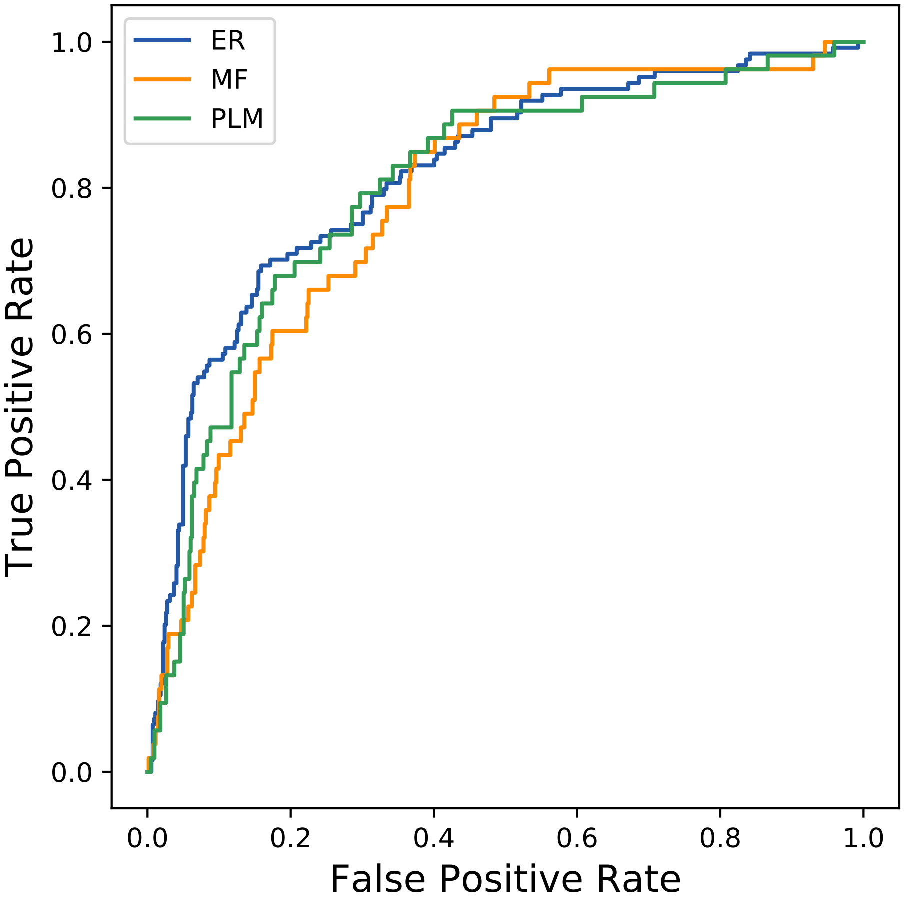

### 30884seq_92col_6izk_PF13185_plots.pdf

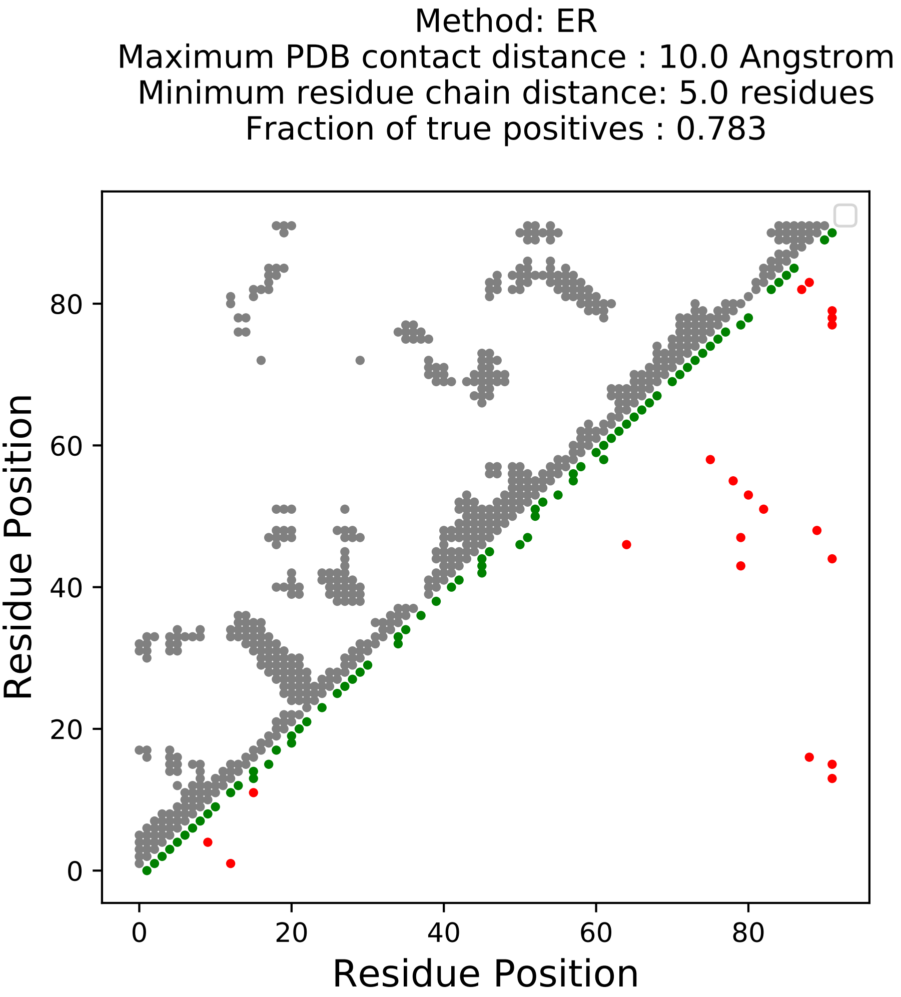

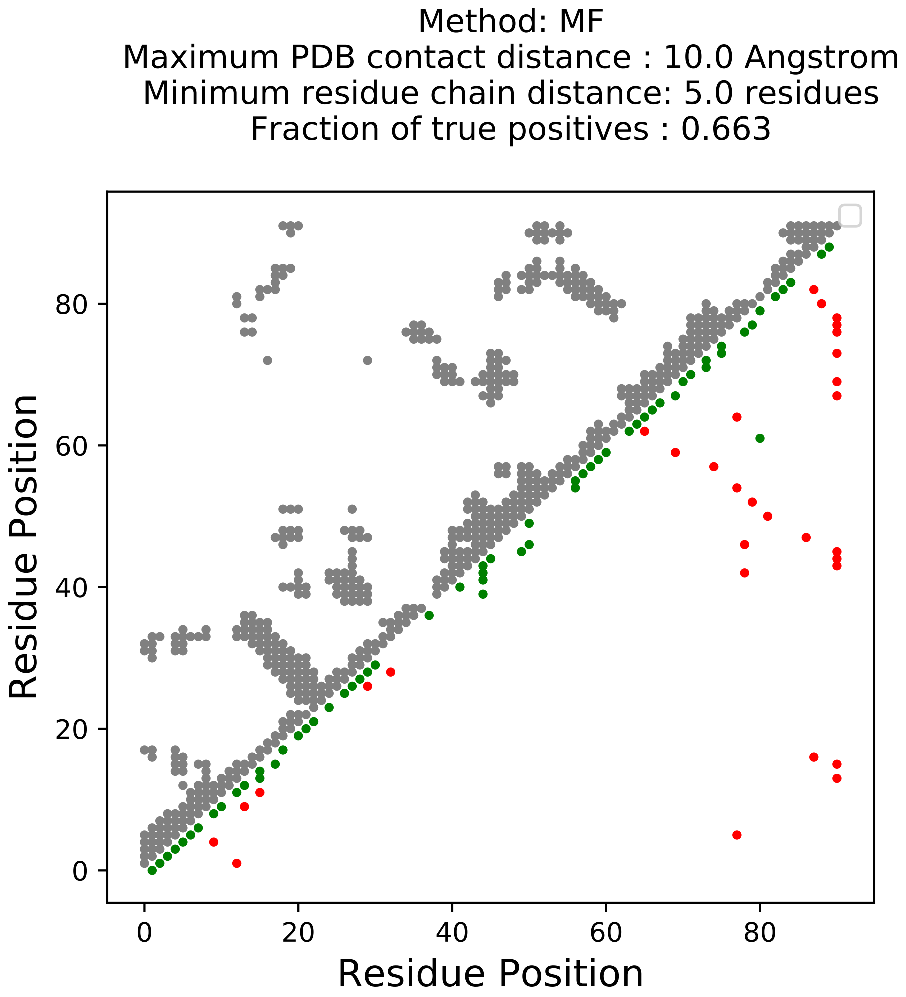

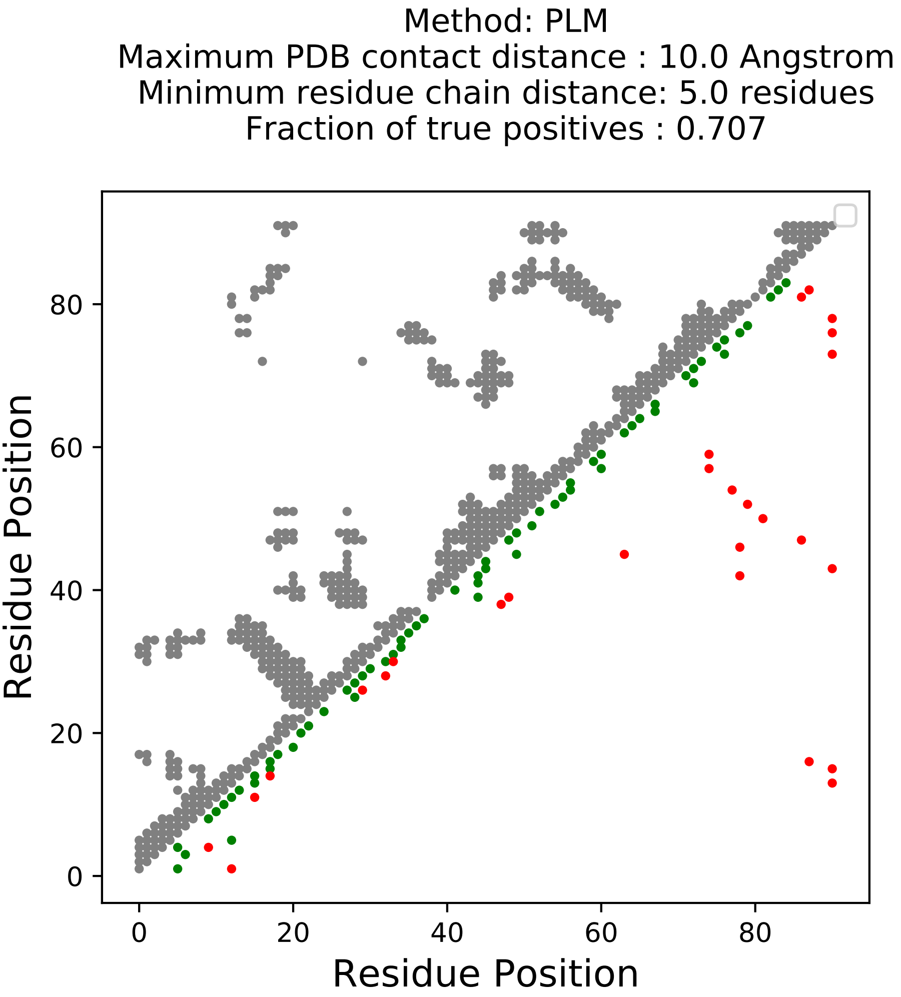

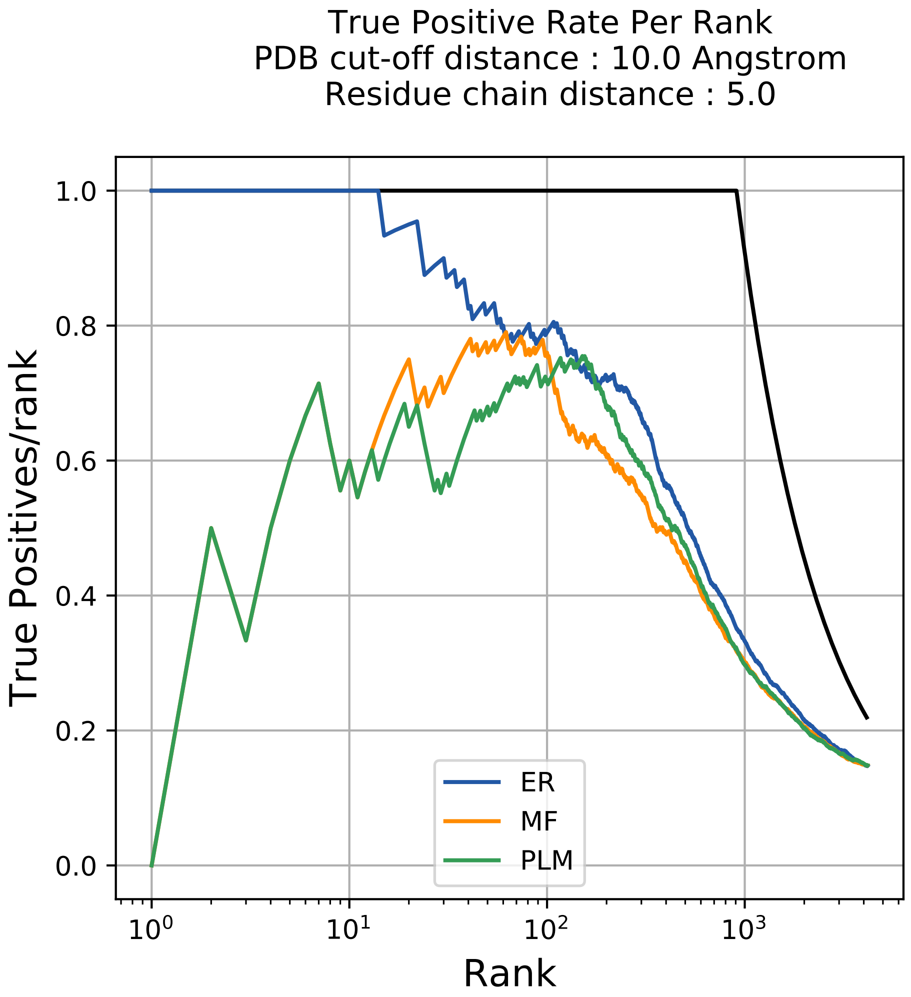

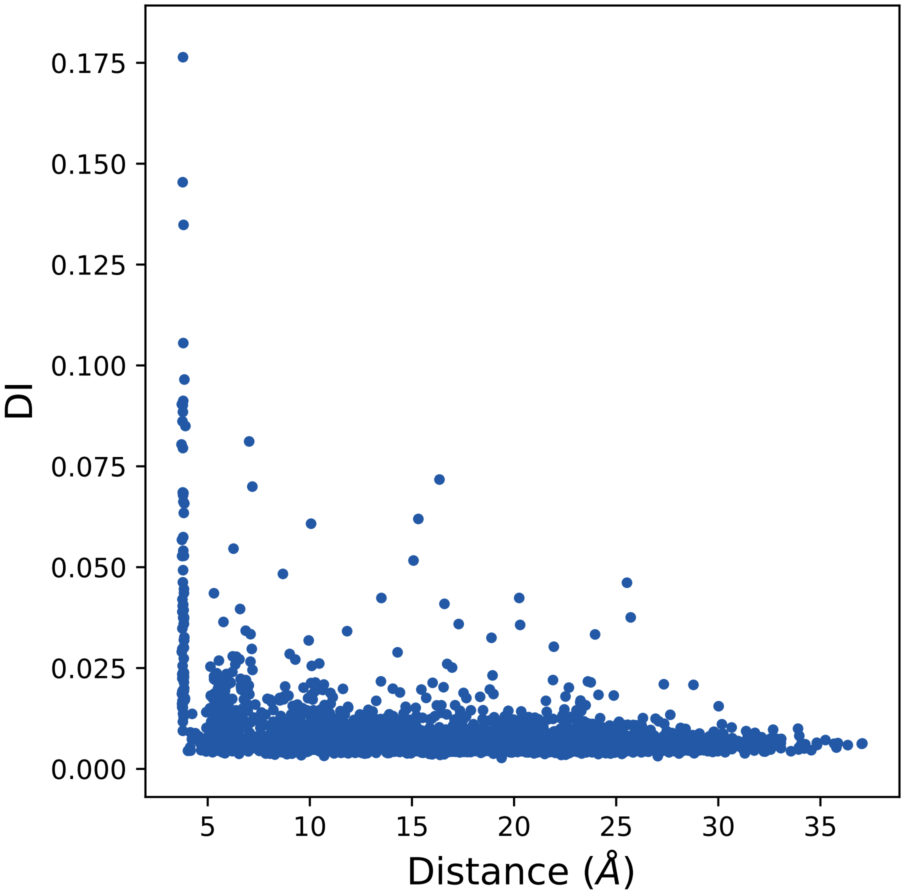

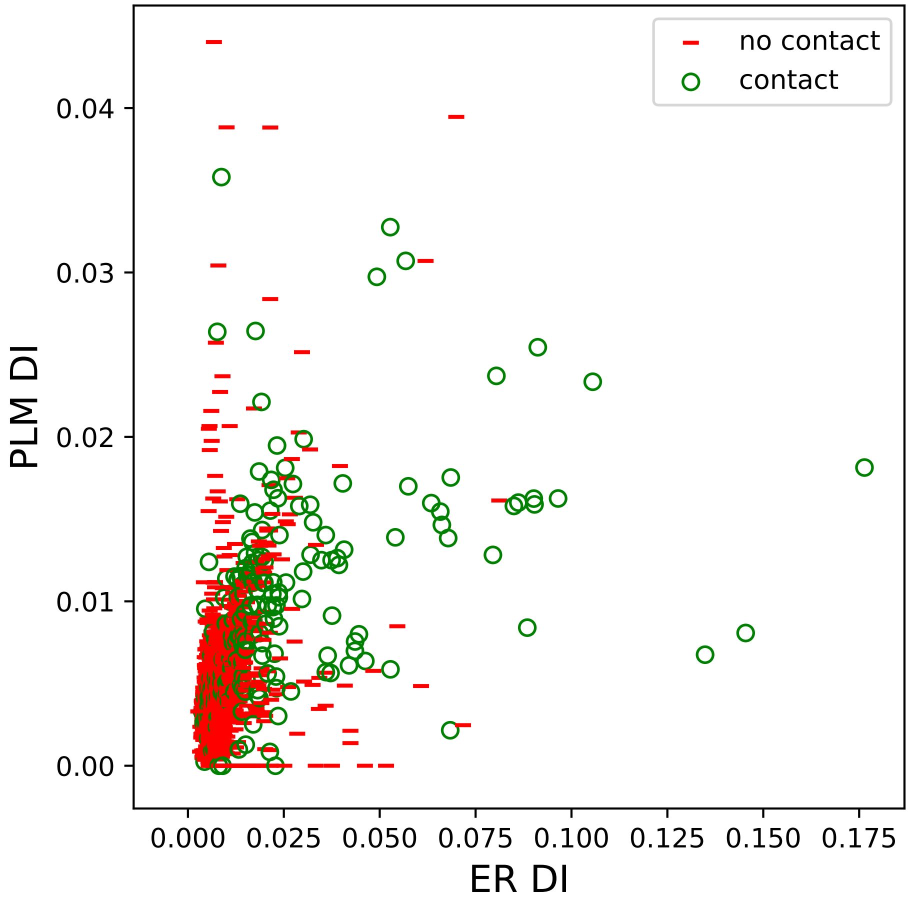

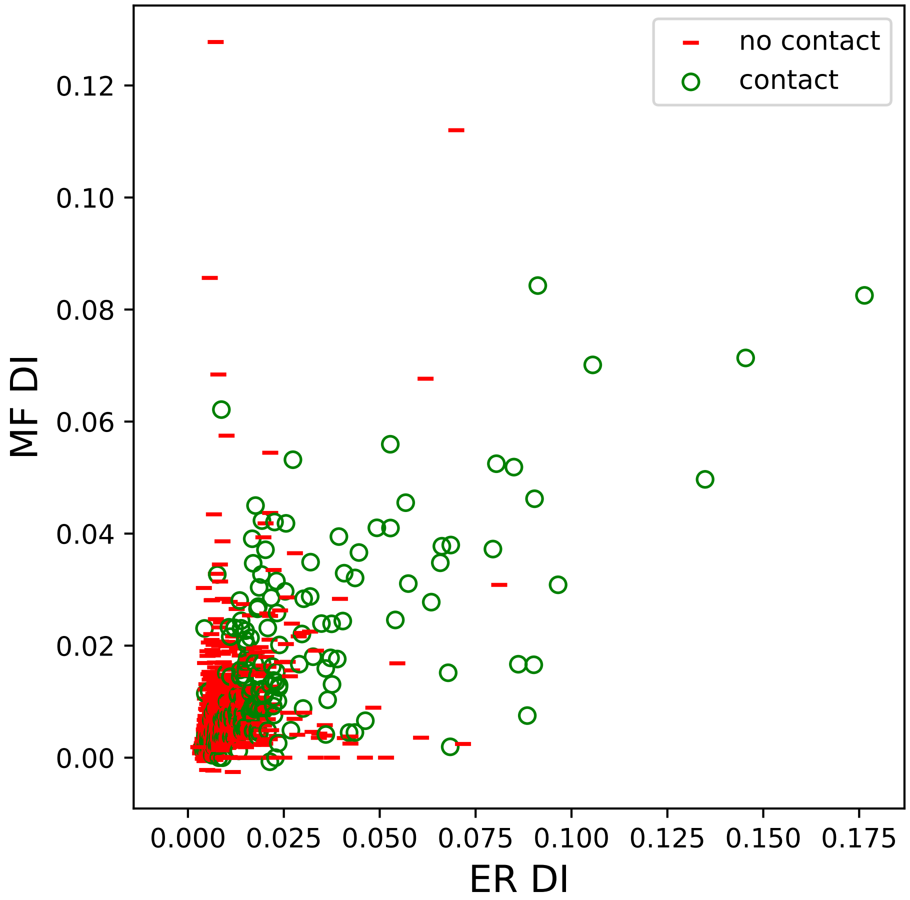

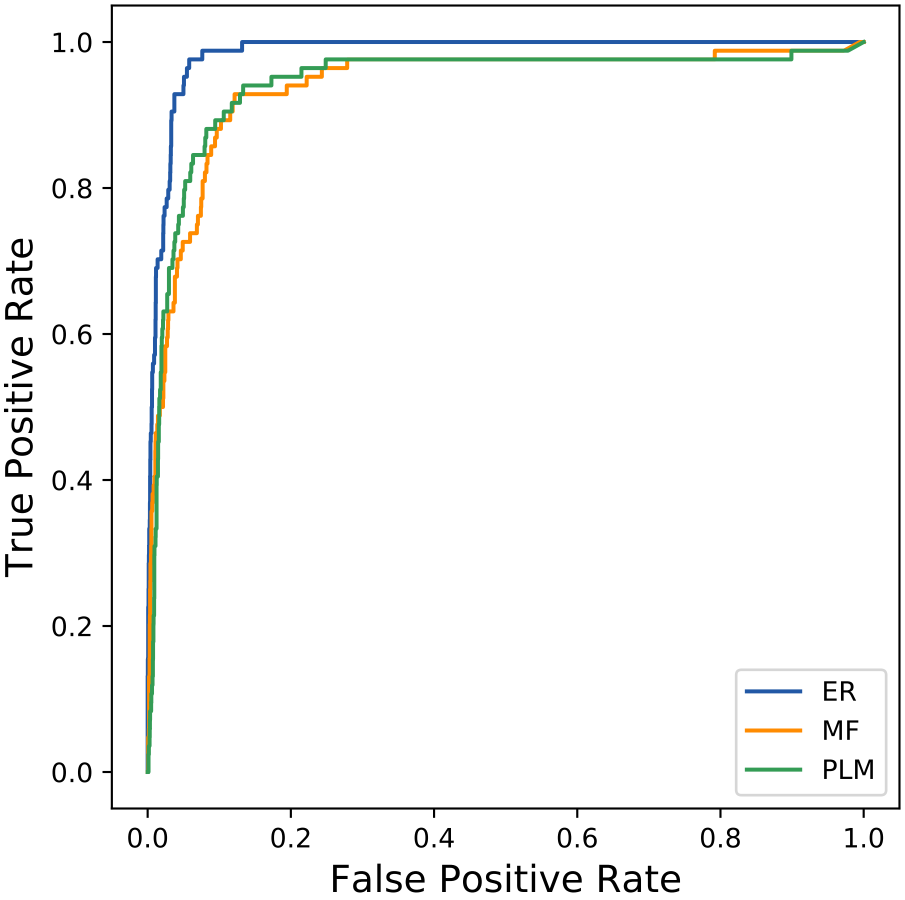

### 33933seq_81col_6pq3_PF00071_plots.pdf

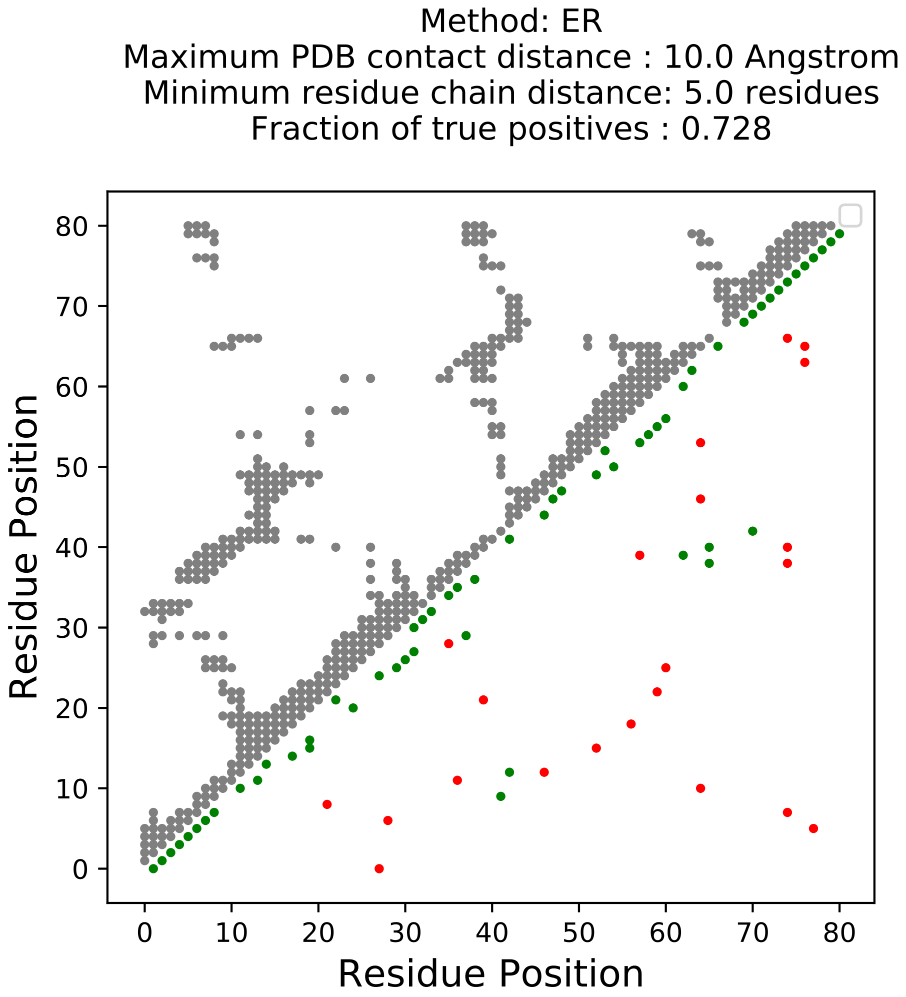

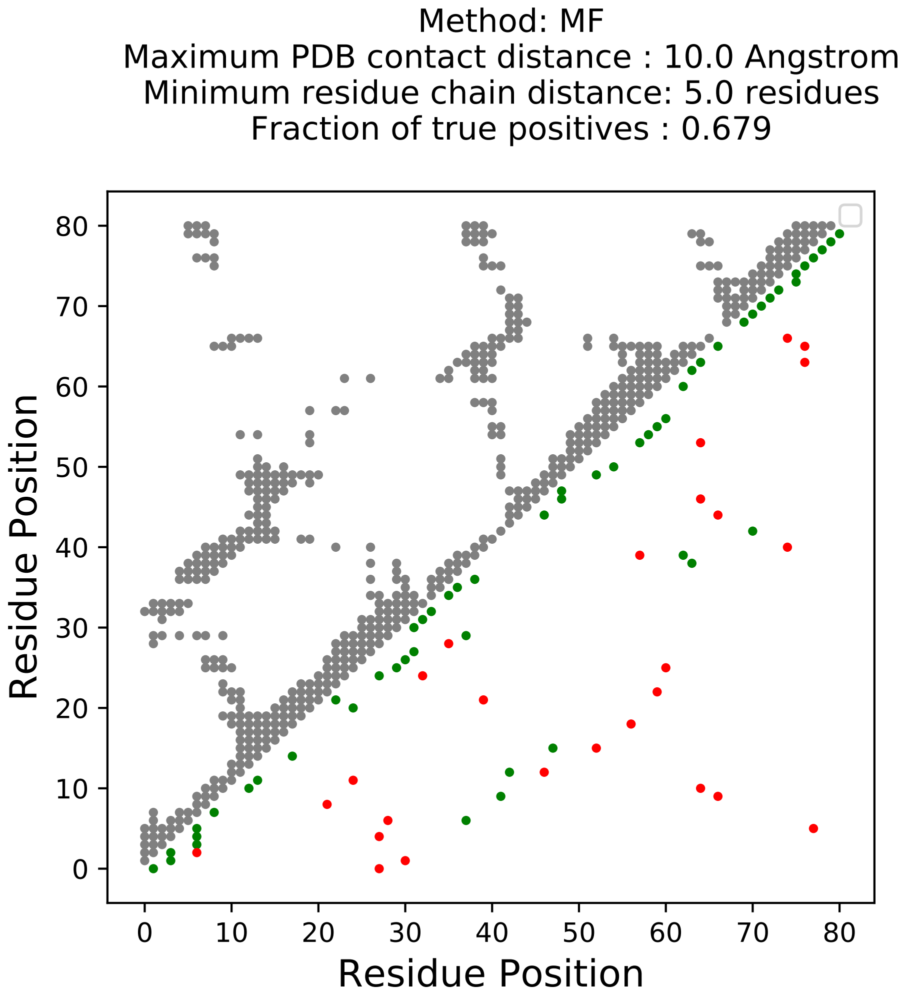

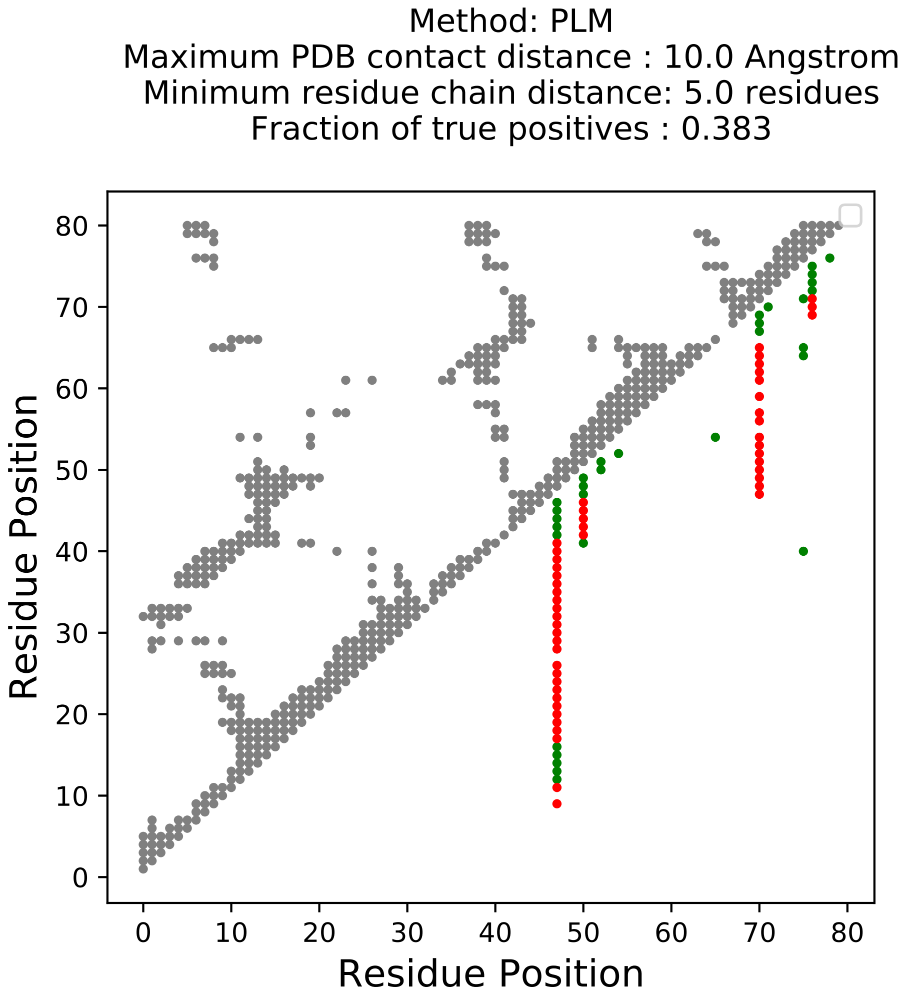

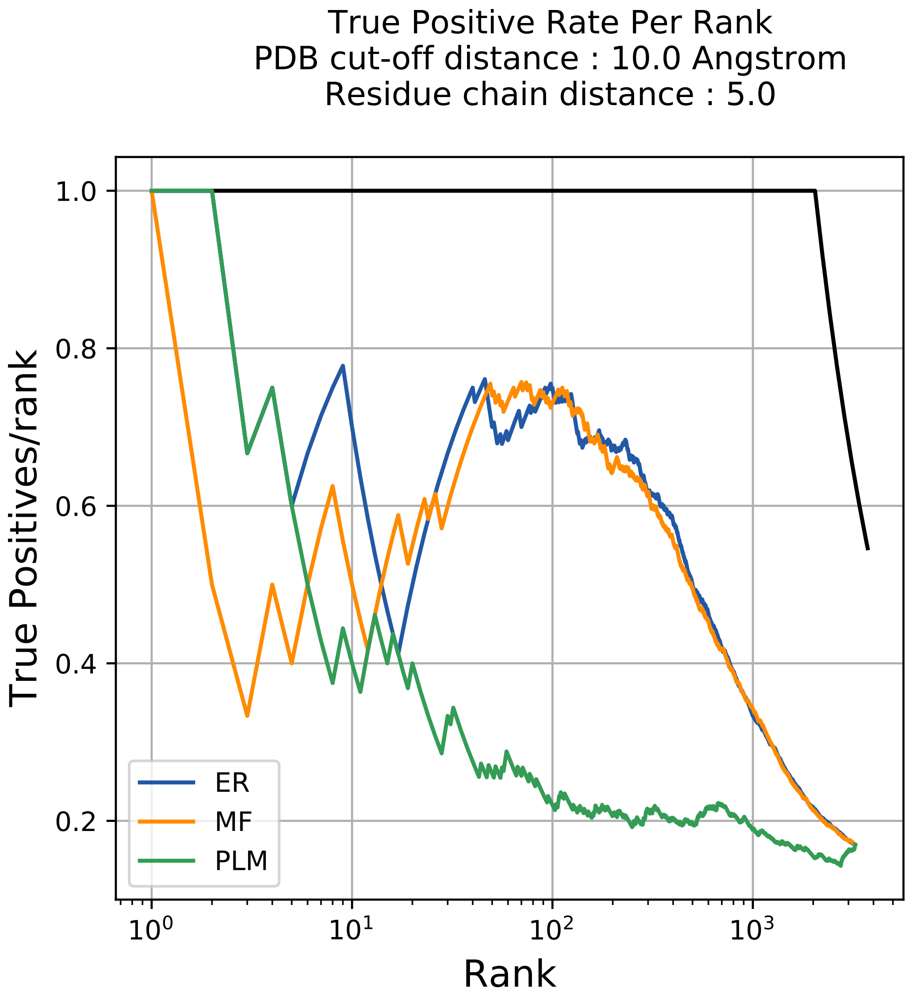

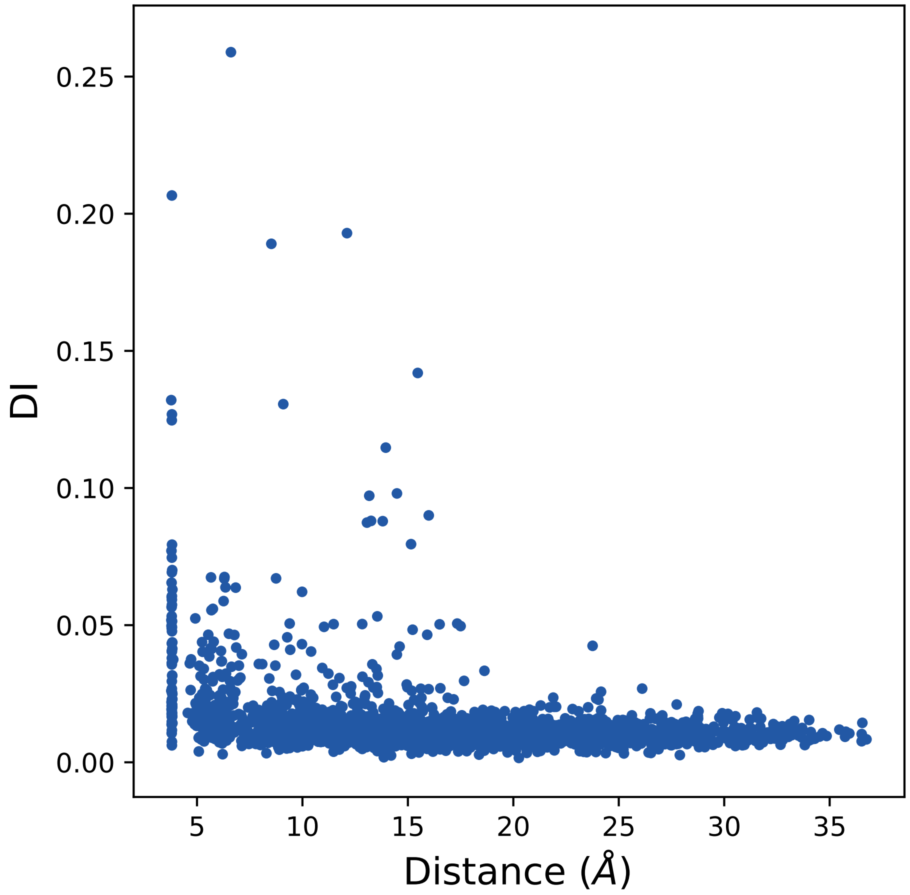

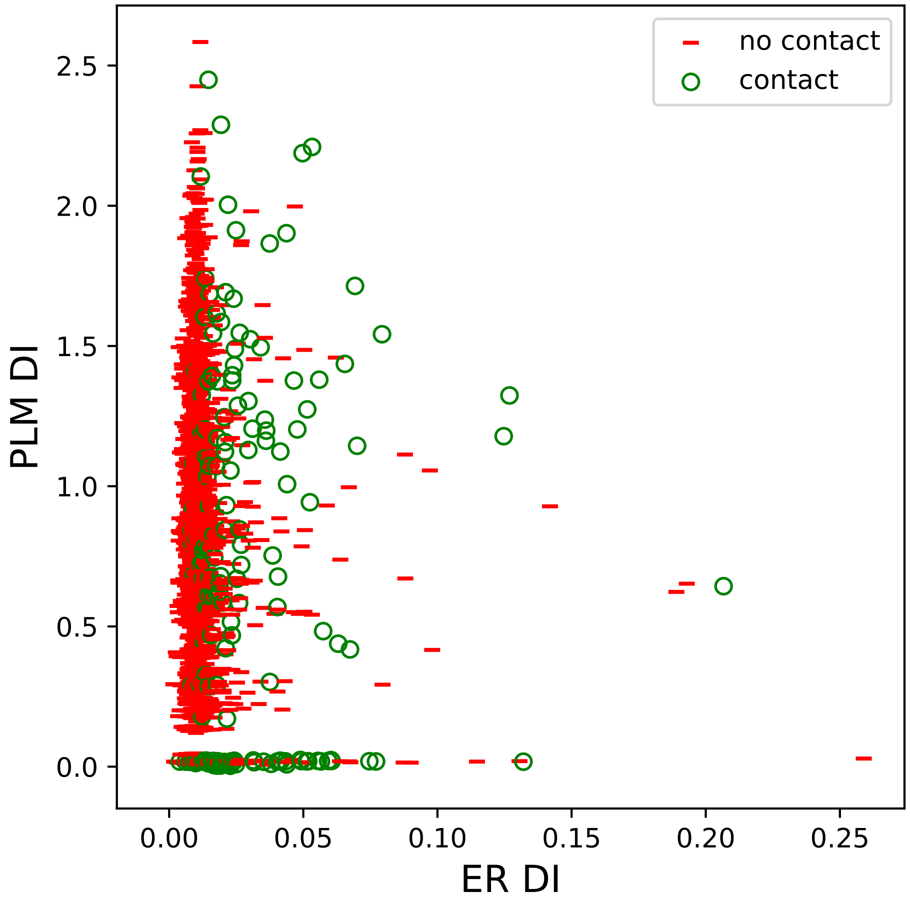

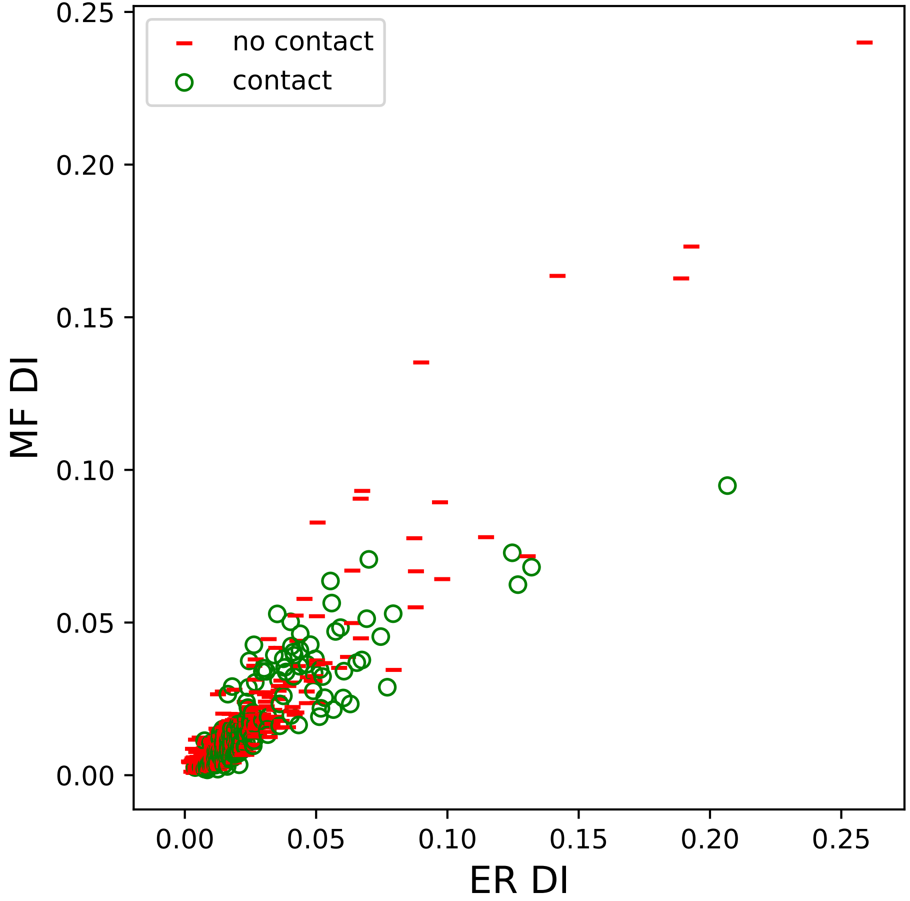

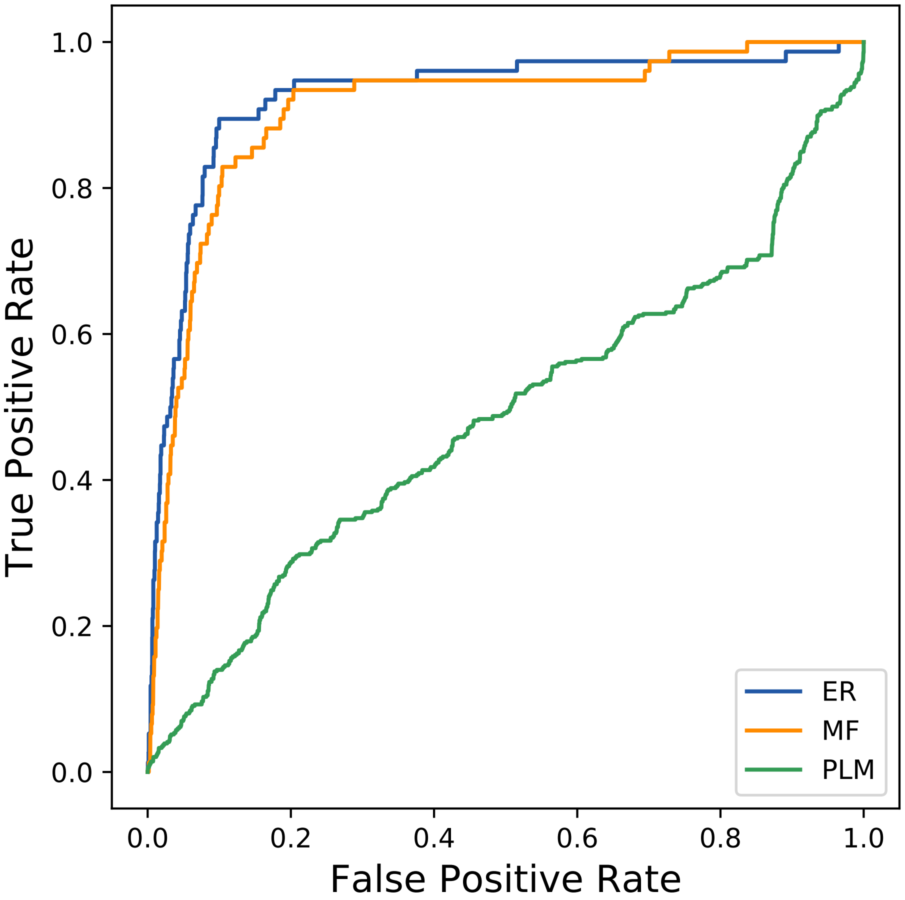

### 34104seq_86col_3rs4_PF00071_plots.pdf

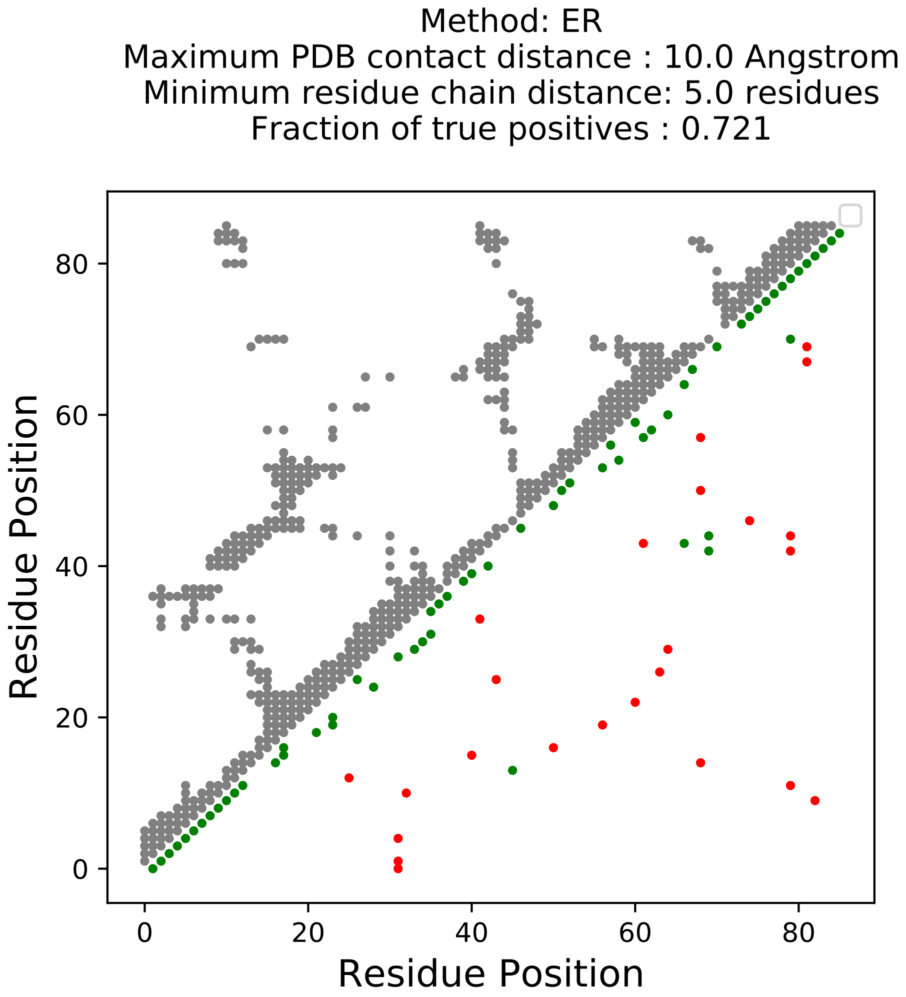

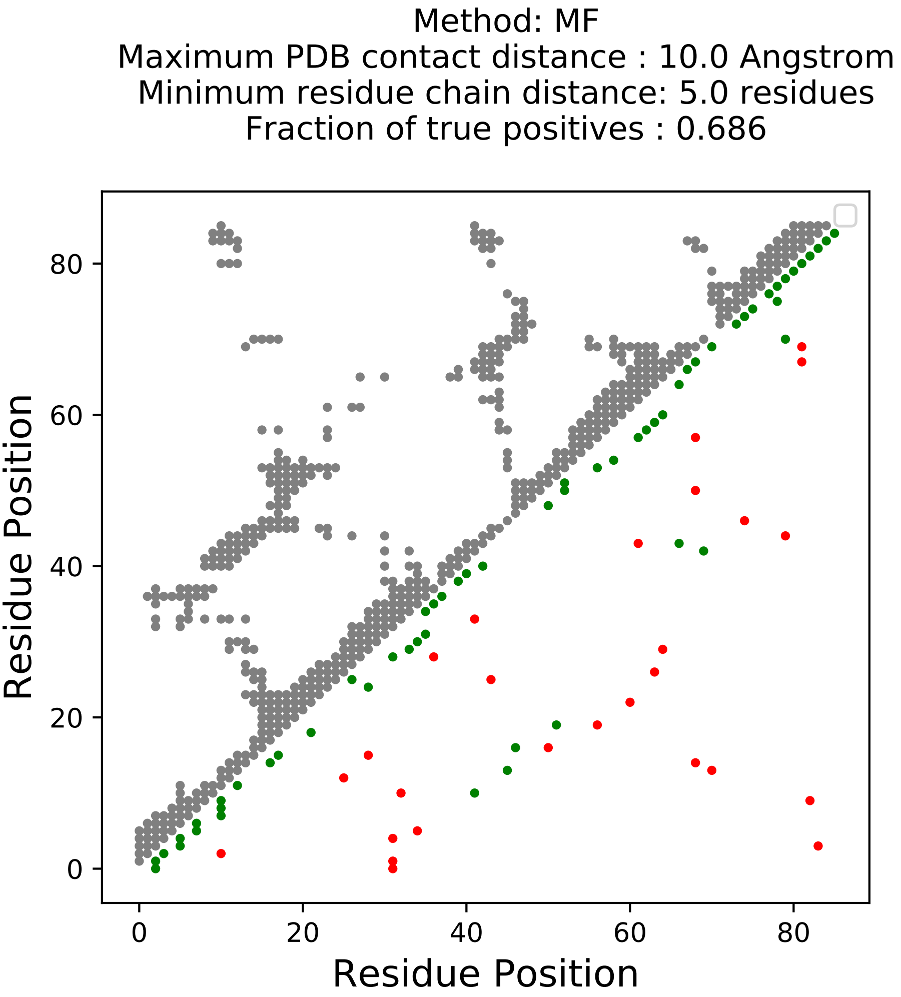

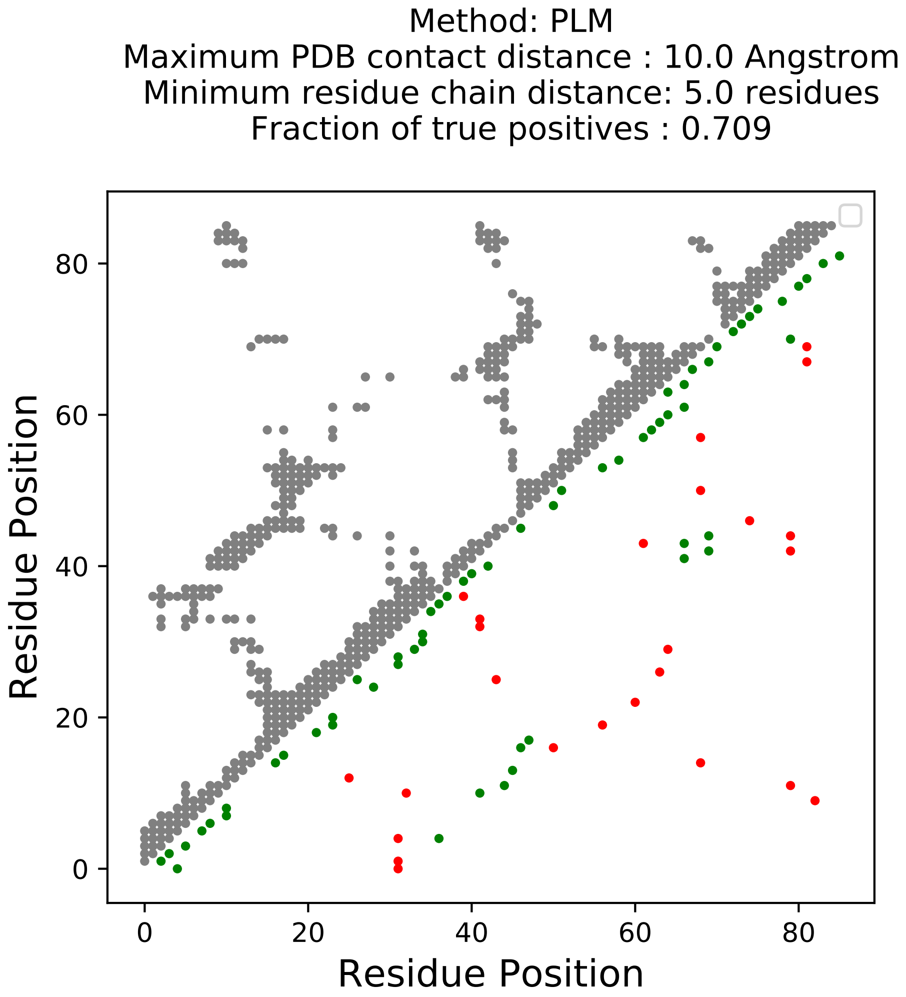

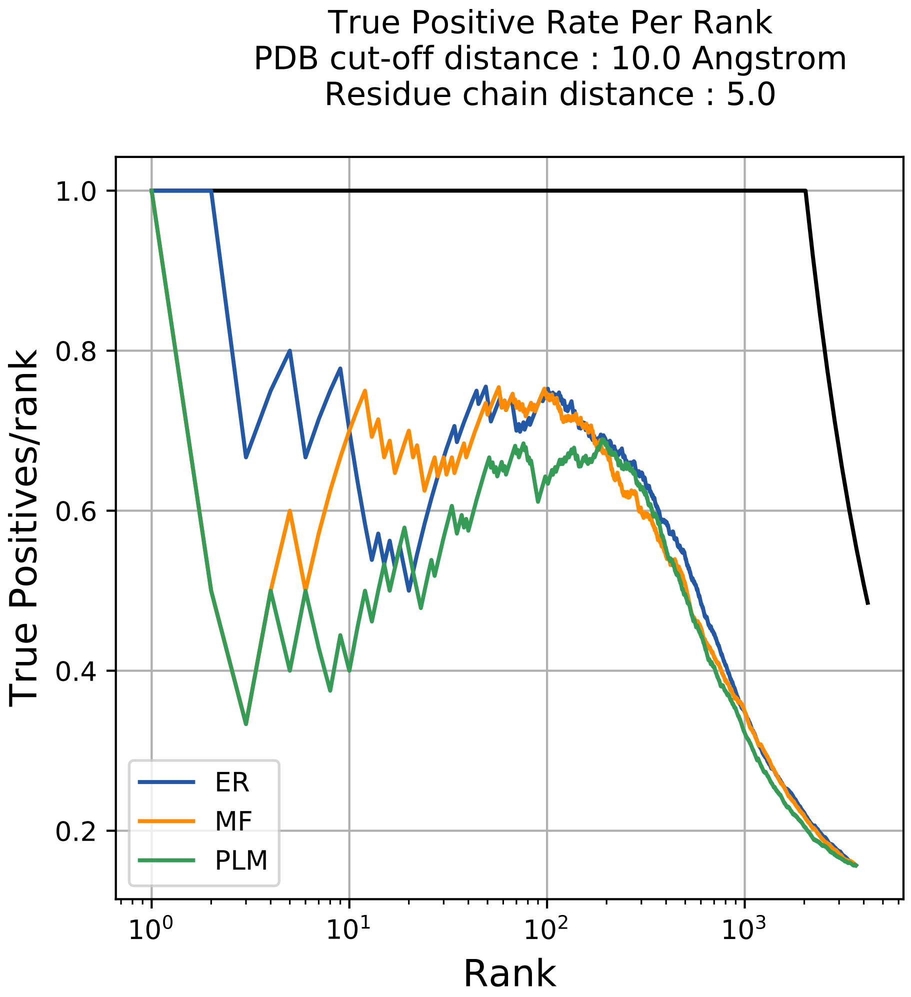

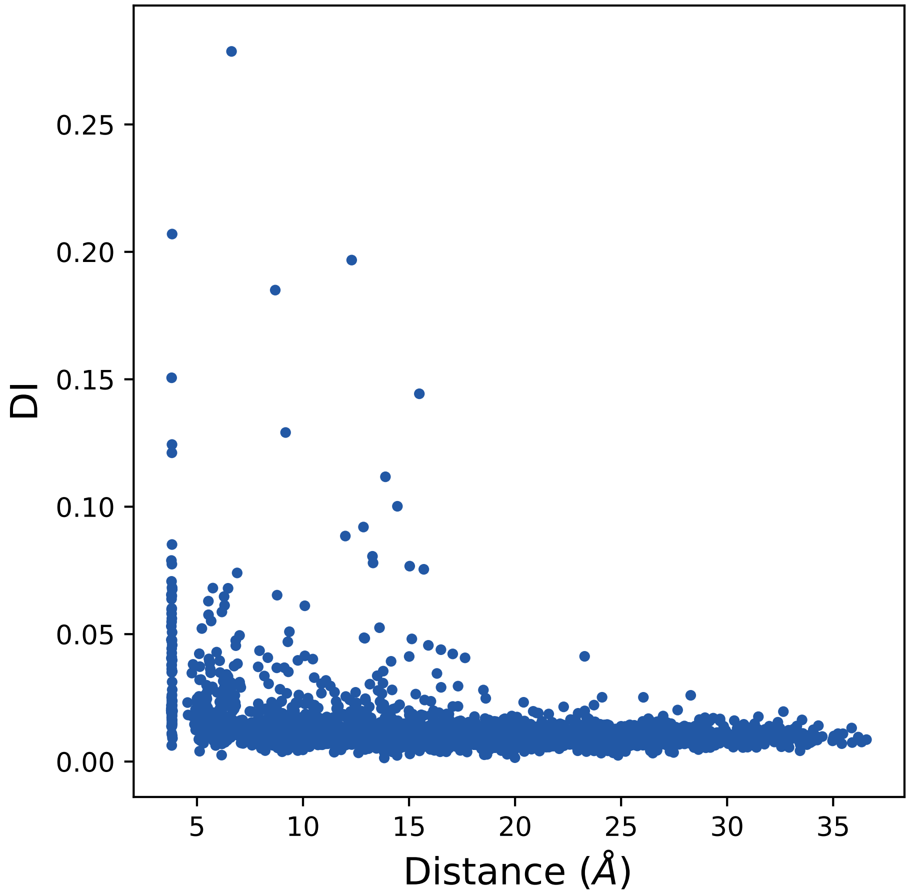

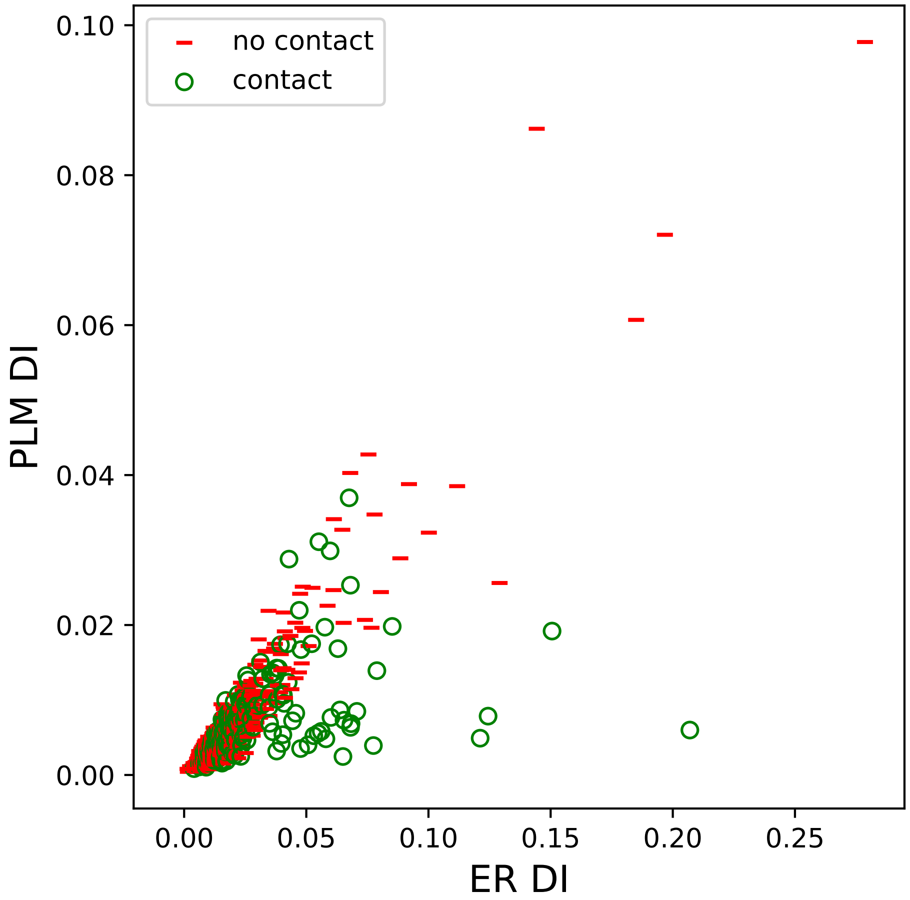

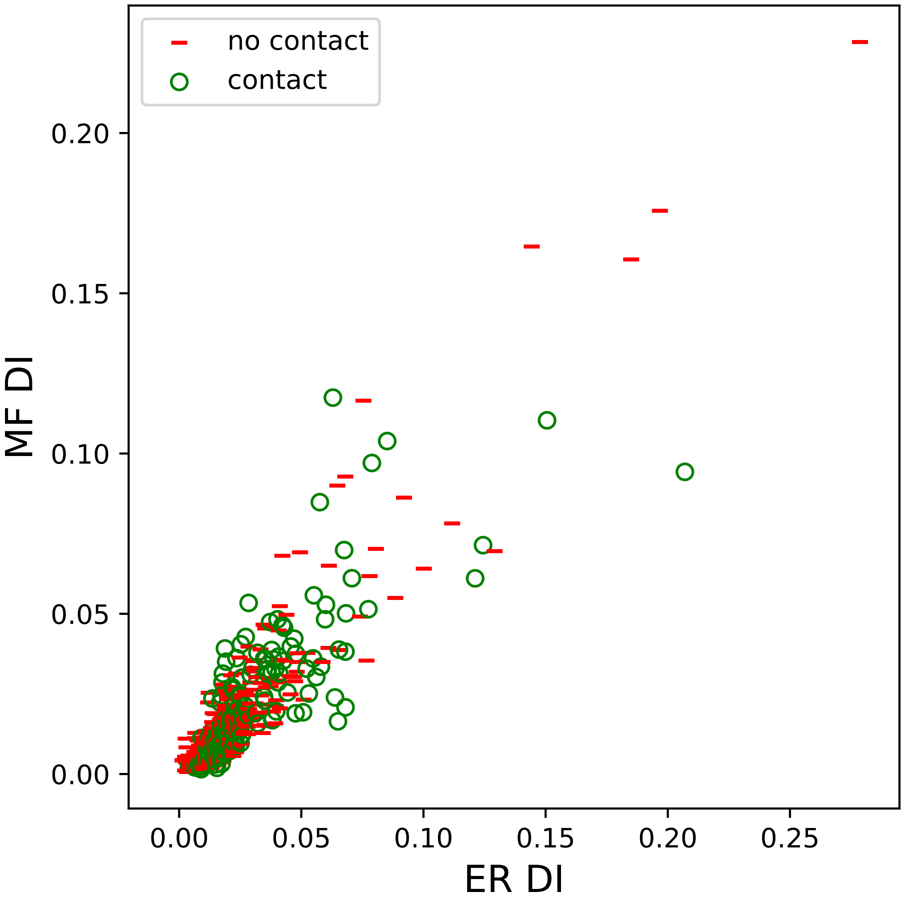

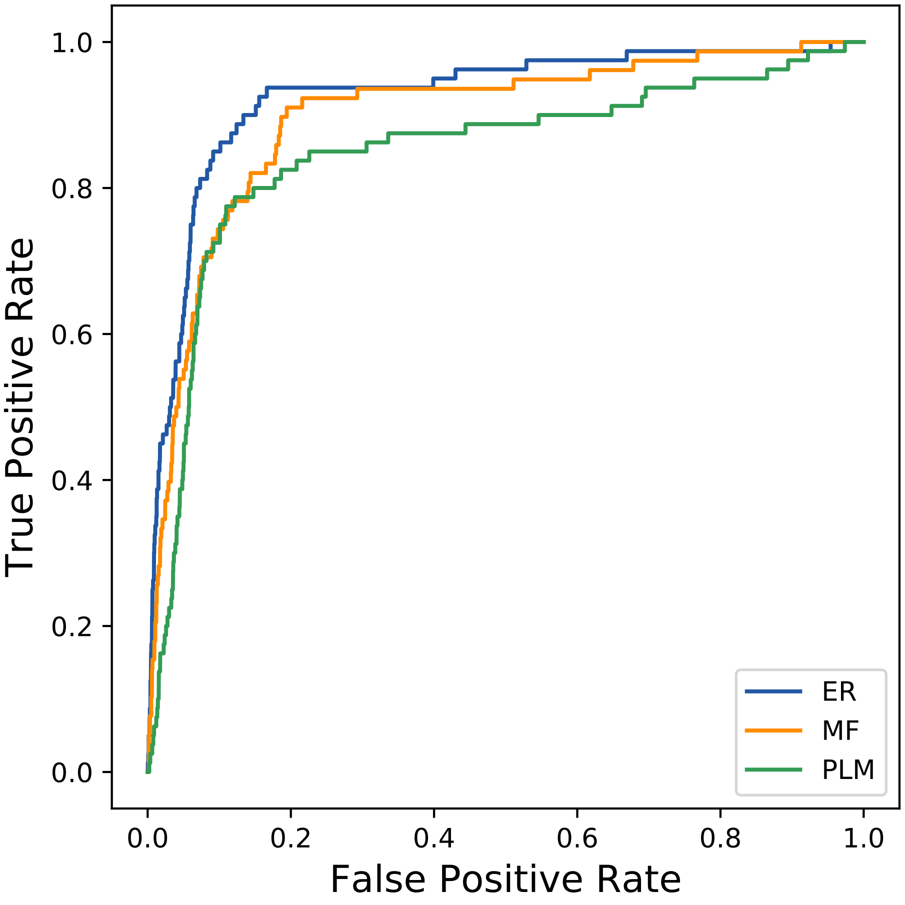

### 35617seq_52col_1rwy_PF13499_plots.pdf

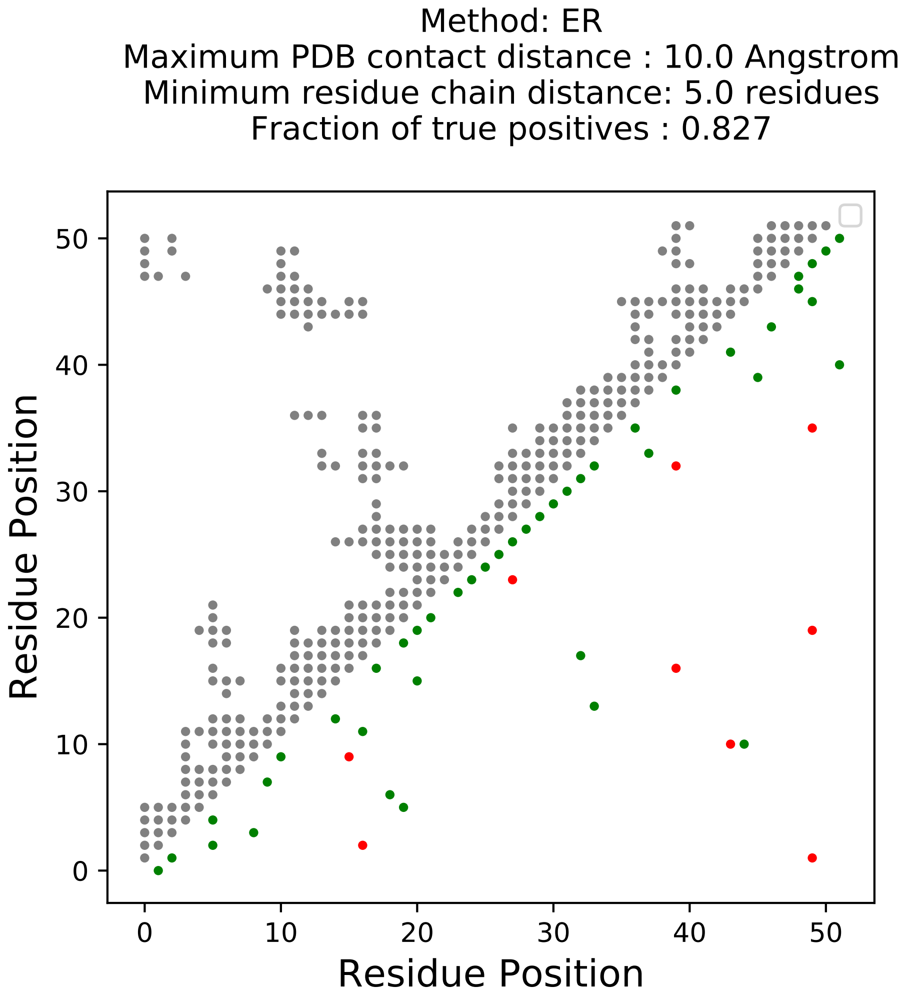

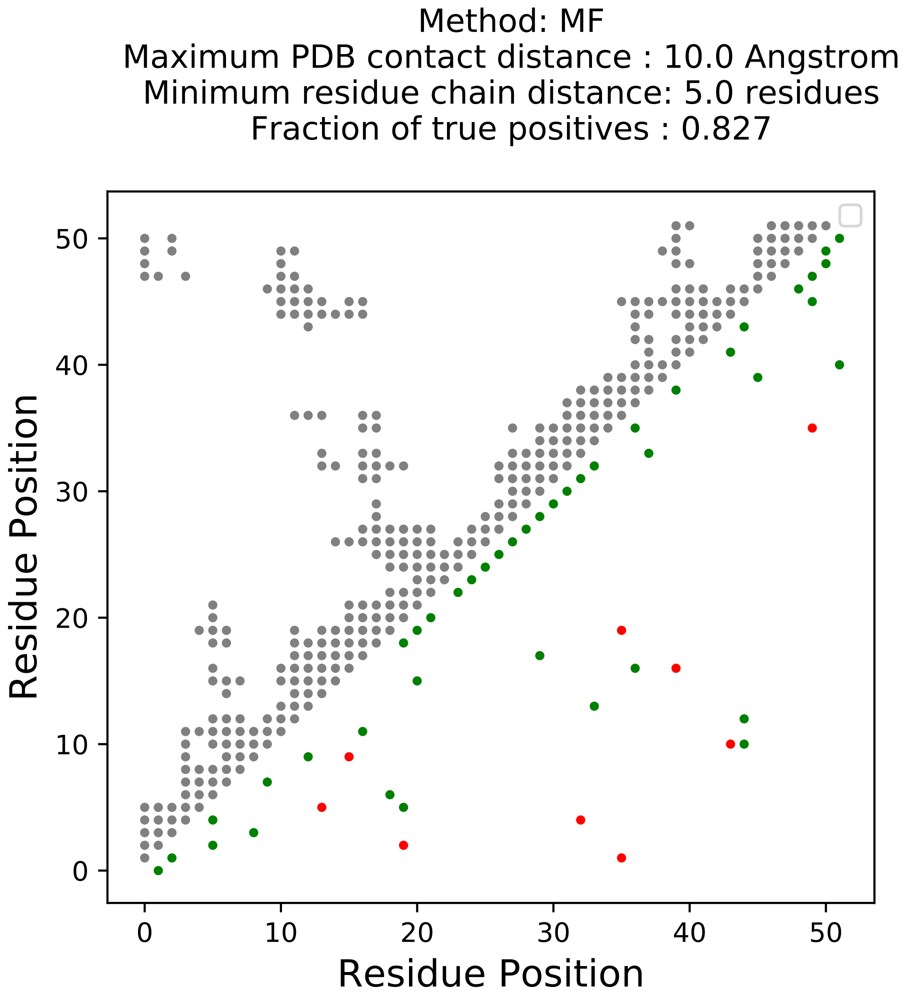

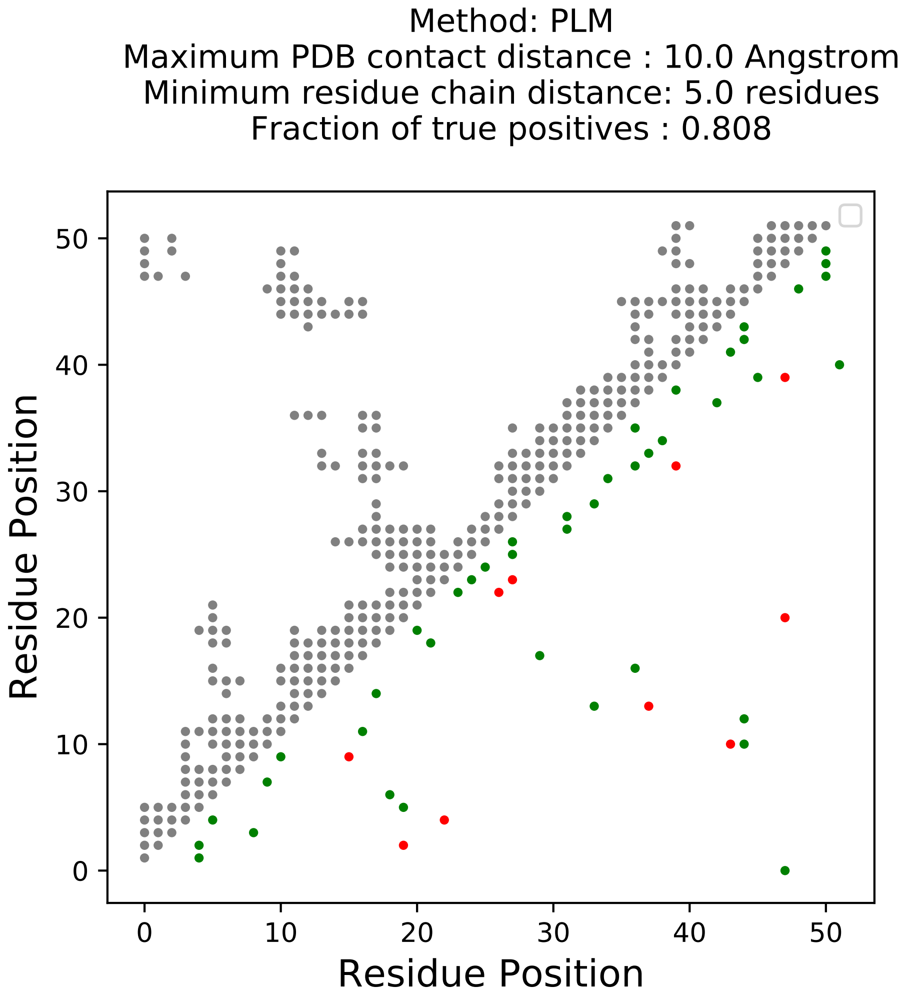

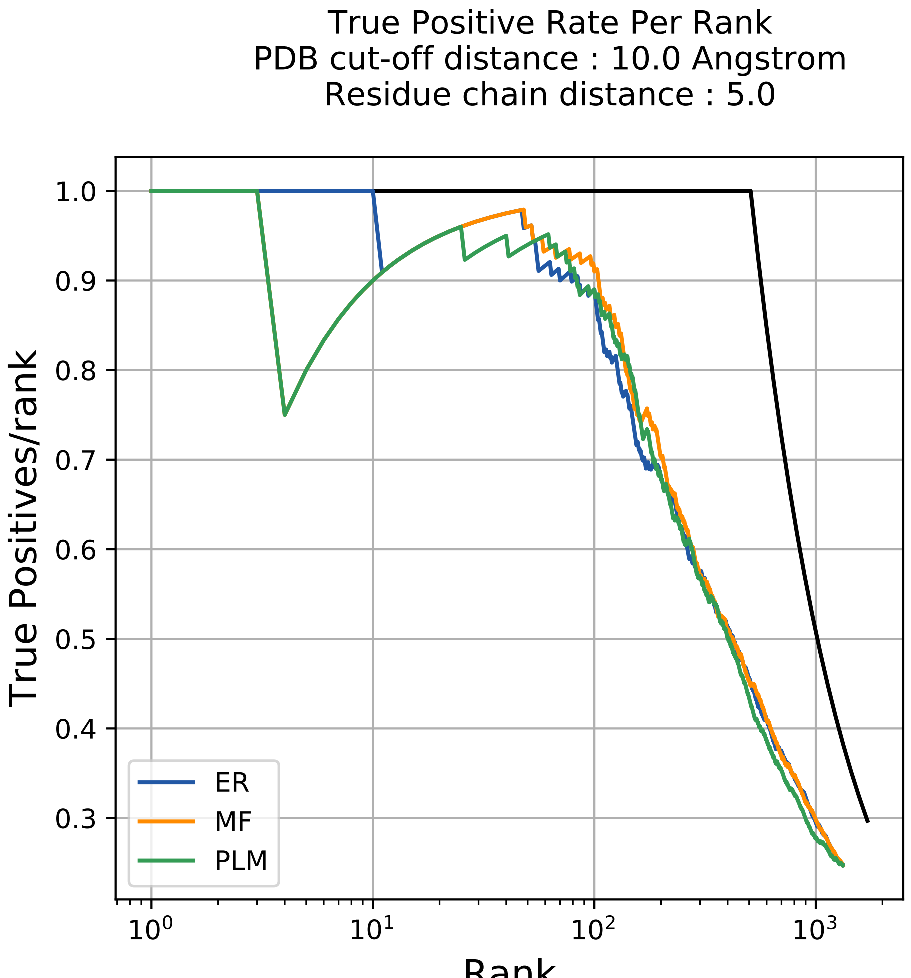

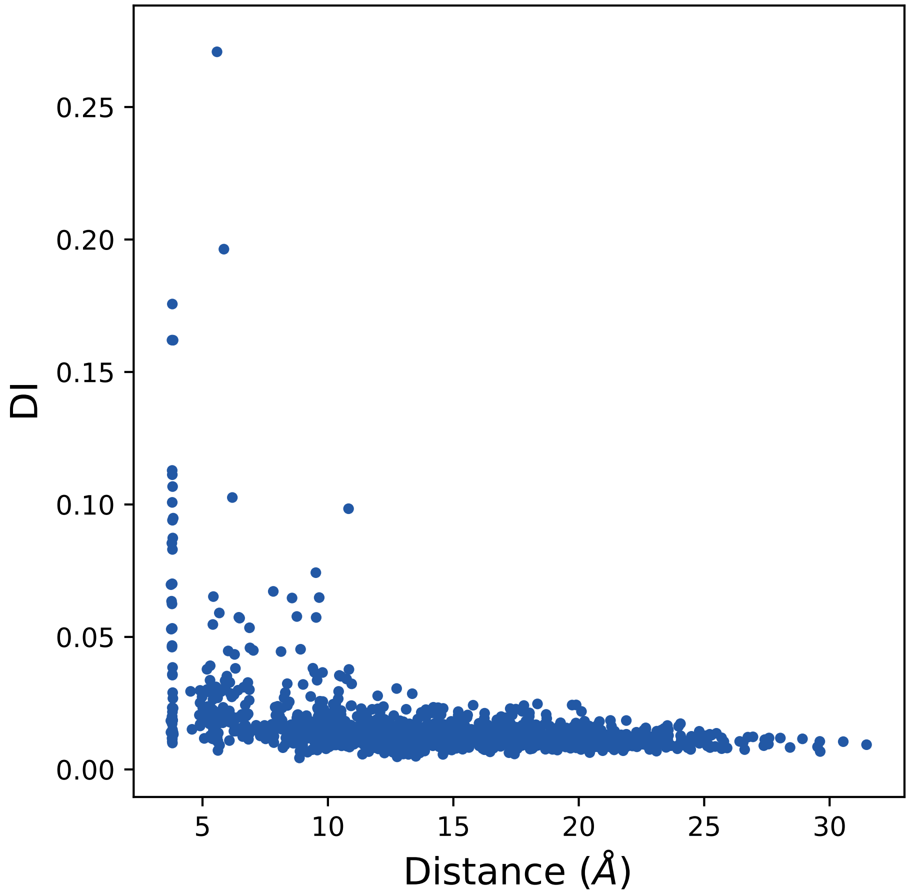

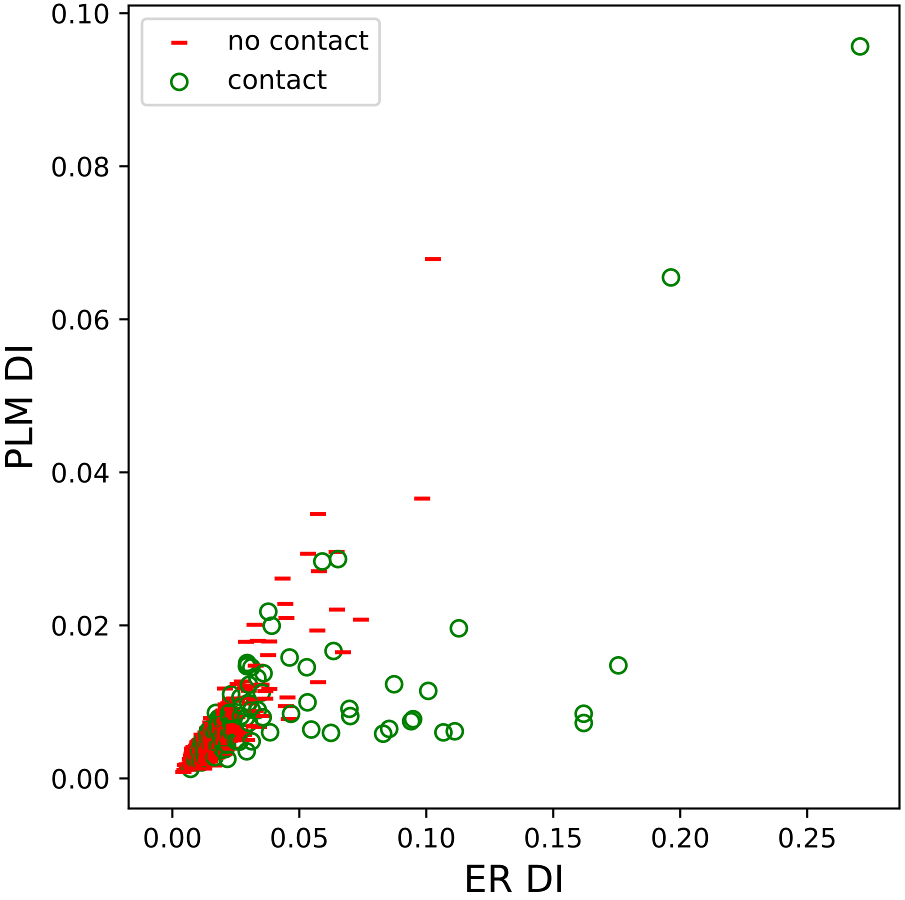

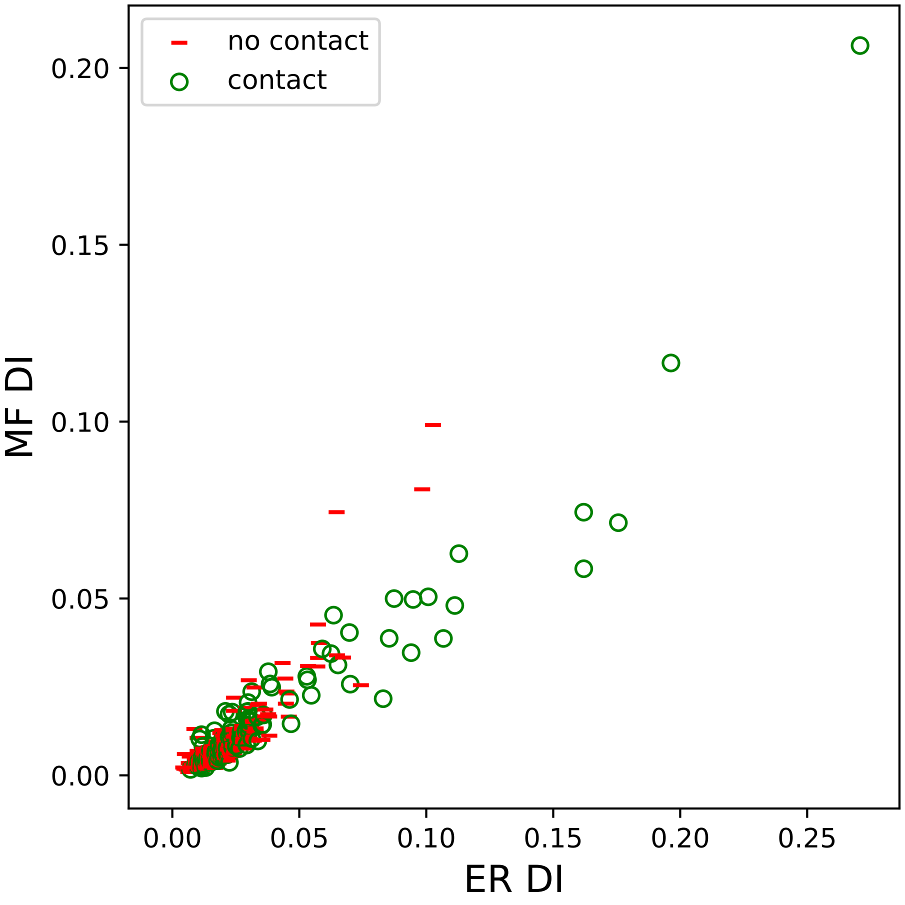

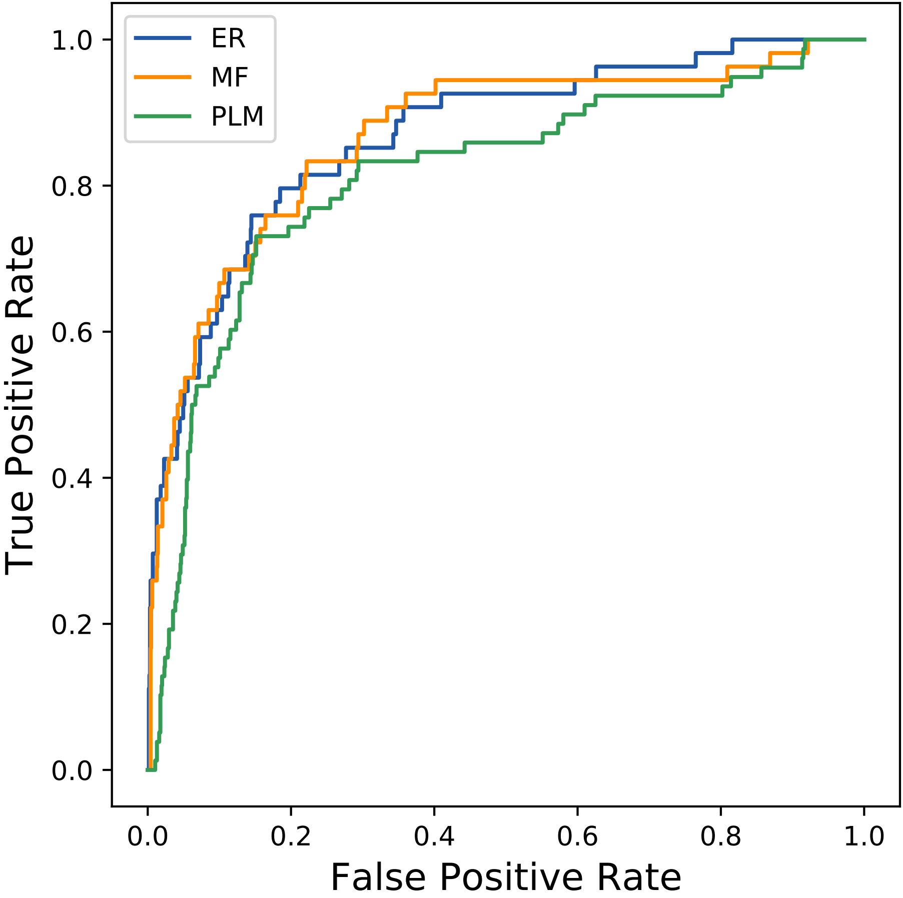

### 36387seq_101col_3a2w_PF00578_plots.pdf

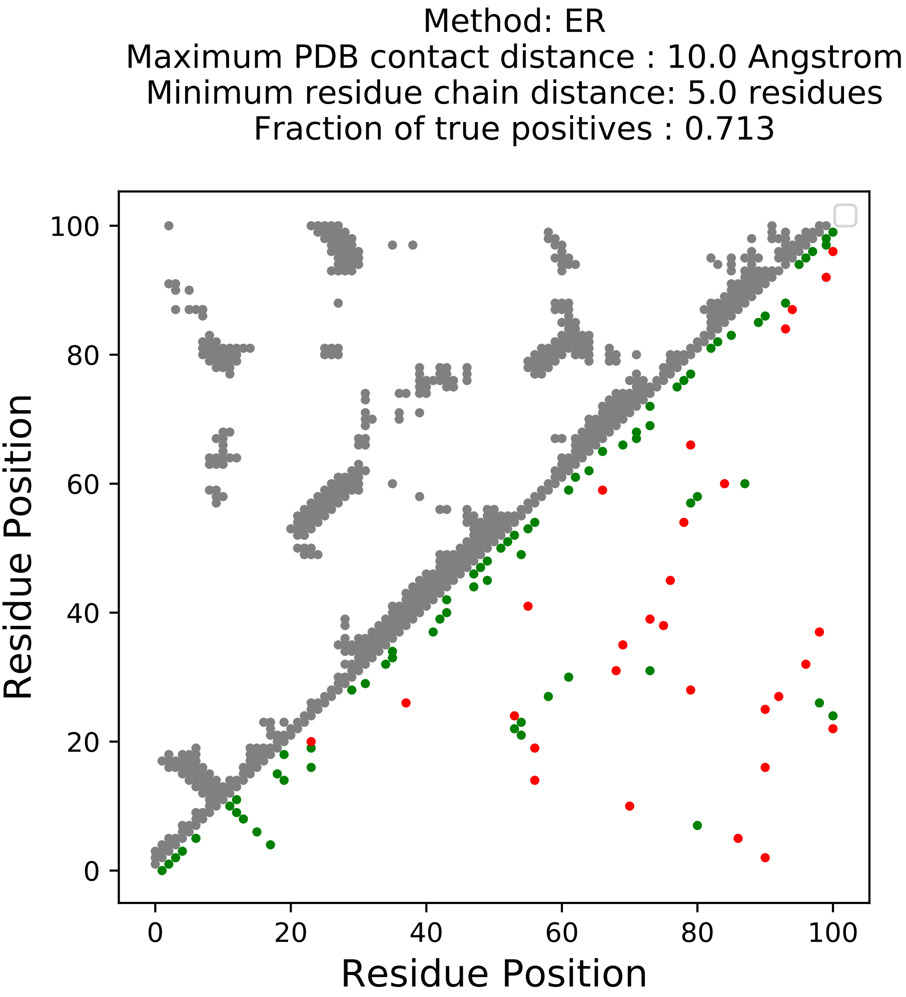

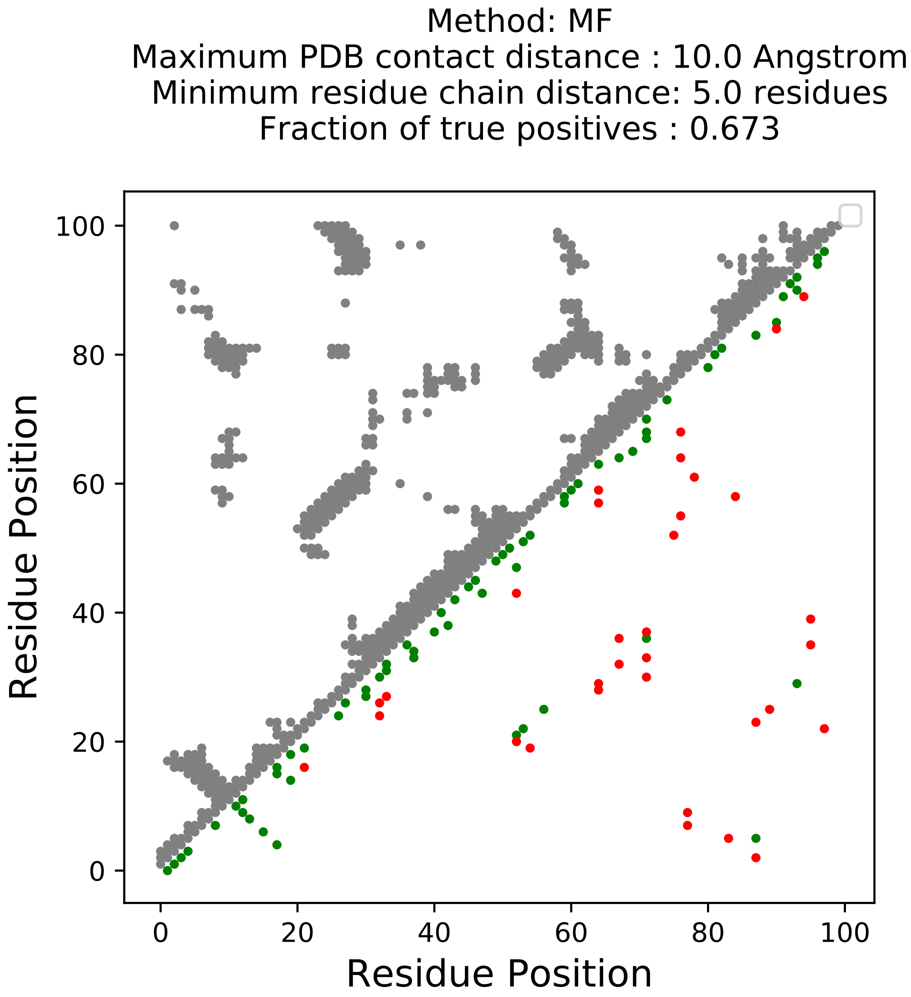

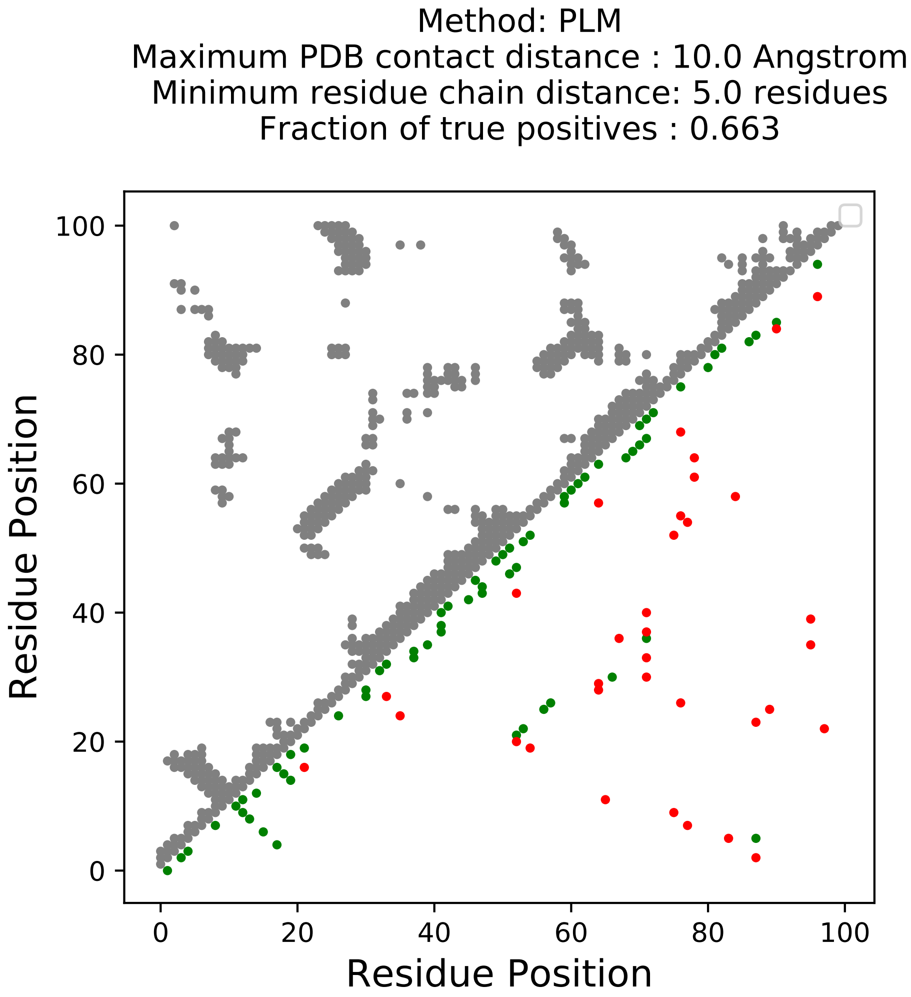

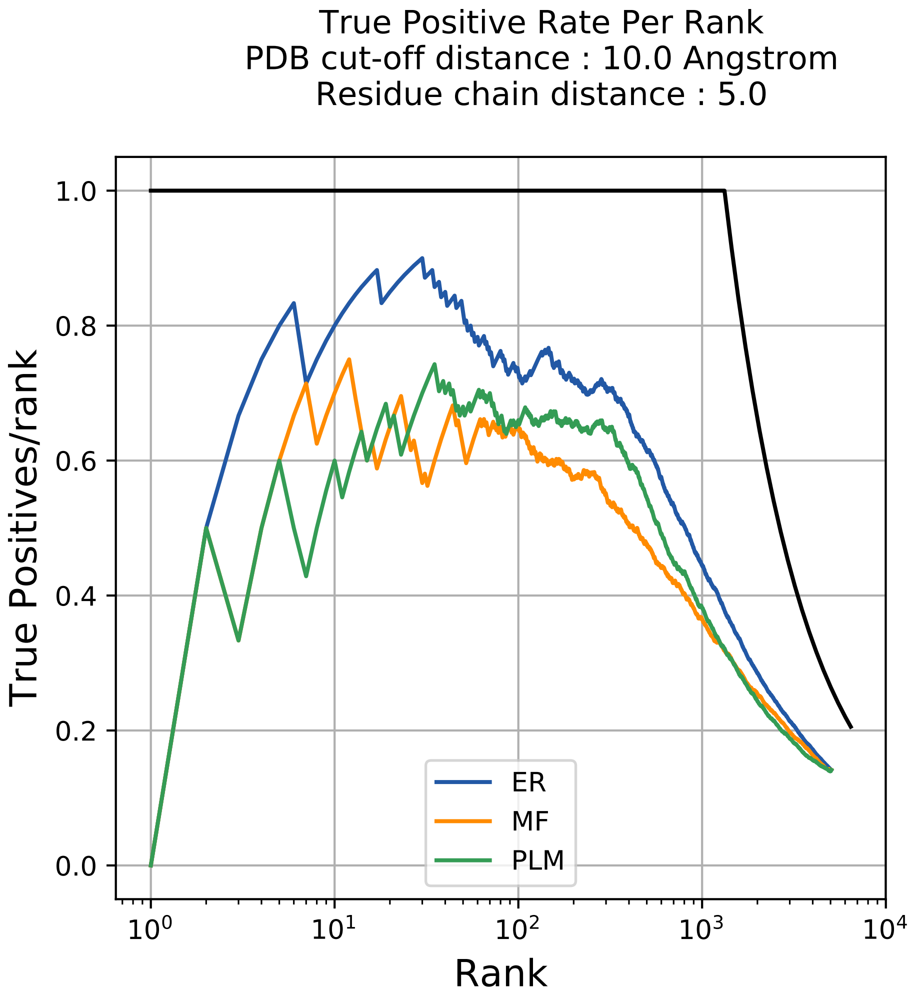

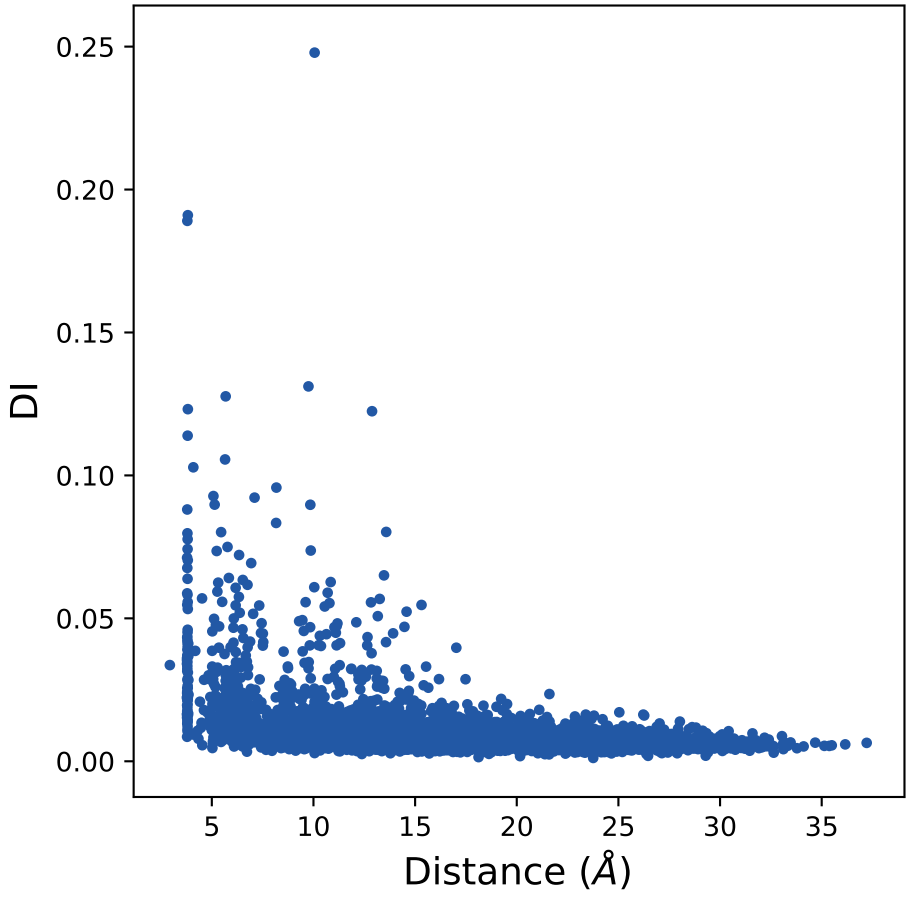

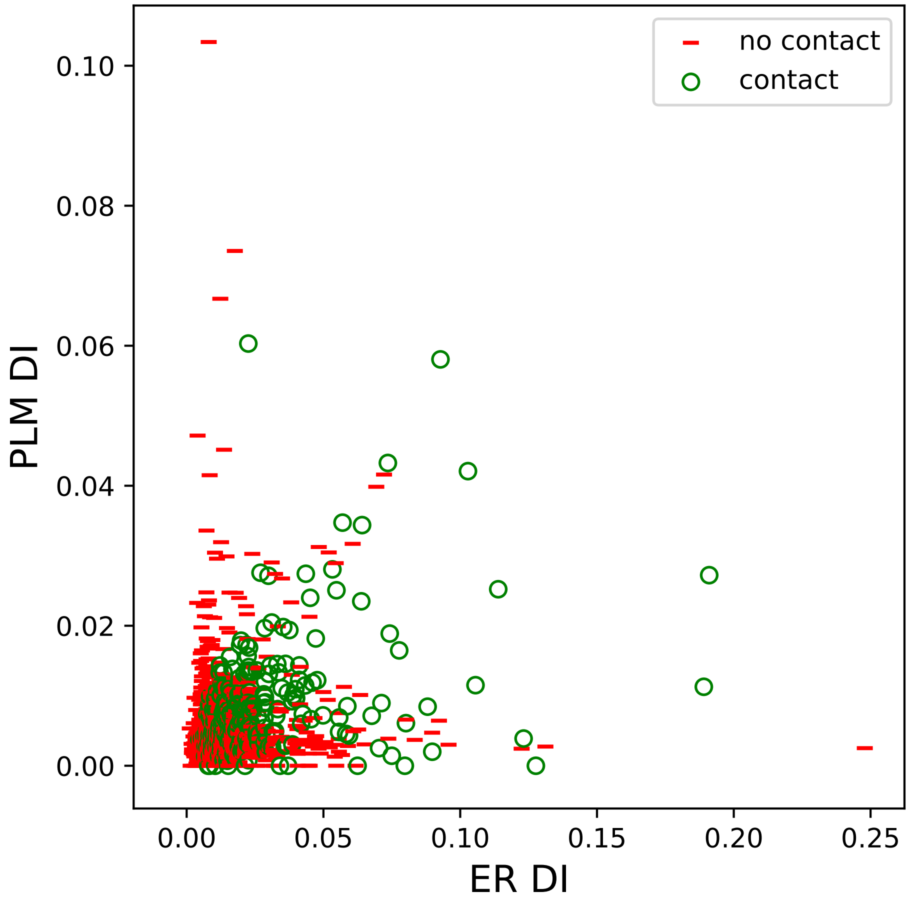

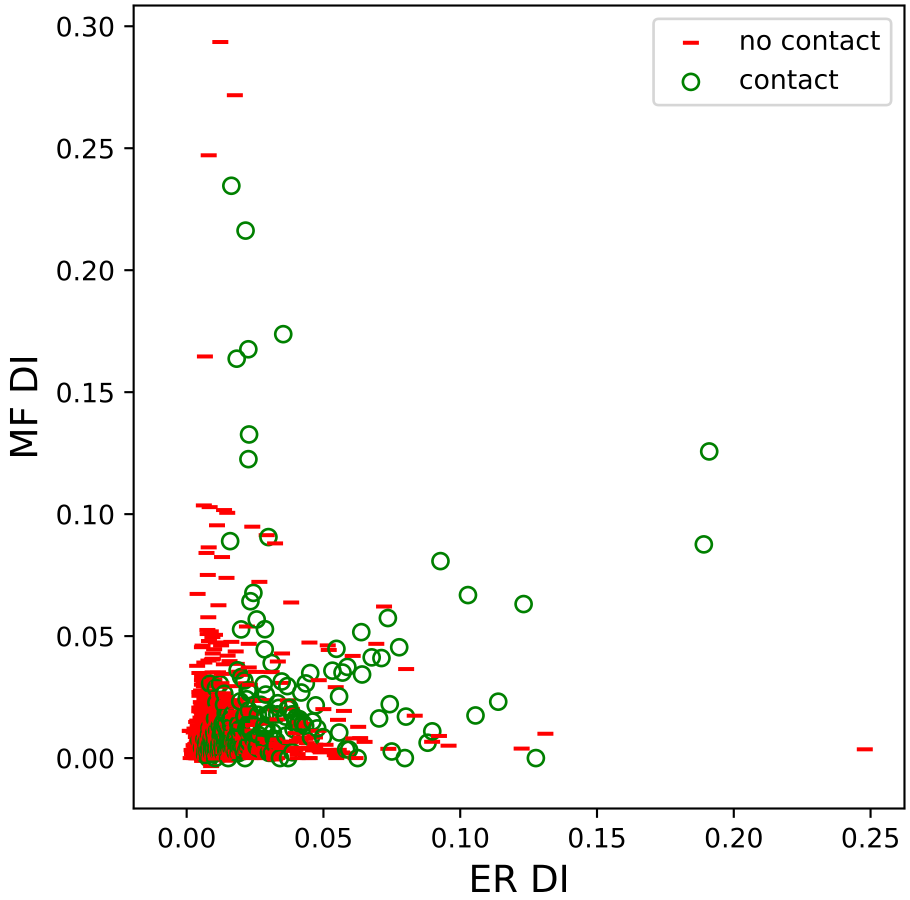

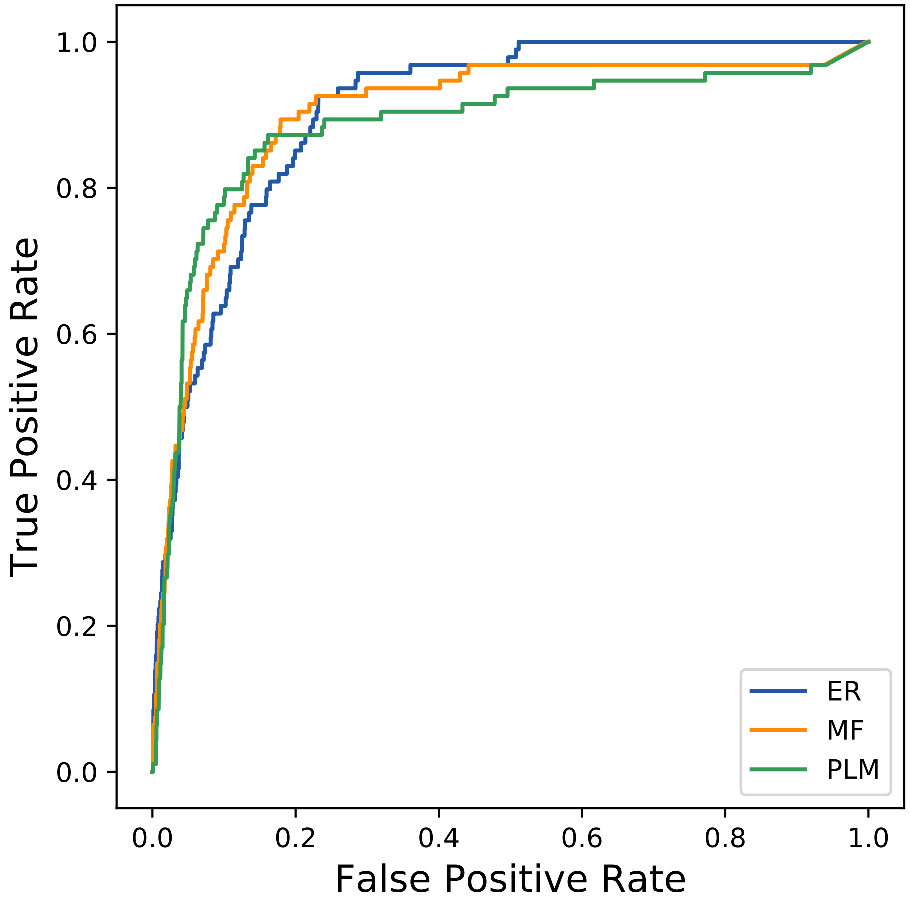

### 37011seq_186col_5b7u_PF00266_plots.pdf

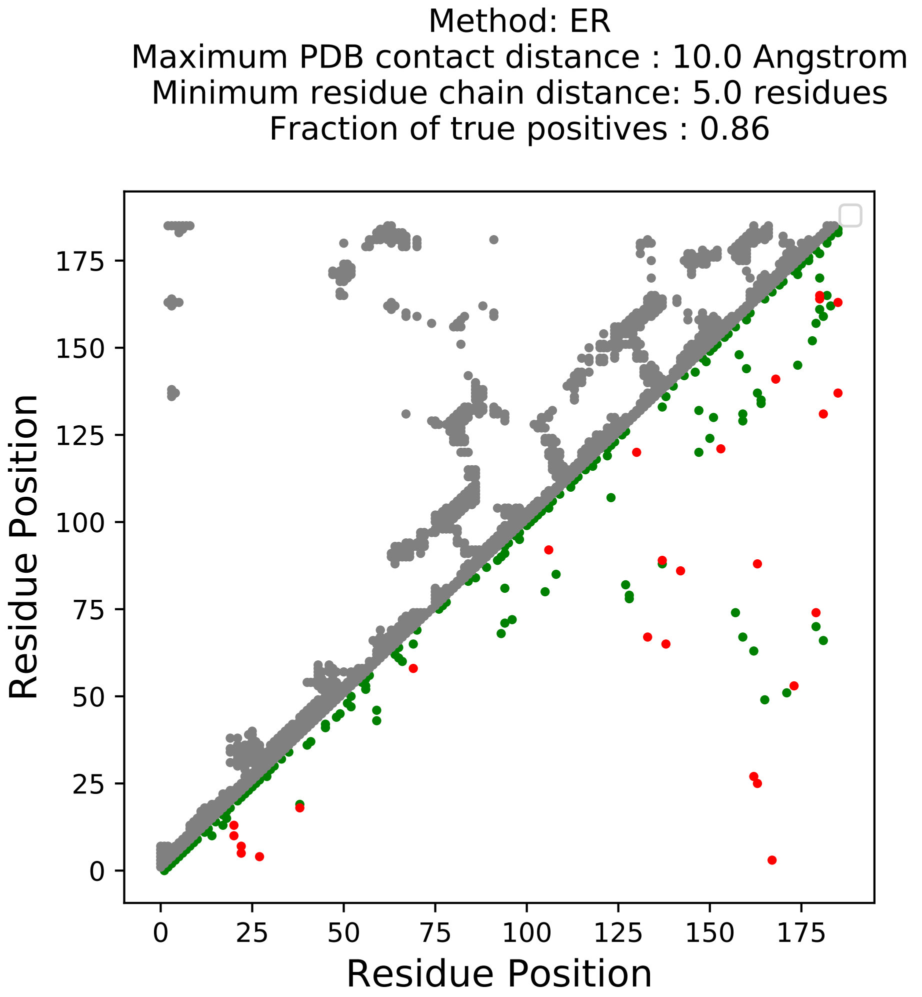

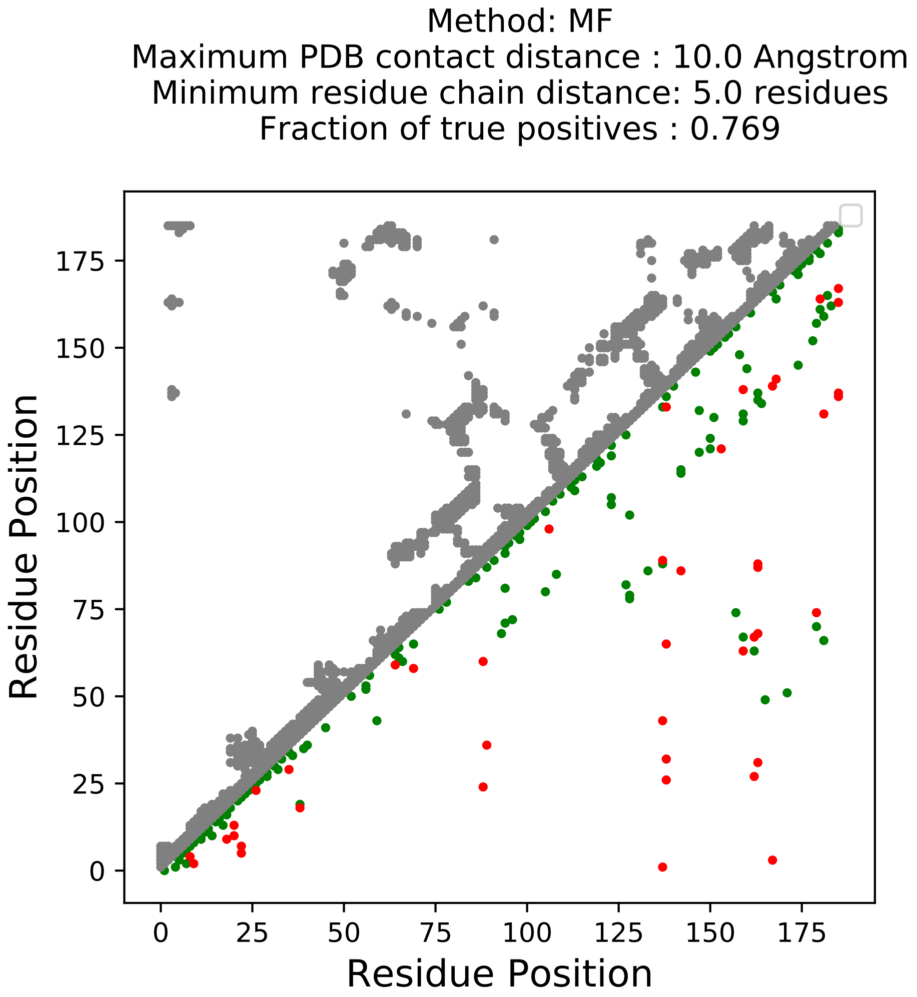

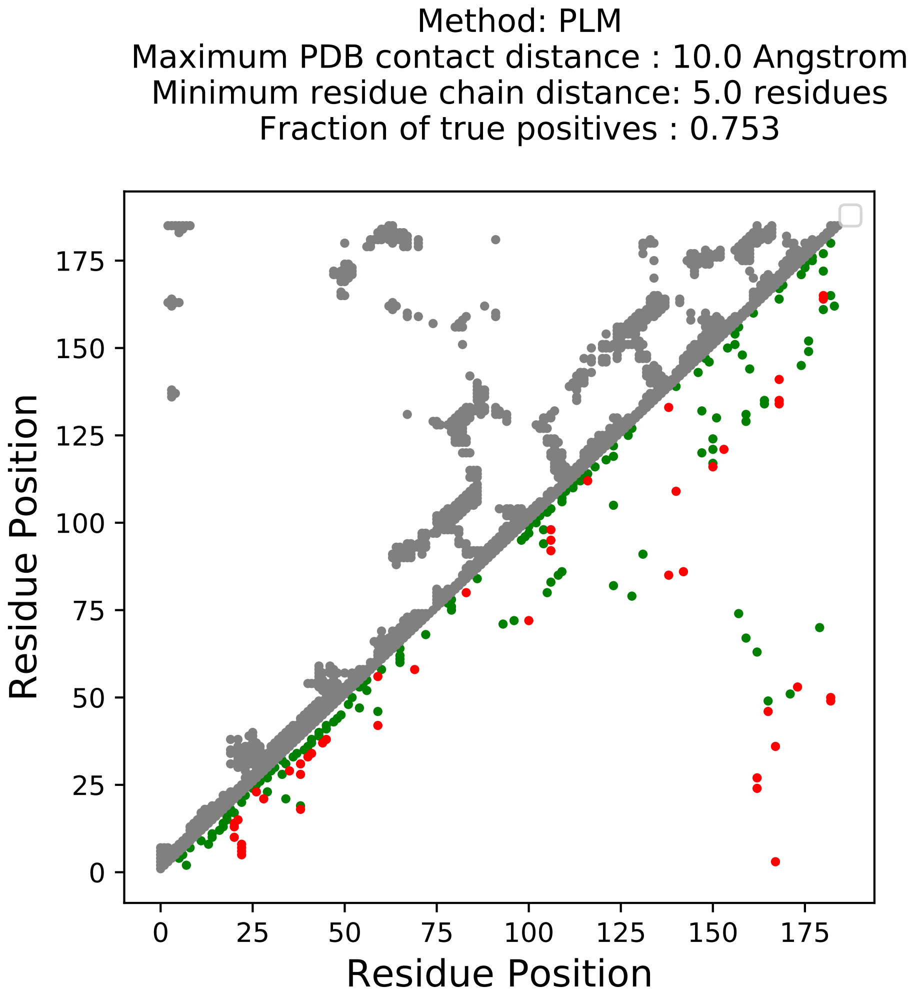

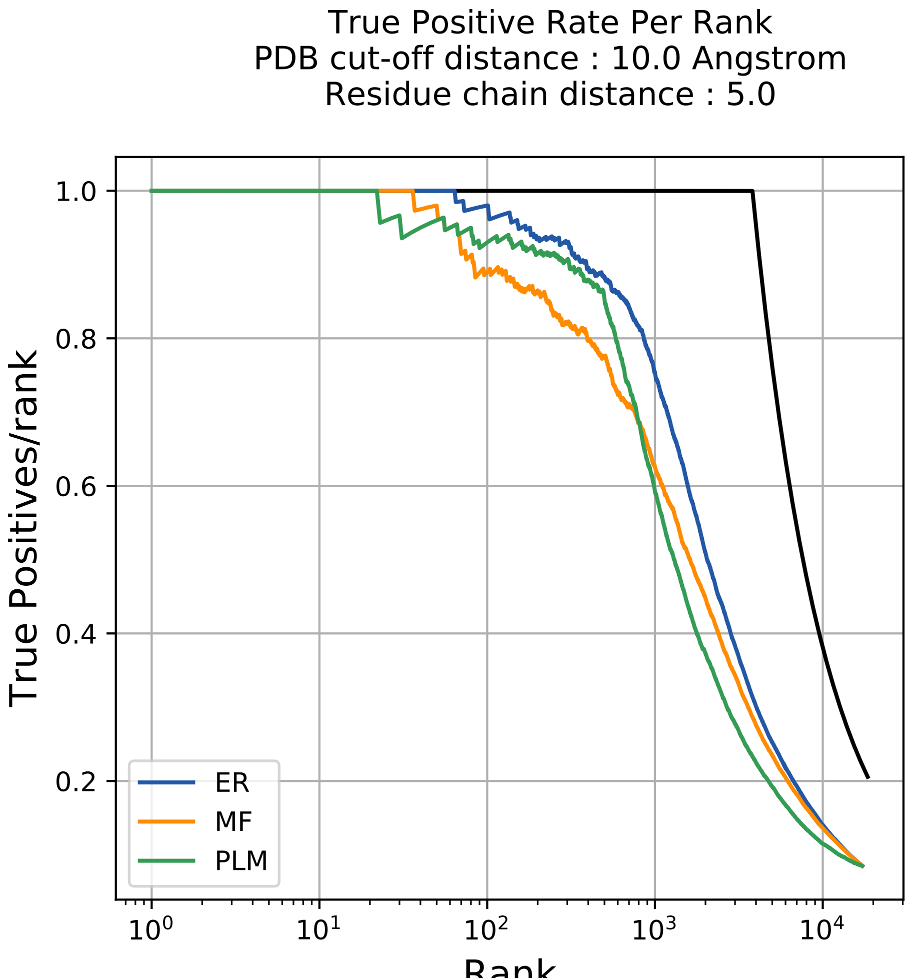

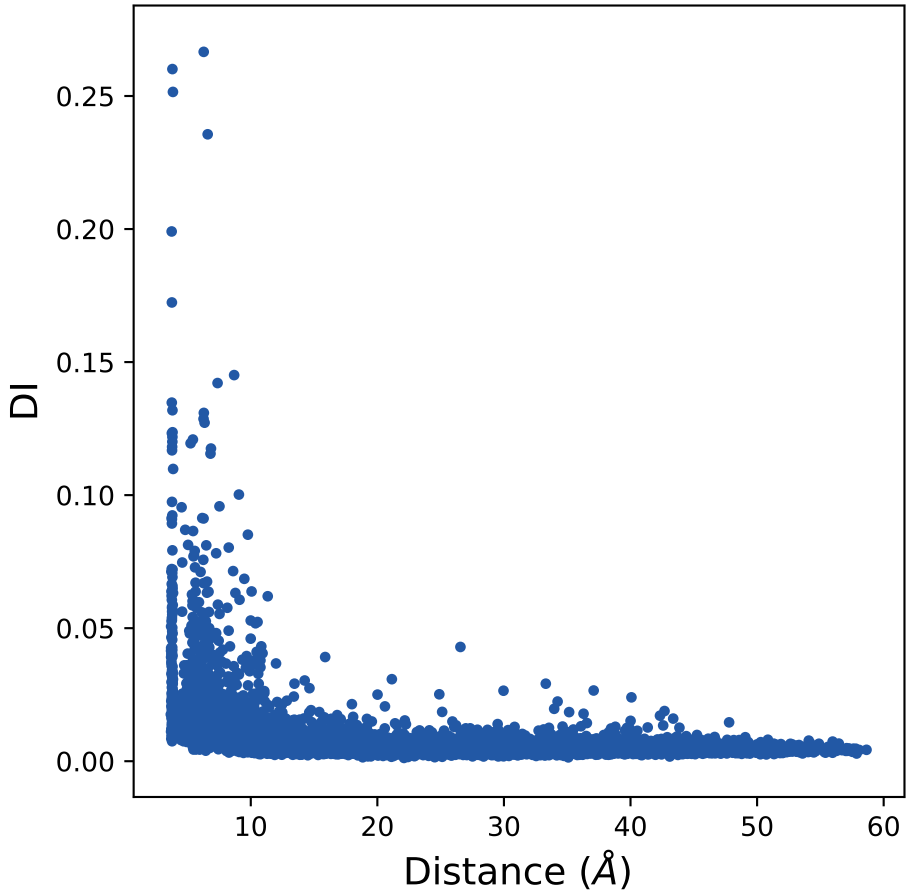

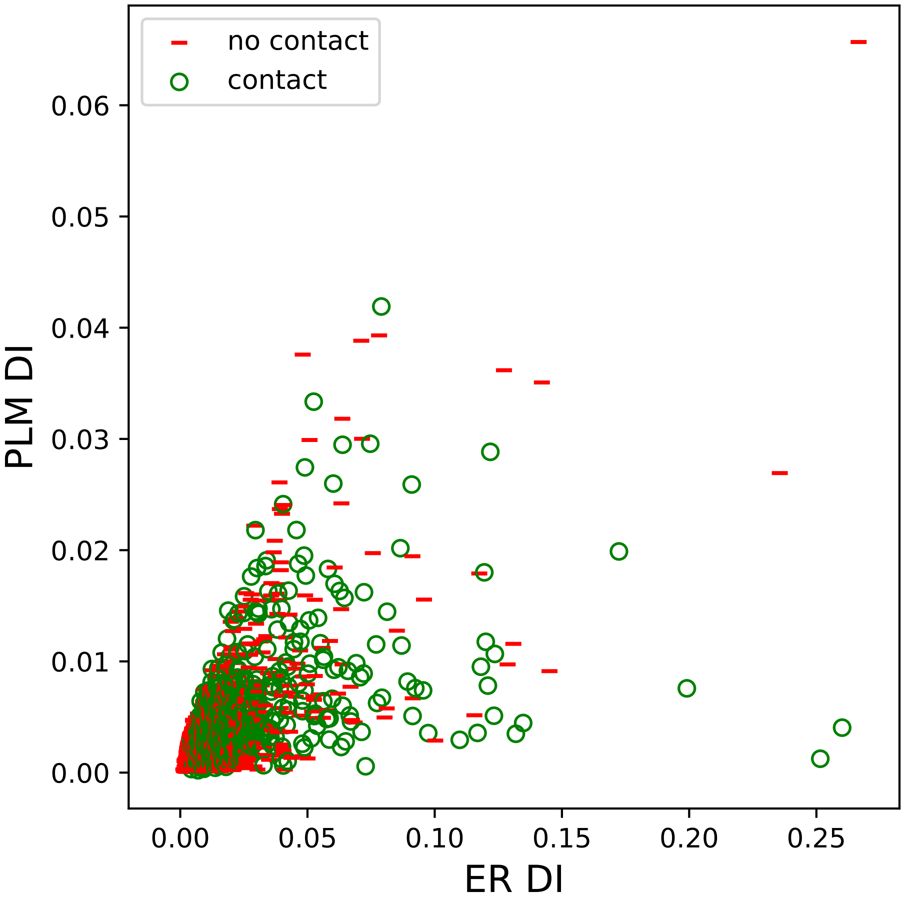

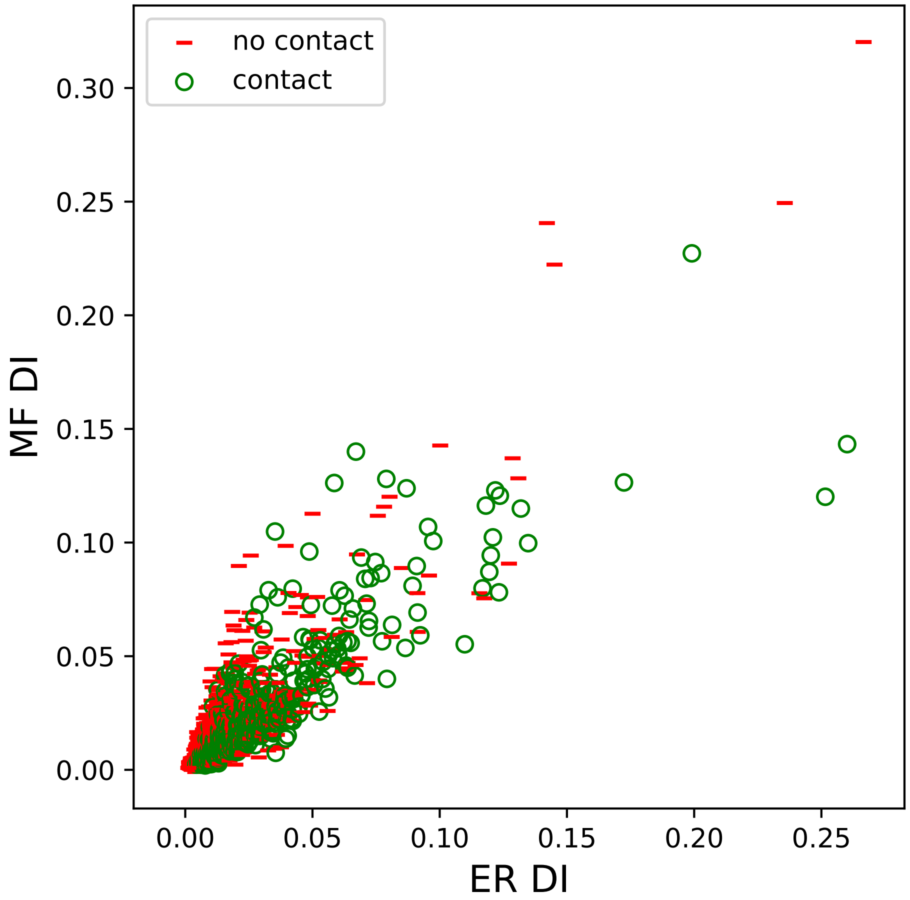

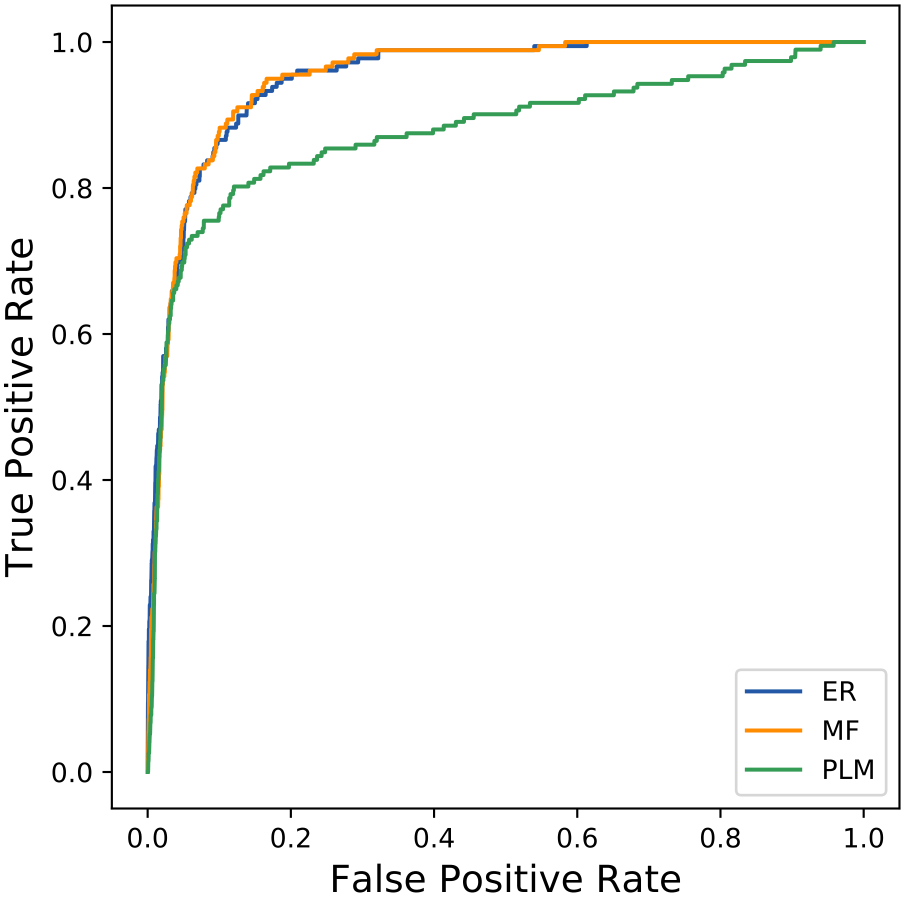
